# An allele-resolved nanopore-guided tour of the human placental methylome

**DOI:** 10.1101/2023.02.13.528289

**Authors:** Michaela Kindlova, Hannah Byrne, Jade M Kubler, Sarah E Steane, Jessica M Whyte, Danielle J Borg, Vicki L Clifton, Adam D Ewing

## Abstract

The placenta is a temporary organ present during pregnancy that is responsible for coordinating all aspects of pregnancy between the mother and fetus. It has a distinct epigenetic, transcriptomic, and mutational landscape with low levels of methylation, high numbers of transcribed loci, and a high mutational burden relative to somatic tissues. We present this landscape through the application of nanopore sequencing technology to provide a more comprehensive picture of female placental genomics and methylomics along with integrated haplotype-resolved transcriptomic analyses across eight trios. Whole genome sequencing of trios allows robust phasing, permitting comprehensive genome-wide investigation of parent-of-origin methylation and transcription. This enhanced view facilitates identifications of many new differentially methylated regions (DMRs), both conserved and differing between individuals, as well as novel imprinted genes including ILDR2 and RASA1 which are potentially important for healthy placental and fetal development.

## Introduction

The placenta is a unique cohort of cell types that collectively present an unusual epigenomic and transcriptomic profile, one that is required for establishing and maintaining healthy fetal development in placental mammals. It has been observed that the placenta has a lower level of DNA methylation when compared to other somatic tissues (Ehrlich et al. 1982; Robinson and Price 2015), with hypomethylation concentrated in partially methylated domains (Schroeder et al. 2013, 2015). The placental transcriptome includes a greater number of genes that are uniquely expressed (Gong et al. 2021) and uniquely depleted (Gong et al. 2022) than other tissues, perhaps in part owing to the distinctive epigenomic landscape. This landscape includes many imprinted genes, which are expressed predominantly from either the maternal or paternal allele and are a feature of organisms that transfer nutrients directly to their developing offspring (Moore and Haig 1991). The placenta mediates this interface in mammals and so is a natural epicentre for imprinted genes, where the balance between nutrient consumption and fetal growth is ultimately controlled (Frost and Moore 2010; Coan et al. 2005; Hanna 2020).

Here, we take advantage of technological and methodological advancements in nanopore sequencing to better understand the unique epigenomic landscape of the human placental genome. Nanopore sequencing was initially conceived in 1989 (Deamer et al. 2016) and has conceptual origins in patch clamp technology developed in the late 1970s for electrophysiology (Neher and Sakmann 1976). In recent years, the technology has been subject to rapid development and commercialisation by Oxford Nanopore Technologies Ltd. (ONT). ONT devices offer arbitrarily long and increasingly accurate reads that directly encode both canonical and modified nucleic acid base calls. Through the incorporation of genetic variation, nanopore sequence data can readily be phased into its constituent alleles via read-backed phasing of variants followed by tagging of individual reads (Patterson et al. 2015). In combination with one of the available highly accurate methods for discriminating between 5mC and canonical cytosine (Yuen et al. 2021), phased long reads enable examination of differentially methylated regions (DMRs) with unprecedented fidelity (Kolmogorov et al. 2023).

We carried out whole-genome sequencing (WGS) on ONT platforms at greater than 20-fold coverage with a median per-sample N50 of 34kbp, along with Illumina WGS at 30-fold coverage on eight female placentas (Supplemental Table 1). We opted to focus this study on female placentas as they are typically diploid throughout the genome (i.e. two copies of the X chromosome). Previous studies suggest the female placenta induces a greater genomic response to adverse environmental impacts (Clifton 2010). All placentas were obtained from participants in the Queensland Family Cohort (Borg et al. 2021) with uncomplicated pregnancies. Human research ethics approval for the study was granted by the Mater Misericordiae Limited HREC (HREC/MML/73929). Parental DNA was obtained from PBMCs and sequenced on the Illumina NovaSeq 6000 platform to a minimum 30-fold coverage. RNA sequencing was carried out via Illumina sequencing on the same eight placentas with a target throughput of 100M paired-end reads per sample. Sequencing metrics are presented as Supplemental Table 1.

### Comprehensive allele-resolved methylation profiling

Backed by long-read genome sequence data, we used informative germline variants to phase the placental genome into maternal and paternal alleles, enabling downstream studies of allele-specific methylation and transcription. This also enables a direct assessment of maternal cellularity in the bulk homogenised placental sample through examination of the variant allele fraction (VAF) of maternally inherited alleles, as substantial deviation upwards from a mean VAF of 0.5 would indicate excess maternal cells in the sample. This was not observed to a notable degree, with all samples showing less than 1% deviation from expectation in VAF for maternally or paternally inherited alleles (Supplemental Table 2), indicating little to no anticipated impact of maternal cells on subsequent analyses.

Comparison of methylation profiles derived from direct modified basecalling of nanopore sequence data enables both a comprehensive genome-wide view of placental methylation patterns and a highly detailed view at individual loci (Simpson et al. 2017). The placental genome is substantially demethylated relative to the heart, liver, and hippocampus tissues we have previously analysed (Ewing et al. 2020), with an apparently bimodal distribution of methylation levels (Figure 1a). Visualisation of the chromosome-wide methylation pattern shows that, with the exception of the X chromosome, relative demethylation is not chromosome-wide, but is concentrated in distinct regions (Figure 1b, see Supplemental Figure 1 for all chromosomes). This is consistent with the prior observation of partially methylated domains (PMDs) across the placental methylome (Schroeder et al. 2013, 2015).

**Figure 1:**
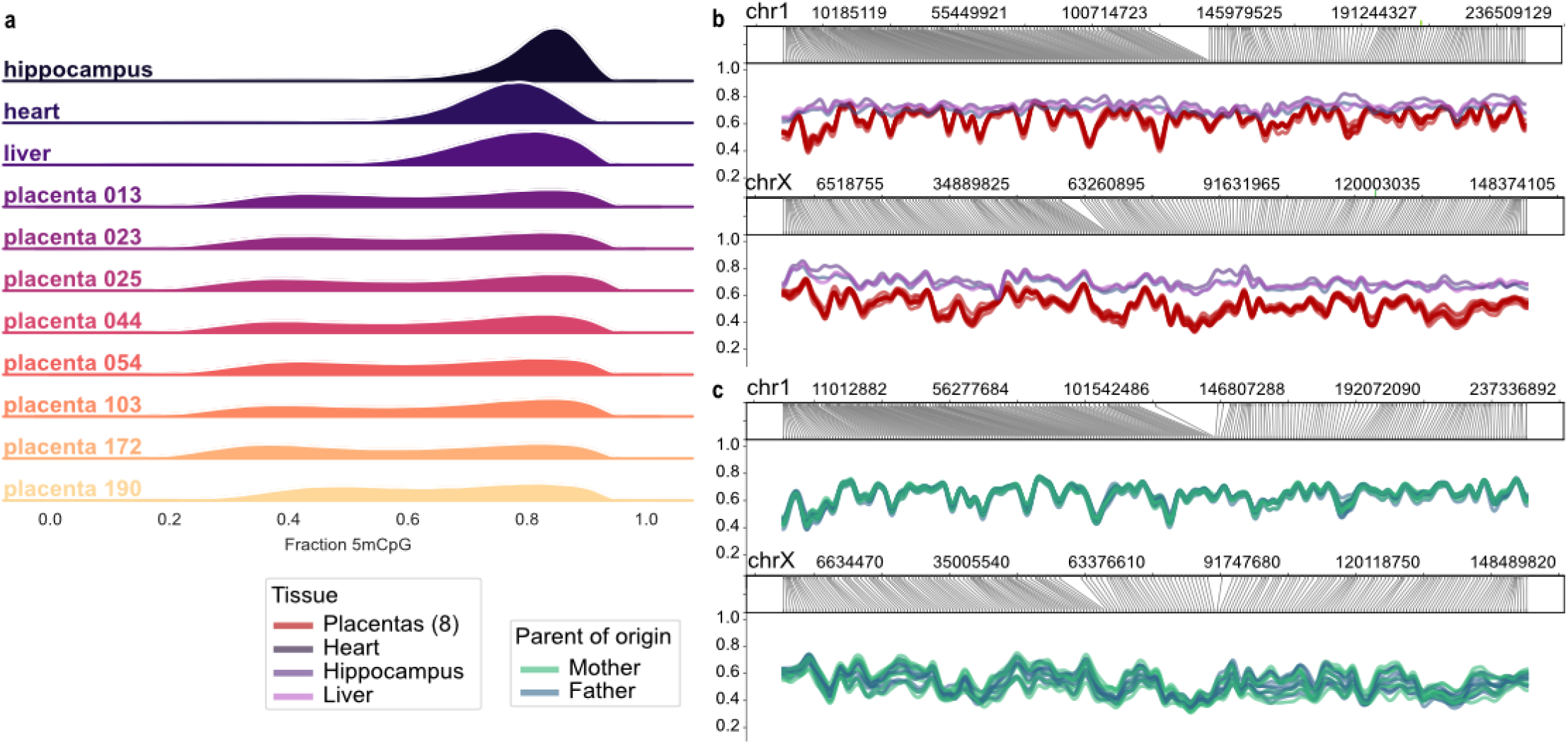
Large-scale methylation patterns in human placentas. (a) The distribution of methylation levels for non-placental tissues versus placentas in this study. (b) The same samples as in (a) showing the distribution of methylation across chromosomes 1 and X. For all chromosomes see Supplemental Figure 1. Profiles for all chromosomes are presented in Supplemental Figure 1. (c) The methylation profiles for placental samples phased into paternal and maternal profiles are shown for chromosomes 1 and X. For all chromosomes see Supplemental Figure 2.

Using read-backed phasing (Patterson et al. 2015) coupled with parental variants, attribution of reads as maternal or paternal in origin is straightforward. Comparison of alleles on a chromosomal scale yields highly similar methylation profiles between maternal and paternal autosomes (Figure 1c, Supplemental Figure 2). The X chromosome is an interesting exception, with some but not all placentas showing globally higher methylation of either the maternal or paternal chromosome. (Figure 1c, Supplemental Figure 3) We ascribe this to clonal X-inactivation yielding a “patchy” geographic profile of maternal or paternal-specific methylation across the X chromosome, as previously described (Phung et al. 2022). The use of bulk tissue captures a stochastic clonal profile when homogenised, yielding the observed profiles.

On a locus-specific level, we observe many known differentially methylated regions (DMRs) such as those associated with IGF2/H19, and GNAS (All DMR visualisations are included as Supplemental Data). Extending this to catalogue all DMRs, we identify 723 DMRs in total, with 184 specific to this study and 539 previously identified DMRs as annotated by a collection of studies providing specific DMR coordinates (Court et al. 2014; Hanna et al. 2016; Sanchez-Delgado et al. 2016; Jima et al. 2022; Monteagudo-Sánchez et al. 2019; Hamada et al. 2016) (Supplemental Figure 4, Supplemental Table 3). Of the 184 newly identified DMRs, 41 are near genes annotated as at least putatively imprinted, while 143 are not near a known imprinted gene. DMRs were observed to include those consistent across all samples and those present in fewer than all eight placentas analysed, consistent with polymorphic DMRs noted in previous studies (Hanna et al. 2016; Hamada et al. 2016; Yuen et al. 2009).

A known visually striking example is the chromosome 19 micro-RNA cluster (“C19MC”), which encodes approximately 50 placentally-expressed paternally-imprinted miRNAs (Noguer-Dance et al. 2010). The miRNAs at this locus may be essential for trophoblast differentiation (Kobayashi et al. 2022), and their deregulation may be linked to pre-eclampsia and other complications (Hromadnikova et al. 2015). The extent and the consistency of the differential methylation, as shown in Figure 2a is remarkable, and we observe a pattern of steadily decreasing differential methylation across the locus.

**Figure 2:**
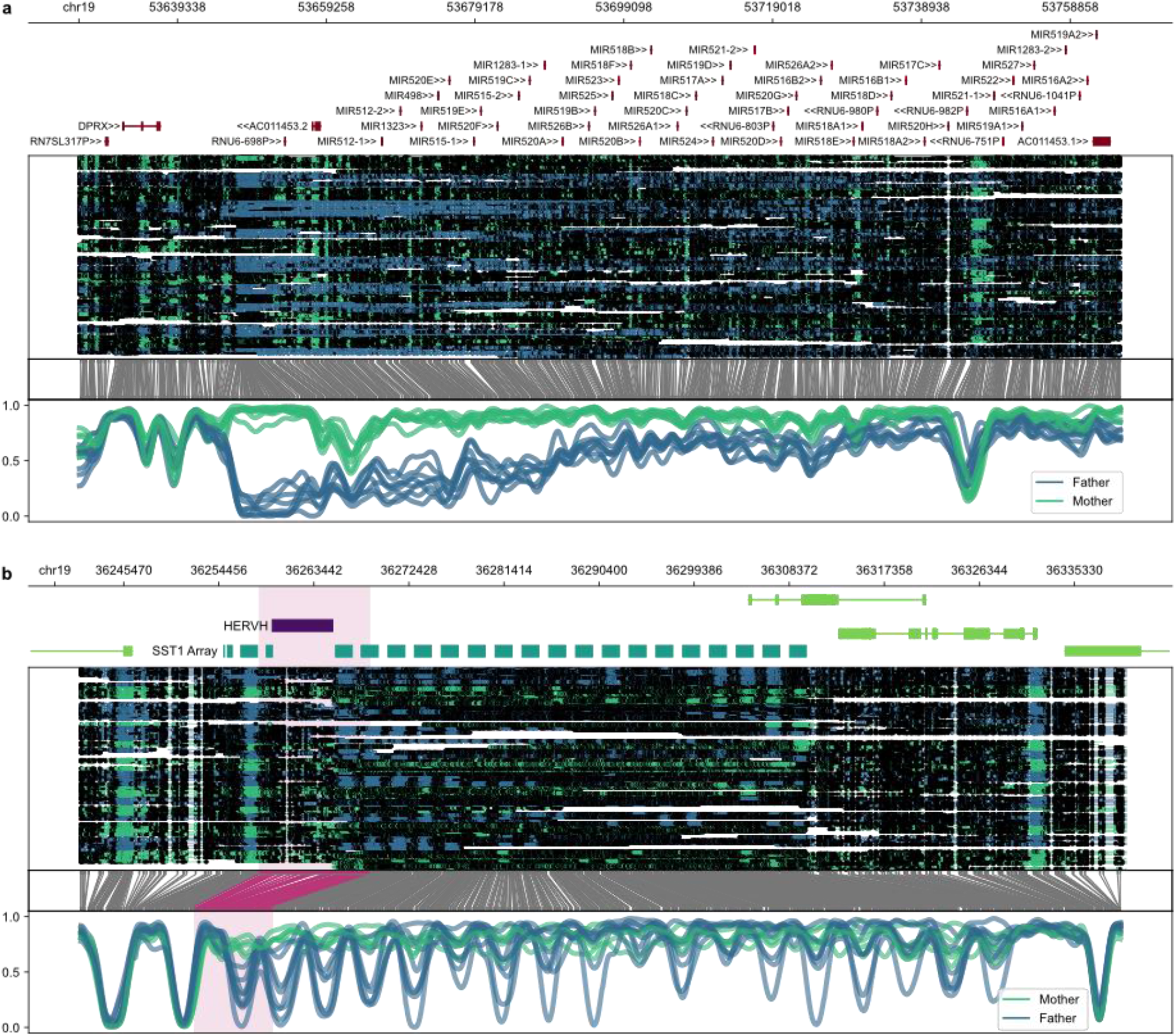
Examples of regions with notable allele-specific methylation patterns in placenta. From the top down, each panel shows the genomic coordinates, relevant annotations, read alignments with methylated CpGs as closed dots, translation from genomic space into CpG-only space, and a smoothed methylation plot at the bottom. The C19MC locus (a) contains many paternally-expressed (Supplementary Figure 5) and paternally-demethylated pri-miRNA loci and is among the best known examples of a placental DMR. Here we can see the pattern of differential methylation across the locus. The SST1/MER22/NBL2 macrosatellite array (b), also on chromosome 19, is a novel example with a periodic pattern of DMRs corresponding to the SST1 repeats, originating at a HERVH element upstream of the array. This locus is also paternally expressed, as shown in Supplemental Figure 6.

To highlight an interesting example of a novel DMR-rich region, we identified that an array of SST1 macrosatellite repeats on chromosome 19 has a curious methylation pattern, which exhibits periodic methylation and demethylation of the paternal allele corresponding to the positions of SST1 repeats (Figure 2b). The locus is also paternally expressed with transcription initiating at an intact HERVH retrotransposon upstream of the differentially methylated portion of the array (Figure 2b). While SST1, also known as MER22 or NBL2, is highly variable in copy number and is present in arrays at multiple locations in the genome, this particular array on chr19q is one of the more ancestral and stable instances (Hoyt et al. 2022).

Finally, our data confirm a strong bias toward paternal imprinting in the placenta (Wang et al. 2013) (Figure 3a), with the large majority of DMRs being demethylated on the paternal allele relative to the maternal allele. Comparison of parent-of-origin between placenta and the NA19240 lymphoblastoid cell line (De Coster et al. 2019; Byrska-Bishop et al. 2022) shows that this bias is not present in NA19240 (Figure 3a). When compared to other tissues and cell lines, most placental DMRs show a greater absolute magnitude of differential methylation in placentas relative to the same site in other tissues or cells (Figure 3b). With the caveat that a limited number of nanopore-sequenced tissue samples are available for comparison at this time, this indicates a high degree of placental specificity for the DMRs identified here, along with some cross-tissue conservation of DMRs, which is also of interest.

**Figure 3:**
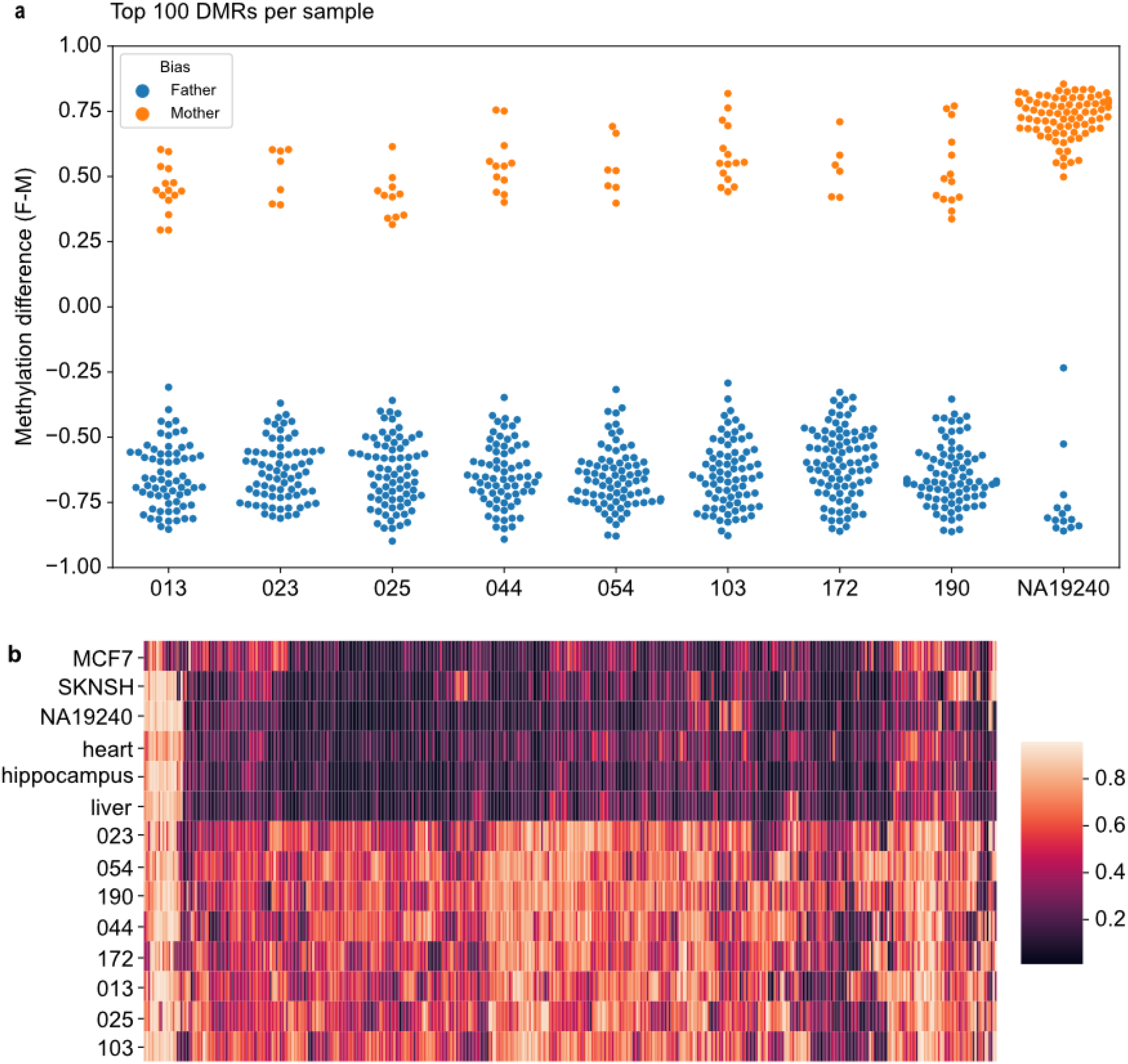
Properties of placental DMRs. **(a)** Placental DMRs are highly biased towards paternal demethylation when compared to a non-placental sample (NA19240, EBV-transformed lymphoblastoid cell line). For each sample, the top 100 DMRs from analysis with DSS are shown. **(b)** Placental DMRs are more differentially methylated in placenta versus other tissues or cell lines. Lighter colour corresponds to a greater absolute difference in methylation fraction between alleles.

### Allele-resolved expression

Deep cDNA sequencing was carried out on the placental samples and separated into maternal and paternal components where possible through the presence of one or more phased variants. Multidimensional scaling of the overall transcriptional profile suggests the variance is largely between individuals rather than between parents of origin, as one might expect (Figure 4a). In contrast, when this analysis is carried out on loci immediately surrounding DMRs, parent of origin emerges as a substantial source of variance, reflected in the separation of maternal and paternal expression (Figure 4b). As phasing is limited to reads that contain informative variants, the ability to phase short-read RNA-seq data is limited, particularly for RNA-seq where coverage is biased towards exons. Furthermore, estimates of differential allelic expression over a given locus may be possible for some individuals but not others given that informative variants are segregating. Nonetheless, imprinted genes are readily identifiable through differential gene expression between maternal and paternal alleles; an analysis of allelic differential expression of genes identifies 74 imprinted genes at FDR < 0.25 (Figure 4c, Supplemental Table 4).

**Figure 4:**
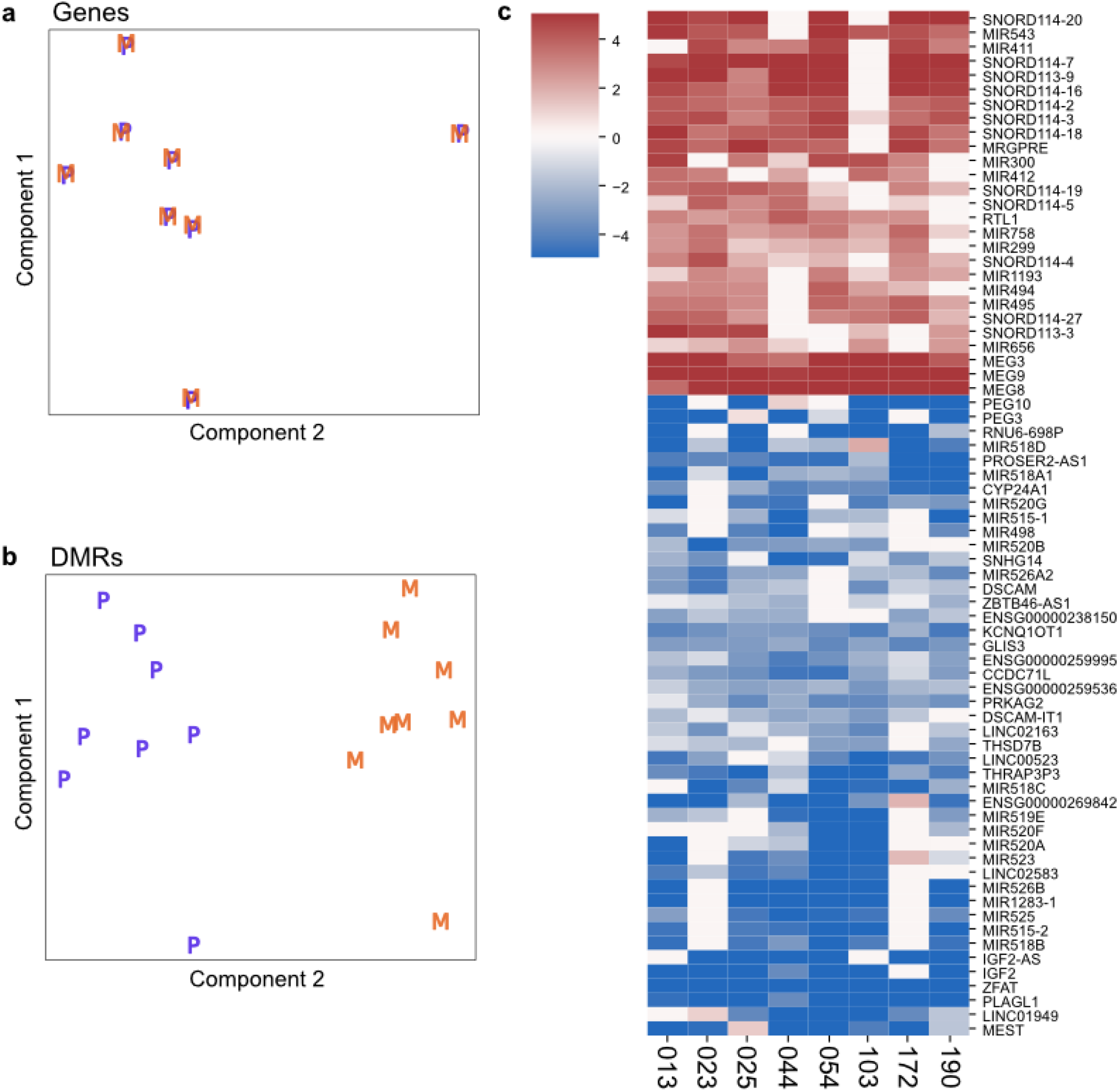
Allele-specific expression via read-backed phasing. Overall **(a)**, multidimensional scaling shows variability in gene expression is attributable to differences between individuals with negligible variance due to maternal versus paternal allele. This is the expected result given most expressed loci are not imprinted or expressed in an allele-specific manner. When focusing specifically on expression surrounding DMRs (+/- 5kbp) **(b)**, parent-of-origin becomes a major driver of variance in gene expression at DMR loci (M = maternal, P = paternal). Statistical analysis of allele-specific expression **(c)** shows many known imprinted loci - most of the maternally expressed examples are non-coding and short RNAs clustered around the MEG3/8/9 locus on chromosome 14. The paternally-expressed loci are more heterogeneous but show a plurality of paternal expressed miRNAs at the C19MC locus (Figure 2a). As genetic variability involving informative phased SNVs induces considerable dropout (white cells in heatmap), and consequently high variability in expression, the FDR is set at 0.25 to capture the most differentially expressed loci for this figure.

### Discovery of novel imprinted genes

Intersecting the DMRs described here with our allele-specific expression results has the potential to uncover previously unrecognised imprinted genes. The most prominently notable examples are ILDR2, which appears to be paternally imprinted, and RASA1, which is maternally imprinted.

ILDR2 is a member of the Ig superfamily and is a B7-like protein known to modulate T cell activity, thereby potentially serving an important function in the regulation of autoimmune responses (Hecht et al. 2018; Podojil et al. 2018). To our knowledge, ILDR2 has not previously been identified as imprinted. Our data indicate a strong paternally-demethylated DMR corresponding to ENCODE-derived CTCF-bound cis-regulatory elements around the penultimate exon of most ILDR2 isoforms, with essentially all transcription derived from the paternal allele (Figure 5a). Together, this suggests ILDR2 is a novel paternally imprinted gene that may play a role in suppression of active T cells in the human placenta.

**Figure 5:**
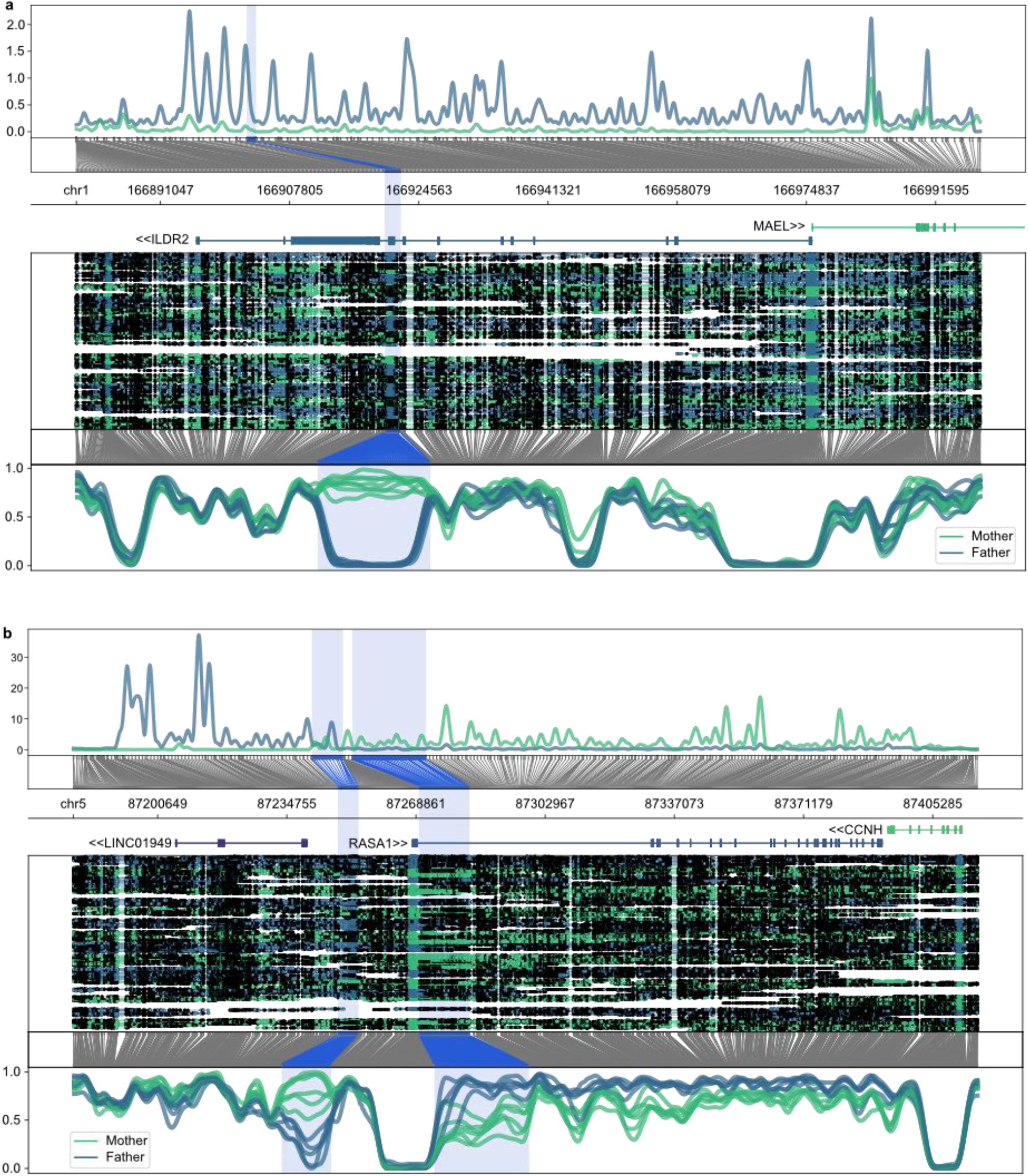
Previously unreported imprinted genes **(a)** ILDR and **(b)** RASA1. In each panel, the top subpanel shows allele- specific expression merged across all samples reduced to a space that includes only bases with a minimum coverage (smoothed CPM > 0.1) in the RNA-seq data. Here, ILDR2 is clearly paternally expressed while RASA1 is maternally expressed. The lncRNA upstream of RASA1 appears to be paternally expressed. The correspondence between this coordinate space and genomic coordinate space is shown below the allele-specific expression plot, followed by coordinates and gene annotations. Highlights correspond to DMR locations. The remaining subpanels (alignment, % mCpG) are as described in Figure 2.

RASA1 (RAS p21 activator 1 or p120-RasGAP) is a novel example of a maternally imprinted gene (Figure 5b). It is part of the RAS/MAPK pathway where studies in trophoblasts suggest it plays an inhibitory role, and its expression may attenuate trophoblast proliferation and invasion in the placenta (Zhang et al. 2021a). The maternally imprinted gene MEG3 has been suggested as an upstream silencer of RASA1 in this context via recruiting of EZH2, leading to H3K27me3-mediated silencing of RASA1 (Zhang et al. 2021b). Beyond the placenta, mutations impacting RASA1 are implicated in an autosomal dominant disorder with variable presentation including abnormal vascular growth and limb hypertrophy (Eerola et al. 2003; Revencu et al. 2013). The maternally imprinted DMR is downstream of a nearby paternally imprinted DMR, which seems to correspond to imprinted expression of the lncRNA LINC01949 (Figure 5b). Together, the region encompassing the 5’ end and upstream of RASA1 appears to represent an imprinting control region for multiple genes exhibiting both paternal and maternal imprinting as has been observed at other ICRs.

Tumour suppressor candidate 3 (TUSC3) is an example of “epipolymorphism”, where a previous study remarked on the variability of maternal-specific methylation and a potential association with preeclampsia (Yuen et al. 2009). Here, we can confirm the specific location of the DMR at the TUSC3 5’UTR as well as show a corresponding bias towards expression of the paternal allele (Supplemental Figure 7). It is interesting to note that gene body methylation of TUSC3 has an inverse relationship with DMR methylation in that while the paternal allele is demethylated relative to the maternal allele in 4 of 8 samples, the opposite is true of the gene body. This is consistent with a high level of methylation of transcribed gene bodies previously observed in the placental methylome (Schroeder et al. 2015). WNT2 is another epipolymorphic gene noted in the same study (Yuen et al. 2009) with a comparable pattern of inconsistent differential methylation with differential parent-of-origin expression (Supplemental Figure 8).

### Genome variation

It has been noted that an unusually high number of somatic mutations are detectable from bulk placenta (Coorens et al. 2021; Hannibal et al. 2014). Comparison of placental genomes to parental genomes indicated an average of 186 point mutations (single base substitutions and short insertions/deletions) per sample not detectable in the parental genome or in gnomAD at an allele frequency greater than 10^−5^ (Figure 6a, Supplemental Table 5). On average, 102 of these have a VAF significantly below 0.5 as assessed by a one-sided binomial test; a VAF lower than 0.5 suggests a *de novo* somatic mutation rather than a *de novo* constitutive mutation. This finding is consistent with a recent report of widespread somatic mutation in placental genomes (Coorens et al. 2021).

**Figure 6:**
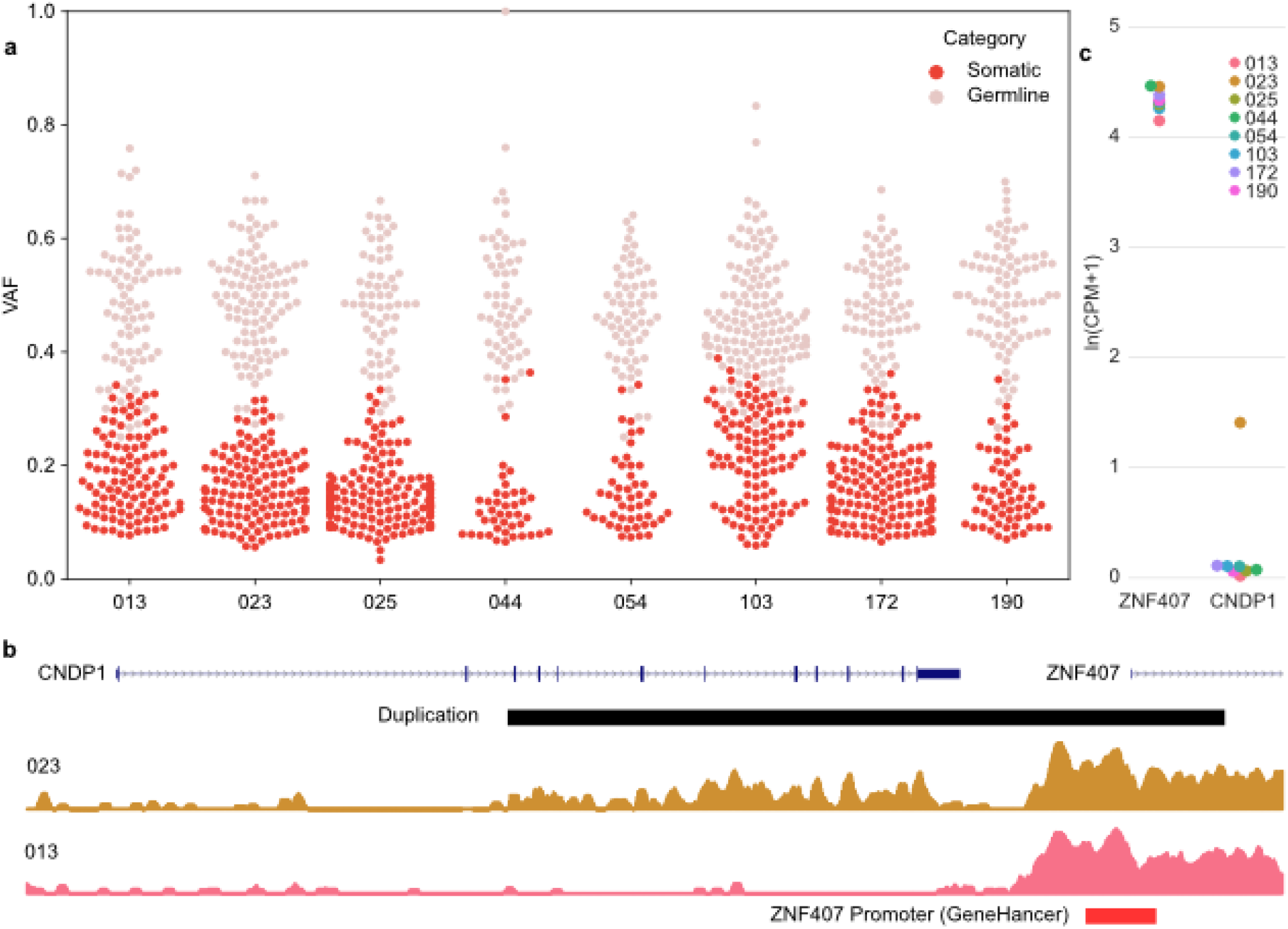
Detection of *de novo* placental variants. (a) Variant allele fractions of point mutations identified from Illumina sequencing in one placenta only, absent from all parental samples, and confirmed in Nanopore sequence data. VAFs significantly below 0.5 via proportion test using read counts are highlighted in dark red. **(b)** Identification of a duplication involving most of CNDP1 and the promoter of ZNF407 (red box). RNA-seq read depth is shown for the sample with the duplication (023, orange) and a representative sample without the duplication (013, pink). **(c)** Elevated CDNP1 expression is unique to the duplicated sample and may be due to the rearrangement bringing the promoter of ZNF407, which is expressed in all 8 placentas, upstream of the duplicated portion of CDNP, consistent with the read depth plot.

Placenta-specific structural variants were considerably less numerous, with only 4 identified concordantly in both short and long read data (Supplemental Table 5b). The most notable among these is a 44.7 kbp duplication involving most of CNDP1 and the promoter of a downstream gene, ZNF407 (Figure 6b). While CNDP1 is not expressed in the placentas without this duplication, the sample with the duplication expresses CNDP1 downstream of the duplication 5’ junction. The promoter of ZNF407, which is highly expressed in all eight placentas, presumably drives transcription of CDCP1 in the sample with this structural rearrangement. Based on VAF (∼0.1 for both Illumina and Nanopore), absence from the parental genomes, and a relatively low level of ZNF407-driven CNDP1 expression as compared to ZNF407, it seems likely that this duplication is somatic. While there are examples of placenta-specific duplications associated with adverse fetal outcomes (Del Gobbo et al. 2021), that was not the case with the duplication reported here. However, this does add to evidence that a small minority of somatically acquired mutations affect the placental transcriptome and a subset of these instances may impact fetal development.

## Discussion

In this study, we have applied the advantages of nanopore sequencing technology to the unusual epigenomic environments of the human placenta, yielding insights and avenues for further study. Through the application of this technology to the samples available through the Queensland Family Cohort, we have been able to explore the allele-specific methylation profiles of a placental cohort to identify a substantial number of DMRs not previously described. We have confirmed and provided better visualisation of hundreds of known DMRs alongside identification of many new DMRs and novel imprinted genes. Overall, differential methylation between parental alleles is more prevalent than differential gene expression between alleles. Speculatively, this could indicate loci primed for differential gene expression between parental alleles, but perhaps occurring in a different cell type than the placenta or at an earlier developmental time point.

The genetic conflict hypothesis is a leading theoretical explanation for genomic imprinting that posits a motive for regulation of genes, particularly those involved in growth, in a manner specific to either the maternal or paternal allele (Moore and Haig 1991). In this paradigm, the regulation of the maternal genome has evolved to optimise the distribution of resources between her and her offspring in a way that is compatible with multiple offspring over time. This conflicts with the paternal genome, which has evolved a regulatory regime to support as much growth as possible in each offspring. Therefore, the argument follows that genes driving placental growth will tend to be paternally imprinted while genes that inhibit growth will tend to be maternally imprinted, as reviewed in (Wilkins and Haig 2003). The novel imprinted genes we highlight here are compatible with this model. RASA1, which is maternally imprinted, inhibits the activity of the RAS/MAPK pathway, thereby inhibiting trophoblast proliferation and invasion. The paternally imprinted ILDR2 modulates activated T cells which is consistent with promoting placental growth through reduction of maternal immune response towards the placenta.

This study highlights the technical capabilities and utility of nanopore sequencing technology and associated software tools with their combined ability to capture modified bases (5mCpG in this manuscript) and genomic variation. What has been uncovered here forms the basis for using a similar approach to study the function of these loci in a targeted manner and the impetus for expanding this cohort such that association between variation in DMR status and relevant outcomes is adequately powered. Male placentas are a notable omission from the present study, and their future inclusion may enable additional insights into why females are better able to adapt to environmental stressors *in utero* than males (Clifton 2010). The unique epigenomic profile of the placenta surely contains many yet unrecognised signals linked to the developmental origins of health and disease.

## Methods

### DNA Sequencing

Placental samples and parental blood samples were obtained from the Queensland Family Cohort (QFC) (Borg et al. 2021). High molecular weight (HMW) DNA for sequencing was extracted from the placental tissue using the Nanobind Tissue Big DNA kit (Circulomics). For each placenta, approximately 40mg of tissue was homogenized using a pre-sterilised pestle in lysis buffer at 4°C and DNA was extracted as per manufacturer’s protocol. Where required, samples were additionally purified with Ampure Beads XP and 80% EtOH. Samples were treated with the Short Read Eliminator (SRE) kit (Circulomics) as per manufacturer’s protocol to eliminate fragments <25 kb prior to sequencing.

DNA for short read whole genome sequencing was extracted using a DNEasy kit (Qiagen). For the placental samples approx. 35mg of tissue was used as the starting input and DNA was extracted as per manufacturer’s protocol with a tissue lysis time of 90 minutes. For the parental genomes, 250ul of PBMCs in 10% DMSO was washed twice with PBS and then resuspended in 250ul of PBS before DNA was extracted as per the manufacturer’s protocol. Concentration and purity of all DNA samples was assessed by qubit and nanodrop, respectively.

Short read whole genome sequencing was carried out on all samples by Macrogen Oceania on an Illumina NovaSeq 6000. Short reads were mapped to the reference genome (hg38, GATK Resource Bundle) using bwa-mem2 2.0pre2 (Vasimuddin et al. 2019) and duplicates were marked via samblaster 0.1.26 (Faust and Hall 2014).

Long read nanopore sequencing was carried out on the placenta samples at the Australian Centre for Ecogenomics on an Oxford Nanopore Technologies (ONT) PromethION on version 9.4.1 flowcells. Where required to obtain comparable coverage across all samples, Additional top-up sequencing was carried out on ONT Mk1c devices using version 9.4.1 flowcells. Bases (A, C, G, T, 5mCpG) were called with Megalodon 2.5.0 using the models included with guppy 5.0.7 dna_r9.4.1_450bps_modbases_5mc_hac_prom for Promethion data and dna_r9.4.1_450bps_modbases_5mc_hac for MinION data. Reads were aligned to the reference genome (hg38, GATK Resource Bundle) using minimap2 2.22 (Li 2018). Visualisations were created using methylartist 1.0.6 (Cheetham et al. 2022).

### cDNA Sequencing

Total RNA was isolated from the frozen placenta samples using an RNeasy extraction kit (Qiagen). Placental tissue was kept frozen by liquid nitrogen and powdered using mortar and pestle. Aliquots of approx. 30 mg of tissue were resuspended in a lysis buffer containing 1% *β*-mercaptoethanol and immediately homogenised using ceramic beads (MP Biomedicals) on a Bead Ruptor homogeniser (Omni Inc). Samples were subsequently spun at 13000 g for 10 min, supernatant was moved to a new tube, and RNA was isolated according to the manufacturer’s protocol. All samples were treated with Turbo DNase (Invitrogen) to avoid DNA contamination. Purity and concentration of RNA samples were checked on an Agilent Bioanalyser (RIN scores of samples used for further analysis ranged from 5.8 to 7).

Synthesis of cDNA using a RiboZero Plus protocol and sequencing were carried out by the Australian Genome Research Facility (AGRF) on an Illumina NovaSeq 6000 (150bp PE) with a total target depth of 100M PE reads per sample. Reads were aligned to the reference genome (hg38, GATK Resource Bundle) using STAR 2.7.10a (Dobin et al. 2012), duplicate reads were marked via samblaster 0.1.26 (Faust and Hall 2014). Aligned reads were assessed against quality control metrics using rnaseqc 2.4.2 (Graubert et al. 2021) and CollectRnaSeqMetrics from picard 2.27.5. Reads were phased using whatshap-phased variants as per long read DNA sequencing. Reads were counted against DMR locations and against Ensembl gene build 106 using featureCounts from Rsubread 4.1.2 (Liao et al. 2014). Read counts were assessed for differential expression using edgeR 4.1.2 (Robinson et al. 2010).

### Mutation detection SNVs/INDELs

To detect germline variants used for phasing, variants for each trio were called jointly with GATK 4.1.9 HaplotypeCaller and quality scores were recalibrated using GATK VQSR. Putative *de novo* and somatic variants were detected using GATK Mutect2 (Benjamin et al. 2019), with gnomAD 3.1.2 (Karczewski et al. 2020) as a germline resource, using the set of all parental genomes as a “panel of normals”. Mutect calls were filtered via GetPileupSummaries, CalculateContamination, and FilterMutectCalls and screened using a script to remove variants with a gnomAD allele frequency > 1e^-5^ and total read depths < 10. Putative *de novo* and somatic mutations detected in placentas from the Illumina WGS data were verified in the nanopore data from the same placenta.

### SVs

Structural variants were detected from short-read data via delly v1.1.6 (Rausch et al. 2012) on a per-trio basis, filtered for somatic variants in the placenta sample, genotyped against the whole cohort and further filtered for somatic variants versus the whole-cohort genotyping results. SVs from long-read data were detected via sniffles2 version 2.0.4 (Sedlazeck et al. 2018) using the –non-germline and –phased options, and delly-derived somatic SVs confirmed via these results were reported. Additional tools were used to screen for transposable element insertions on a per-trio basis: TEBreak for short-read data (Carreira et al. 2016) and TLDR for long-read data (Ewing et al. 2020).

### Phasing

Variants called from Illumina along with pedigree information (trios) were phased using nanopore reads via WhatsHap v1.4 (Patterson et al. 2015). Phased VCFs were used to tag nanopore reads using a combination of WhatsHap and customised methods. Manual examination of known DMRs (e.g. IGF2, GNAS, PEG3) was used as a quality control check.

### DMR detection

Genome-wide phased methylation values at each CpG were output via “methylartist wgmeth --dss”, which yields outputs suitable for import to the DSS differential methylation package implemented in R. DSS was used on both a per-individual and whole-cohort basis to identify differentially methylated loci. These were plotted with a +/- 20kbp window using “methylartist locus” and visually inspected to filter DMRs. Known DMRs were intersected via “bedtools intersect” to construct a single list of known DMRs and these were screened against DMRs detected via our approach. Previously reported DMRs that were not identified by our approach were also plotted via “methylartist locus” and visually inspected to assess whether the known DMR was apparent in our cohort.

### Data Availability

Sequence data generated in this study will be distributed through the UQ Research Data Management system (https://rdm.uq.edu.au/)

Nanopore sequence data from Heart, Liver, and Hippocampus tissues (Ewing et al. 2020) was obtained from SRA Bioproject PRJNA629858. Nanopore sequence data from NA19240 (De Coster et al. 2019) was obtained via ENA accession PRJEB26791. MCF-7 sequence data (Cheetham et al. 2022) was obtained from SRA Bioproject PRJNA748257.

## Acknowledgements

We gratefully acknowledge the families participating in the Queensland Family Cohort study which enabled this study. We acknowledge the Translational Research Institute (TRI) for research space, equipment, and facilities. This study was funded by the Australian Department of Health Medical Research Future Fund (MRFF) (MRF1175457 to A.D.E.), by the Advance Queensland Women’s Research Assistance Program (WRAP086-2019RD1 to M.K.), by a UQ Research Stimulus Package Fellowship (D.J.B.), by the National Health and Medical Research Council (NHMRC) (APP113100 to V.L.C.), and by the Mater Foundation. We acknowledge the sequencing service providers used in this study: The Australian Centre for Ecogenomics (ACE), Macrogen Oceania, and the Australian Genome Research Facility (AGRF).

## Author Contributions

ADE designed the study and analysed sequencing data. ADE, MK, and HB wrote the manuscript with feedback and input from all authors. MK, HB, JMK, SES, and JMW carried out sample preparation and quality control. JMK and SES provided valuable advice and expertise concerning the preparation of placental RNA. VLC and DJB facilitated provision of samples on behalf of the Queensland Family Cohort. VLC provided expertise and advice concerning study design and data analysis.

