## Supplemental Figures for "An allele-resolved nanopore-guided tour of the human placental methylome"

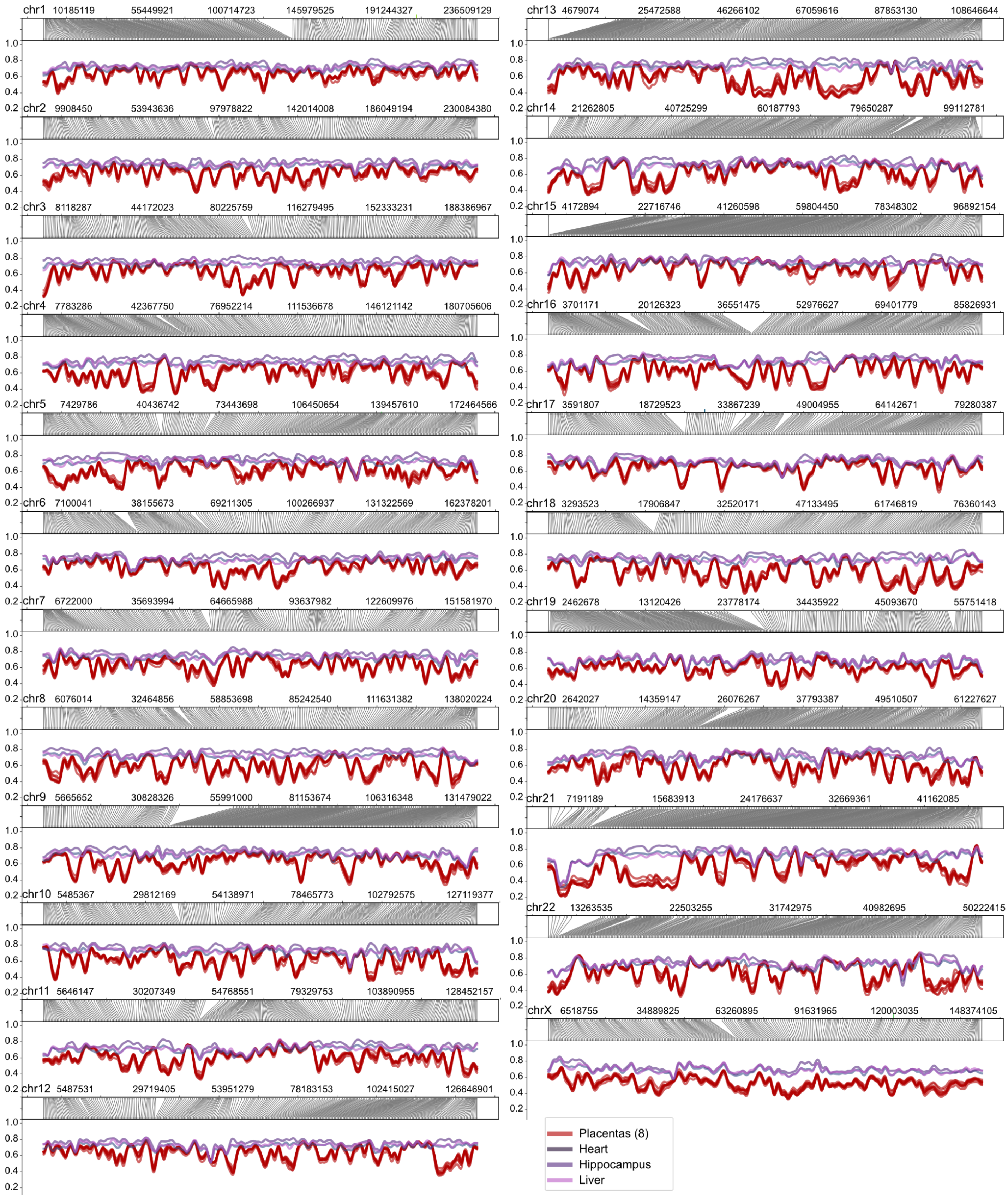


**Supplemental Figure 1:** Methylation profiles for placentas (red) as compared to other tissues (heart, hippocampus, liver). Each panel shows genome coordinate space translated to CpG coordinate space, chromosomes are not scaled by length.

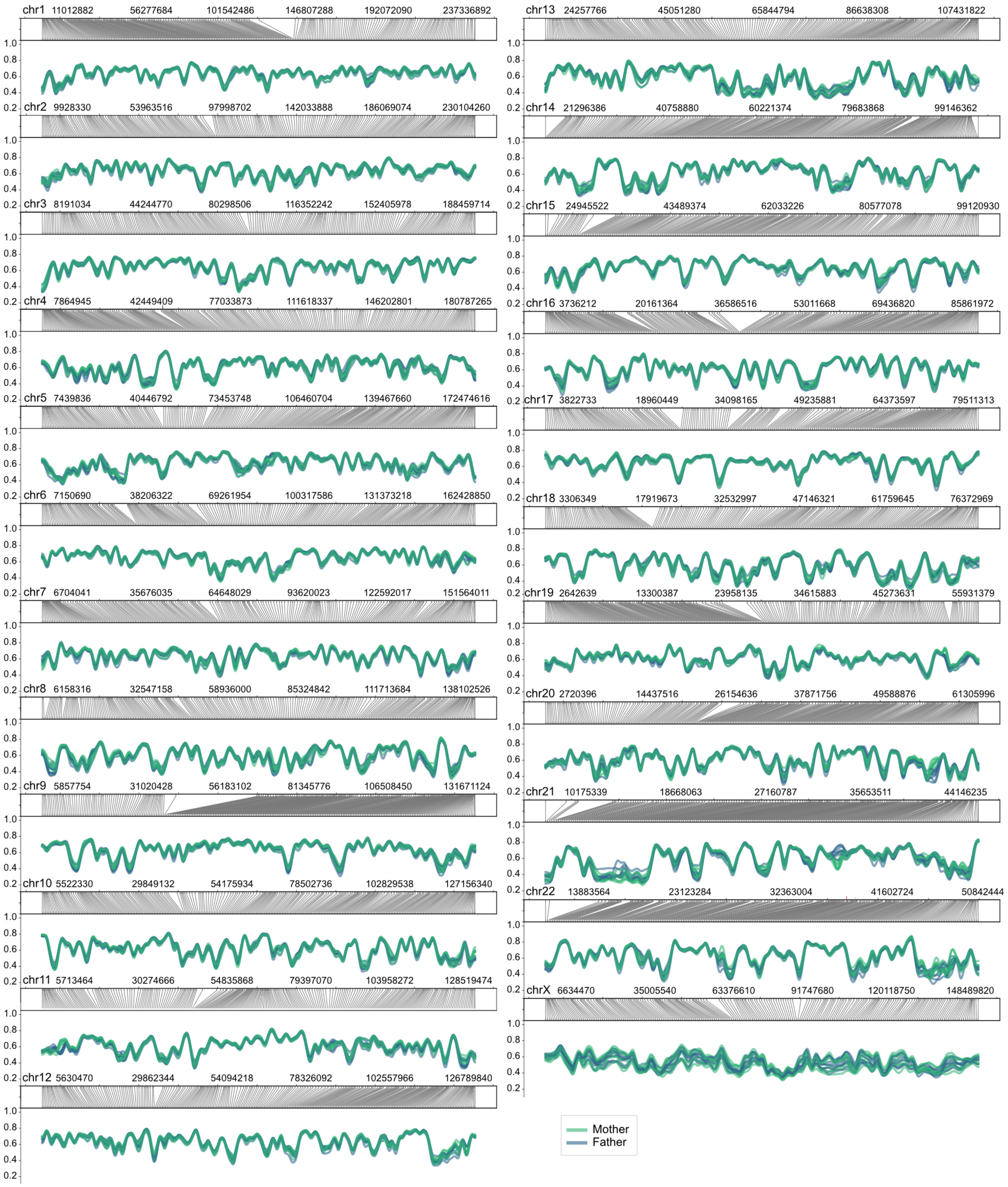


**Supplemental Figure 2:** Haplotype-resolved methylation by parent-of-origin. Maternal and paternal alleles are broadly similar with the exception of the X chromosome (see Supplemental Figure 3).

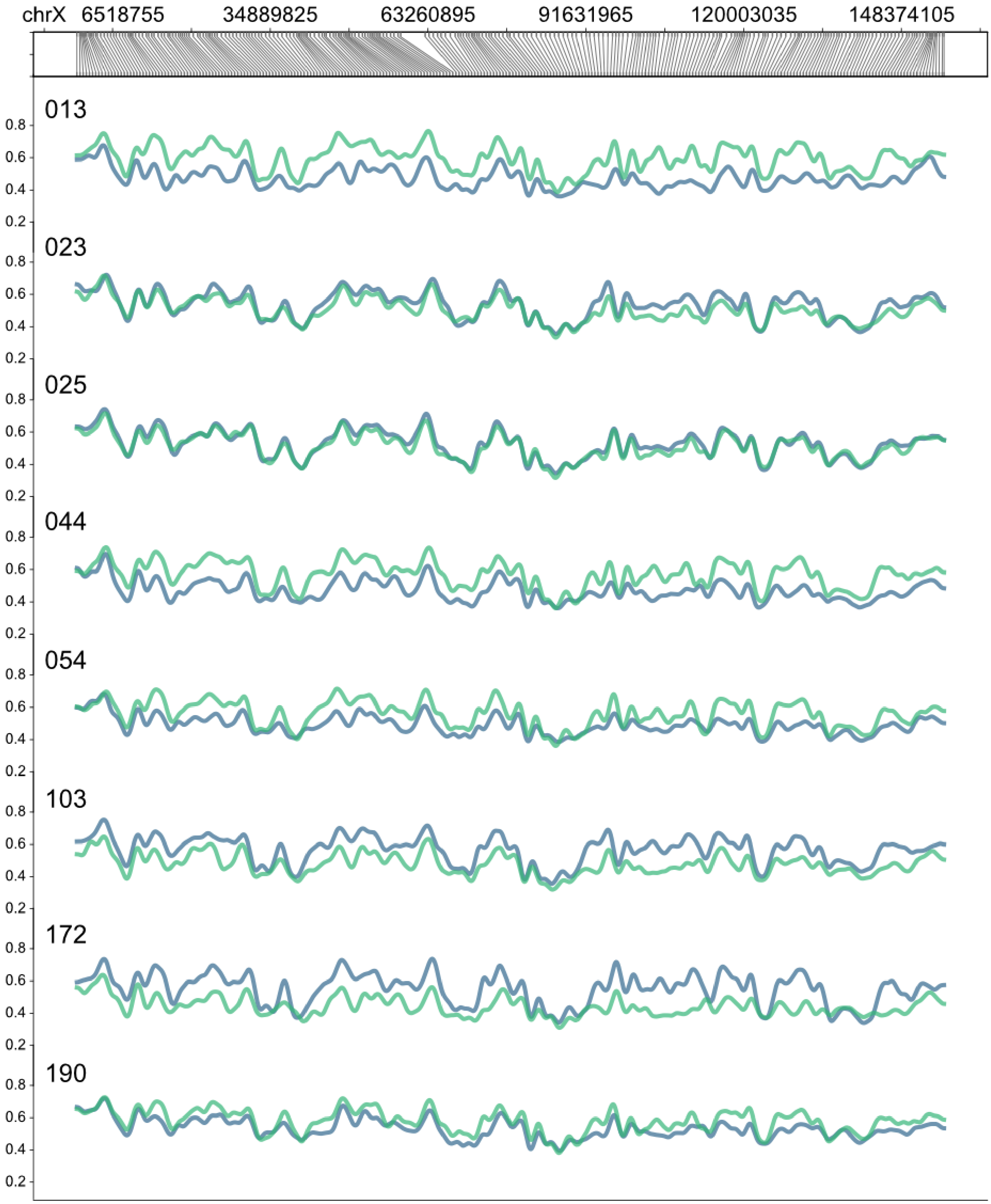


**Supplemental Figure 3:** Comparison of maternal (green) and paternal (blue) methylation profiles on chromosome X. In some samples the maternal chromosome is generally more methylated (013, 044, 045, 190), in some the paternal chromosome is more methylated (103, 172) and in others methylation levels are comparable (023, 025). As noted, this is likely the result of the clonal development of placental tissue.

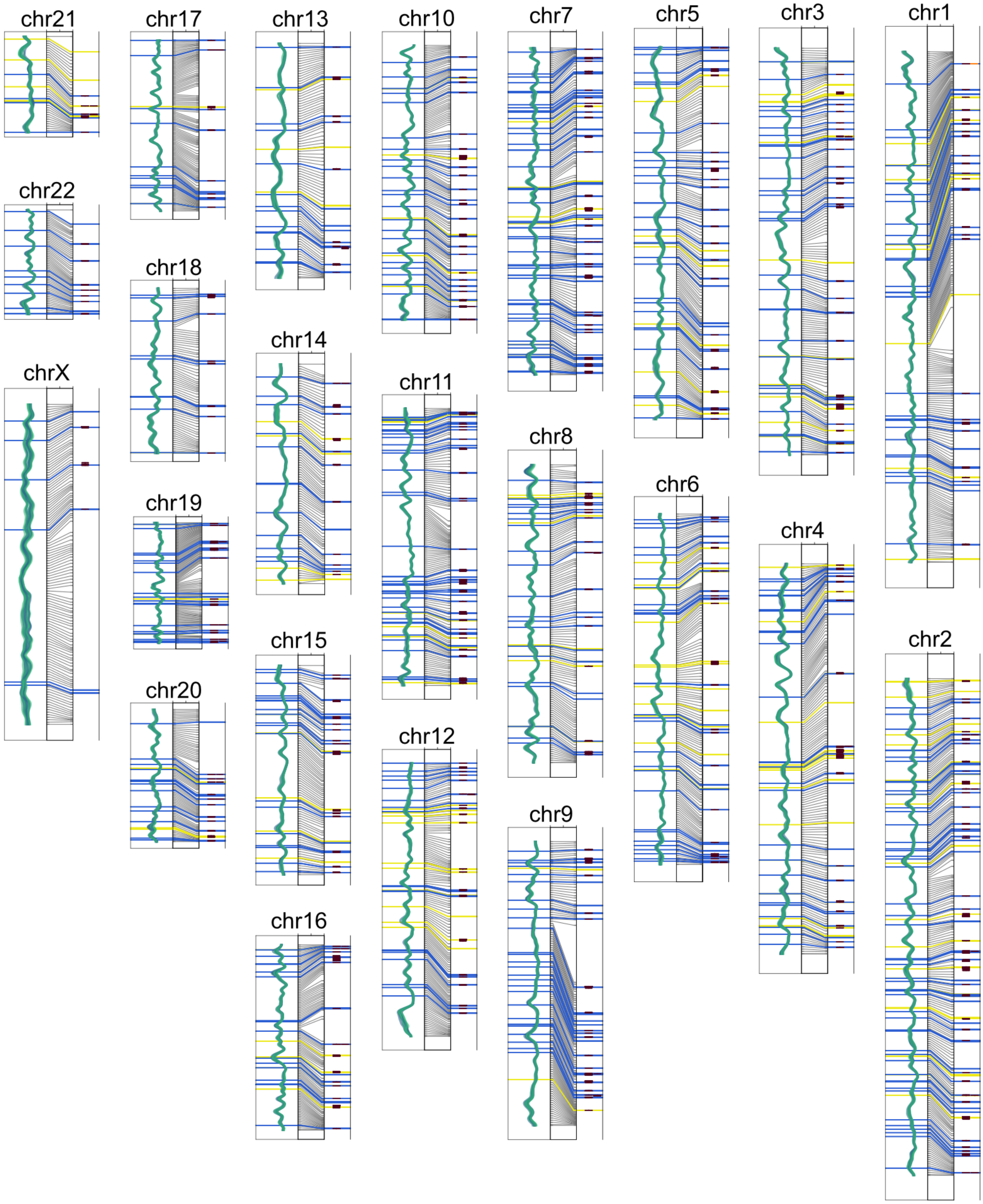


**Supplemental Figure 4:** Novel and previously described DMRs distributed across the genome. Novel DMRs are indicated as yellow highlights and those previously described are highlighted in blue. Associated protein-coding genes are included in the rightmost track and the parent-specific methylation profiles are included on the left track of each chromosome plot with the translation between genome and CpG coordinate space in between.

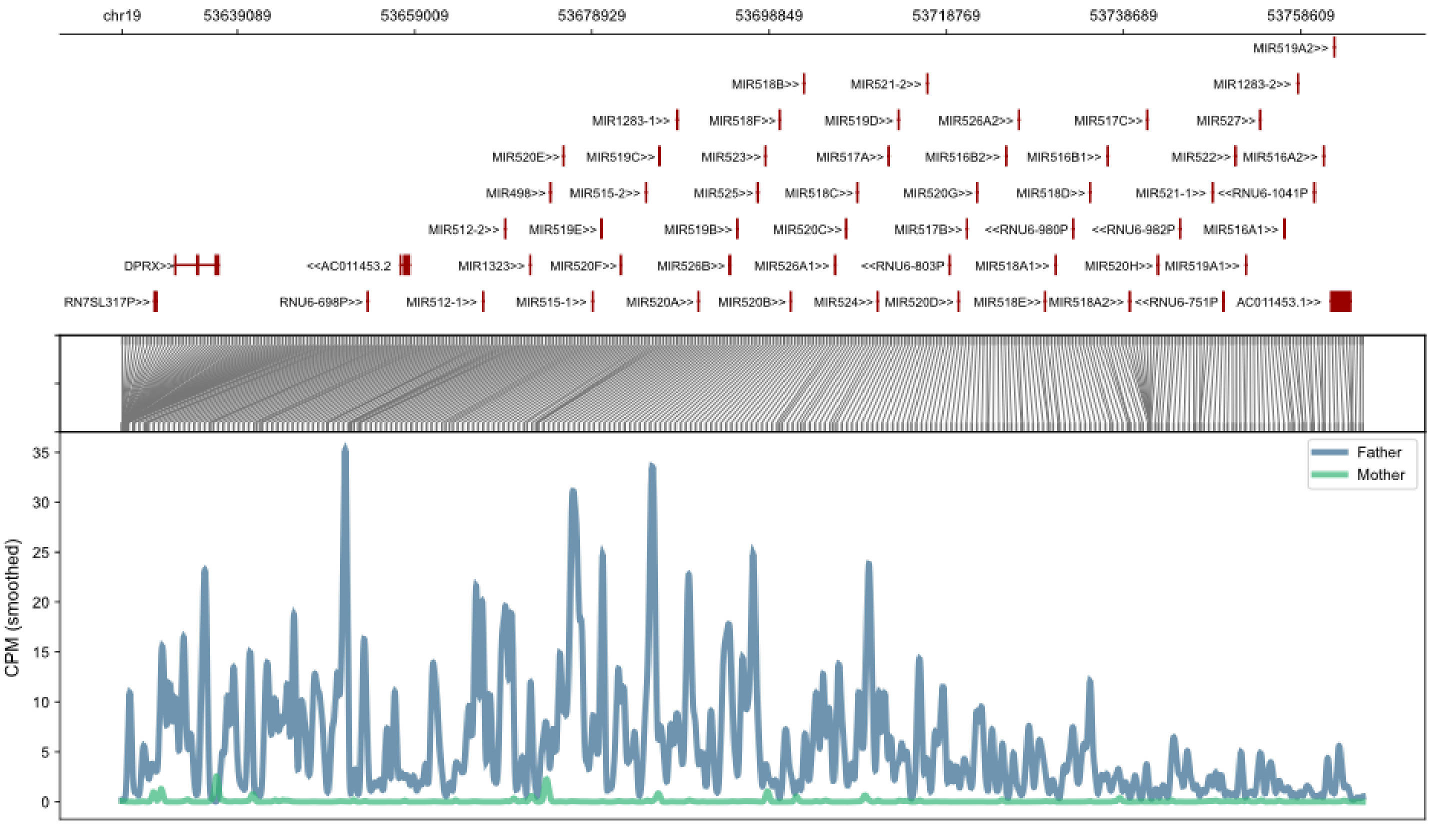


**Supplemental Figure 5:** Allele-specific expression profiles of the C19MC region containing many paternally-imprinted miRNAs. The top panel shows coordinates and gene locations, followed by a translation from genome coordinate space into expressed coordinate space.

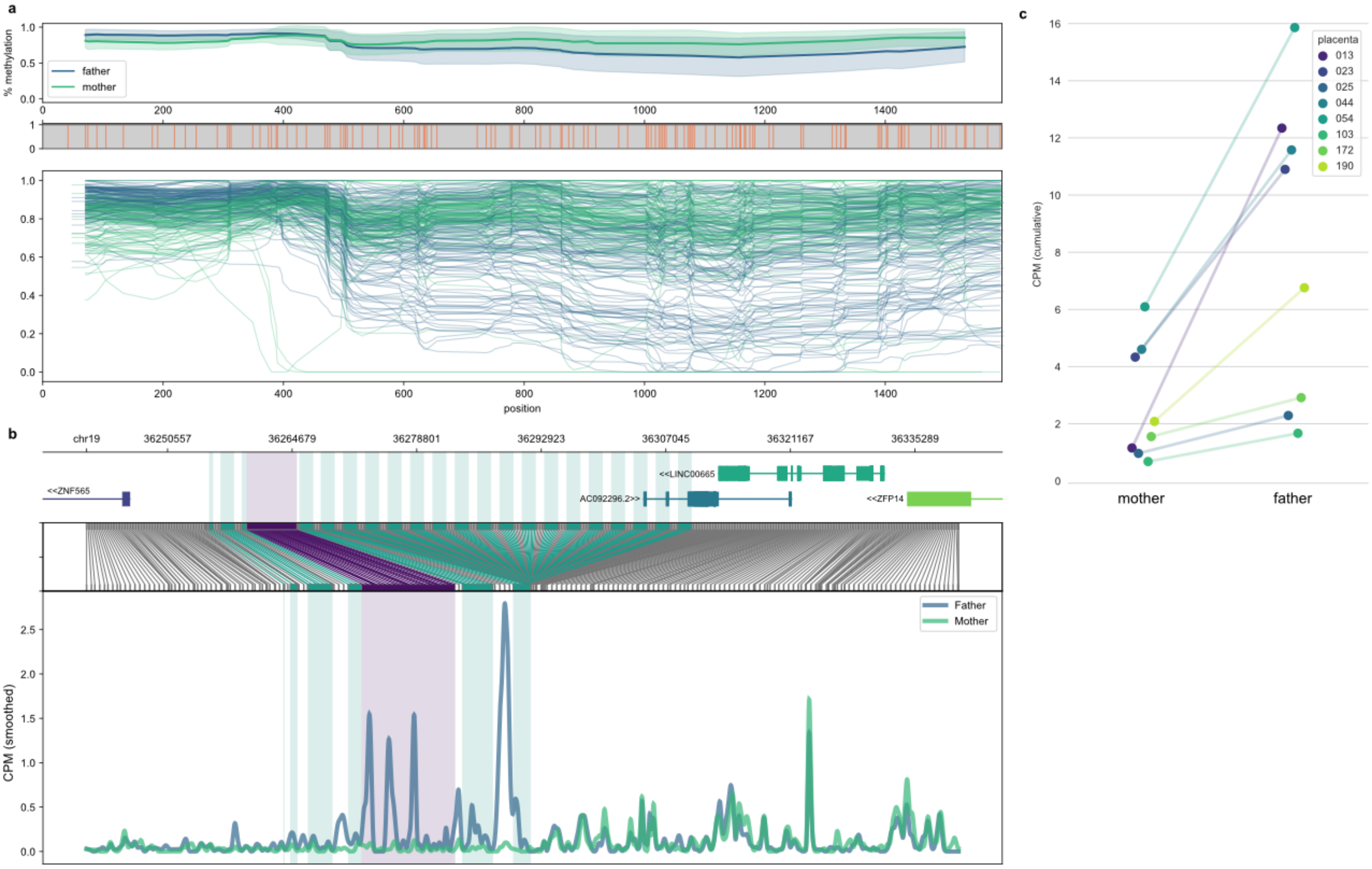


**Supplemental Figure 6:** Panel (a) shows the parent-specific methylation profile of each SST1 repeat in the left chr19 SST1 array, with the averaged methylation plot on top. The allele-specific expression detectable at the SST1 array is shown in panel (b), with the caveat that allele-specific expression is only apparent over informative variants which are few in number across this highly repetitive locus. The purple highlight indicates the position of the HERVH element that seems to be driving allele-specific expression across the locus.

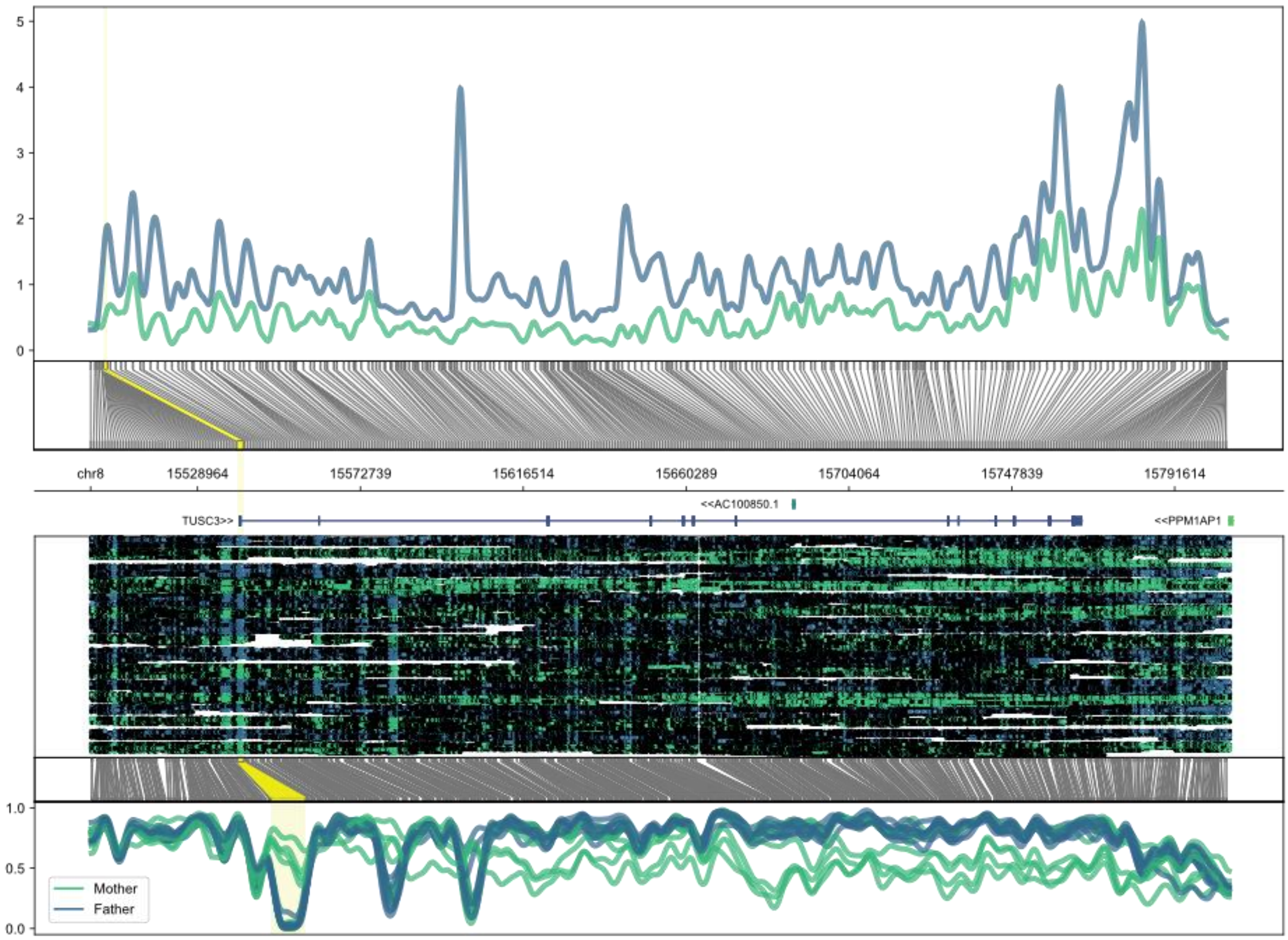


**Supplemental Figure 7:** Polymorphic allele-specific methylation (bottom panel) and paternal-biased expression (top panel) of TUSC3. The DMR location is highlighted (yellow) across the panels.

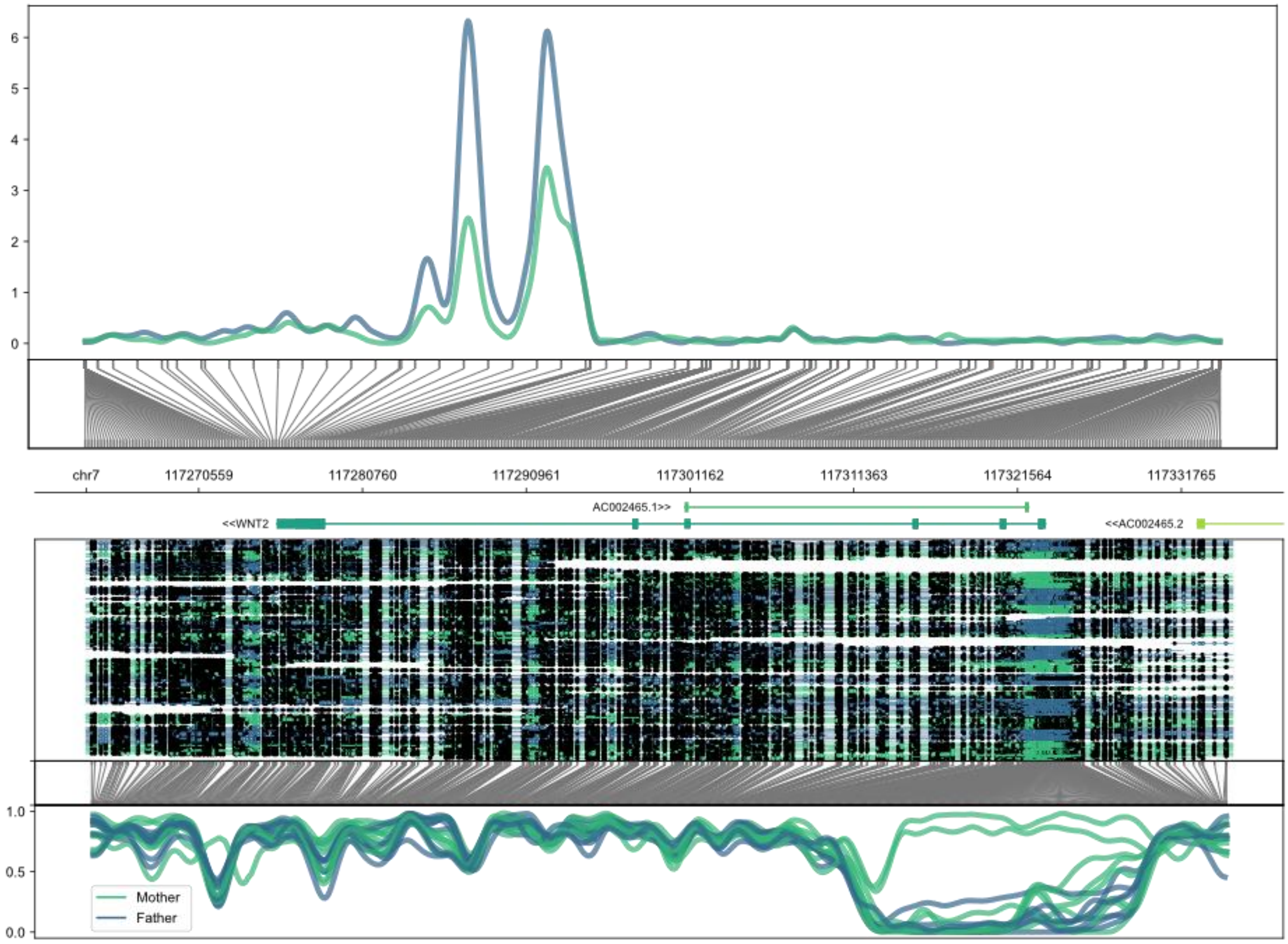


**Supplemental Figure 8:** Polymorphic allele-specific methylation (bottom panel) and paternal-biased expression (top panel) of WNT2.
