## Supplemental Data for "An allele-resolved nanopore-guided tour of the human placental methylome"

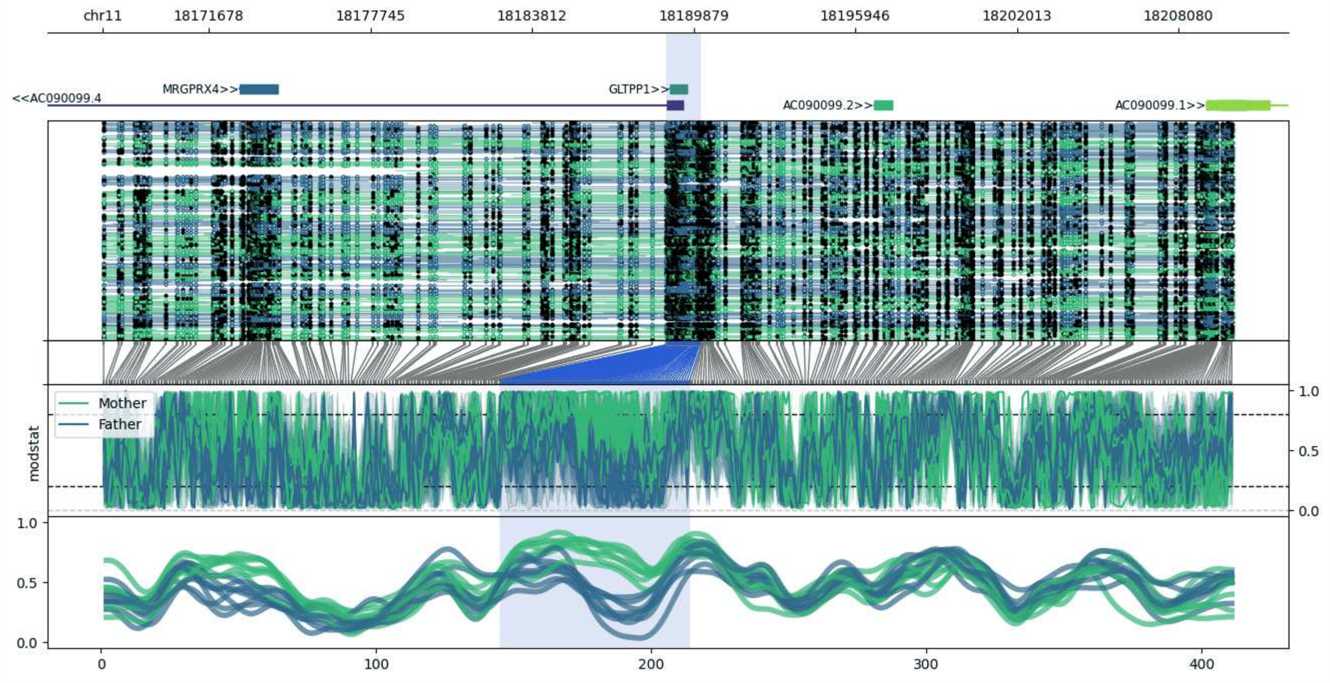


**chr11:18167703-18210172:** Region with 1 DMR containing genes: AC090099.1, AC090099.2, AC090099.4, GLTPP1, MRGPRX4.

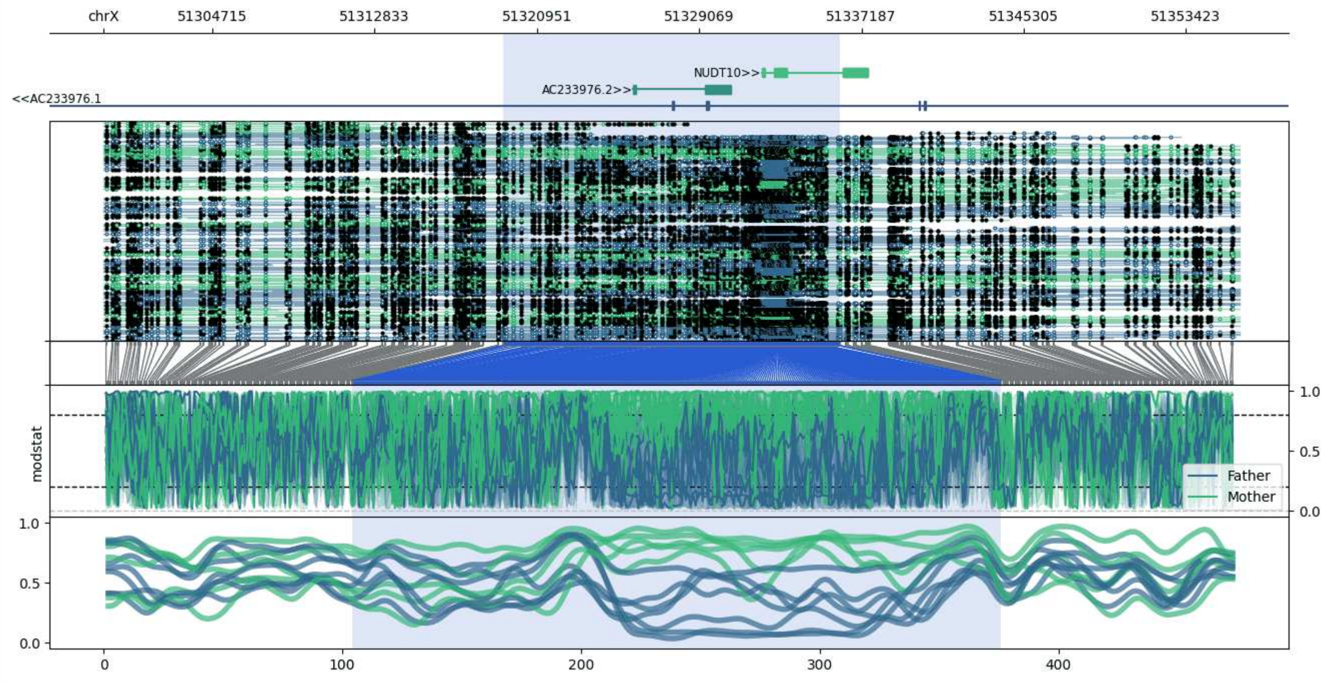


**chrX:51299280-51356109:** Region with 1 DMR containing genes: AC233976.1, AC233976.2, NUDT10.

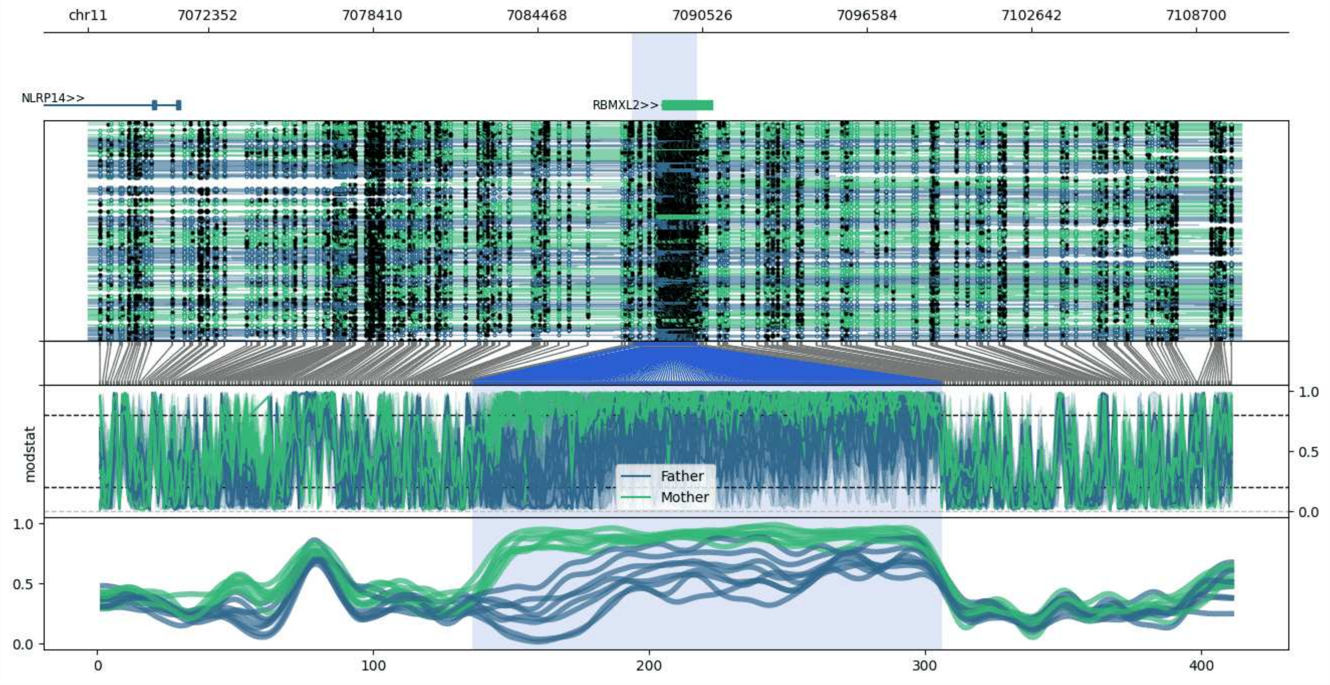


**chr11:7067926-7110335:** Region with 1 DMR containing genes: NLRP14, RBMXL2.

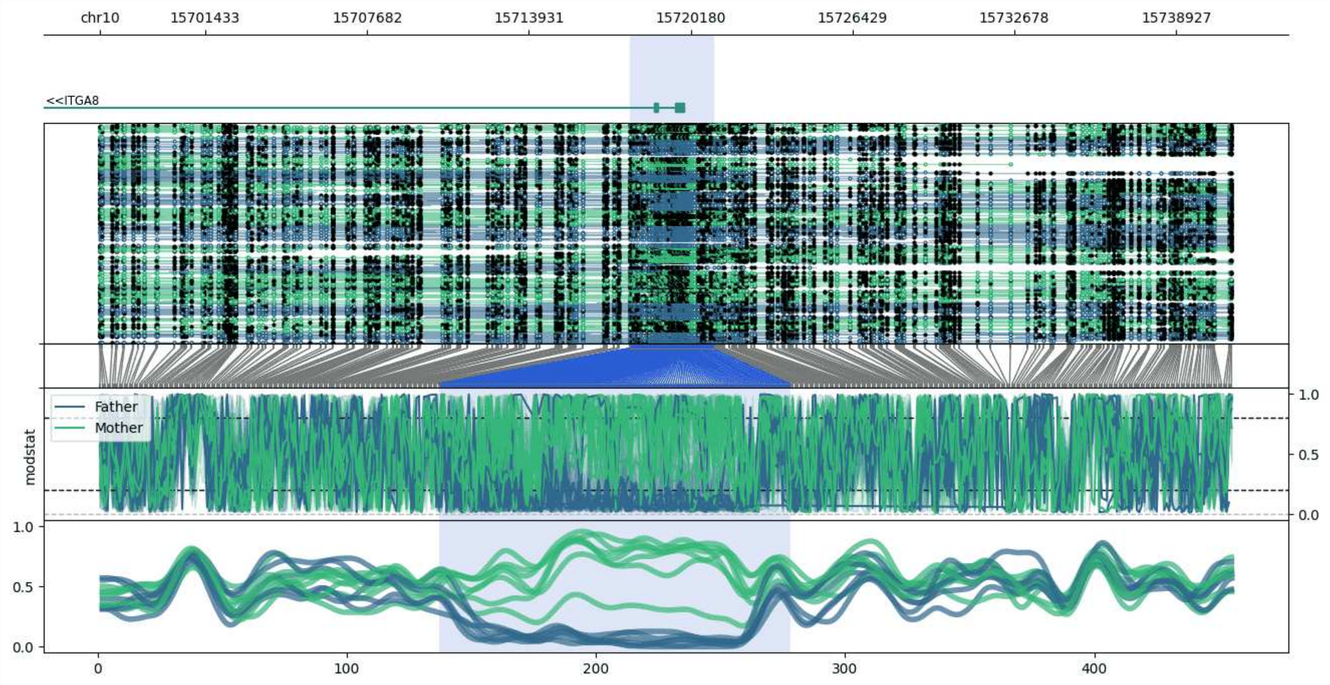


**chr10:15697357-15741106:** Region with 1 DMR containing genes: ITGA8.

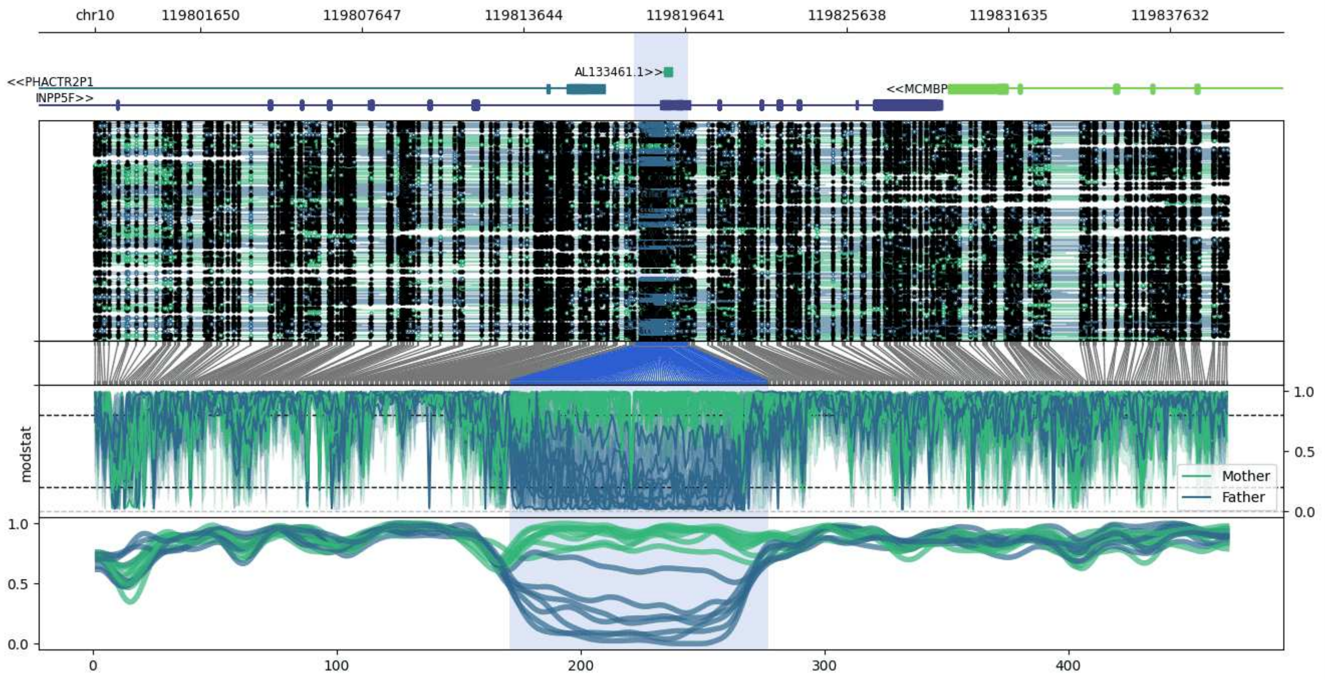


**chr10:119797751-119839735:** Region with 1 DMR containing genes: AL133461.1, INPP5F, MCMBP, PHACTR2P1.

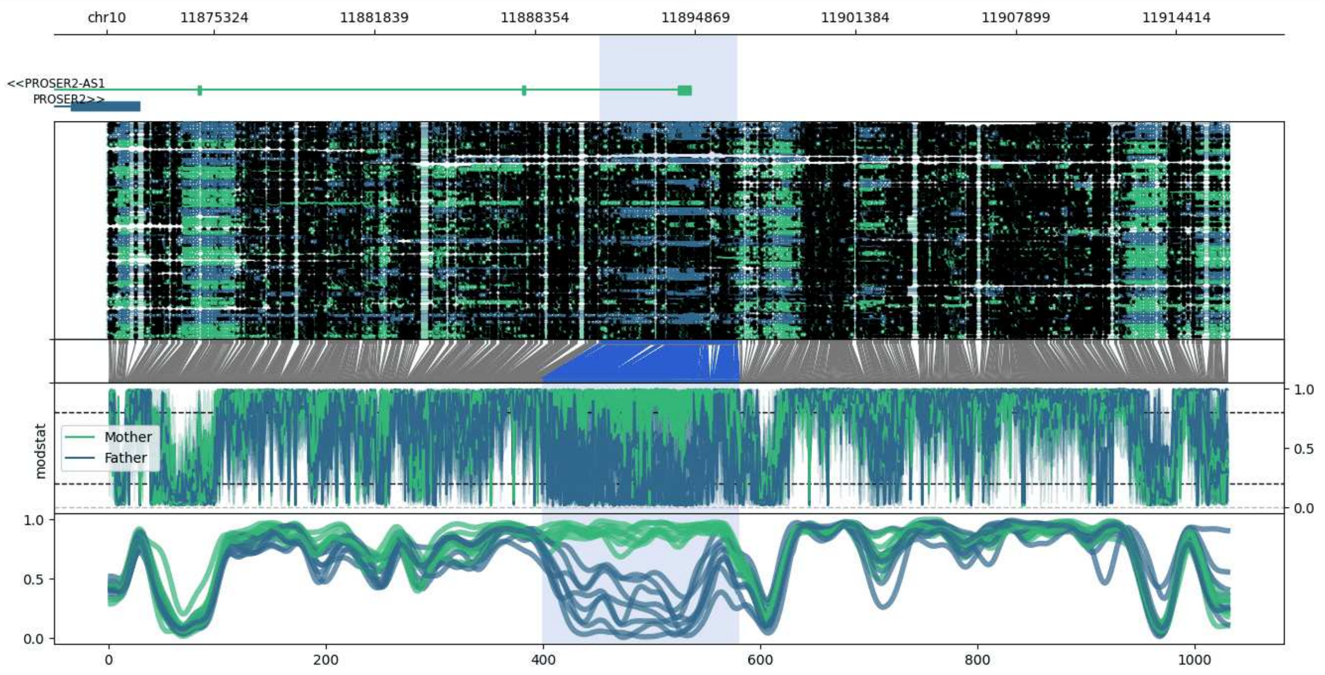


**chr10:11870965-11916572:** Region with 1 DMR containing genes: PROSER2, PROSER2-AS1, RNU6-1095P, UPF2.

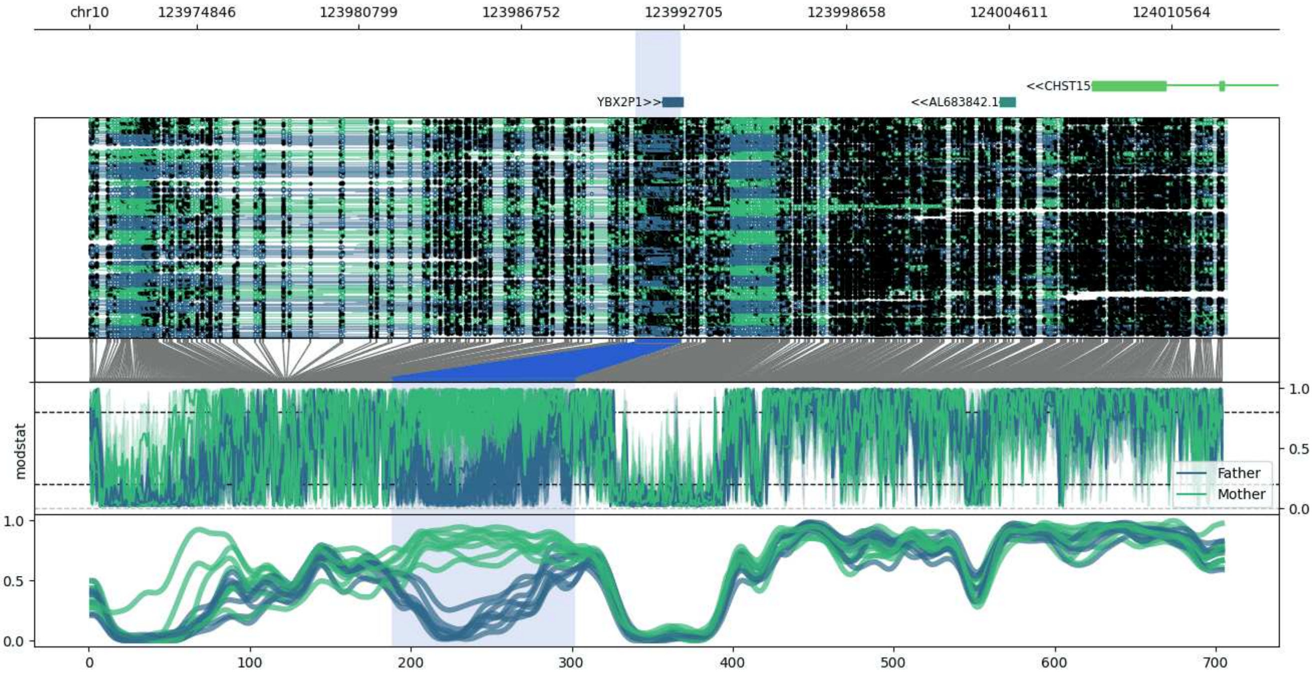


**chr10:123970931-124012606:** Region with 1 DMR containing genes: AL683842.1, CHST15, YBX2P1.

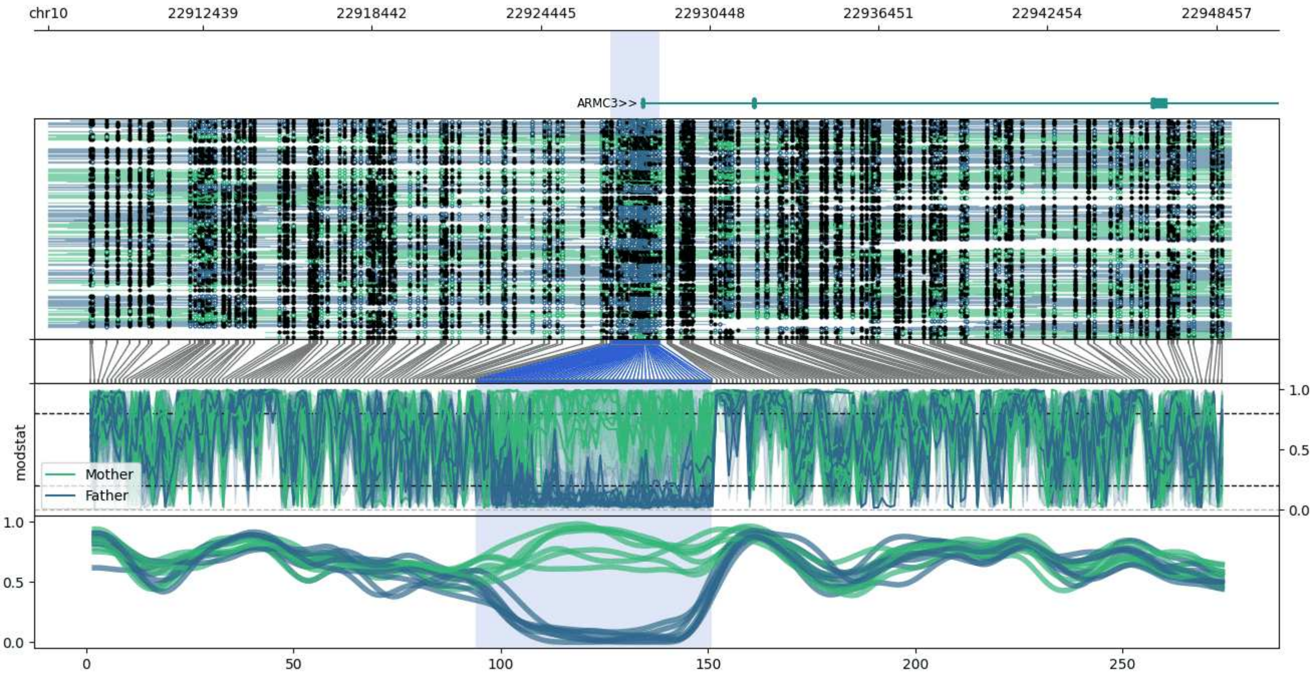


**chr10:22906932-22948954:** Region with 1 DMR containing genes: ARMC3.

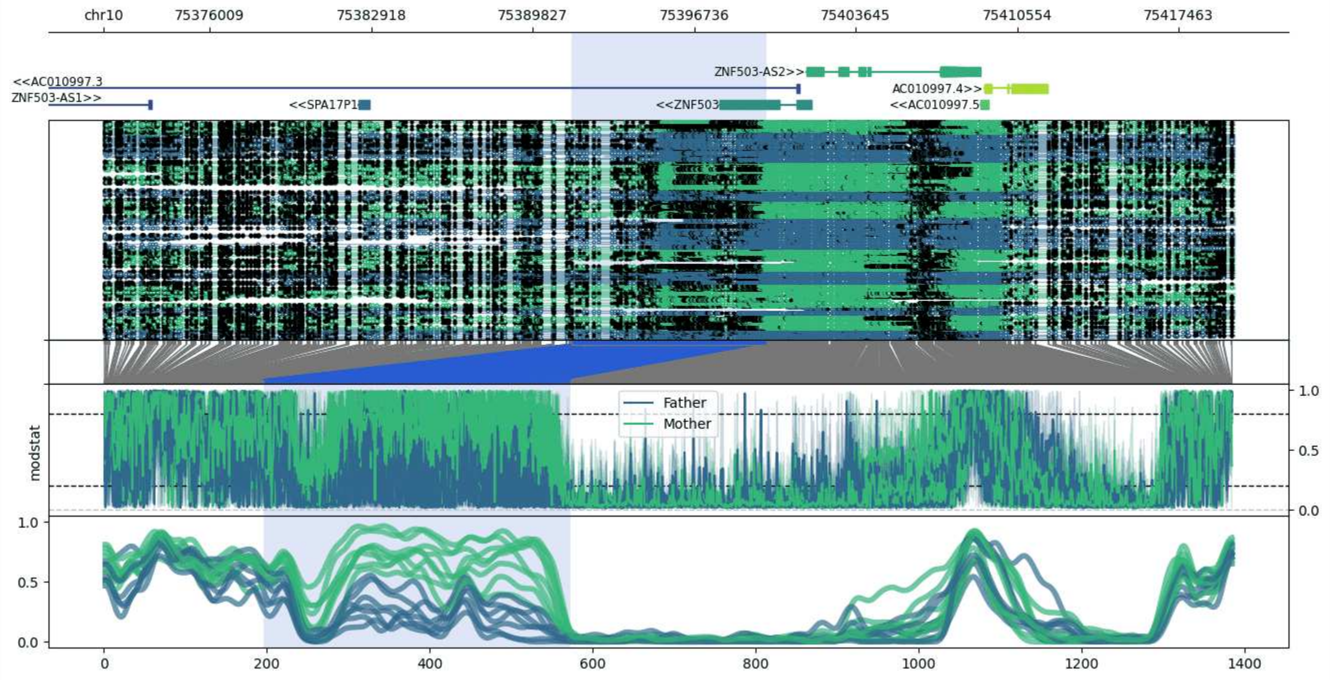


**chr10:75371462-75419830:** Region with 1 DMR containing genes: AC010997.3, AC010997.4, AC010997.5, SPA17P1, ZNF503, ZNF503-AS1, ZNF503-AS2.

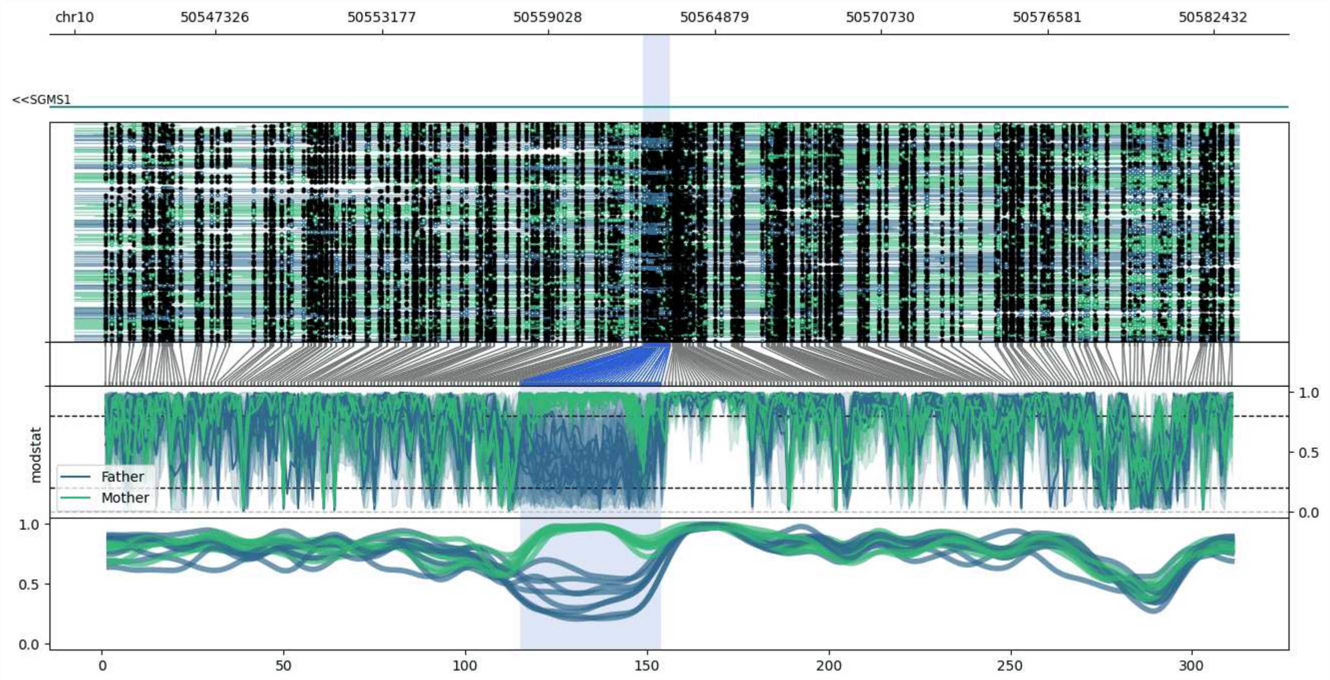


**chr10:50542356-50583313:** Region with 1 DMR containing genes: RNU7-107P, SGMS1.

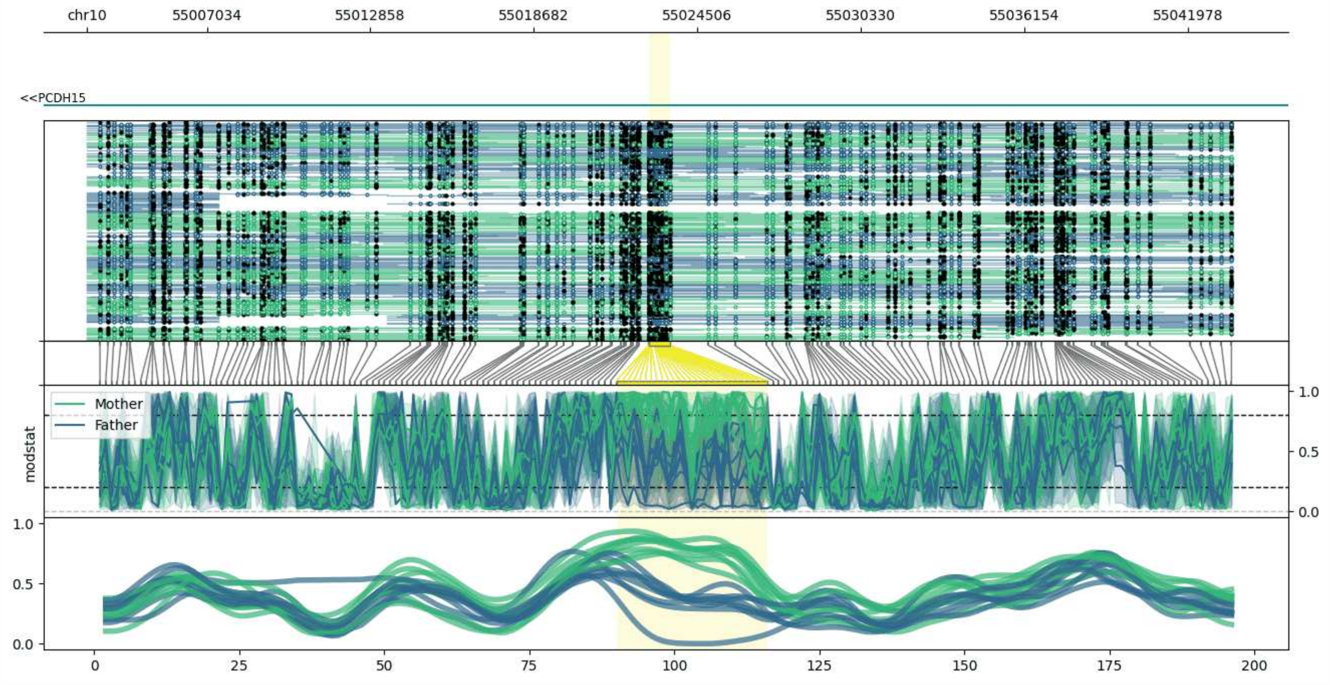


**chr10:55002770-55043541:** Region with 1 DMR containing genes: PCDH15.

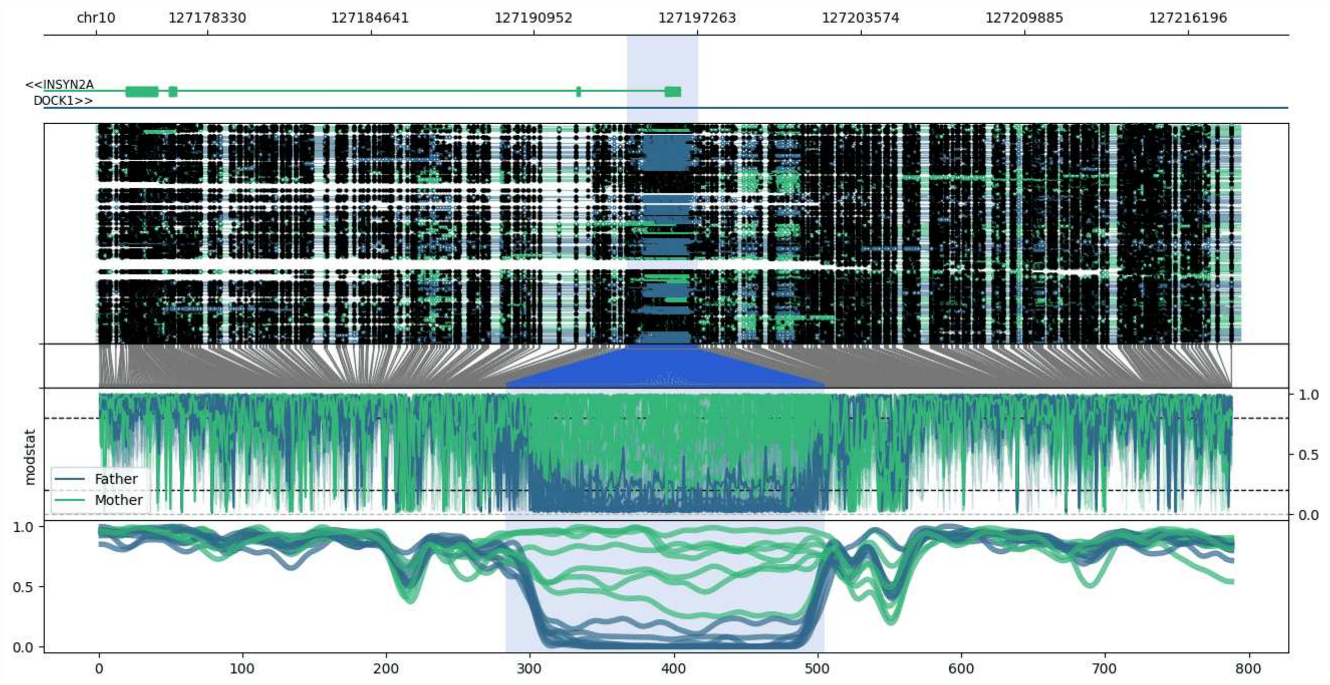


**chr10:127174031-127218210:** Region with 1 DMR containing genes: DOCK1, INSYN2A.

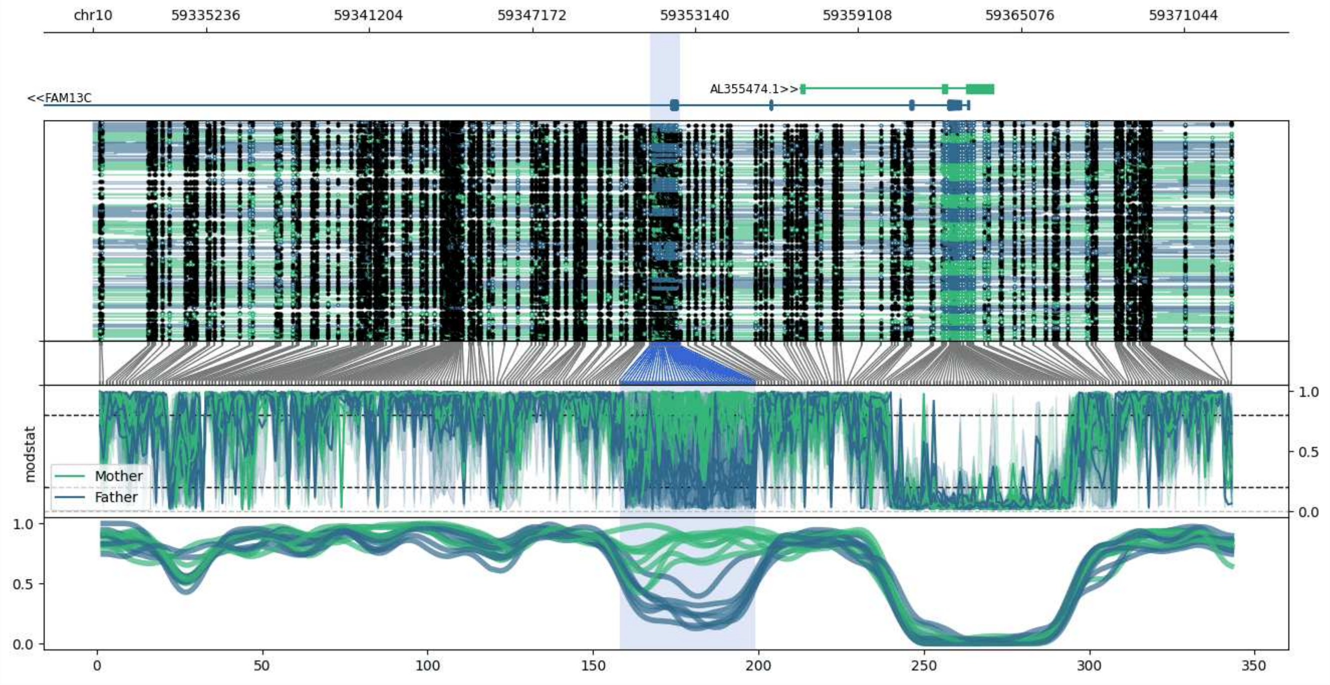


**chr10:59331086-59372868:** Region with 1 DMR containing genes: AL355474.1, FAM13C.

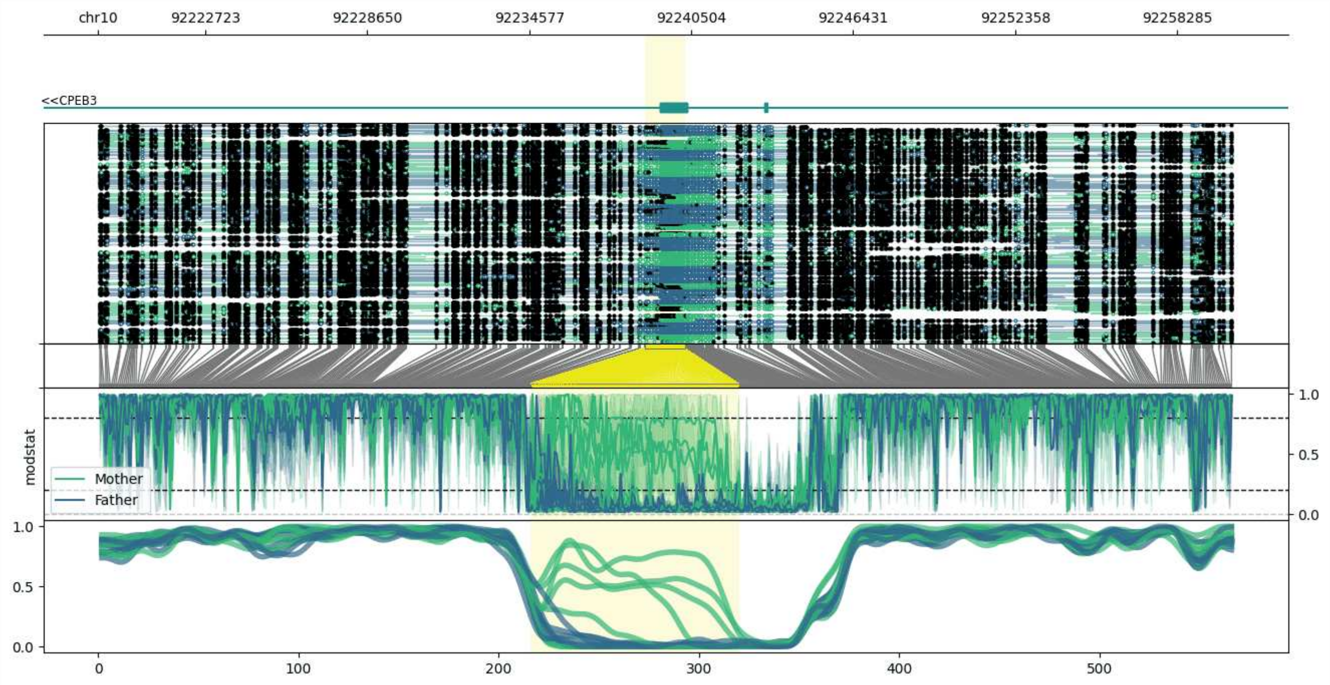


**chr10:92218799-92260288:** Region with 1 DMR containing genes: CPEB3, NHP2P1.

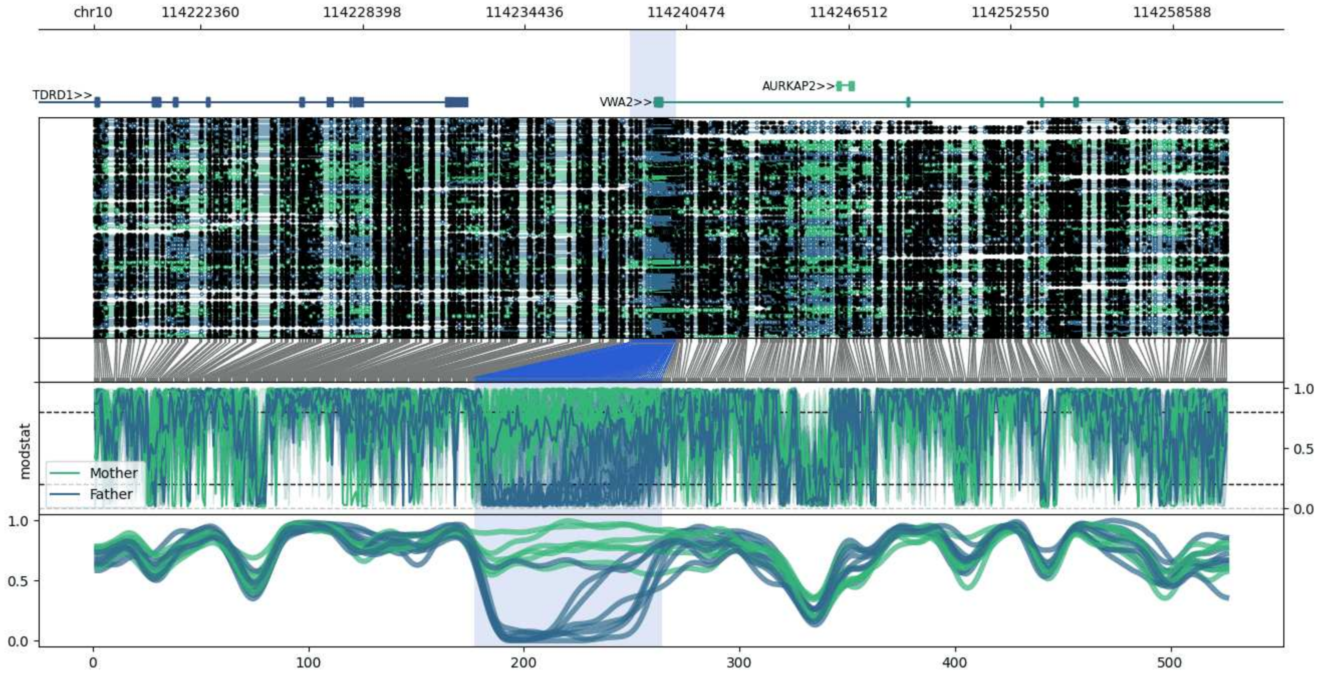


**chr10:114218372-114260638:** Region with 1 DMR containing genes: AURKAP2, TDRD1, VWA2.

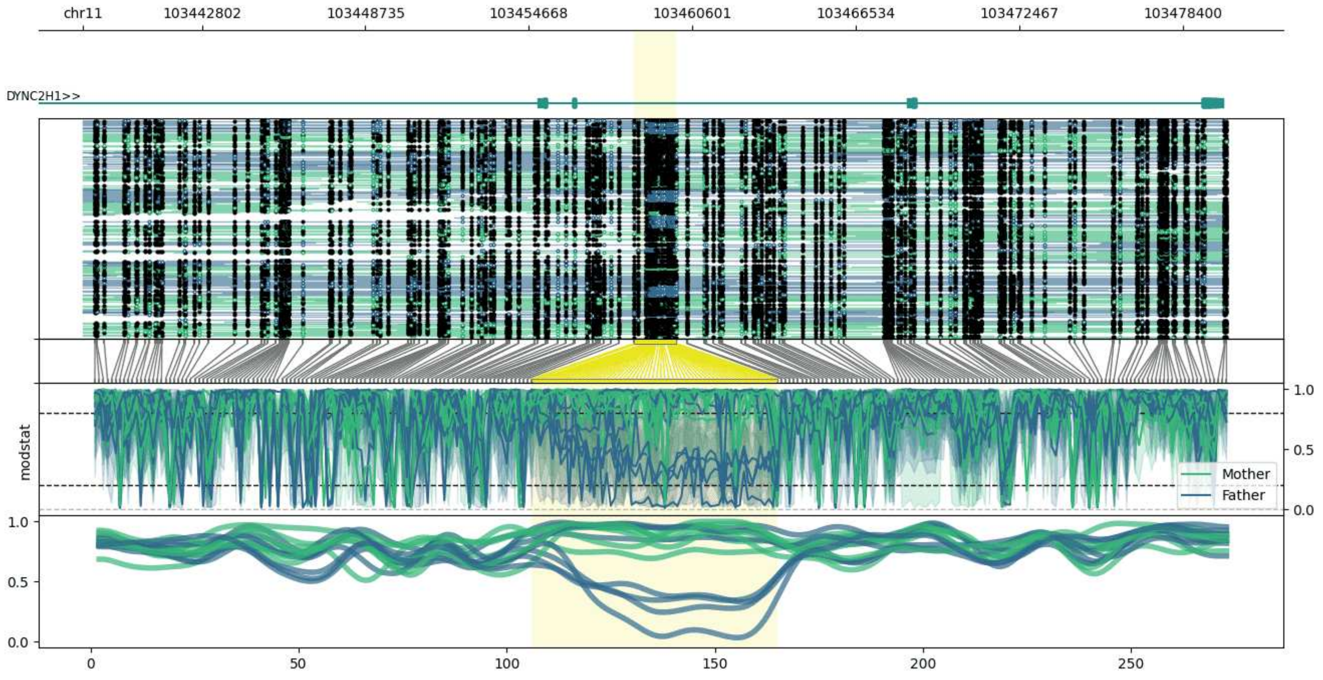


**chr11:103438492-103480023:** Region with 1 DMR containing genes: DYNC2H1.

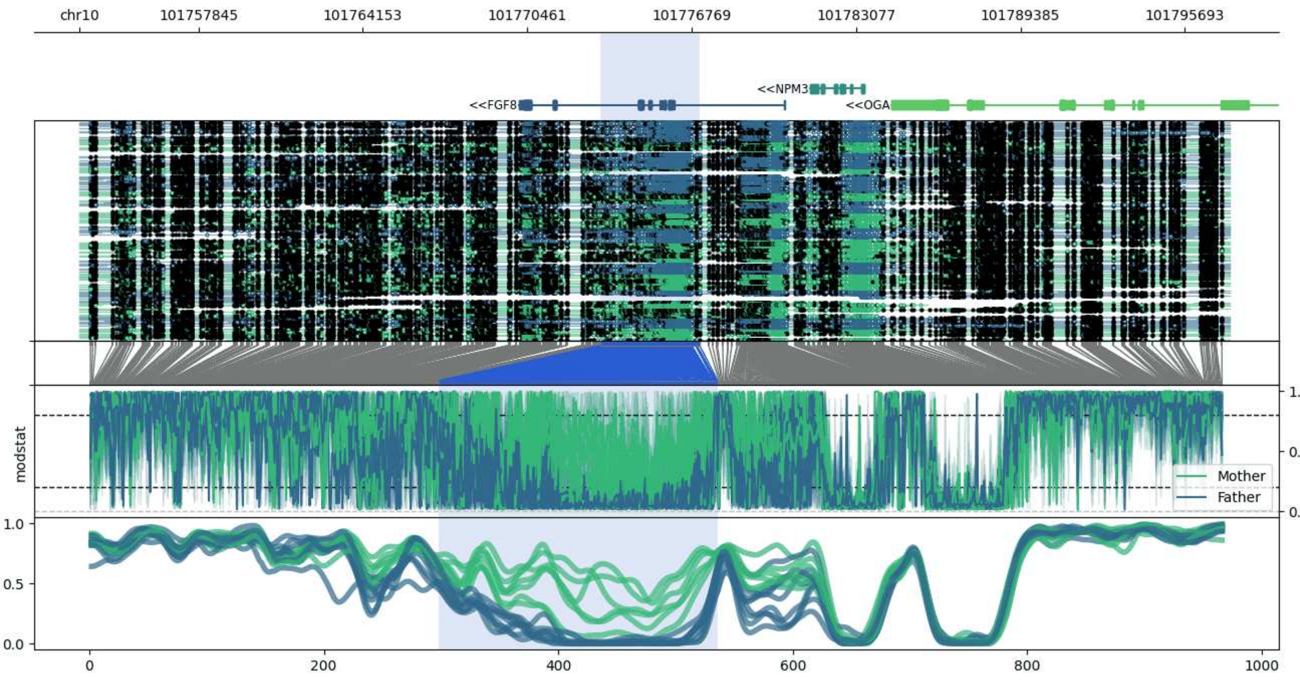


**chr10:101753270-101797428:** Region with 1 DMR containing genes: FGF8, NPM3, OGA.

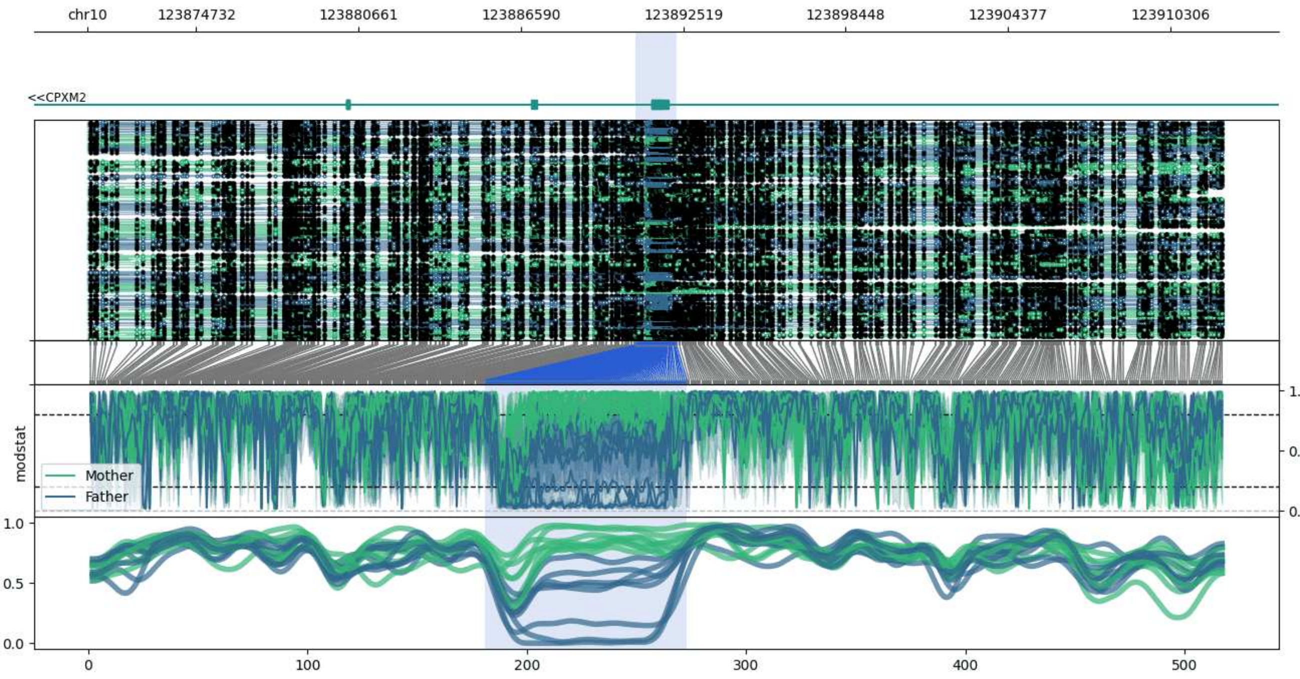


**chr10:123870768-123912273:** Region with 1 DMR containing genes: AC068058.1, CPXM2.

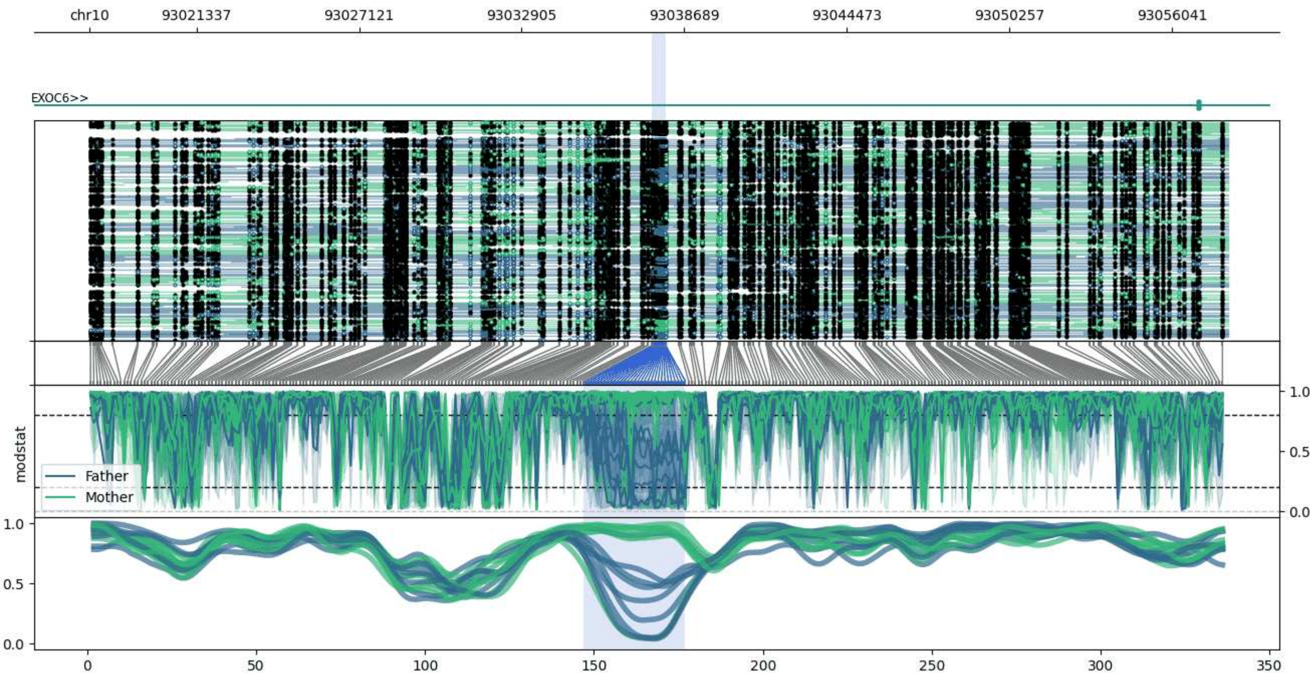


**chr10:93017542-93058034:** Region with 1 DMR containing genes: AL358613.1, AL358613.3, CYP26C1, EXOC6.

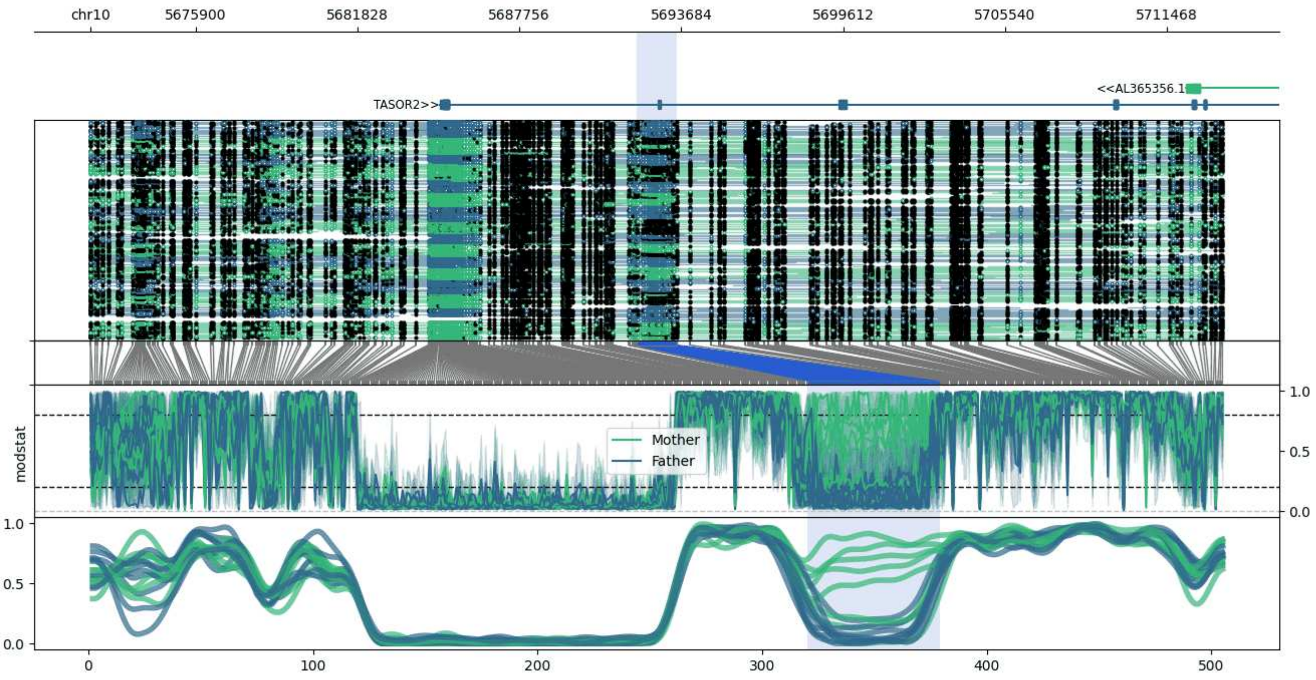


**chr10:5672034-5713534:** Region with 1 DMR containing genes: AL365356.1, ASB13, TASOR2.

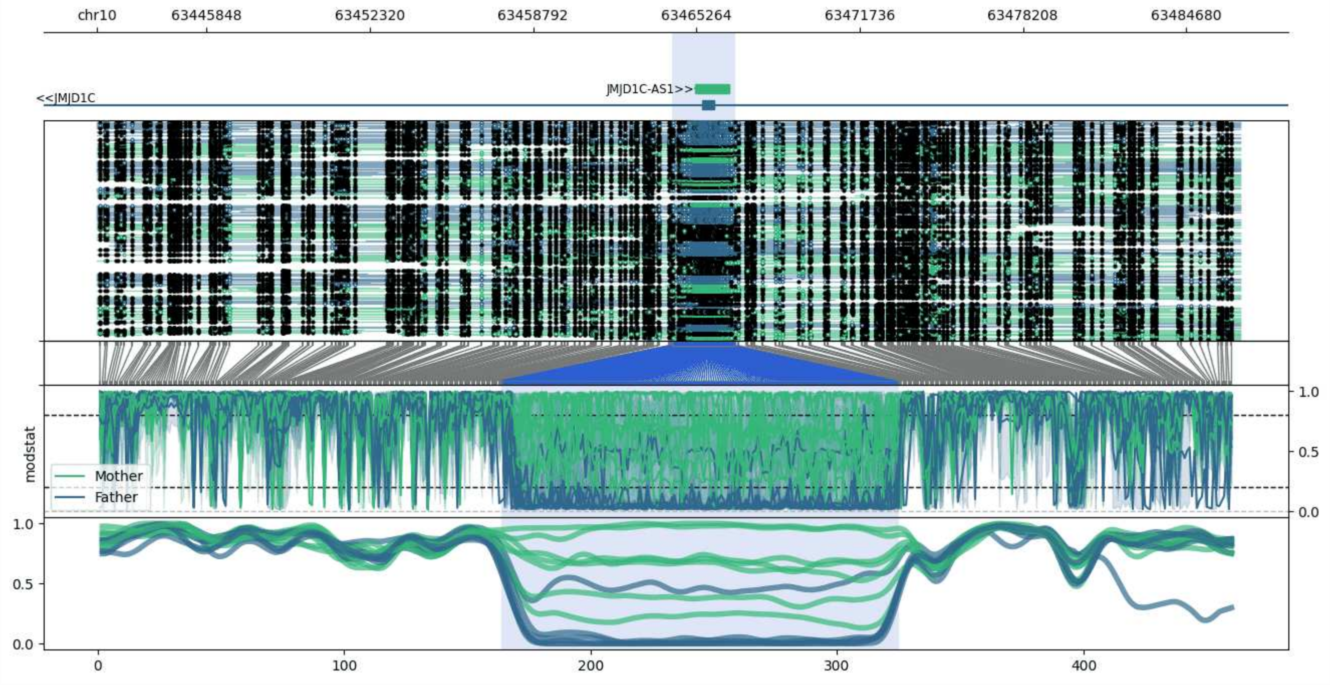


**chr10:63441511-63486815:** Region with 1 DMR containing genes: AC022022.1, JMJD1C, JMJD1C-AS1.

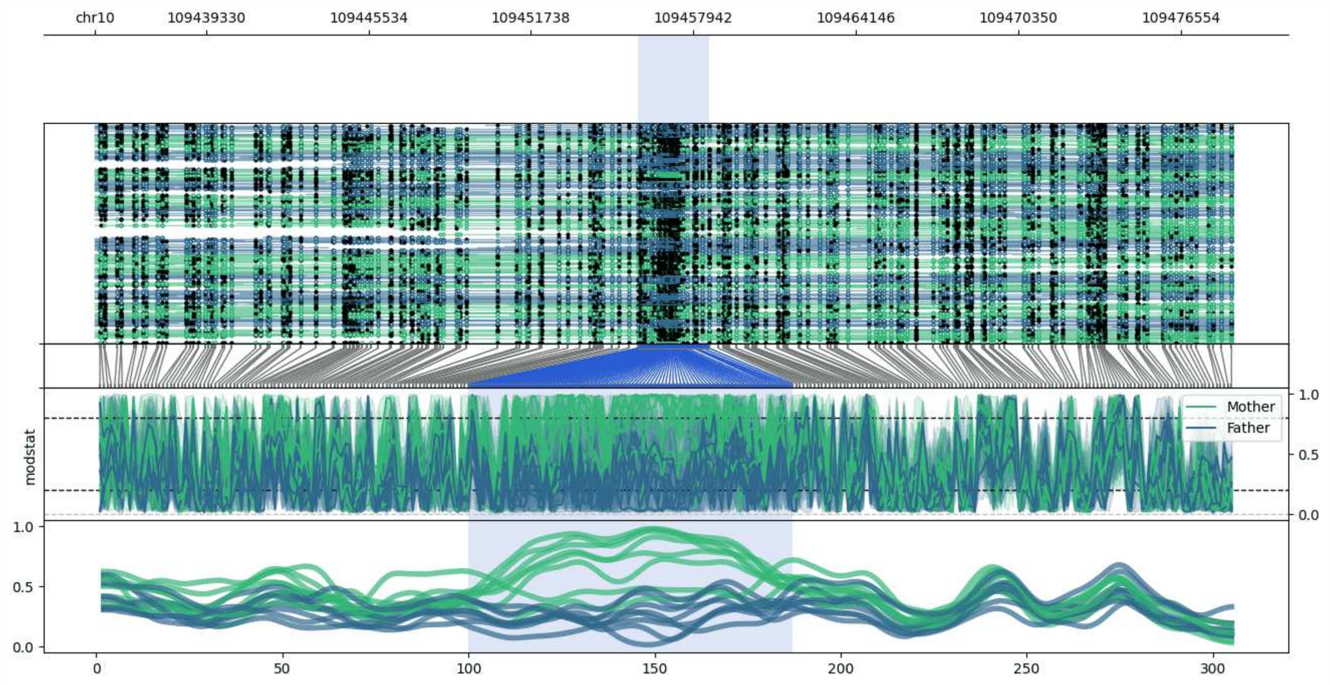


**chr10:109435101-109478533:** Region with 1 DMR containing no genes.

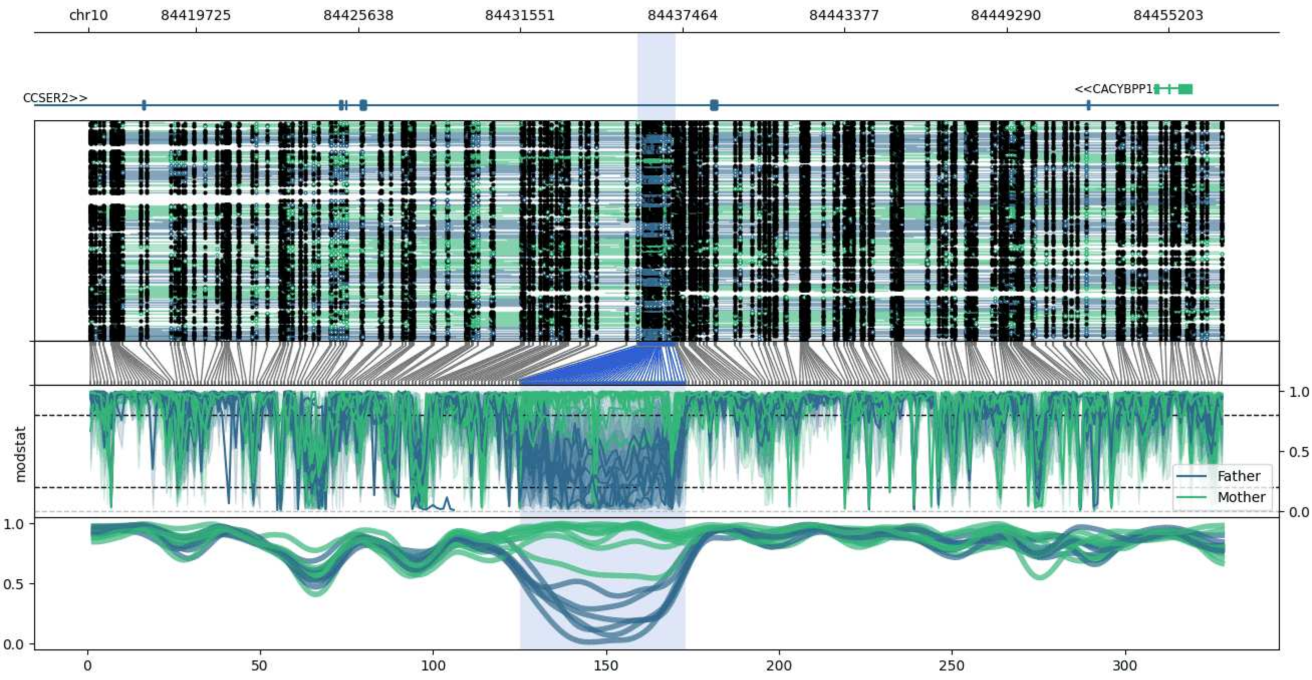


**chr10:84415810-84457204:** Region with 1 DMR containing genes: CACYBPP1, CCSER2.

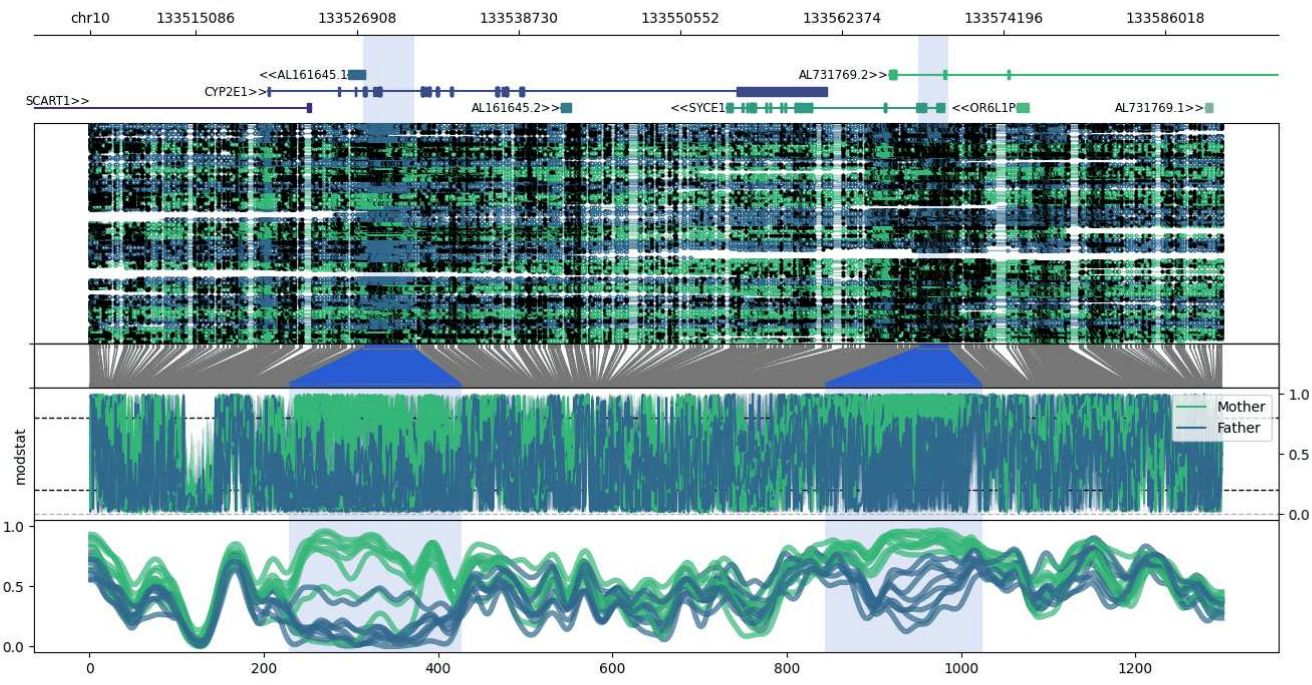


**chr10:133507388-133590146:** Region with 2 DMRs containing genes: AL161645.1, AL161645.2, AL731769.1, AL731769.2, CYP2E1, OR6L1P, SCART1, SYCE1.

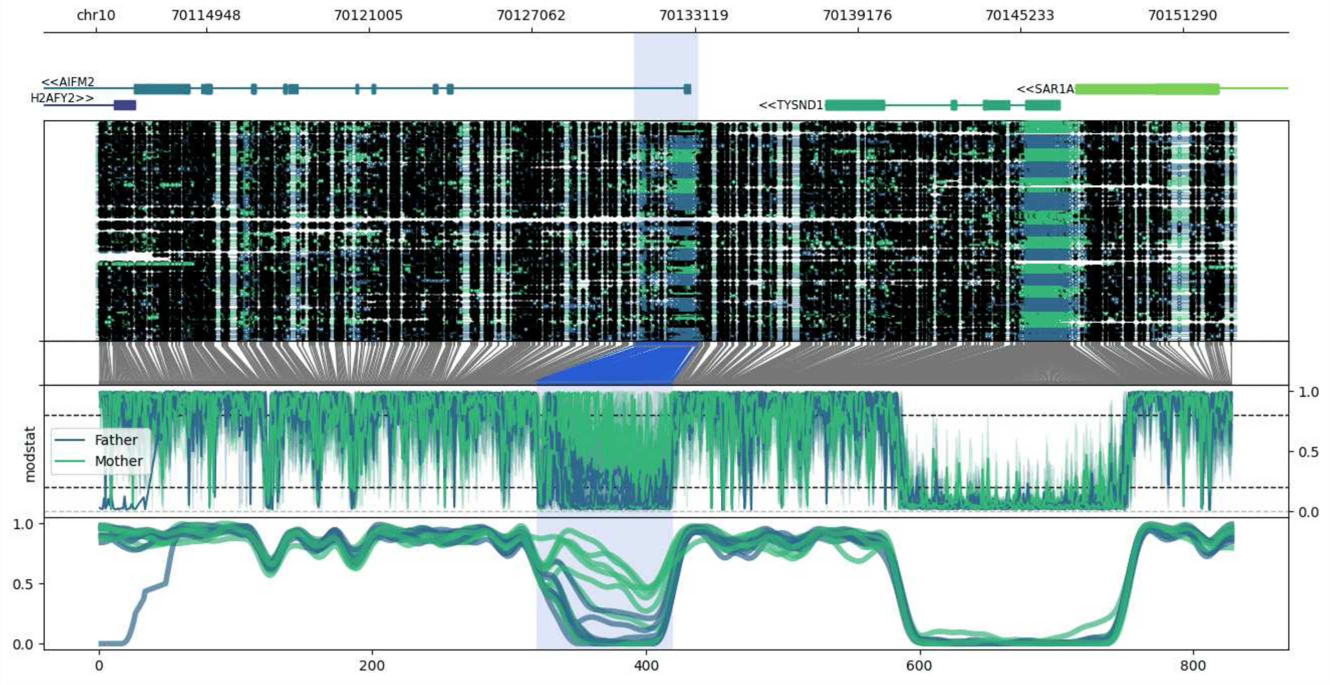


**chr10:70110855-70153260:** Region with 1 DMR containing genes: AIFM2, H2AFY2, SAR1A, TYSND1.

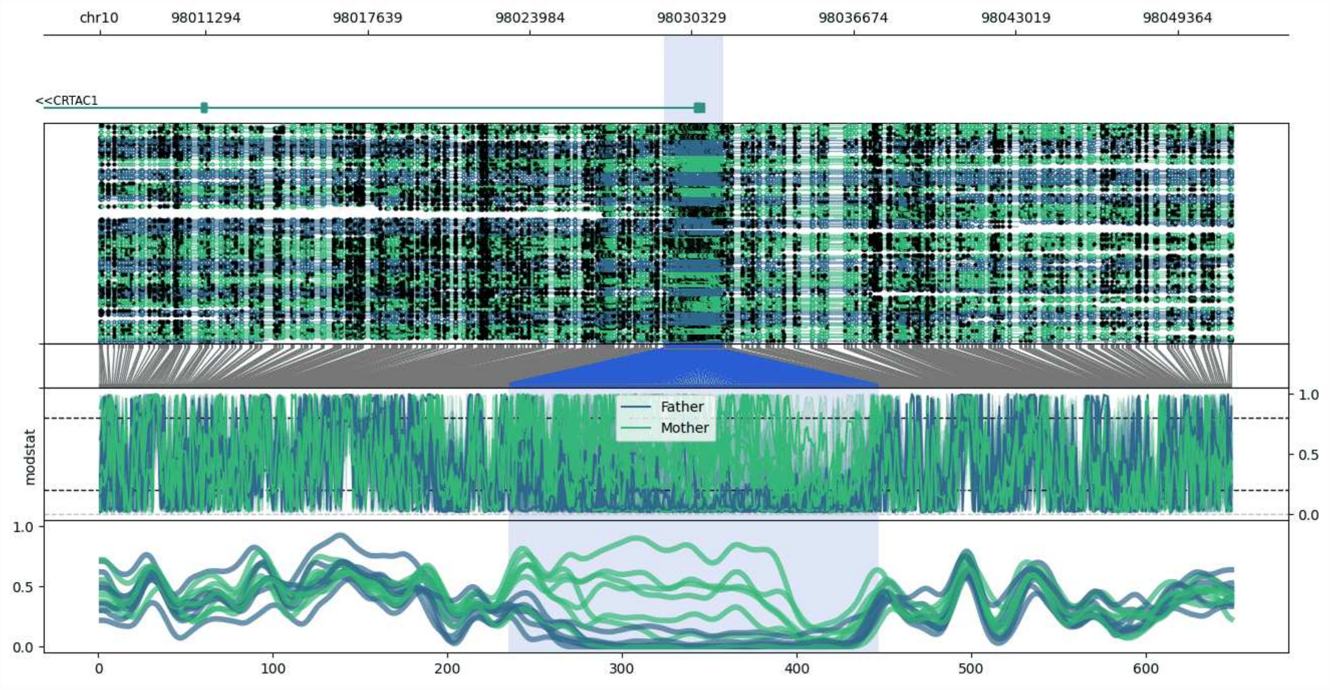


**chr10:98007145-98051561:** Region with 1 DMR containing genes: CRTAC1.

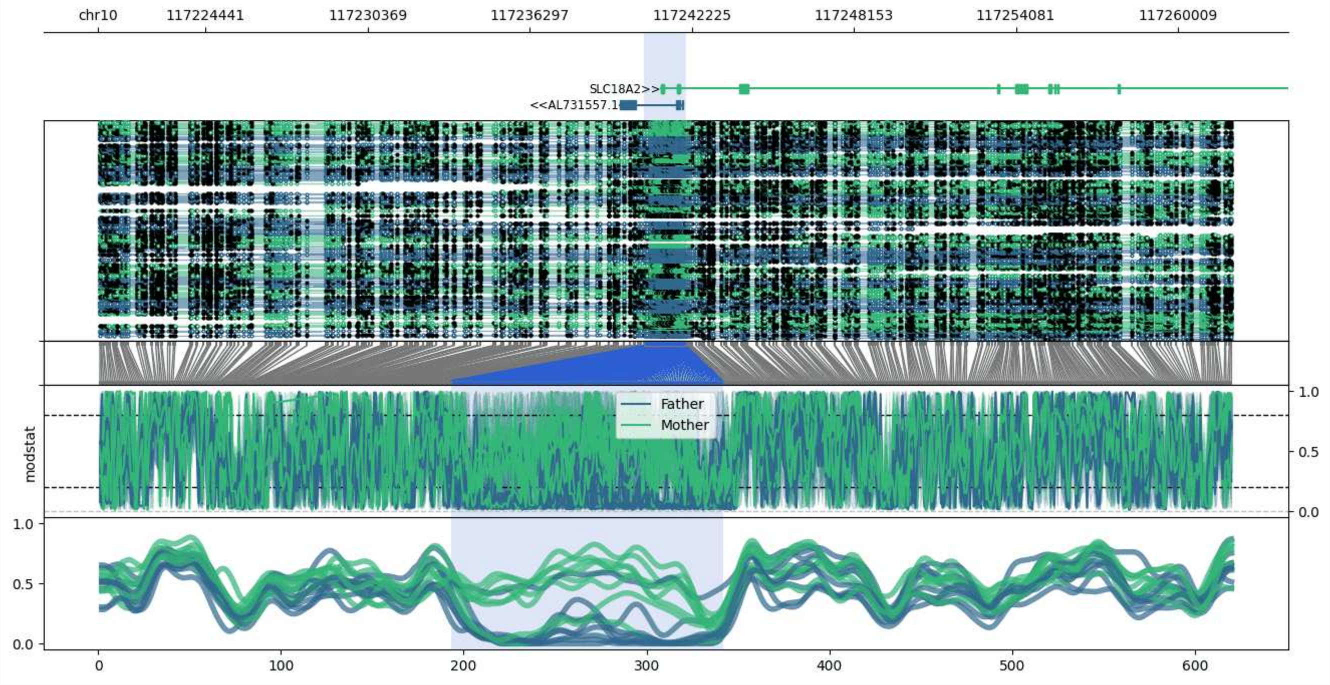


**chr10:117220493-117261989:** Region with 1 DMR containing genes: AL391988.1, AL731557.1, SLC18A2.

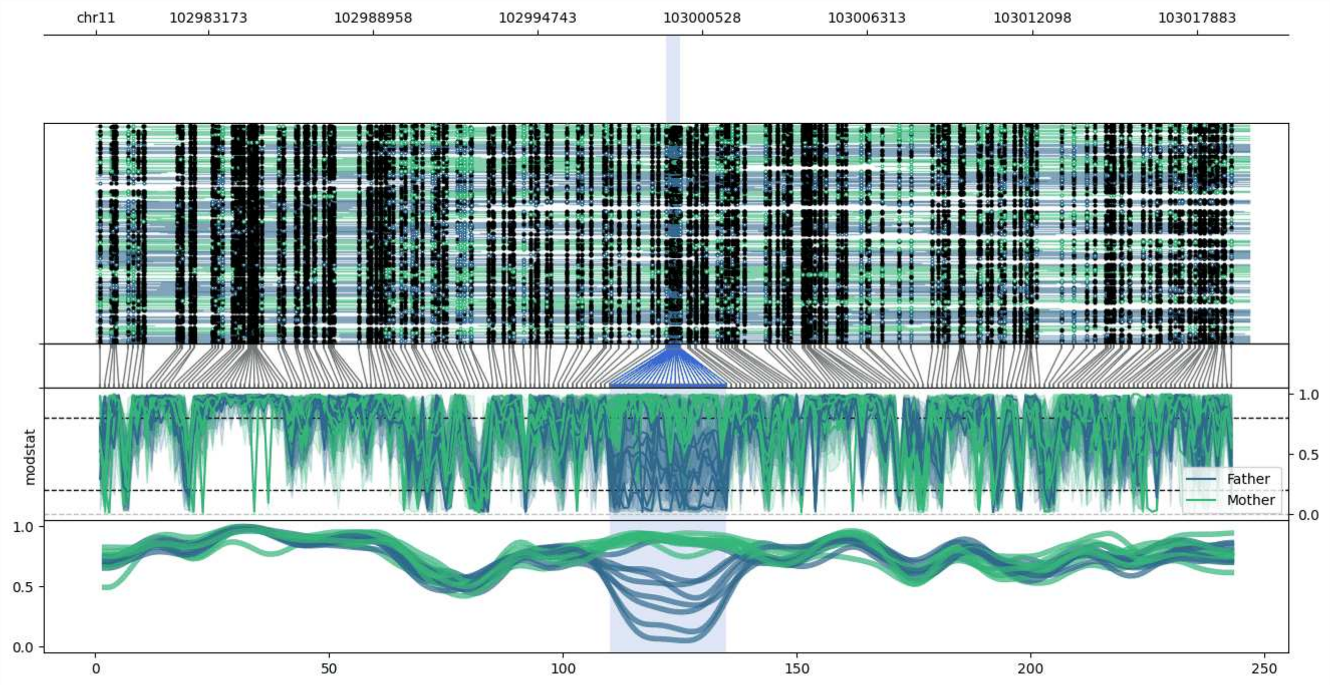


**chr11:102979240-103019741:** Region with 1 DMR containing no genes.

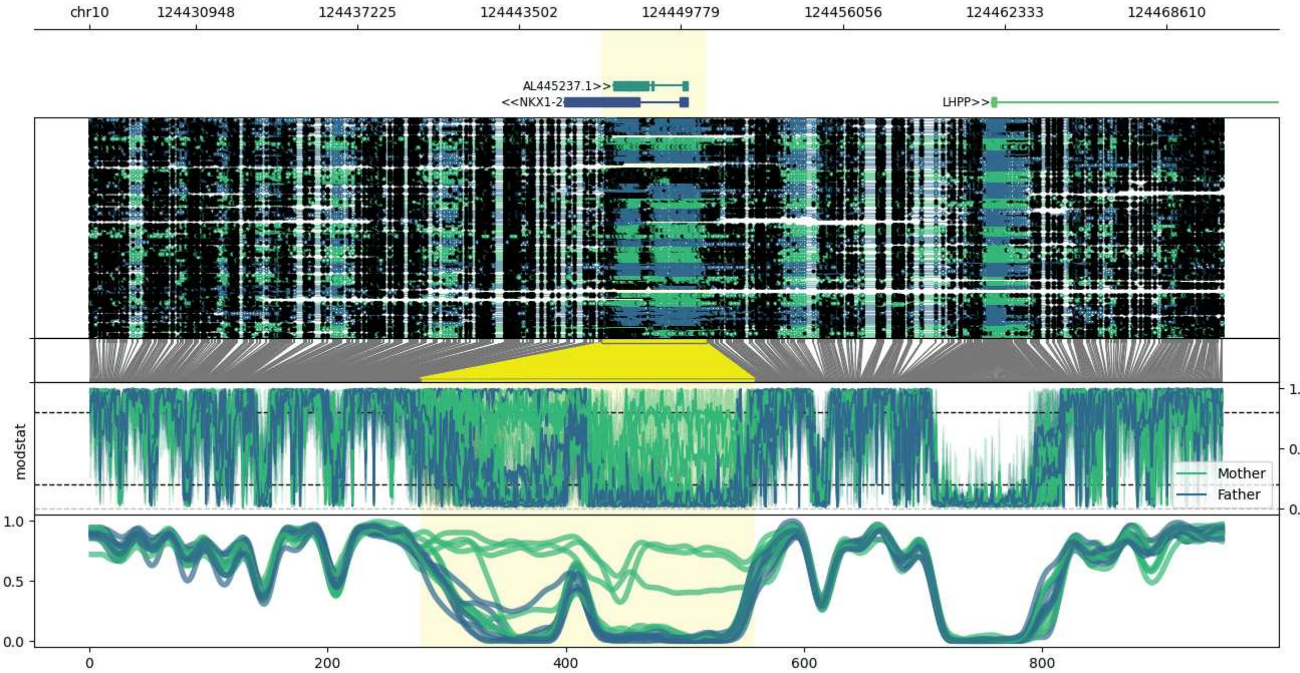


**chr10:124426835-124470774:** Region with 1 DMR containing genes: AL445237.1, LHPP, NKX1-2, OAT.

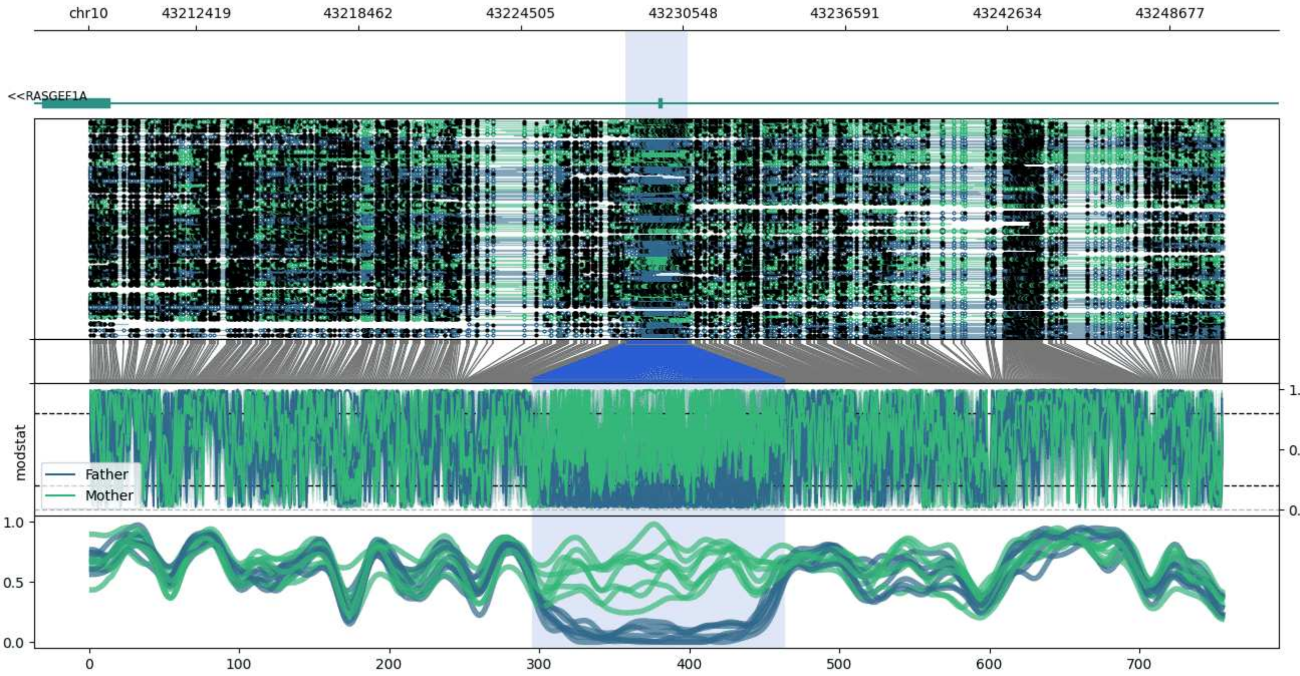


**chr10:43208410-43250715:** Region with 1 DMR containing genes: RASGEF1A.

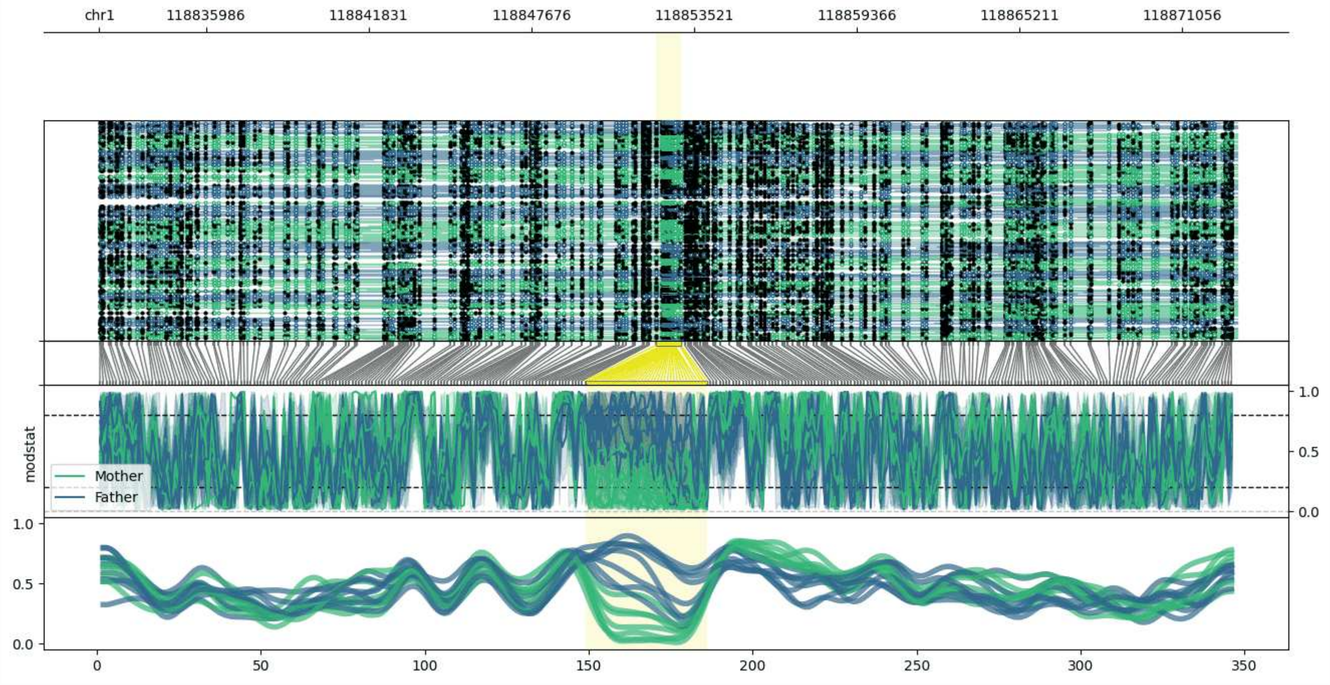


**chr1:118832149-118873064:** Region with 1 DMR containing genes: TBX15.

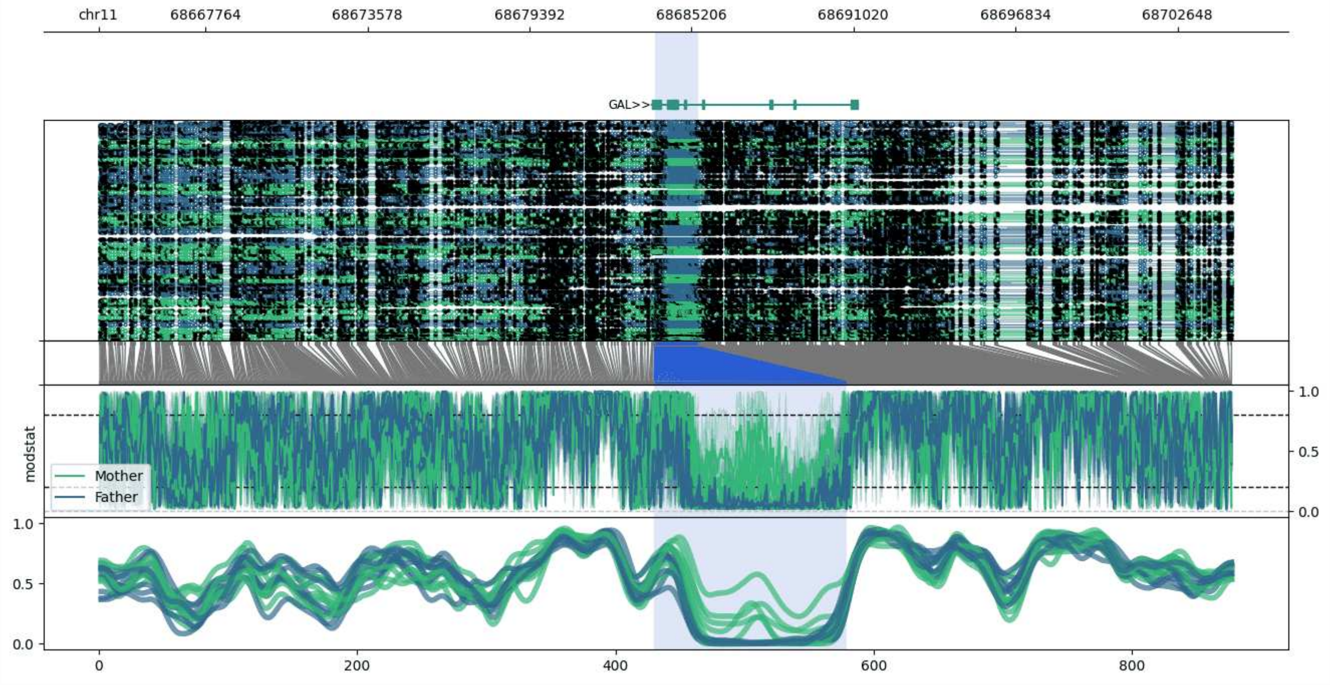


**chr11:68663928-68704629:** Region with 1 DMR containing genes: GAL, TESMIN.

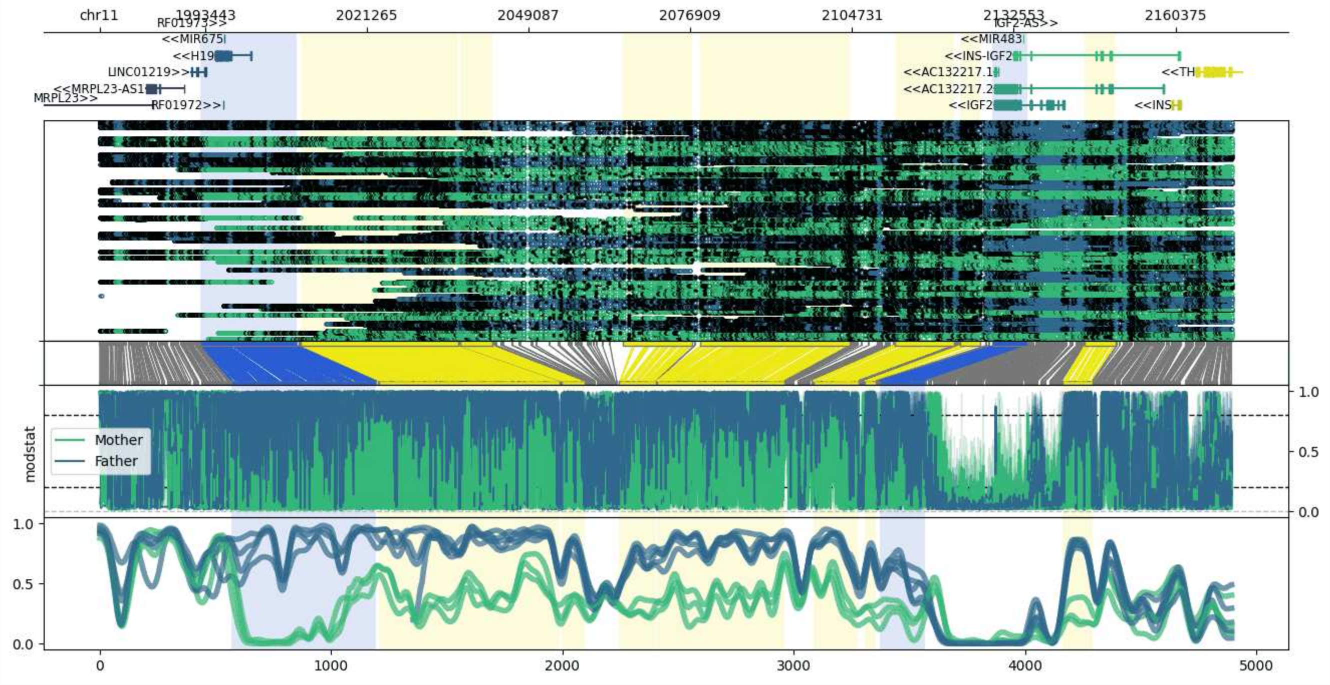


**chr11:1975283-2170042:** Region with 9 DMRs containing genes: AC132217.1, AC132217.2, H19, IGF2, IGF2-AS, INS, INS-IGF2, LINC01219, MIR4686, MIR483, MIR675, MRPL23, MRPL23-AS1, RF01972, RF01973, TH.

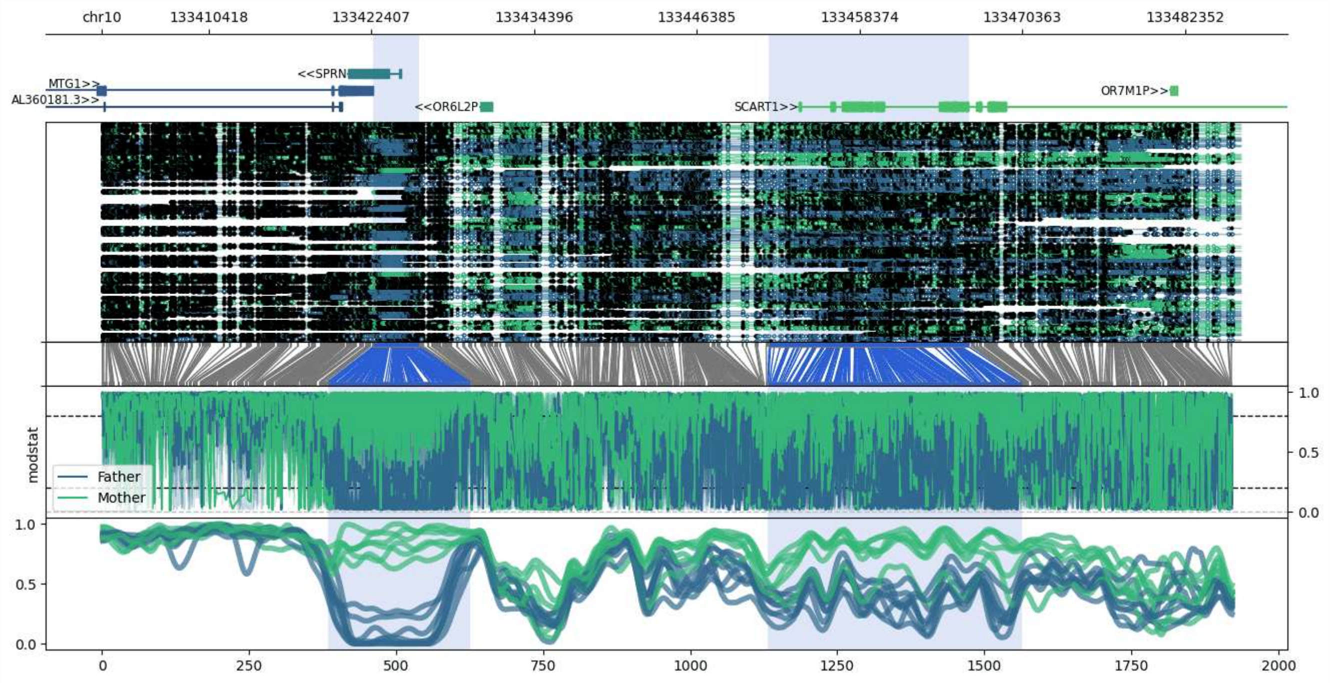


**chr10:133402546-133486470:** Region with 2 DMRs containing genes: AL360181.3, MTG1, OR6L2P, OR7M1P, SCART1, SPRN.

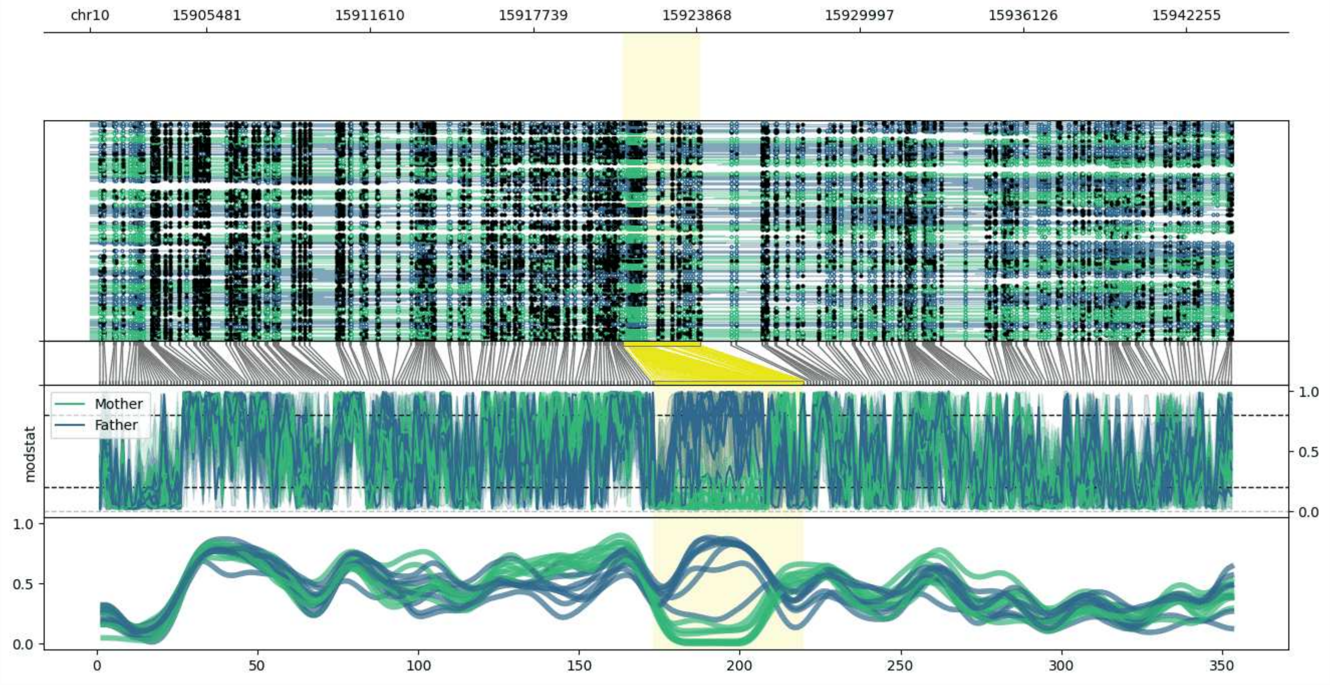


**chr10:15901108-15944013:** Region with 1 DMR containing no genes.

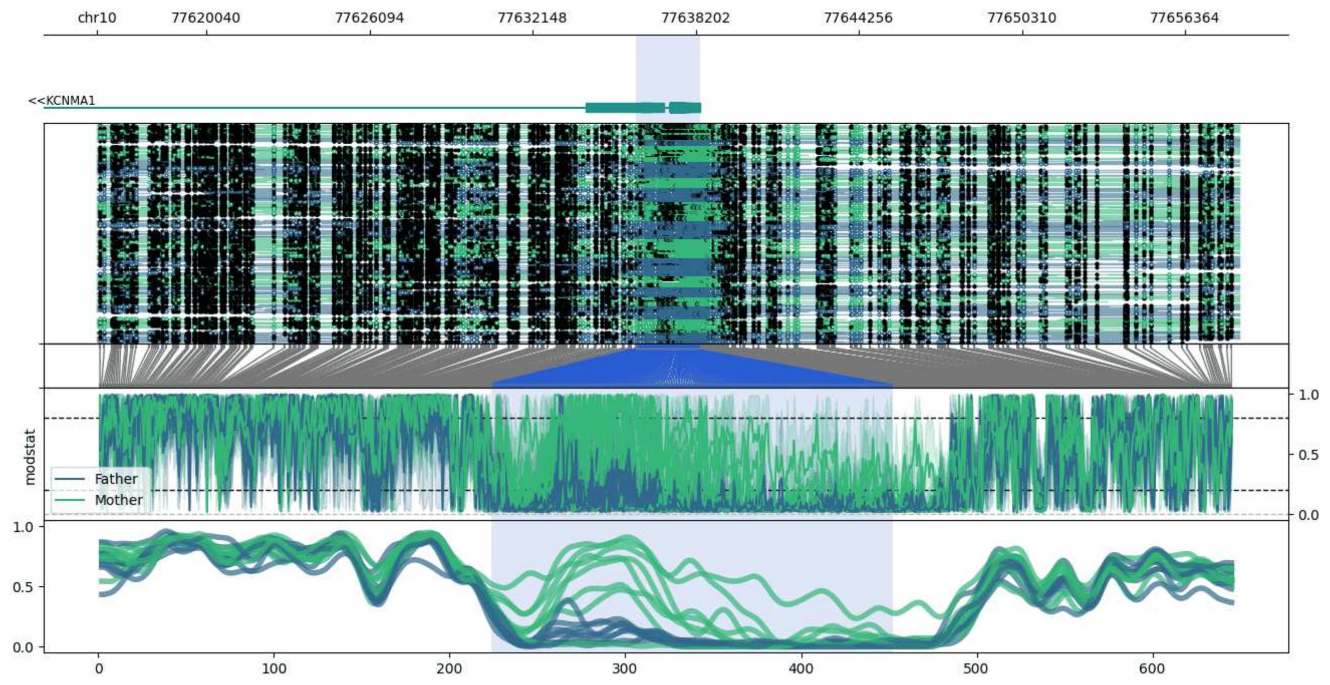


**chr10:77615988-77658366:** Region with 1 DMR containing genes: KCNMA1.

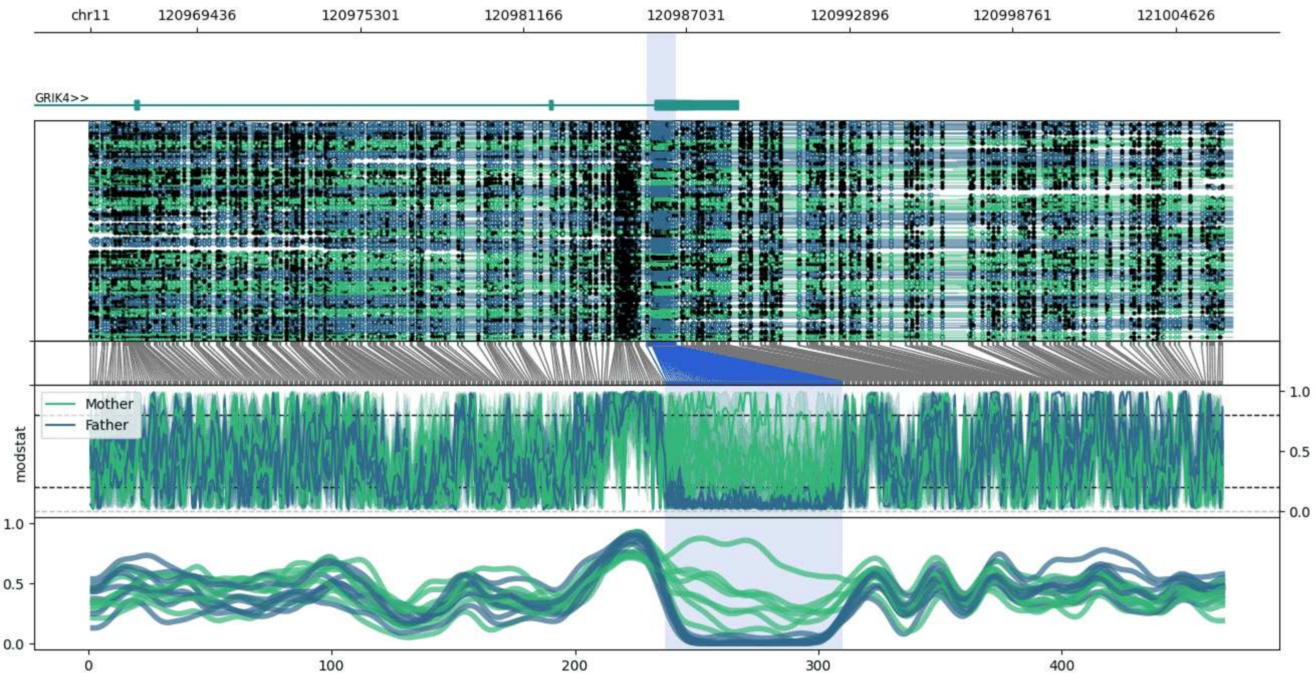


**chr11:120965593-121006649:** Region with 1 DMR containing genes: GRIK4.

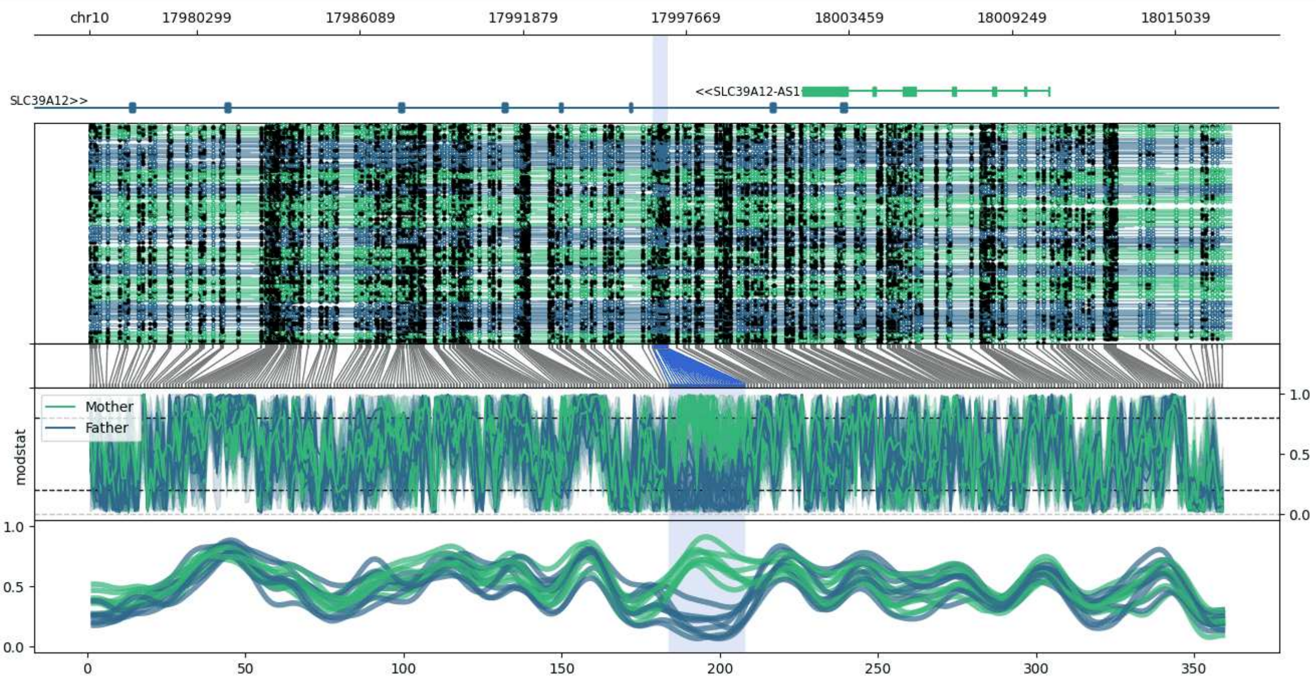


**chr10:17976481-18017016:** Region with 1 DMR containing genes: SLC39A12, SLC39A12-AS1.

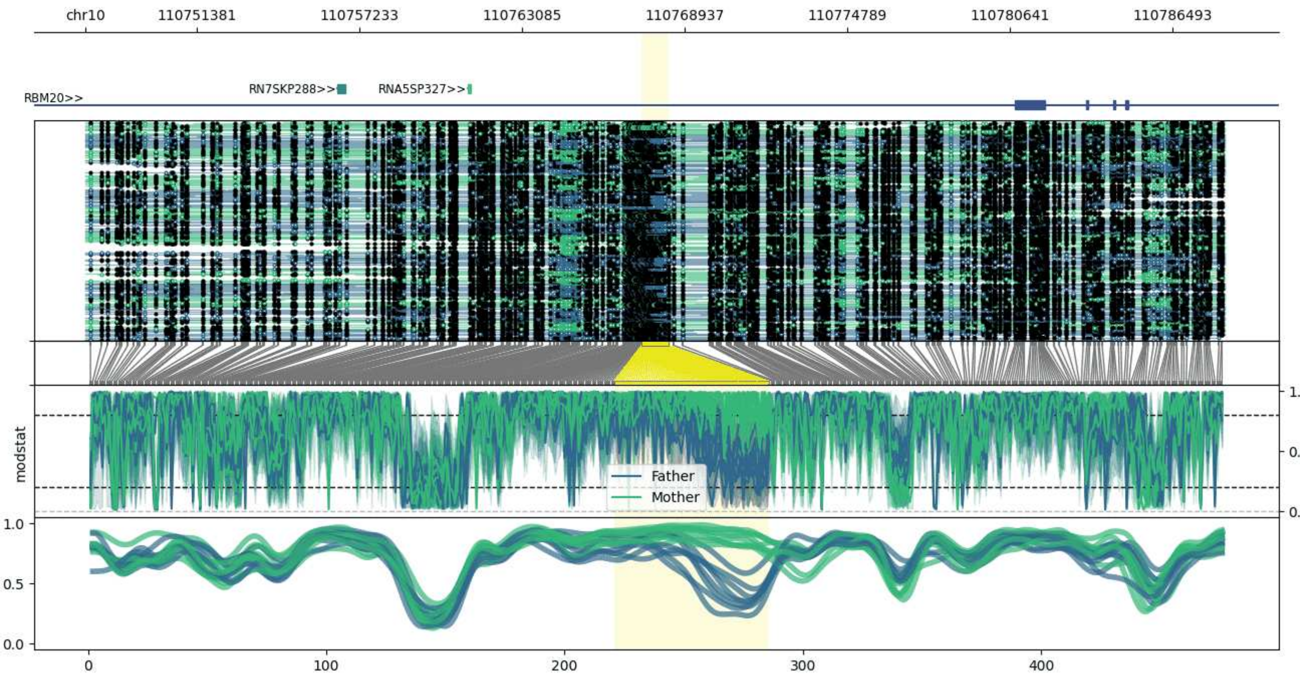


**chr10:110747382-110788349:** Region with 1 DMR containing genes: RBM20, RN7SKP288, RNA5SP327.

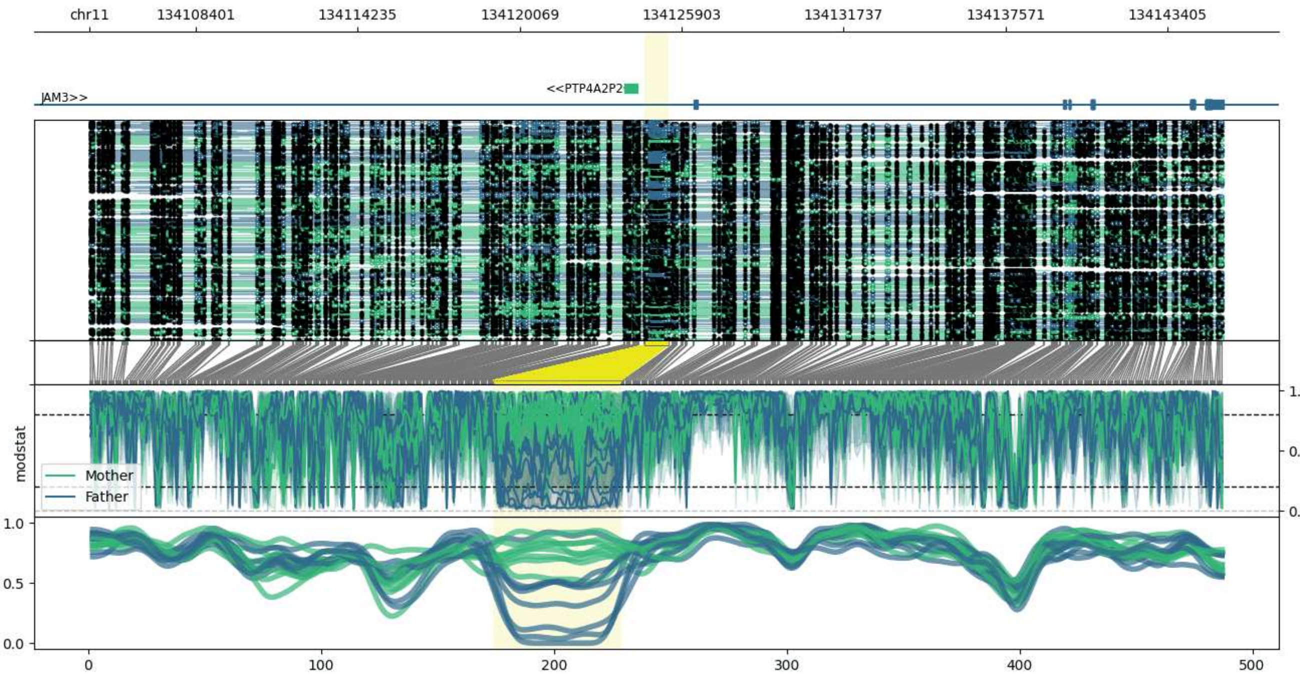


**chr11:134104574-134145417:** Region with 1 DMR containing genes: JAM3, NCAPD3, PTP4A2P2.

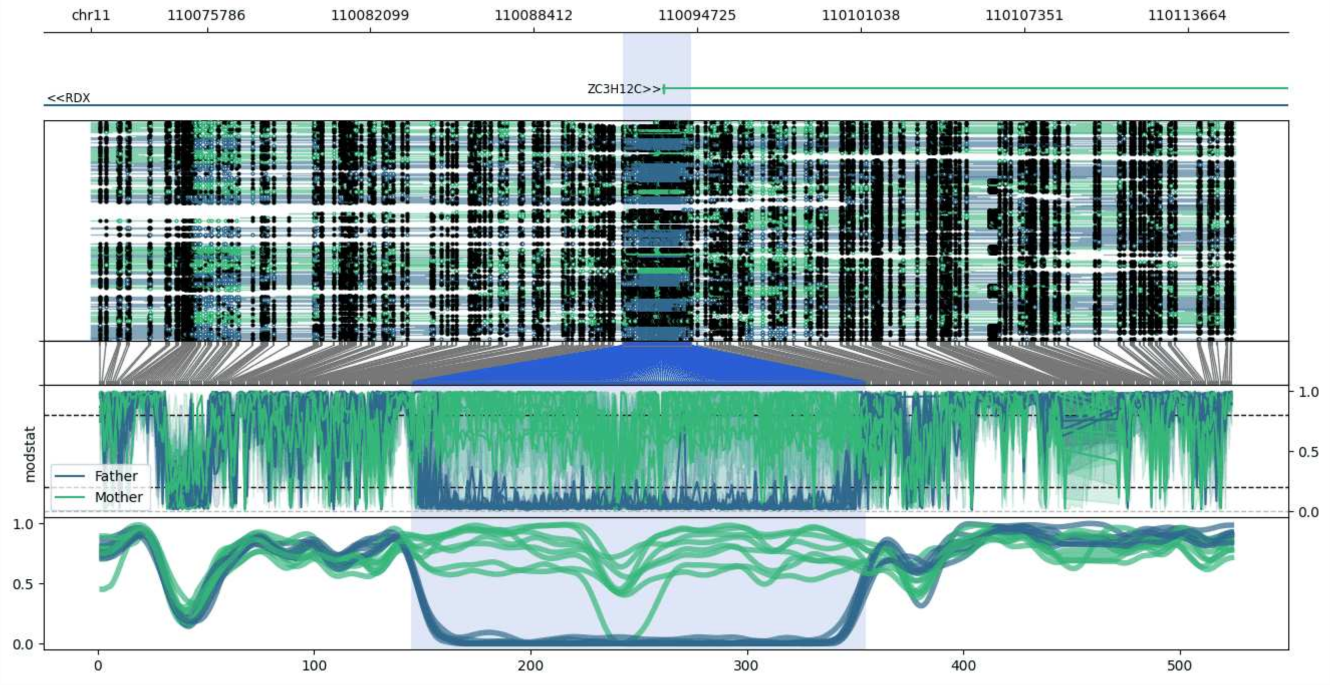


**chr11:110071320-110115514:** Region with 1 DMR containing genes: RDX, ZC3H12C.

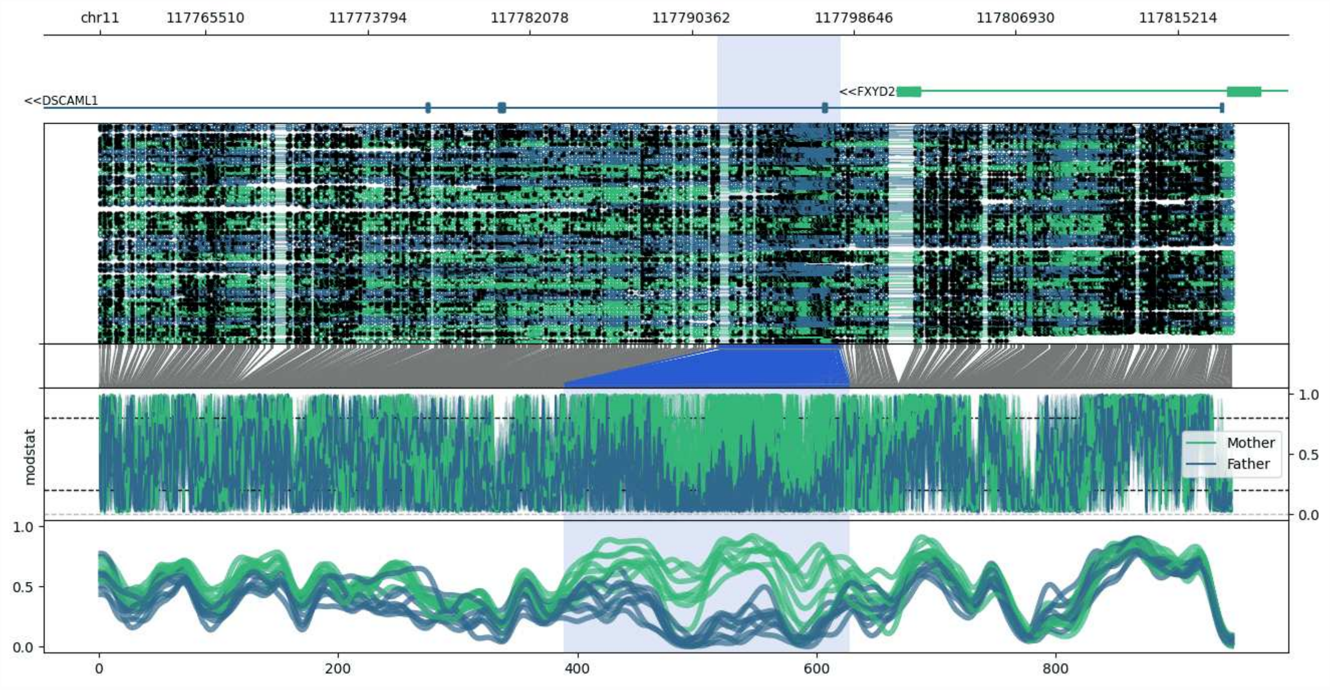


**chr11:117760105-117818099:** Region with 1 DMR containing genes: AP000757.2, DSCAML1, FXYD2, FXYD6-FXYD2.

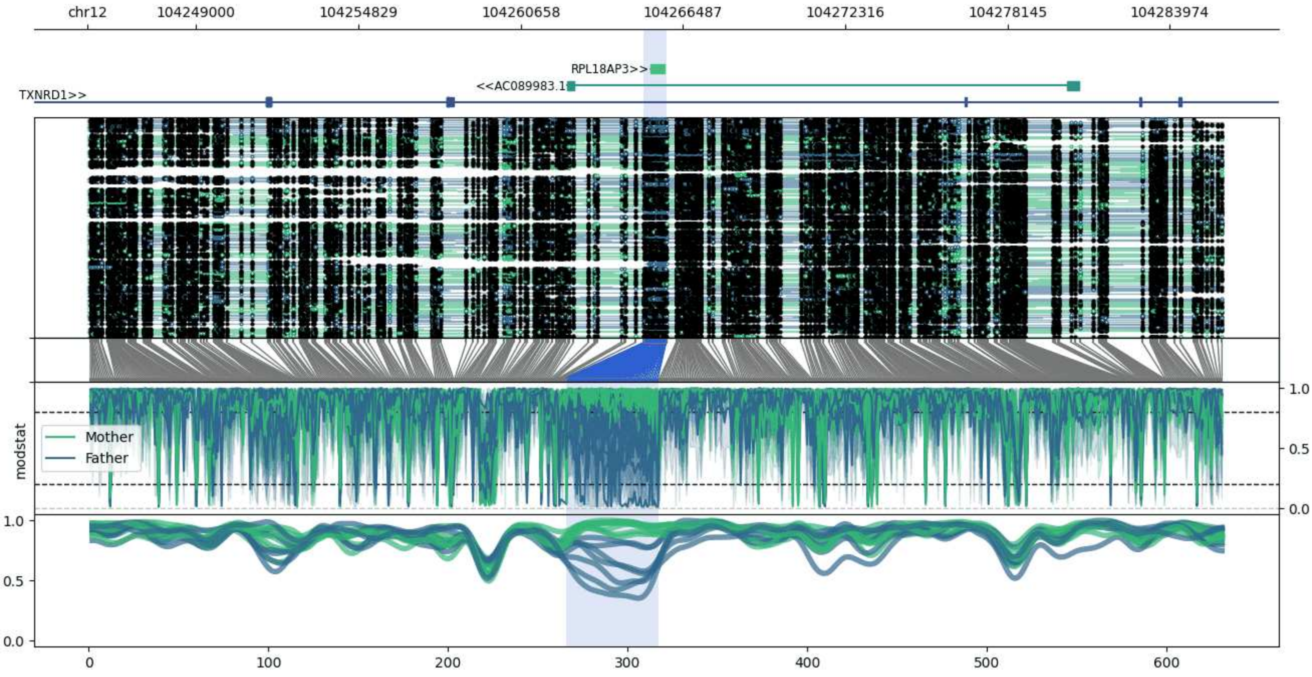


**chr12:104245079-104285887:** Region with 1 DMR containing genes: AC089983.1, RPL18AP3, TXNRD1.

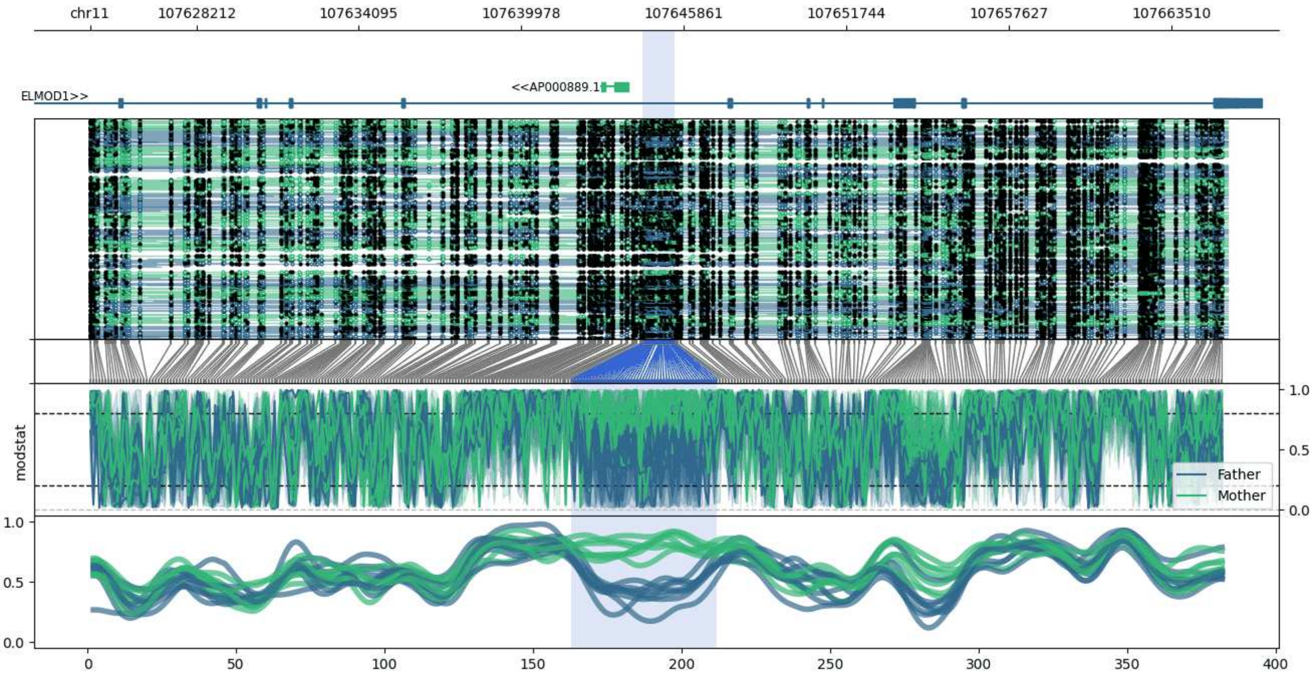


**chr11:107624360-107665543:** Region with 1 DMR containing genes: AP000889.1, ELMOD1.

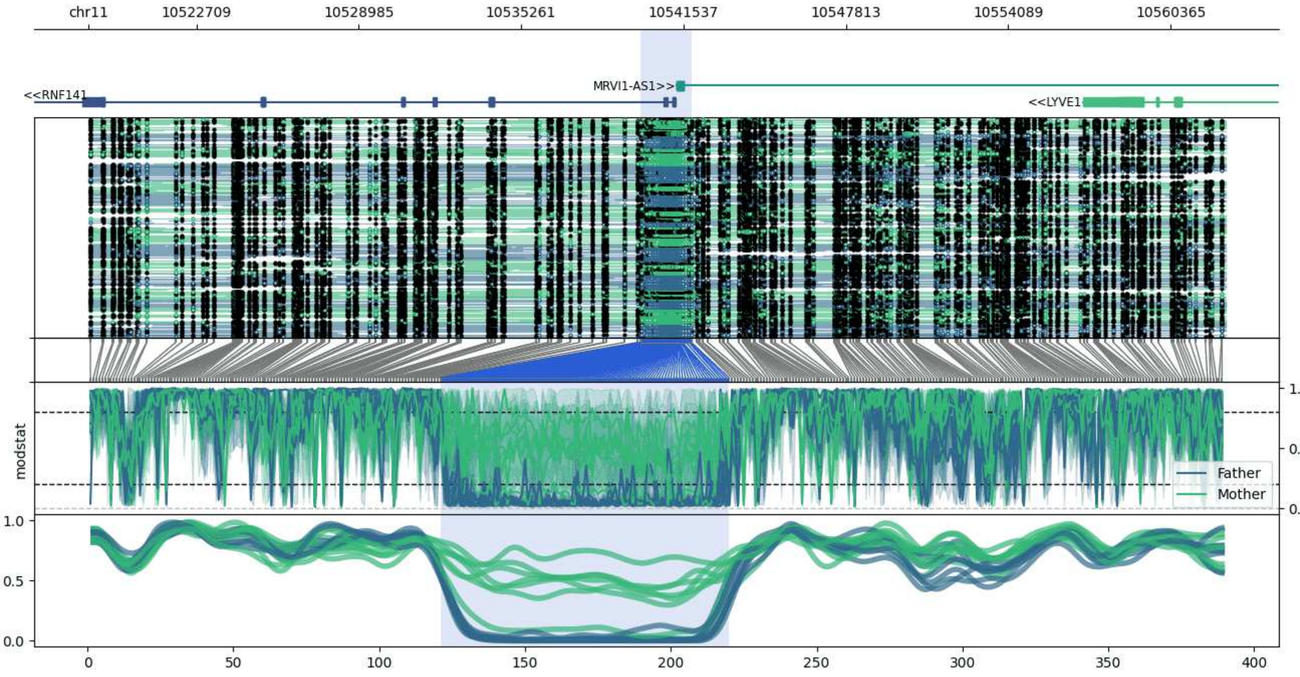


**chr11:10518544-10562482:** Region with 1 DMR containing genes: LYVE1, MRVI1-AS1, MTRNR2L8, RNF141.

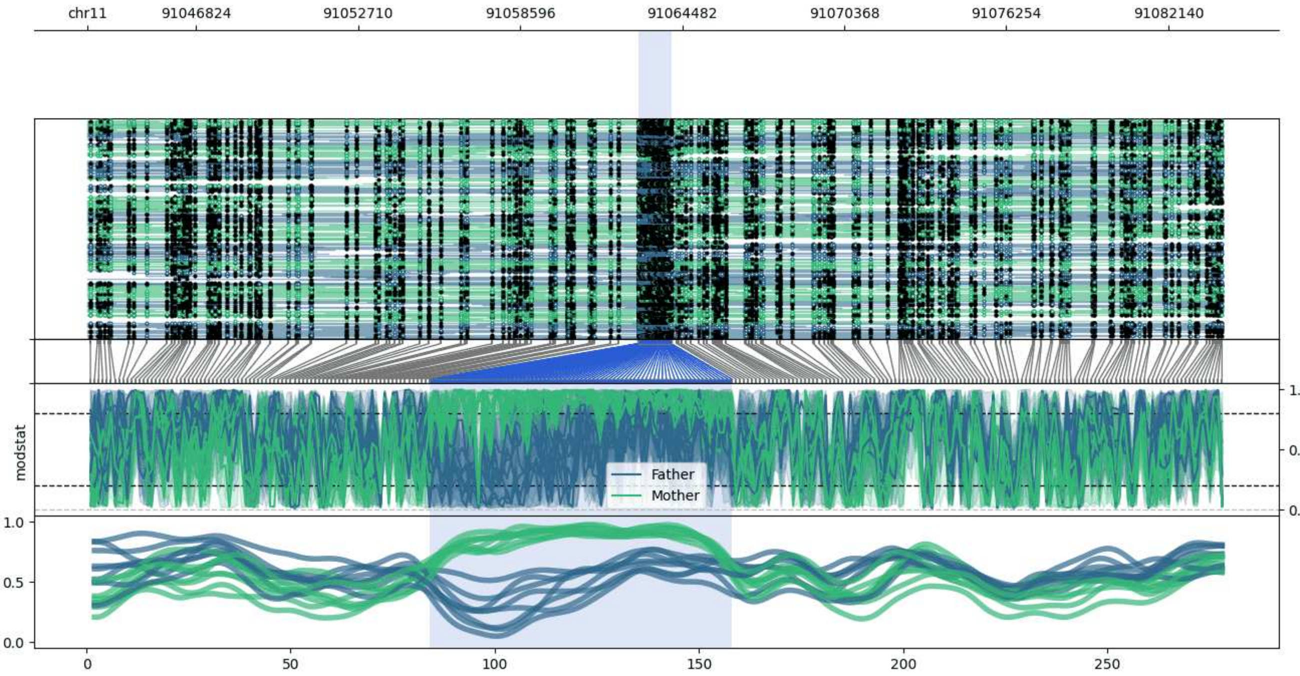


**chr11:91042885-91084088:** Region with 1 DMR containing genes: AP001002.2.

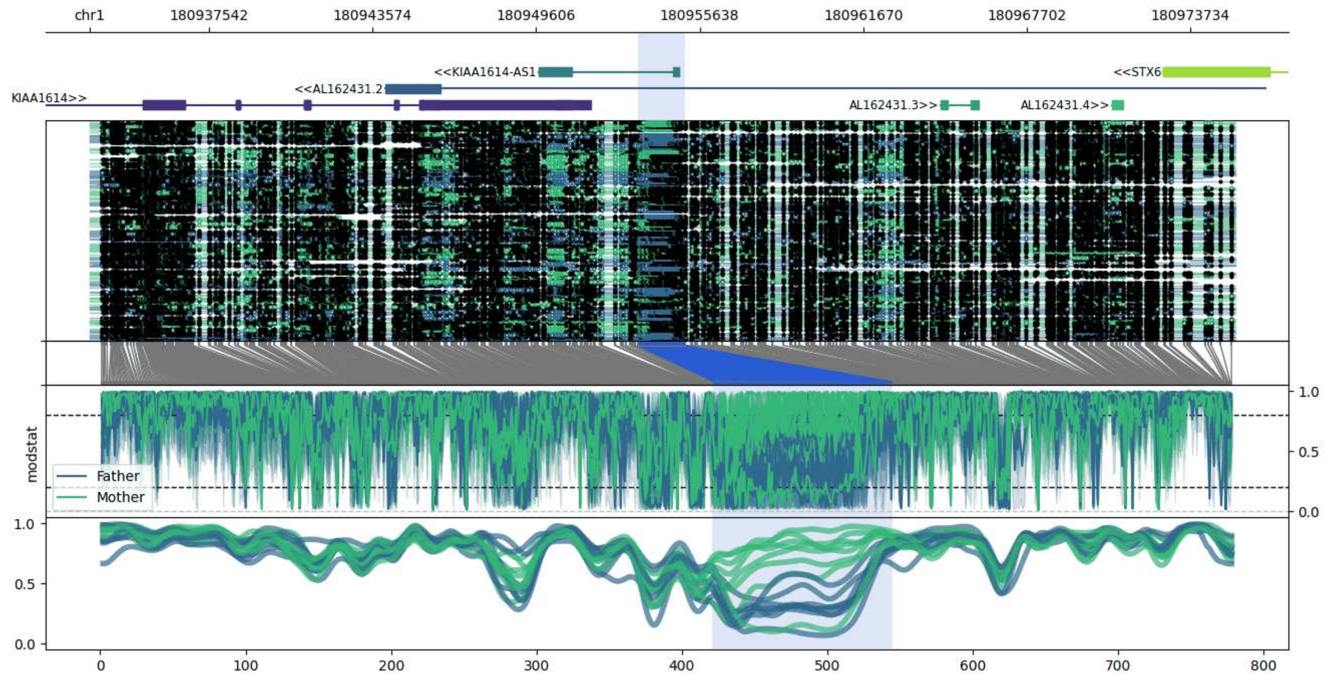


**chr1:180933141-180975370:** Region with 1 DMR containing genes: AL162431.2, AL162431.3, AL162431.4, KIAA1614, KIAA1614-AS1, STX6.

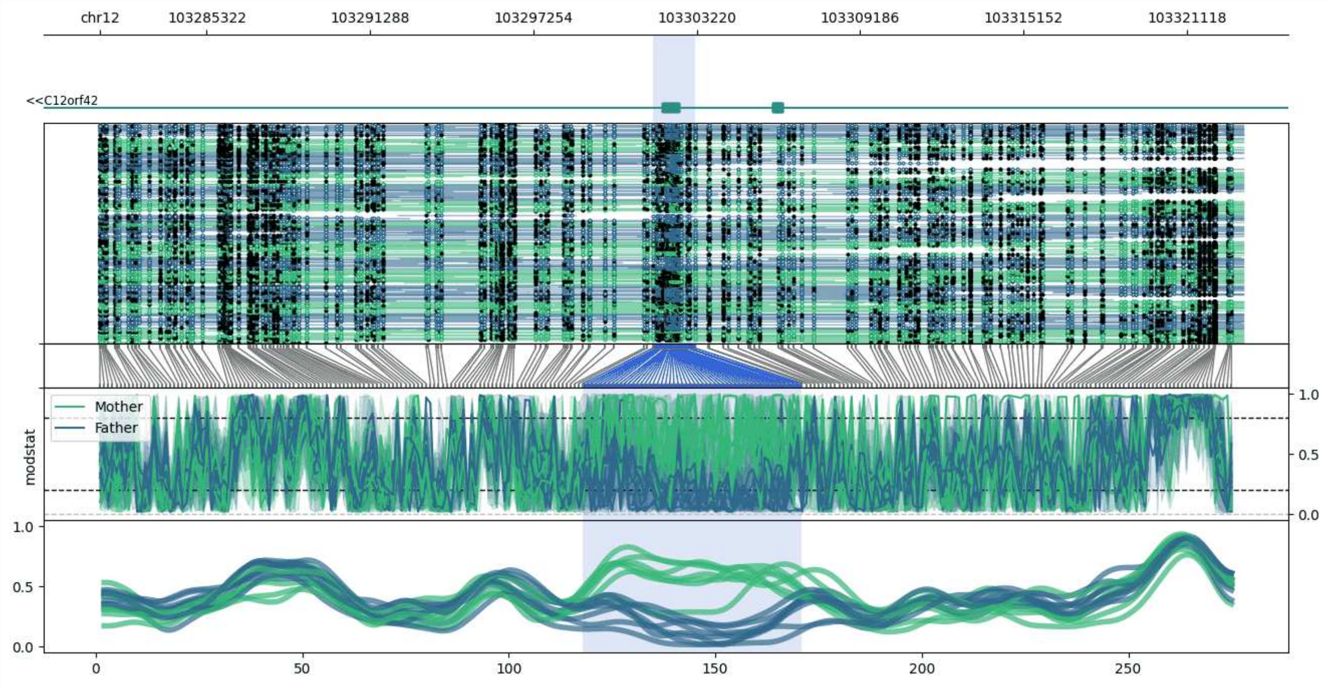


**chr12:103281413-103323177:** Region with 1 DMR containing genes: C12orf42.

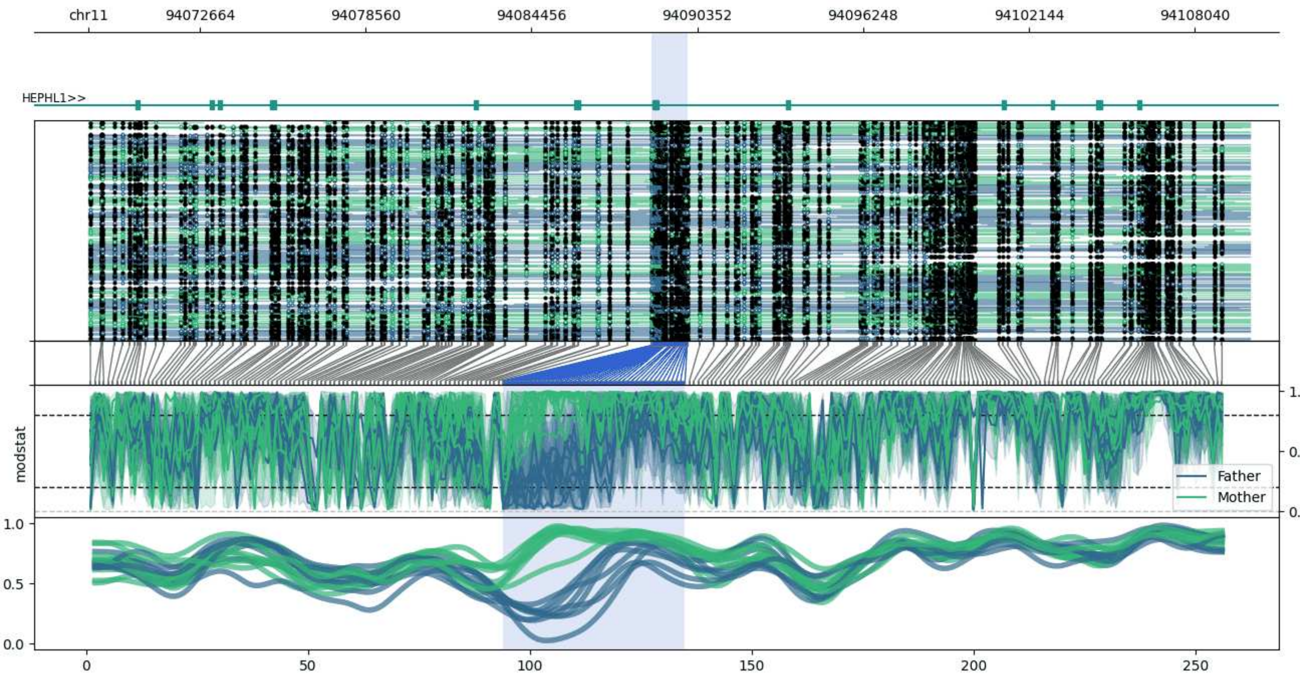


**chr11:94068716-94109994:** Region with 1 DMR containing genes: HEPHL1.

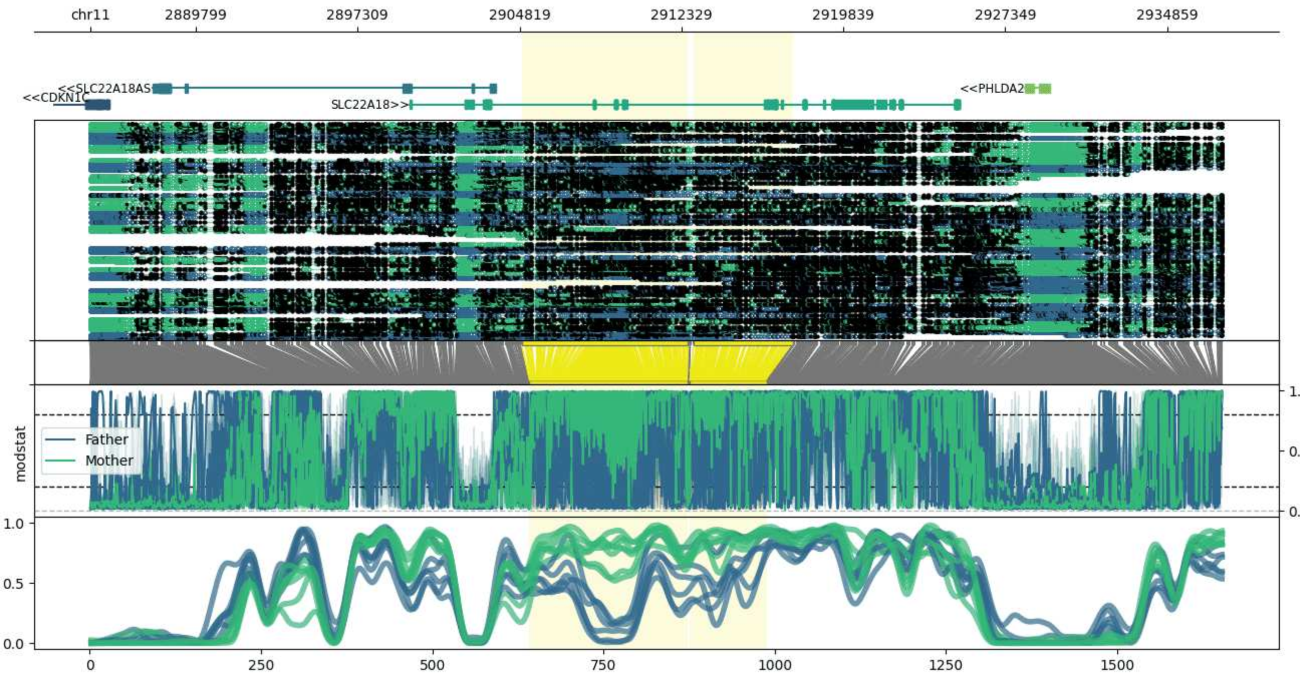


**chr11:2884912-2937482:** Region with 2 DMRs containing genes: CDKN1C, NAP1L4, PHLDA2, SLC22A18, SLC22A18AS.

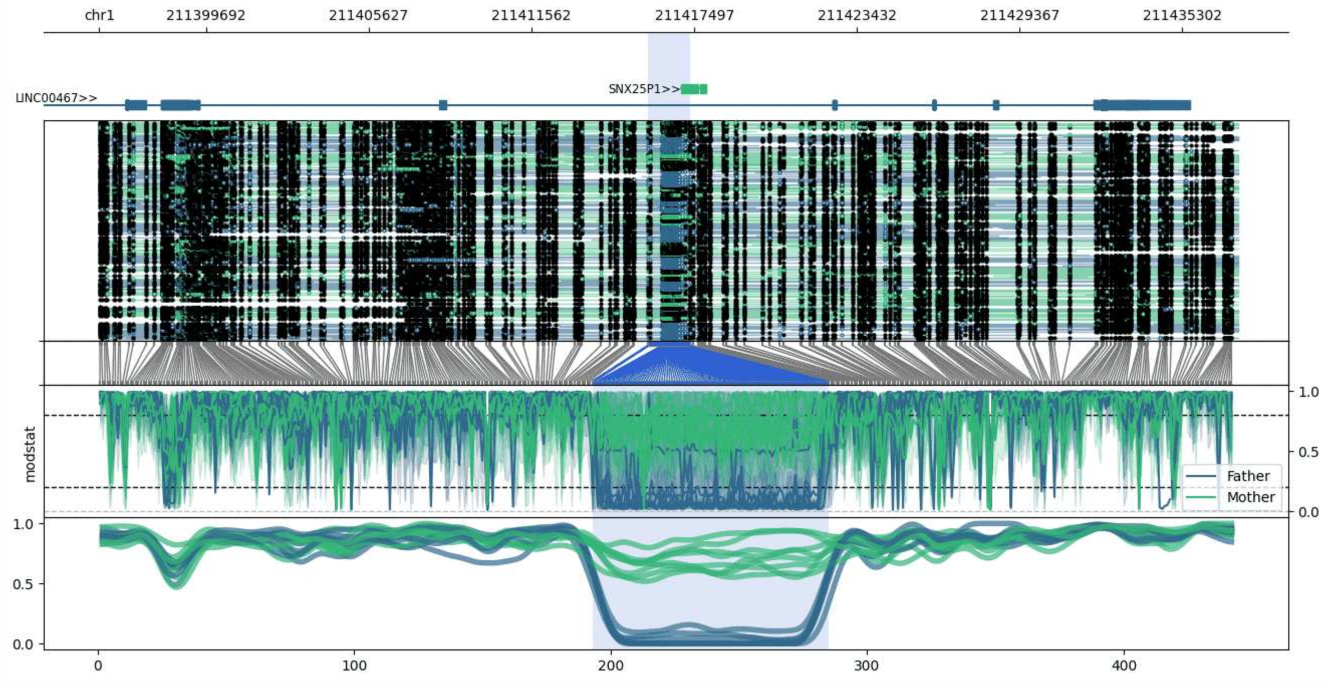


**chr1:211395796-211437345:** Region with 1 DMR containing genes: ARPC3P2, LINC00467, SNX25P1.

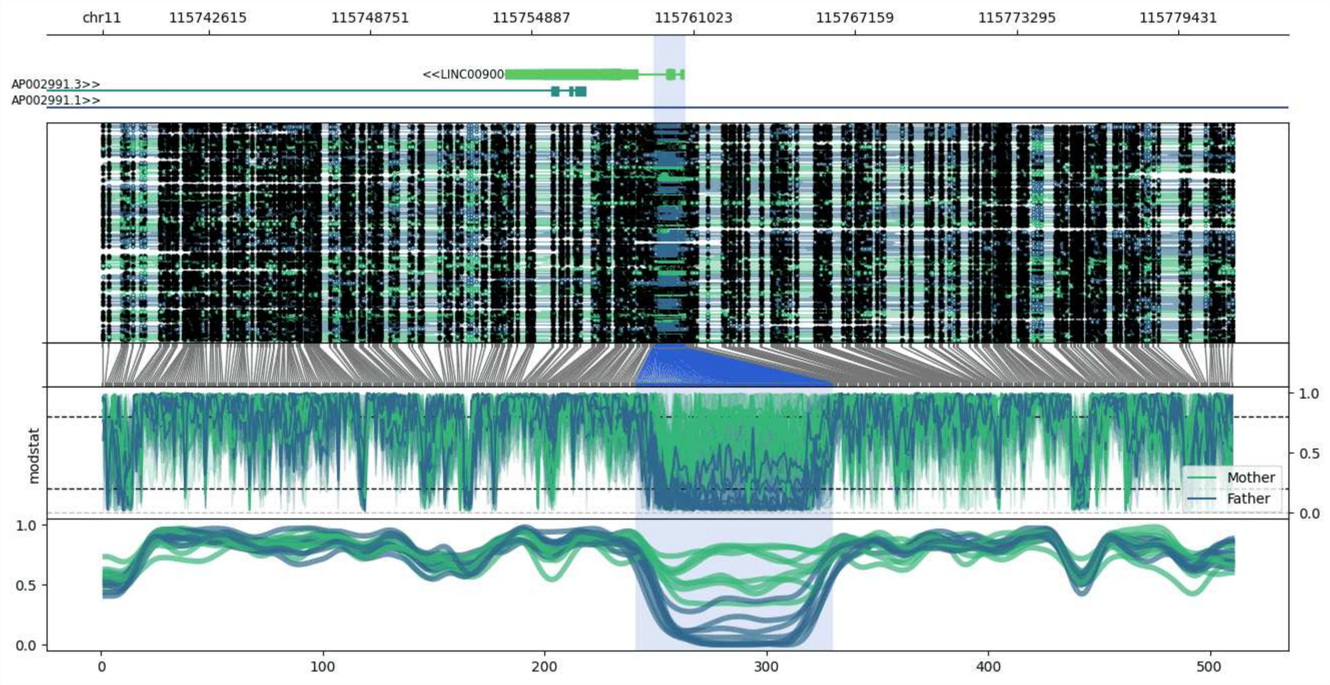


**chr11:115738591-115781545:** Region with 1 DMR containing genes: AP002991.1, AP002991.2, AP002991.3, LINC00900.

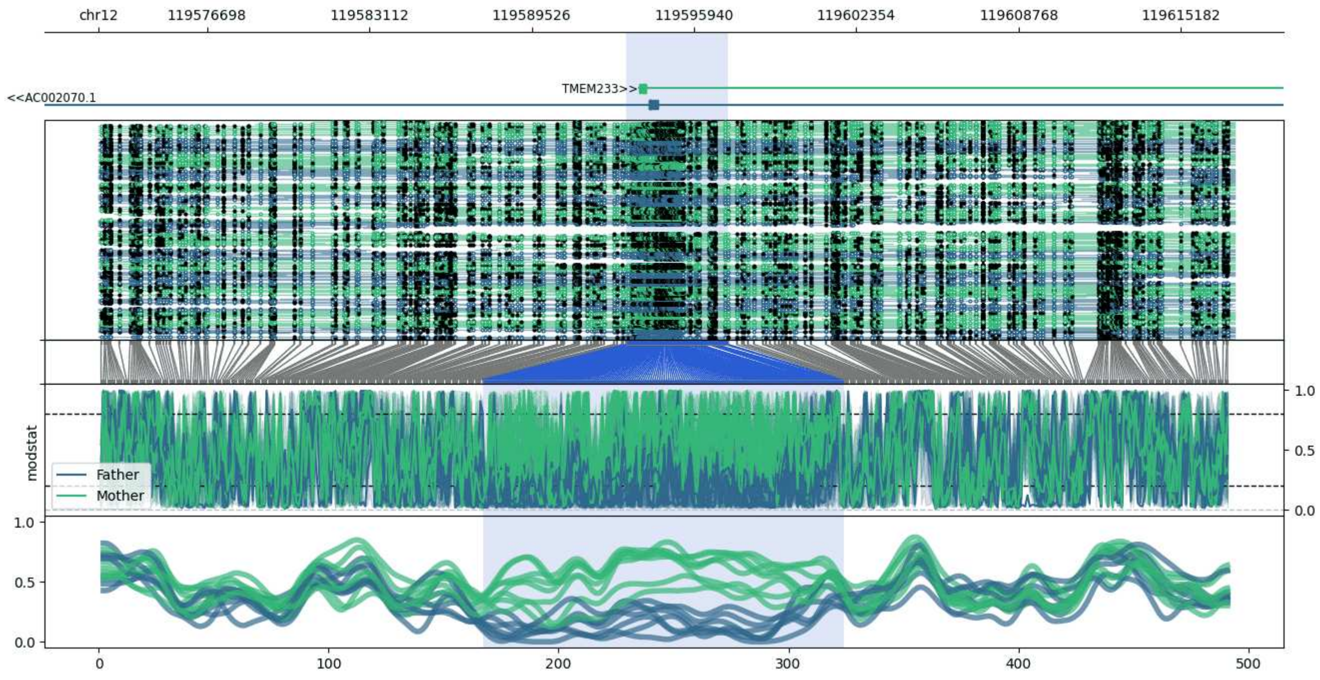


**chr12:119572398-119617298:** Region with 1 DMR containing genes: AC002070.1, TMEM233.

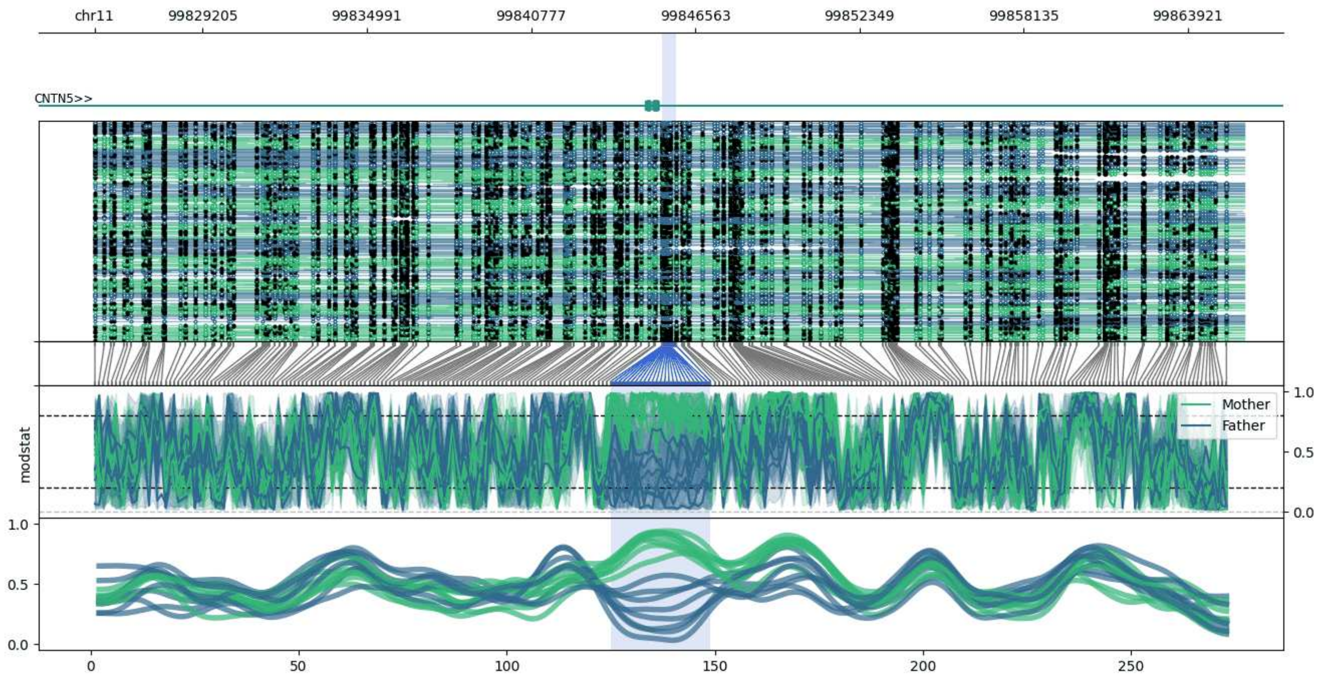


**chr11:99825394-99865898:** Region with 1 DMR containing genes: CNTN5.

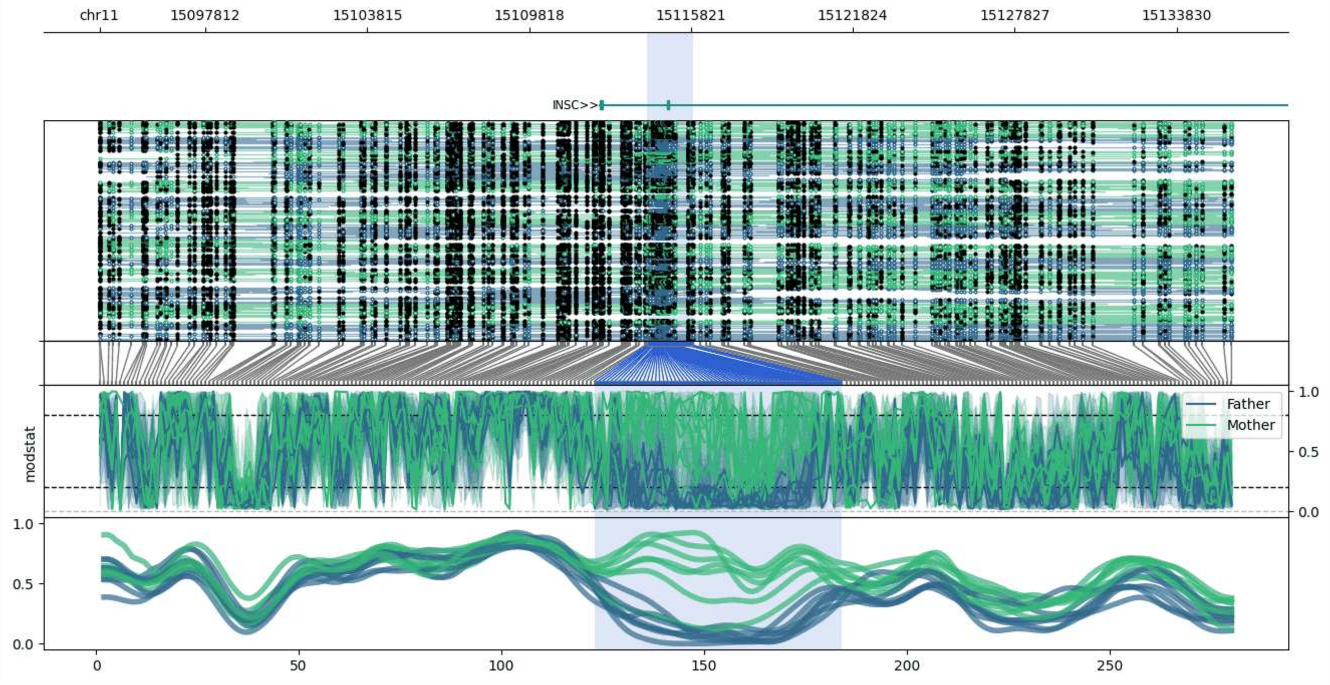


**chr11:15093885-15135908:** Region with 1 DMR containing genes: INSC.

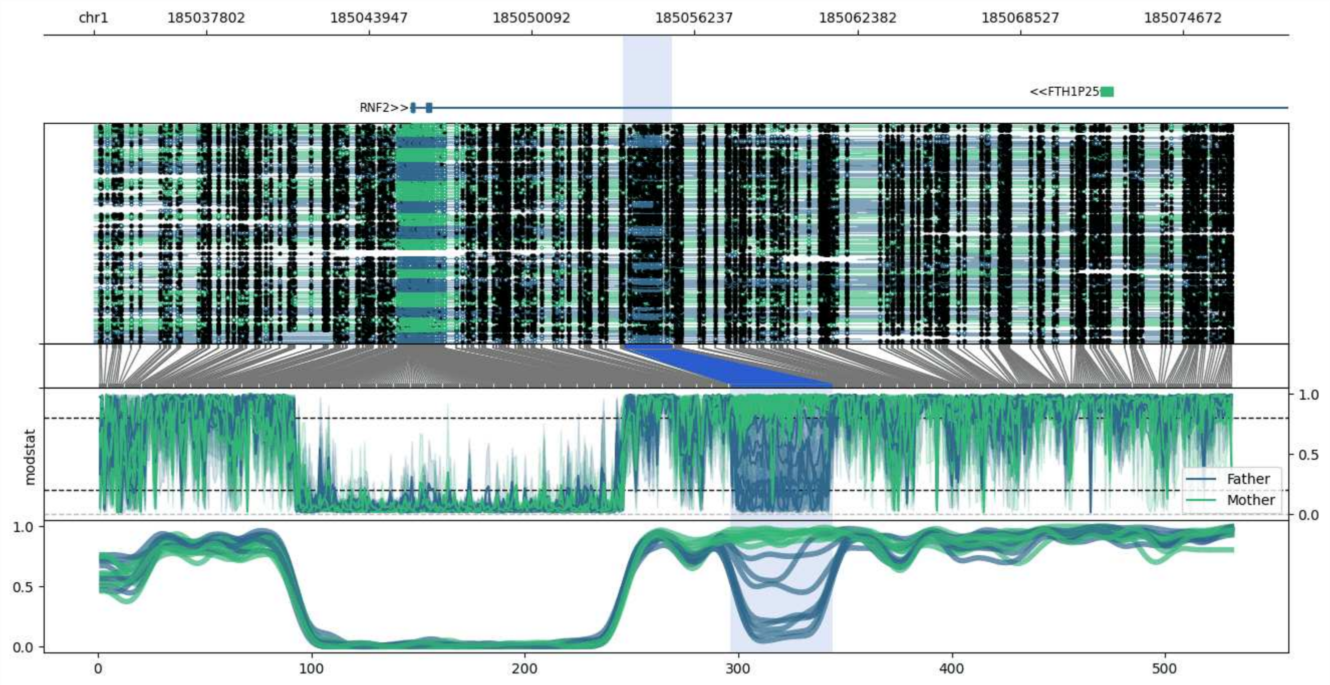


**chr1:185033543-185076559:** Region with 1 DMR containing genes: FTH1P25, RNF2.

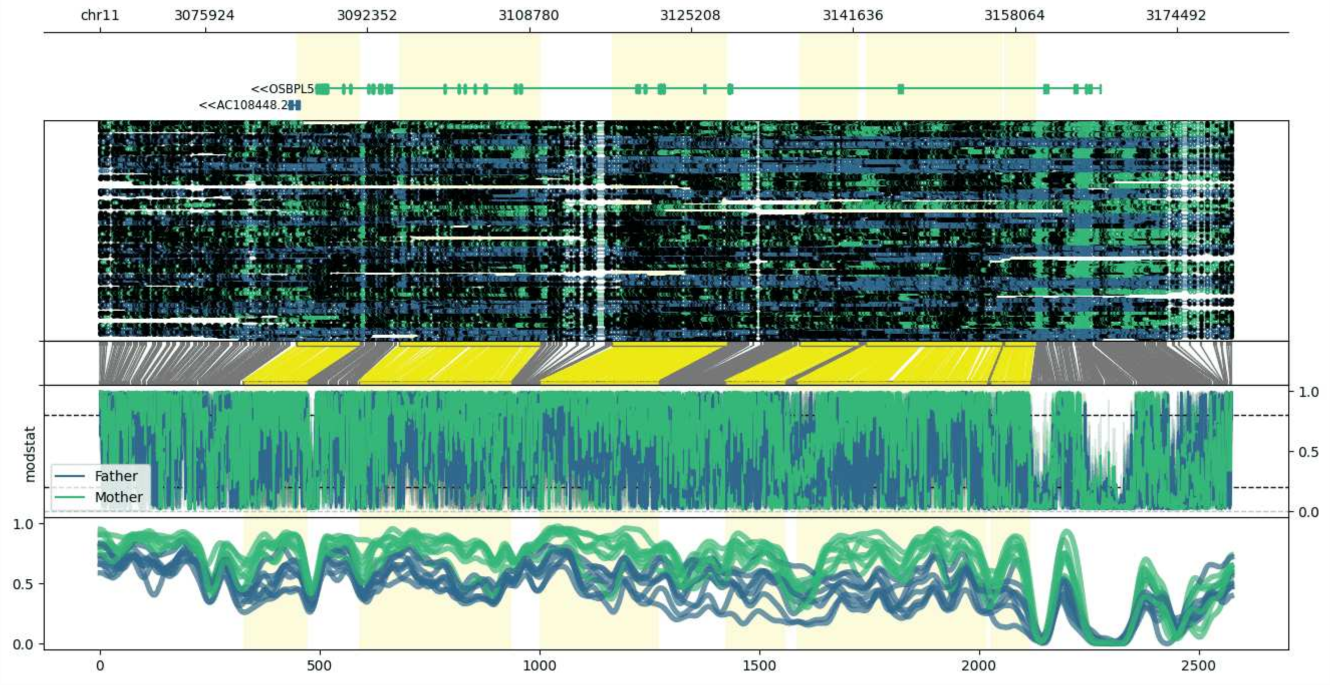


**chr11:3065194-3180194:** Region with 6 DMRs containing genes: AC108448.1, AC108448.2, AC108448.3, CARS, OSBPL5.

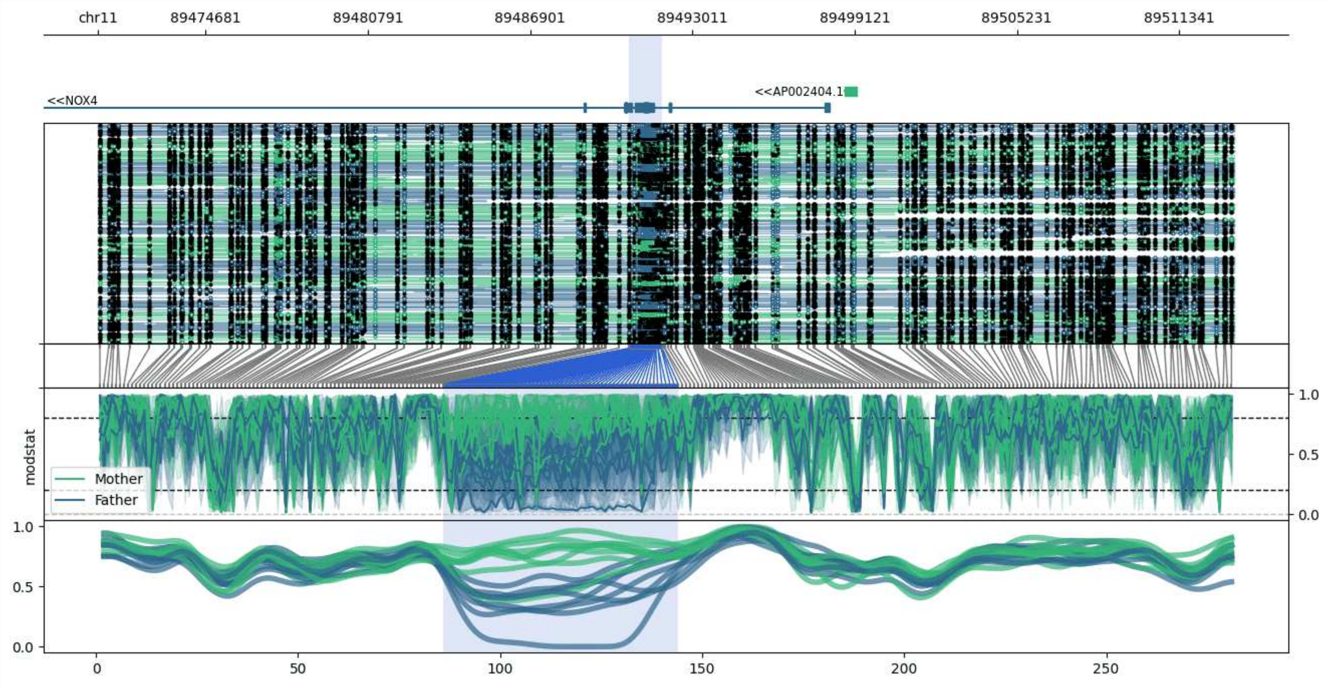


**chr11:89470636-89513412:** Region with 1 DMR containing genes: AP002404.1, NOX4.

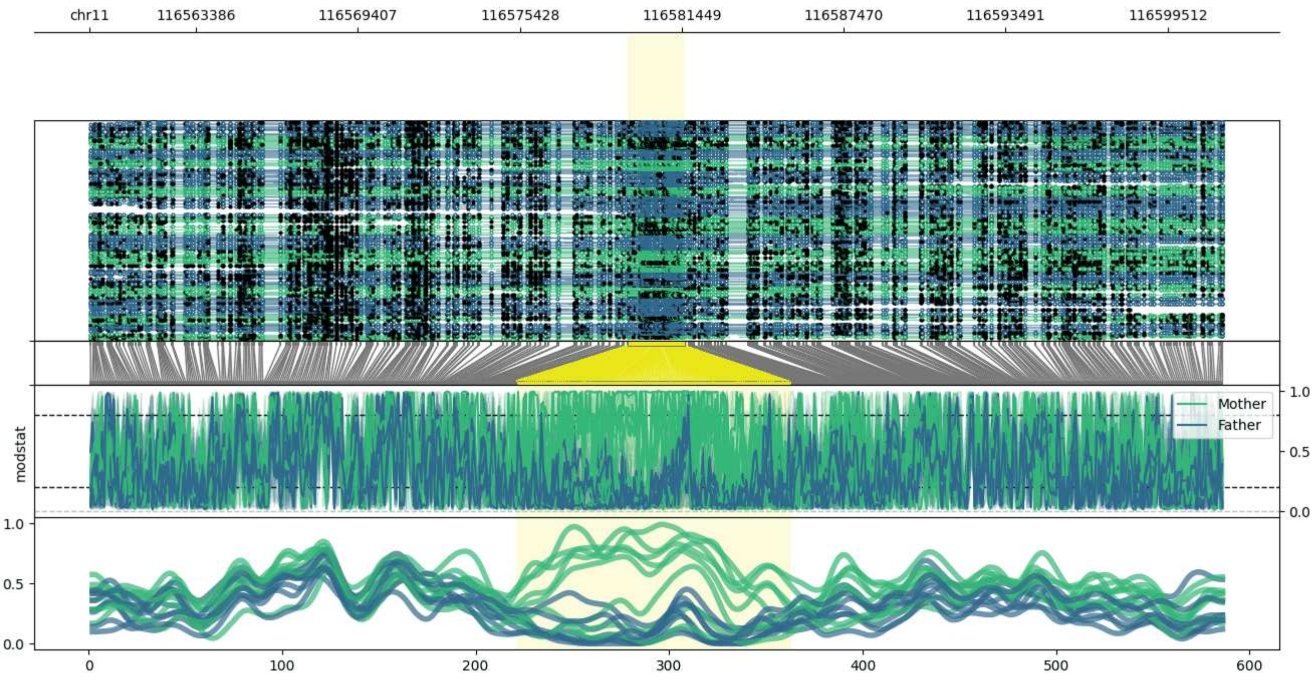


**chr11:116559436-116601583:** Region with 1 DMR containing genes: AP000770.2.

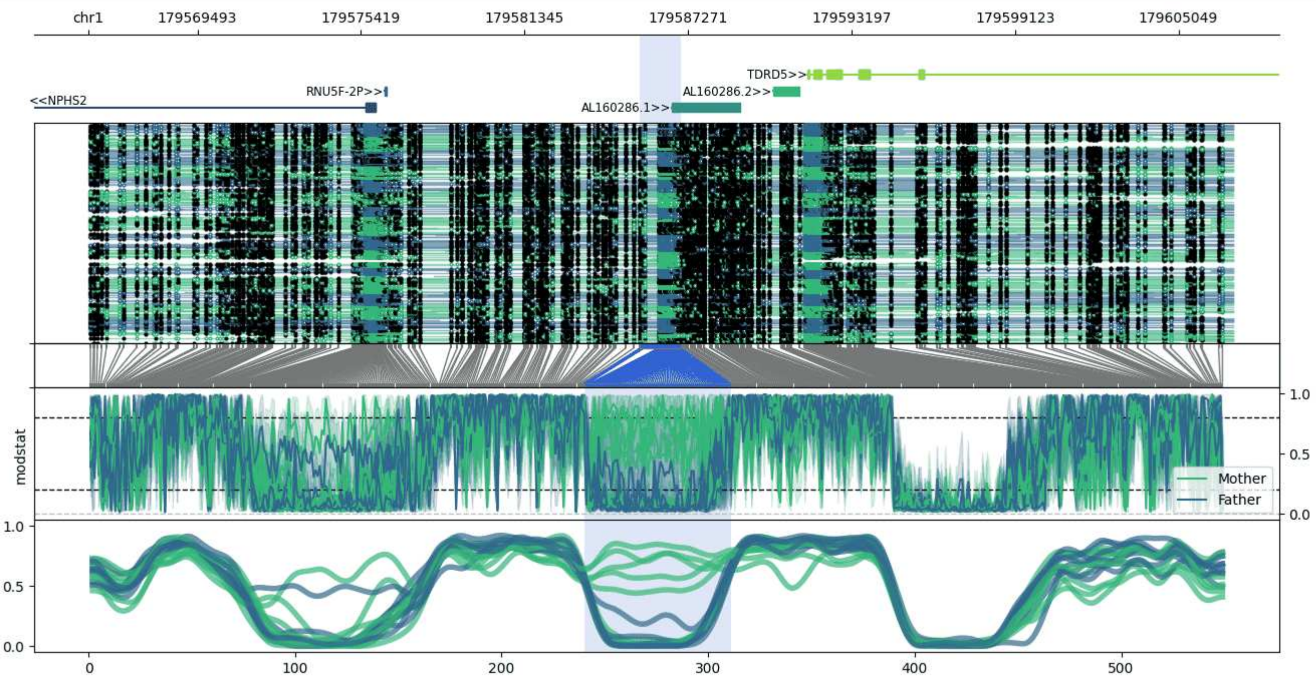


**chr1:179565540-179607023:** Region with 1 DMR containing genes: AL160286.1, AL160286.2, NPHS2, RNU5F-2P, TDRD5.

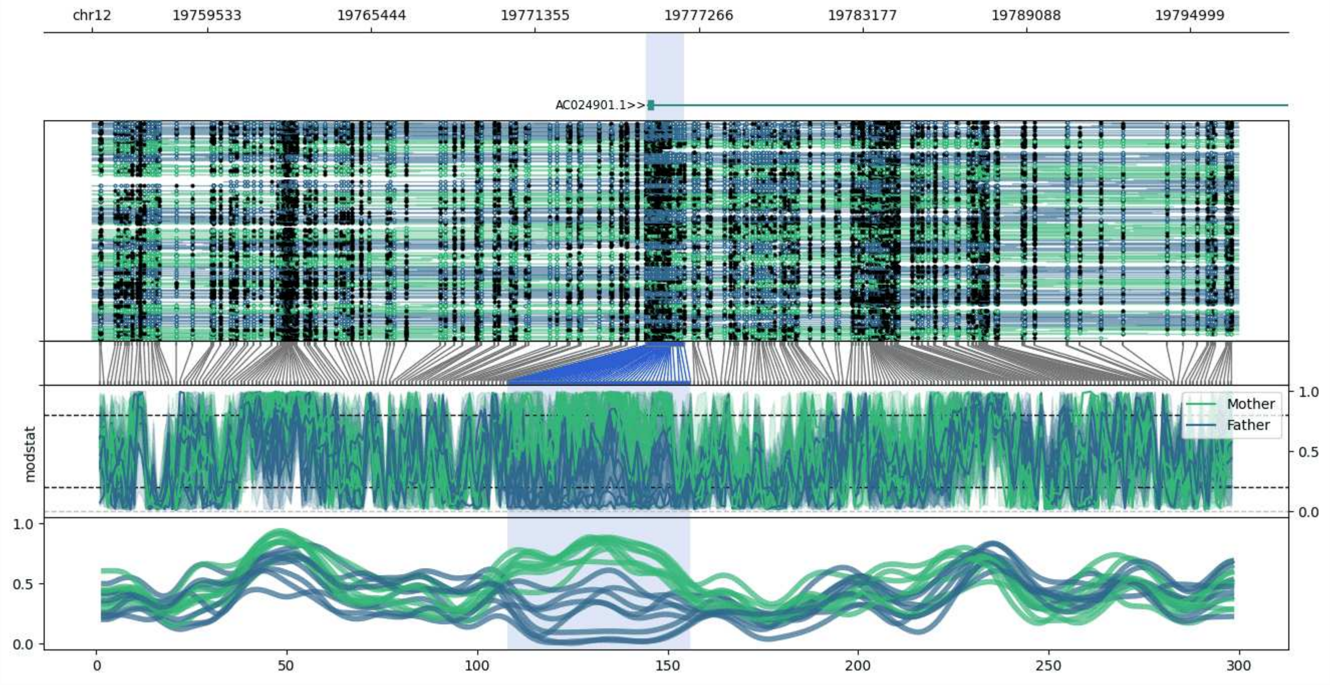


**chr12:19755357-19796738:** Region with 1 DMR containing genes: AC024901.1.

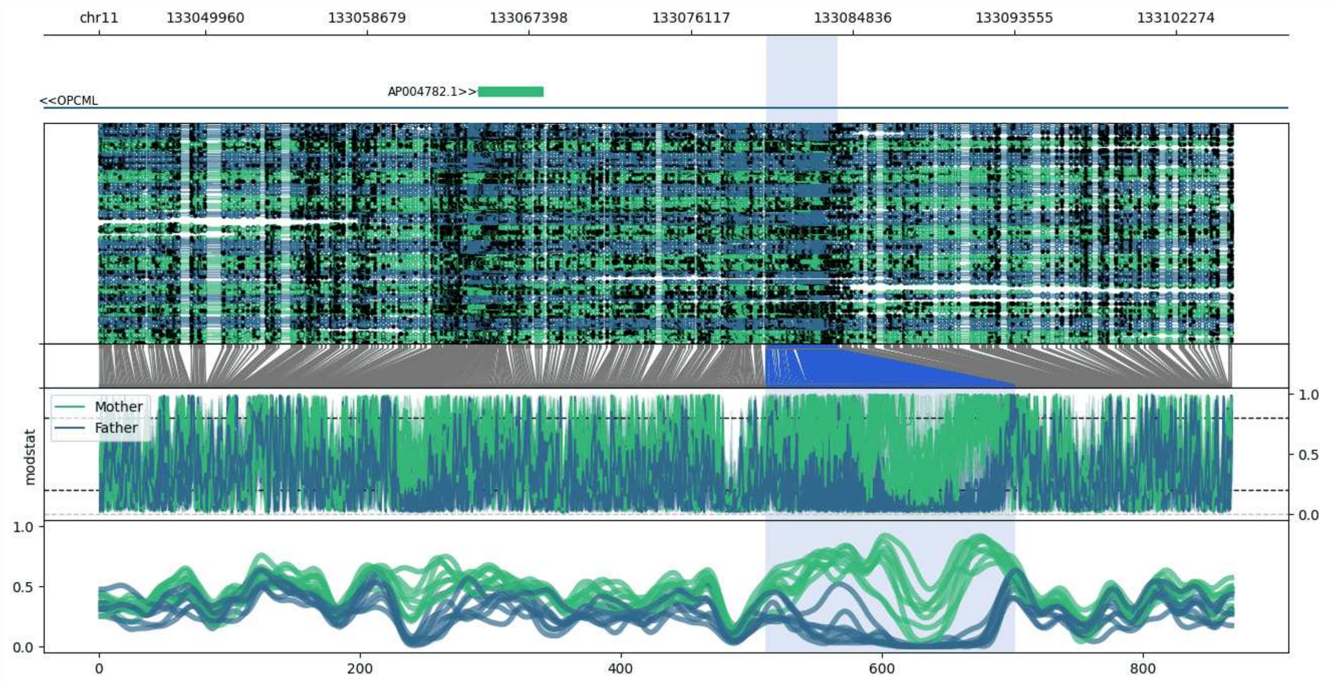


**chr11:133044250-133105285:** Region with 1 DMR containing genes: AP004782.1, OPCML.

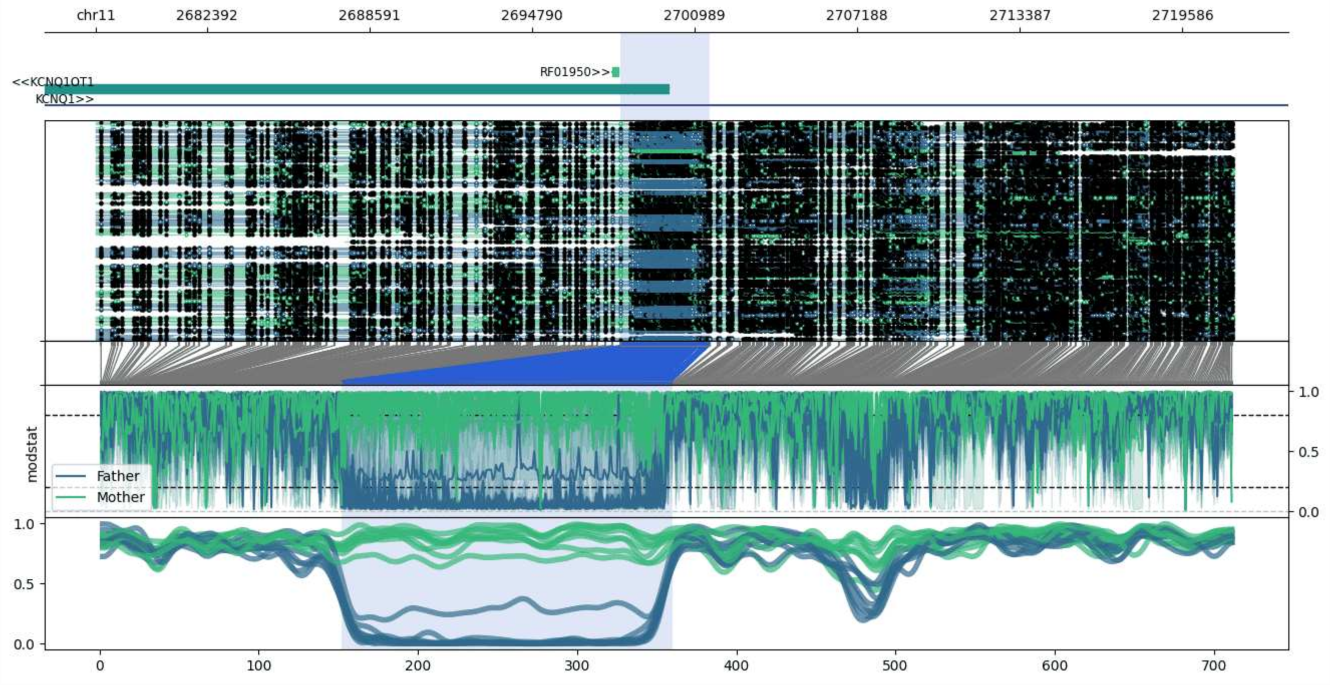


**chr11:2678156-2721554:** Region with 1 DMR containing genes: KCNQ1, KCNQ1OT1, RF01947, RF01948, RF01950.

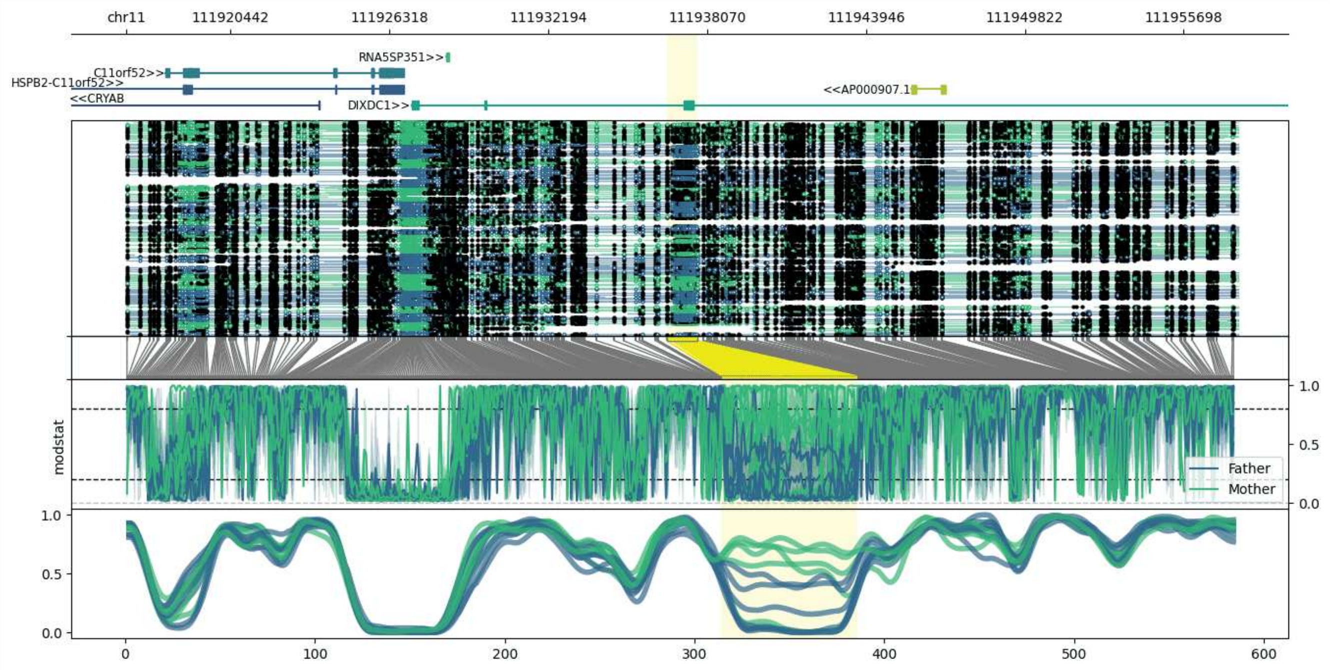


**chr11:111916573-111957706:** Region with 1 DMR containing genes: AP000907.1, C11orf52, CRYAB, DIXDC1, HSPB2, HSPB2-C11orf52, RNA5SP351.

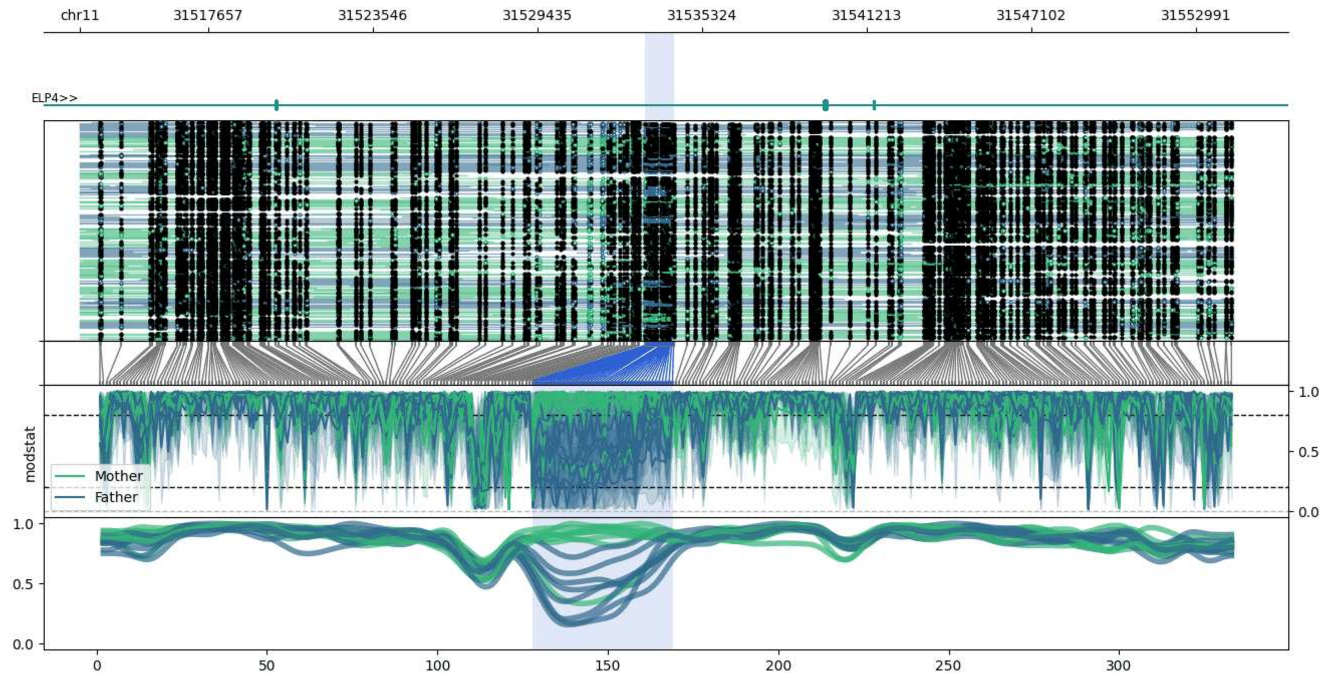


**chr11:31513071-31554300:** Region with 1 DMR containing genes: ELP4, IMMP1L.

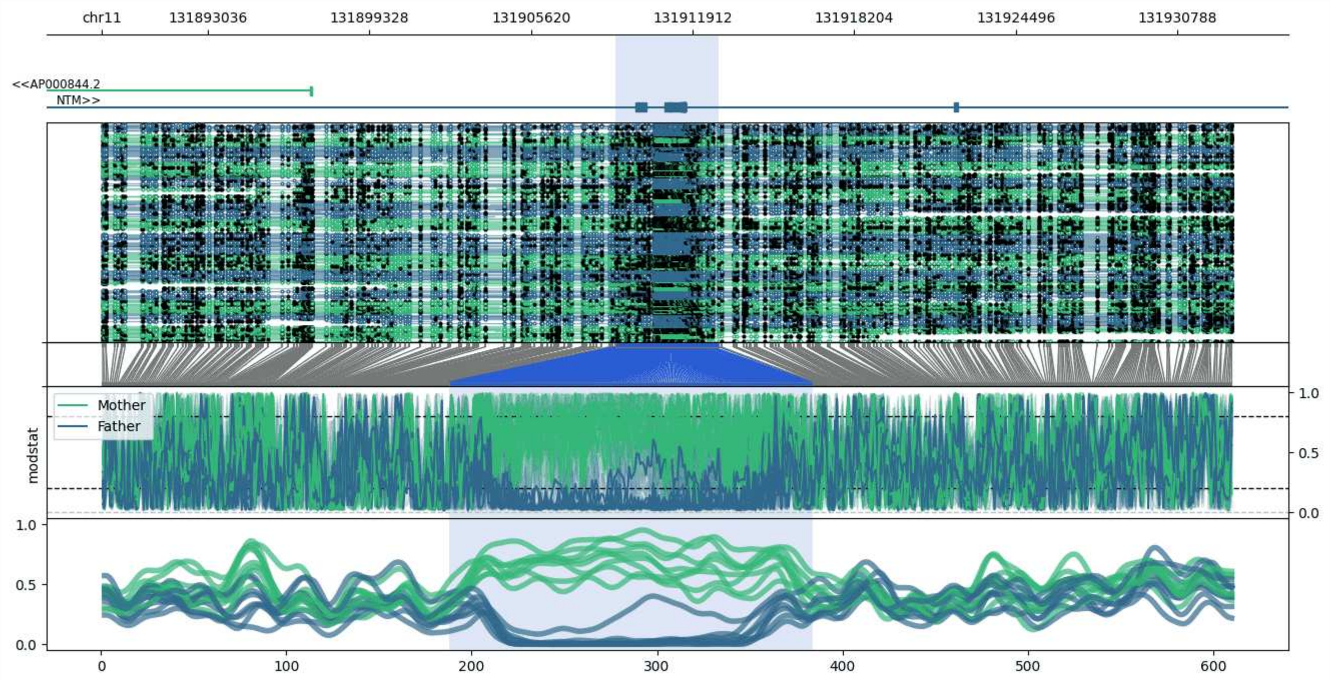


**chr11:131888898-131932943:** Region with 1 DMR containing genes: AP000844.2, NTM.

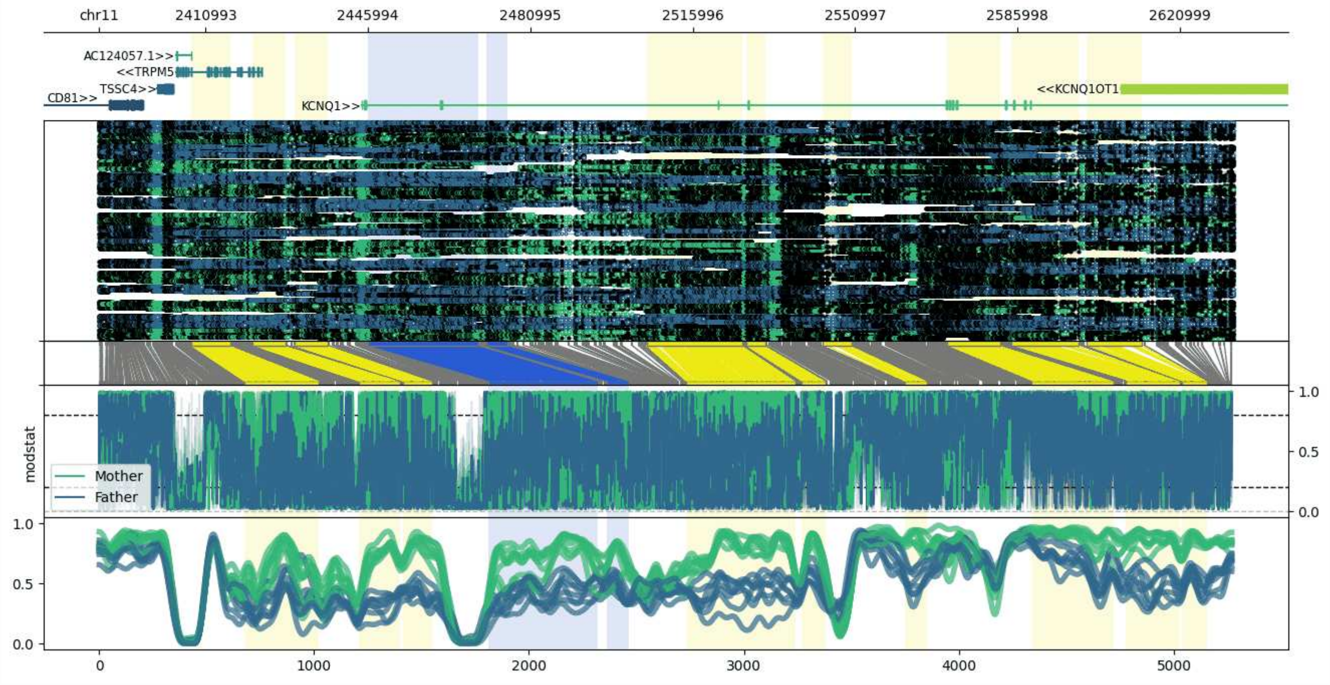


**chr11:2387905-2632912:** Region with 11 DMRs containing genes: AC124057.1, CD81, CD81-AS1, KCNQ1, KCNQ1OT1, TRPM5, TSSC4.

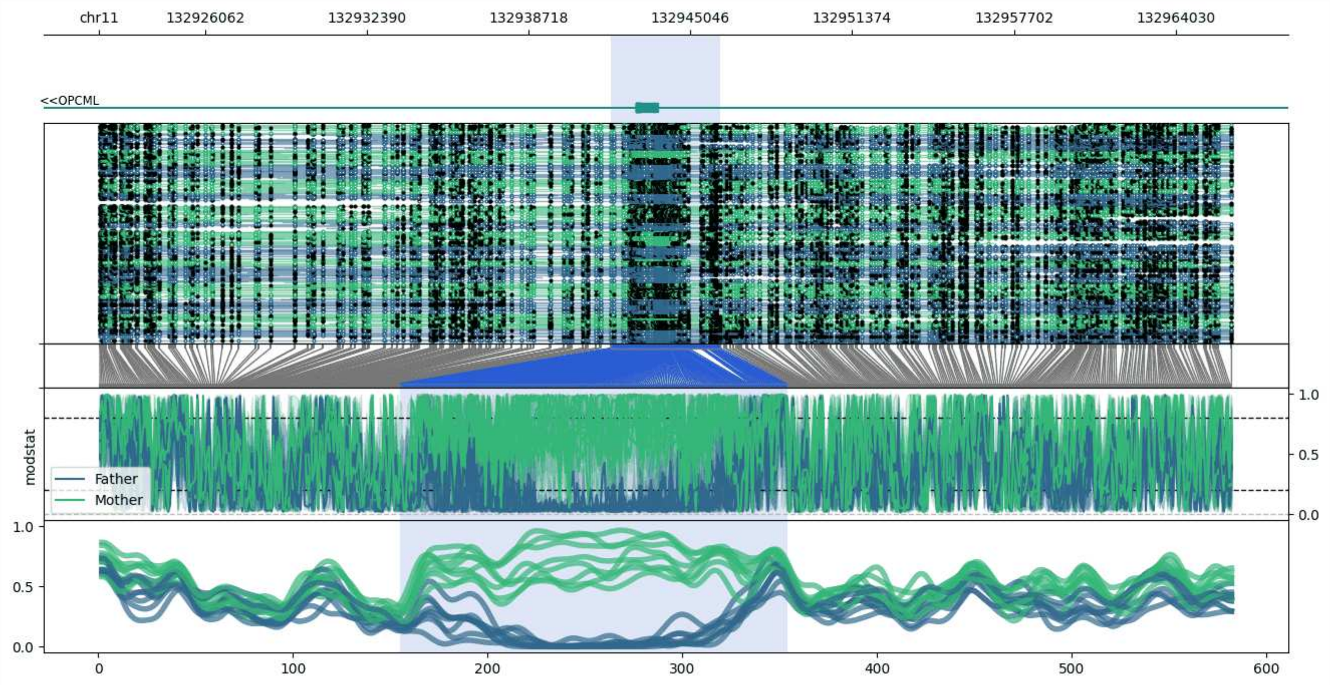


**chr11:132921923-132966225:** Region with 1 DMR containing genes: OPCML.

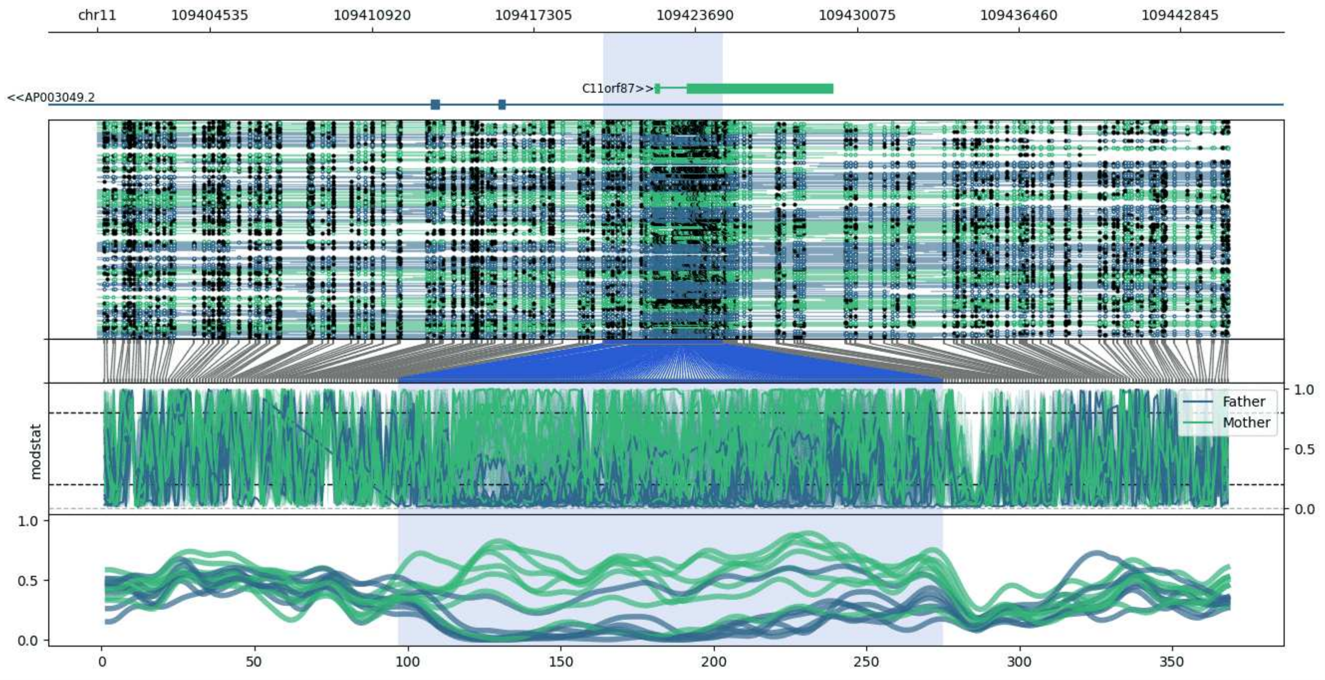


**chr11:109400063-109444764:** Region with 1 DMR containing genes: AP003049.2, C11orf87.

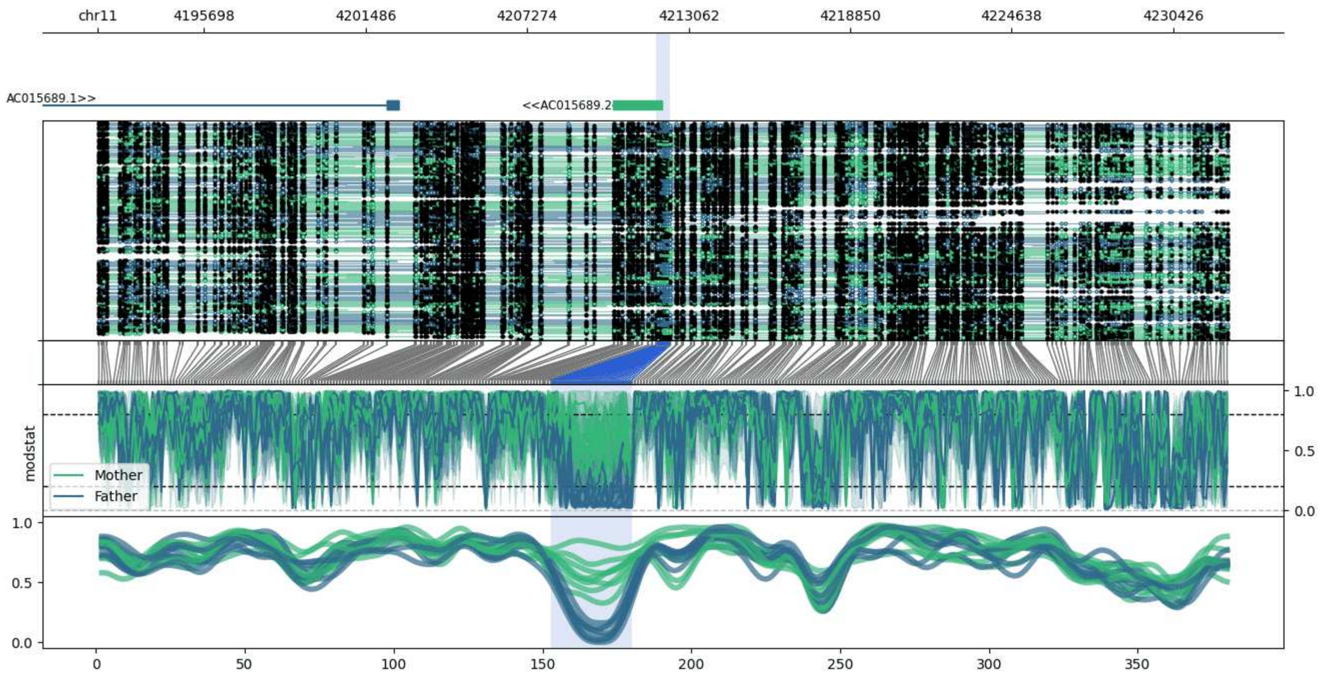


**chr11:4191892-4232408:** Region with 1 DMR containing genes: AC015689.1, AC015689.2, SSU72P2, SSU72P5.

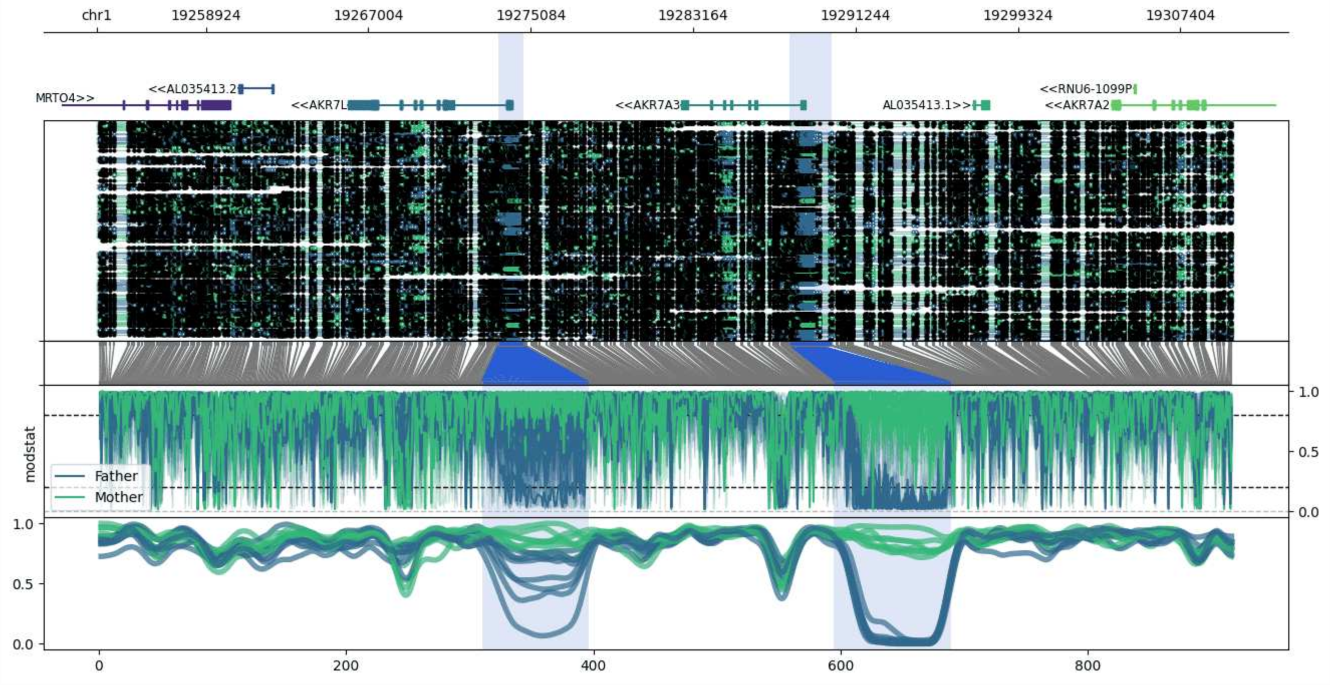


**chr1:19253486-19310050:** Region with 2 DMRs containing genes: AKR7A2, AKR7A3, AKR7L, AL035413.1, AL035413.2, EMC1, MRTO4, PQLC2, RNU6-1099P.

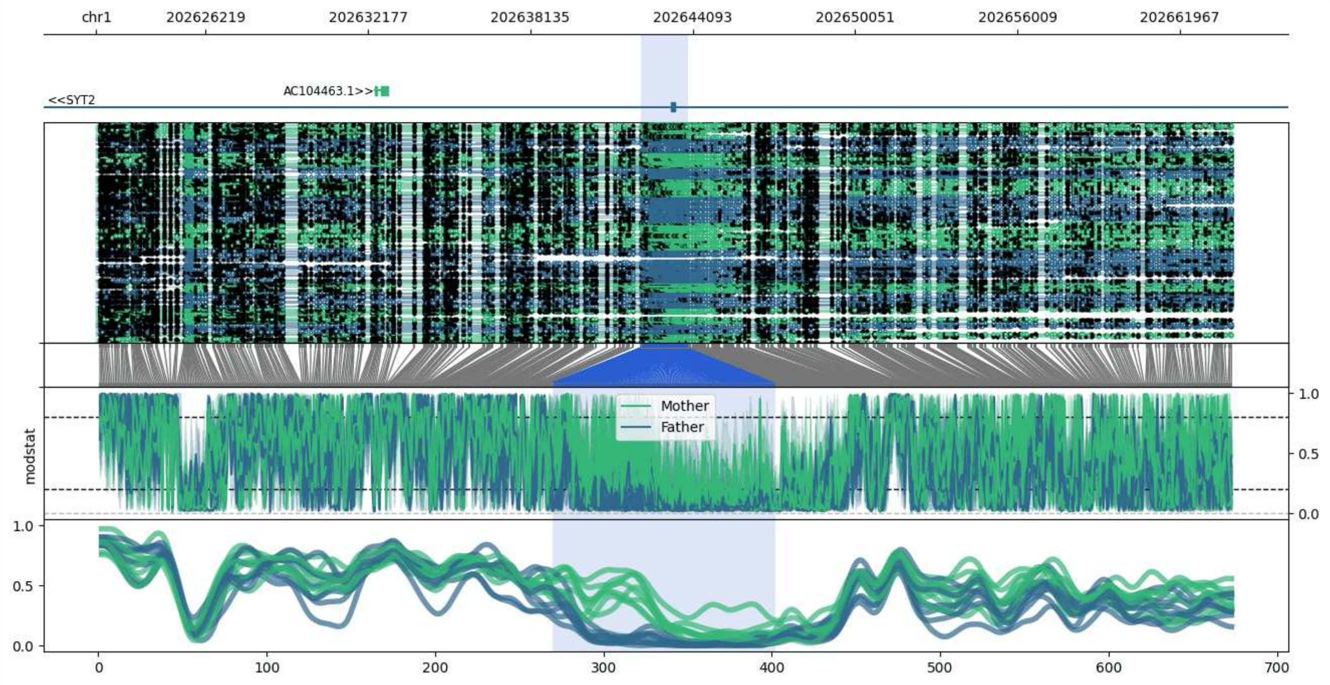


**chr1:202622201-202663909:** Region with 1 DMR containing genes: AC104463.1, SYT2.

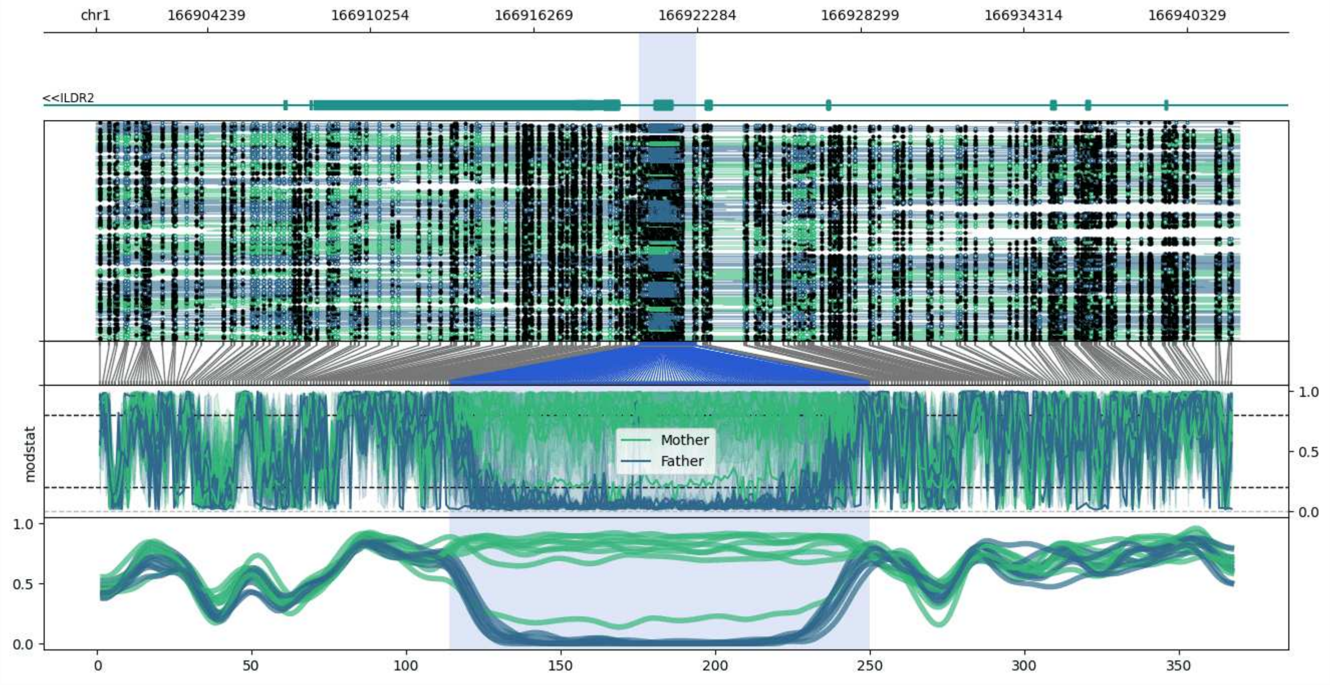


**chr1:166900144-166942253:** Region with 1 DMR containing genes: ILDR2.

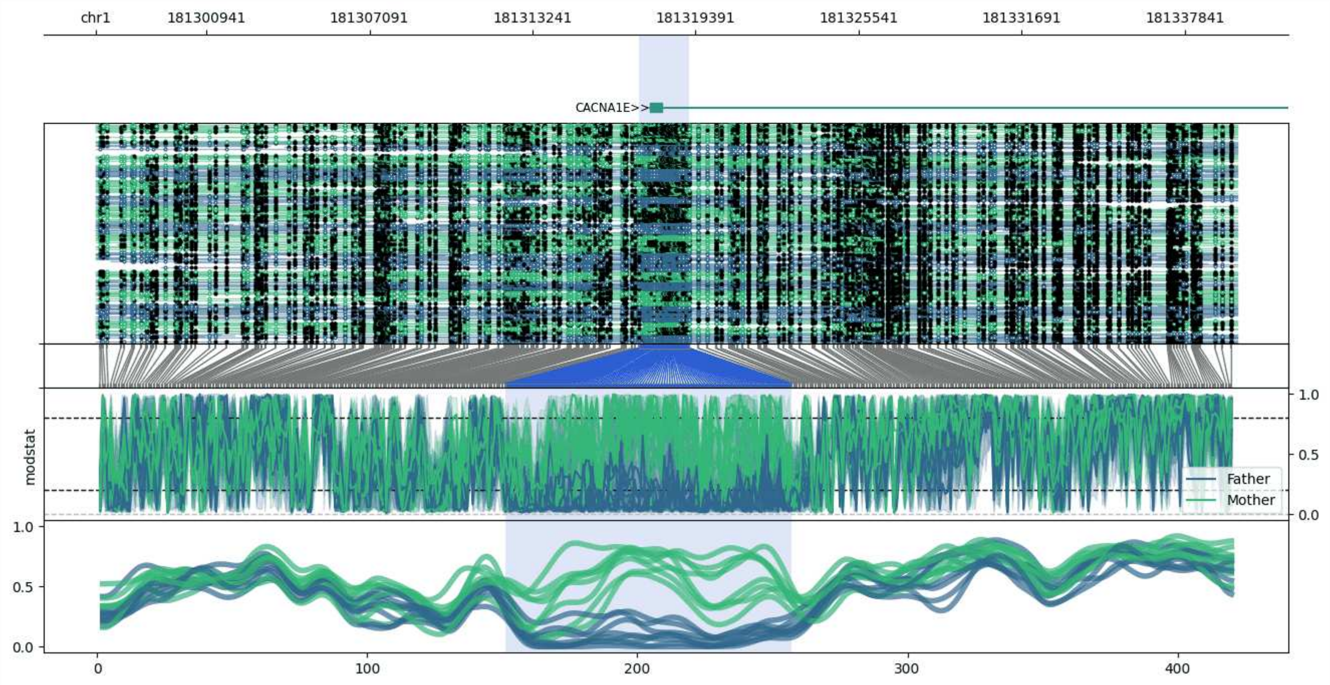


**chr1:181296778-181339828:** Region with 1 DMR containing genes: CACNA1E.

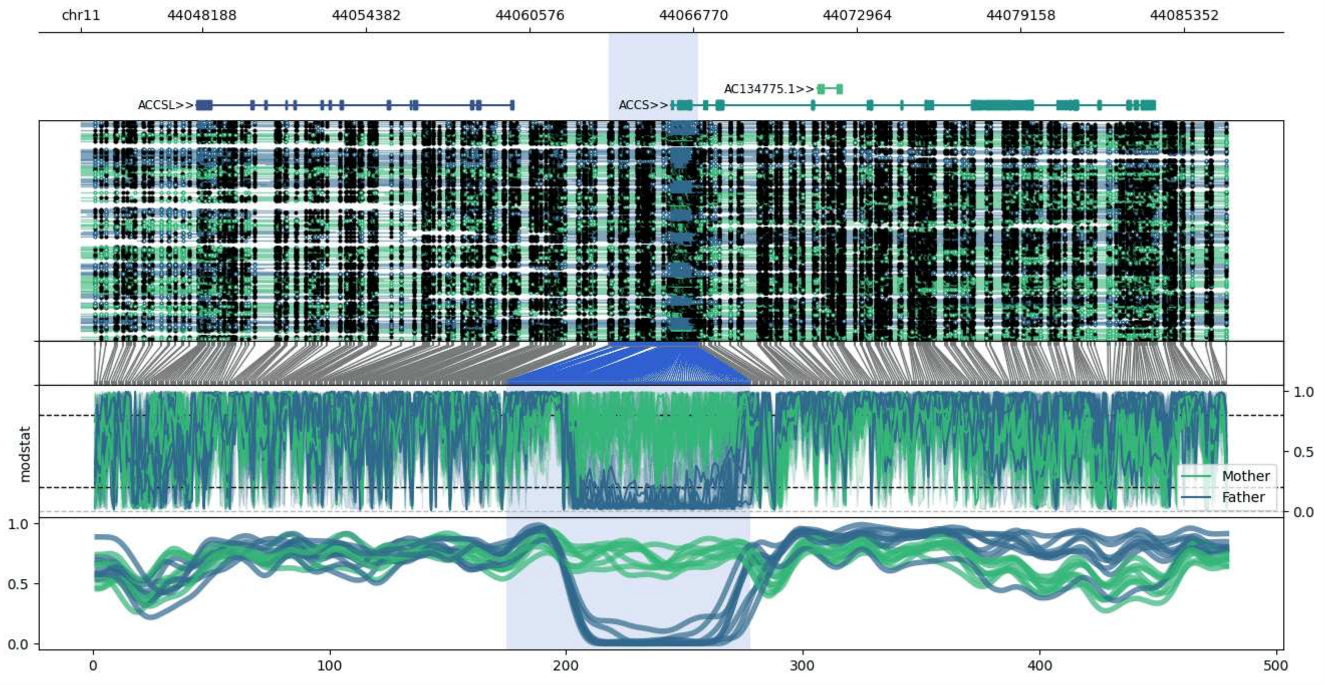


**chr11:44043594-44086953:** Region with 1 DMR containing genes: AC134775.1, ACCS, ACCSL, EXT2.

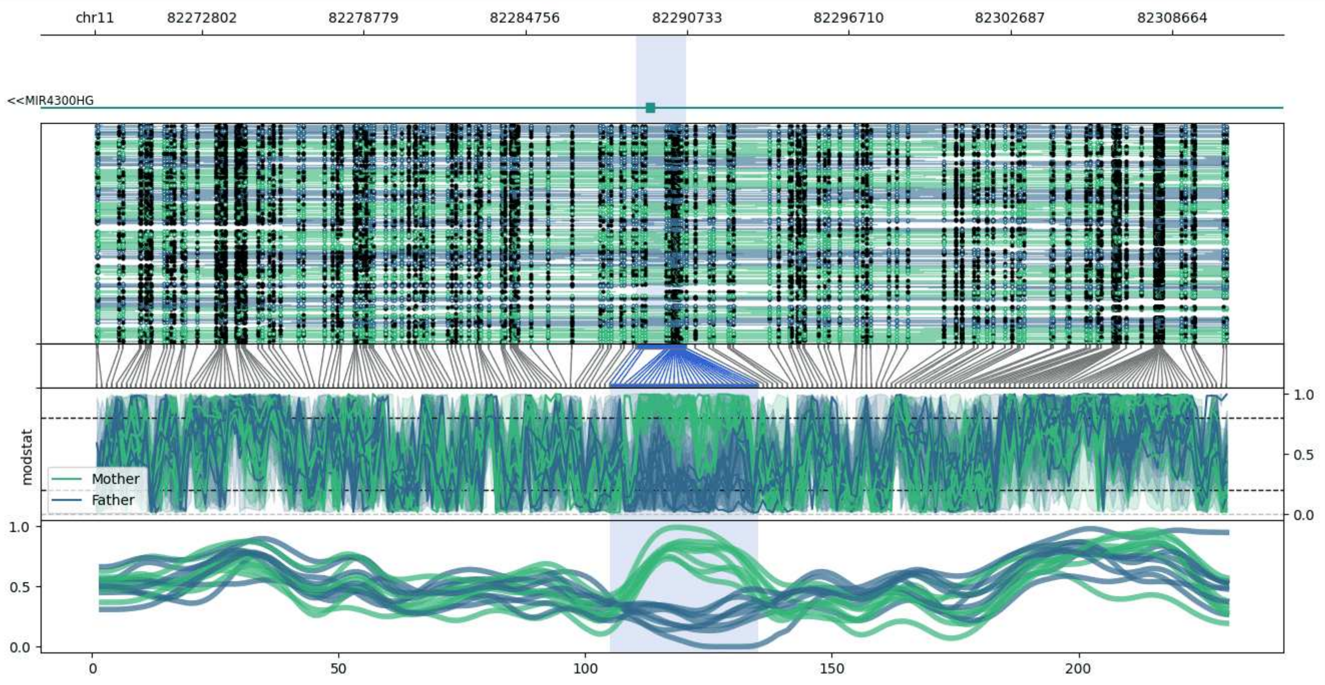


**chr11:82268855-82310696:** Region with 1 DMR containing genes: MIR4300HG.

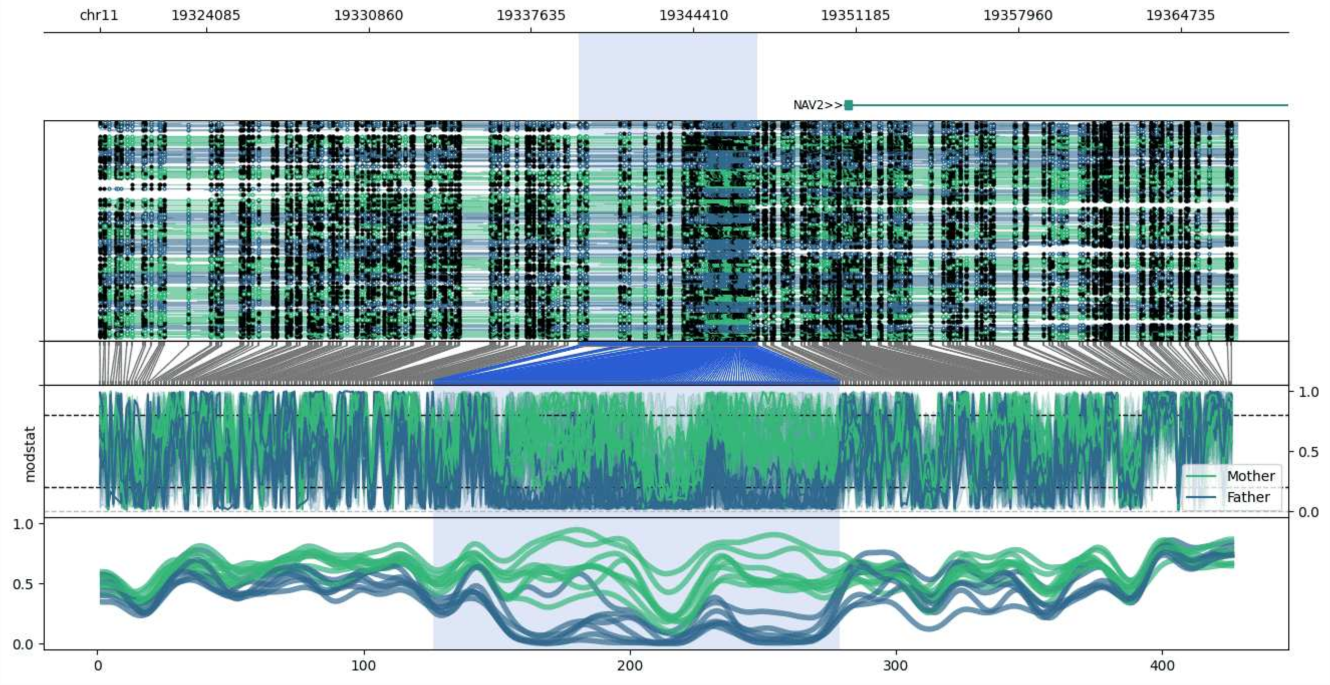


**chr11:19319645-19367072:** Region with 1 DMR containing genes: NAV2.

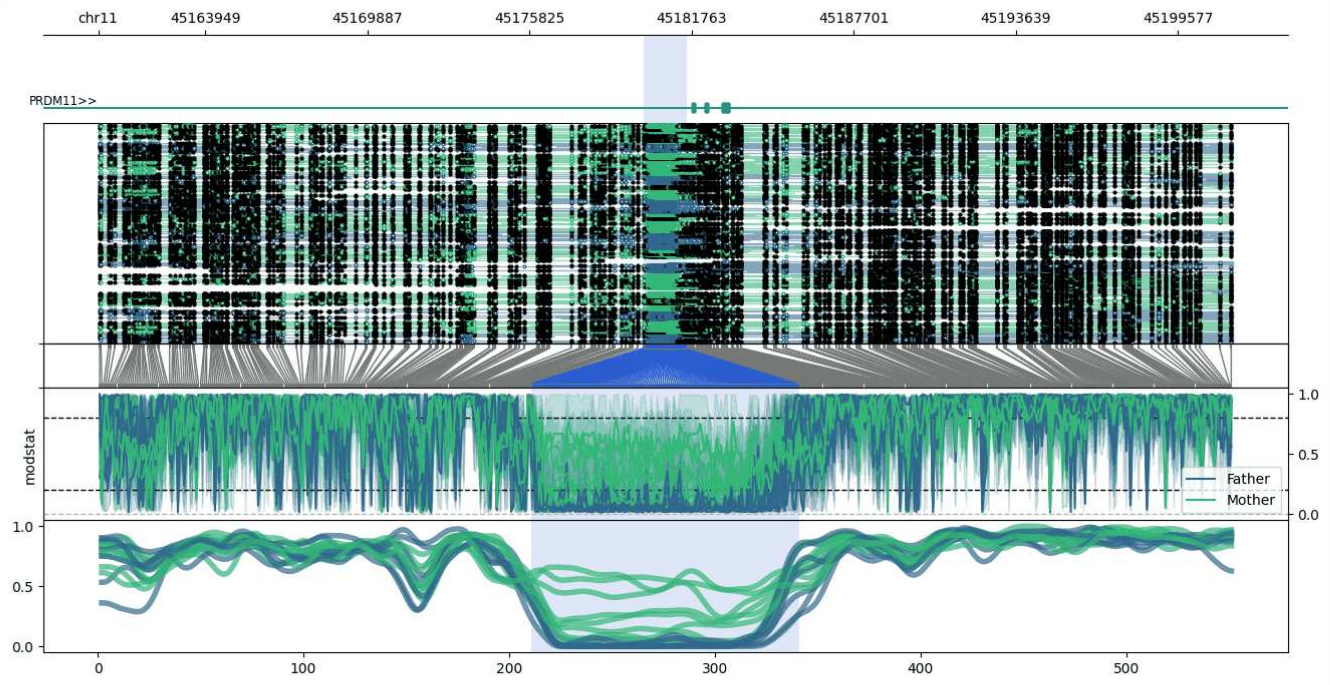


**chr11:45160025-45201593:** Region with 1 DMR containing genes: PRDM11.

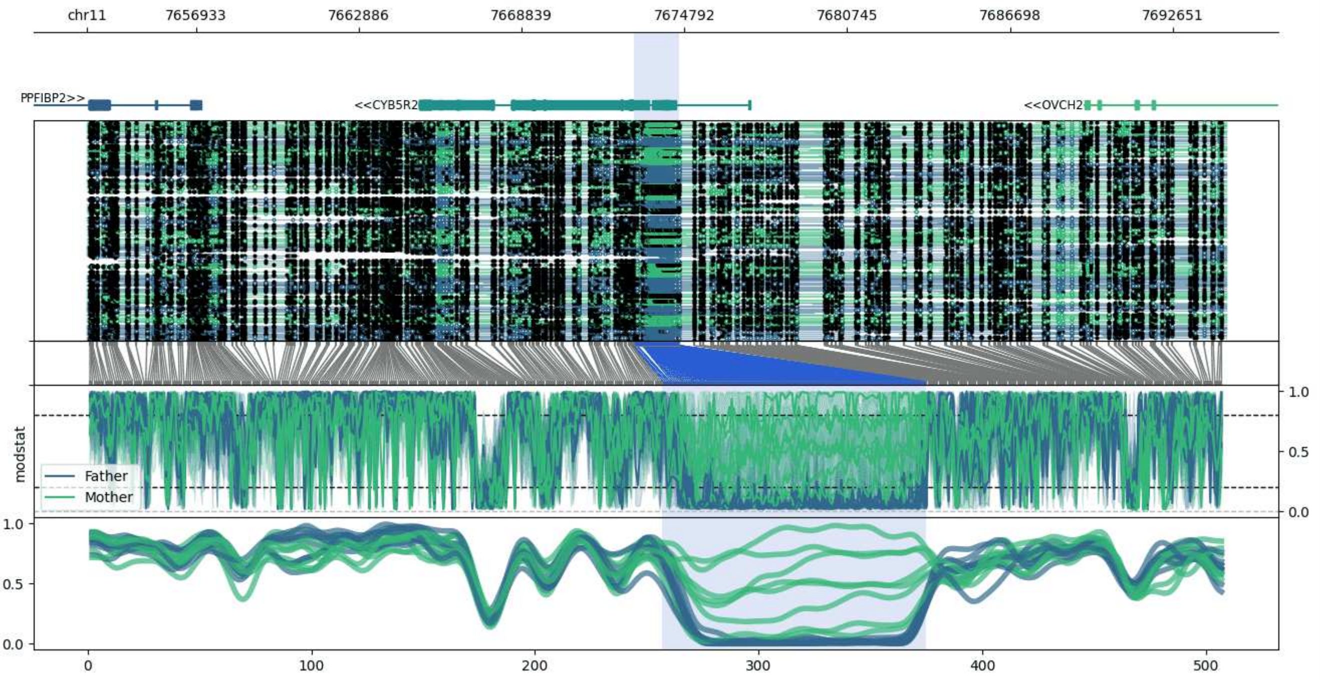


**chr11:7652931-7694605:** Region with 1 DMR containing genes: AC104237.2, AC104237.3, CYB5R2, OVCH2, PPFIBP2.

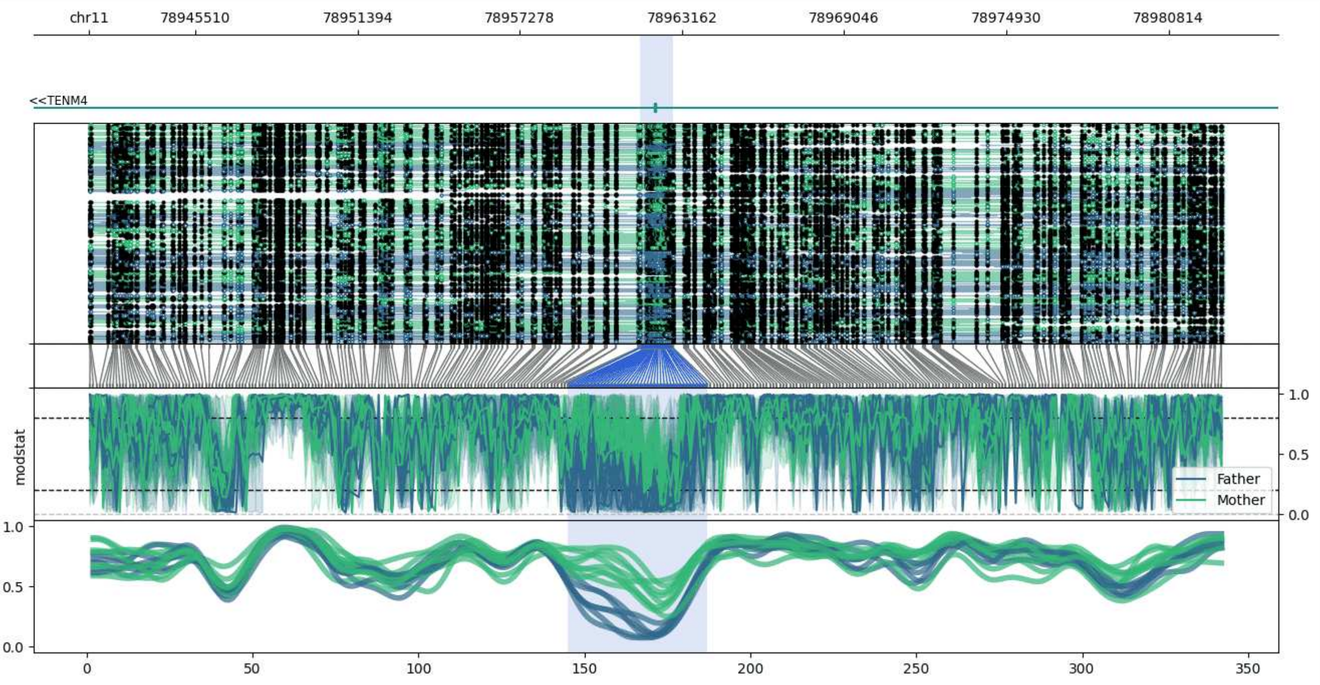


**chr11:78941631-78982825:** Region with 1 DMR containing genes: TENM4.

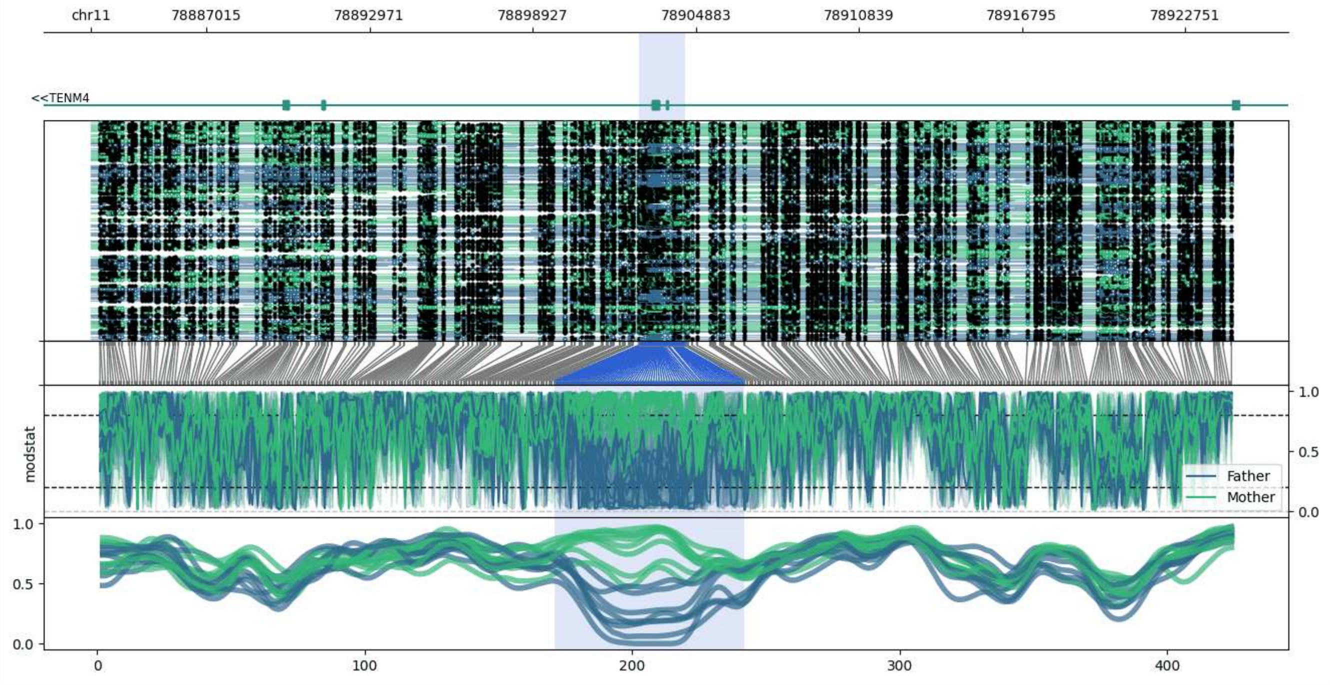


**chr11:78882795-78924488:** Region with 1 DMR containing genes: TENM4.

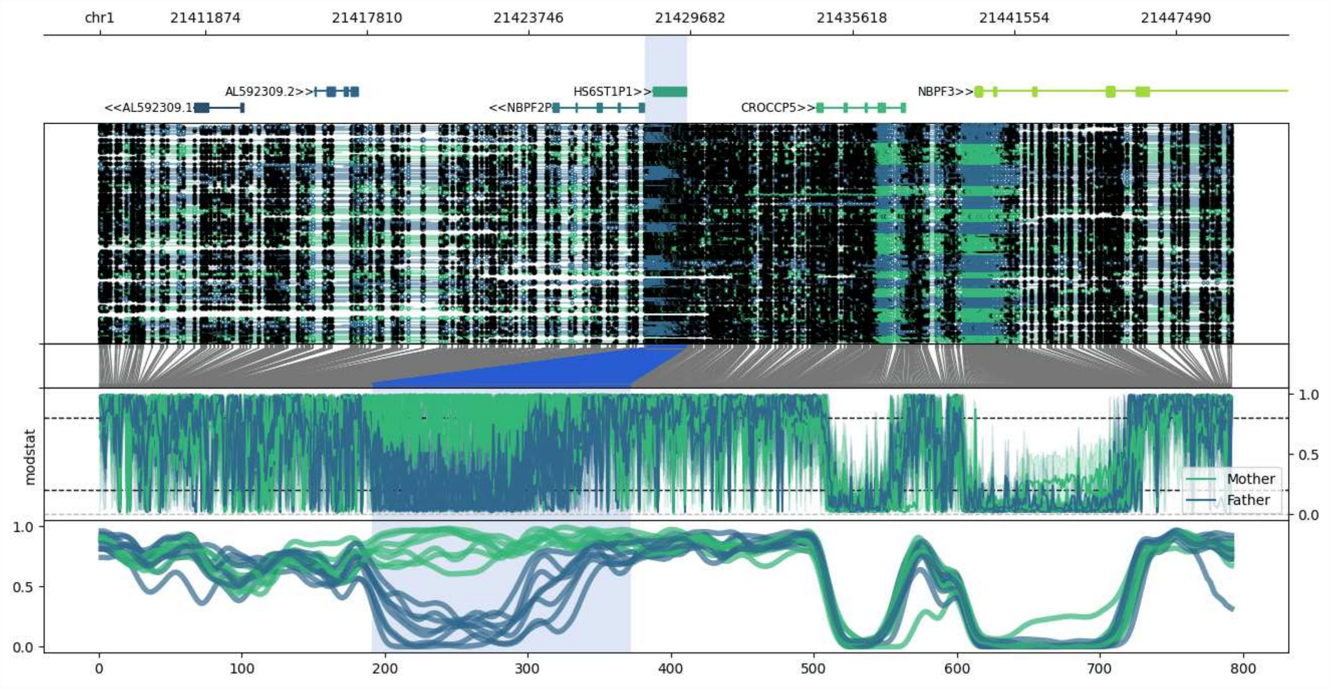


**chr1:21407995-21449552:** Region with 1 DMR containing genes: AL592309.1, AL592309.2, CROCCP5, HS6ST1P1, NBPF2P, NBPF3, PPP1R11P1.

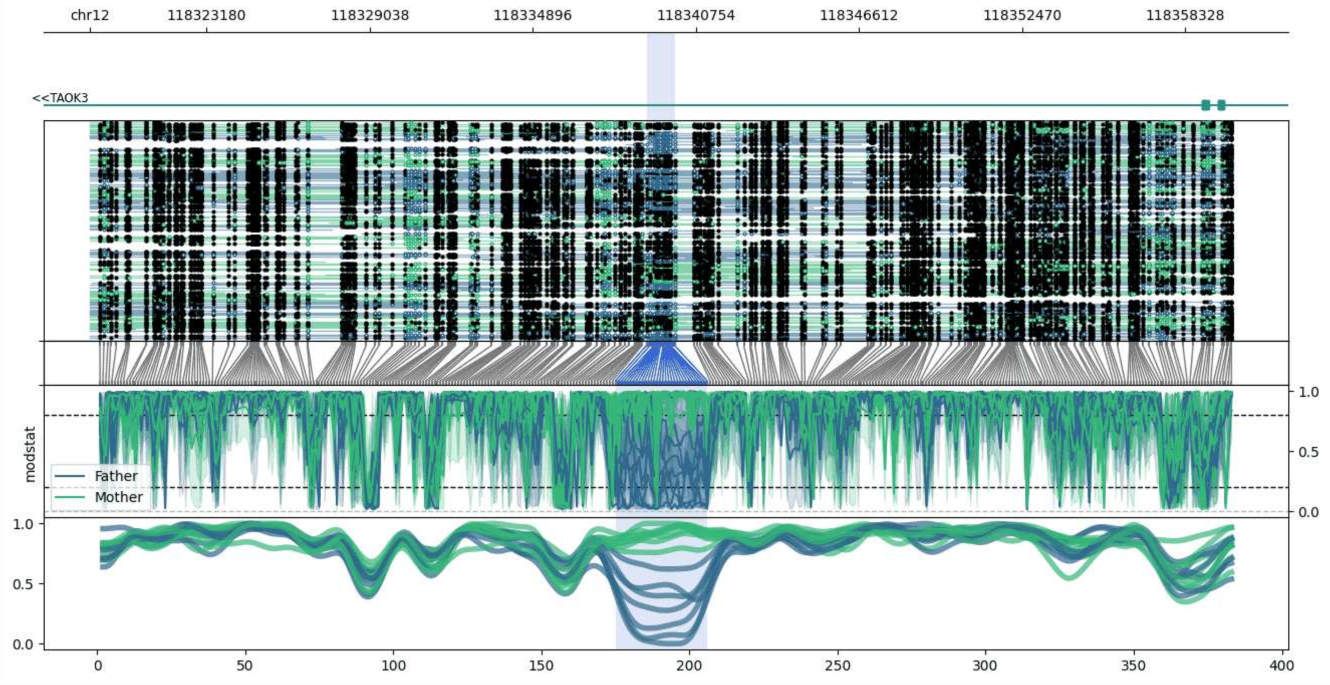


**chr12:118318984-118359991:** Region with 1 DMR containing genes: TAOK3.

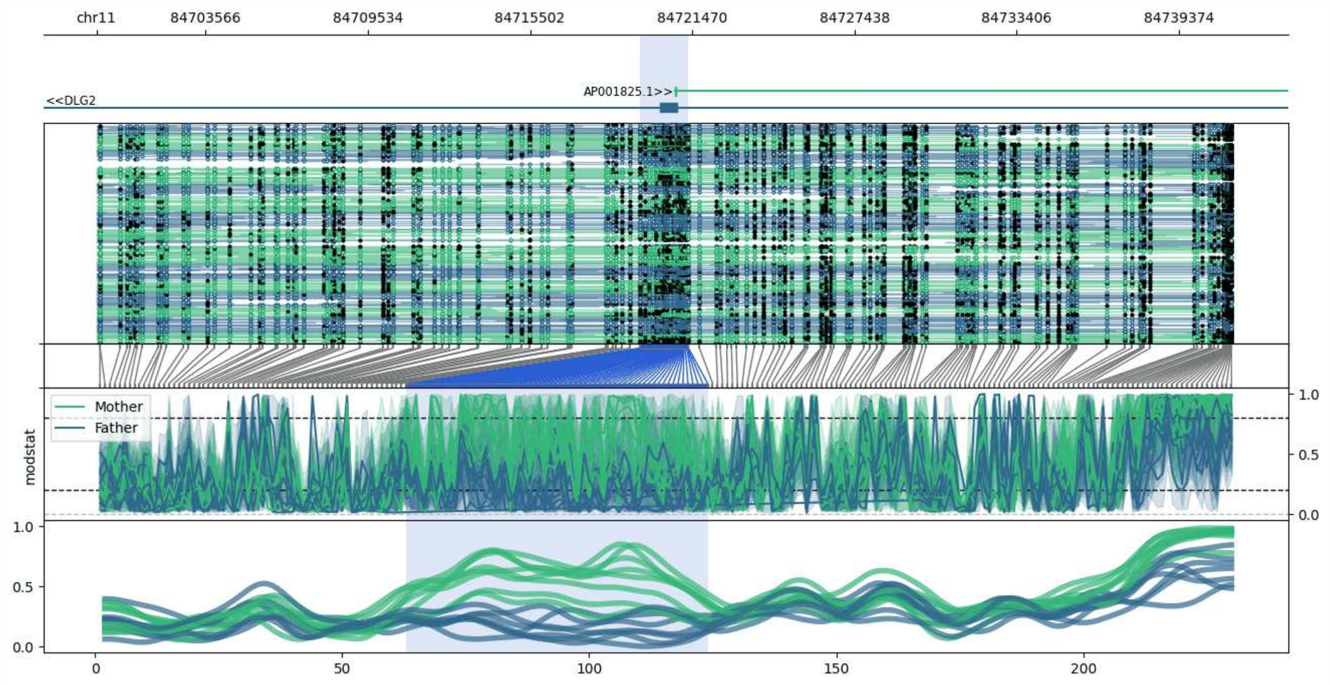


**chr11:84699550-84741331:** Region with 1 DMR containing genes: AP001825.1, DLG2.

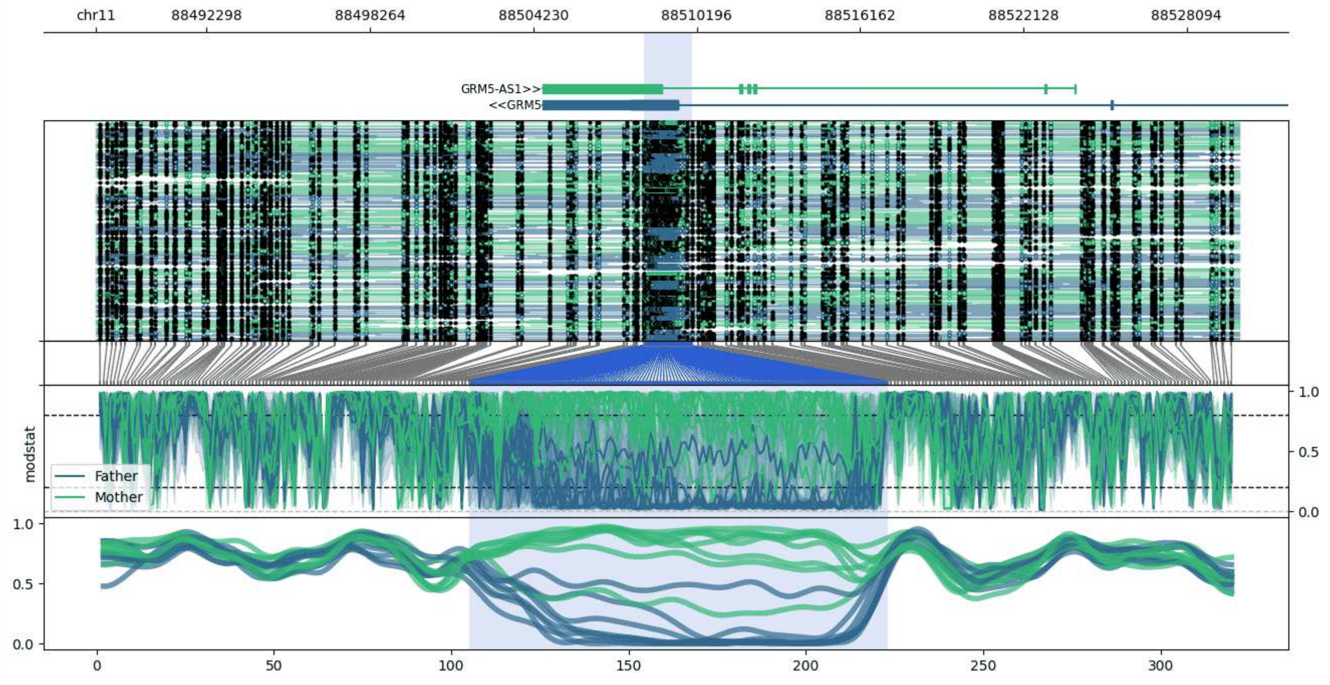


**chr11:88488256-88530020:** Region with 1 DMR containing genes: GRM5, GRM5-AS1.

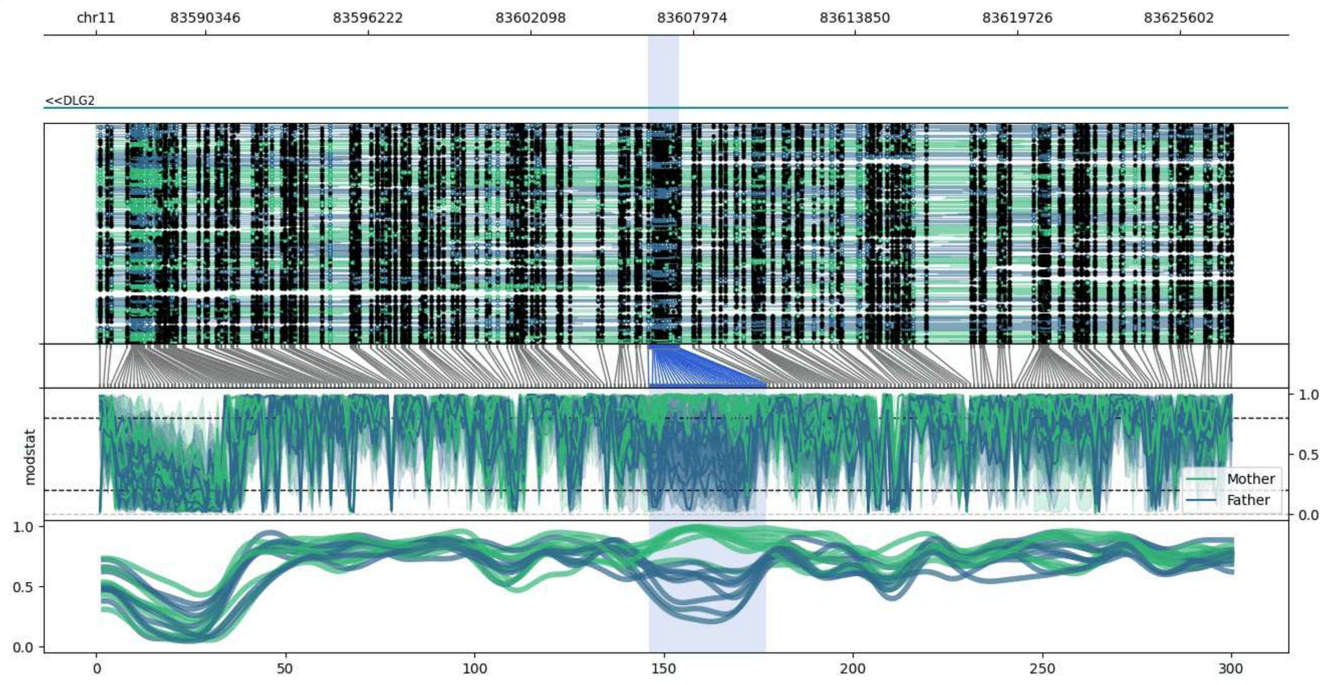


**chr11:83586360-83627497:** Region with 1 DMR containing genes: DLG2.

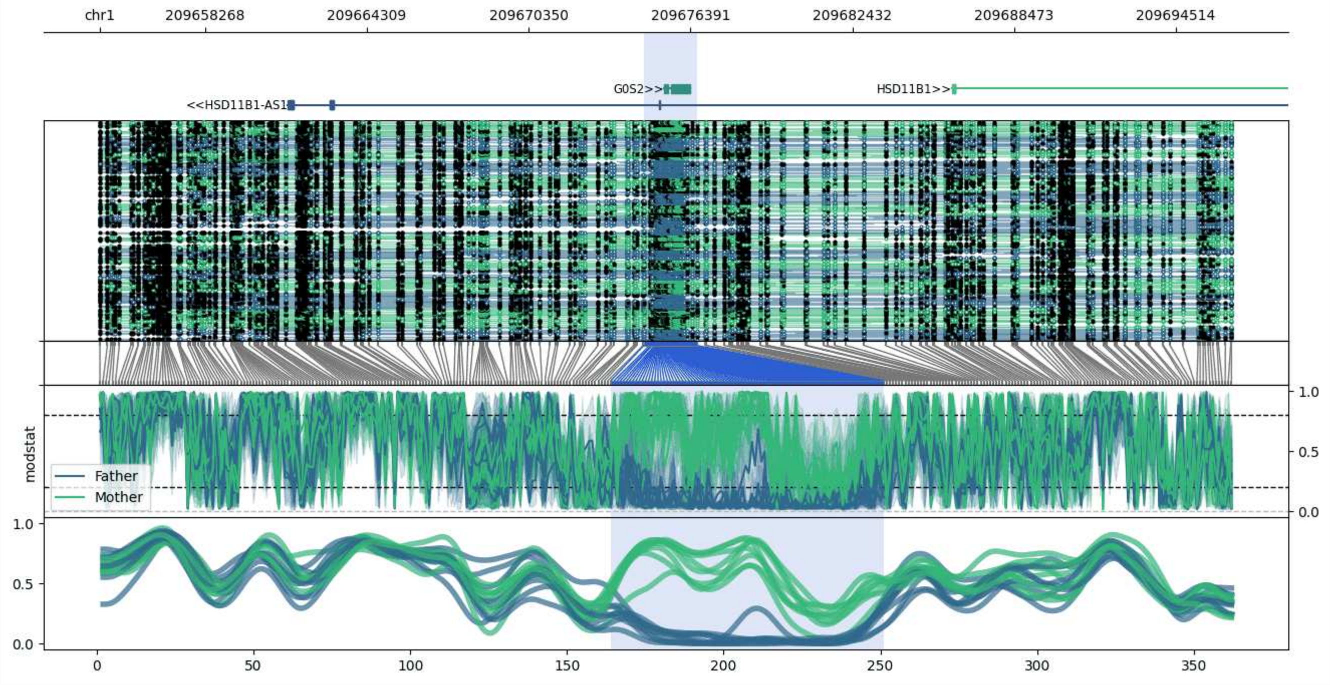


**chr1:209654335-209696625:** Region with 1 DMR containing genes: G0S2, HSD11B1, HSD11B1-AS1, LAMB3.

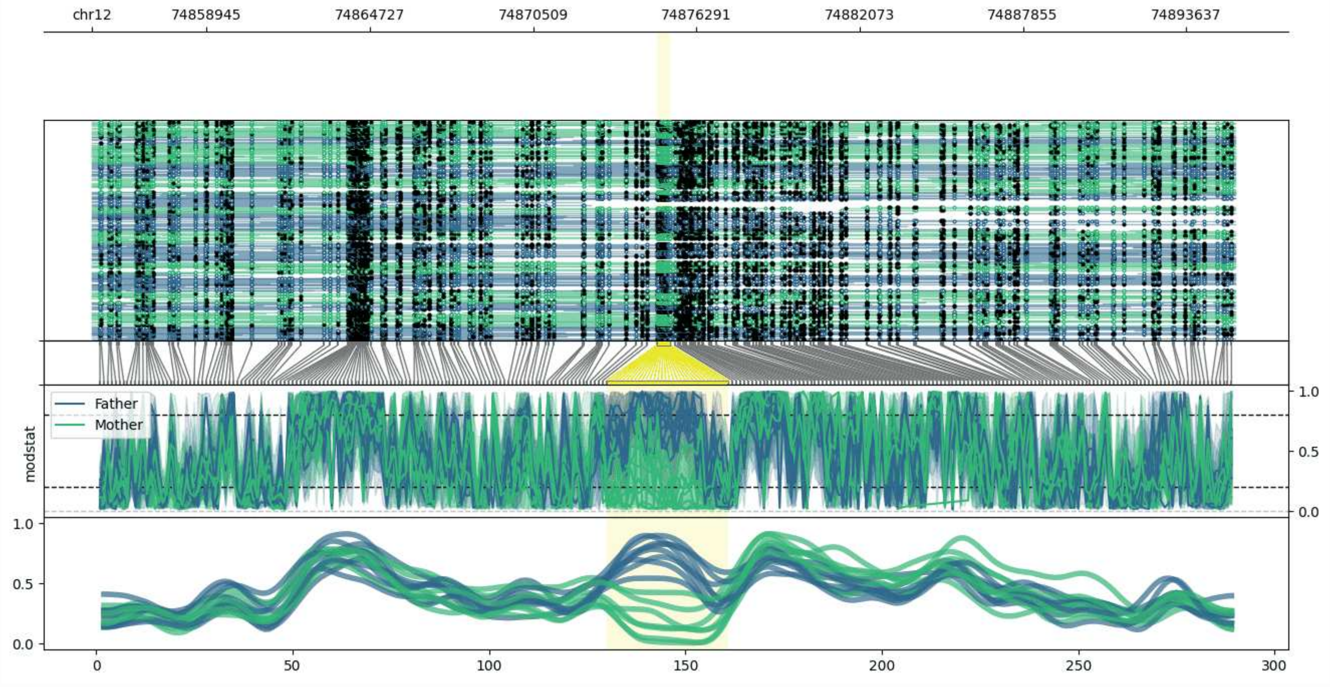


**chr12:74854888-74895363:** Region with 1 DMR containing no genes.

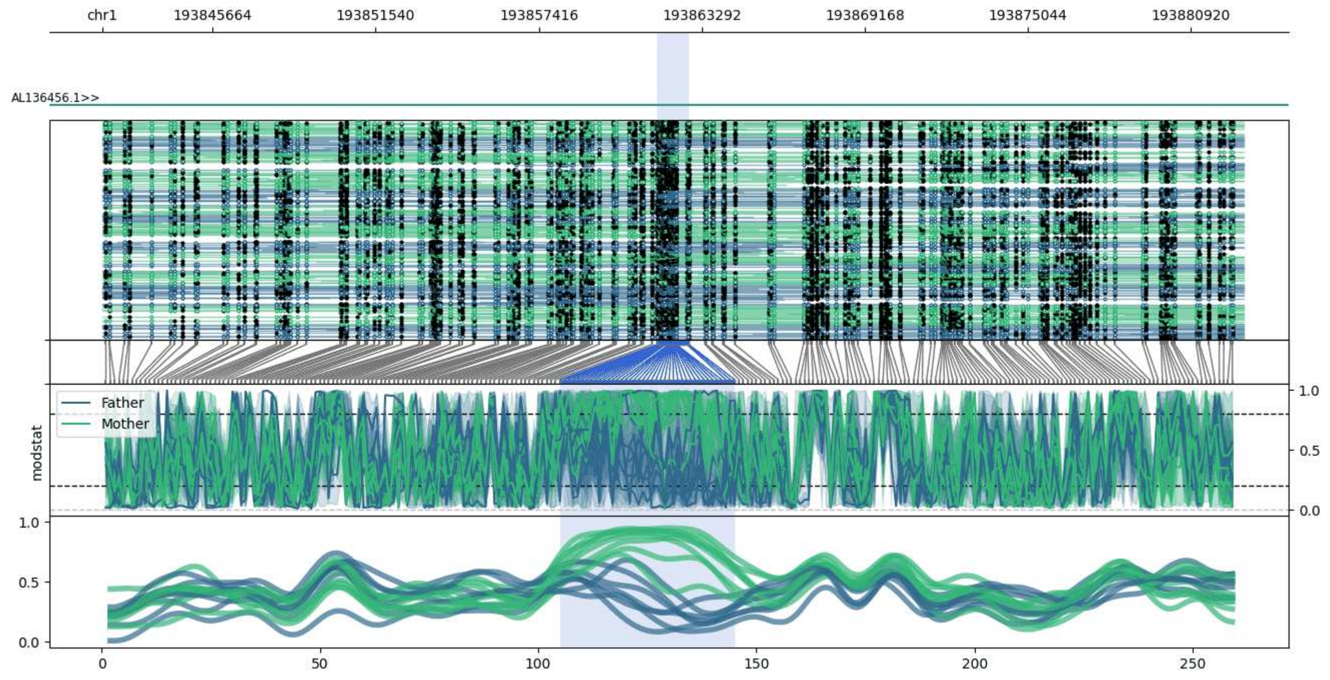


**chr1:193841683-193882821:** Region with 1 DMR containing genes: AL136456.1.

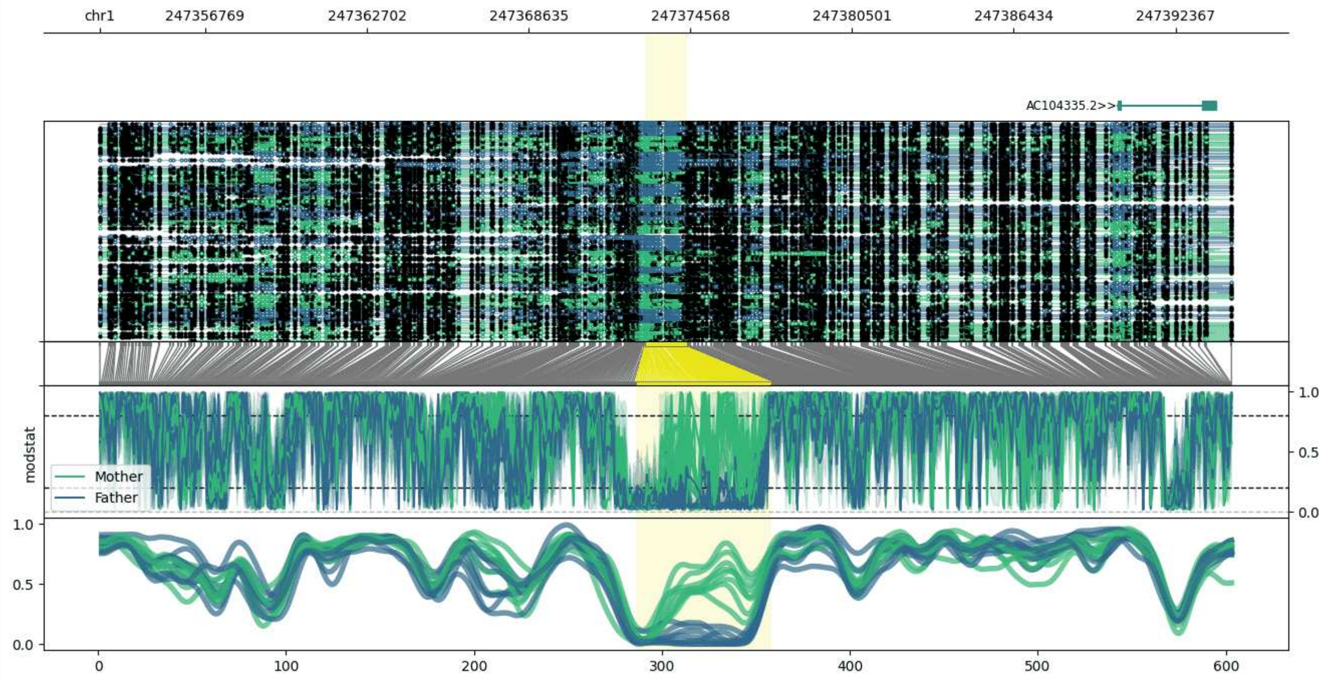


**chr1:247352909-247394443:** Region with 1 DMR containing genes: AC104335.1, AC104335.2.

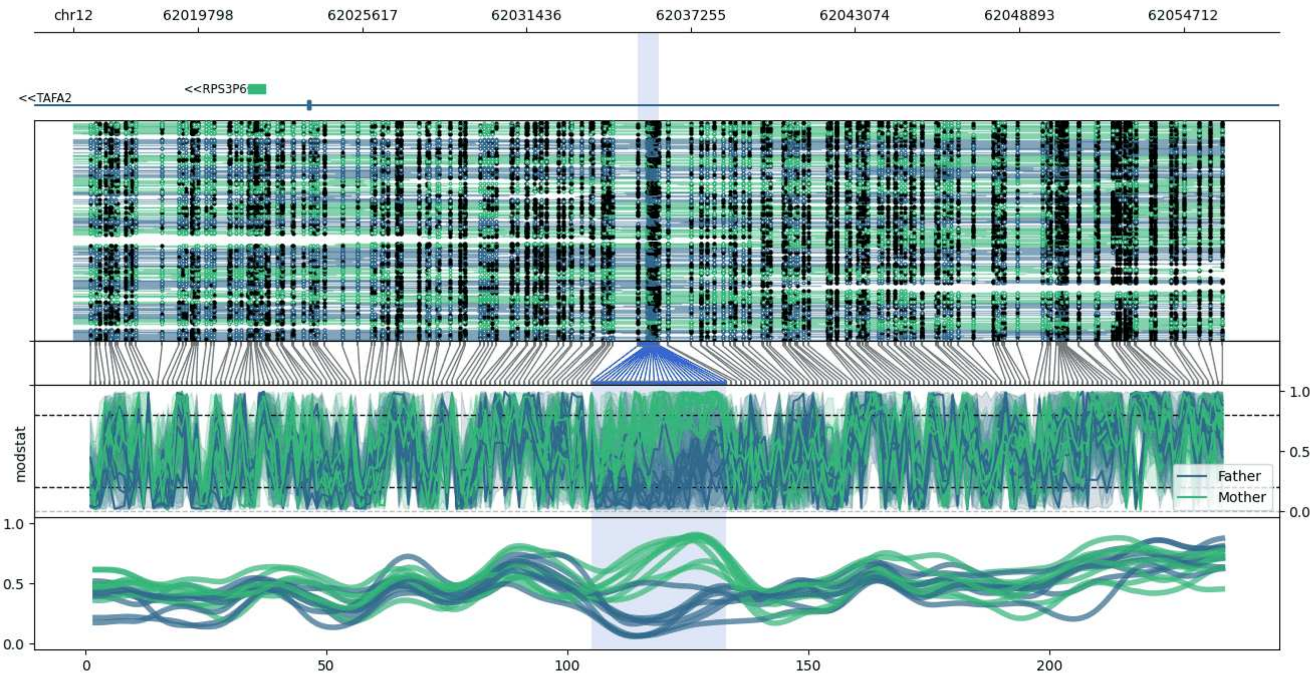


**chr12:62015389-62056128:** Region with 1 DMR containing genes: RPS3P6, TAFA2.

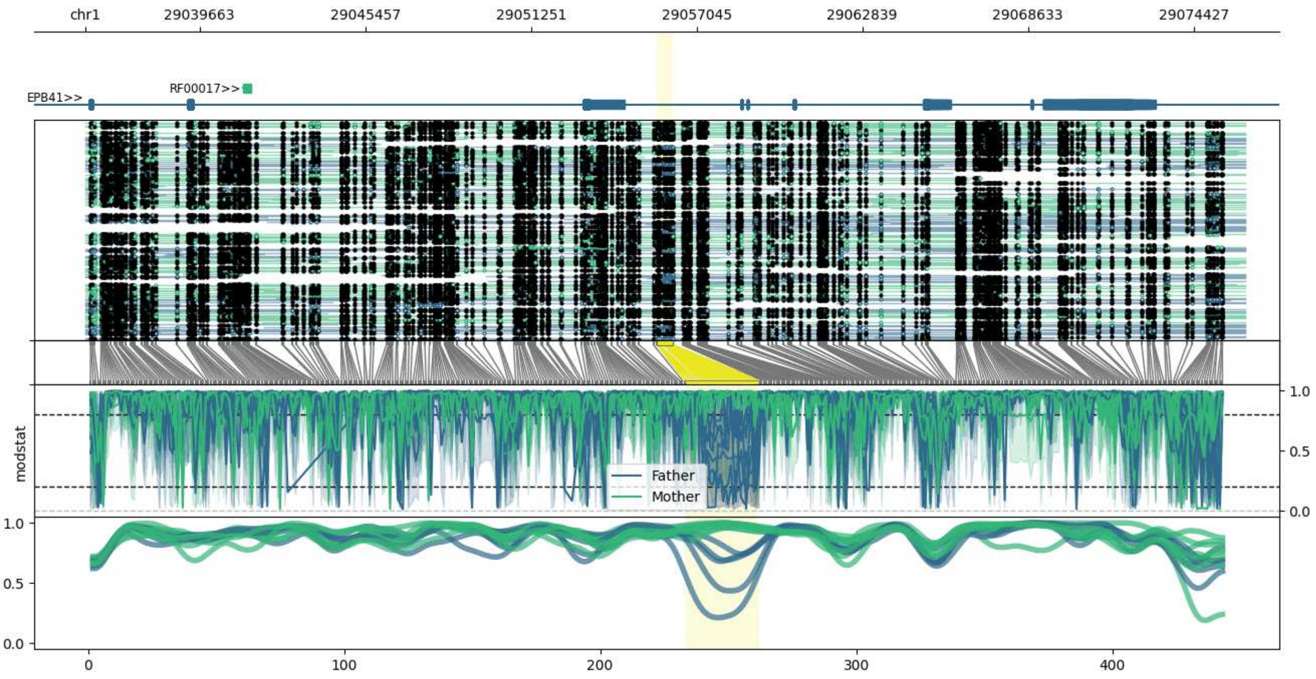


**chr1:29035667-29076227:** Region with 1 DMR containing genes: EPB41, RF00017.

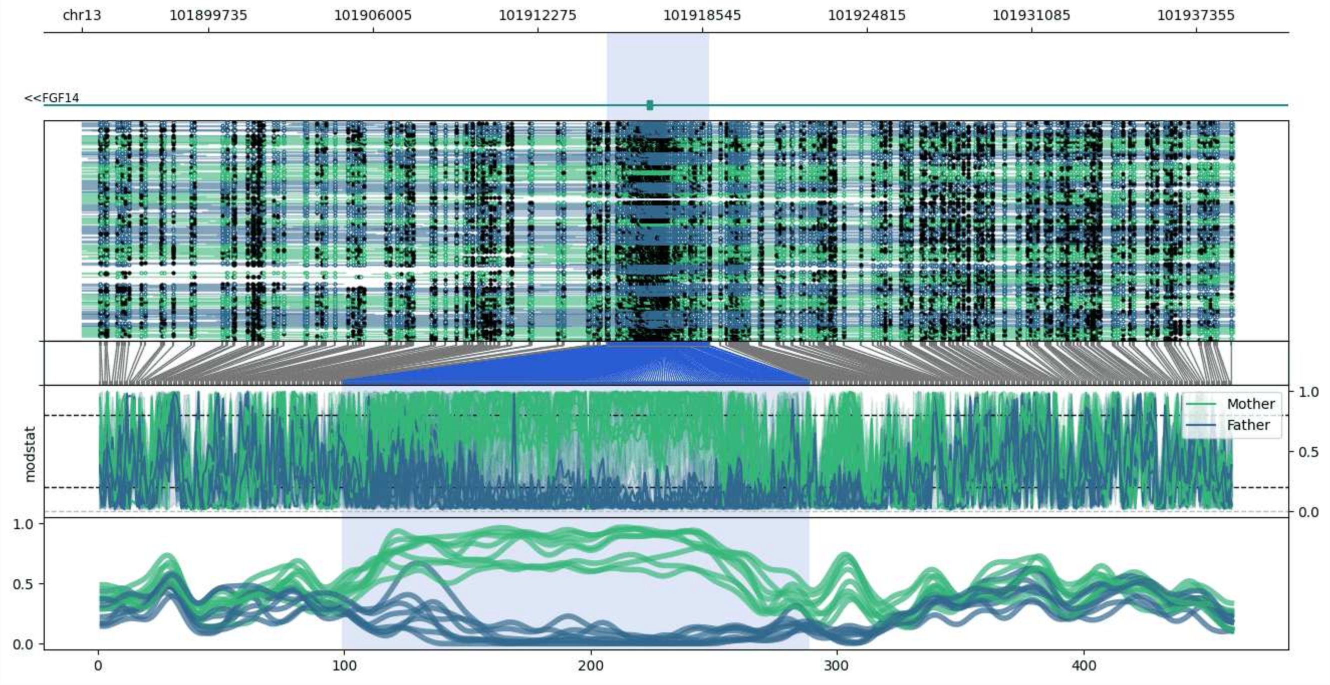


**chr13:101894916-101938812:** Region with 1 DMR containing genes: FGF14.

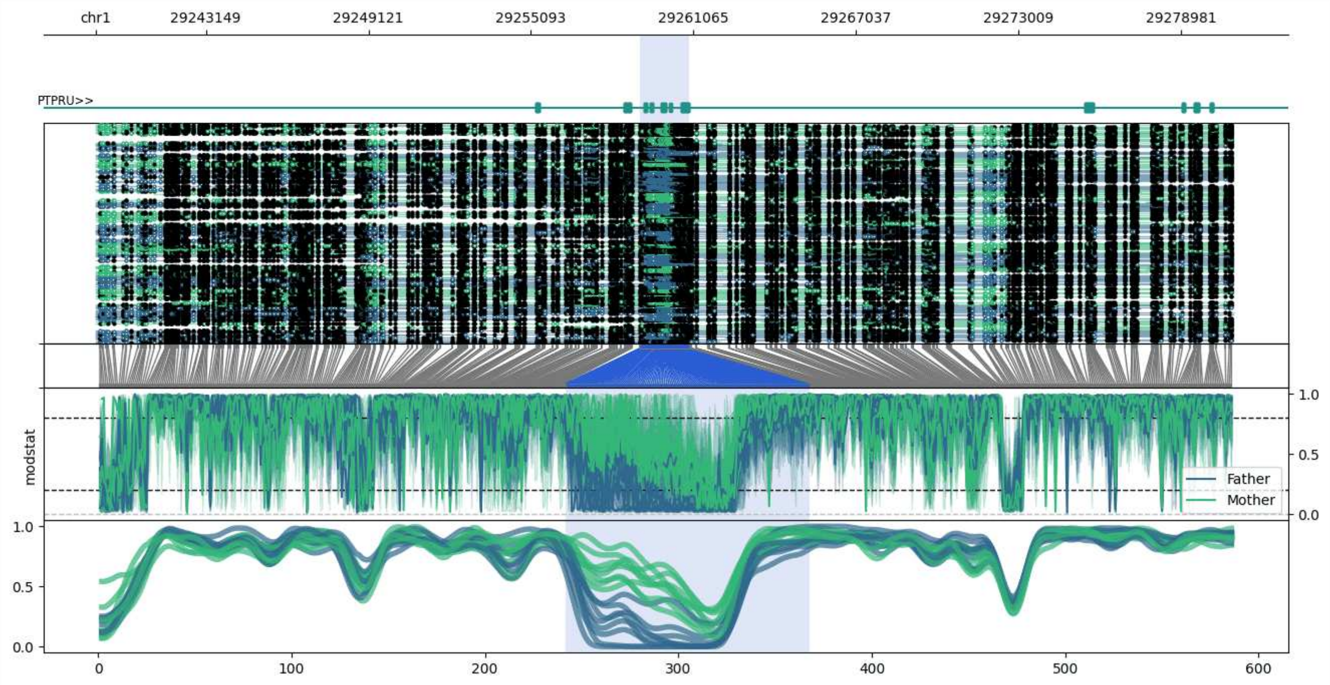


**chr1:29239091-29280898:** Region with 1 DMR containing genes: MECR, PTPRU.

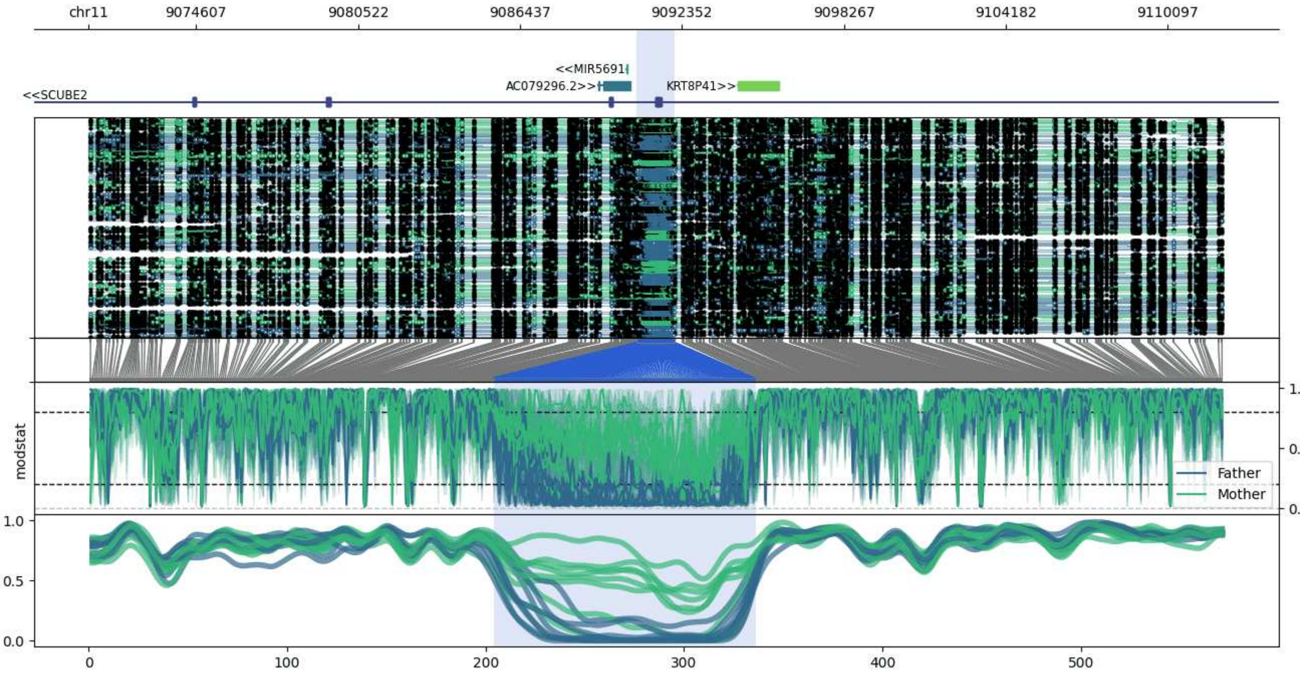


**chr11:9070692-9112102:** Region with 1 DMR containing genes: AC079296.1, AC079296.2, KRT8P41, MIR5691, SCUBE2.

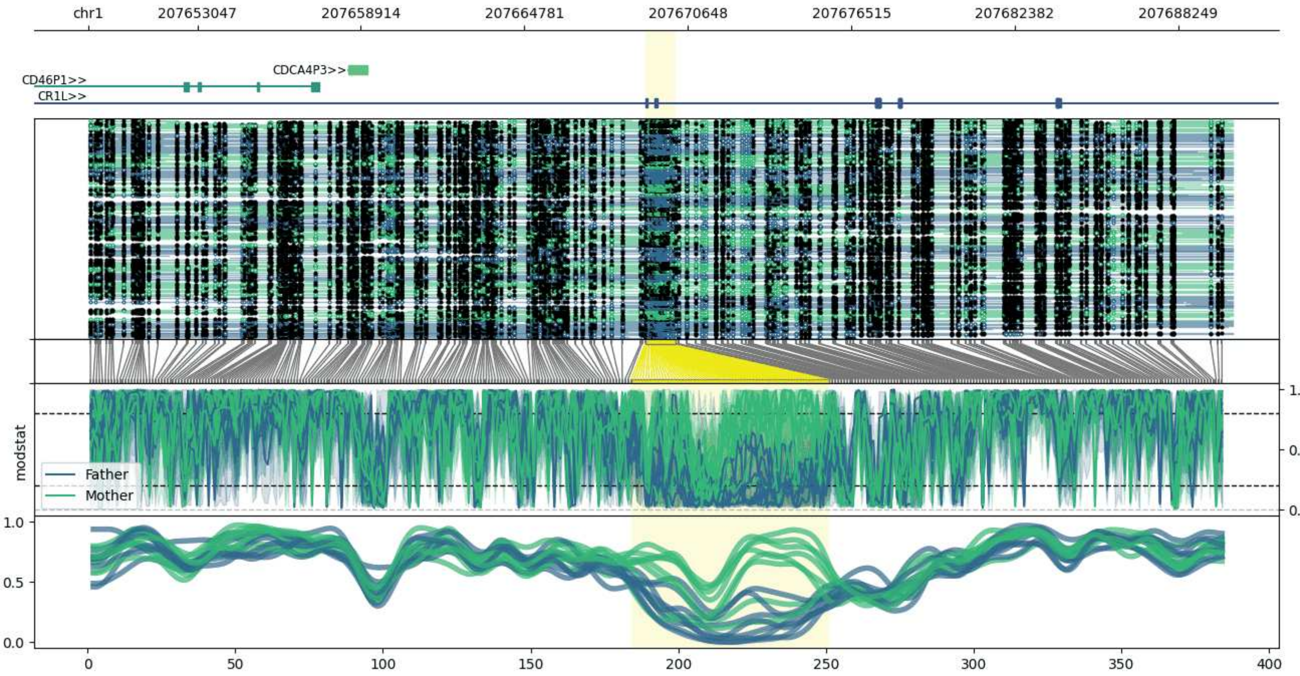


**chr1:207649121-207690191:** Region with 1 DMR containing genes: CD46P1, CDCA4P3, CR1, CR1L.

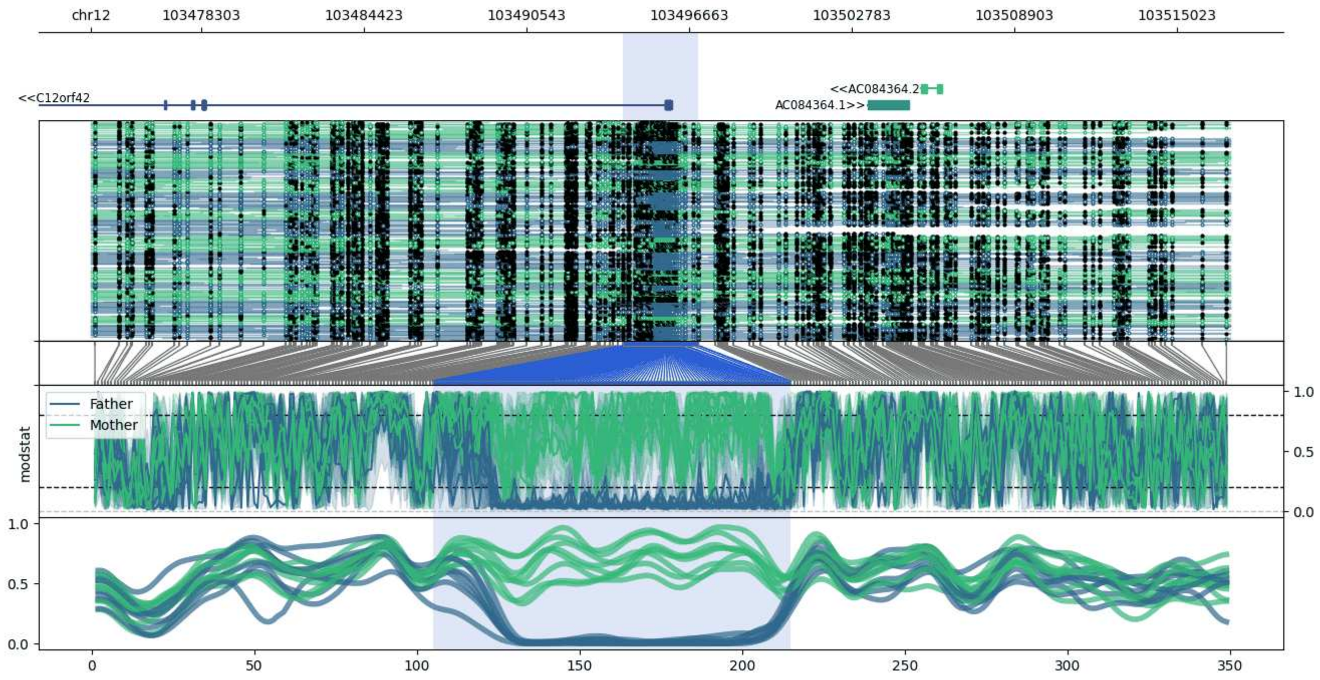


**chr12:103474174-103517019:** Region with 1 DMR containing genes: AC084364.1, AC084364.2, AC084364.3, C12orf42.

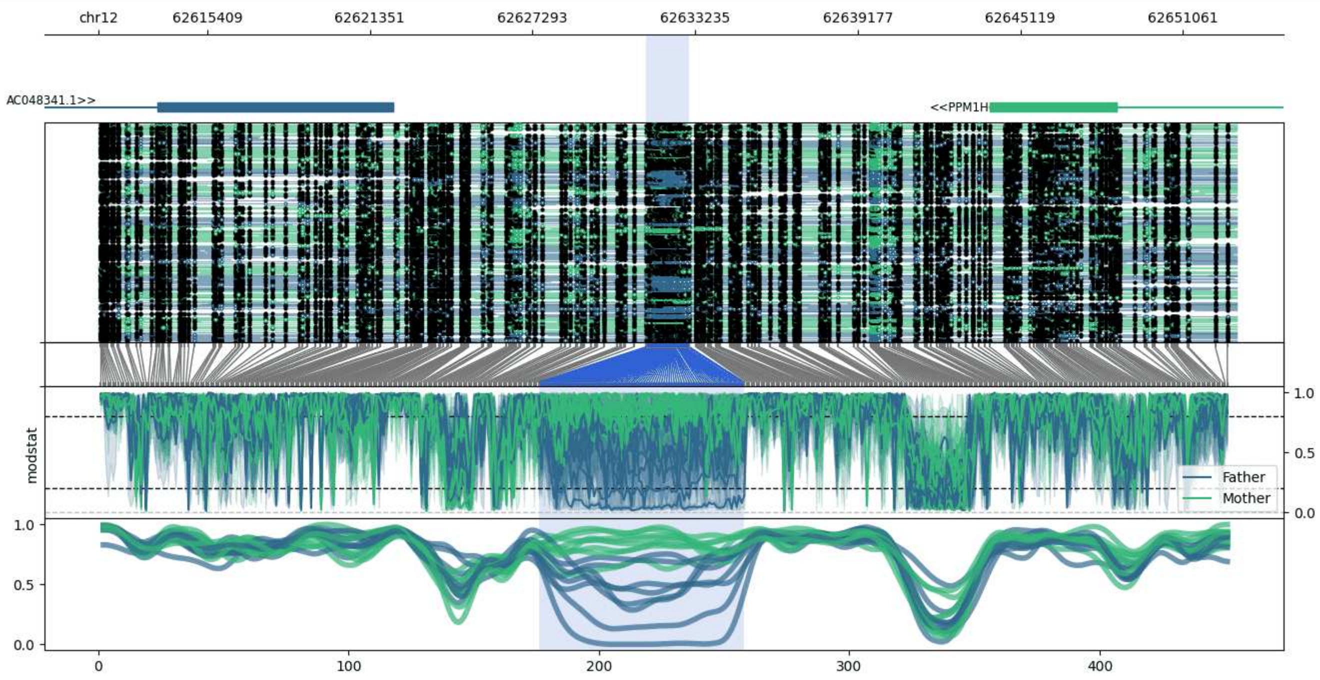


**chr12:62611430-62653026:** Region with 1 DMR containing genes: AC048341.1, AC048341.2, LINC01465, MIRLET7I, PPM1H, RPL32P26.

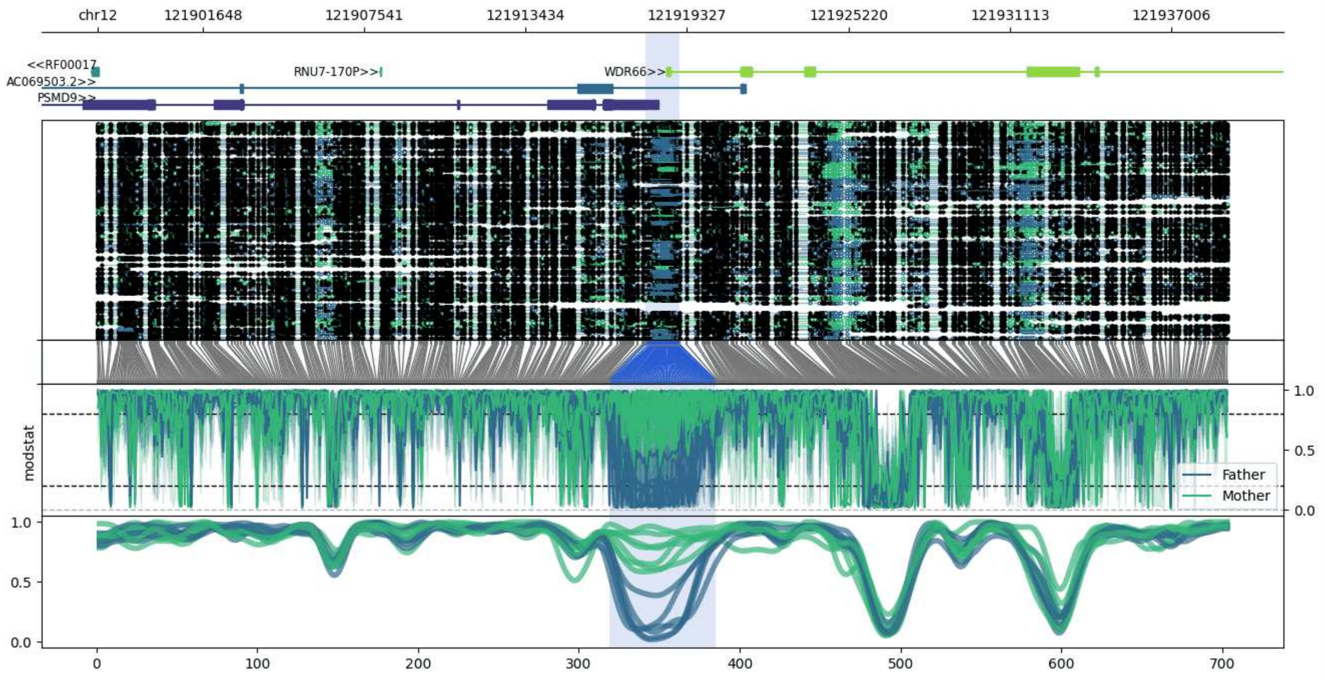


**chr12:121897804-121939057:** Region with 1 DMR containing genes: AC069503.2, AC069503.4, PSMD9, RF00017, RNU7-170P, WDR66.

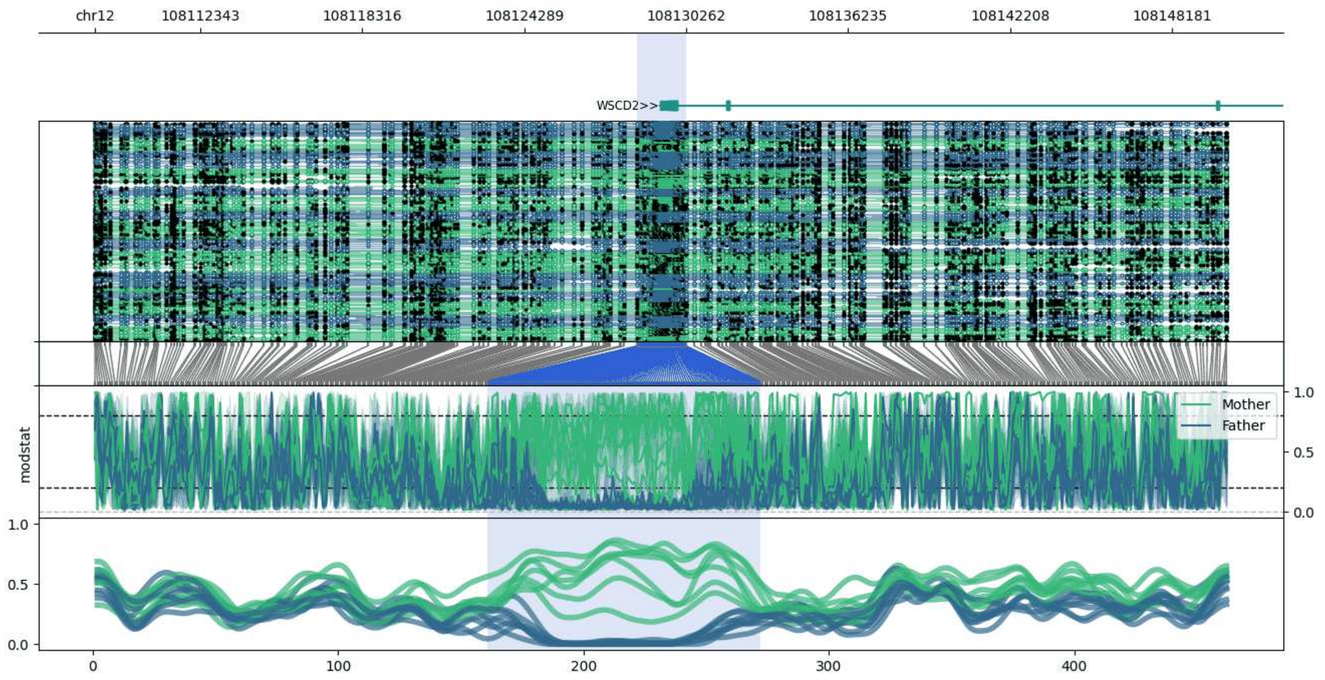


**chr12:108108444-108150259:** Region with 1 DMR containing genes: WSCD2.

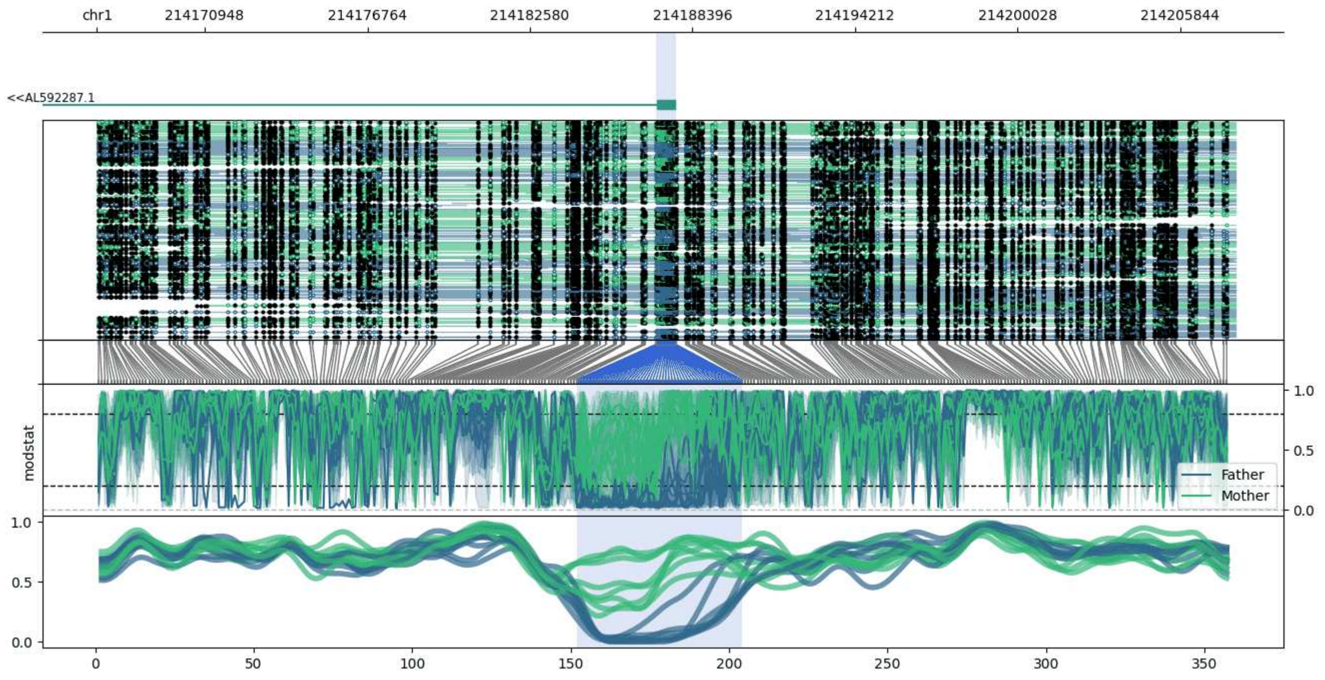


**chr1:214167090-214207803:** Region with 1 DMR containing genes: AL592287.1.

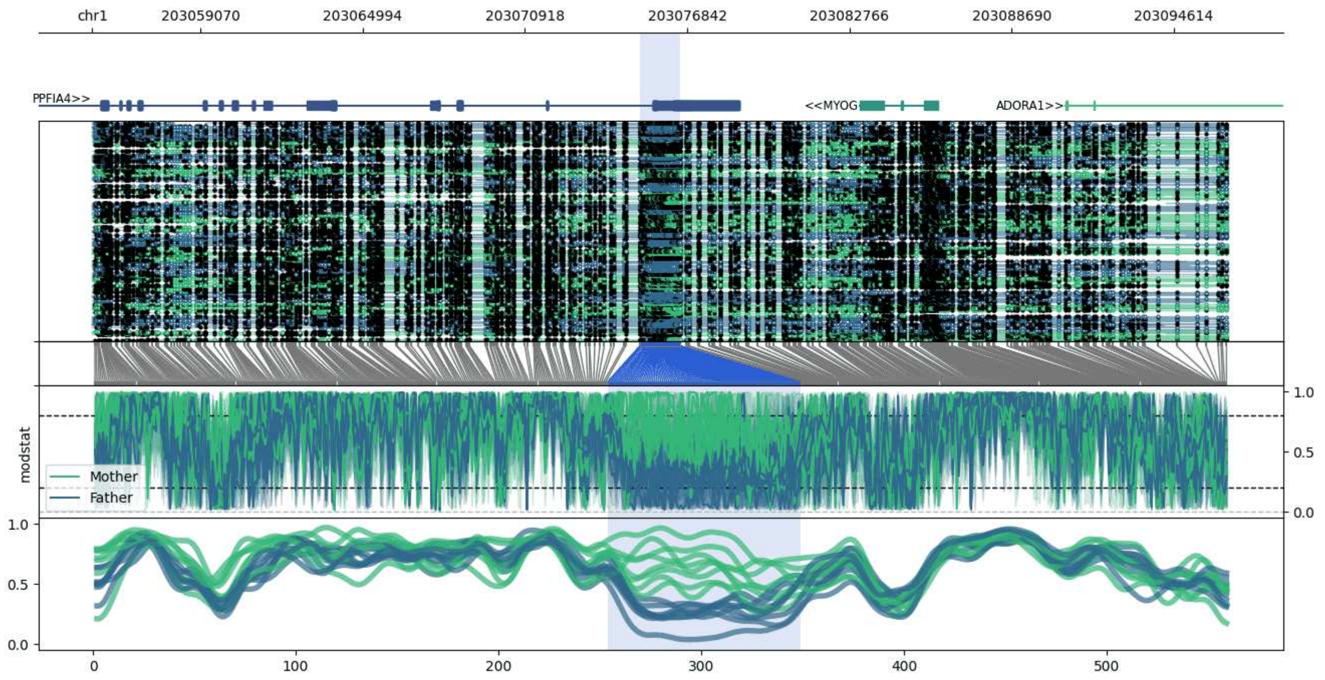


**chr1:203055116-203096587:** Region with 1 DMR containing genes: ADORA1, MYOG, PPFIA4.

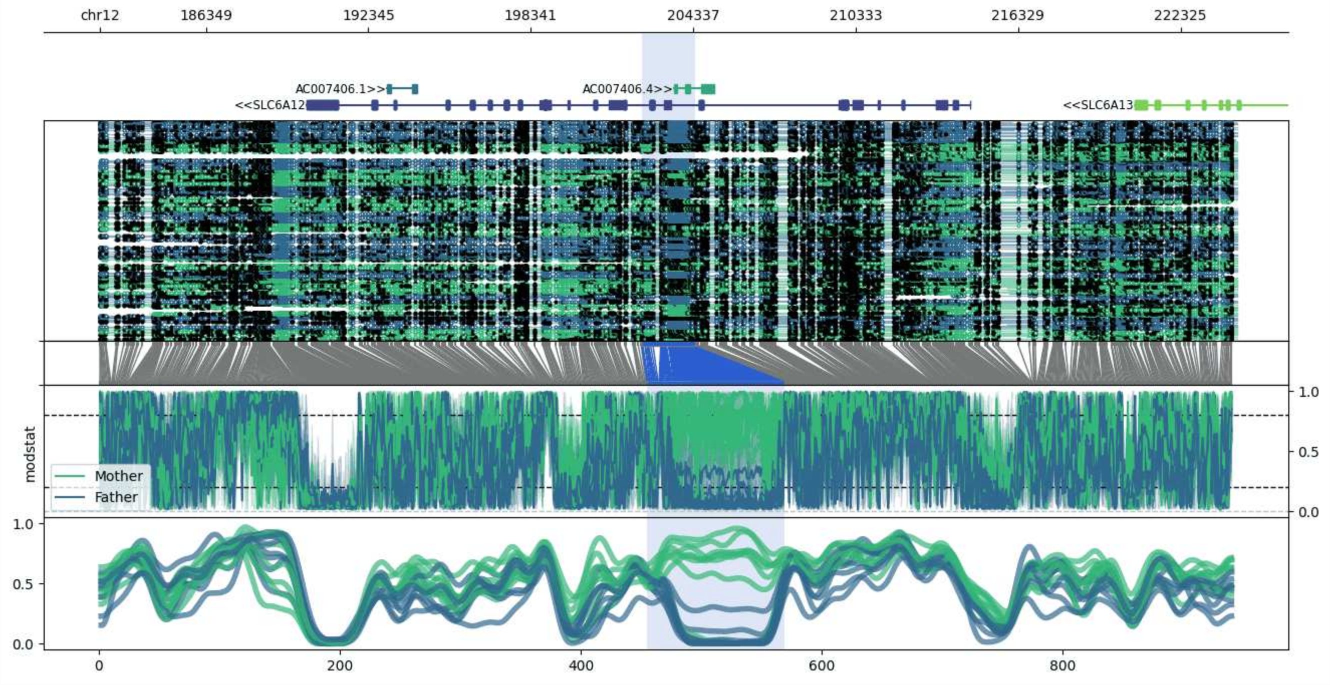


**chr12:182438-224411:** Region with 1 DMR containing genes: AC007406.1, AC007406.3, AC007406.4, IQSEC3, SLC6A12, SLC6A13.

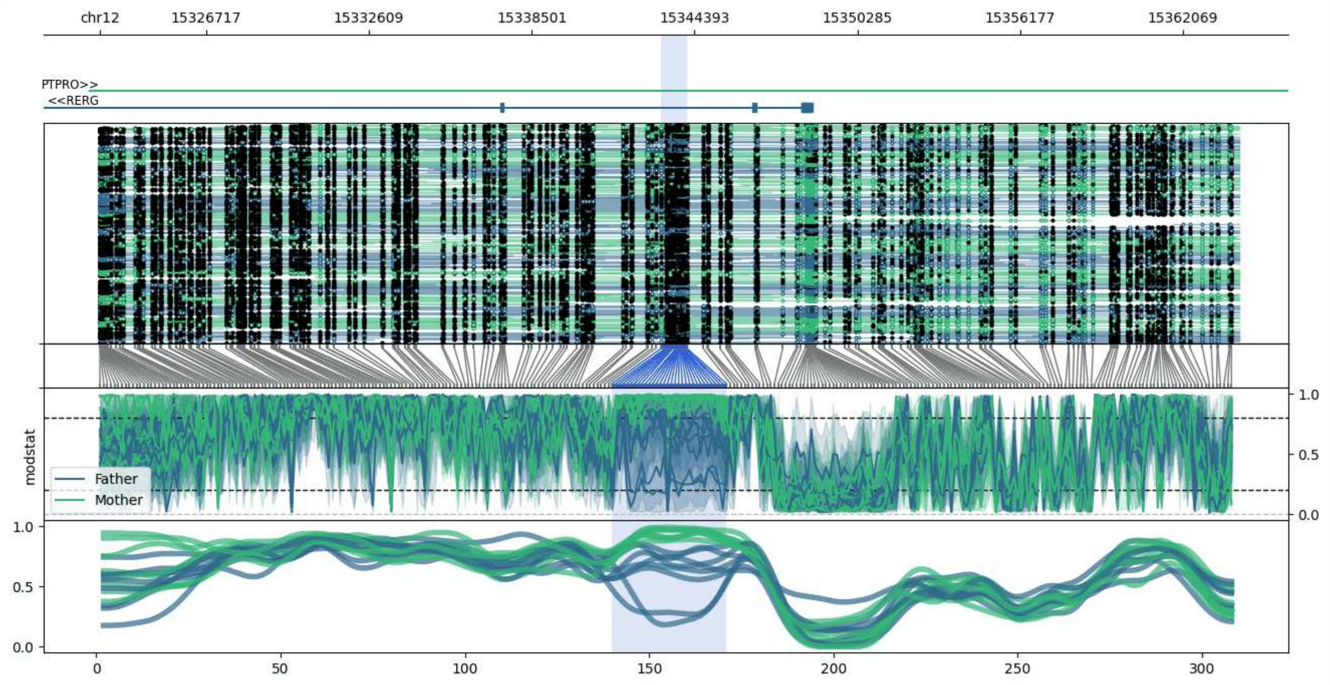


**chr12:15322866-15364114:** Region with 1 DMR containing genes: PTPRO, RERG.

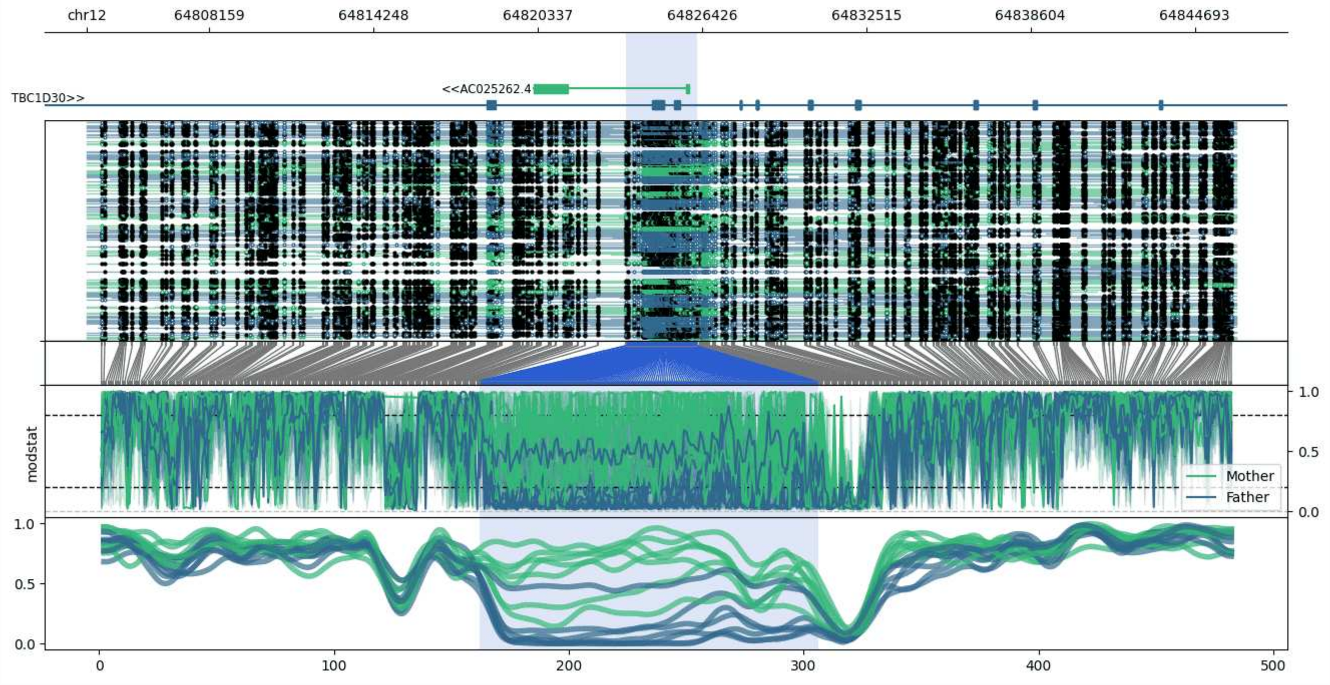


**chr12:64803615-64846244:** Region with 1 DMR containing genes: AC025262.4, TBC1D30.

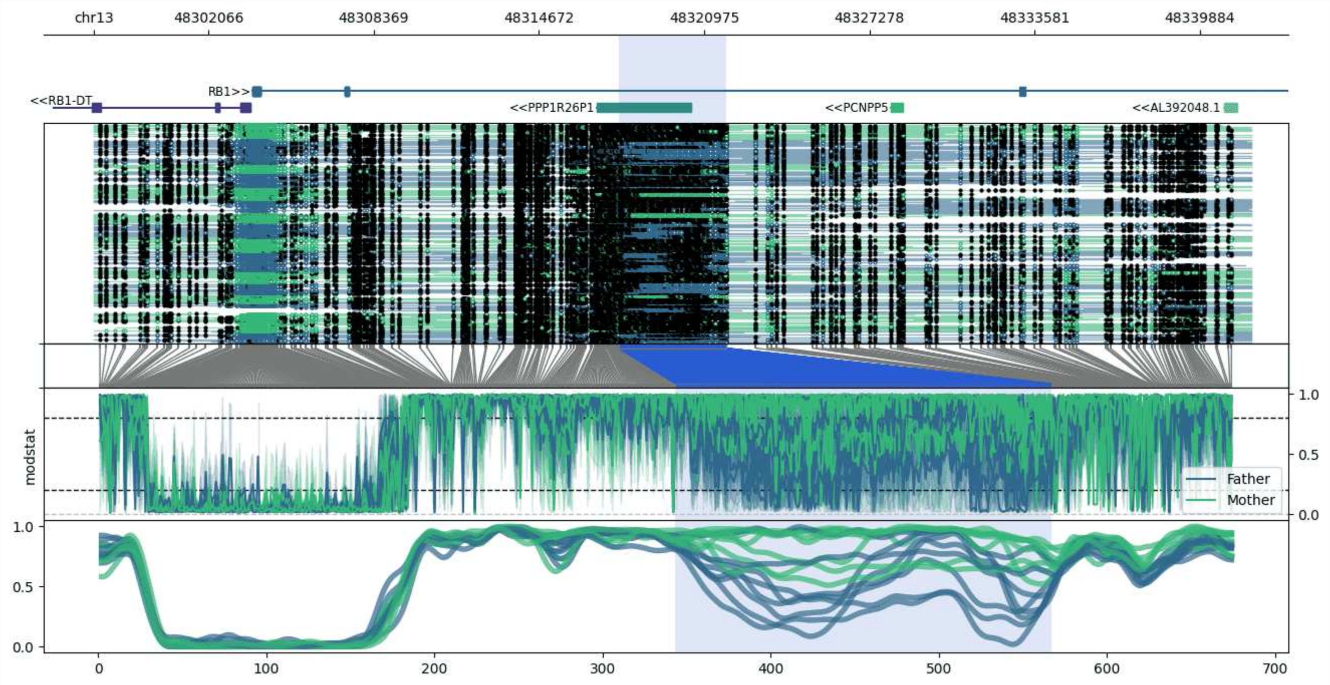


**chr13:48297707-48341833:** Region with 1 DMR containing genes: AL392048.1, PCNPP5, PPP1R26P1, RB1, RB1-DT.

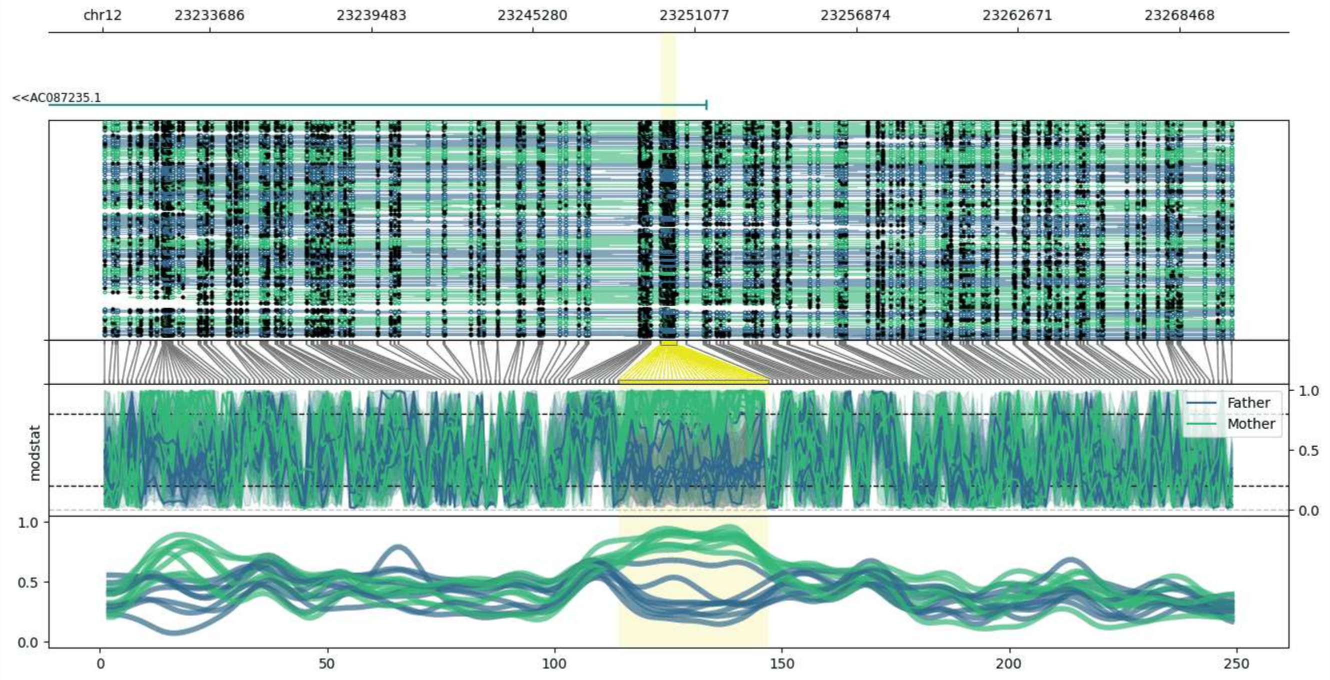


**chr12:23229846-23270425:** Region with 1 DMR containing genes: AC087235.1.

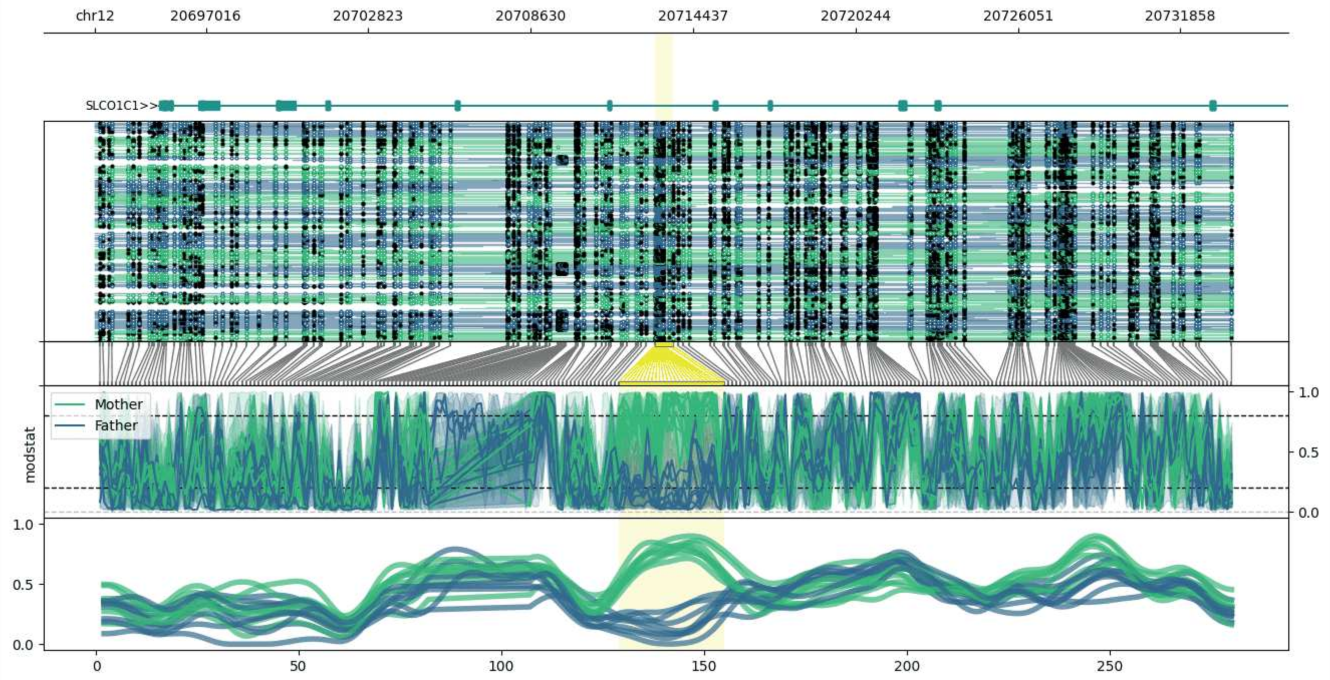


**chr12:20693063-20733712:** Region with 1 DMR containing genes: PDE3A, SLCO1C1.

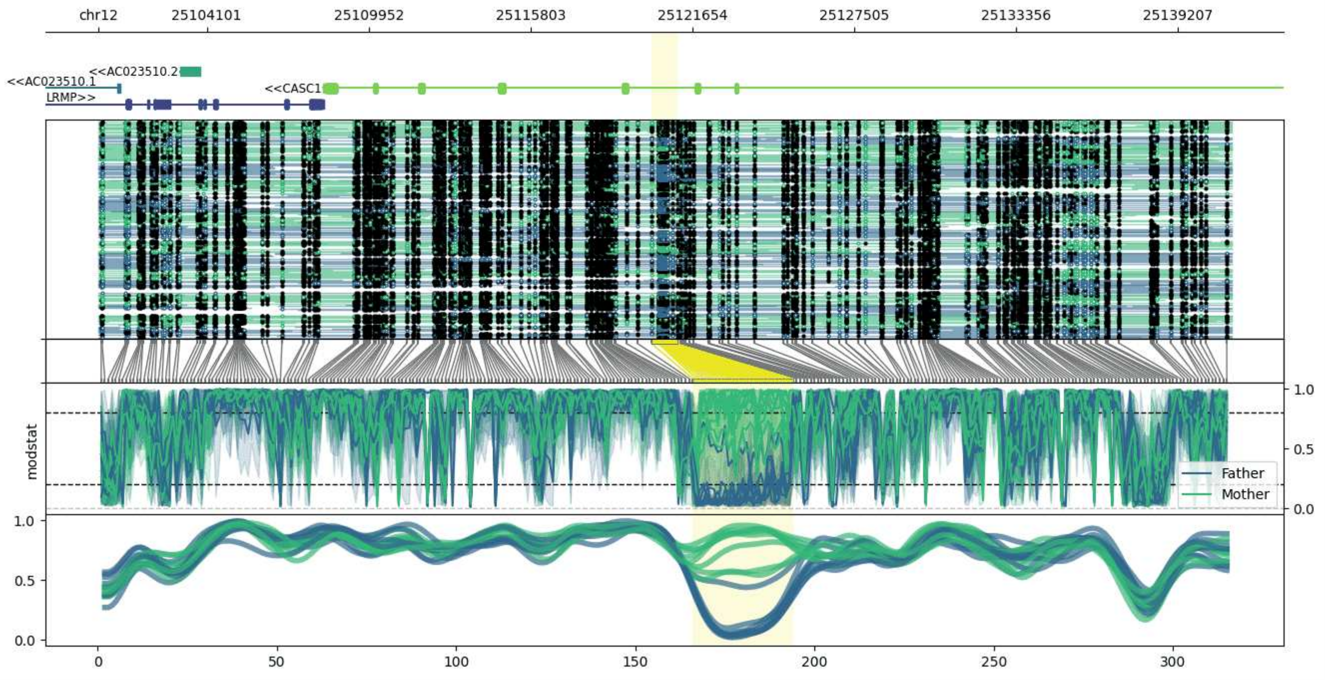


**chr12:25100167-25141129:** Region with 1 DMR containing genes: AC023510.1, AC023510.2, CASC1, CENPUP2, LRMP.

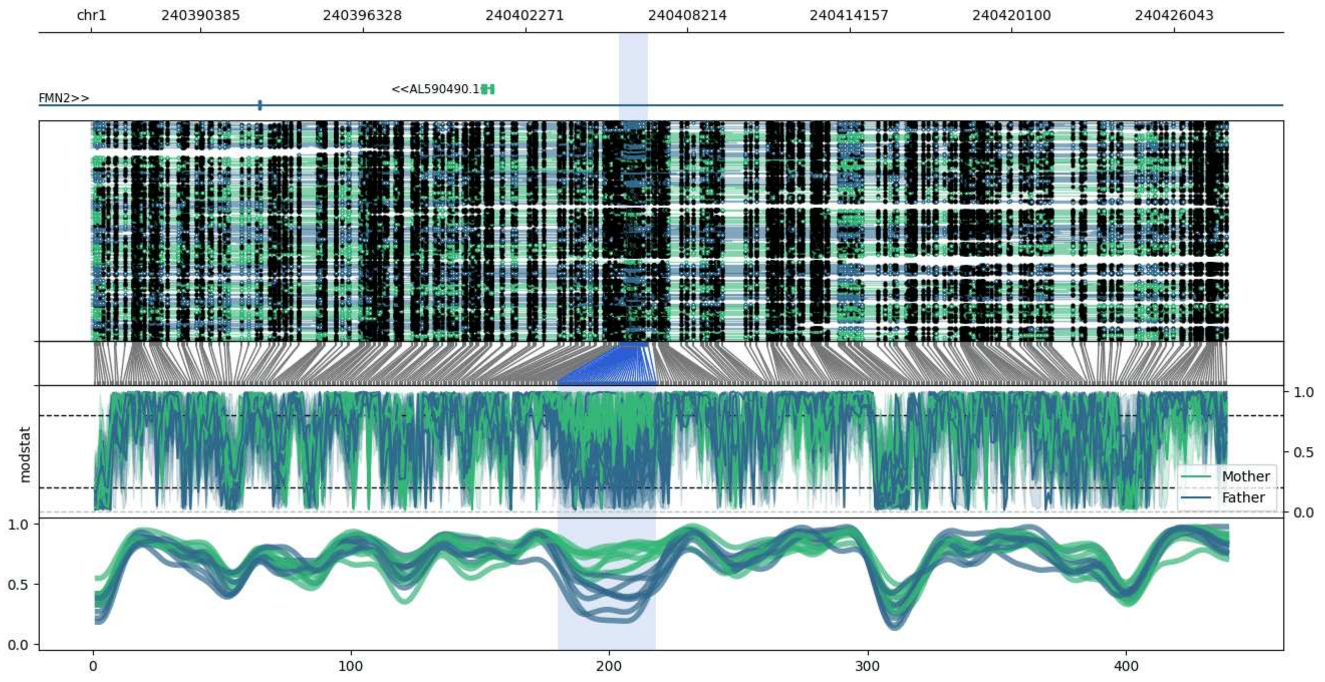


**chr1:240386378-240427982:** Region with 1 DMR containing genes: AL590490.1, FMN2.

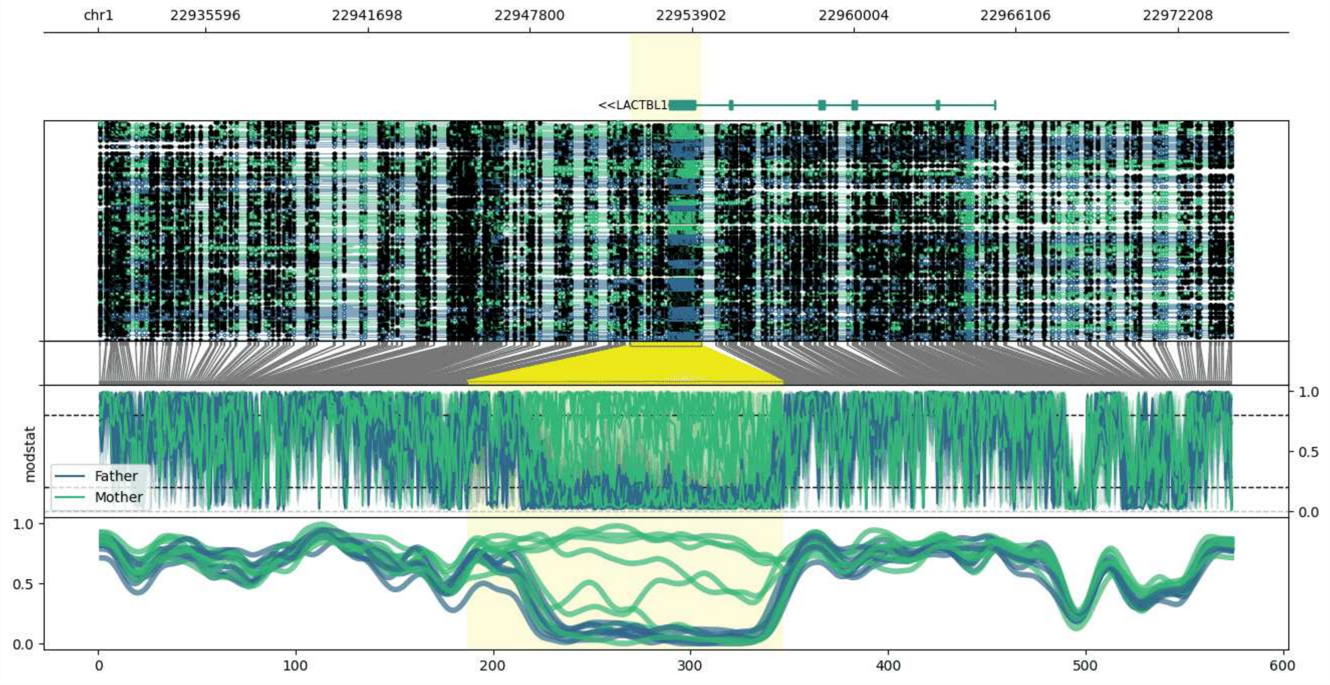


**chr1:22931559-22974276:** Region with 1 DMR containing genes: LACTBL1.

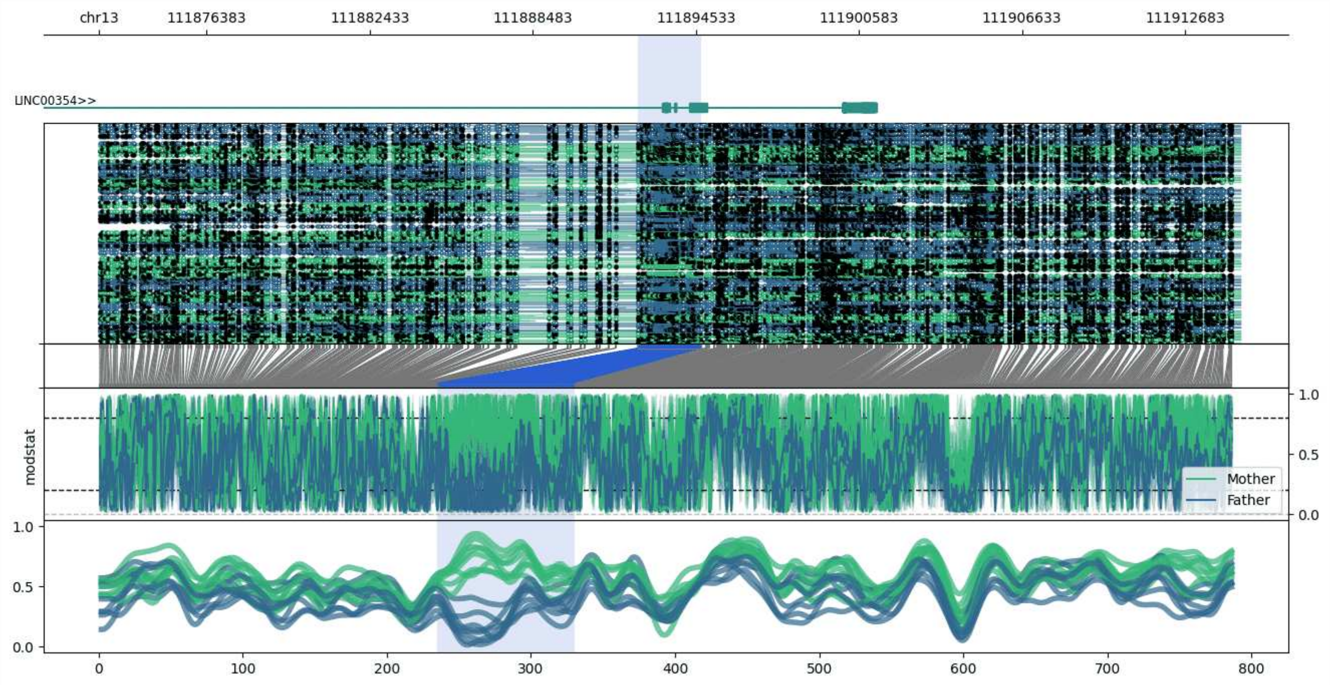


**chr13:111872383-111914733:** Region with 1 DMR containing genes: LINC00354.

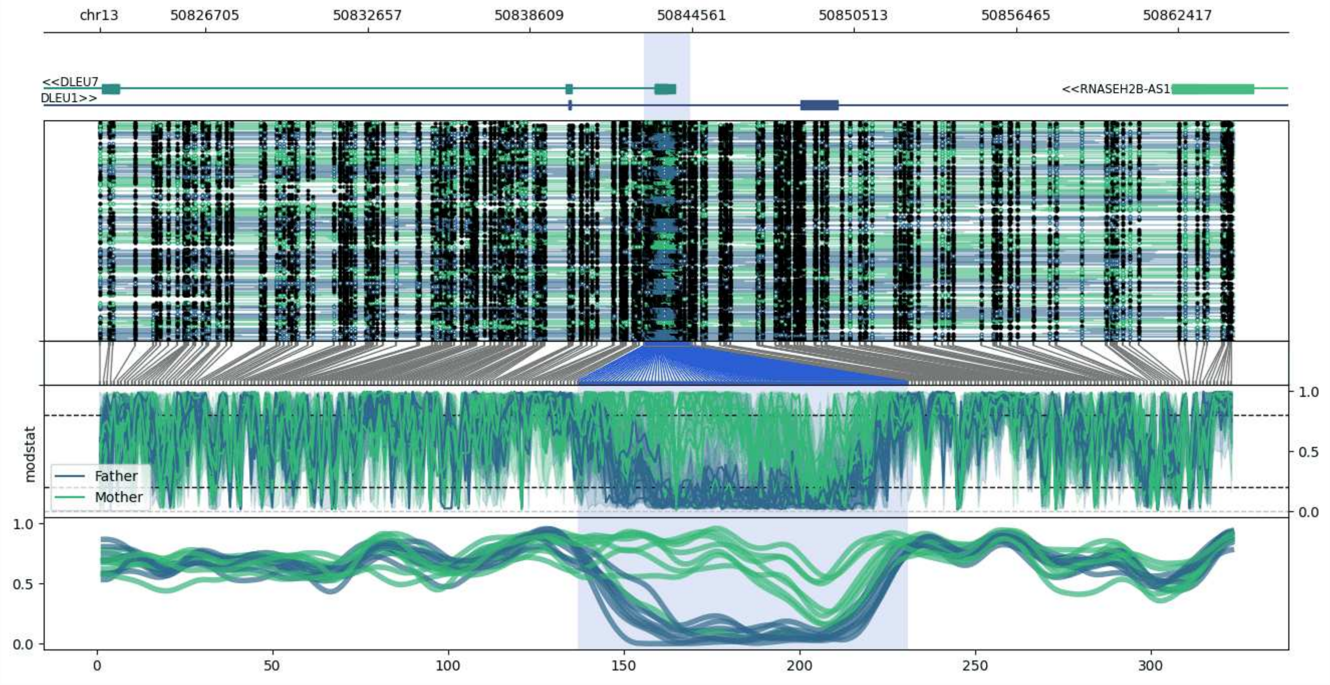


**chr13:50822811-50864478:** Region with 1 DMR containing genes: DLEU1, DLEU7, RNA5SP28, RNASEH2B-AS1.

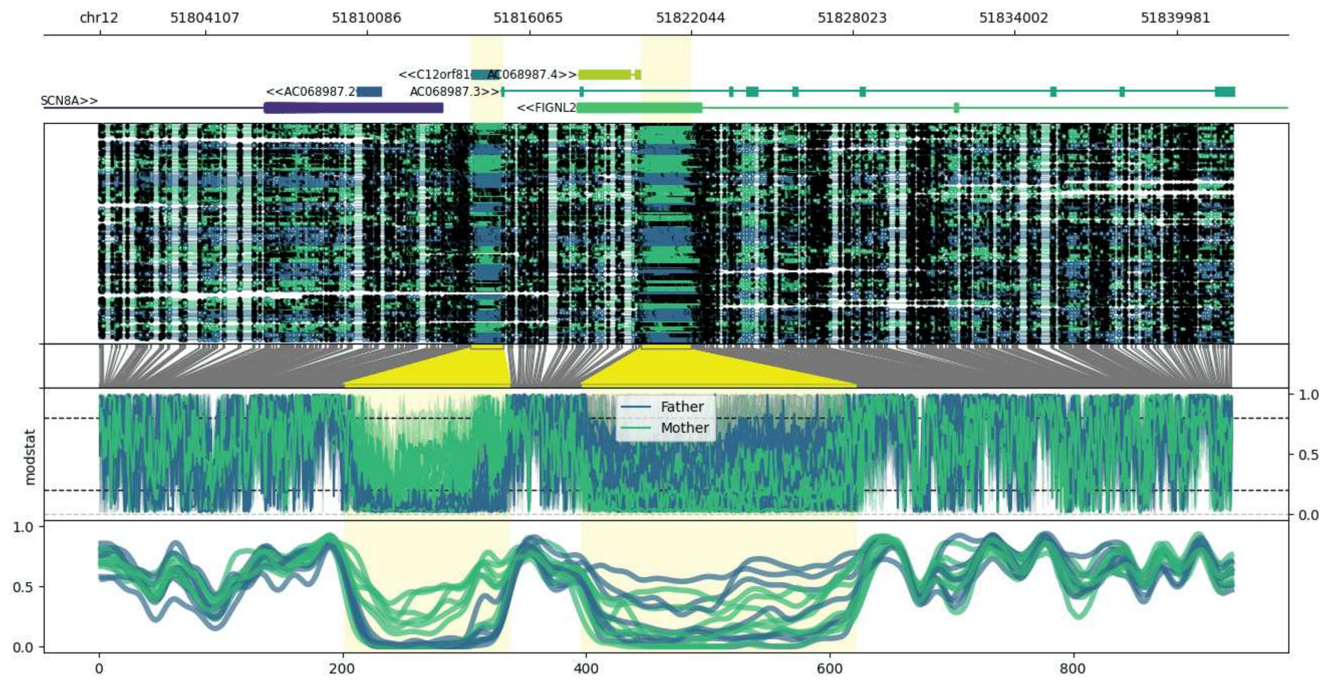


**chr12:51800198-51842053:** Region with 2 DMRs containing genes: AC068987.1, AC068987.2, AC068987.3, AC068987.4, C12orf81, FIGNL2, SCN8A.

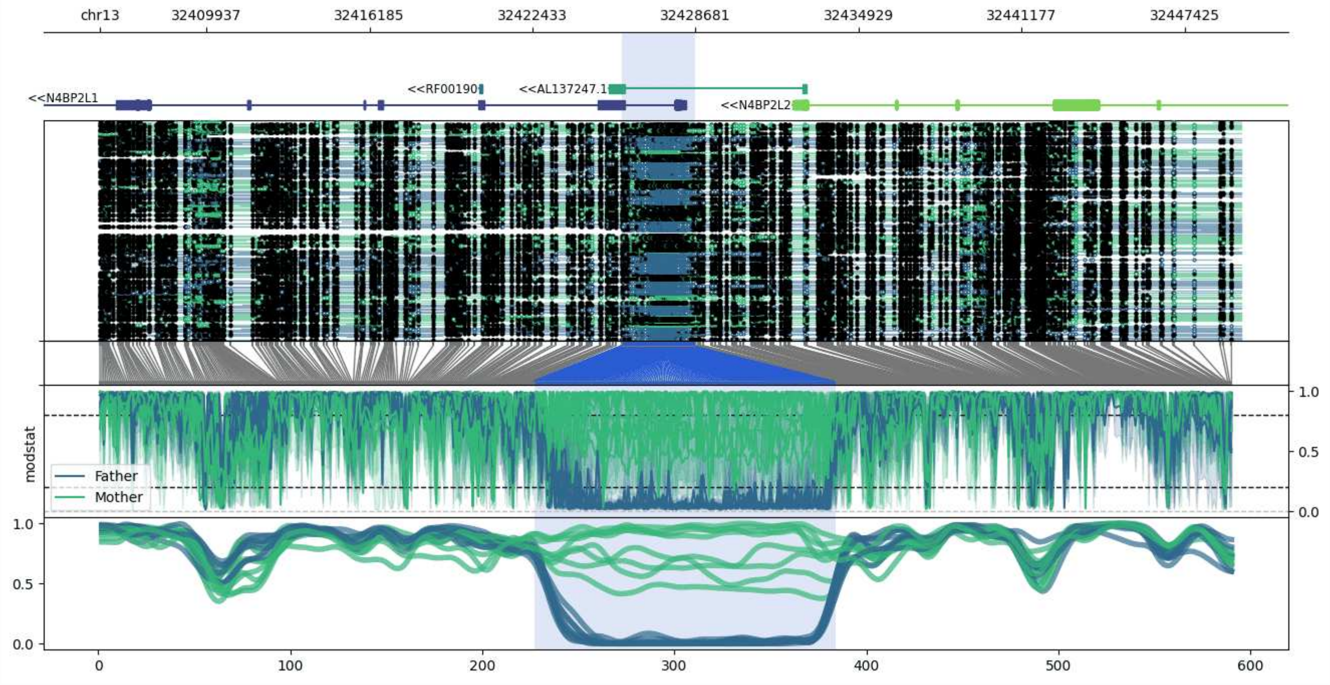


**chr13:32405851-32449593:** Region with 1 DMR containing genes: AL137247.1, BRCA2, N4BP2L1, N4BP2L2, RF00190.

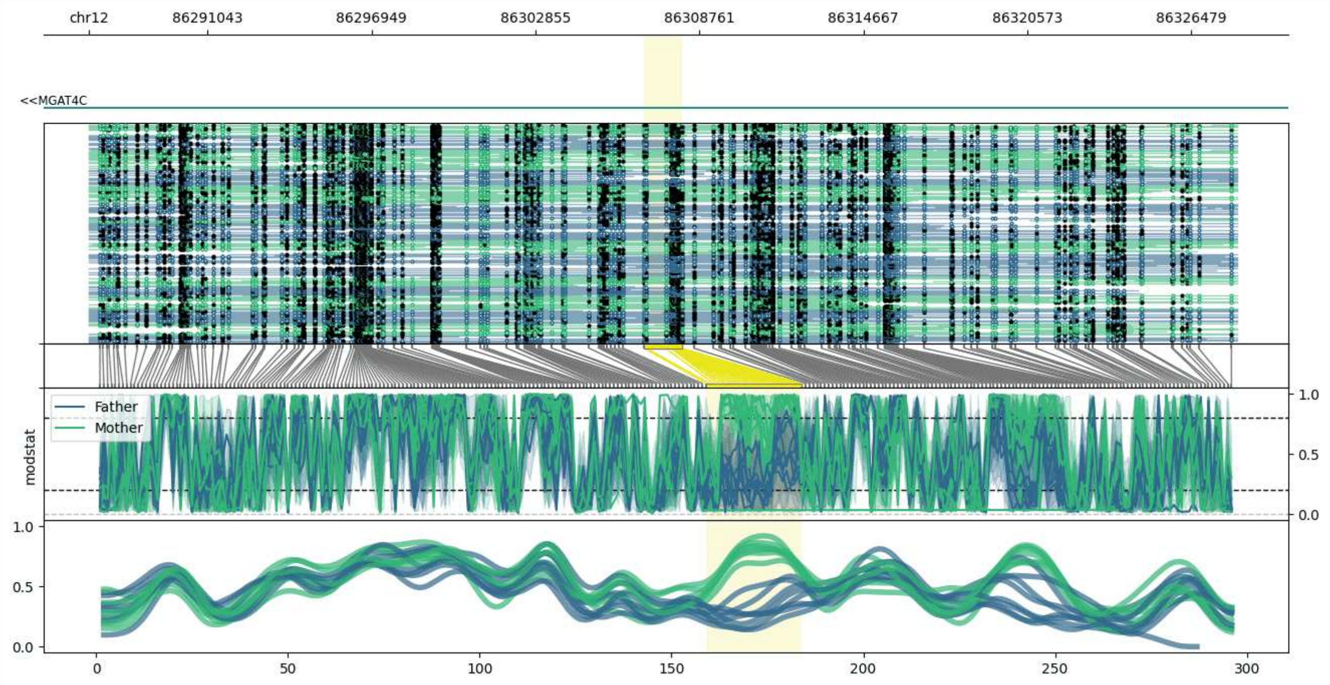


**chr12:86286776-86328123:** Region with 1 DMR containing genes: MGAT4C.

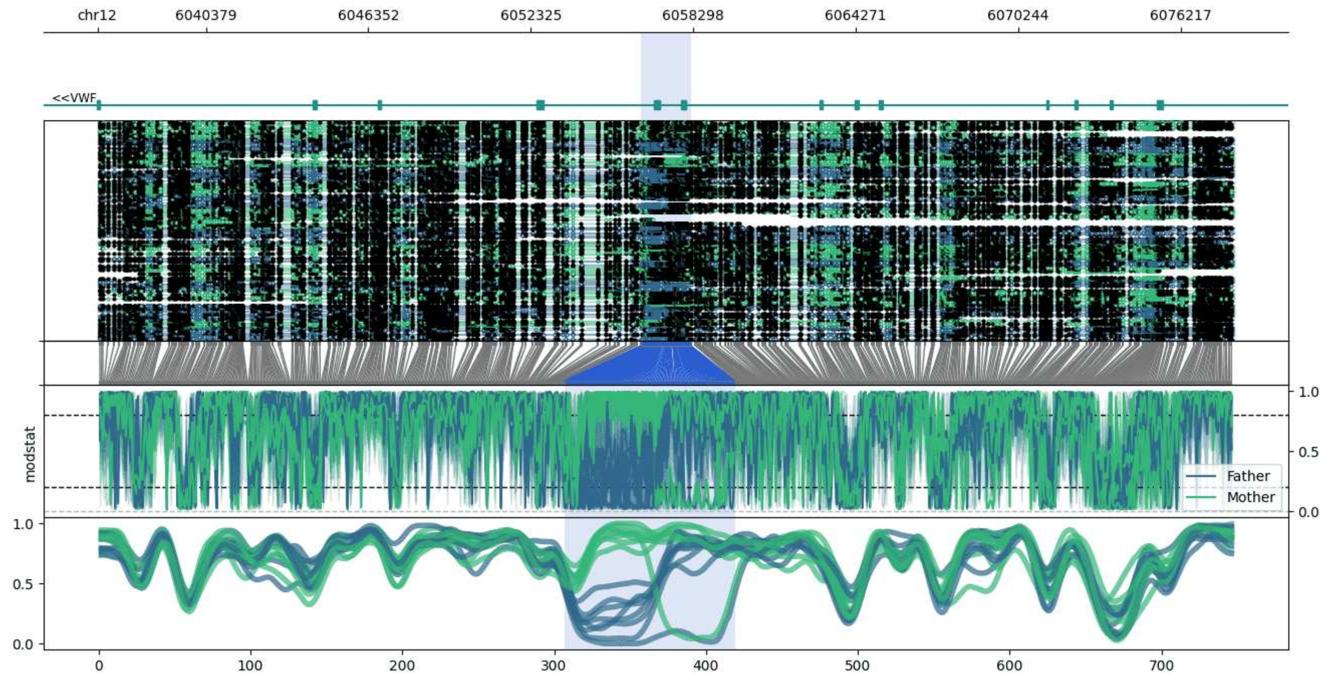


**chr12:6036392-6078206:** Region with 1 DMR containing genes: VWF.

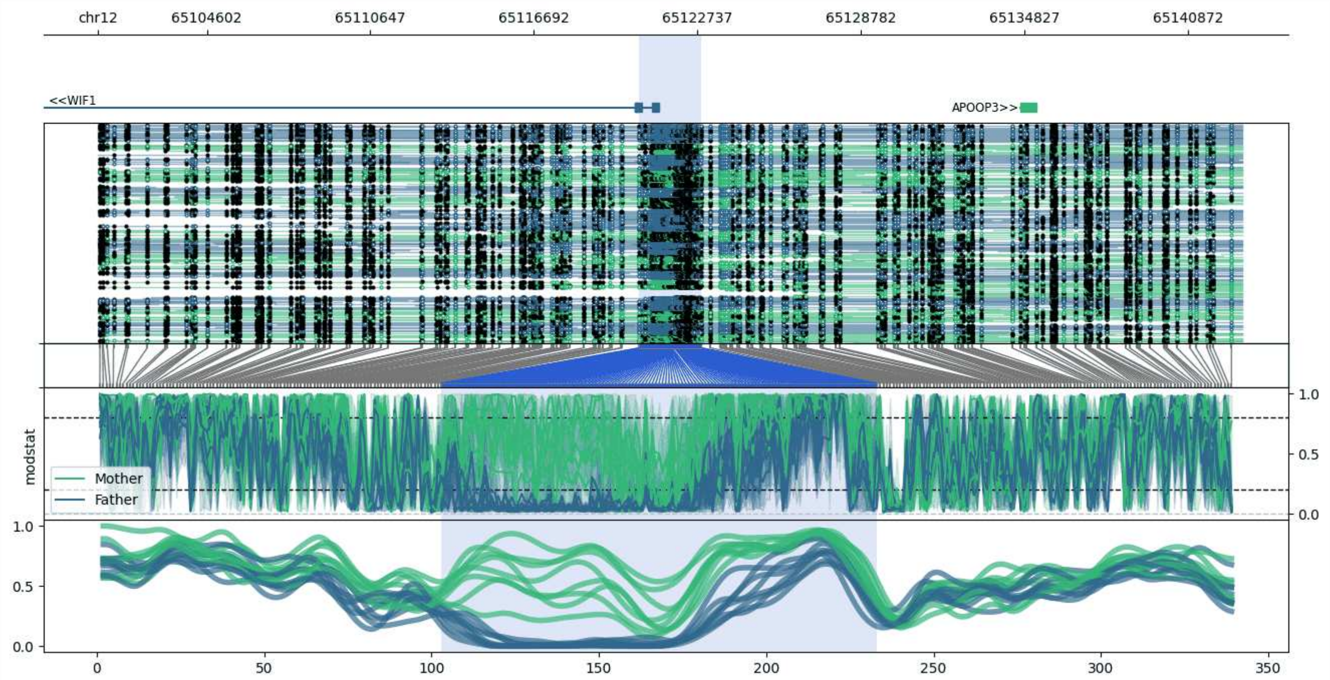


**chr12:65100577-65142895:** Region with 1 DMR containing genes: APOOP3, RNU6-166P, WIF1.

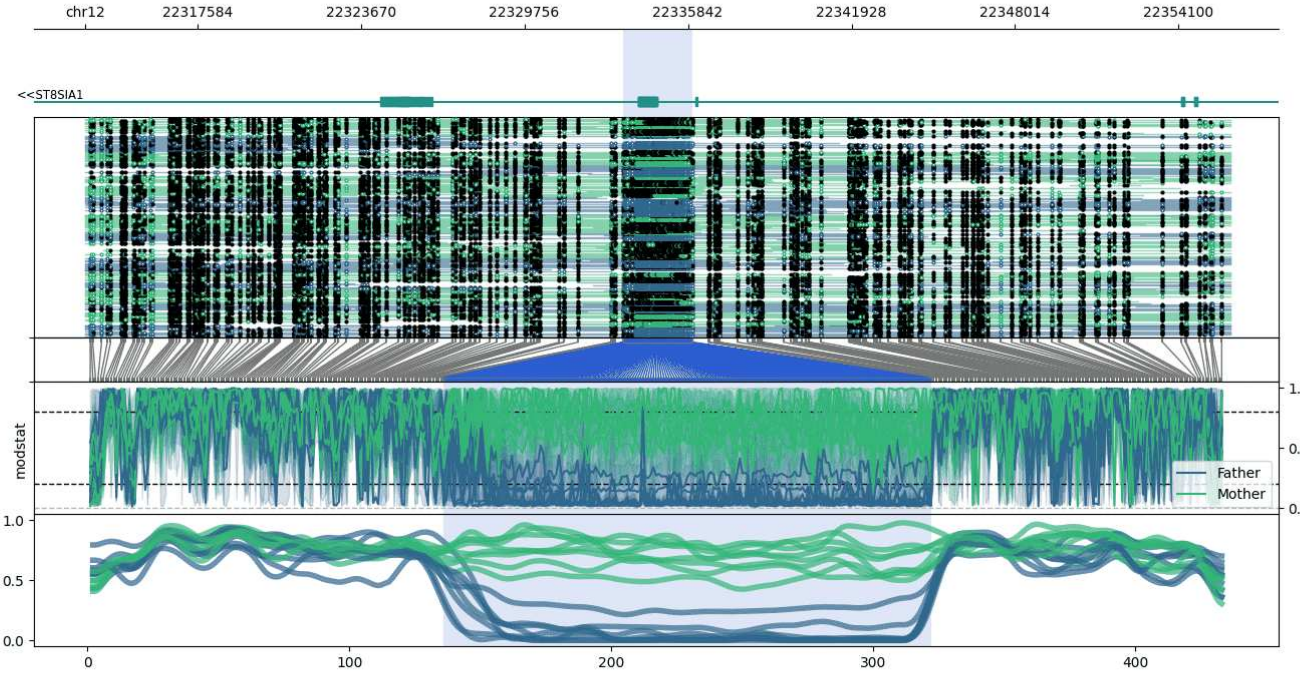


**chr12:22313421-22356026:** Region with 1 DMR containing genes: ST8SIA1.

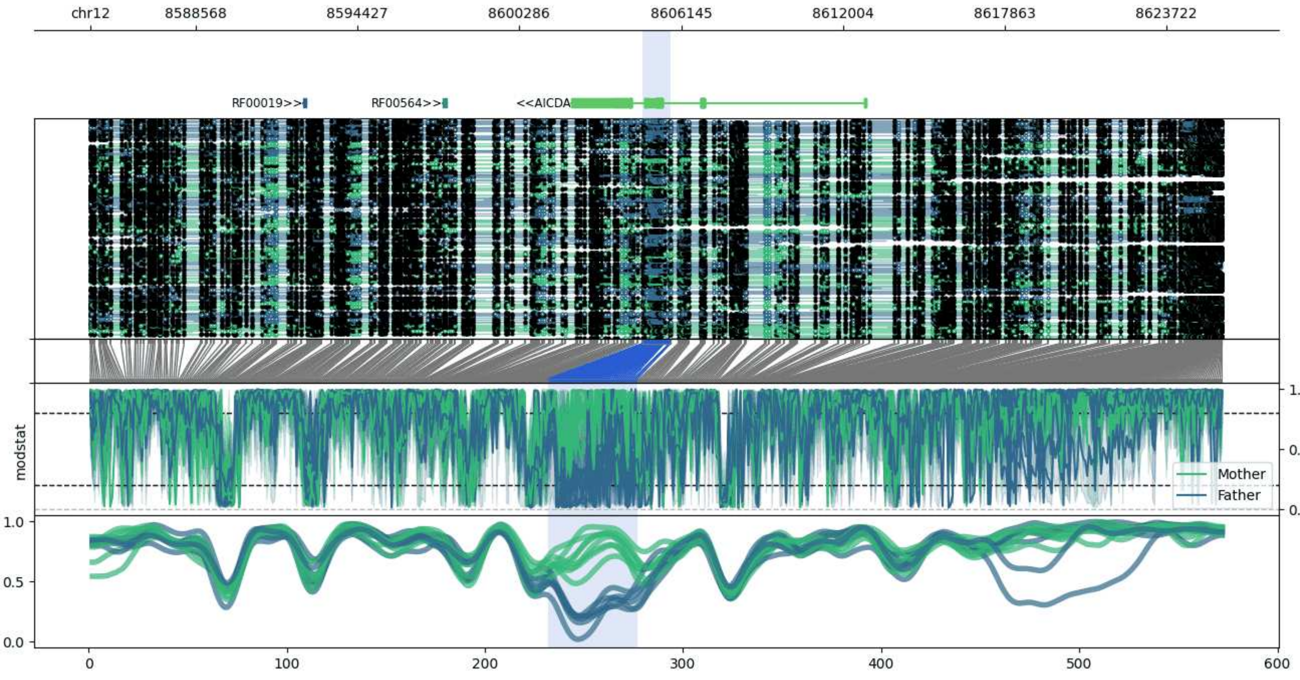


**chr12:8584744-8625763:** Region with 1 DMR containing genes: AC092184.1, AICDA, HADHAP2, RF00019, RF00564.

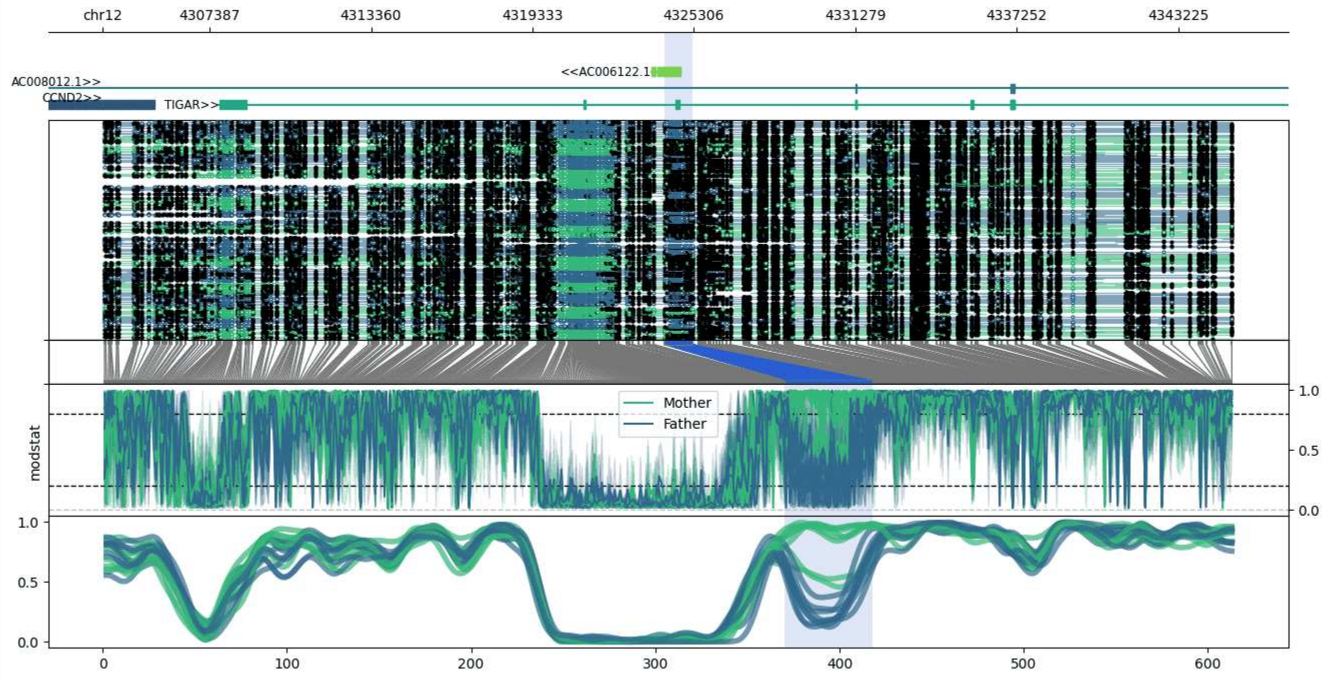


**chr12:4303428-4345240:** Region with 1 DMR containing genes: AC006122.1, AC008012.1, CCND2, TIGAR.

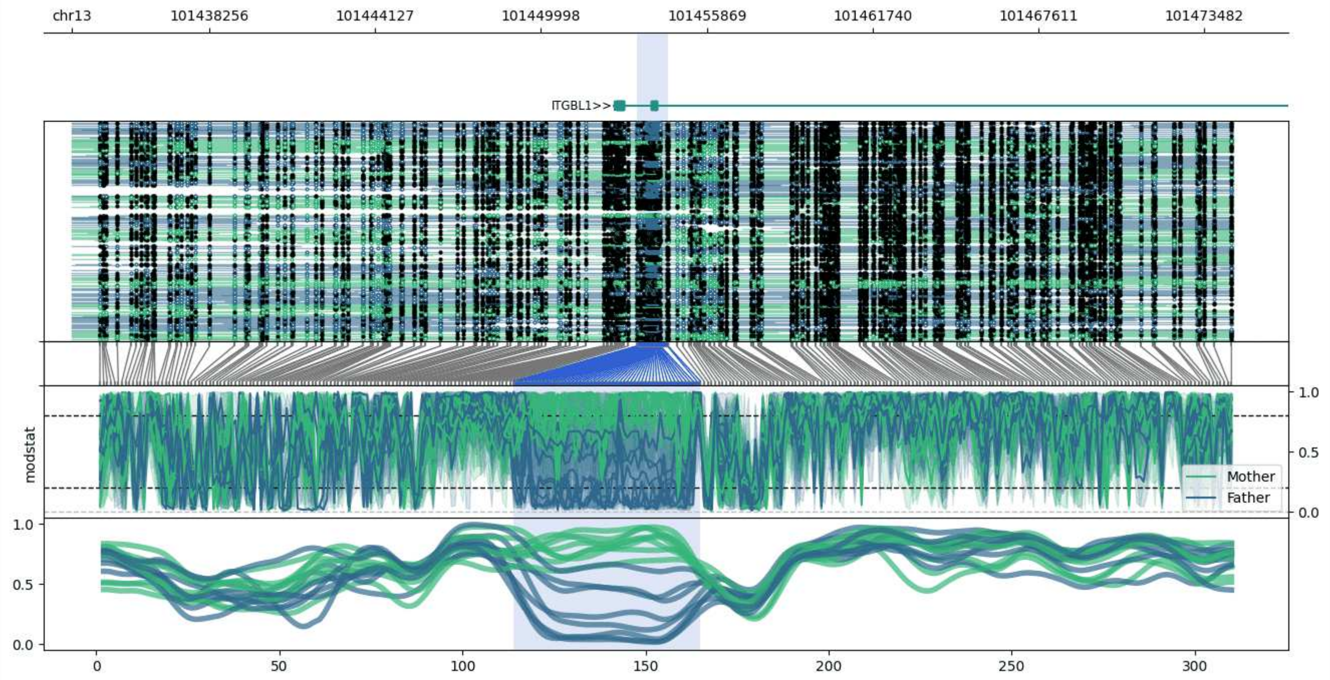


**chr13:101433394-101474493:** Region with 1 DMR containing genes: ITGBL1.

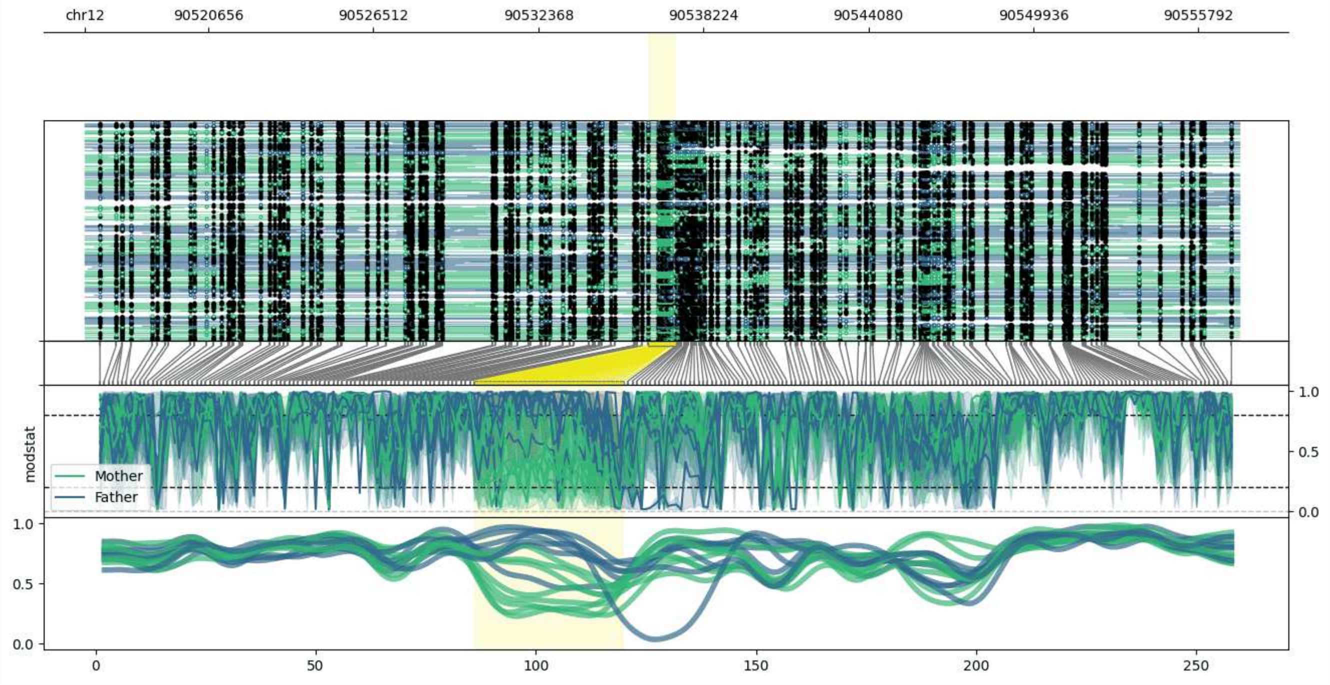


**chr12:90516259-90557254:** Region with 1 DMR containing no genes.

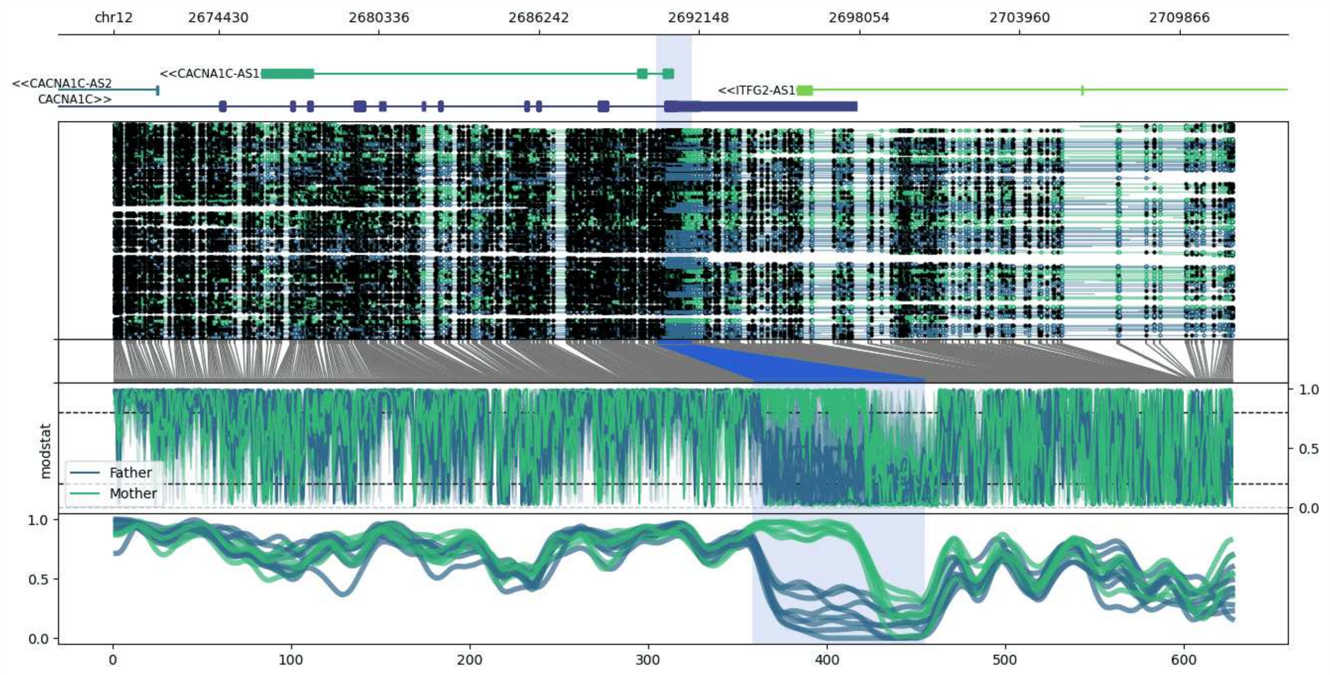


**chr12:2670552-2711894:** Region with 1 DMR containing genes: AC007618.1, CACNA1C, CACNA1C-AS1, CACNA1C-AS2, ITFG2-AS1.

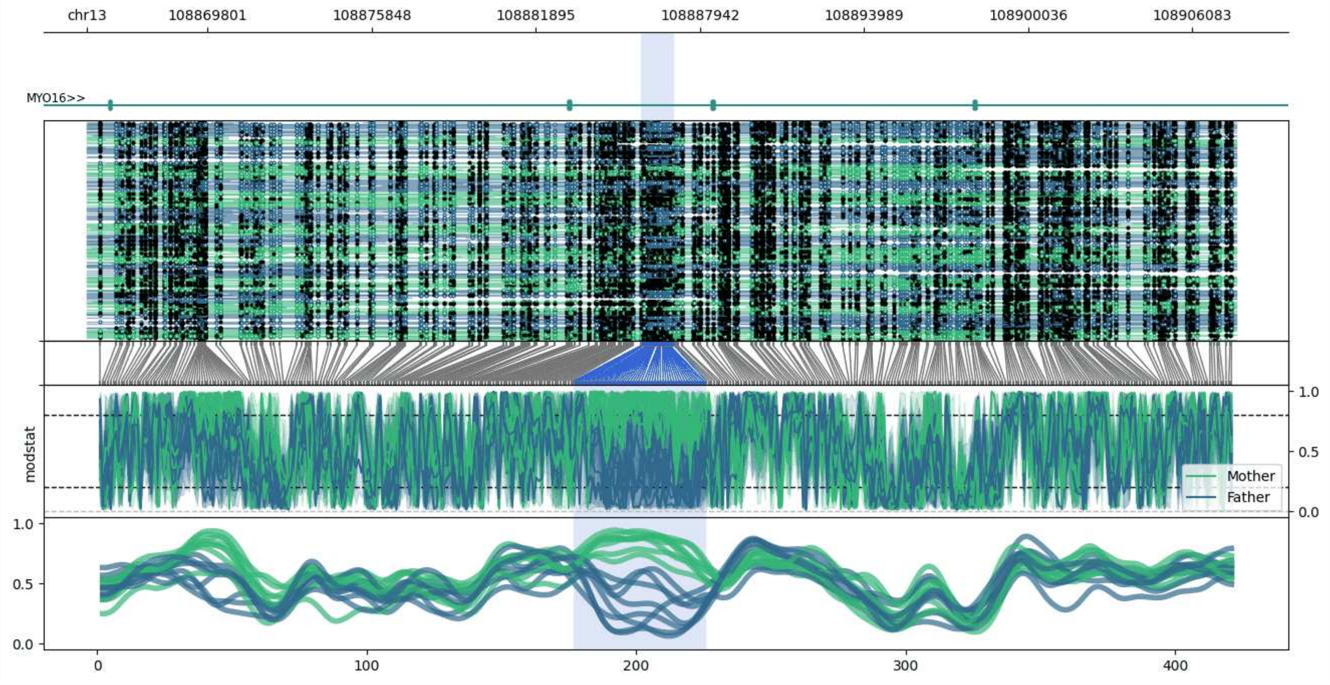


**chr13:108865371-108907700:** Region with 1 DMR containing genes: MYO16.

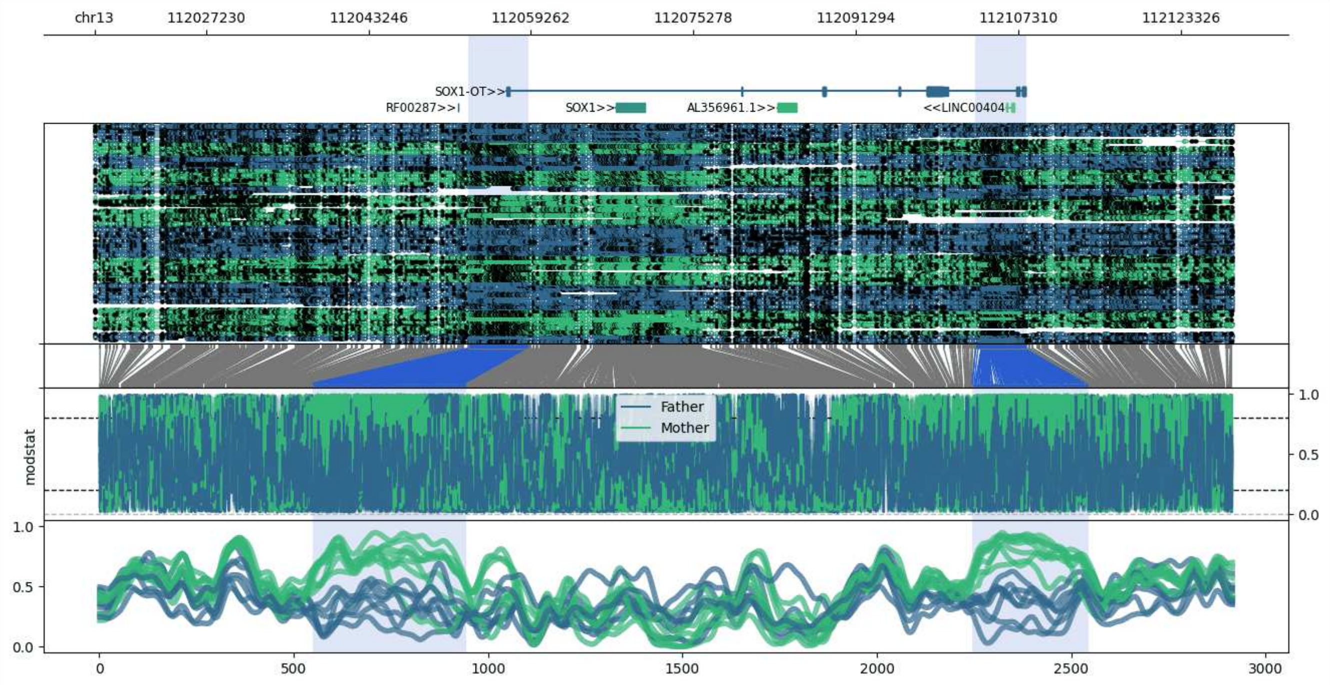


**chr13:112016237-112128353:** Region with 2 DMRs containing genes: AL356961.1, LINC00404, RF00287, SOX1, SOX1-OT.

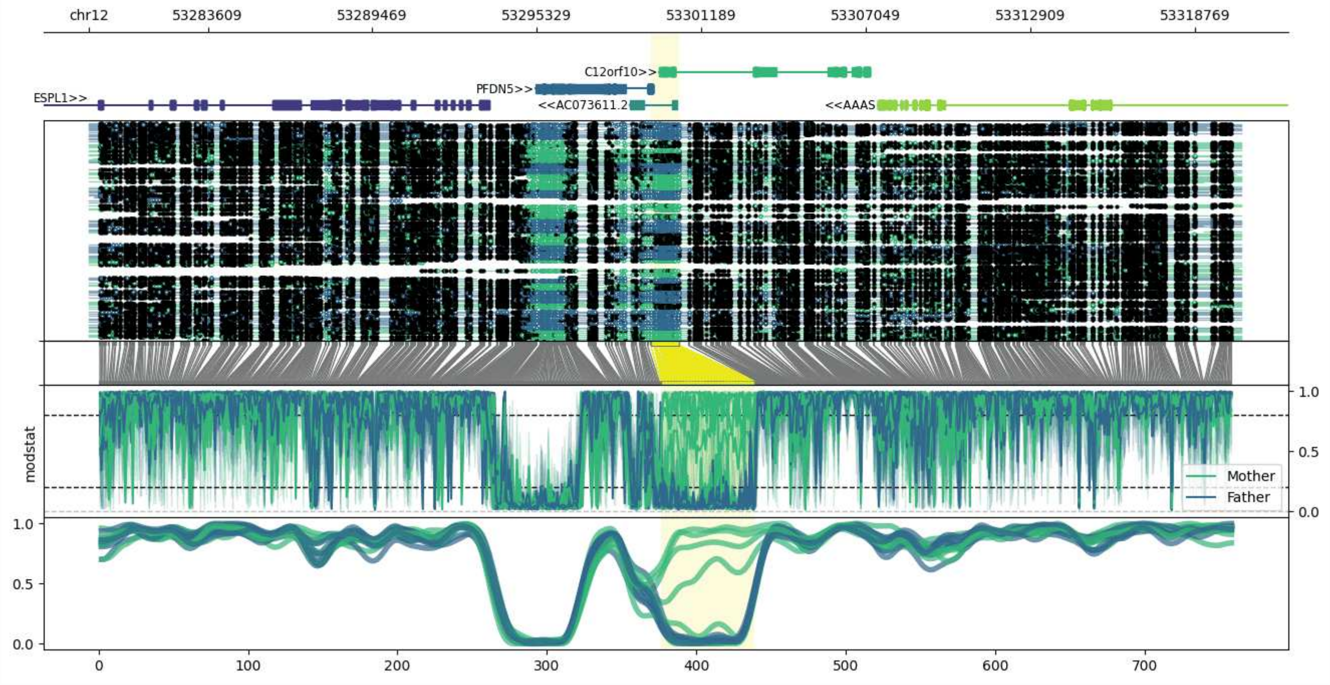


**chr12:53279384-53320410:** Region with 1 DMR containing genes: AAAS, AC073611.1, AC073611.2, C12orf10, ESPL1, PFDN5.

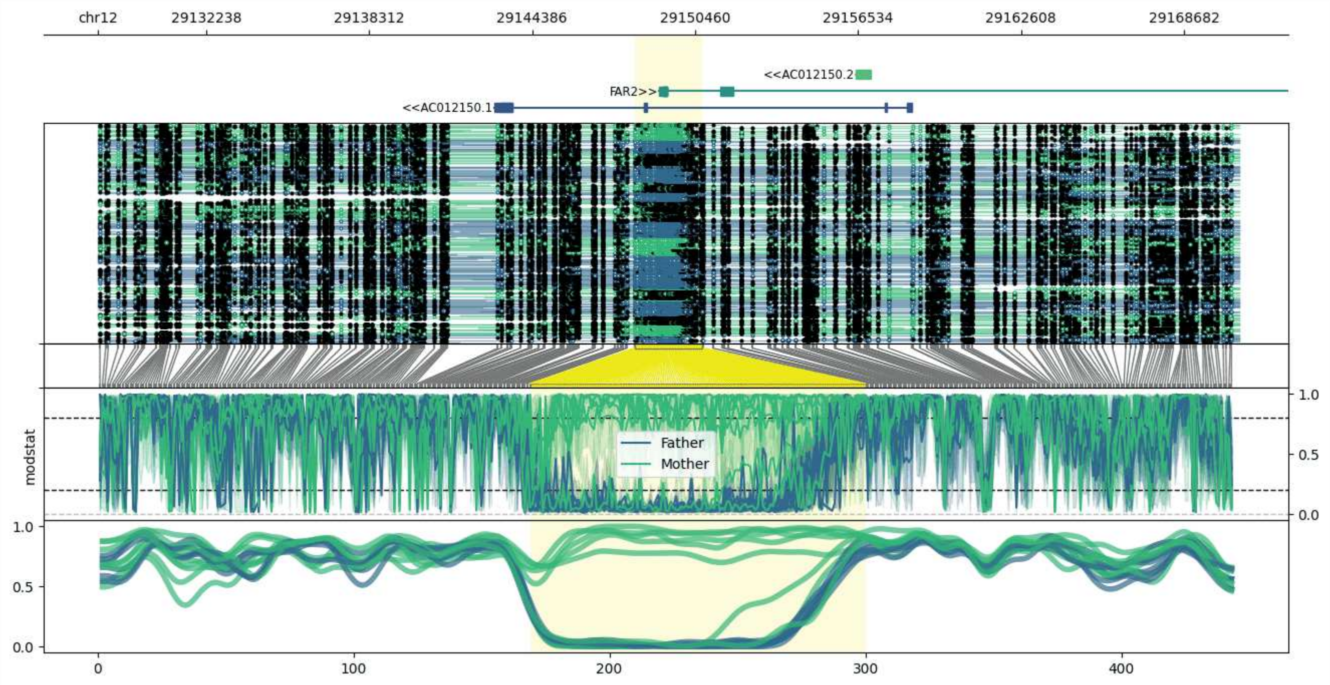


**chr12:29128211-29170731:** Region with 1 DMR containing genes: AC012150.1, AC012150.2, FAR2.

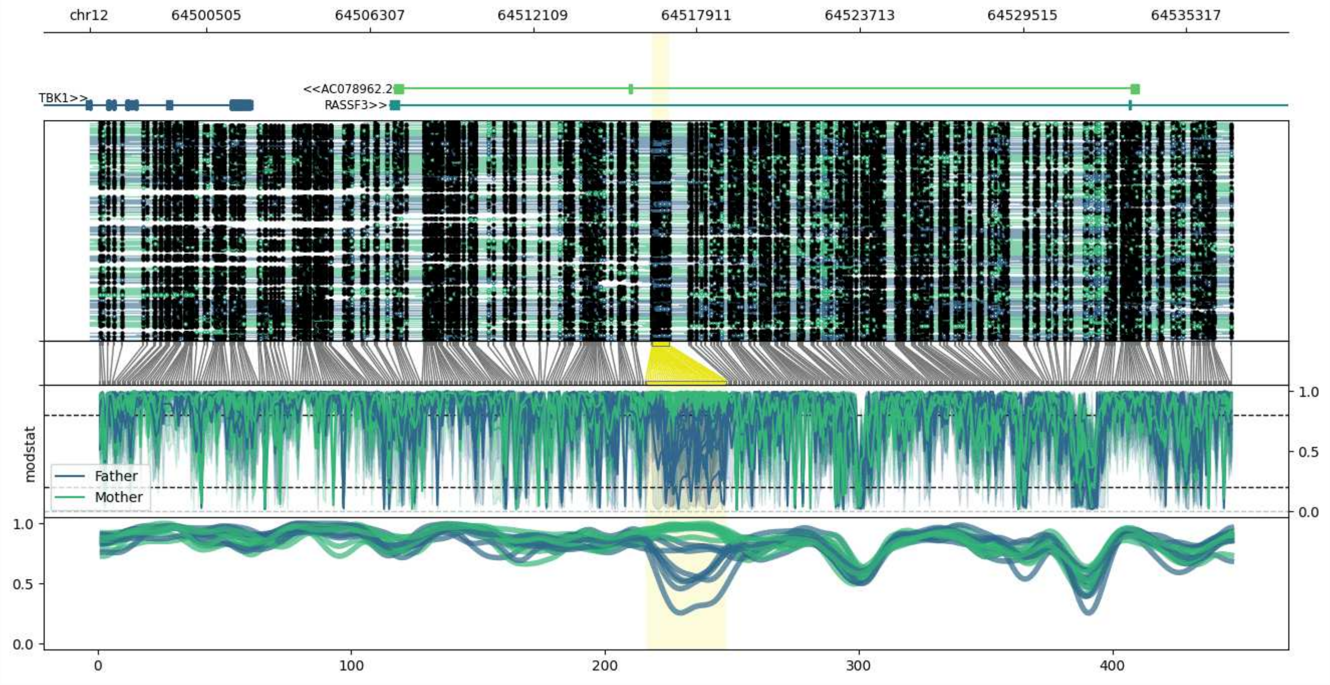


**chr12:64496336-64536952:** Region with 1 DMR containing genes: AC078962.2, RASSF3, TBK1.

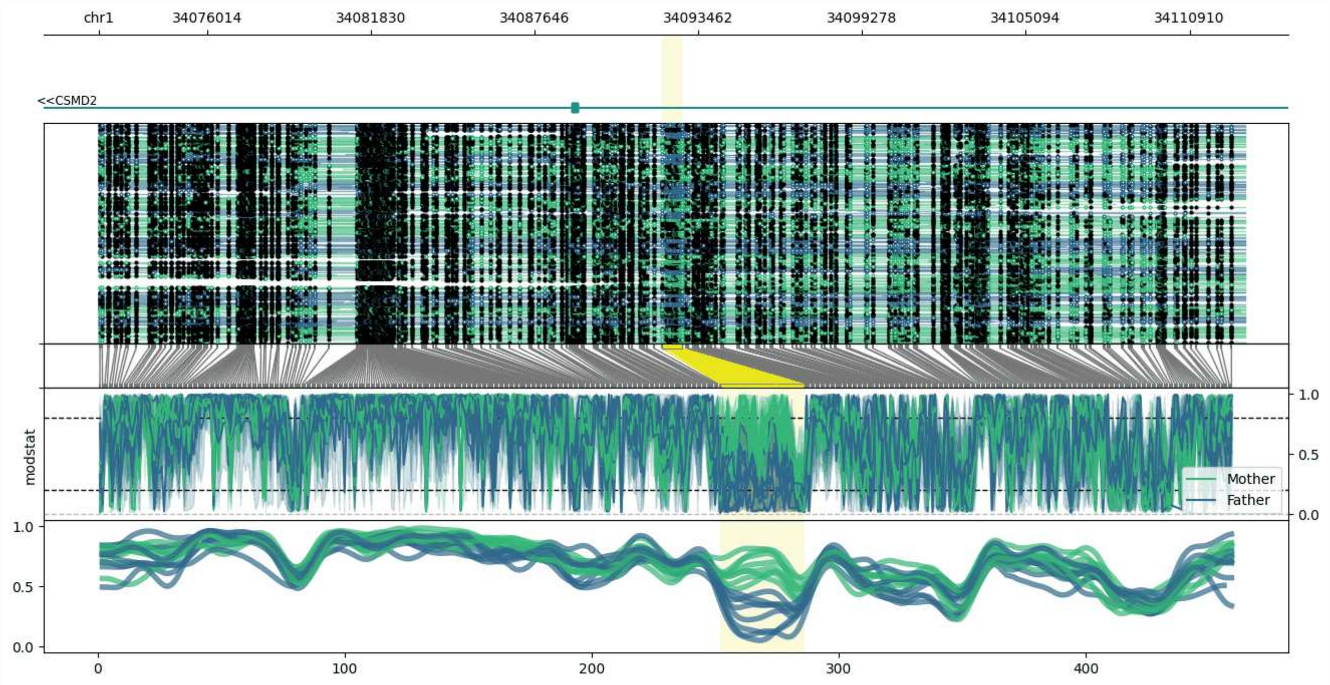


**chr1:34072171-34112885:** Region with 1 DMR containing genes: CSMD2, RNA5SP42.

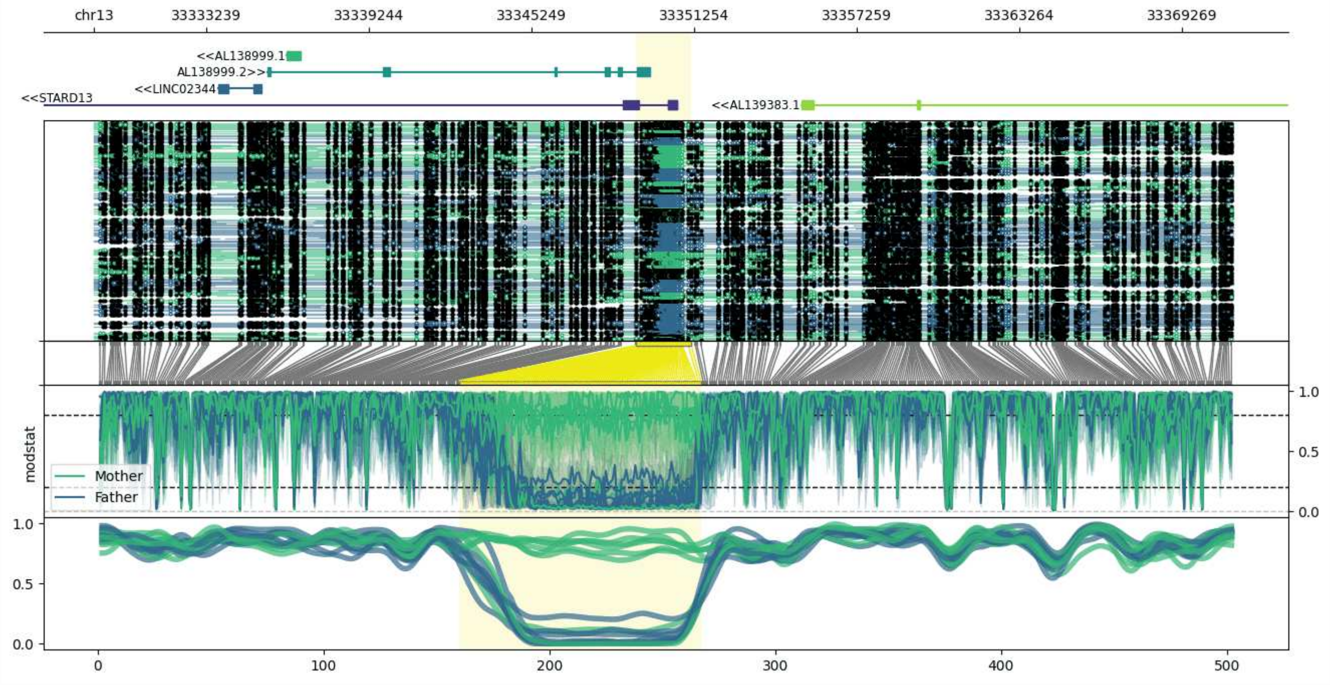


**chr13:33329111-33371152:** Region with 1 DMR containing genes: AL138999.1, AL138999.2, AL139383.1, LINC02344, STARD13.

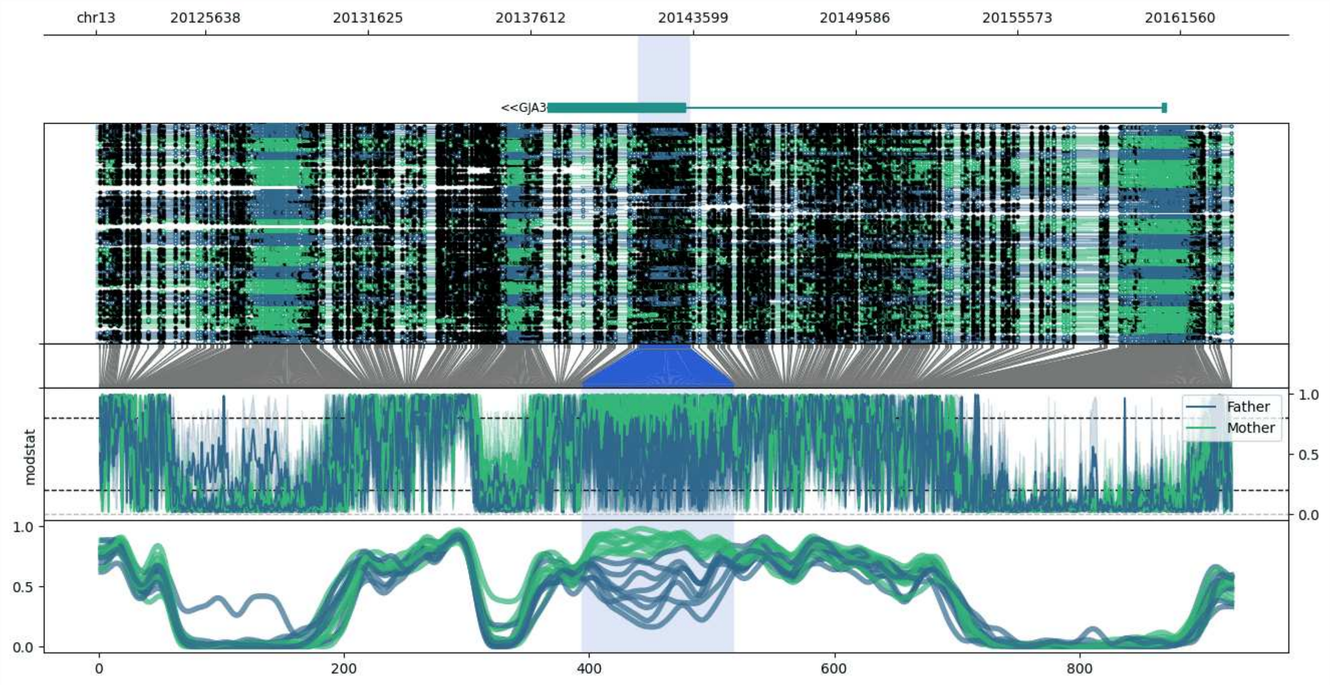


**chr13:20121573-20163482:** Region with 1 DMR containing genes: GJA3.

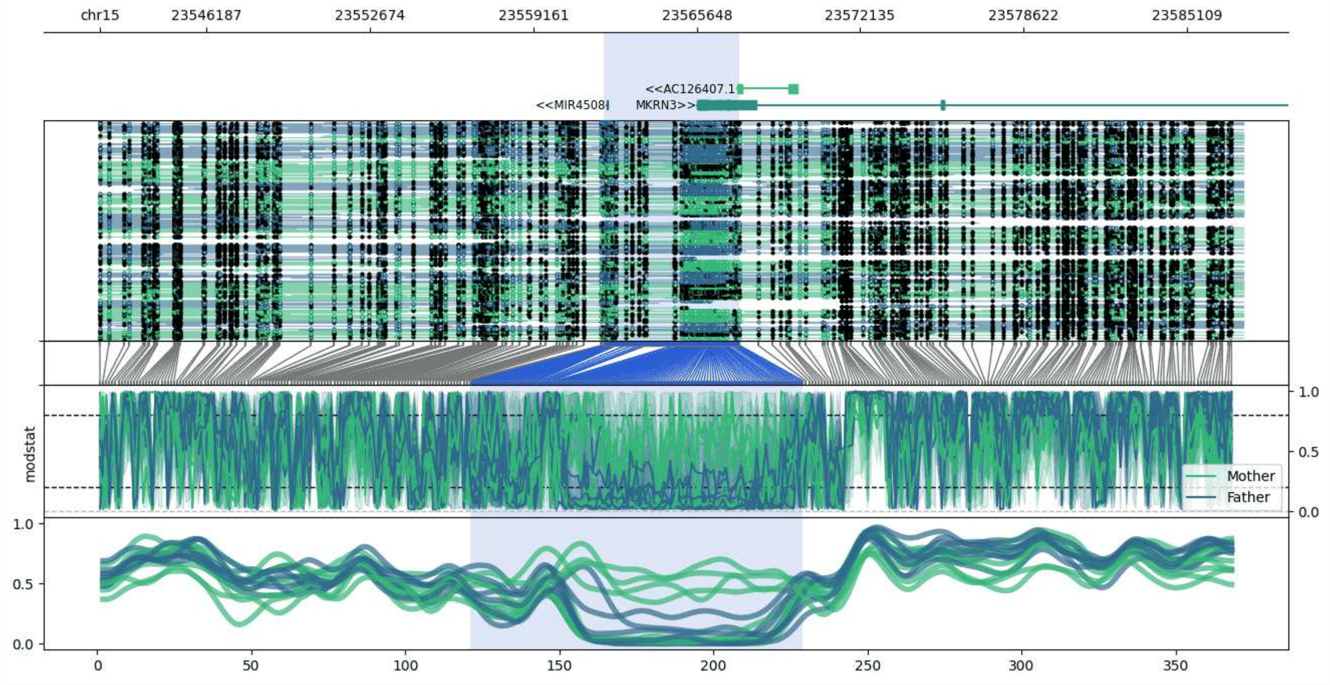


**chr15:23541939-23587348:** Region with 1 DMR containing genes: AC126407.1, MIR4508, MKRN3.

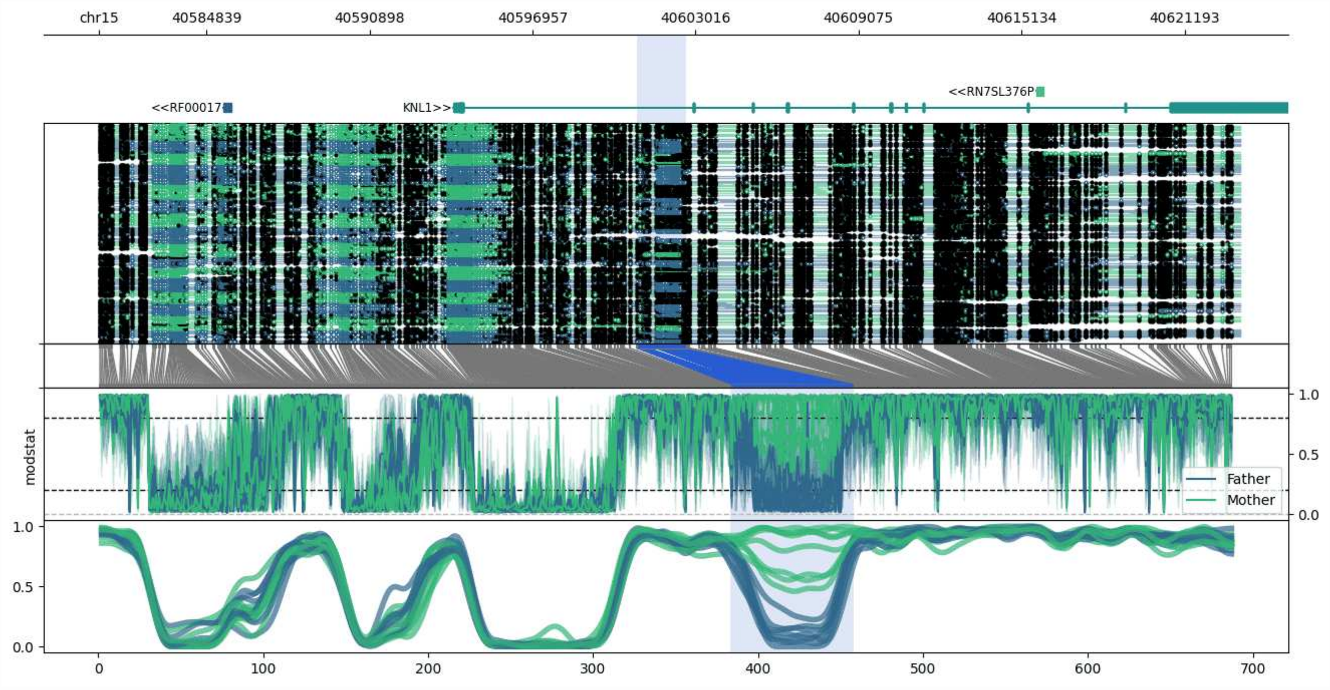


**chr15:40580842-40623260:** Region with 1 DMR containing genes: KNL1, RF00017, RN7SL376P, RPUSD2.

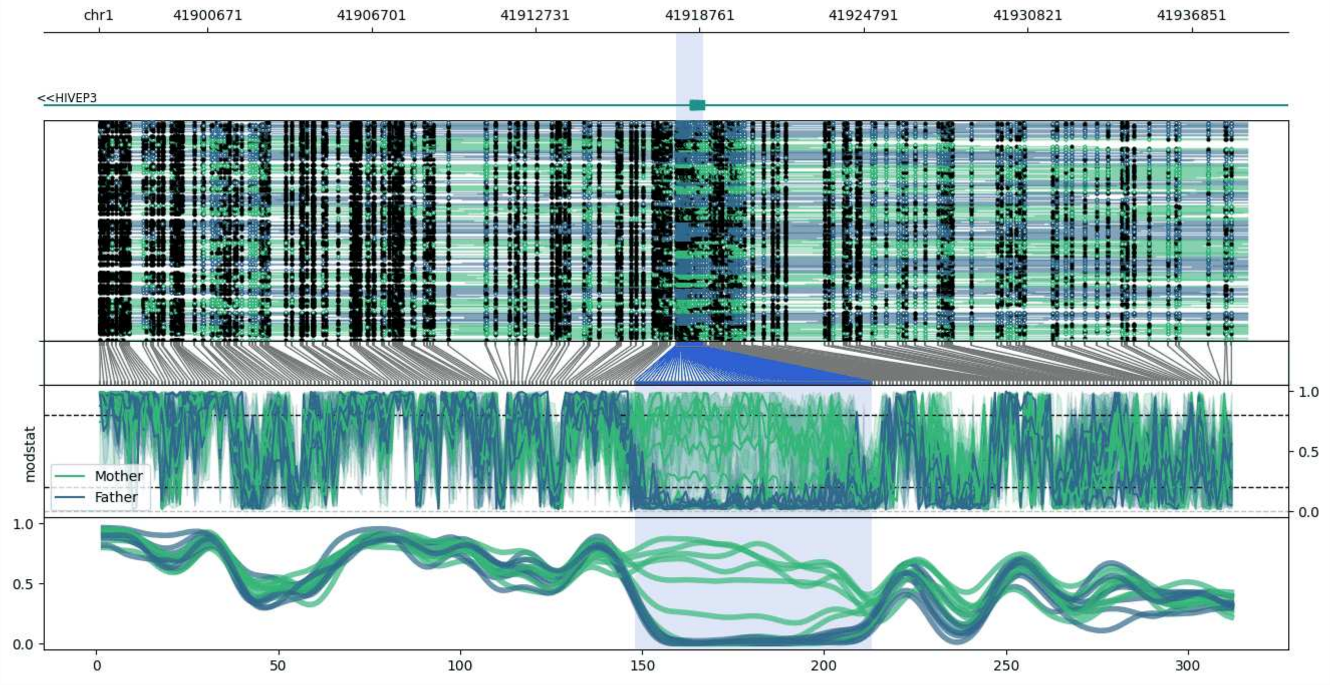


**chr1:41896681-41938892:** Region with 1 DMR containing genes: HIVEP3.

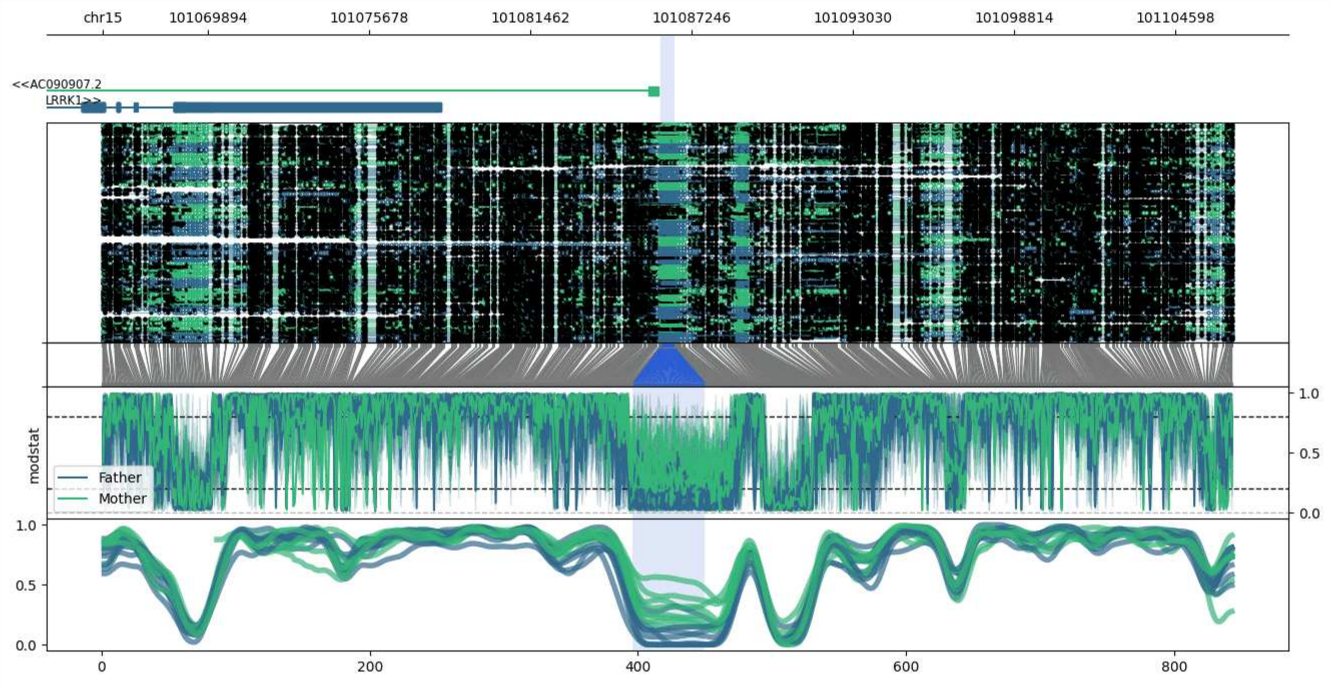


**chr15:101066130-101106619:** Region with 1 DMR containing genes: AC019254.2, AC090907.2, AC090907.3, LRRK1.

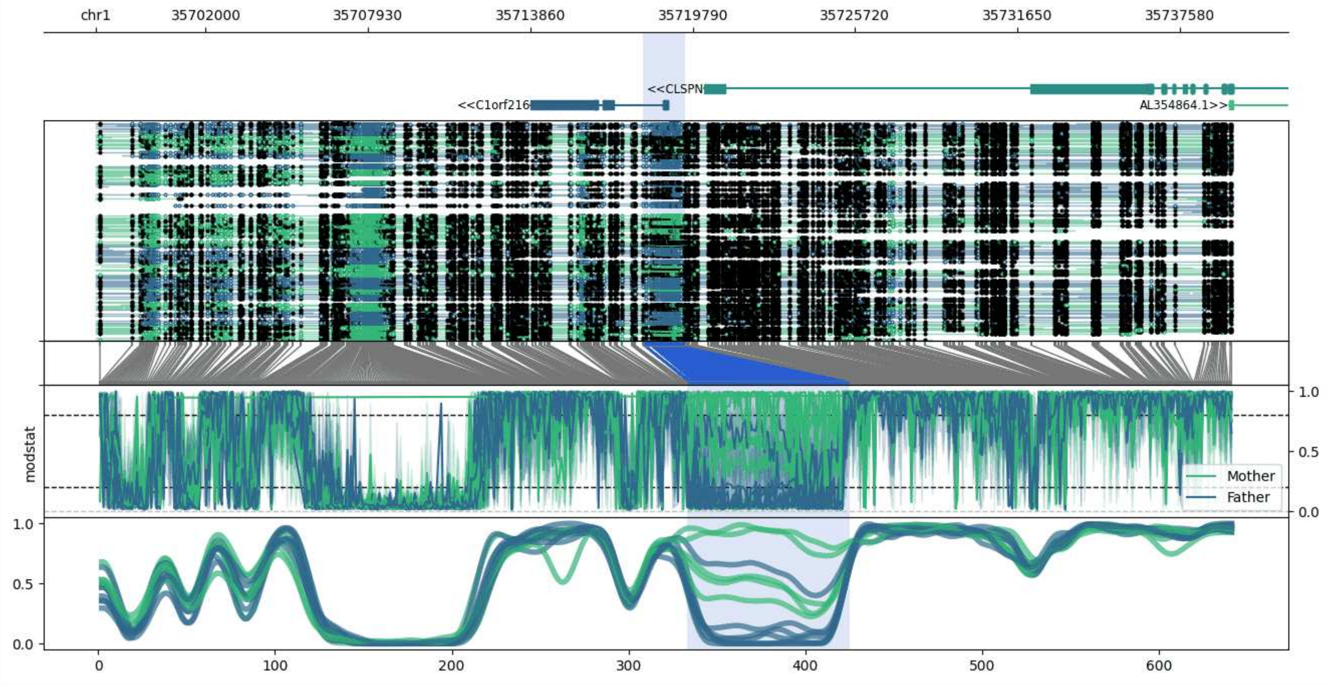


**chr1:35697984-35739497:** Region with 1 DMR containing genes: AL354864.1, C1orf216, CLSPN.

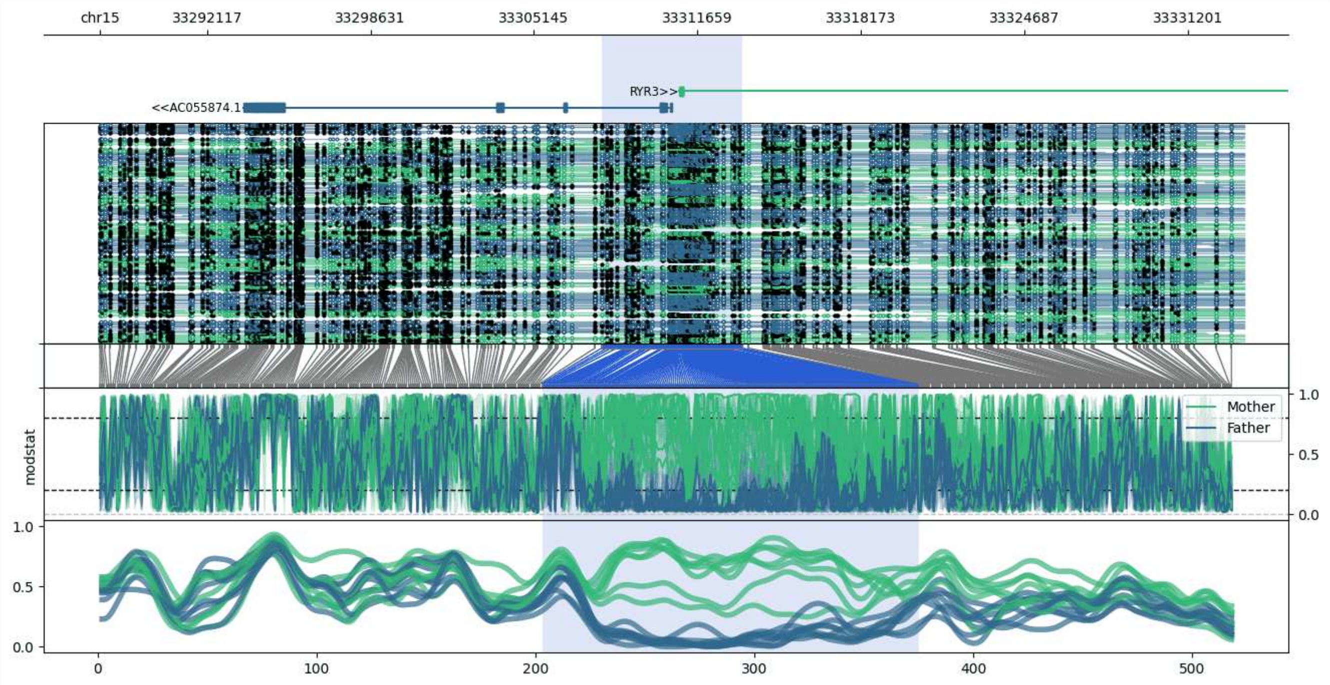


**chr15:33287841-33333444:** Region with 1 DMR containing genes: AC055874.1, RYR3.

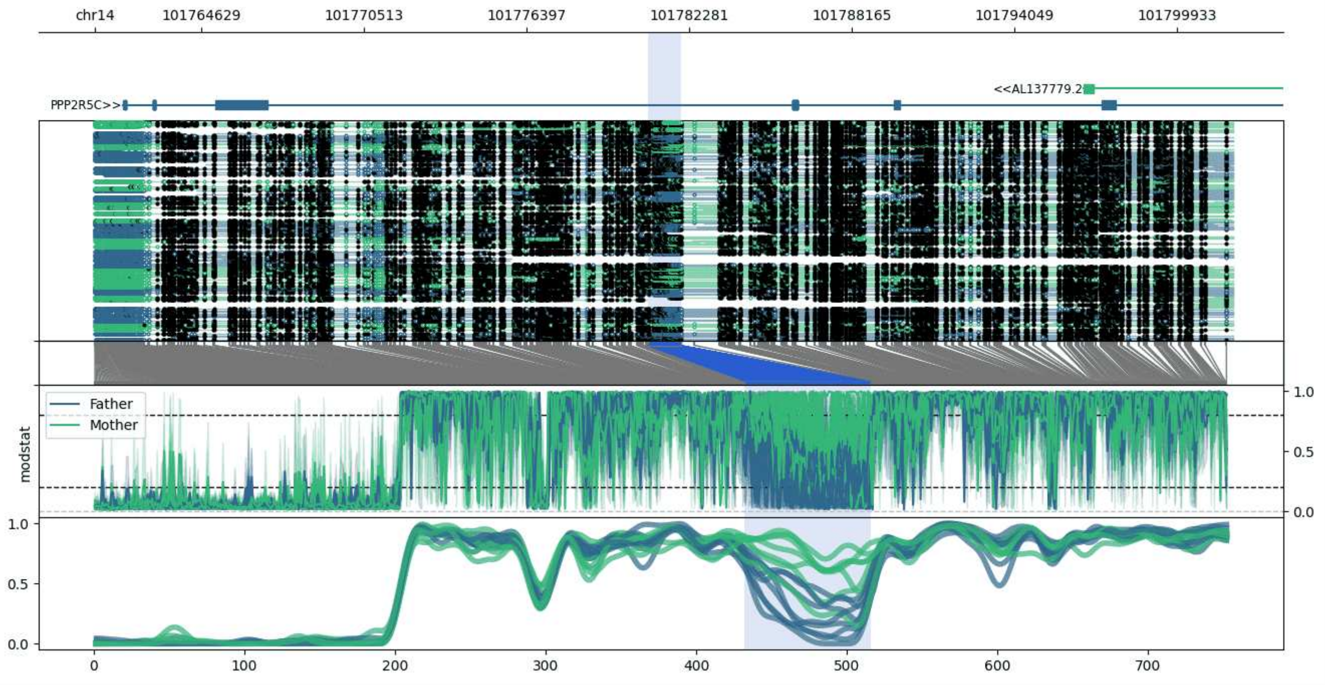


**chr14:101760783-101801972:** Region with 1 DMR containing genes: AL137779.2, PPP2R5C, RNU6-790P.

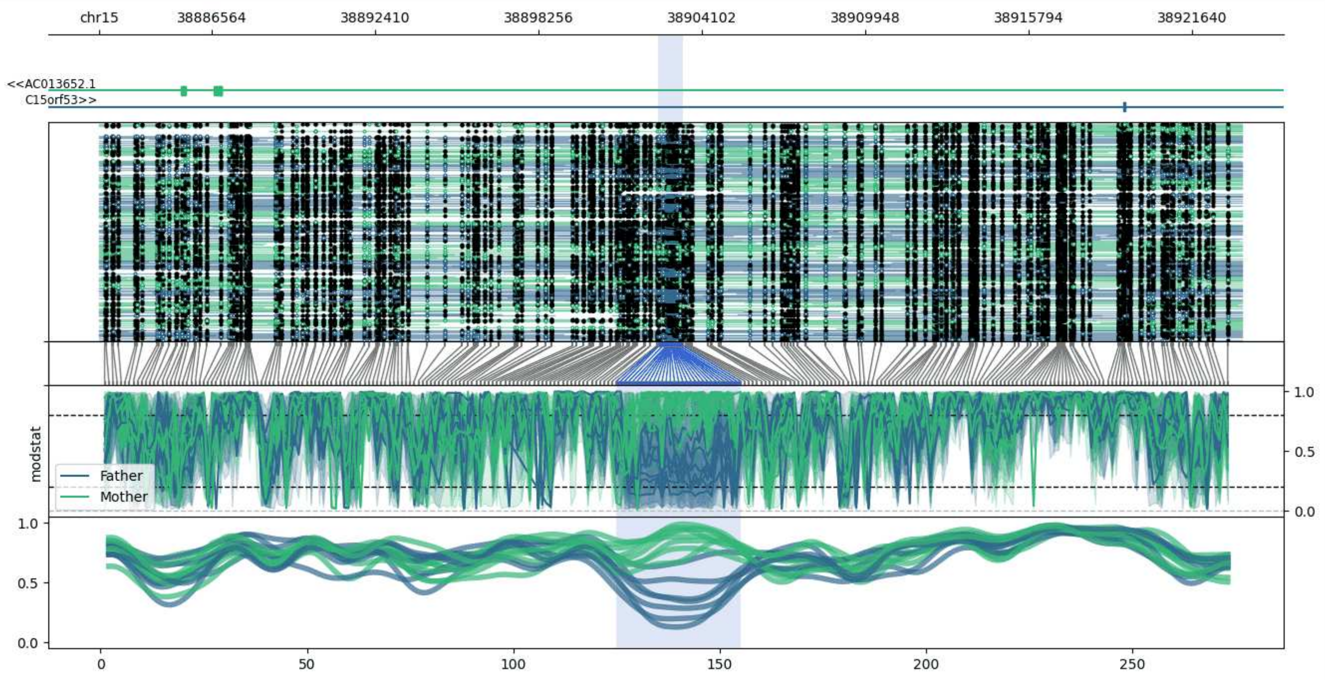


**chr15:38882519-38923444:** Region with 1 DMR containing genes: AC013652.1, C15orf53.

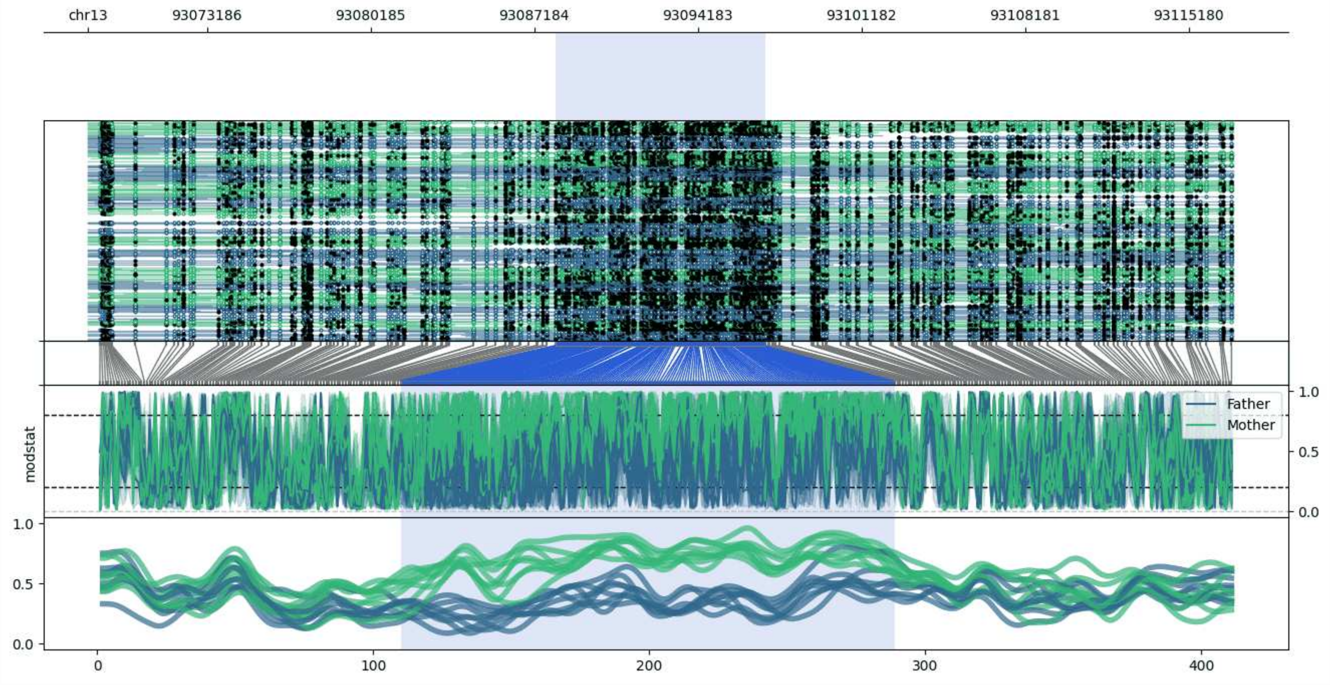


**chr13:93068085-93117079:** Region with 1 DMR containing no genes.

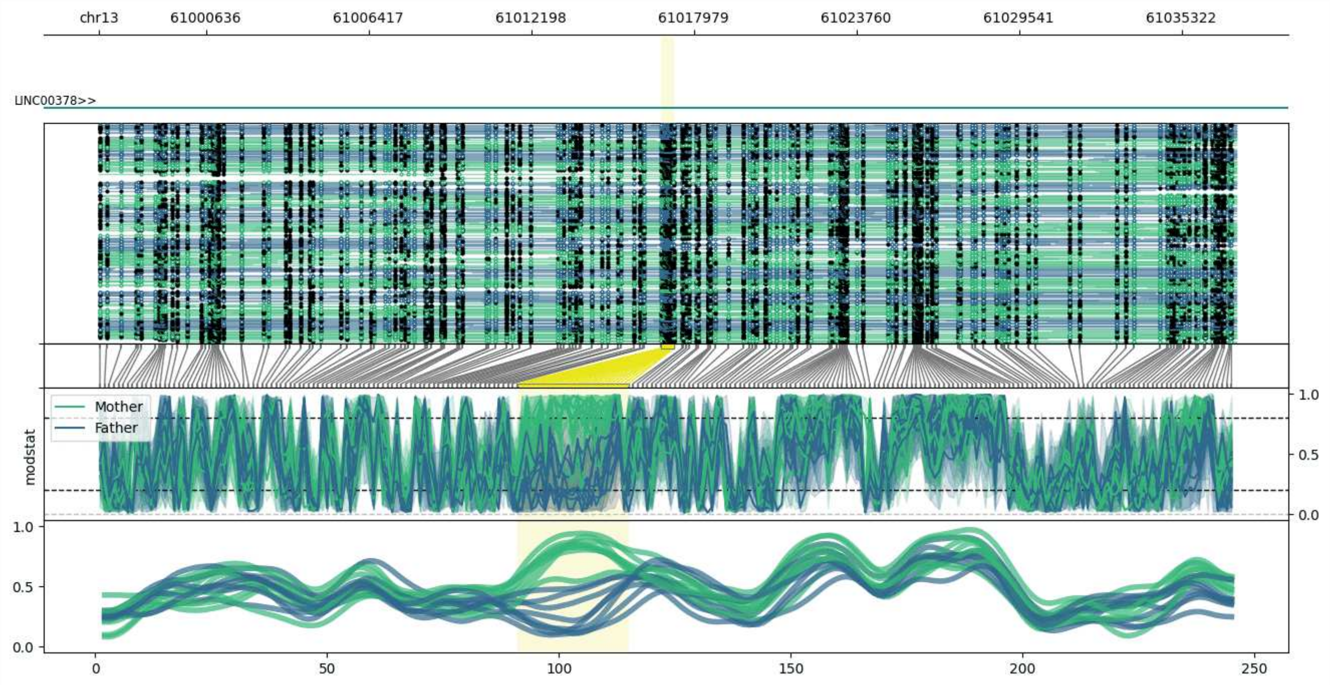


**chr13:60996815-61037282:** Region with 1 DMR containing genes: LINC00378.

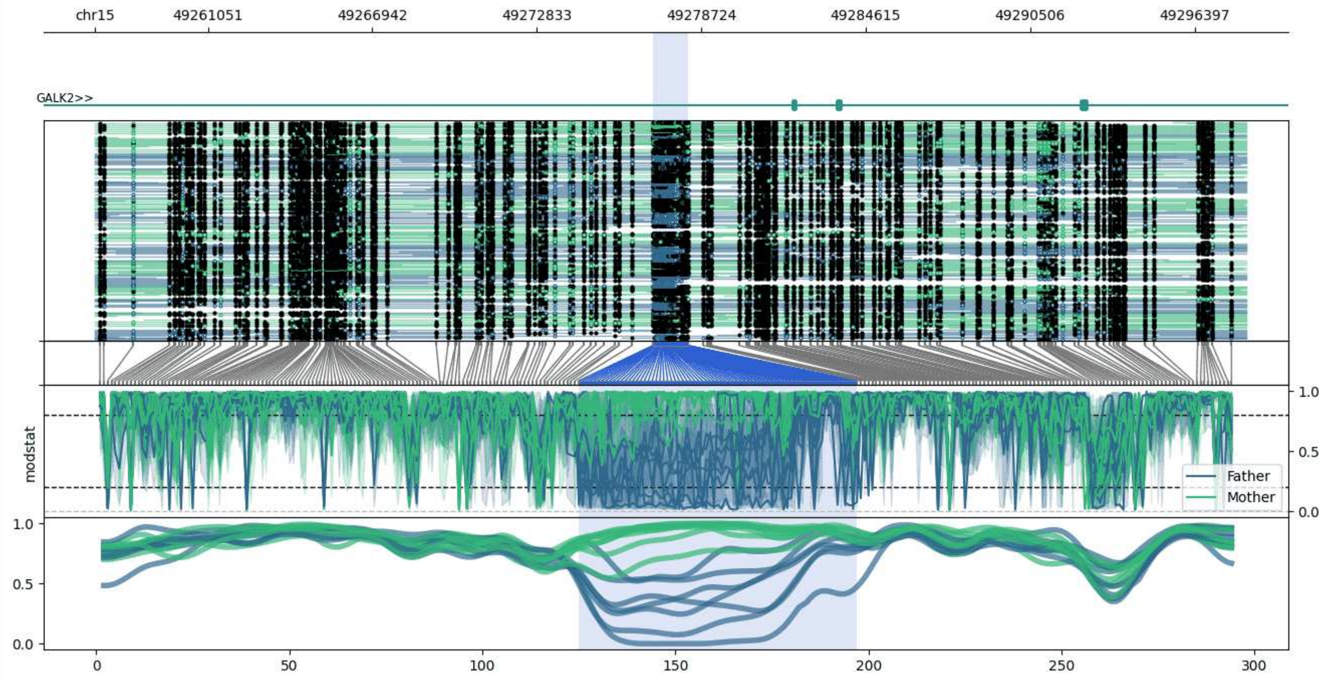


**chr15:49256999-49298238:** Region with 1 DMR containing genes: GALK2.

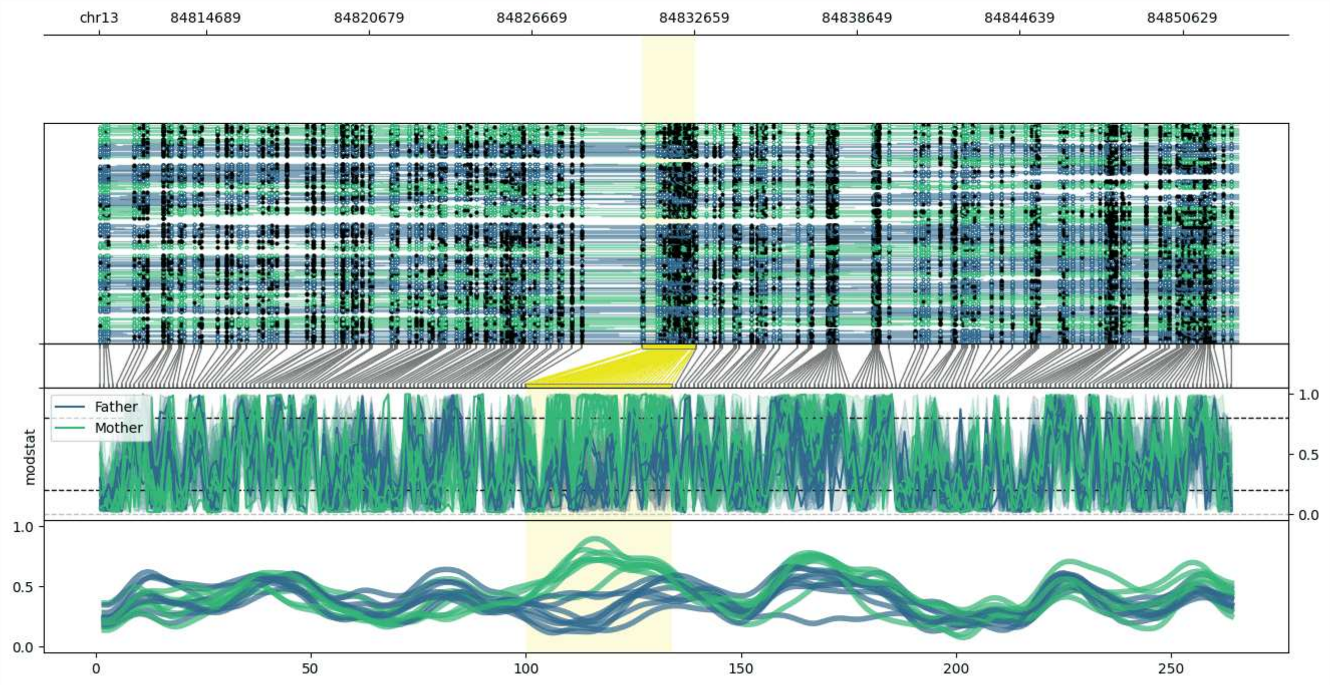


**chr13:84810744-84852678:** Region with 1 DMR containing no genes.

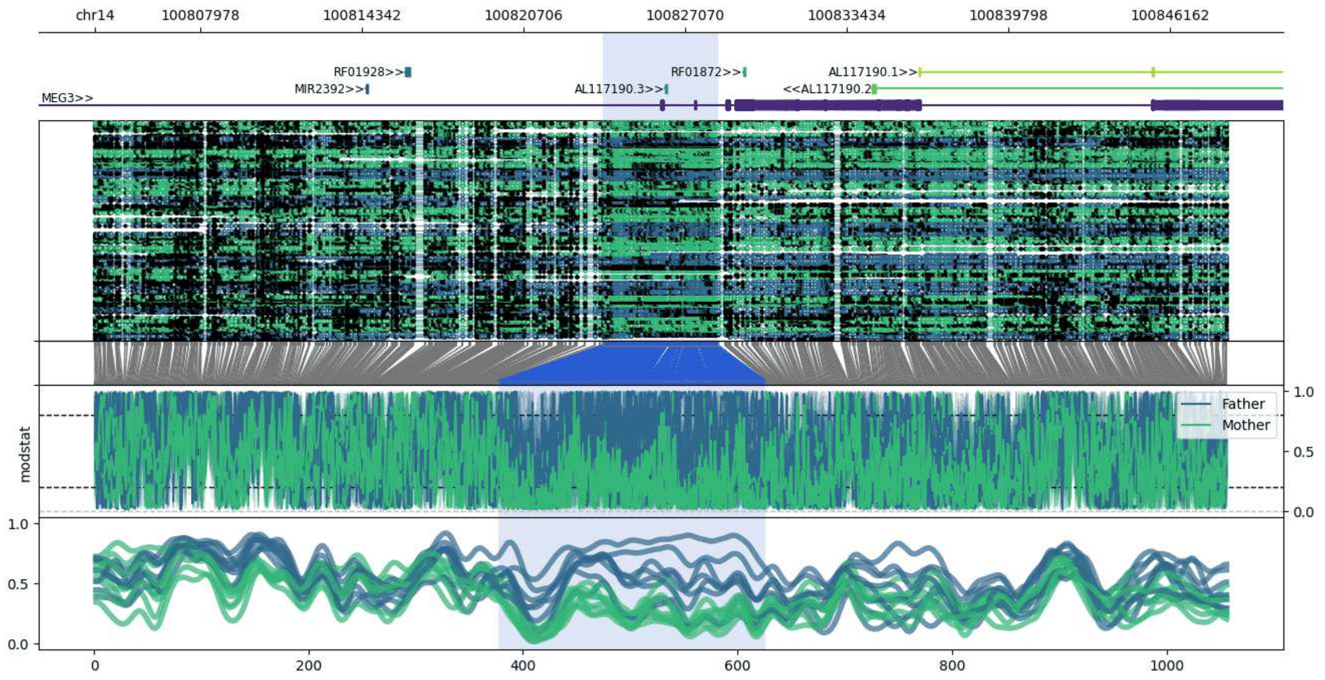


**chr14:100803840-100848392:** Region with 1 DMR containing genes: AL117190.1, AL117190.2, AL117190.3, MEG3, MIR2392, MIR770, RF01872, RF01928.

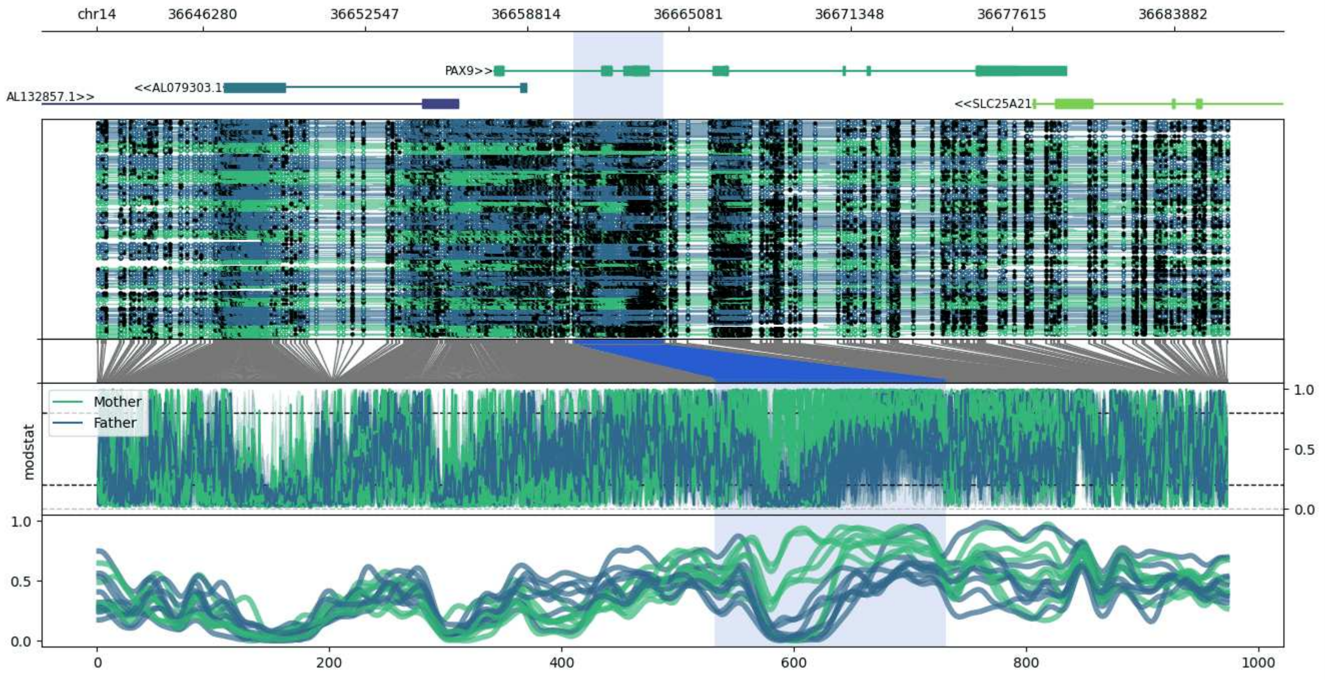


**chr14:36642143-36686016:** Region with 1 DMR containing genes: AL079303.1, AL132857.1, PAX9, SLC25A21.

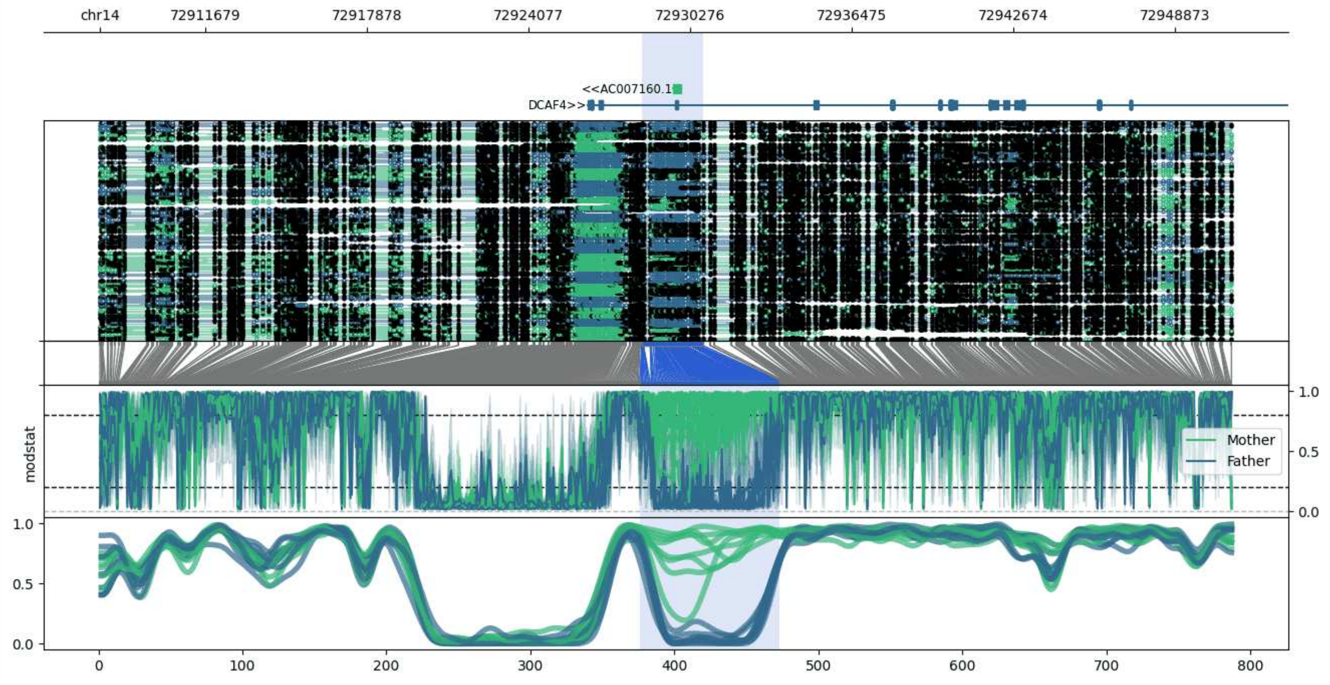


**chr14:72907642-72951035:** Region with 1 DMR containing genes: AC007160.1, AL442663.4, DCAF4.

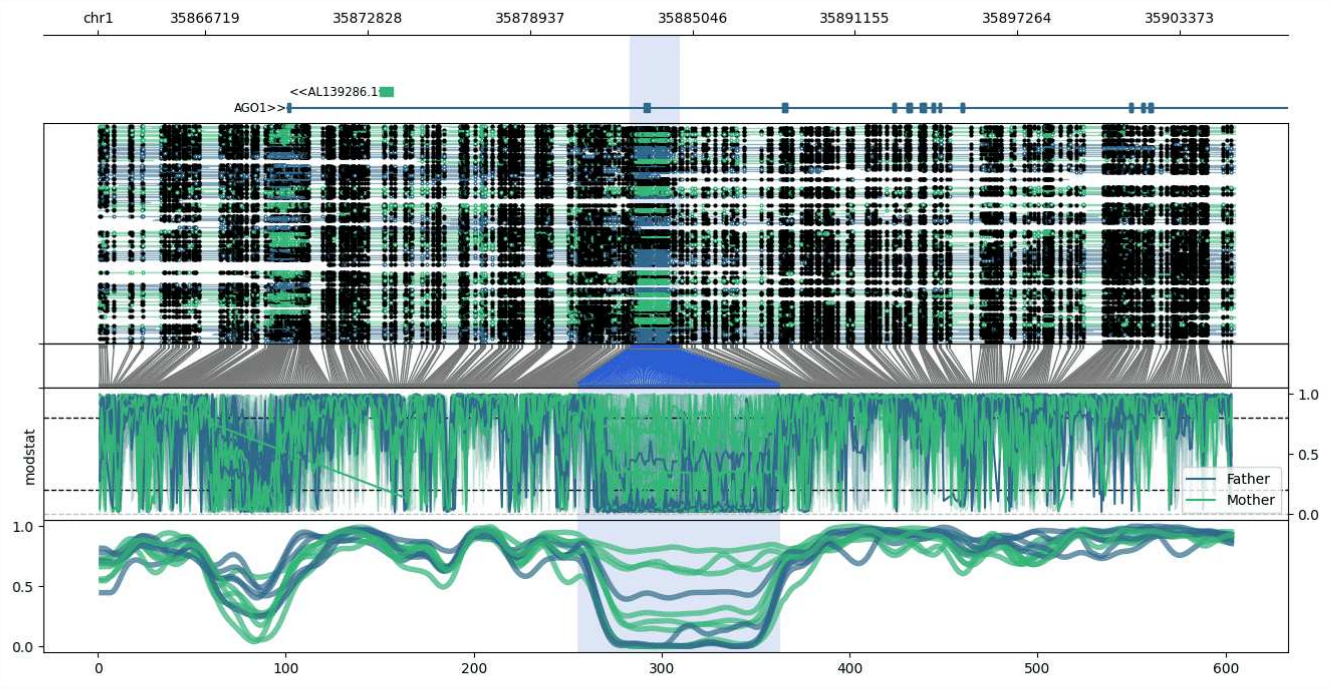


**chr1:35862674-35905440:** Region with 1 DMR containing genes: AGO1, AGO4, AL139286.1, AL139286.3.

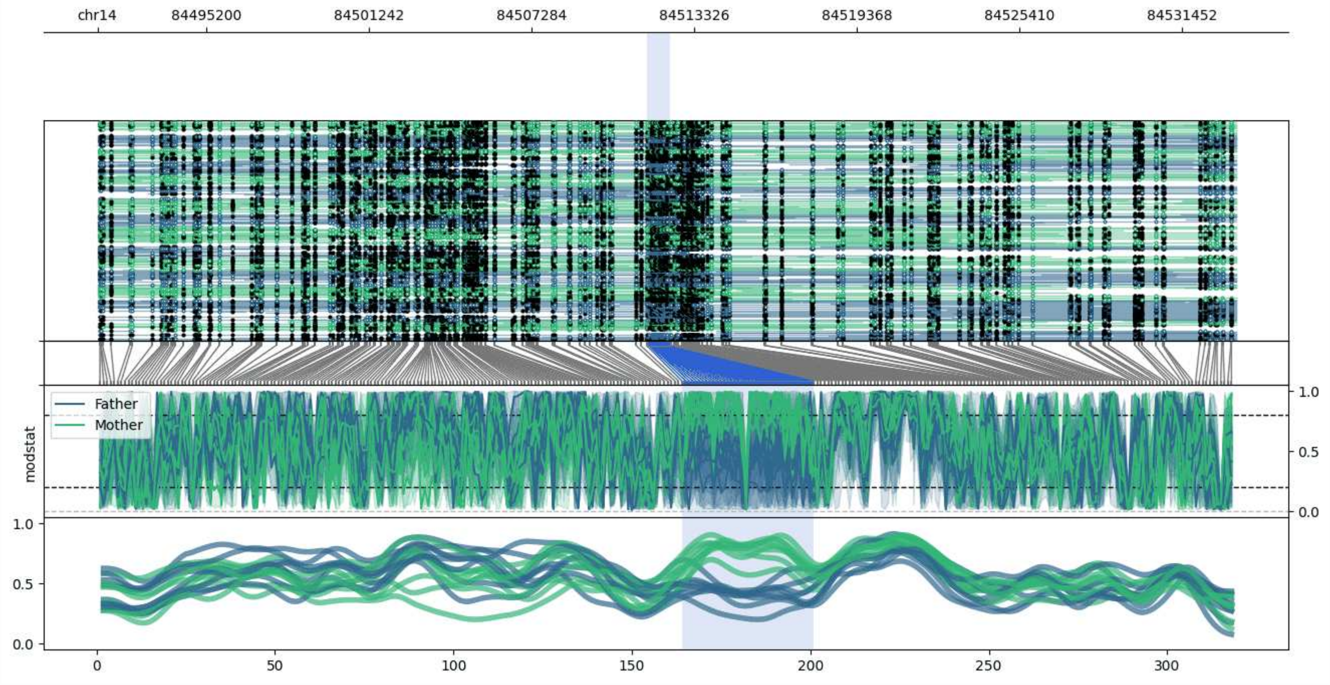


**chr14:84491164-84533459:** Region with 1 DMR containing no genes.

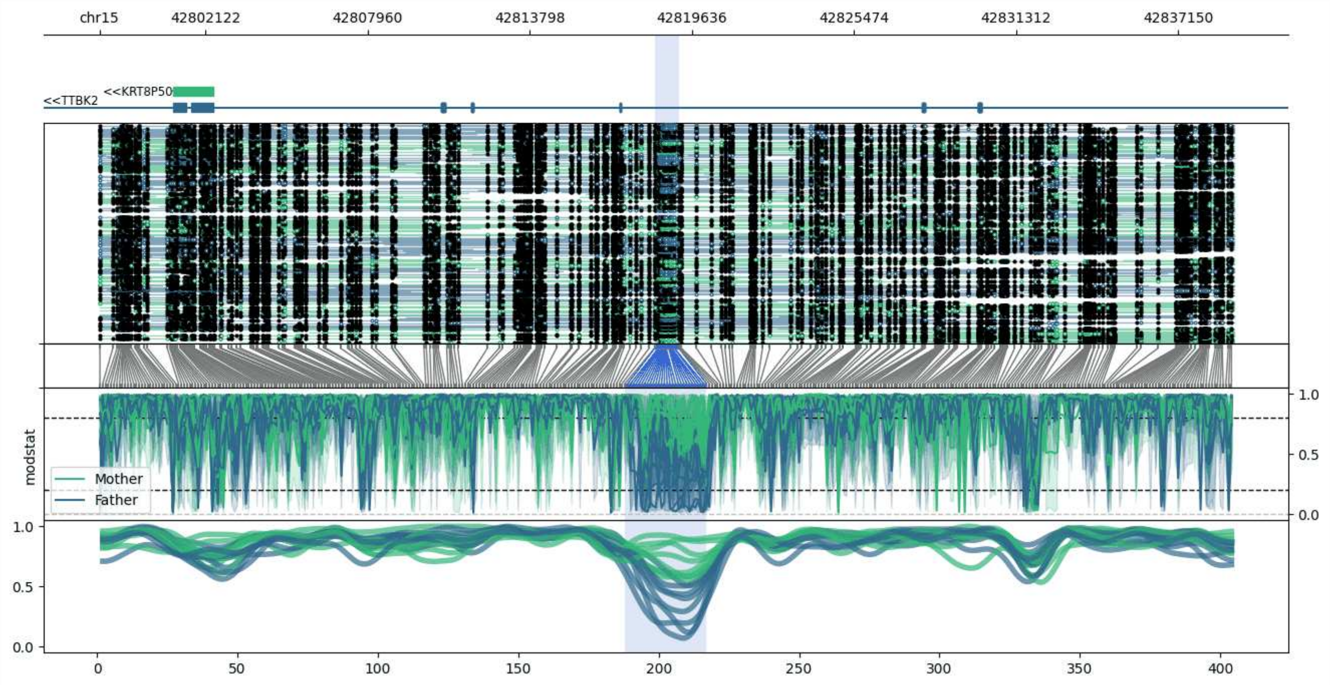


**chr15:42798301-42839170:** Region with 1 DMR containing genes: KRT8P50, TTBK2.

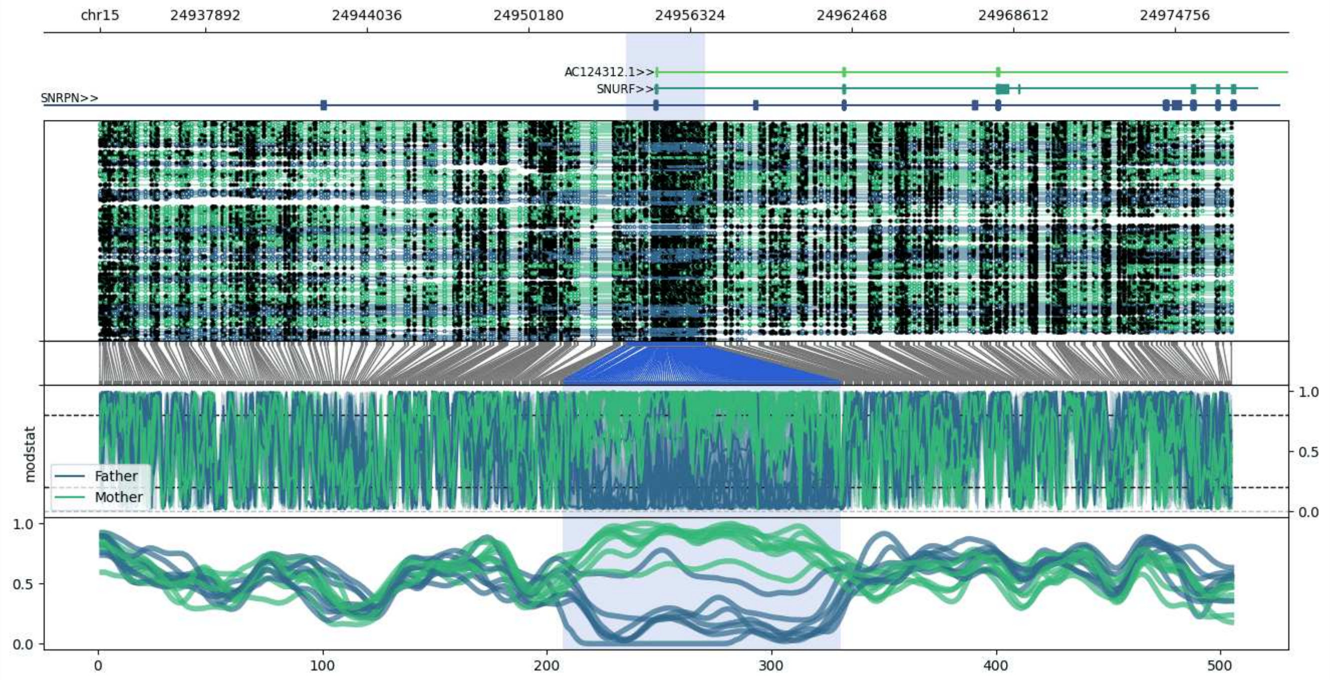


**chr15:24933880-24976894:** Region with 1 DMR containing genes: AC124312.1, PWAR5, SNHG14, SNORD107, SNORD64, SNRPN, SNURF.

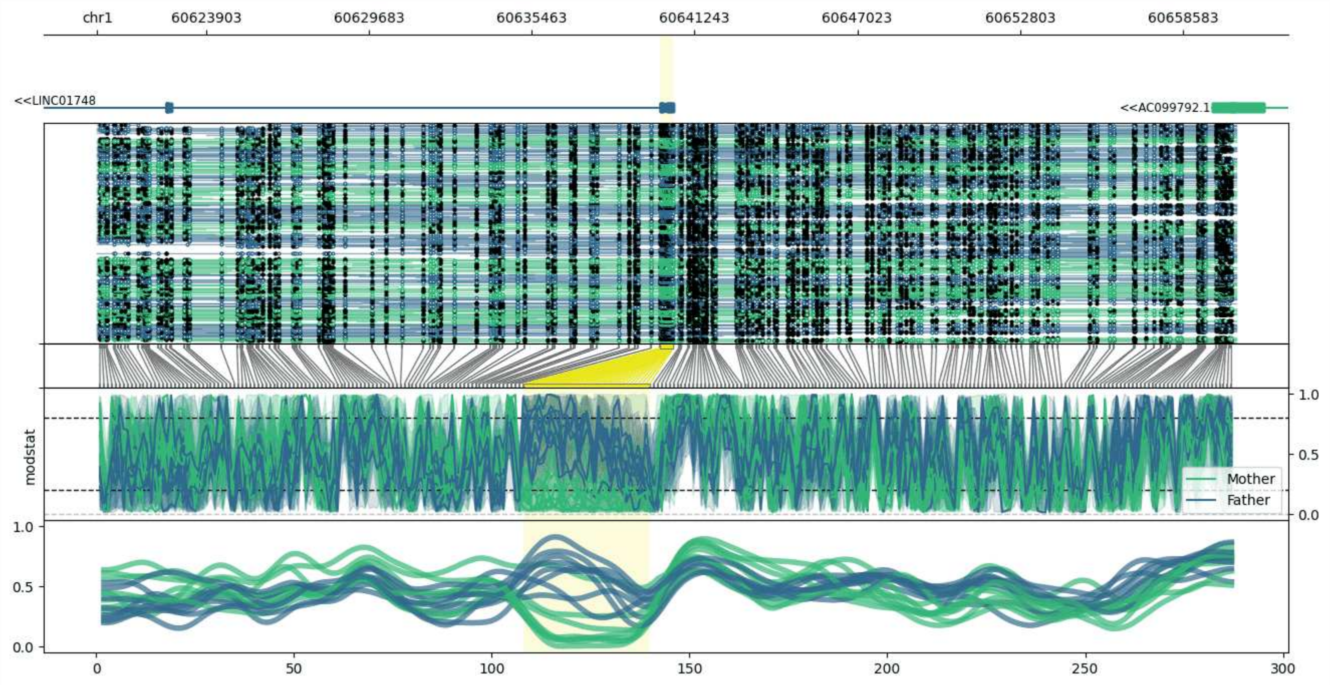


**chr1:60620031-60660492:** Region with 1 DMR containing genes: AC099792.1, LINC01748.

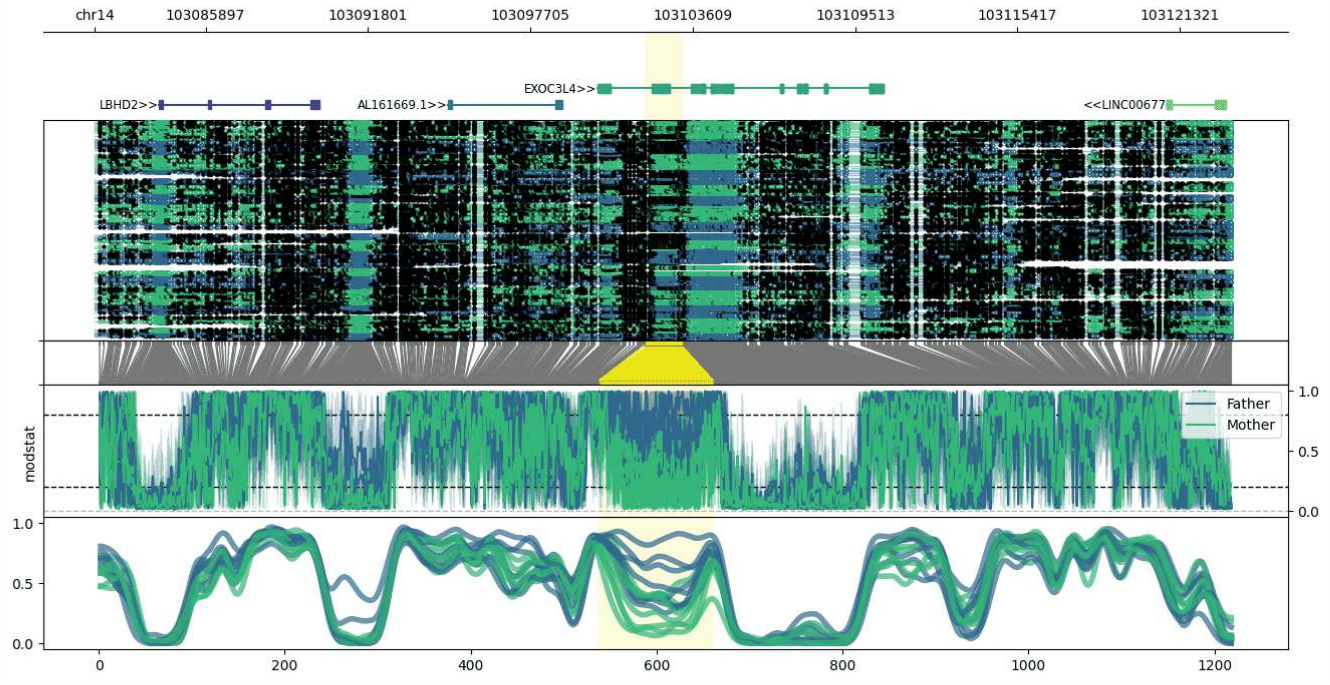


**chr14:103081857-103123191:** Region with 1 DMR containing genes: AL161669.1, EXOC3L4, LBHD2, LINC00677, TNFAIP2.

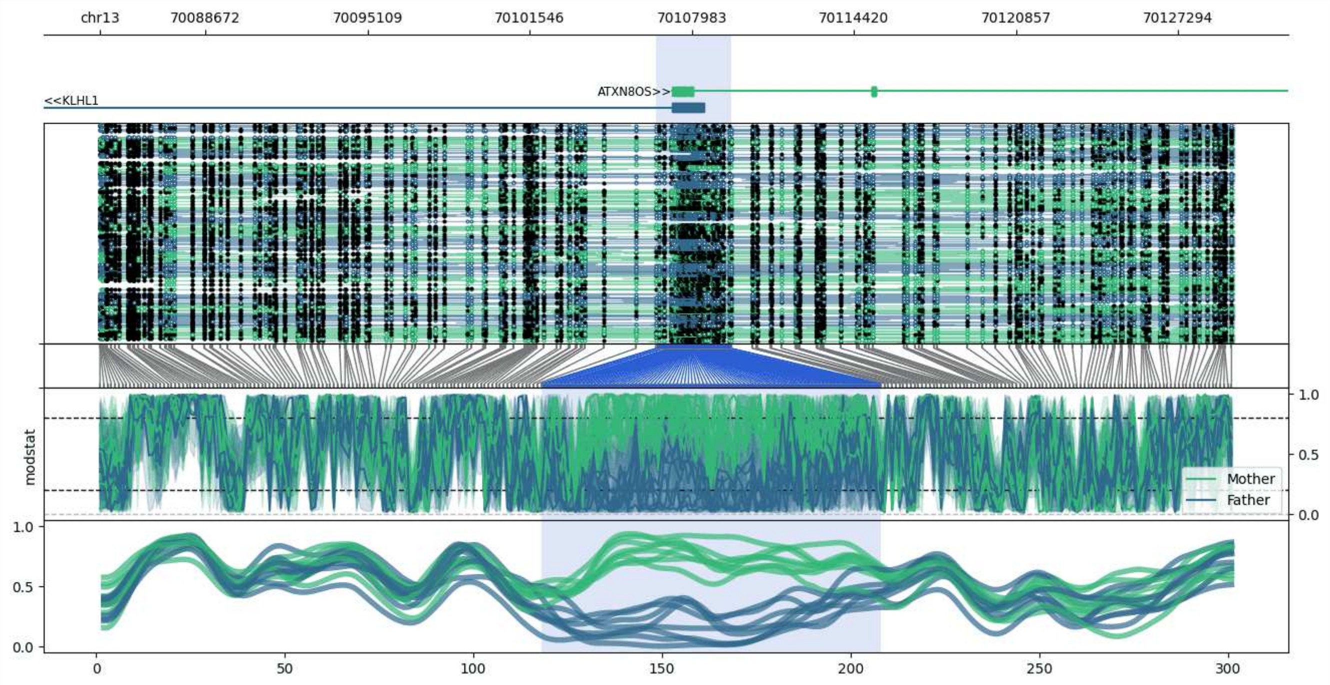


**chr13:70084477-70129539:** Region with 1 DMR containing genes: ATXN8OS, KLHL1.

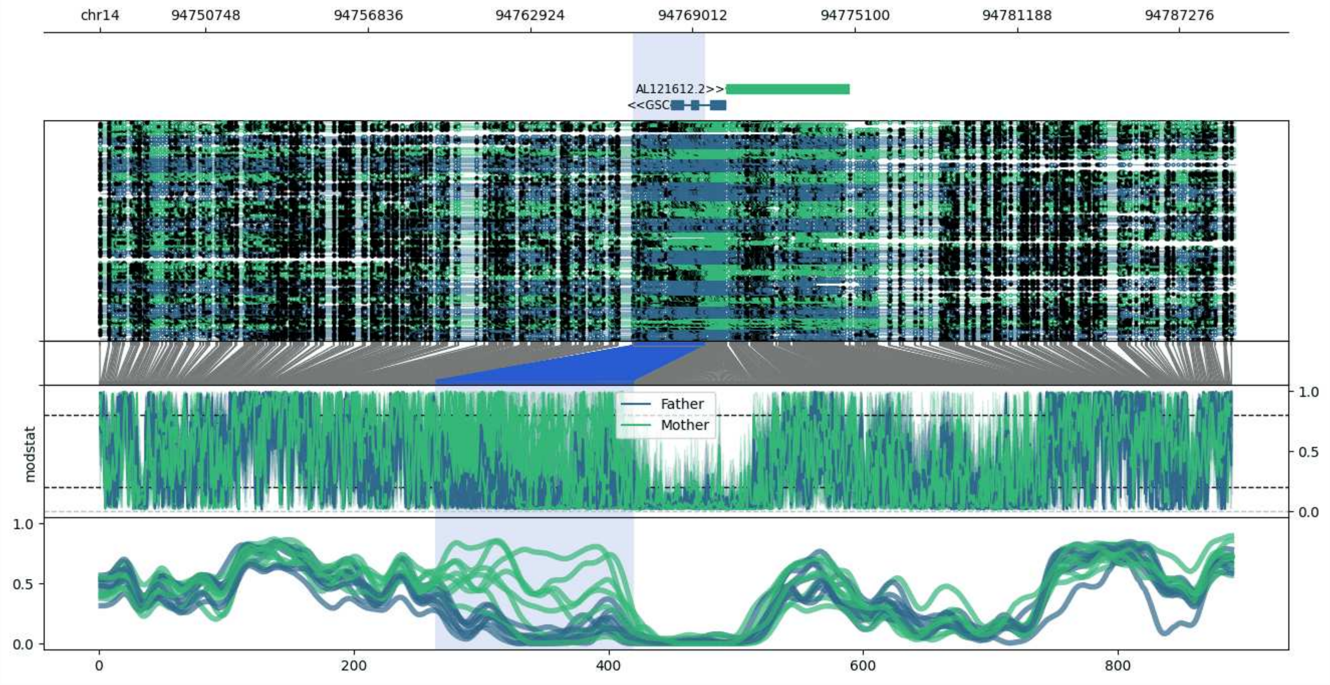


**chr14:94746772-94789393:** Region with 1 DMR containing genes: AL121612.2, GSC.

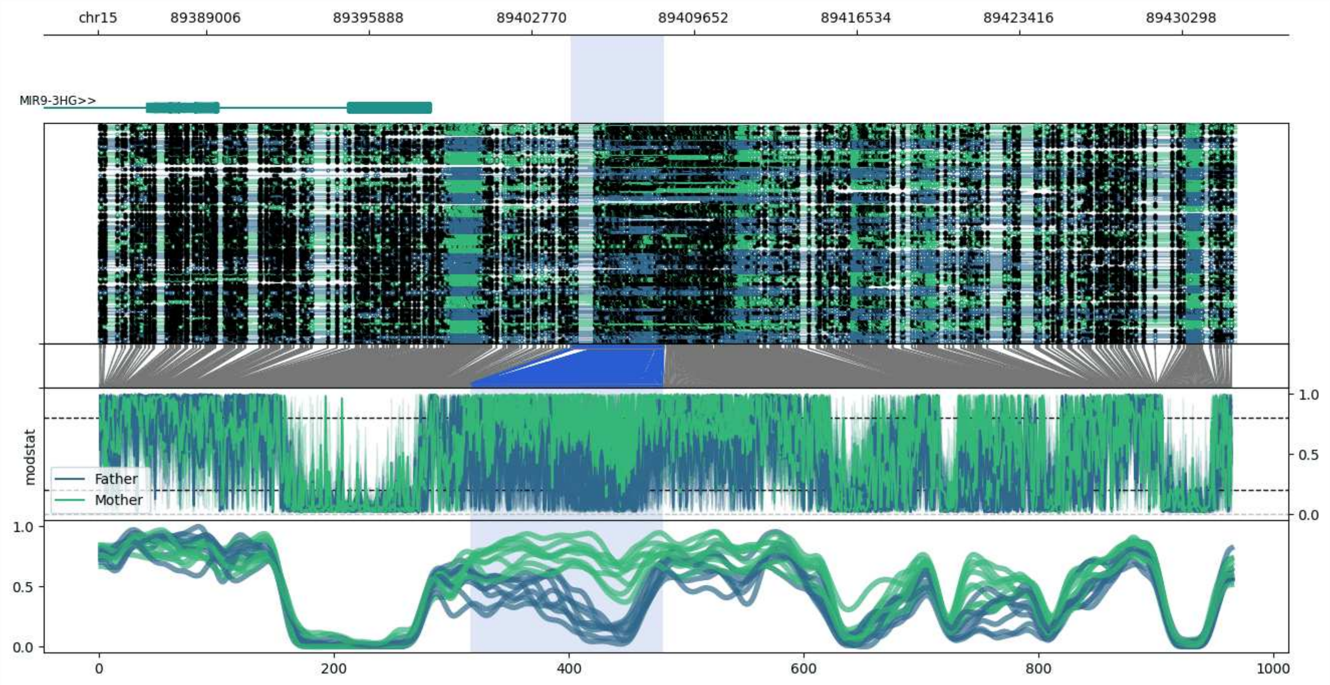


**chr15:89384434-89432608:** Region with 1 DMR containing genes: MIR9-3HG.

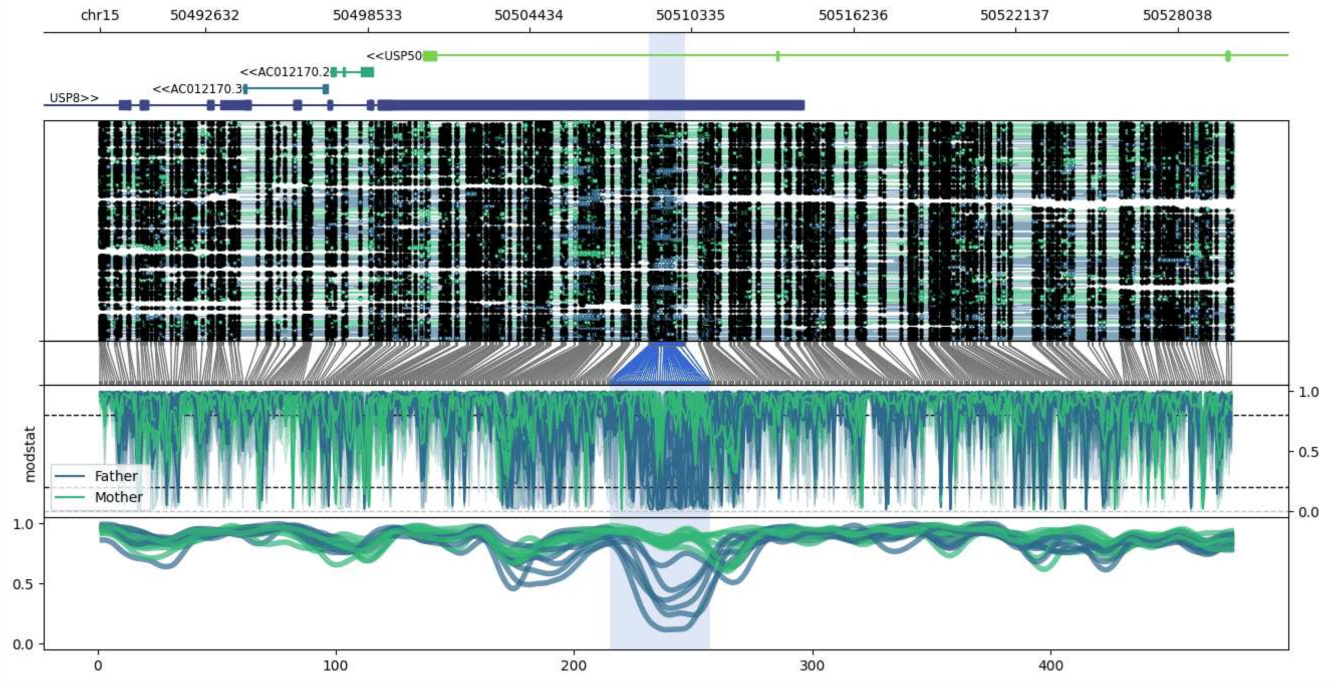


**chr15:50488776-50530089:** Region with 1 DMR containing genes: AC012170.2, AC012170.3, USP50, USP8.

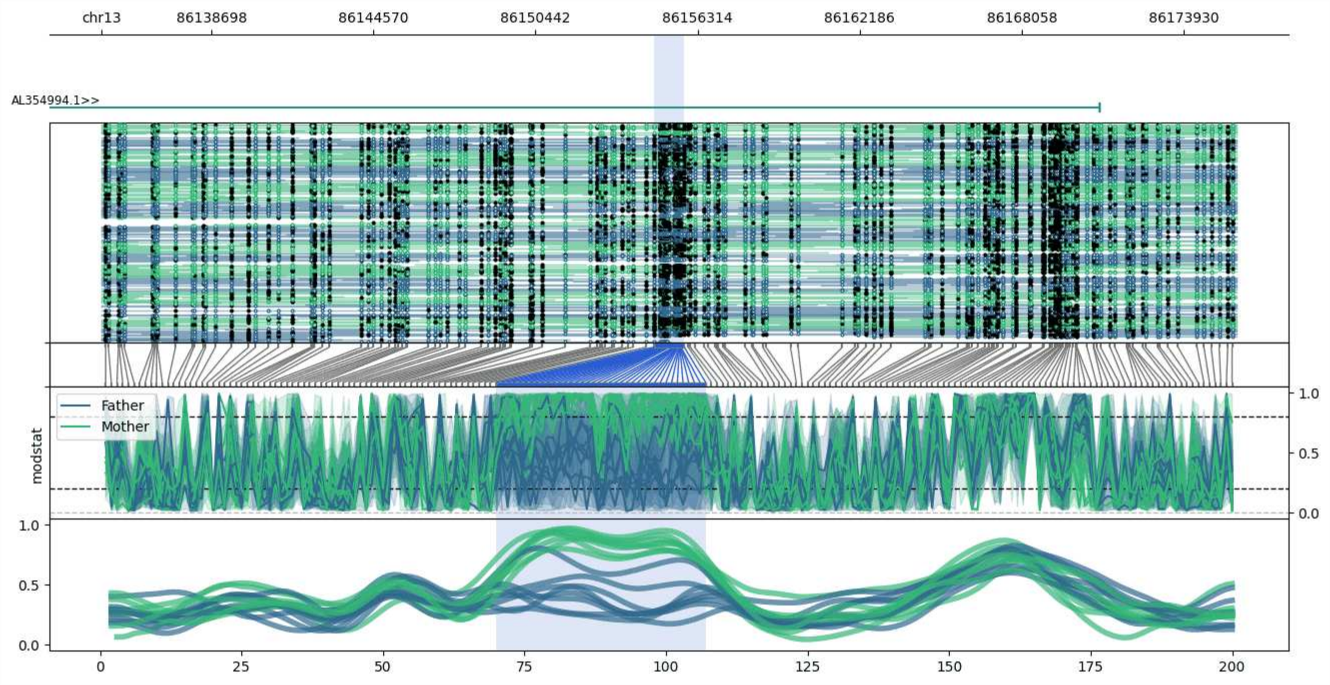


**chr13:86134723-86175831:** Region with 1 DMR containing genes: AL354994.1.

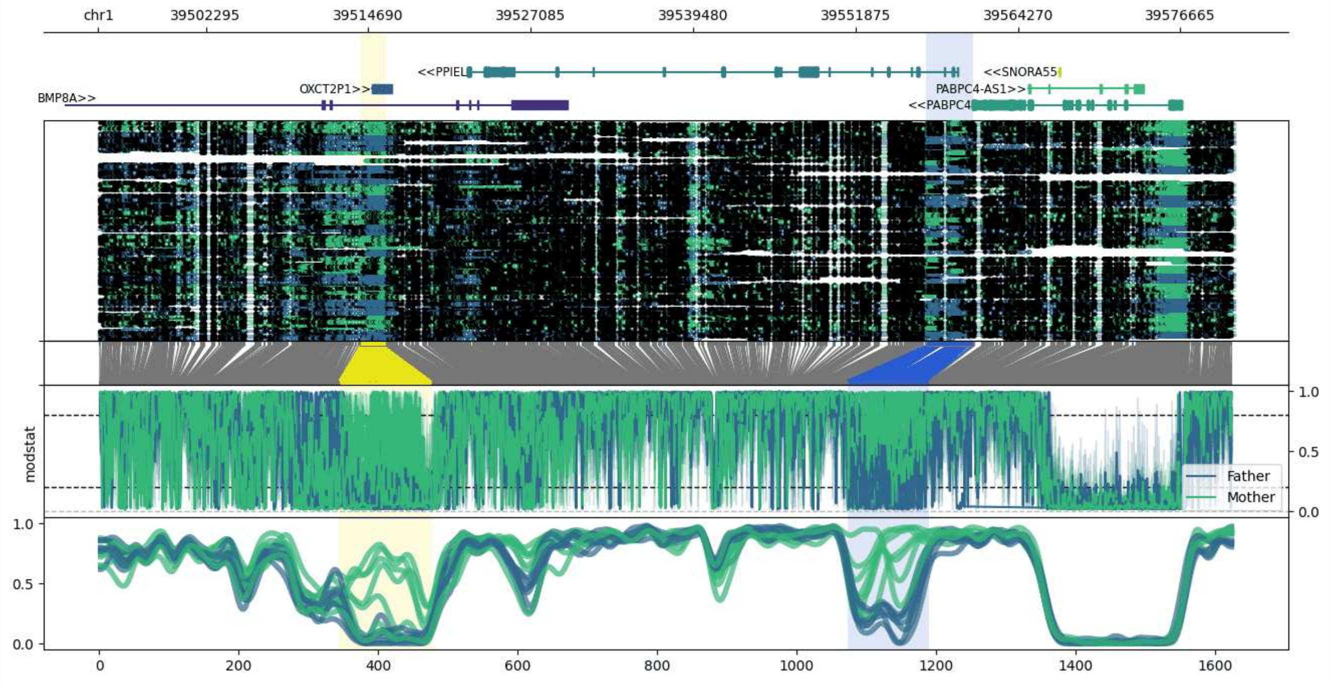


**chr1:39494077-39580848:** Region with 2 DMRs containing genes: BMP8A, MACF1, OXCT2P1, PABPC4, PABPC4-AS1, PPIEL, SNORA55.

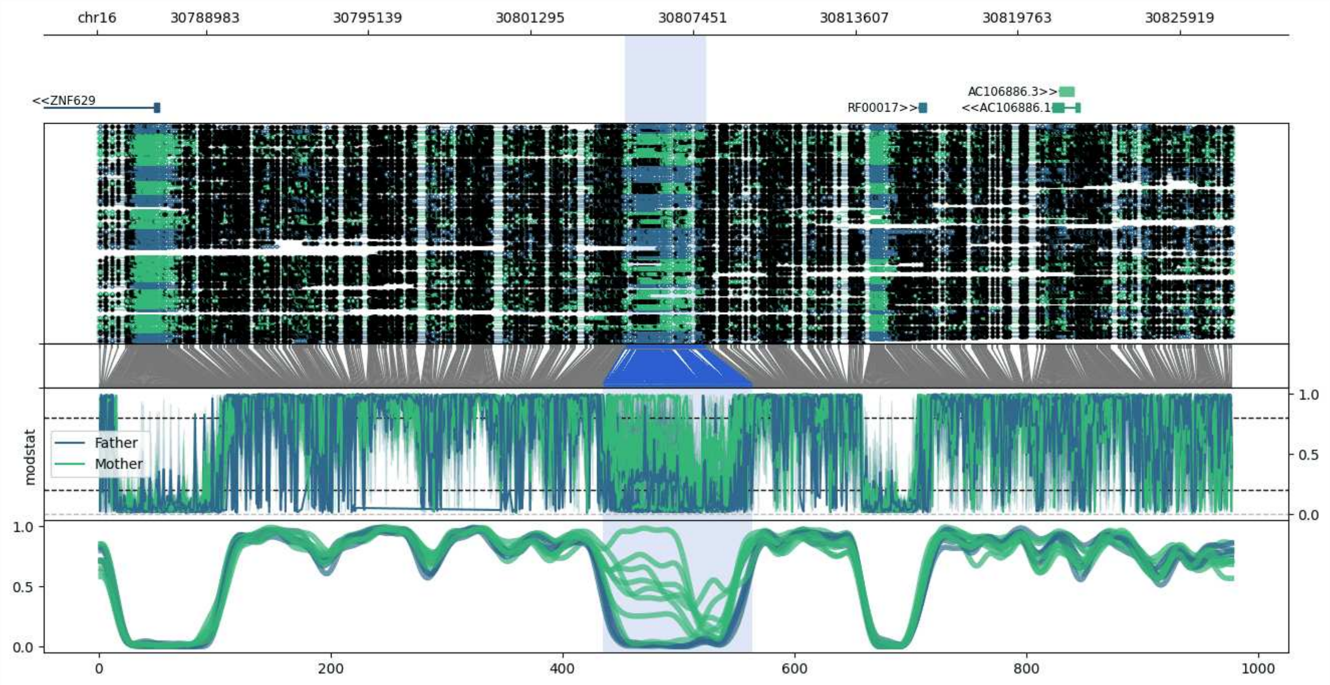


**chr16:30784867-30827960:** Region with 1 DMR containing genes: AC106886.1, AC106886.3, AC106886.4, BCL7C, RF00017, RNF40, ZNF629.

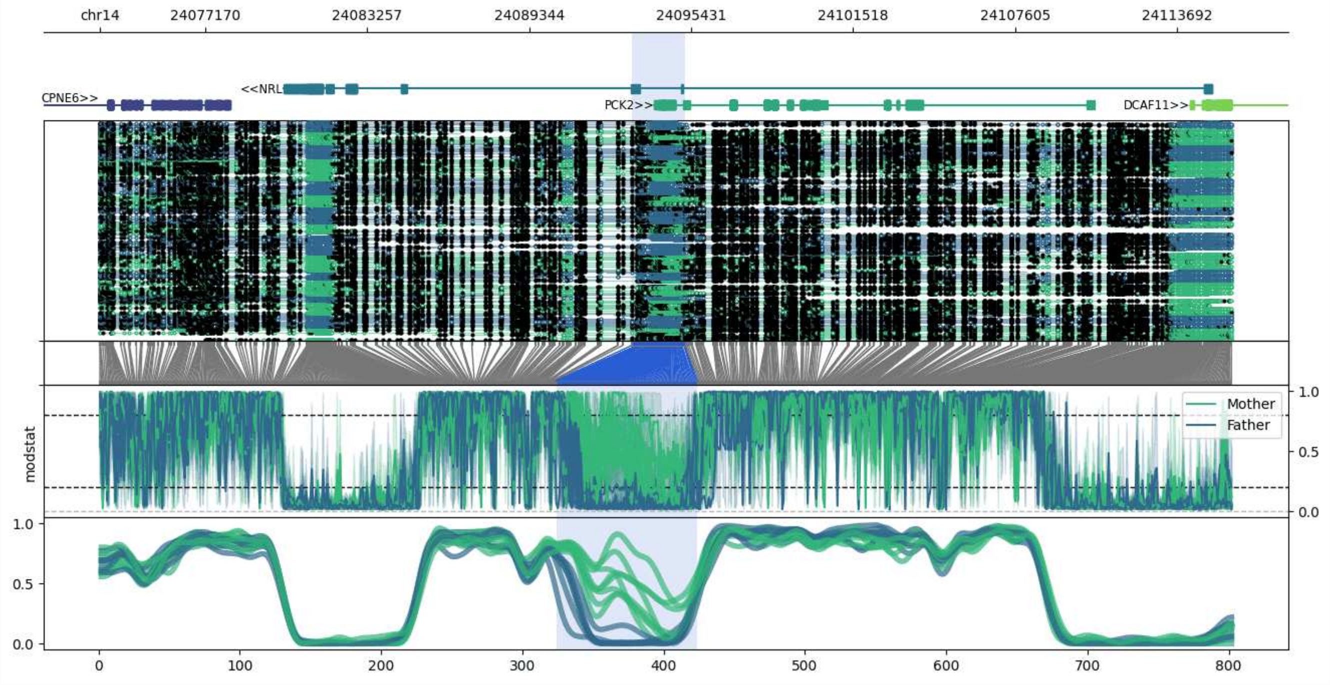


**chr14:24073193-24115803:** Region with 1 DMR containing genes: AL136295.3, CARMIL3, CPNE6, DCAF11, NRL, PCK2.

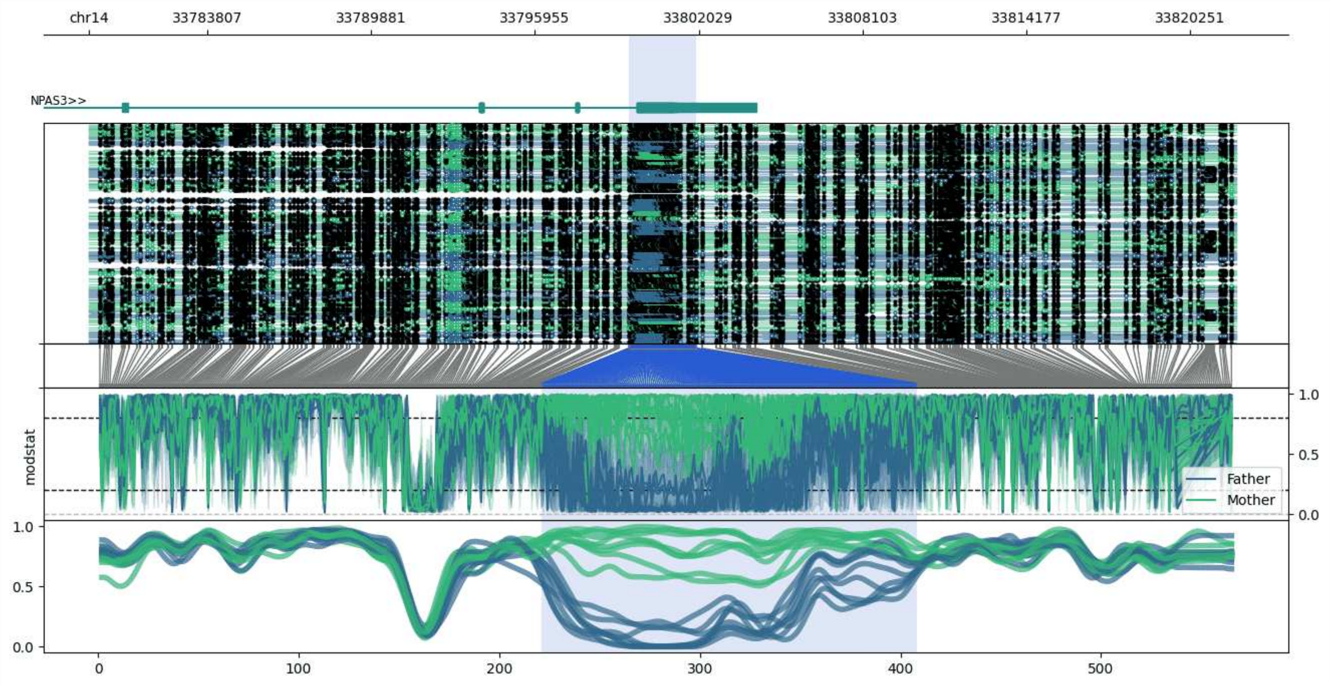


**chr14:33779425-33821943:** Region with 1 DMR containing genes: NPAS3.

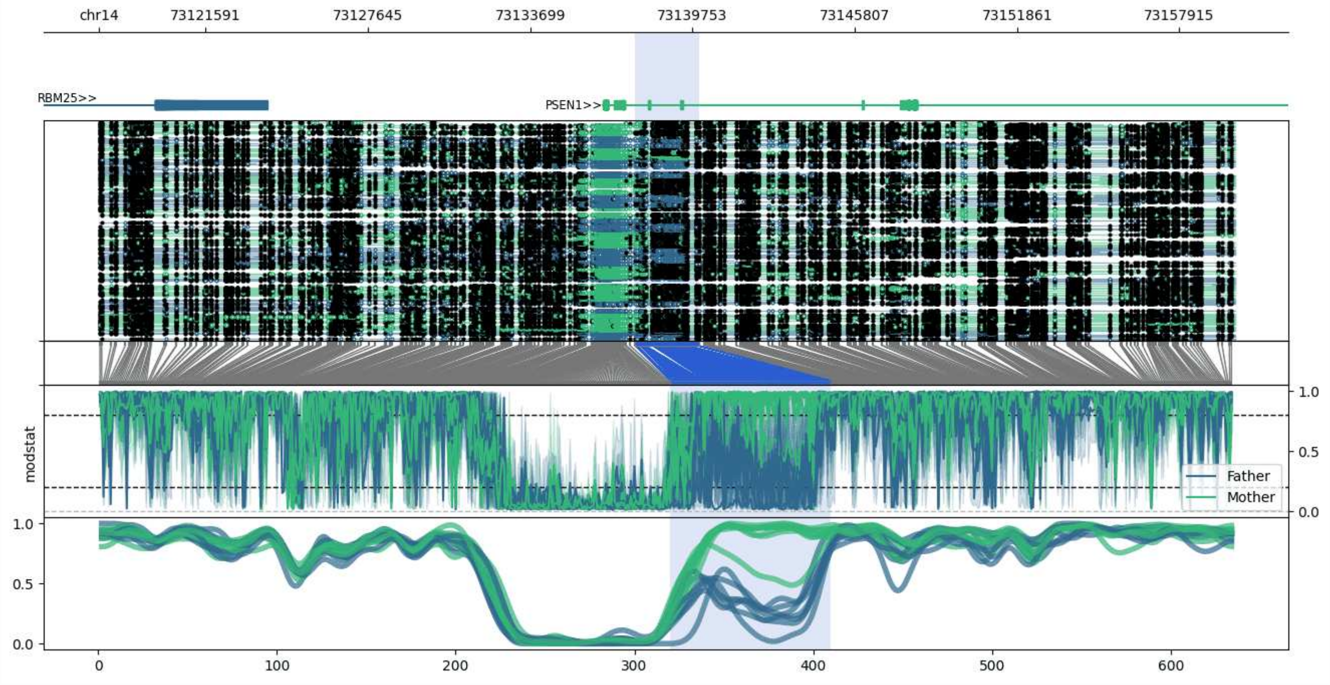


**chr14:73117597-73159980:** Region with 1 DMR containing genes: PSEN1, RBM25.

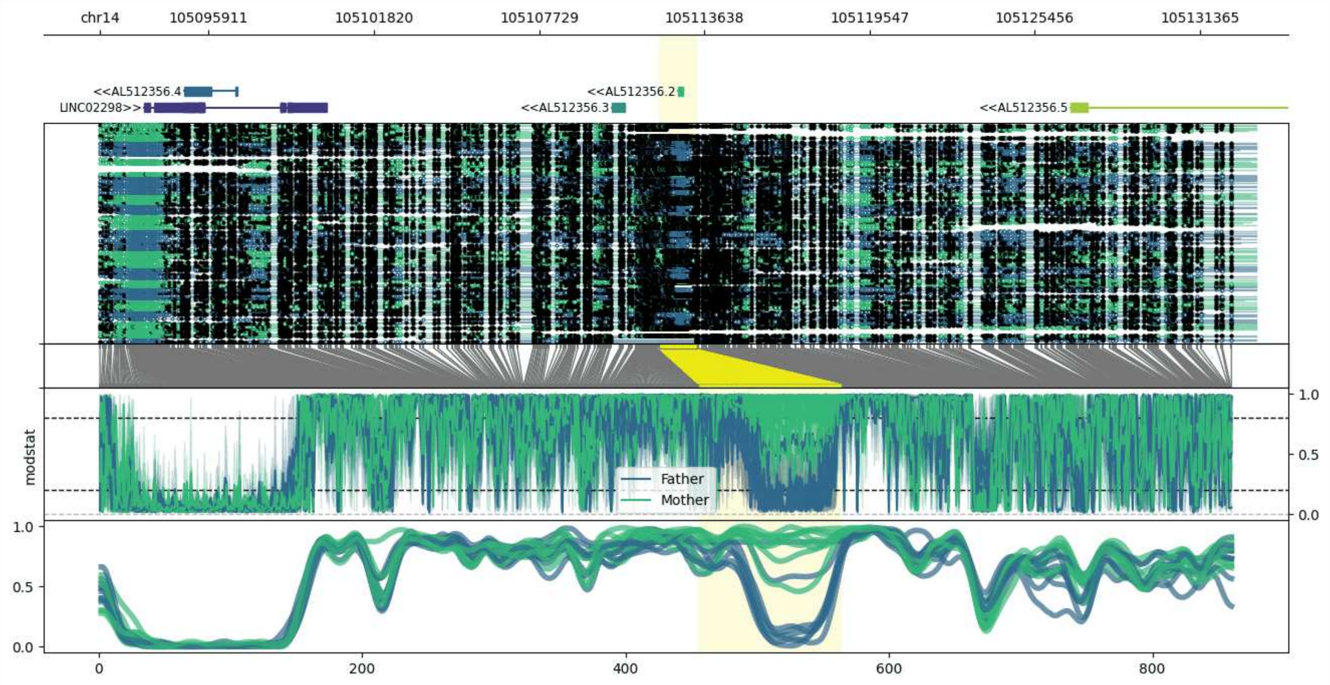


**chr14:105092010-105133376:** Region with 1 DMR containing genes: AL512356.2, AL512356.3, AL512356.4, AL512356.5, JAG2, LINC02298.

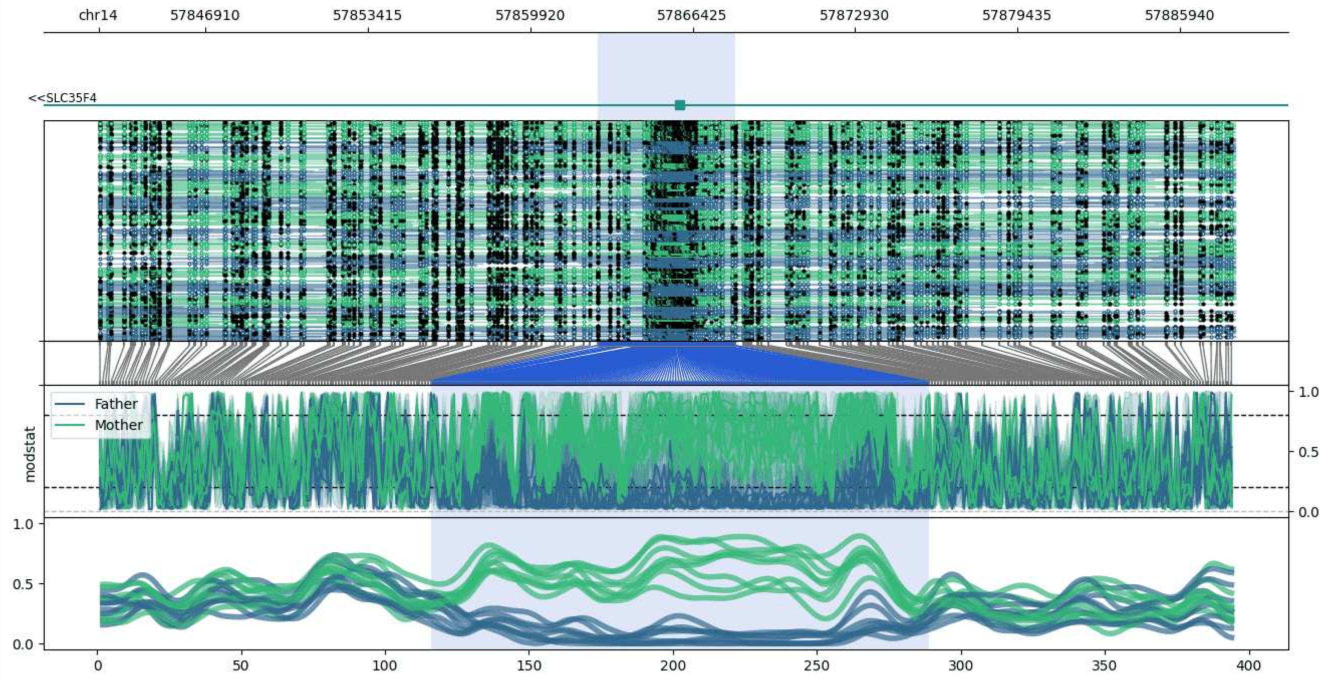


**chr14:57842612-57888149:** Region with 1 DMR containing genes: RN7SKP99, SLC35F4.

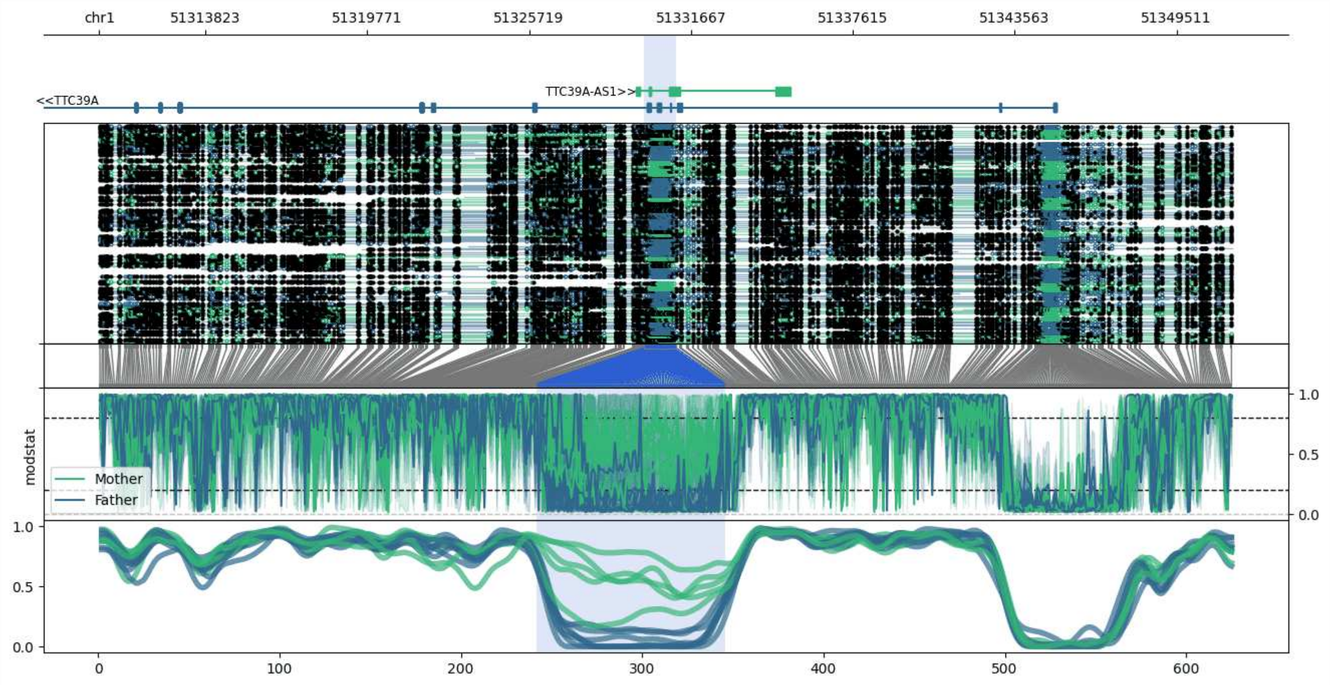


**chr1:51309926-51351565:** Region with 1 DMR containing genes: EPS15, TTC39A, TTC39A-AS1.

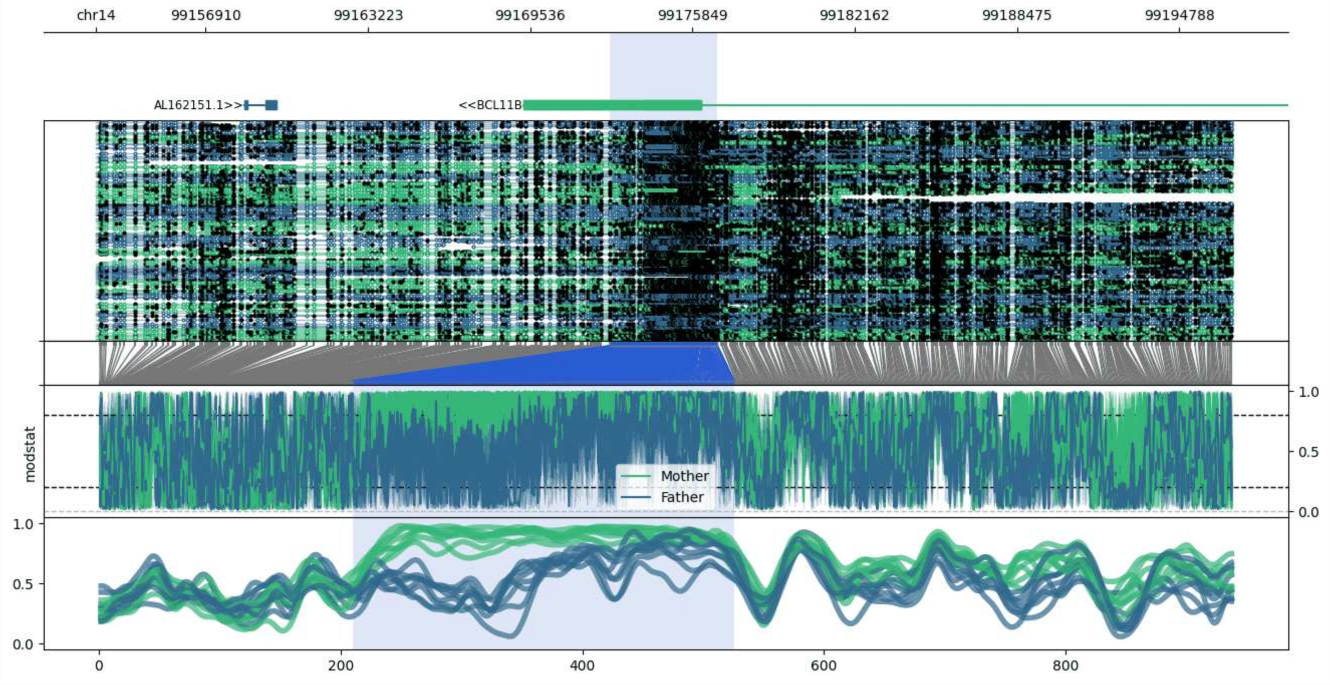


**chr14:99152641-99196832:** Region with 1 DMR containing genes: AL162151.1, BCL11B.

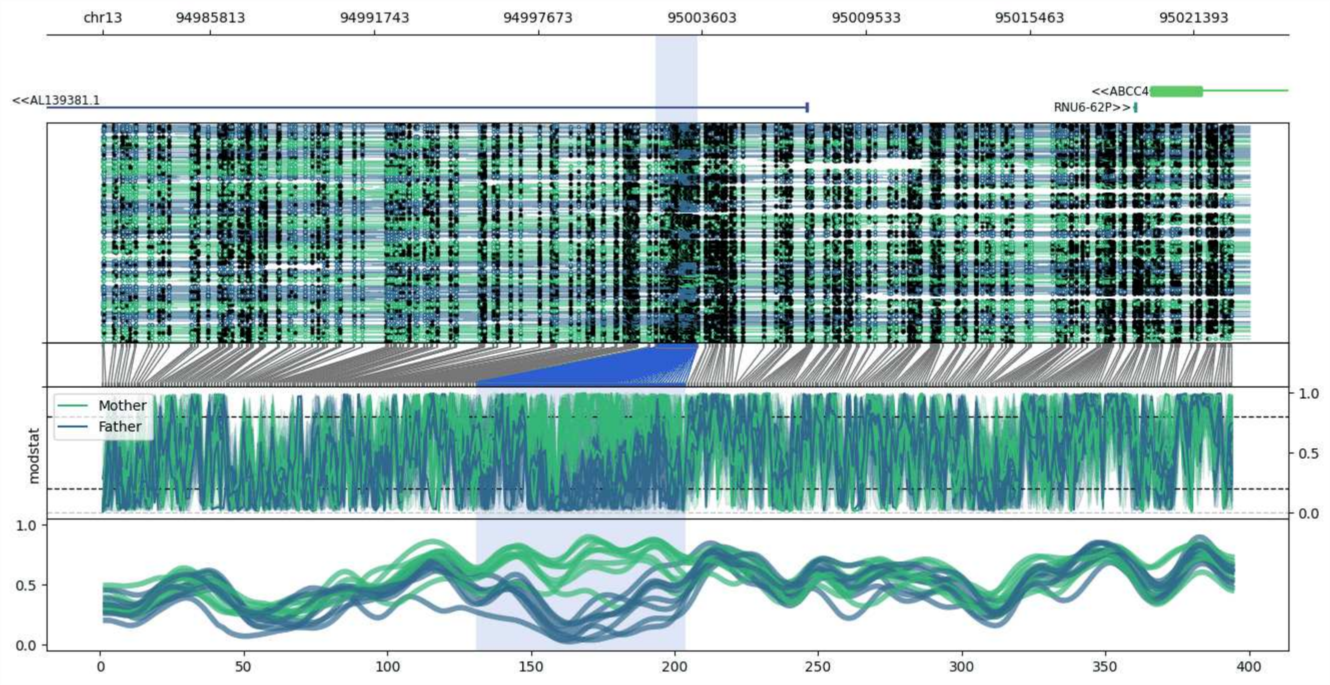


**chr13:94981922-95023434:** Region with 1 DMR containing genes: ABCC4, AL139381.1, RNU6-62P.

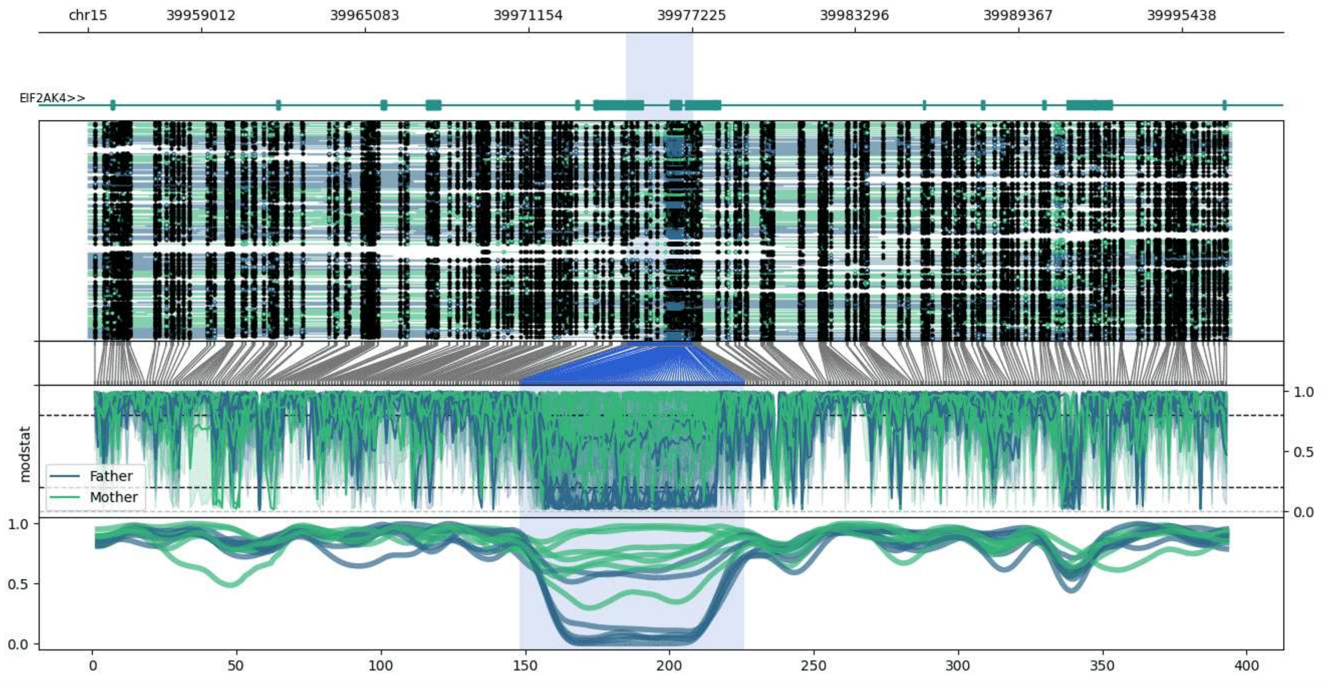


**chr15:39954779-39997280:** Region with 1 DMR containing genes: EIF2AK4, H3F3AP1.

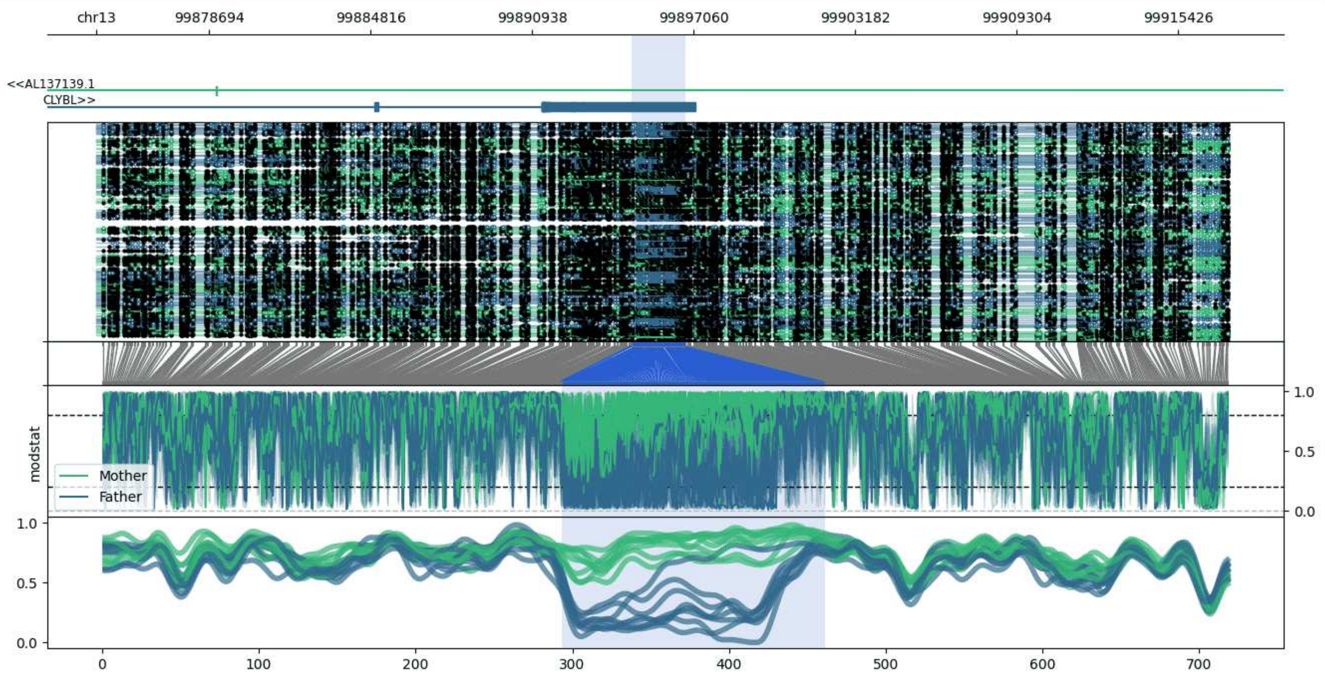


**chr13:99874438-99917295:** Region with 1 DMR containing genes: AL137139.1, CLYBL.

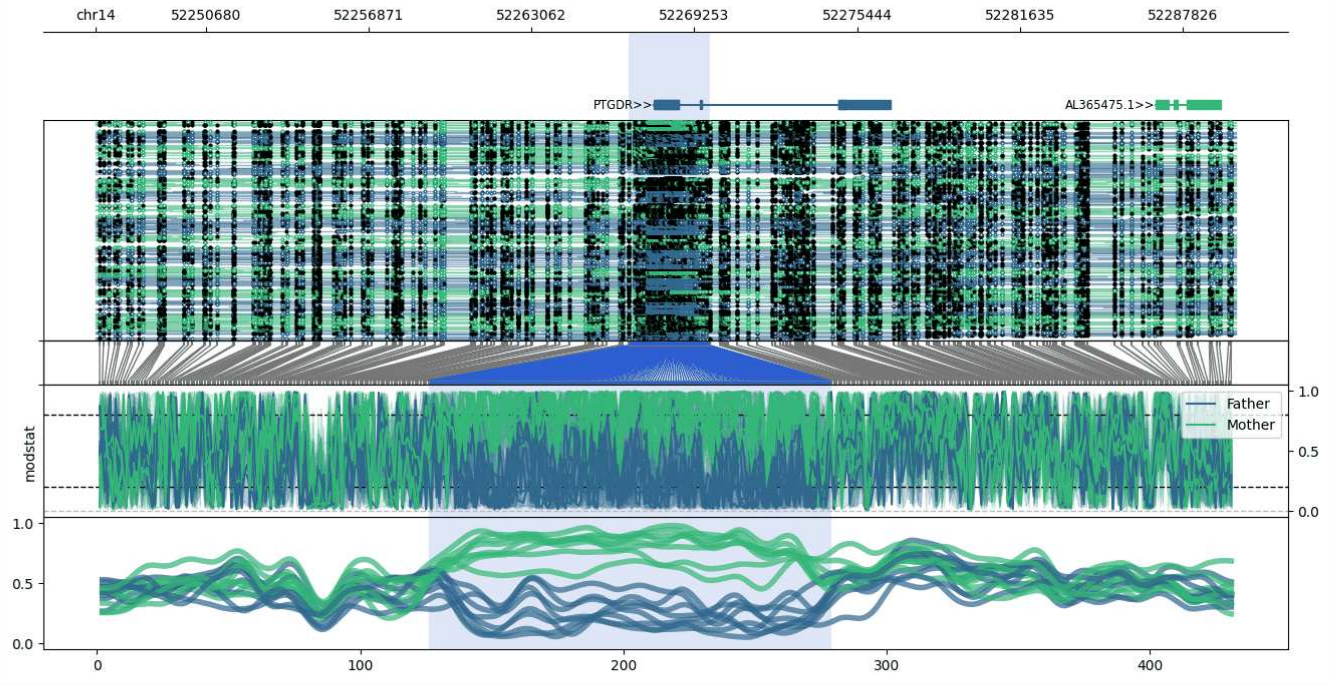


**chr14:52246498-52289839:** Region with 1 DMR containing genes: AL365475.1, PTGDR.

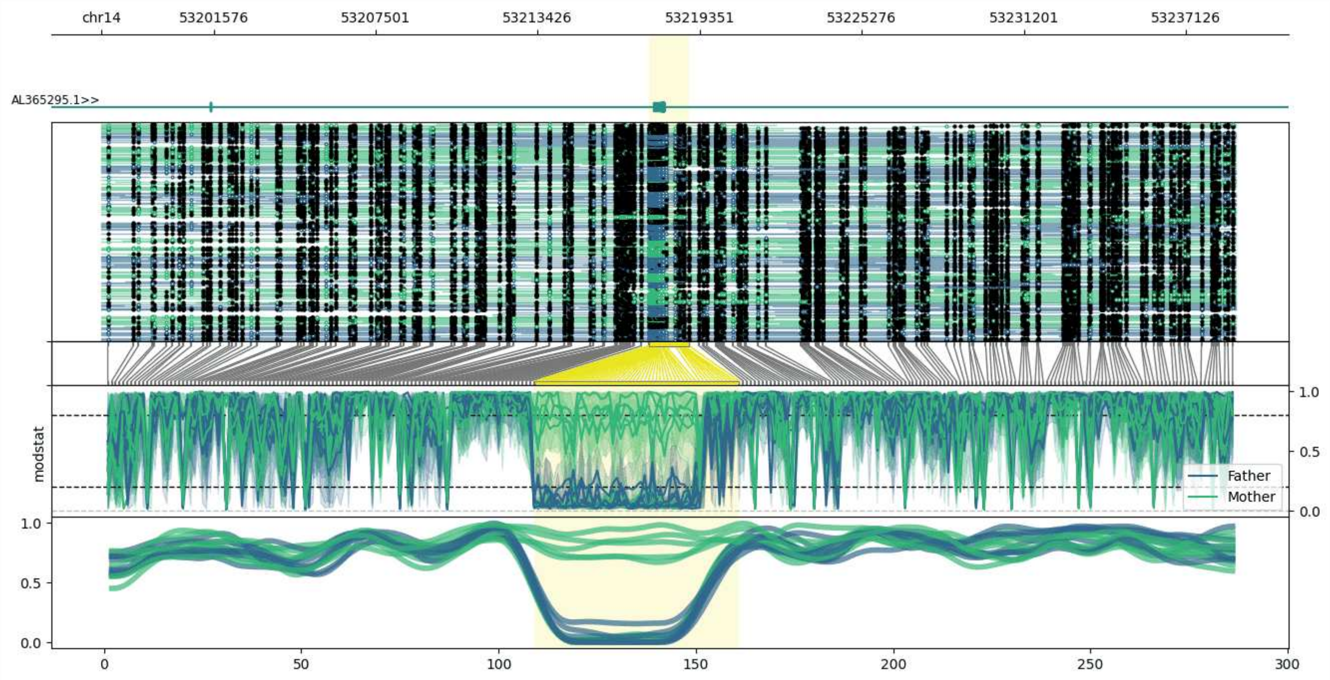


**chr14:53197479-53238958:** Region with 1 DMR containing genes: AL365295.1.

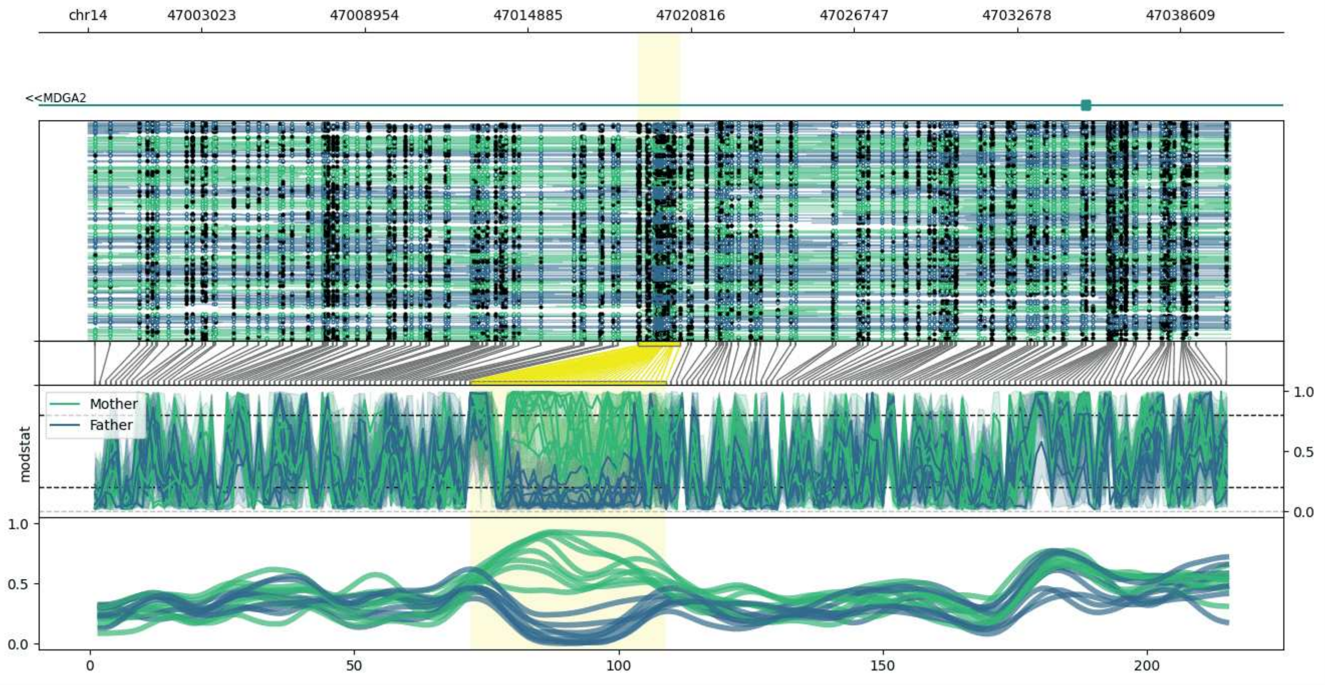


**chr14:46998906-47040425:** Region with 1 DMR containing genes: MDGA2, RPA2P1.

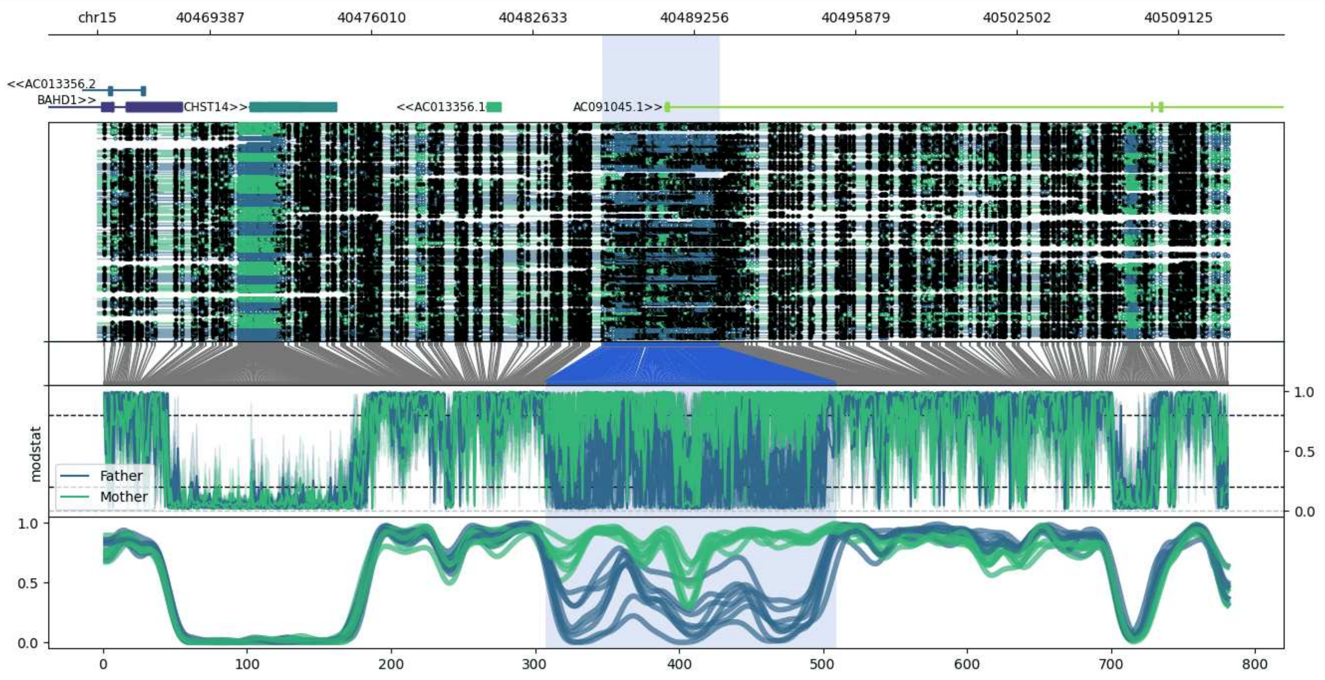


**chr15:40464785-40511152:** Region with 1 DMR containing genes: AC013356.1, AC013356.2, AC091045.1, BAHD1, CHST14.

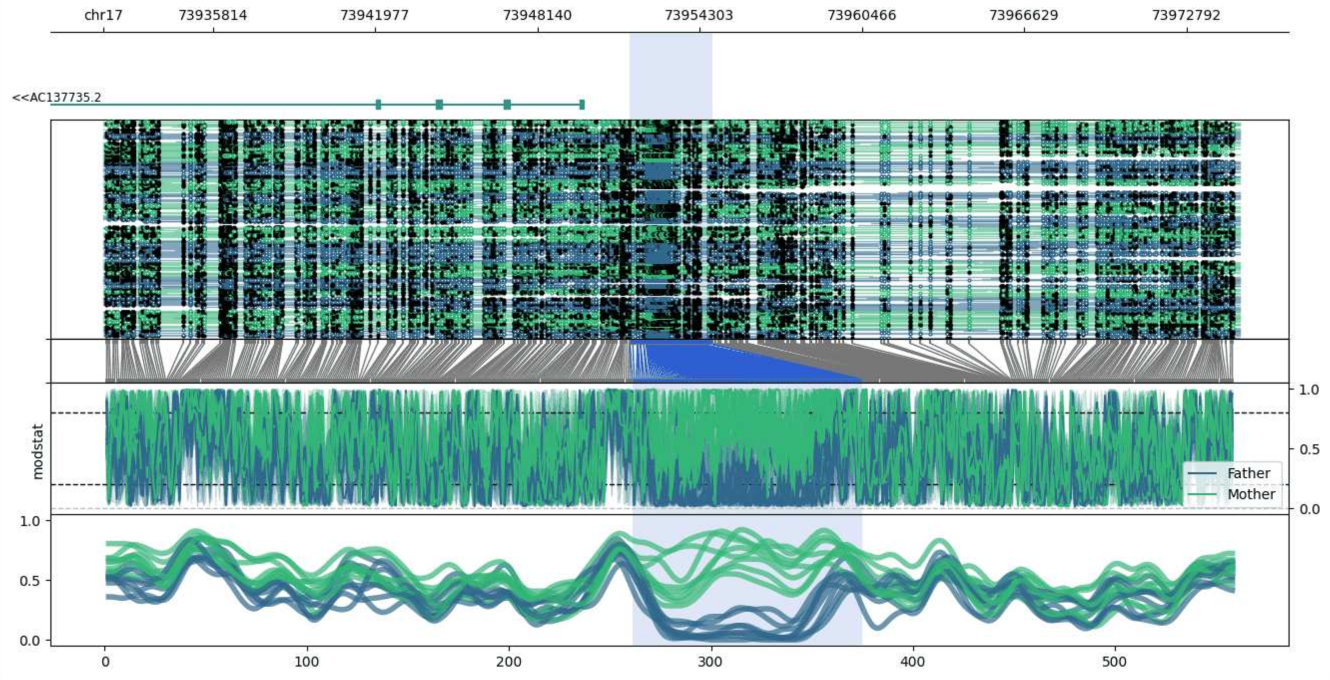


**chr17:73931660-73974801:** Region with 1 DMR containing genes: AC137735.2, AC137735.3.

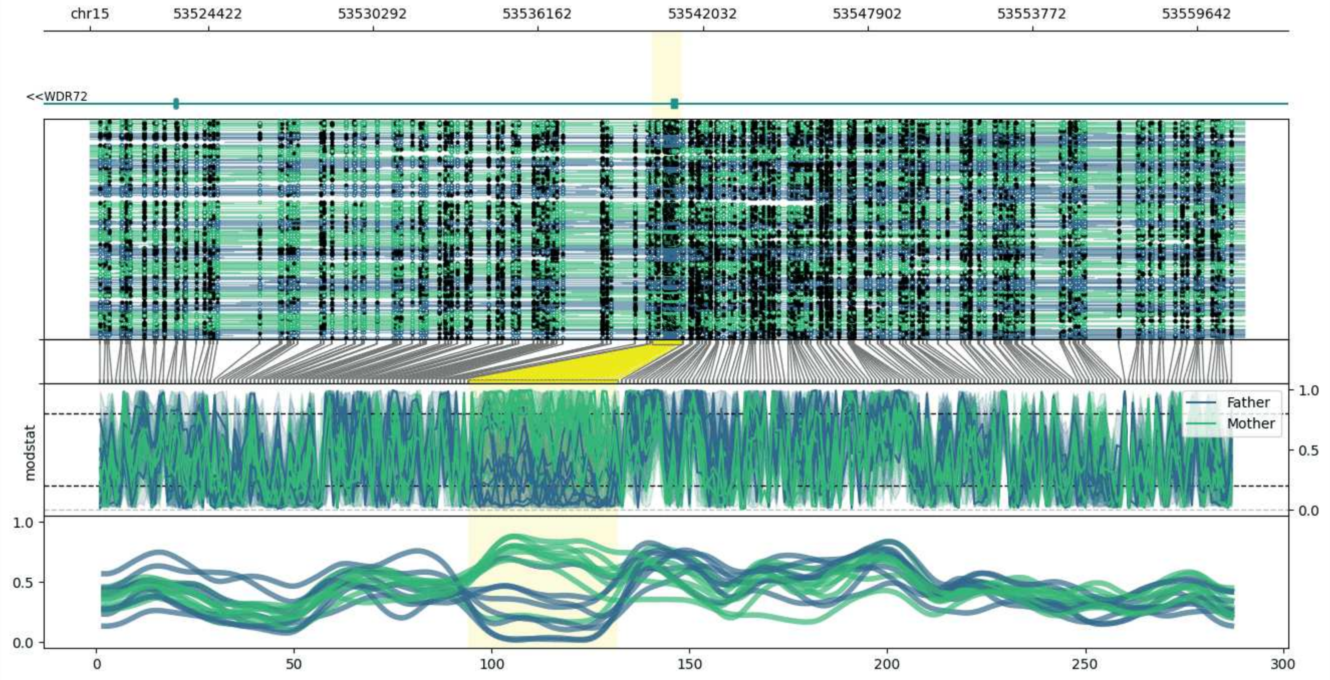


**chr15:53520208-53561300:** Region with 1 DMR containing genes: WDR72.

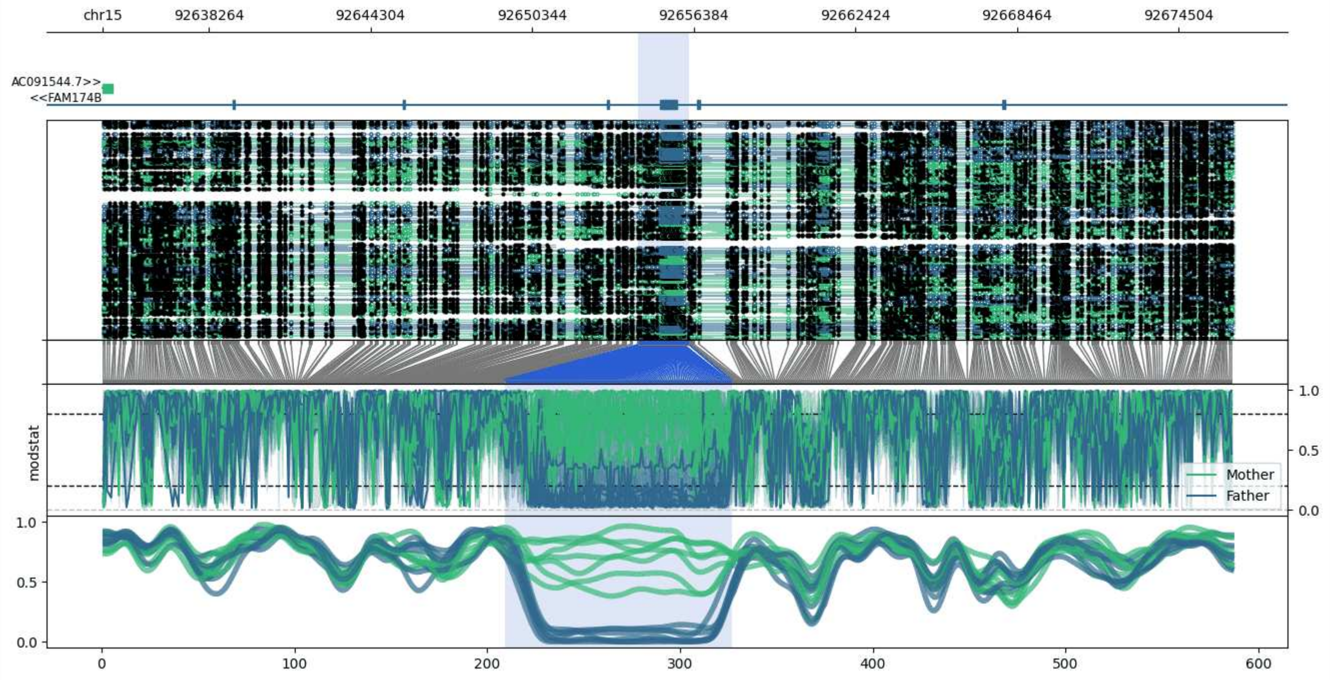


**chr15:92634318-92676599:** Region with 1 DMR containing genes: AC091544.3, AC091544.7, FAM174B.

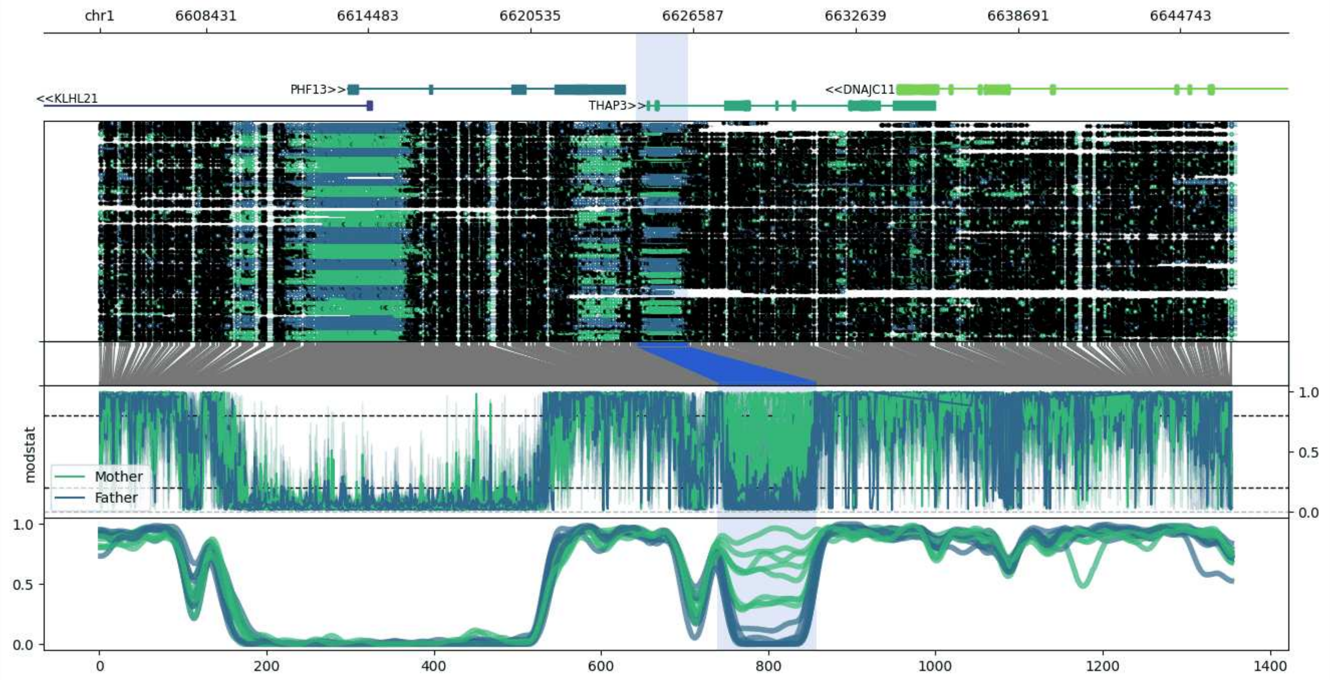


**chr1:6604476-6646840:** Region with 1 DMR containing genes: DNAJC11, KLHL21, PHF13, THAP3.

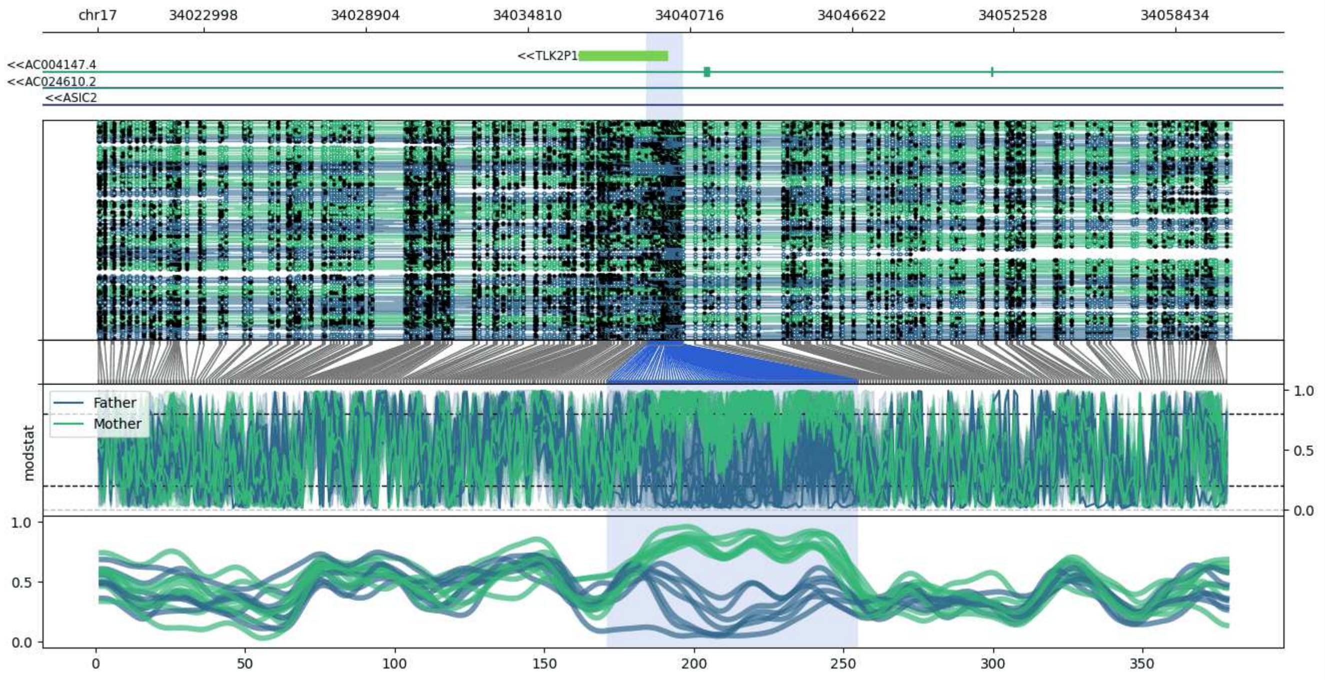


**chr17:34019132-34060476:** Region with 1 DMR containing genes: AC004147.4, AC024610.2, ASIC2, TLK2P1.

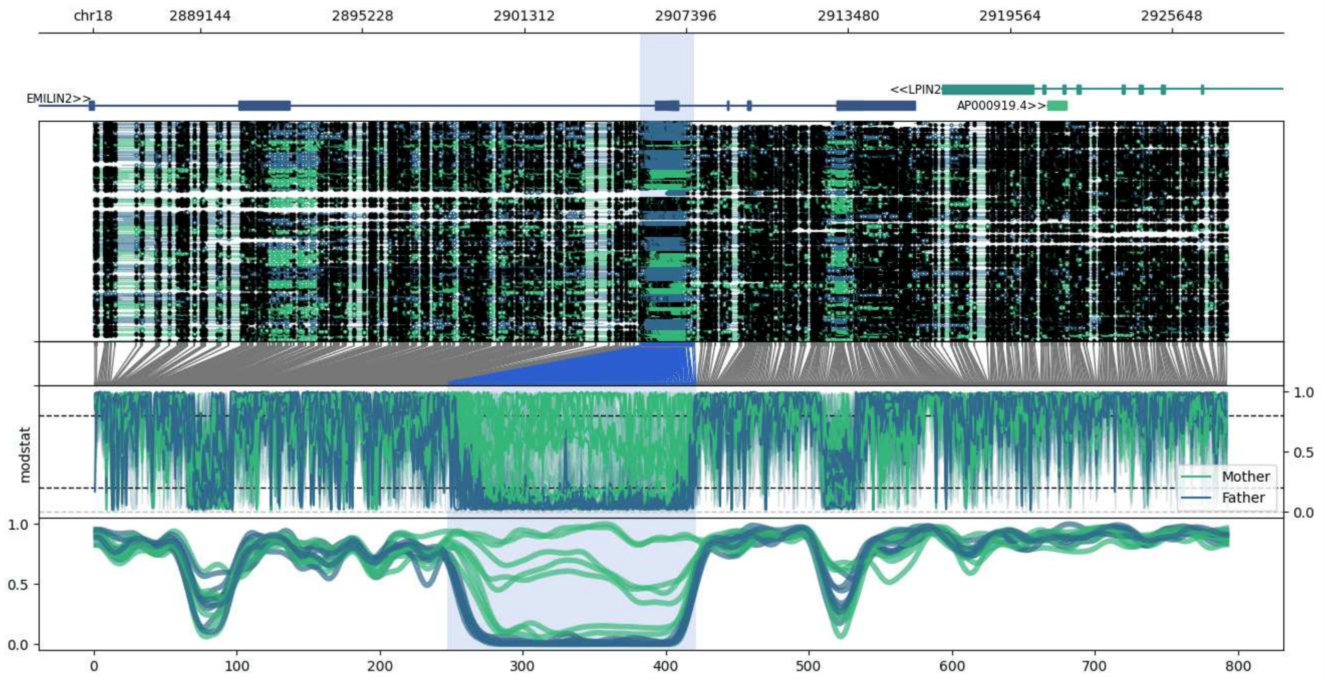


**chr18:2885118-2927711:** Region with 1 DMR containing genes: AP000919.4, EMILIN2, LPIN2.

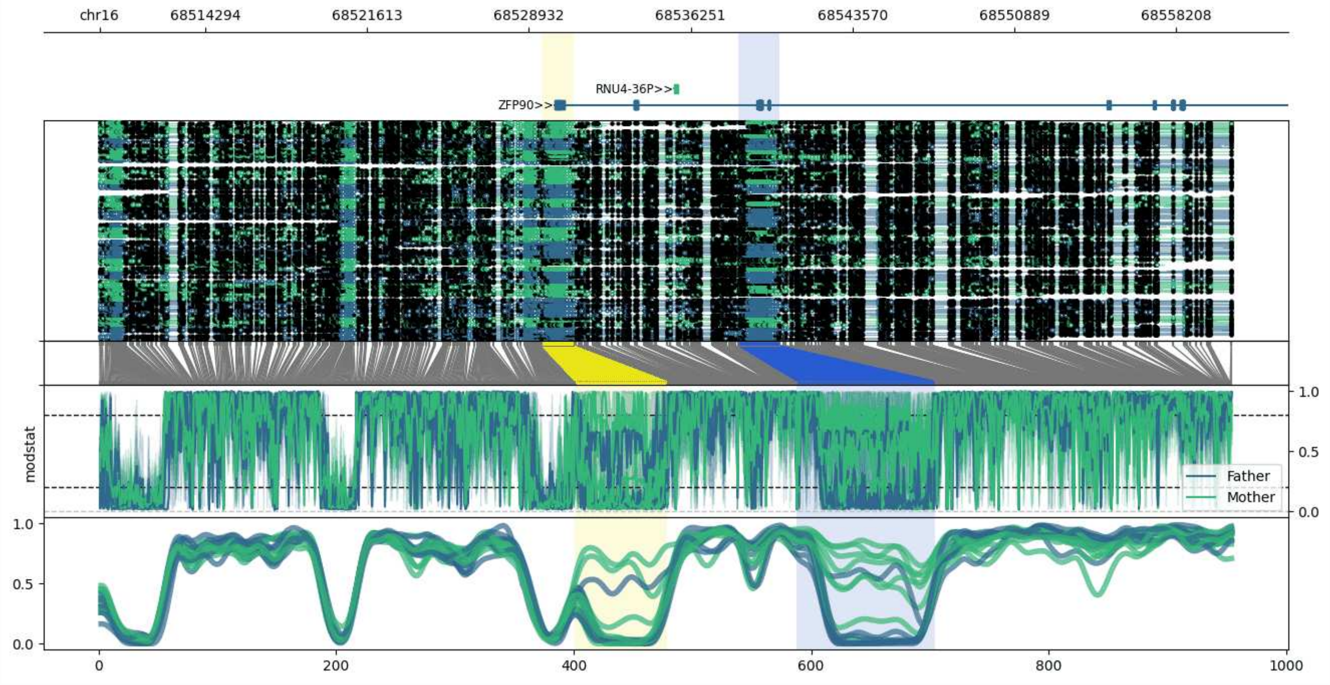


**chr16:68509513-68560751:** Region with 2 DMRs containing genes: RNU4-36P, ZFP90.

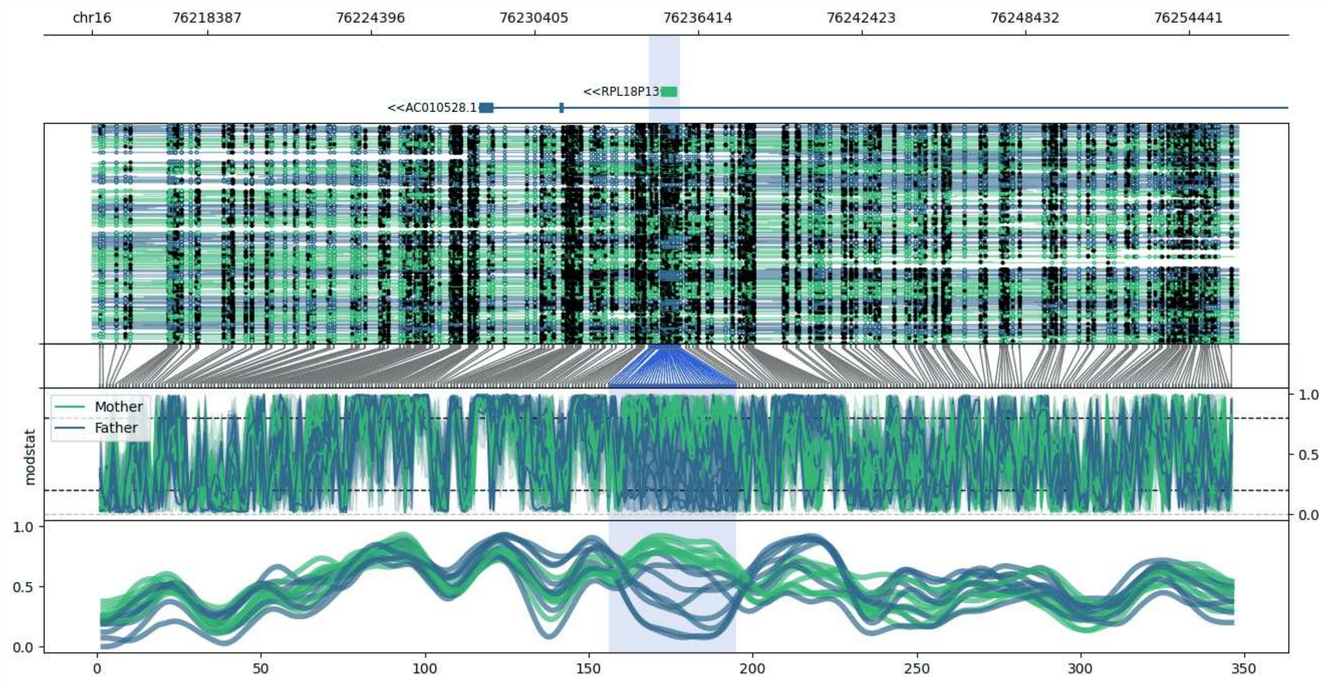


**chr16:76214173-76256239:** Region with 1 DMR containing genes: AC010528.1, AC010528.2, RPL18P13.

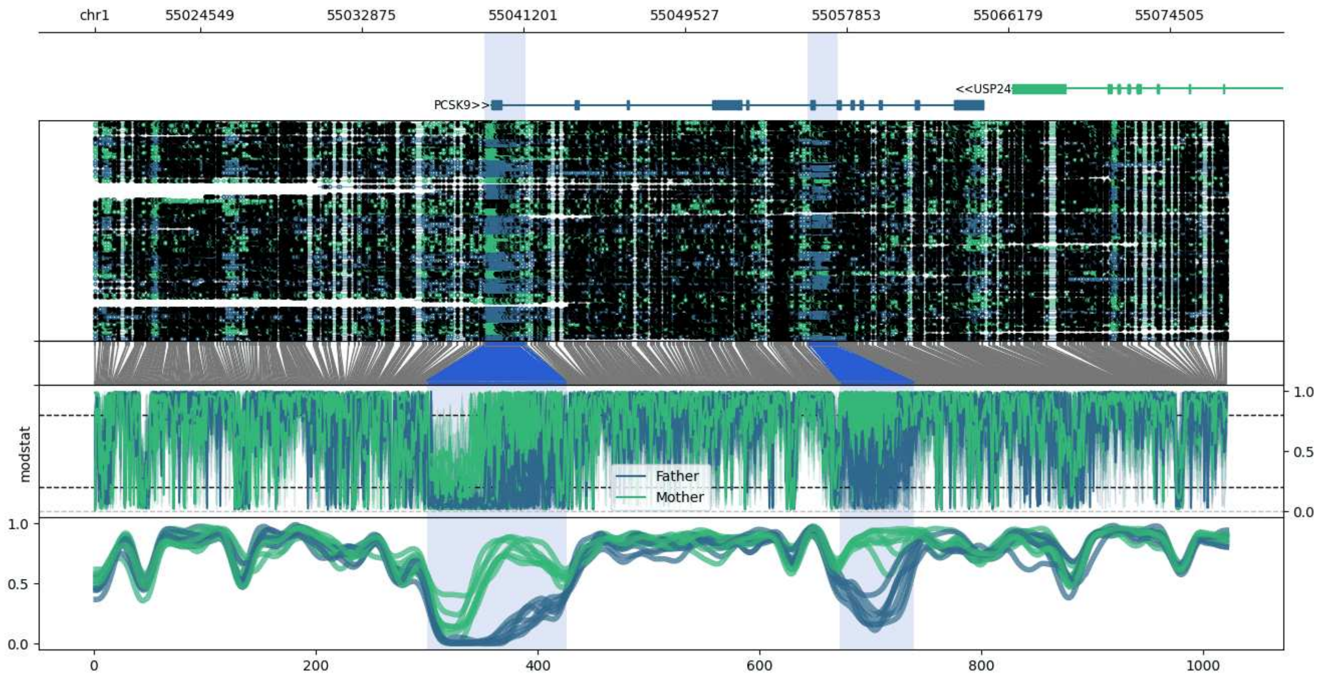


**chr1:55019117-55077403:** Region with 2 DMRs containing genes: BSND, PCSK9, USP24.

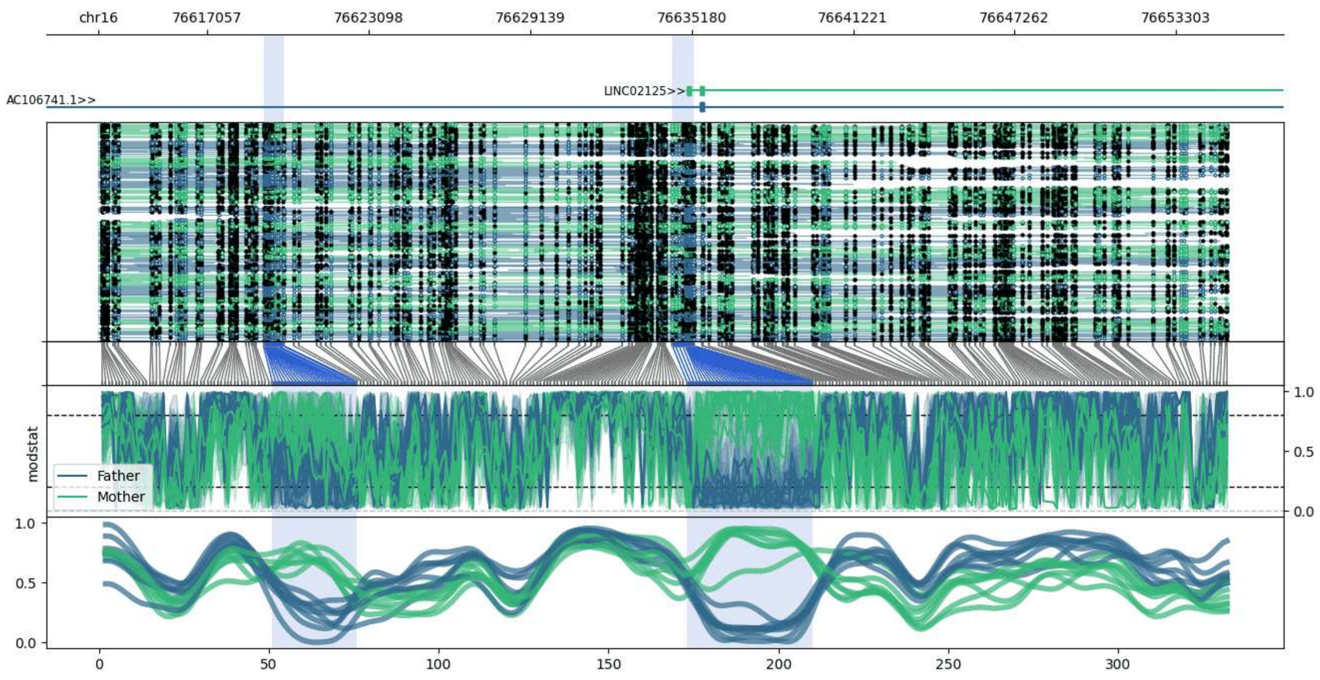


**chr16:76612963-76655253:** Region with 2 DMRs containing genes: AC106741.1, LINC02125.

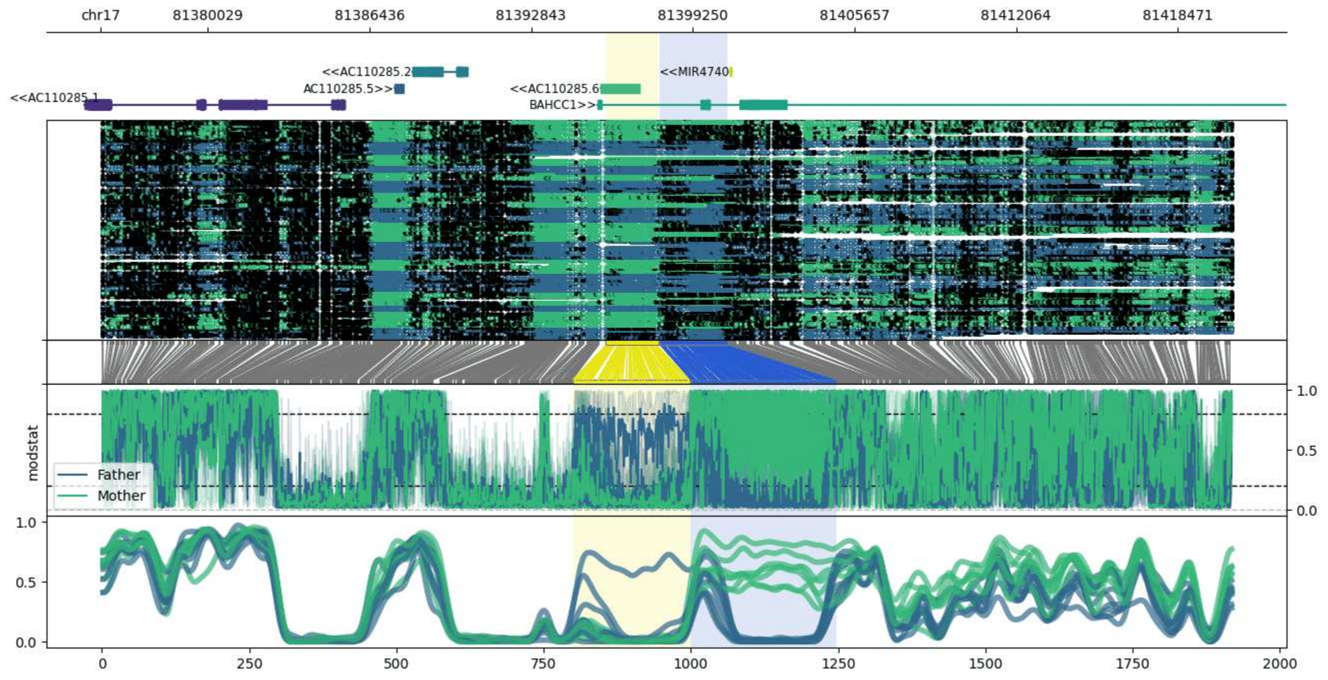


**chr17:81375790-81420643:** Region with 2 DMRs containing genes: AC110285.1, AC110285.2, AC110285.3, AC110285.5, AC110285.6, BAHCC1, MIR4740.

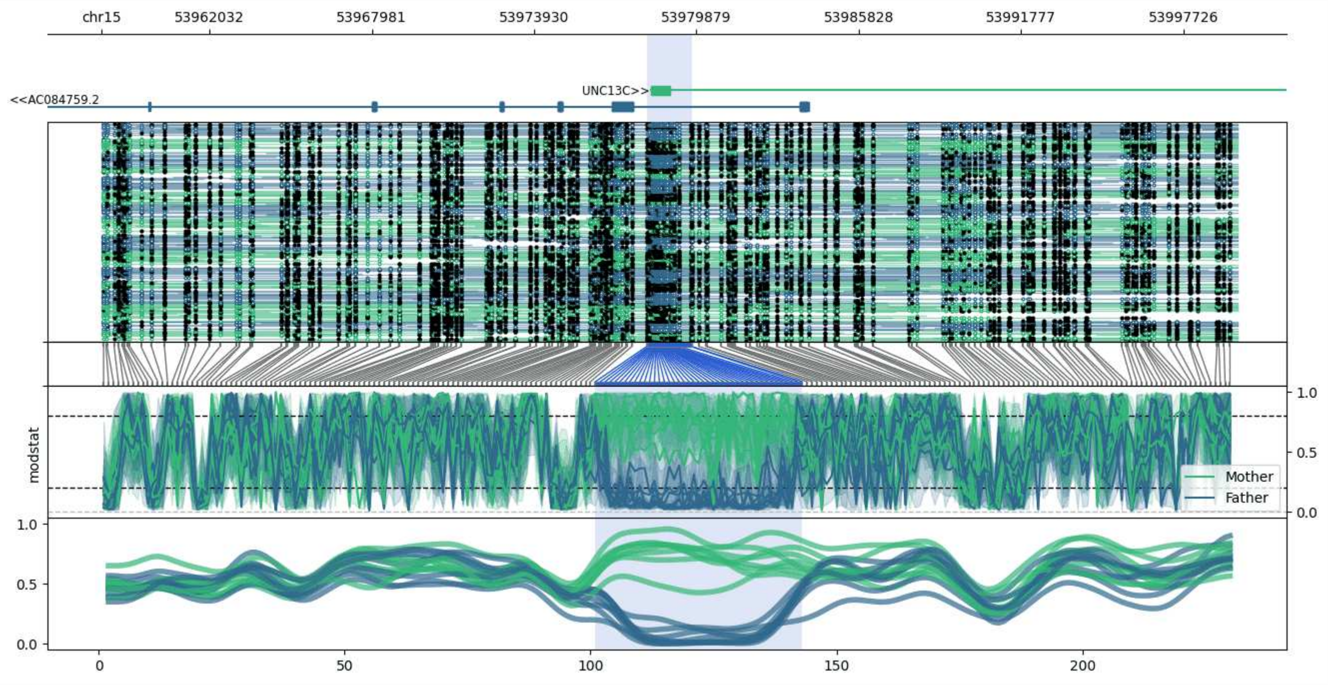


**chr15:53958057-53999701:** Region with 1 DMR containing genes: AC084759.2, UNC13C.

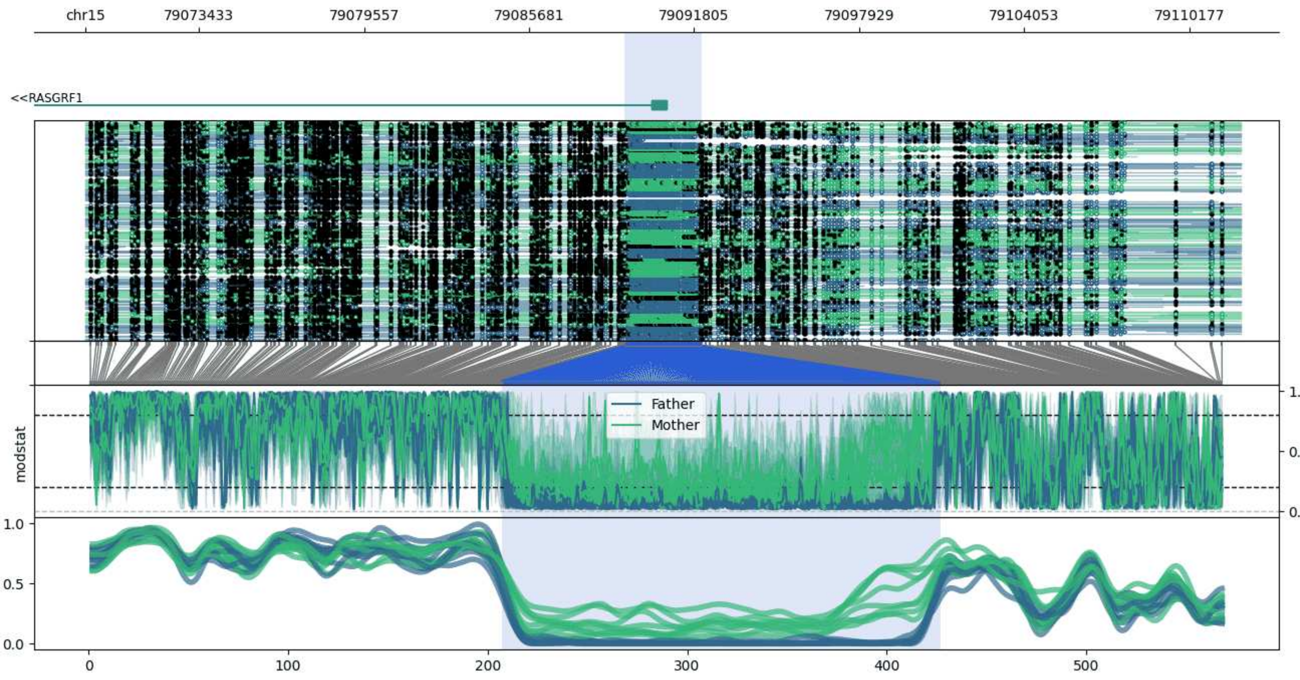


**chr15:79069218-79112088:** Region with 1 DMR containing genes: RASGRF1.

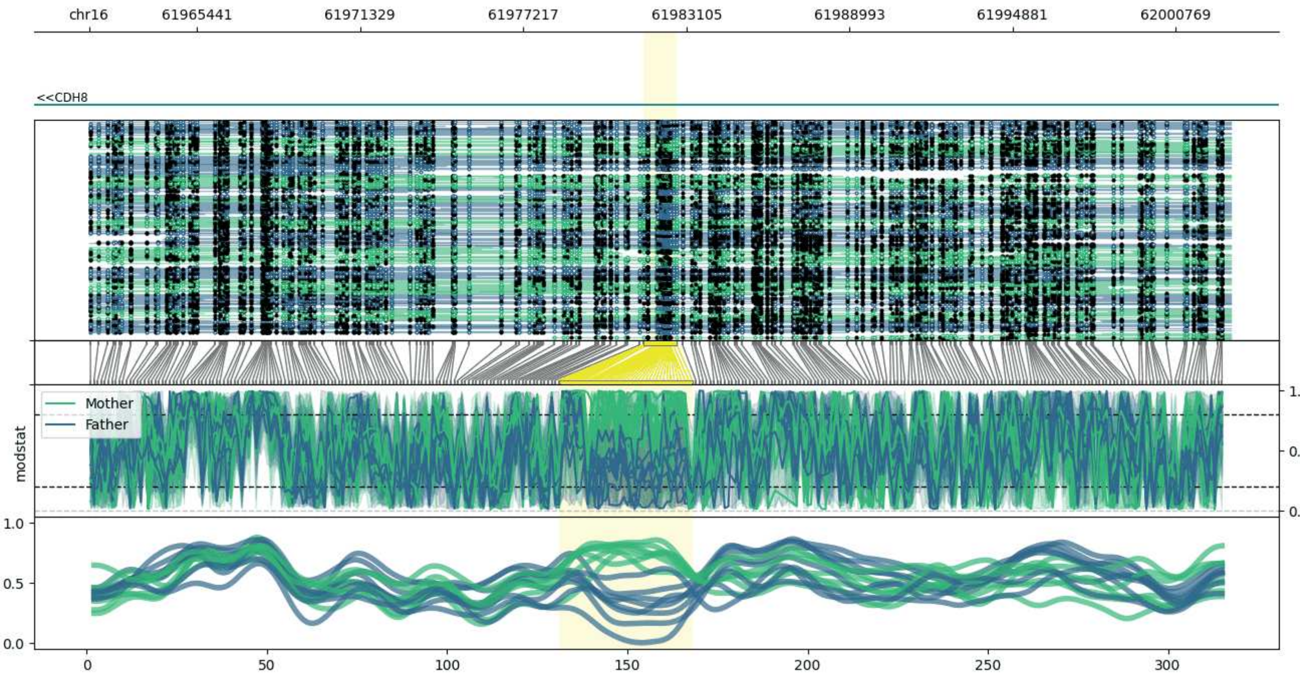


**chr16:61961543-62002760:** Region with 1 DMR containing genes: CDH8.

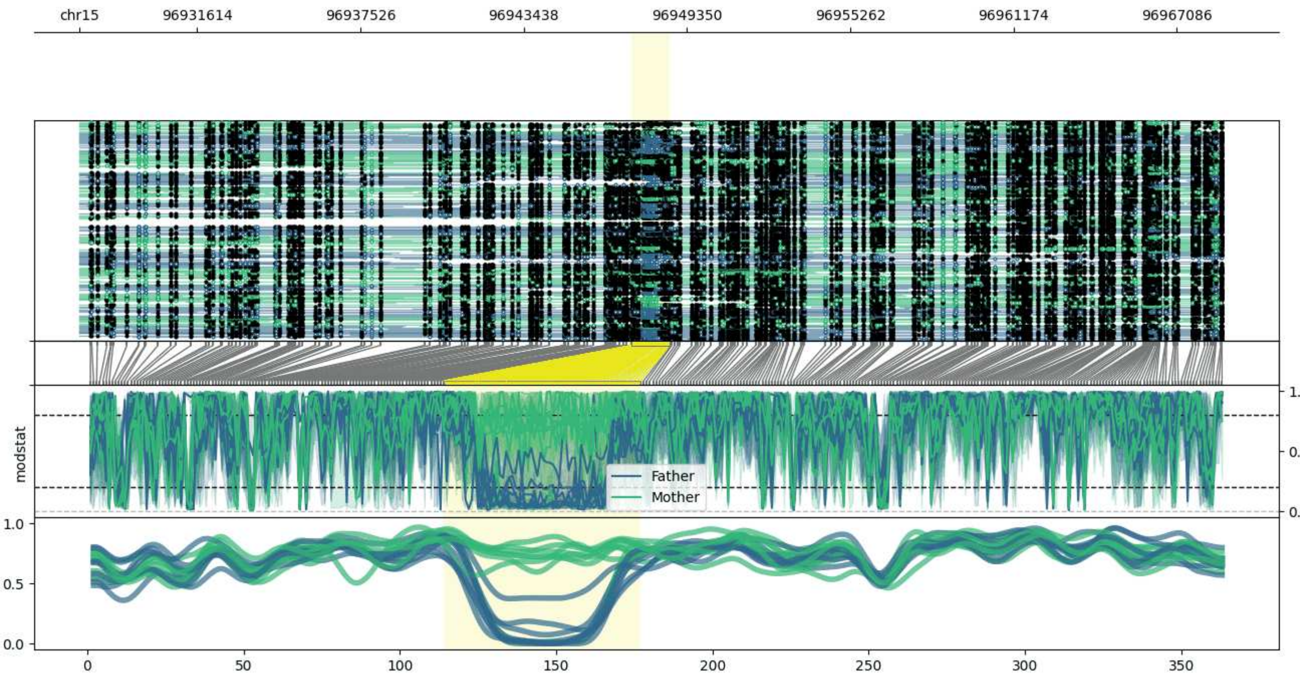


**chr15:96927336-96968721:** Region with 1 DMR containing no genes.

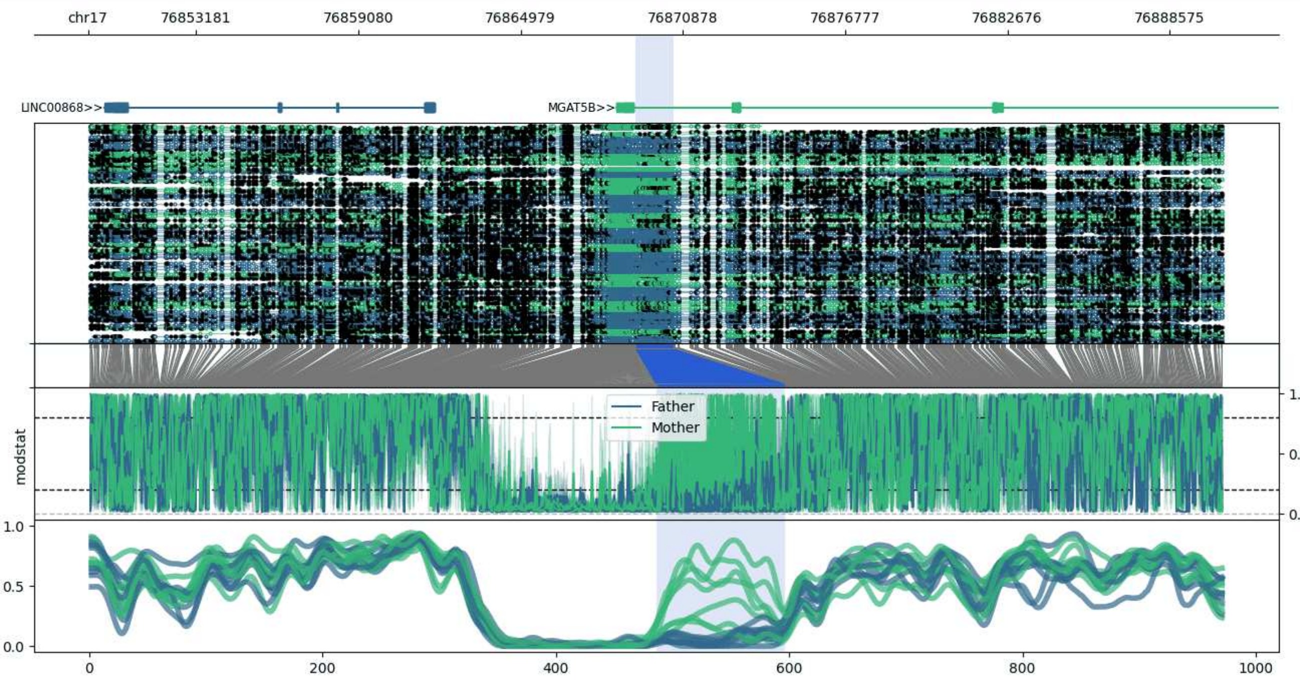


**chr17:76849248-76890544:** Region with 1 DMR containing genes: AC016168.4, LINC00868, MGAT5B.

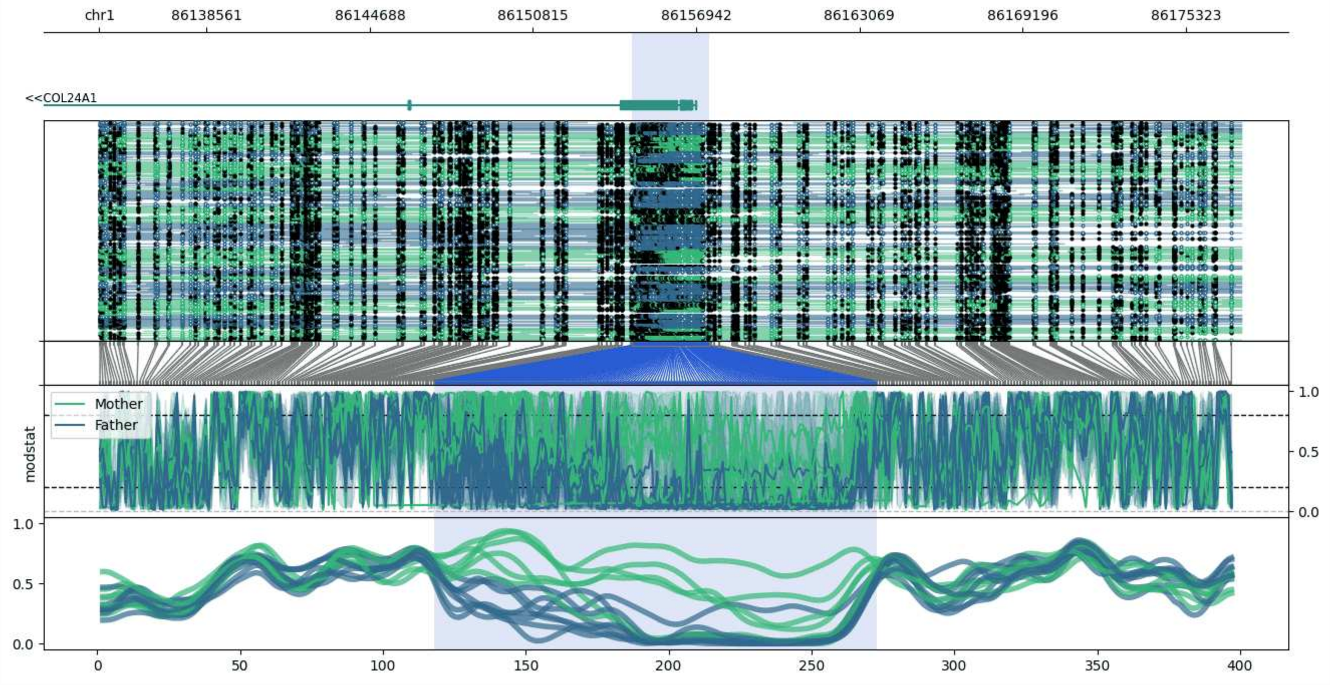


**chr1:86134528-86177417:** Region with 1 DMR containing genes: COL24A1.

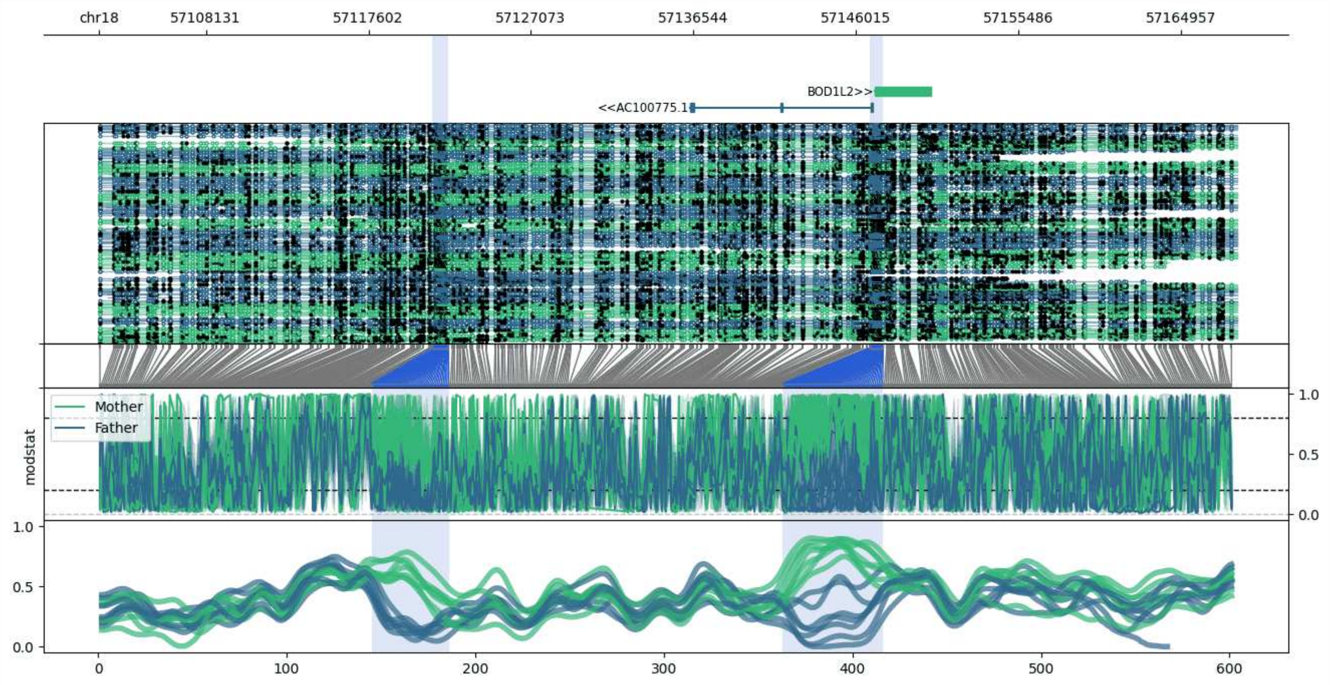


**chr18:57101905-57168202:** Region with 2 DMRs containing genes: AC100775.1, BOD1L2, LINC02565.

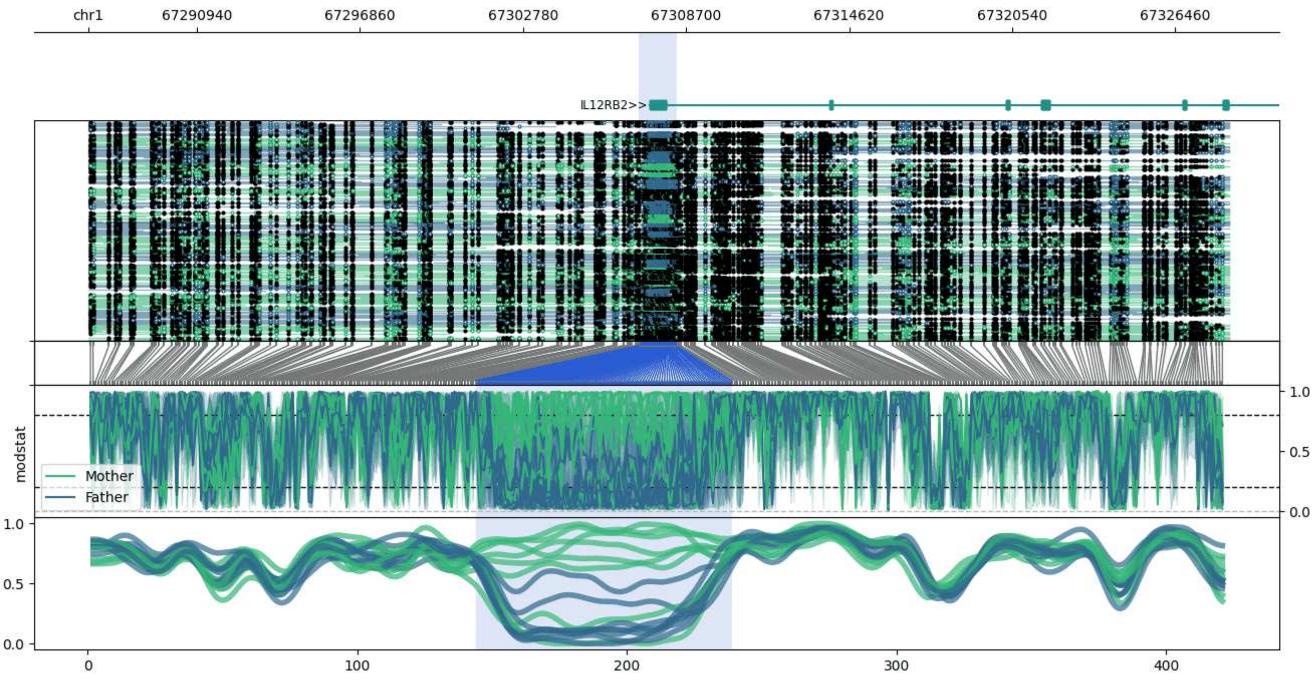


**chr1:67286992-67328433:** Region with 1 DMR containing genes: AL389925.1, IL12RB2.

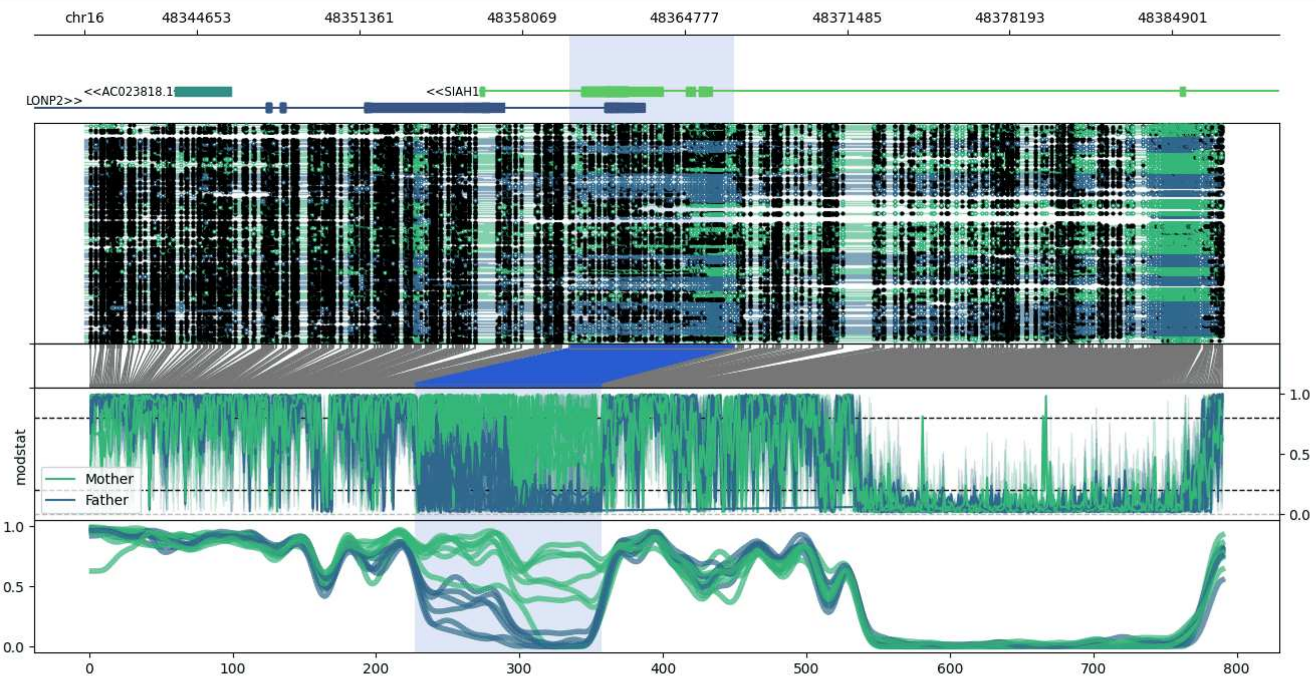


**chr16:48340026-48386983:** Region with 1 DMR containing genes: AC023818.1, LONP2, SIAH1, UBA52P8.

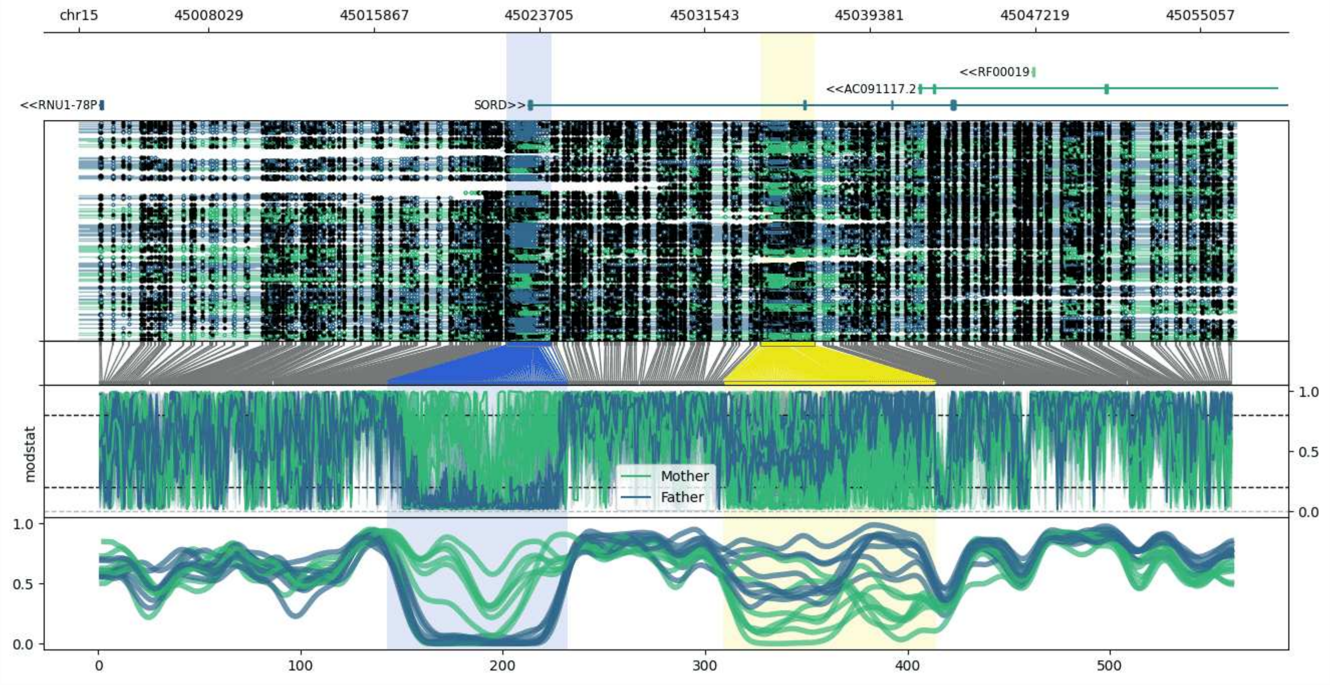


**chr15:45001877-45056743:** Region with 2 DMRs containing genes: AC091117.2, RF00019, RNU1-78P, SORD.

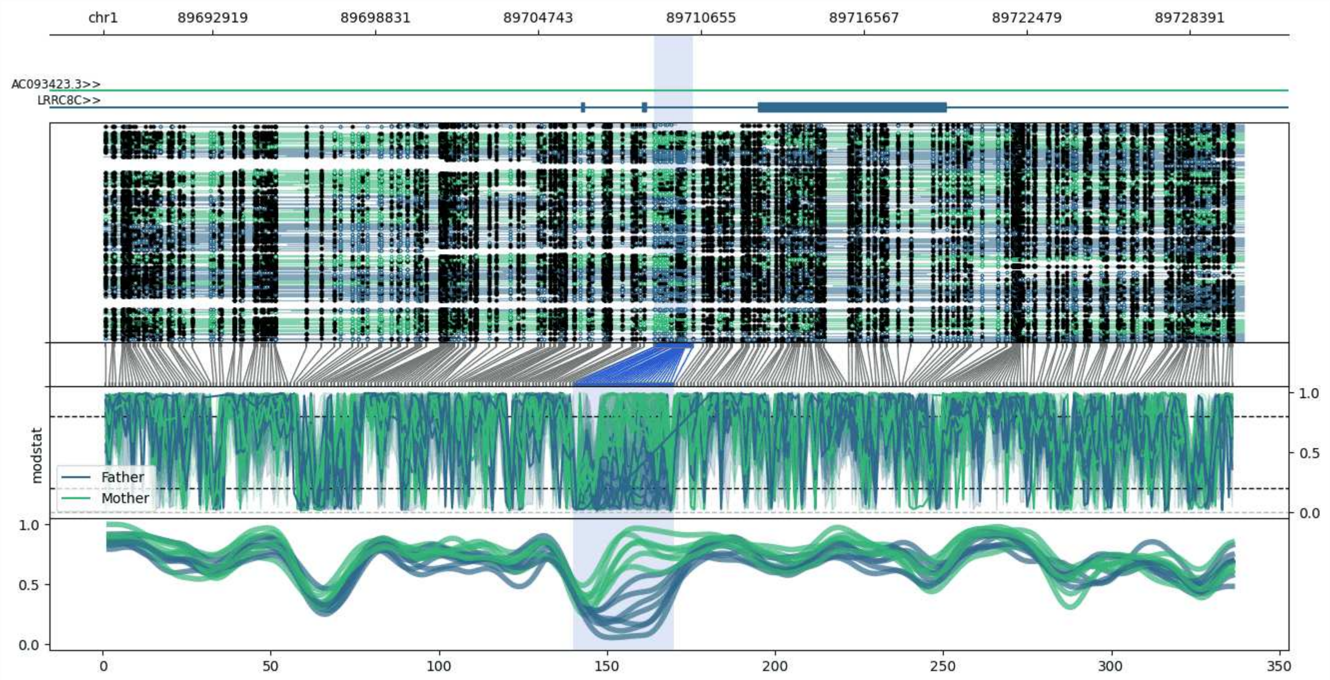


**chr1:89688966-89730352:** Region with 1 DMR containing genes: AC093423.3, LRRC8C.

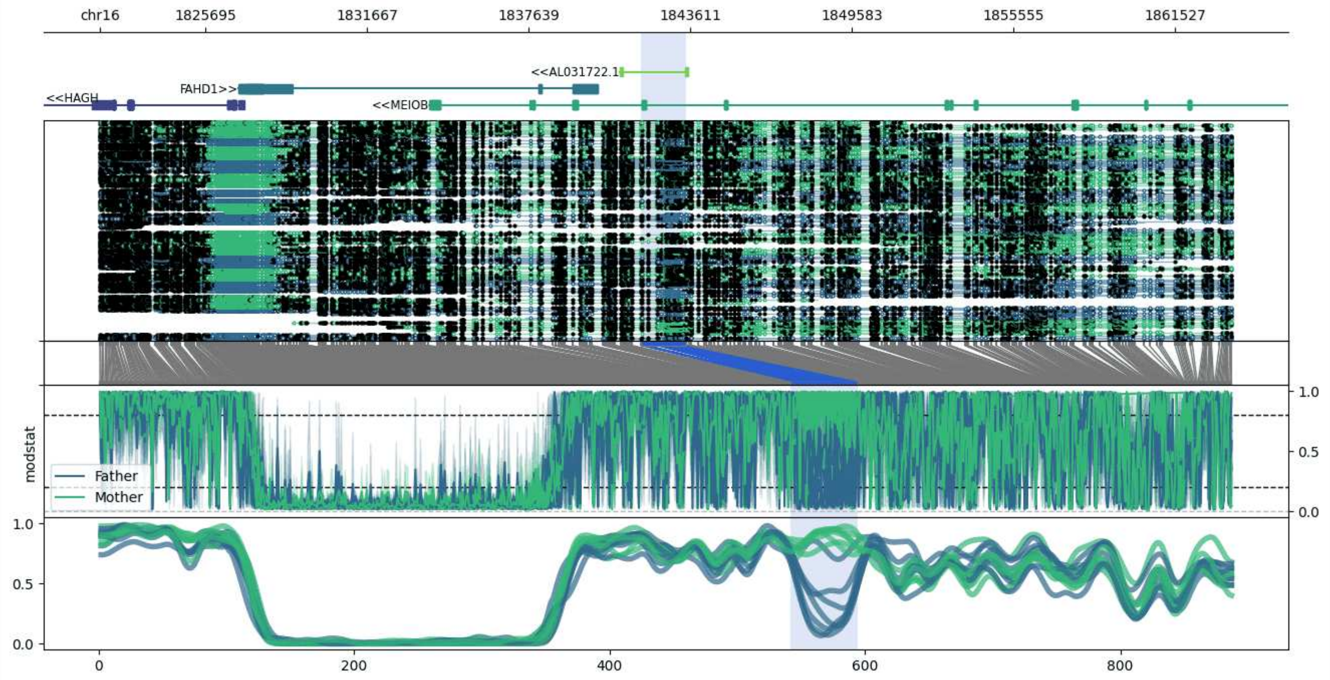


**chr16:1821808-1863612:** Region with 1 DMR containing genes: AL031722.1, FAHD1, HAGH, MEIOB.

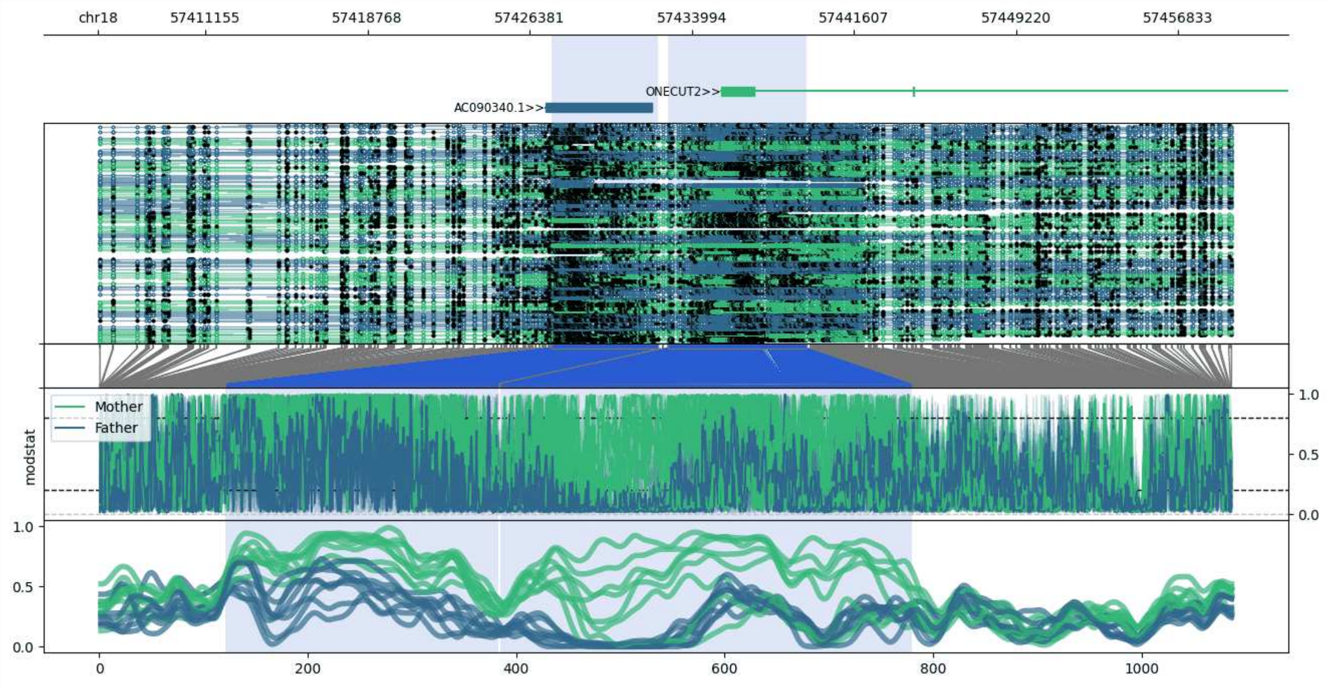


**chr18:57406097-57459393:** Region with 2 DMRs containing genes: AC090340.1, ONECUT2.

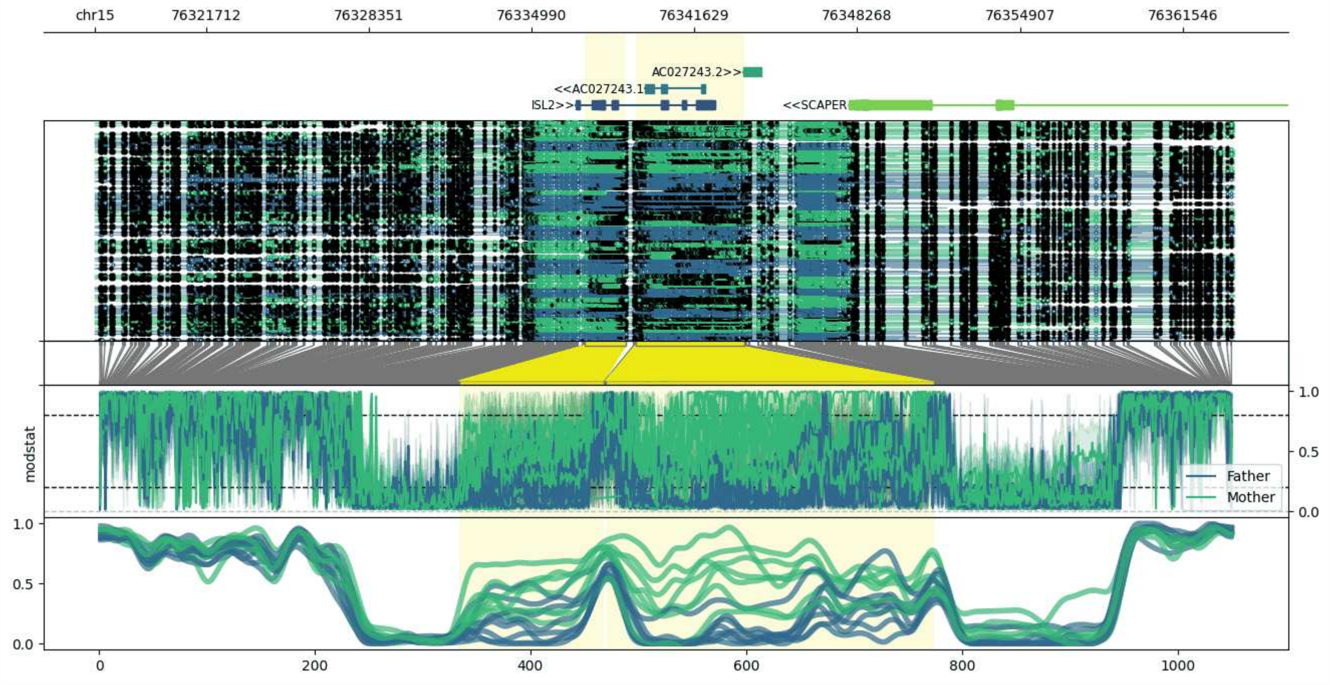


**chr15:76317168-76363646:** Region with 2 DMRs containing genes: AC027243.1, AC027243.2, ETFA, ISL2, SCAPER.

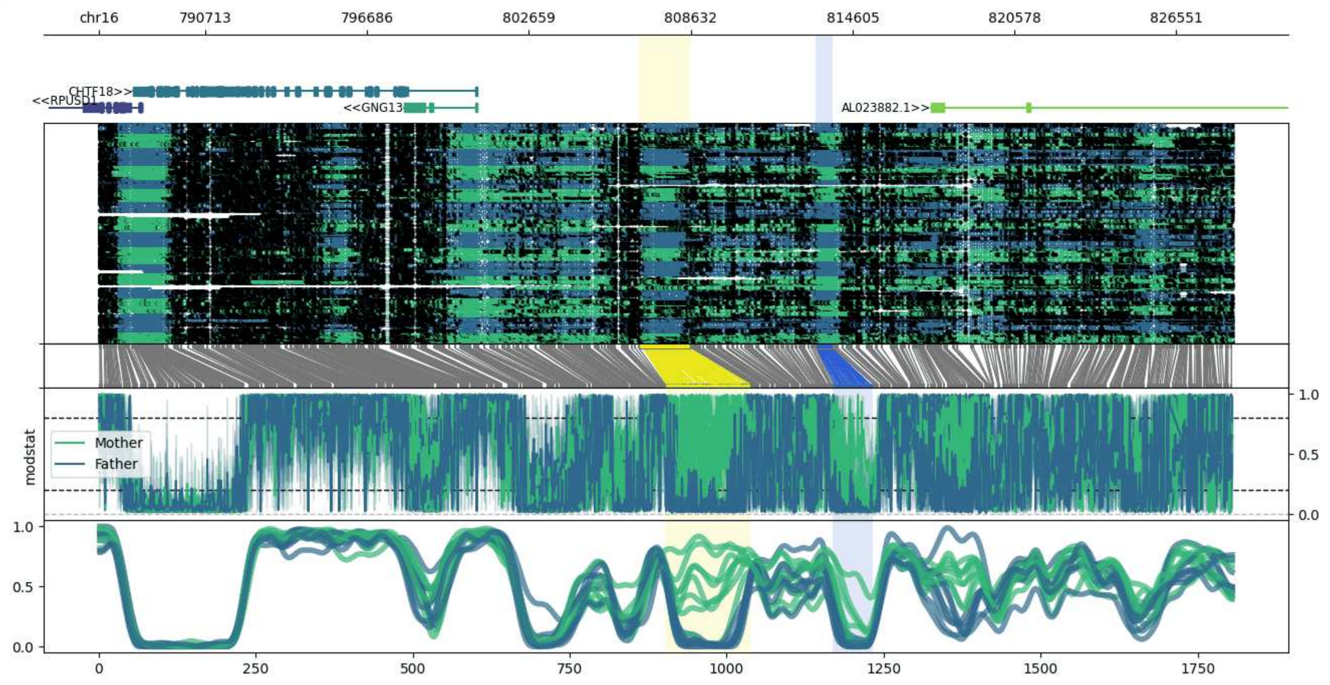


**chr16:786789-828600:** Region with 2 DMRs containing genes: AL023882.1, CHTF18, GNG13, MSLNL, RPUSD1.

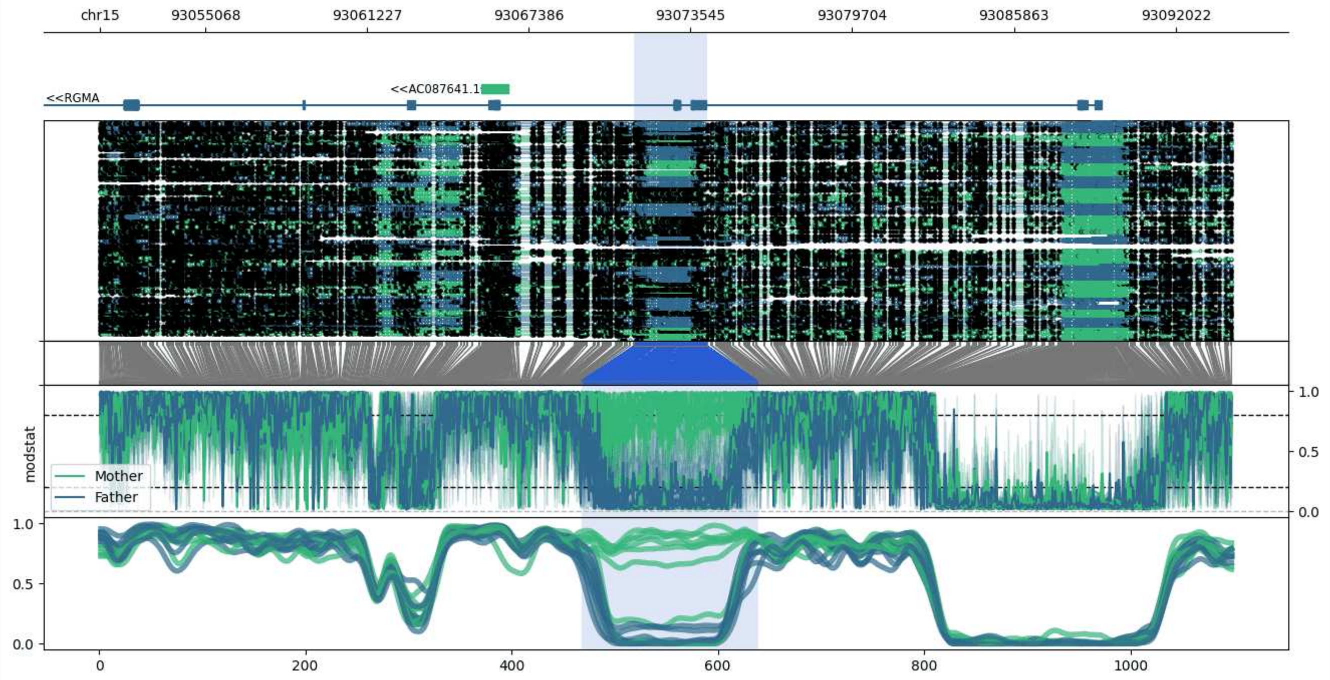


**chr15:93051059-93094172:** Region with 1 DMR containing genes: AC087641.1, RGMA.

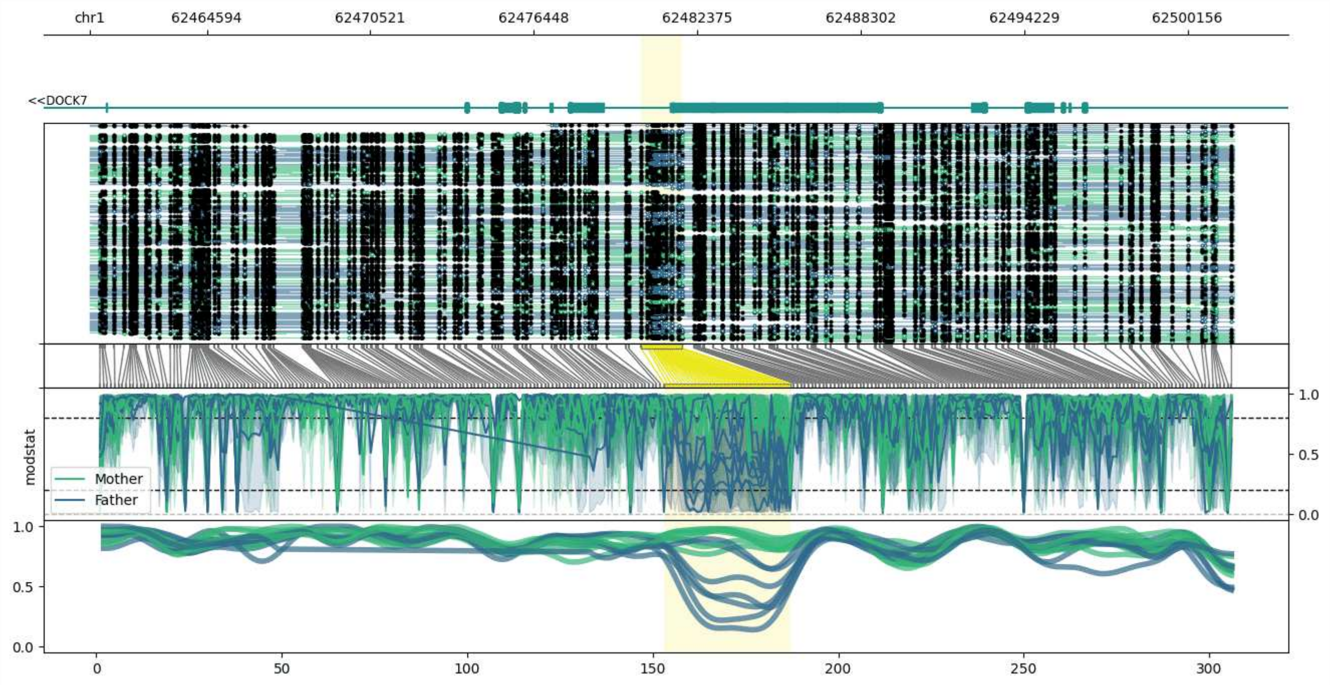


**chr1:62460336-62501827:** Region with 1 DMR containing genes: DOCK7, USP1.

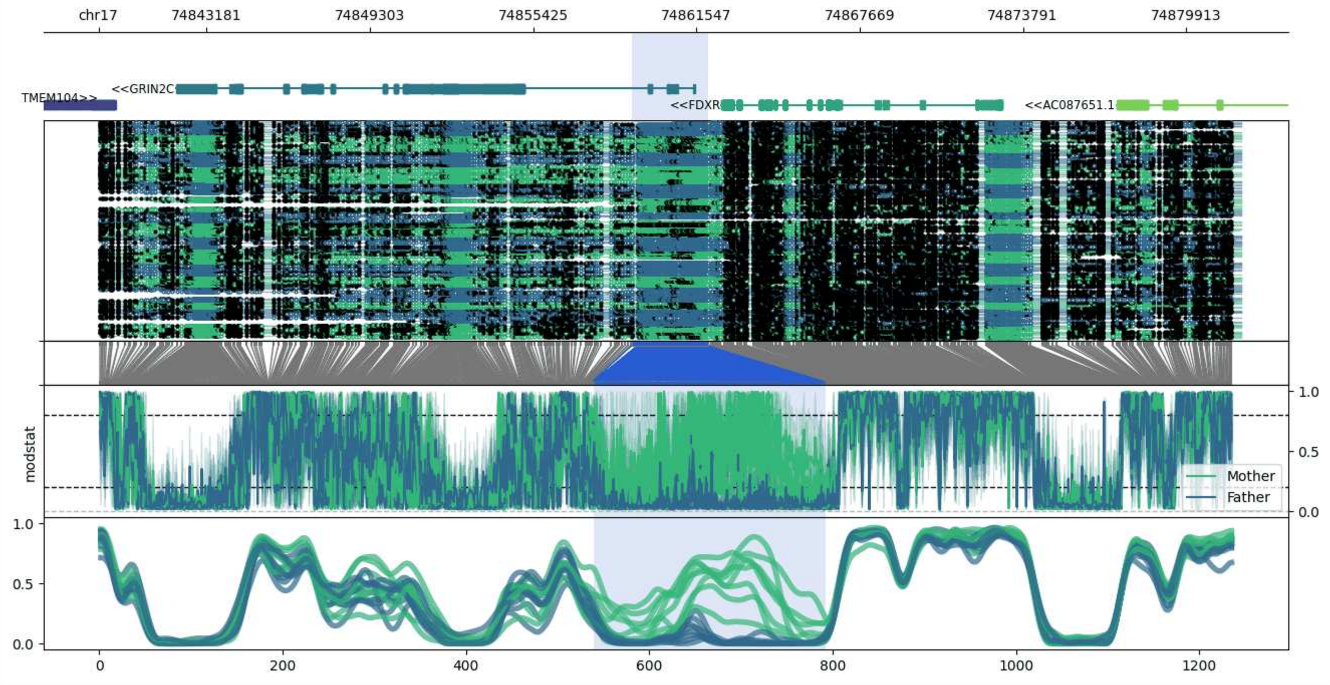


**chr17:74839124-74881984:** Region with 1 DMR containing genes: AC087651.1, FDXR, GRIN2C, TMEM104.

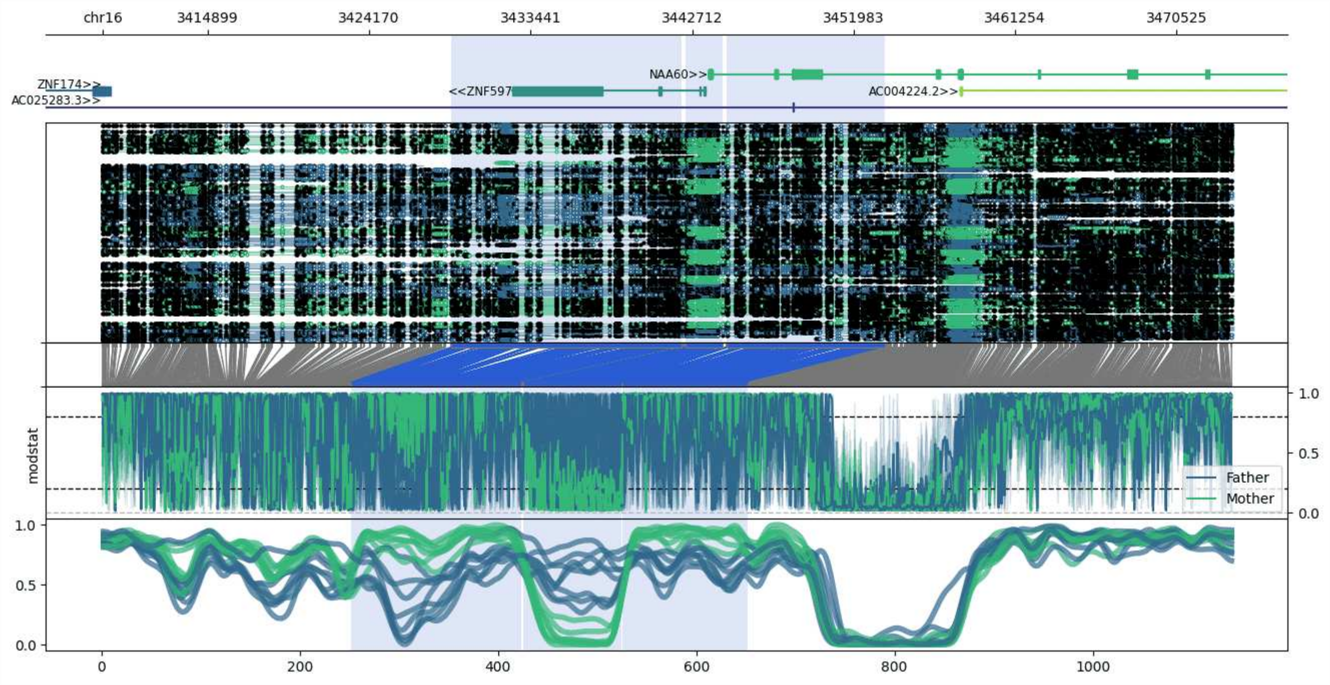


**chr16:3408864-3473763:** Region with 3 DMRs containing genes: AC004224.2, AC025283.3, NAA60, ZNF174, ZNF597, ZSCAN32.

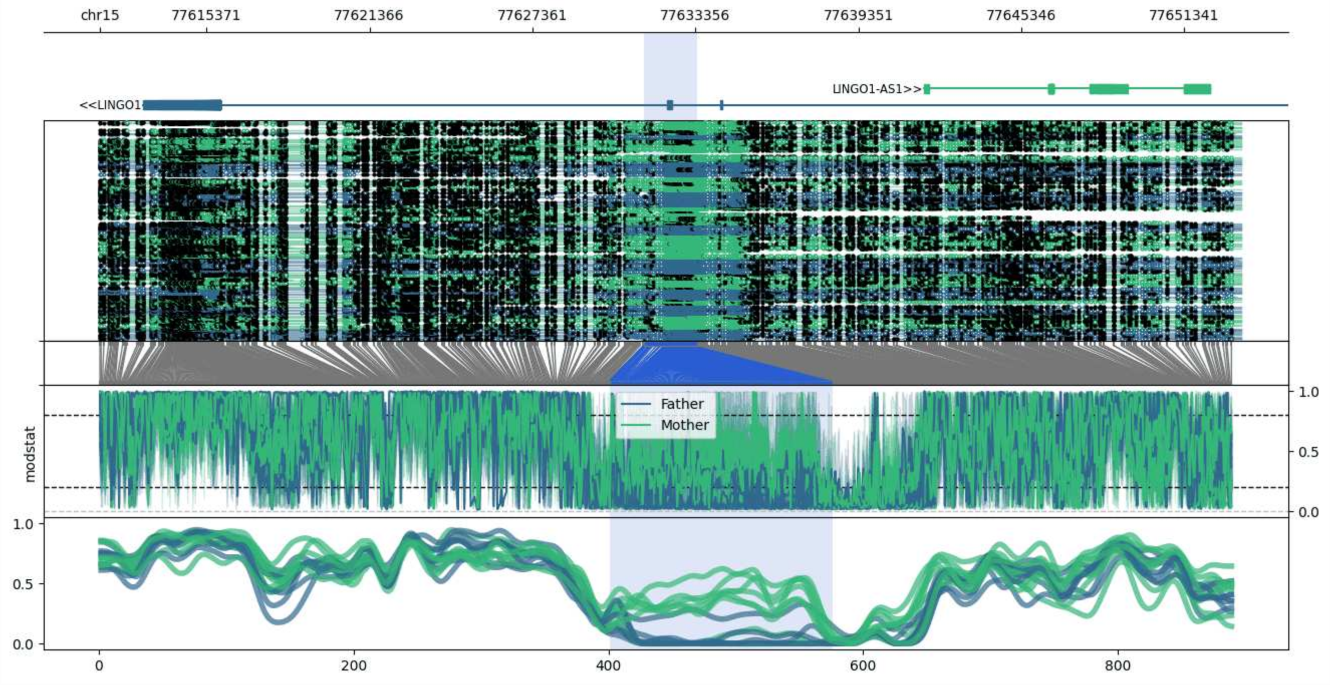


**chr15:77611454-77653421:** Region with 1 DMR containing genes: AC046168.2, LINGO1, LINGO1-AS1, LINGO1-AS2.

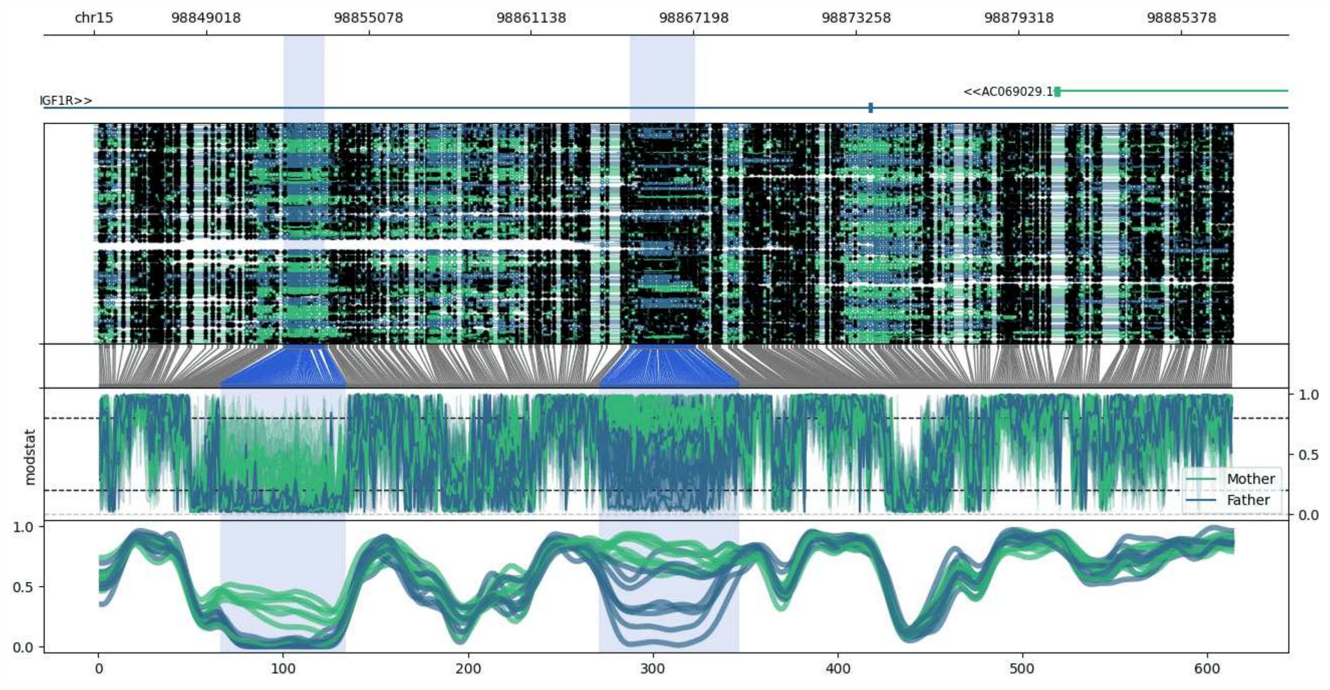


**chr15:98844844-98887267:** Region with 2 DMRs containing genes: AC055807.1, AC069029.1, IGF1R.

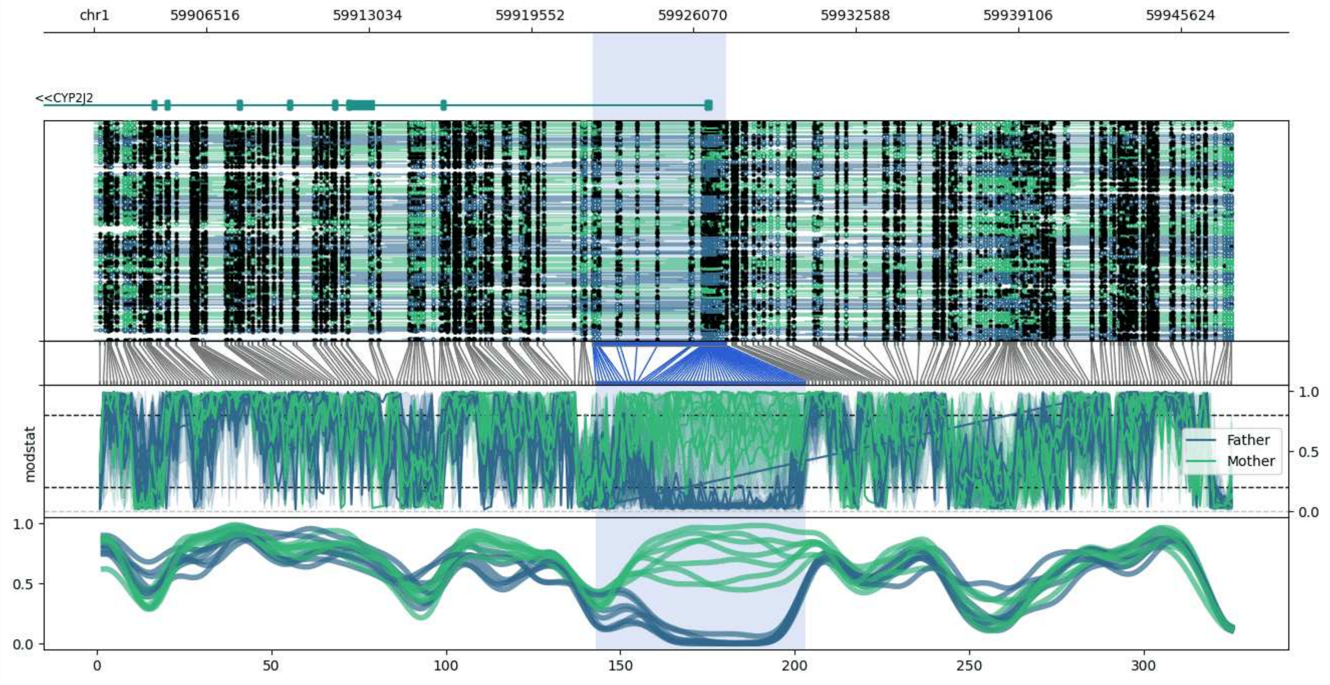


**chr1:59902033-59947665:** Region with 1 DMR containing genes: CYP2J2.

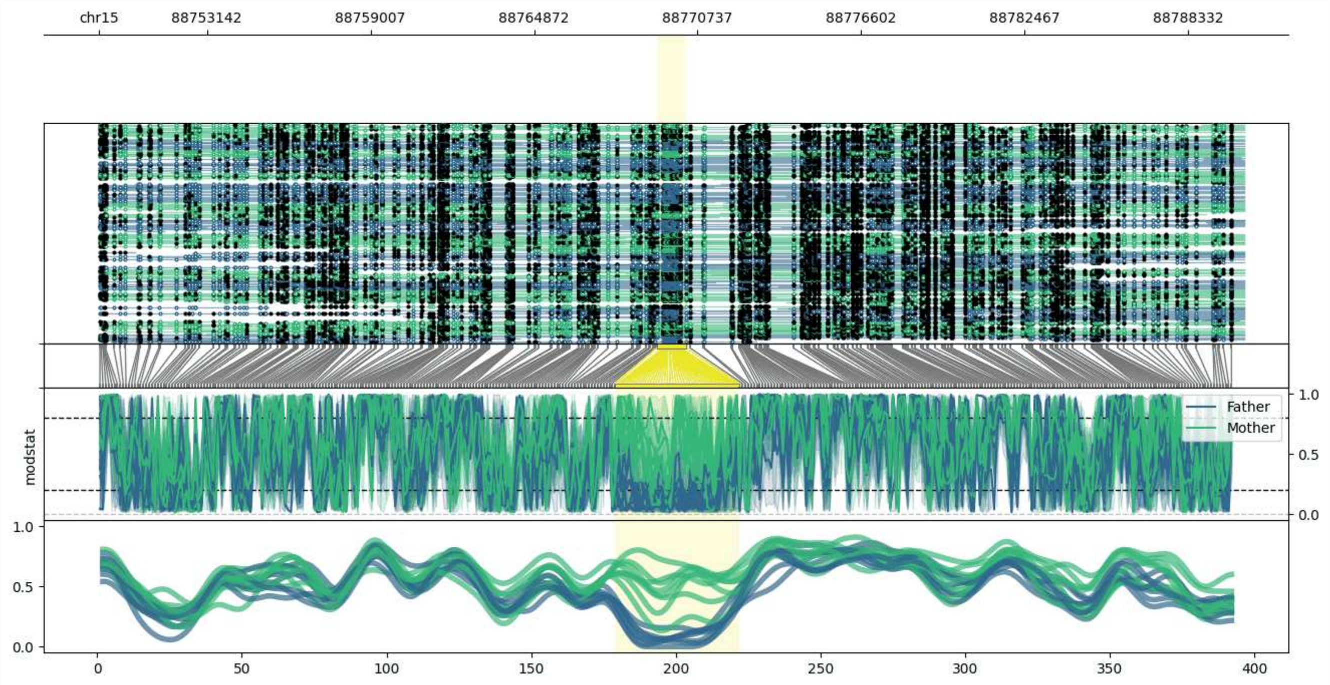


**chr15:88749266-88790321:** Region with 1 DMR containing genes: AC103982.1.

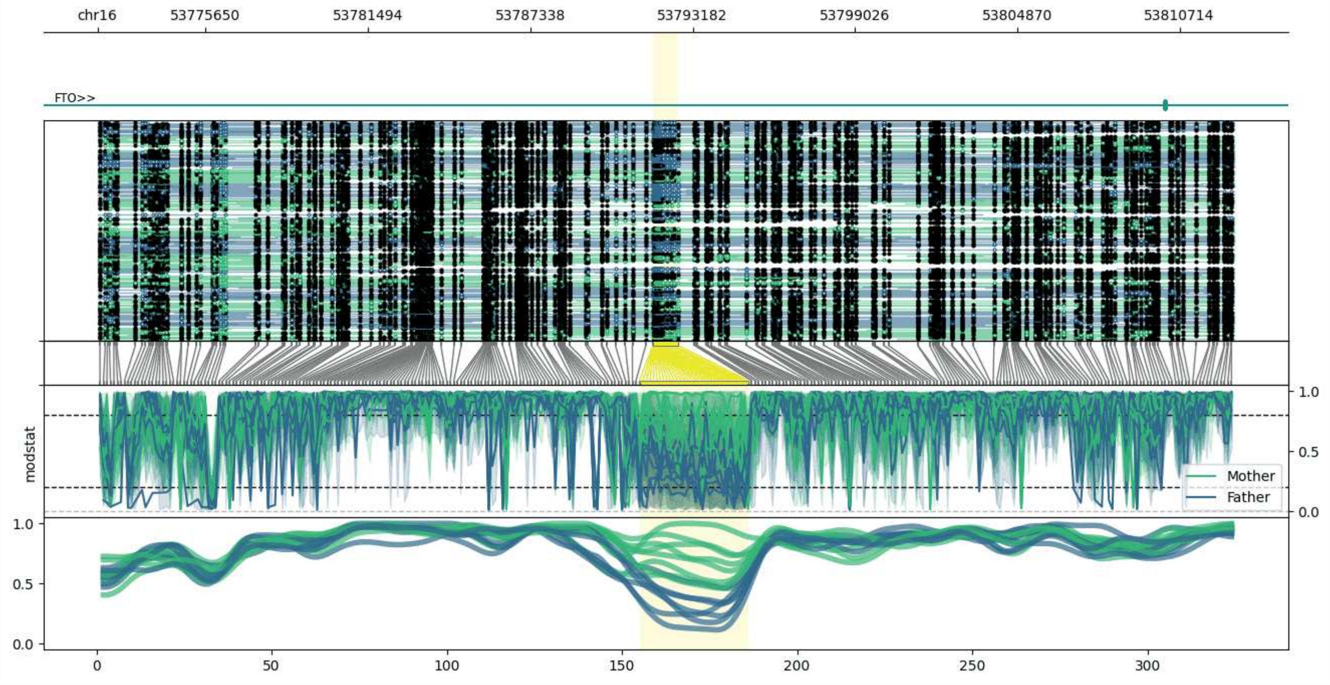


**chr16:53771753-53812667:** Region with 1 DMR containing genes: FTO.

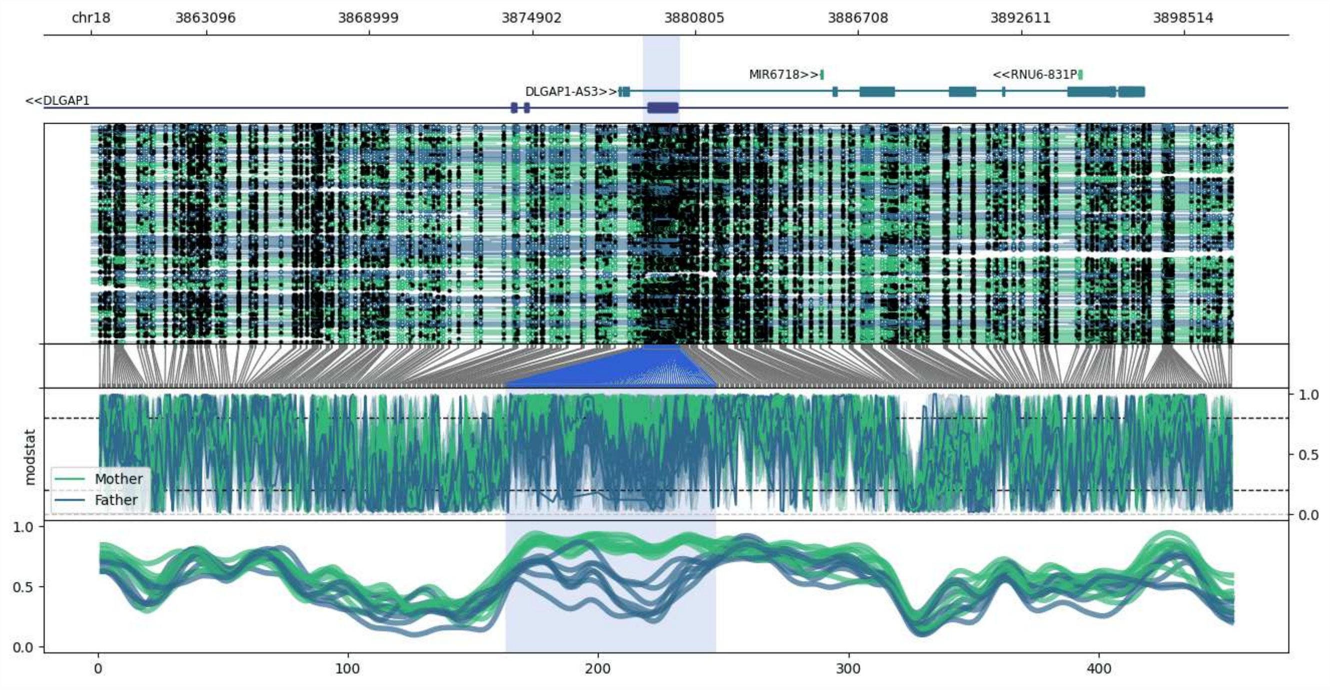


**chr18:3858917-3900243:** Region with 1 DMR containing genes: DLGAP1, DLGAP1-AS3, MIR6718, RNU6-831P.

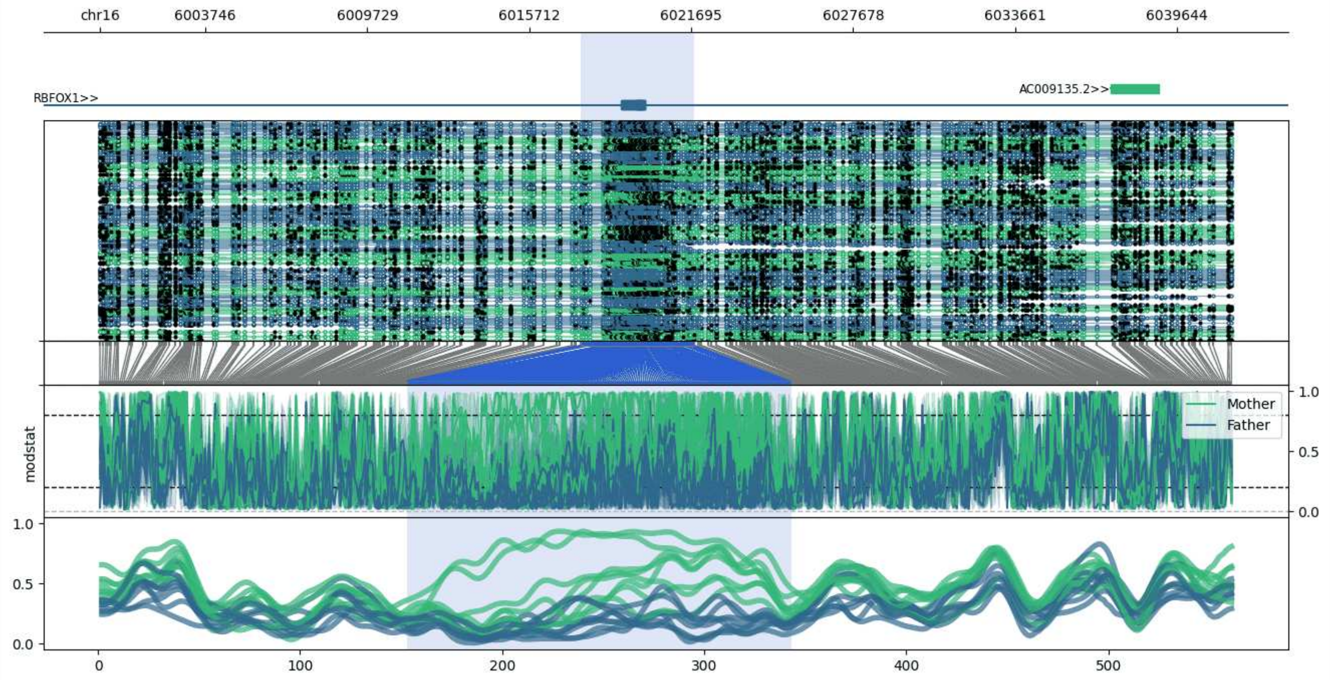


**chr16:5999852-6041734:** Region with 1 DMR containing genes: AC009135.2, RBFOX1.

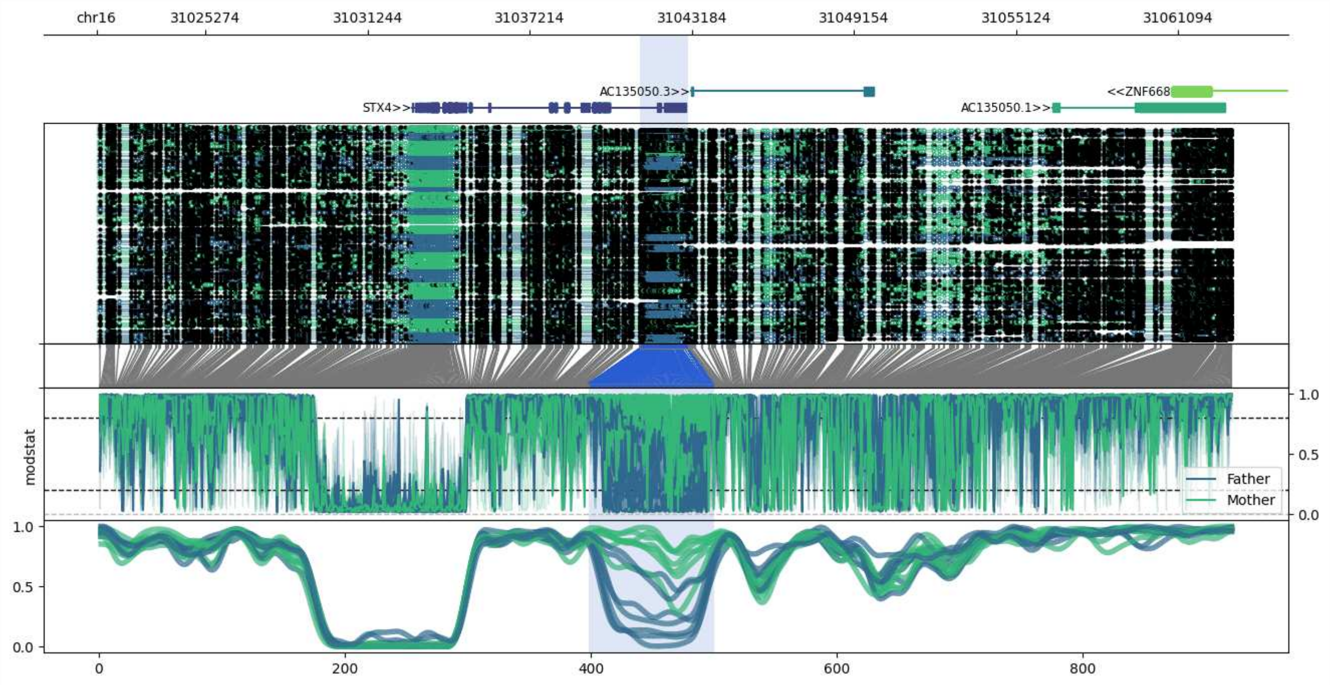


**chr16:31021256-31063049:** Region with 1 DMR containing genes: AC135050.1, AC135050.3, AC135050.4, STX4, ZNF668.

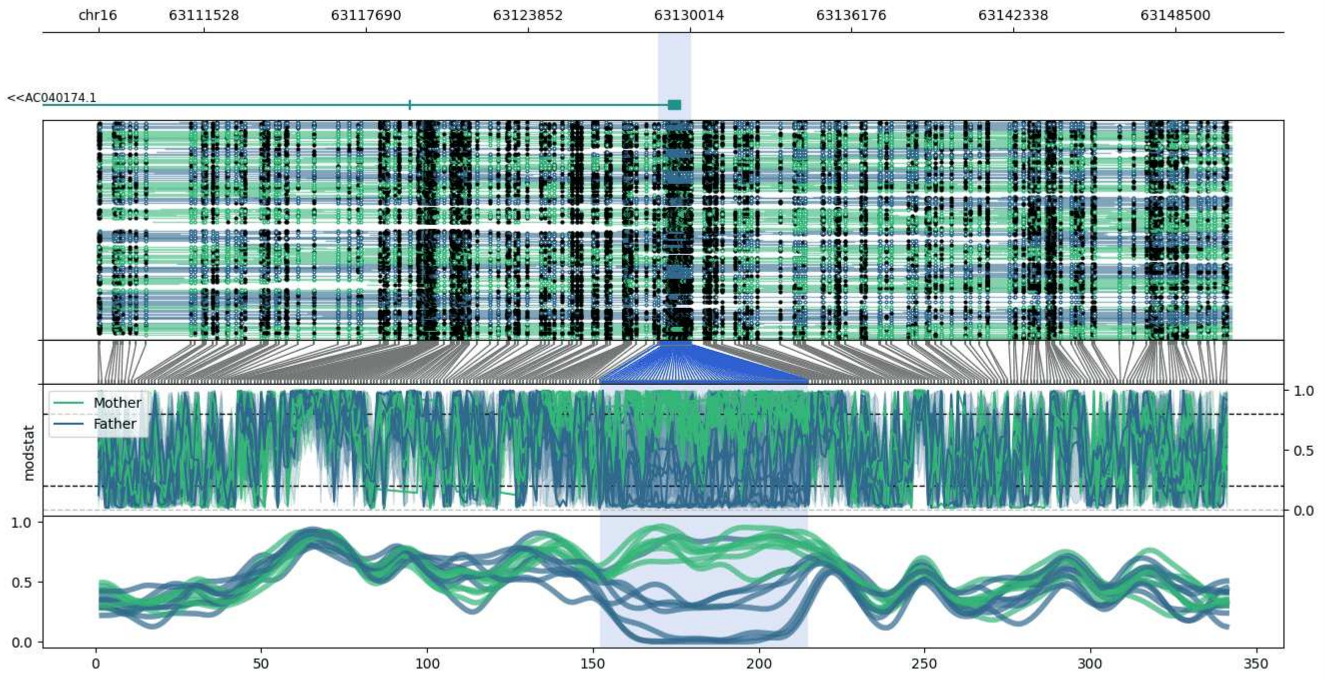


**chr16:63107501-63150641:** Region with 1 DMR containing genes: AC040174.1.

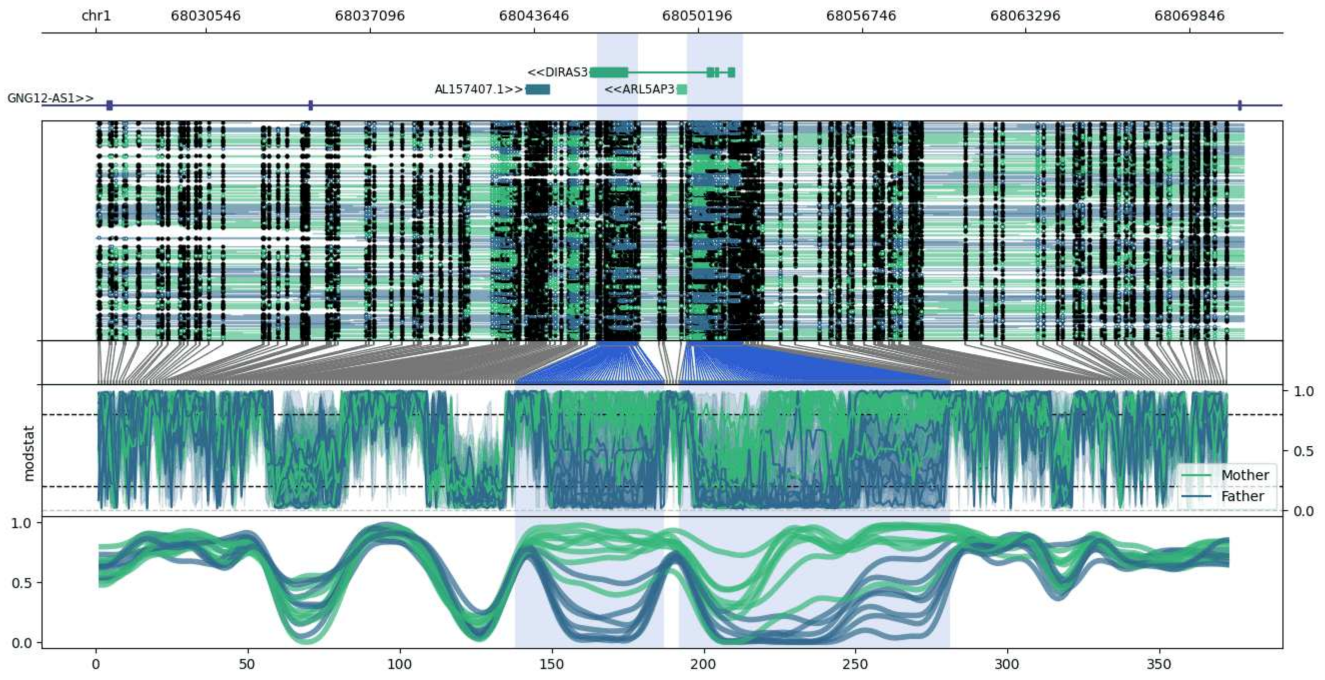


**chr1:68026152-68072007:** Region with 2 DMRs containing genes: AL157407.1, ARL5AP3, DIRAS3, GNG12-AS1.

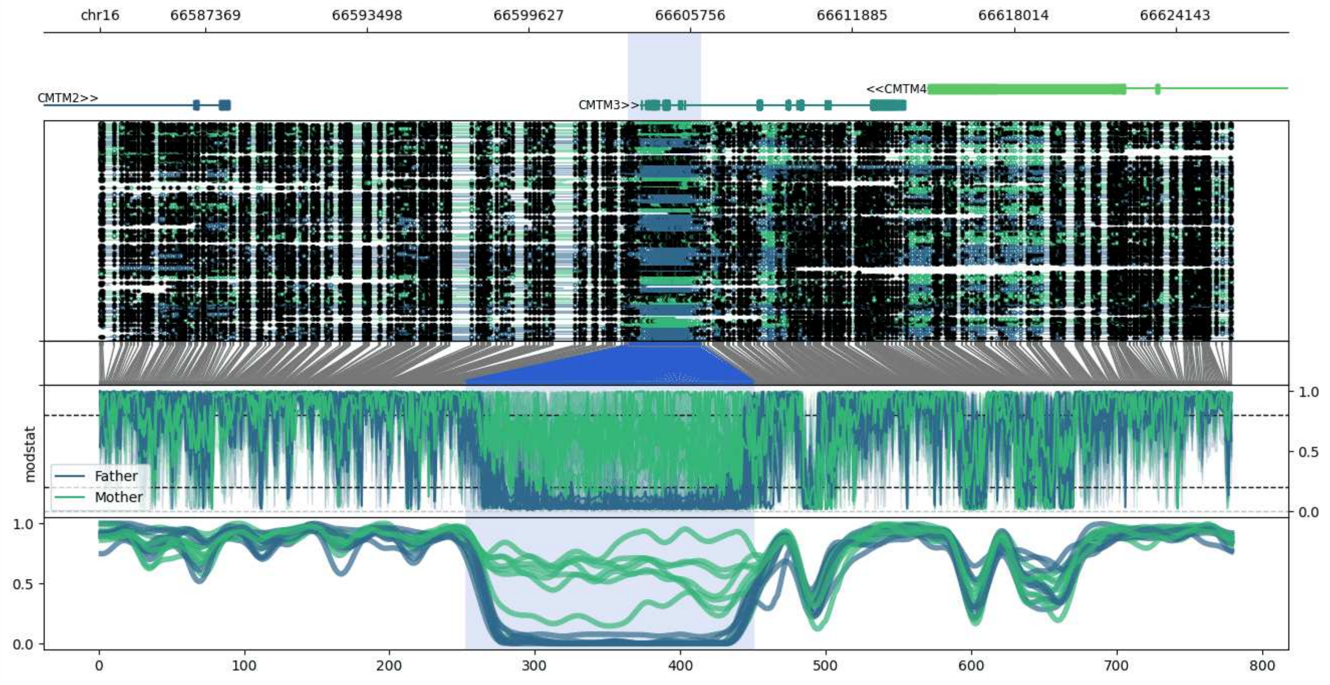


**chr16:66583379-66626283:** Region with 1 DMR containing genes: AC010542.1, CKLF-CMTM1, CMTM1, CMTM2, CMTM3, CMTM4.

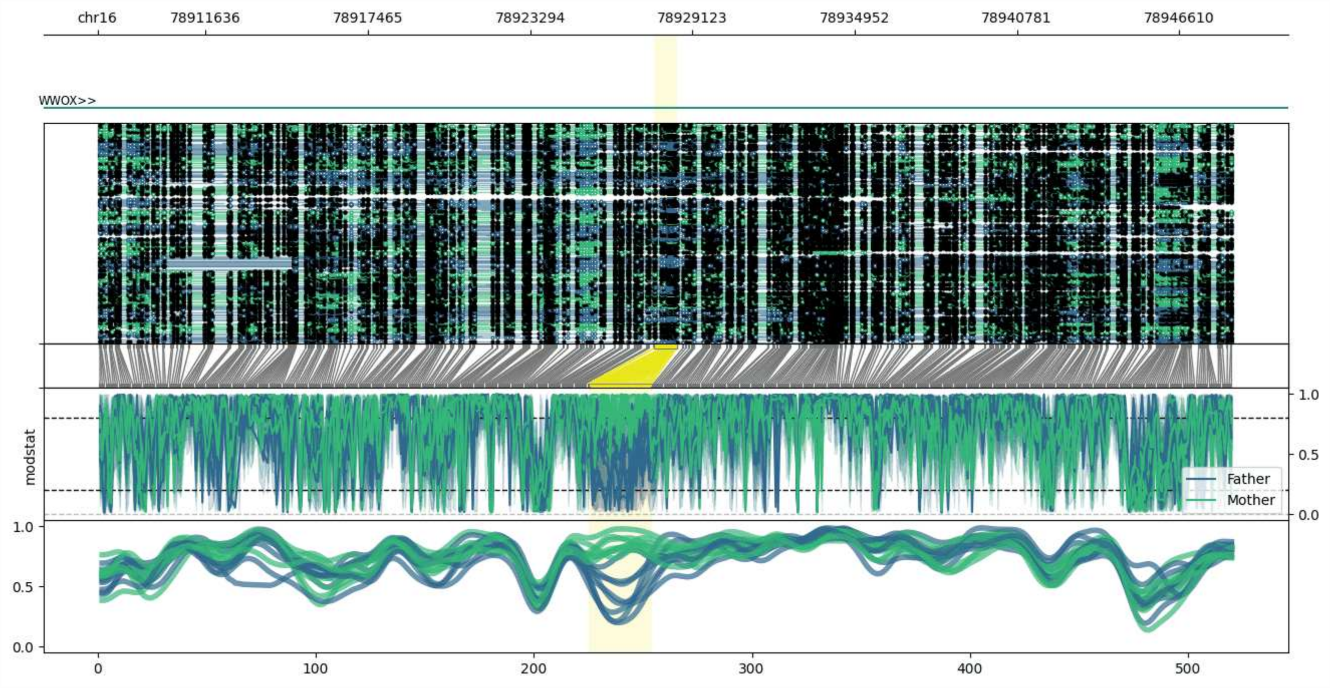


**chr16:78907751-78948555:** Region with 1 DMR containing genes: AC027279.1, WWOX.

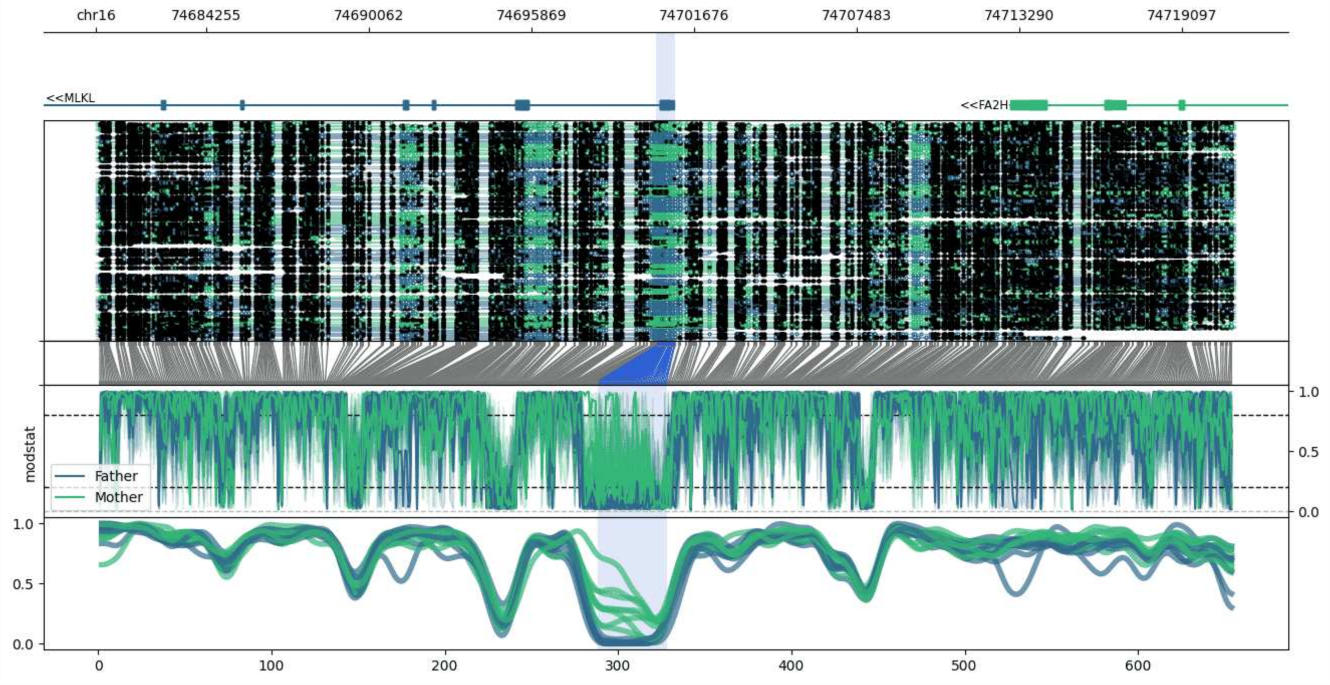


**chr16:74680332-74720987:** Region with 1 DMR containing genes: FA2H, MLKL.

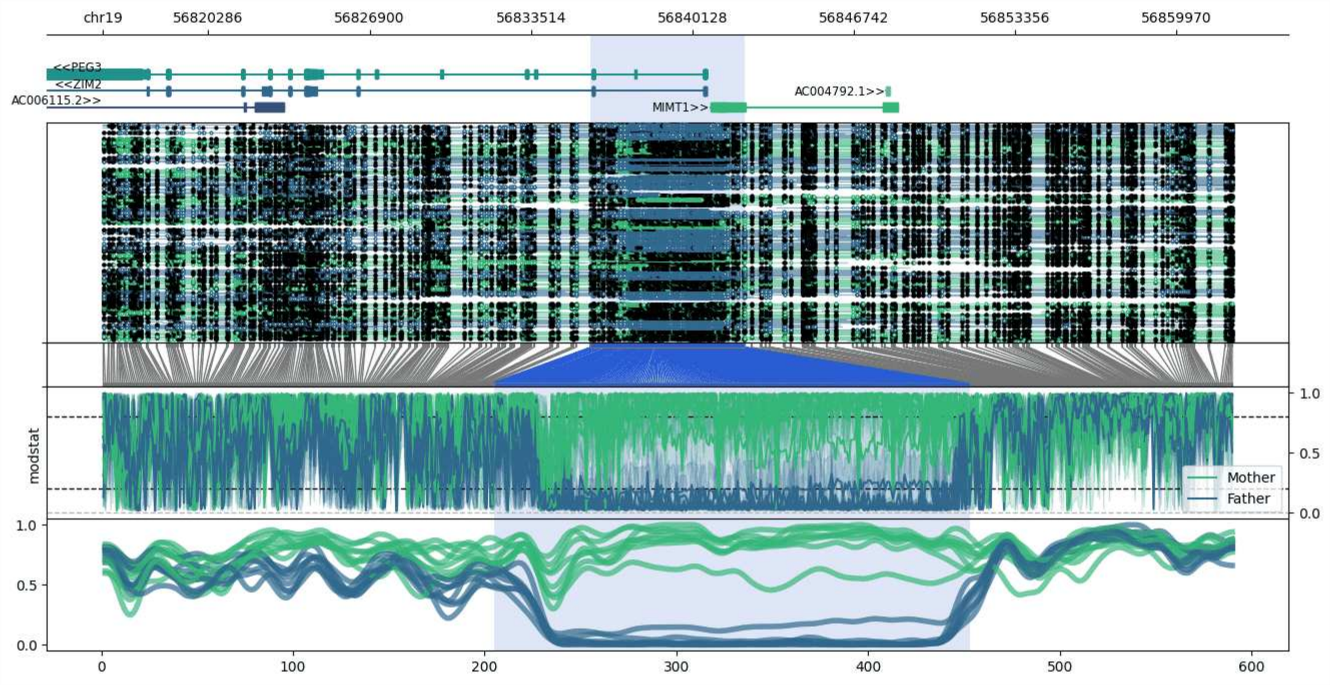


**chr19:56815964-56862264:** Region with 1 DMR containing genes: AC004792.1, AC006115.2, MIMT1, PEG3, ZIM2.

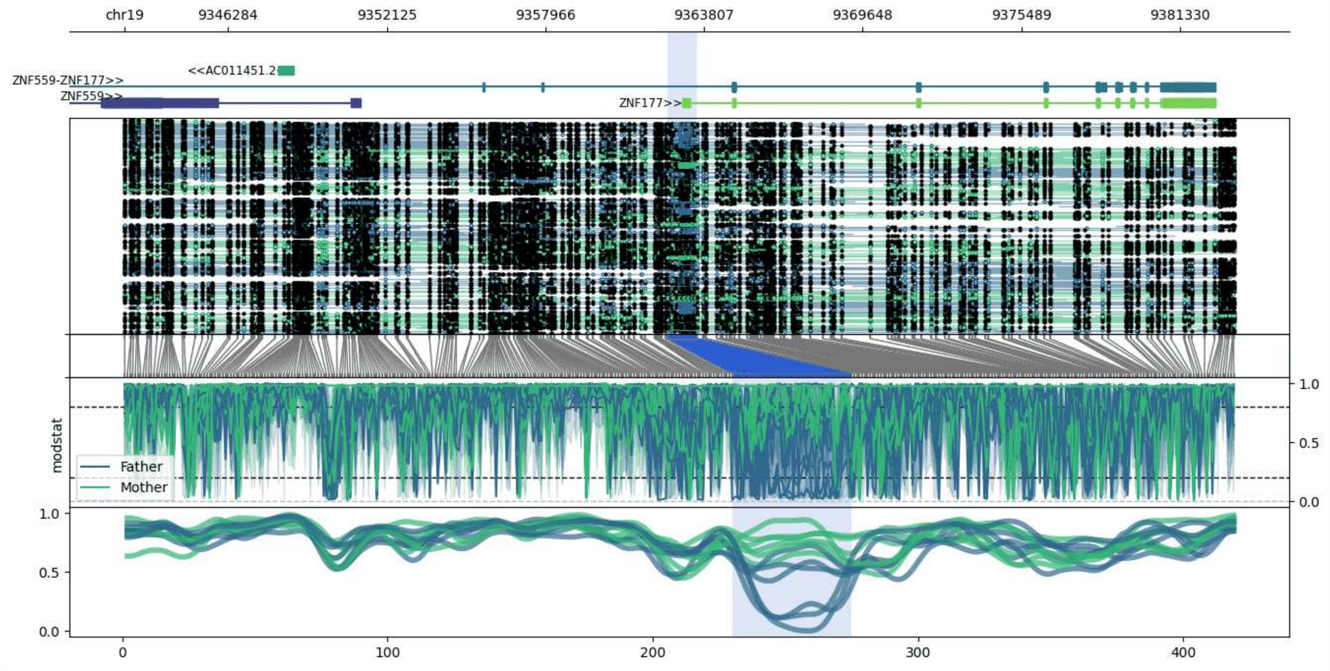


**chr19:9342474-9383366:** Region with 1 DMR containing genes: AC011451.2, AC011451.3, ZNF177, ZNF559, ZNF559-ZNF177.

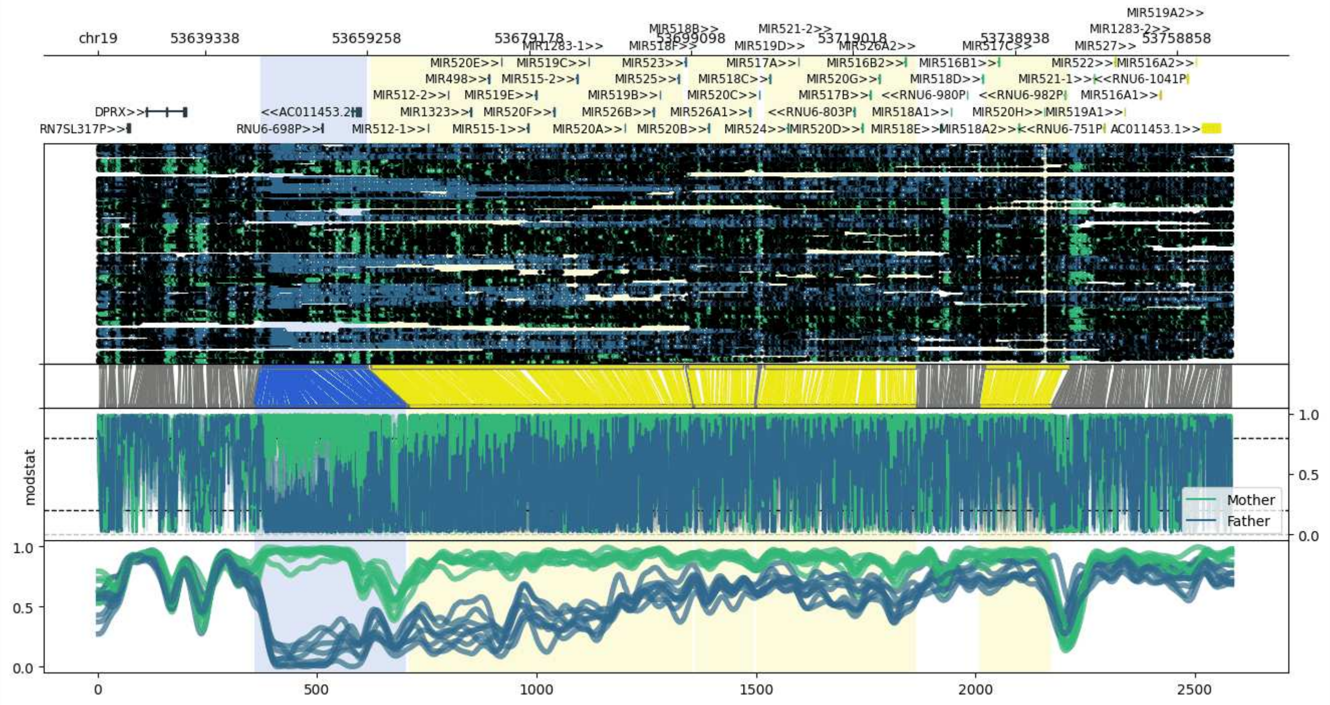


**chr19:53626141-53765585:** Region with 5 DMRs containing genes: AC011453.1, AC011453.2, AC011487.3, DPRX, HMGN1P32, MIR1283-1, MIR1283-2, MIR1323, MIR498, MIR512-1, MIR512-2, MIR515-1, MIR515-2, MIR516A1, MIR516A2, MIR516B1, MIR516B2, MIR517A, MIR517B, MIR517C, MIR518A1, MIR518A2, MIR518B, MIR518C, MIR518D, MIR518E, MIR518F, MIR519A1, MIR519A2, MIR519B, MIR519C, MIR519D, MIR519E, MIR520A, MIR520B, MIR520C, MIR520D, MIR520E, MIR520F, MIR520G, MIR520H, MIR521-1, MIR521-2, MIR522, MIR523, MIR524, MIR525, MIR526A1, MIR526A2, MIR526B, MIR527, RN7SL317P, RNU6-1041P, RNU6-165P, RNU6-698P, RNU6-751P, RNU6-803P, RNU6-980P, RNU6-982P, SEPT7P8.

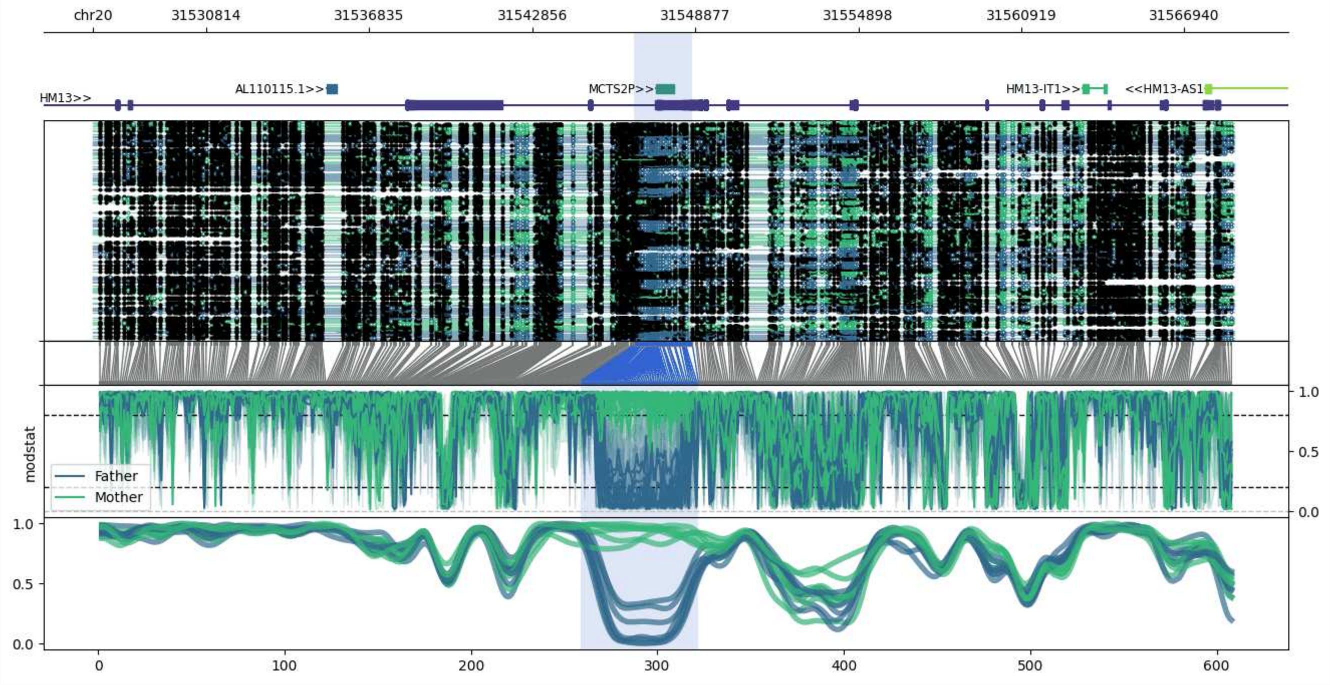


**chr20:31526621-31568772:** Region with 1 DMR containing genes: AL110115.1, HM13, HM13-AS1, HM13-IT1, MCTS2P.

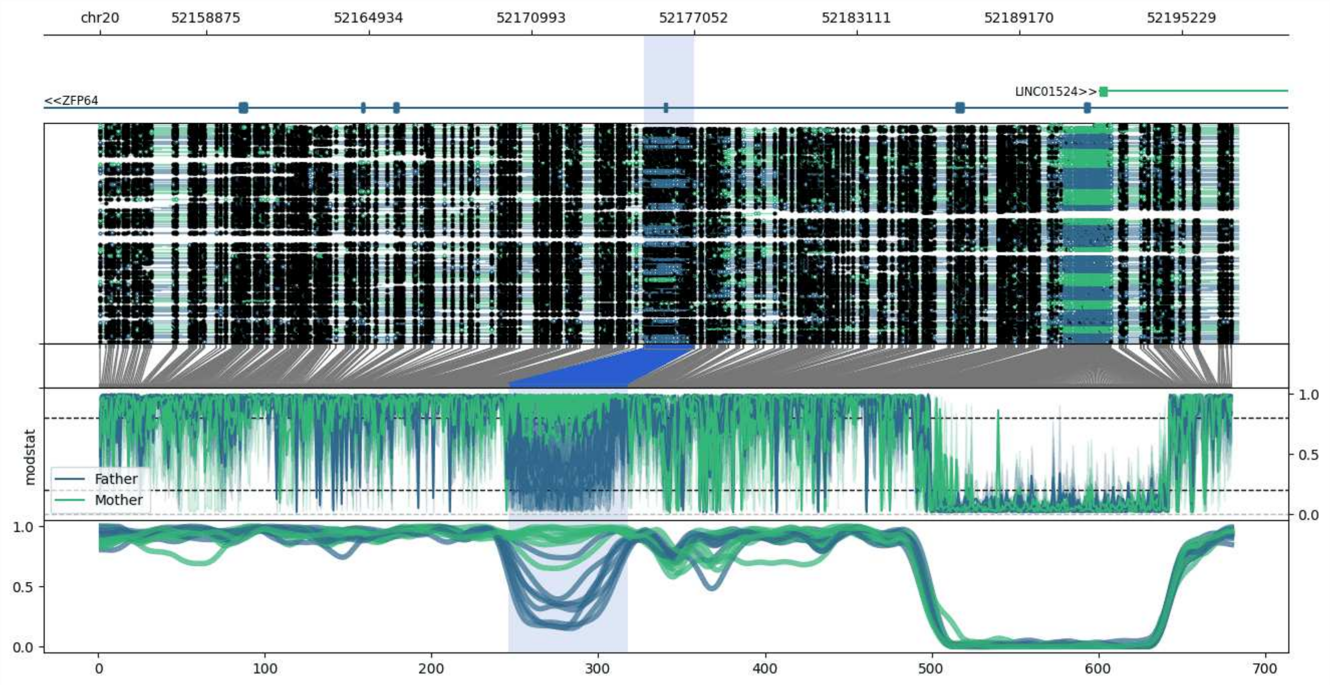


**chr20:52154904-52197320:** Region with 1 DMR containing genes: LINC01524, ZFP64.

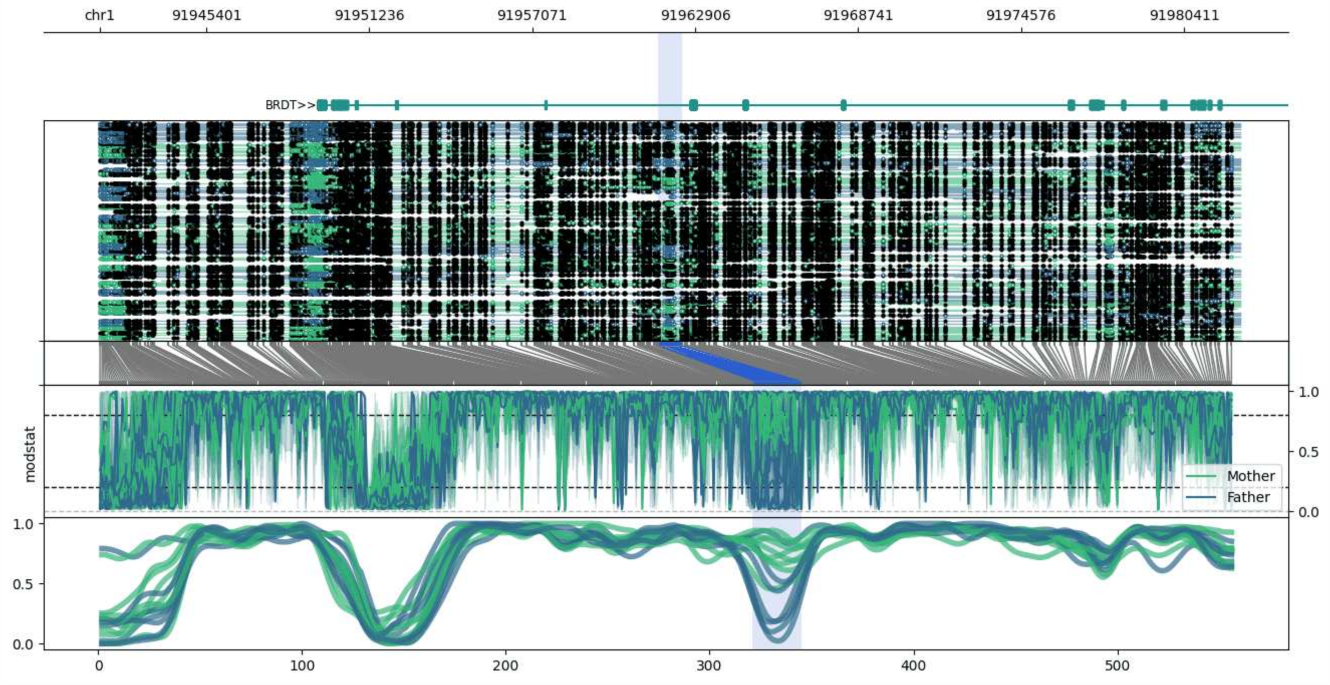


**chr1:91941577-91982426:** Region with 1 DMR containing genes: BRDT.

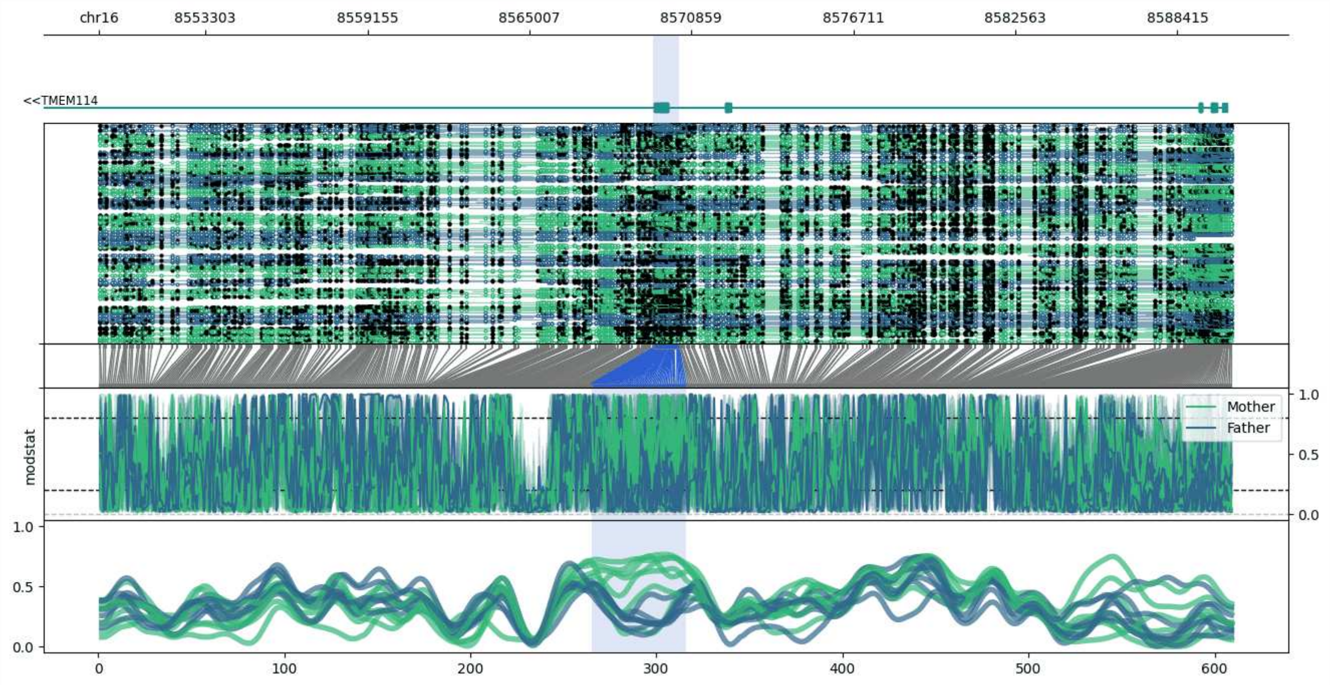


**chr16:8549458-8590423:** Region with 1 DMR containing genes: TMEM114.

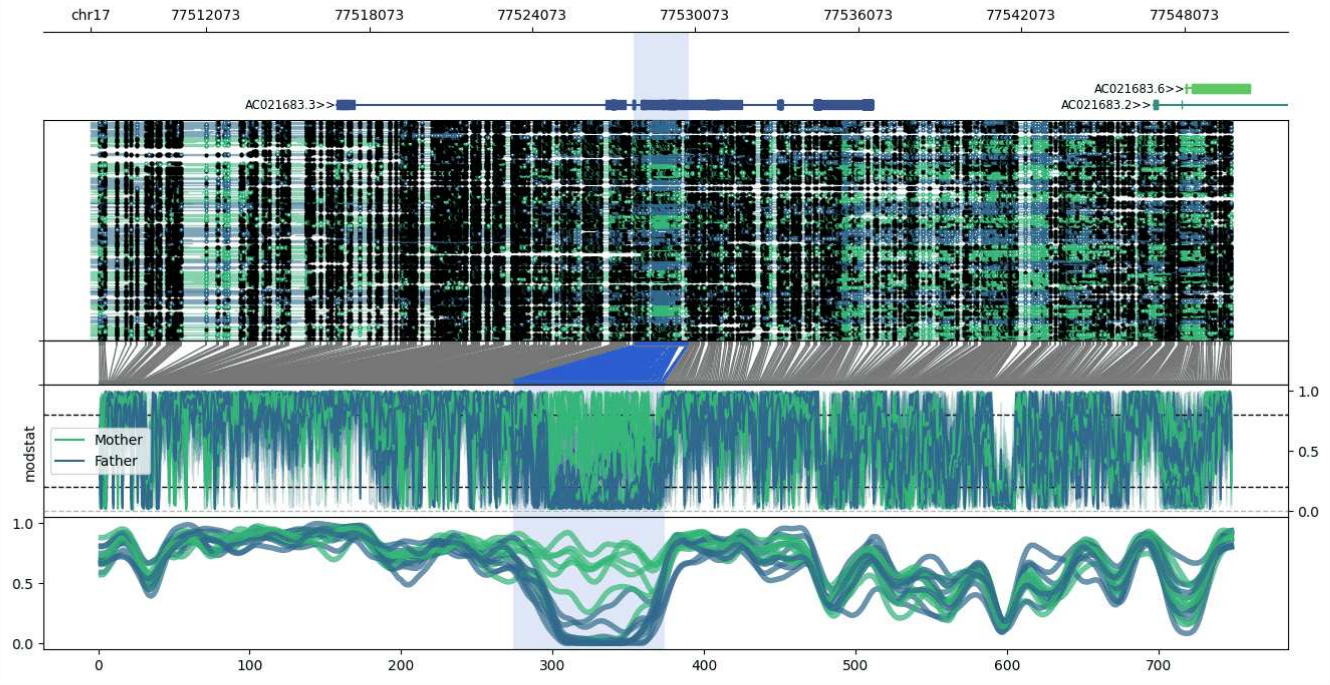


**chr17:77507815-77549821:** Region with 1 DMR containing genes: AC021683.2, AC021683.3, AC021683.6, SEPT9.

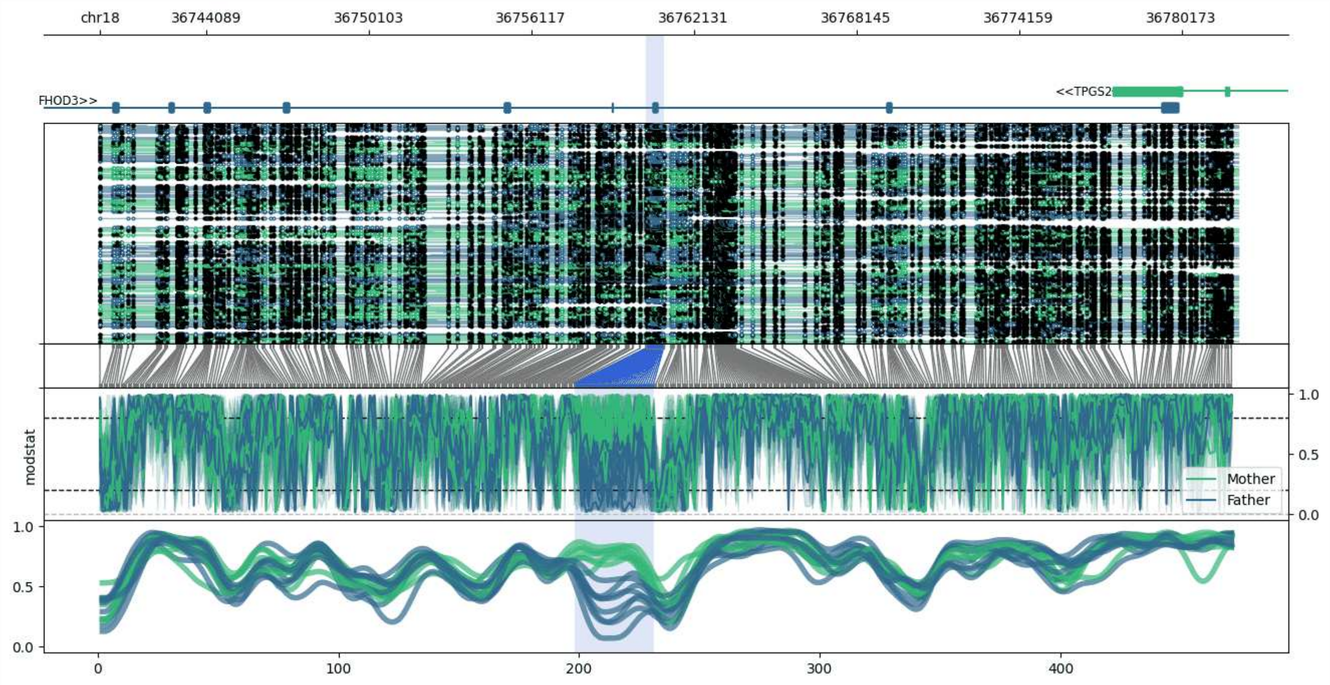


**chr18:36740155-36782255:** Region with 1 DMR containing genes: FHOD3, TPGS2.

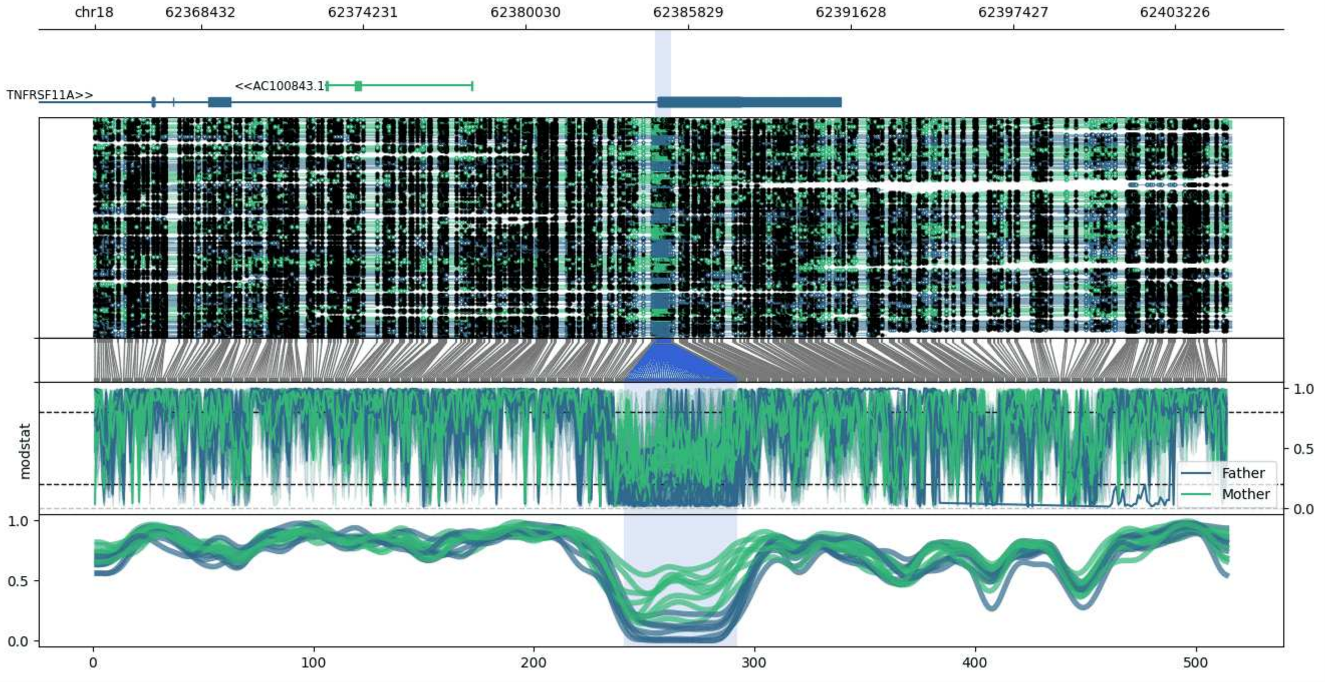


**chr18:62364637-62405231:** Region with 1 DMR containing genes: AC100843.1, TNFRSF11A.

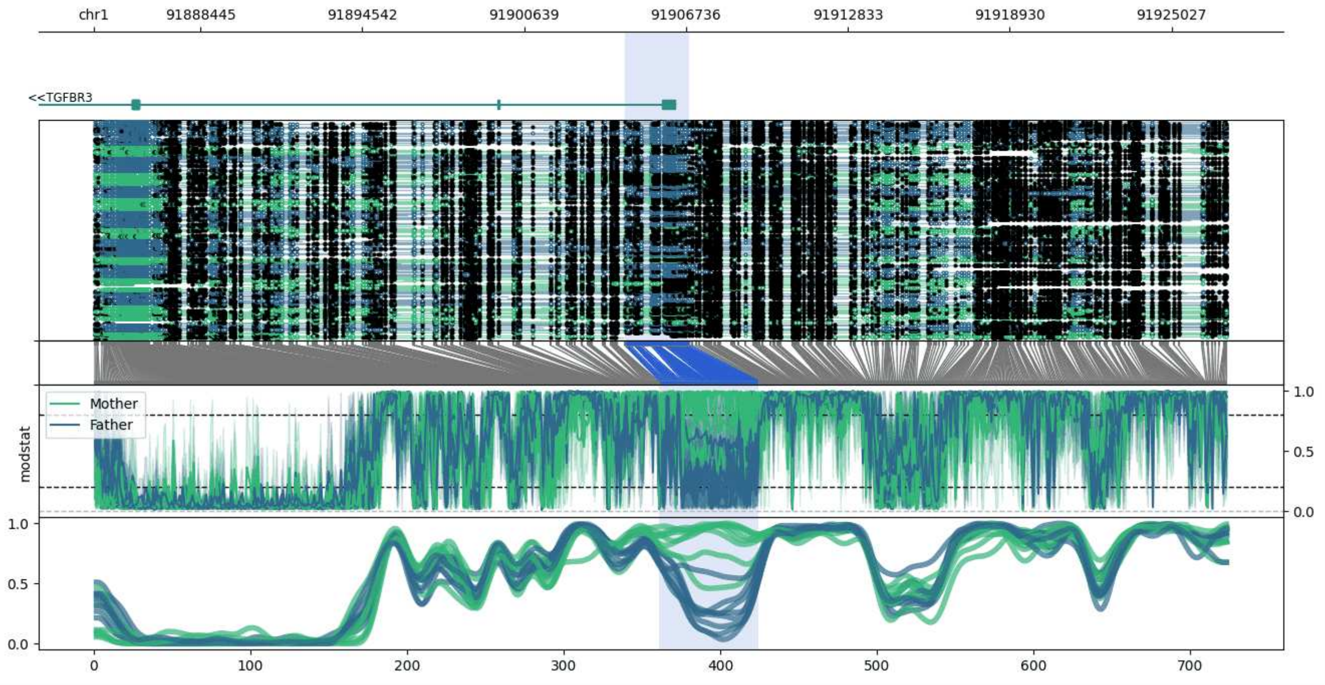


**chr1:91884434-91927115:** Region with 1 DMR containing genes: TGFBR3.

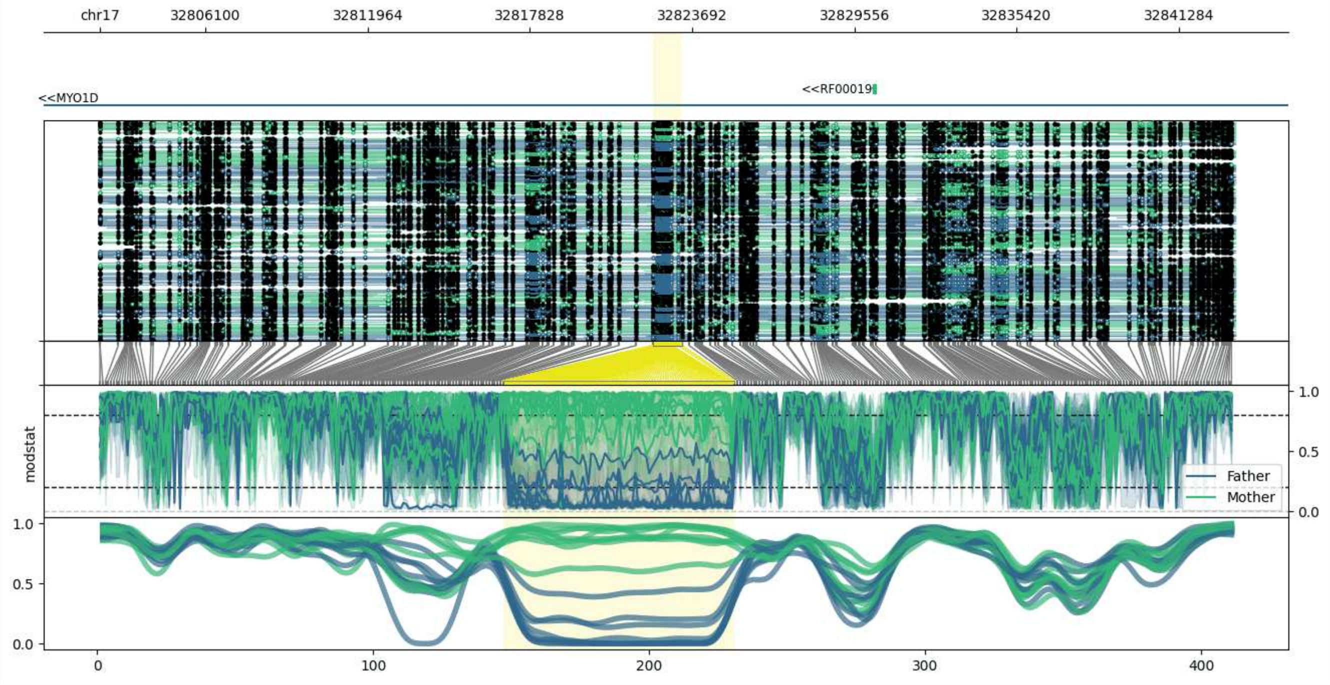


**chr17:32802266-32843316:** Region with 1 DMR containing genes: MYO1D, RF00019.

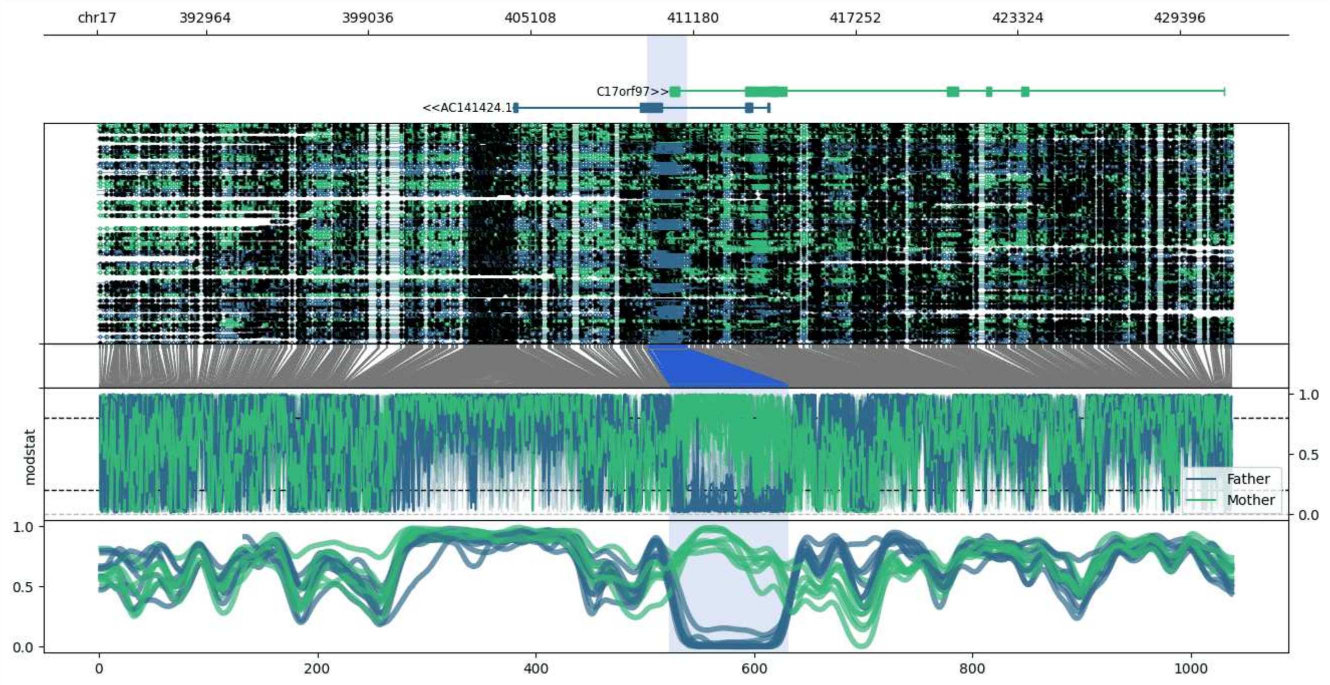


**chr17:388885-431389:** Region with 1 DMR containing genes: AC141424.1, C17orf97, RFLNB, RPH3AL.

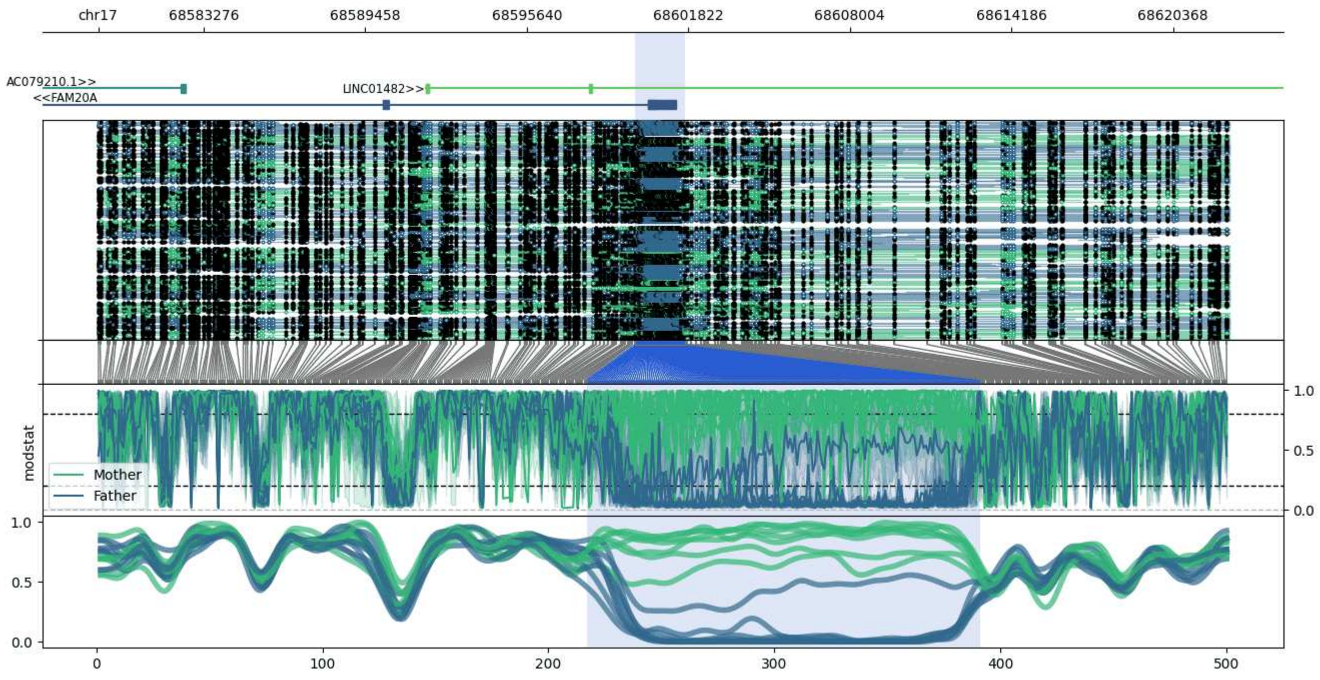


**chr17:68579238-68622517:** Region with 1 DMR containing genes: AC079210.1, FAM20A, LINC01482.

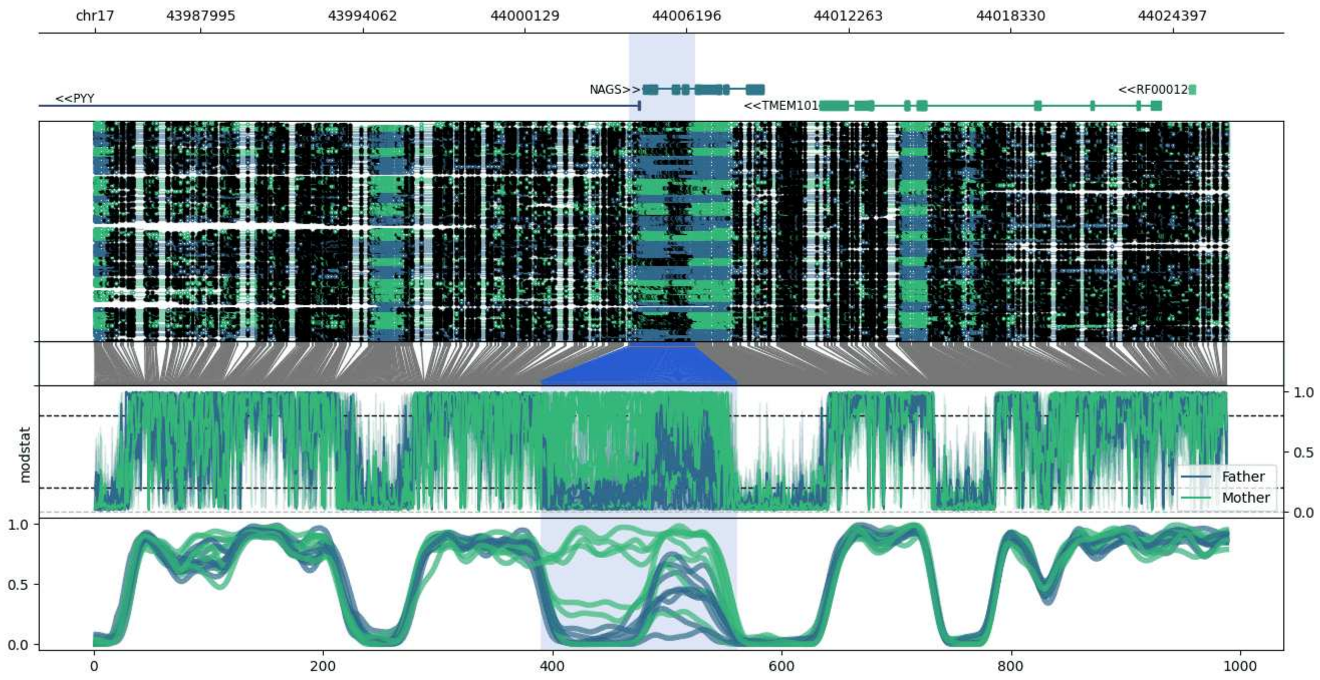


**chr17:43984045-44026519:** Region with 1 DMR containing genes: LSM12, NAGS, PYY, RF00012, TMEM101.

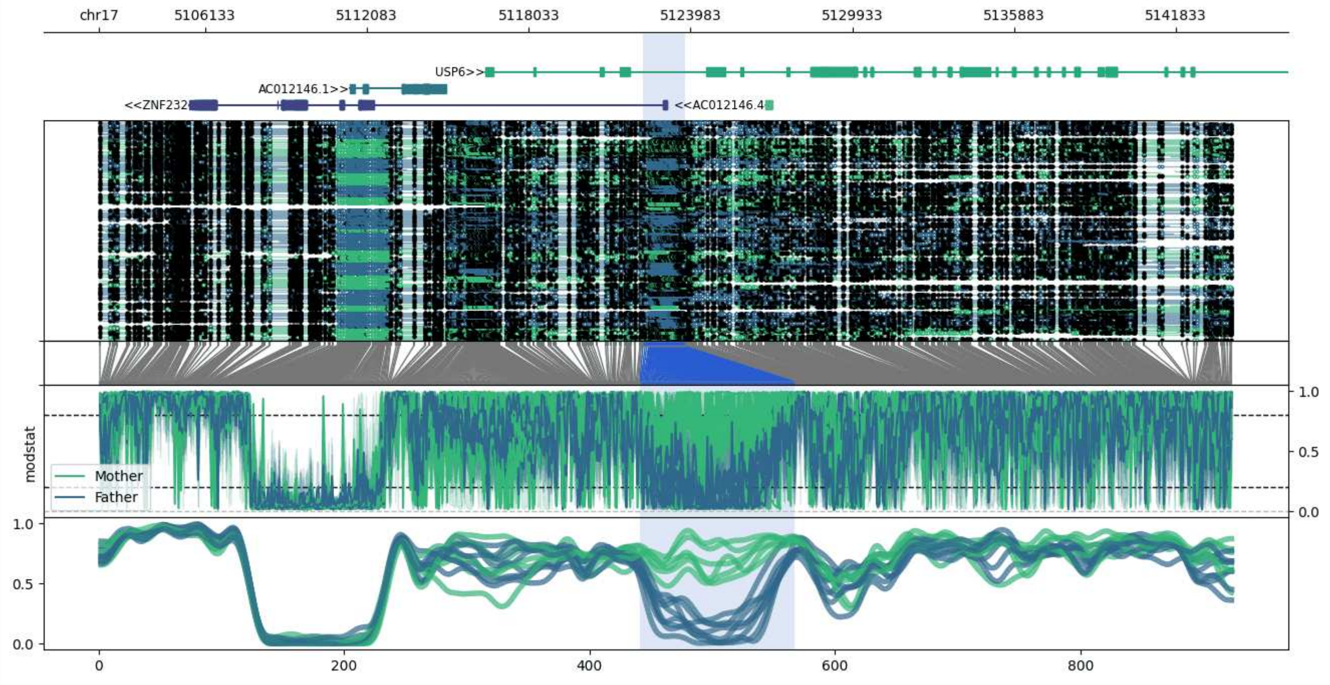


**chr17:5102232-5143888:** Region with 1 DMR containing genes: AC012146.1, AC012146.4, USP6, ZFP3, ZNF232.

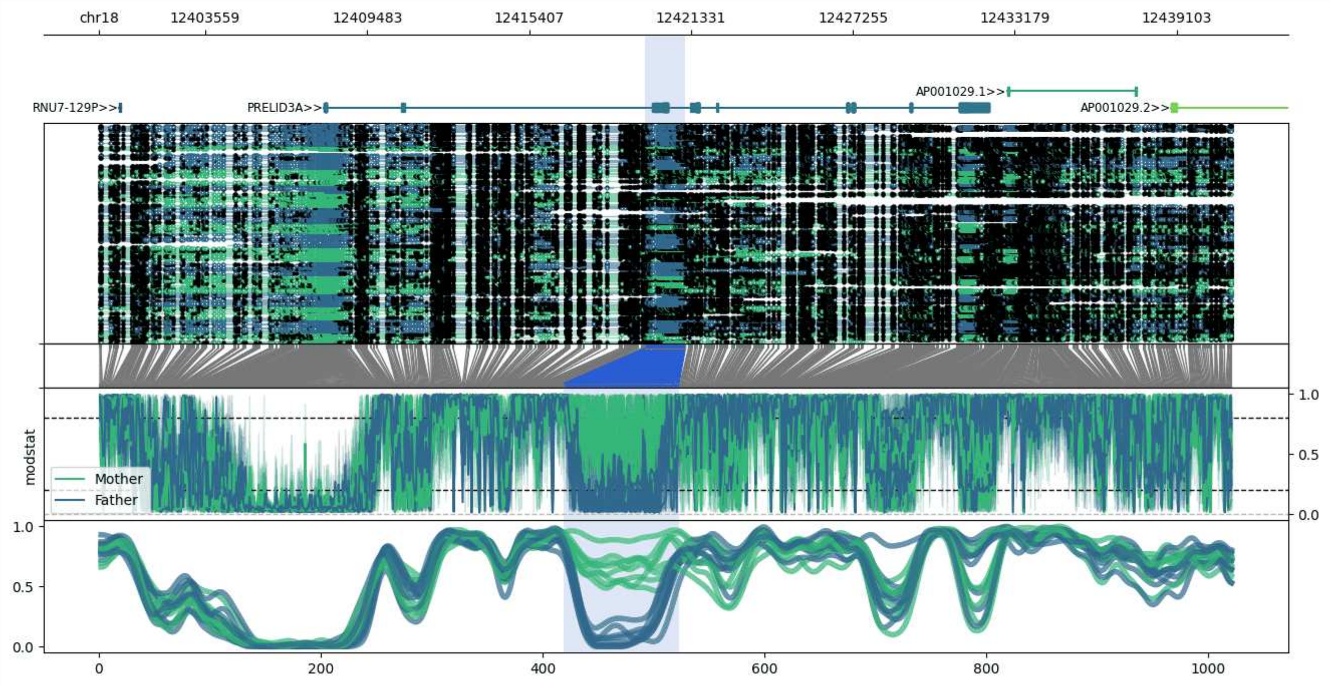


**chr18:12399660-12441133:** Region with 1 DMR containing genes: AP001029.1, AP001029.2, AP001029.3, PRELID3A, RNU7-129P, SPIRE1.

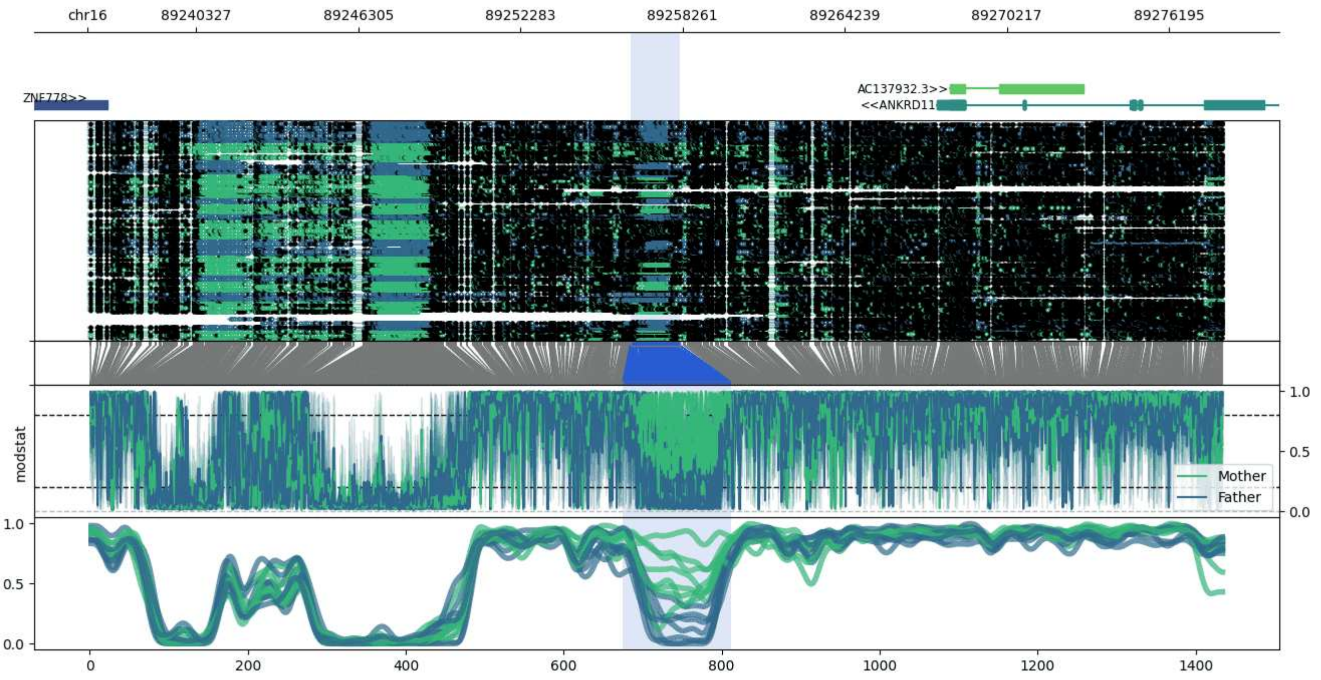


**chr16:89236329-89278177:** Region with 1 DMR containing genes: AC009113.2, AC137932.3, ANKRD11, ZNF778.

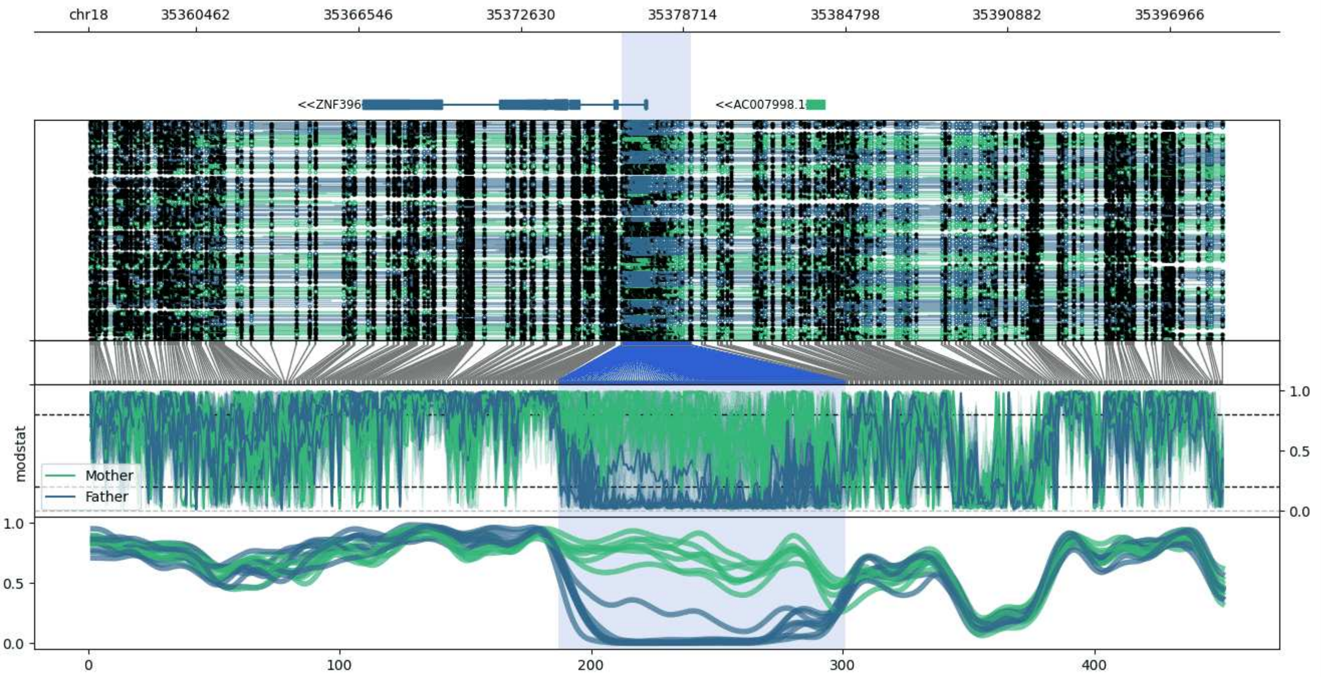


**chr18:35356409-35398997:** Region with 1 DMR containing genes: AC007998.1, AC007998.4, ZNF396.

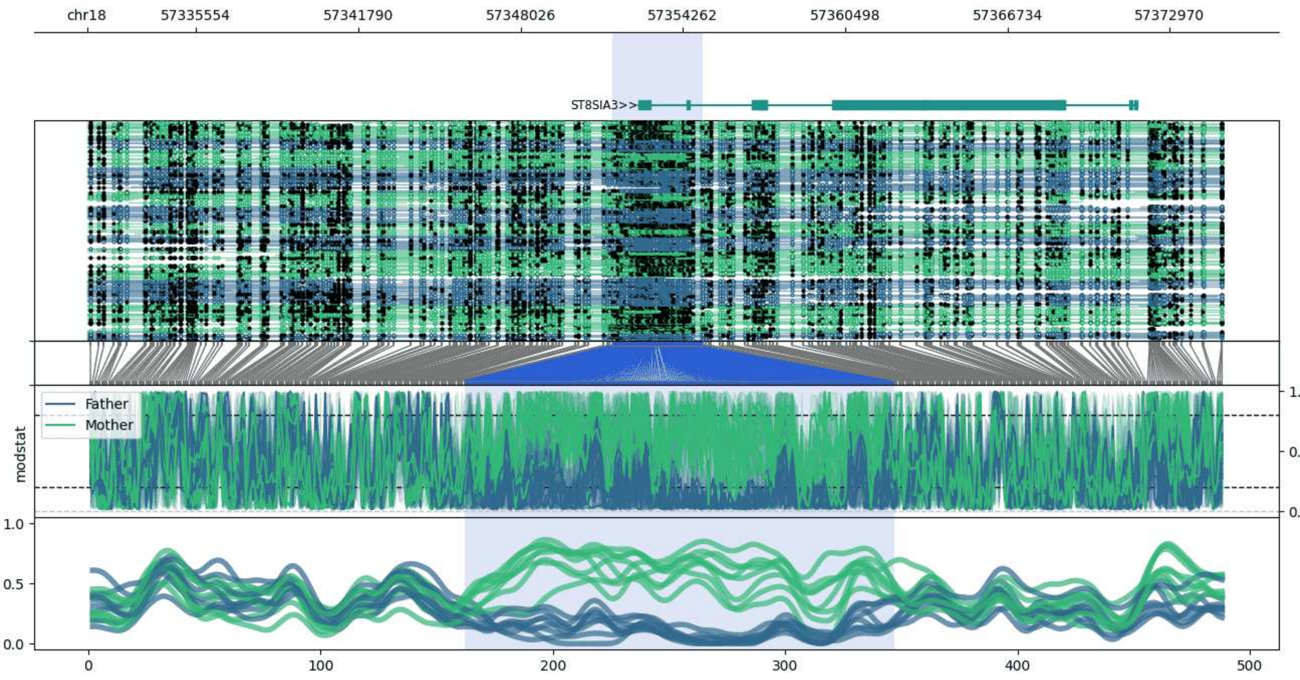


**chr18:57331356-57375008:** Region with 1 DMR containing genes: ST8SIA3.

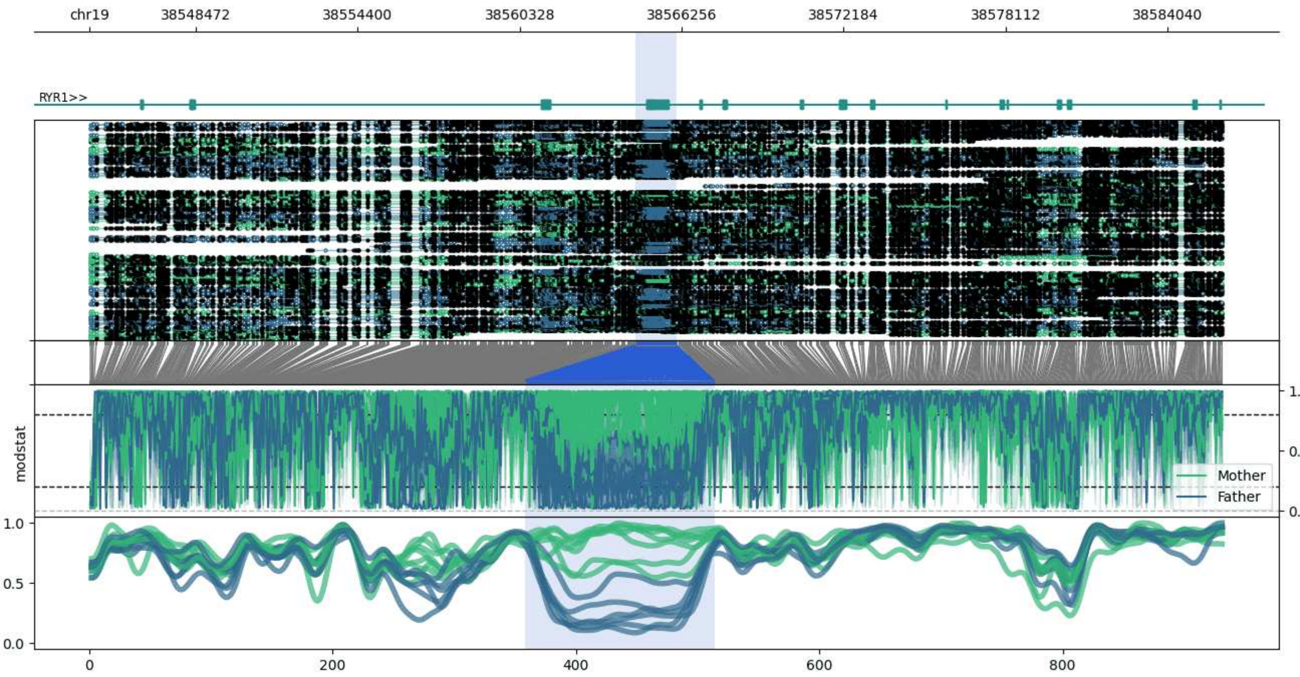


**chr19:38544579-38586079:** Region with 1 DMR containing genes: AC067969.1, MAP4K1, RYR1.

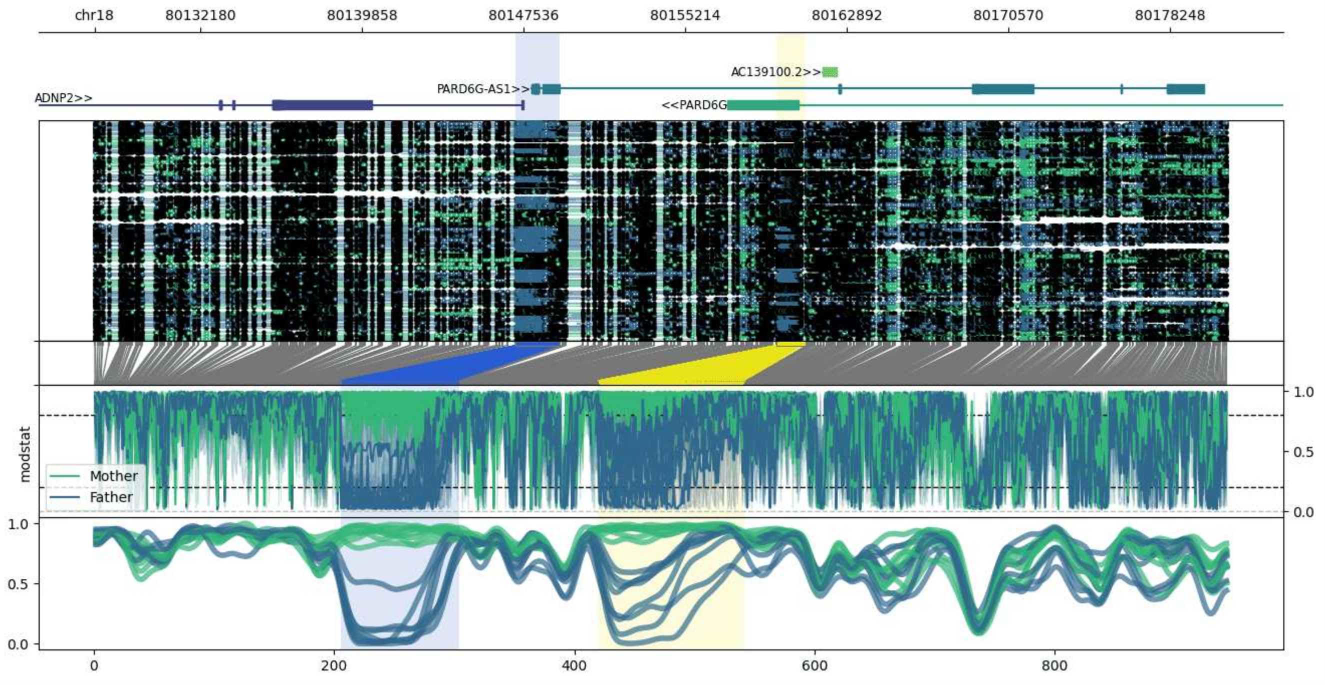


**chr18:80127167-80180916:** Region with 2 DMRs containing genes: AC139100.1, AC139100.2, ADNP2, PARD6G, PARD6G-AS1.

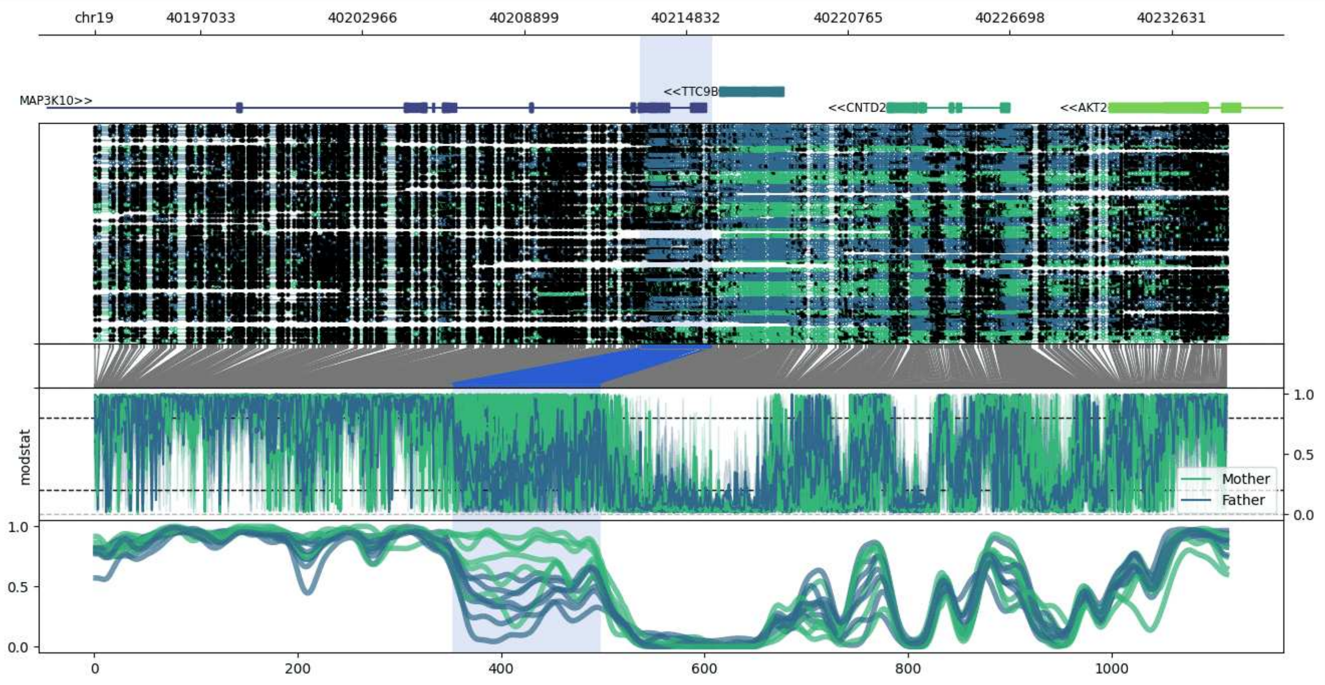


**chr19:40193156-40234688:** Region with 1 DMR containing genes: AKT2, CNTD2, MAP3K10, TTC9B.

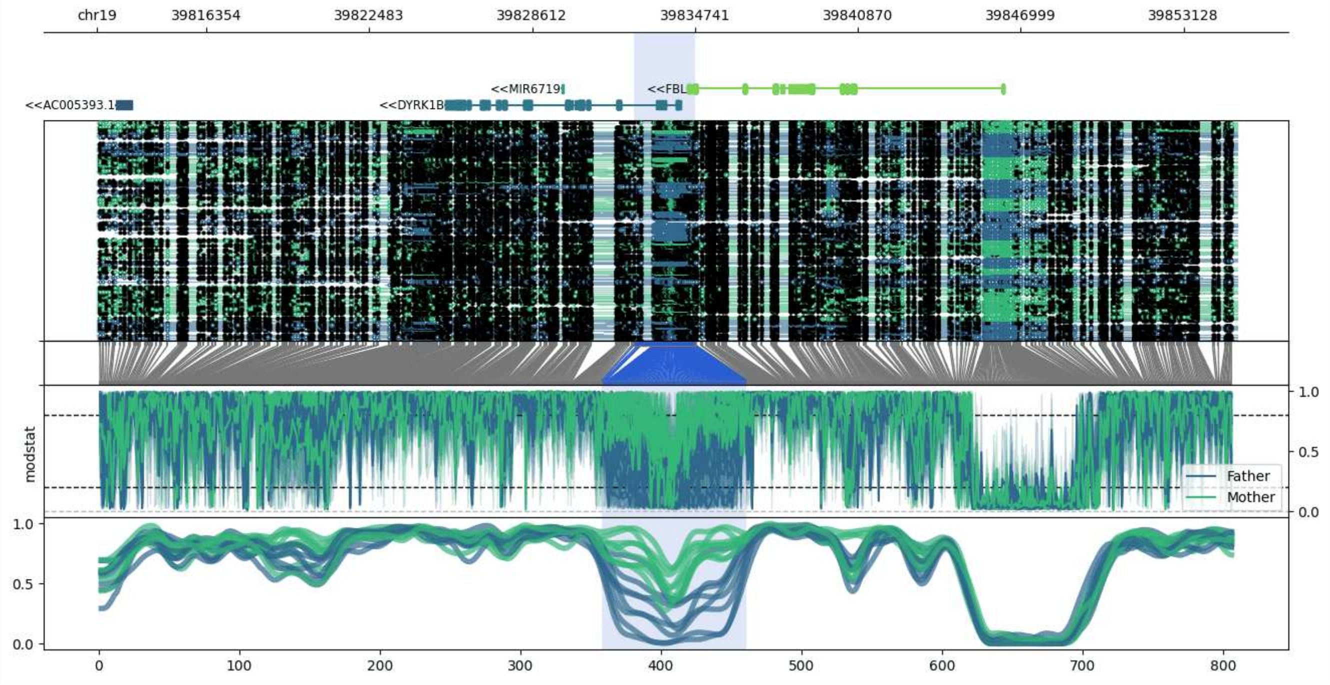


**chr19:39812245-39855149:** Region with 1 DMR containing genes: AC005393.1, DYRK1B, FBL, FCGBP, MIR6719.

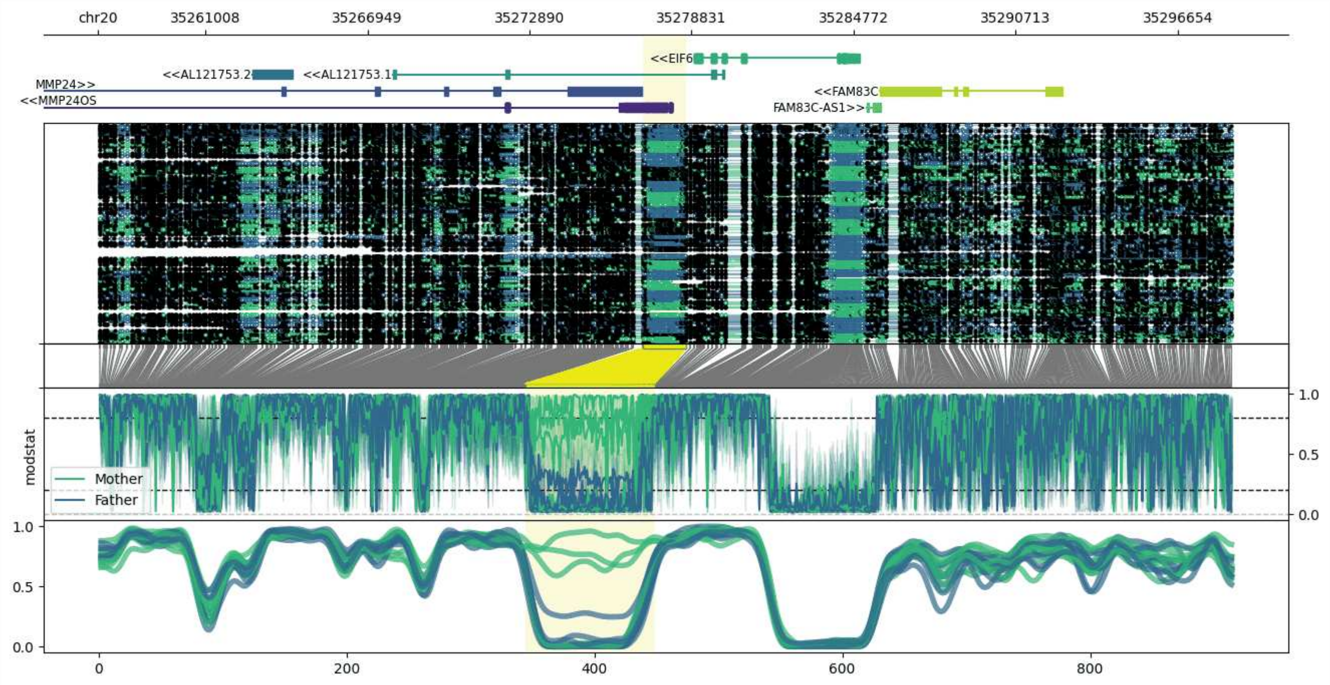


**chr20:35257056-35298648:** Region with 1 DMR containing genes: AL121753.1, AL121753.2, EIF6, FAM83C, FAM83C-AS1, MMP24, MMP24OS, UQCC1.

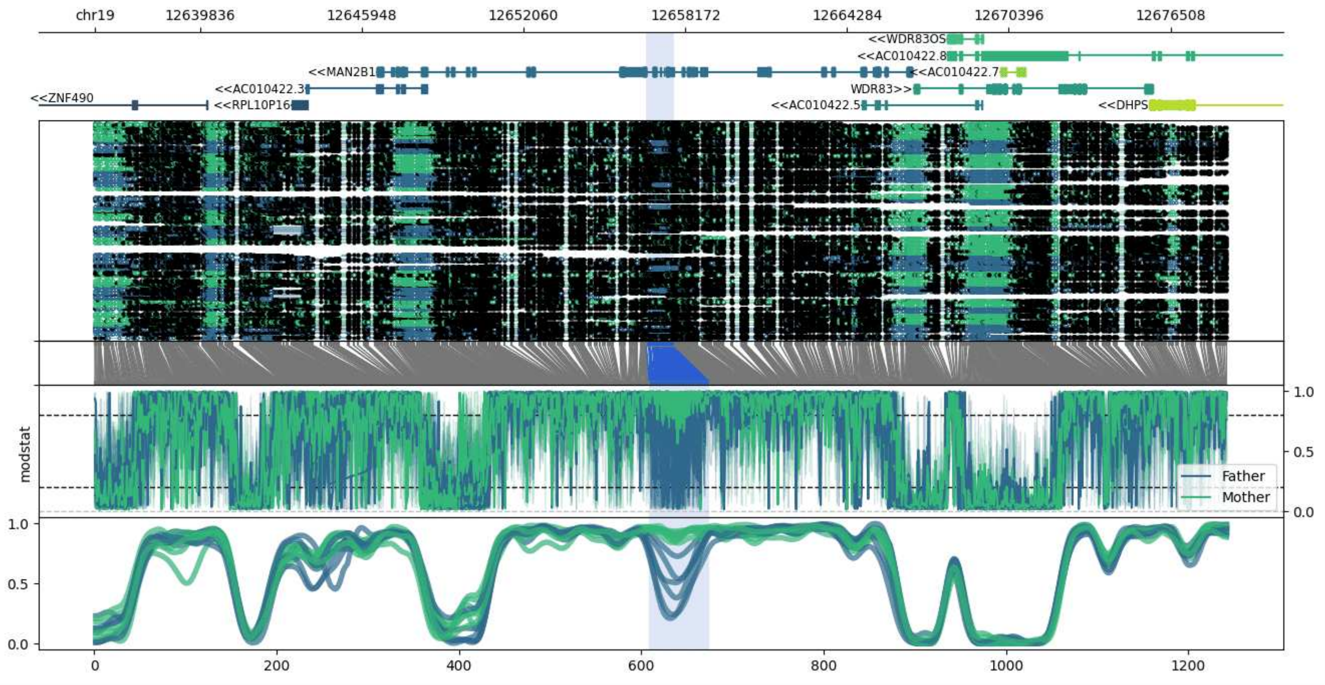


**chr19:12635847-12678633:** Region with 1 DMR containing genes: AC010422.2, AC010422.3, AC010422.5, AC010422.7, AC010422.8, DHPS, FBXW9, GNG14, MAN2B1, RPL10P16, WDR83, WDR83OS, ZNF490, ZNF791.

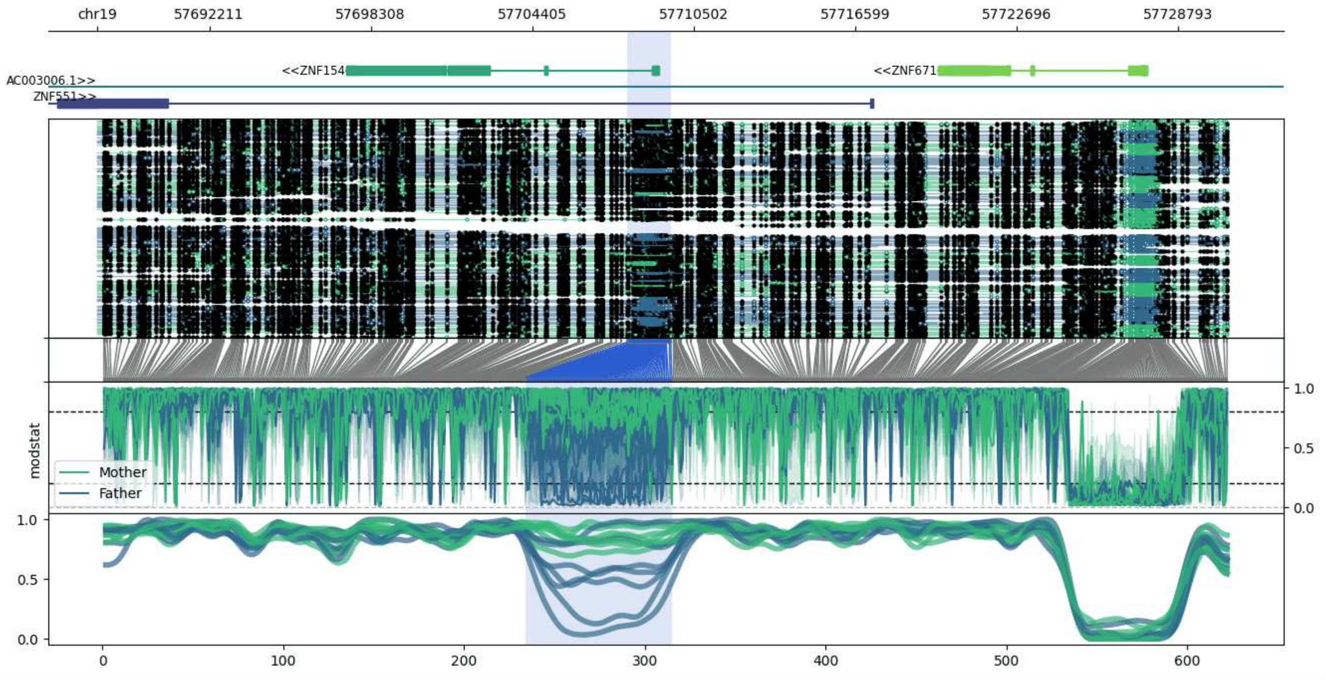


**chr19:57687993-57730676:** Region with 1 DMR containing genes: AC003006.1, ZNF154, ZNF551, ZNF671, ZSCAN4.

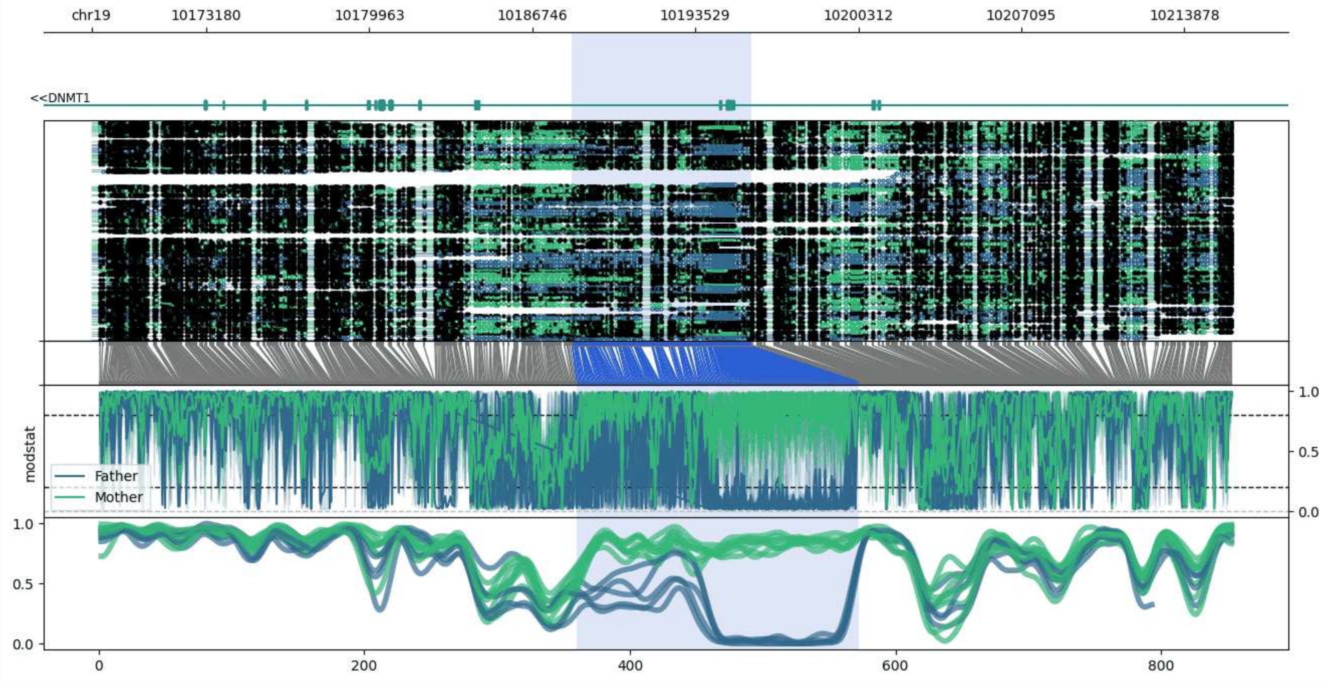


**chr19:10168395-10215876:** Region with 1 DMR containing genes: DNMT1, S1PR2.

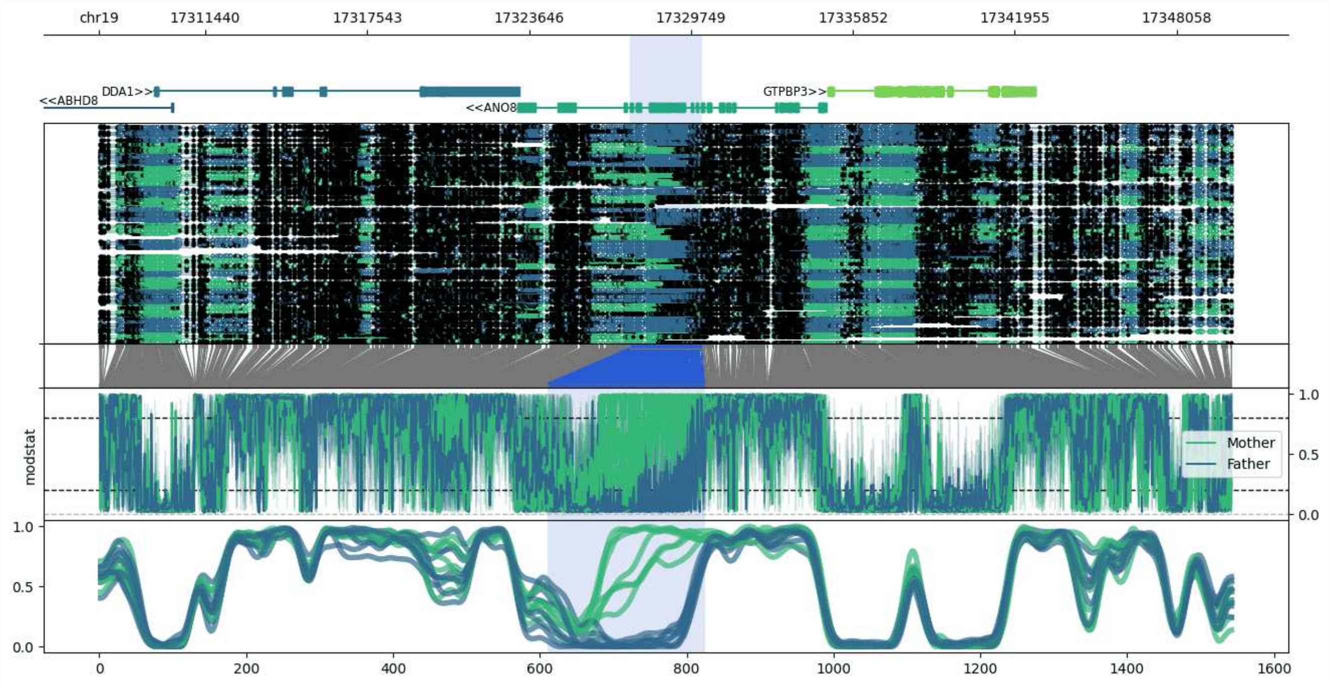


**chr19:17307440-17350163:** Region with 1 DMR containing genes: ABHD8, ANO8, DDA1, GTPBP3, MRPL34, PLVAP.

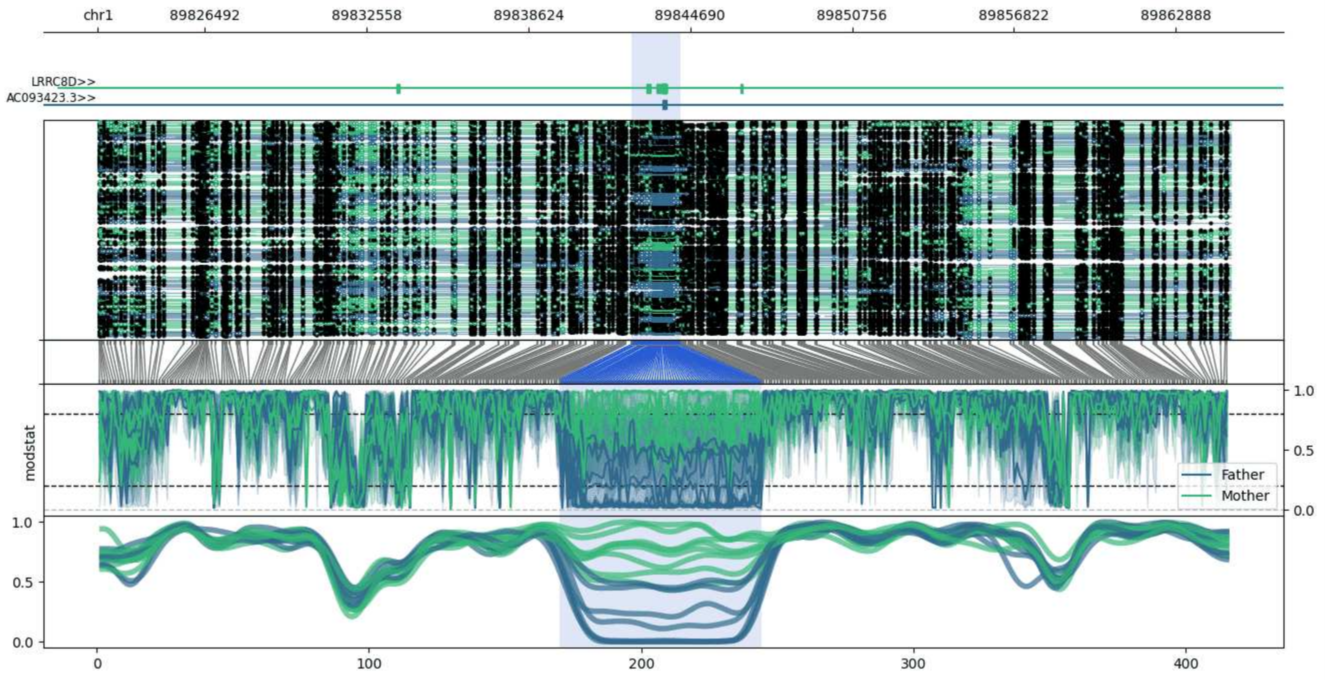


**chr1:89822481-89864944:** Region with 1 DMR containing genes: AC093423.3, AC099568.2, LRRC8D.

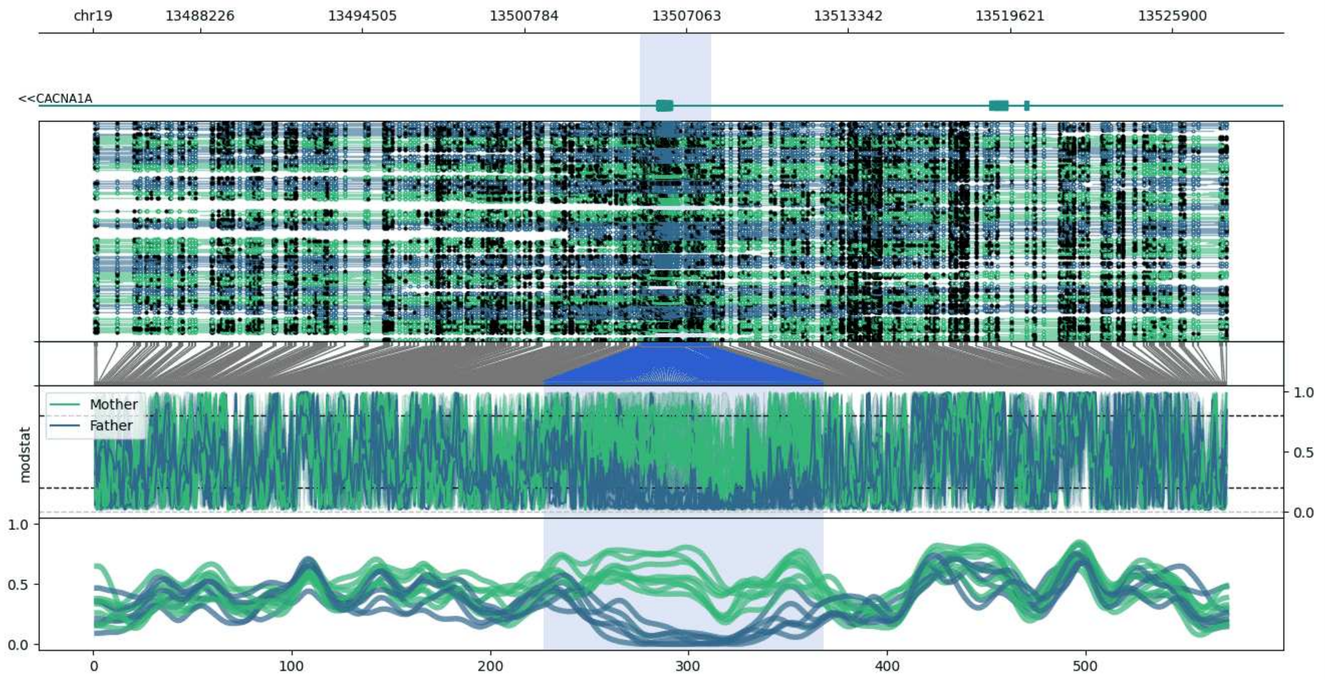


**chr19:13484068-13528023:** Region with 1 DMR containing genes: CACNA1A.

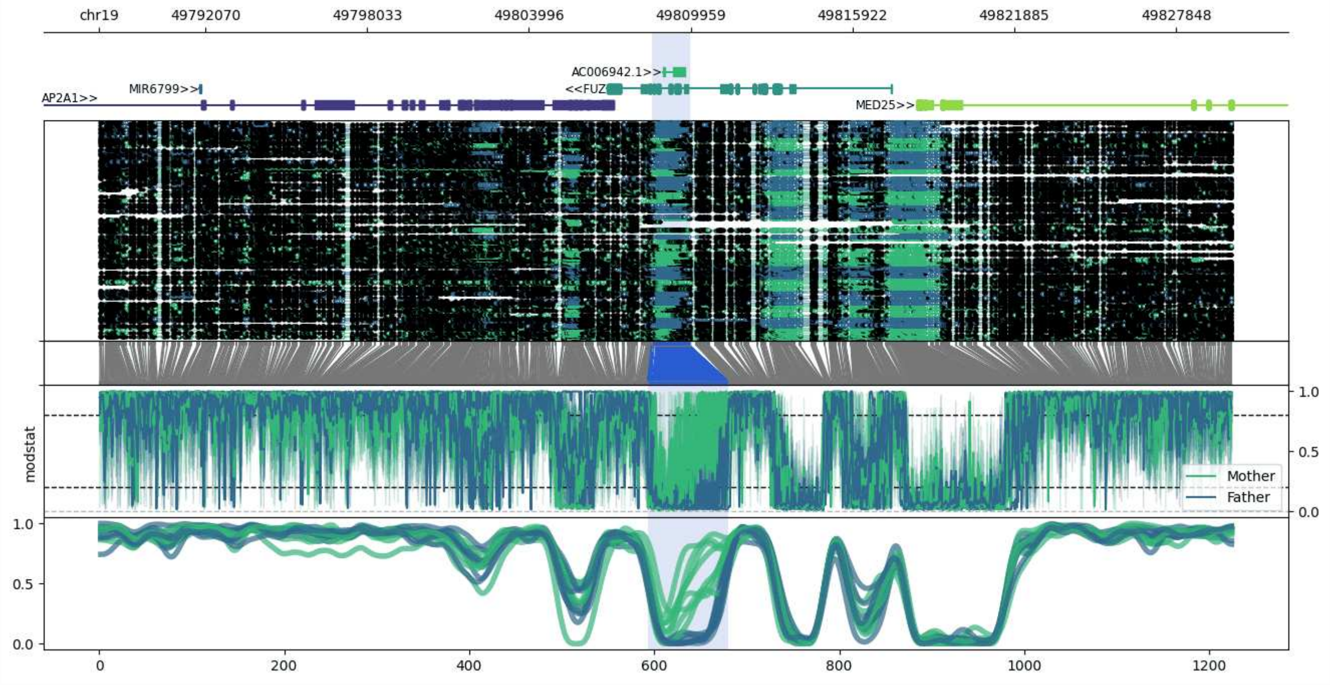


**chr19:49788148-49829892:** Region with 1 DMR containing genes: AC006942.1, AP2A1, FUZ, MED25, MIR6799, MIR6800, PTOV1-AS1.

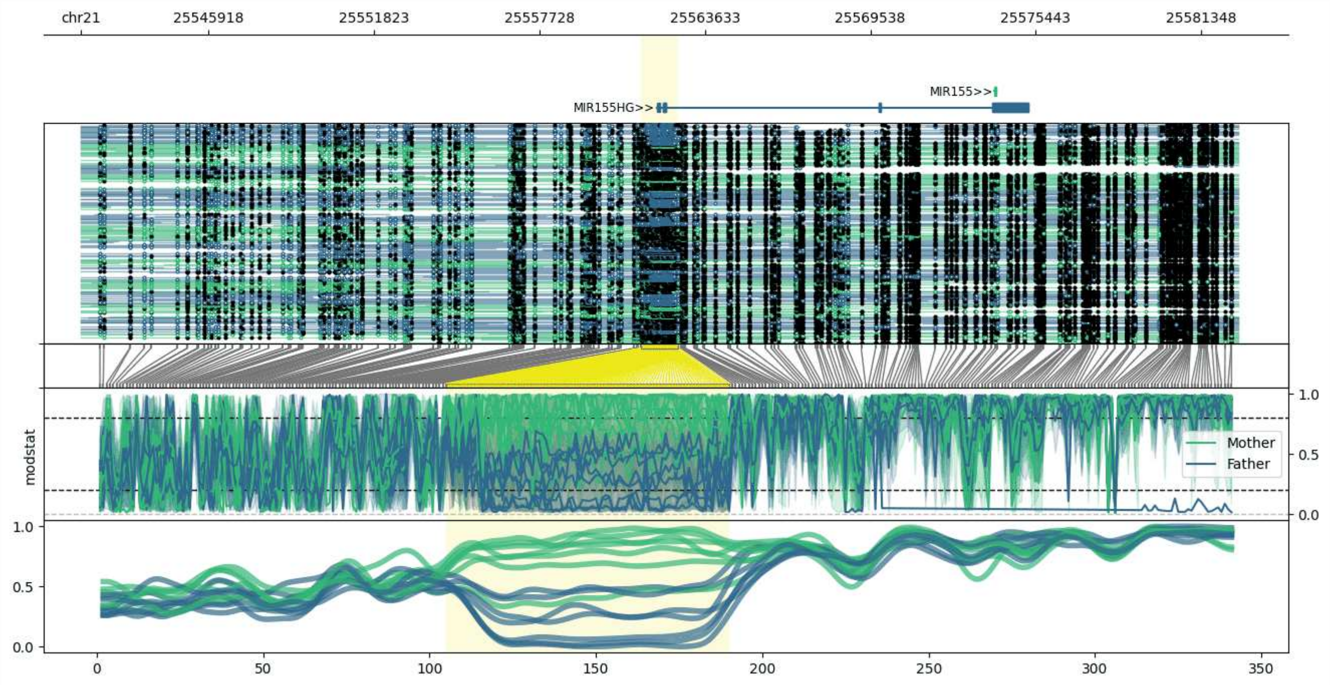


**chr21:25541349-25582688:** Region with 1 DMR containing genes: AP000223.1, MIR155, MIR155HG, MRPL39.

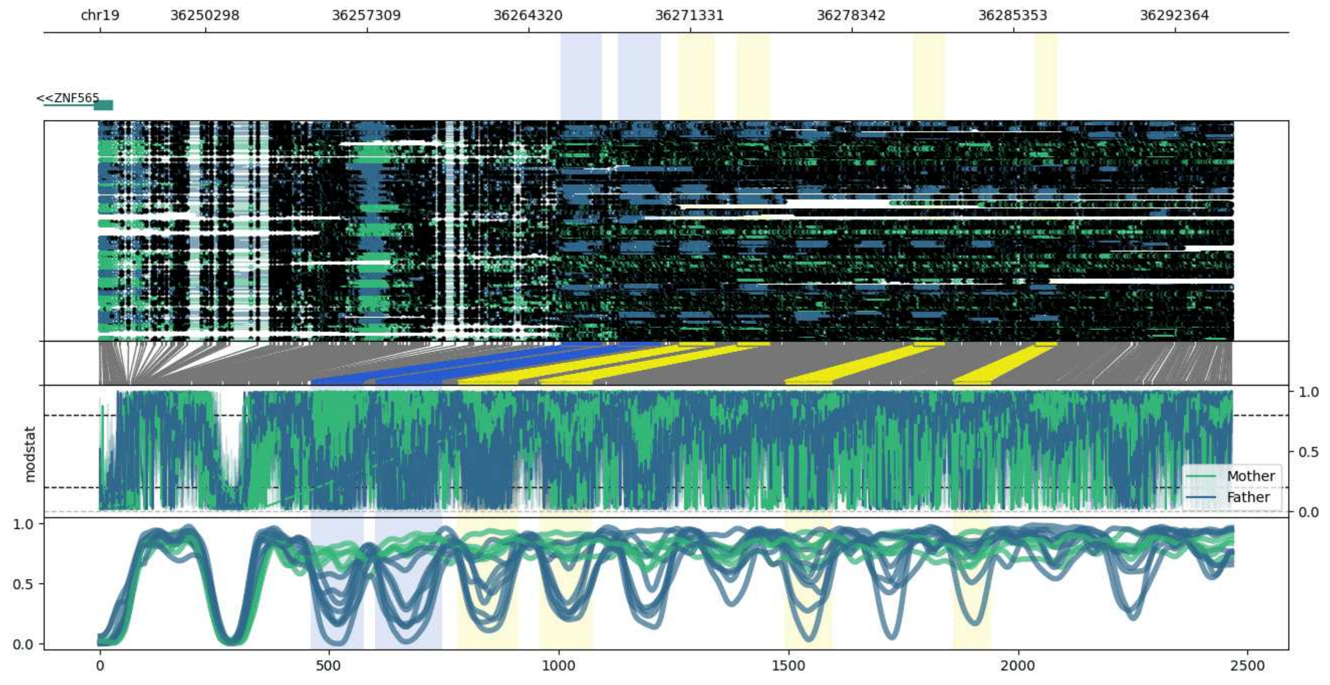


**chr19:36245738-36294817:** Region with 6 DMRs containing genes: AC092296.2, ZNF146, ZNF565.

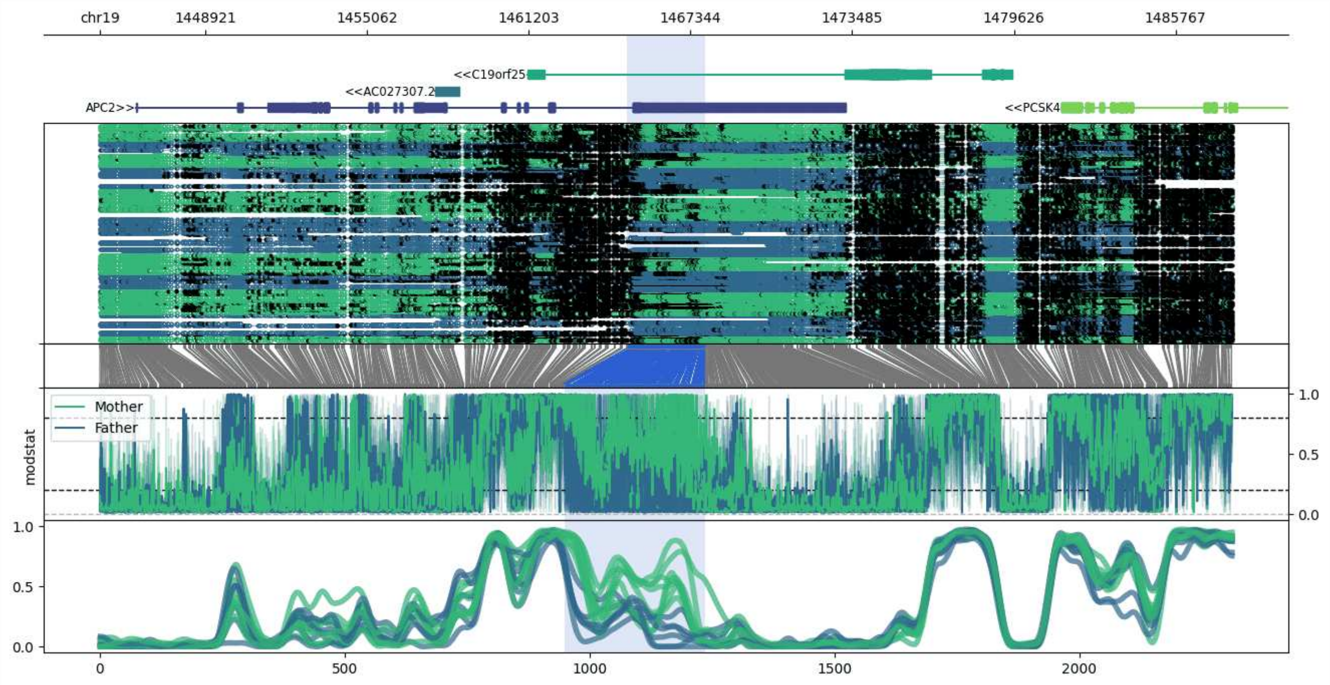


**chr19:1444927-1487915:** Region with 1 DMR containing genes: AC027307.2, AC027307.3, APC2, C19orf25, DAZAP1, PCSK4, REEP6, RPS15.

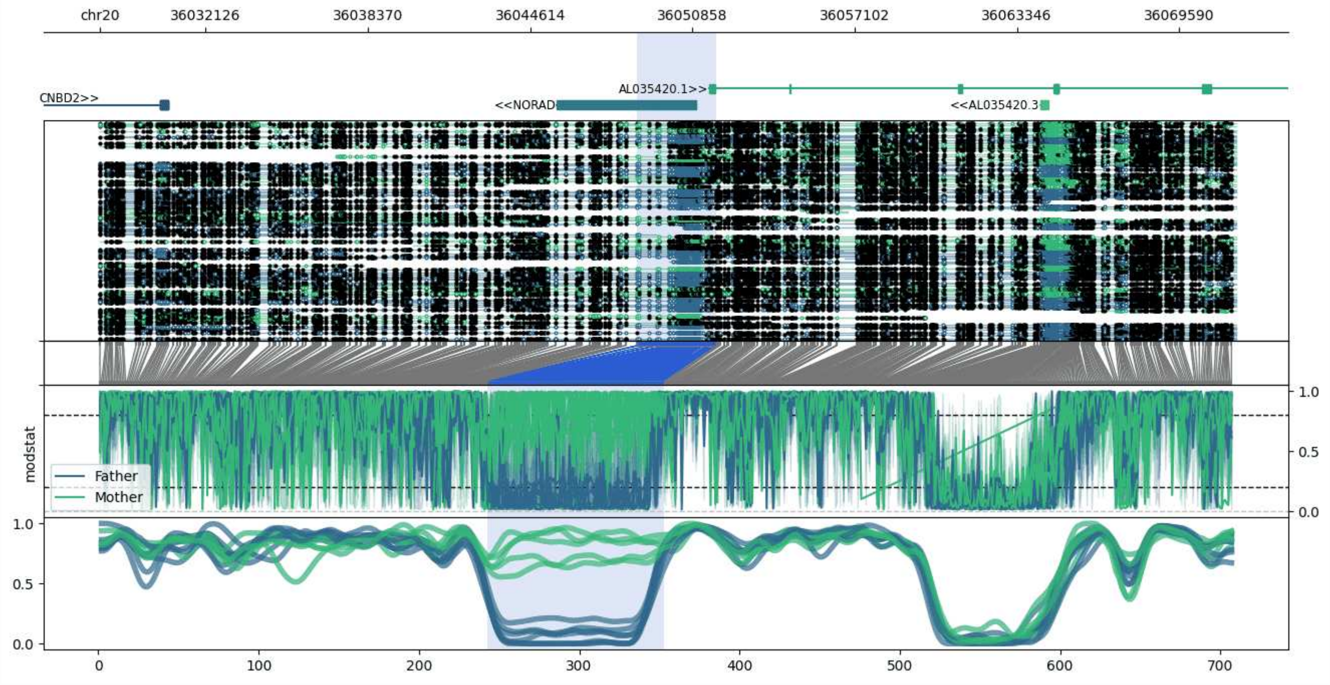


**chr20:36028053-36071764:** Region with 1 DMR containing genes: AL035420.1, AL035420.3, CNBD2, HMGB3P2, NORAD.

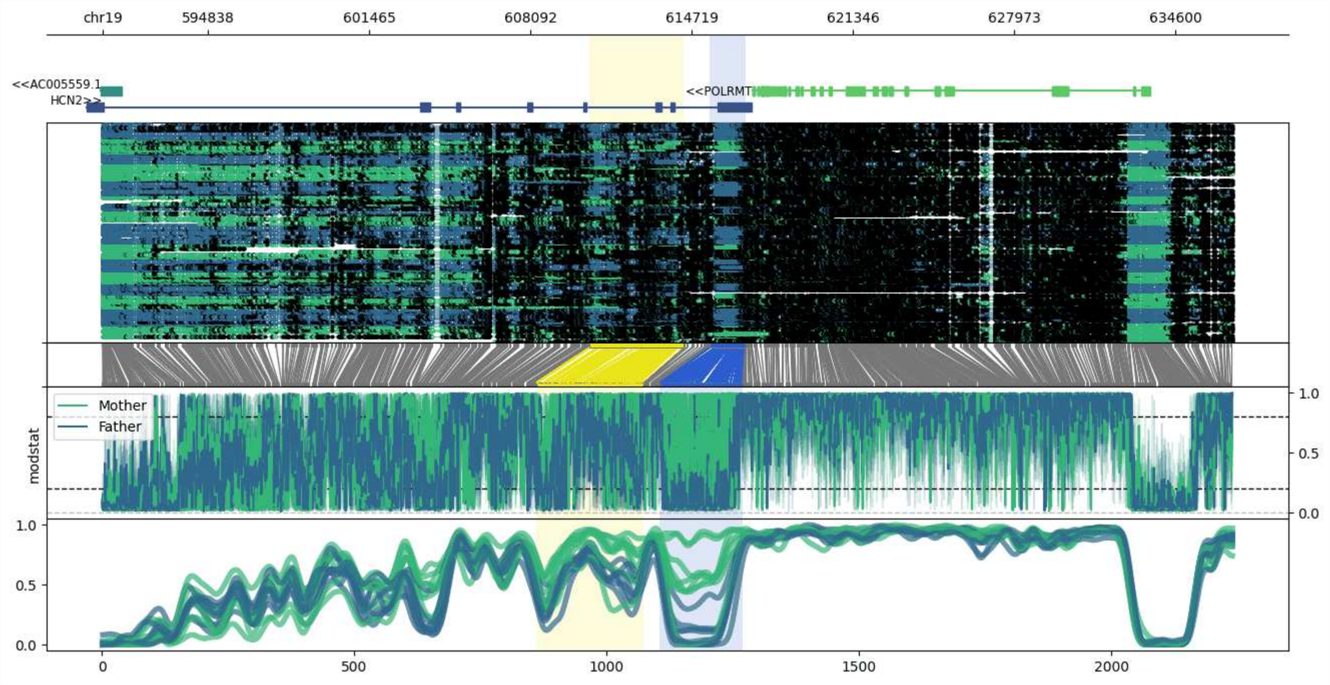


**chr19:590525-636919:** Region with 2 DMRs containing genes: AC004449.1, AC005559.1, BSG, FGF22, HCN2, POLRMT.

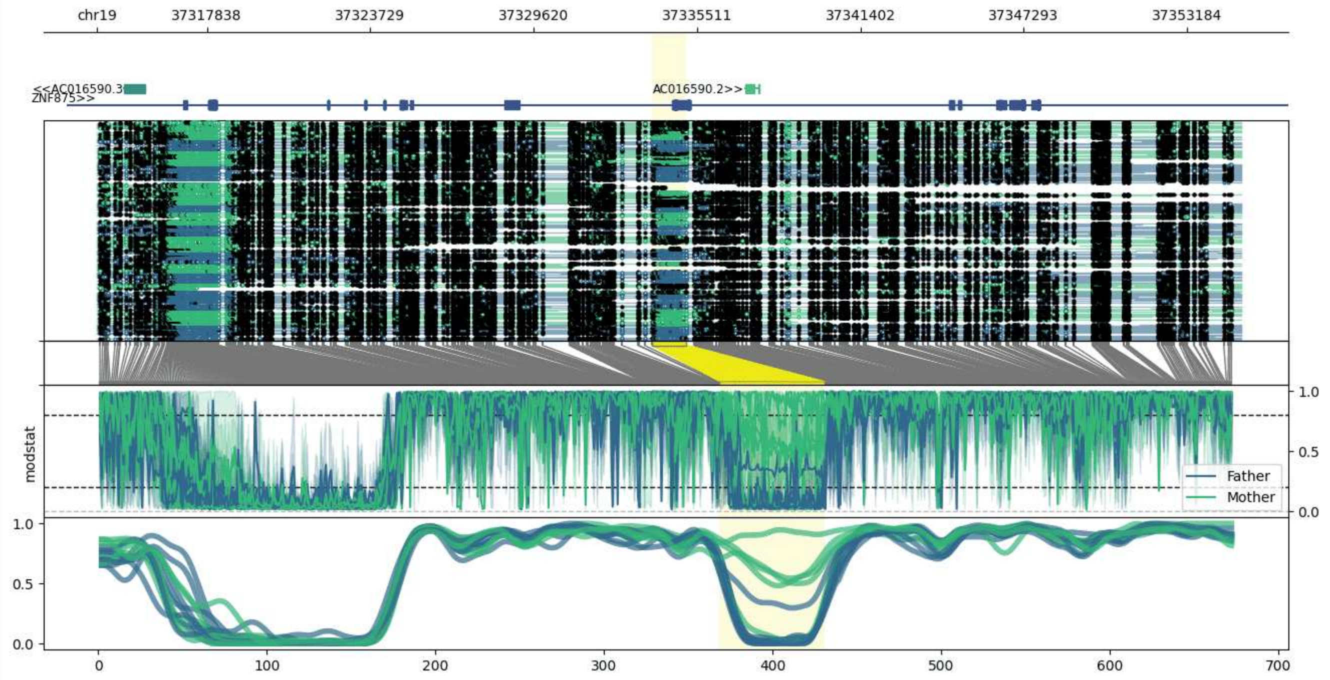


**chr19:37313878-37355115:** Region with 1 DMR containing genes: AC016590.1, AC016590.2, AC016590.3, AC016590.4, ZNF875.

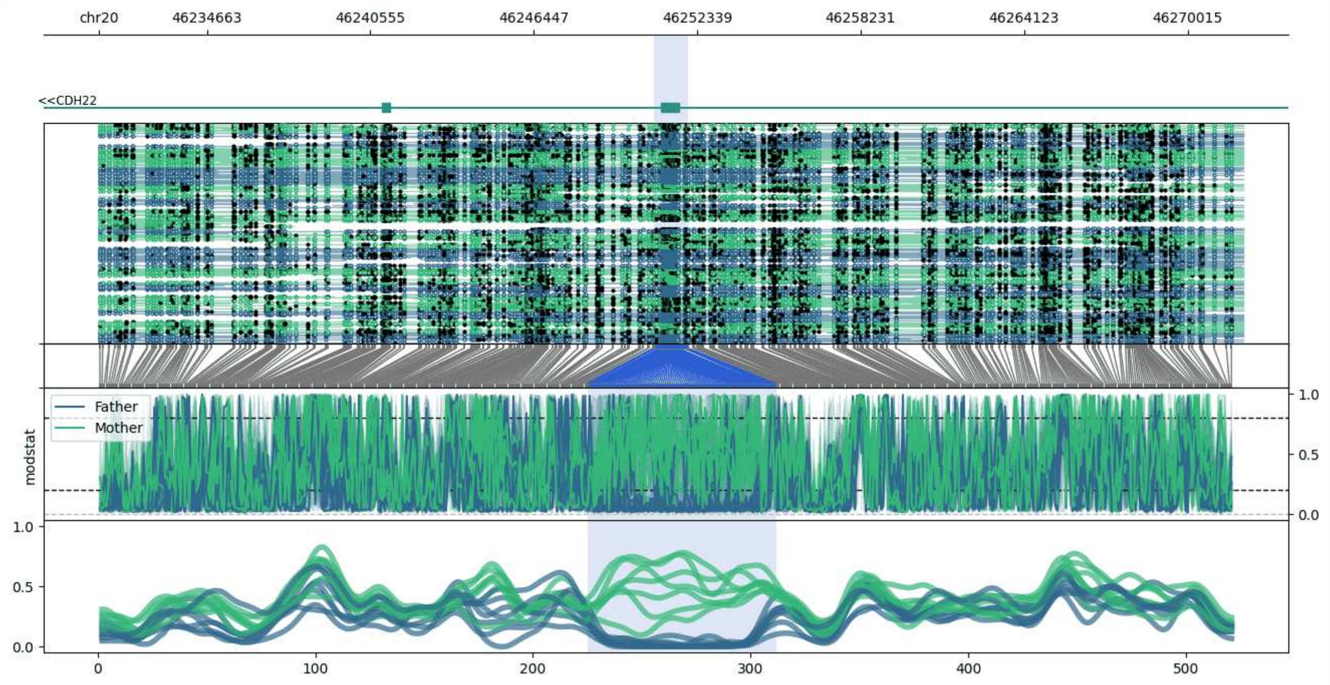


**chr20:46230763-46272008:** Region with 1 DMR containing genes: CDH22.

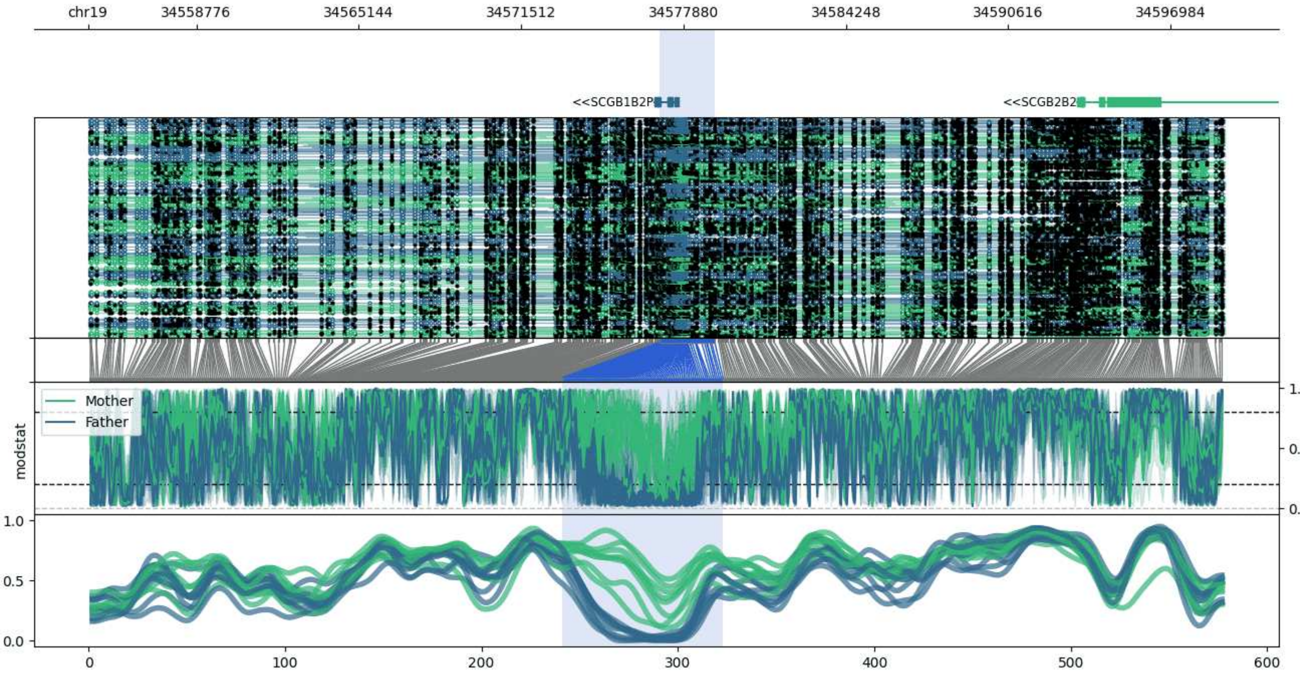


**chr19:34554527-34599105:** Region with 1 DMR containing genes: SCGB1B2P, SCGB2B2, ZNF807.

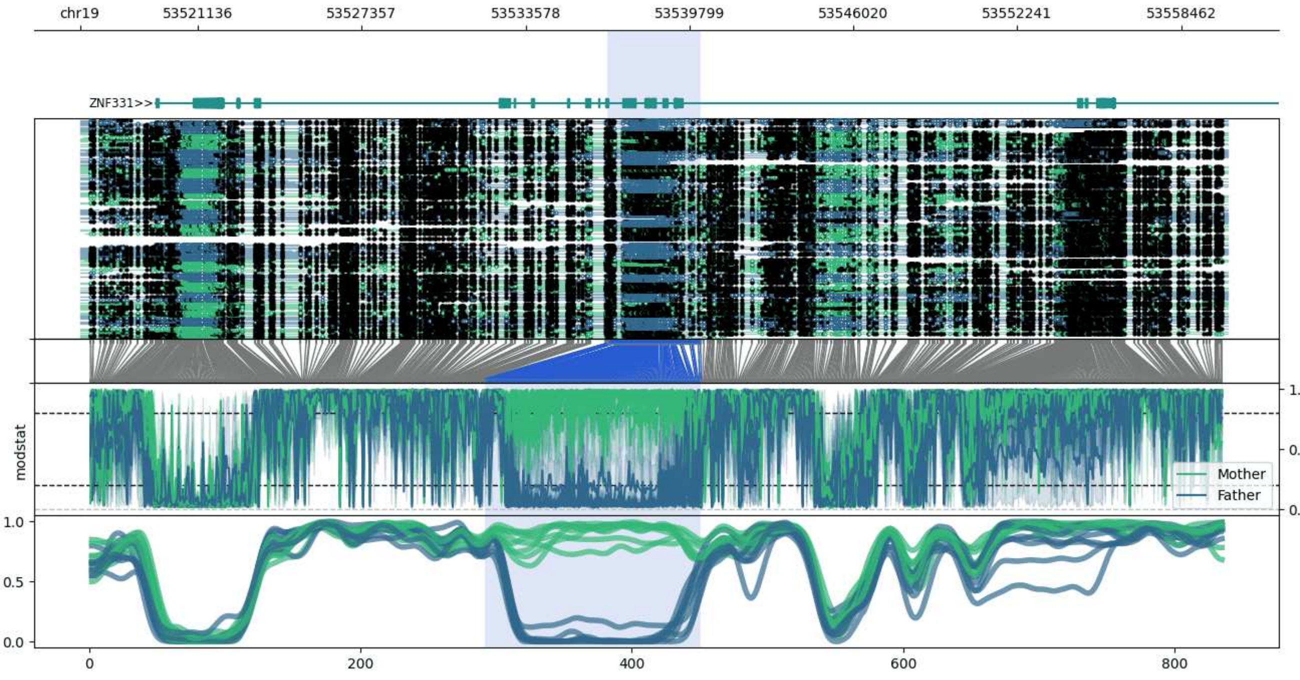


**chr19:53516664-53560216:** Region with 1 DMR containing genes: AC011487.1, AC011487.4, ZNF331.

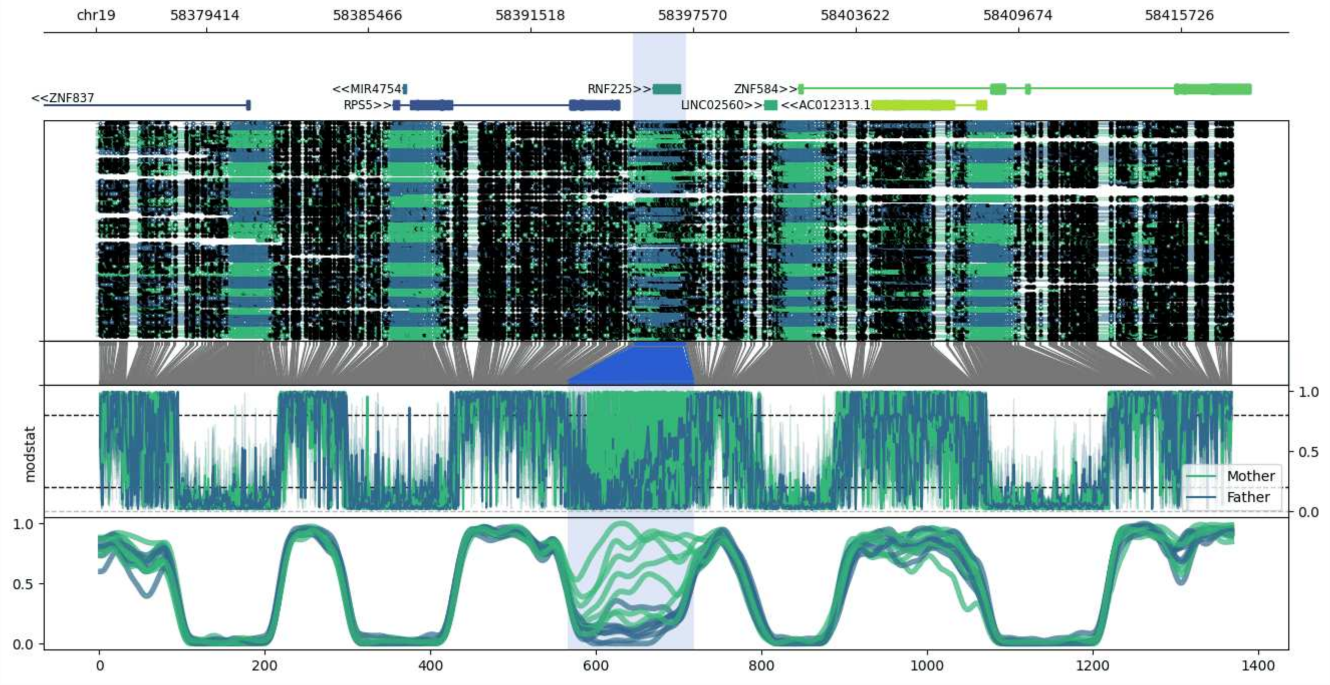


**chr19:58375320-58417690:** Region with 1 DMR containing genes: AC012313.1, AC012313.4, AC012313.6, LINC02560, MIR4754, RNF225, RPS5, ZNF584, ZNF837.

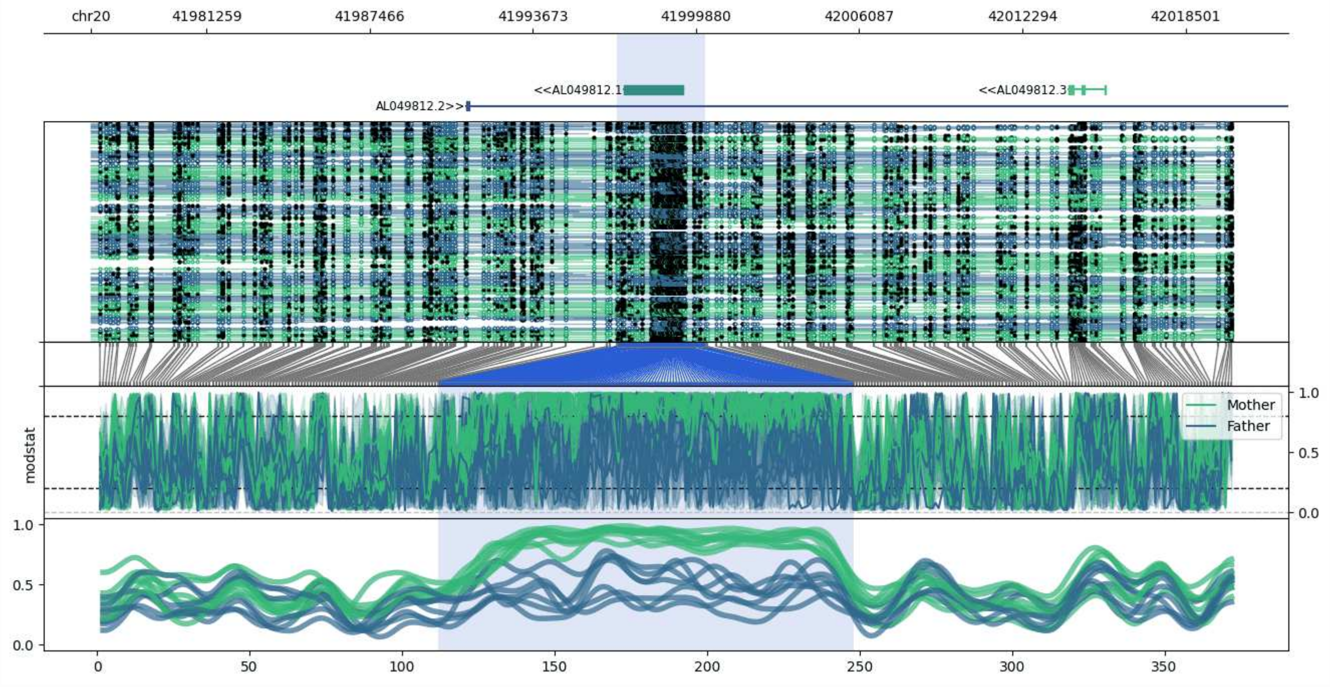


**chr20:41976856-42020311:** Region with 1 DMR containing genes: AL049812.1, AL049812.2, AL049812.3.

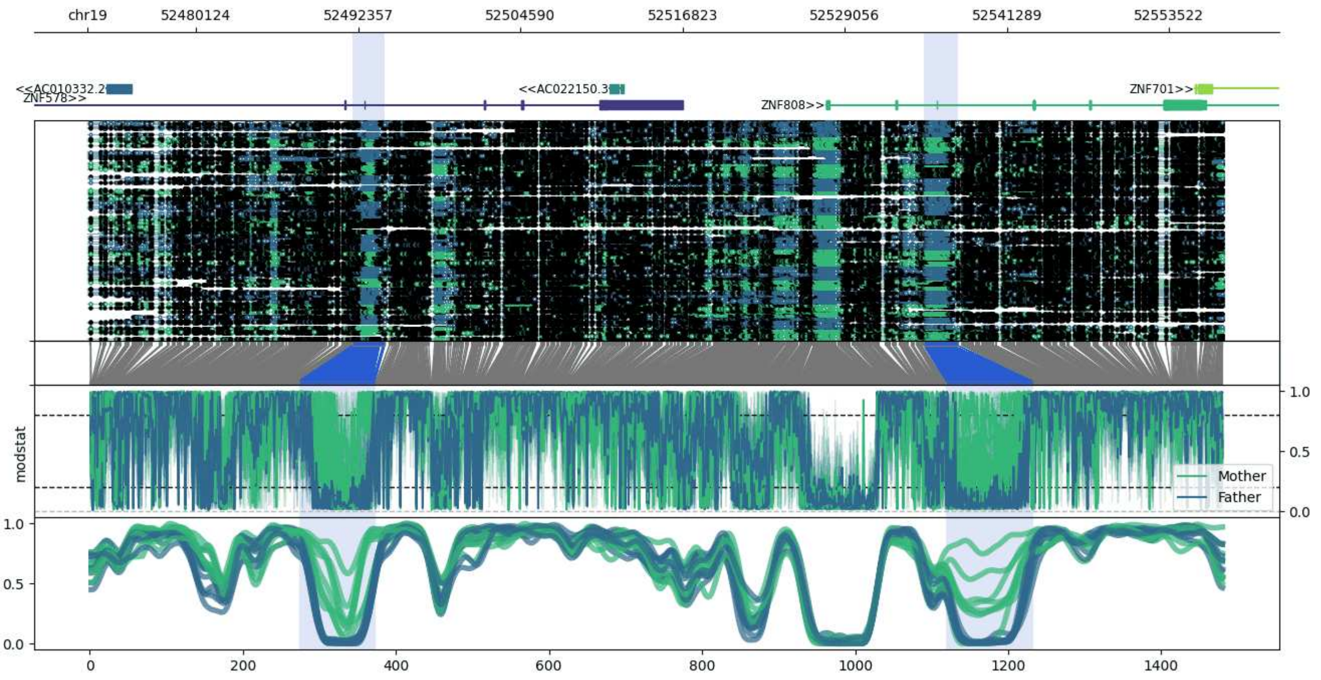


**chr19:52471955-52557586:** Region with 2 DMRs containing genes: AC010332.2, AC022150.3, RPL39P34, ZNF578, ZNF701, ZNF808.

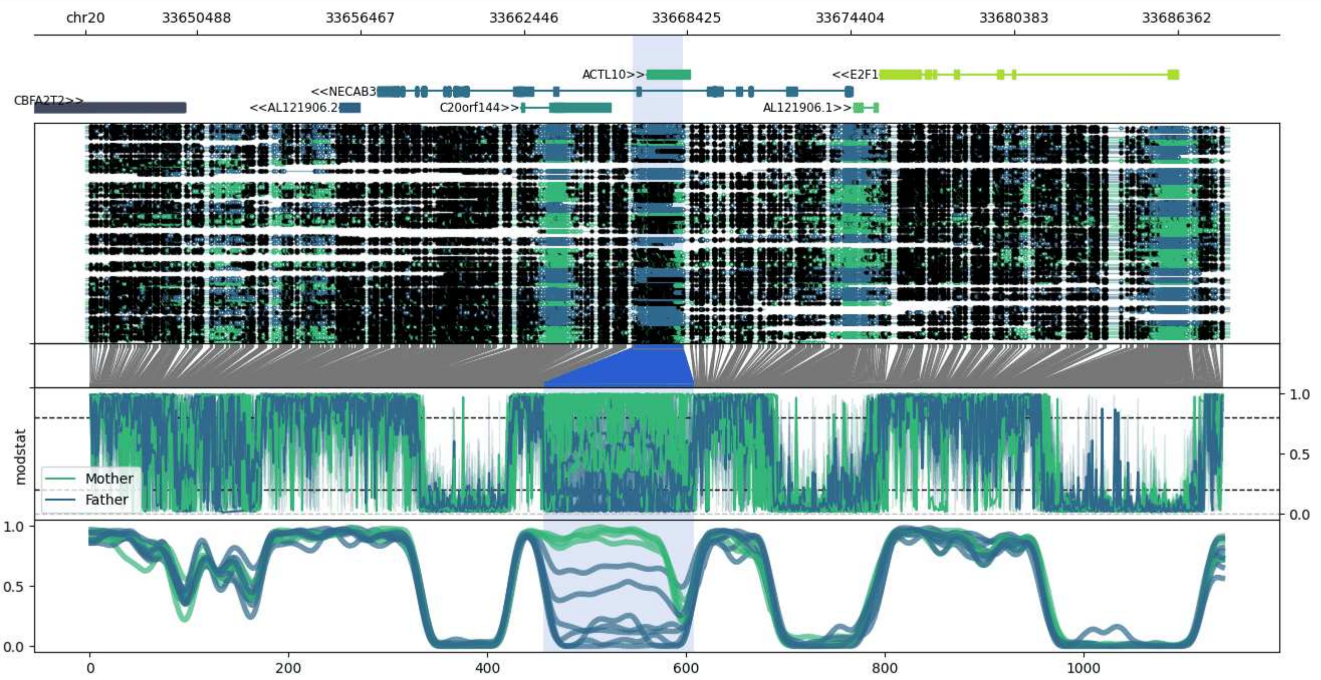


**chr20:33646409-33688264:** Region with 1 DMR containing genes: ACTL10, AL121906.1, AL121906.2, C20orf144, CBFA2T2, E2F1, NECAB3.

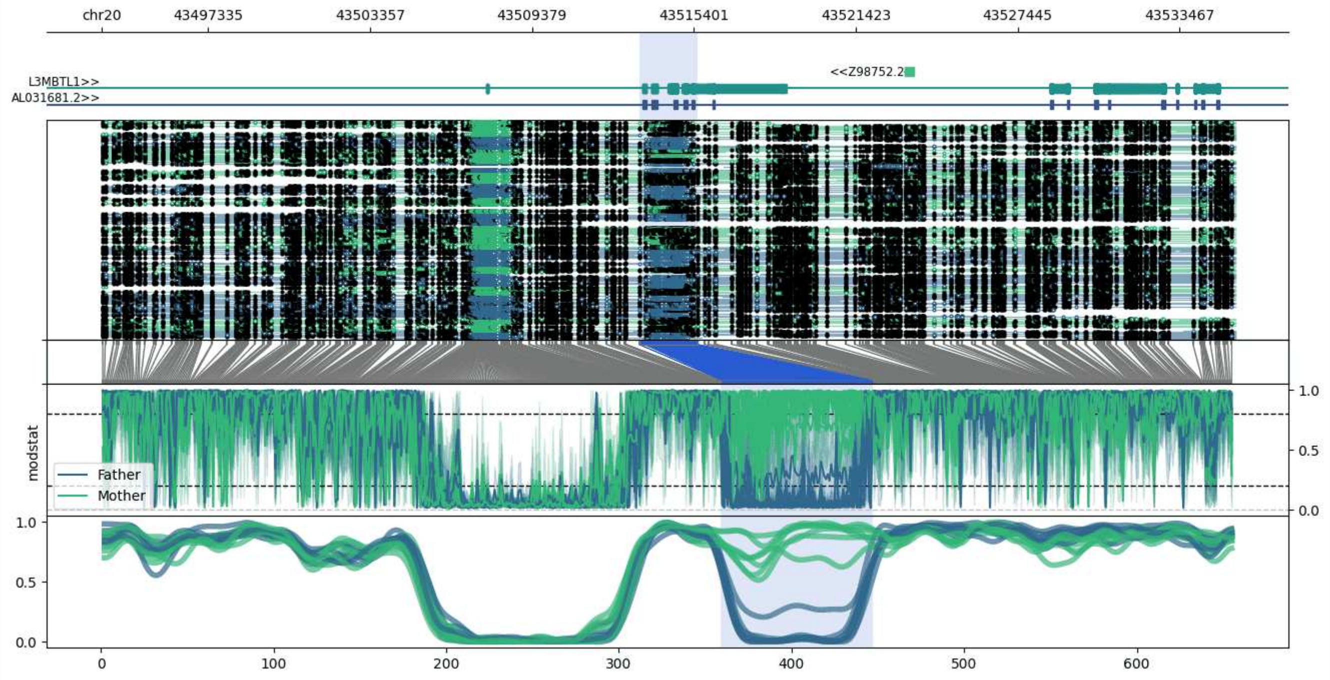


**chr20:43493364-43535520:** Region with 1 DMR containing genes: AL031681.2, L3MBTL1, Z98752.2, Z98752.3.

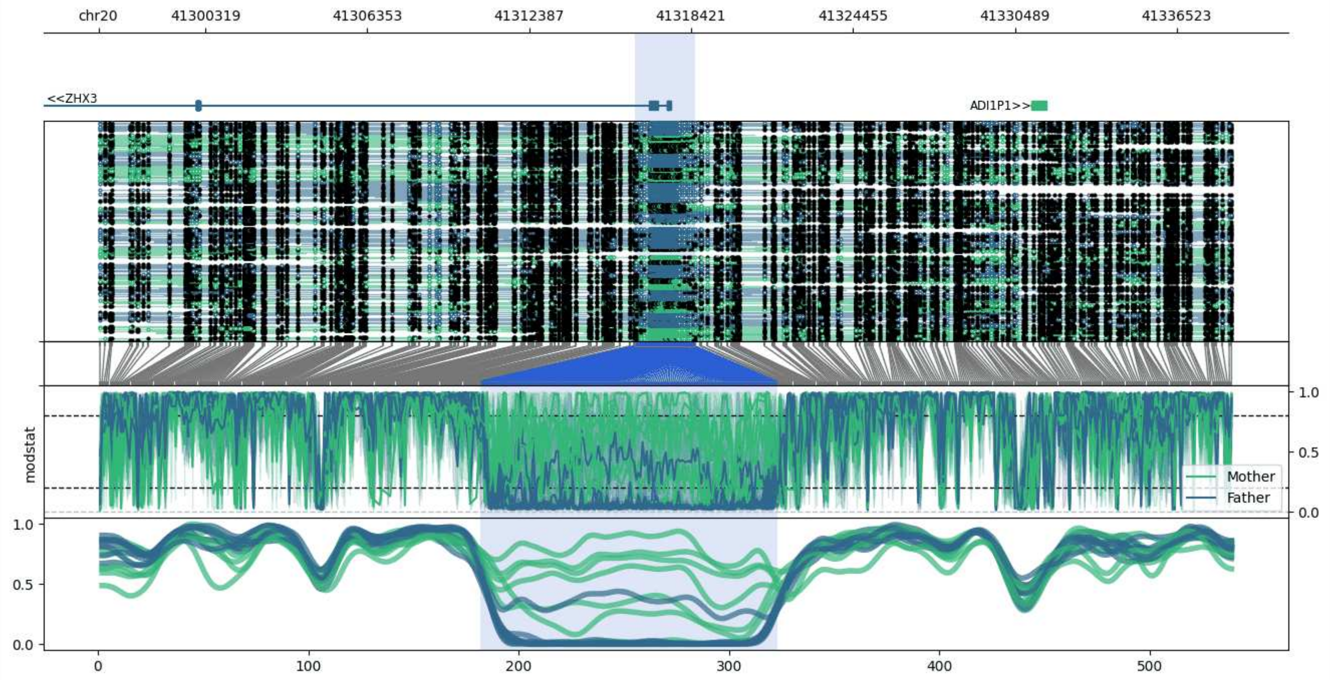


**chr20:41296335-41338574:** Region with 1 DMR containing genes: ADI1P1, LPIN3, RN7SL615P, ZHX3.

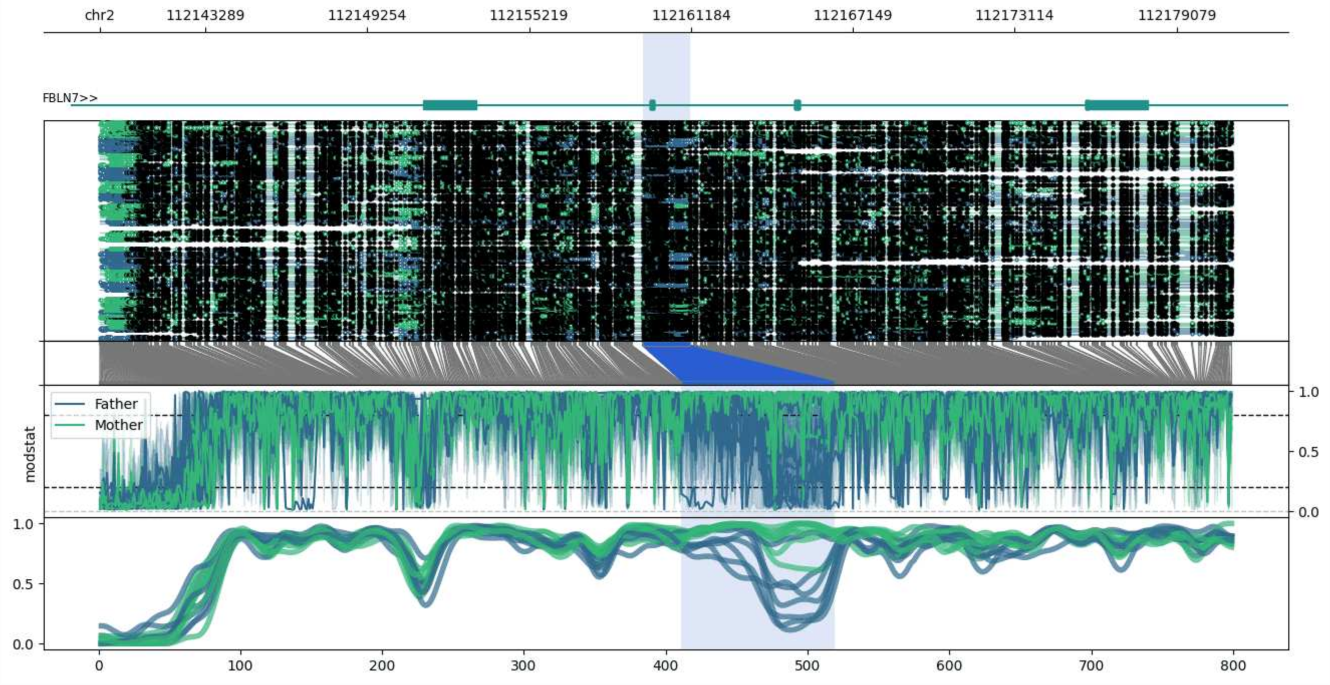


**chr2:112139397-112181153:** Region with 1 DMR containing genes: AC092645.1, FBLN7.

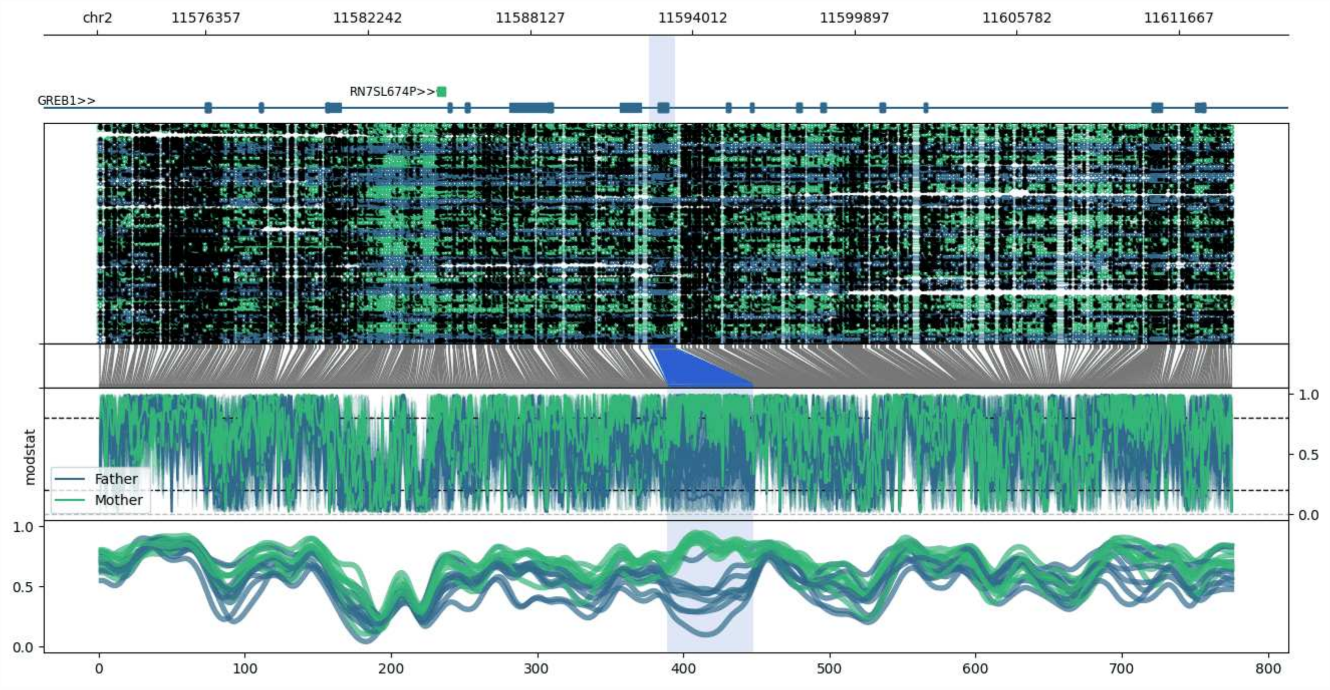


**chr2:11572426-11613626:** Region with 1 DMR containing genes: GREB1, RN7SL674P.

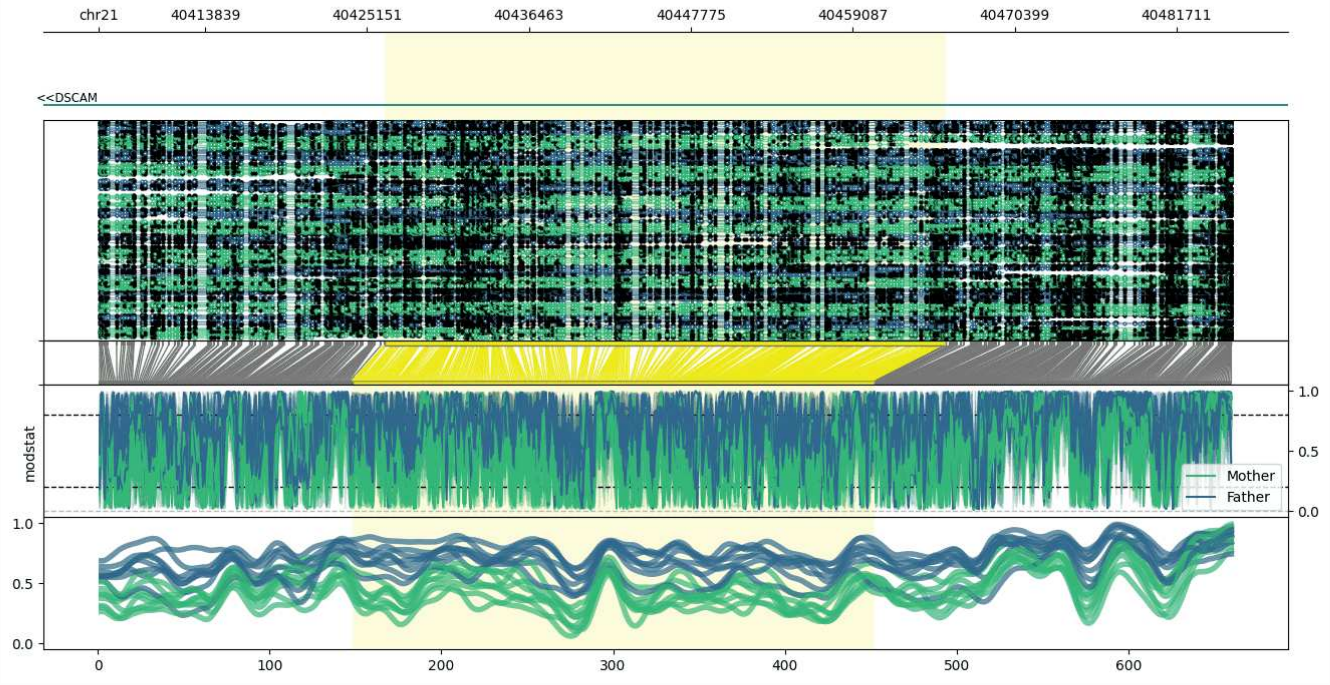


**chr21:40406380-40485570:** Region with 1 DMR containing genes: DSCAM.

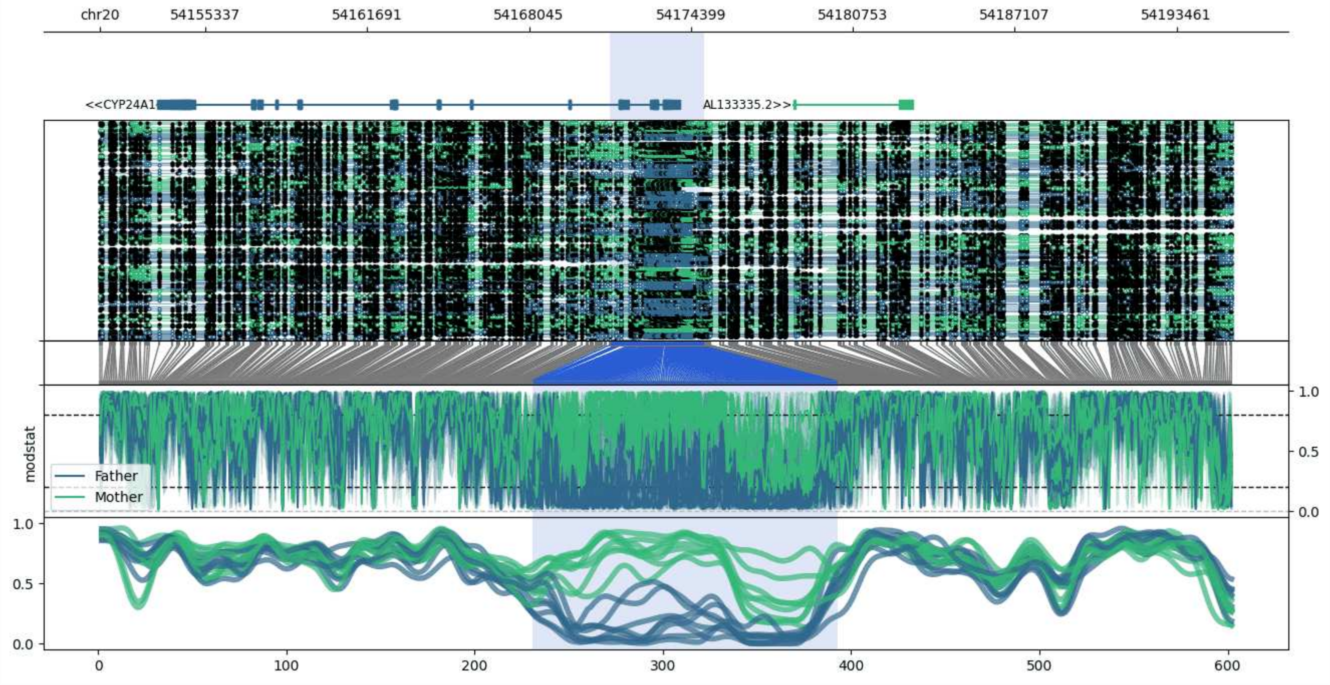


**chr20:54151201-54195685:** Region with 1 DMR containing genes: AL133335.2, CYP24A1.

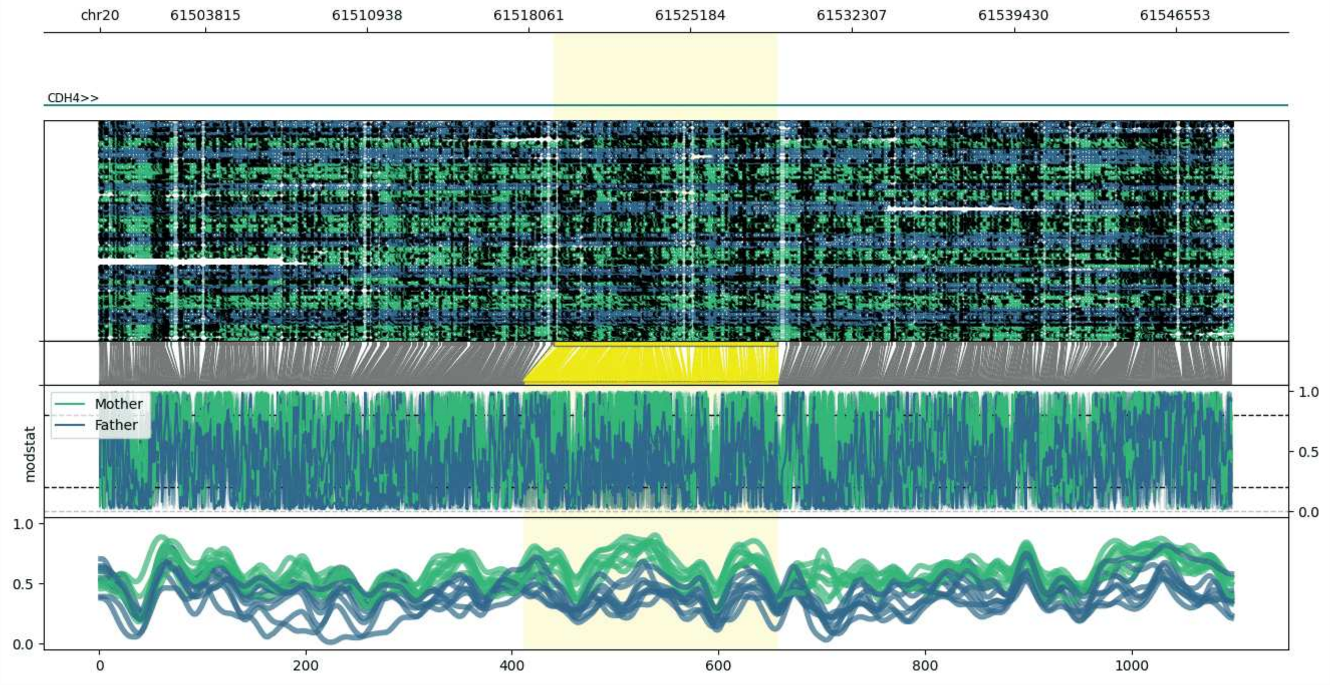


**chr20:61499171-61549032:** Region with 1 DMR containing genes: CDH4.

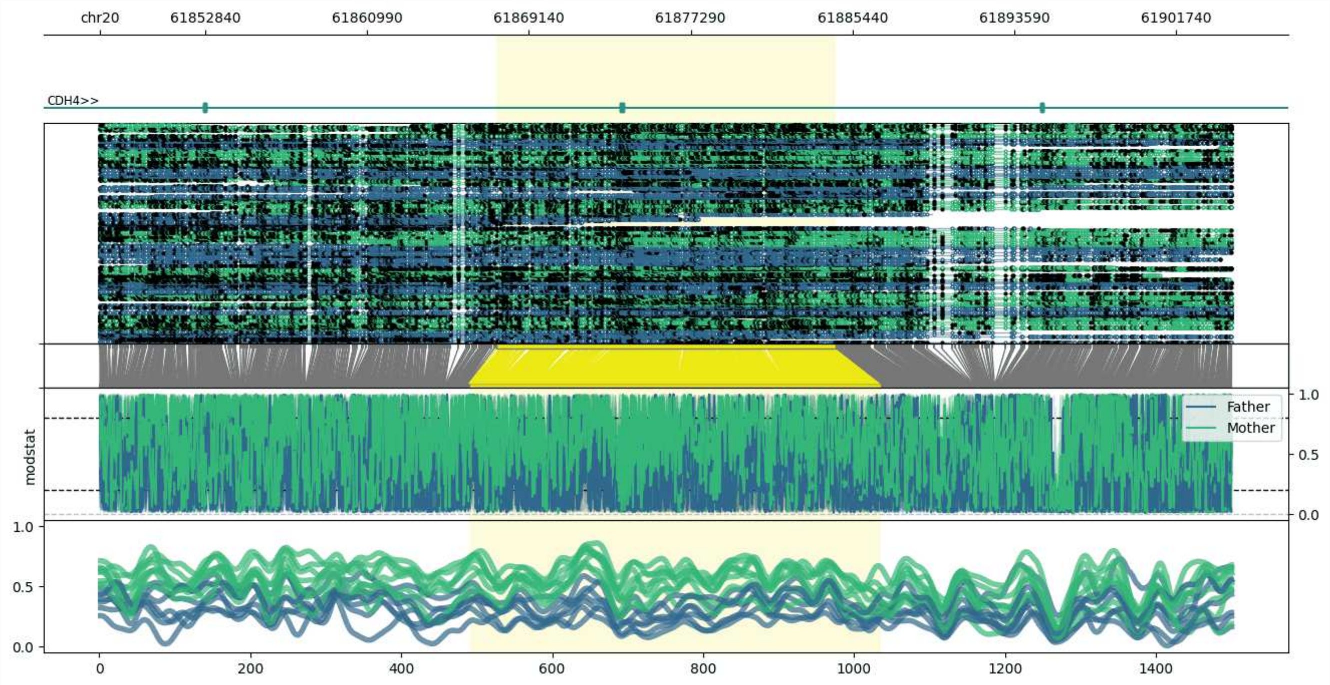


**chr20:61847527-61904579:** Region with 1 DMR containing genes: CDH4.

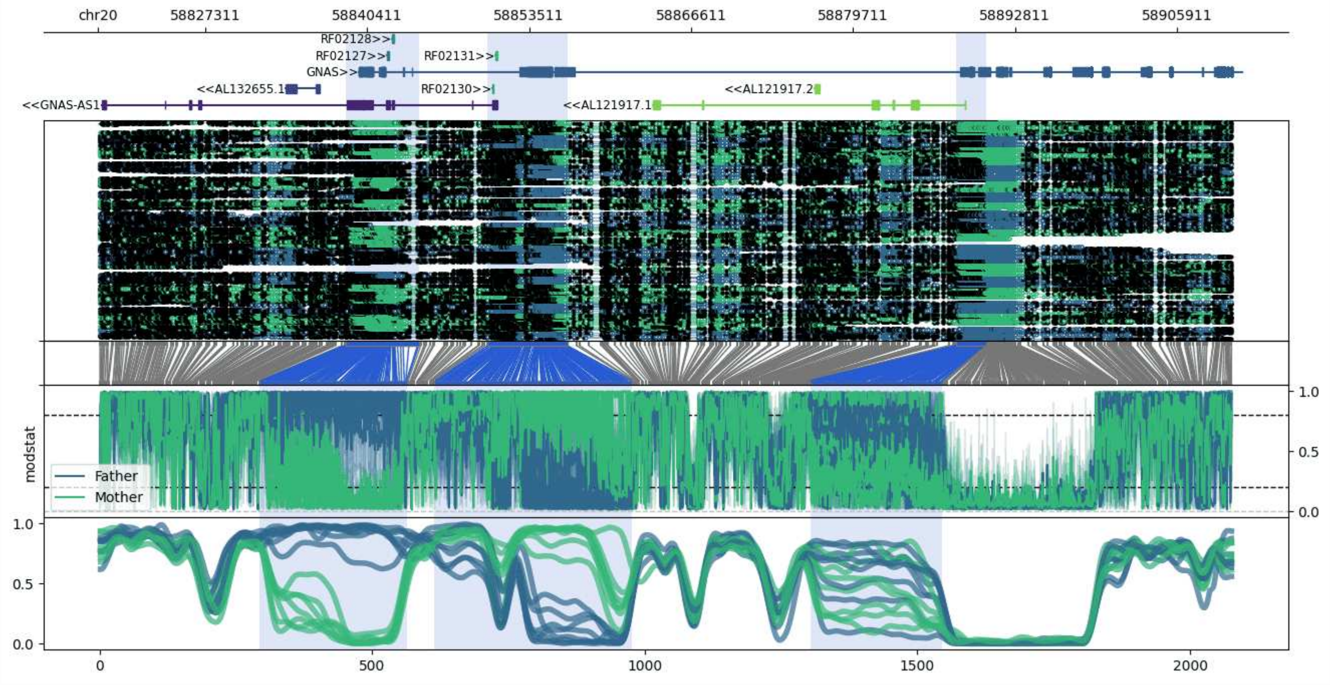


**chr20:58818639-58910343:** Region with 3 DMRs containing genes: AL109840.2, AL121917.1, AL121917.2, AL132655.1, AL132655.2, GNAS, GNAS-AS1, MIR296, MIR298, RF02127, RF02128, RF02130, RF02131.

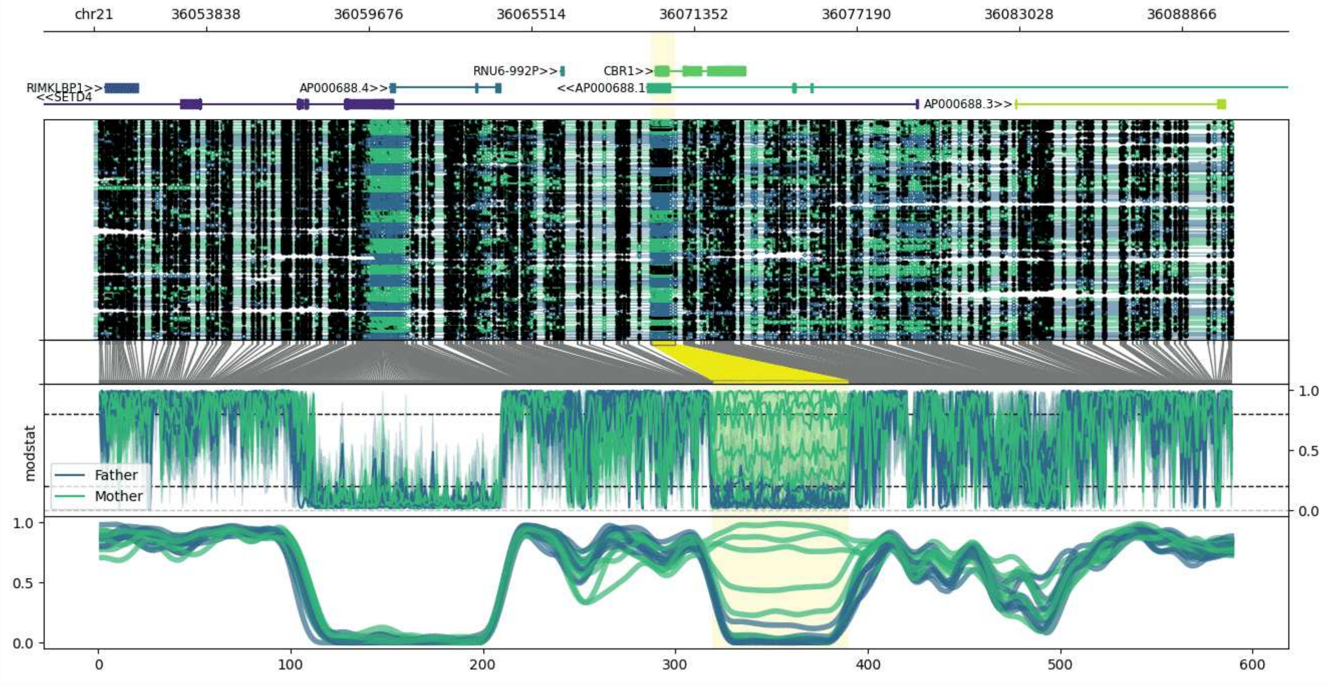


**chr21:36049805-36090677:** Region with 1 DMR containing genes: AP000688.1, AP000688.3, AP000688.4, CBR1, RIMKLBP1, RNU6-992P, SETD4.

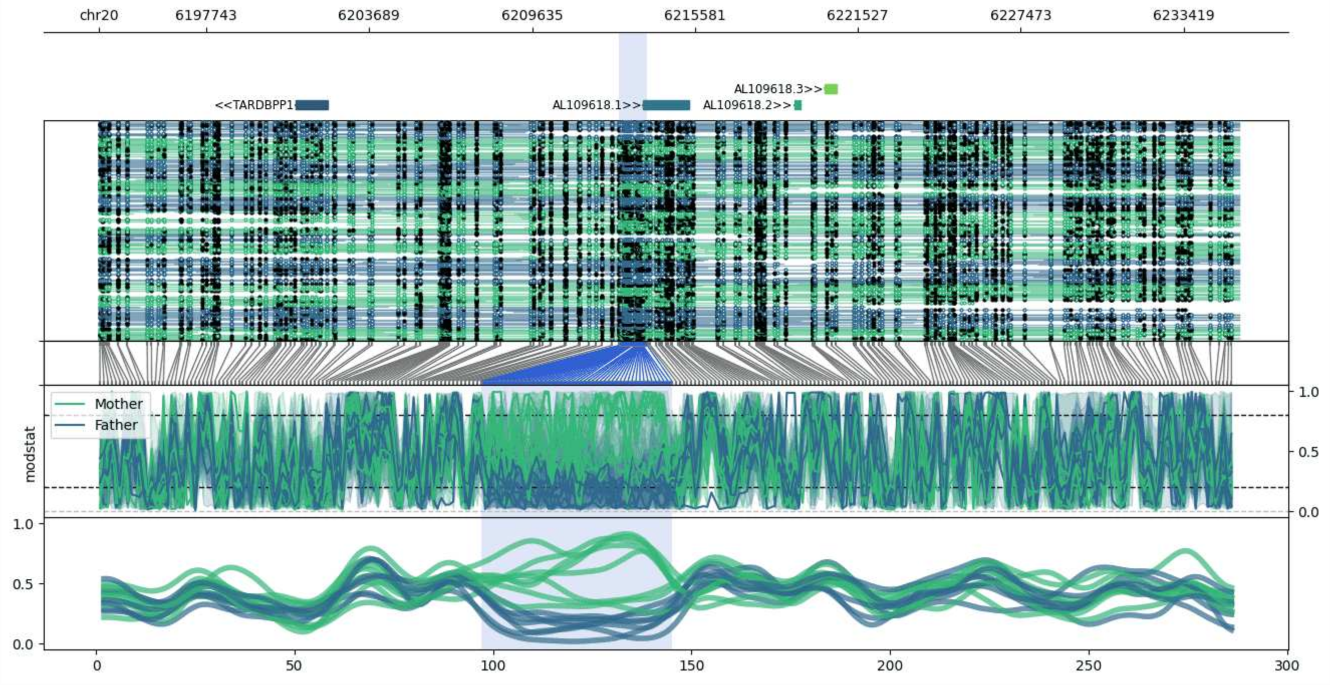


**chr20:6193839-6235461:** Region with 1 DMR containing genes: AL109618.1, AL109618.2, AL109618.3, TARDBPP1.

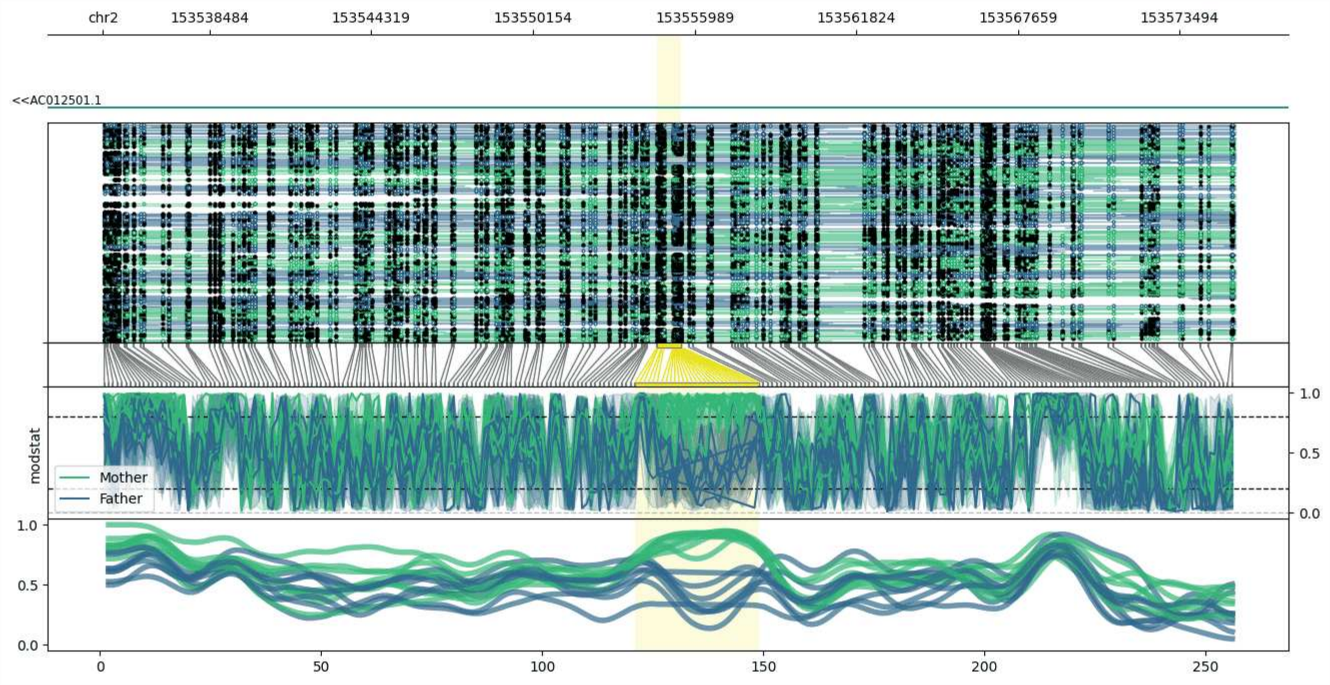


**chr2:153534642-153575488:** Region with 1 DMR containing genes: AC012501.1.

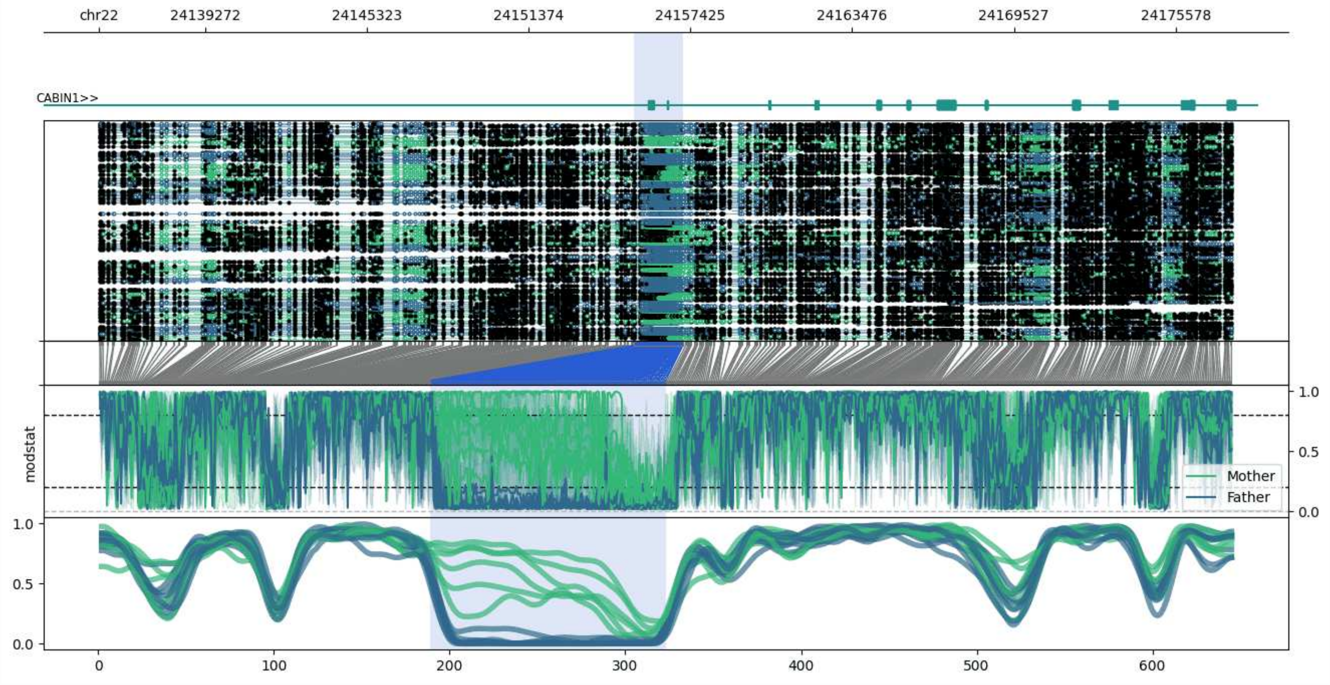


**chr22:24135312-24177672:** Region with 1 DMR containing genes: CABIN1, SUSD2.

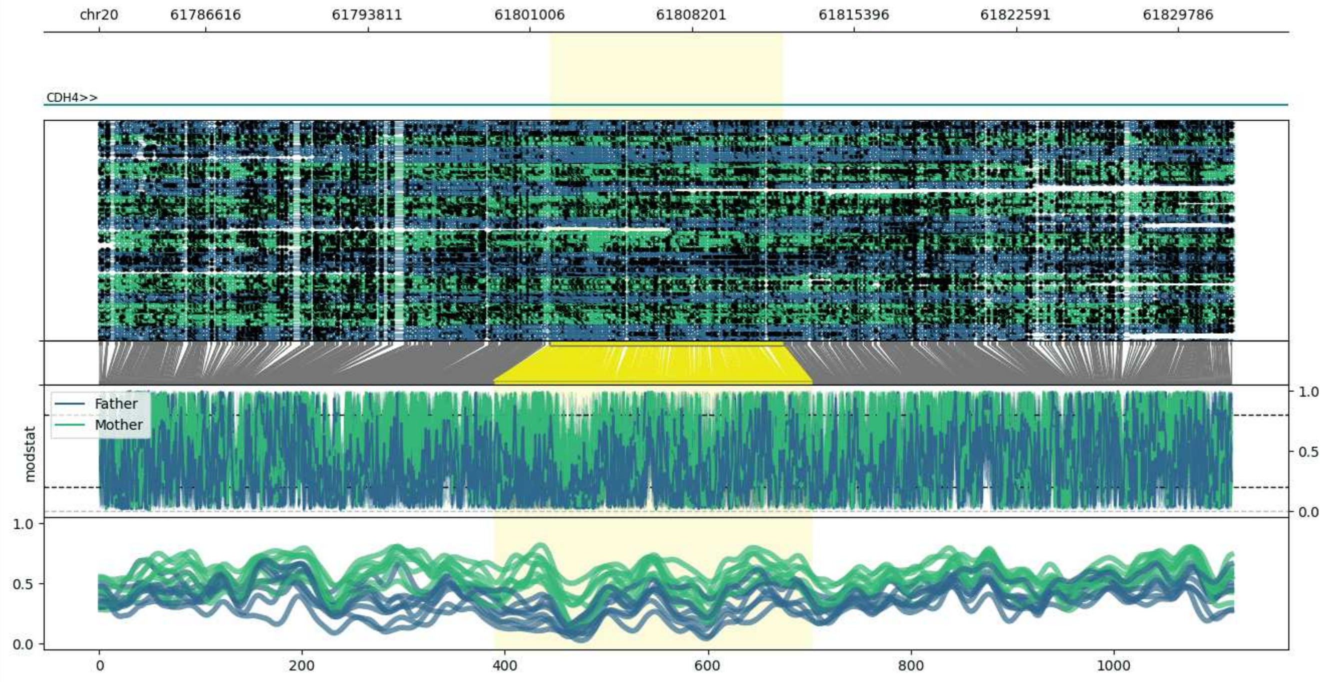


**chr20:61781894-61832260:** Region with 1 DMR containing genes: CDH4.

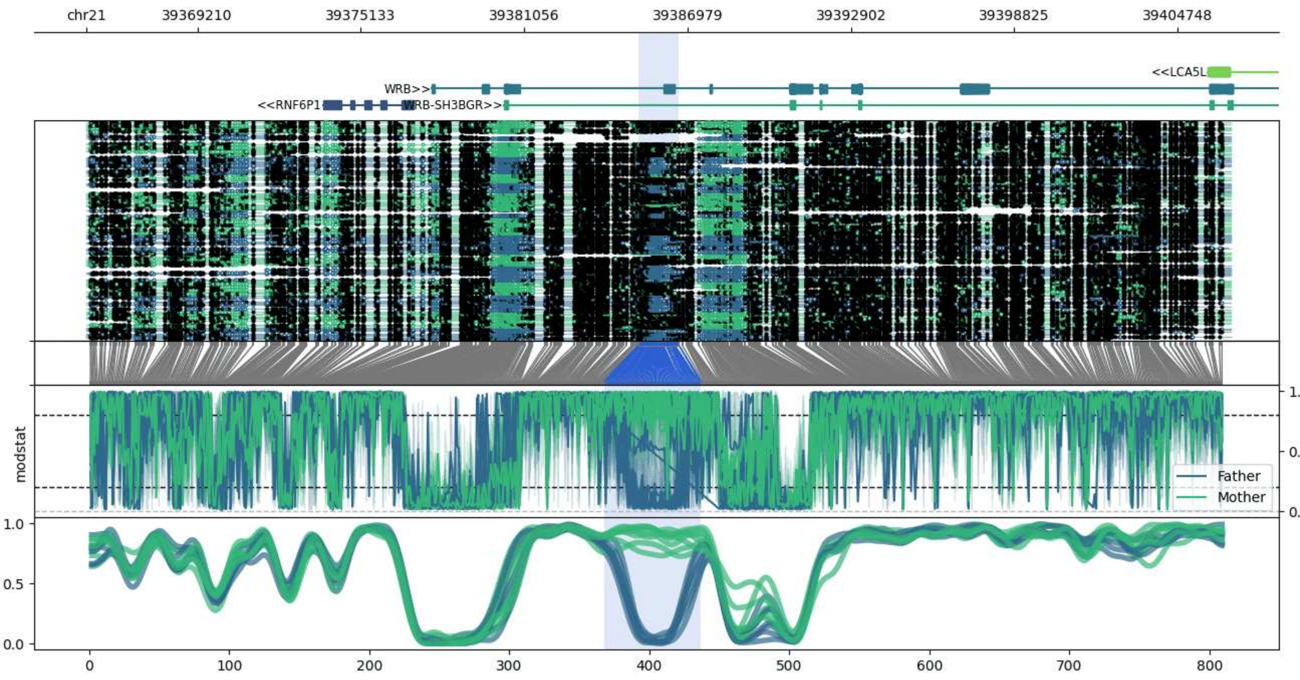


**chr21:39365206-39406673:** Region with 1 DMR containing genes: LCA5L, RNF6P1, WRB, WRB-SH3BGR.

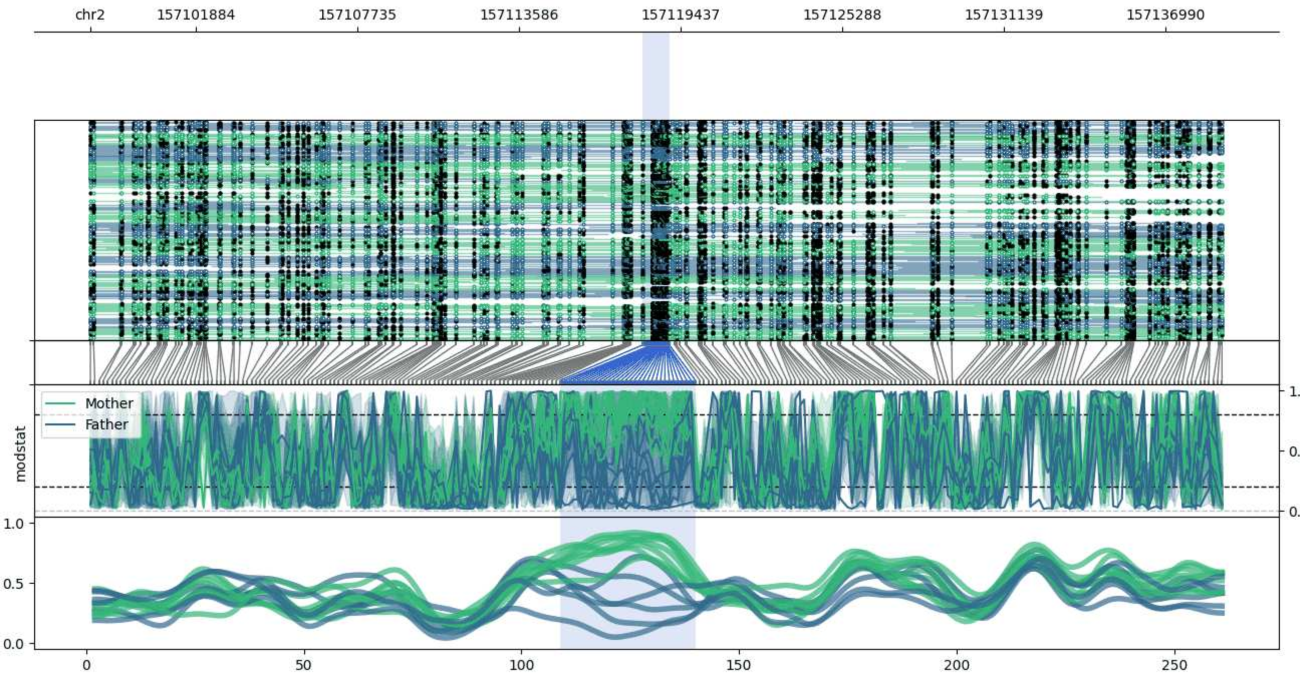


**chr2:157098065-157139027:** Region with 1 DMR containing no genes.

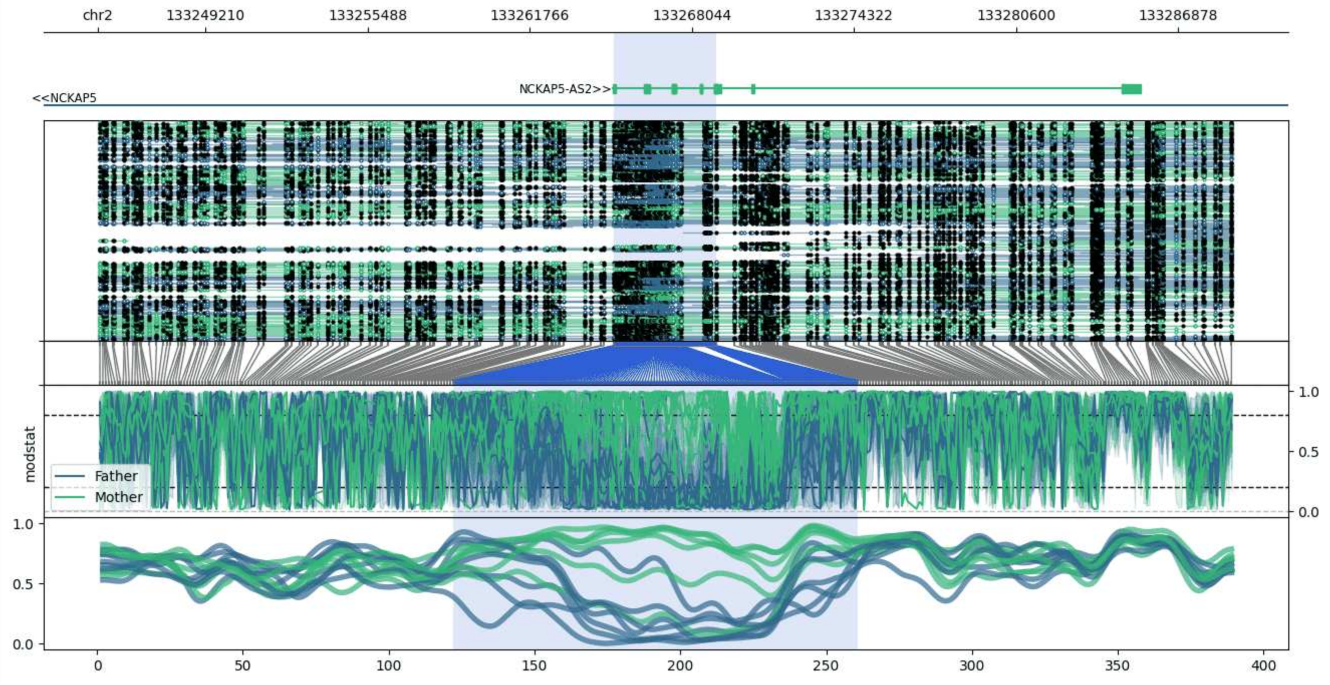


**chr2:133245042-133288994:** Region with 1 DMR containing genes: NCKAP5, NCKAP5-AS2.

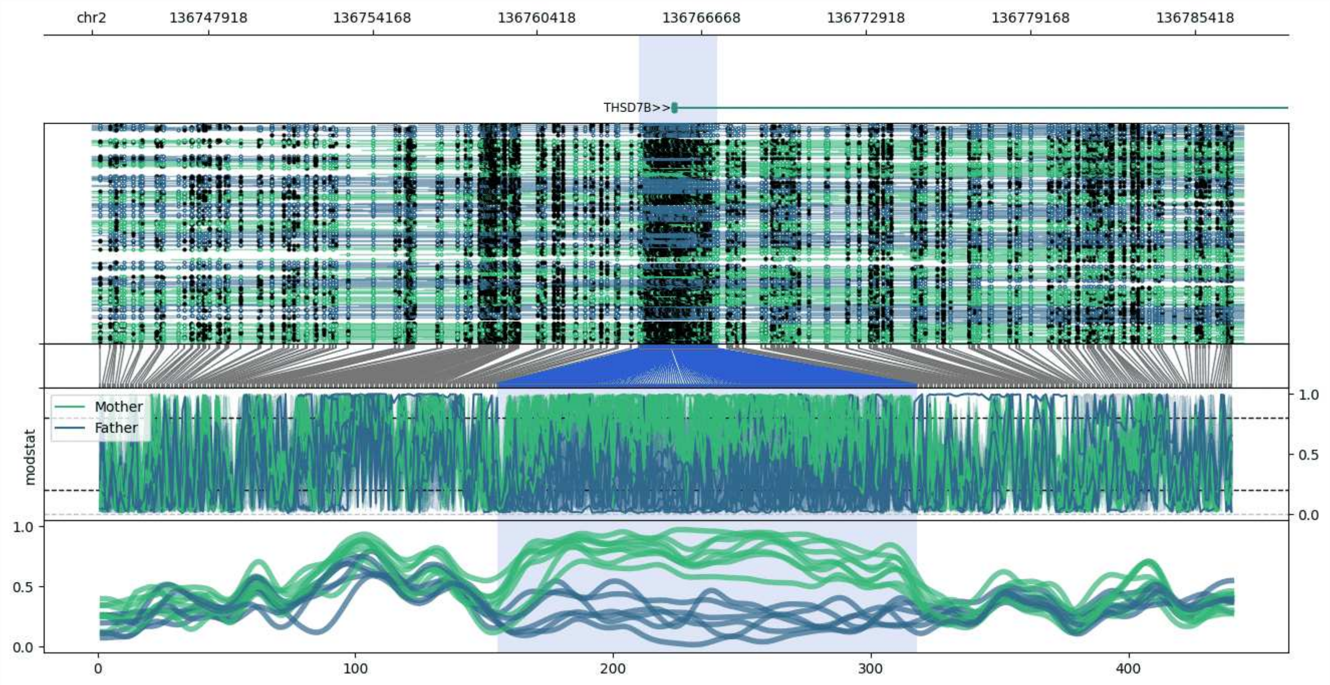


**chr2:136743502-136787255:** Region with 1 DMR containing genes: THSD7B.

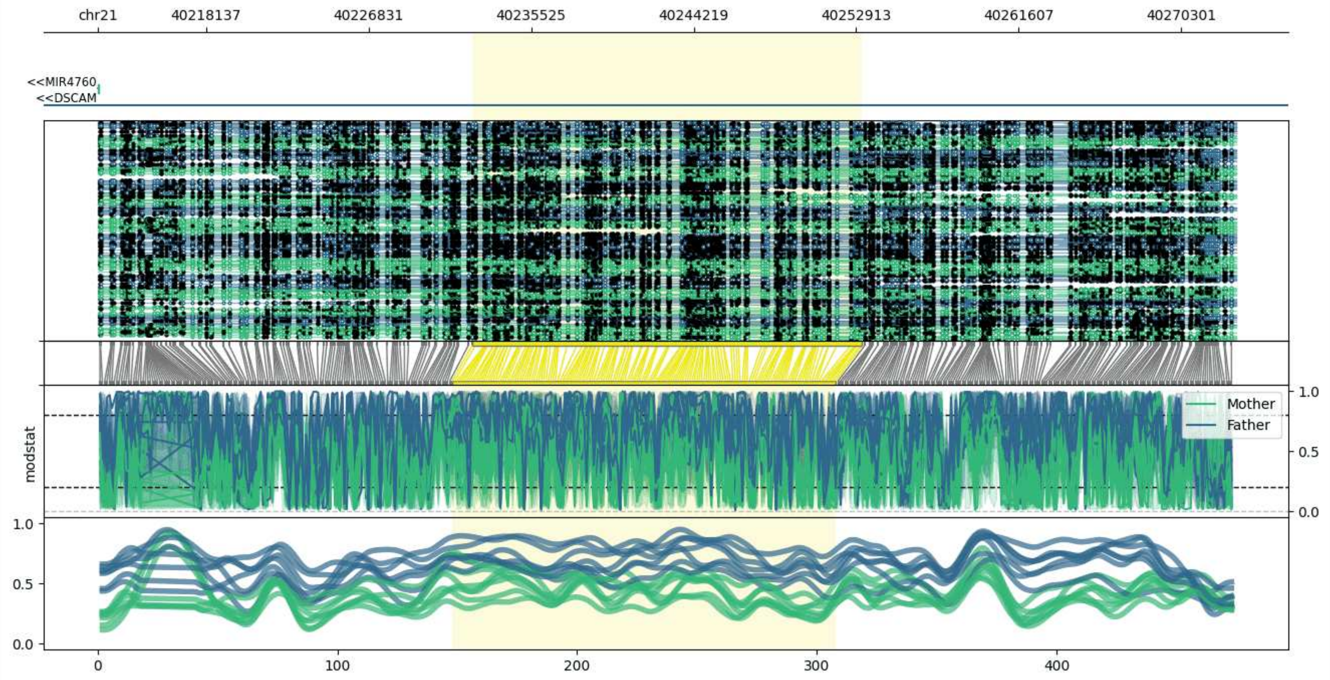


**chr21:40212365-40273228:** Region with 1 DMR containing genes: DSCAM, MIR4760.

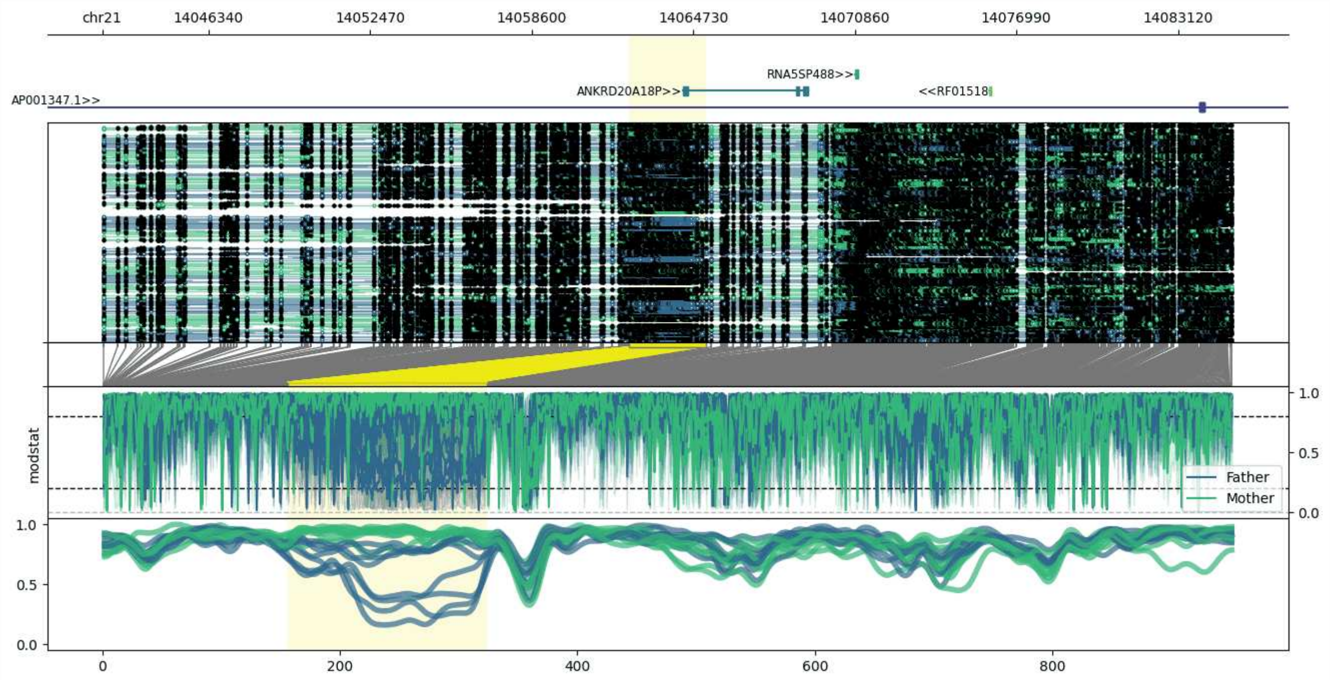


**chr21:14042294-14085205:** Region with 1 DMR containing genes: ANKRD20A18P, AP001347.1, RF01518, RNA5SP488.

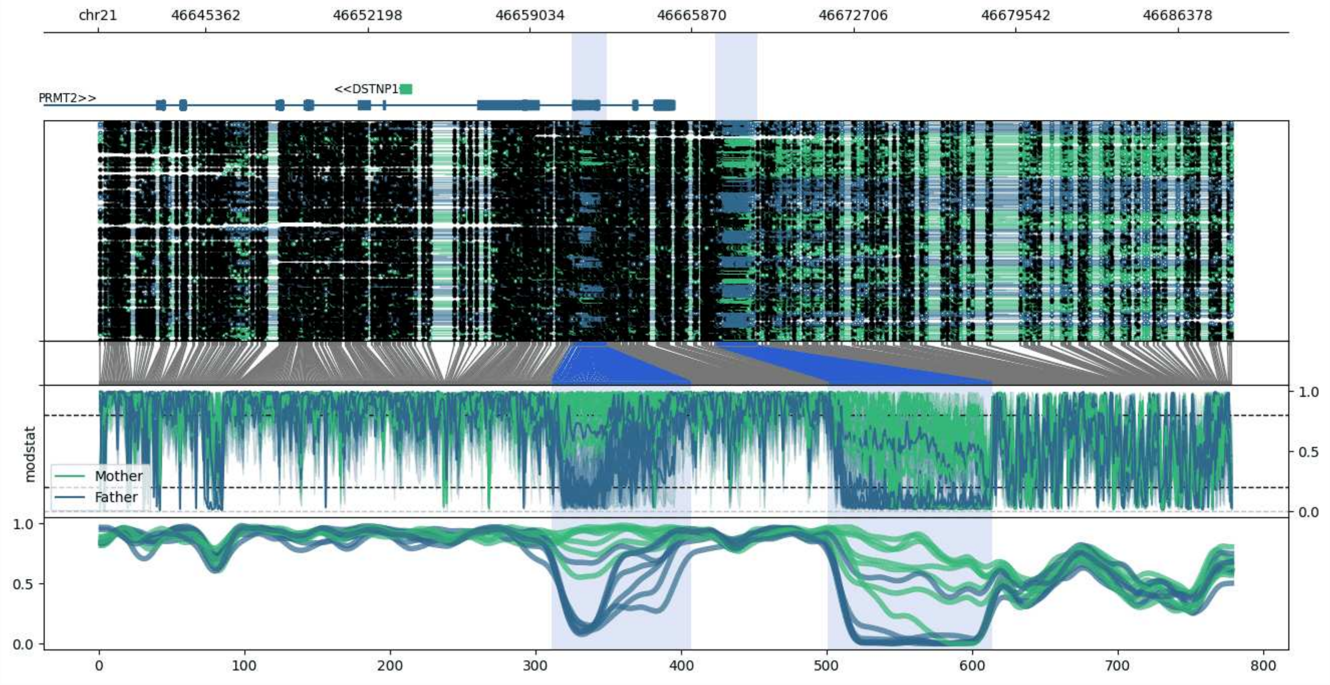


**chr21:46640811-46688665:** Region with 2 DMRs containing genes: DSTNP1, PRMT2, RPL23AP4.

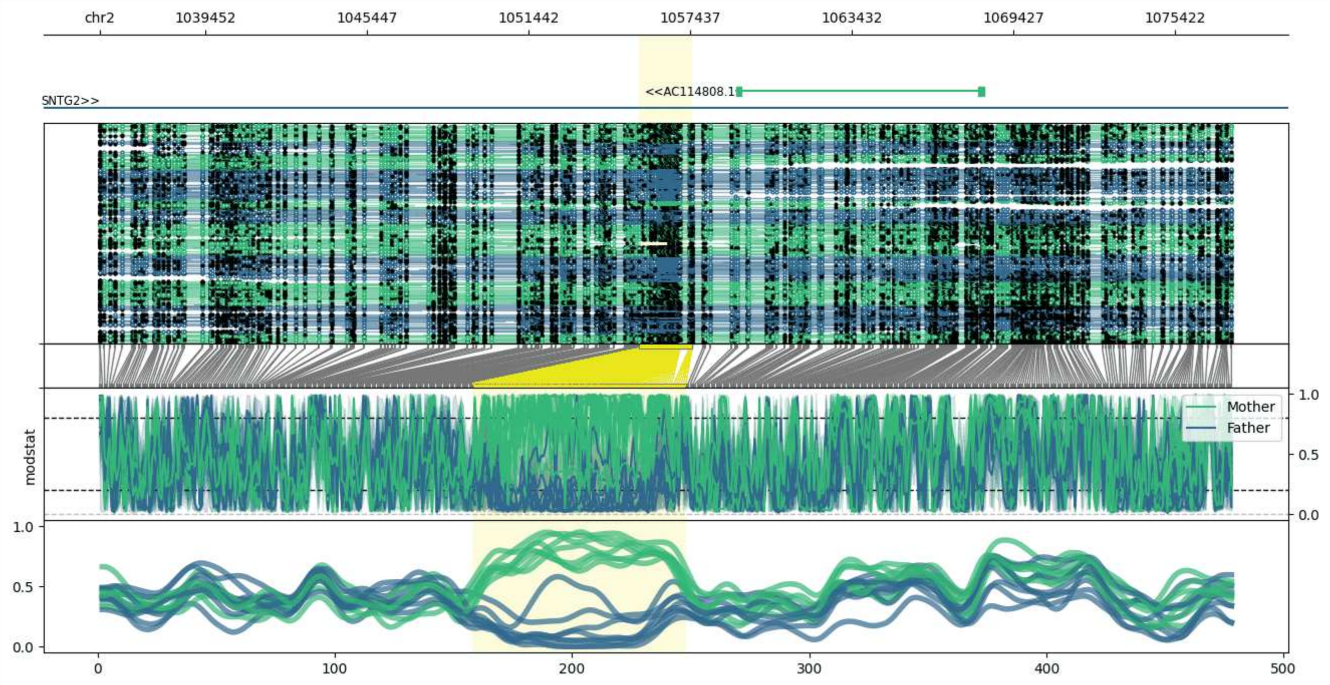


**chr2:1035549-1077519:** Region with 1 DMR containing genes: AC114808.1, SNTG2.

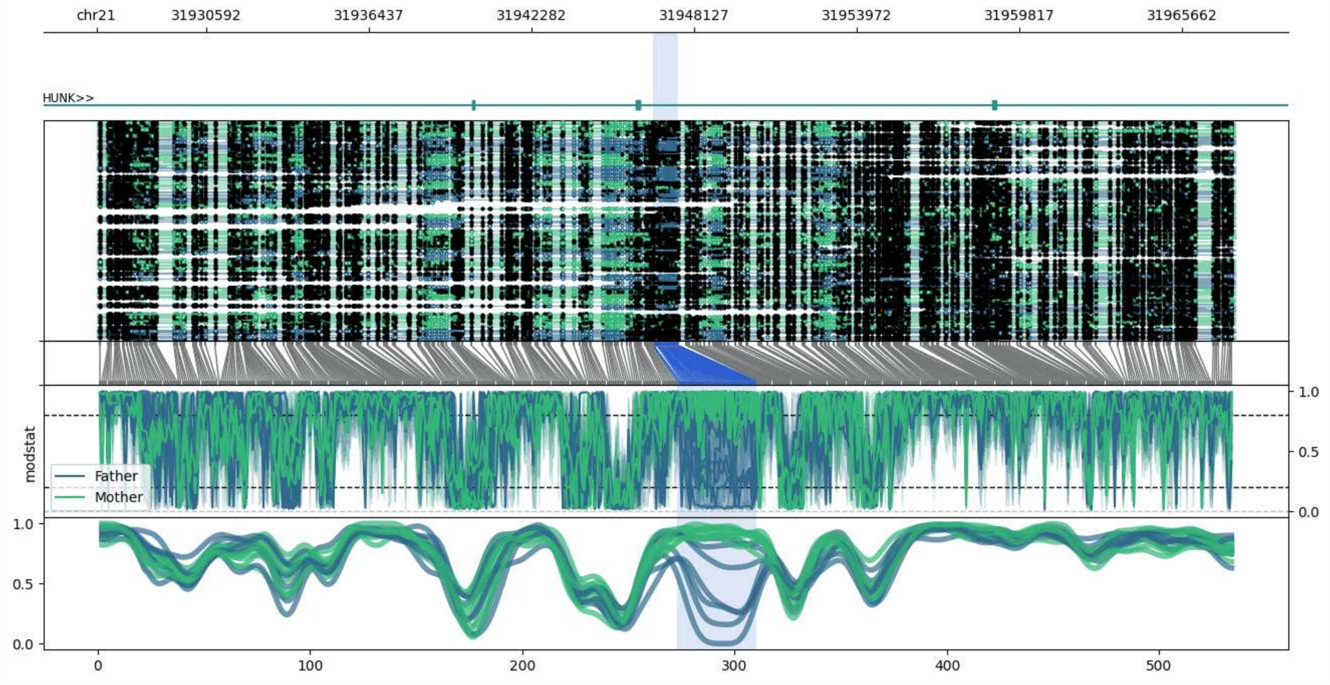


**chr21:31926652-31967569:** Region with 1 DMR containing genes: HUNK.

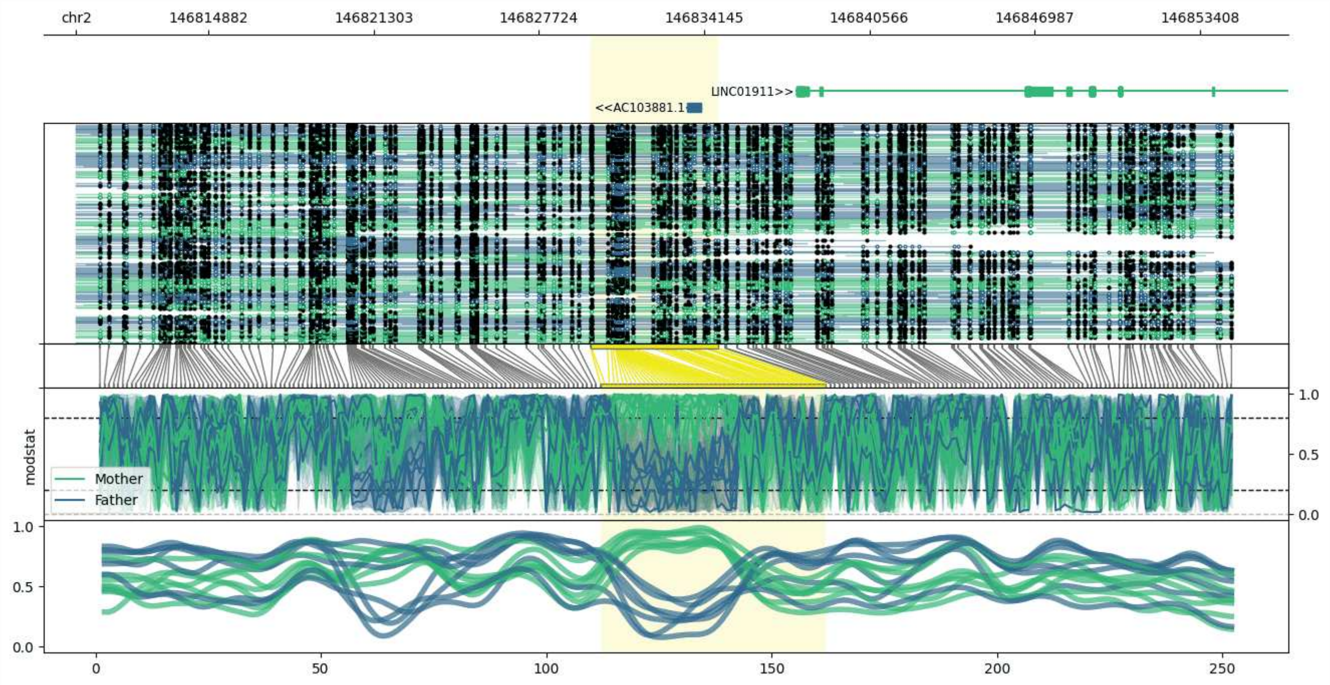


**chr2:146809730-146854679:** Region with 1 DMR containing genes: AC103881.1, LINC01911.

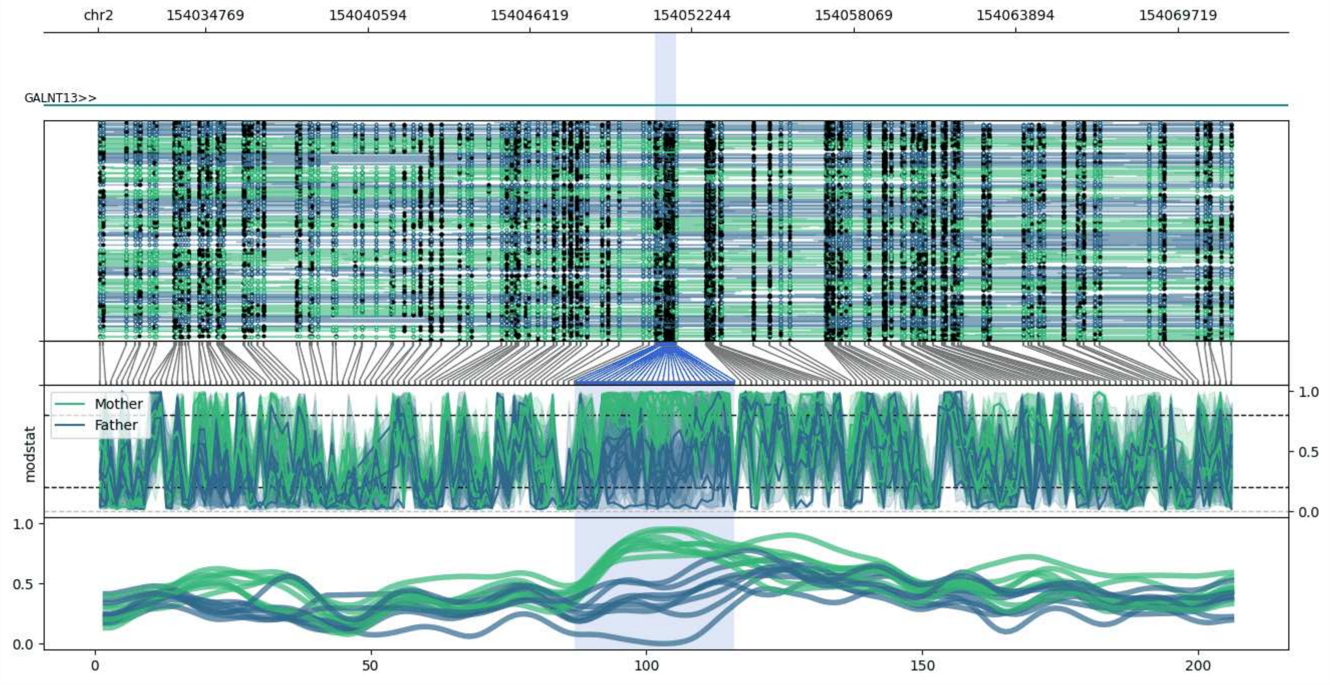


**chr2:154030913-154071691:** Region with 1 DMR containing genes: GALNT13.

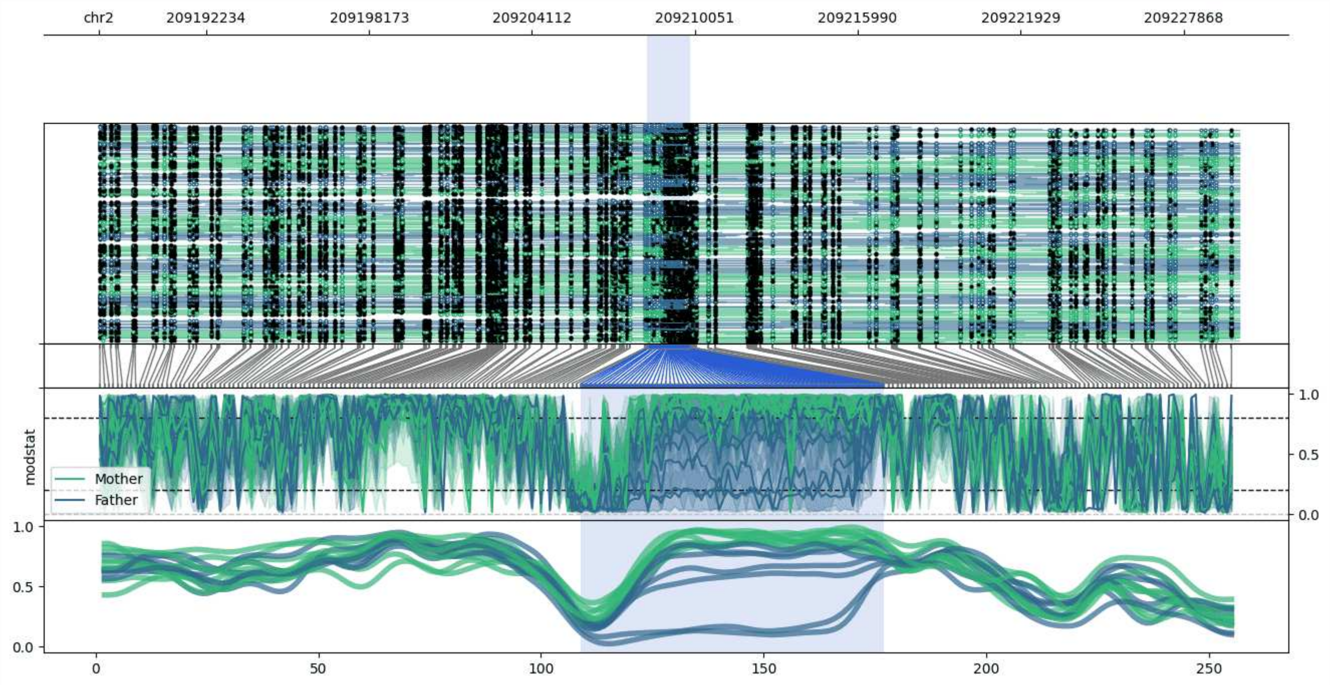


**chr2:209188311-209229889:** Region with 1 DMR containing genes: MEAF6P1, PKP4P1.

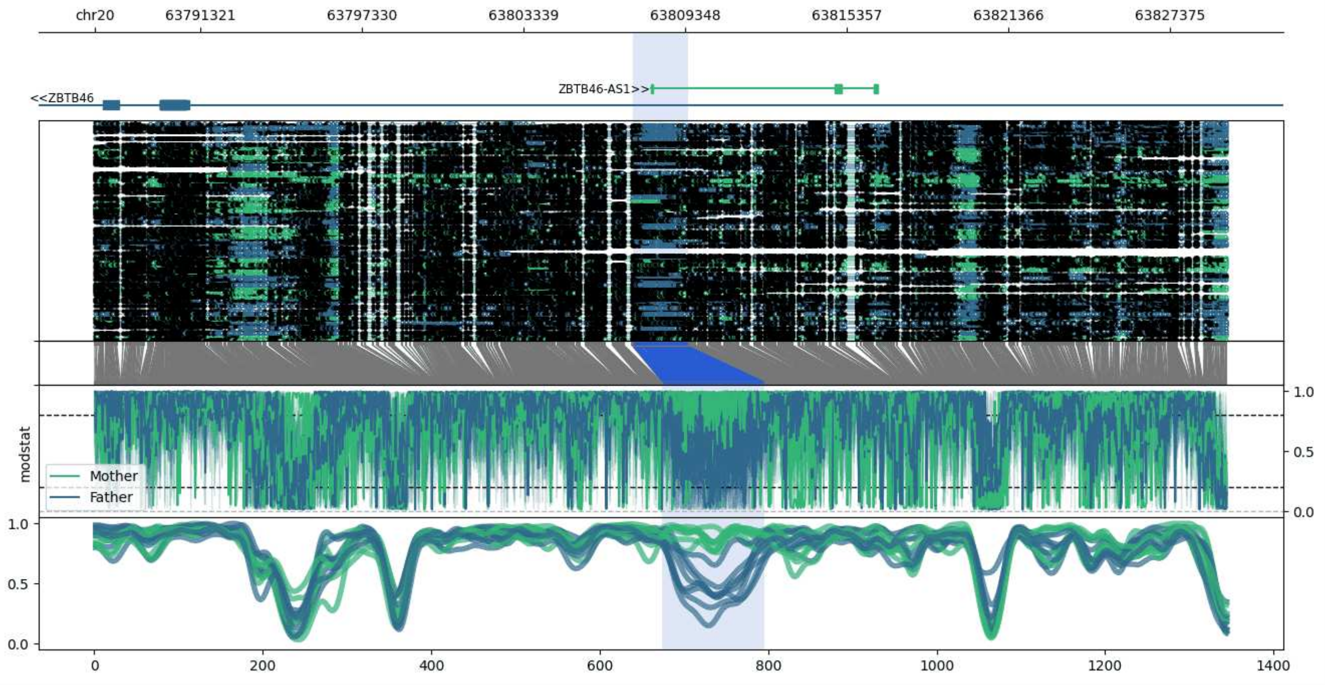


**chr20:63787404-63829469:** Region with 1 DMR containing genes: ZBTB46, ZBTB46-AS1.

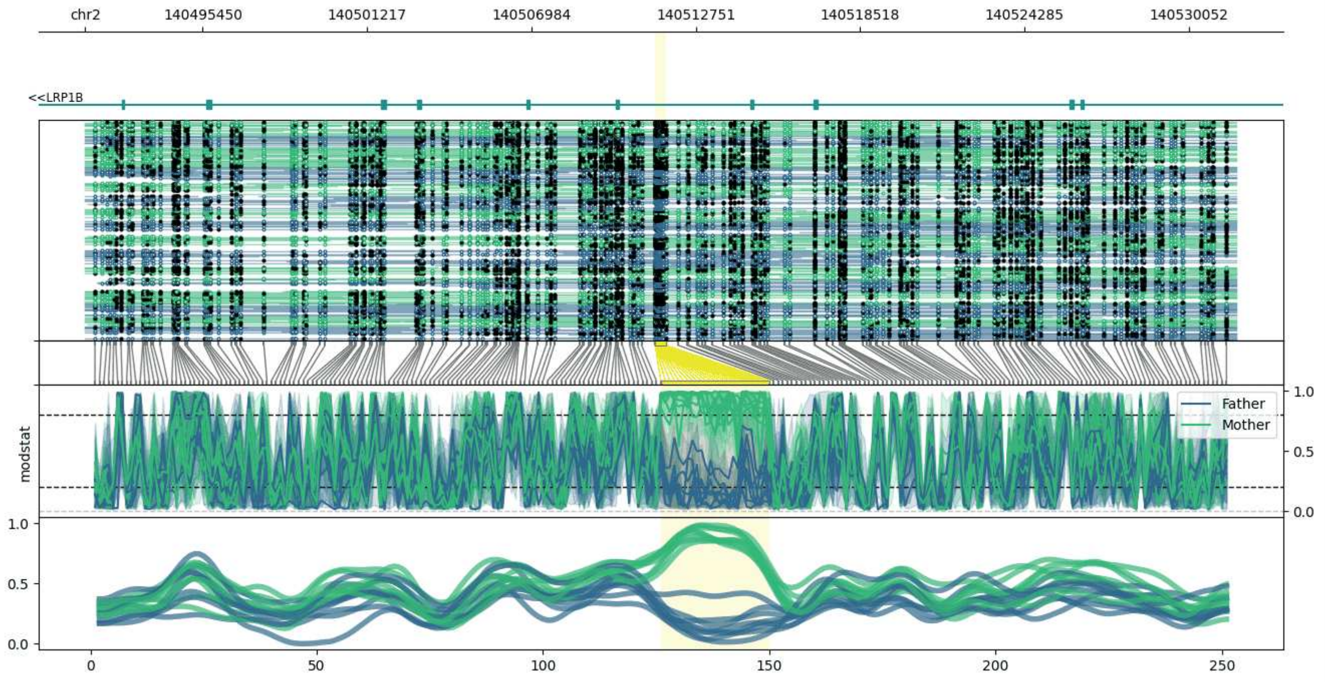


**chr2:140491323-140531695:** Region with 1 DMR containing genes: LRP1B.

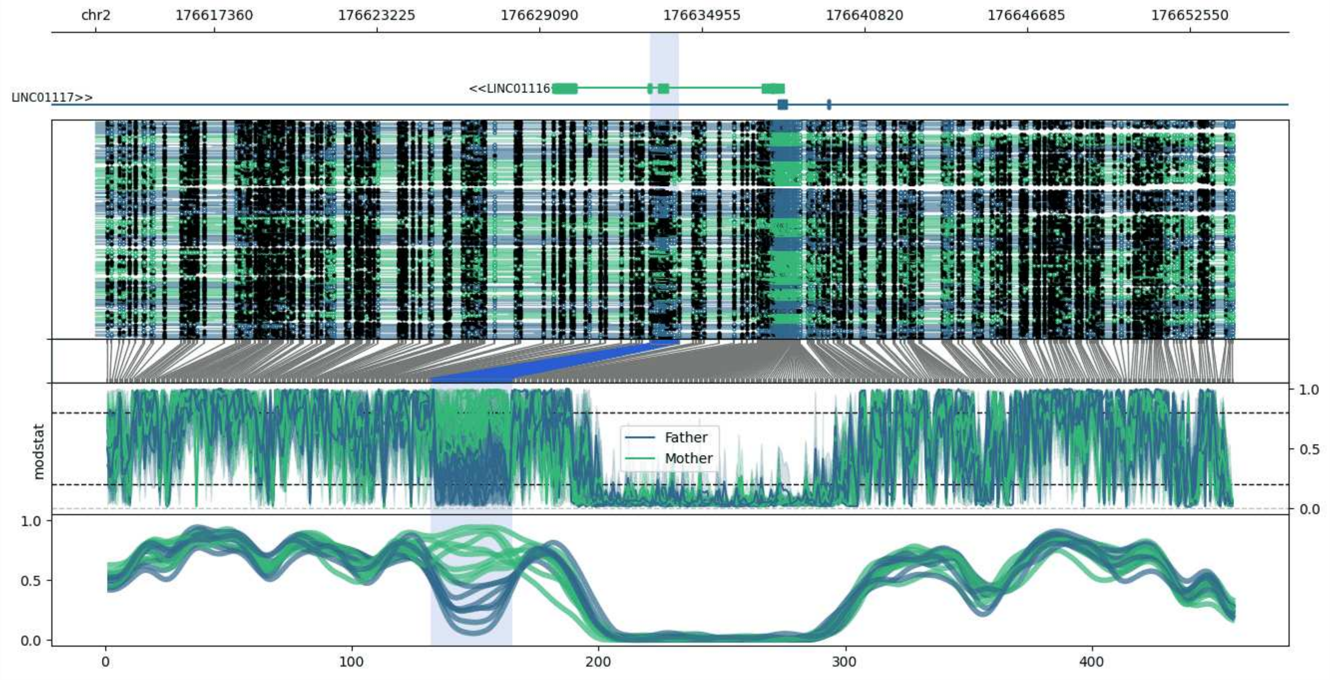


**chr2:176613093-176654153:** Region with 1 DMR containing genes: AC017048.3, LINC01116, LINC01117.

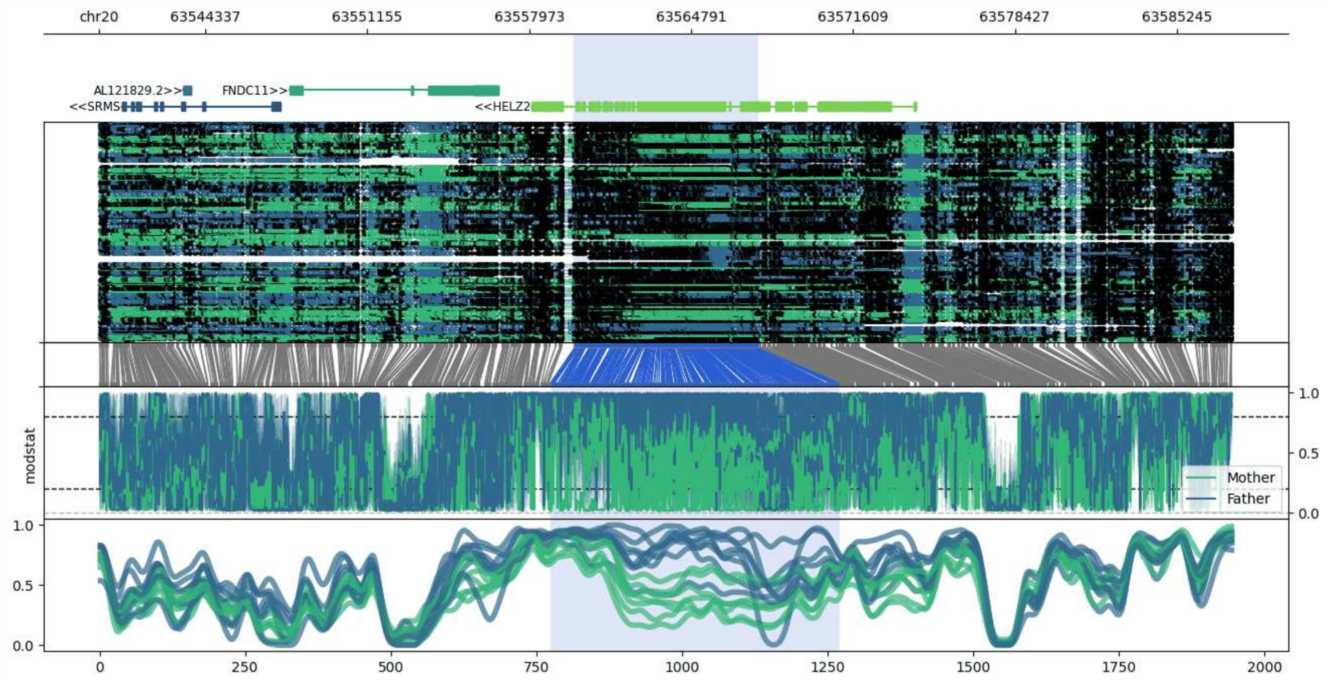


**chr20:63539857-63587586:** Region with 1 DMR containing genes: AL121829.2, FNDC11, GMEB2, HELZ2, PTK6, SRMS.

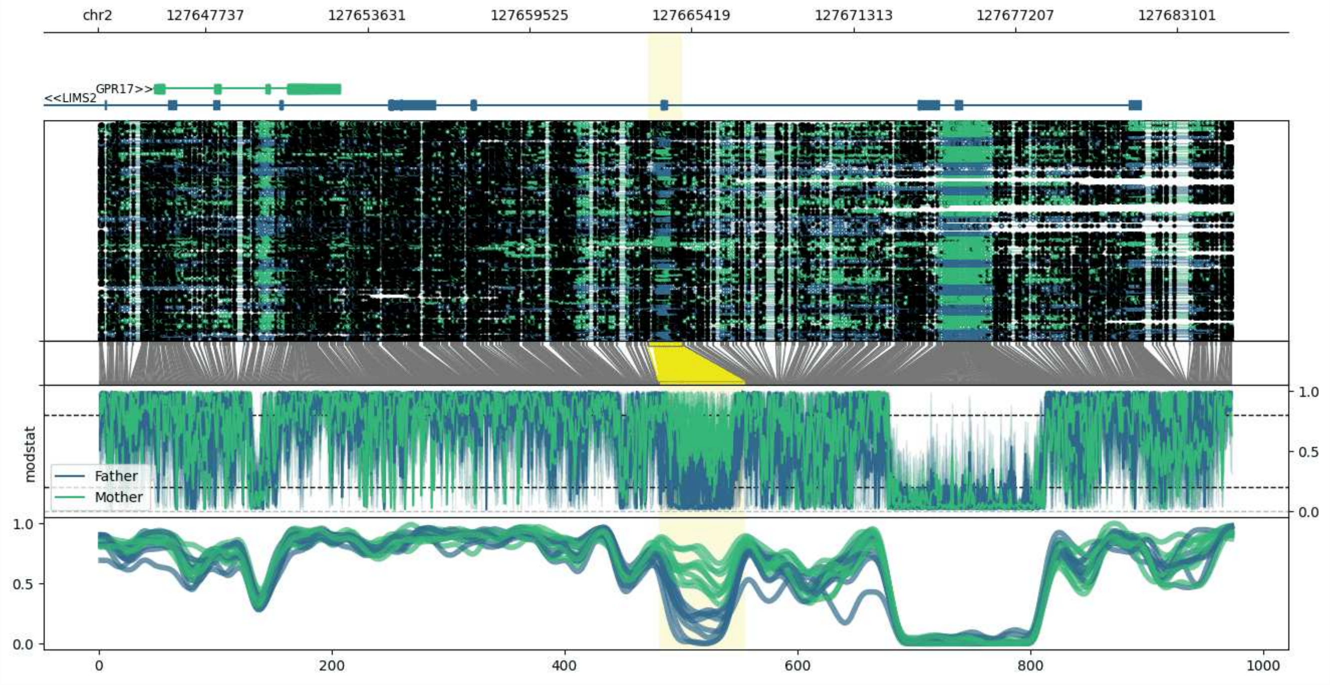


**chr2:127643831-127685089:** Region with 1 DMR containing genes: GPR17, LIMS2, MYO7B.

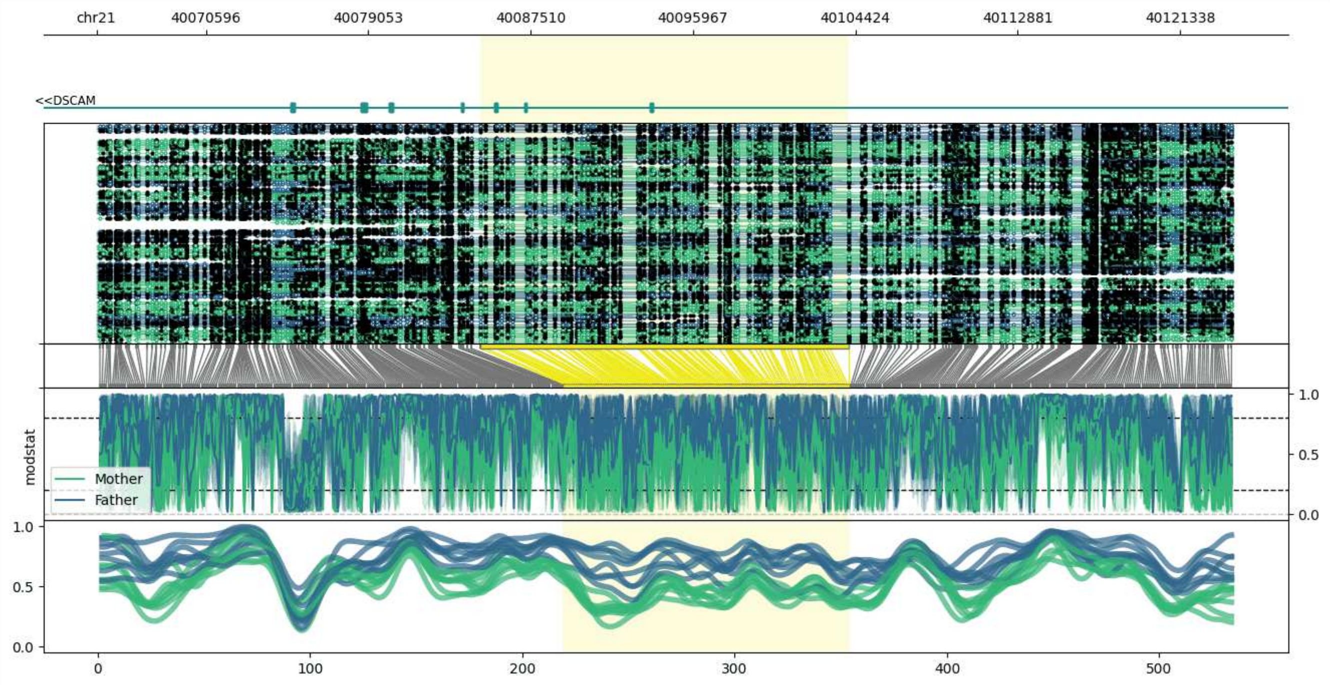


**chr21:40064899-40124103:** Region with 1 DMR containing genes: DSCAM.

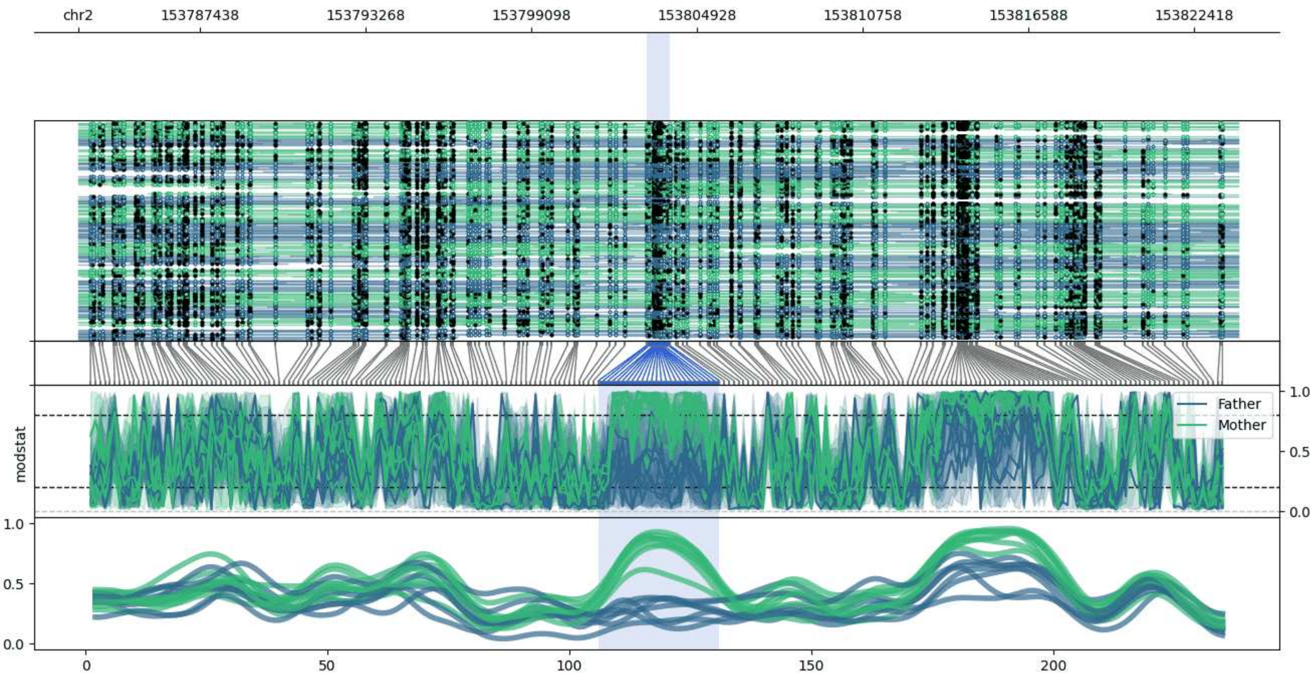


**chr2:153783169-153823983:** Region with 1 DMR containing no genes.

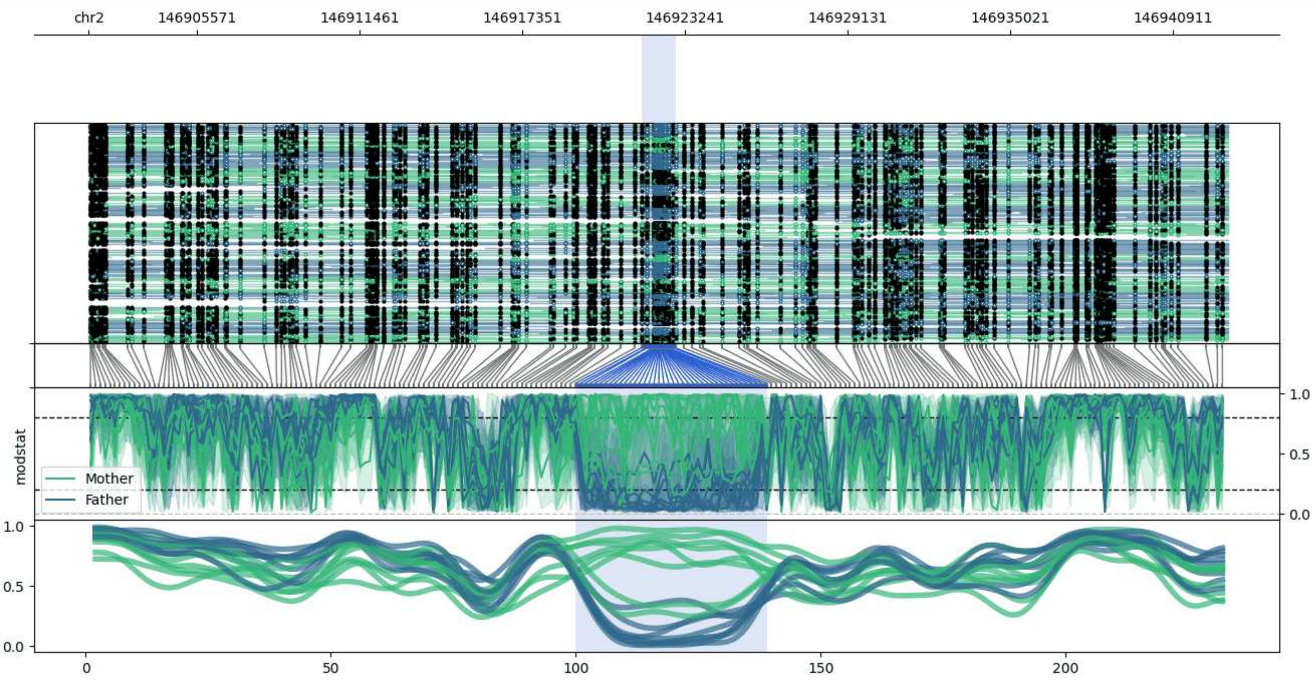


**chr2:146901670-146942904:** Region with 1 DMR containing genes: AC062032.1.

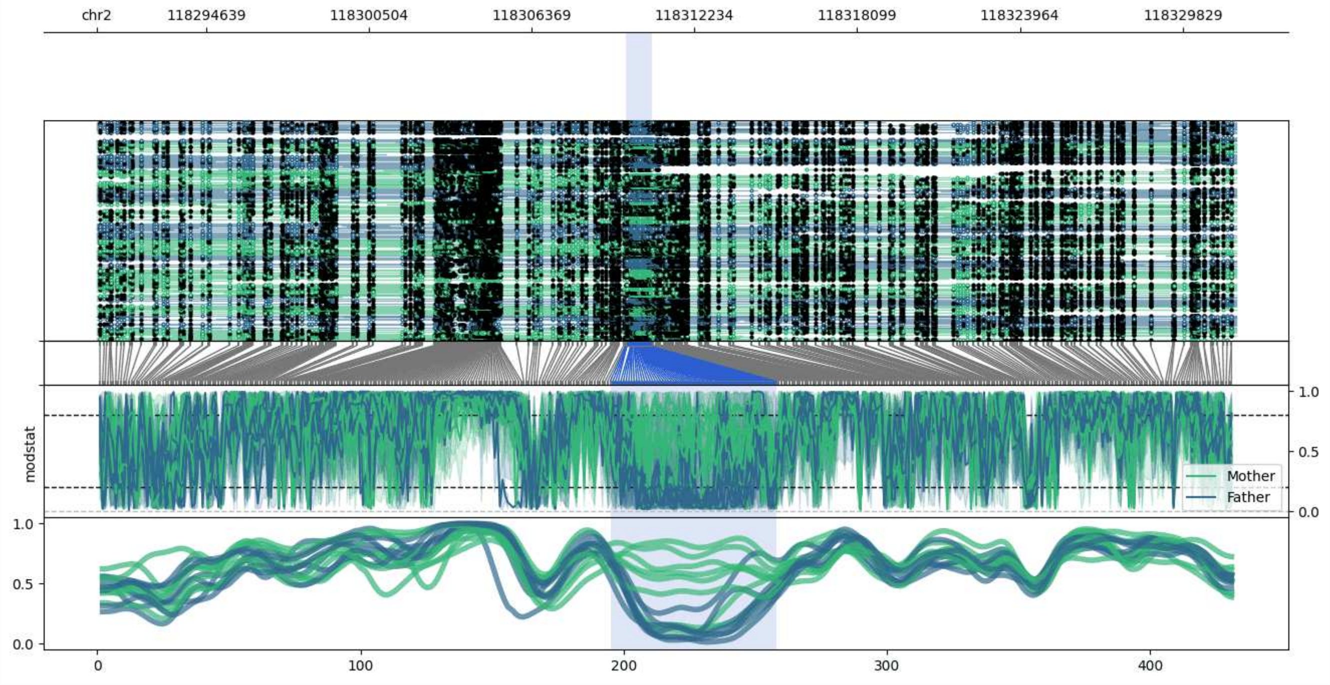


**chr2:118290699-118331756:** Region with 1 DMR containing no genes.

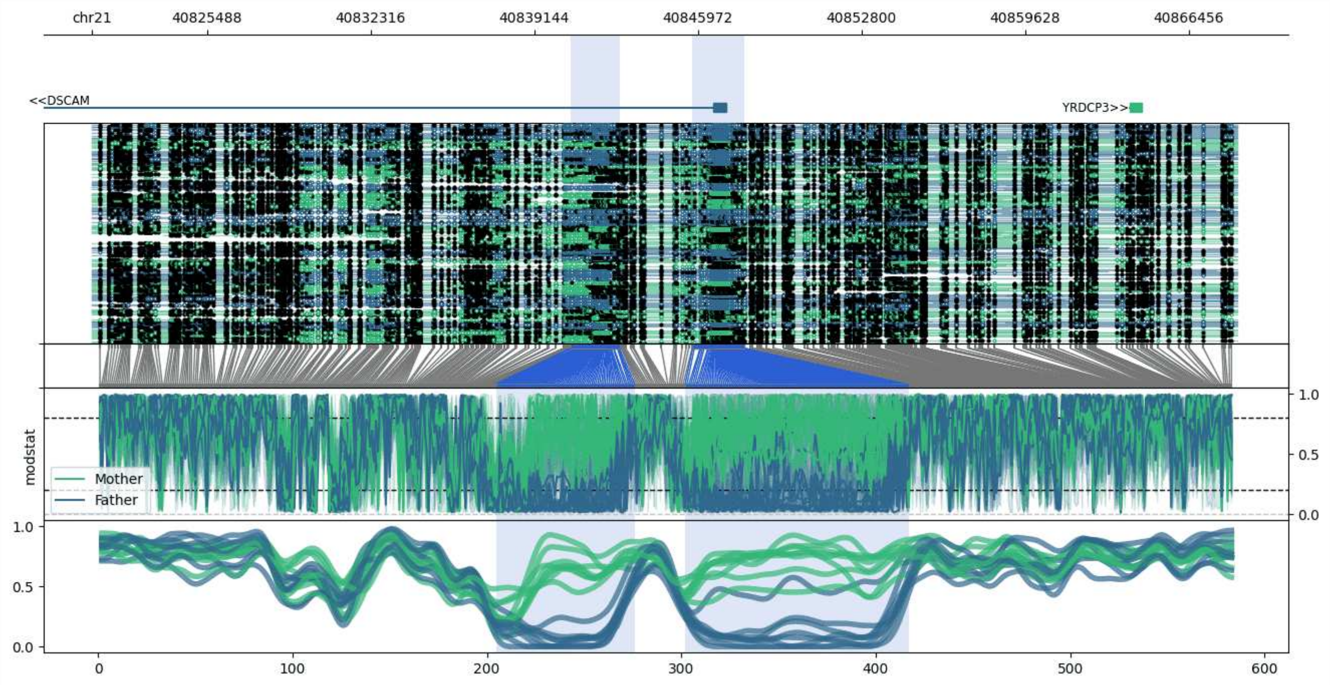


**chr21:40820674-40868473:** Region with 2 DMRs containing genes: DSCAM, YRDCP3.

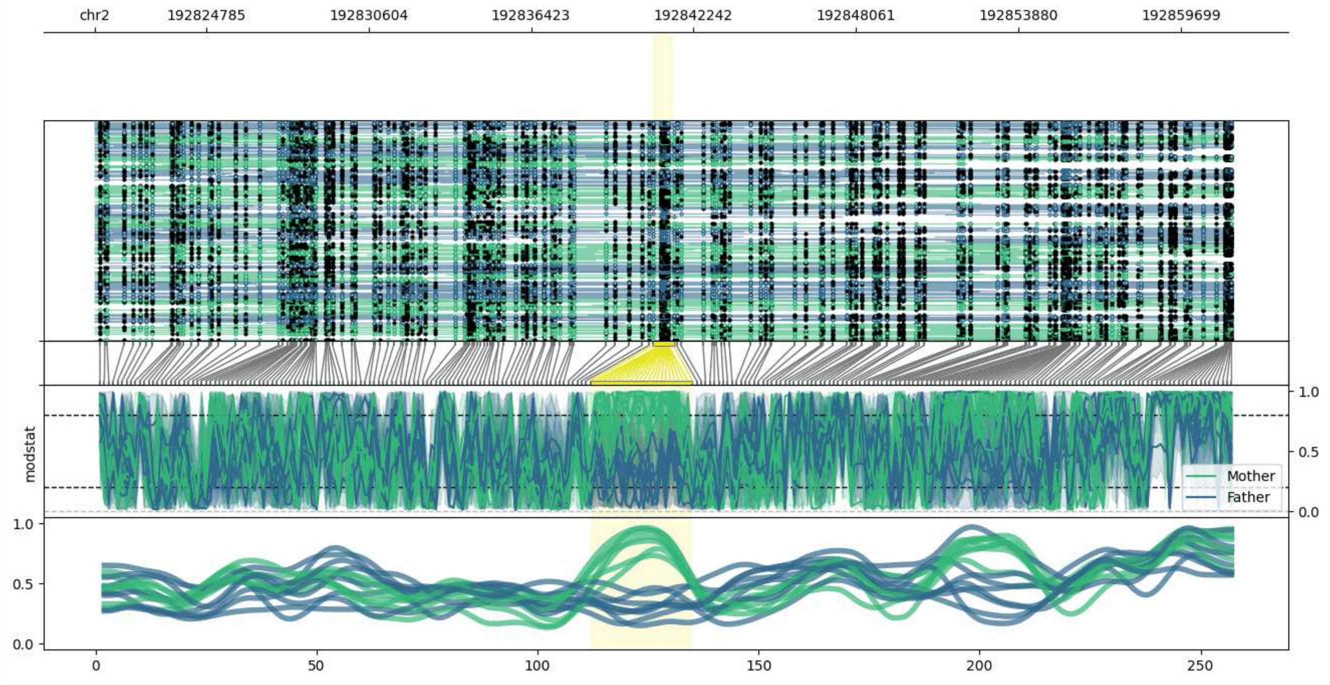


**chr2:192820792-192861529:** Region with 1 DMR containing no genes.

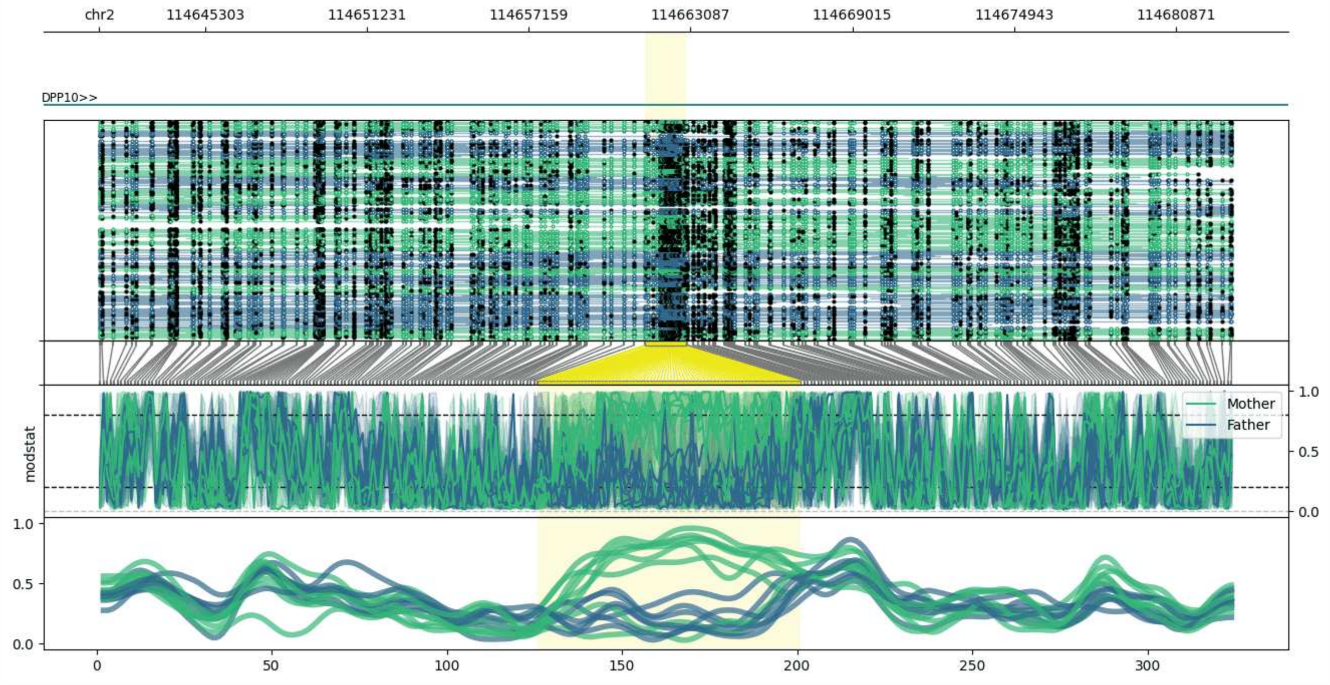


**chr2:114641422-114682922:** Region with 1 DMR containing genes: DPP10.

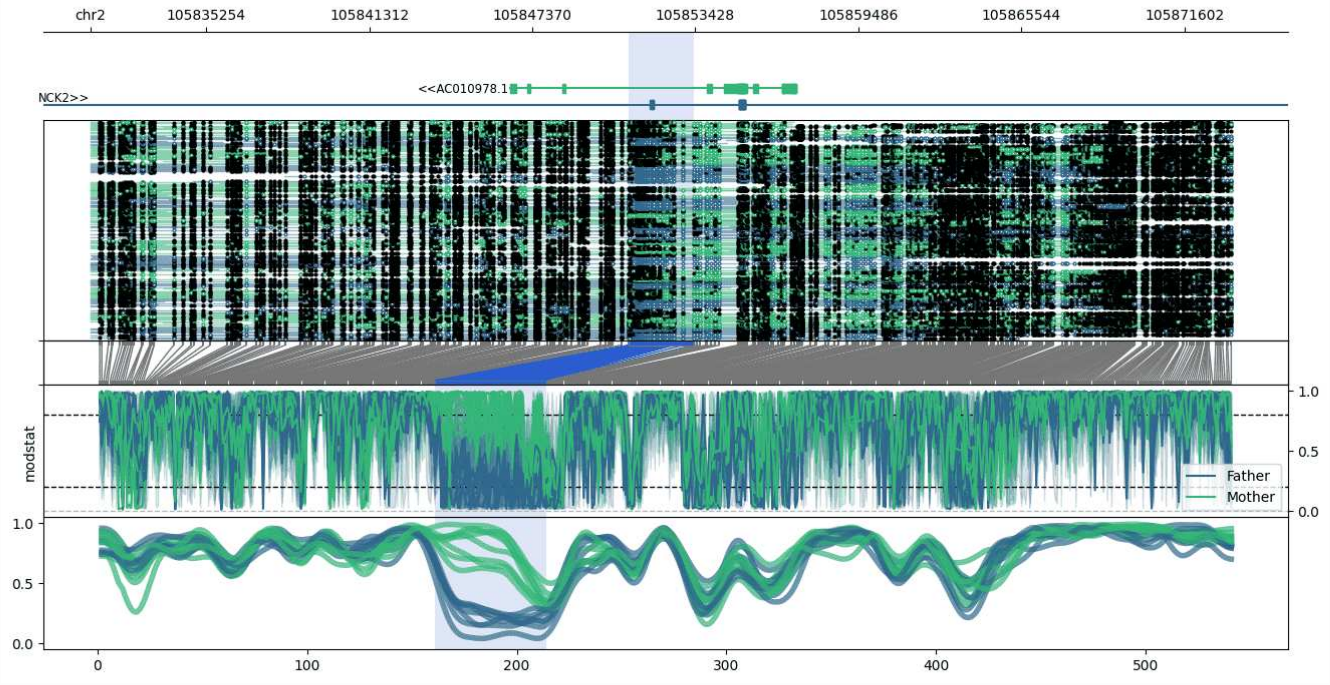


**chr2:105830953-105873362:** Region with 1 DMR containing genes: AC009505.1, AC010978.1, NCK2.

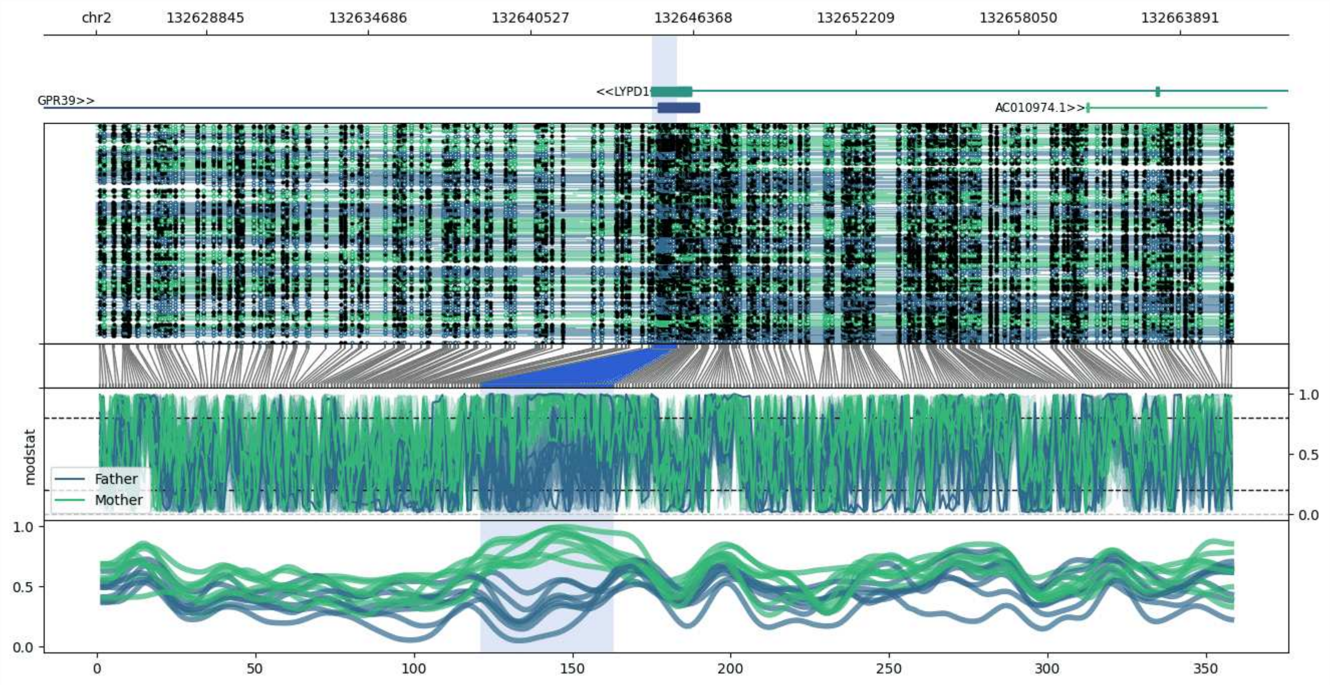


**chr2:132624901-132665788:** Region with 1 DMR containing genes: AC010974.1, GPR39, LYPD1, NCKAP5.

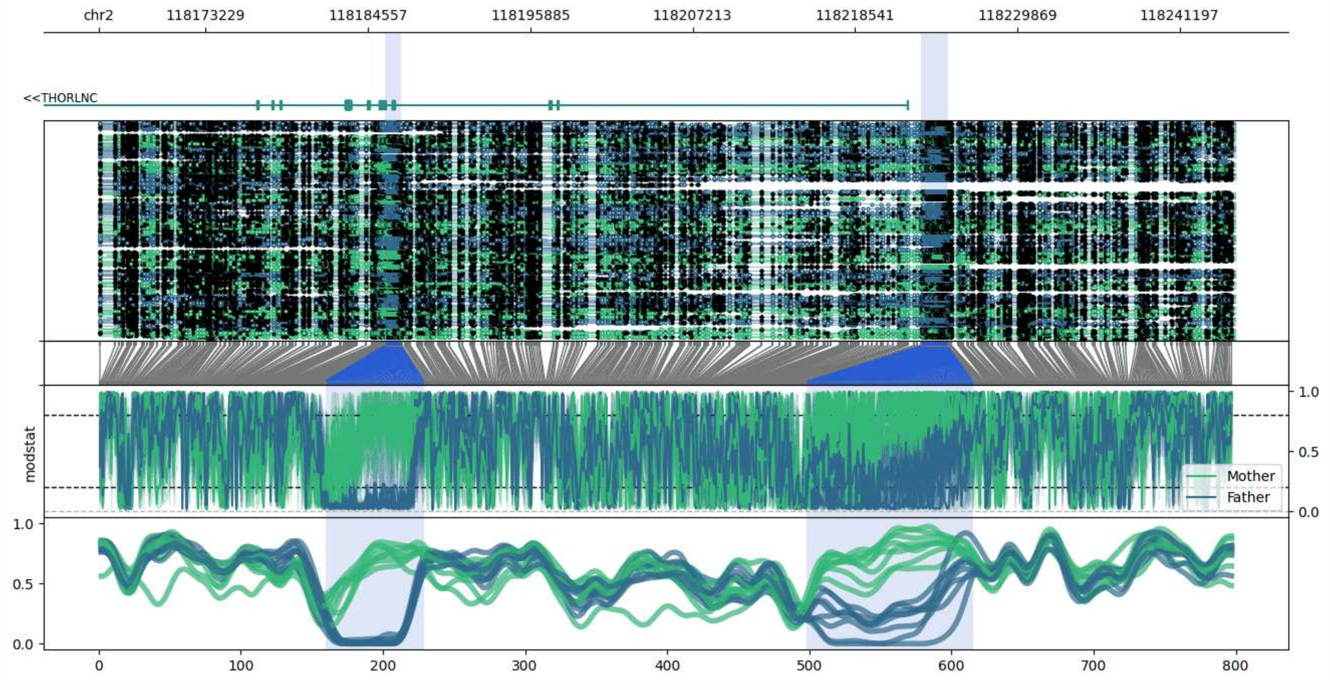


**chr2:118165752-118245052:** Region with 2 DMRs containing genes: THORLNC.

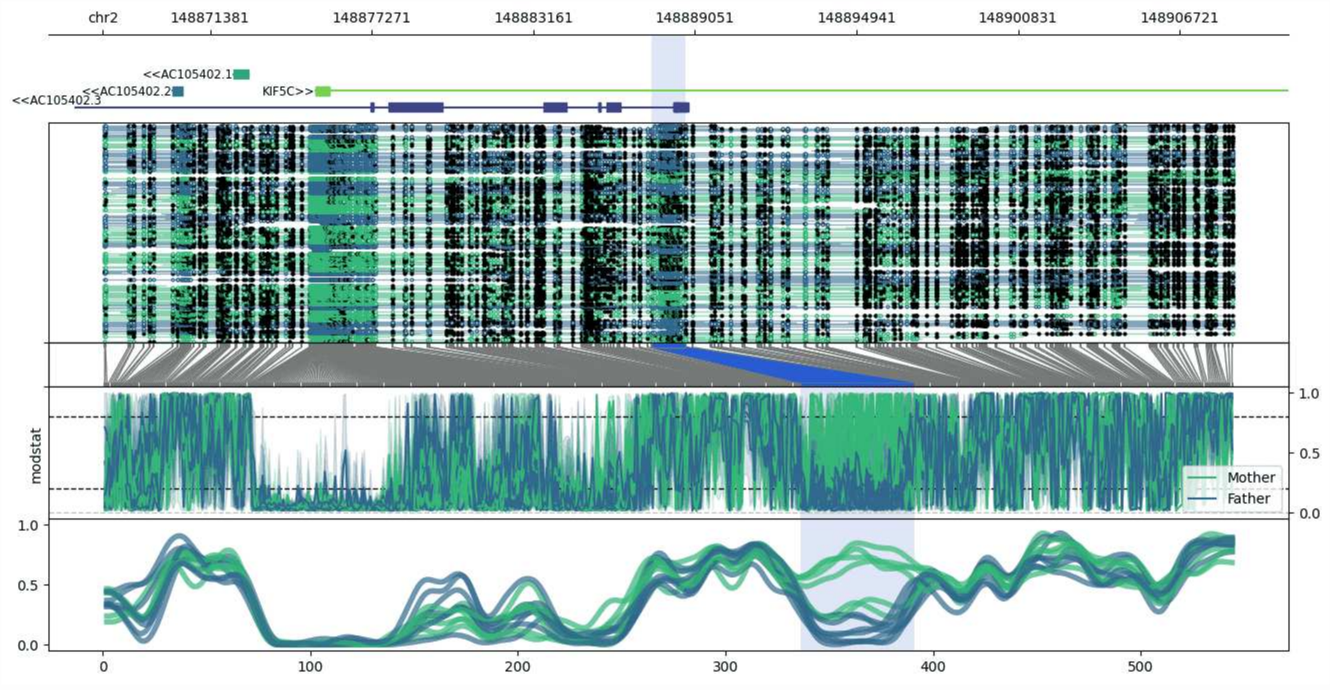


**chr2:148867471-148908707:** Region with 1 DMR containing genes: AC105402.1, AC105402.2, AC105402.3, AC105402.4, KIF5C, UBBP3.

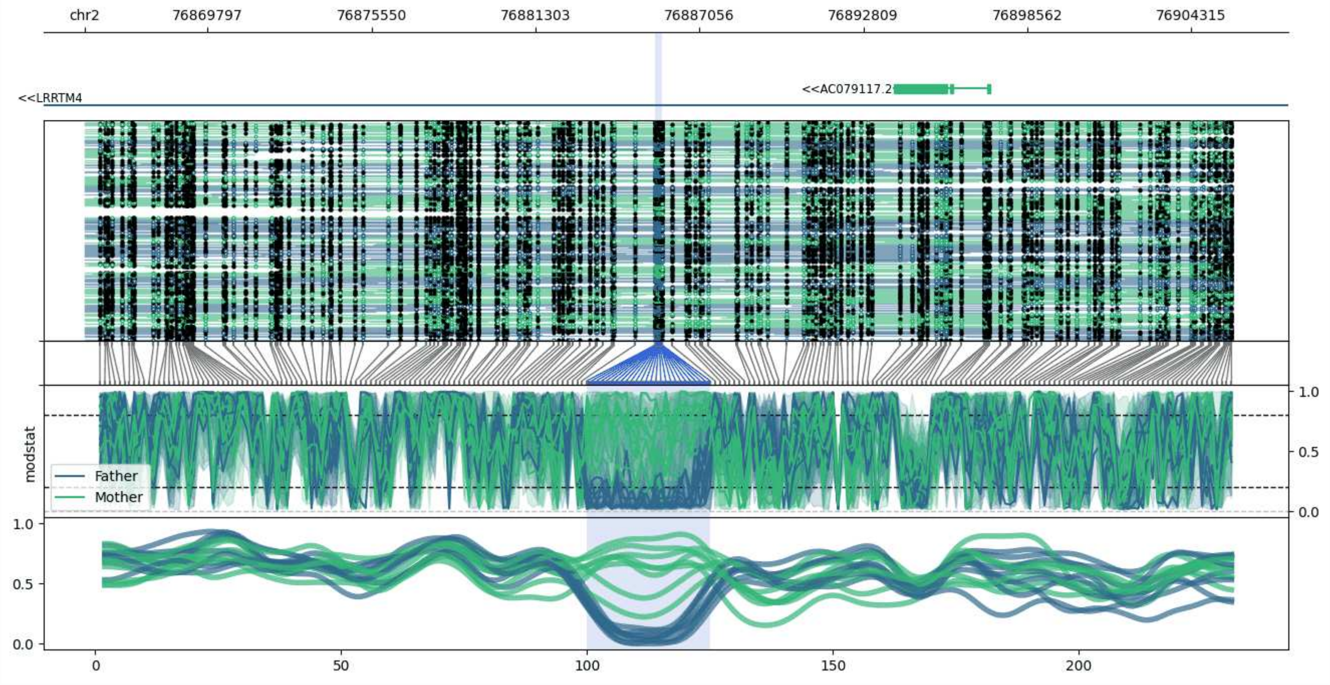


**chr2:76865484-76905759:** Region with 1 DMR containing genes: AC079117.2, LRRTM4.

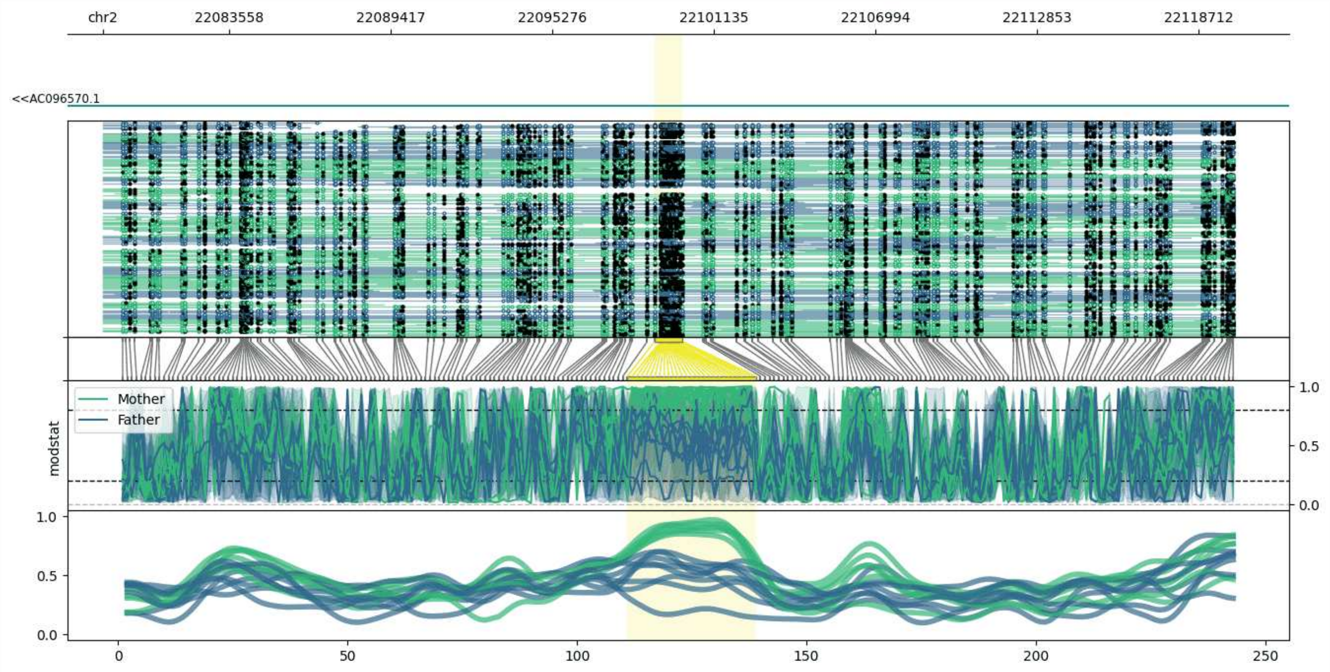


**chr2:22078994-22120007:** Region with 1 DMR containing genes: AC096570.1.

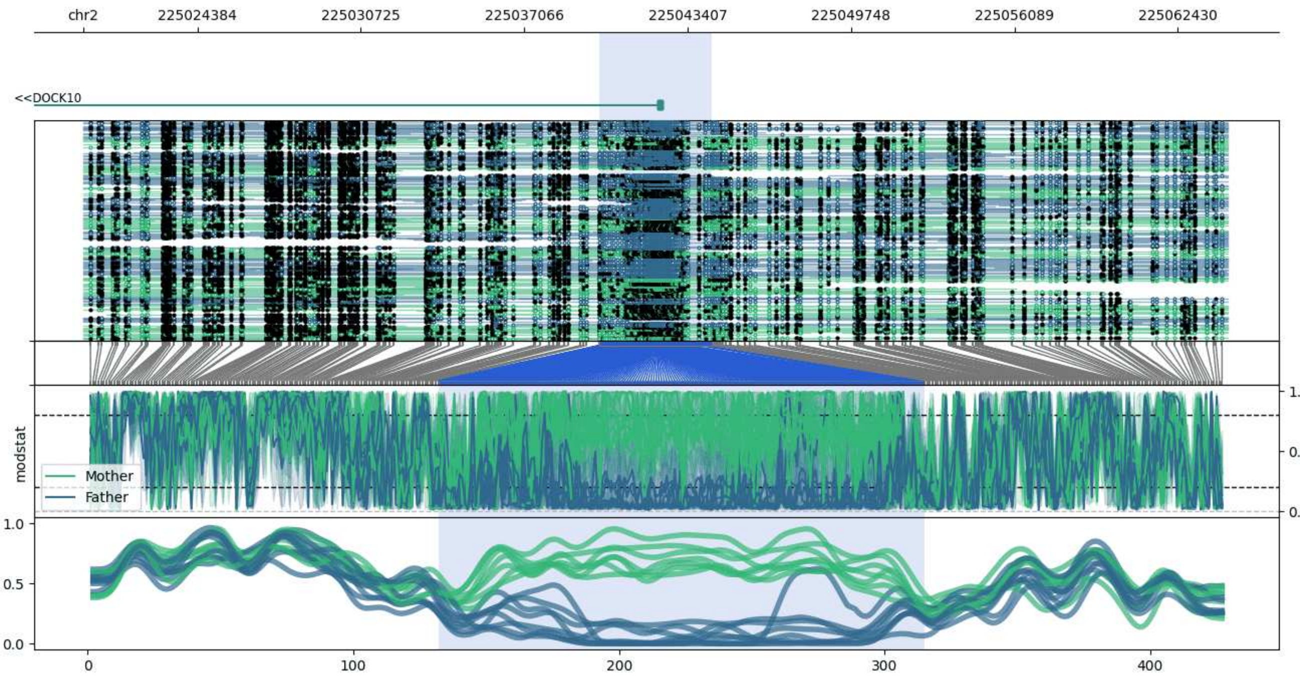


**chr2:225019965-225064353:** Region with 1 DMR containing genes: DOCK10, MIR4439.

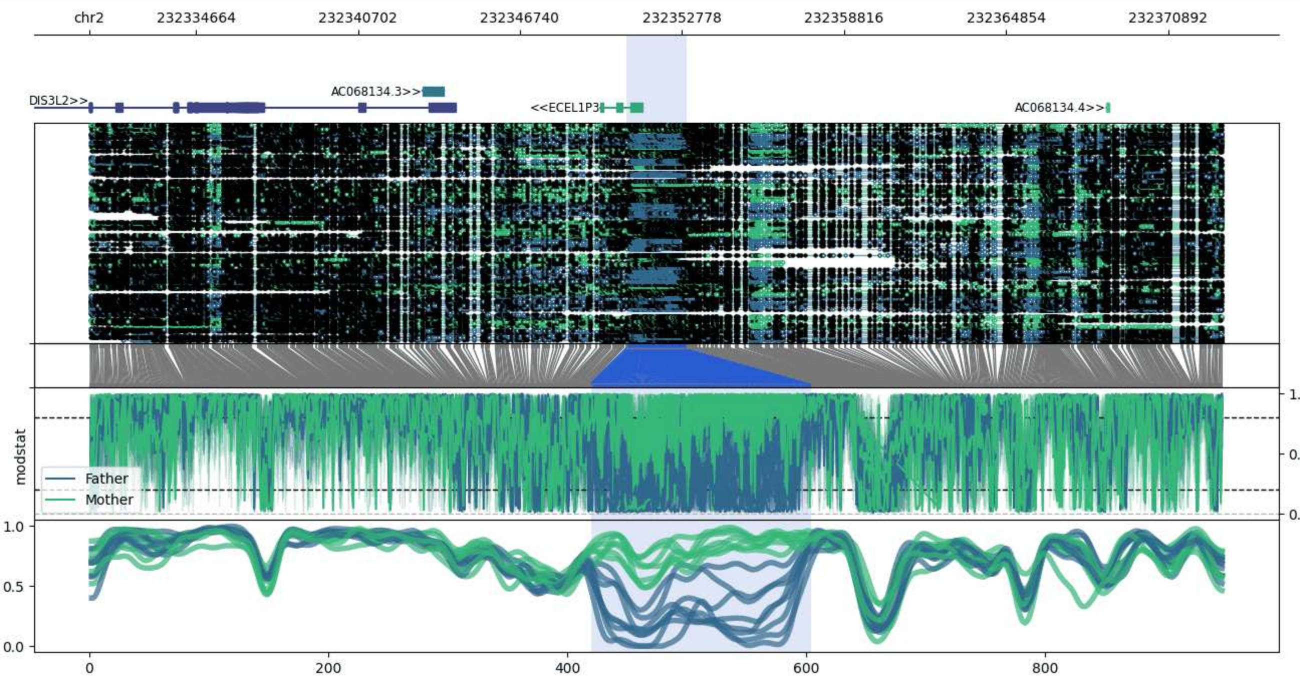


**chr2:232330700-232372970:** Region with 1 DMR containing genes: AC068134.1, AC068134.3, AC068134.4, ALPP, DIS3L2, ECEL1P3.

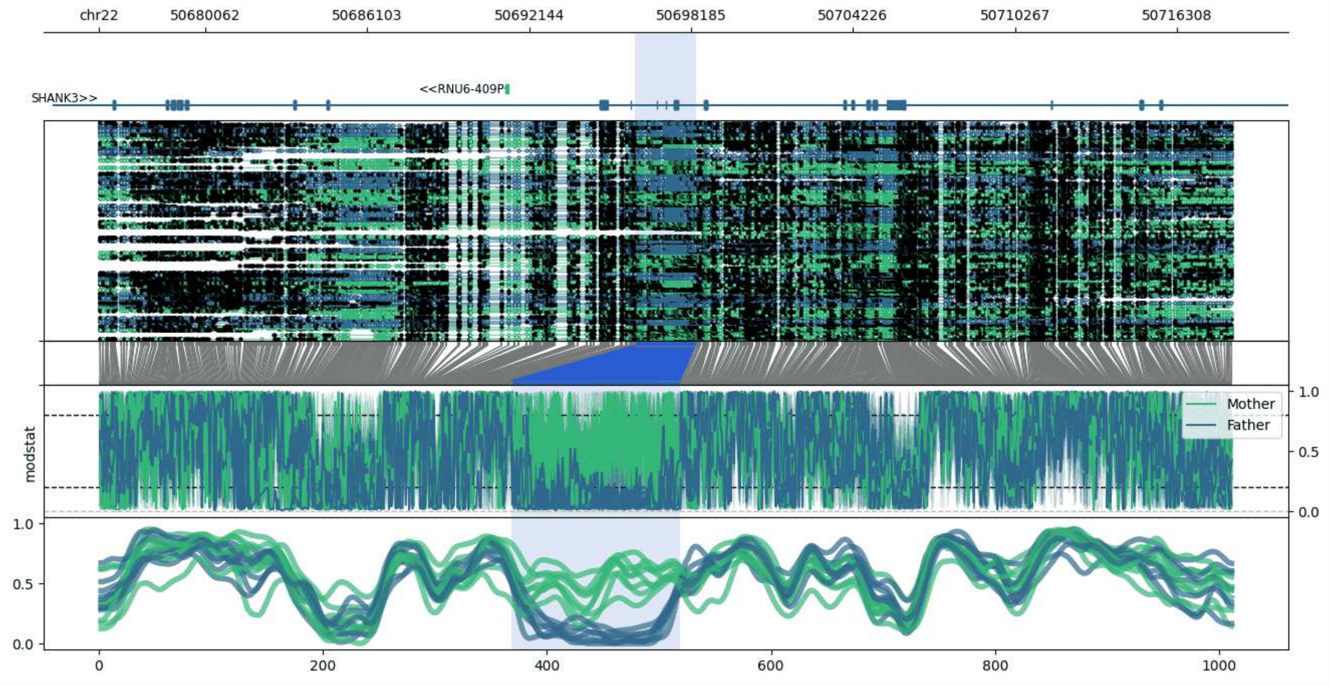


**chr22:50676077-50718368:** Region with 1 DMR containing genes: AC000050.1, RF00019, RNU6-409P, SHANK3.

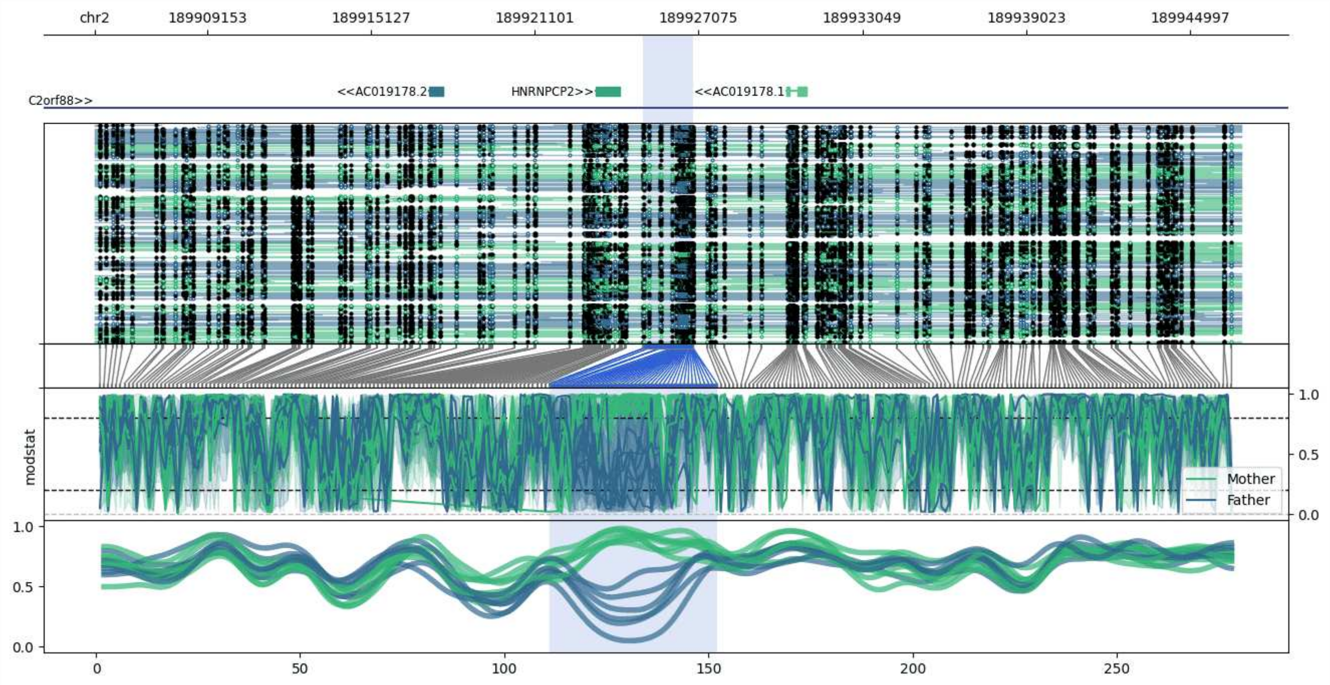


**chr2:189905059-189946877:** Region with 1 DMR containing genes: AC019178.1, AC019178.2, C2orf88, HNRNPCP2.

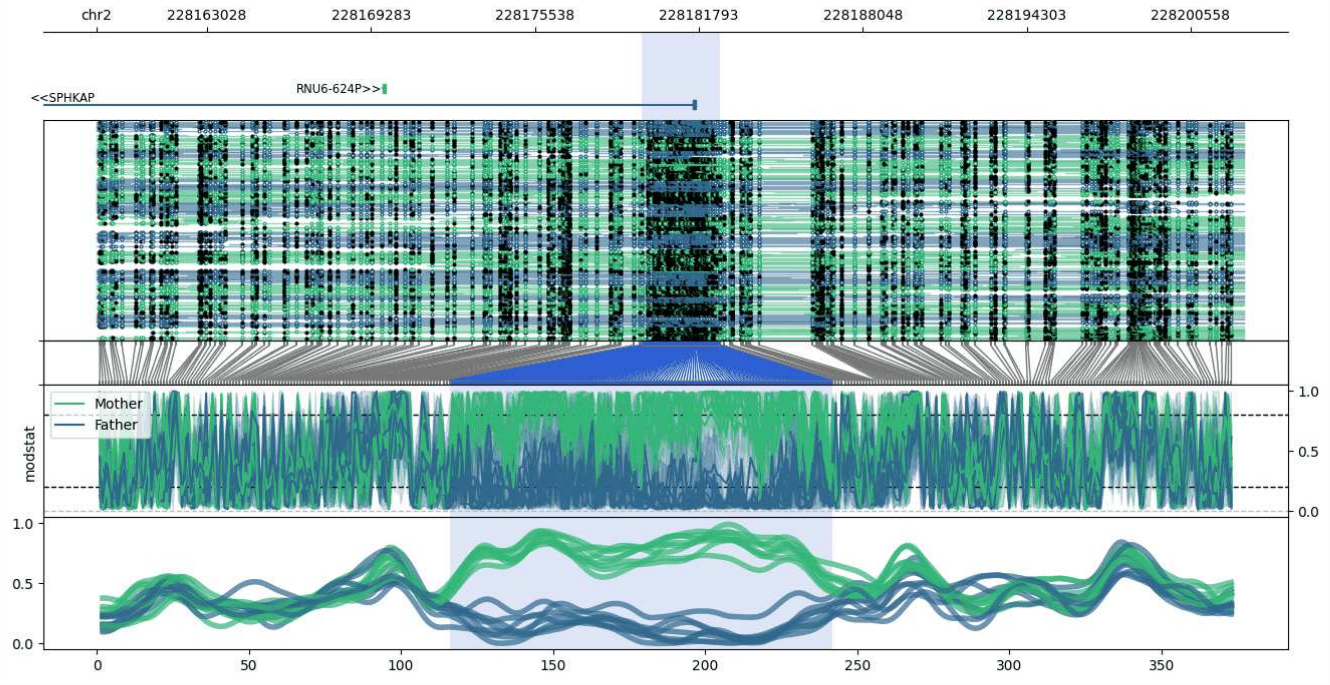


**chr2:228158814-228202605:** Region with 1 DMR containing genes: RNU6-624P, SPHKAP.

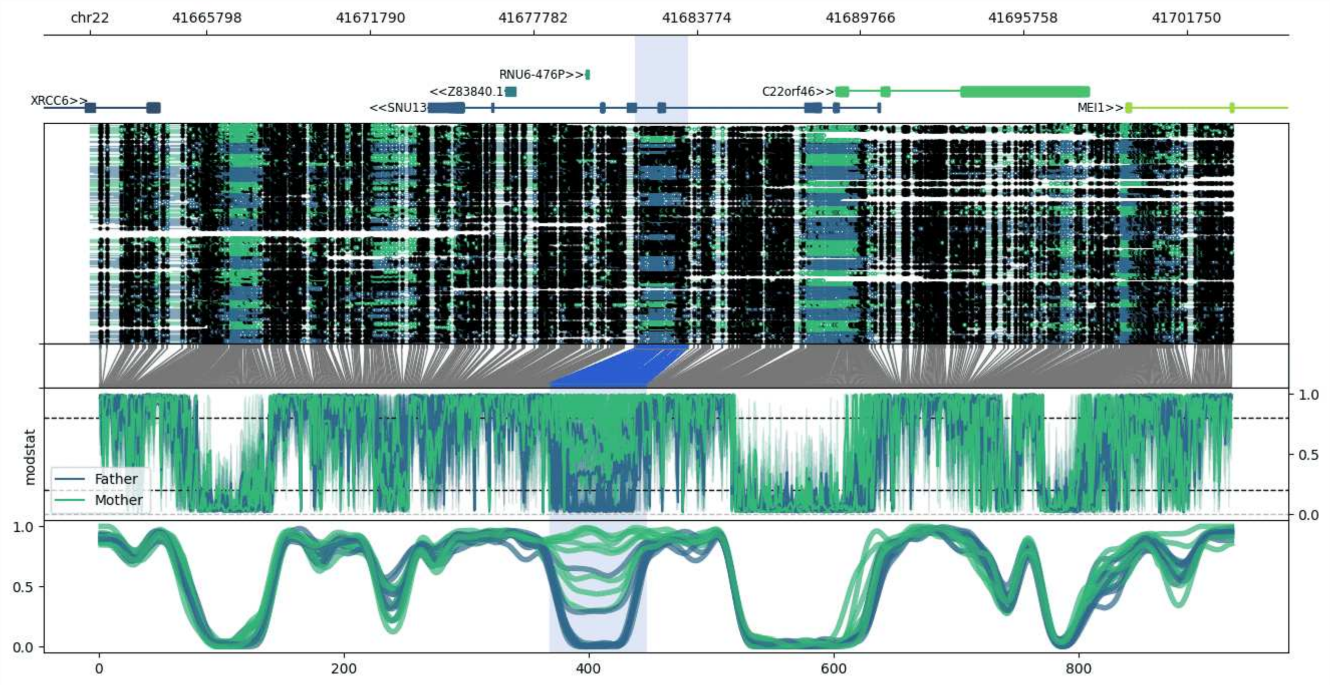


**chr22:41661511-41703457:** Region with 1 DMR containing genes: C22orf46, HMGN2P10, MEI1, RNU6-476P, SNU13, XRCC6, Z83840.1.

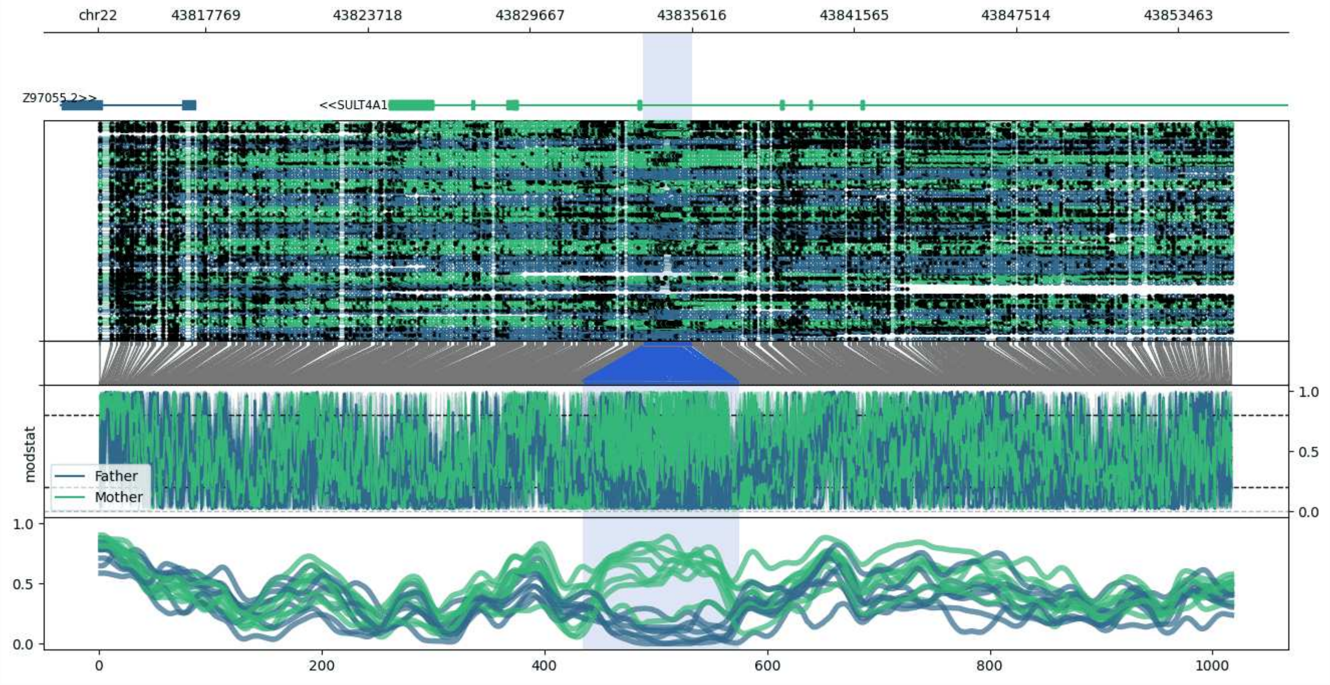


**chr22:43813836-43855484:** Region with 1 DMR containing genes: EFCAB6, SULT4A1, Z97055.2.

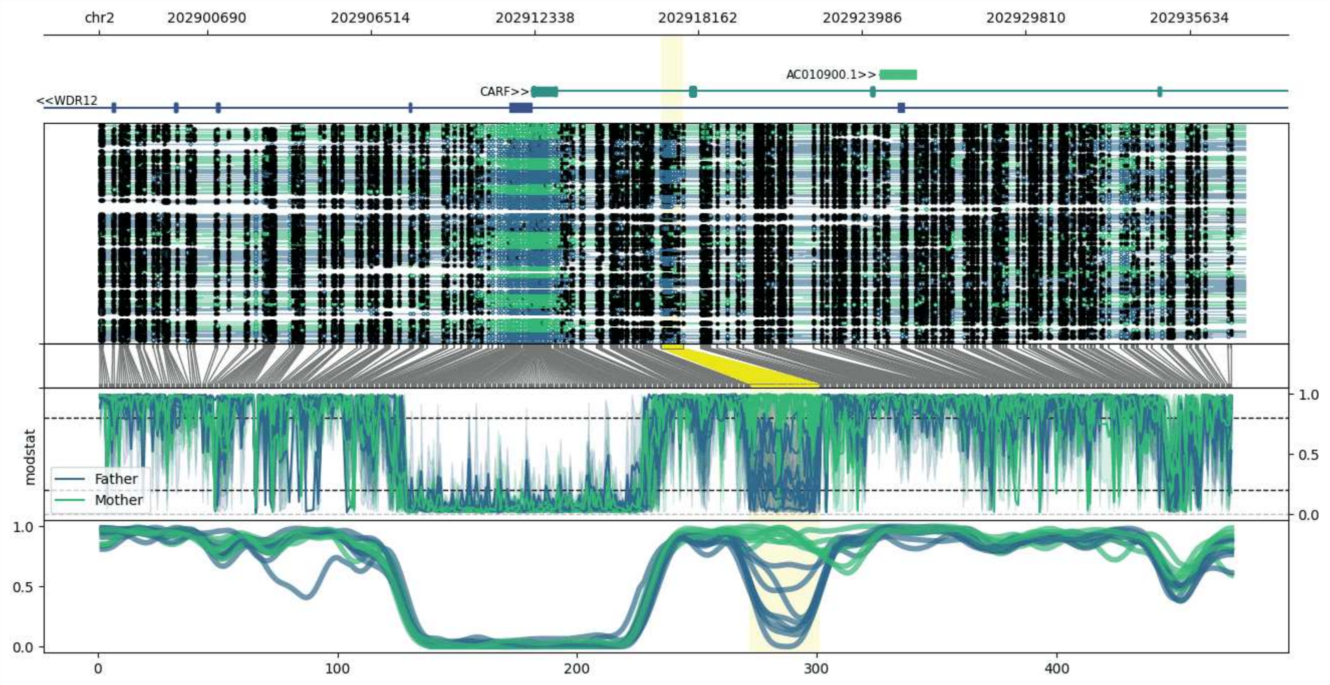


**chr2:202896849-202937620:** Region with 1 DMR containing genes: AC010900.1, CARF, WDR12.

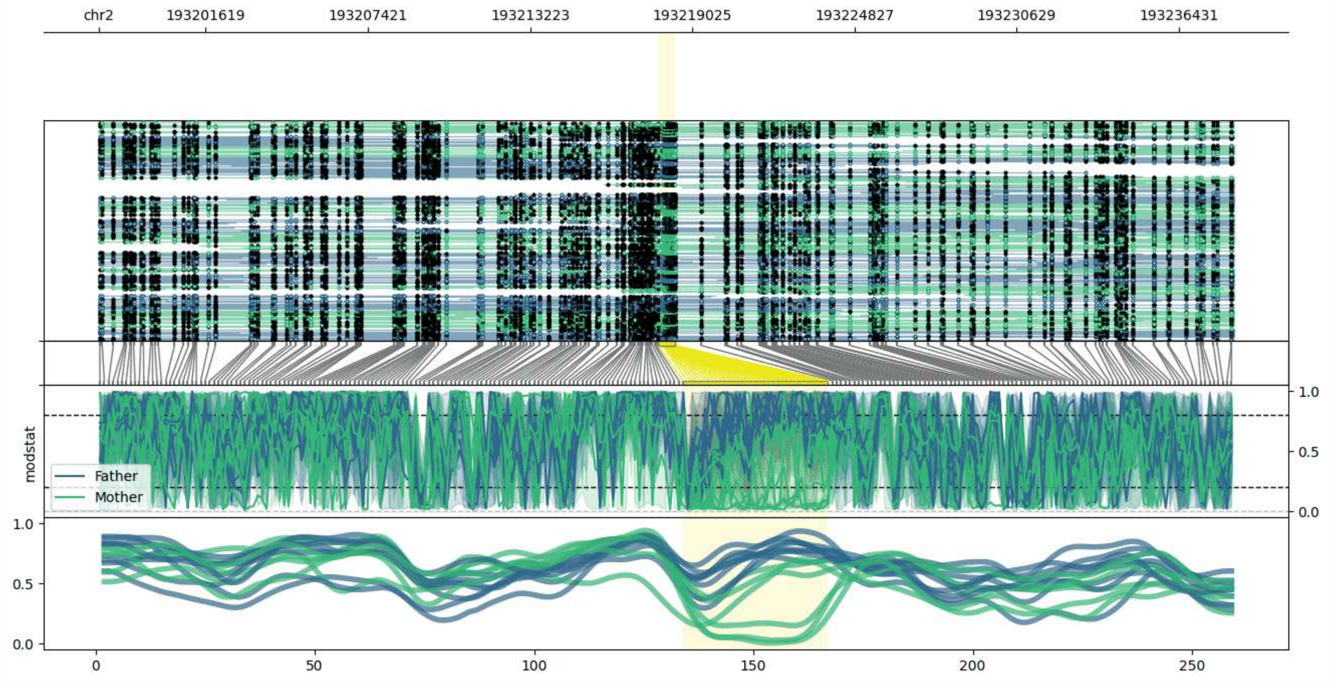


**chr2:193197795-193238410:** Region with 1 DMR containing no genes.

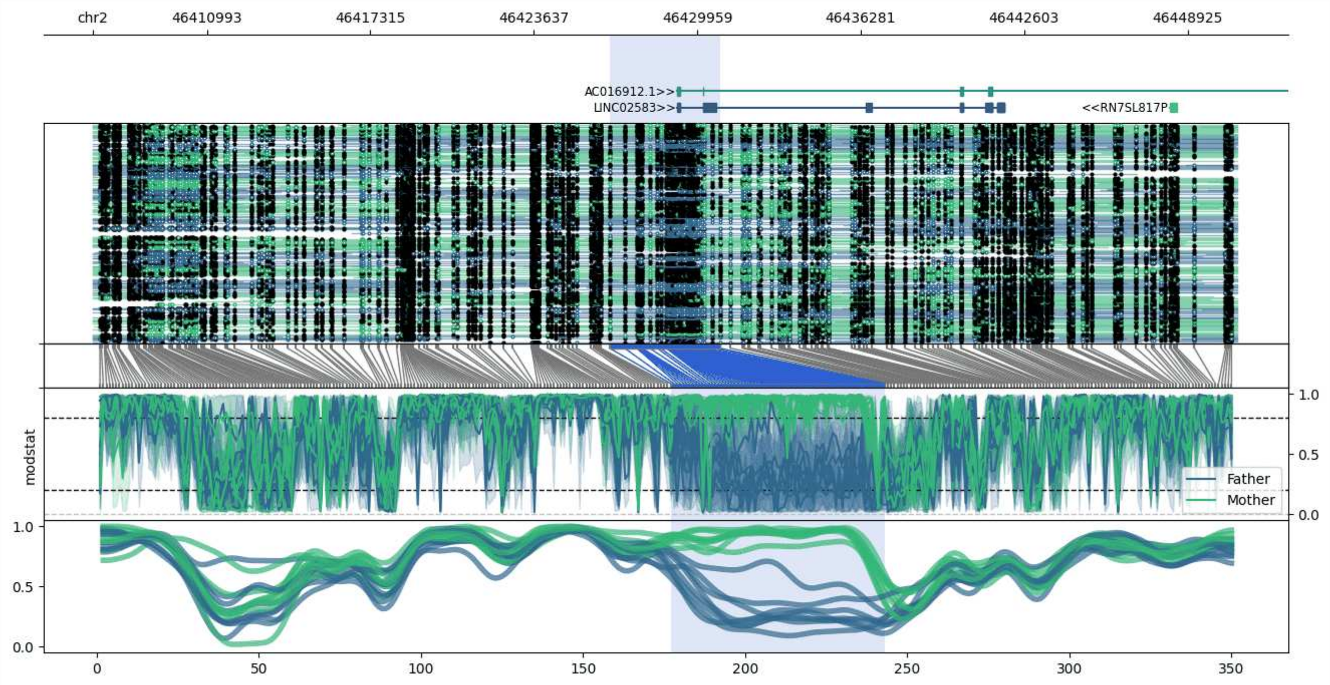


**chr2:46406571-46450829:** Region with 1 DMR containing genes: AC016912.1, LINC02583, RN7SL817P.

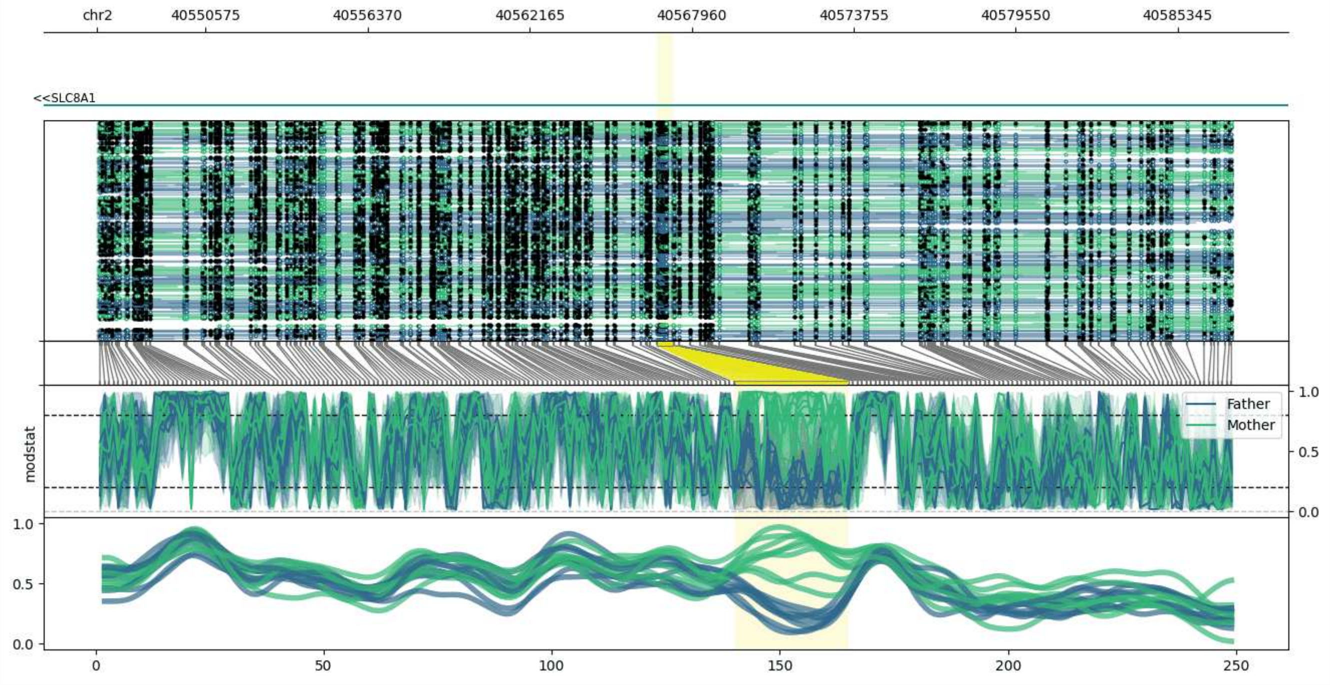


**chr2:40546707-40587278:** Region with 1 DMR containing genes: AC007317.1, AC007877.1, SLC8A1.

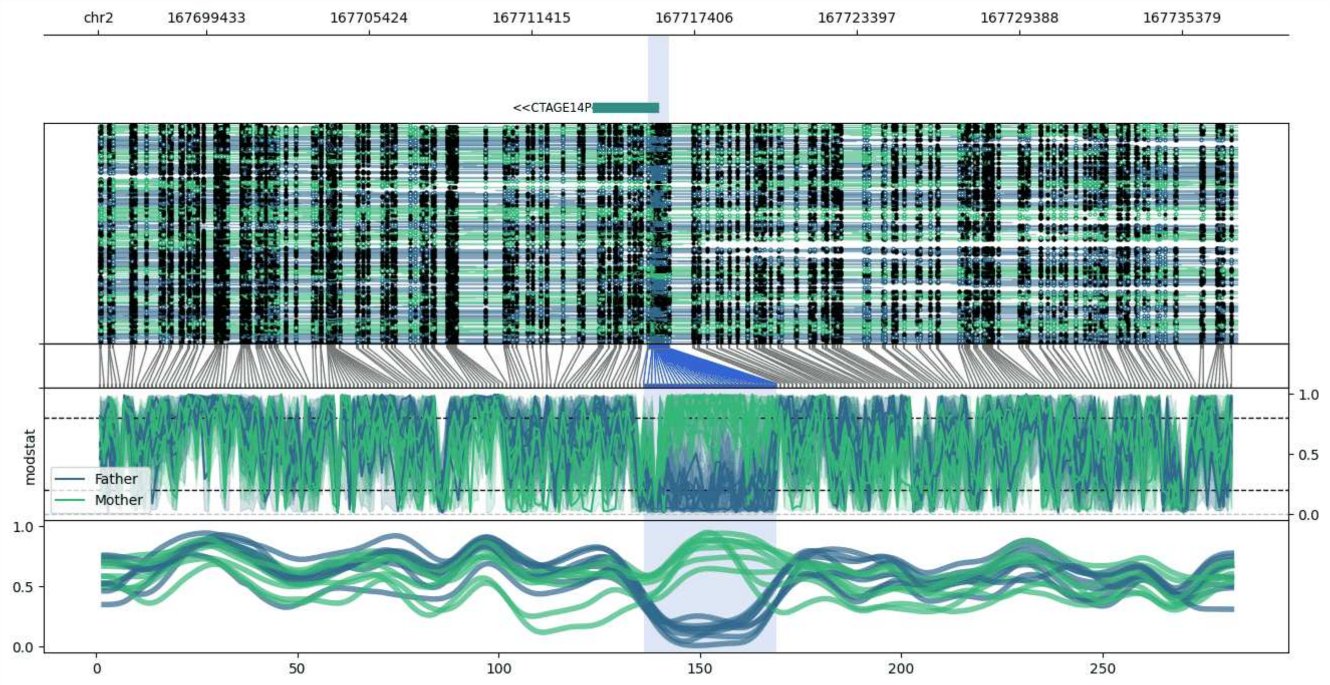


**chr2:167695457-167737396:** Region with 1 DMR containing genes: CTAGE14P.

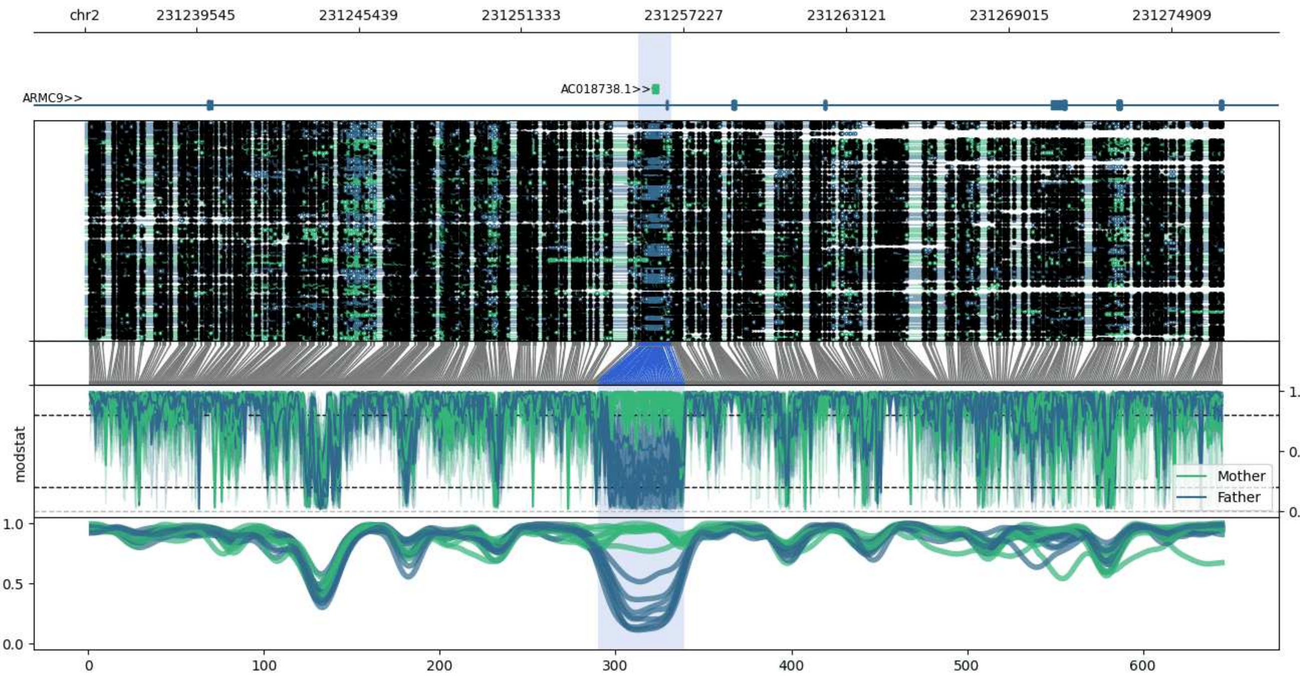


**chr2:231235505-231276764:** Region with 1 DMR containing genes: AC018738.1, ARMC9.

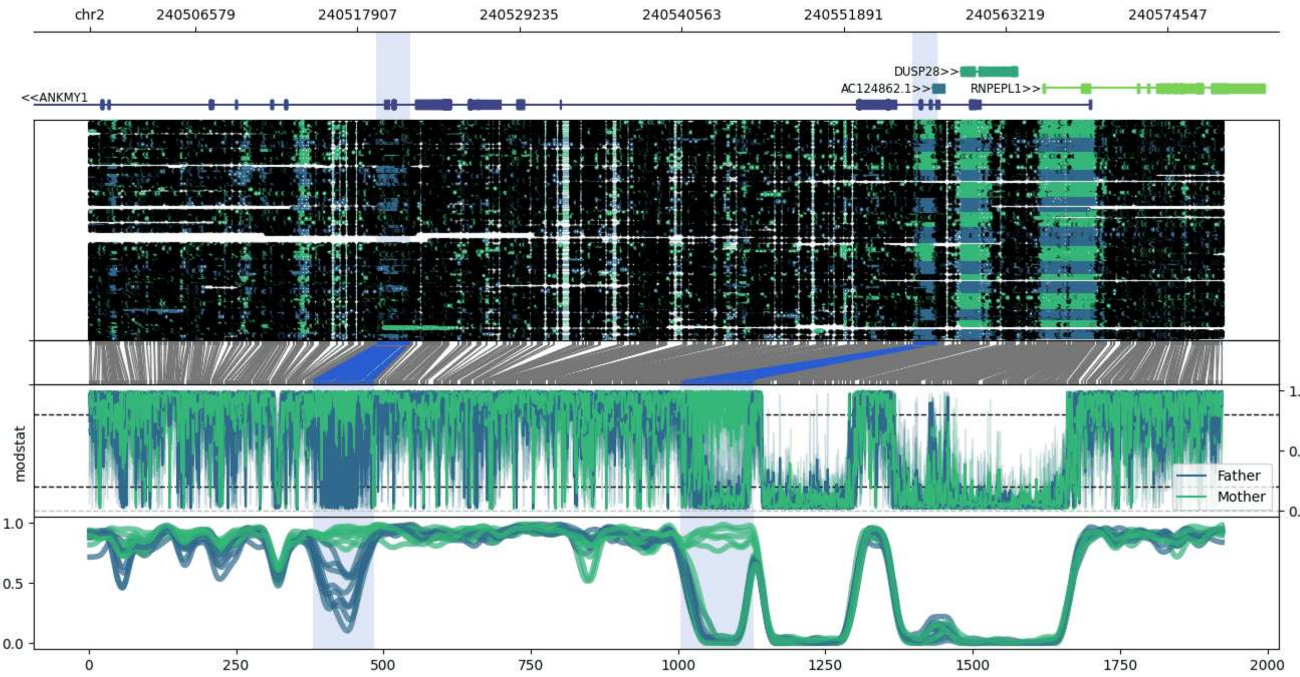


**chr2:240499183-240578483:** Region with 2 DMRs containing genes: AC124862.1, ANKMY1, CAPN10, CAPN10-DT, DUSP28, RNPEPL1.

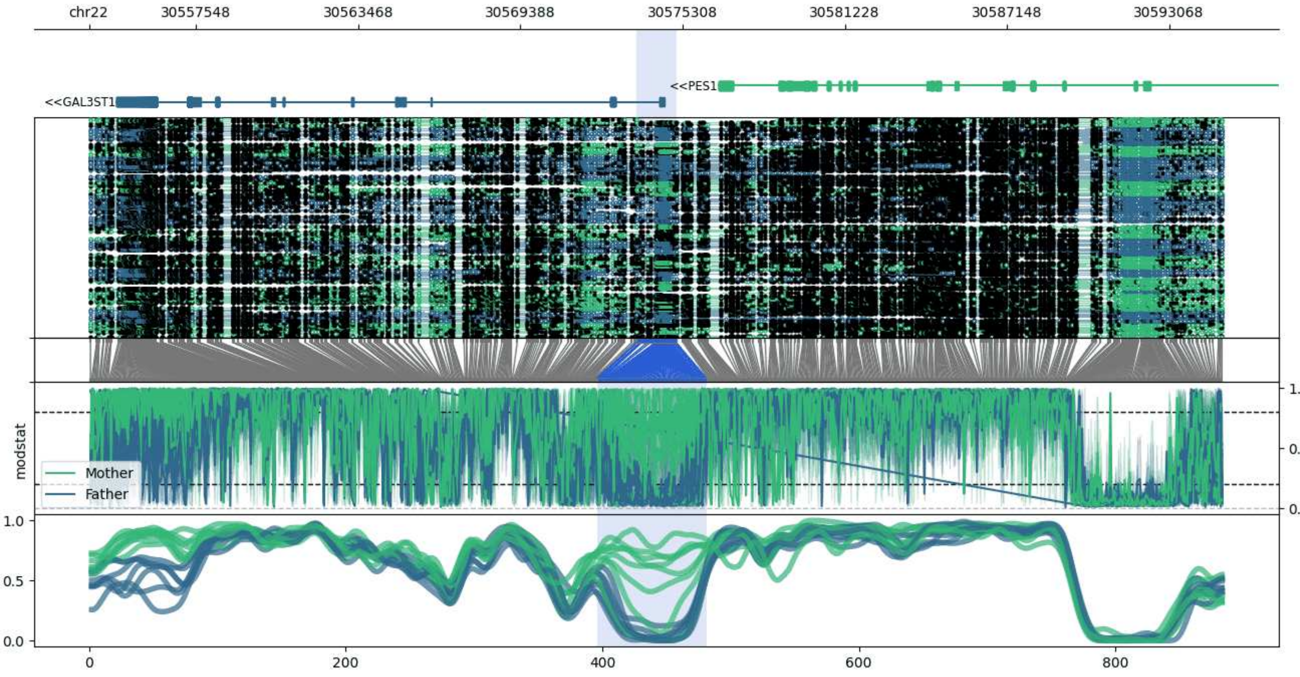


**chr22:30553637-30595078:** Region with 1 DMR containing genes: AC005006.1, GAL3ST1, PES1, SEC14L6, SIRPAP1.

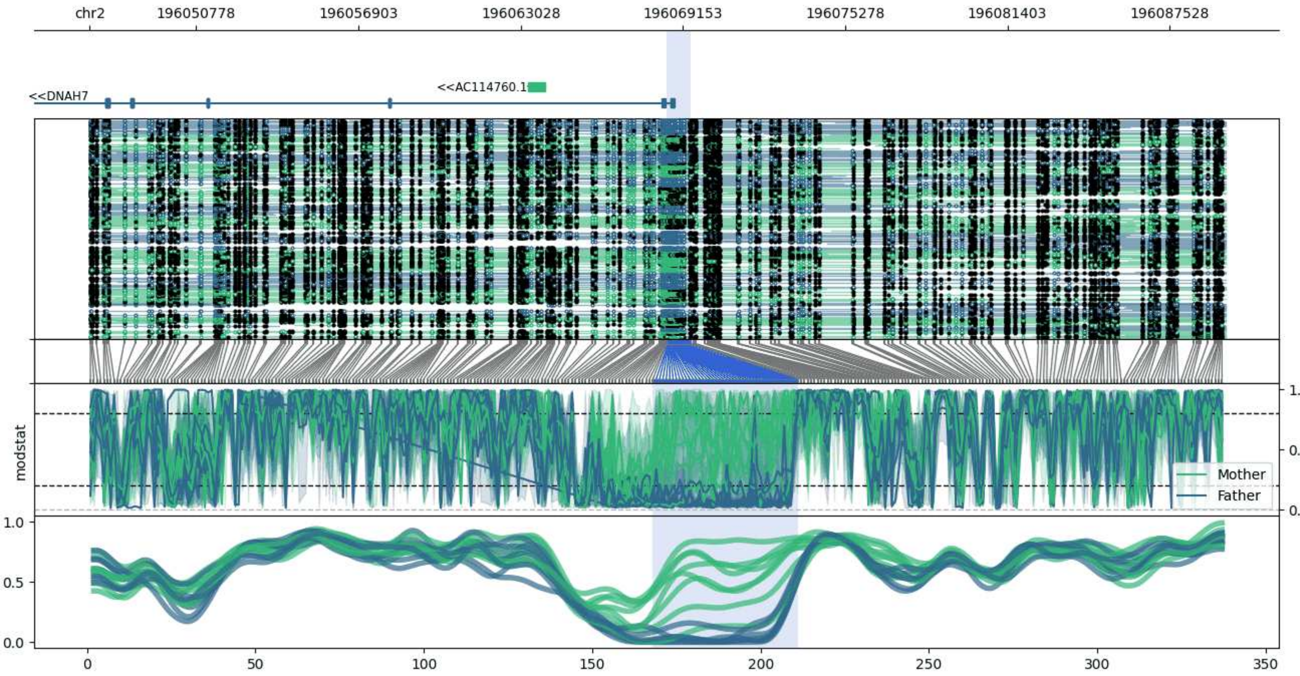


**chr2:196046765-196089646:** Region with 1 DMR containing genes: AC114760.1, DNAH7.

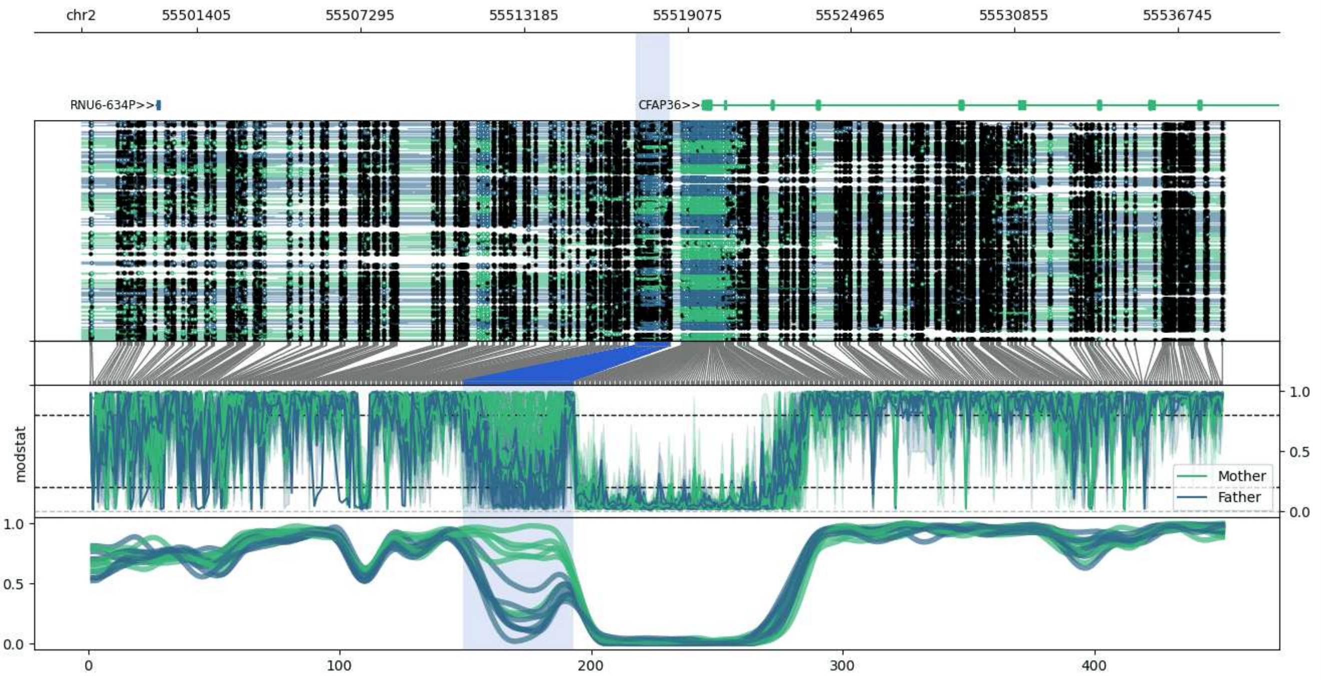


**chr2:55497219-55538449:** Region with 1 DMR containing genes: CFAP36, PPP4R3B, RNU6-634P.

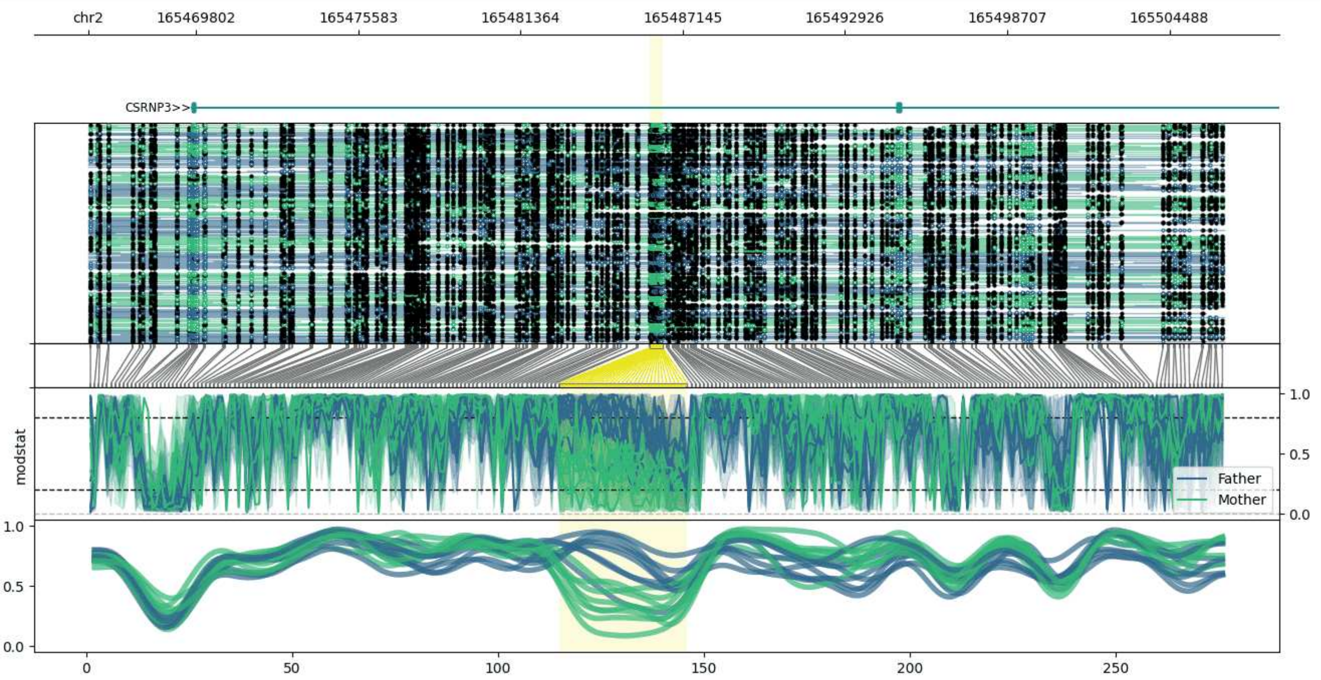


**chr2:165465951-165506422:** Region with 1 DMR containing genes: CSRNP3.

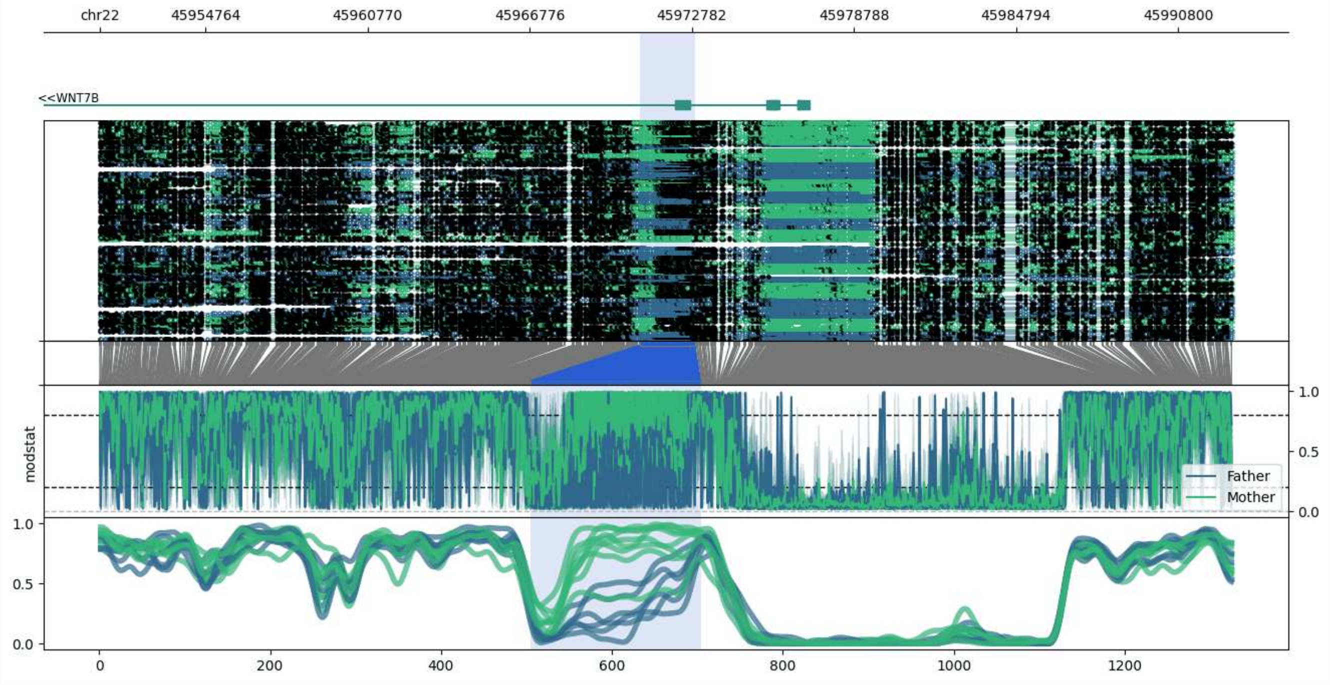


**chr22:45950846-45992892:** Region with 1 DMR containing genes: WNT7B.

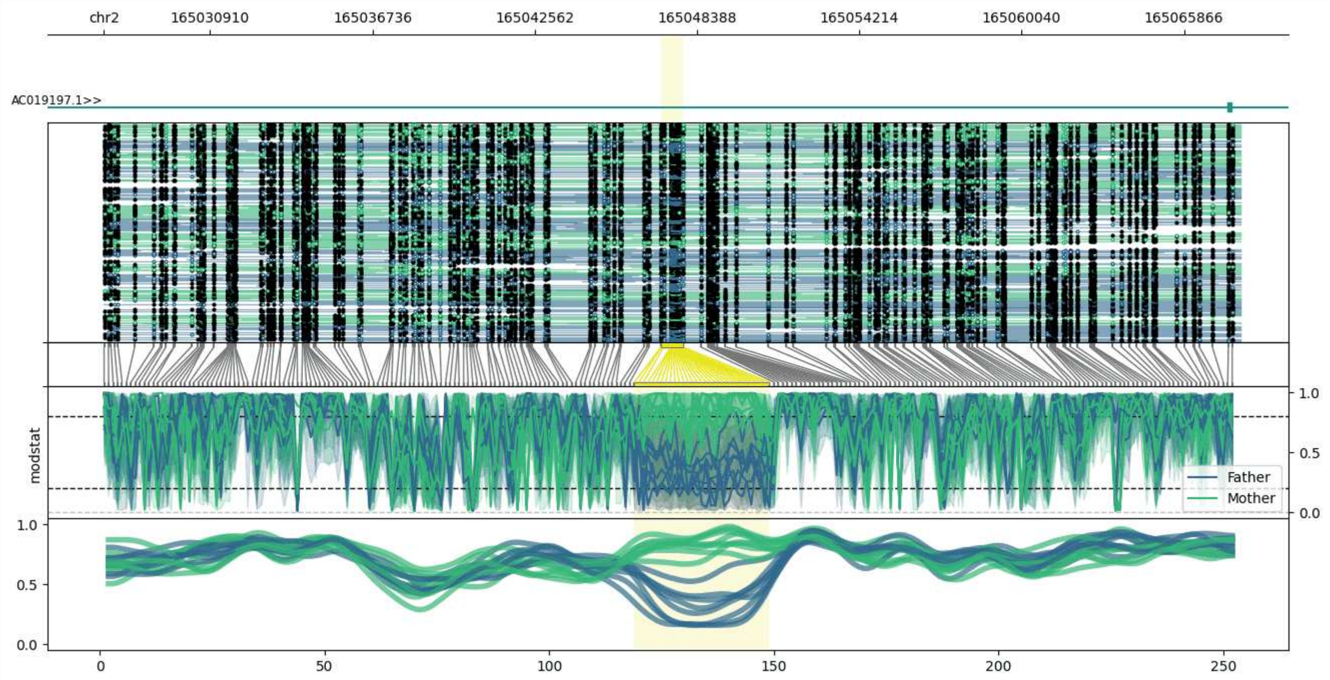


**chr2:165027097-165067881:** Region with 1 DMR containing genes: AC019197.1.

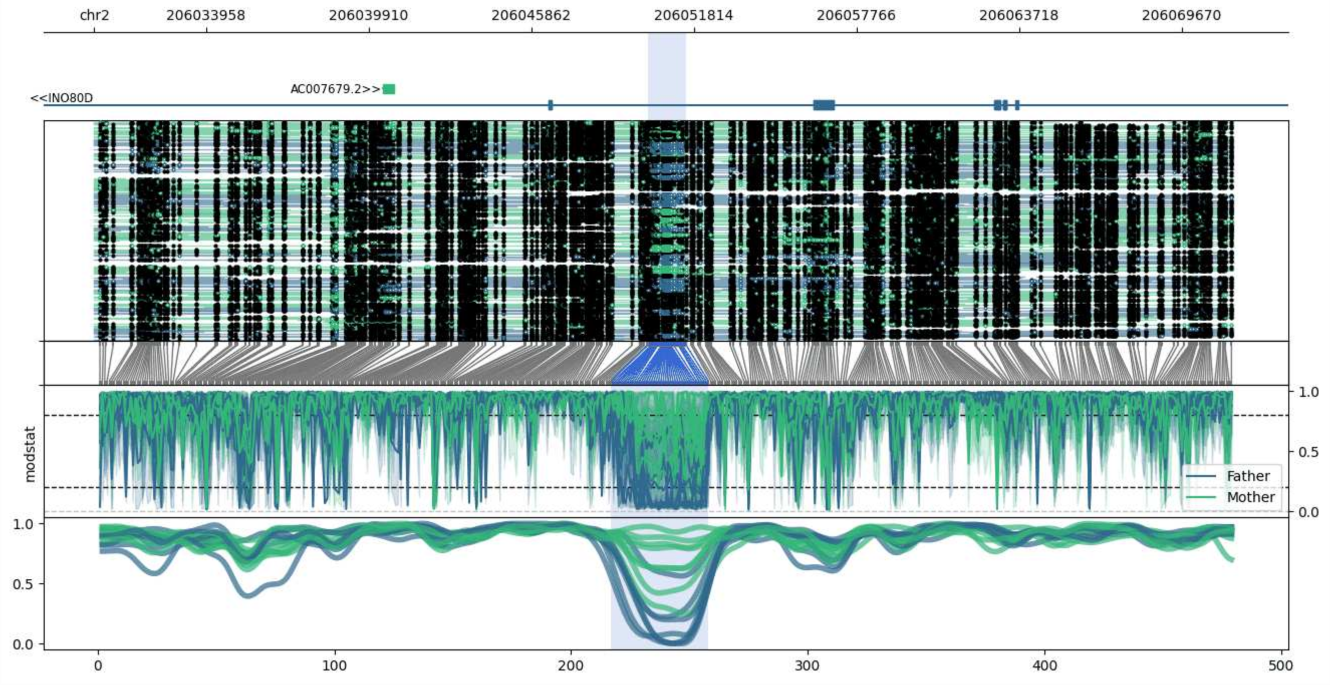


**chr2:206029869-206071539:** Region with 1 DMR containing genes: AC007679.2, INO80D, RF00019.

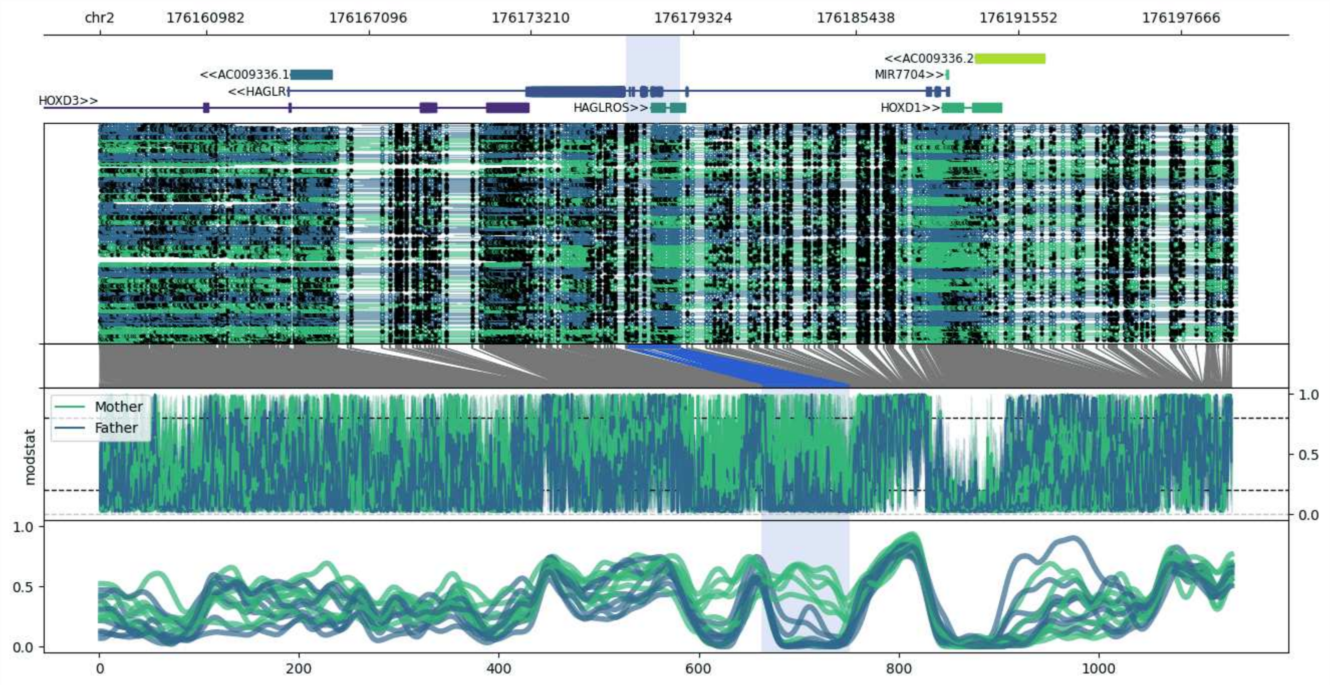


**chr2:176156992-176199793:** Region with 1 DMR containing genes: AC009336.1, AC009336.2, AC016739.1, HAGLR, HAGLROS, HOXD1, HOXD3, HOXD4, MIR10B, MIR7704.

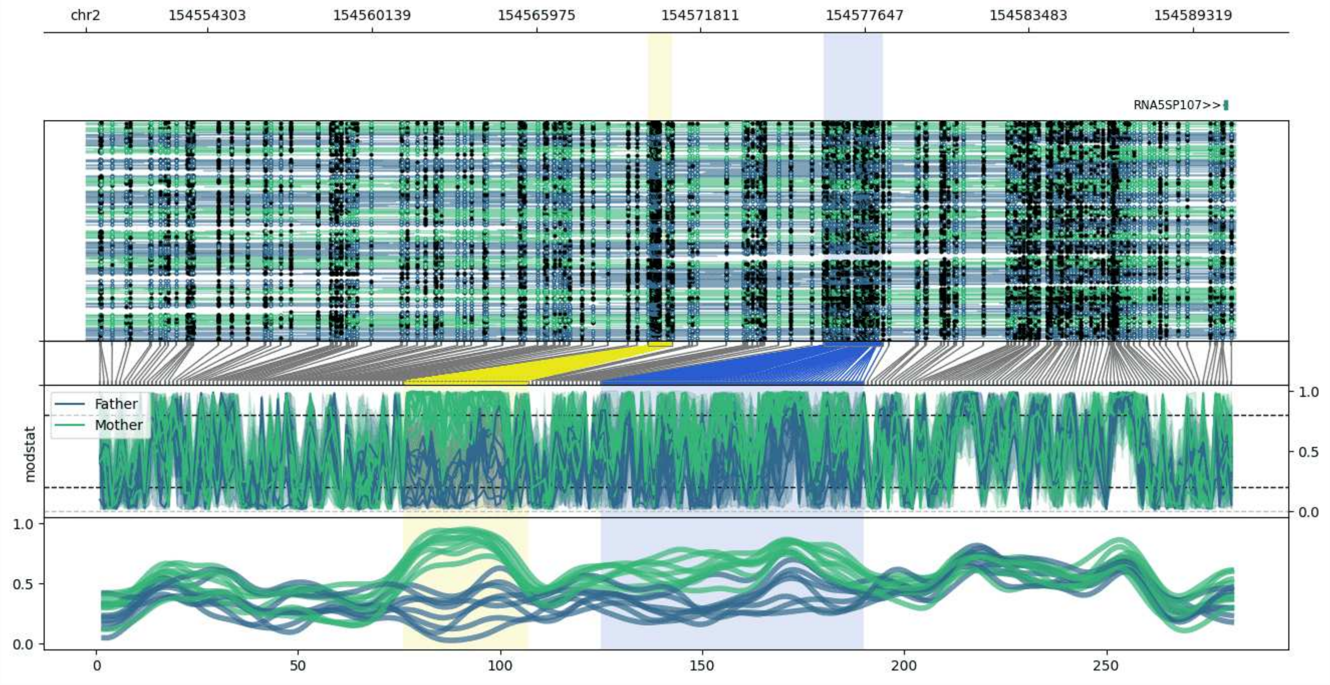


**chr2:154549962-154590815:** Region with 2 DMRs containing genes: RNA5SP107.

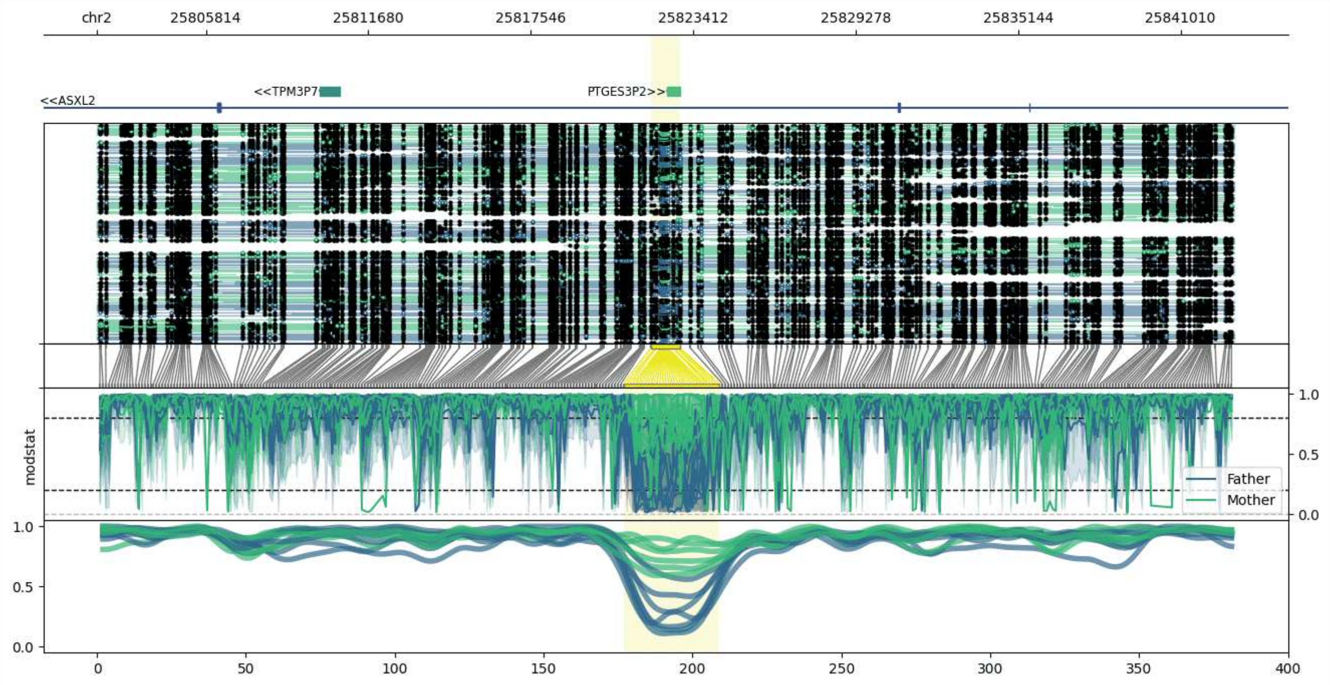


**chr2:25801877-25842941:** Region with 1 DMR containing genes: ASXL2, PTGES3P2, TPM3P7.

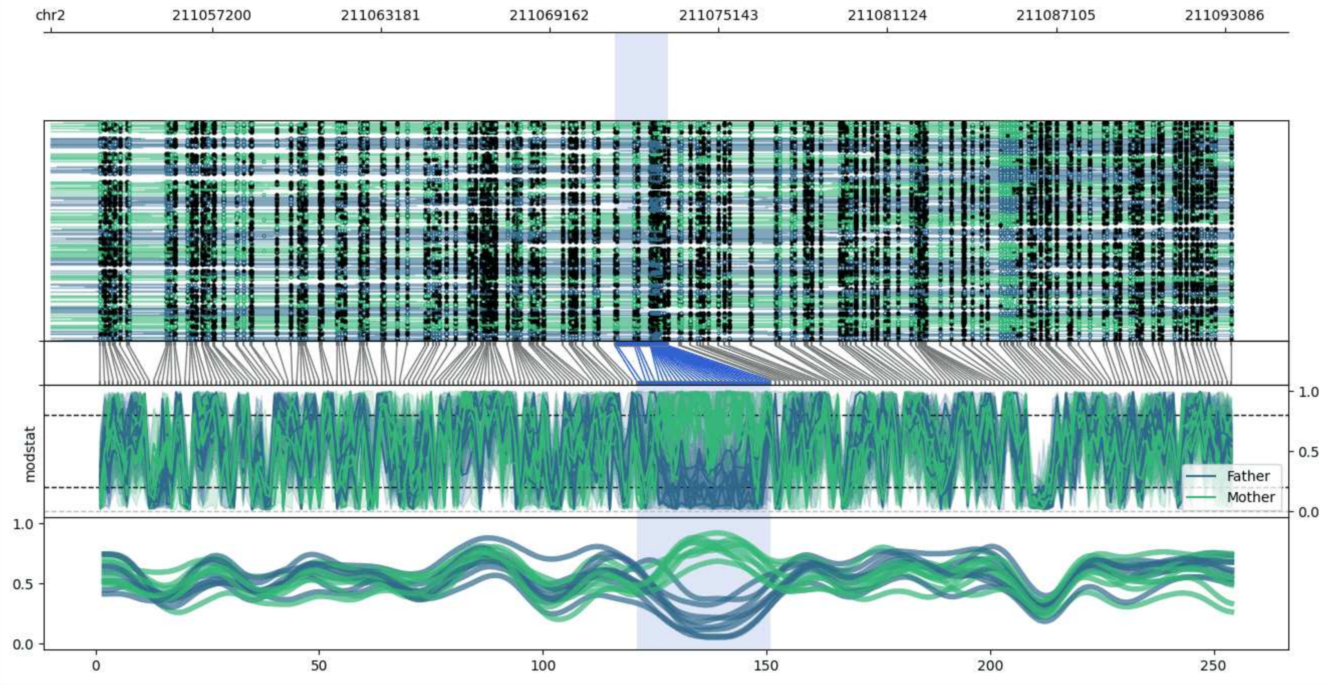


**chr2:211051472-211093345:** Region with 1 DMR containing no genes.

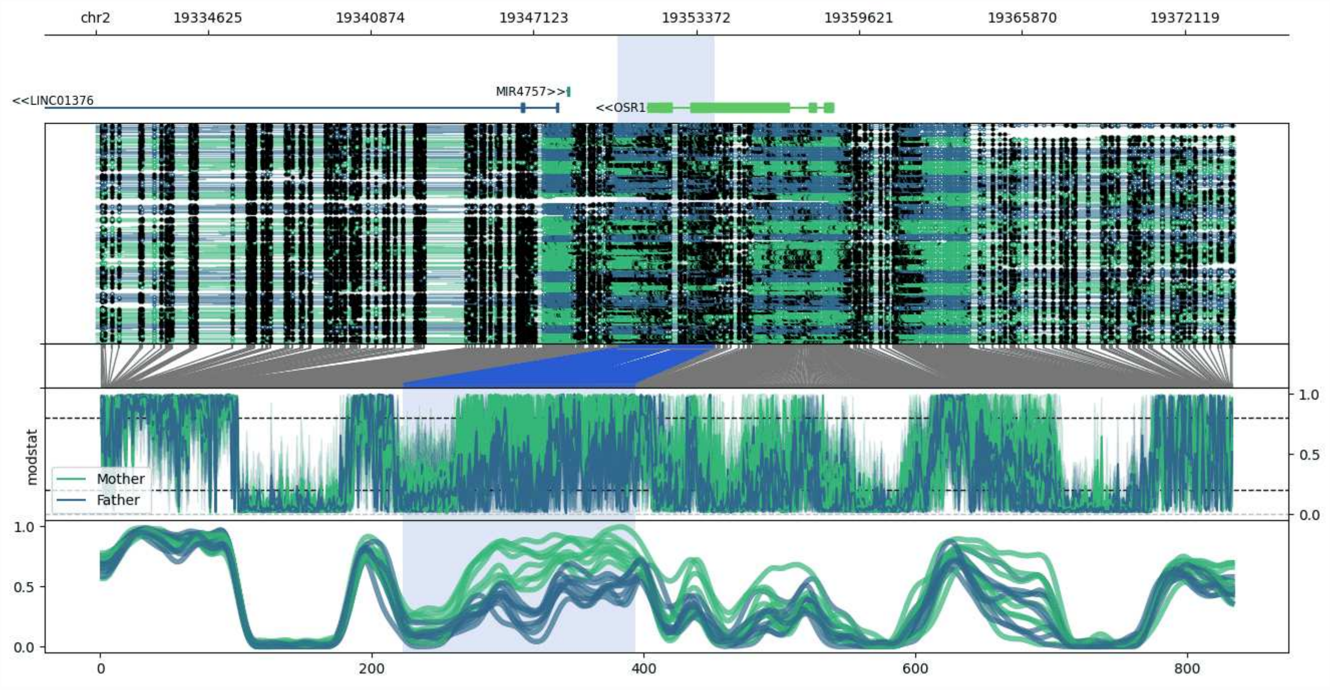


**chr2:19330325-19374071:** Region with 1 DMR containing genes: LINC01376, MIR4757, OSR1.

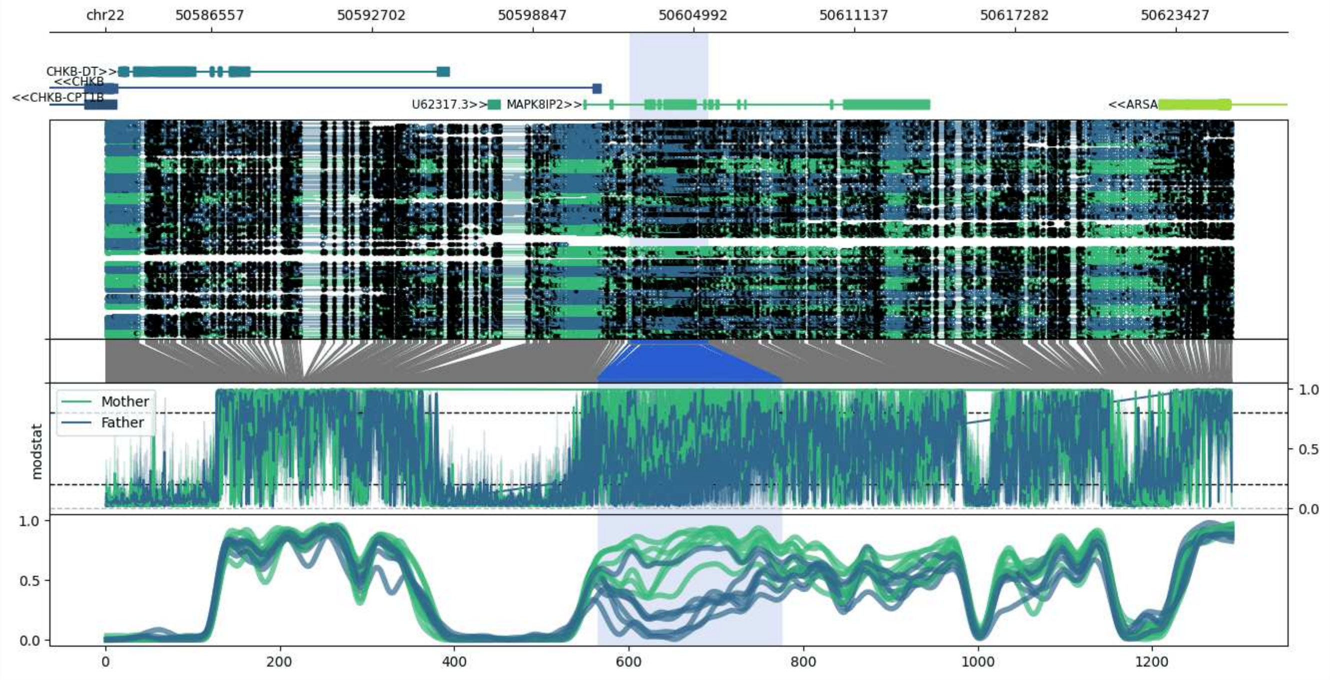


**chr22:50582538-50625555:** Region with 1 DMR containing genes: ARSA, CHKB, CHKB-CPT1B, CHKB-DT, CPT1B, MAPK8IP2, U62317.3.

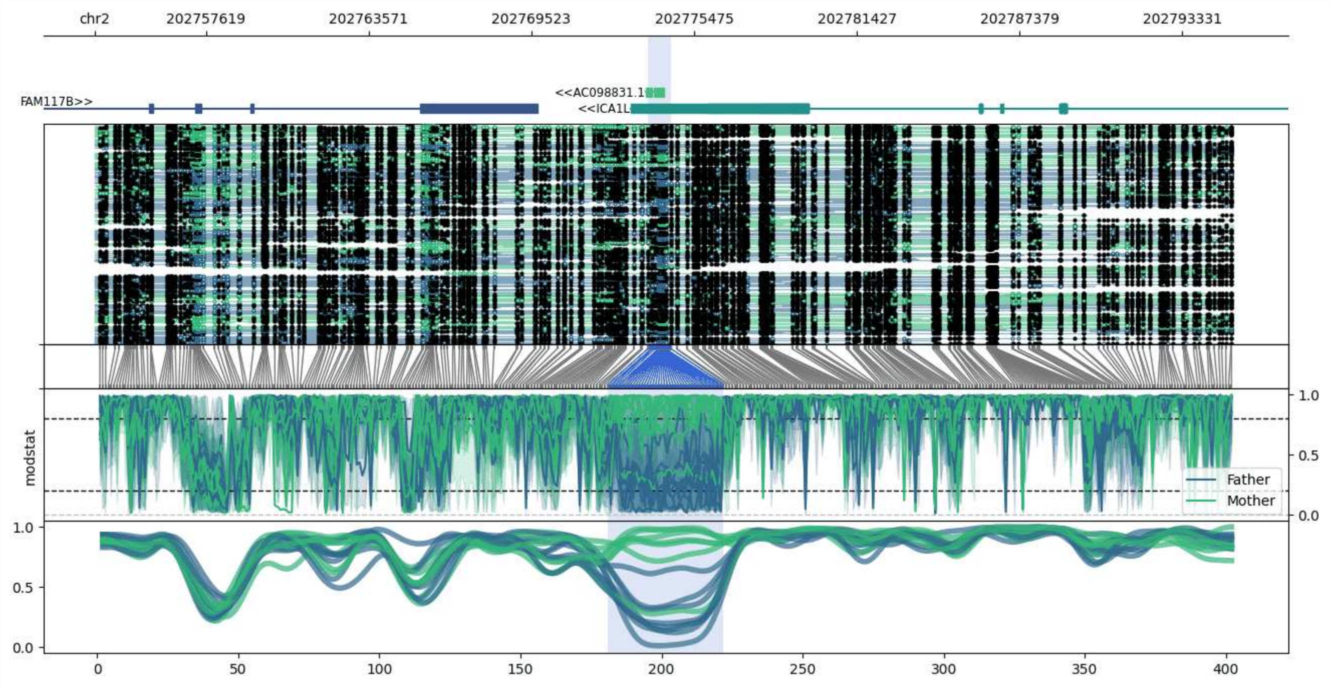


**chr2:202753541-202795207:** Region with 1 DMR containing genes: AC098831.1, FAM117B, ICA1L.

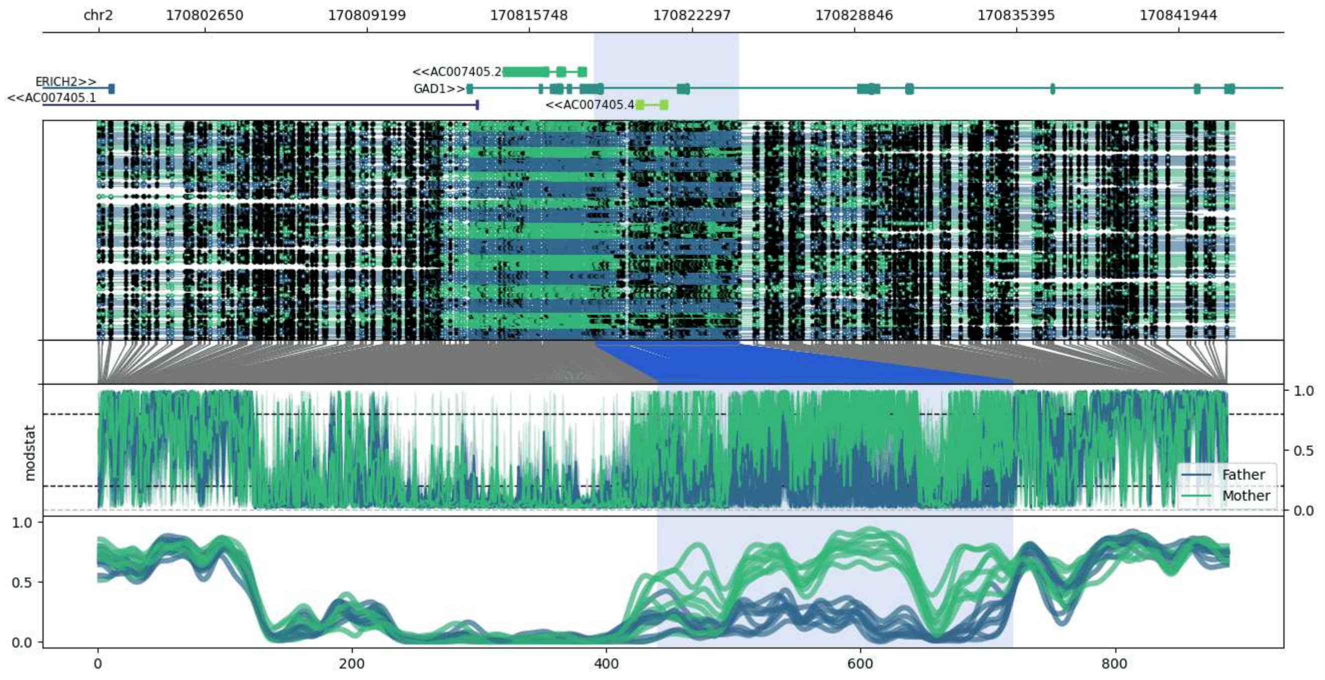


**chr2:170798357-170844204:** Region with 1 DMR containing genes: AC007405.1, AC007405.2, AC007405.4, ERICH2, GAD1.

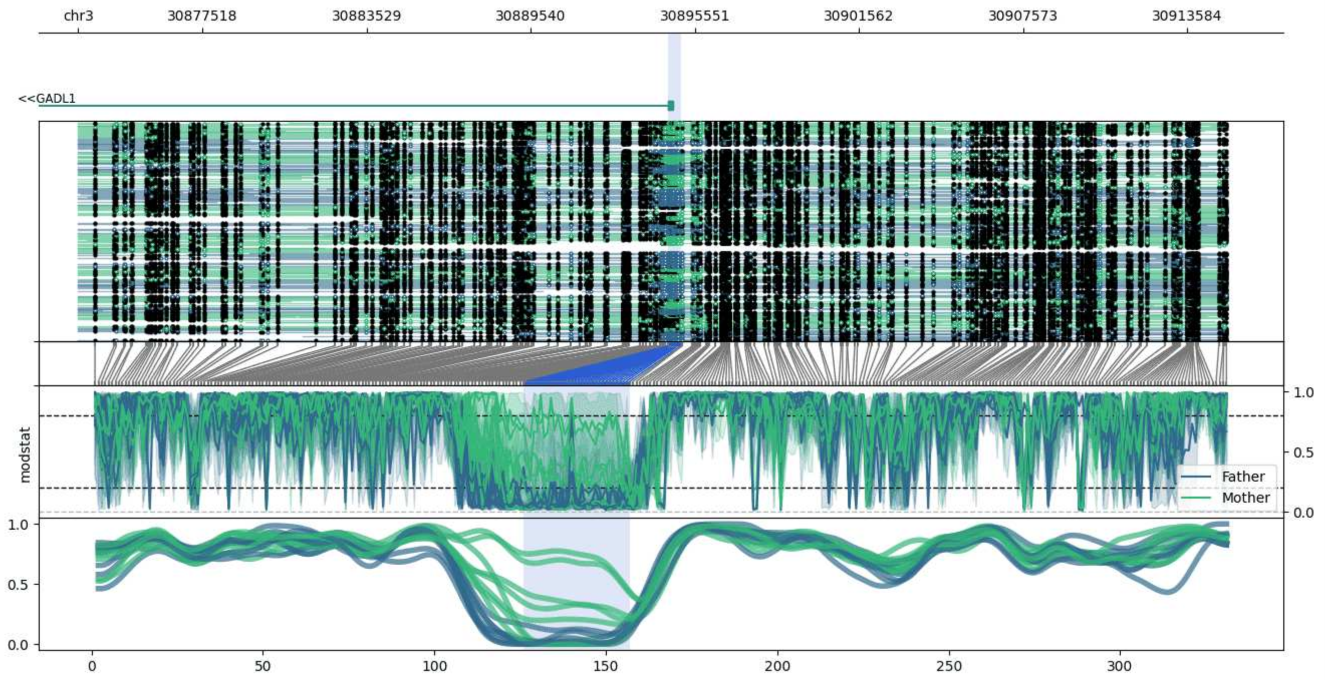


**chr3:30872959-30915039:** Region with 1 DMR containing genes: GADL1.

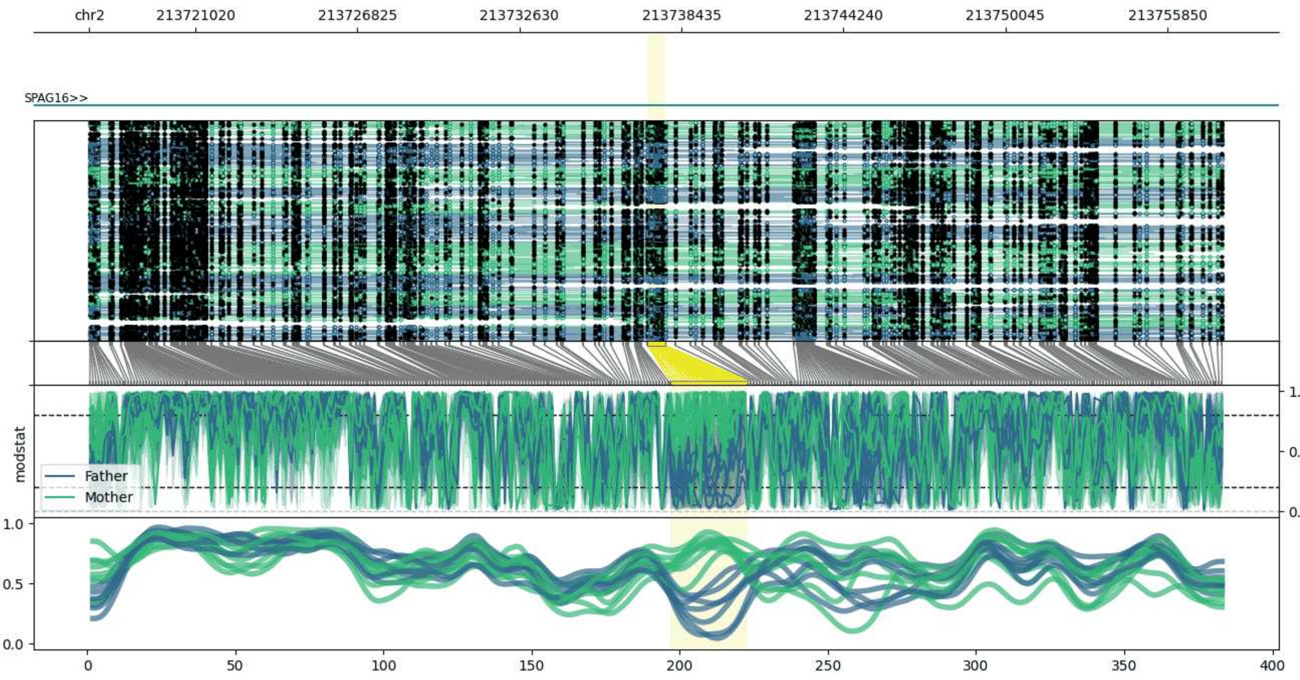


**chr2:213717219-213757856:** Region with 1 DMR containing genes: MIR4438, SPAG16.

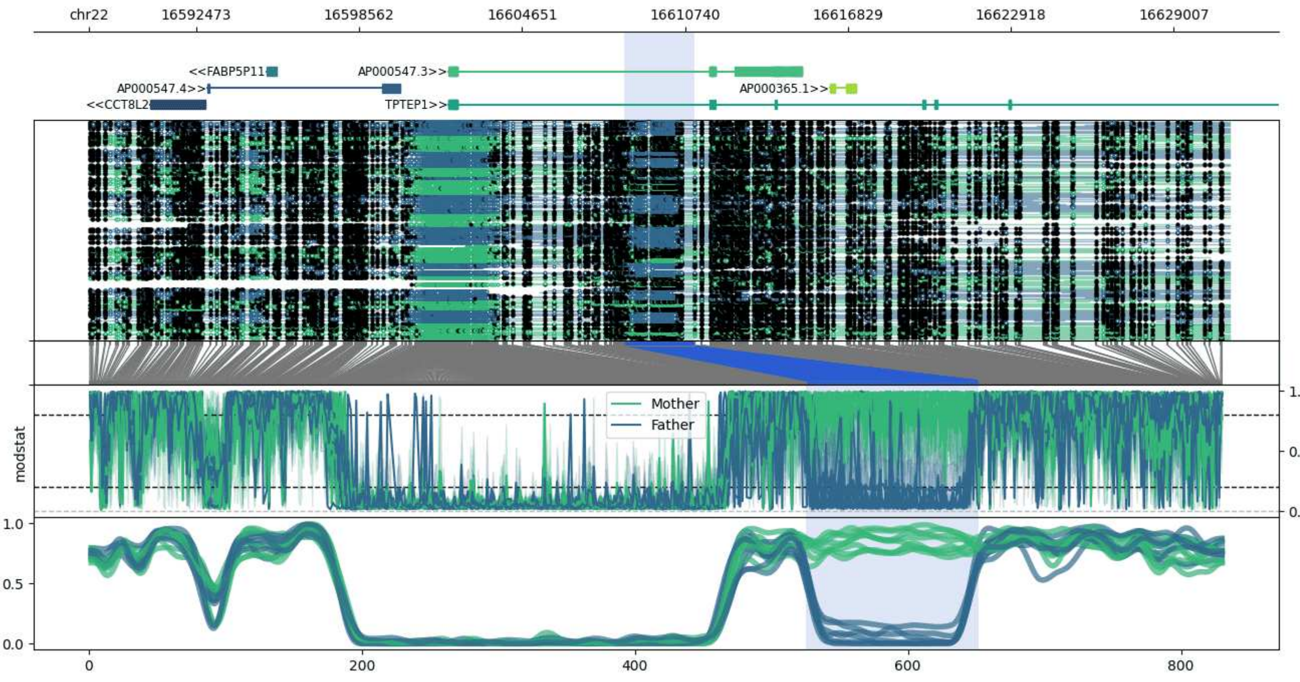


**chr22:16588452-16631080:** Region with 1 DMR containing genes: AP000365.1, AP000547.3, AP000547.4, CCT8L2, FABP5P11, KCNMB3P1, TPTEP1.

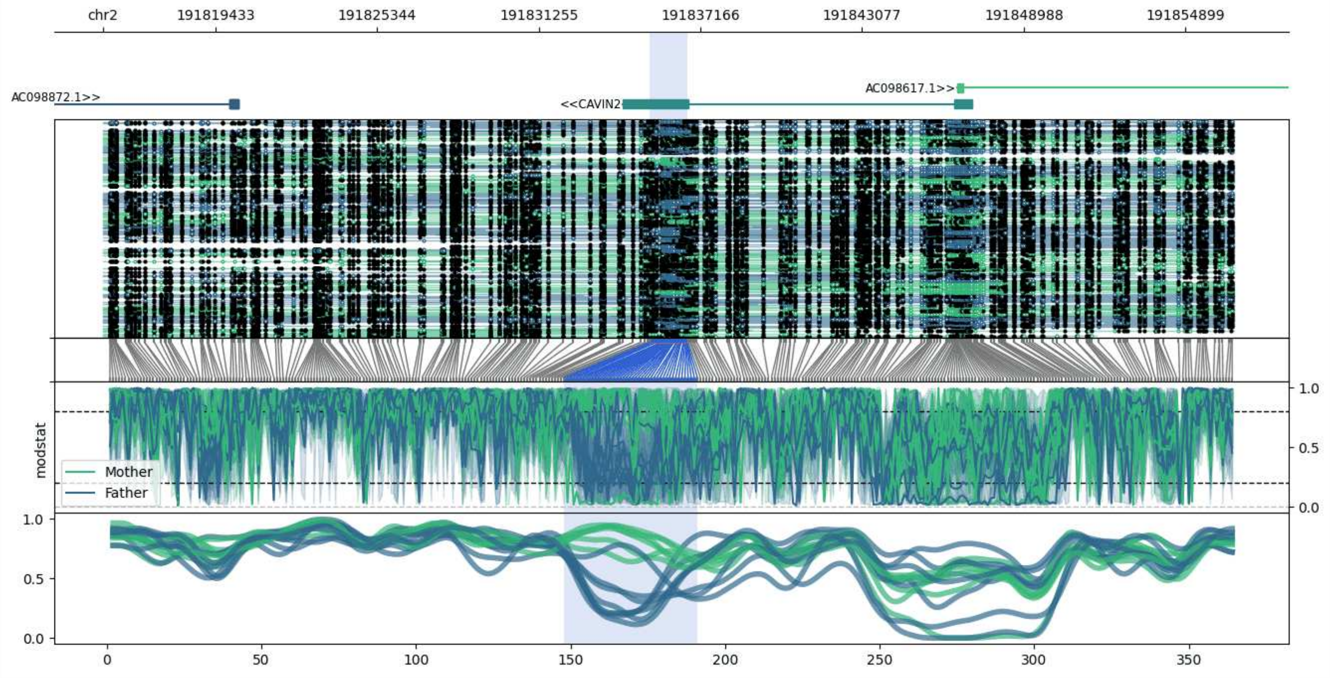


**chr2:191815309-191856692:** Region with 1 DMR containing genes: AC098617.1, AC098872.1, CAVIN2.

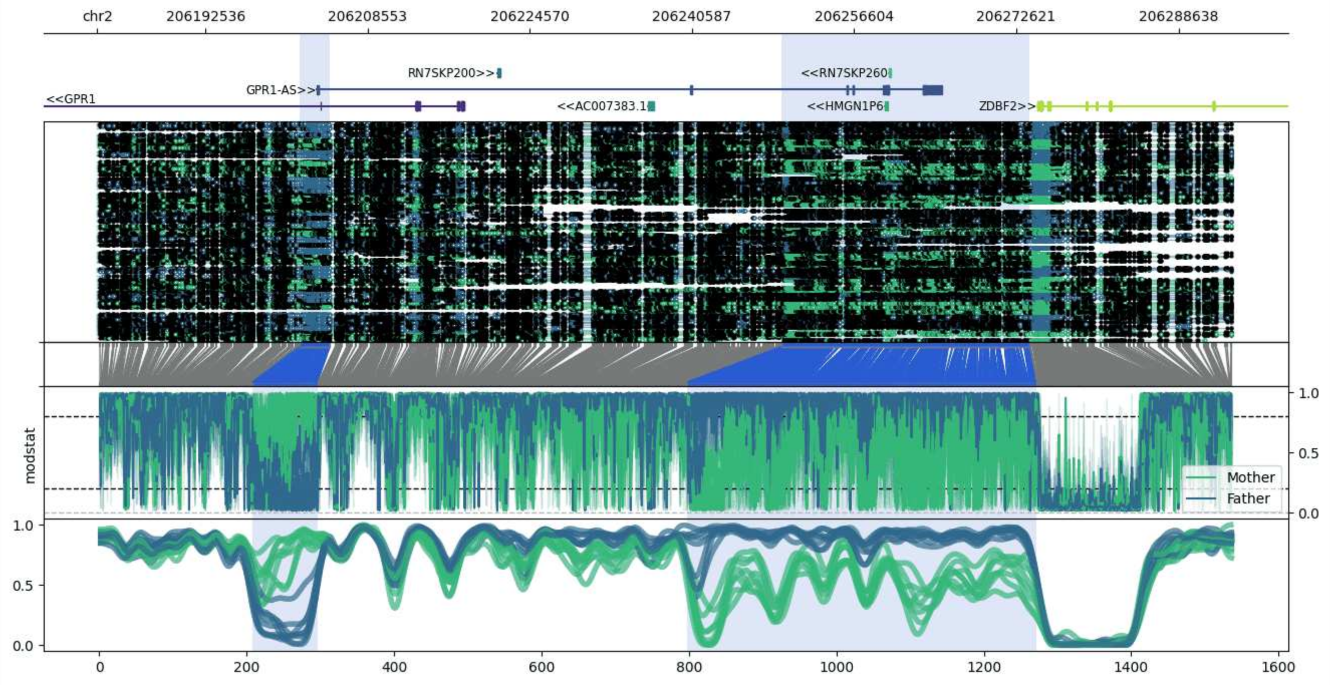


**chr2:206181827-206293948:** Region with 2 DMRs containing genes: AC007383.1, GPR1, GPR1-AS, HMGN1P6, RN7SKP200, RN7SKP260, ZDBF2.

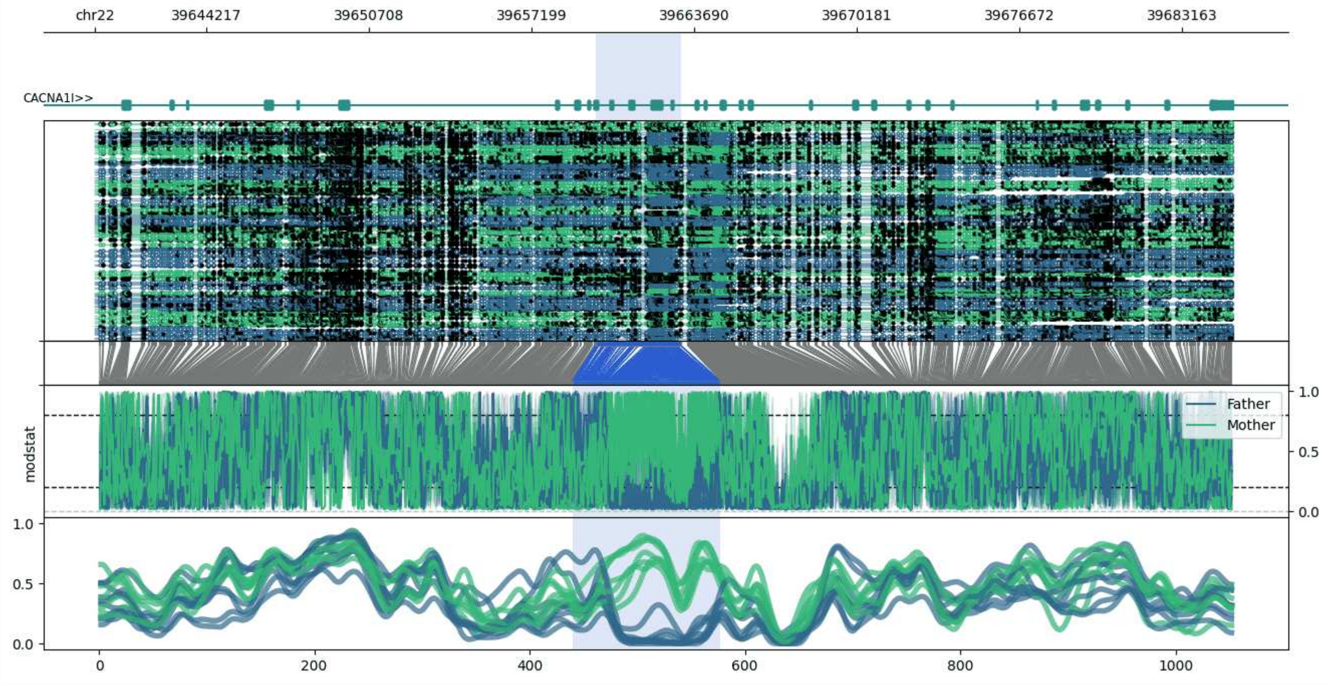


**chr22:39639776-39685218:** Region with 1 DMR containing genes: CACNA1I.

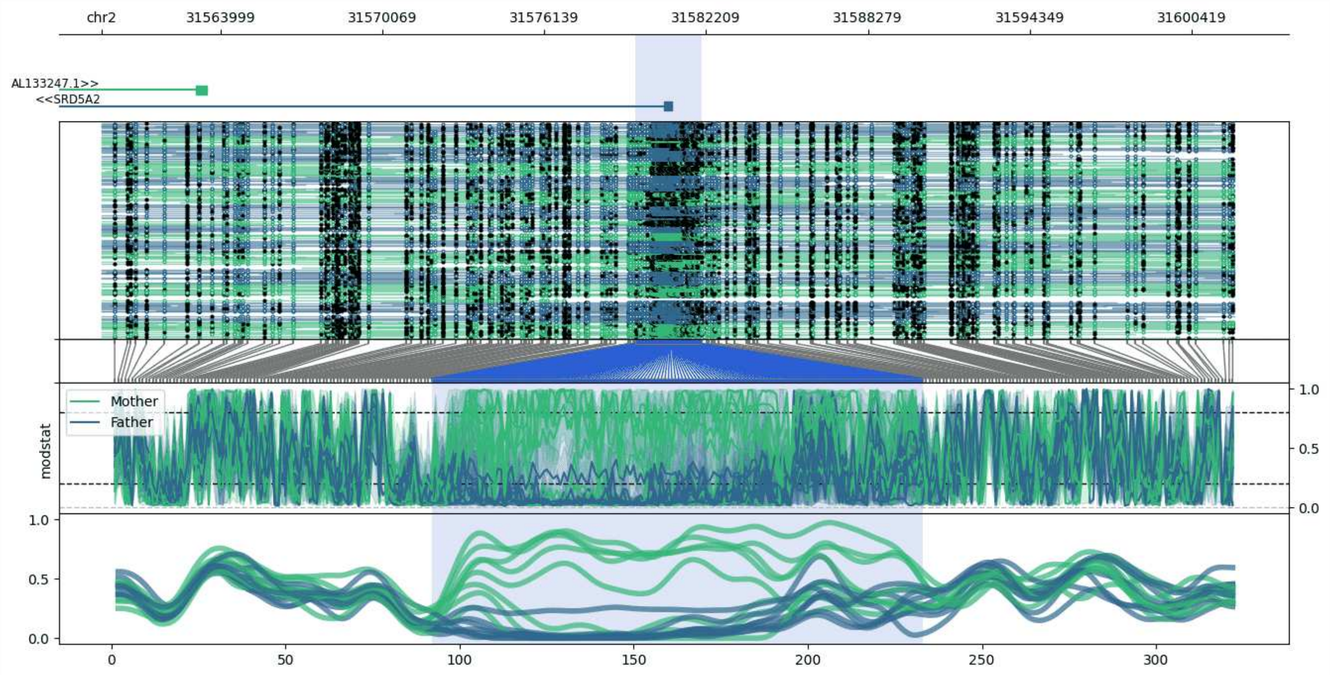


**chr2:31559532-31602027:** Region with 1 DMR containing genes: AL133247.1, SRD5A2.

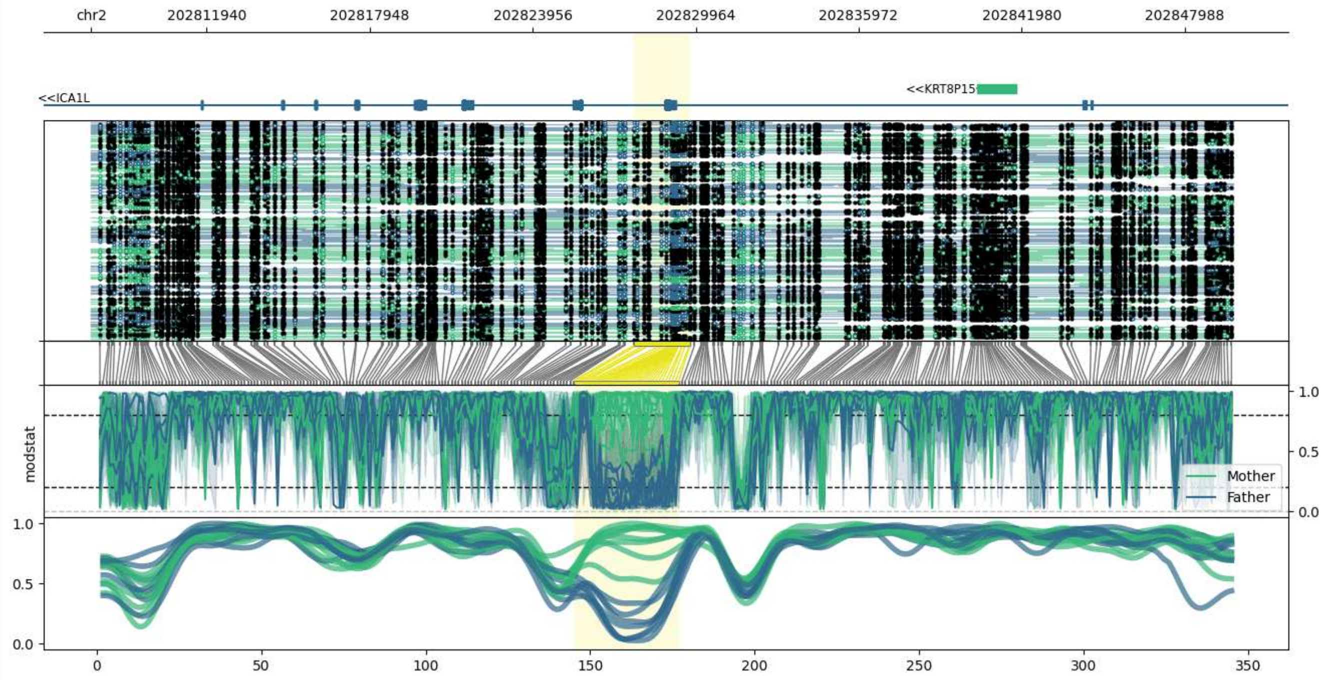


**chr2:202807694-202849753:** Region with 1 DMR containing genes: ICA1L, KRT8P15.

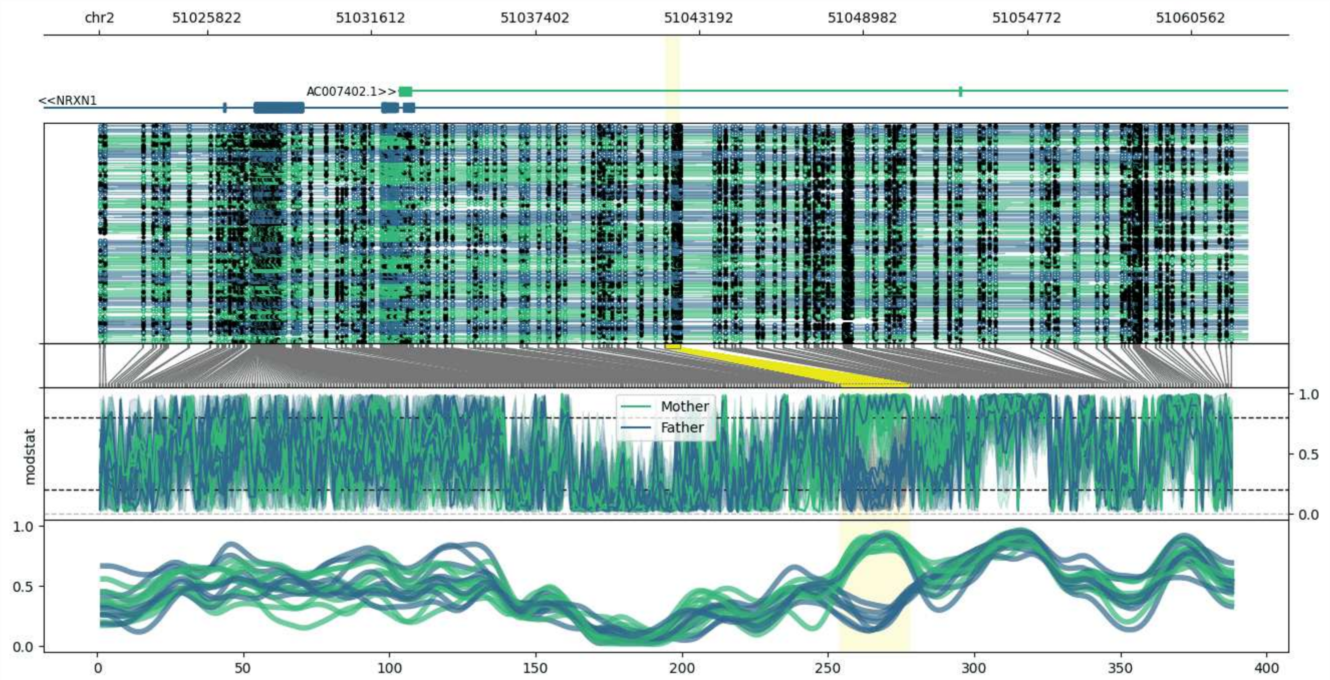


**chr2:51022002-51062534:** Region with 1 DMR containing genes: AC007402.1, AC007682.1, NRXN1.

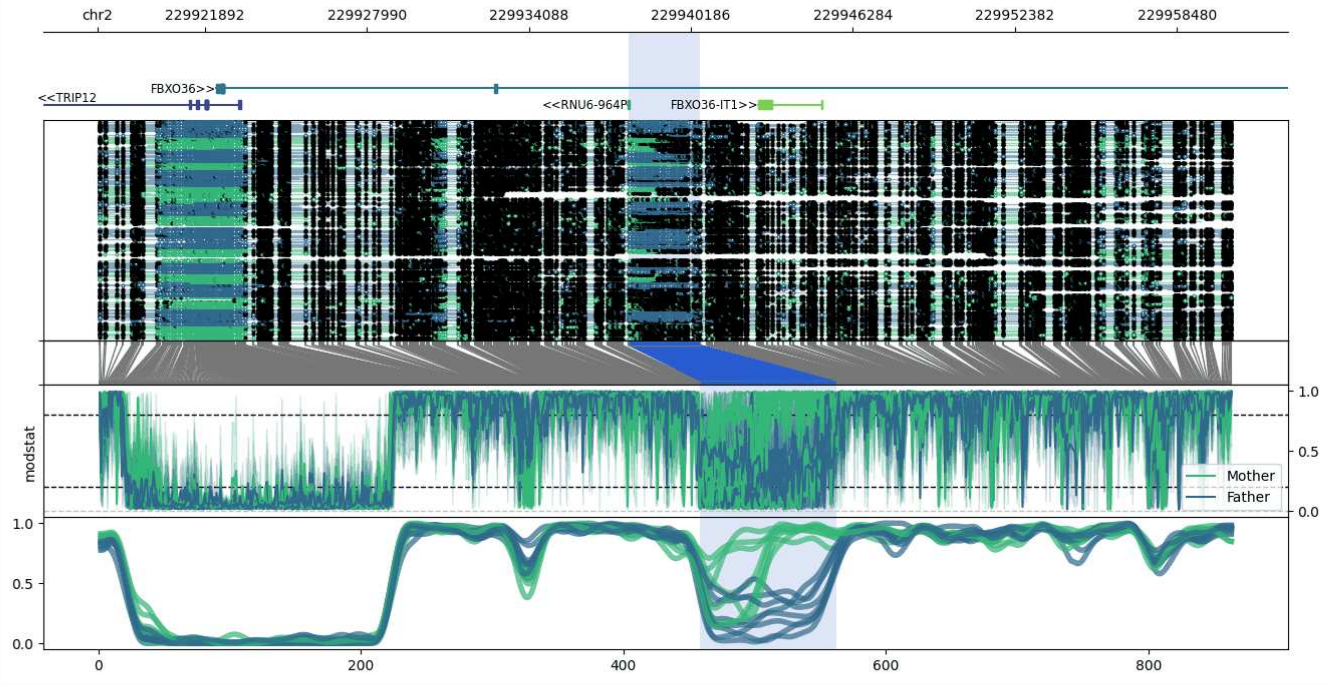


**chr2:229917843-229960533:** Region with 1 DMR containing genes: FBXO36, FBXO36-IT1, RNU6-964P, TRIP12.

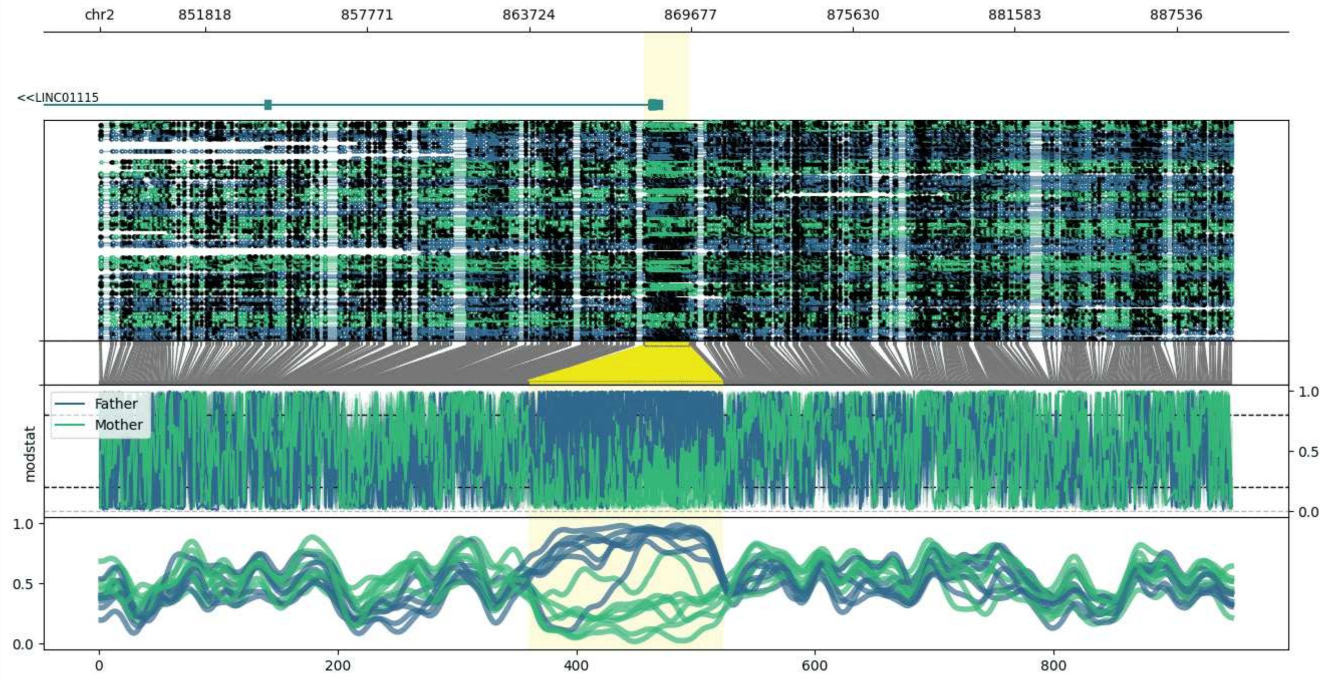


**chr2:847944-889618:** Region with 1 DMR containing genes: LINC01115, LINC01939.

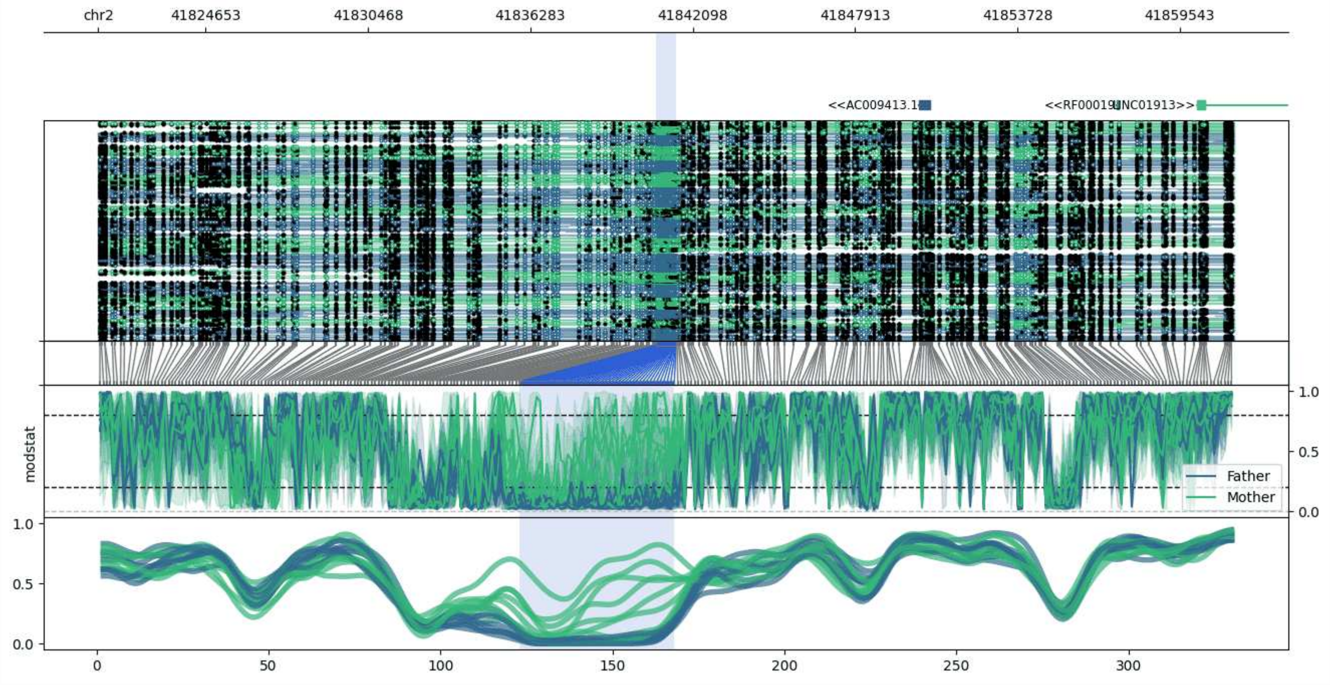


**chr2:41820798-41861508:** Region with 1 DMR containing genes: AC009413.1, LDHAP3, LINC01913, RF00019, RNU4-63P.

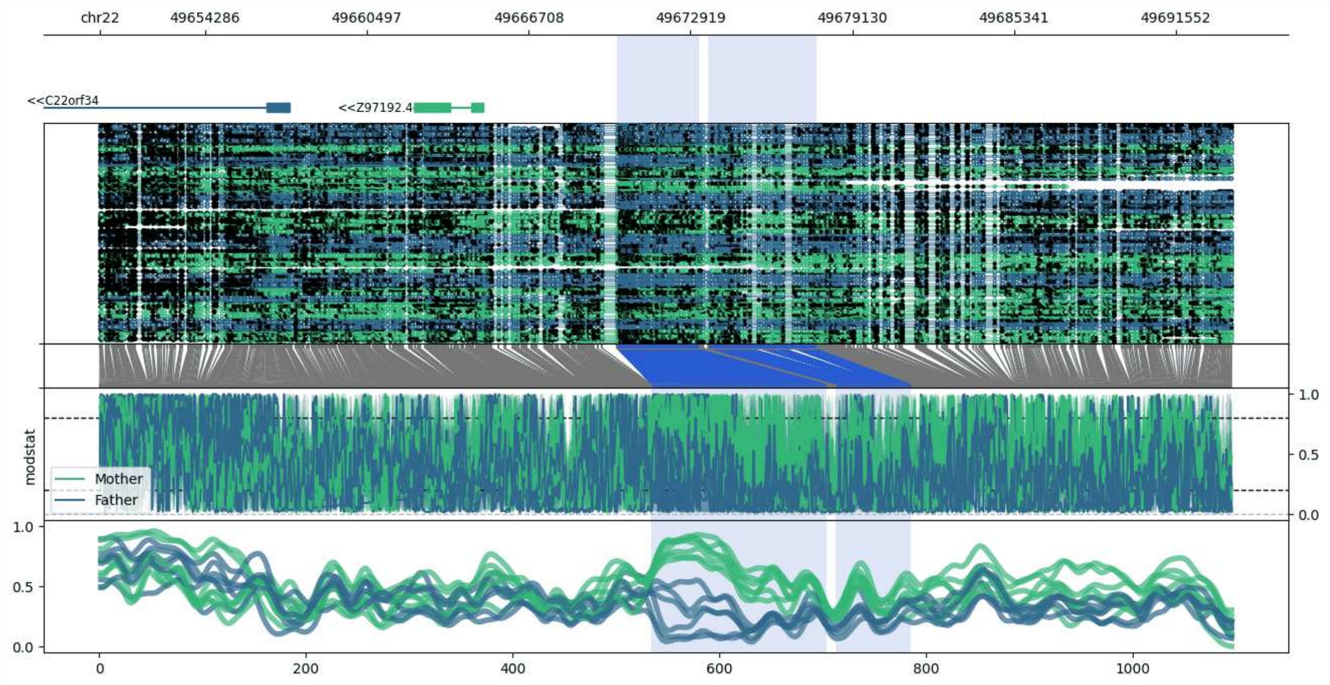


**chr22:49650242-49693722:** Region with 2 DMRs containing genes: C22orf34, Z97192.4.

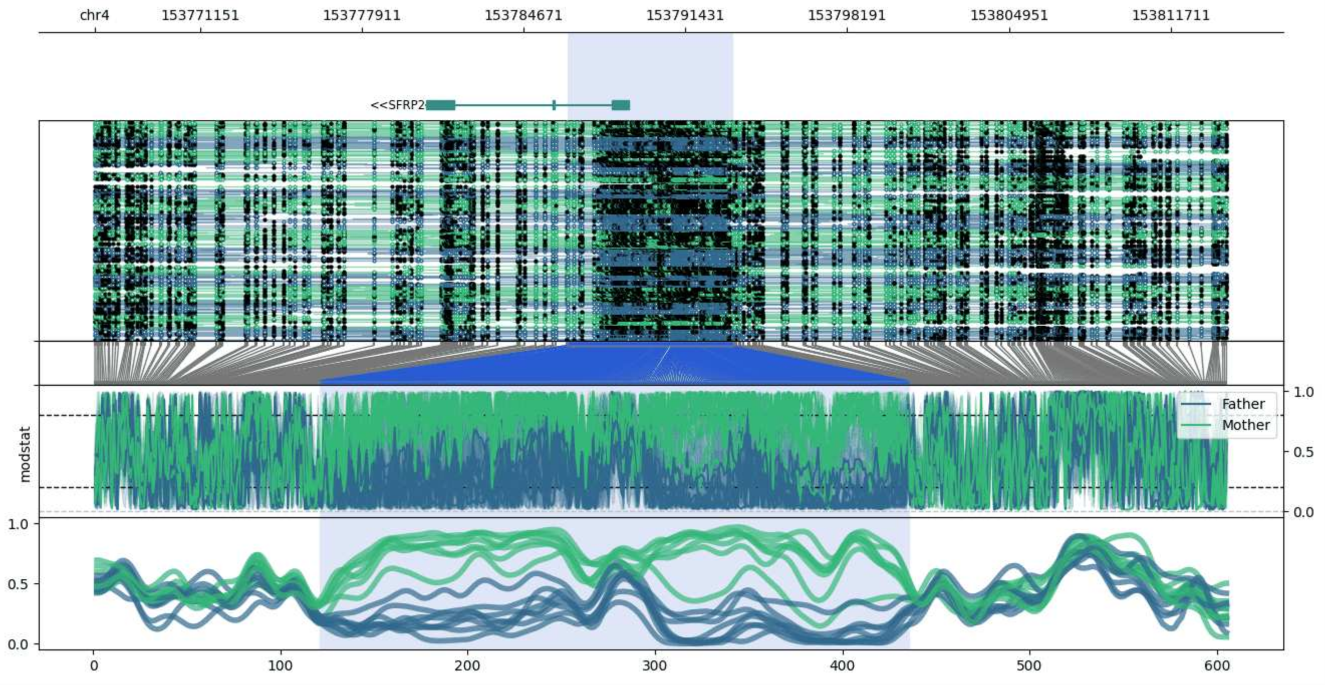


**chr4:153766747-153814068:** Region with 1 DMR containing genes: RNF175, SFRP2.

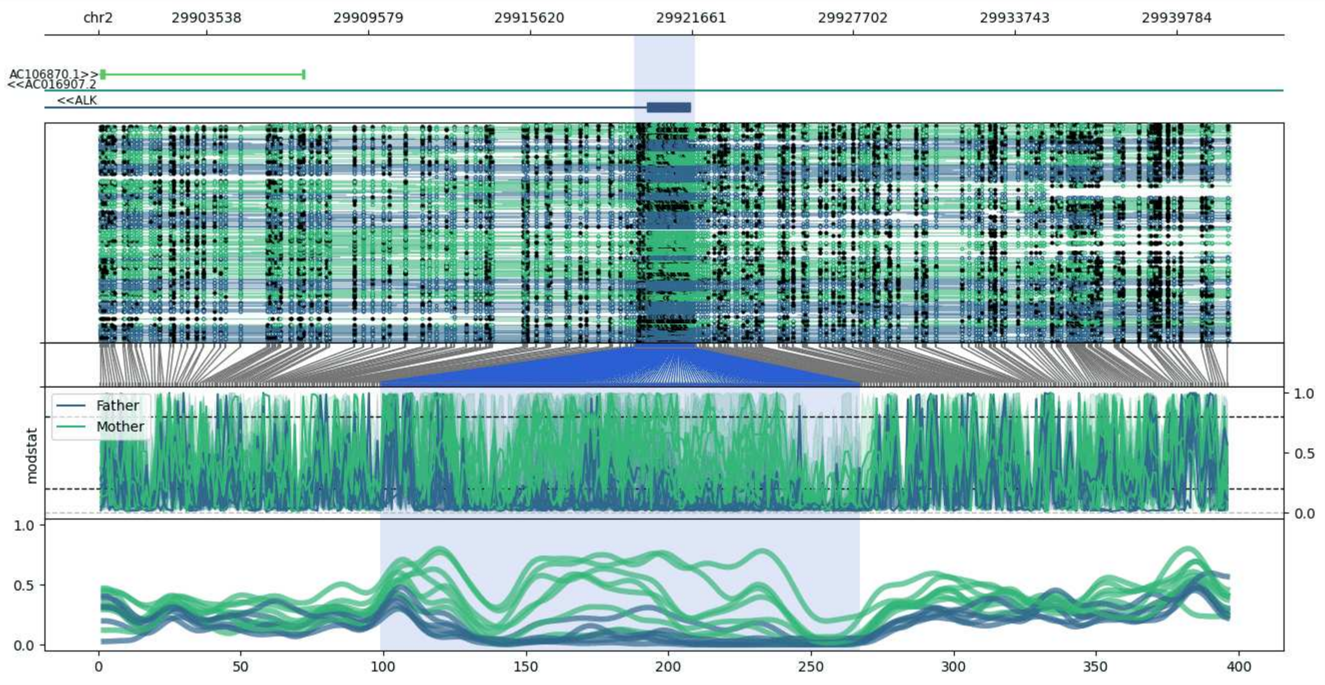


**chr2:29899496-29941784:** Region with 1 DMR containing genes: AC016907.2, AC106870.1, AC106870.2, ALK.

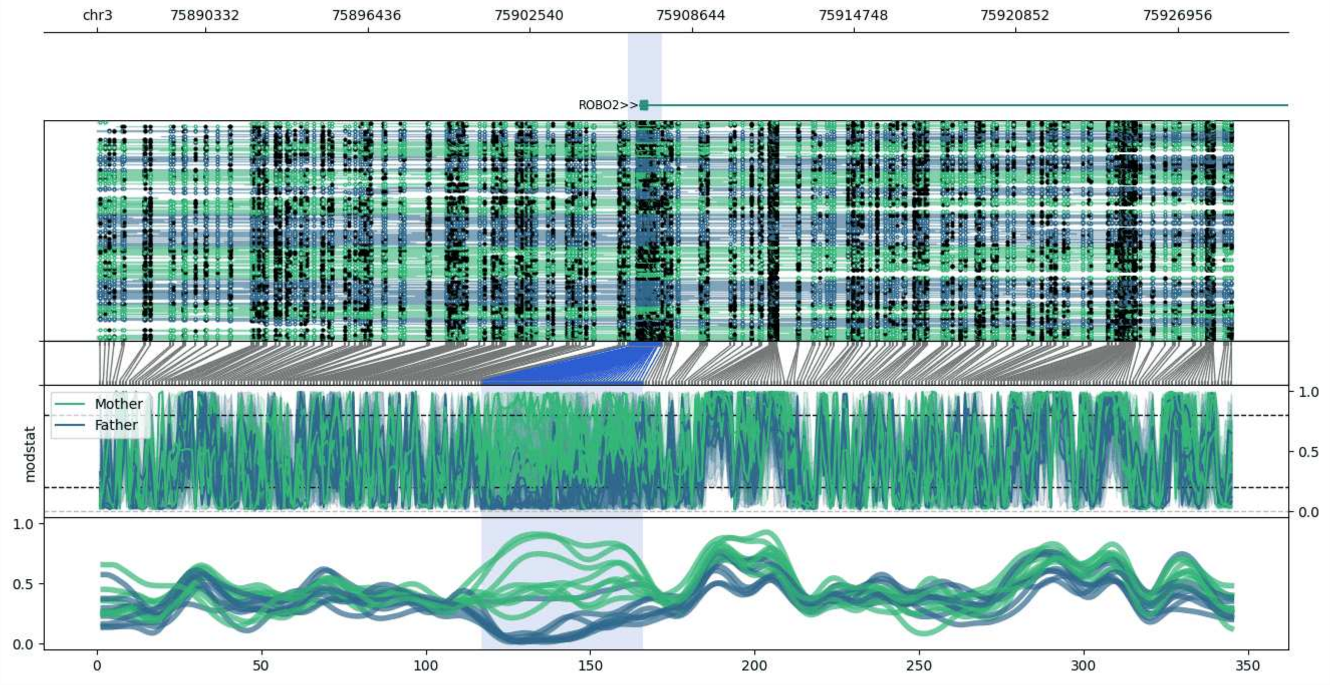


**chr3:75886251-75928983:** Region with 1 DMR containing genes: ROBO2.

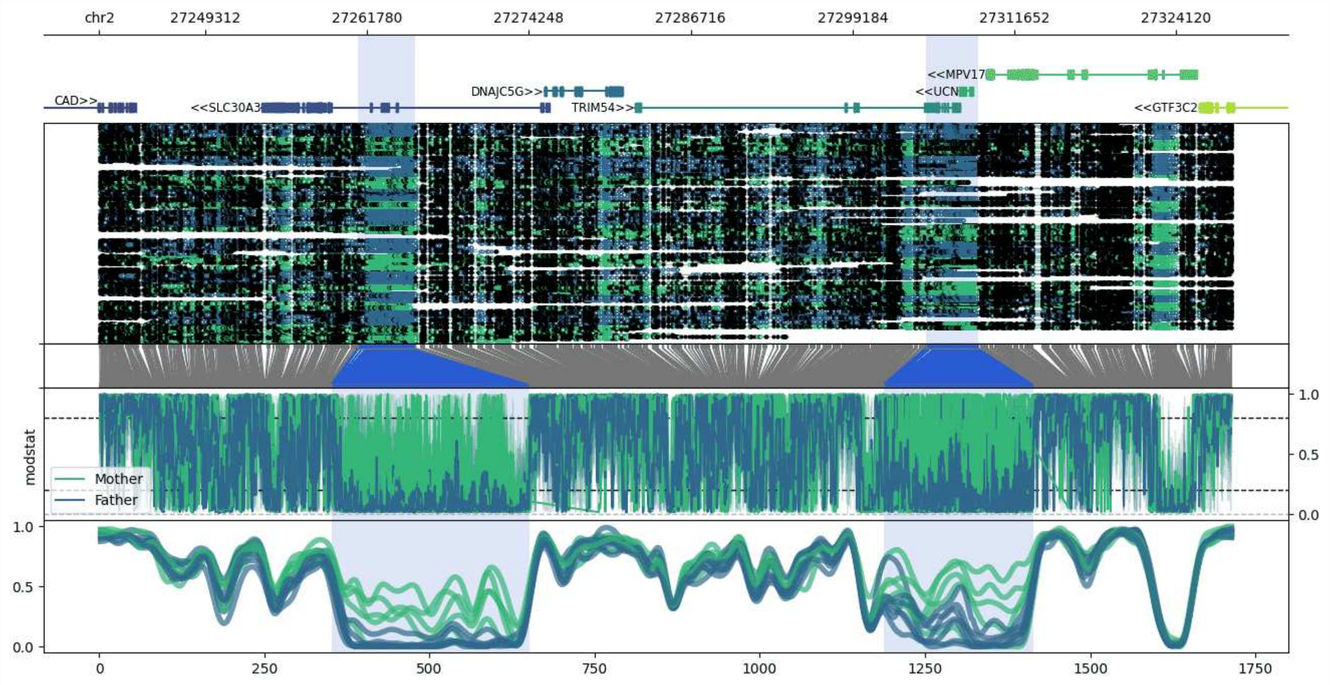


**chr2:27241143-27328422:** Region with 2 DMRs containing genes: CAD, DNAJC5G, GTF3C2, GTF3C2-AS1, MPV17, SLC30A3, TRIM54, UCN.

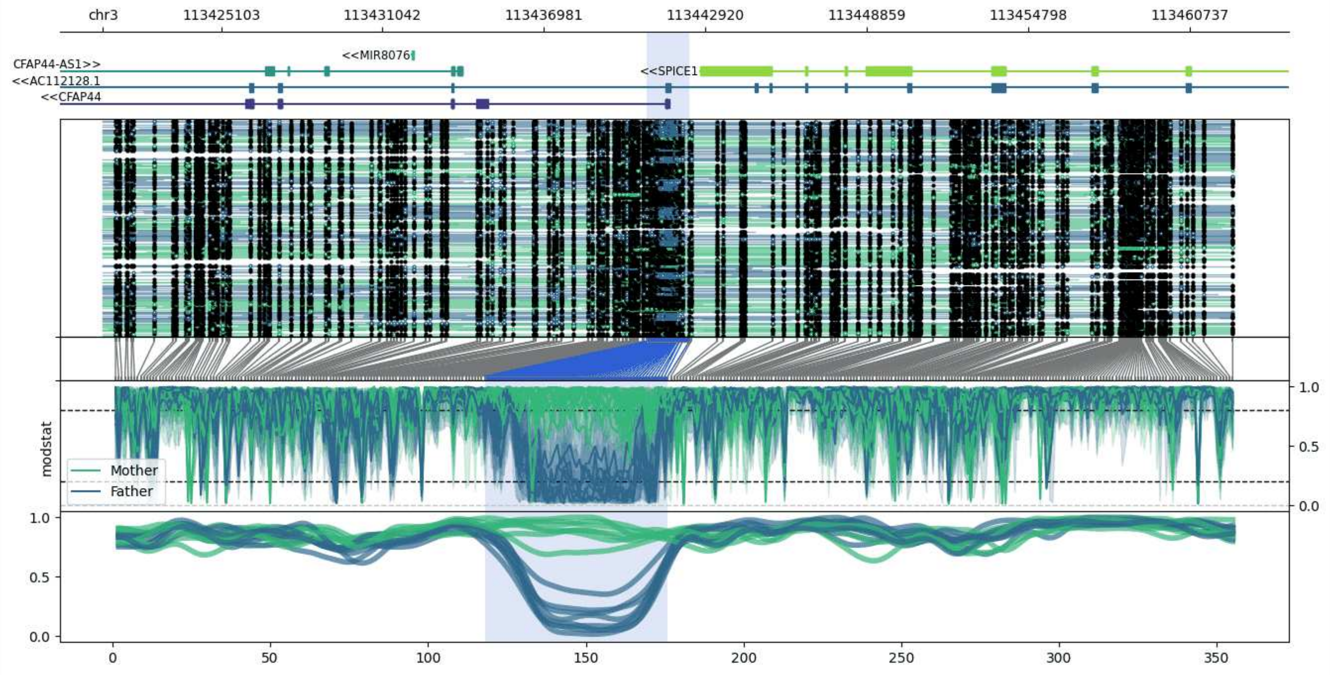


**chr3:113420750-113462329:** Region with 1 DMR containing genes: AC112128.1, CFAP44, CFAP44-AS1, MIR8076, SPICE1.

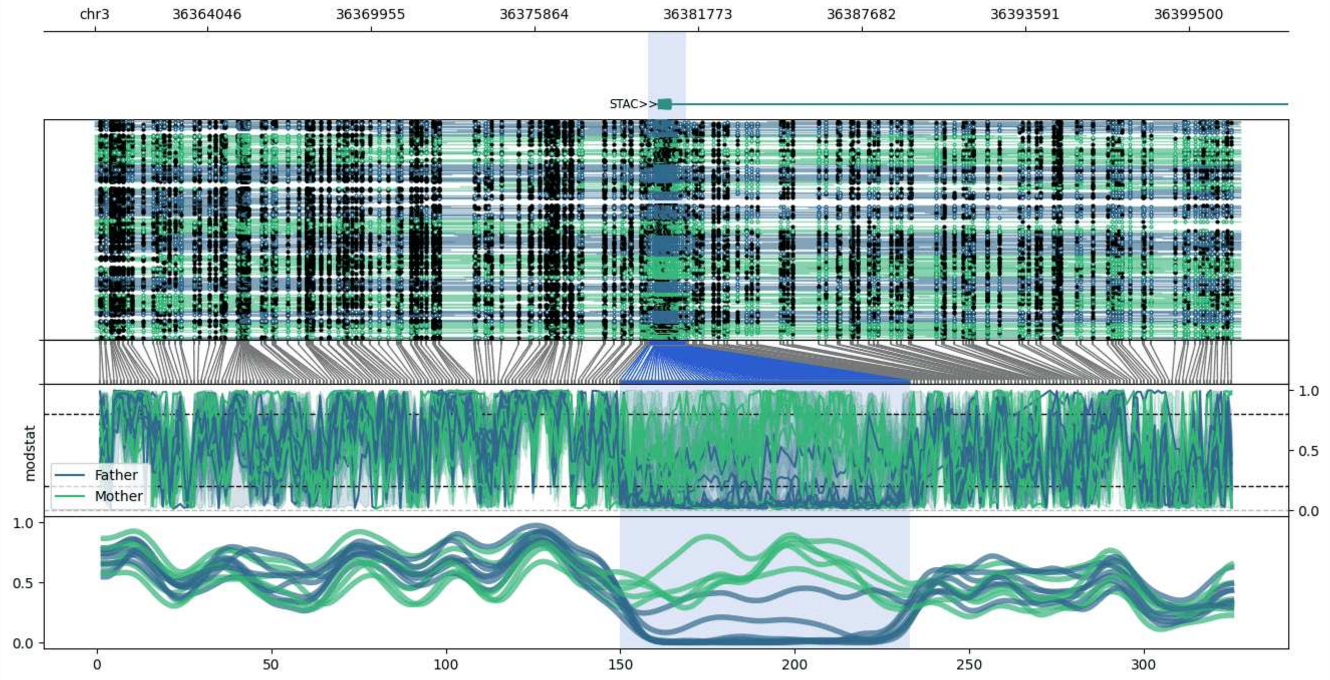


**chr3:36359981-36401346:** Region with 1 DMR containing genes: STAC.

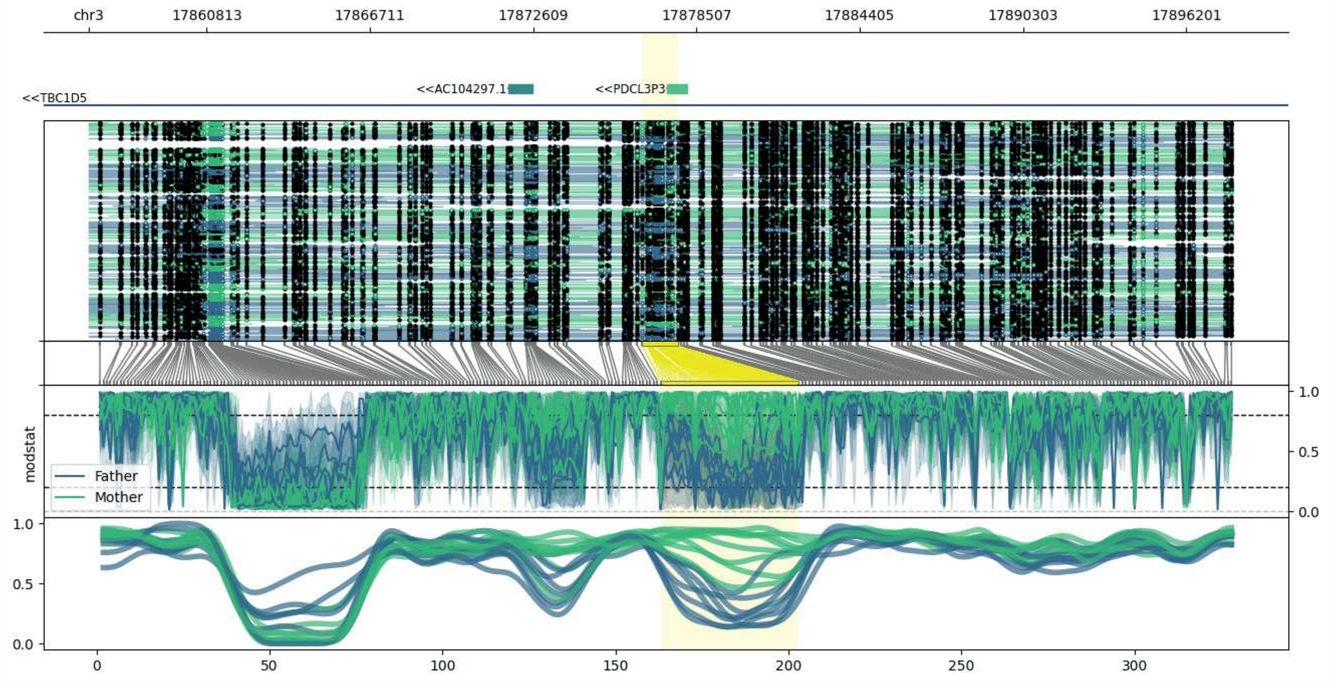


**chr3:17856550-17897842:** Region with 1 DMR containing genes: AC104297.1, PDCL3P3, TBC1D5.

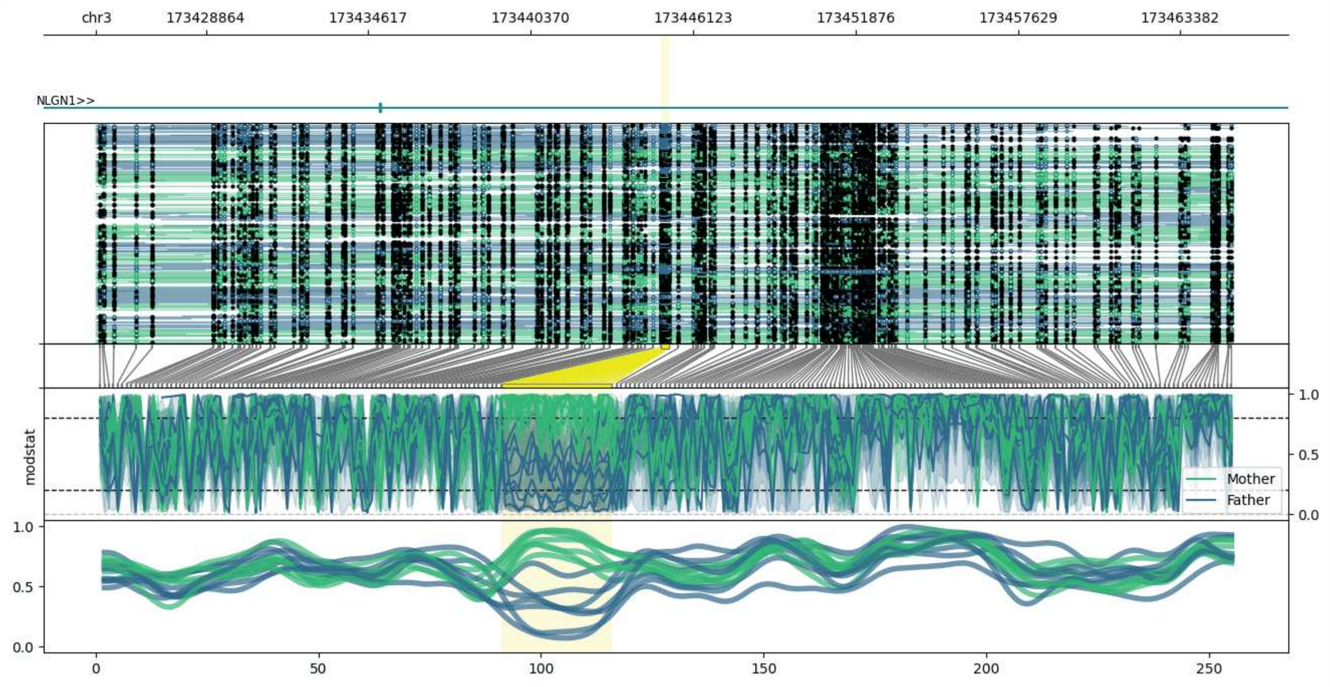


**chr3:173424986-173465261:** Region with 1 DMR containing genes: NLGN1.

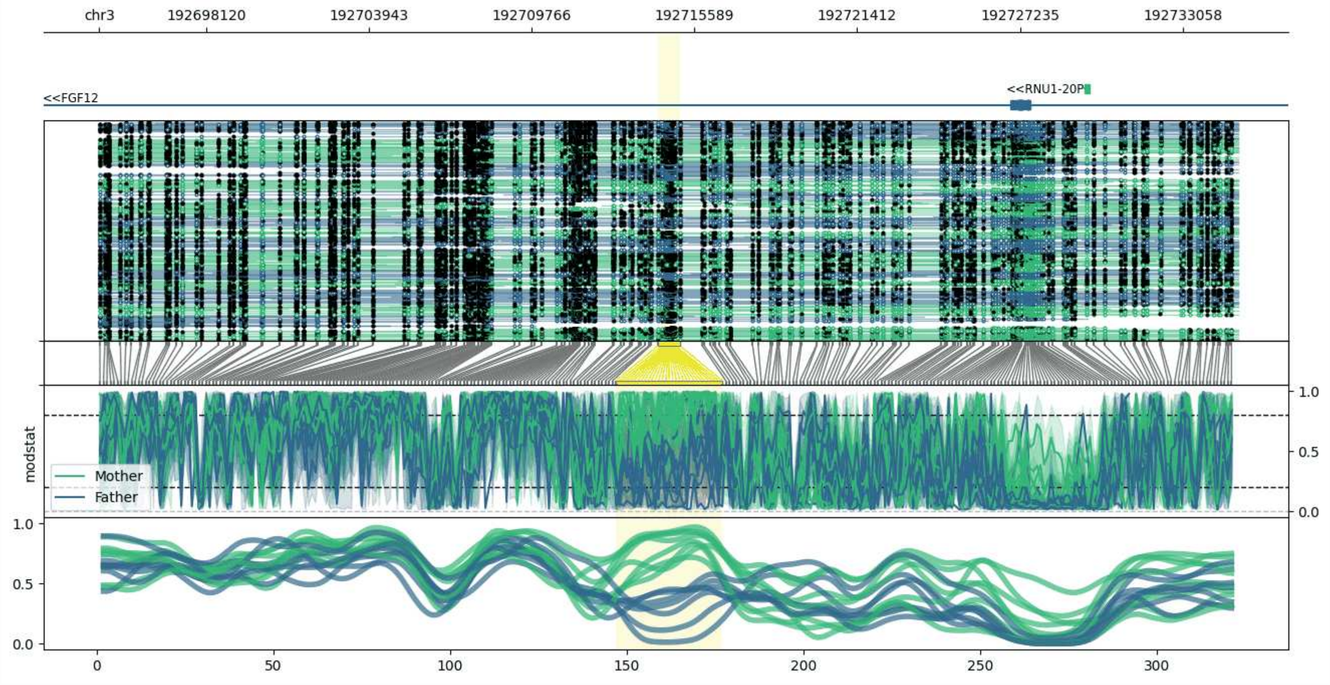


**chr3:192694294-192735058:** Region with 1 DMR containing genes: FGF12, RNU1-20P.

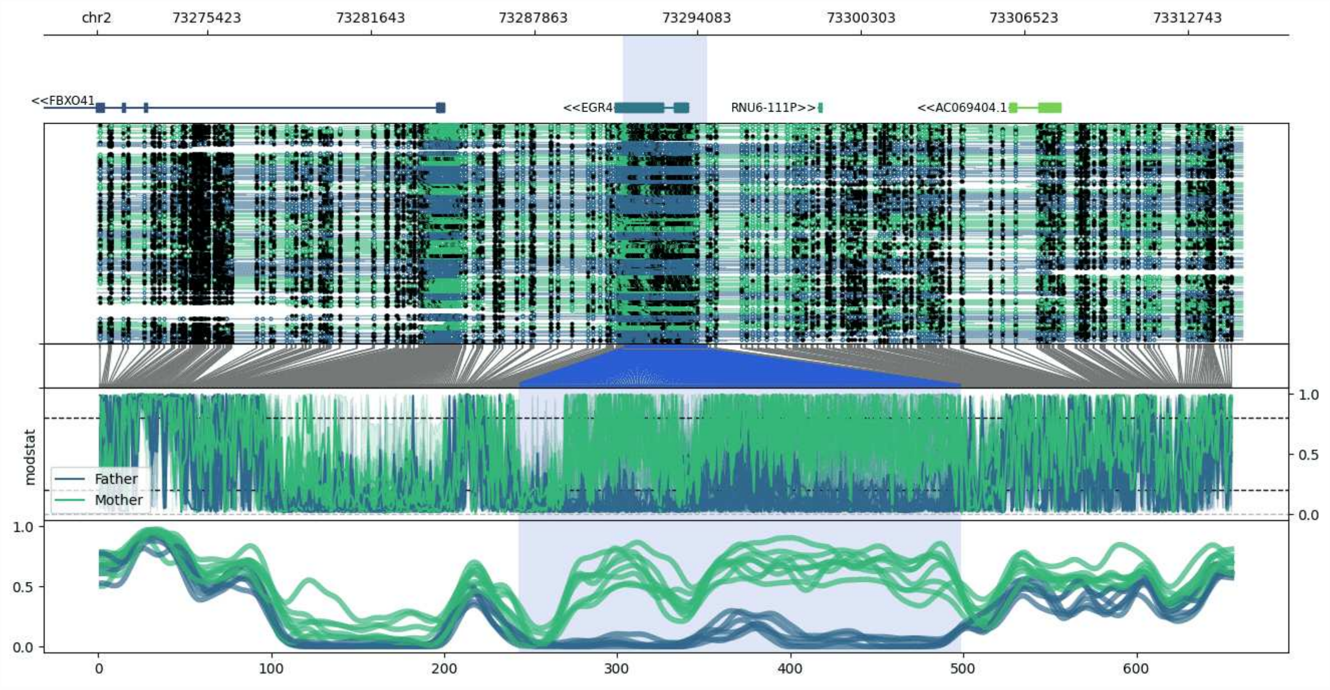


**chr2:73271236-73314777:** Region with 1 DMR containing genes: AC069404.1, EGR4, FBXO41, RNU6-111P.

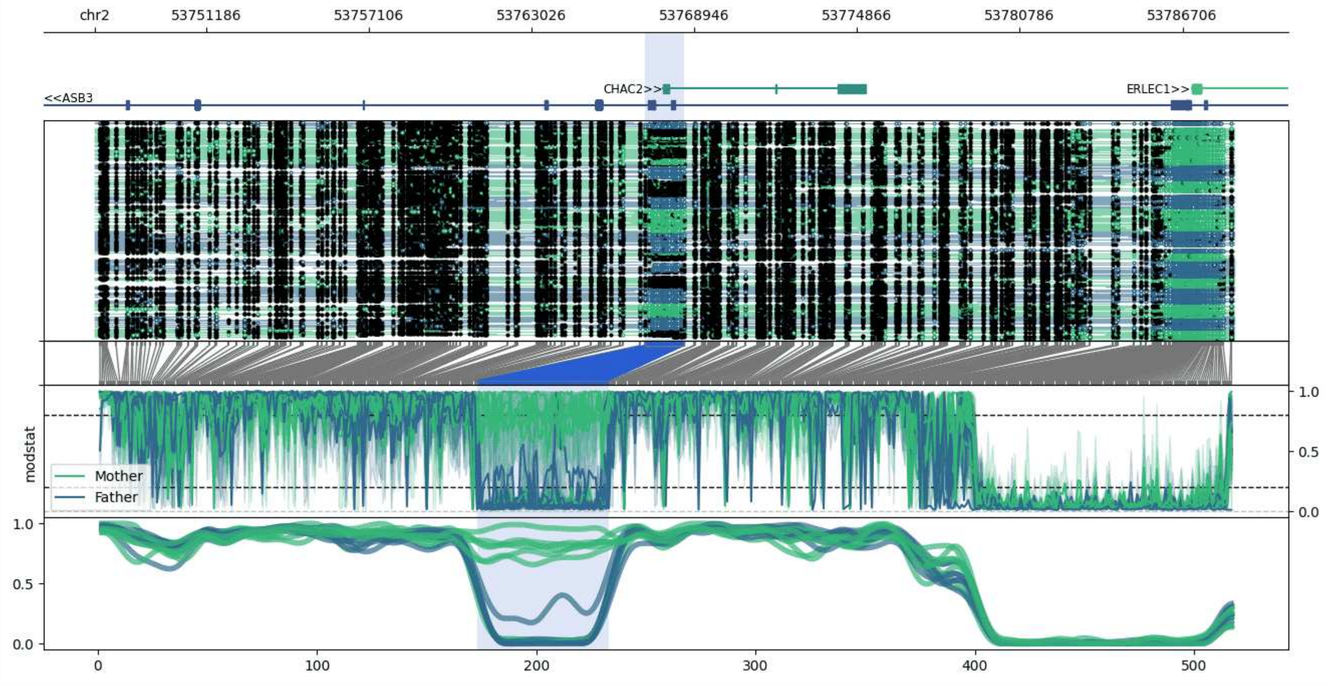


**chr2:53747131-53788572:** Region with 1 DMR containing genes: ASB3, CHAC2, ERLEC1.

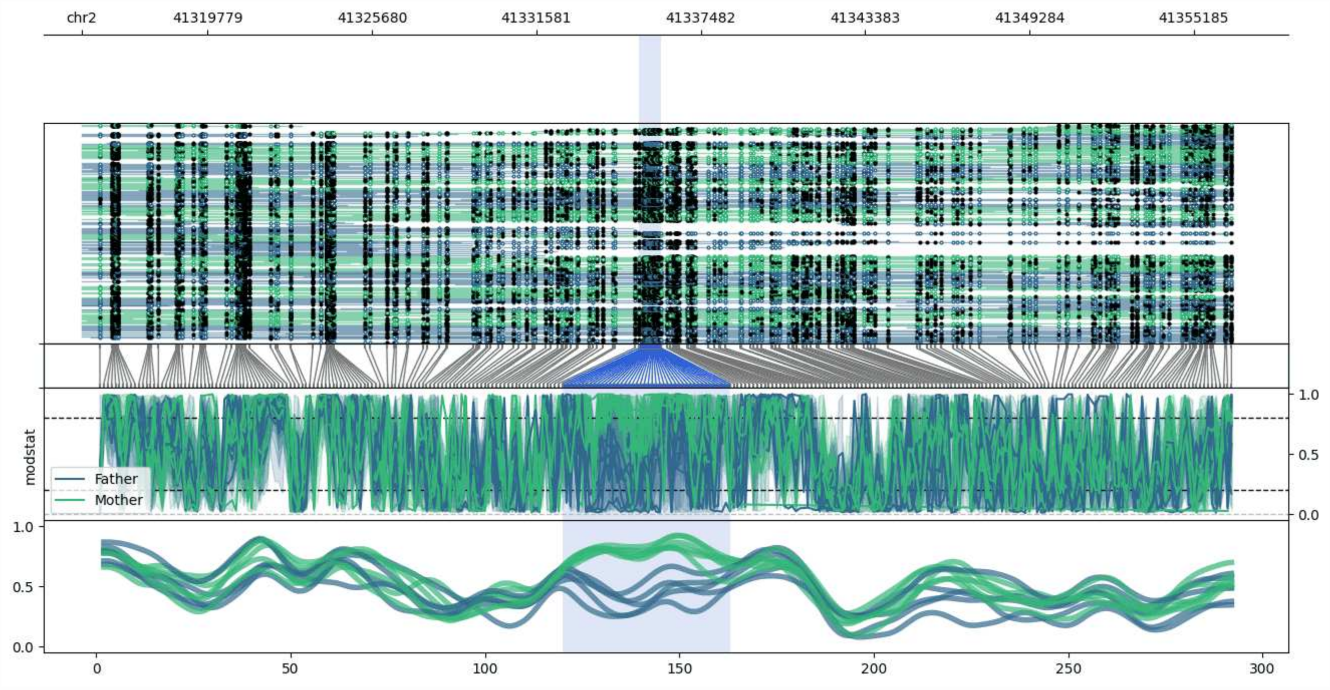


**chr2:41315258-41356565:** Region with 1 DMR containing no genes.

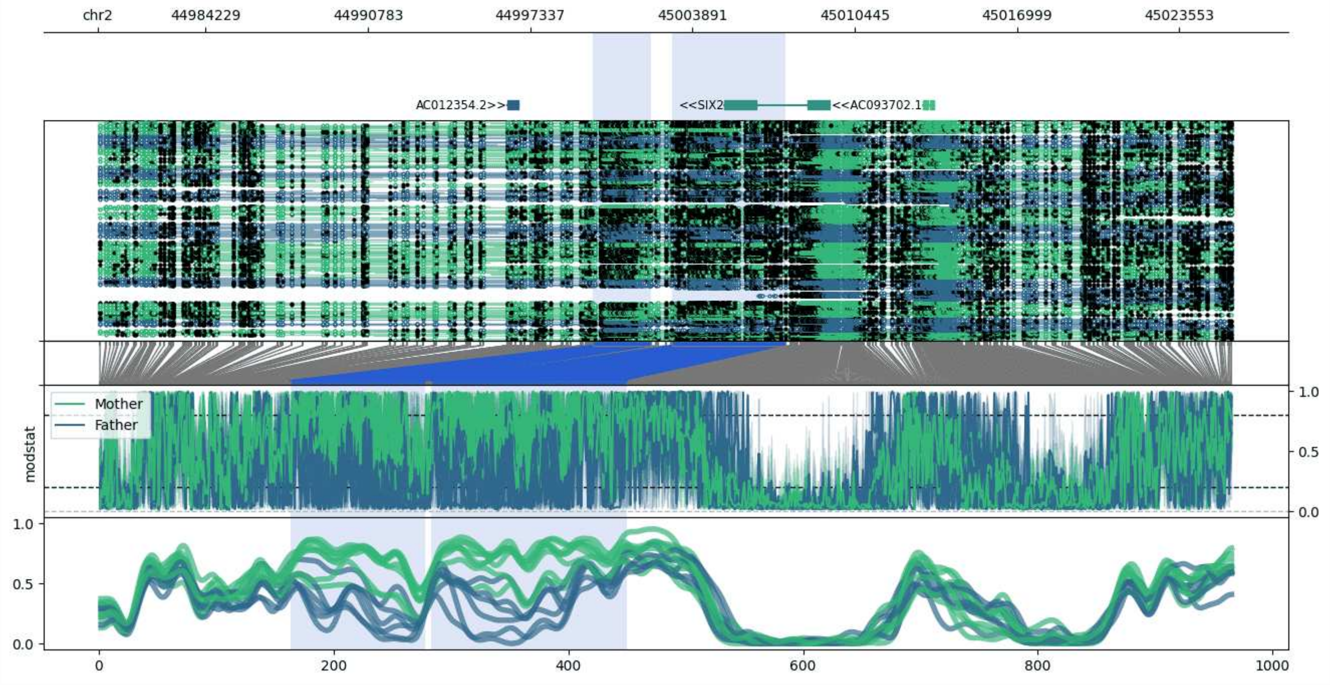


**chr2:44979869-45025749:** Region with 2 DMRs containing genes: AC012354.2, AC093702.1, SIX2.

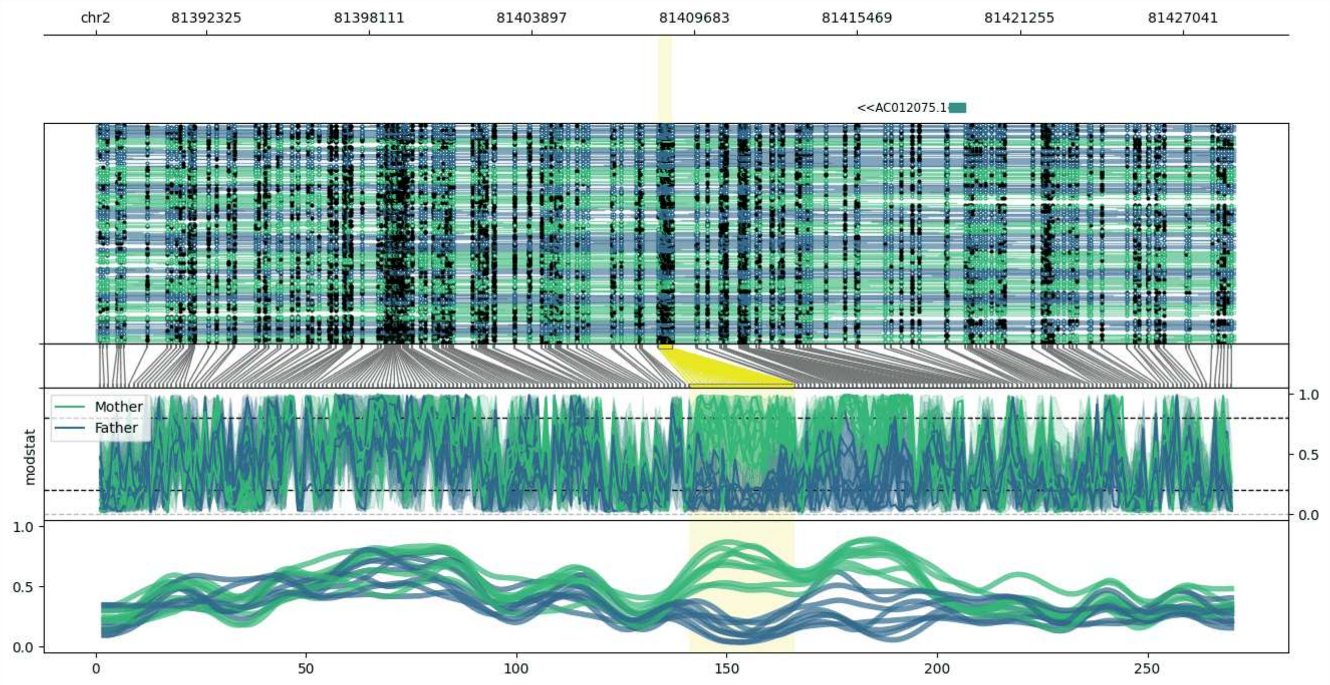


**chr2:81388394-81428900:** Region with 1 DMR containing genes: AC012075.1.

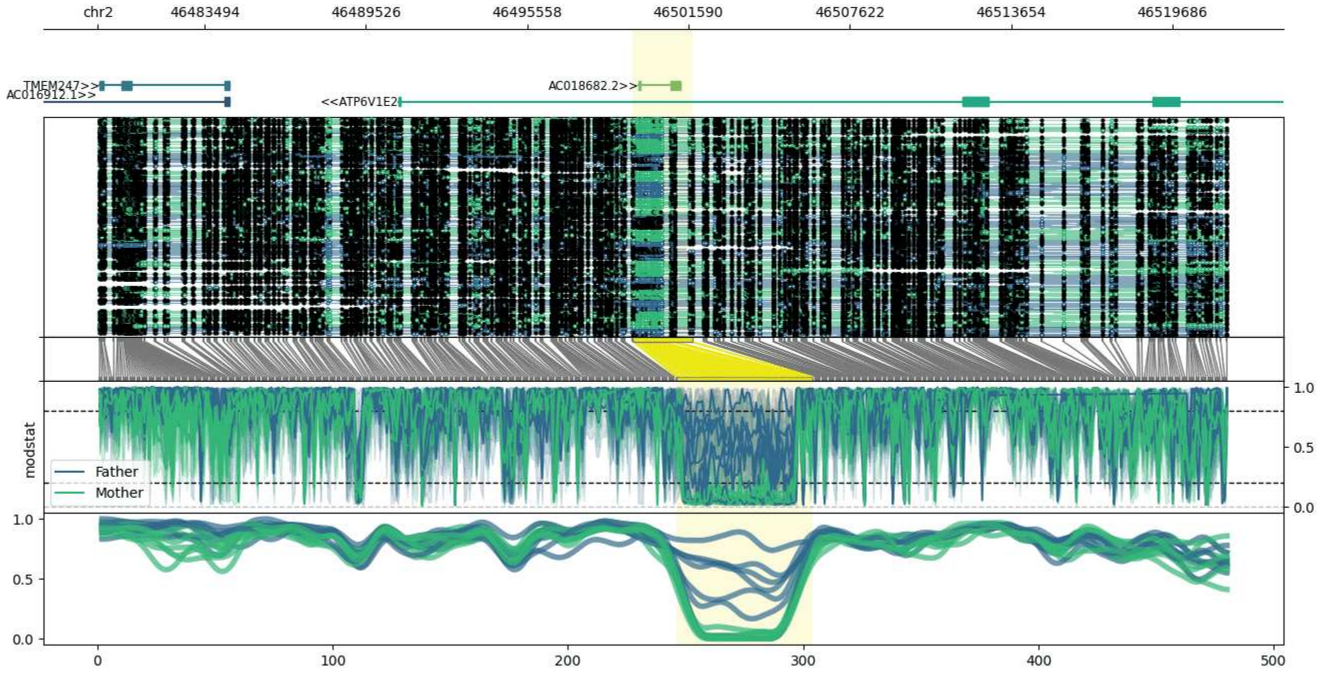


**chr2:46479505-46521730:** Region with 1 DMR containing genes: AC016912.1, AC018682.2, ATP6V1E2, TMEM247.

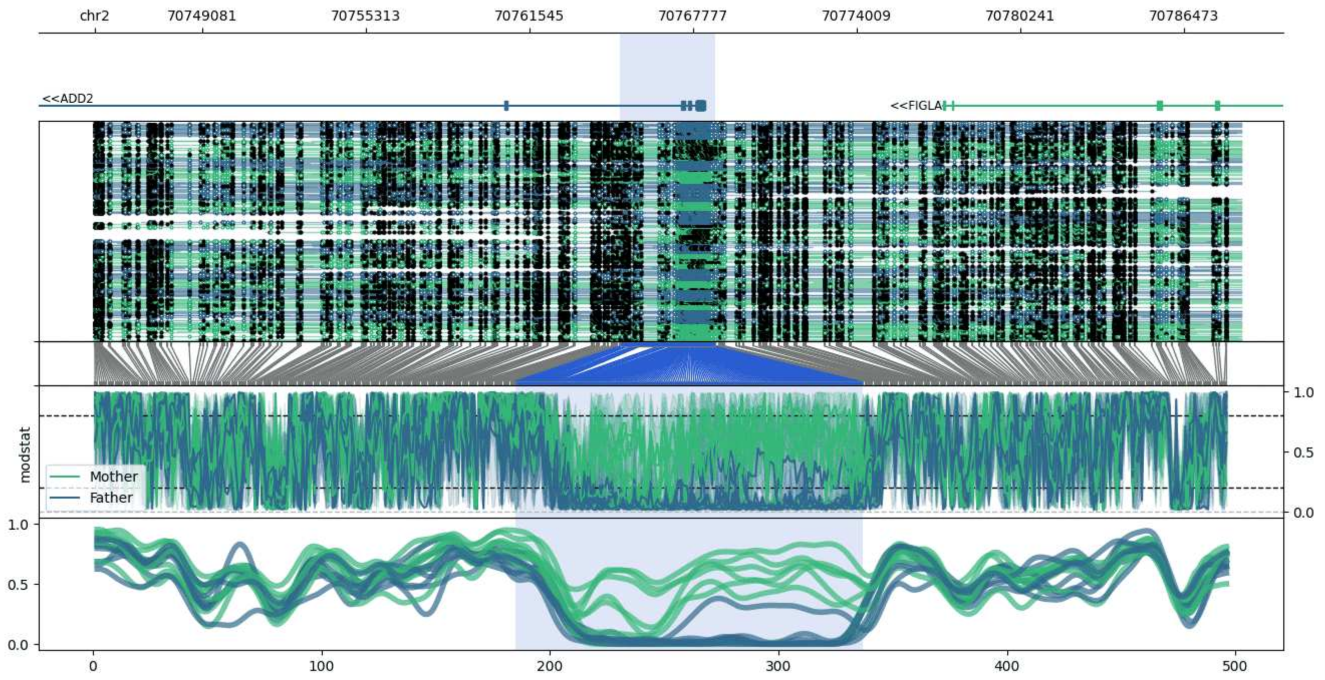


**chr2:70744998-70788625:** Region with 1 DMR containing genes: ADD2, FIGLA.

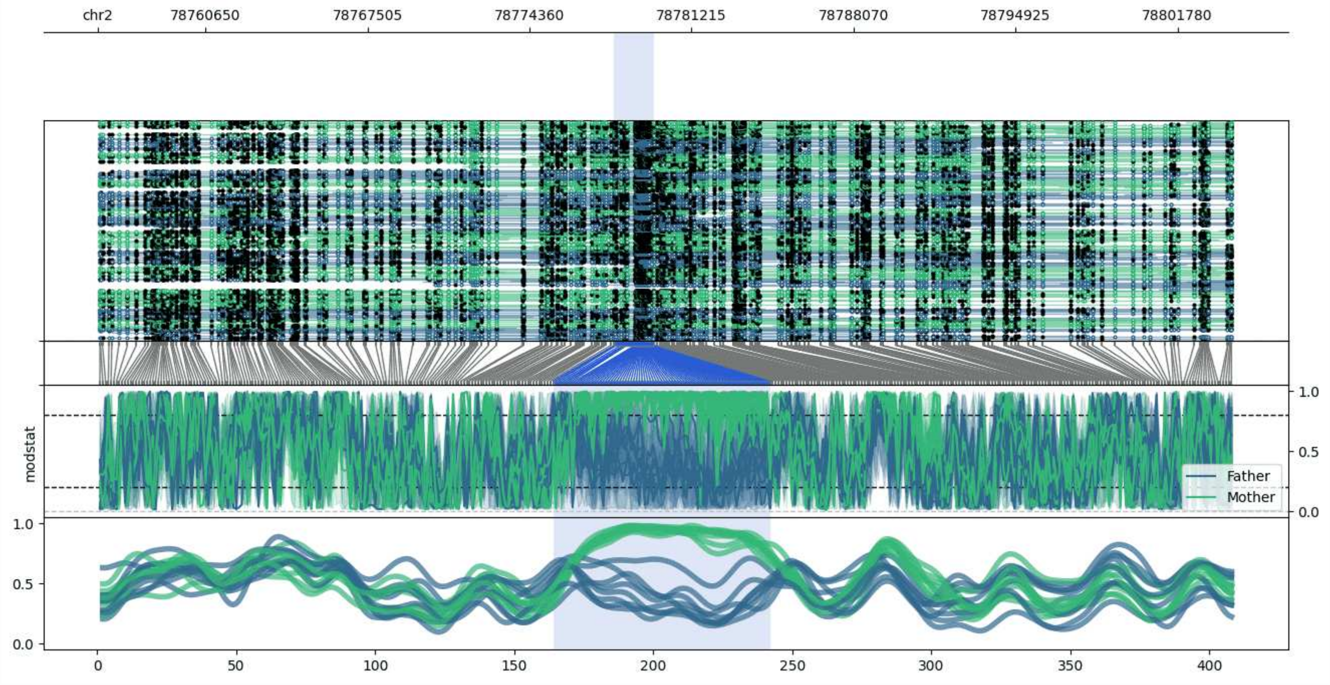


**chr2:78756097-78804085:** Region with 1 DMR containing no genes.

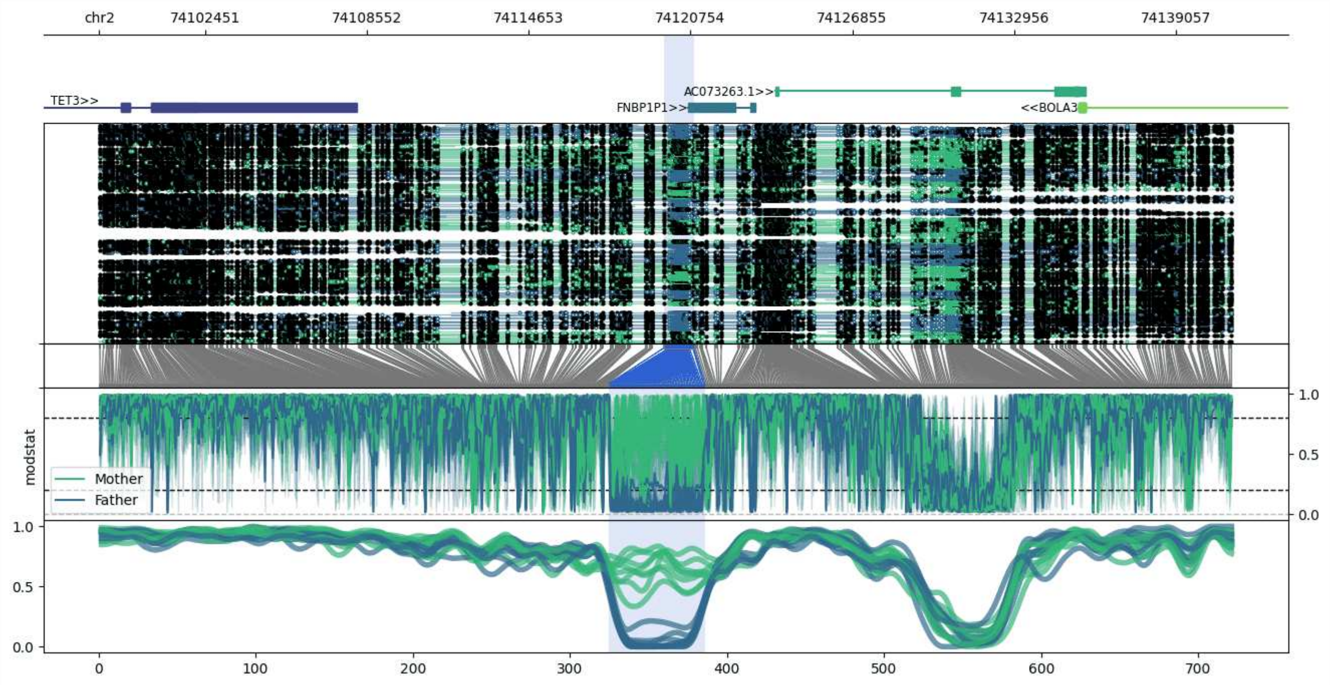


**chr2:74098460-74141171:** Region with 1 DMR containing genes: AC073263.1, BOLA3, BOLA3-AS1, FNBP1P1, TET3.

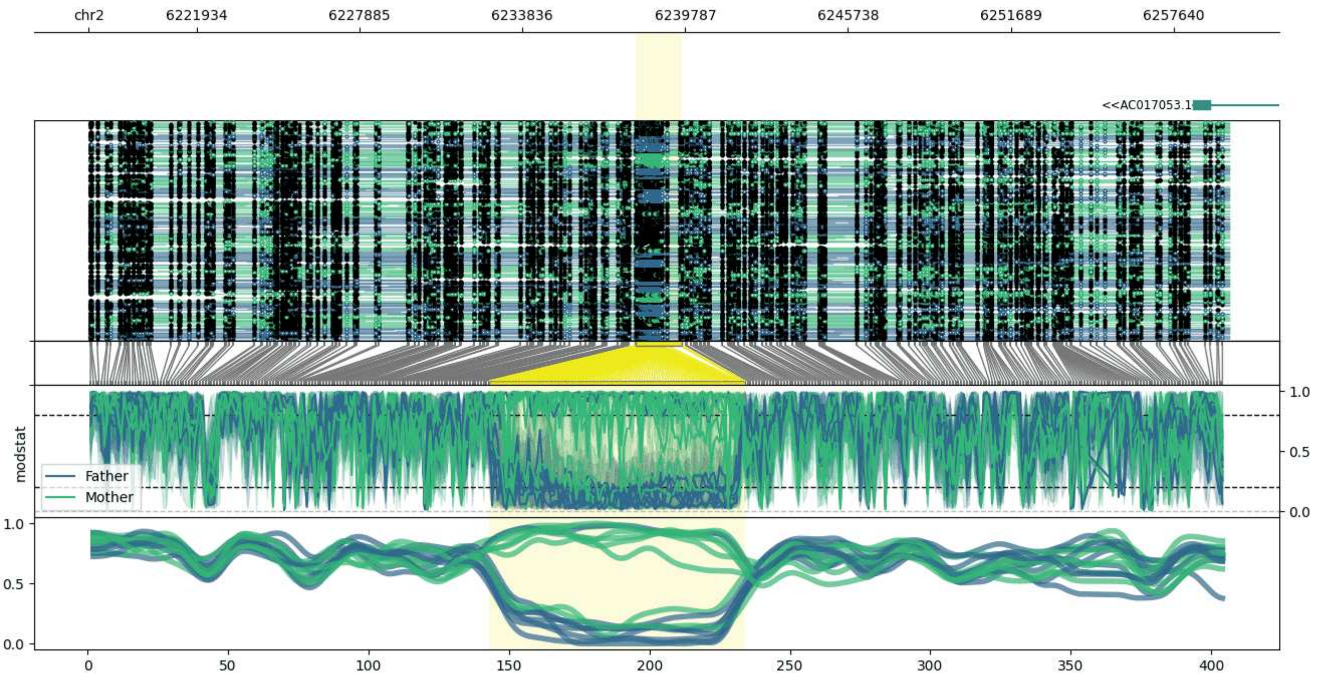


**chr2:6217991-6259650:** Region with 1 DMR containing genes: AC017053.1.

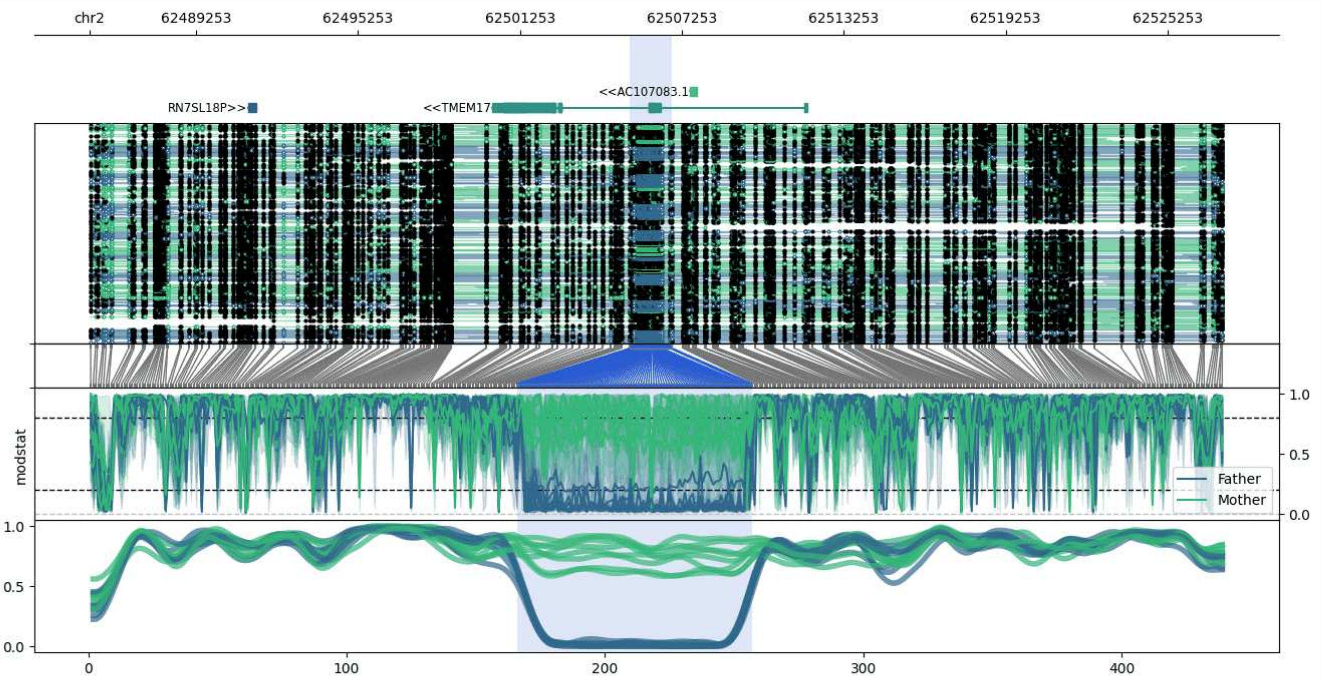


**chr2:62485314-62527319:** Region with 1 DMR containing genes: AC092155.1, AC092155.2, AC107083.1, RN7SL18P, TMEM17.

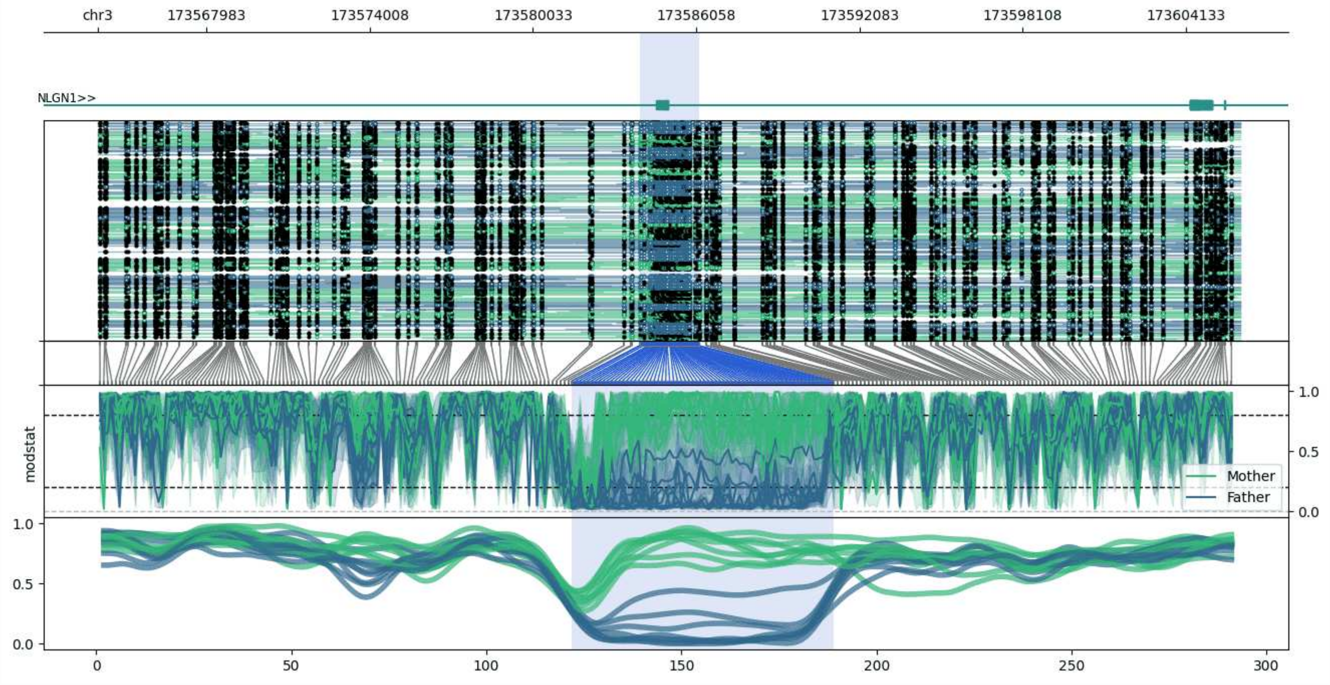


**chr3:173563971-173606152:** Region with 1 DMR containing genes: NLGN1.

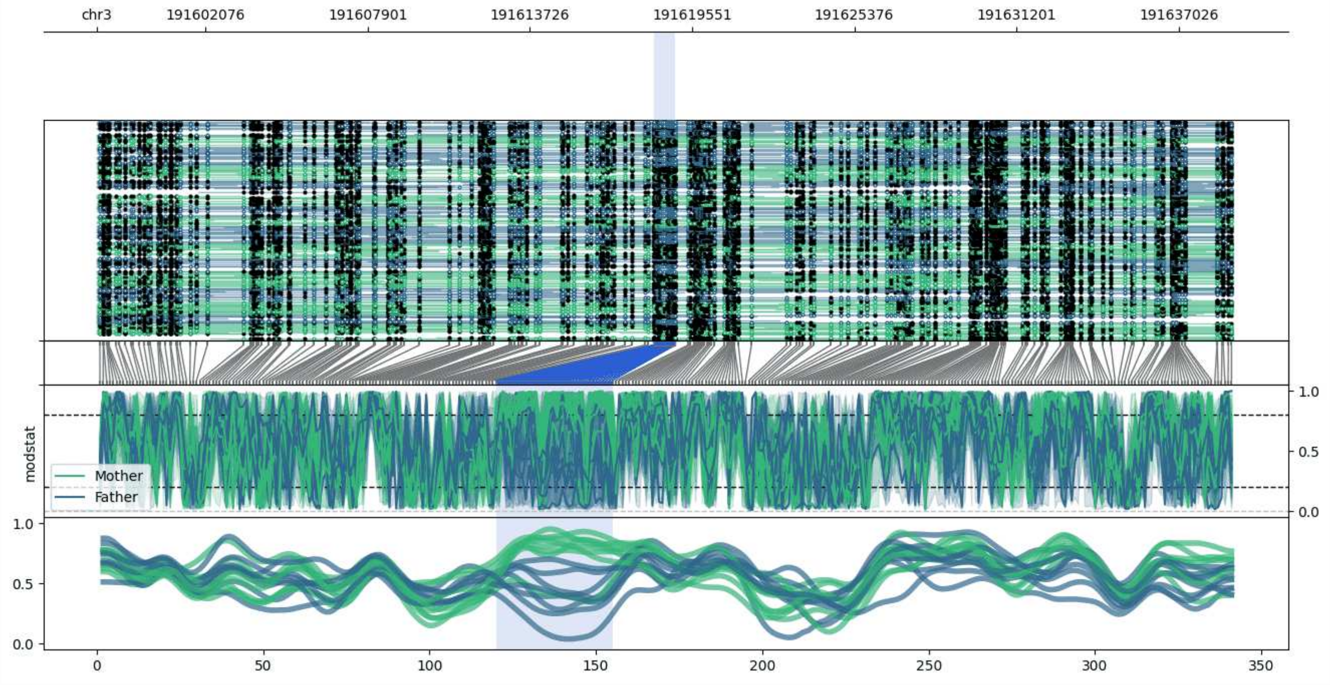


**chr3:191598164-191638939:** Region with 1 DMR containing genes: AC073365.1, RF00019.

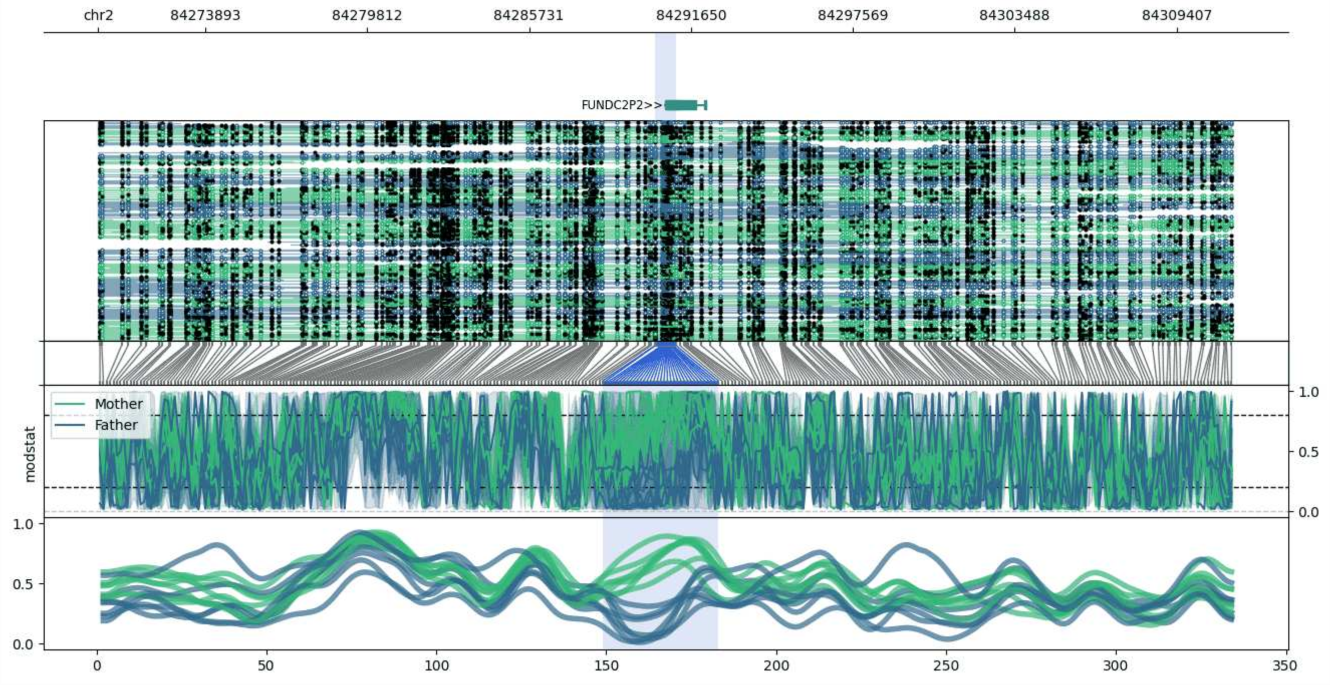


**chr2:84269977-84311416:** Region with 1 DMR containing genes: AC106874.1, FUNDC2P2.

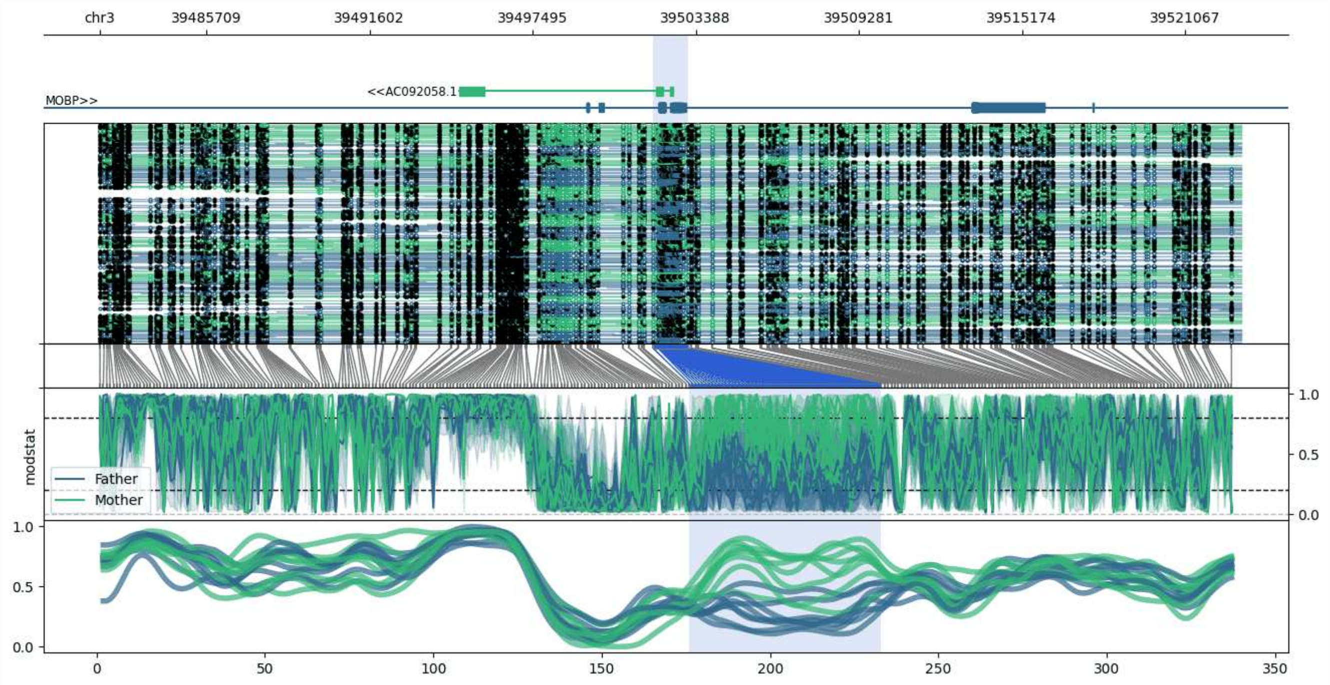


**chr3:39481847-39523099:** Region with 1 DMR containing genes: AC092058.1, MOBP.

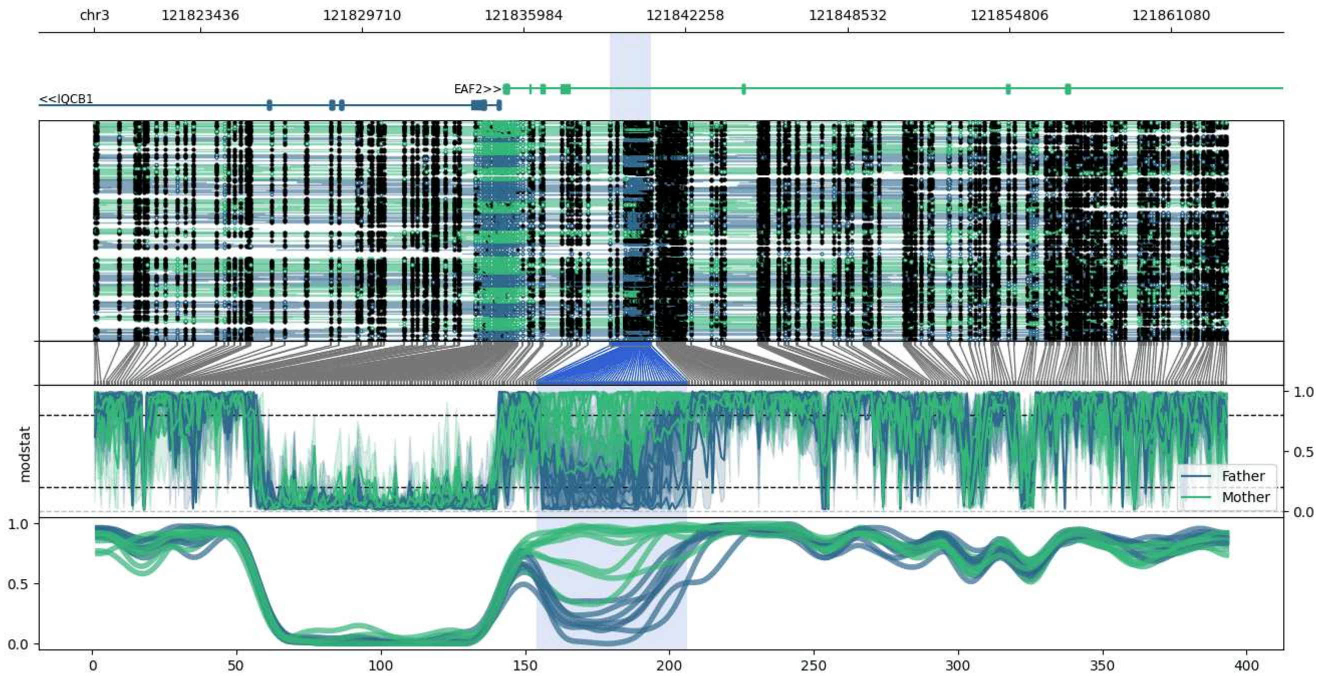


**chr3:121819328-121863246:** Region with 1 DMR containing genes: EAF2, IQCB1.

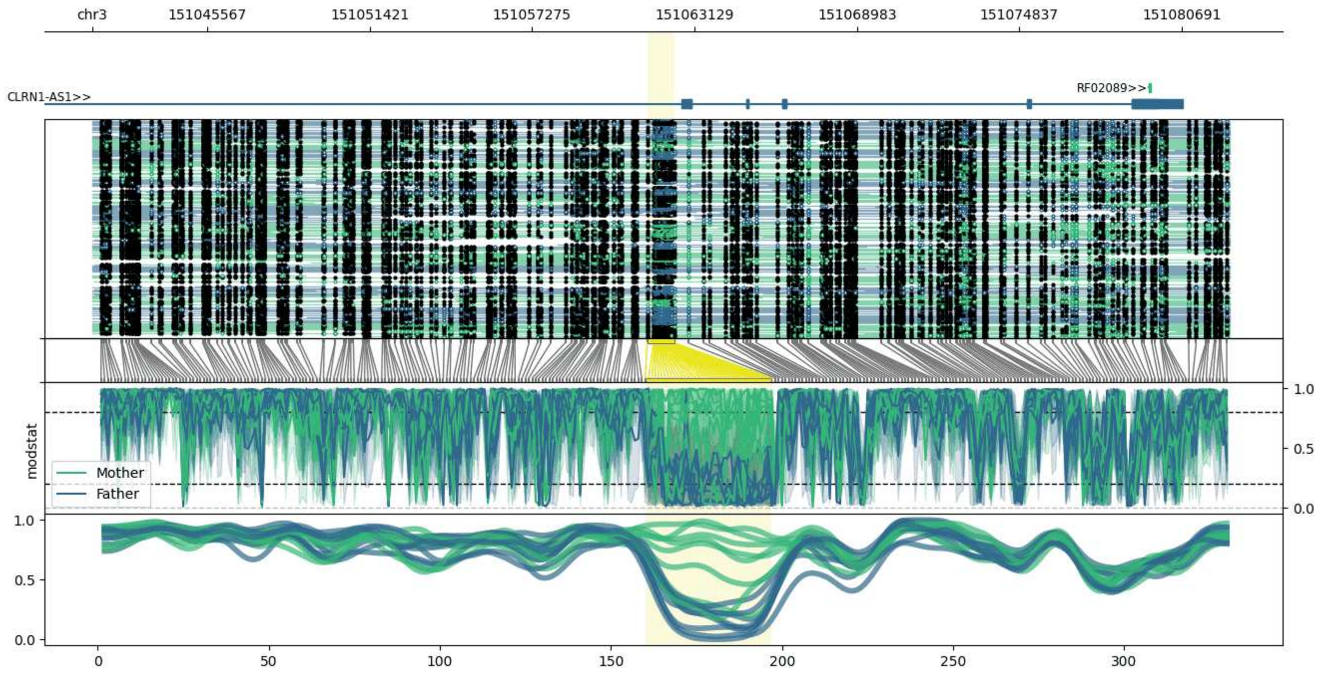


**chr3:151041425-151082408:** Region with 1 DMR containing genes: AC020636.1, CLRN1-AS1, MED12L, RF02089.

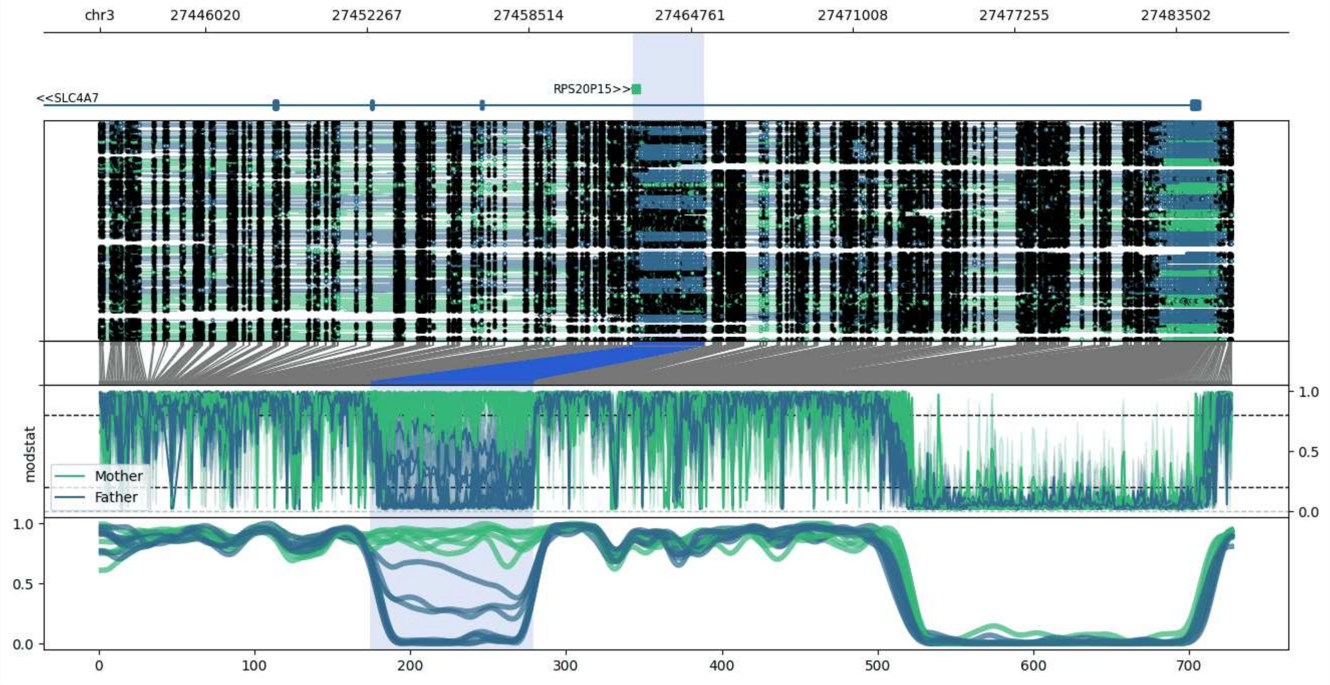


**chr3:27441948-27485682:** Region with 1 DMR containing genes: AC099535.1, RPS20P15, SLC4A7.

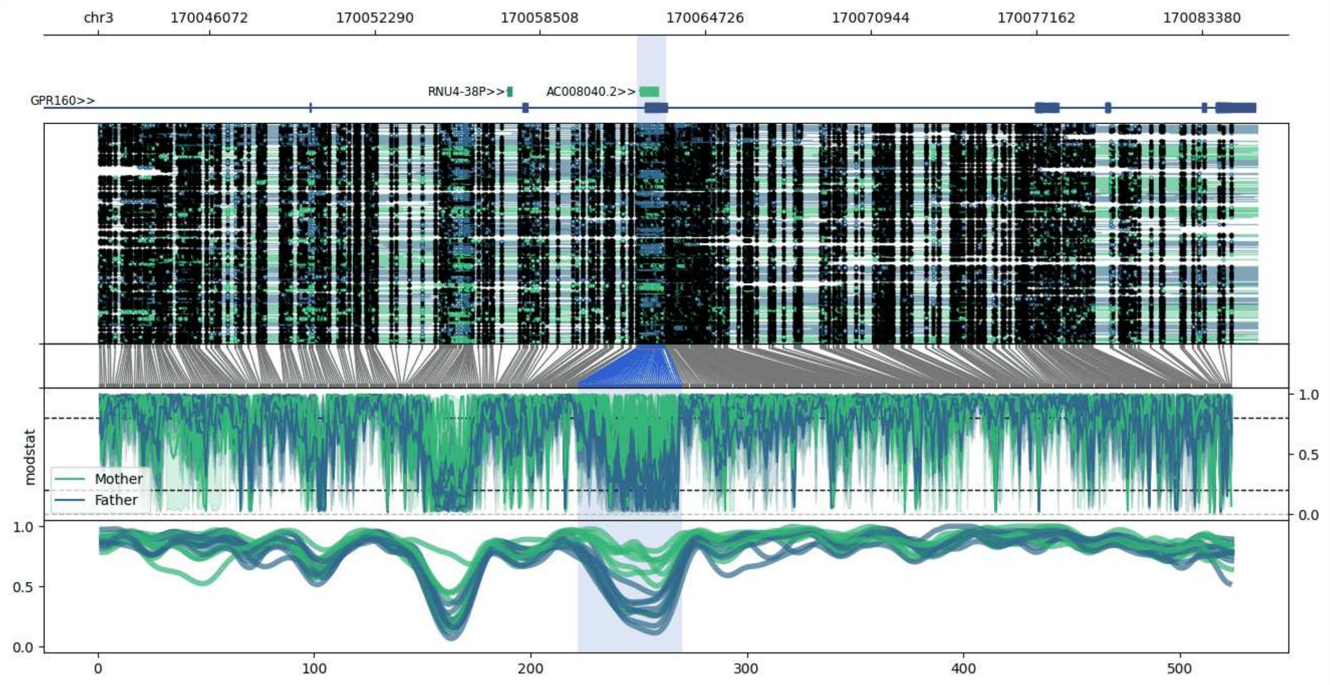


**chr3:170041913-170085445:** Region with 1 DMR containing genes: AC008040.2, GPR160, PHC3, RNU4-38P.

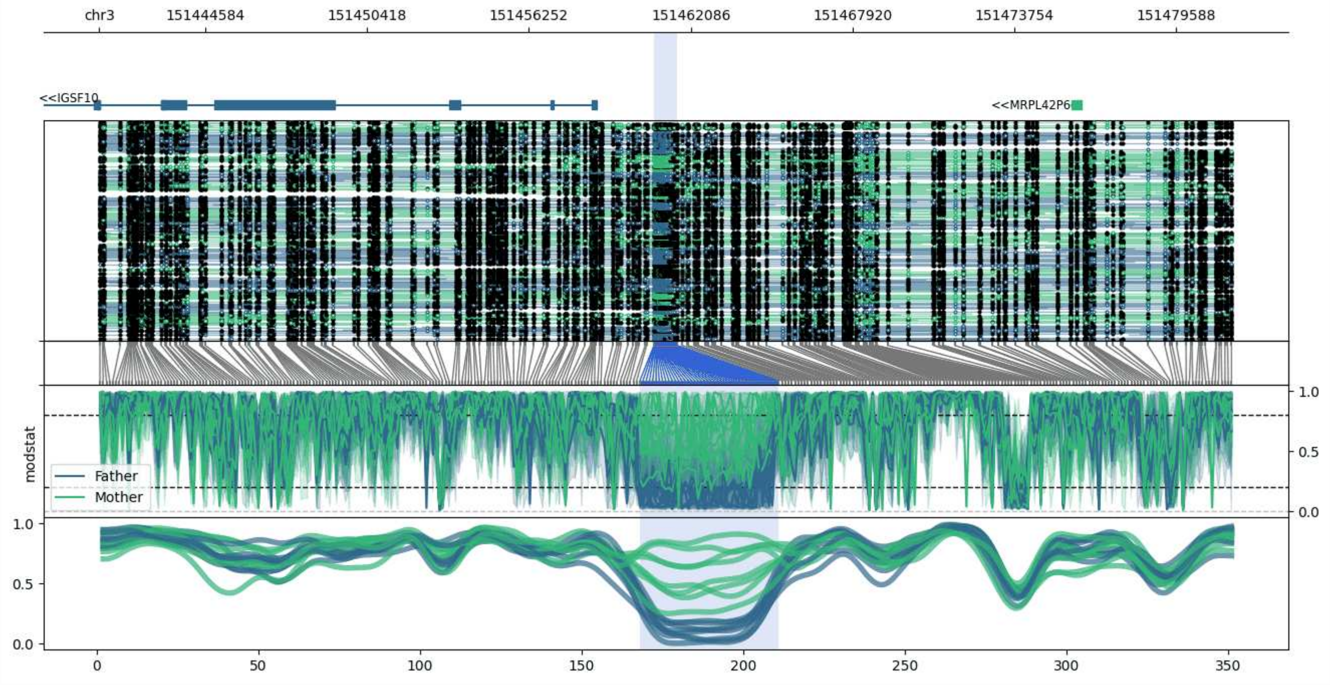


**chr3:151440761-151481600:** Region with 1 DMR containing genes: IGSF10, MED12L, MRPL42P6.

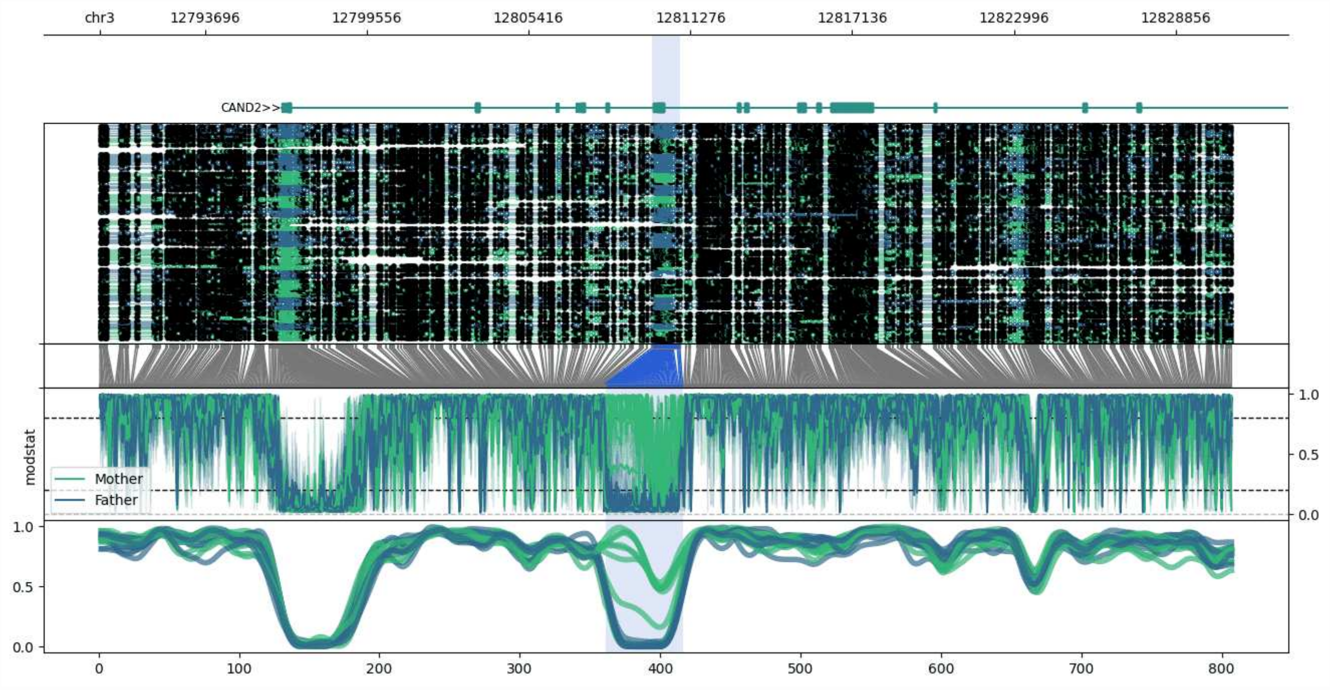


**chr3:12789869-12830895:** Region with 1 DMR containing genes: AC034198.2, CAND2, KRT18P17, RPL32, SNORA7A.

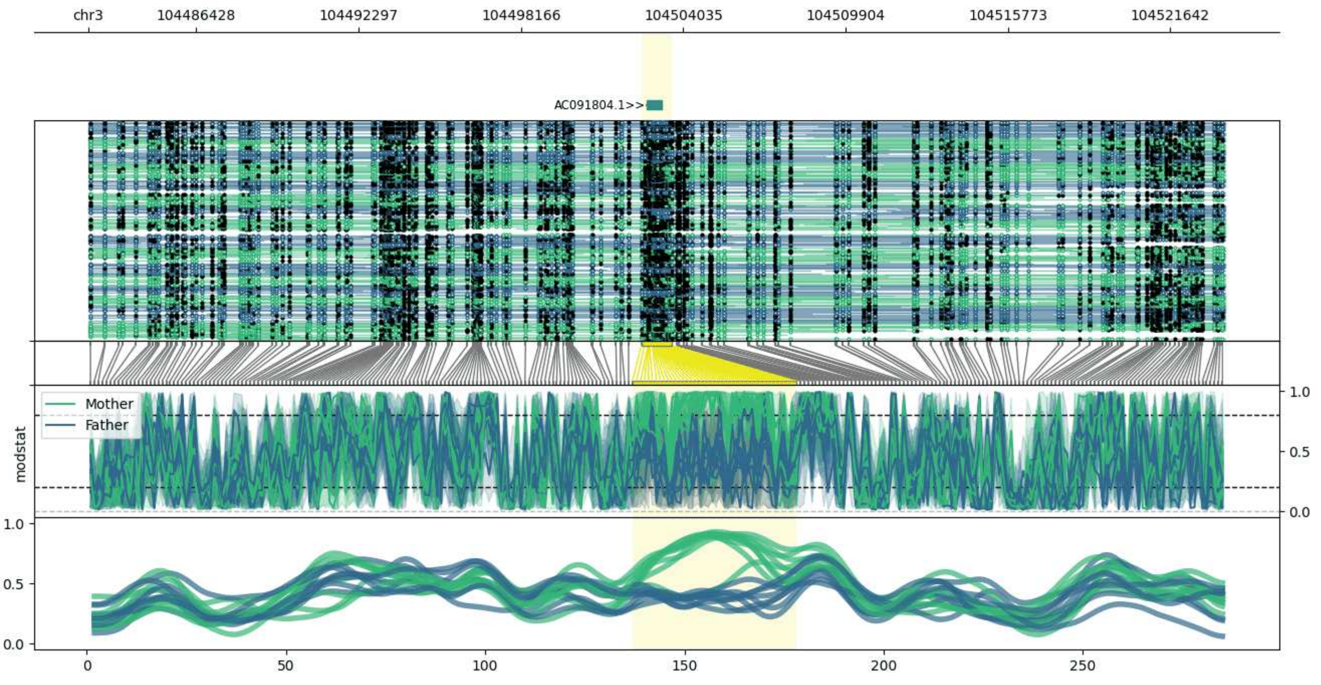


**chr3:104482519-104523608:** Region with 1 DMR containing genes: AC091804.1.

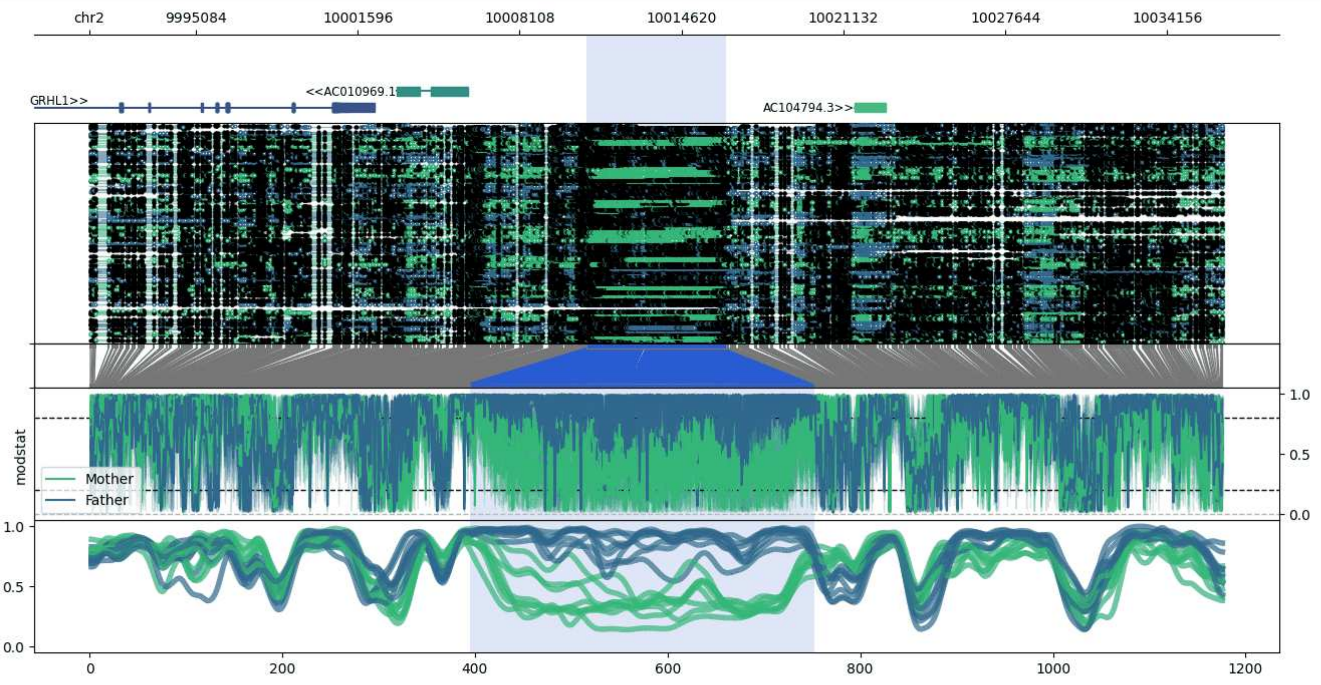


**chr2:9990811-10036401:** Region with 1 DMR containing genes: AC010969.1, AC104794.2, AC104794.3, GRHL1, KLF11.

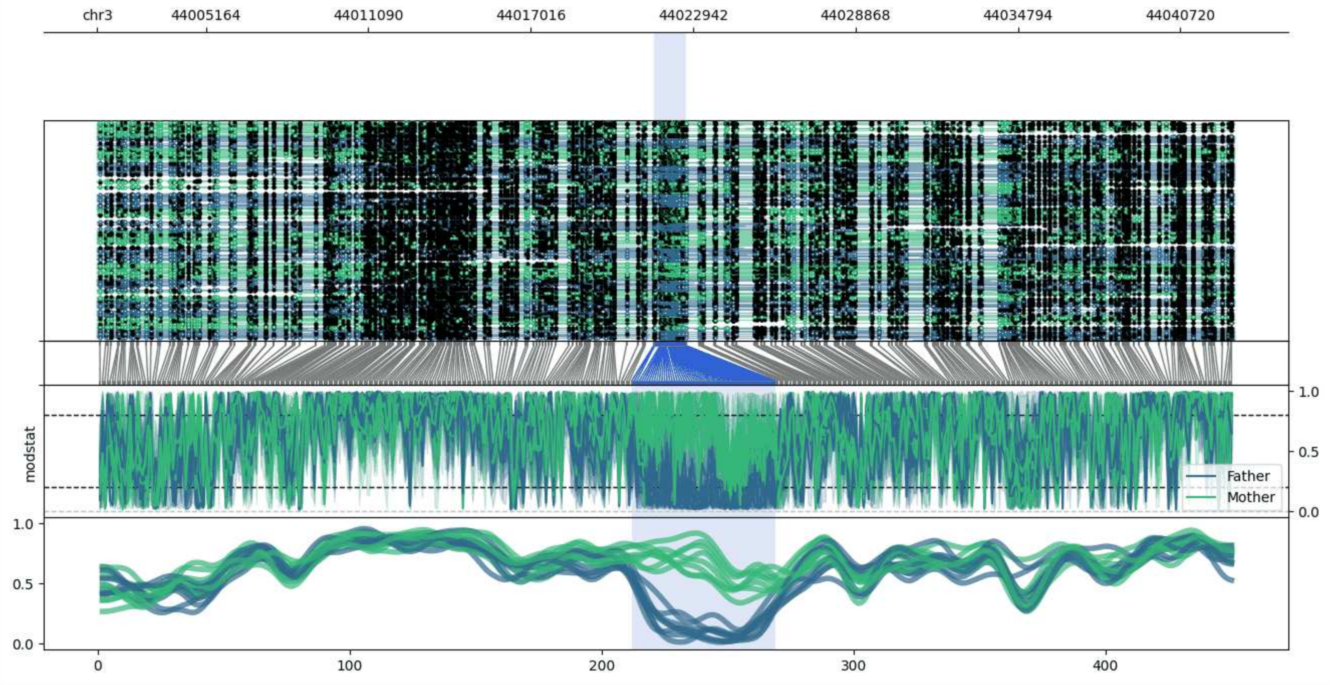


**chr3:44001198-44042683:** Region with 1 DMR containing genes: AC006058.2, AC006058.3.

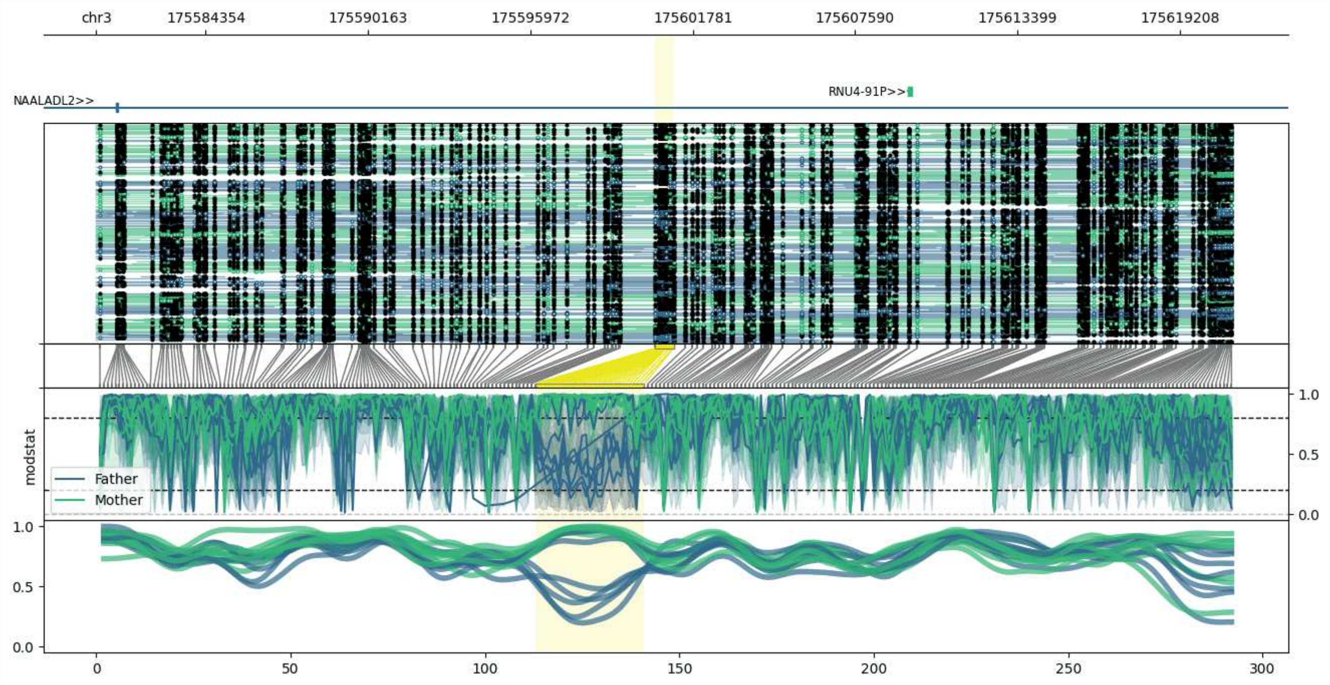


**chr3:175580431-175621094:** Region with 1 DMR containing genes: NAALADL2, RNU4-91P.

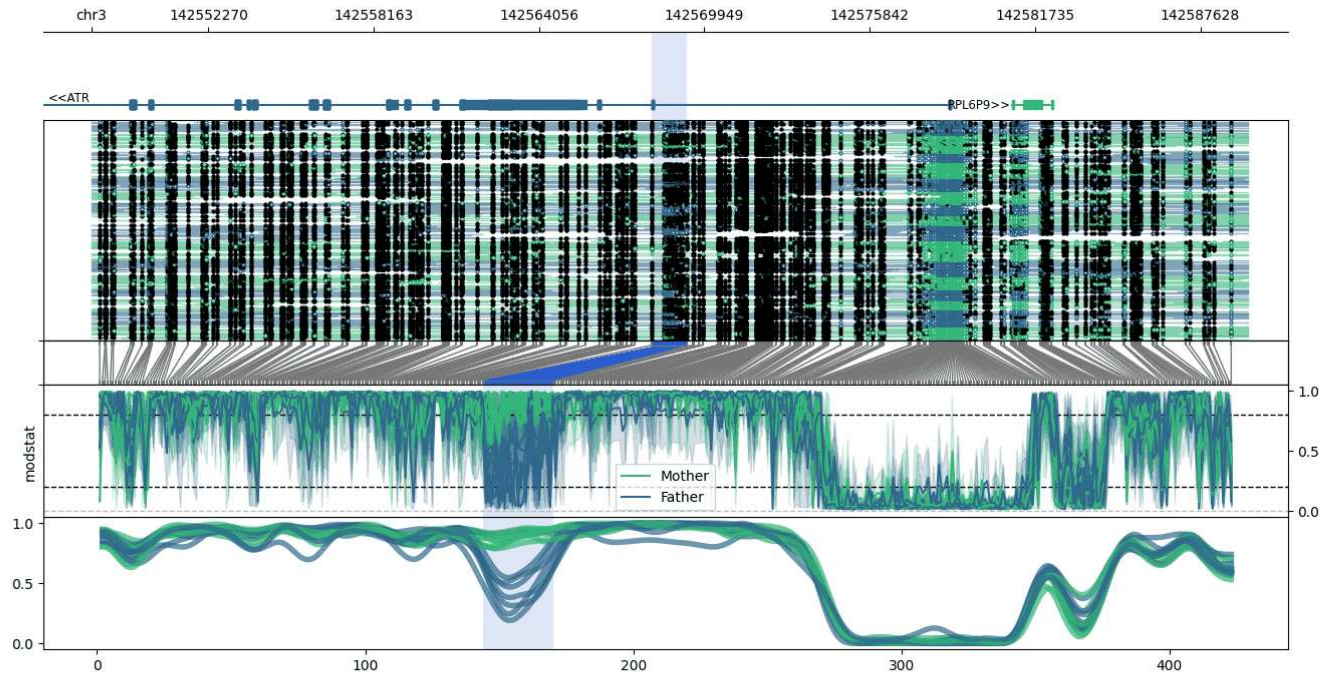


**chr3:142548086-142589340:** Region with 1 DMR containing genes: ATR, PLS1, RNA5SP143, RPL6P9.

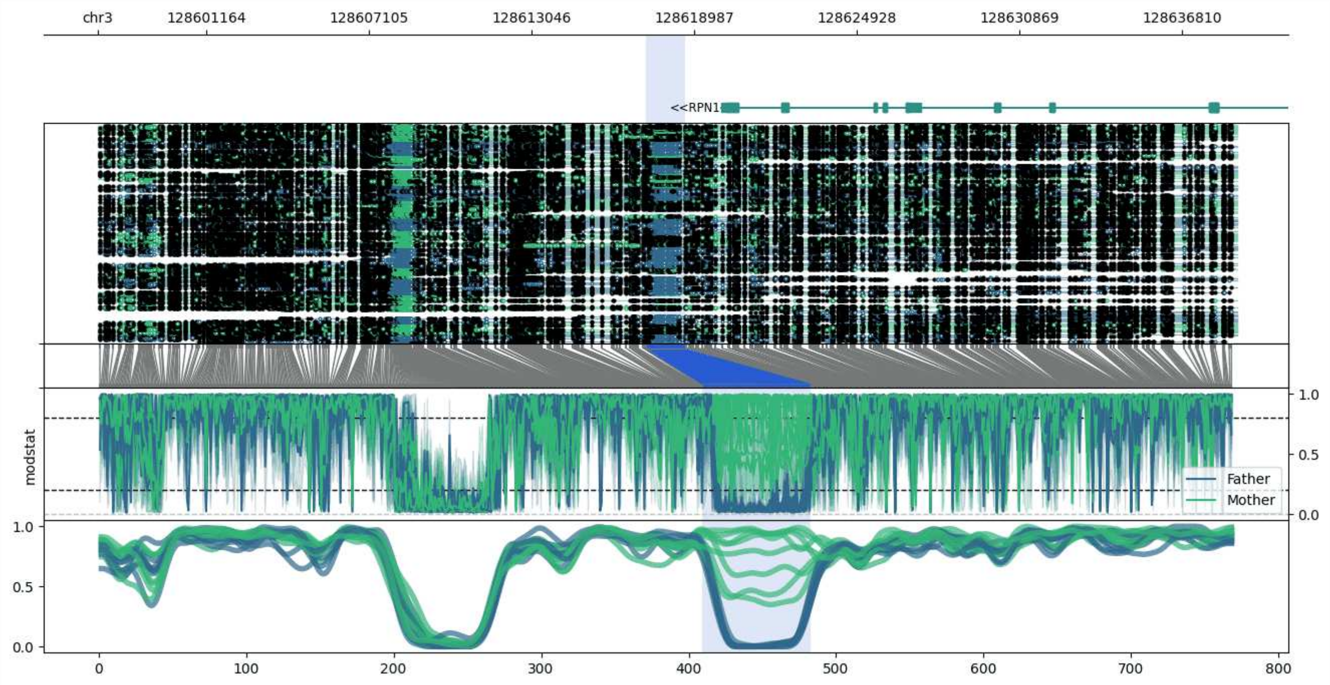


**chr3:128597209-128638799:** Region with 1 DMR containing genes: RPN1.

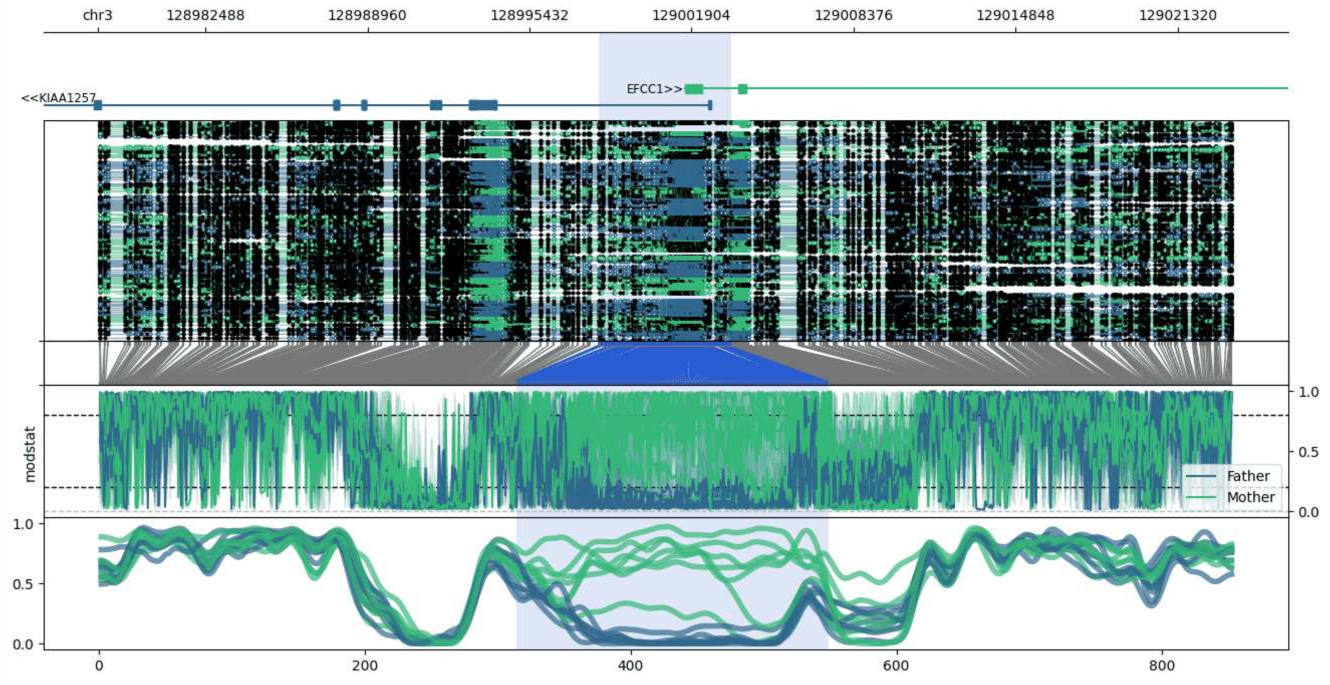


**chr3:128978186-129023490:** Region with 1 DMR containing genes: EFCC1, KIAA1257.

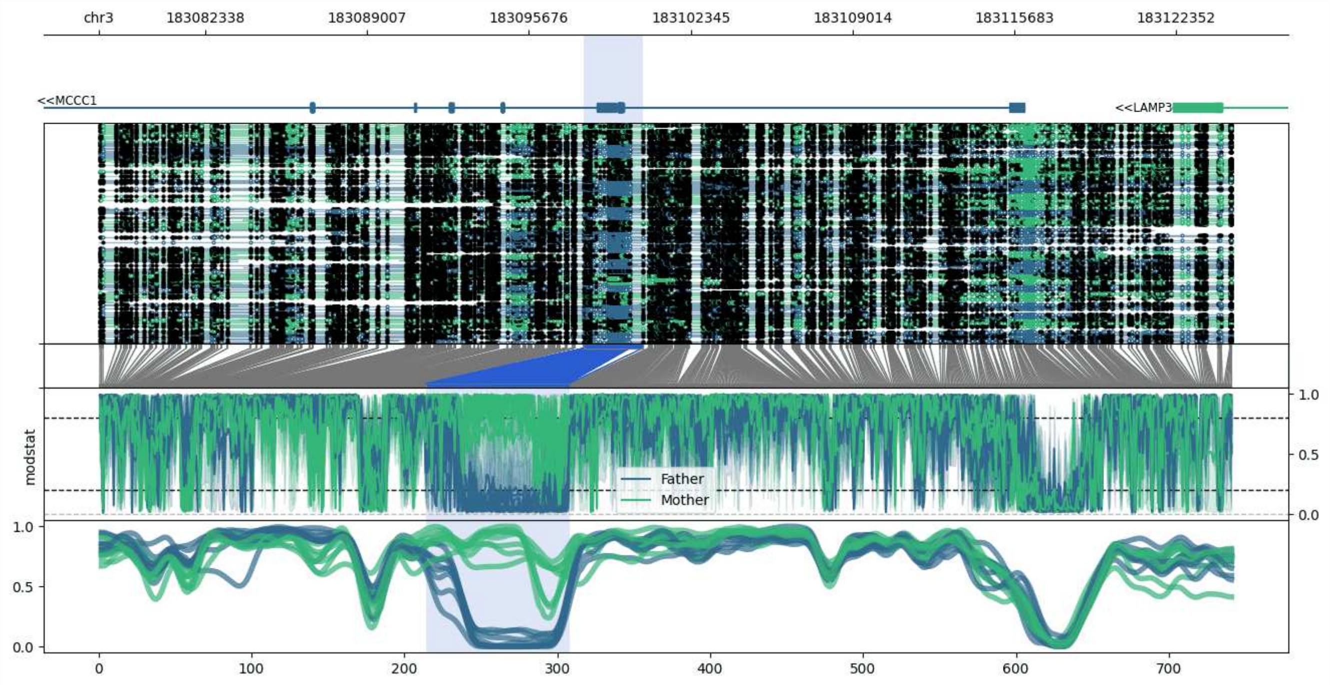


**chr3:183077936-183124622:** Region with 1 DMR containing genes: LAMP3, MCCC1.

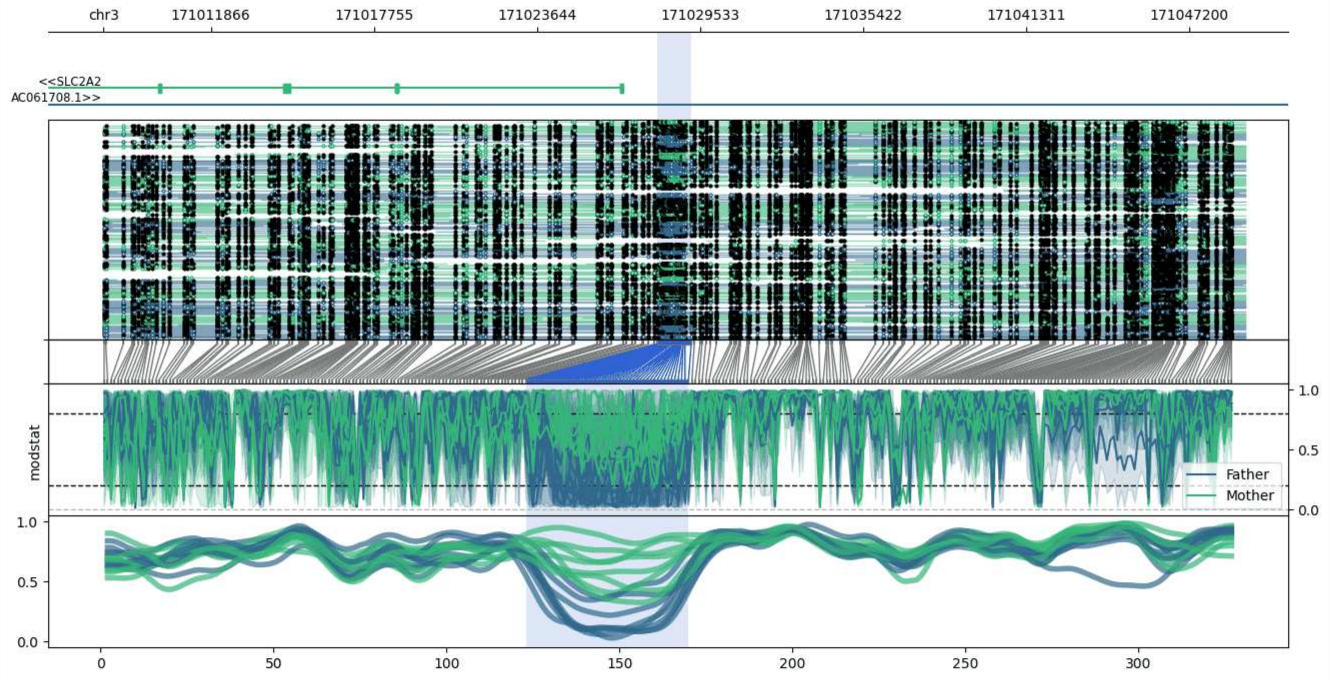


**chr3:171007986-171049214:** Region with 1 DMR containing genes: AC061708.1, SLC2A2, TNIK.

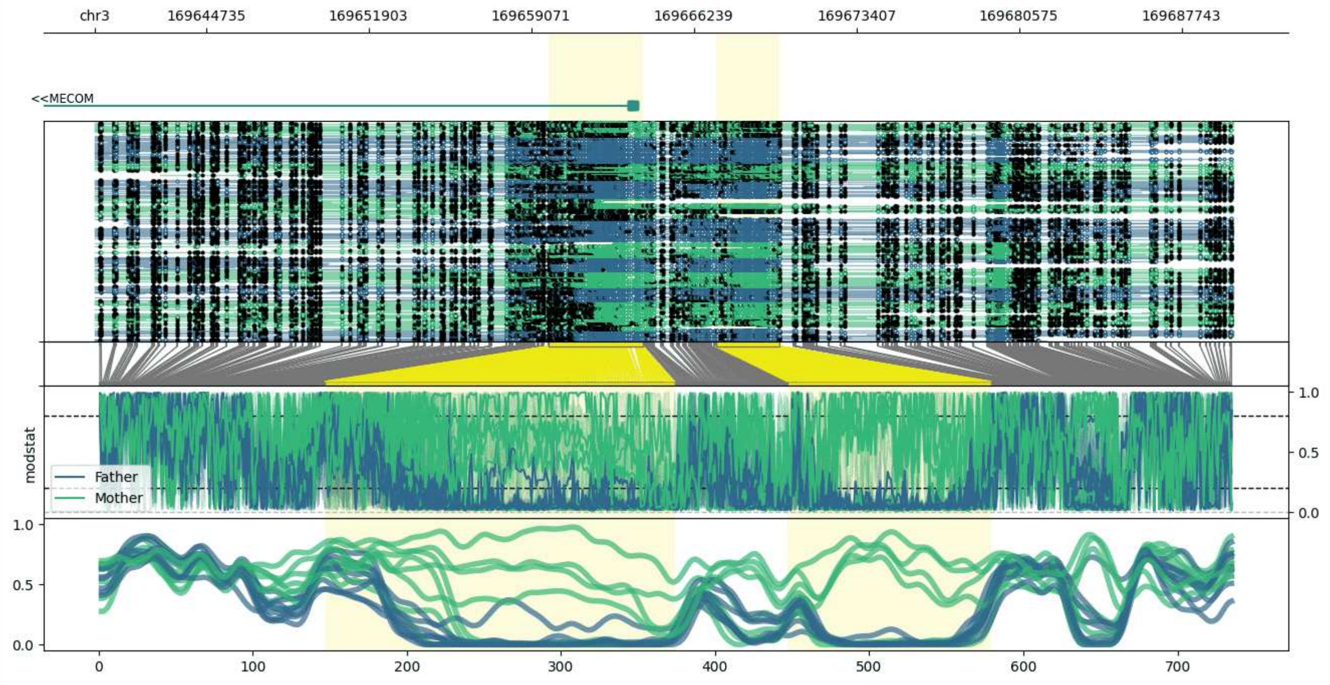


**chr3:169639832-169690010:** Region with 2 DMRs containing genes: MECOM.

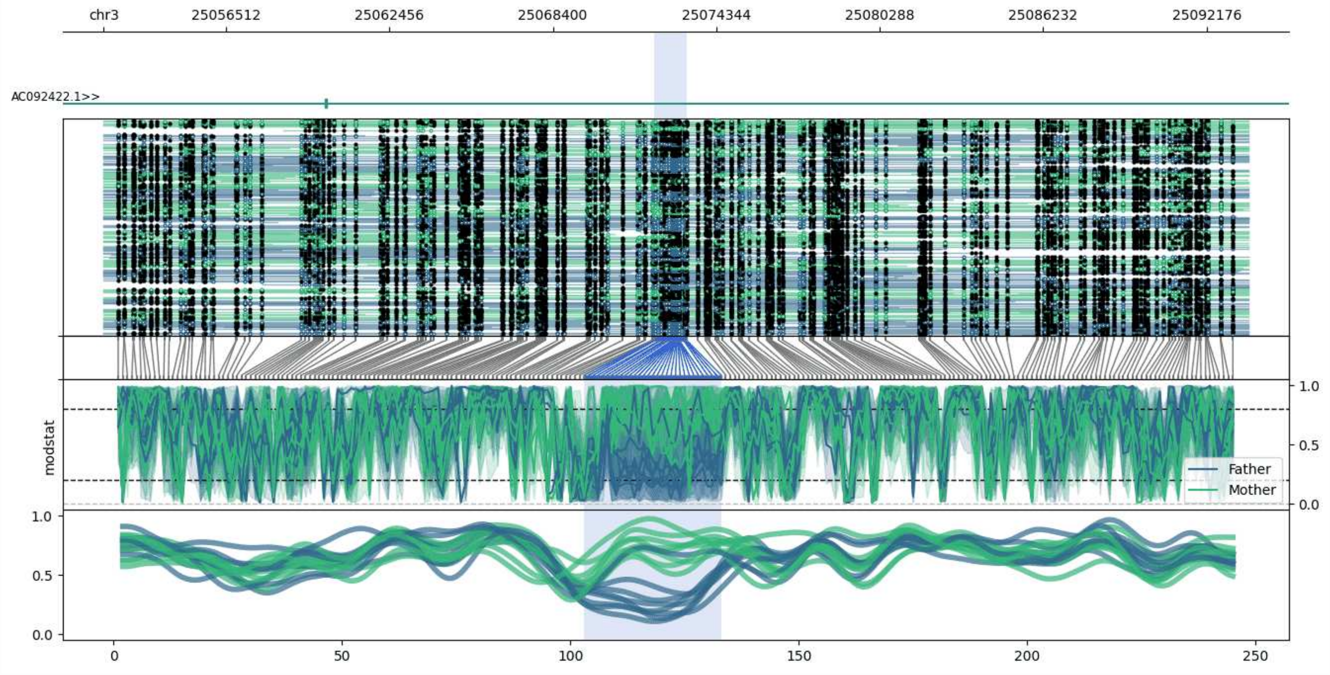


**chr3:25052070-25093679:** Region with 1 DMR containing genes: AC092422.1.

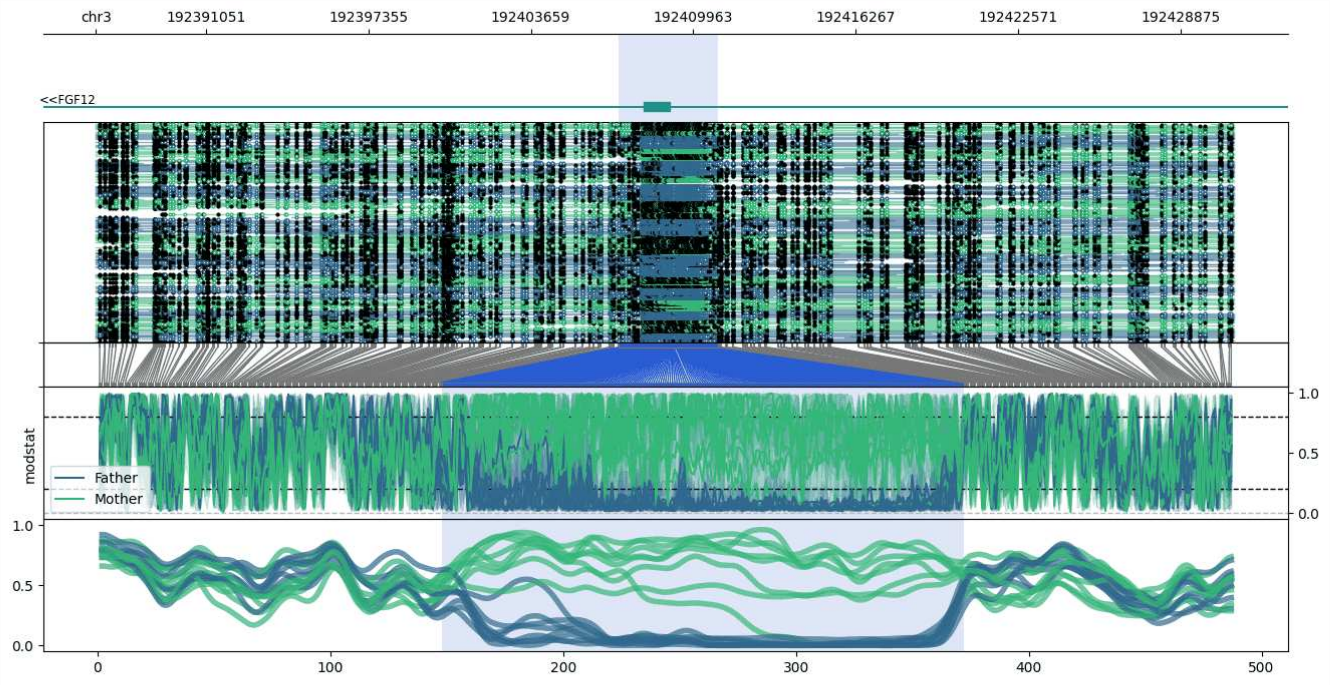


**chr3:192386800-192430932:** Region with 1 DMR containing genes: FGF12.

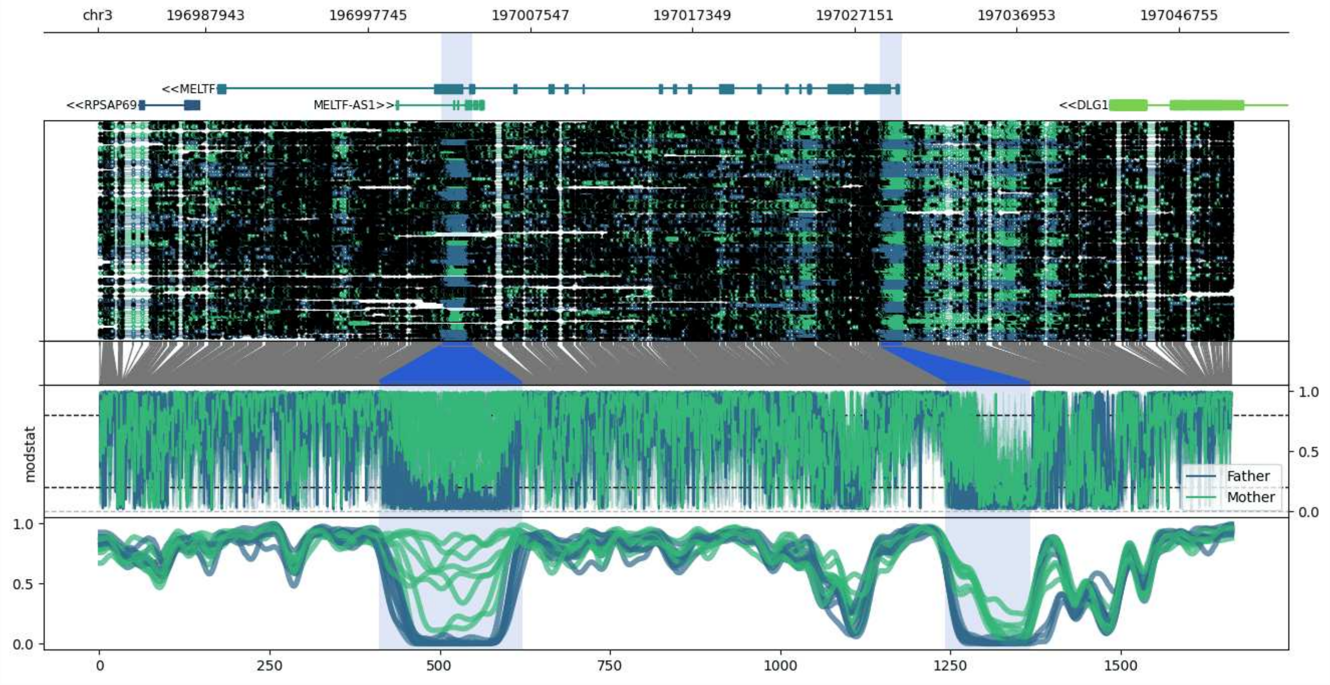


**chr3:196981410-197050028:** Region with 2 DMRs containing genes: AC011322.1, DLG1, MELTF, MELTF-AS1, RPSAP69.

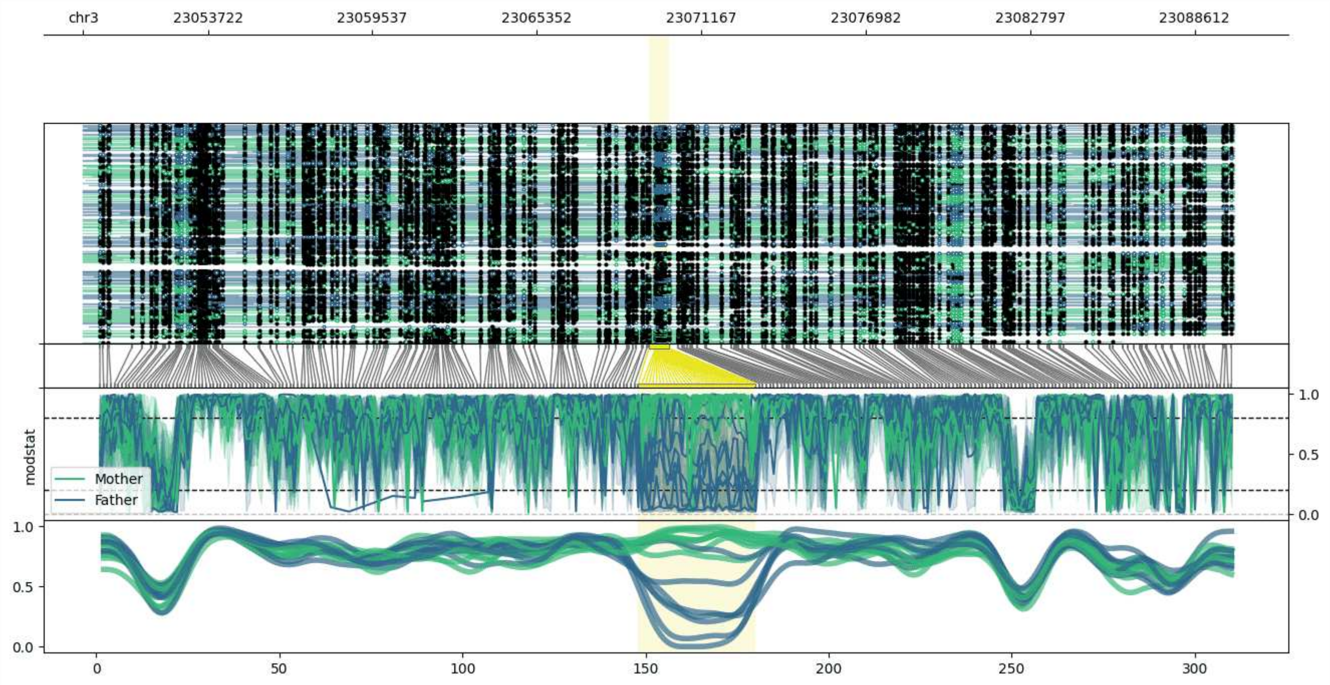


**chr3:23049315-23090020:** Region with 1 DMR containing no genes.

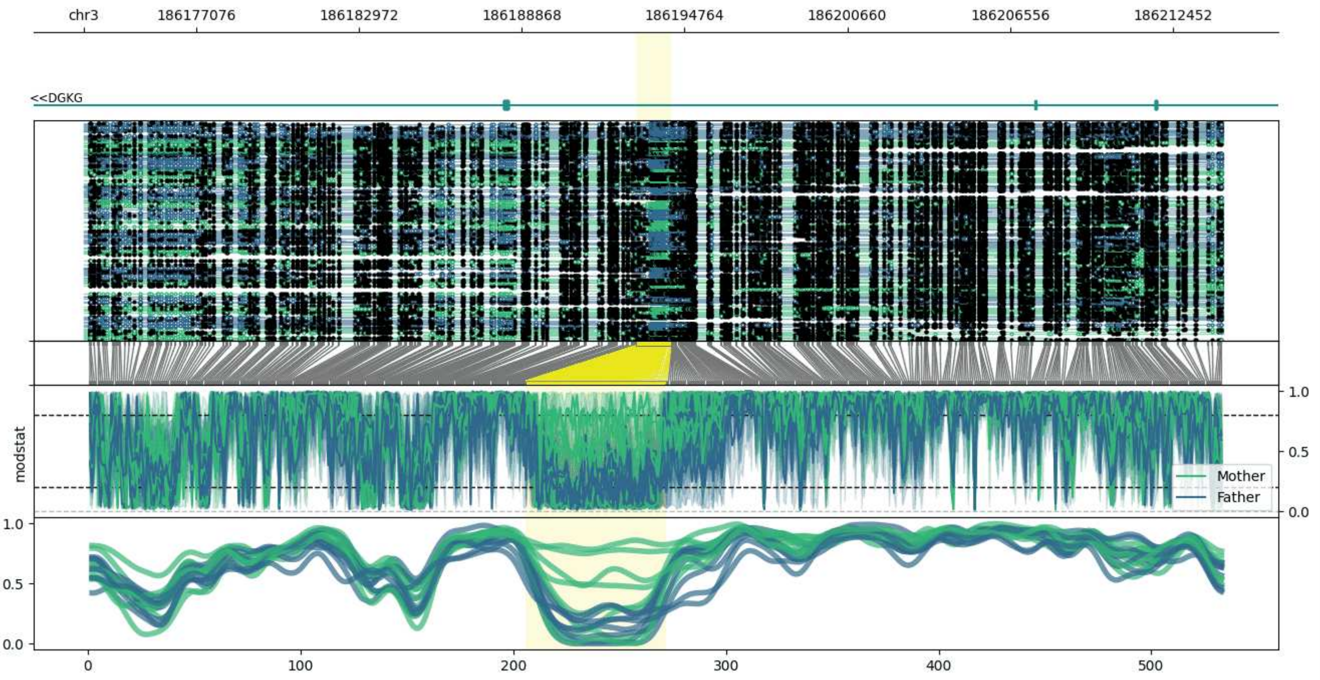


**chr3:186173011-186214288:** Region with 1 DMR containing genes: DGKG.

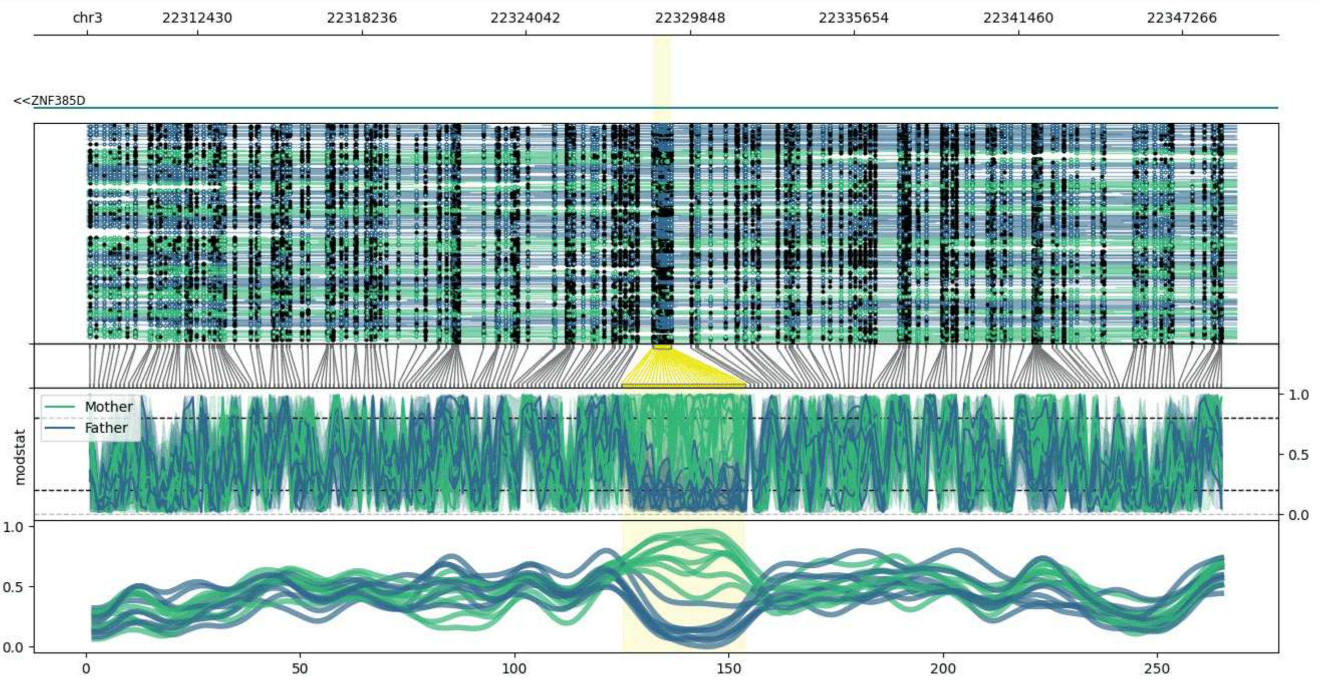


**chr3:22308529-22349177:** Region with 1 DMR containing genes: ZNF385D.

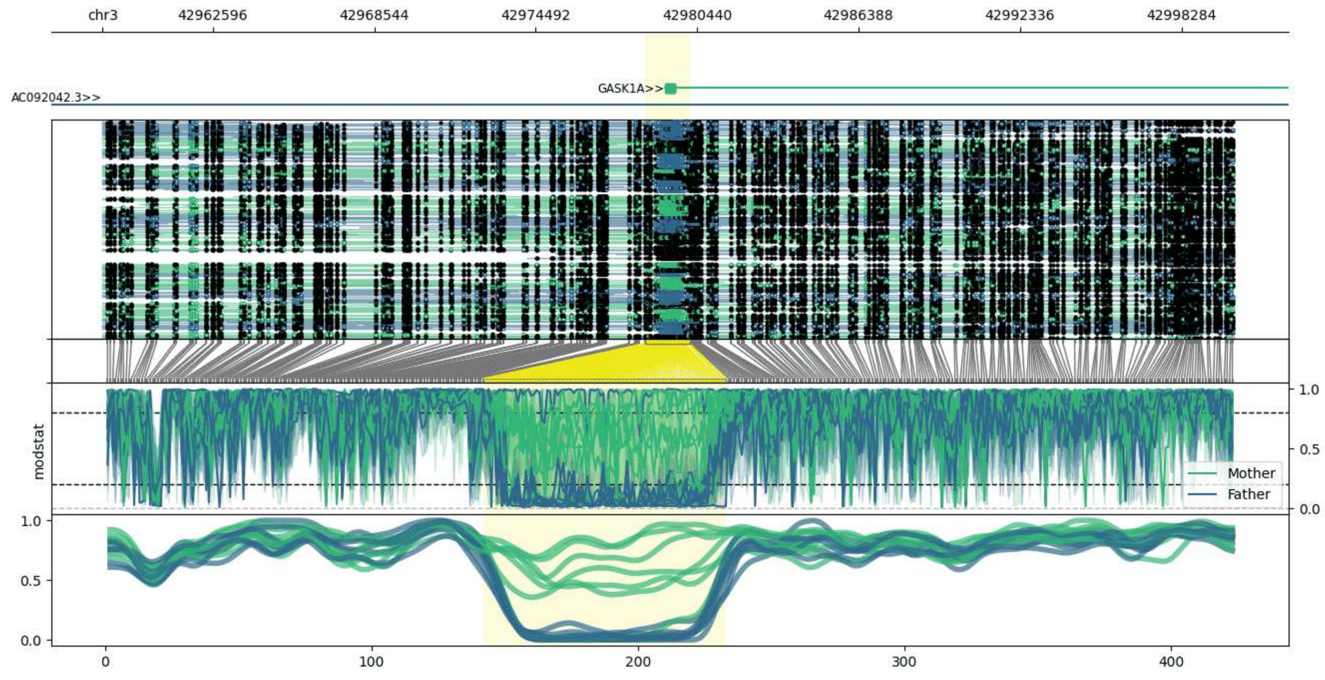


**chr3:42958539-43000180:** Region with 1 DMR containing genes: AC092042.3, GASK1A.

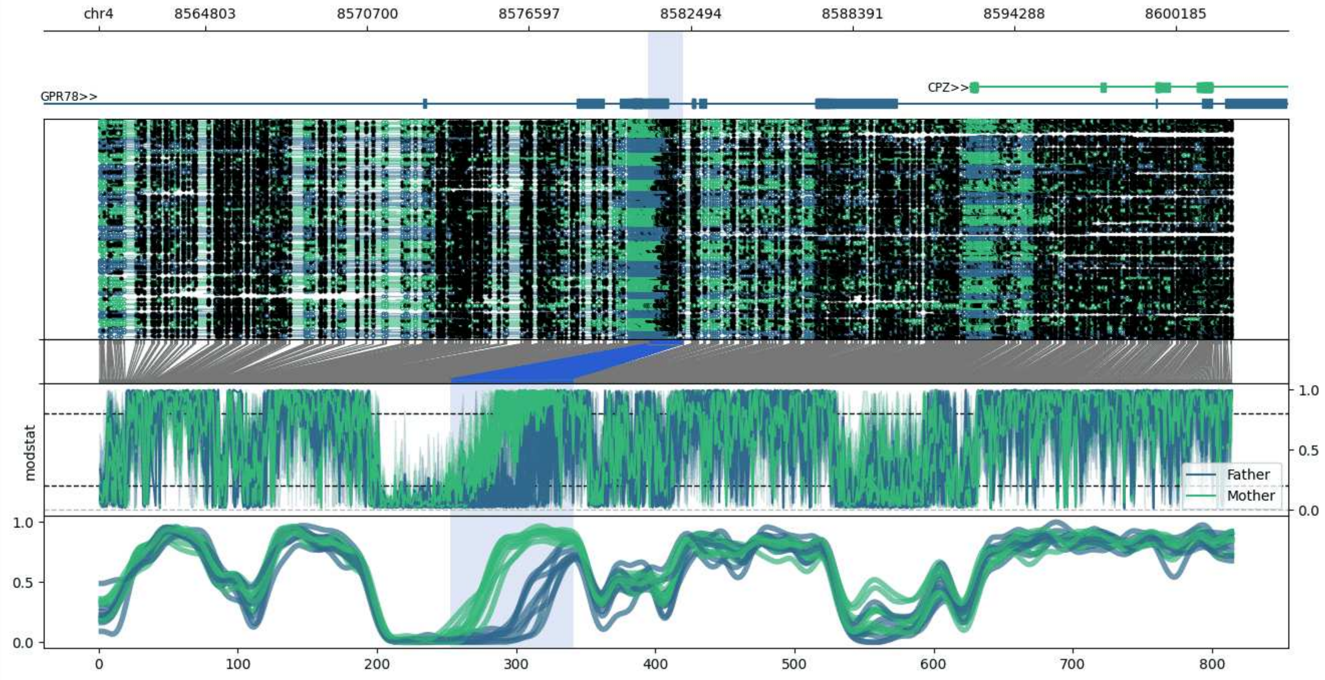


**chr4:8560927-8602209:** Region with 1 DMR containing genes: CPZ, GPR78.

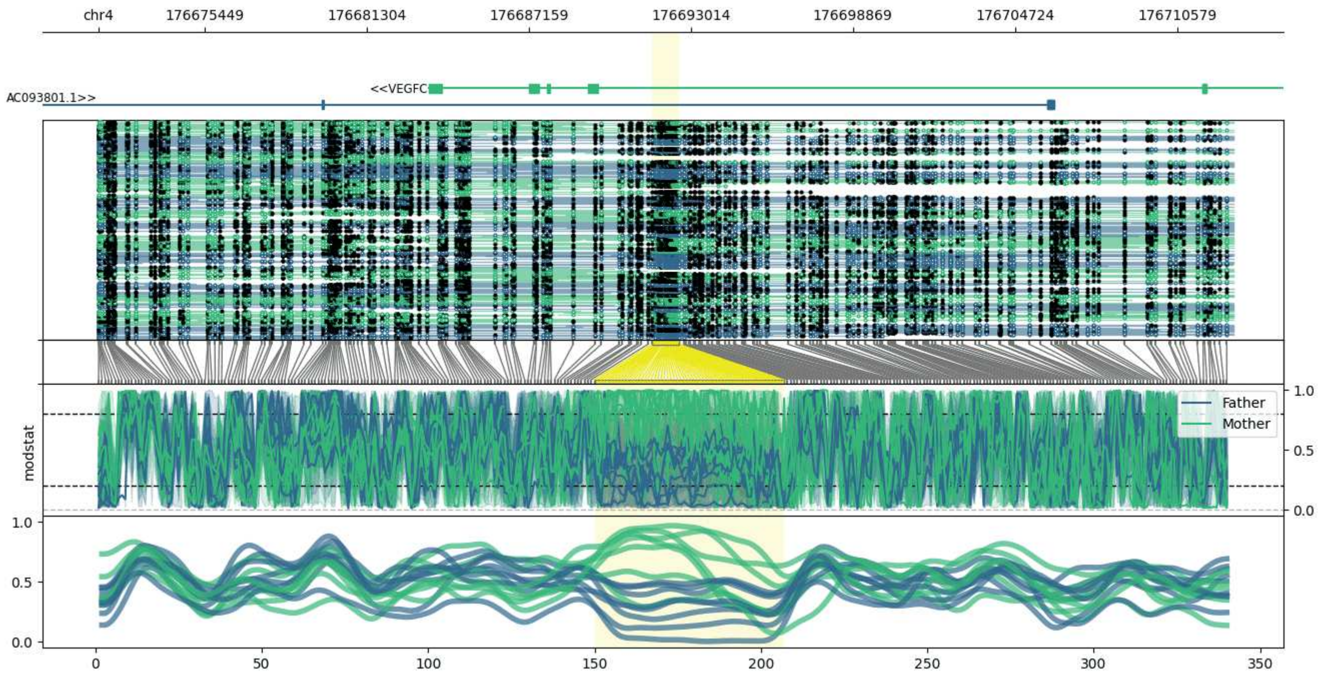


**chr4:176671614-176712604:** Region with 1 DMR containing genes: AC093801.1, VEGFC.

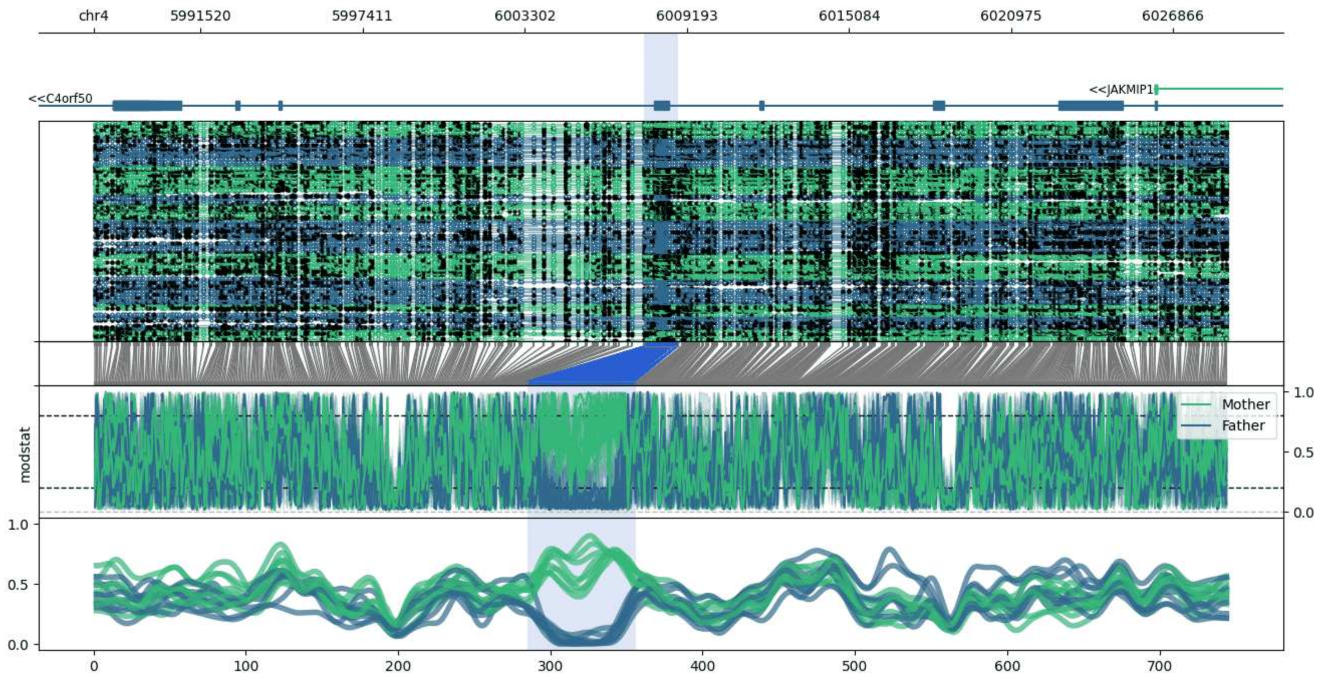


**chr4:5987636-6028873:** Region with 1 DMR containing genes: C4orf50, JAKMIP1.

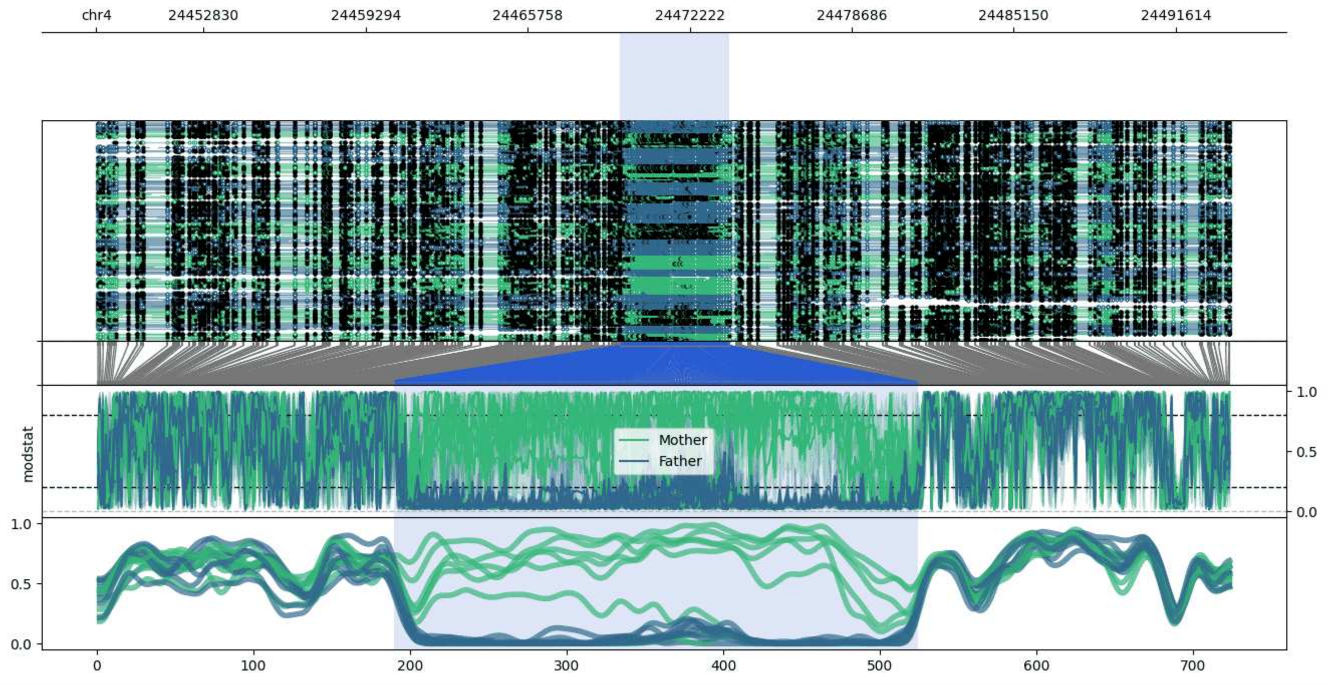


**chr4:24448558-24493806:** Region with 1 DMR containing no genes.

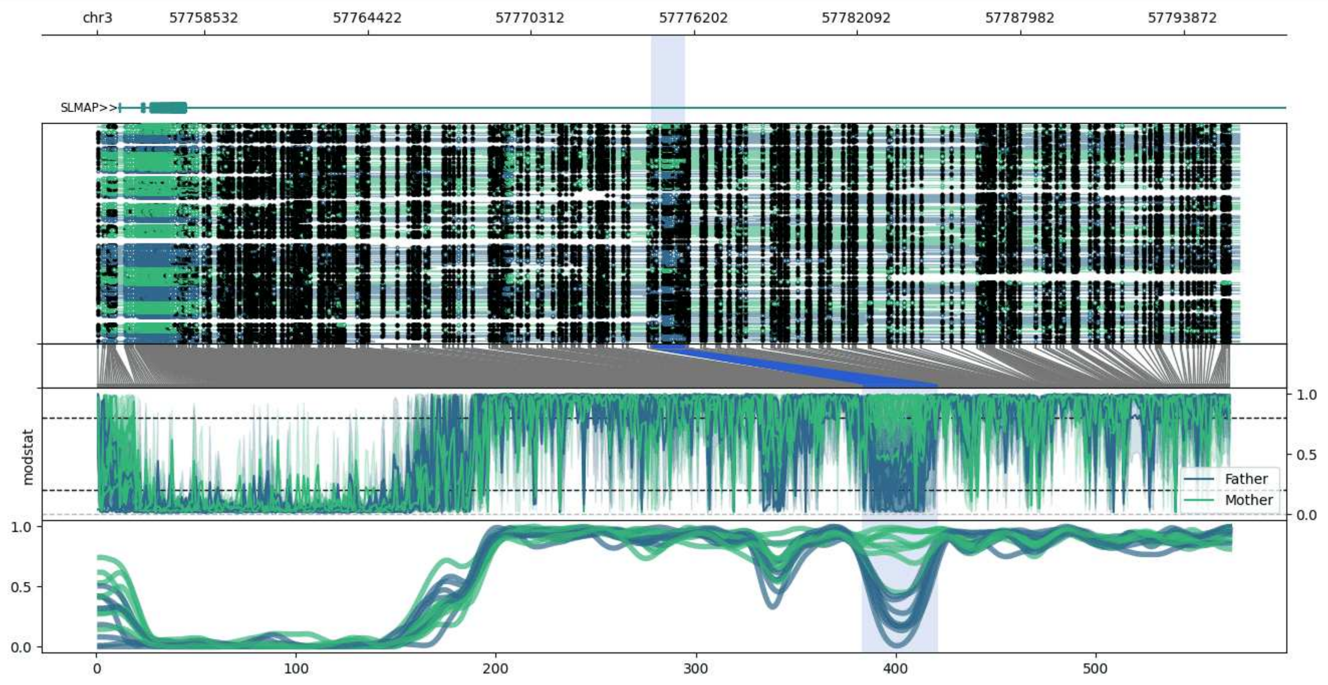


**chr3:57754655-57795889:** Region with 1 DMR containing genes: PDHA1P1, SLMAP.

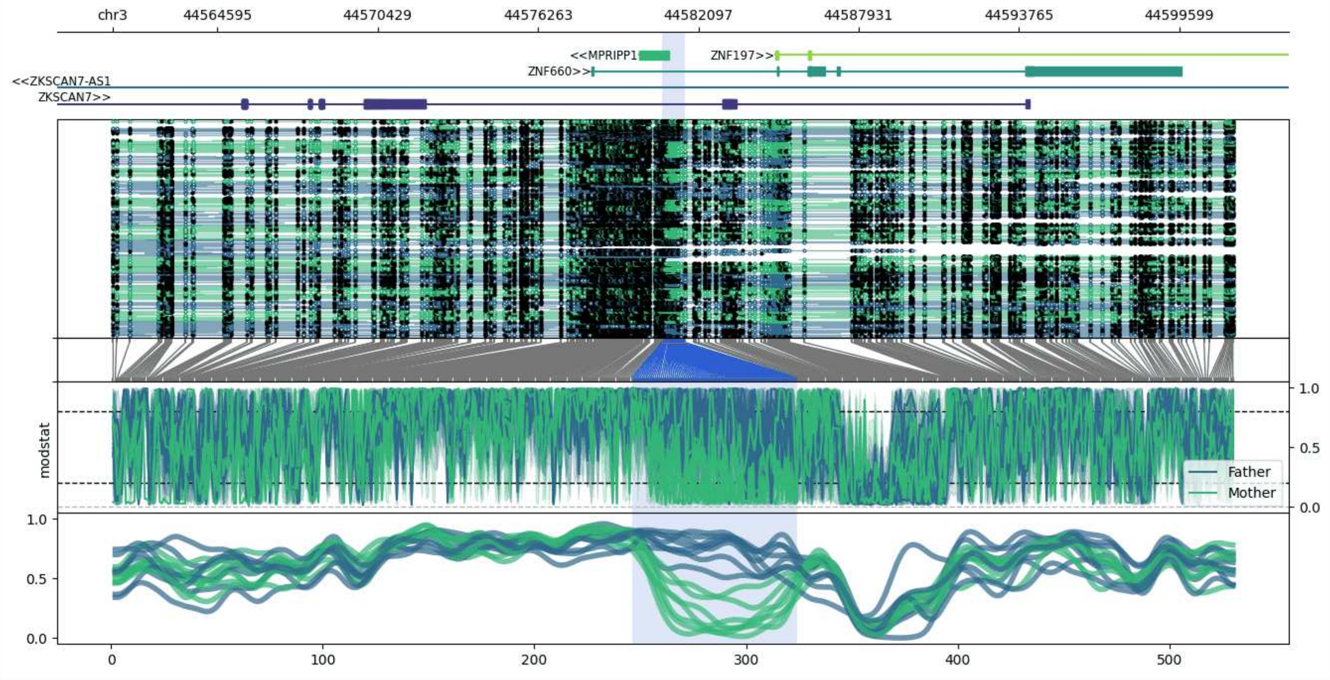


**chr3:44560773-44601612:** Region with 1 DMR containing genes: MPRIPP1, ZKSCAN7, ZKSCAN7-AS1, ZNF197, ZNF660.

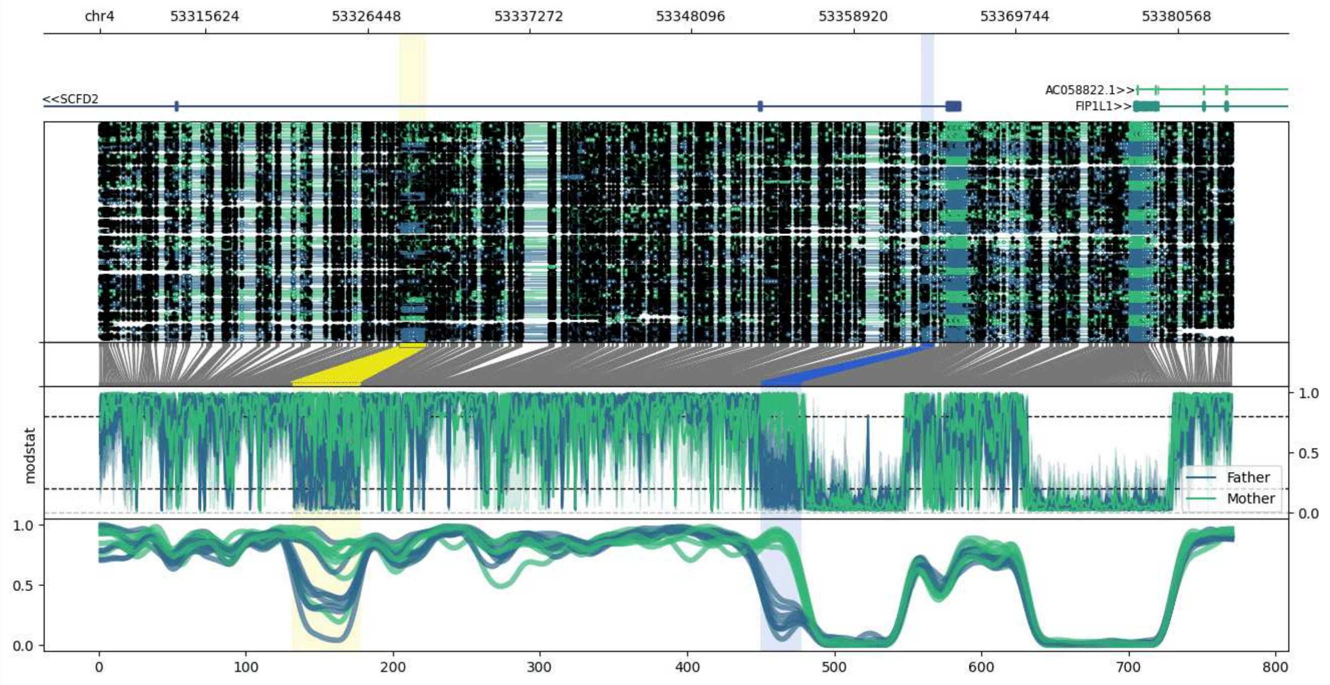


**chr4:53308549-53384319:** Region with 2 DMRs containing genes: AC058822.1, FIP1L1, SCFD2.

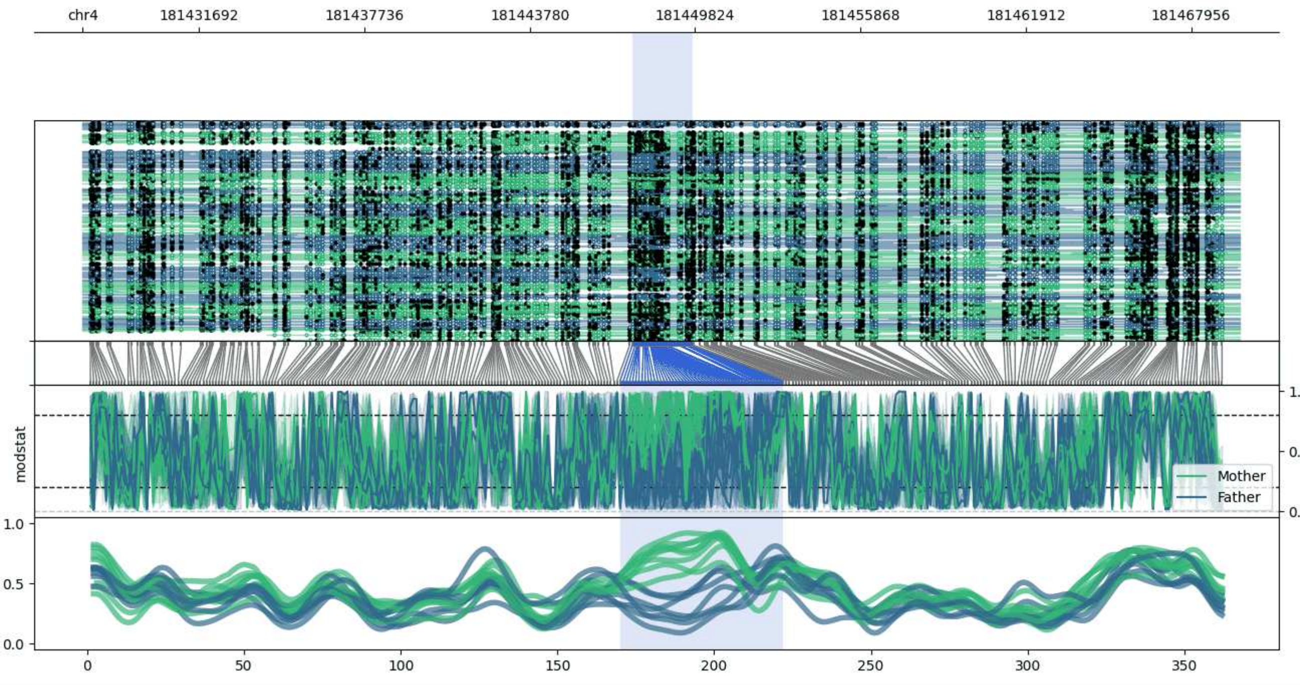


**chr4:181427430-181469740:** Region with 1 DMR containing no genes.

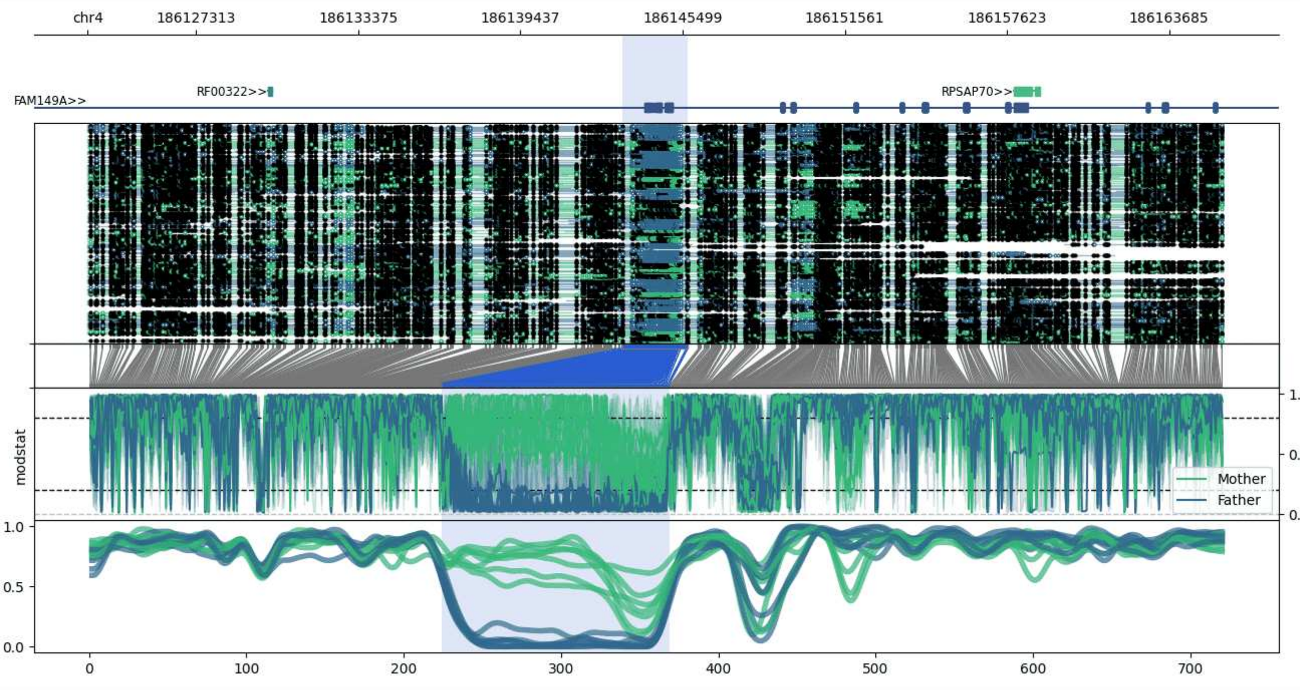


**chr4:186123260-186165699:** Region with 1 DMR containing genes: FAM149A, ORAOV1P1, RF00322, RPSAP70.

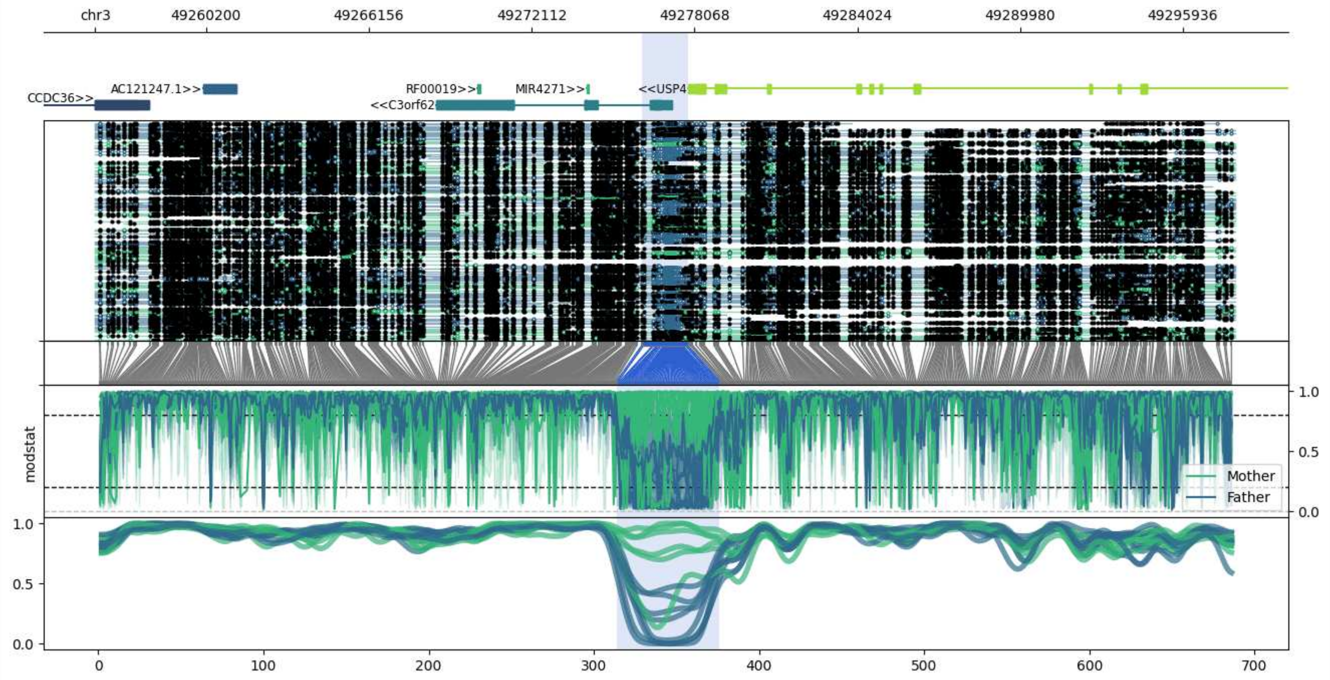


**chr3:49256138-49297831:** Region with 1 DMR containing genes: AC121247.1, AC121247.2, C3orf62, CCDC36, MIR4271, RF00019, USP4.

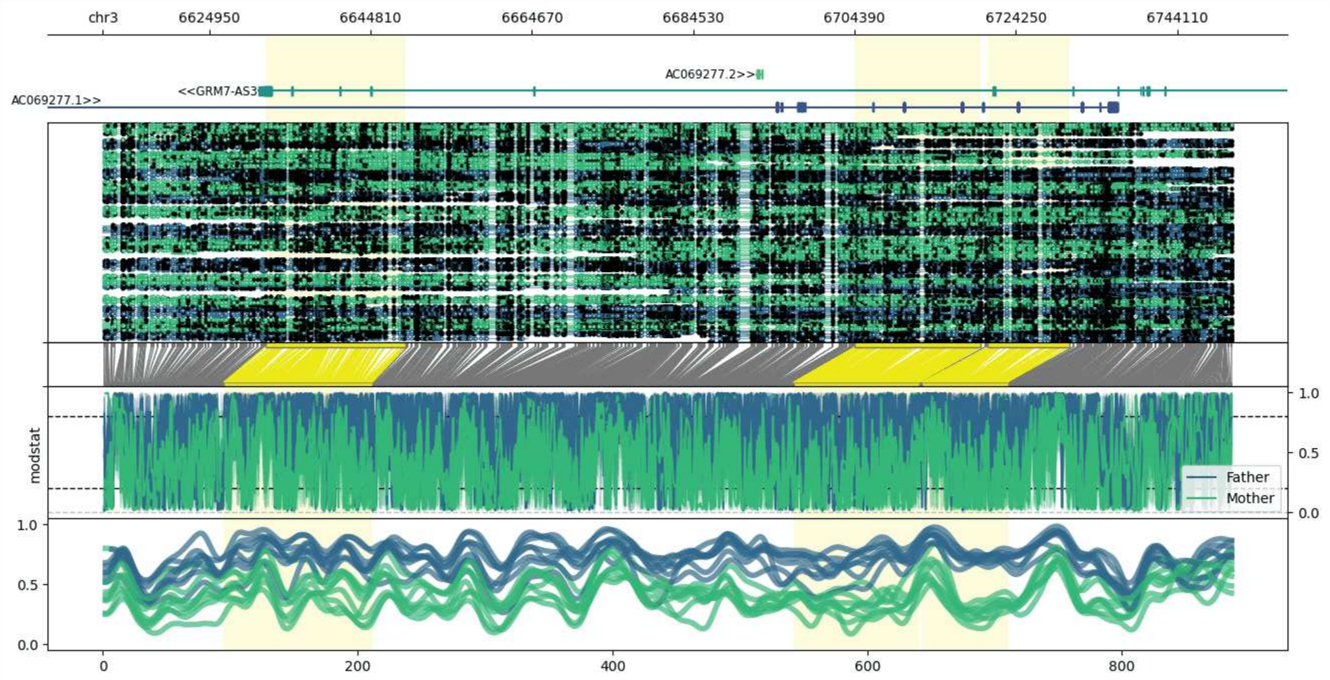


**chr3:6611873-6750897:** Region with 3 DMRs containing genes: AC069277.1, AC069277.2, GRM7-AS3.

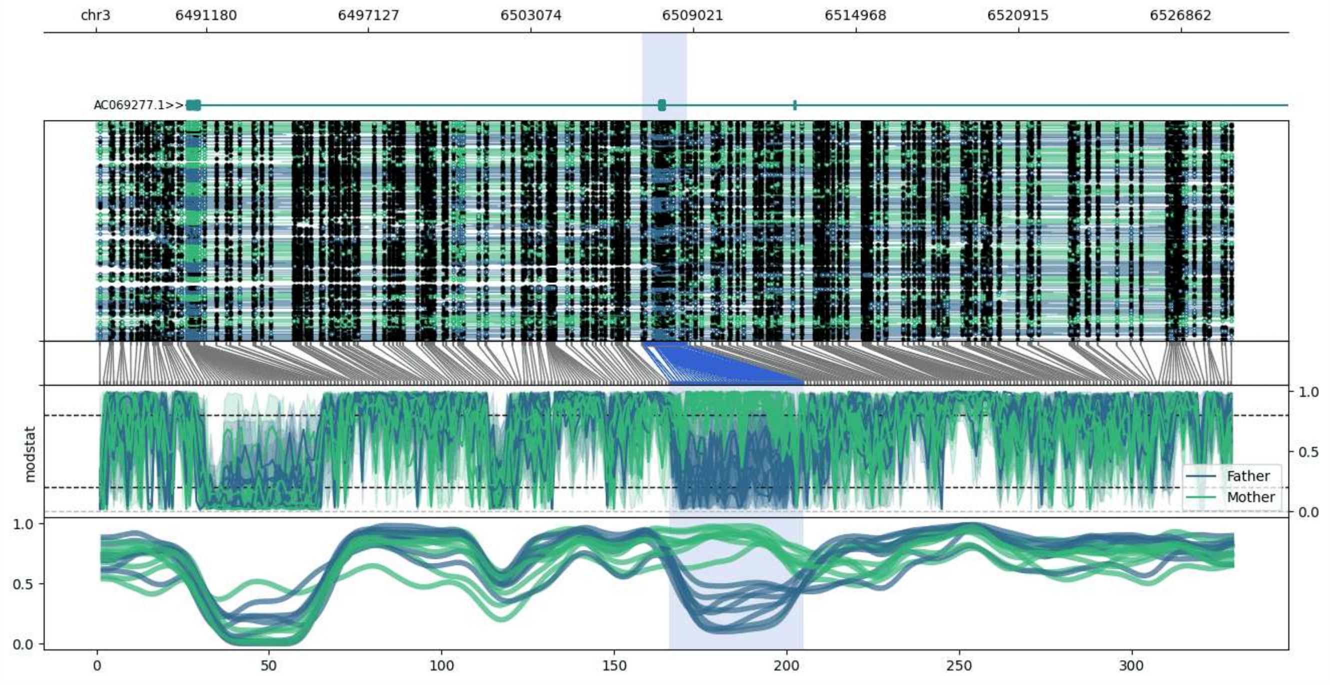


**chr3:6487146-6528777:** Region with 1 DMR containing genes: AC069277.1.

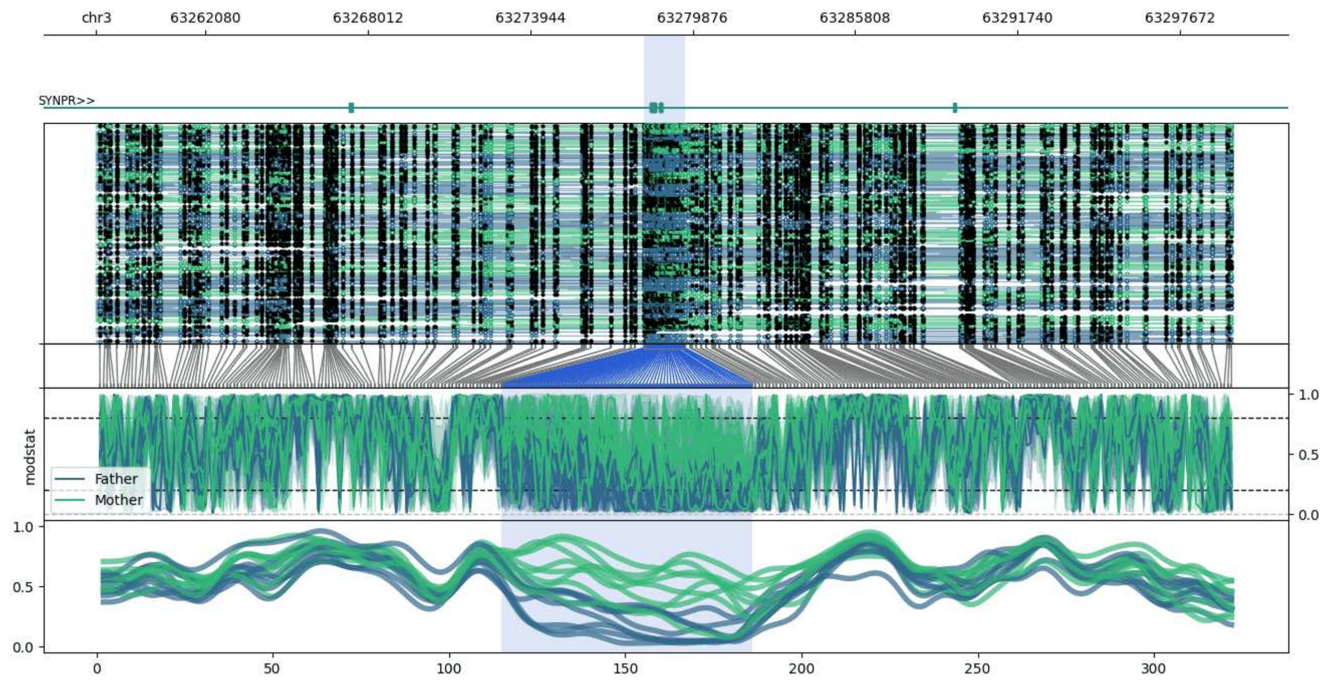


**chr3:63258075-63299602:** Region with 1 DMR containing genes: SYNPR.

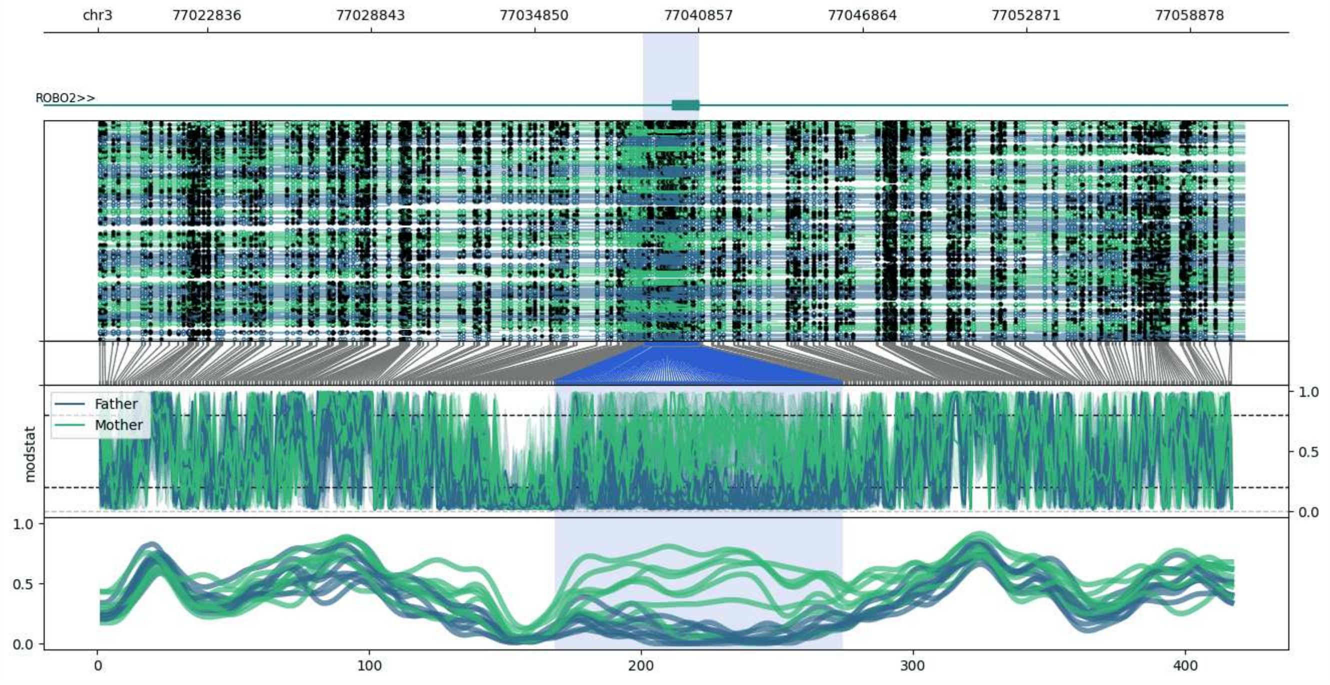


**chr3:77018824-77060876:** Region with 1 DMR containing genes: ROBO2.

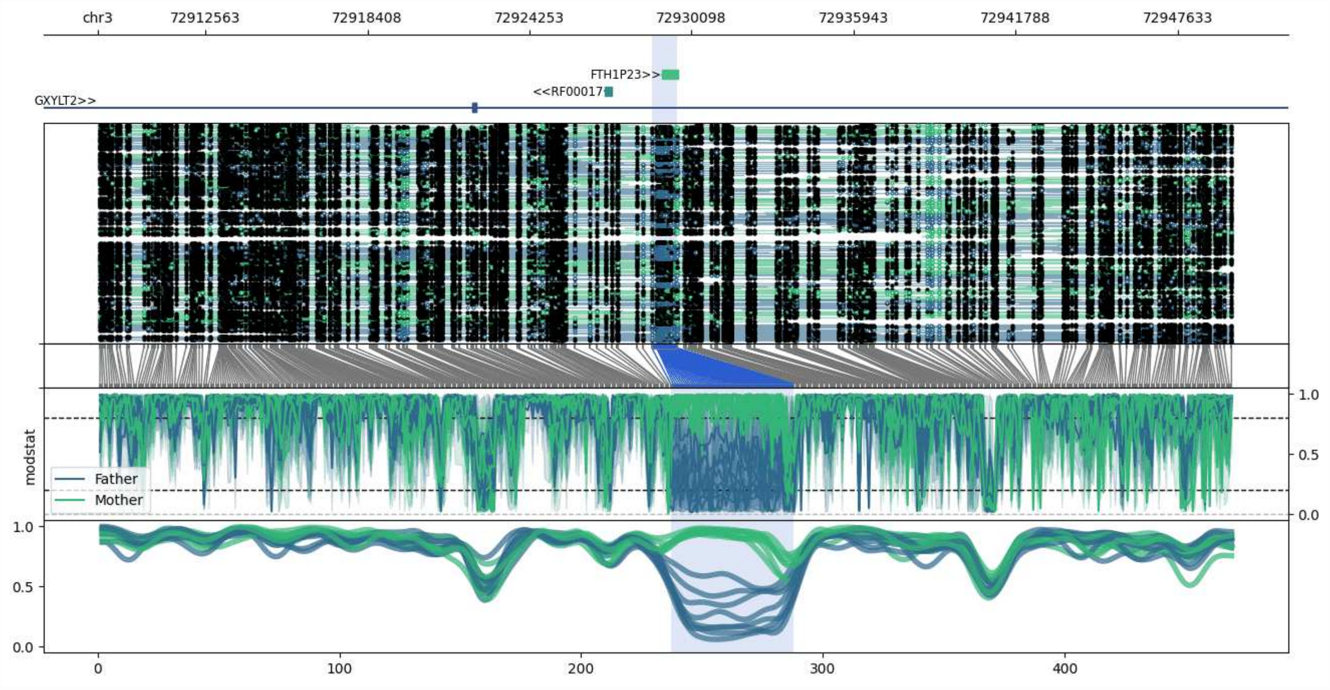


**chr3:72908668-72949587:** Region with 1 DMR containing genes: FTH1P23, GXYLT2, RF00017.

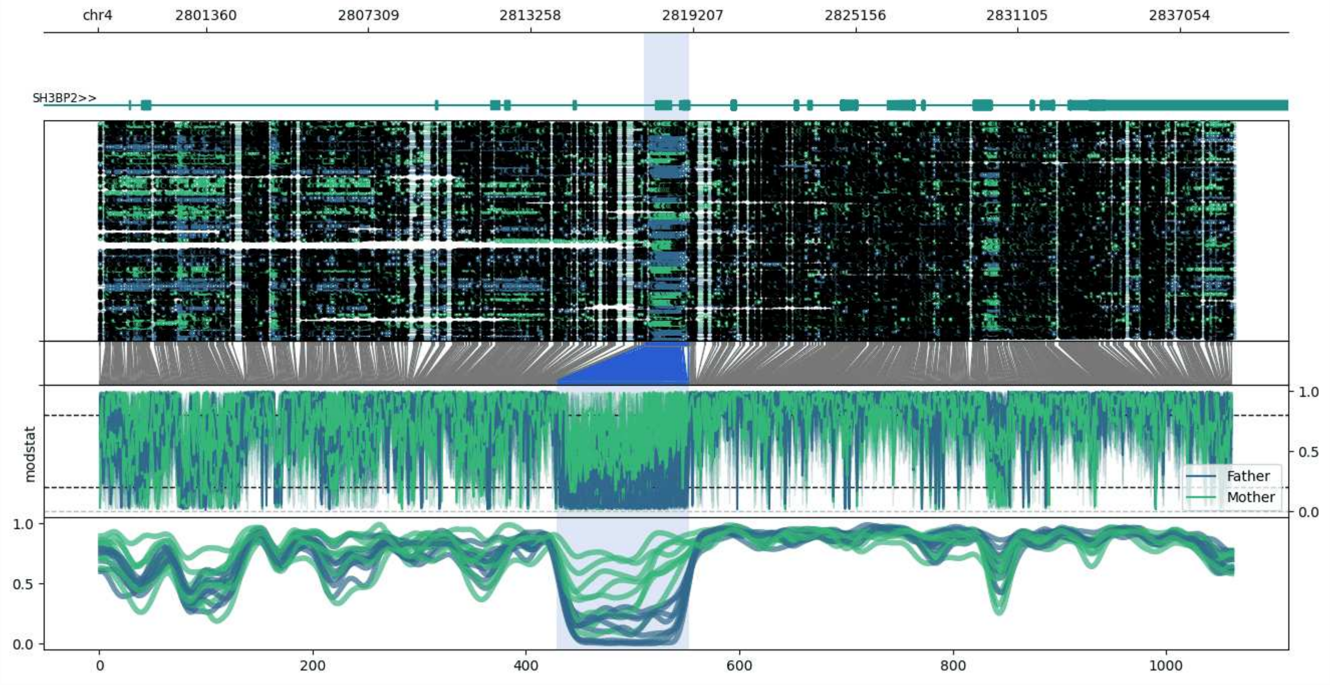


**chr4:2797413-2839058:** Region with 1 DMR containing genes: ADD1, SH3BP2.

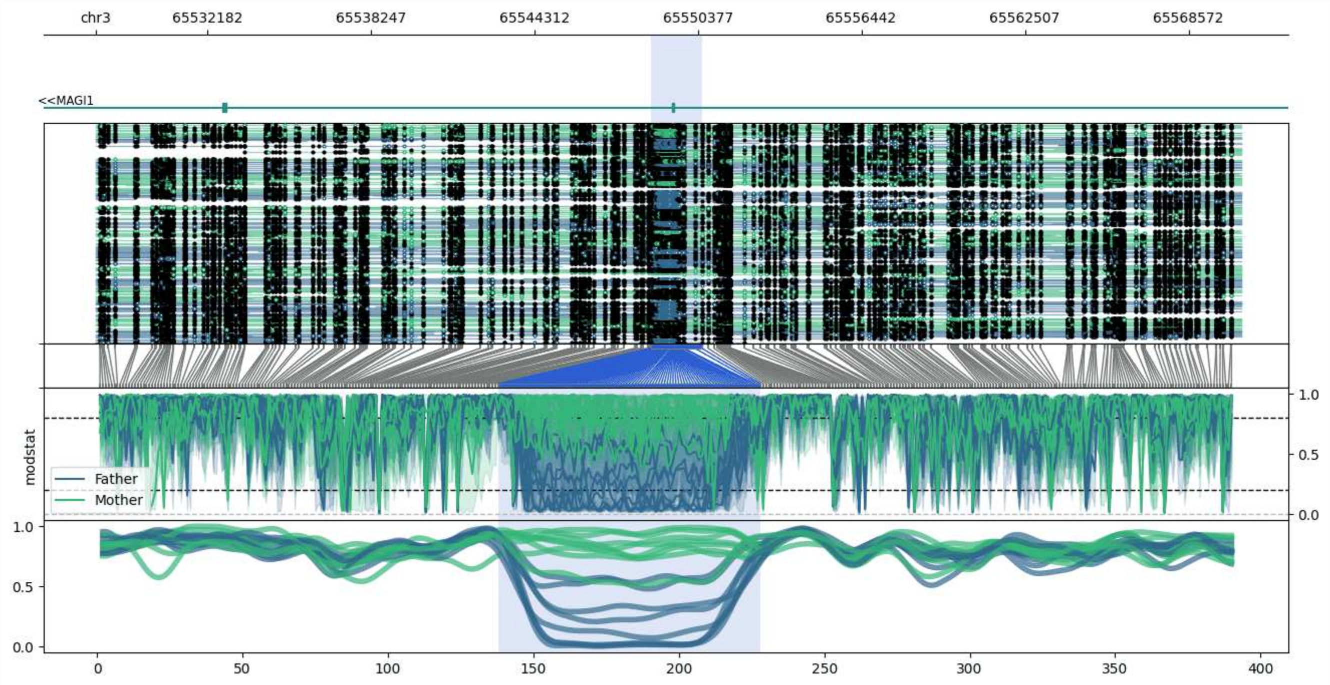


**chr3:65528064-65570525:** Region with 1 DMR containing genes: MAGI1.

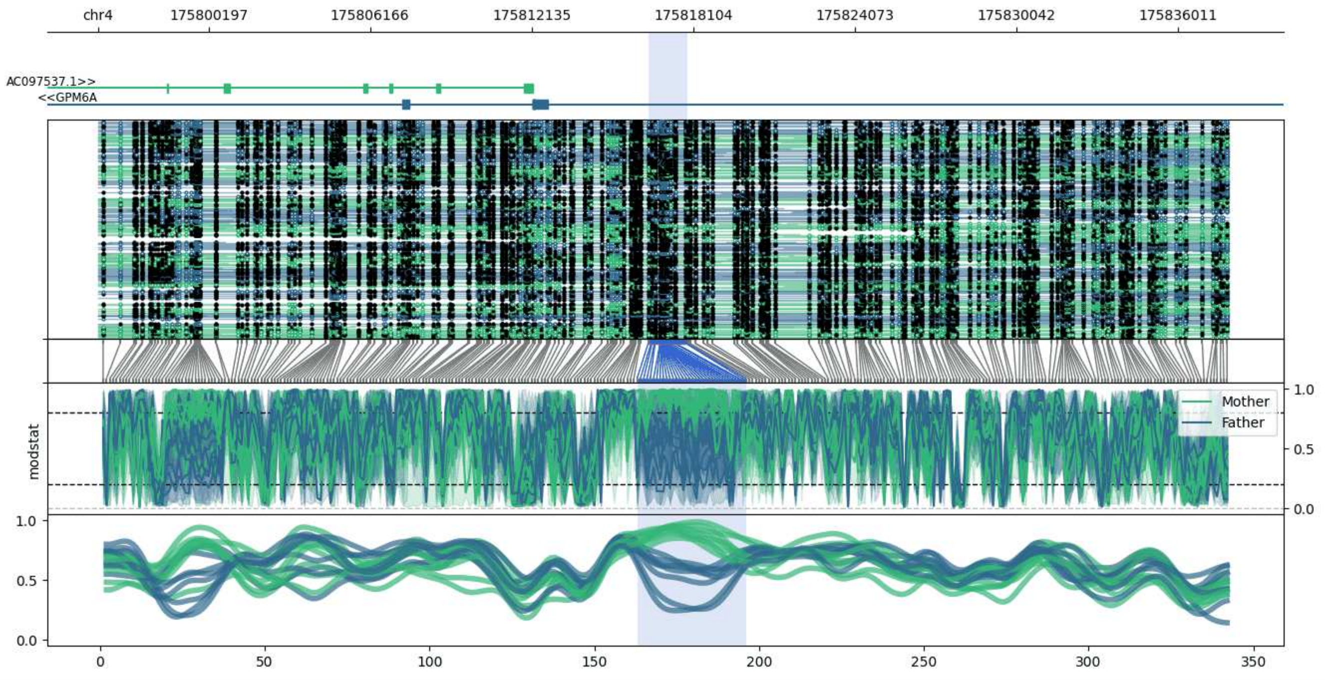


**chr4:175796115-175837900:** Region with 1 DMR containing genes: AC097537.1, GPM6A.

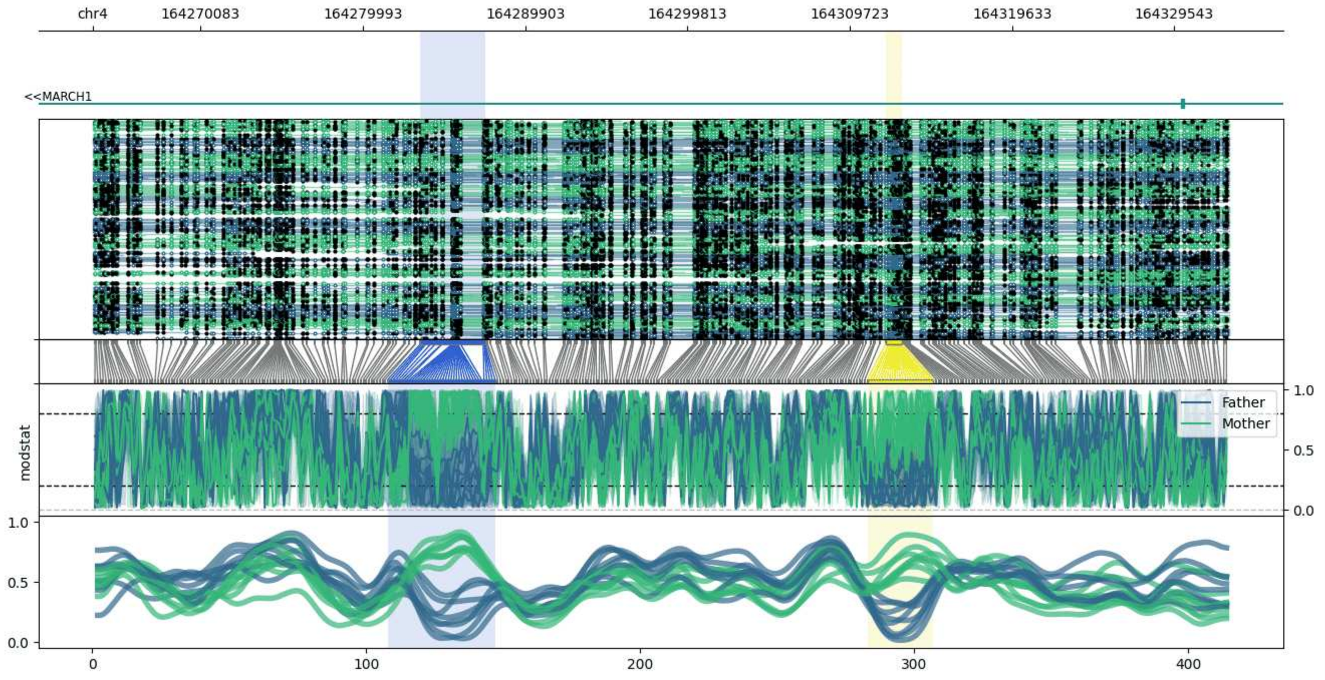


**chr4:164263505-164332877:** Region with 2 DMRs containing genes: MARCH1.

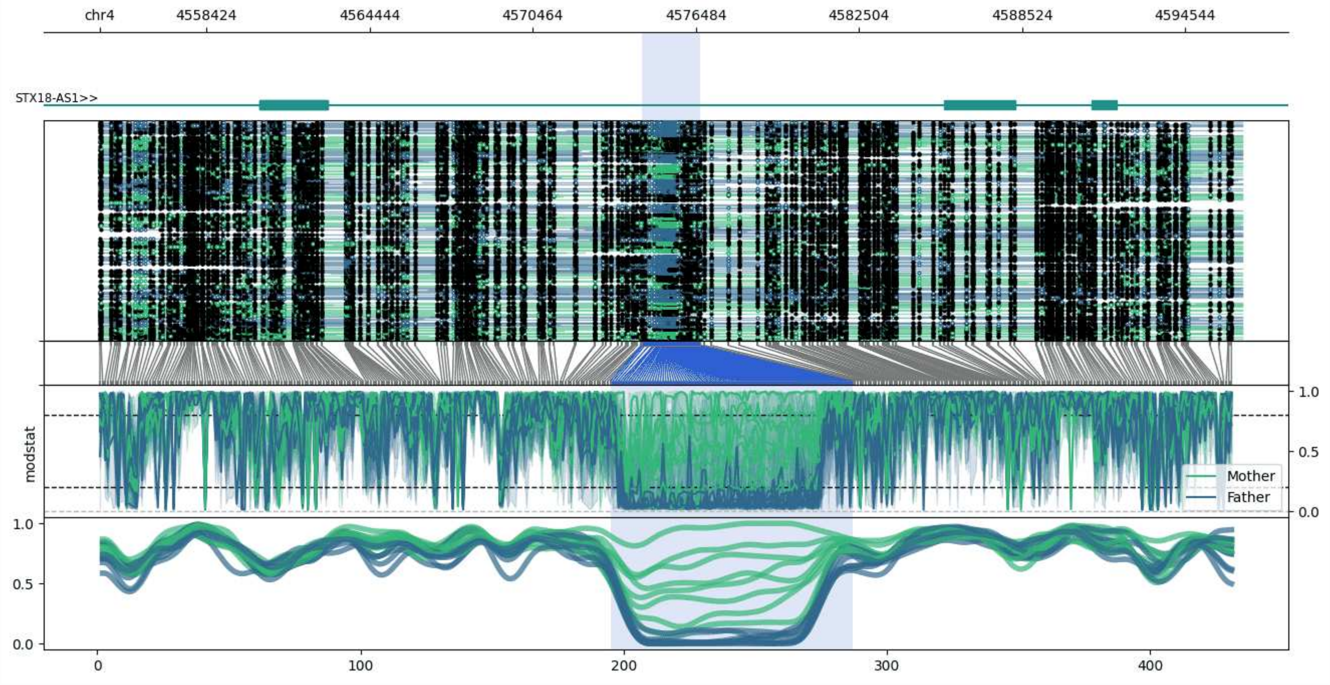


**chr4:4554492-4596634:** Region with 1 DMR containing genes: STX18-AS1.

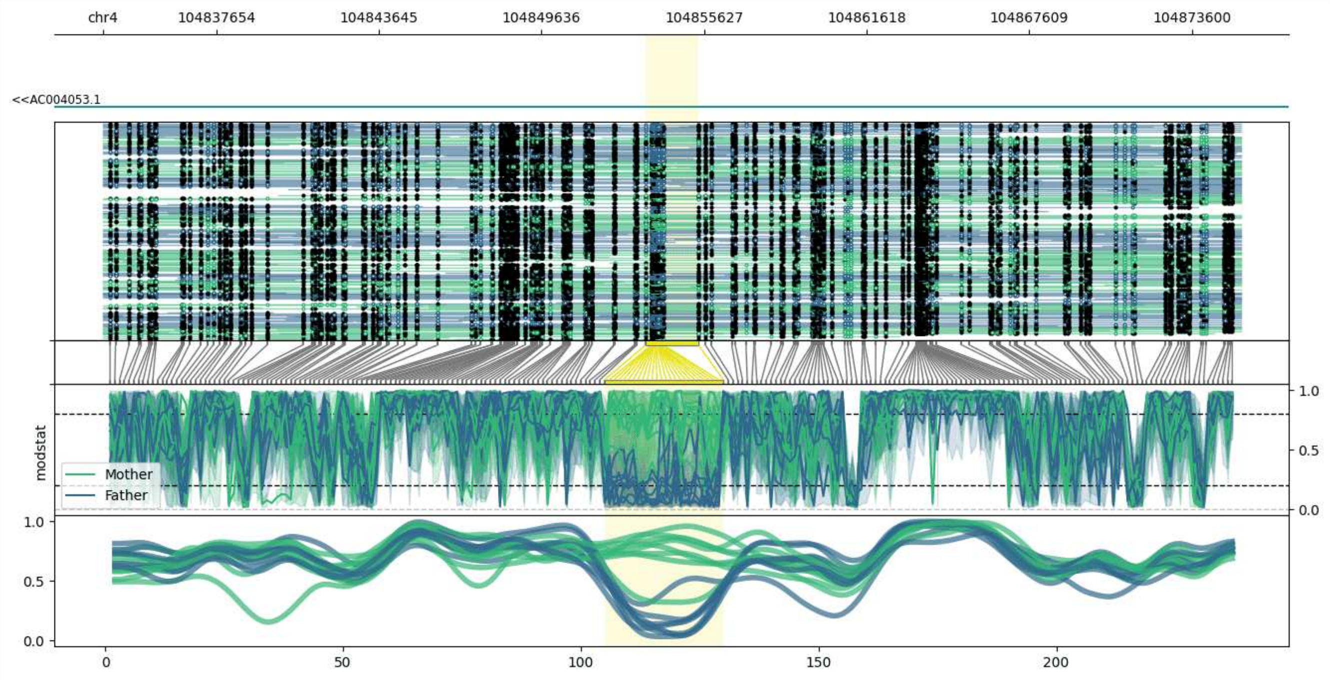


**chr4:104833479-104875417:** Region with 1 DMR containing genes: AC004053.1.

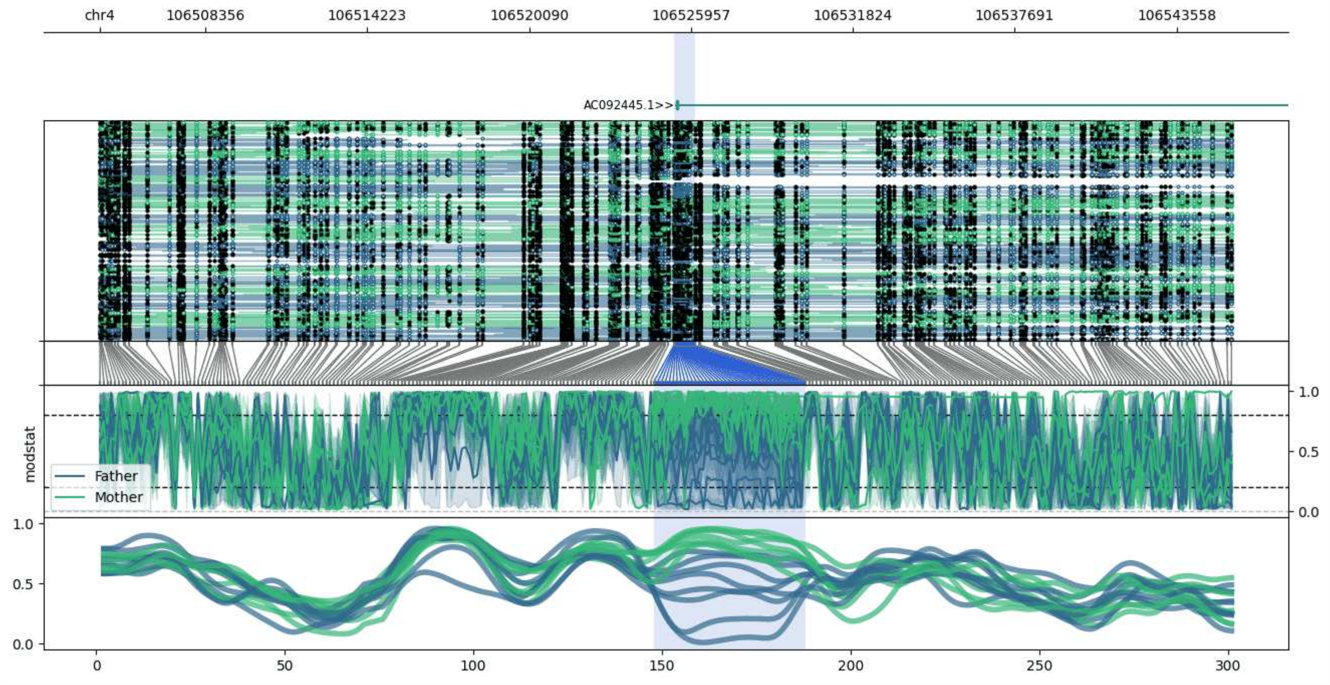


**chr4:106504538-106545613:** Region with 1 DMR containing genes: AC092445.1.

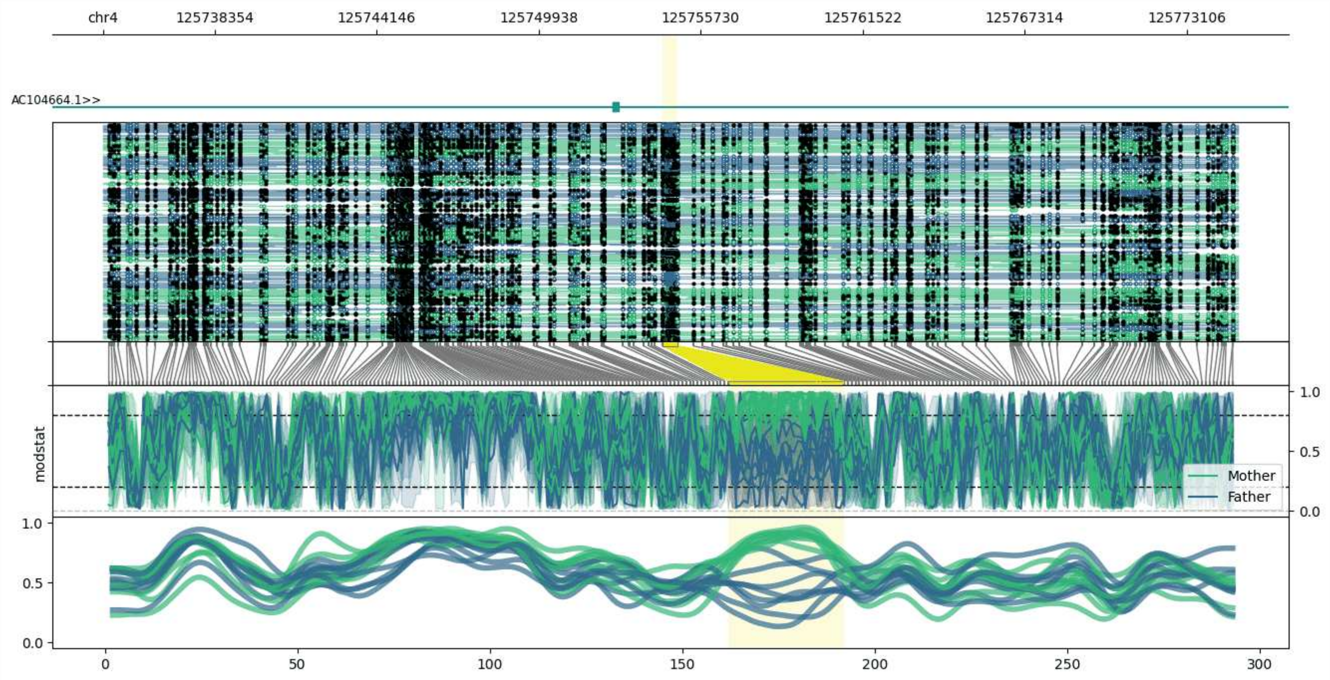


**chr4:125734366-125774915:** Region with 1 DMR containing genes: AC104664.1.

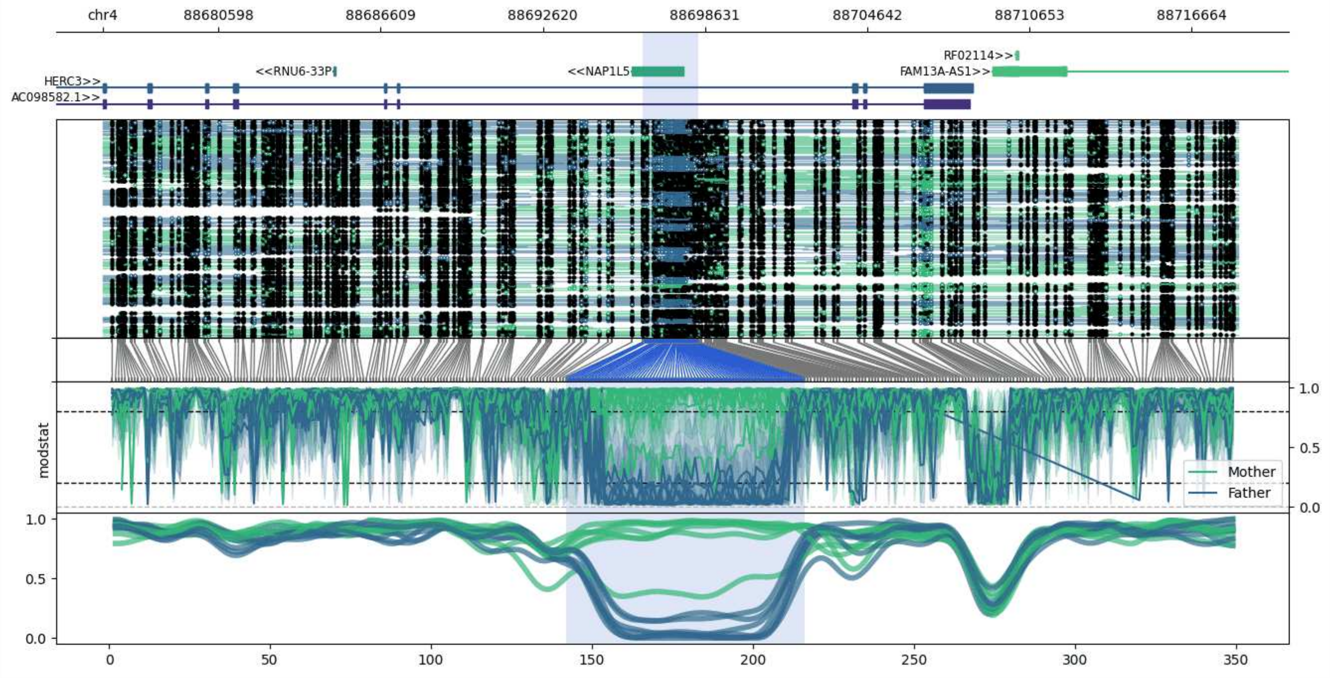


**chr4:88676316-88718397:** Region with 1 DMR containing genes: AC098582.1, FAM13A, FAM13A-AS1, HERC3, NAP1L5, RF02114, RF02115, RNU6-33P.

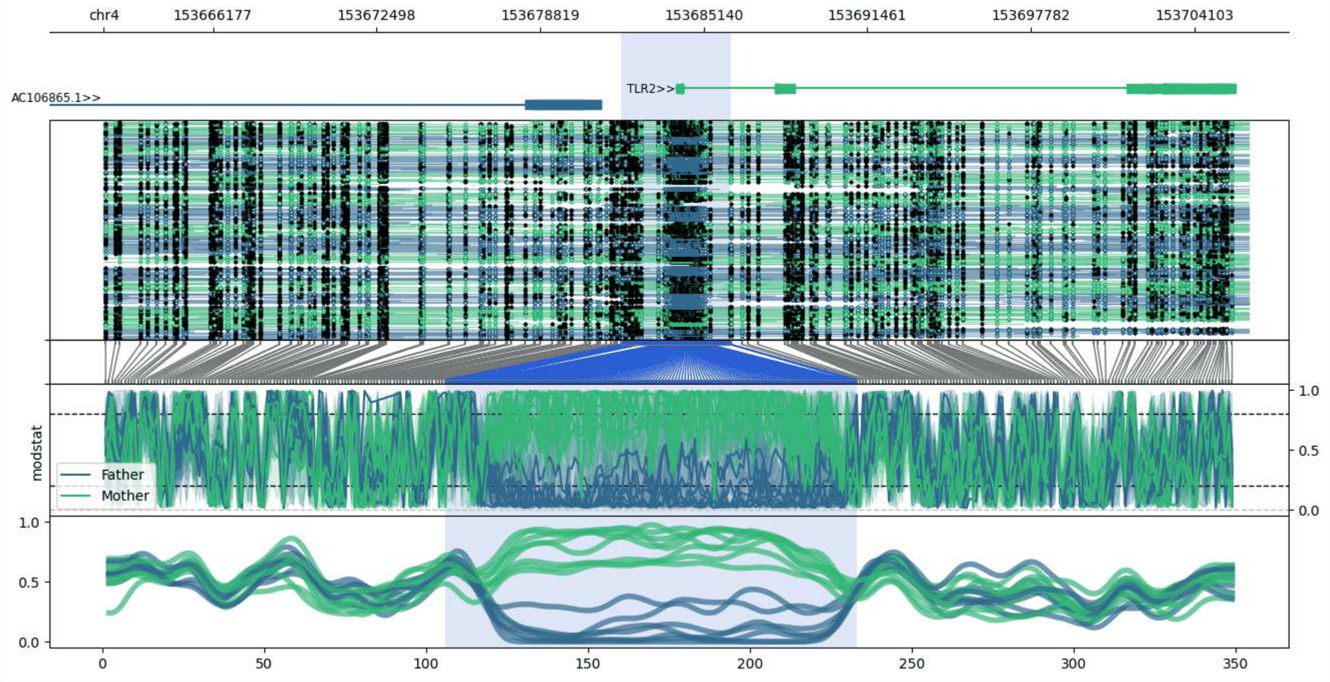


**chr4:153661963-153706214:** Region with 1 DMR containing genes: AC106865.1, AC106865.2, RNF175, TLR2, WDR45P1.

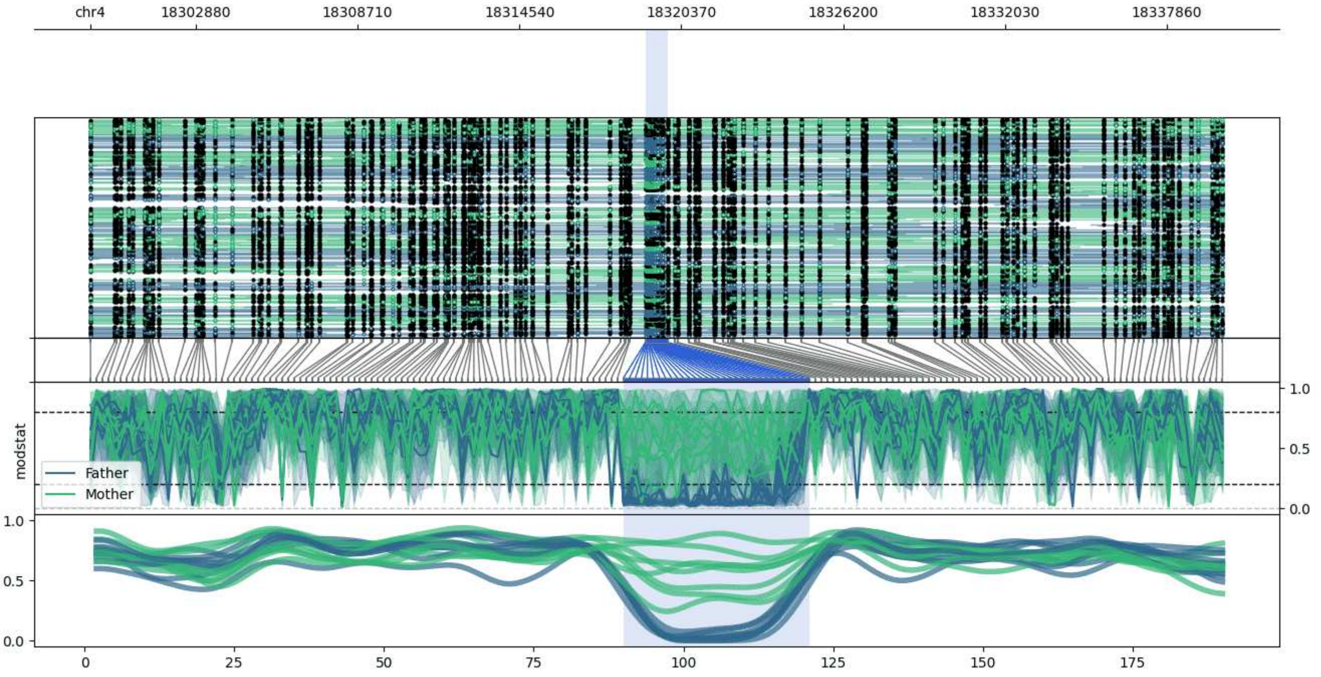


**chr4:18299076-18339888:** Region with 1 DMR containing no genes.

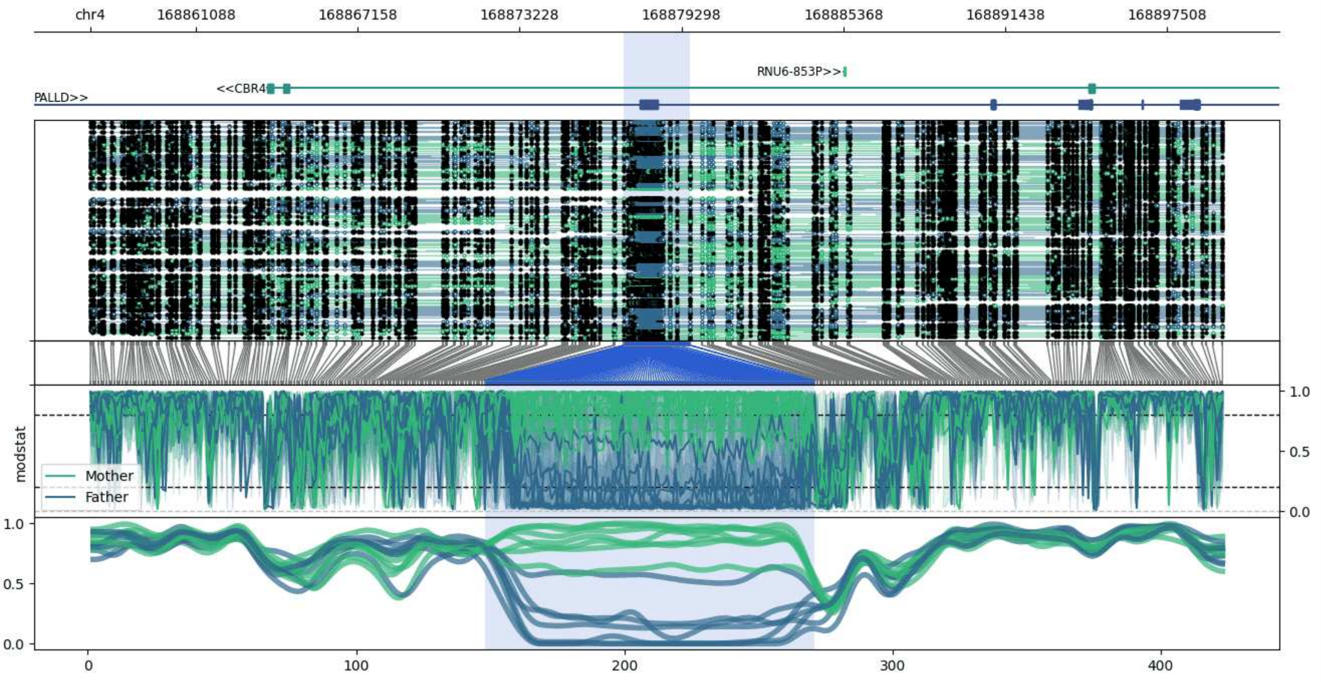


**chr4:168857126-168899616:** Region with 1 DMR containing genes: CBR4, PALLD, RNU6-853P.

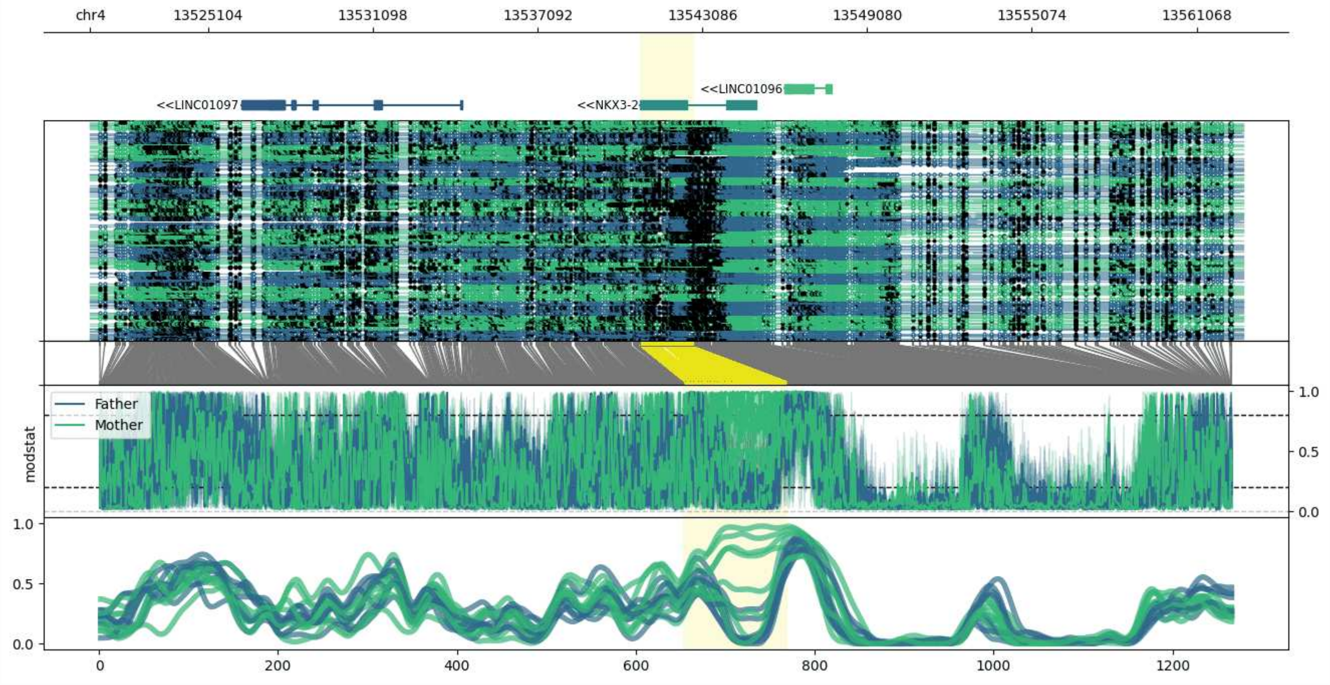


**chr4:13520813-13562774:** Region with 1 DMR containing genes: AC006445.3, BOD1L1, LINC01096, LINC01097, NKX3-2.

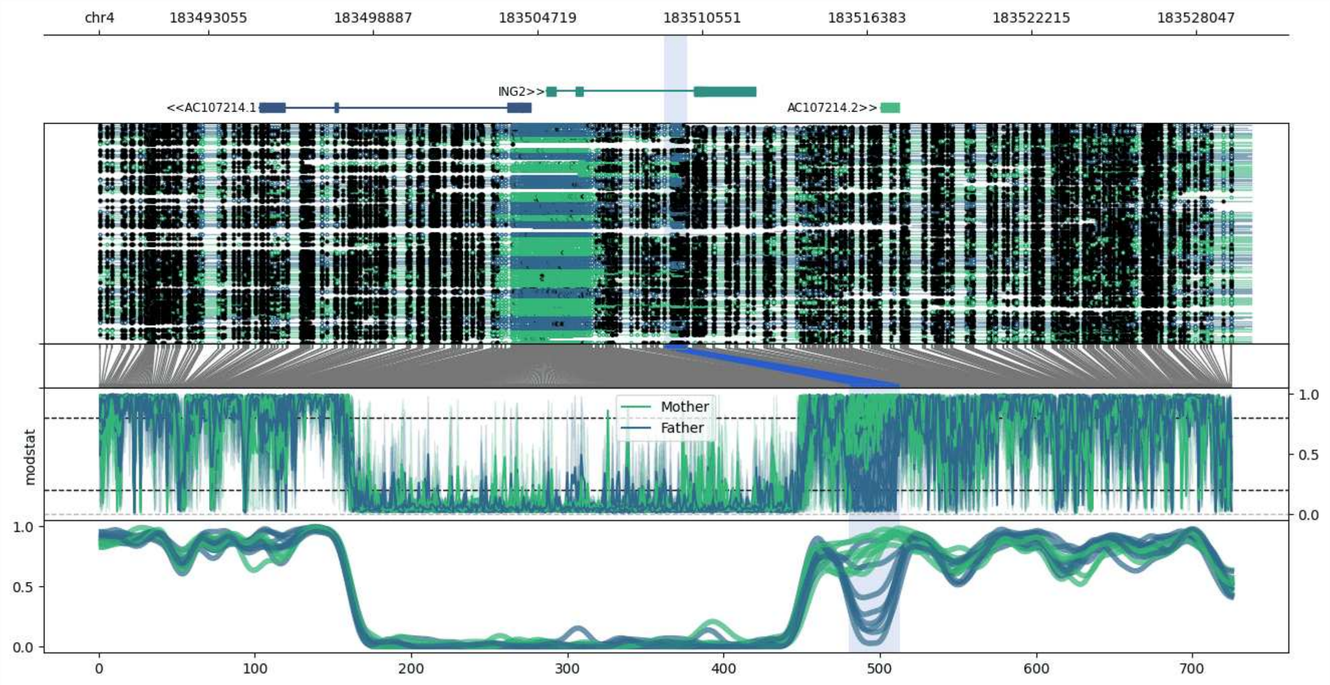


**chr4:183489200-183530026:** Region with 1 DMR containing genes: AC107214.1, AC107214.2, ING2, VTI1BP2.

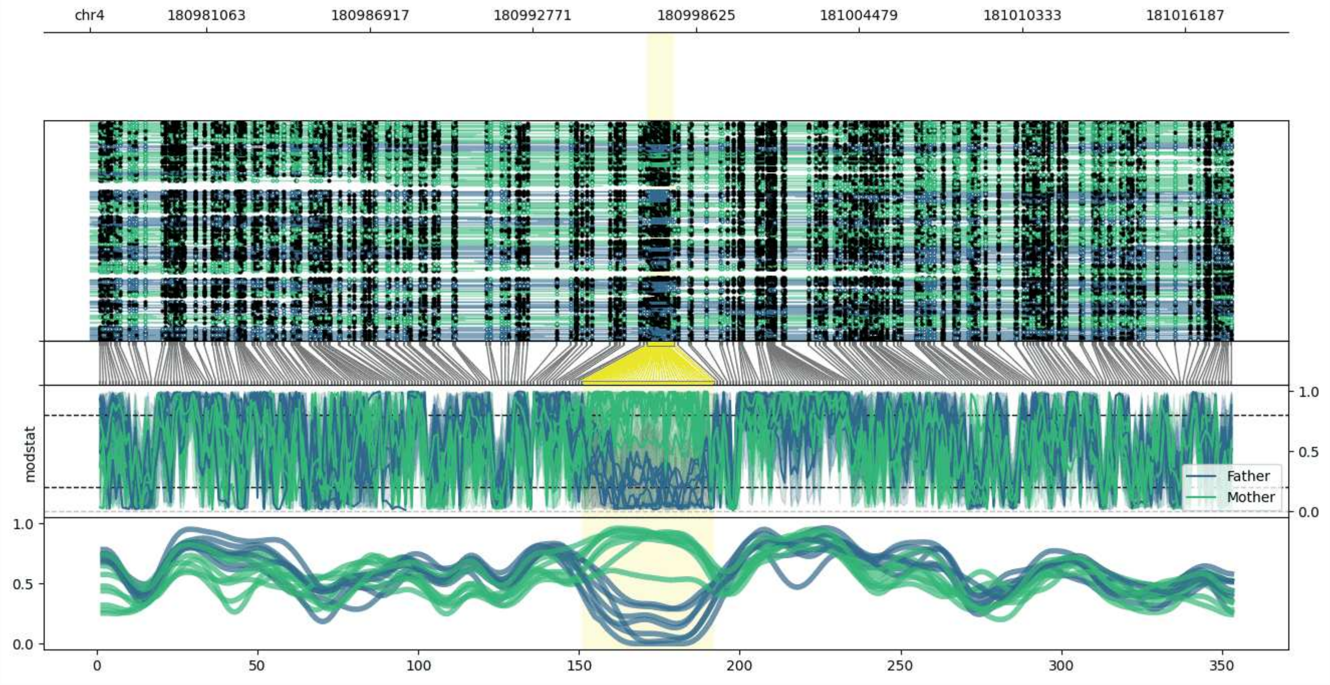


**chr4:180976874-181017856:** Region with 1 DMR containing no genes.

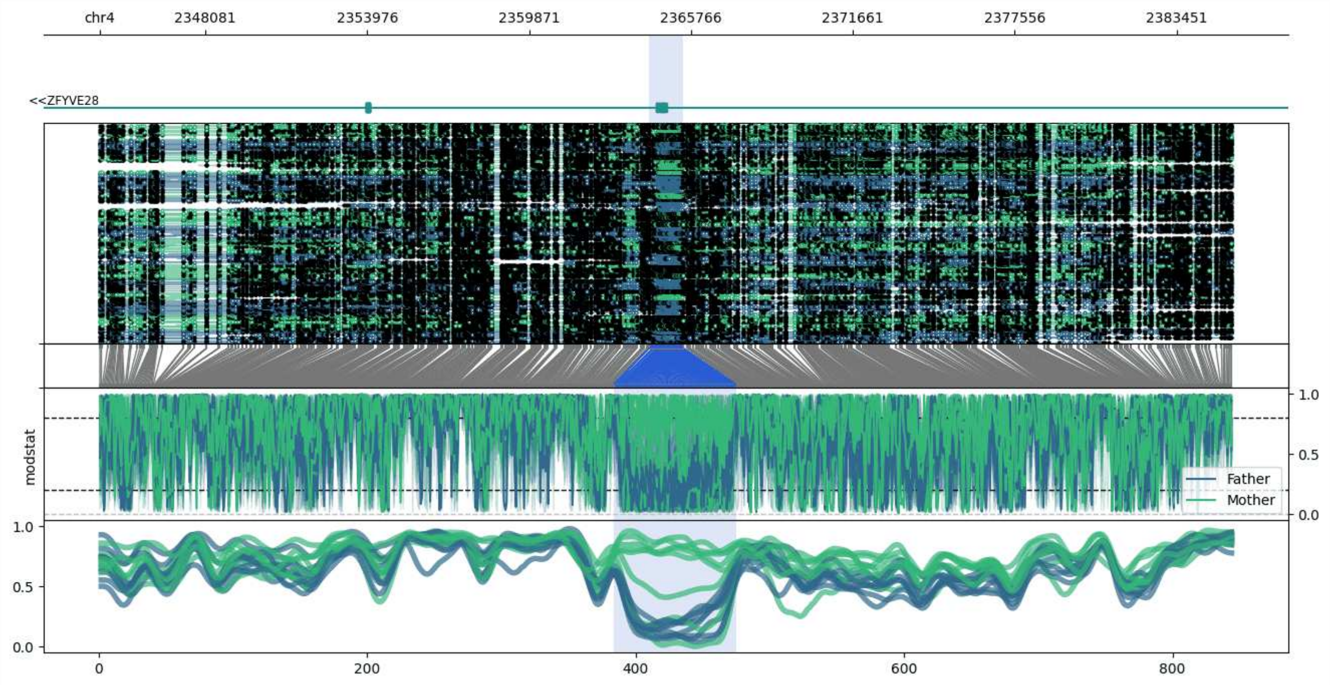


**chr4:2344230-2385498:** Region with 1 DMR containing genes: ZFYVE28.

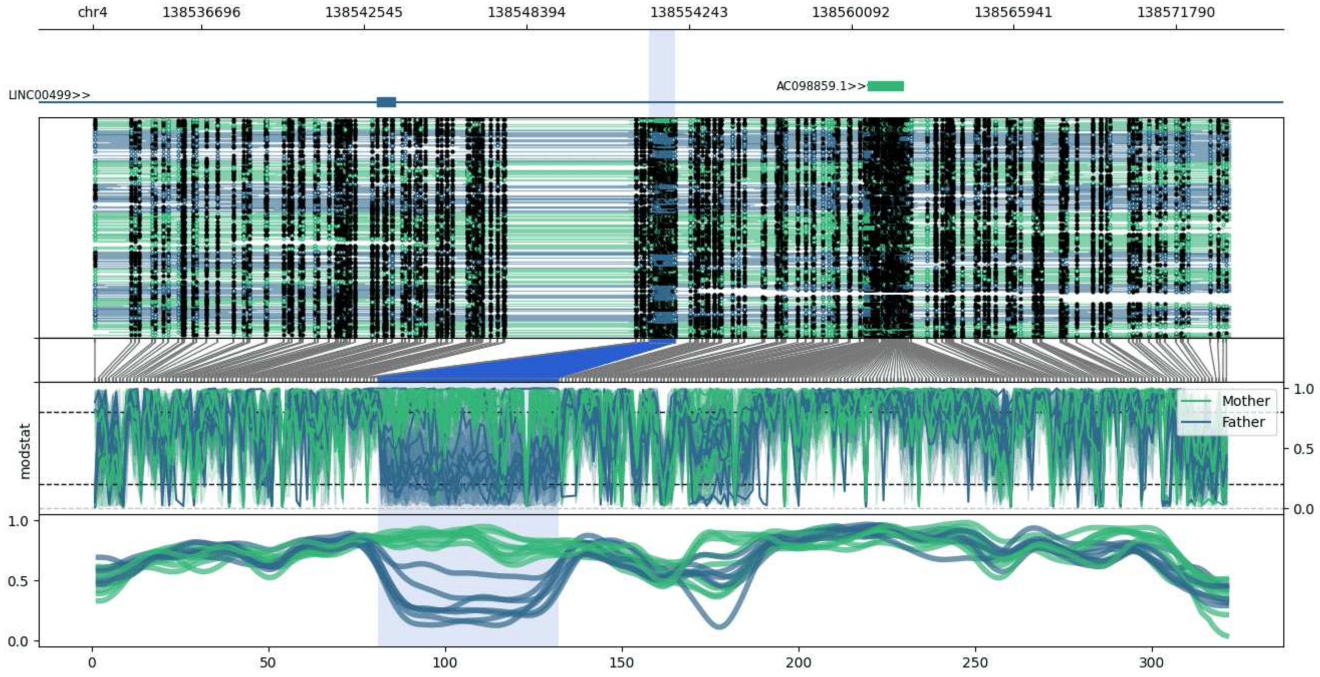


**chr4:138532797-138573742:** Region with 1 DMR containing genes: AC098859.1, LINC00499.

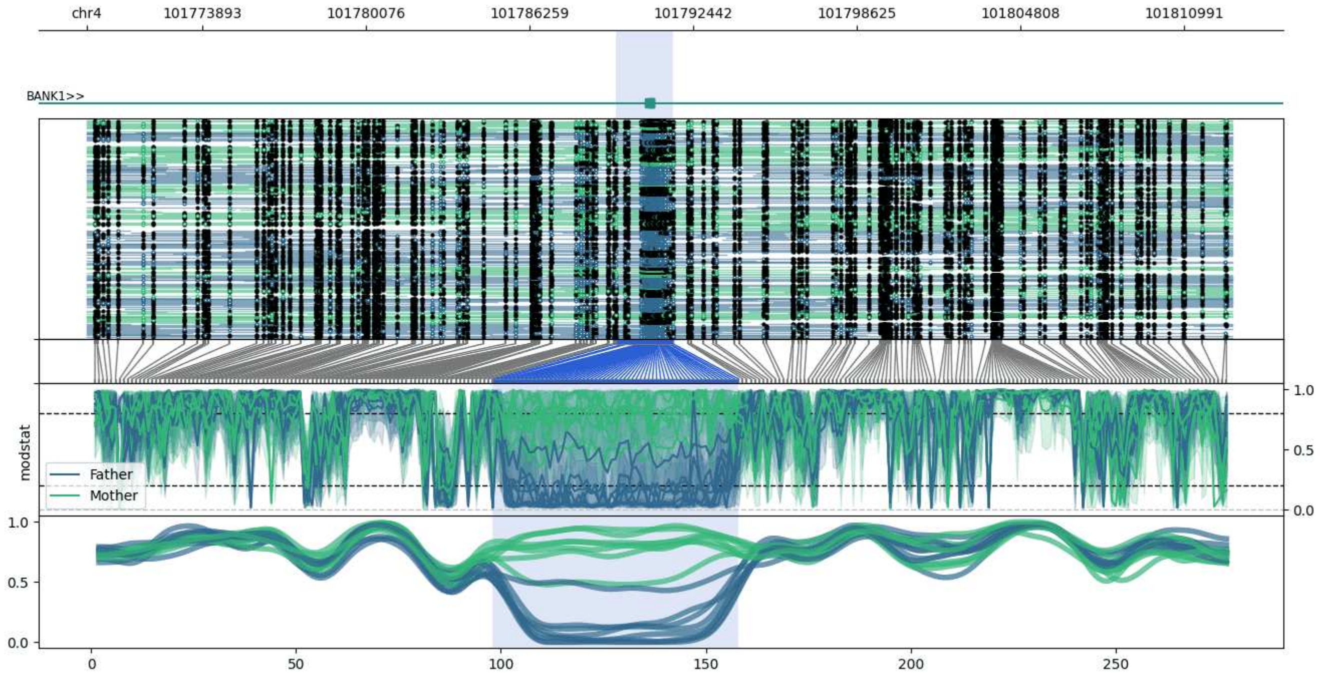


**chr4:101769529-101812810:** Region with 1 DMR containing genes: BANK1.

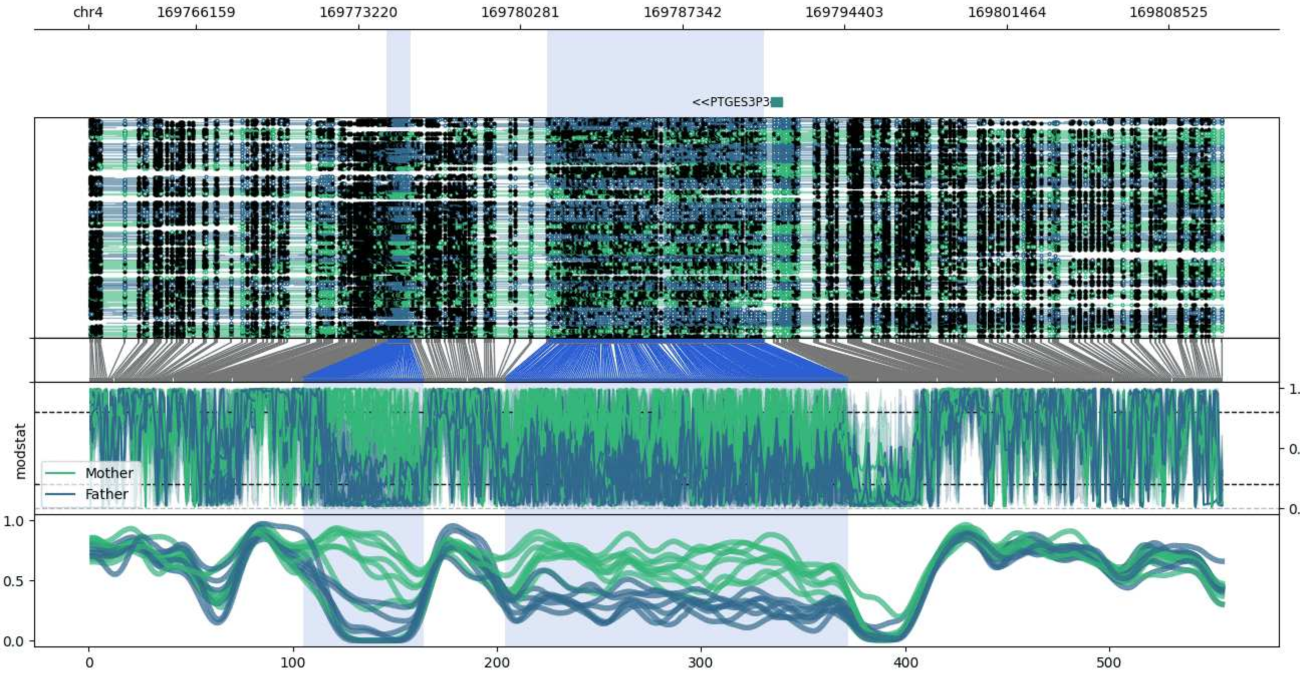


**chr4:169761455-169810883:** Region with 2 DMRs containing genes: HPF1, PTGES3P3.

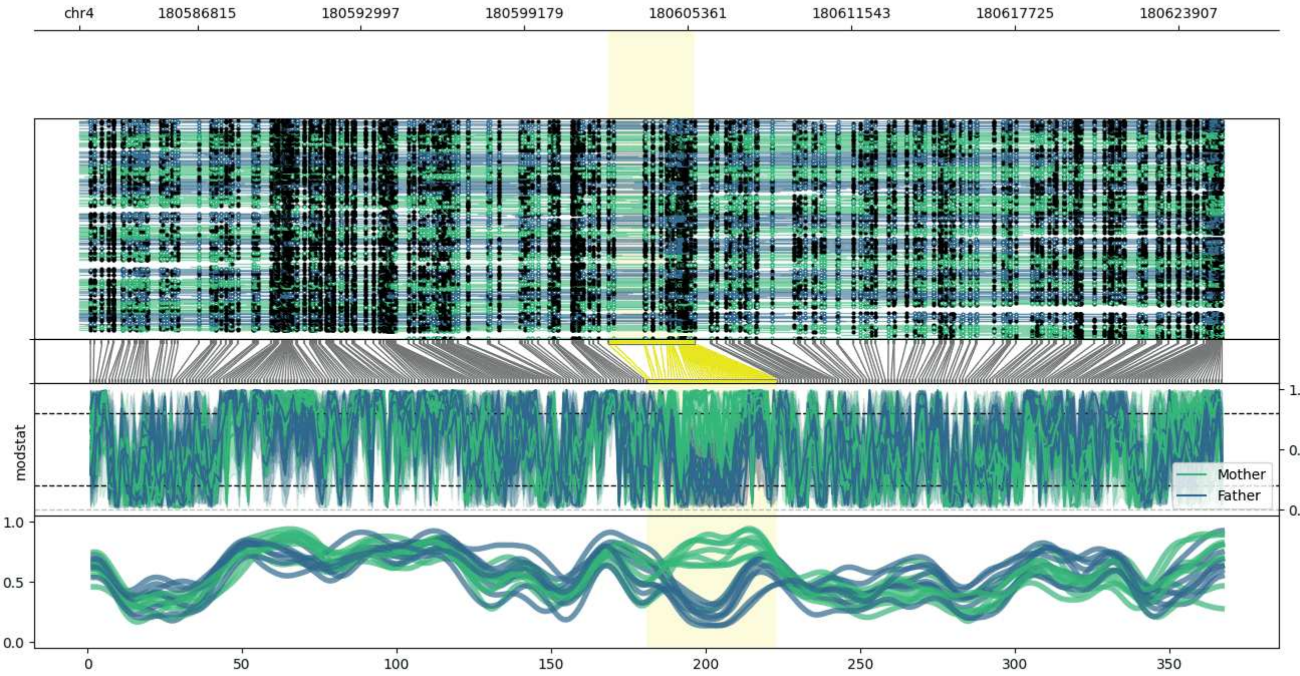


**chr4:180582342-180625619:** Region with 1 DMR containing genes: NDUFB5P1.

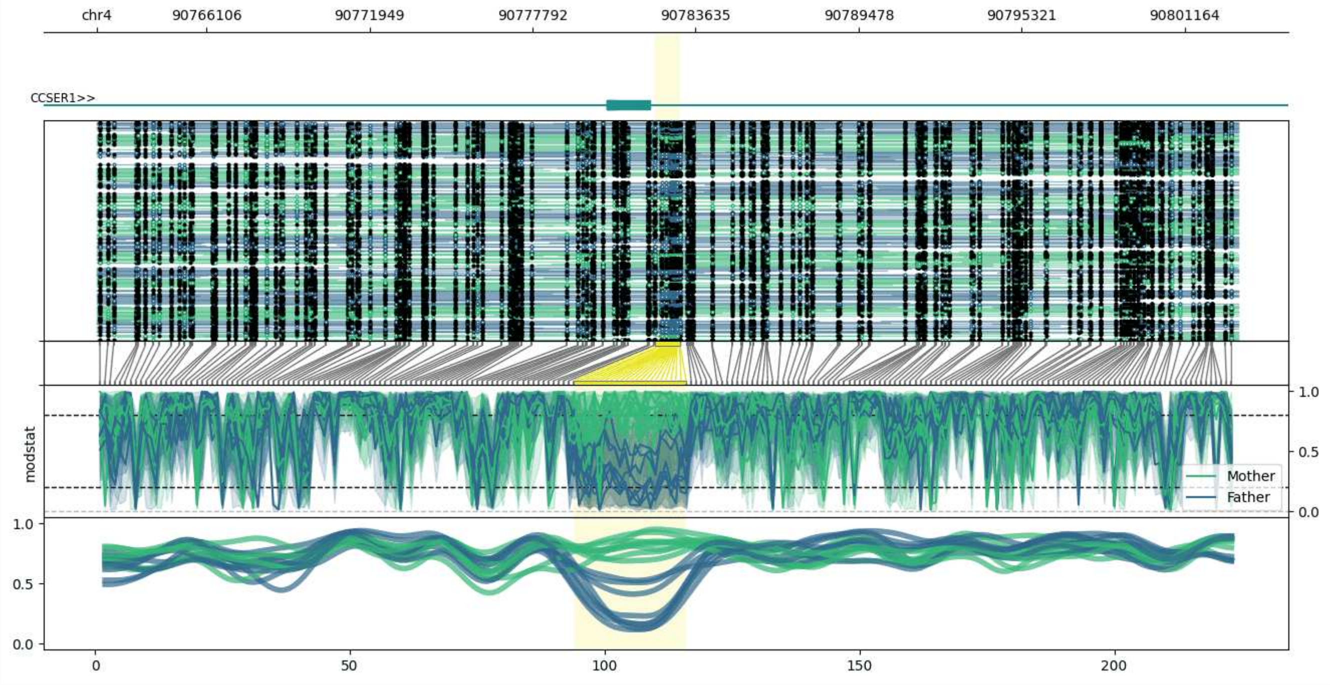


**chr4:90762179-90803081:** Region with 1 DMR containing genes: CCSER1.

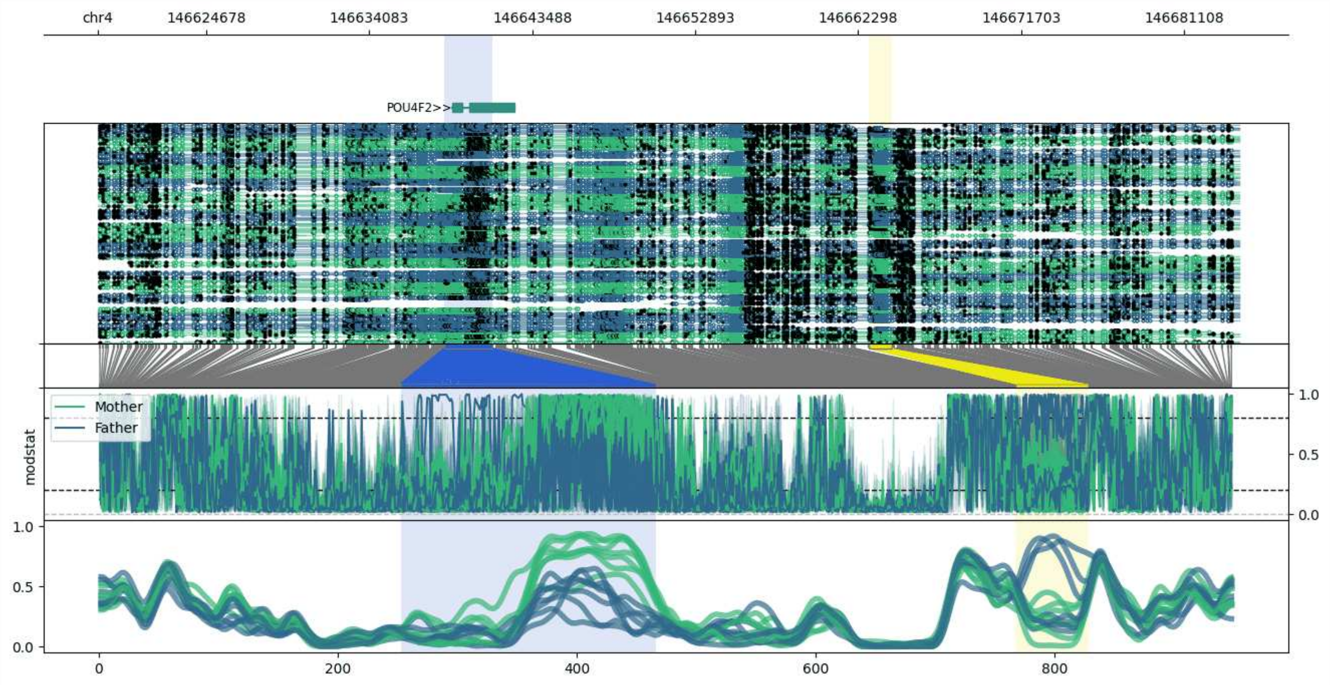


**chr4:146618426-146684262:** Region with 2 DMRs containing genes: POU4F2.

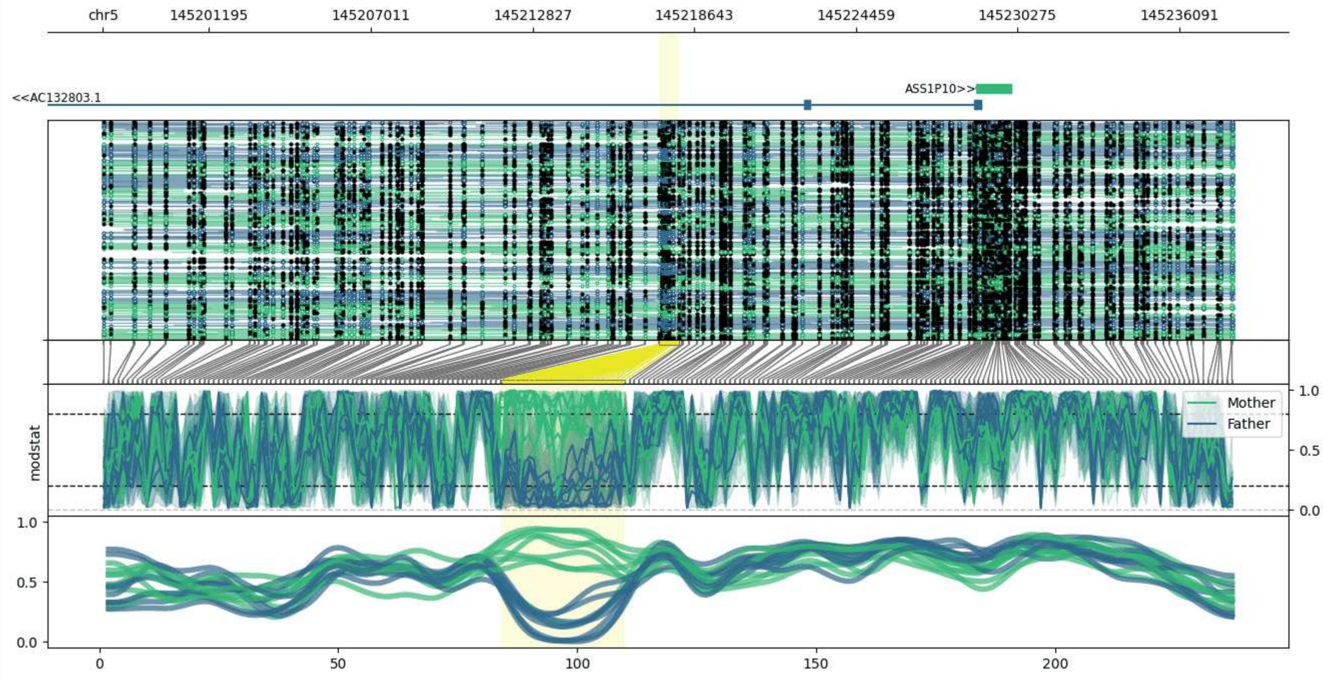


**chr5:145197369-145238086:** Region with 1 DMR containing genes: AC132803.1, ASS1P10.

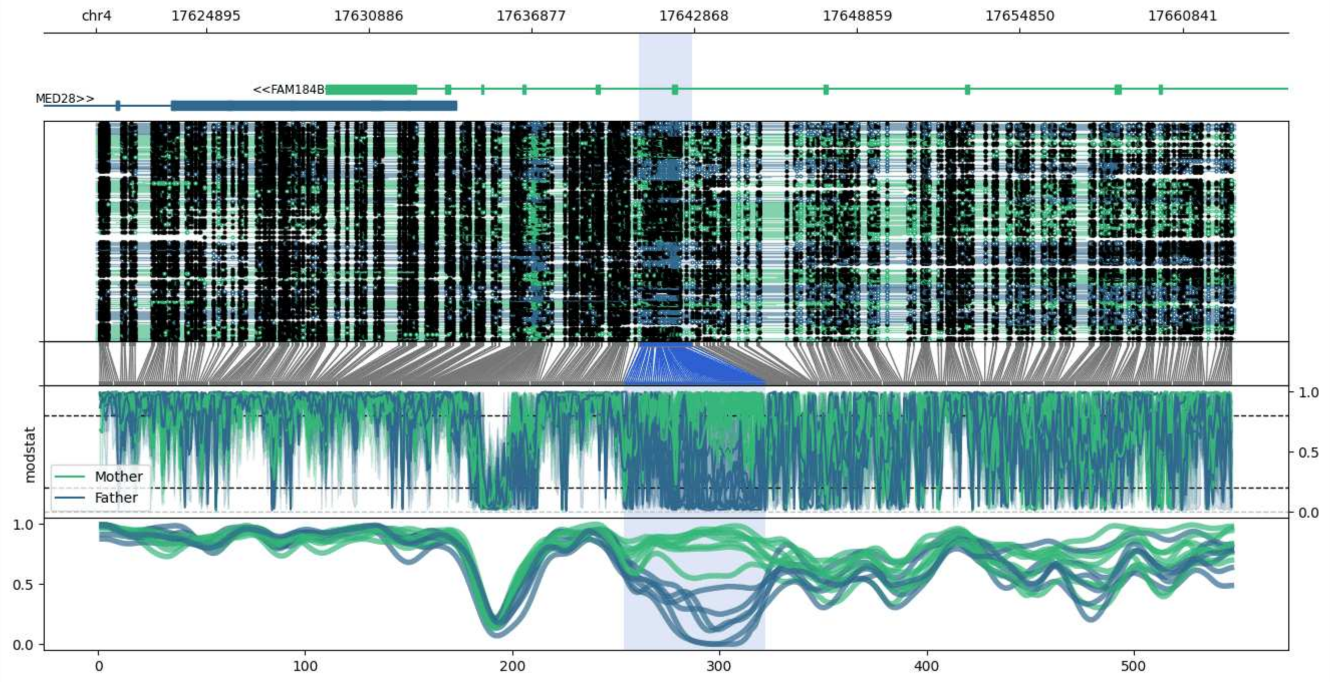


**chr4:17620846-17662788:** Region with 1 DMR containing genes: AC006160.1, FAM184B, MED28.

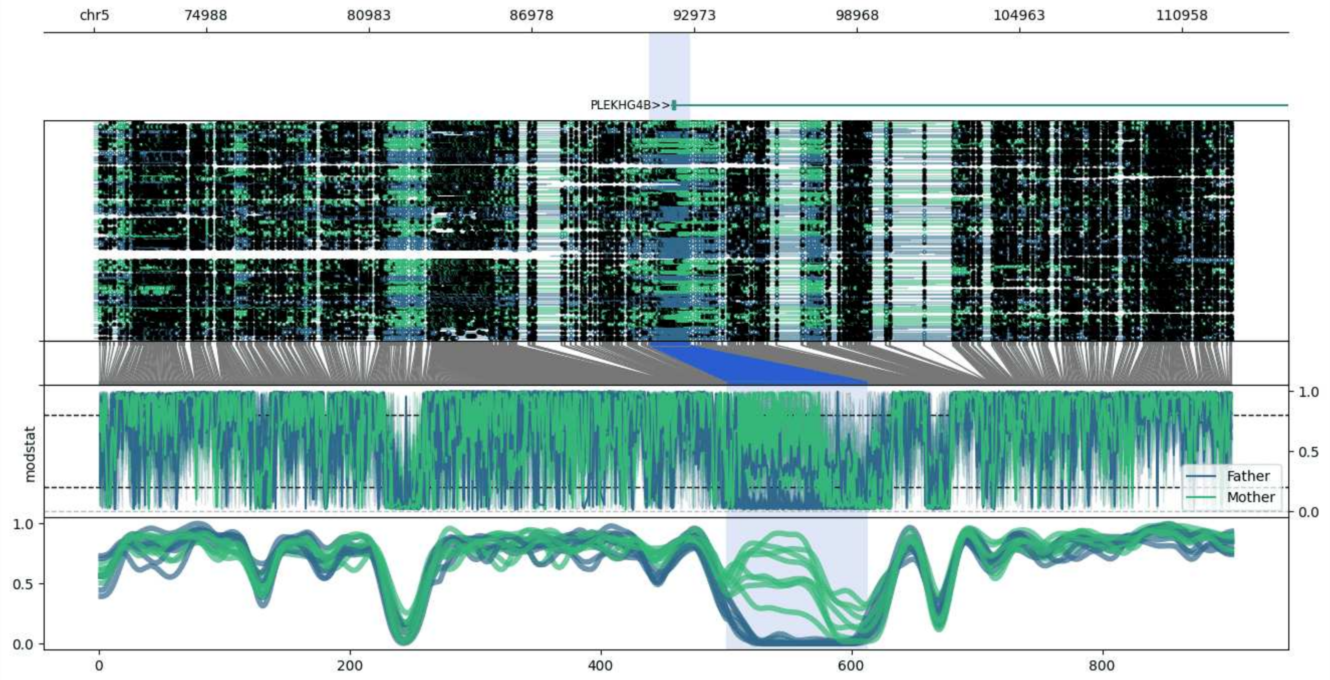


**chr5:70861-112826:** Region with 1 DMR containing genes: PLEKHG4B.

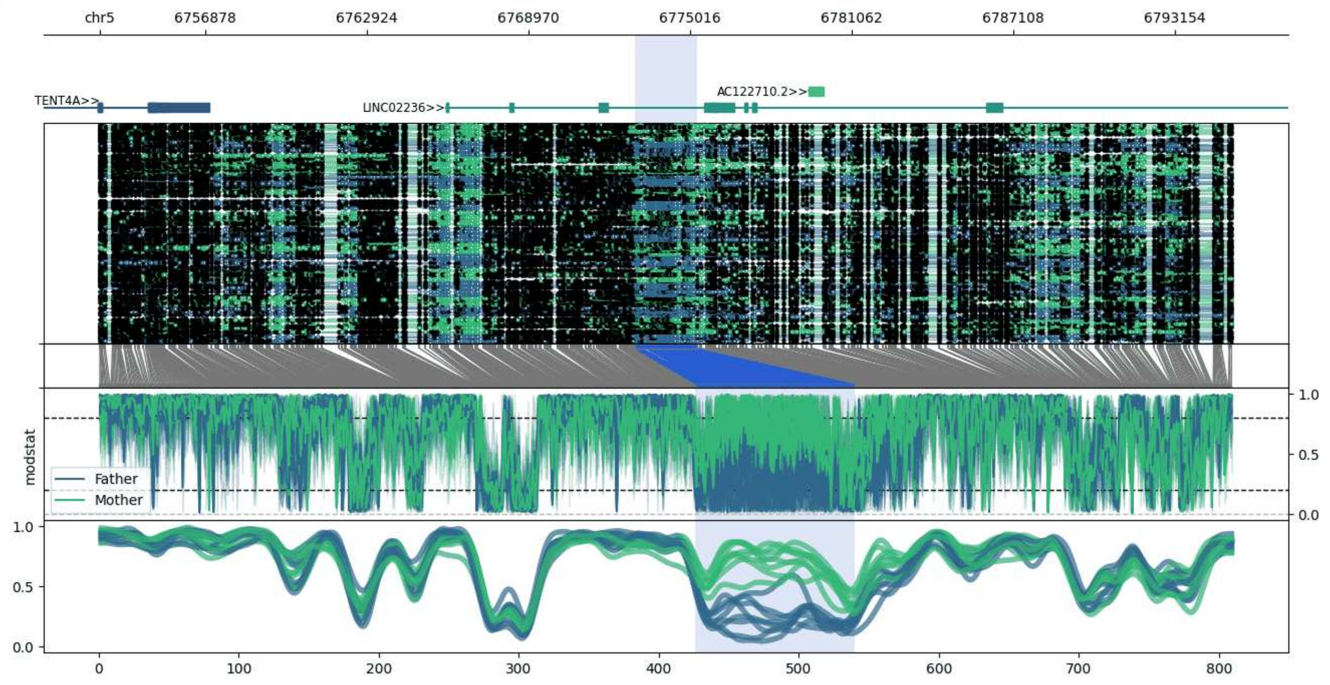


**chr5:6752947-6795272:** Region with 1 DMR containing genes: AC122710.2, LINC02236, TENT4A.

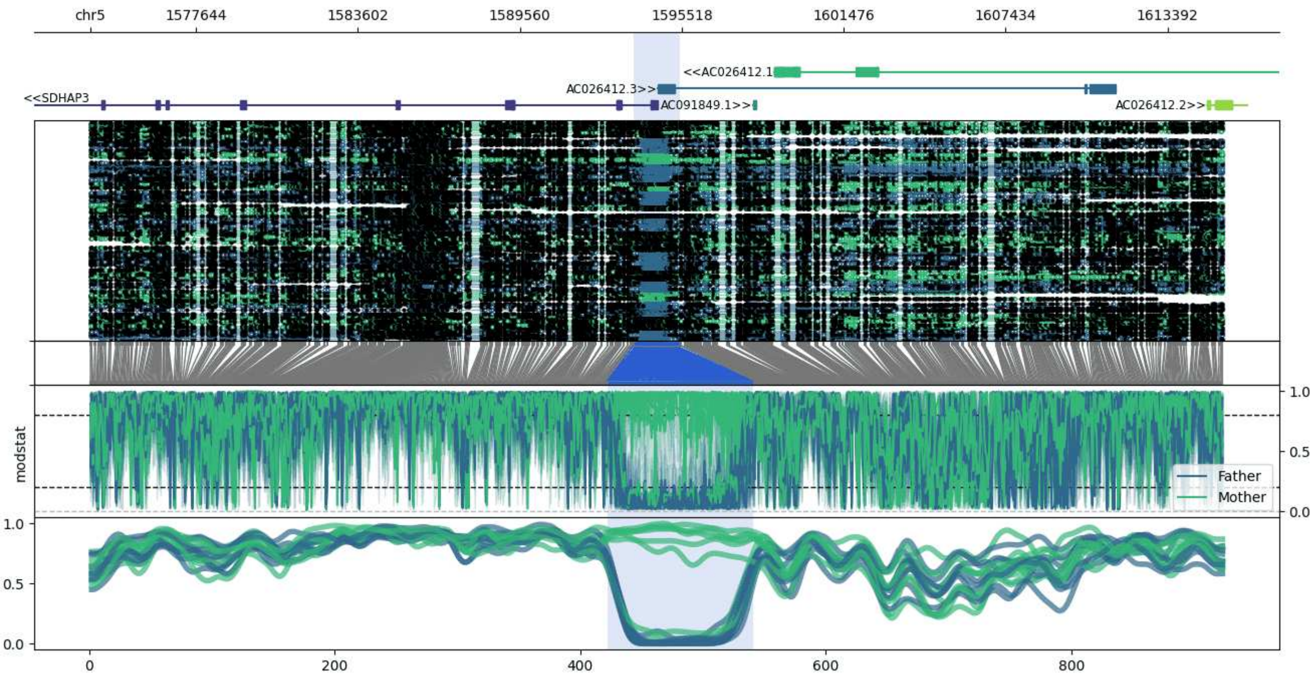


**chr5:1573755-1615463:** Region with 1 DMR containing genes: AC026412.1, AC026412.2, AC026412.3, AC091849.1, SDHAP3.

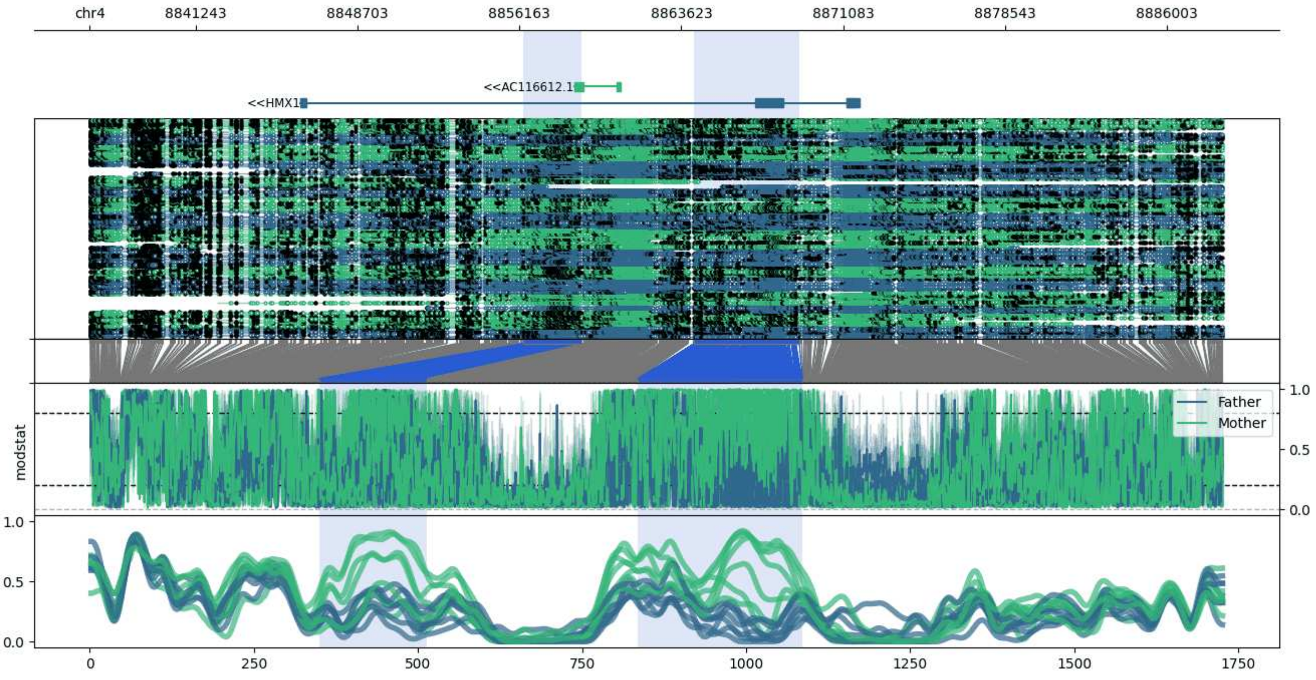


**chr4:8836355-8888578:** Region with 2 DMRs containing genes: AC116612.1, HMX1.

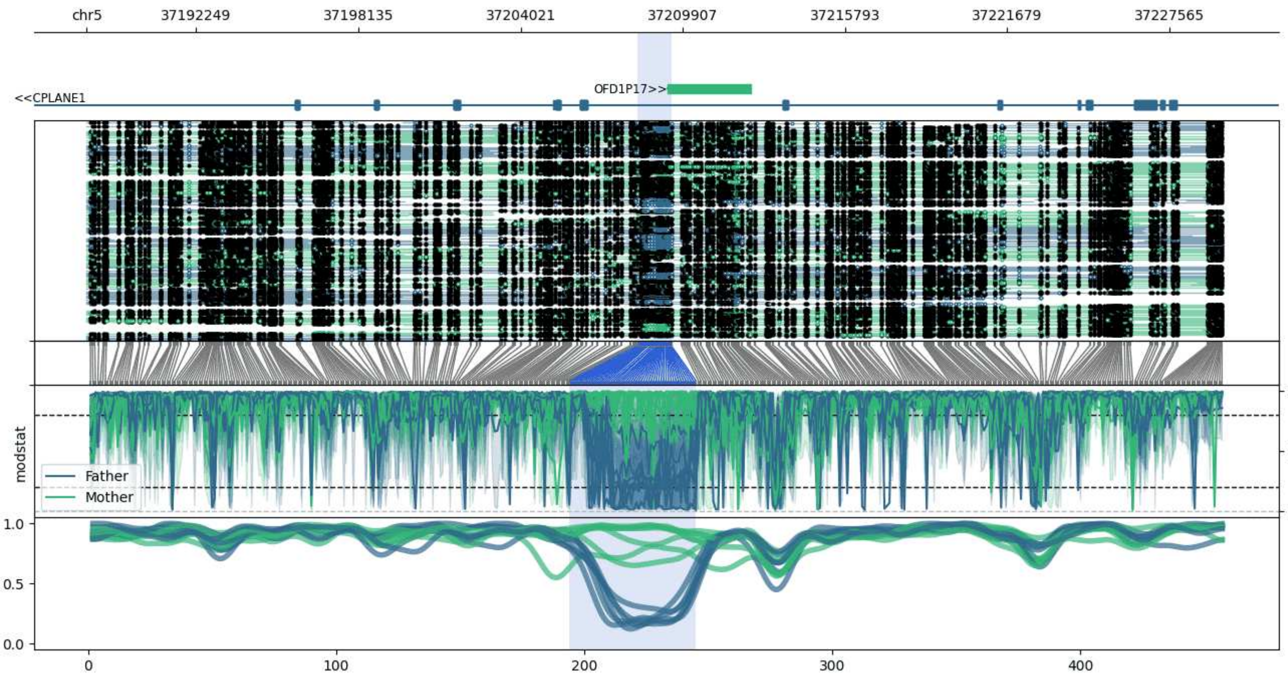


**chr5:37188278-37229483:** Region with 1 DMR containing genes: CPLANE1, OFD1P17.

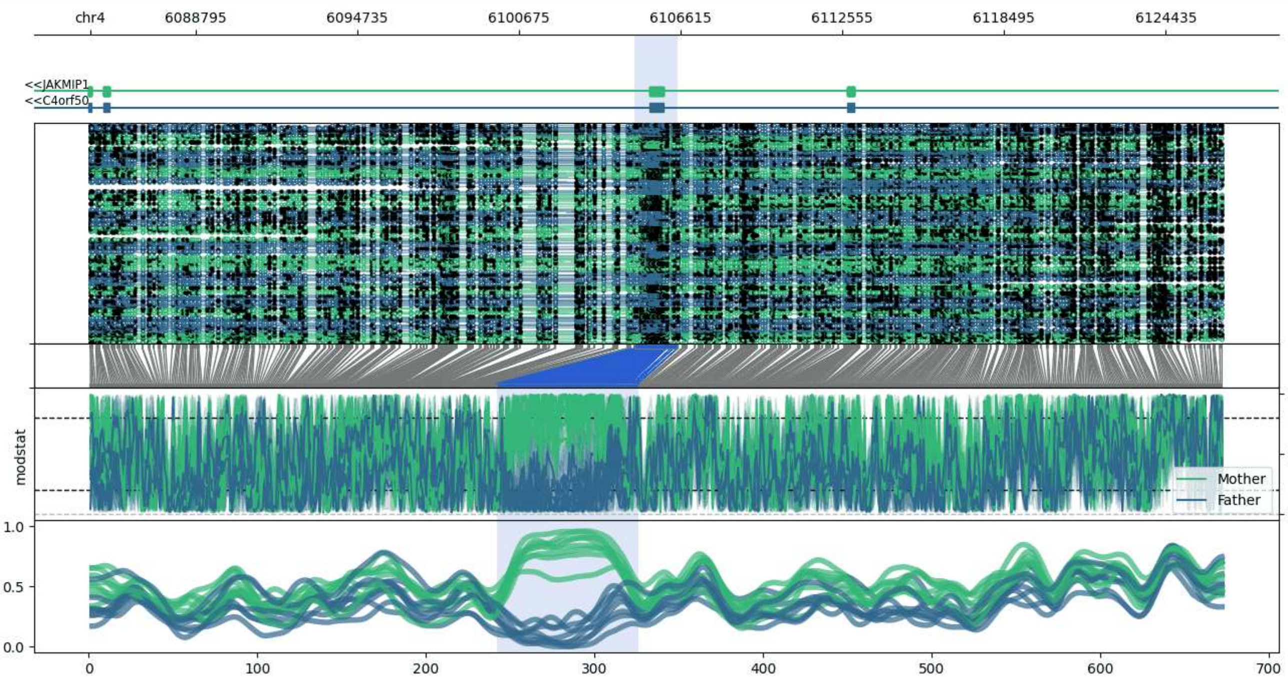


**chr4:6084930-6126512:** Region with 1 DMR containing genes: C4orf50, JAKMIP1.

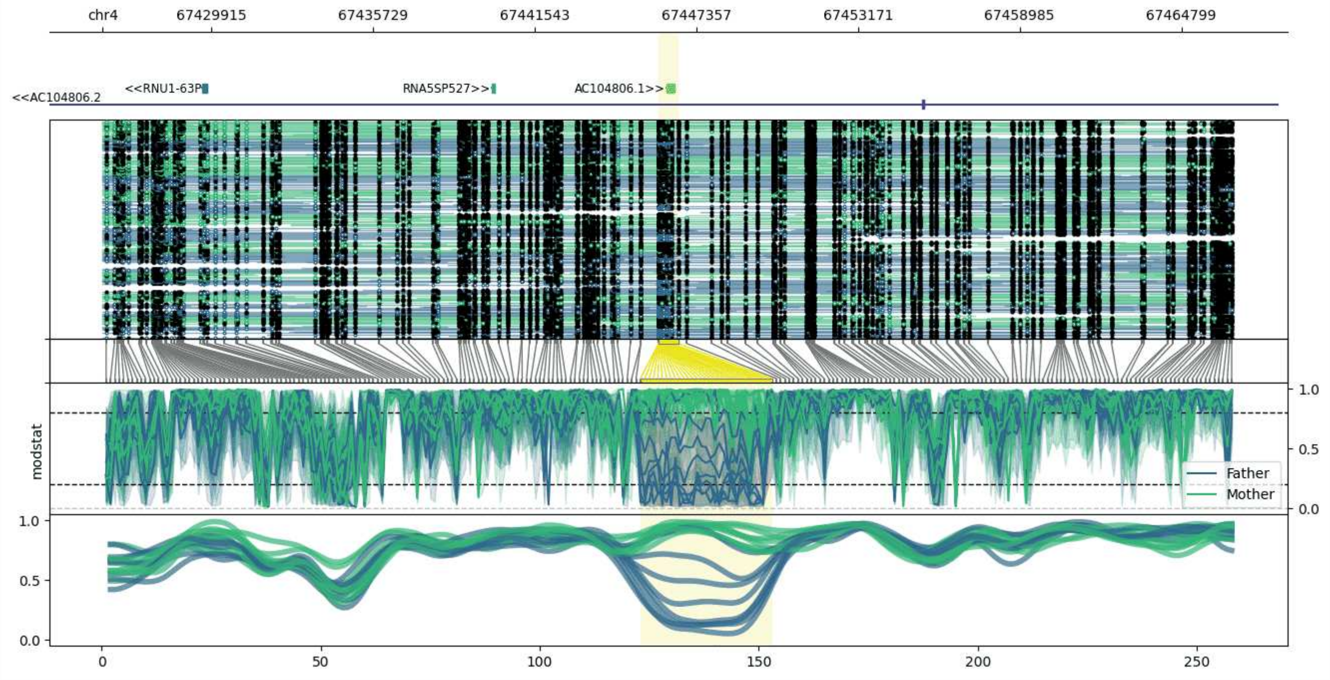


**chr4:67425998-67466696:** Region with 1 DMR containing genes: AC104806.1, AC104806.2, CENPC, RNA5SP527, RNU1-63P.

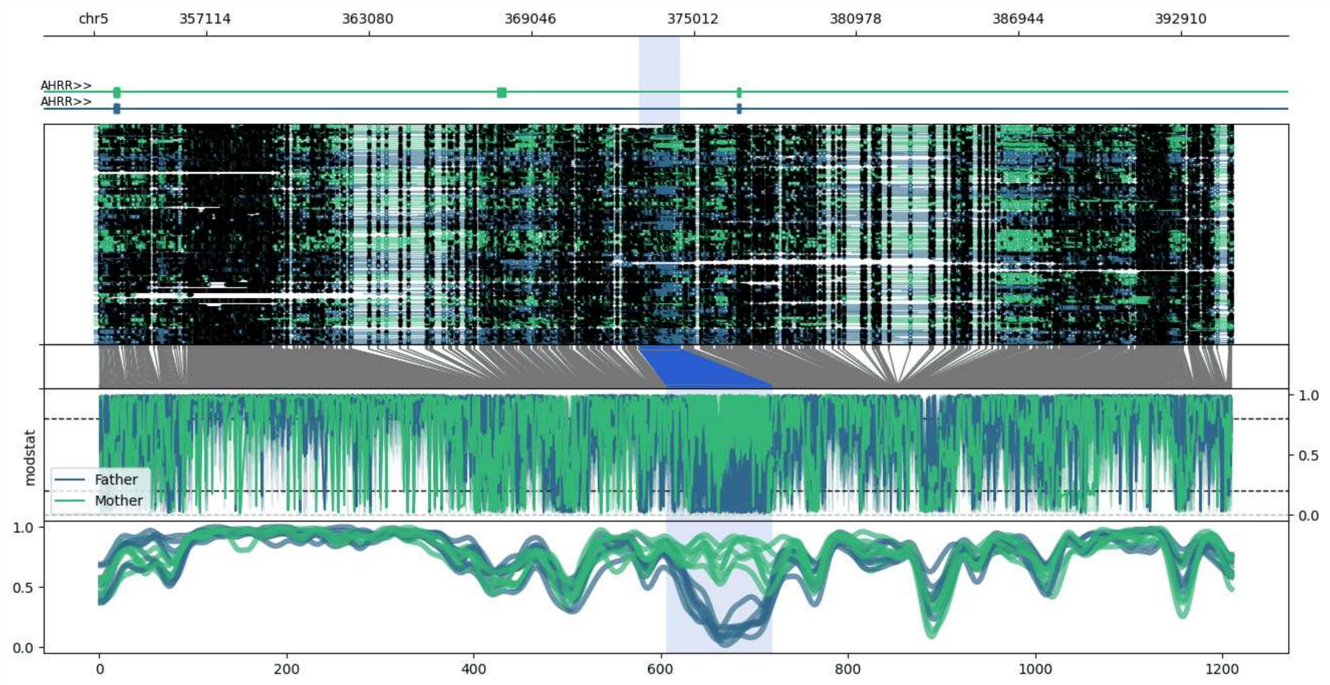


**chr5:352999-394766:** Region with 1 DMR containing genes: AC010442.2, AHRR.

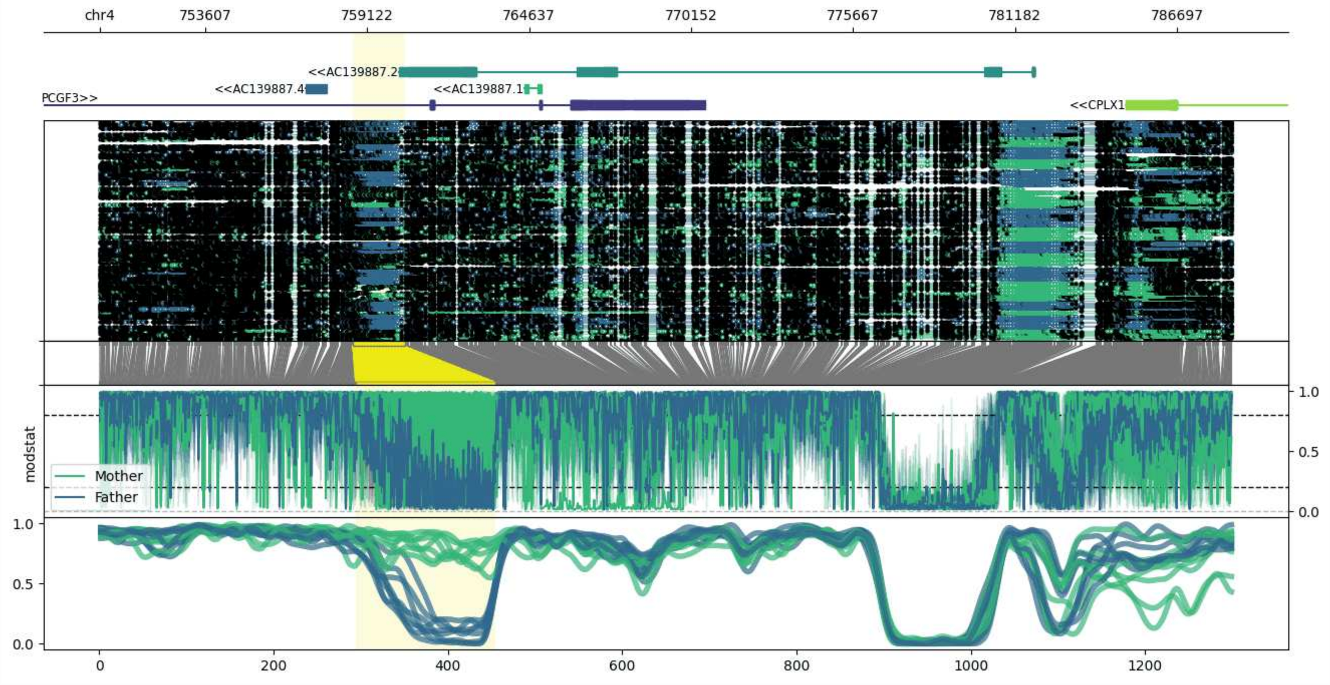


**chr4:750000-788611:** Region with 1 DMR containing genes: AC139887.1, AC139887.2, AC139887.4, CPLX1, PCGF3.

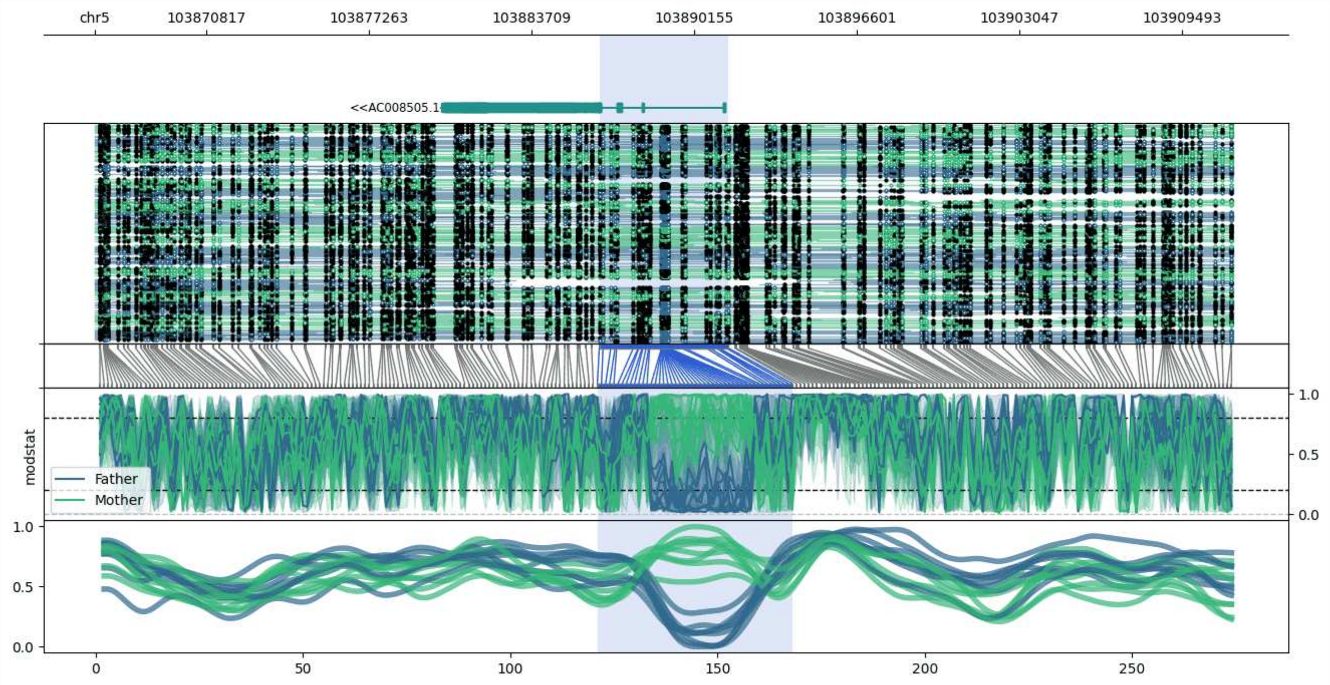


**chr5:103866403-103911530:** Region with 1 DMR containing genes: AC008505.1.

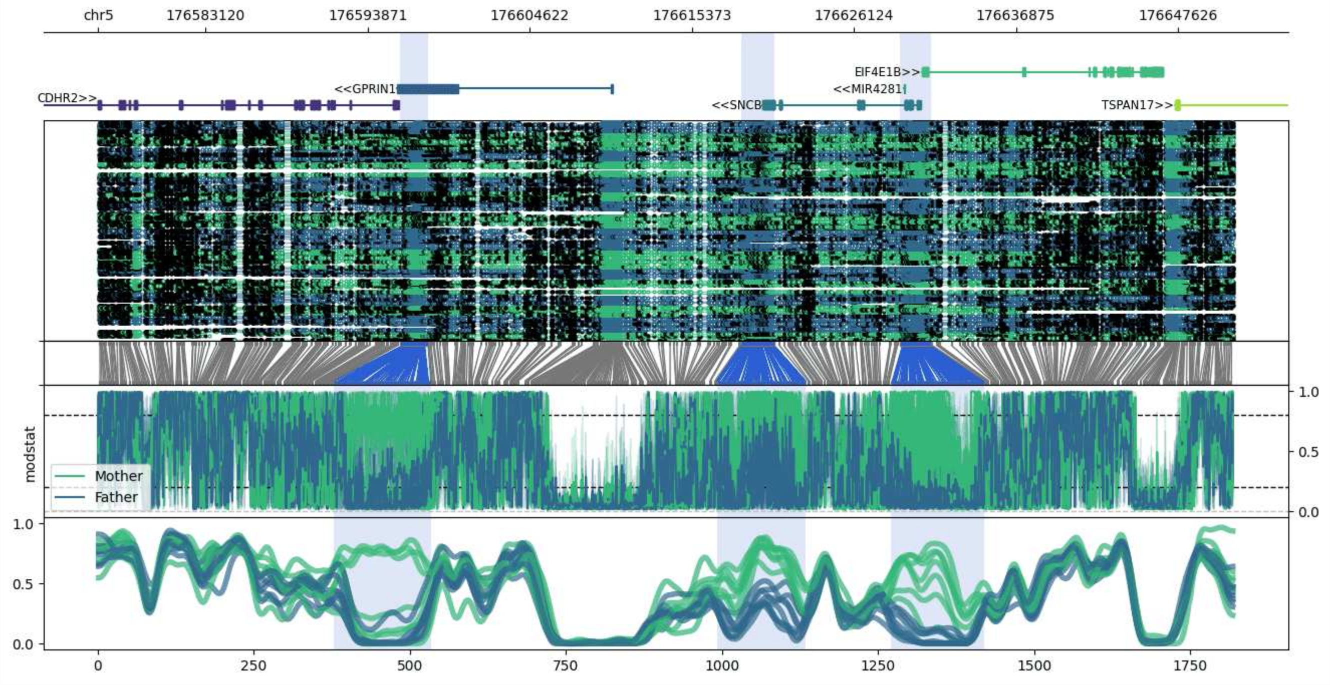


**chr5:176576007-176651268:** Region with 3 DMRs containing genes: CDHR2, EIF4E1B, GPRIN1, MIR4281, SNCB, TSPAN17.

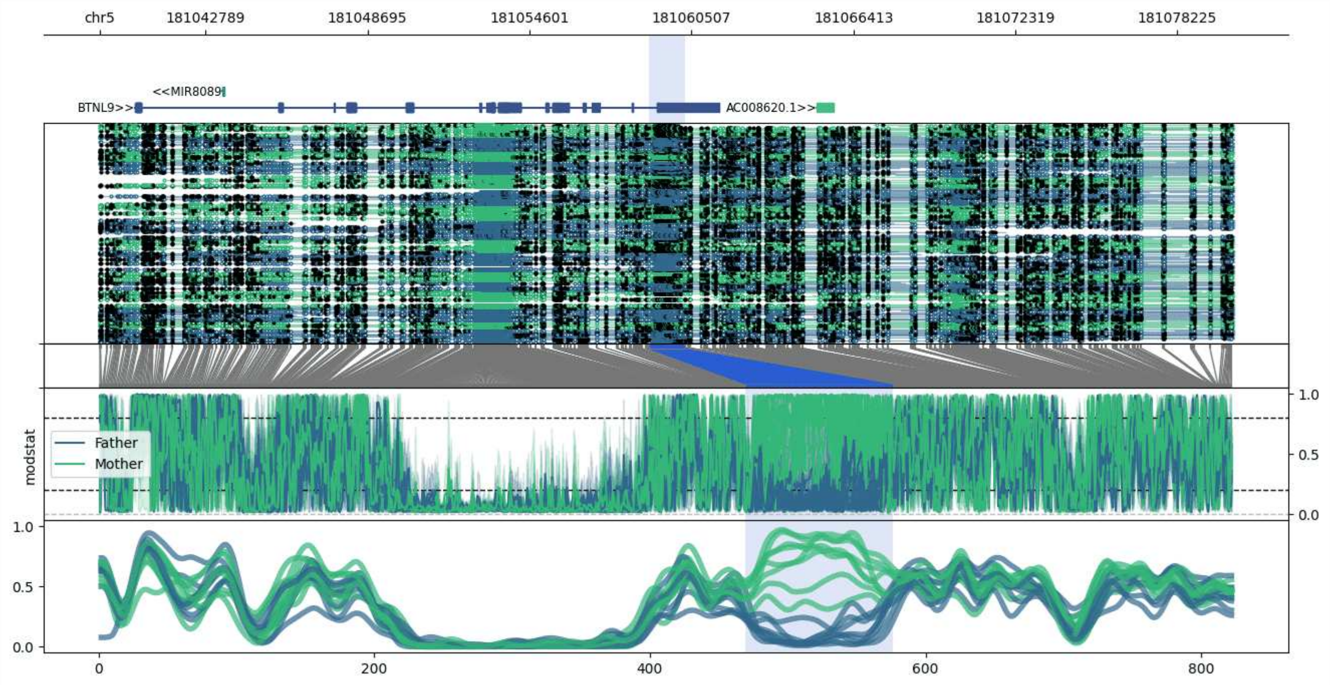


**chr5:181038942-181080284:** Region with 1 DMR containing genes: AC008620.1, BTNL9, MIR8089.

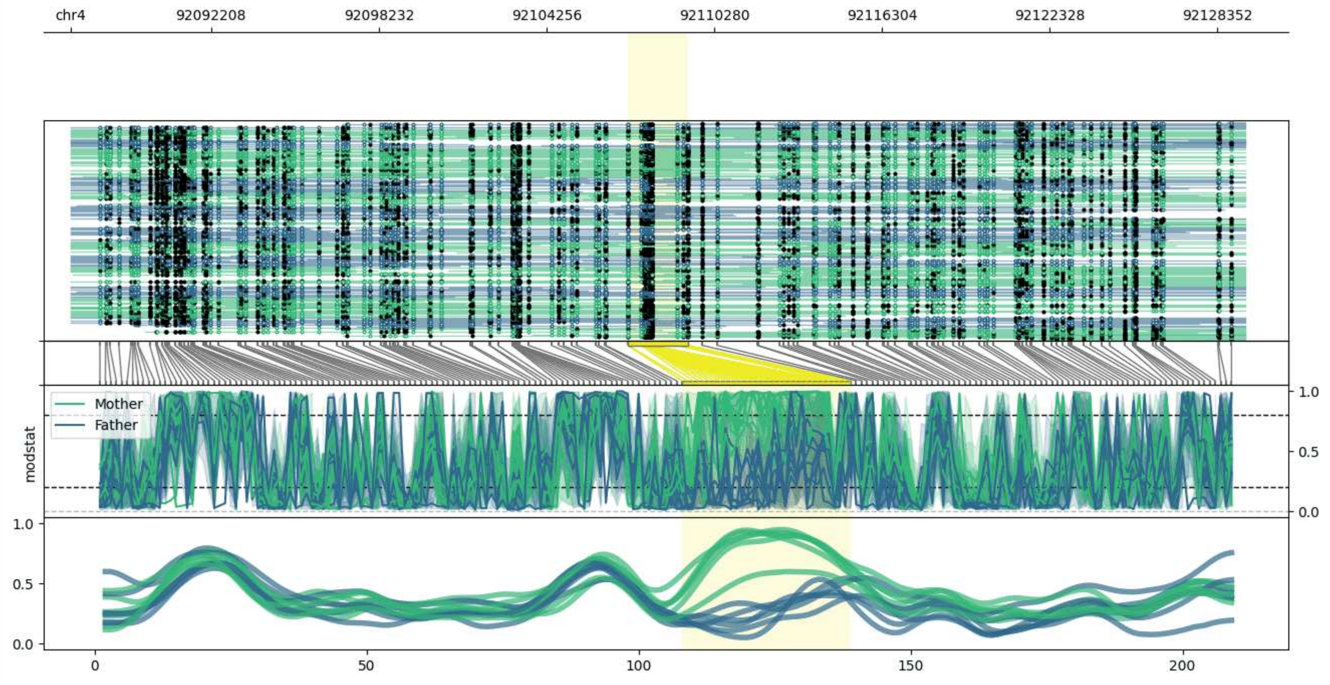


**chr4:92087166-92129337:** Region with 1 DMR containing no genes.

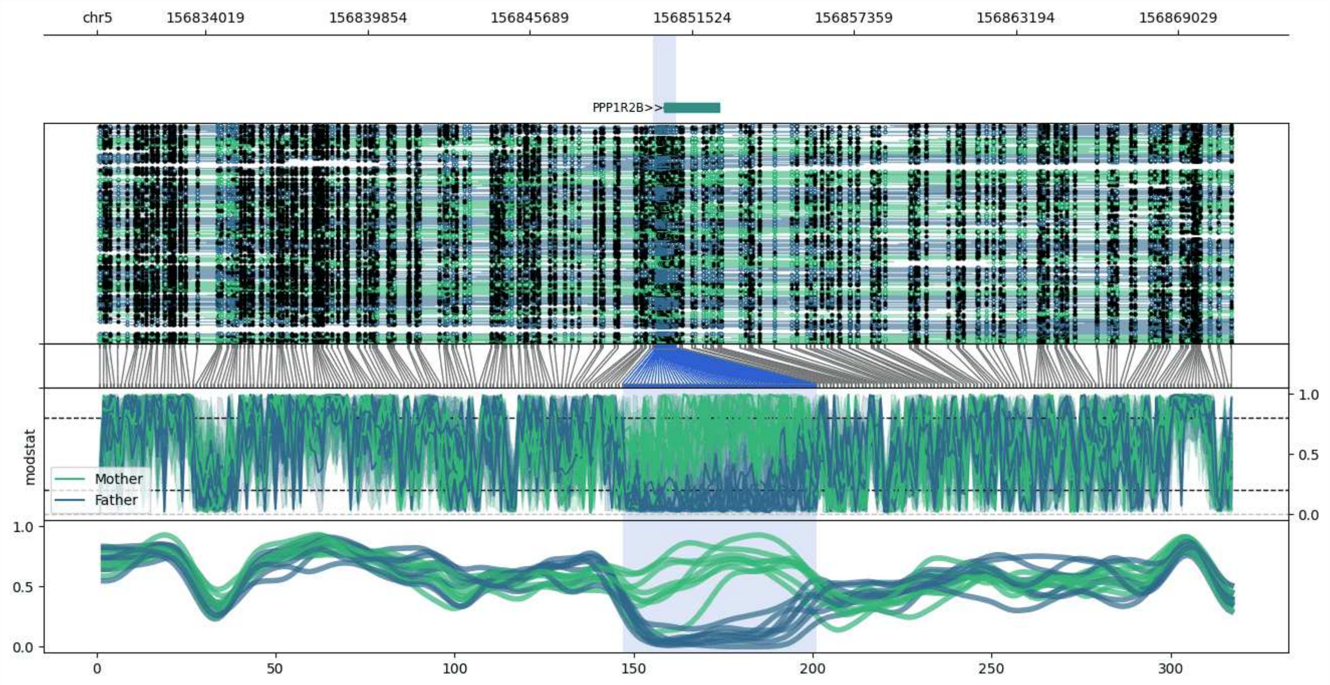


**chr5:156830124-156870973:** Region with 1 DMR containing genes: PPP1R2B.

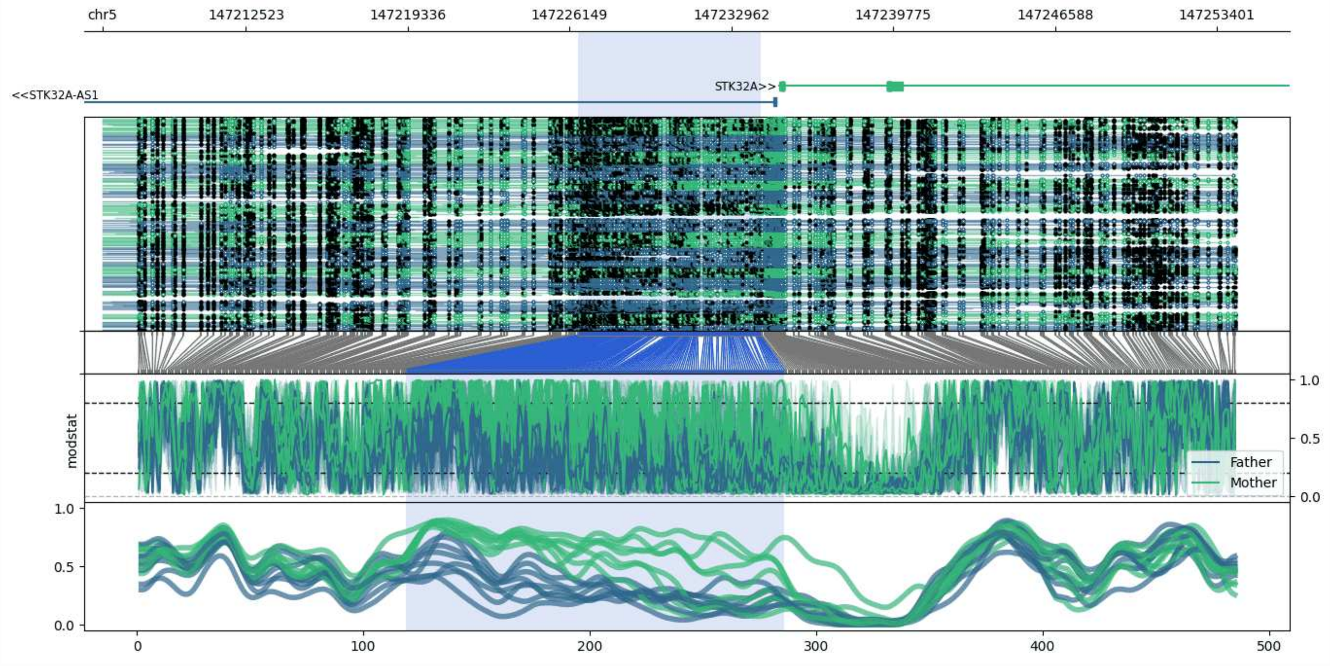


**chr5:147206492-147254183:** Region with 1 DMR containing genes: STK32A, STK32A-AS1.

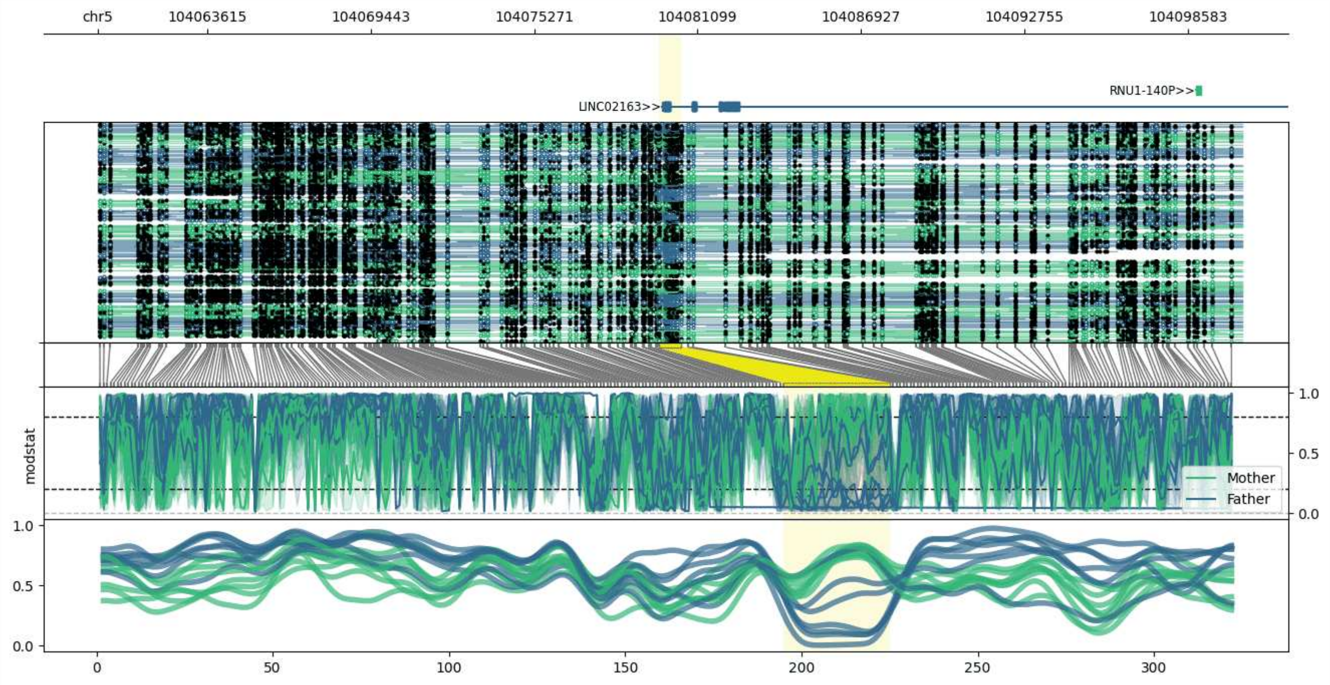


**chr5:104059711-104100510:** Region with 1 DMR containing genes: LINC02163, RNU1-140P.

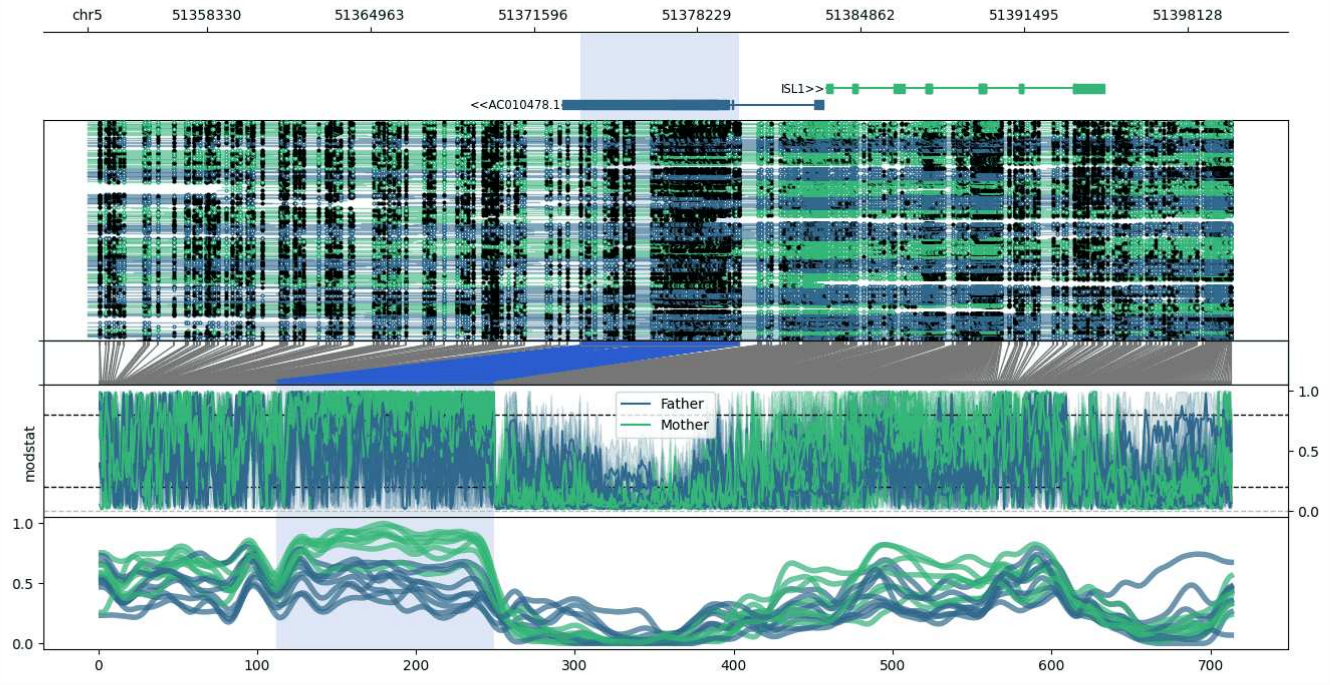


**chr5:51353488-51399921:** Region with 1 DMR containing genes: AC010478.1, HMGB1P47, ISL1.

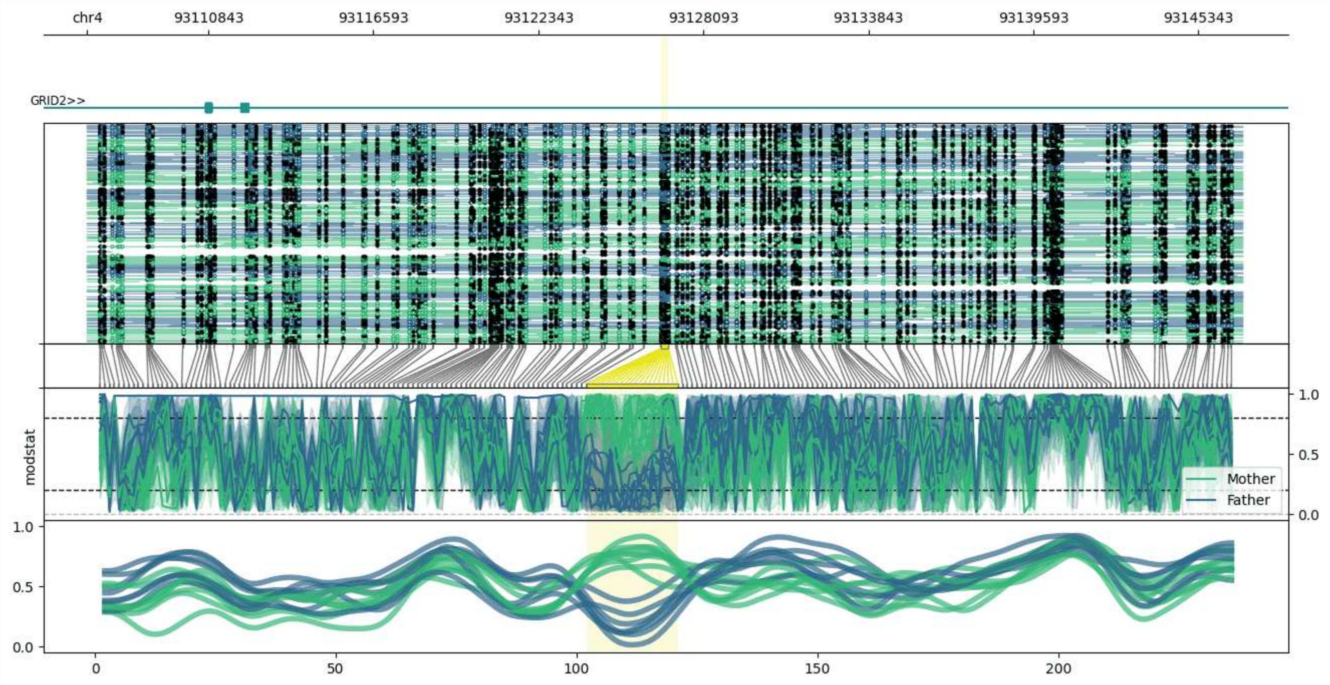


**chr4:93106619-93146875:** Region with 1 DMR containing genes: GRID2.

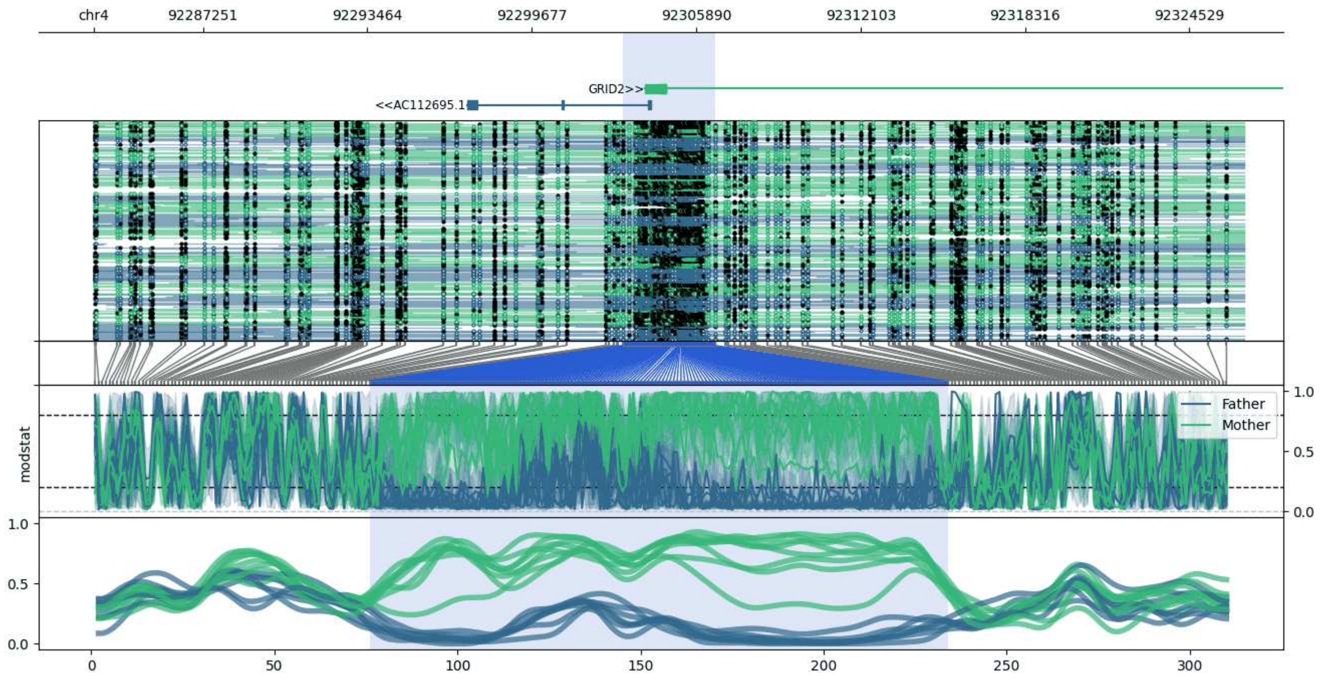


**chr4:92283116-92326607:** Region with 1 DMR containing genes: AC112695.1, GRID2, LNCPRESS2.

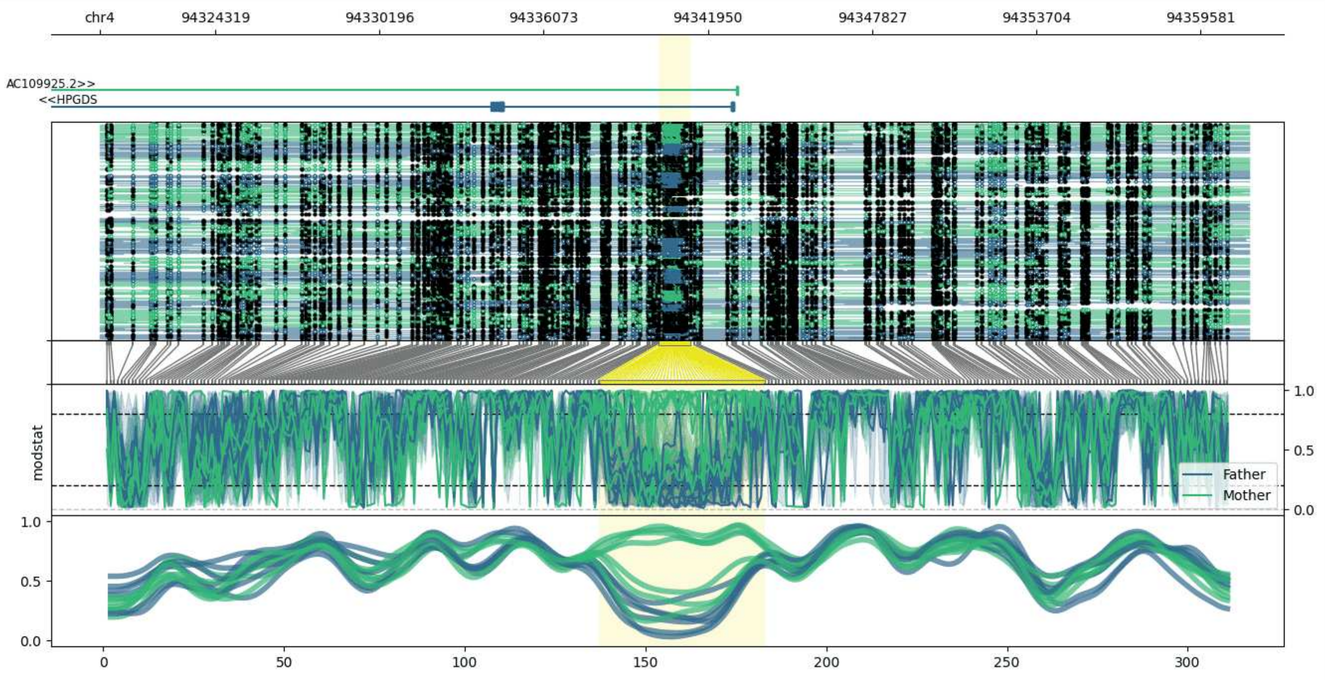


**chr4:94320176-94361315:** Region with 1 DMR containing genes: AC109925.1, AC109925.2, HPGDS.

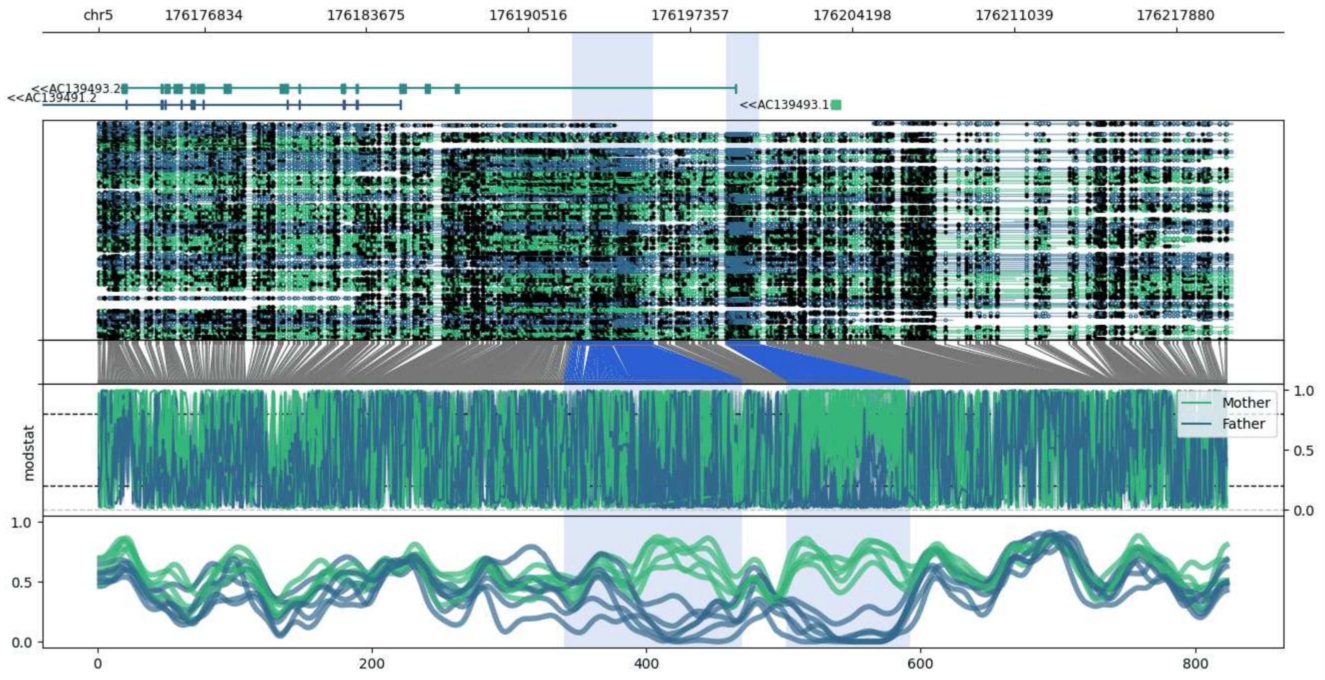


**chr5:176172364-176220255:** Region with 2 DMRs containing genes: AC139491.2, AC139491.8, AC139493.1, AC139493.2.

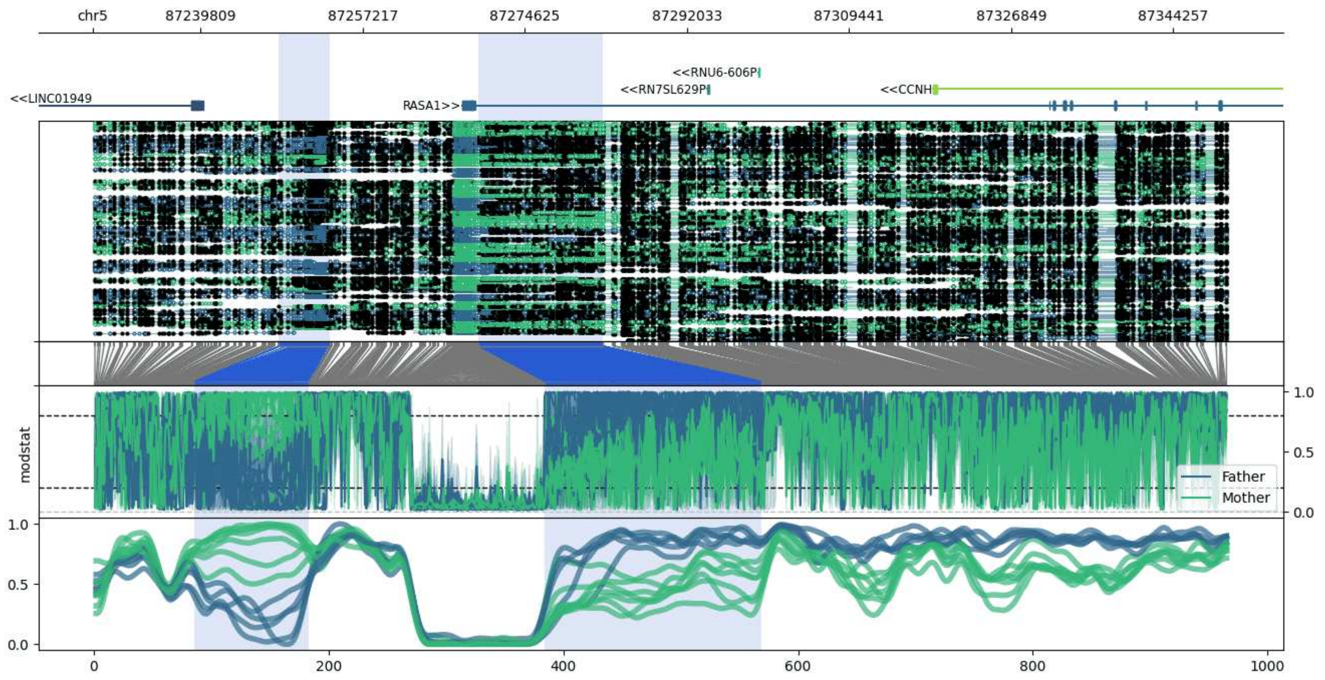


**chr5:87228235-87350097:** Region with 2 DMRs containing genes: CCNH, LINC01949, RASA1, RN7SL629P, RNU6-606P.

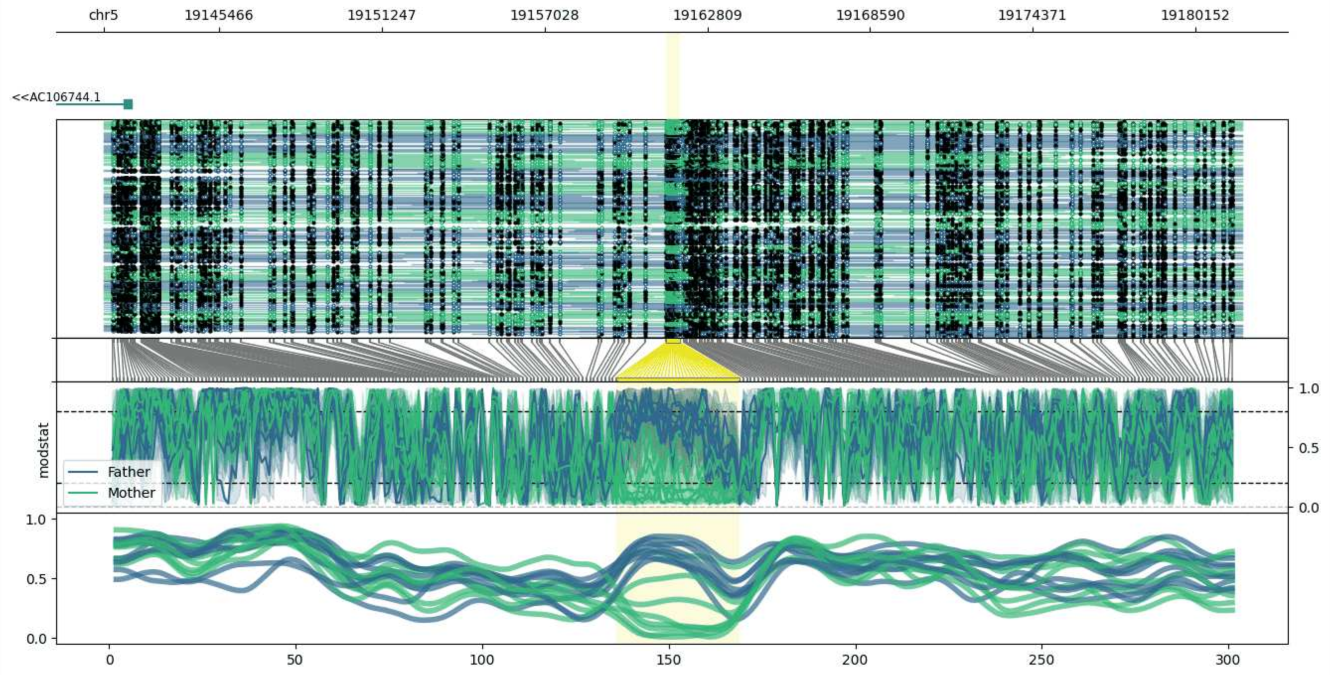


**chr5:19141355-19181826:** Region with 1 DMR containing genes: AC106744.1.

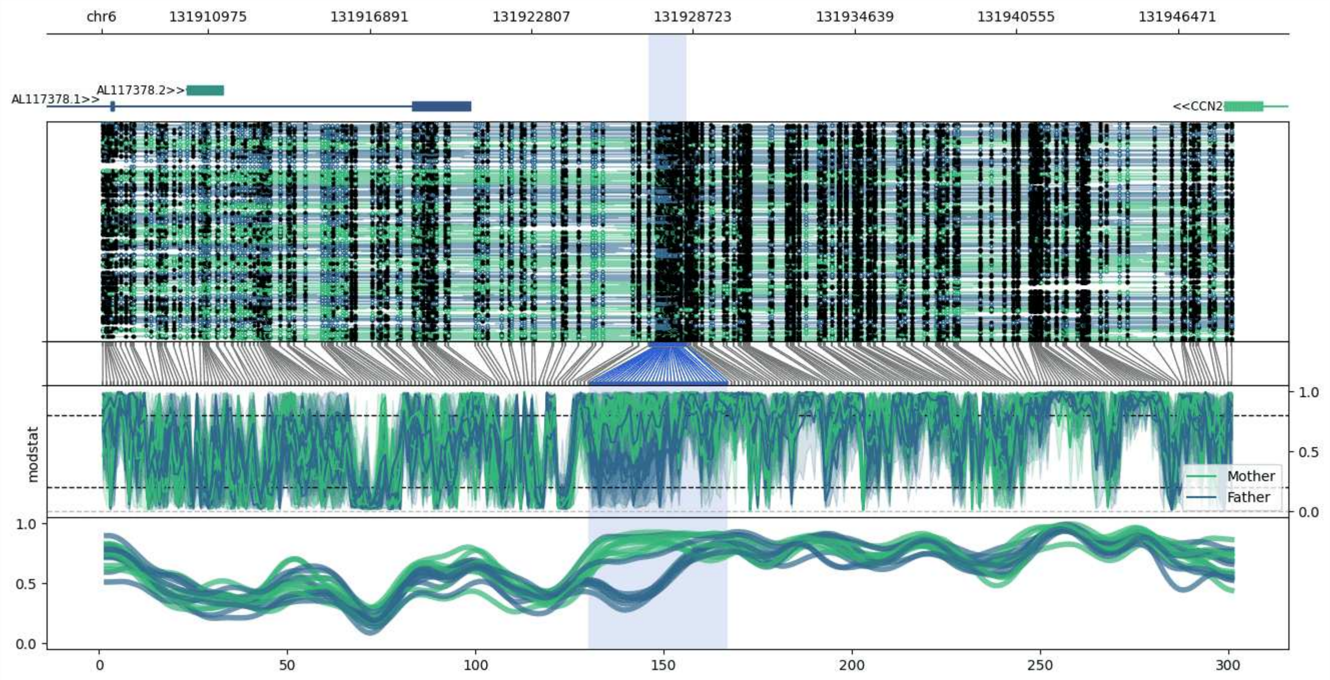


**chr6:131907082-131948496:** Region with 1 DMR containing genes: AL117378.1, AL117378.2, AL133346.1, CCN2.

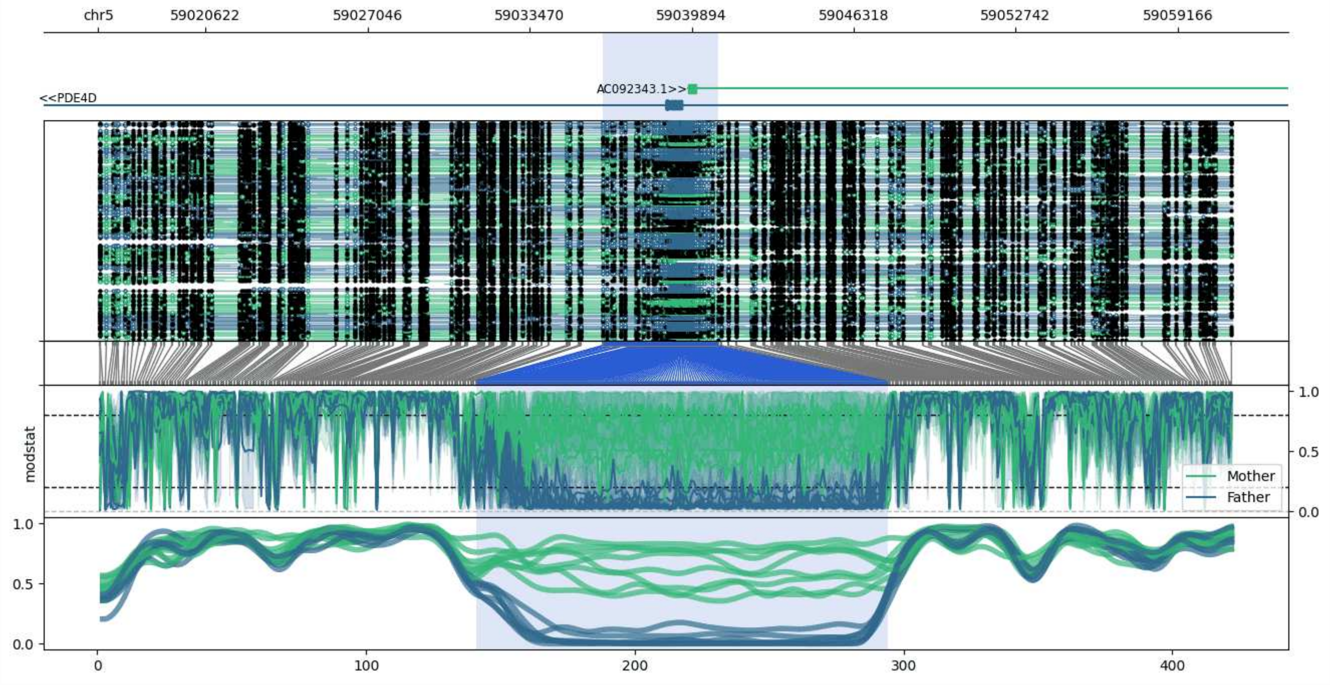


**chr5:59016366-59061339:** Region with 1 DMR containing genes: AC092343.1, PDE4D.

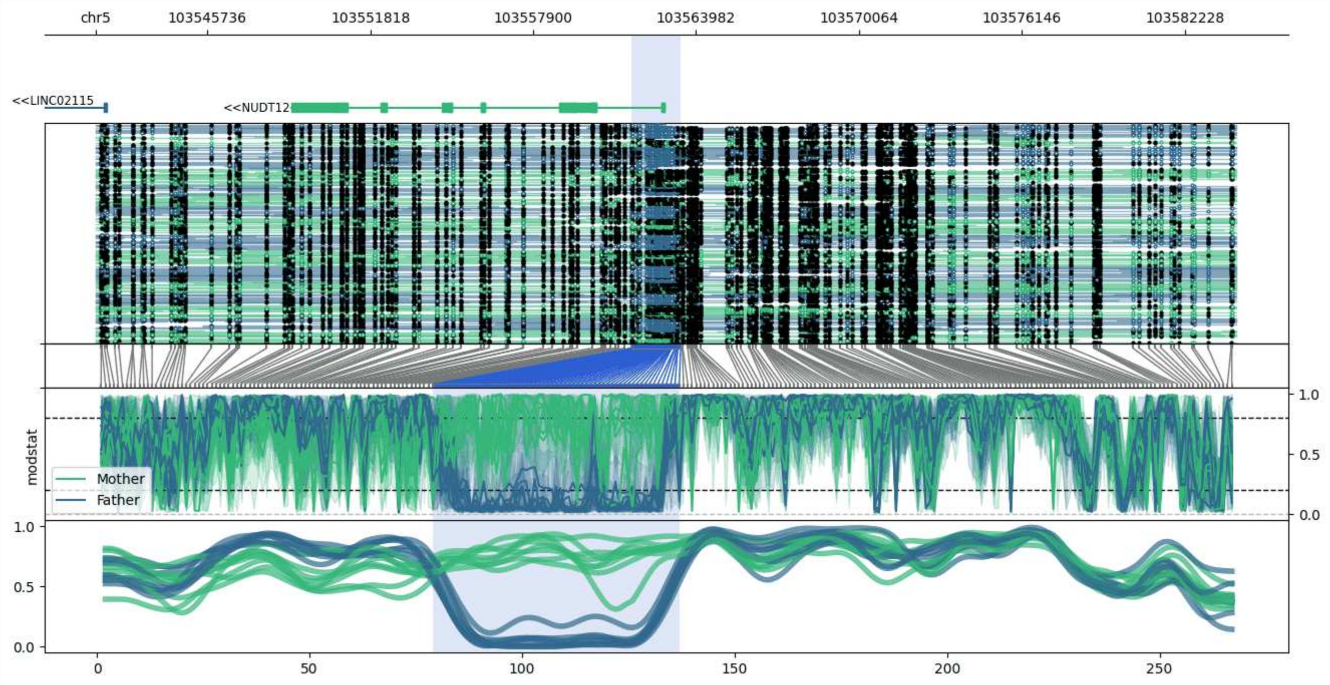


**chr5:103541559-103584137:** Region with 1 DMR containing genes: LINC02115, NUDT12.

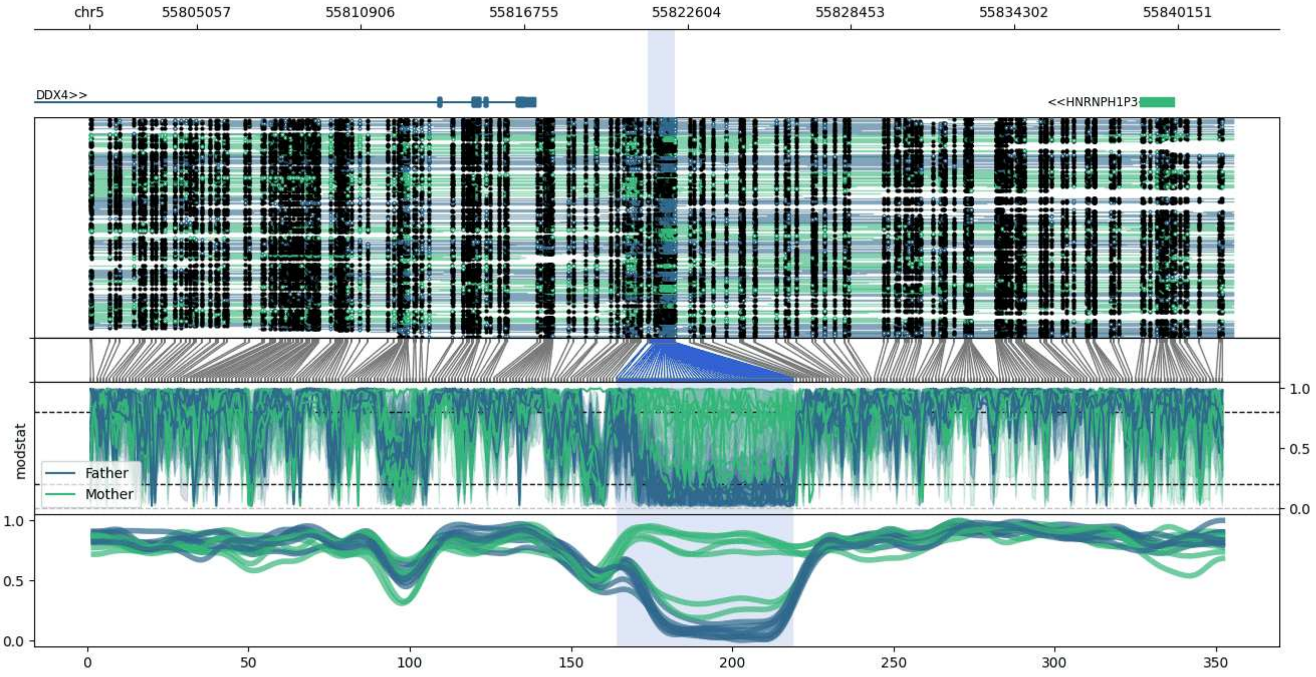


**chr5:55801188-55842136:** Region with 1 DMR containing genes: DDX4, HNRNPH1P3, IL31RA.

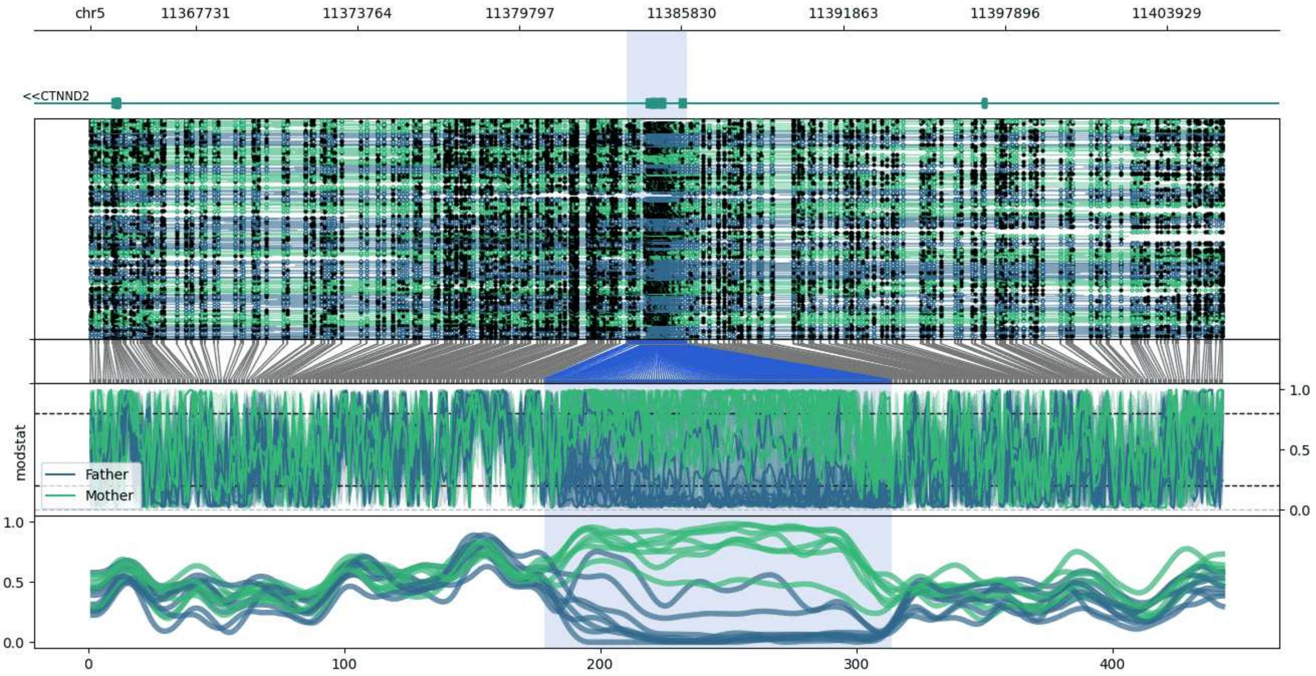


**chr5:11363791-11406026:** Region with 1 DMR containing genes: CTNND2.

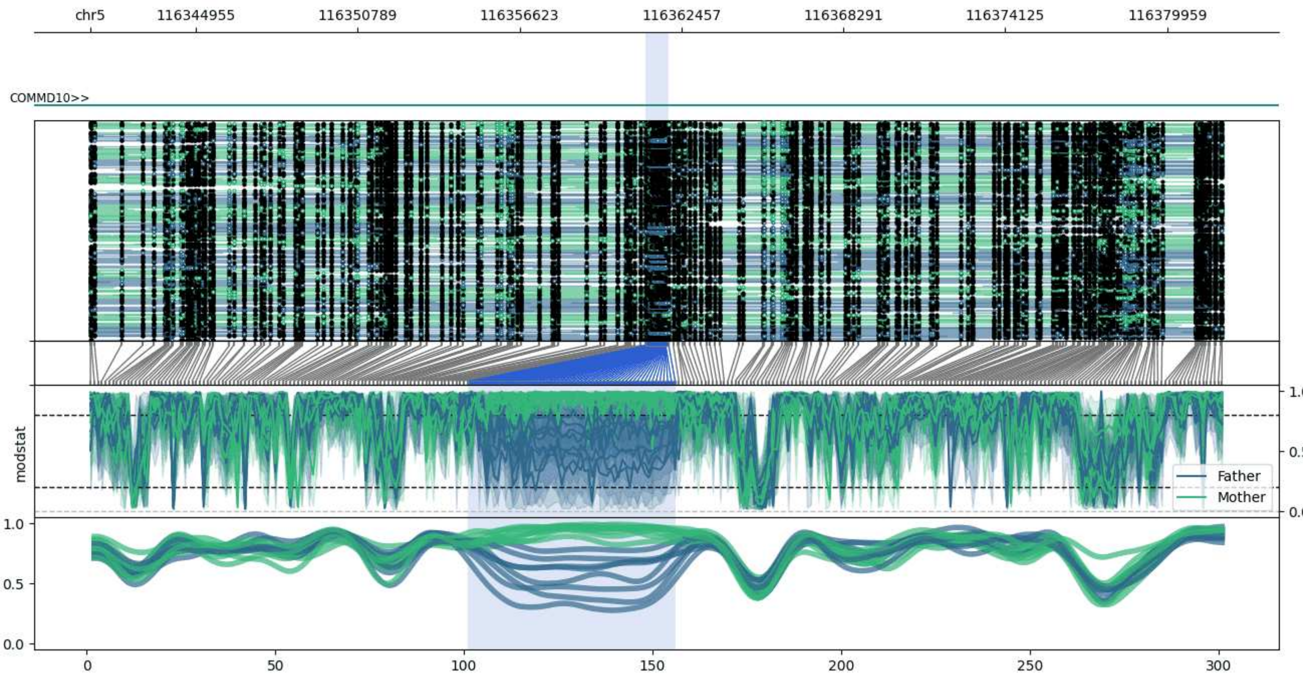


**chr5:116341156-116381998:** Region with 1 DMR containing genes: COMMD10.

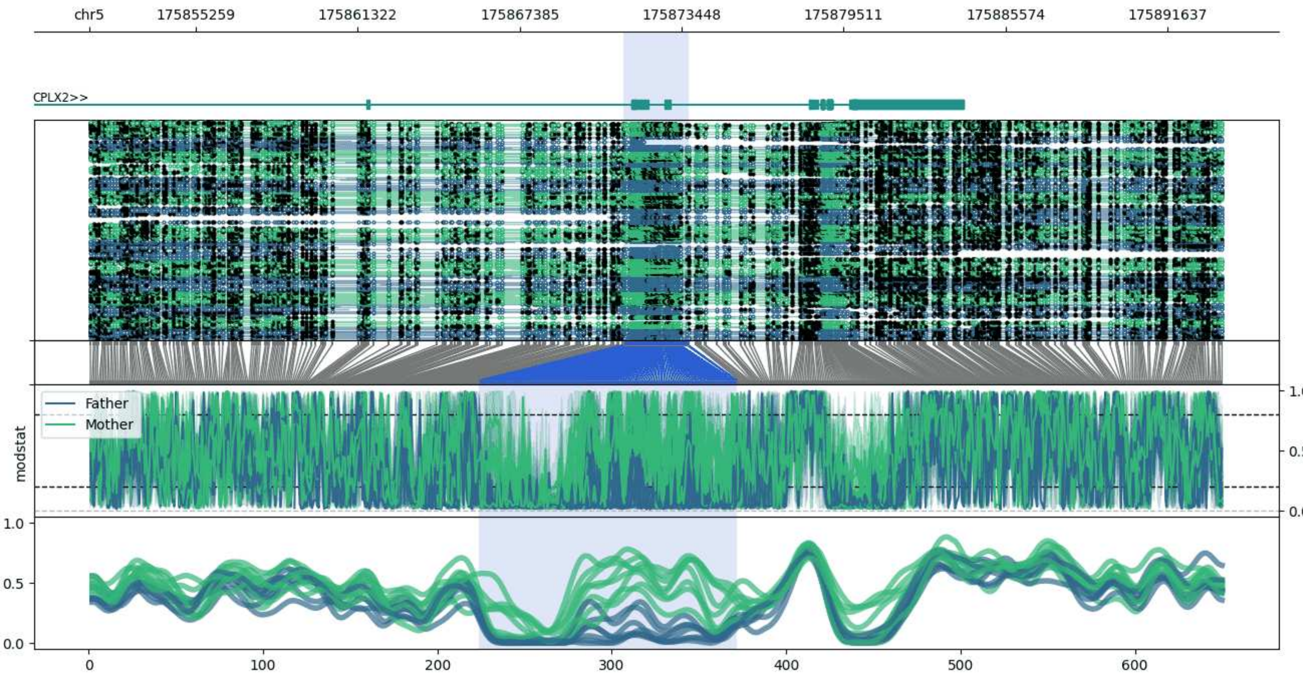


**chr5:175851260-175893701:** Region with 1 DMR containing genes: CPLX2.

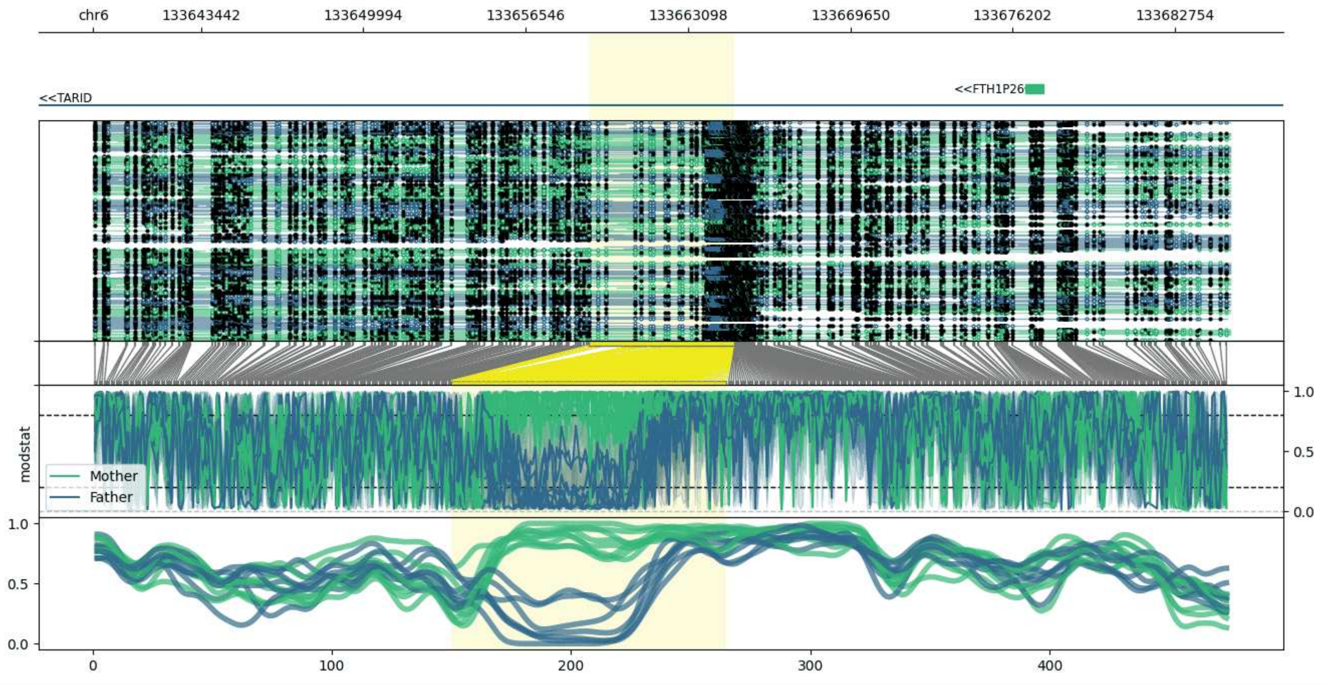


**chr6:133639099-133684967:** Region with 1 DMR containing genes: FTH1P26, TARID.

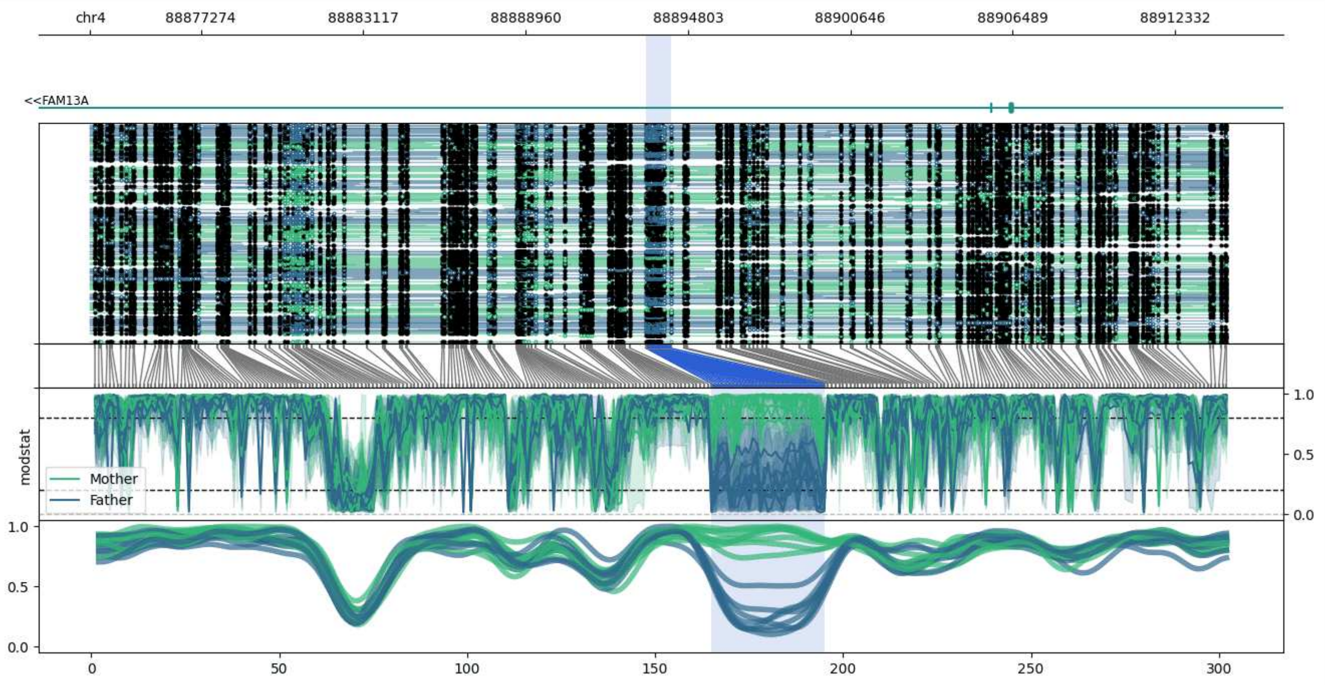


**chr4:88873290-88914194:** Region with 1 DMR containing genes: FAM13A.

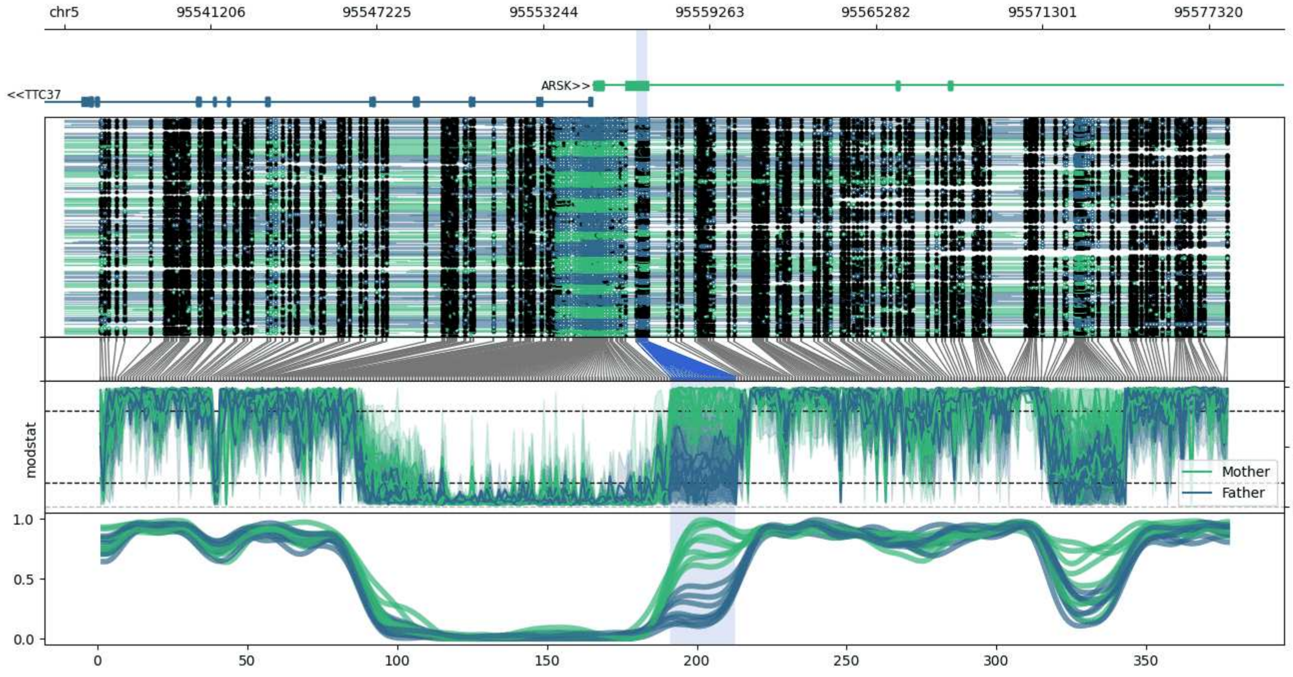


**chr5:95535919-95578055:** Region with 1 DMR containing genes: ARSK, TTC37.

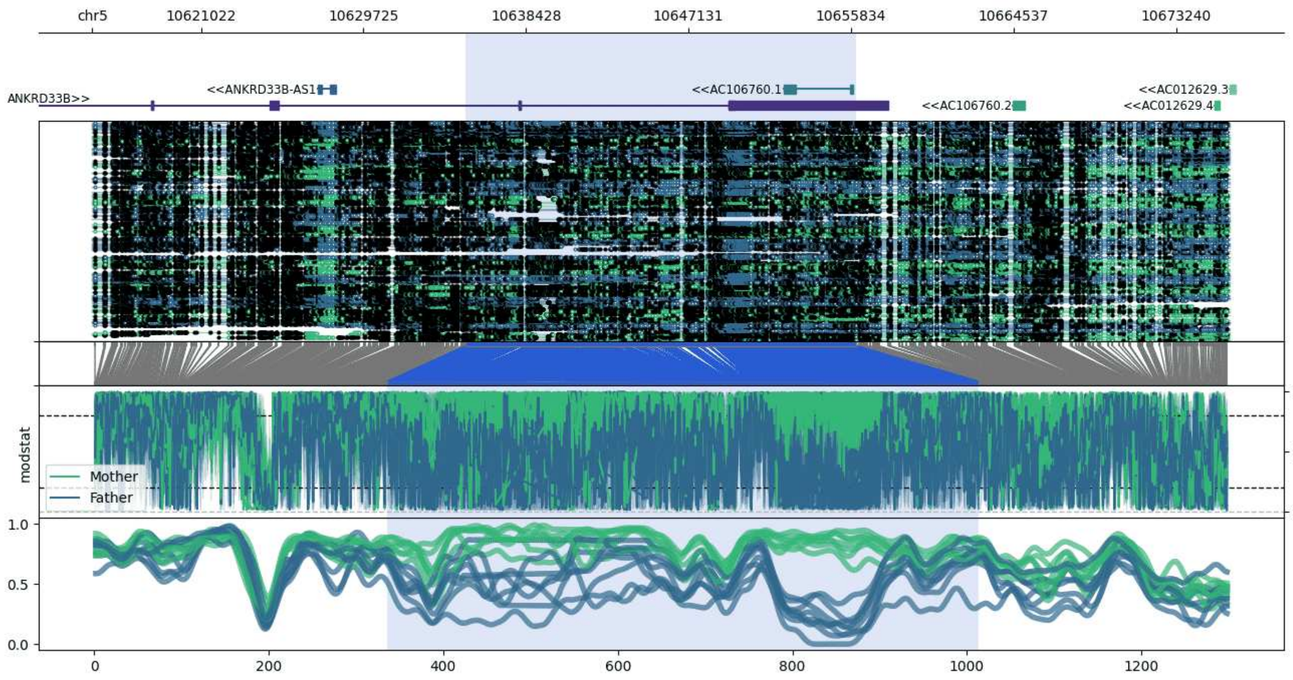


**chr5:10615201-10676124:** Region with 1 DMR containing genes: AC012629.3, AC012629.4, AC106760.1, AC106760.2, ANKRD33B, ANKRD33B-AS1, DAP.

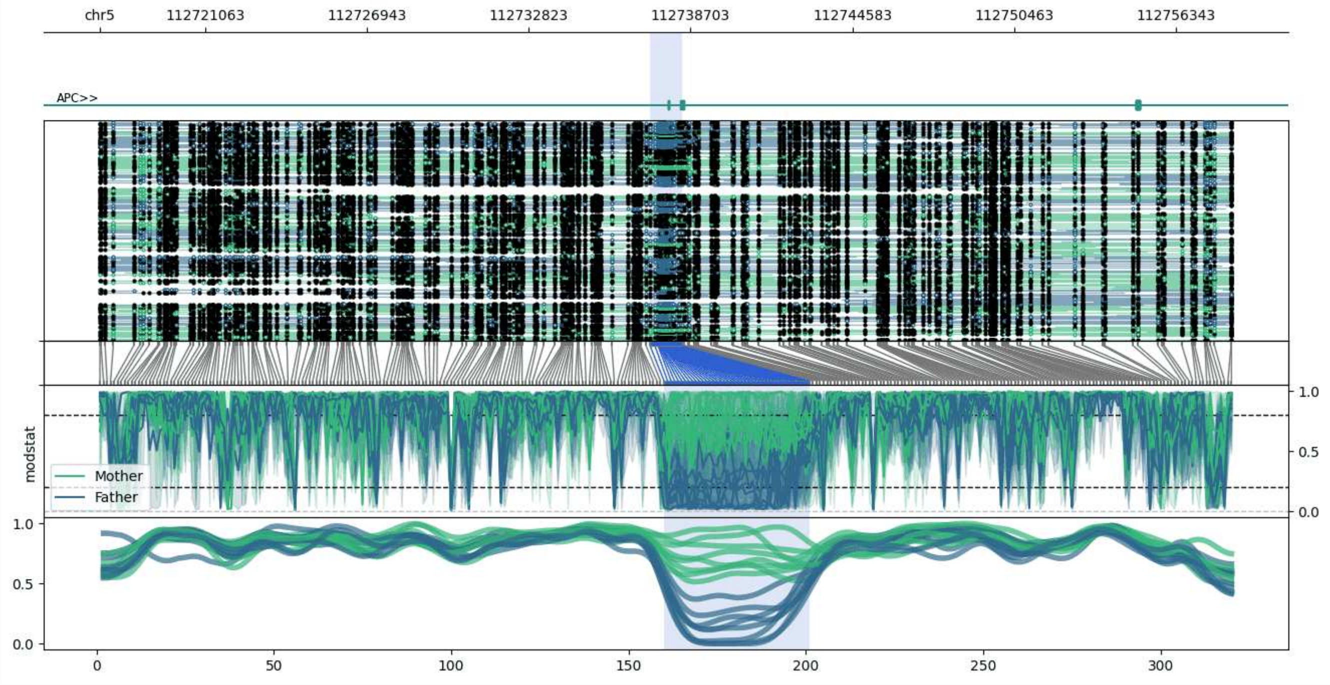


**chr5:112717228-112758392:** Region with 1 DMR containing genes: APC.

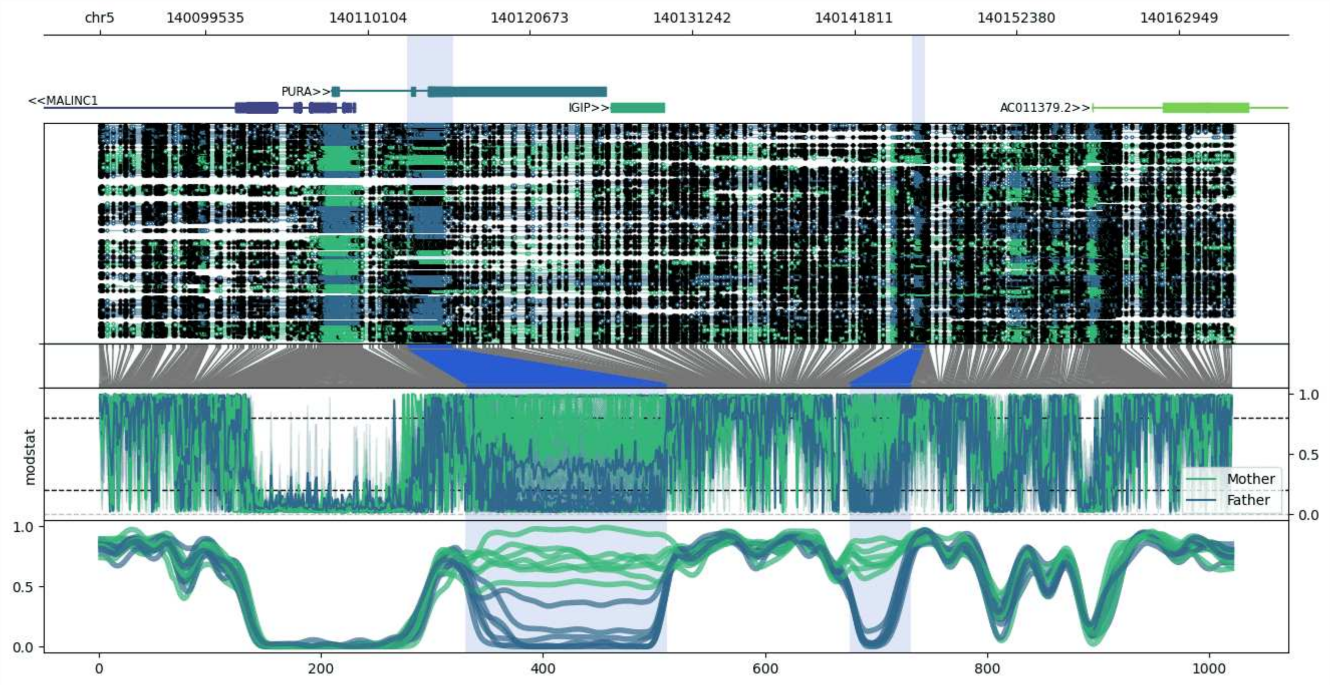


**chr5:140092644-140166633:** Region with 2 DMRs containing genes: AC011379.2, CYSTM1, IGIP, MALINC1, PURA.

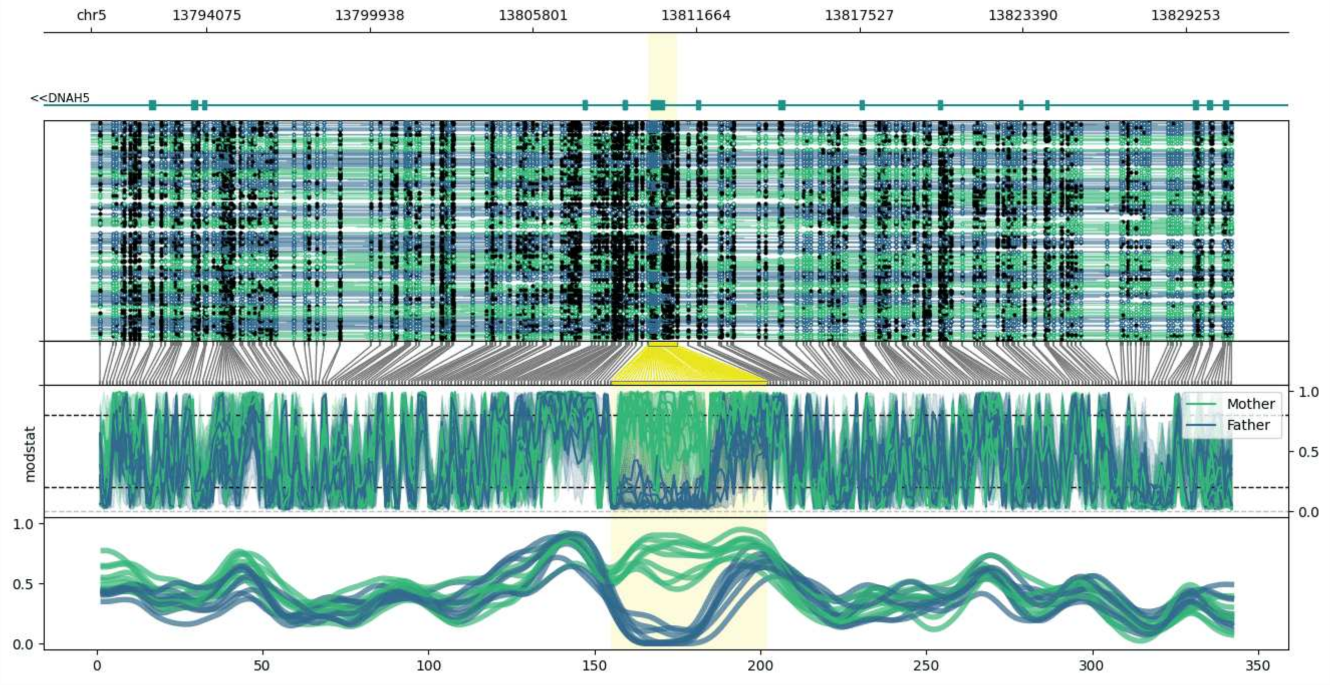


**chr5:13789929-13830976:** Region with 1 DMR containing genes: DNAH5.

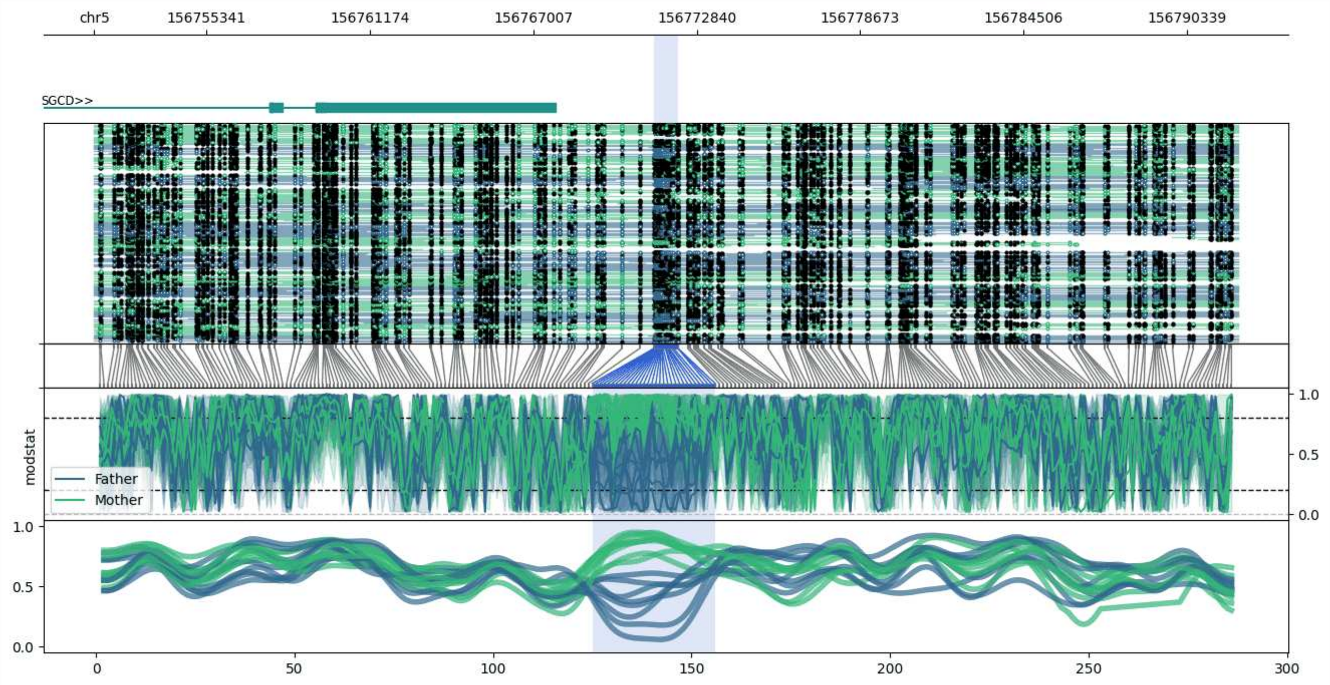


**chr5:156751327-156792163:** Region with 1 DMR containing genes: SGCD.

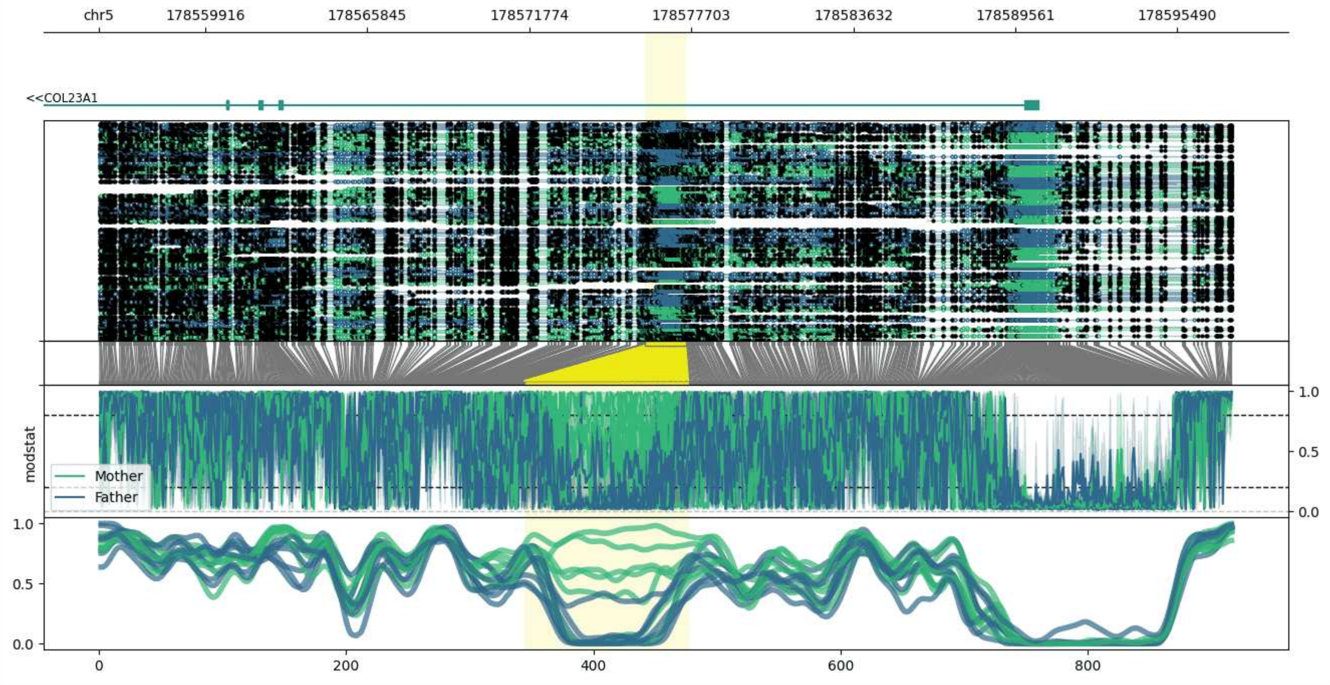


**chr5:178556015-178597524:** Region with 1 DMR containing genes: CLK4, COL23A1.

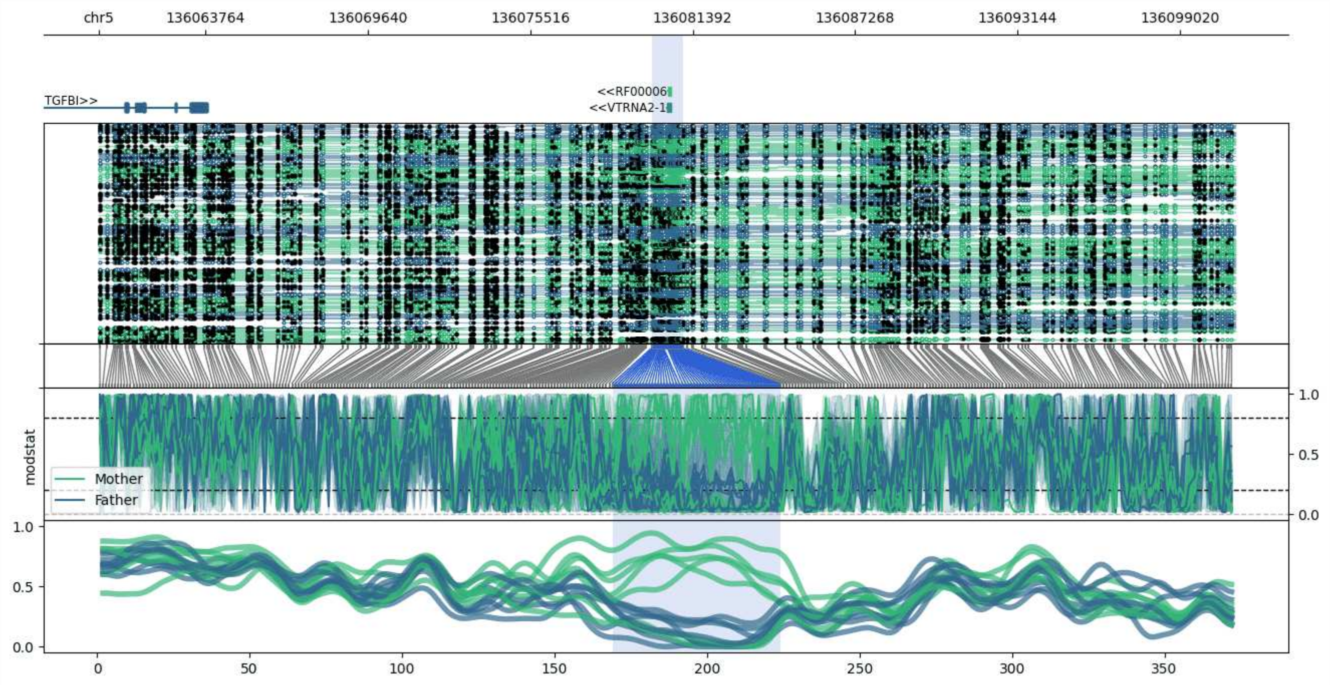


**chr5:136059901-136101035:** Region with 1 DMR containing genes: RF00006, TGFBI, VTRNA2-1.

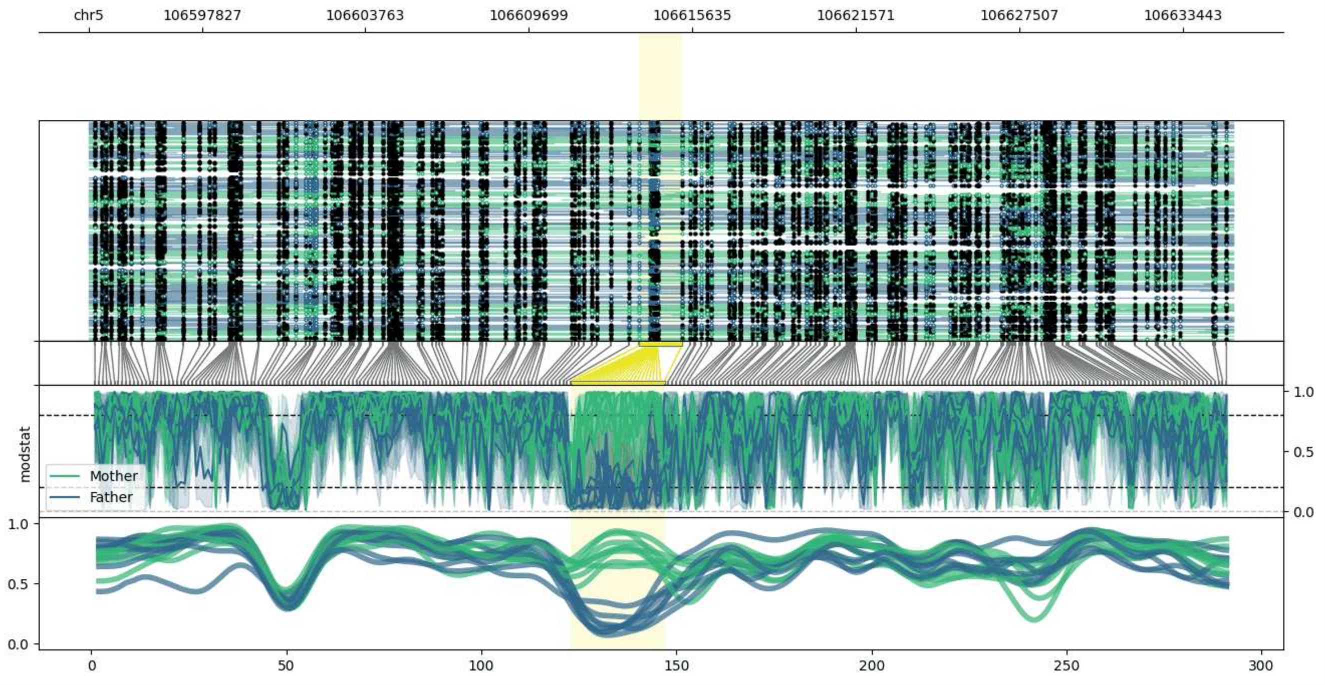


**chr5:106593712-106635267:** Region with 1 DMR containing no genes.

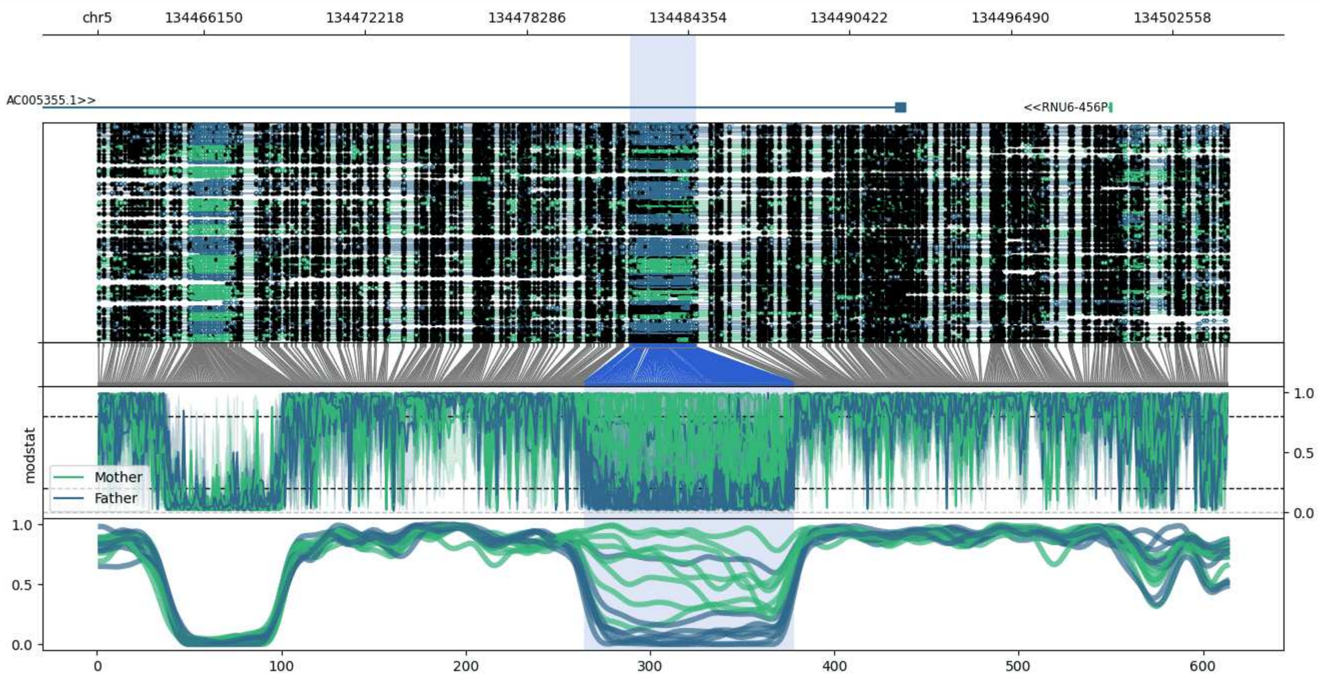


**chr5:134462157-134504637:** Region with 1 DMR containing genes: AC005355.1, AC109454.3, LINC01843, RNU6-456P.

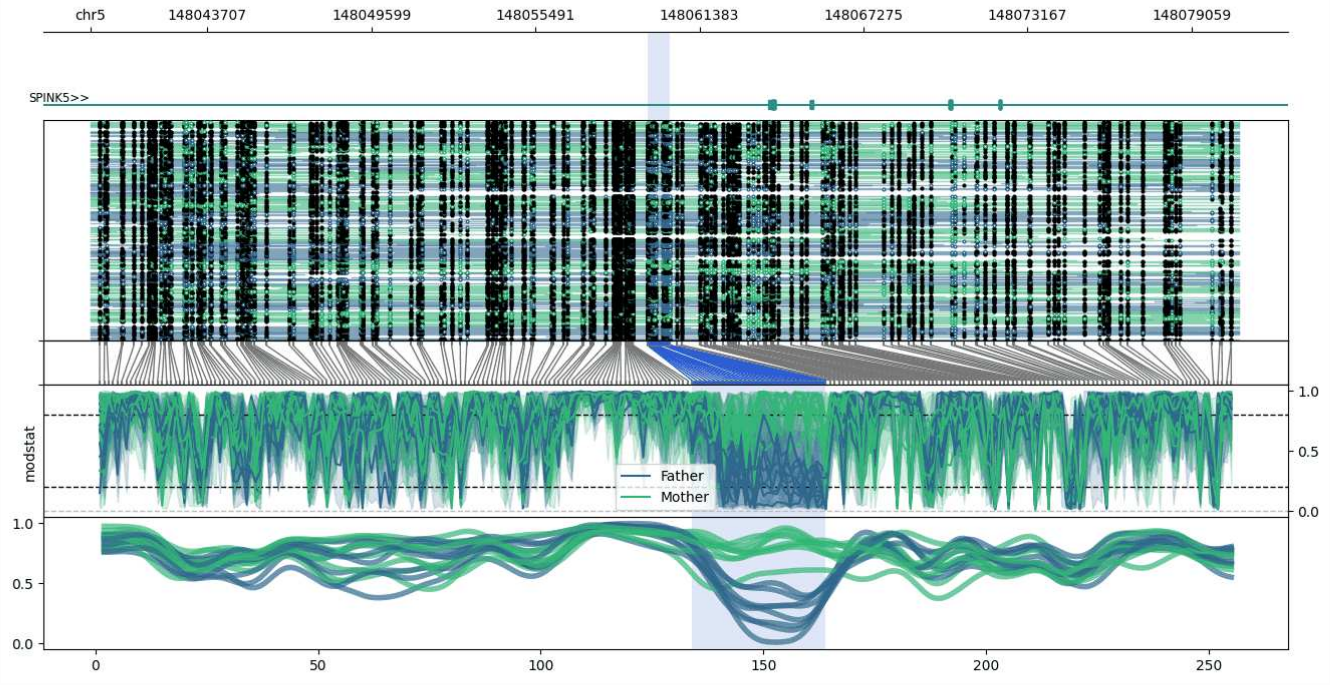


**chr5:148039516-148080761:** Region with 1 DMR containing genes: AC011346.1, SPINK5.

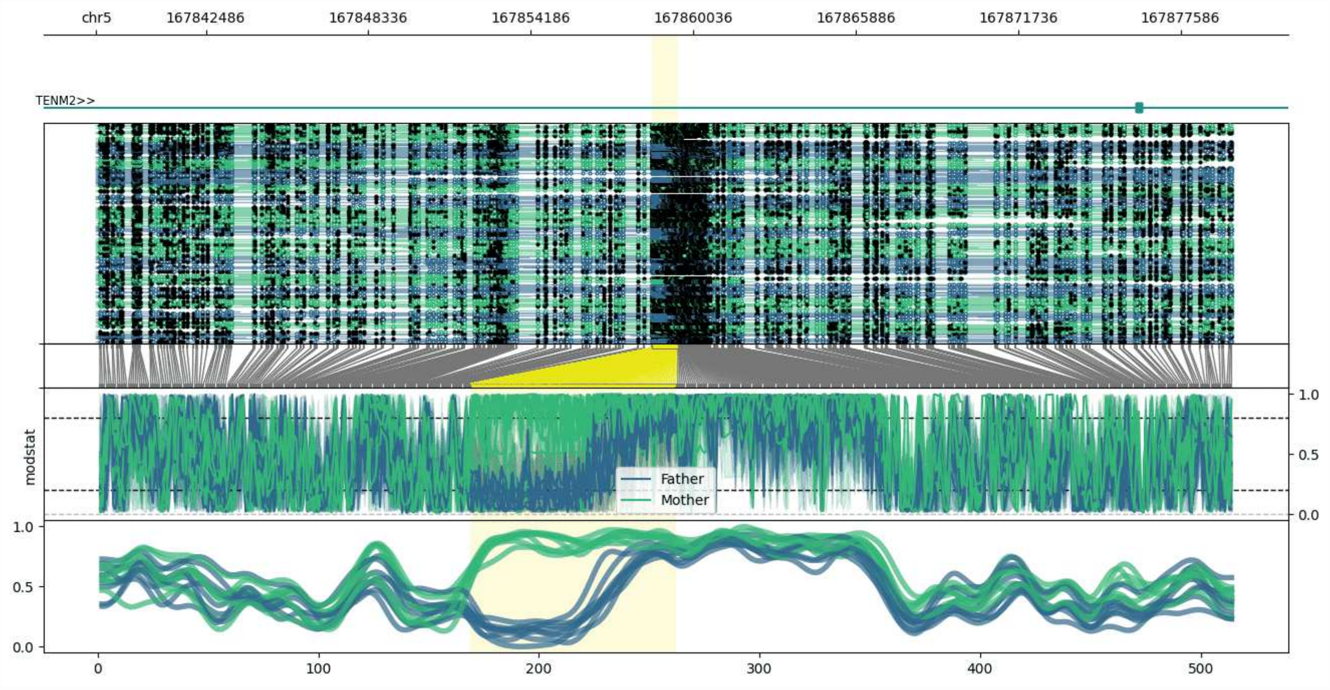


**chr5:167838534-167879490:** Region with 1 DMR containing genes: TENM2.

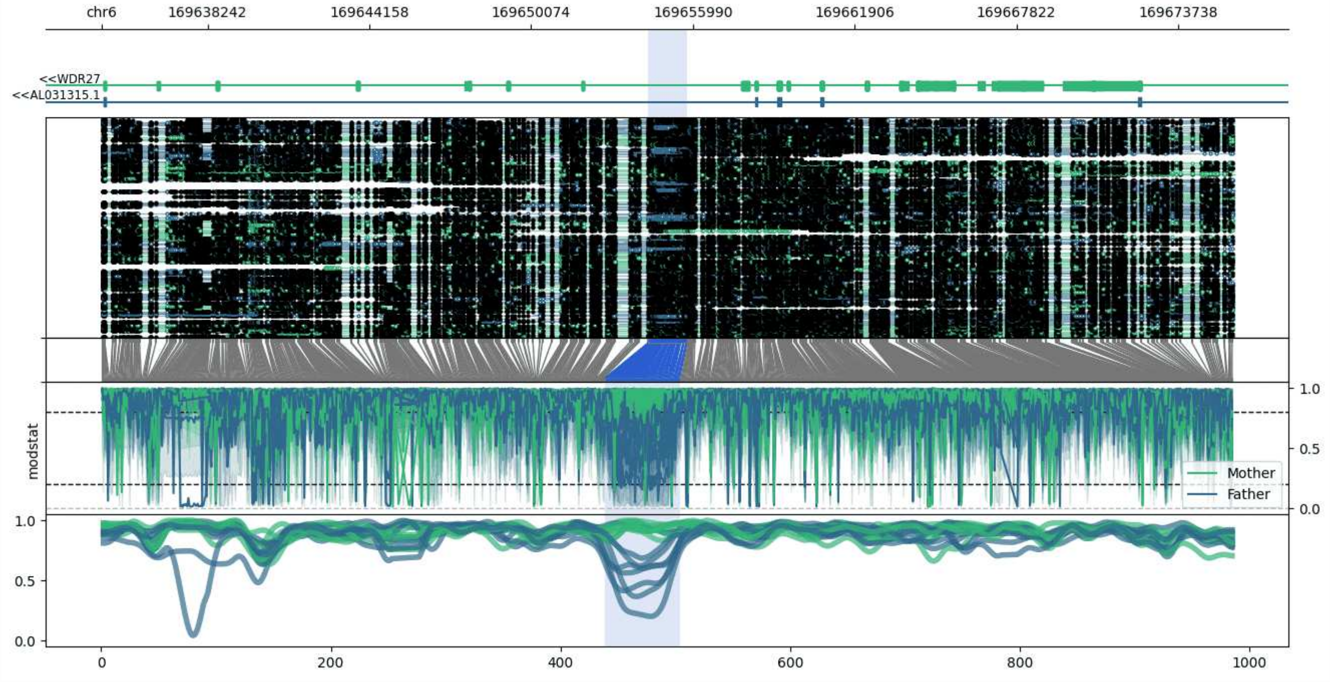


**chr6:169634366-169675781:** Region with 1 DMR containing genes: AL031315.1, WDR27.

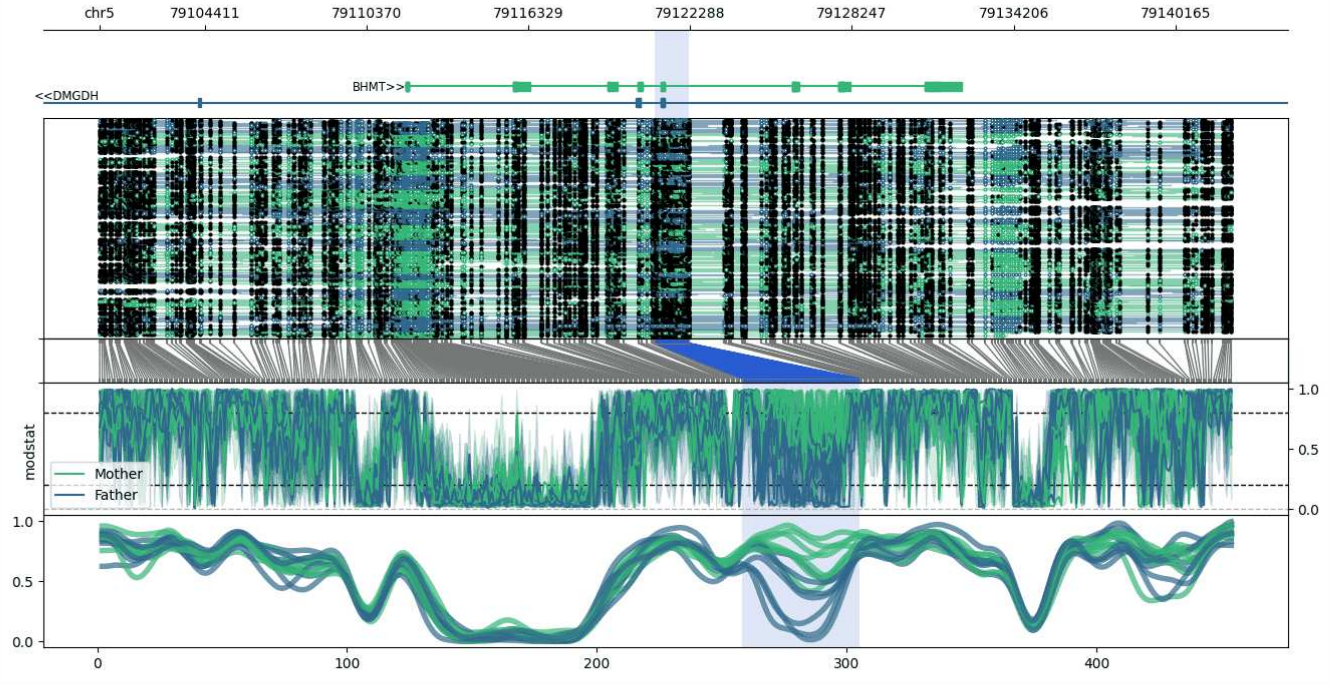


**chr5:79100520-79142235:** Region with 1 DMR containing genes: BHMT, DMGDH.

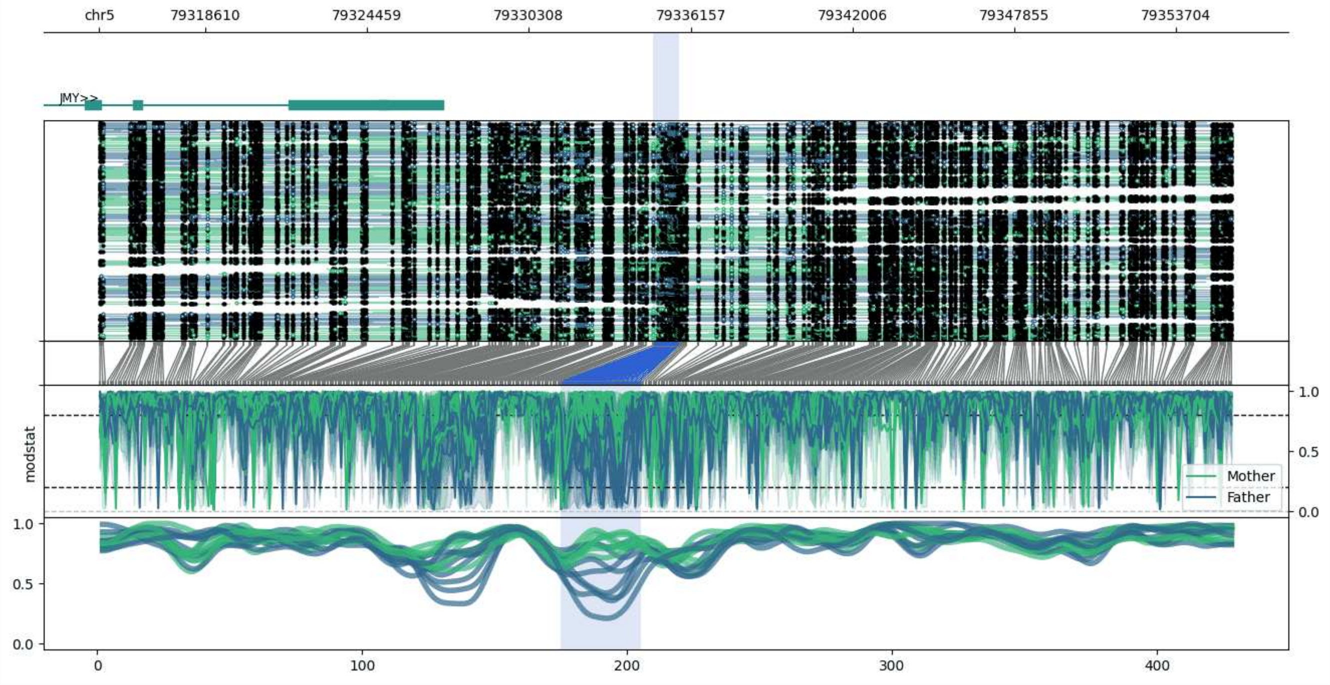


**chr5:79314781-79355725:** Region with 1 DMR containing genes: JMY.

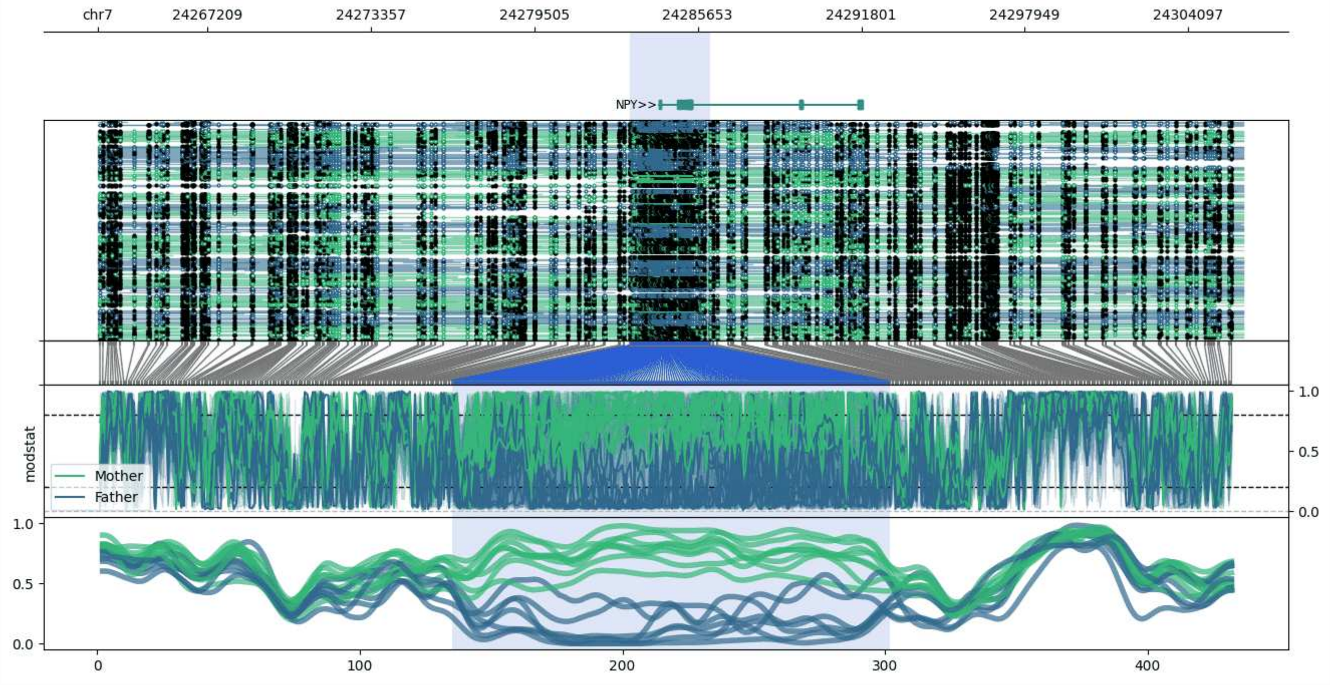


**chr7:24263098-24306140:** Region with 1 DMR containing genes: AC004485.1, NPY.

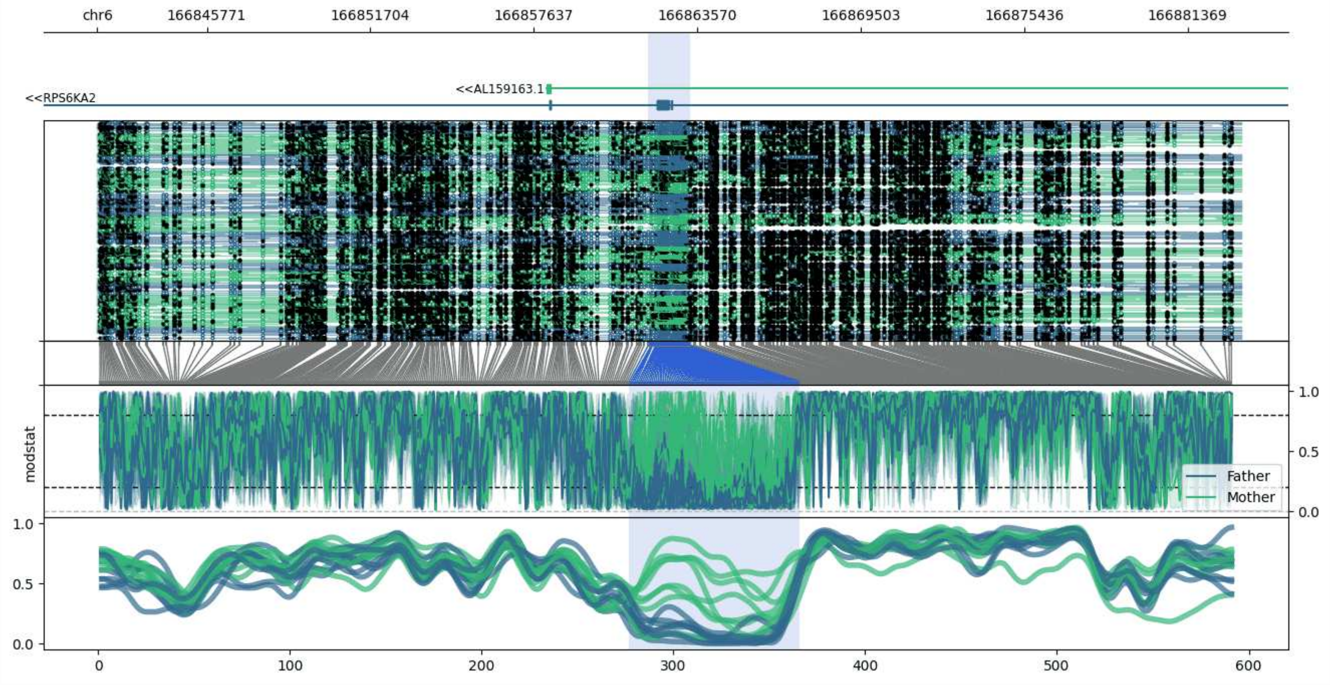


**chr6:166841792-166883324:** Region with 1 DMR containing genes: AL159163.1, RPS6KA2.

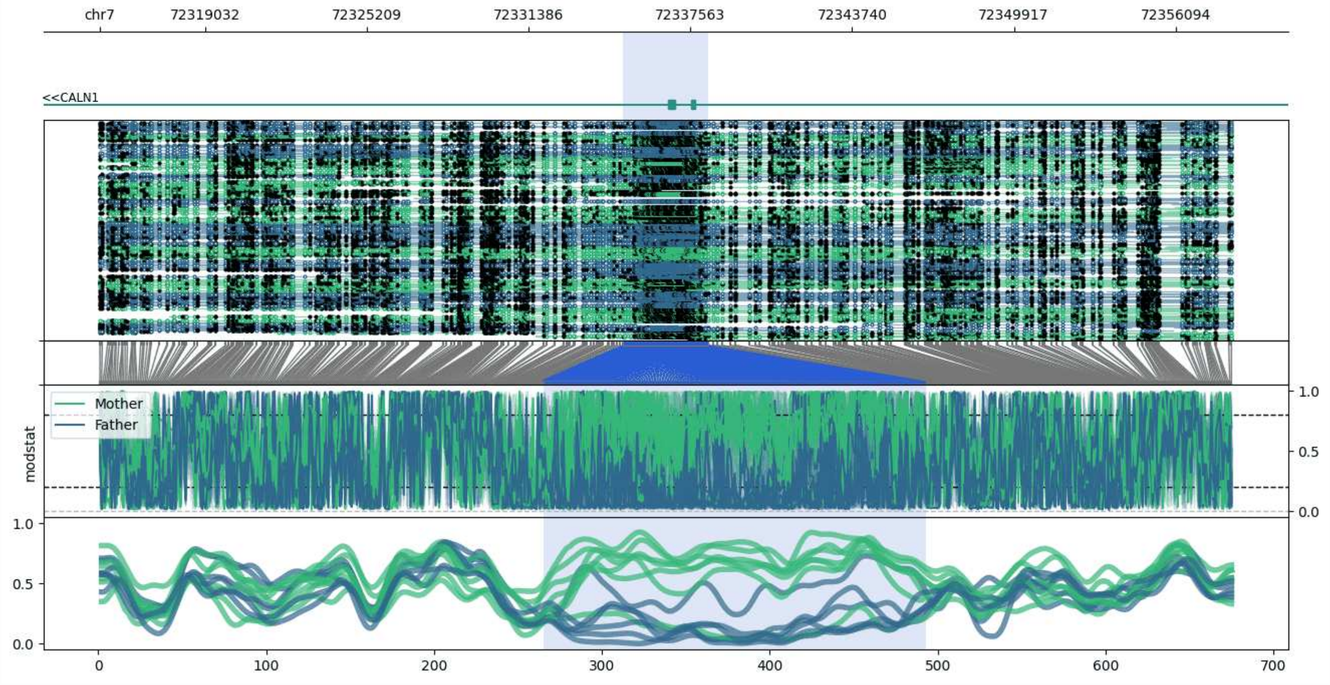


**chr7:72314996-72358241:** Region with 1 DMR containing genes: CALN1.

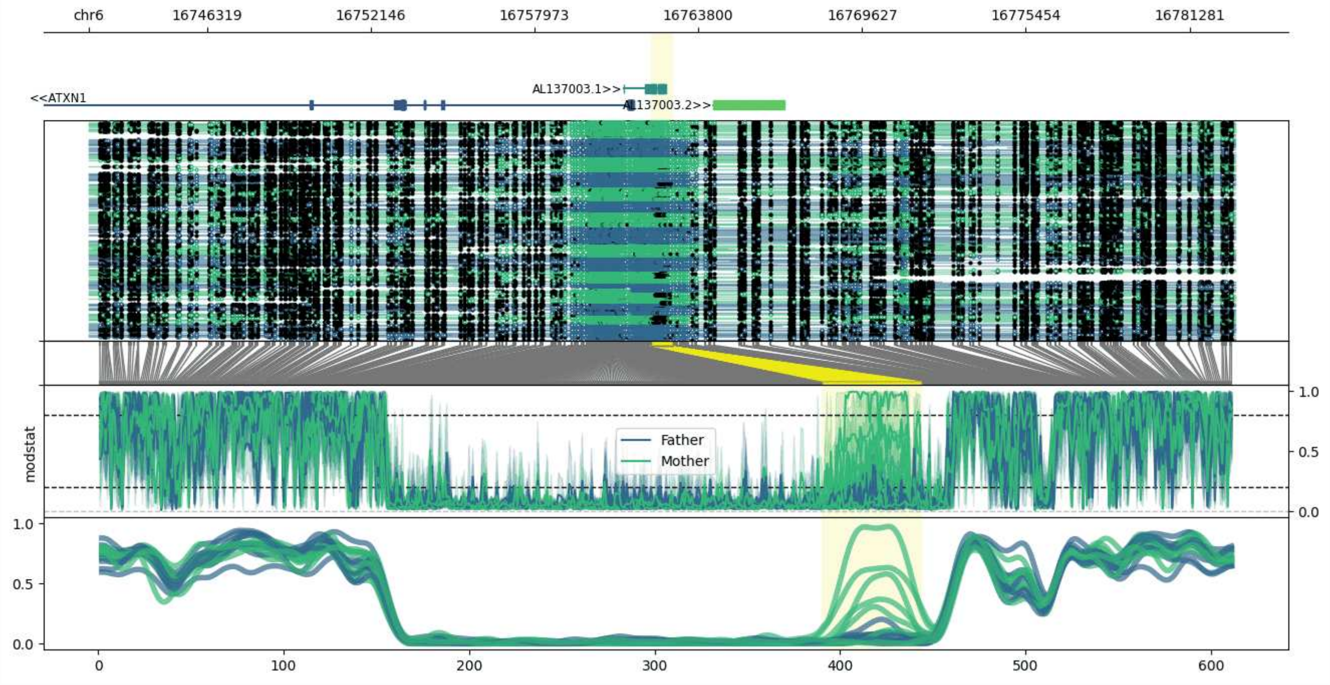


**chr6:16742098-16782893:** Region with 1 DMR containing genes: AL137003.1, AL137003.2, ATXN1.

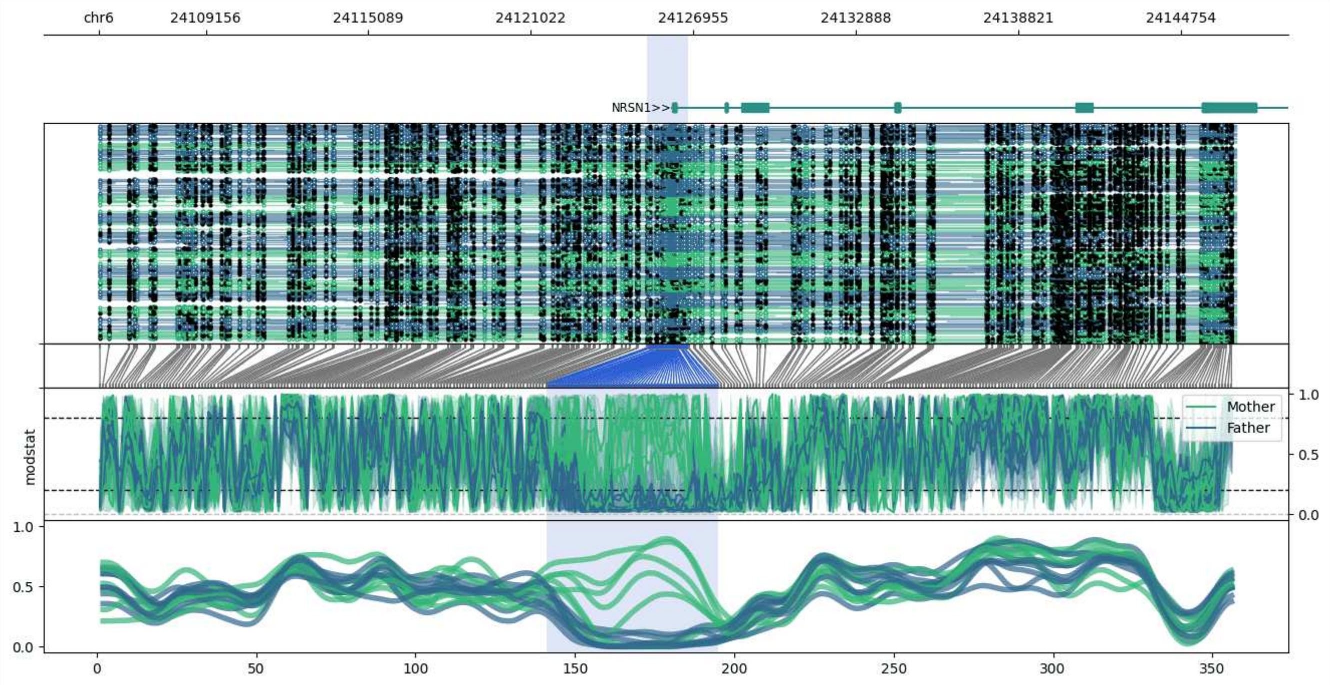


**chr6:24105248-24146782:** Region with 1 DMR containing genes: NRSN1.

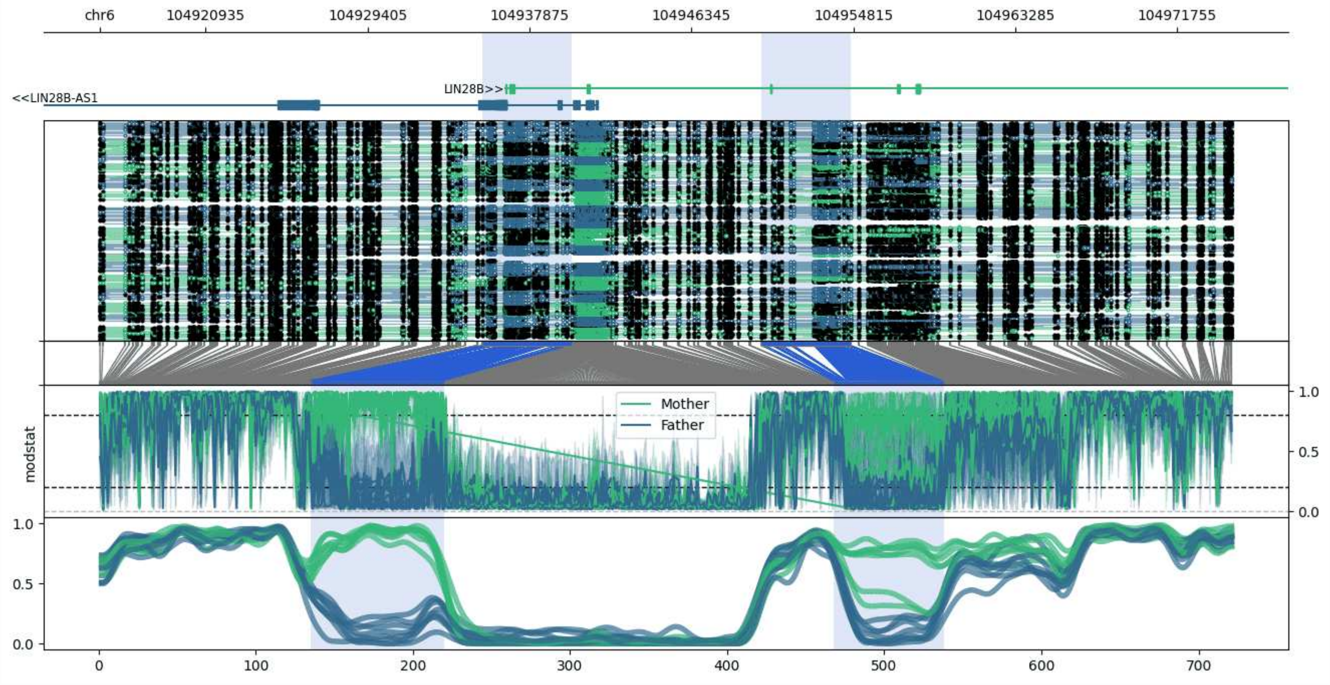


**chr6:104915394-104974684:** Region with 2 DMRs containing genes: LIN28B, LIN28B-AS1.

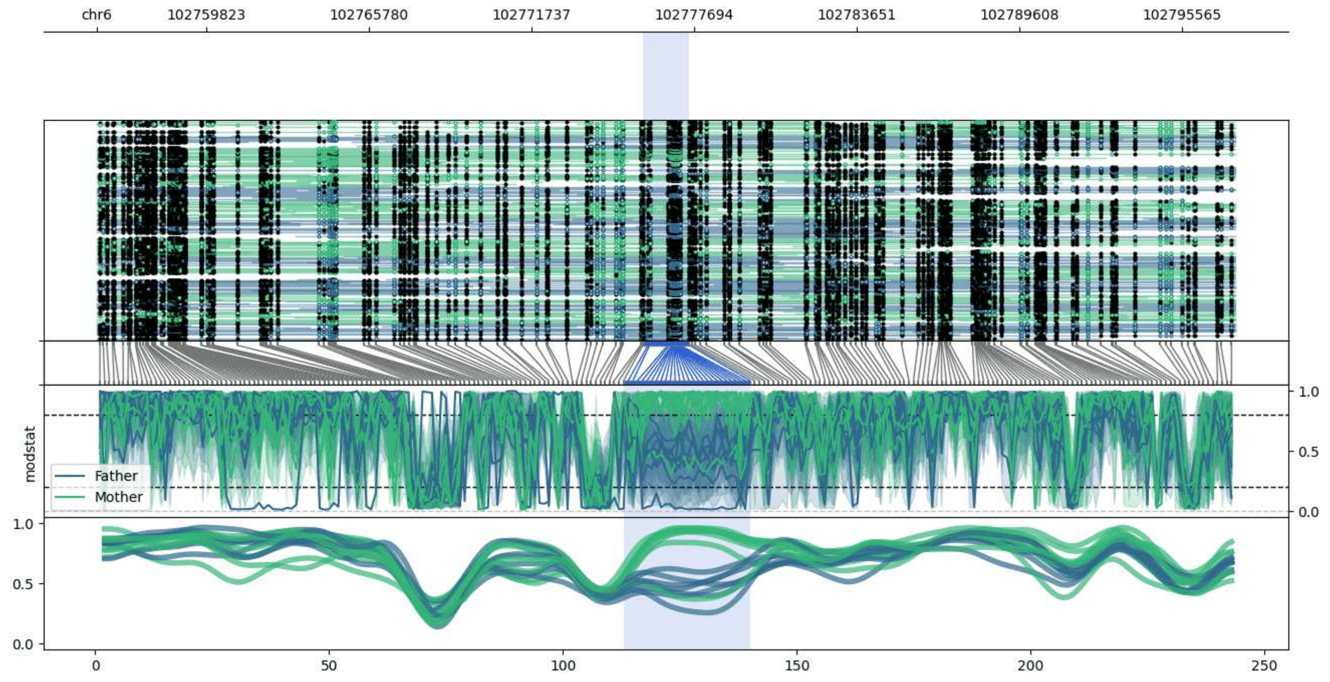


**chr6:102755827-102797529:** Region with 1 DMR containing no genes.

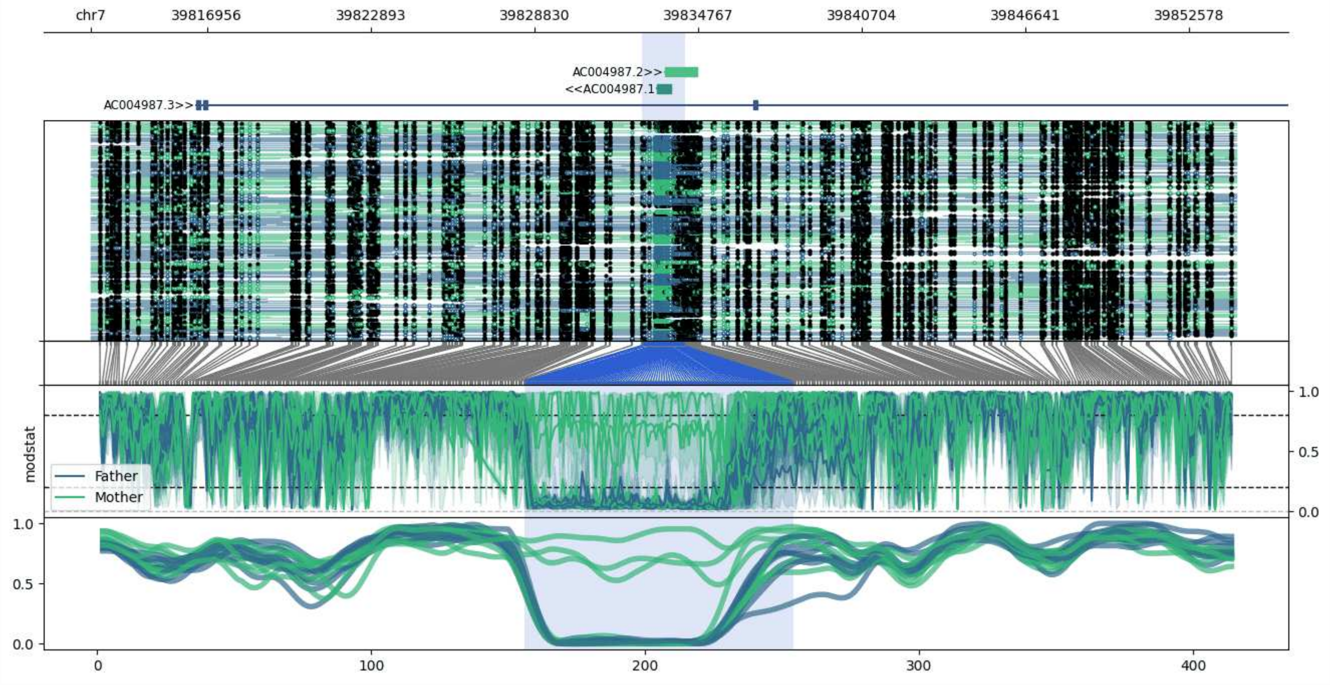


**chr7:39812730-39854290:** Region with 1 DMR containing genes: AC004987.1, AC004987.2, AC004987.3, RWDD4P2.

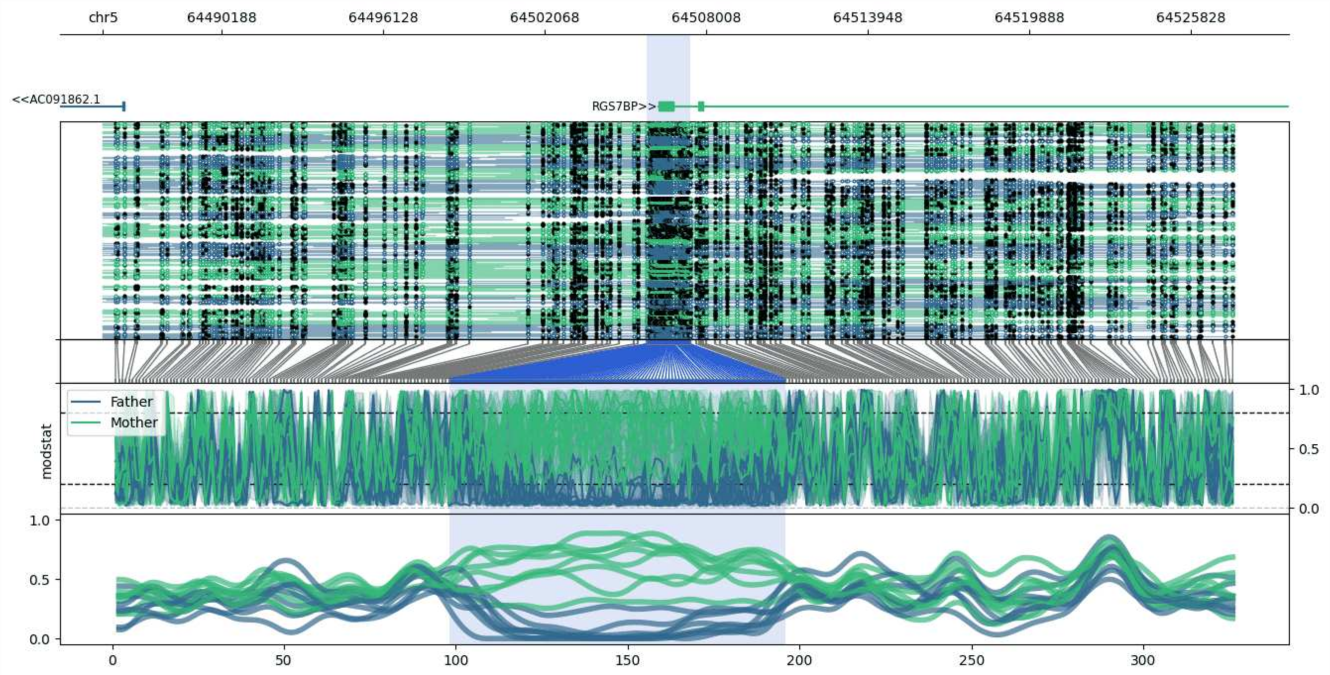


**chr5:64485832-64527416:** Region with 1 DMR containing genes: AC091862.1, RGS7BP.

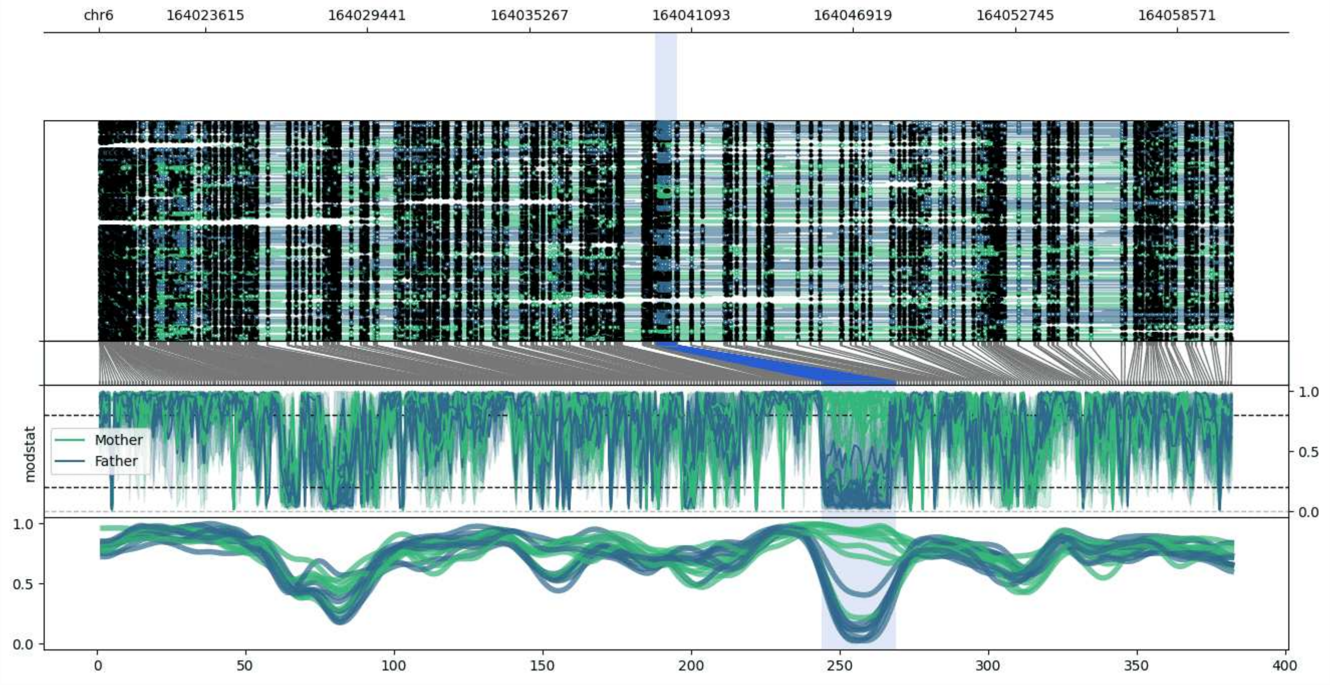


**chr6:164019789-164060575:** Region with 1 DMR containing genes: AL078602.1.

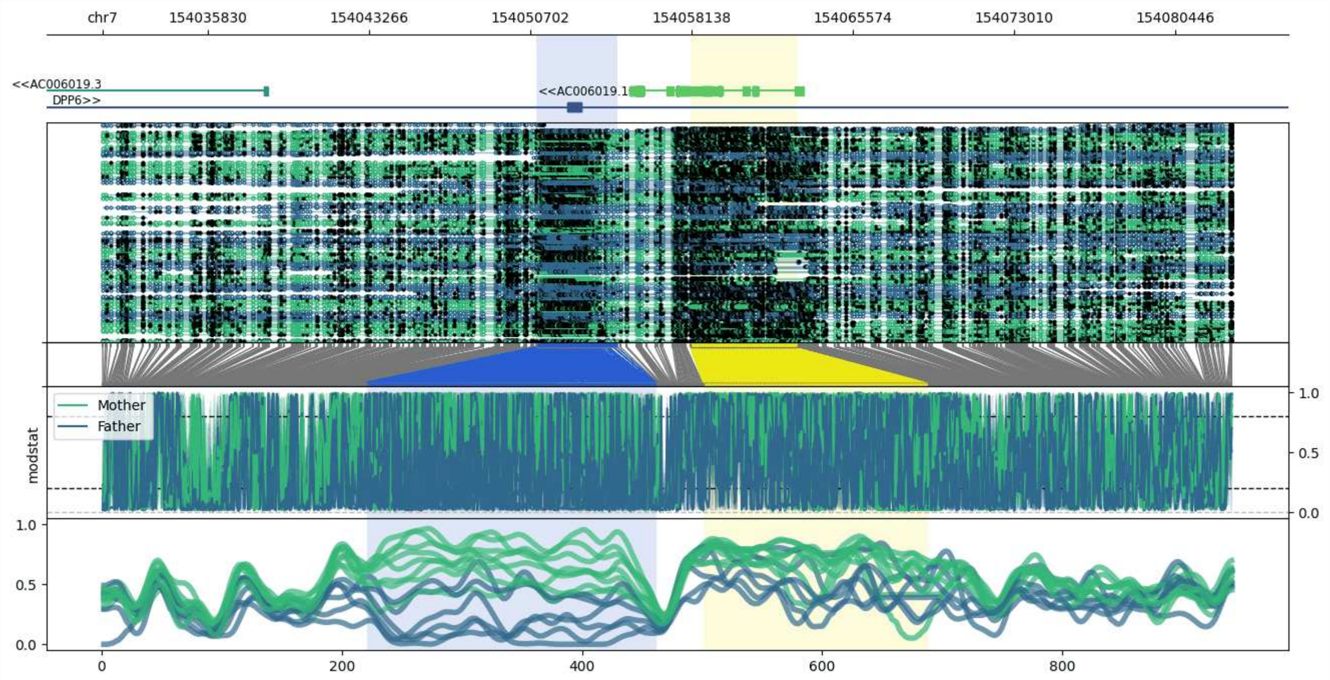


**chr7:154030996-154083054:** Region with 2 DMRs containing genes: AC006019.1, AC006019.2, AC006019.3, DPP6.

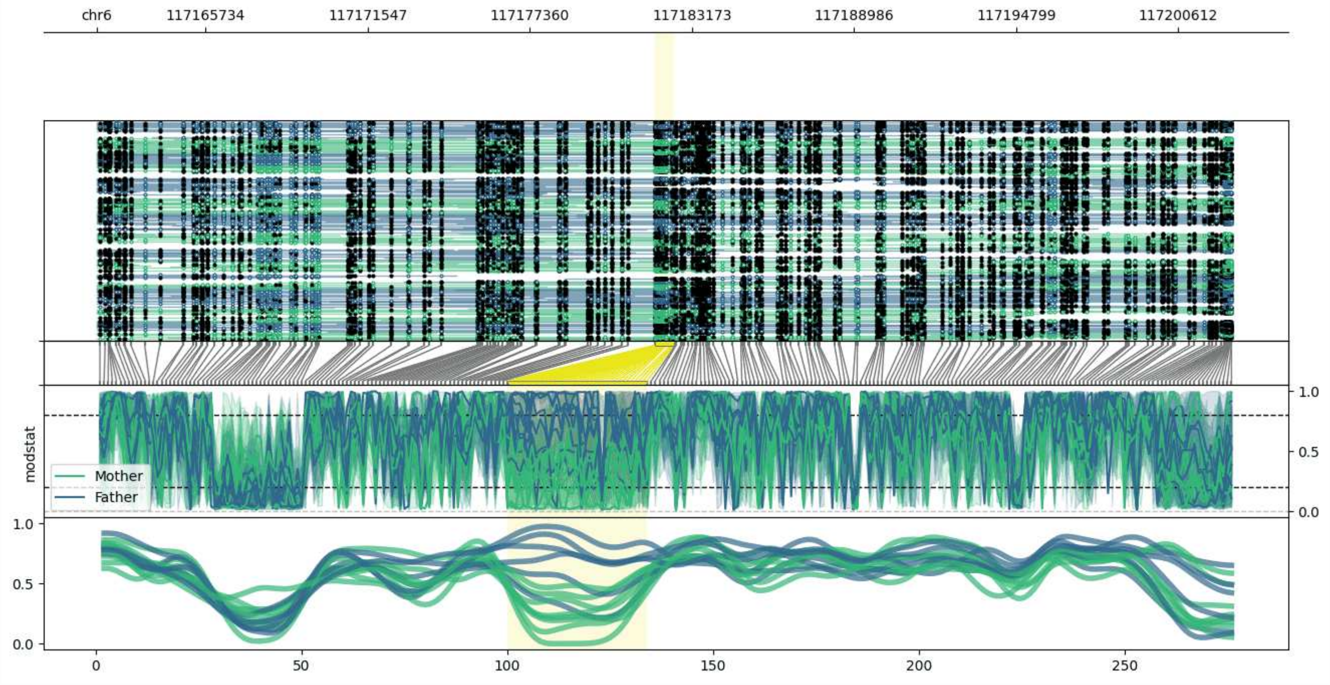


**chr6:117161835-117202529:** Region with 1 DMR containing no genes.

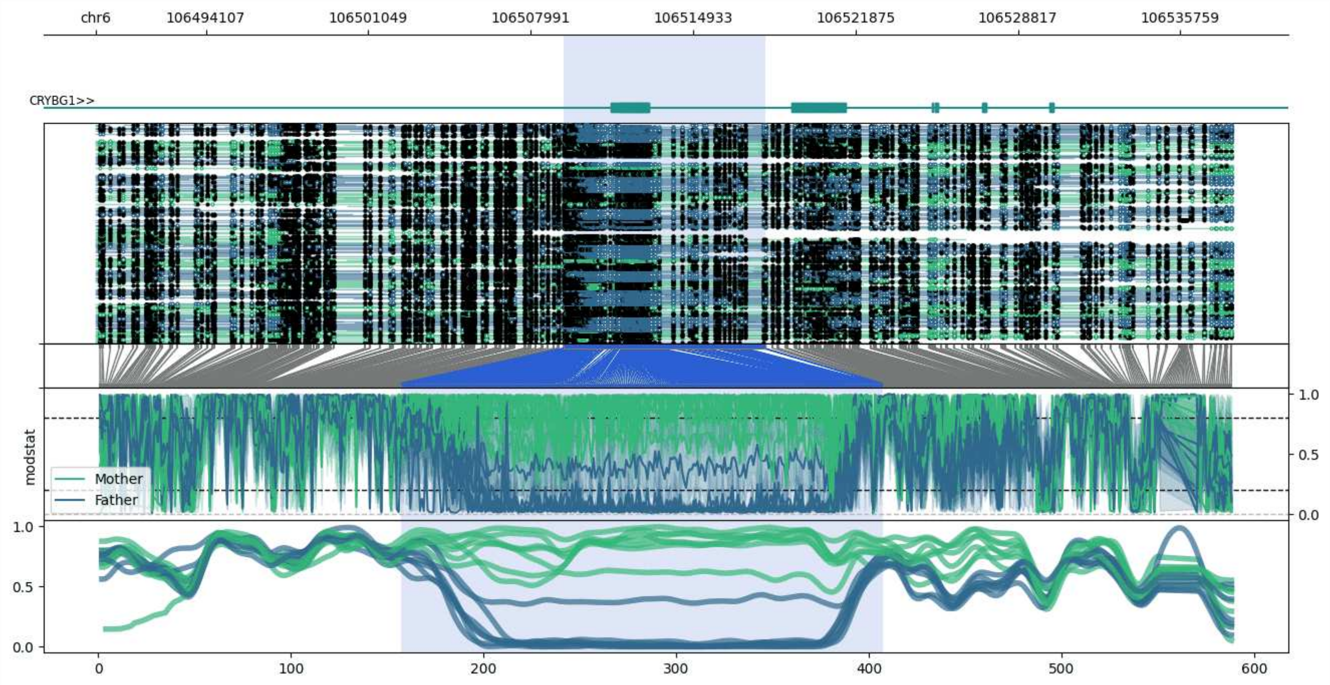


**chr6:106489407-106538002:** Region with 1 DMR containing genes: CRYBG1.

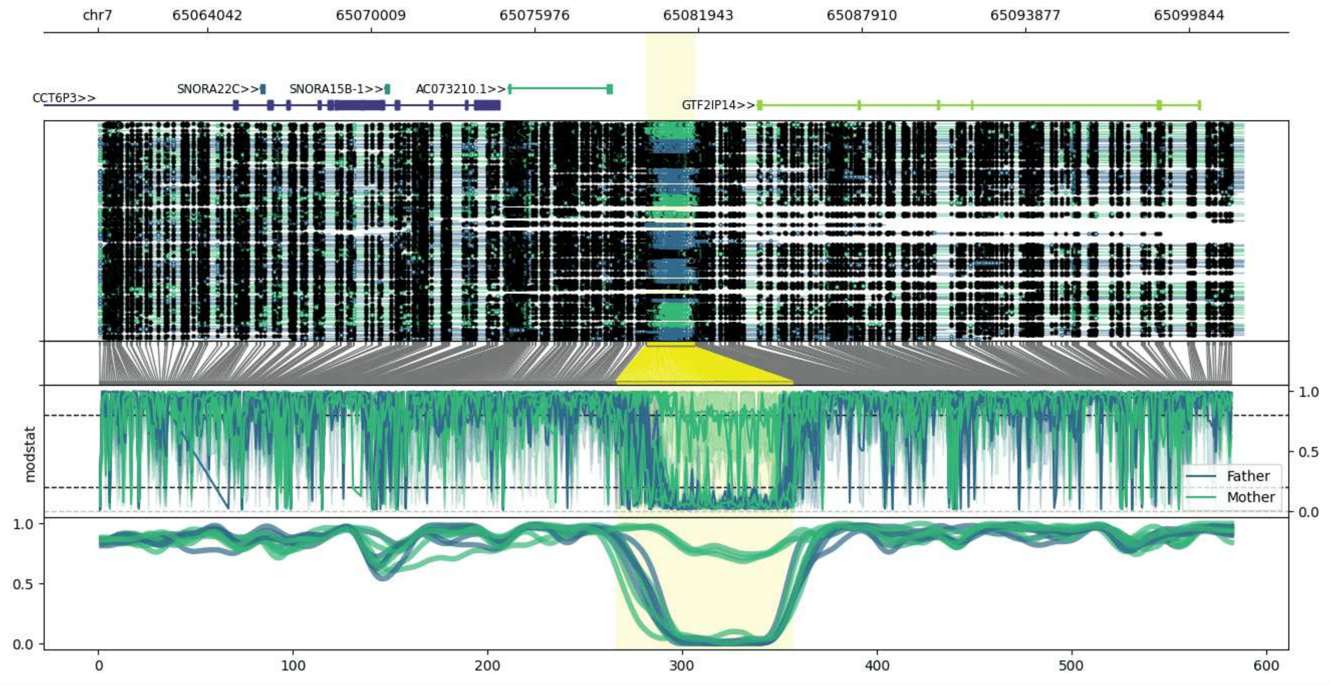


**chr7:65060048-65101821:** Region with 1 DMR containing genes: AC073210.1, CCT6P3, GTF2IP14, SNORA15B-1, SNORA22C.

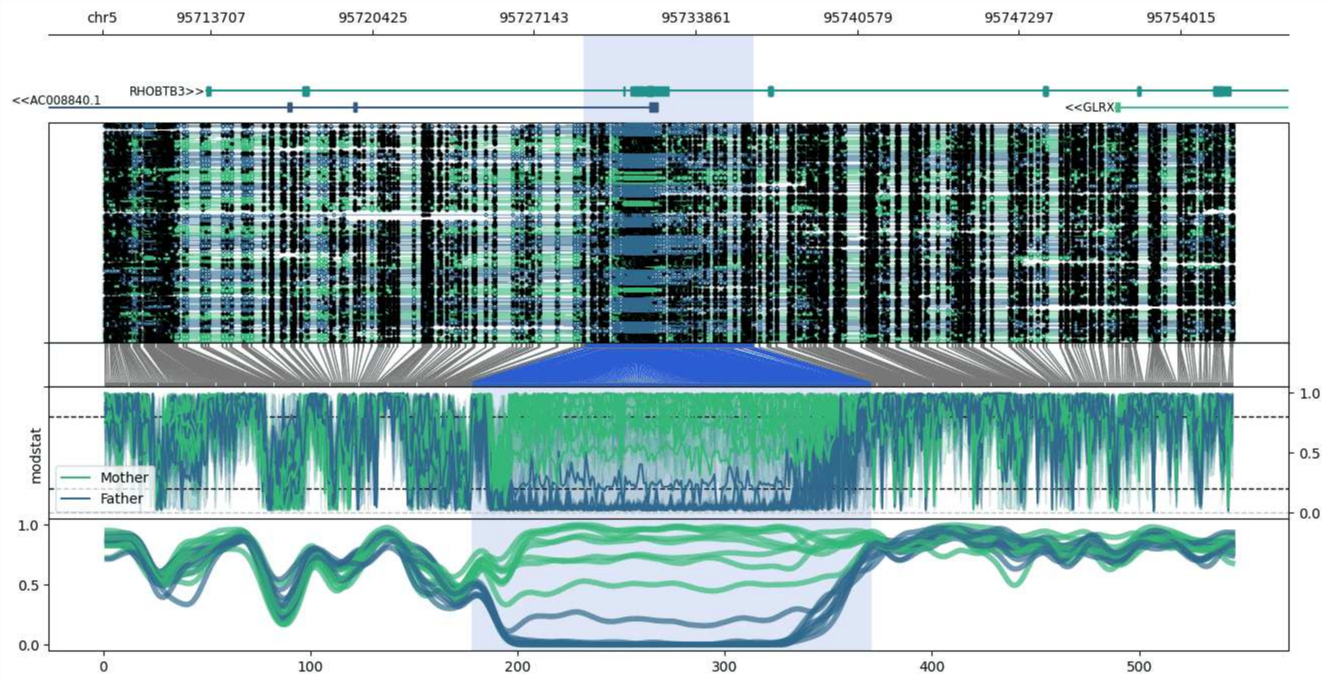


**chr5:95709222-95756248:** Region with 1 DMR containing genes: AC008840.1, GLRX, RHOBTB3.

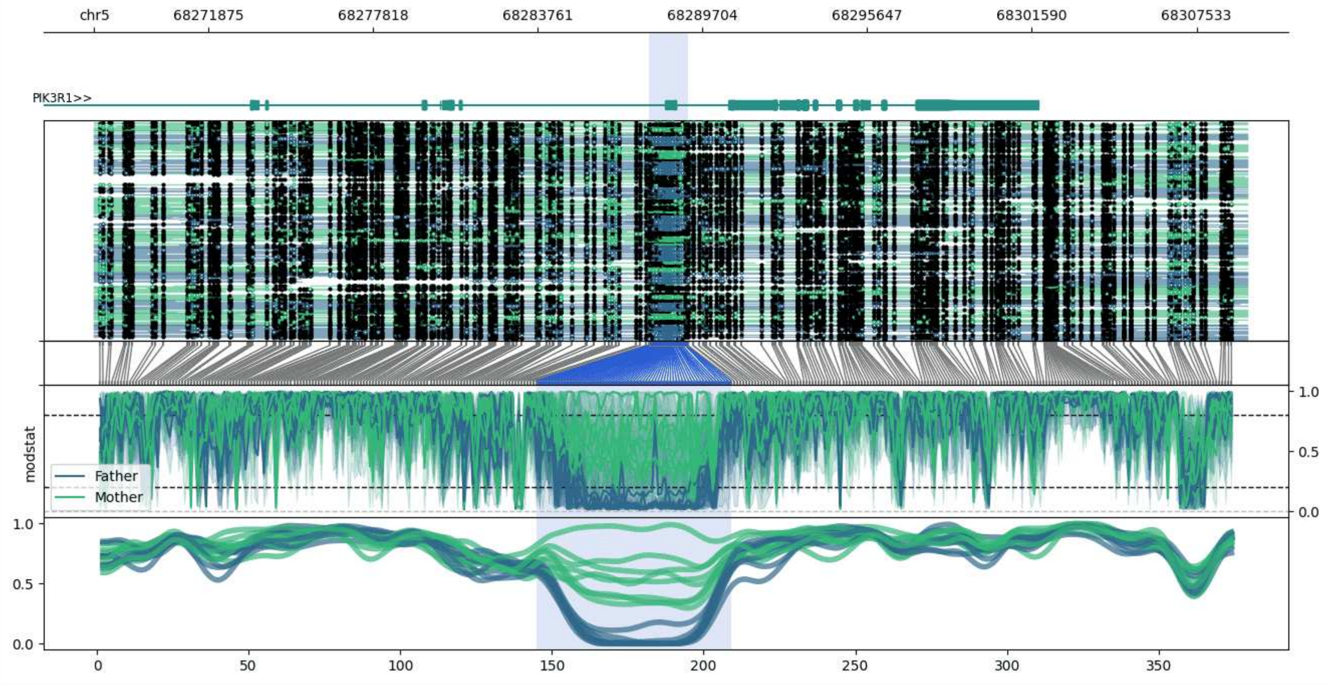


**chr5:68267769-68309374:** Region with 1 DMR containing genes: PIK3R1.

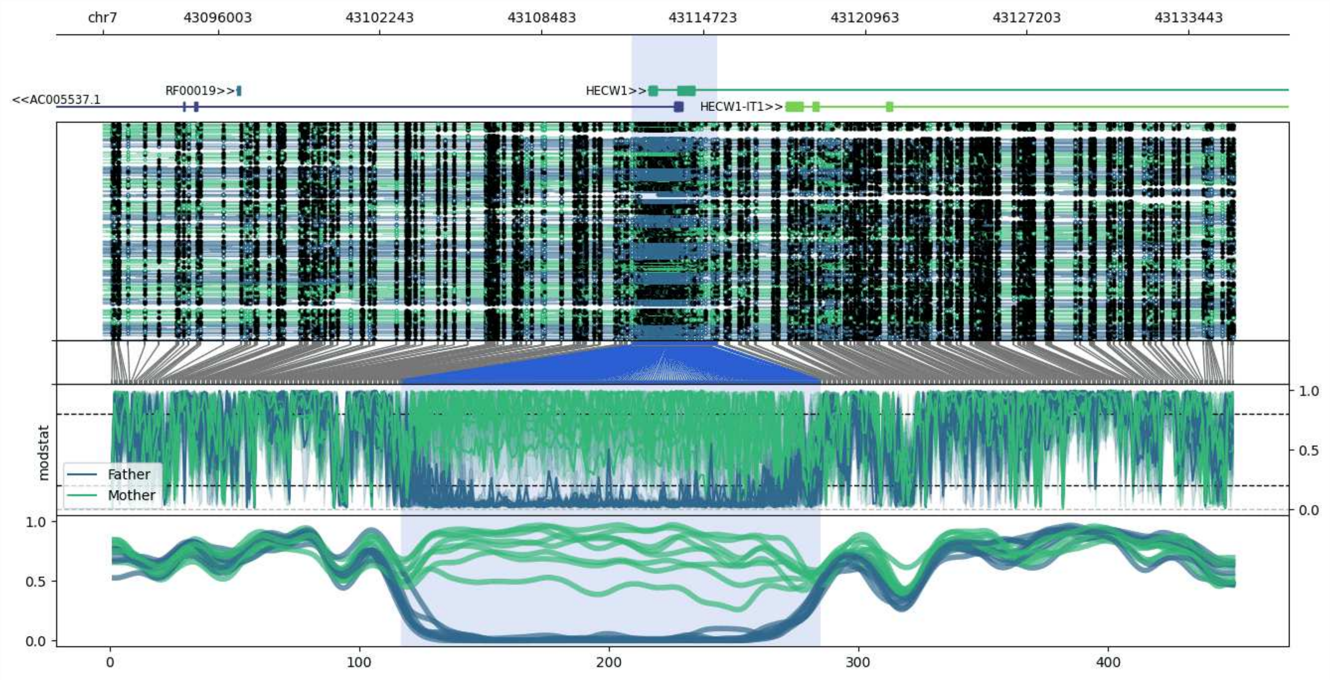


**chr7:43091569-43135250:** Region with 1 DMR containing genes: AC005537.1, HECW1, HECW1-IT1, RF00019.

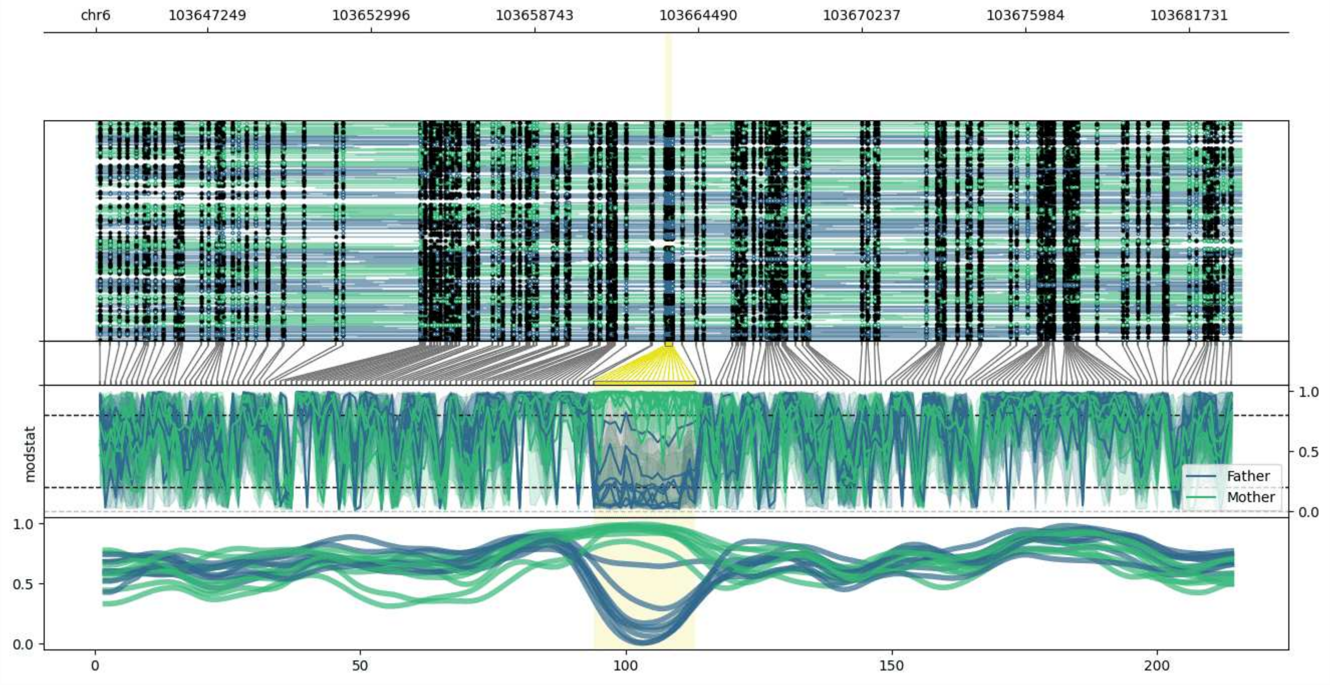


**chr6:103643345-103683574:** Region with 1 DMR containing no genes.

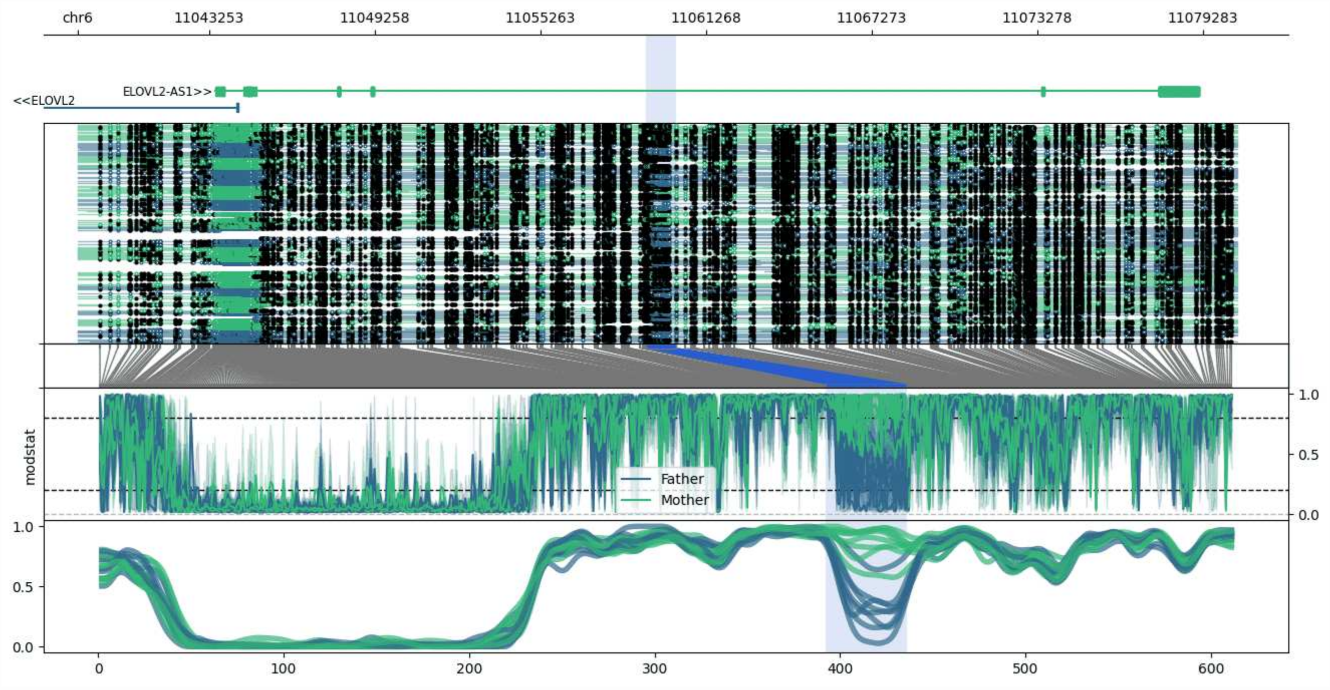


**chr6:11038487-11080523:** Region with 1 DMR containing genes: ELOVL2, ELOVL2-AS1.

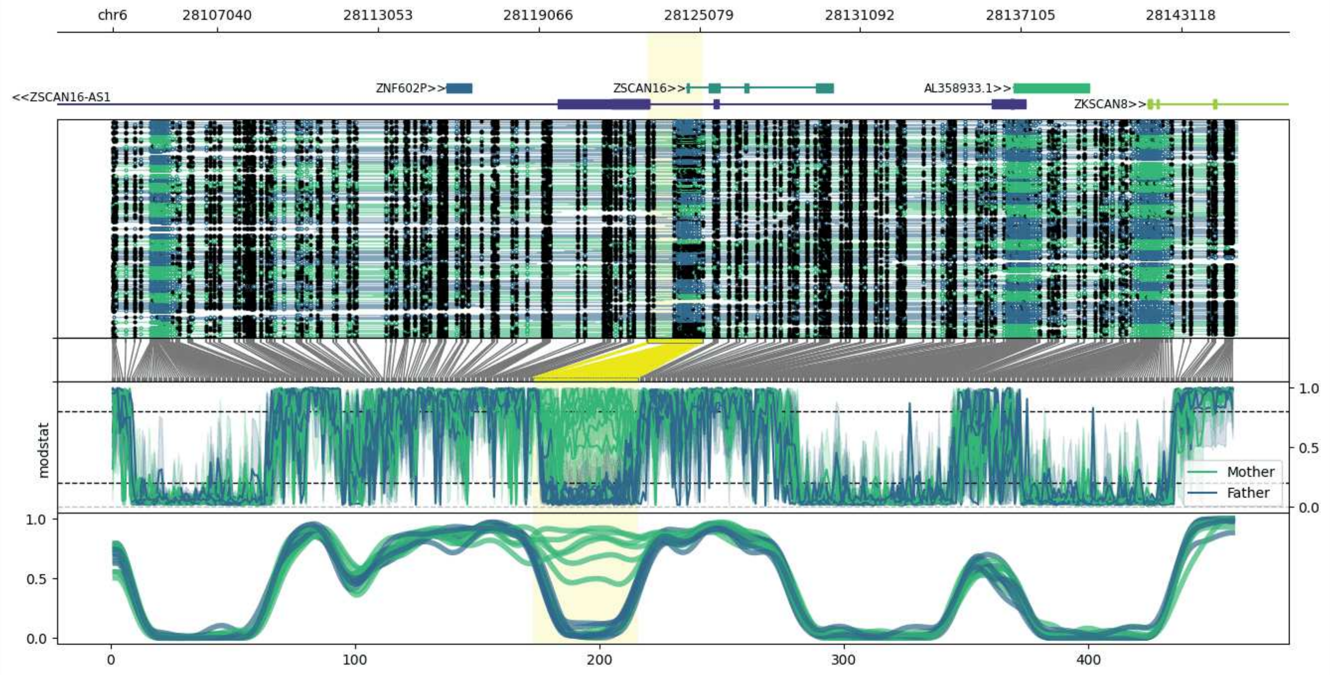


**chr6:28103113-28145205:** Region with 1 DMR containing genes: AL358933.1, ZKSCAN8, ZNF602P, ZSCAN12P1, ZSCAN16, ZSCAN16-AS1.

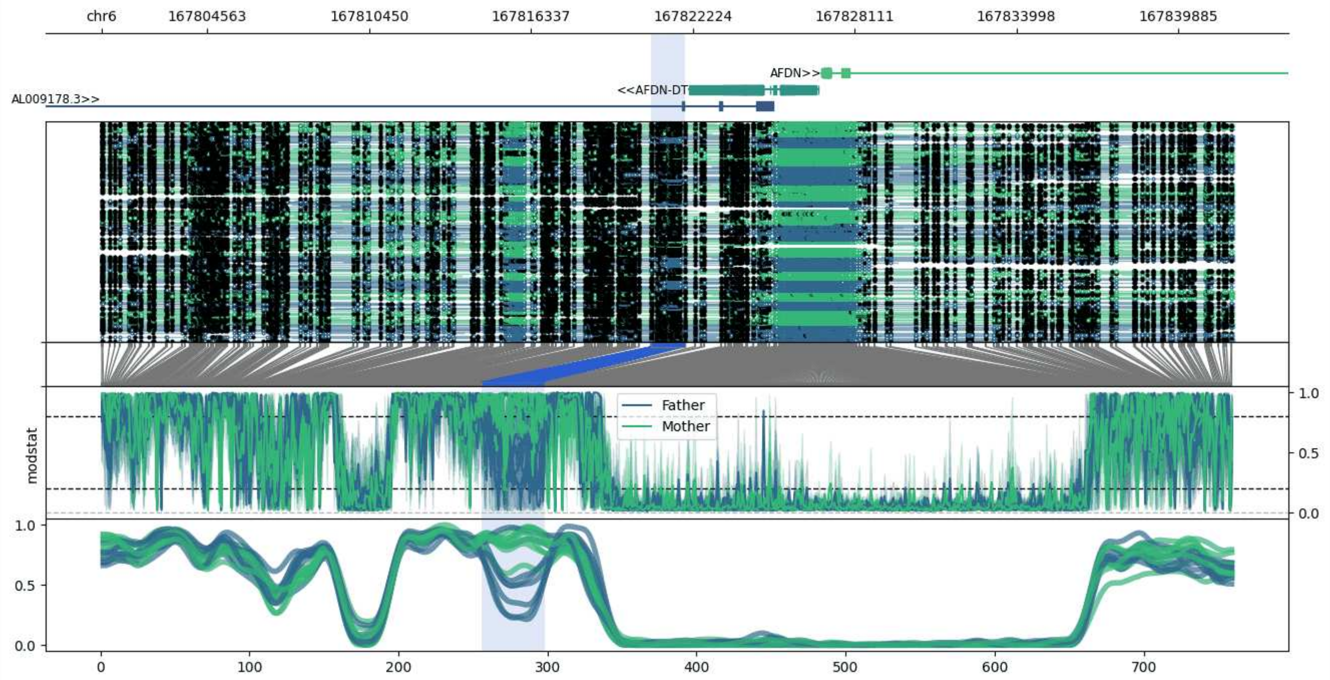


**chr6:167800719-167841931:** Region with 1 DMR containing genes: AFDN, AFDN-DT, AL009178.2, AL009178.3, LINC01558.

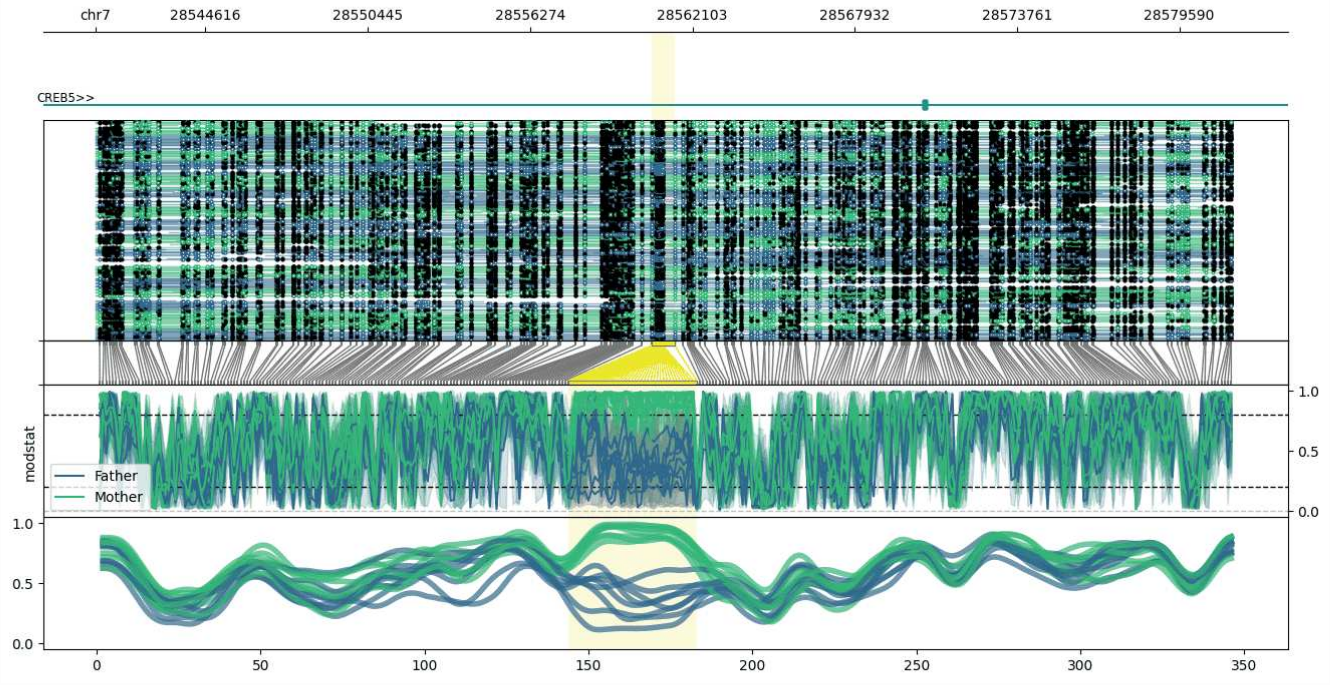


**chr7:28540658-28581461:** Region with 1 DMR containing genes: CREB5.

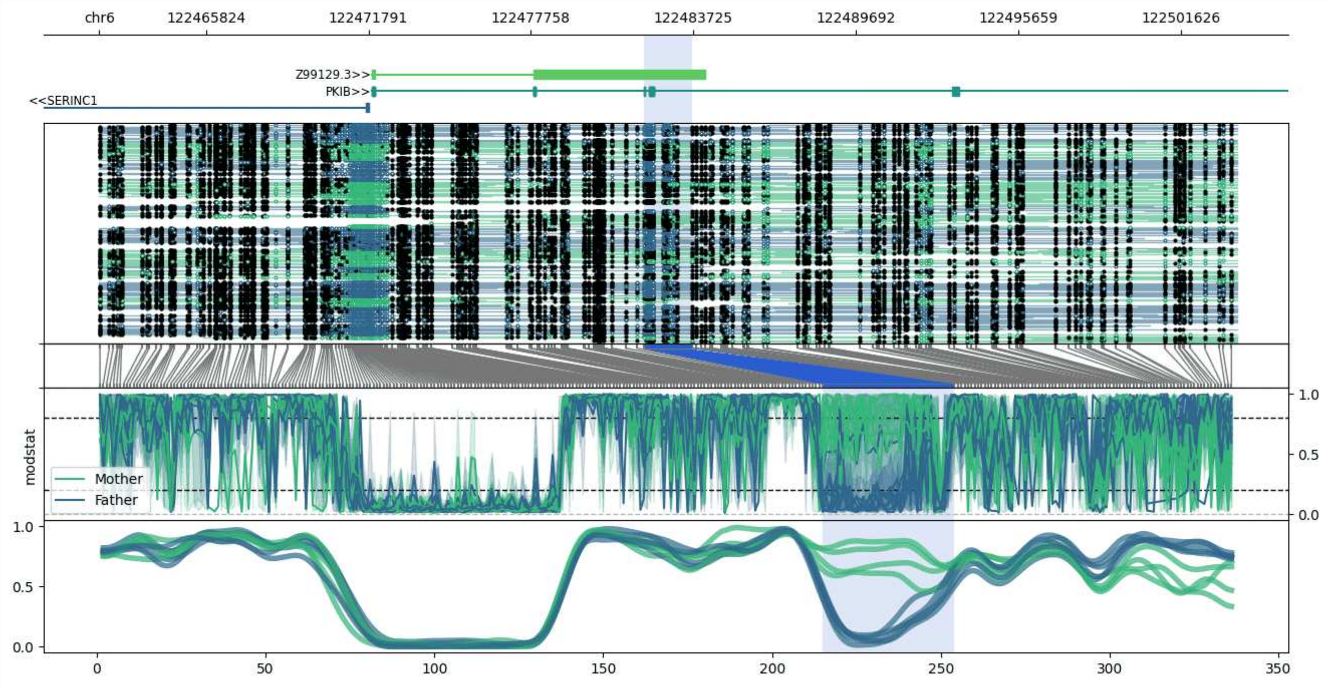


**chr6:122461912-122503685:** Region with 1 DMR containing genes: PKIB, SERINC1, Z99129.1, Z99129.3.

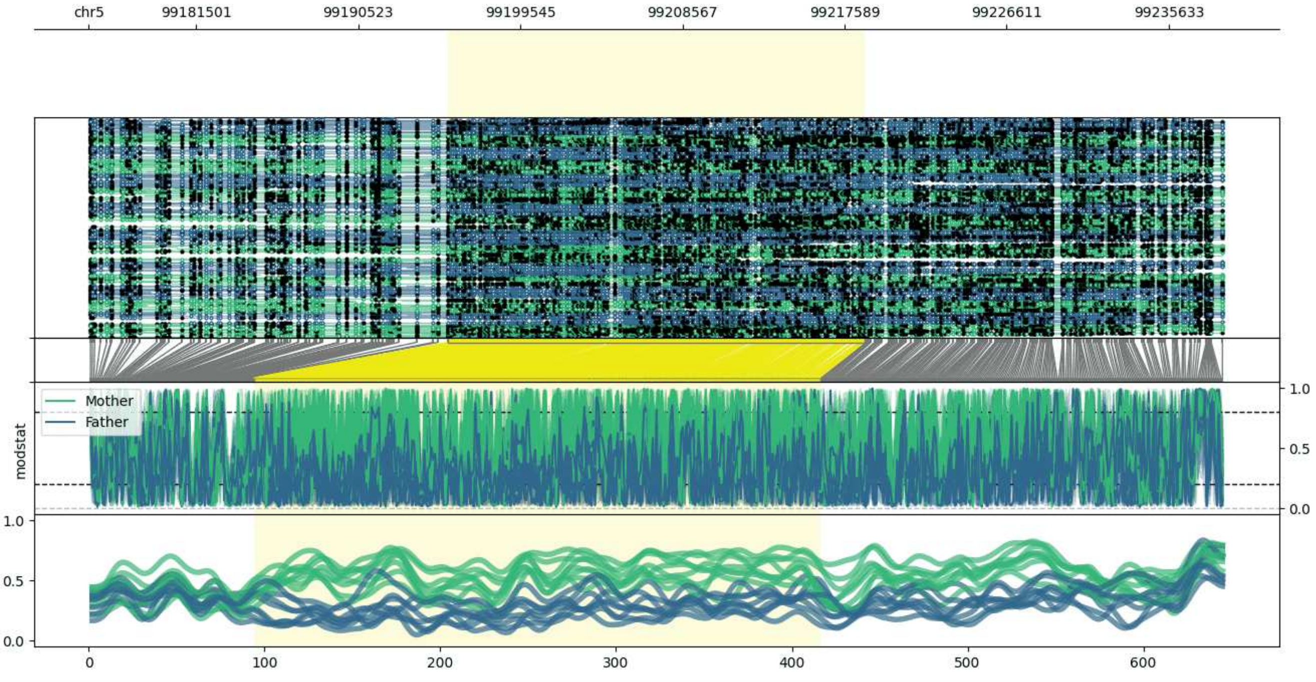


**chr5:99175535-99238692:** Region with 1 DMR containing no genes.

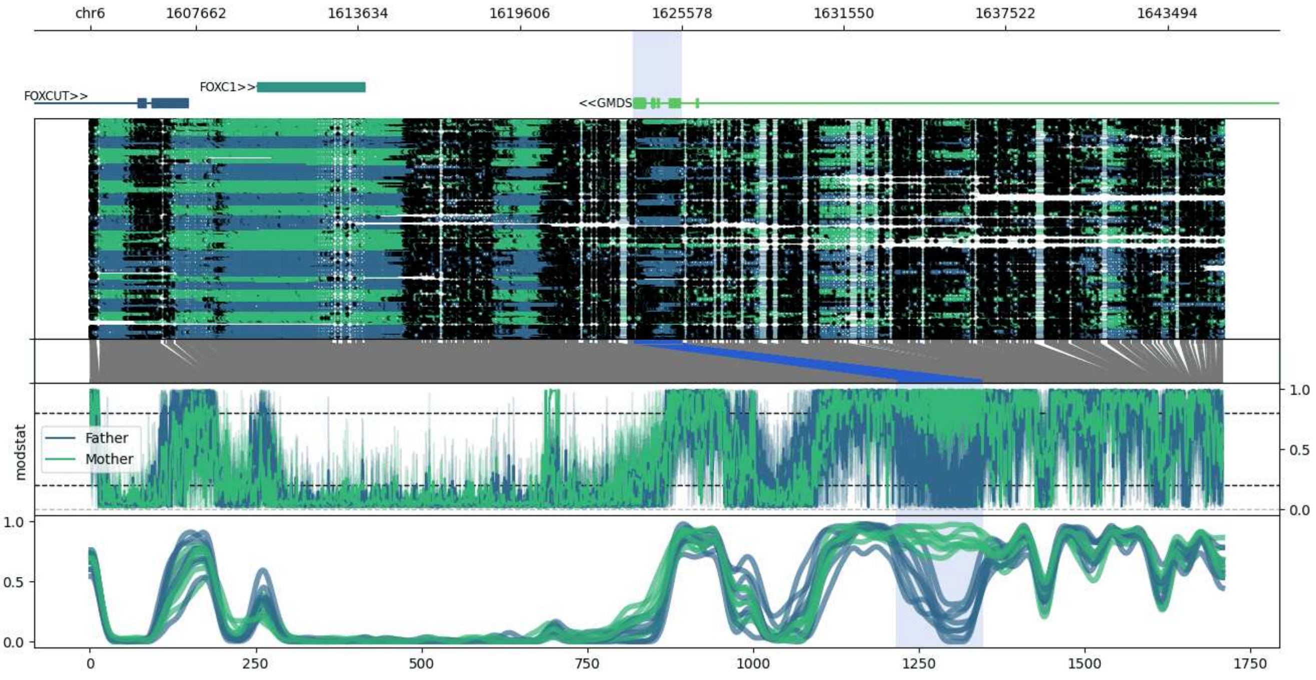


**chr6:1603761-1645568:** Region with 1 DMR containing genes: FOXC1, FOXCUT, GMDS.

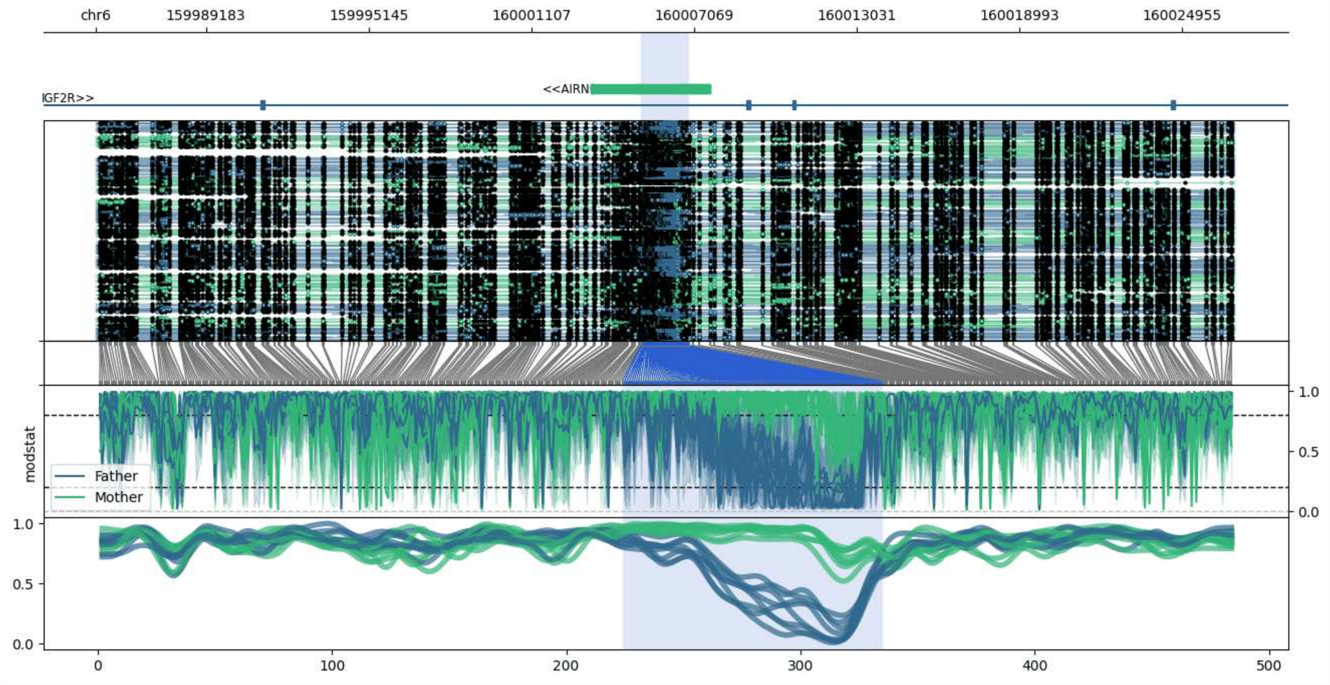


**chr6:159985138-160026878:** Region with 1 DMR containing genes: AIRN, IGF2R.

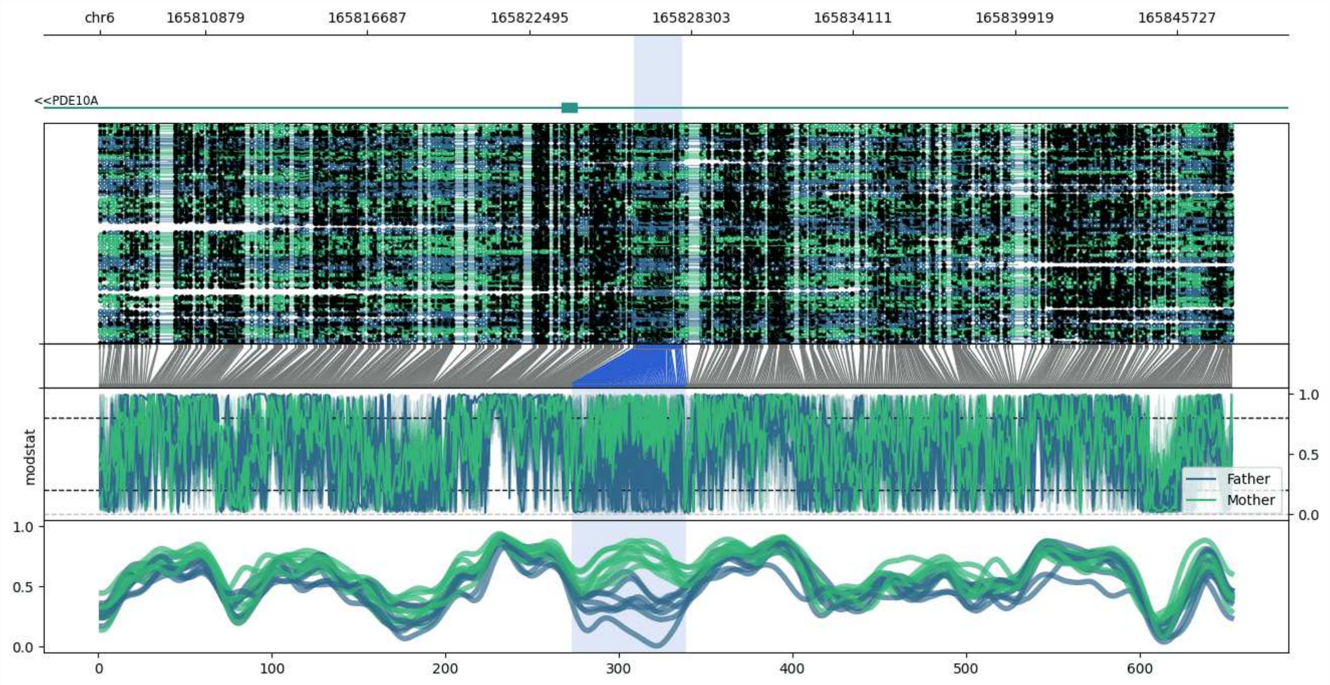


**chr6:165807091-165847750:** Region with 1 DMR containing genes: PDE10A.

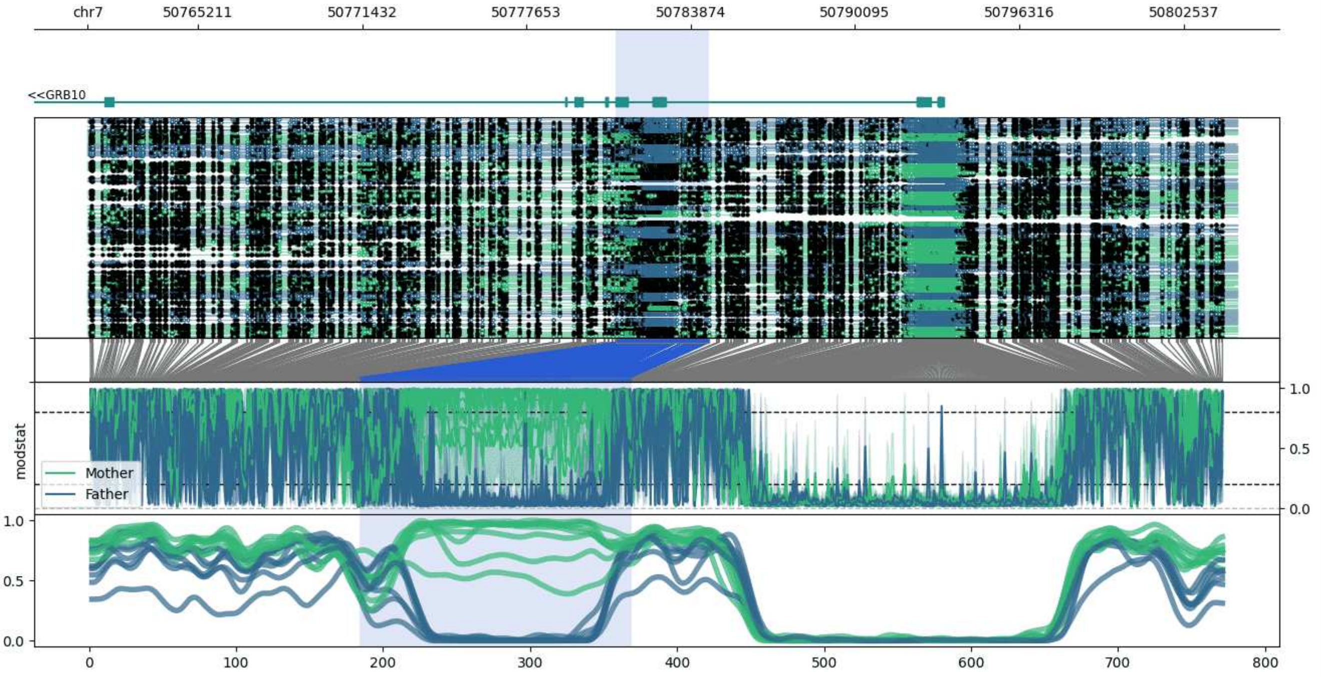


**chr7:50761028-50804580:** Region with 1 DMR containing genes: GRB10.

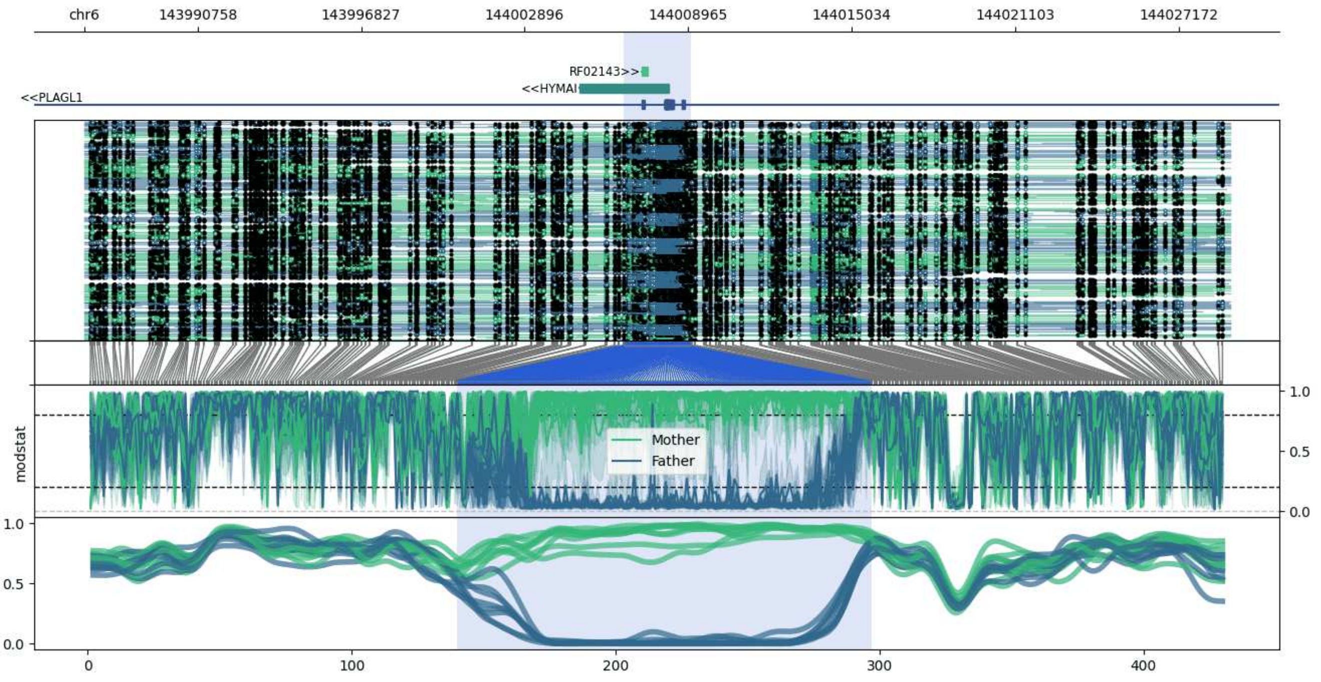


**chr6:143986568-144029053:** Region with 1 DMR containing genes: AL109755.1, HYMAI, PLAGL1, RF02143.

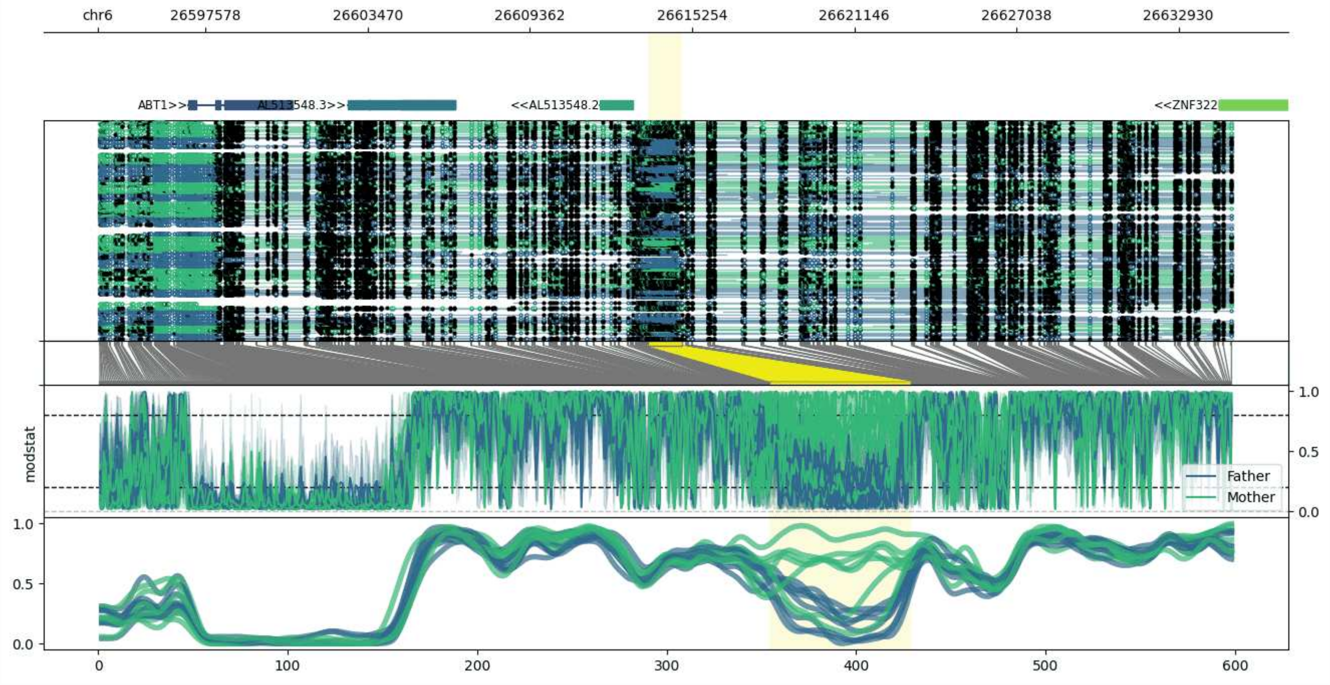


**chr6:26593653-26634899:** Region with 1 DMR containing genes: ABT1, AL513548.2, AL513548.3, ZNF322.

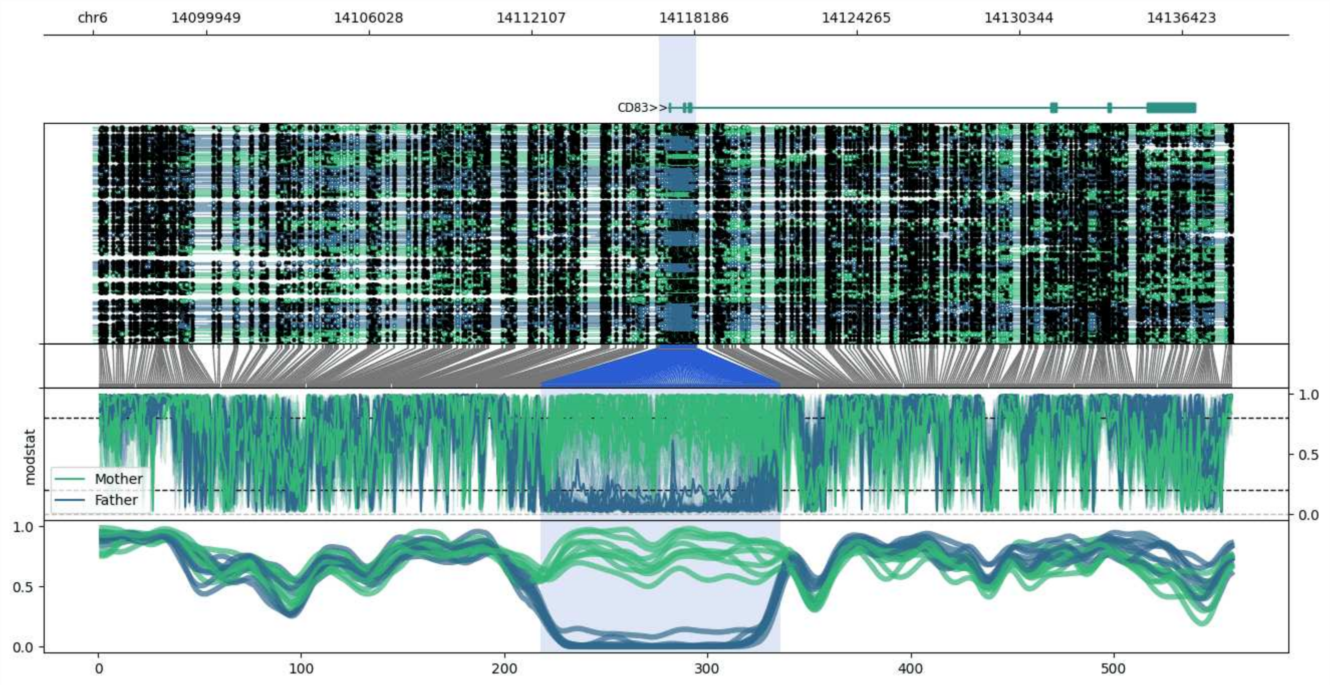


**chr6:14095717-14138272:** Region with 1 DMR containing genes: CD83.

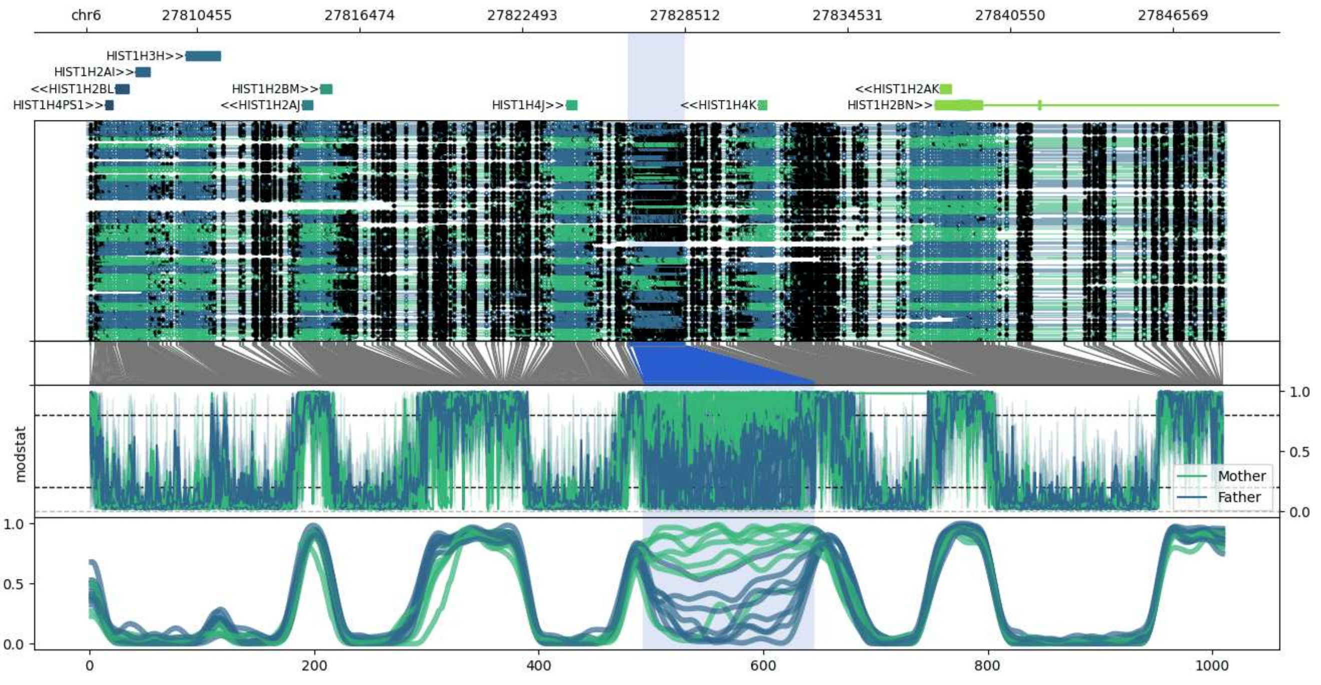


**chr6:27806375-27848511:** Region with 1 DMR containing genes: HIST1H2AI, HIST1H2AJ, HIST1H2AK, HIST1H2BL, HIST1H2BM, HIST1H2BN, HIST1H3H, HIST1H4J, HIST1H4K, HIST1H4PS1.

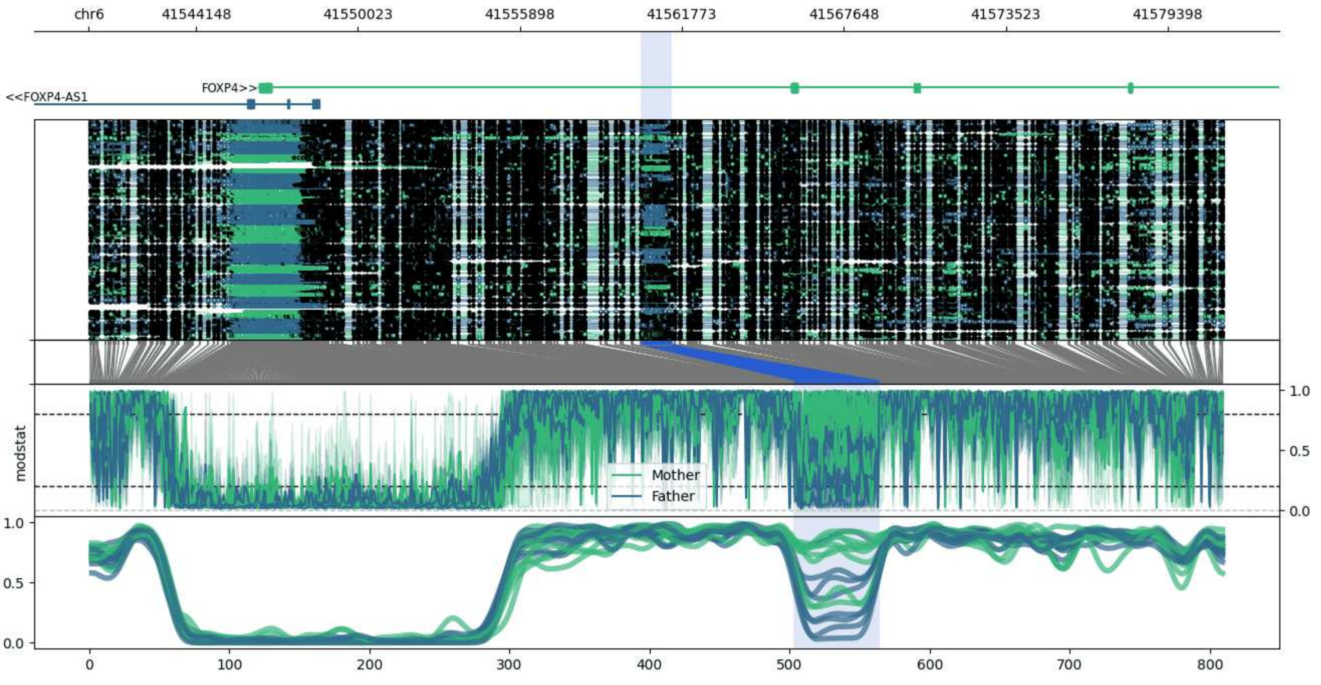


**chr6:41540265-41581395:** Region with 1 DMR containing genes: FOXP4, FOXP4-AS1.

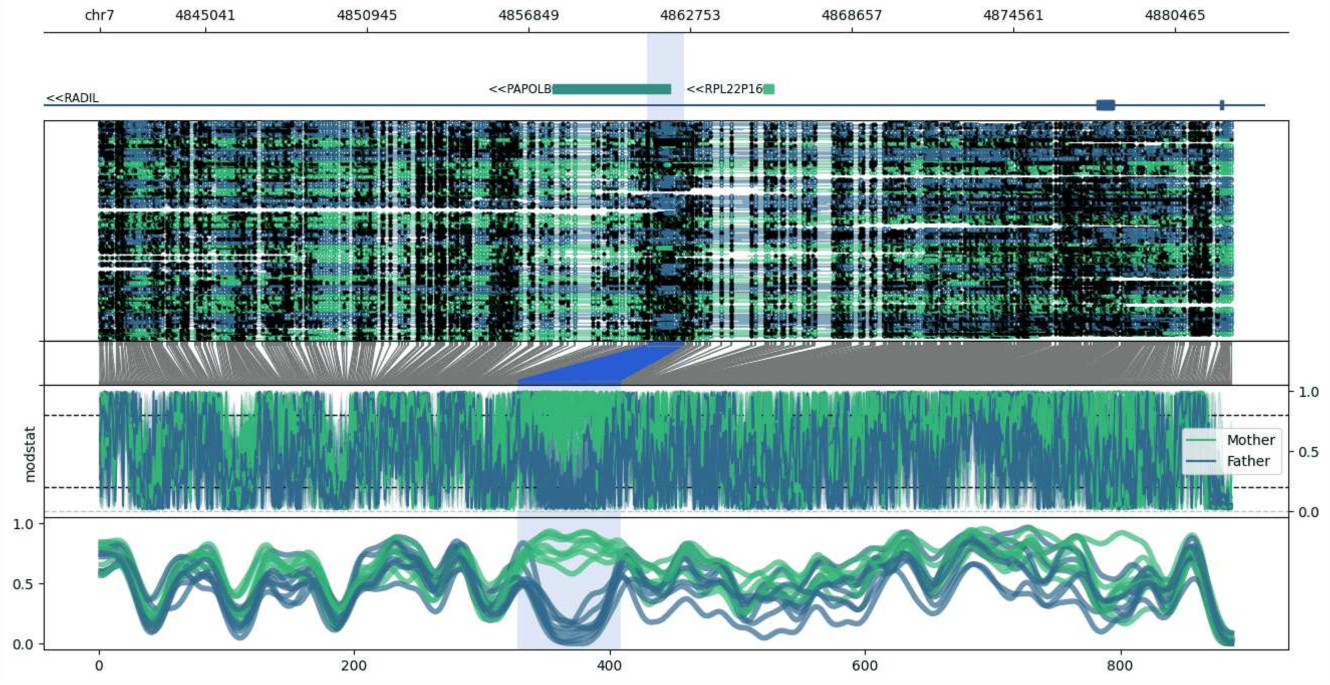


**chr7:4841185-4882514:** Region with 1 DMR containing genes: PAPOLB, RADIL, RPL22P16.

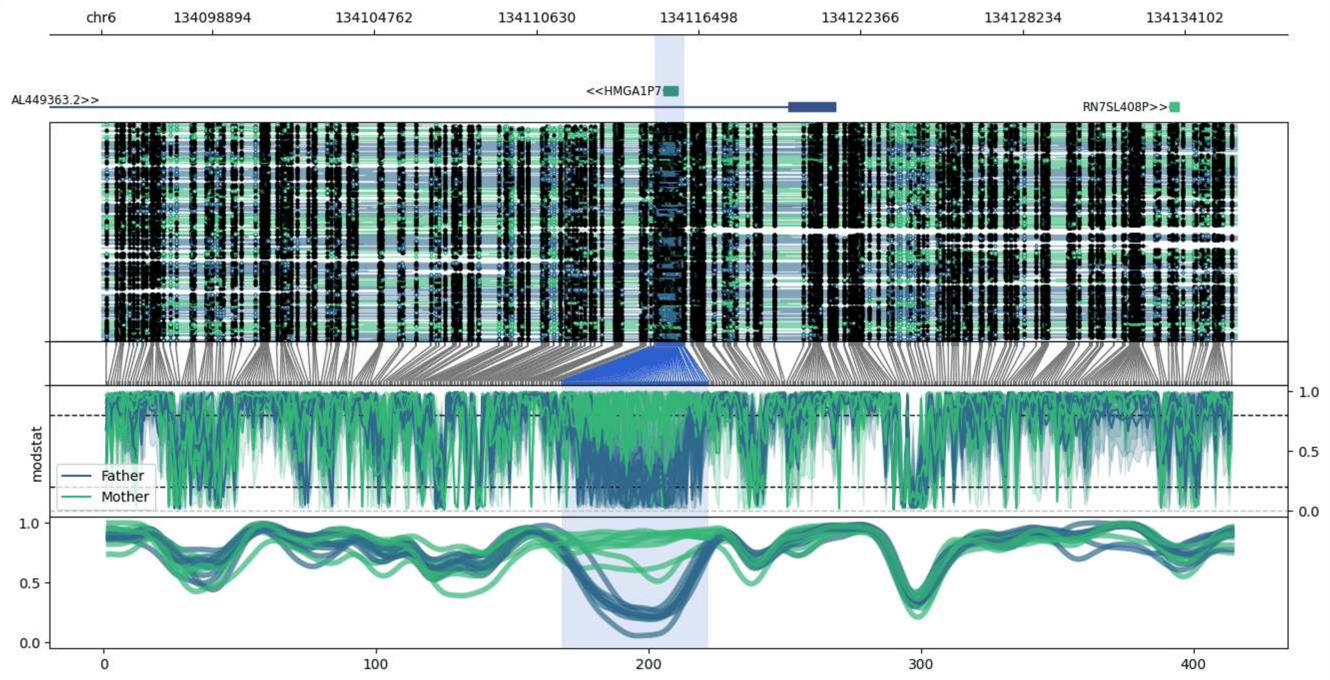


**chr6:134094897-134135978:** Region with 1 DMR containing genes: AL449363.2, HMGA1P7, RN7SL408P.

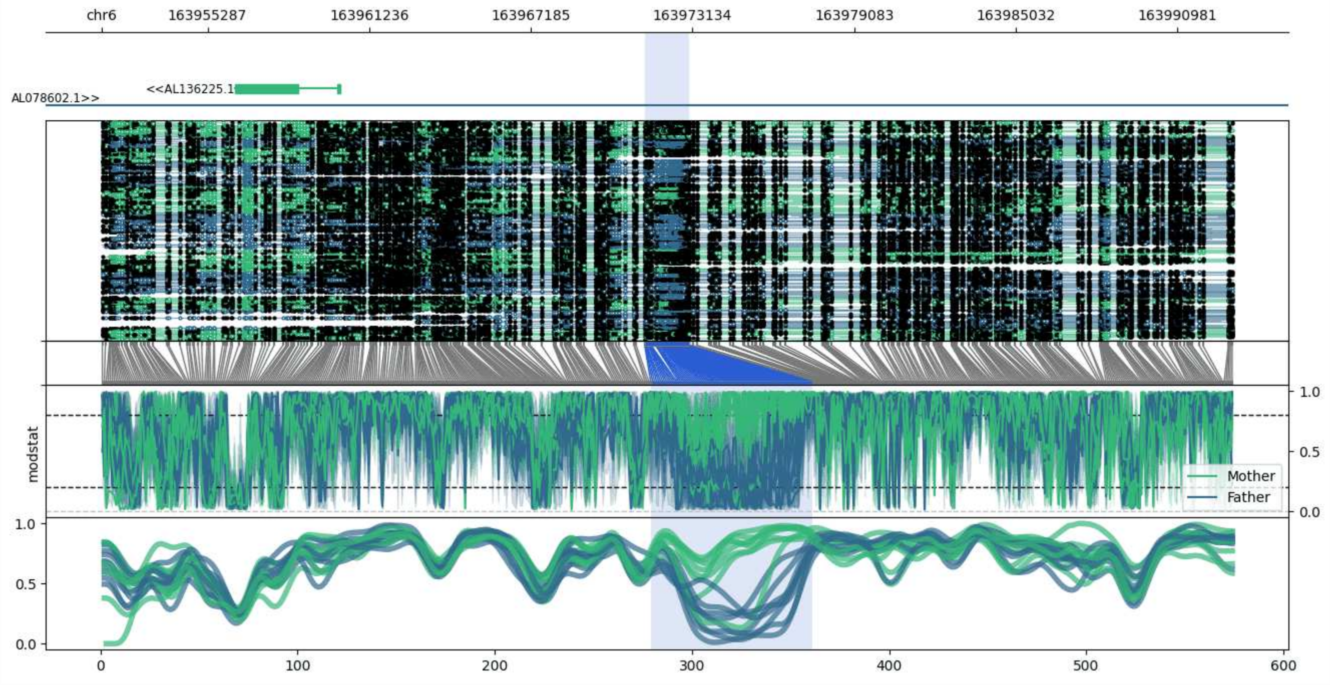


**chr6:163951378-163993025:** Region with 1 DMR containing genes: AL078602.1, AL136225.1.

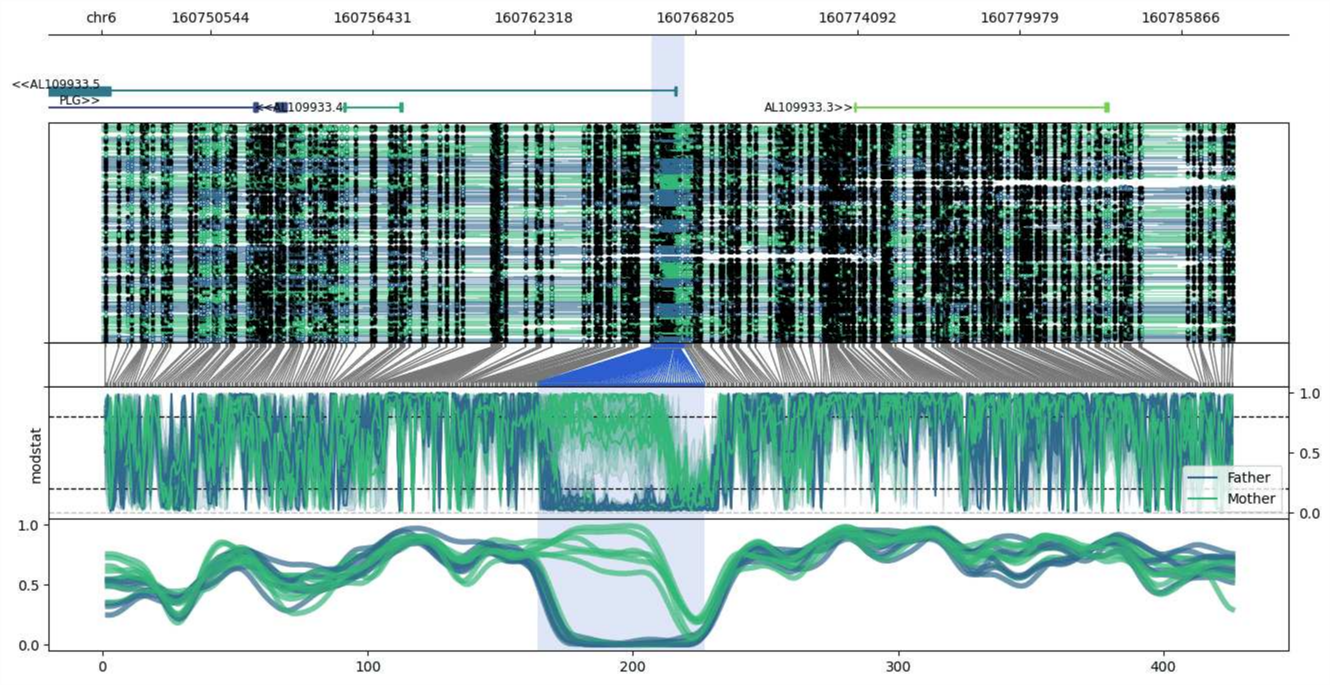


**chr6:160746579-160787790:** Region with 1 DMR containing genes: AL109933.3, AL109933.4, AL109933.5, PLG.

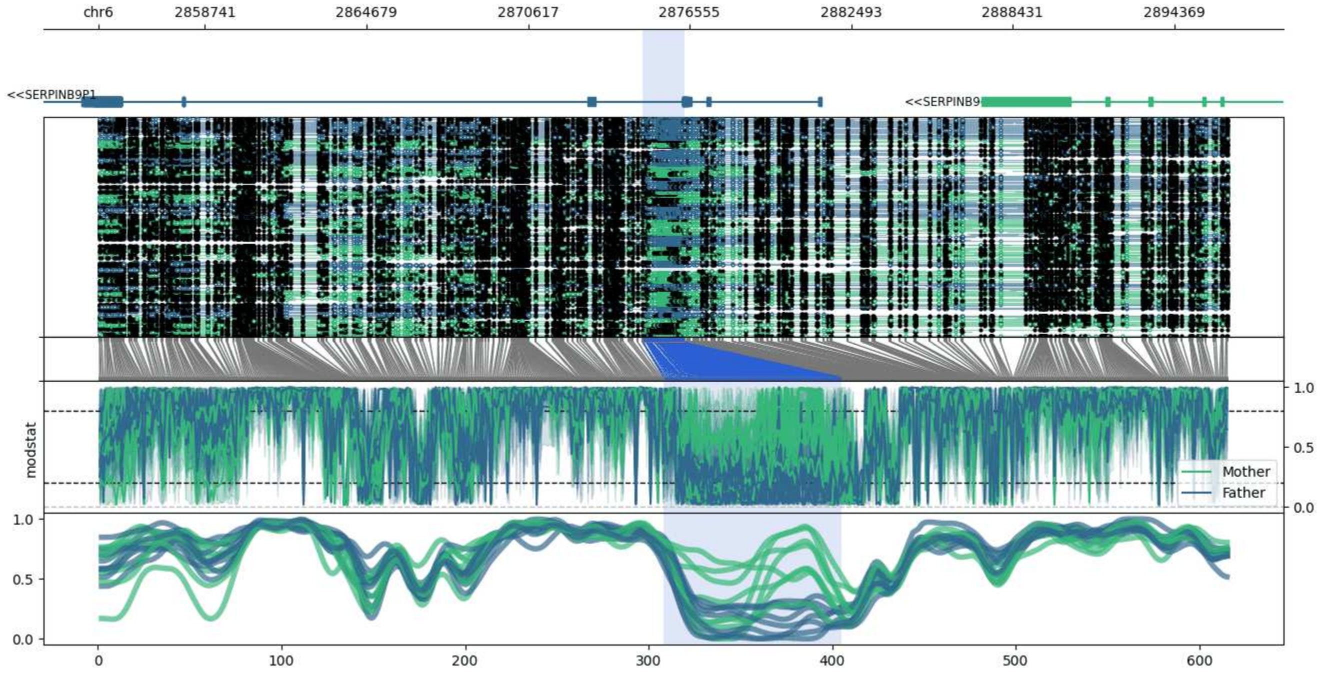


**chr6:2854813-2896380:** Region with 1 DMR containing genes: MIR4645, SERPINB9, SERPINB9P1.

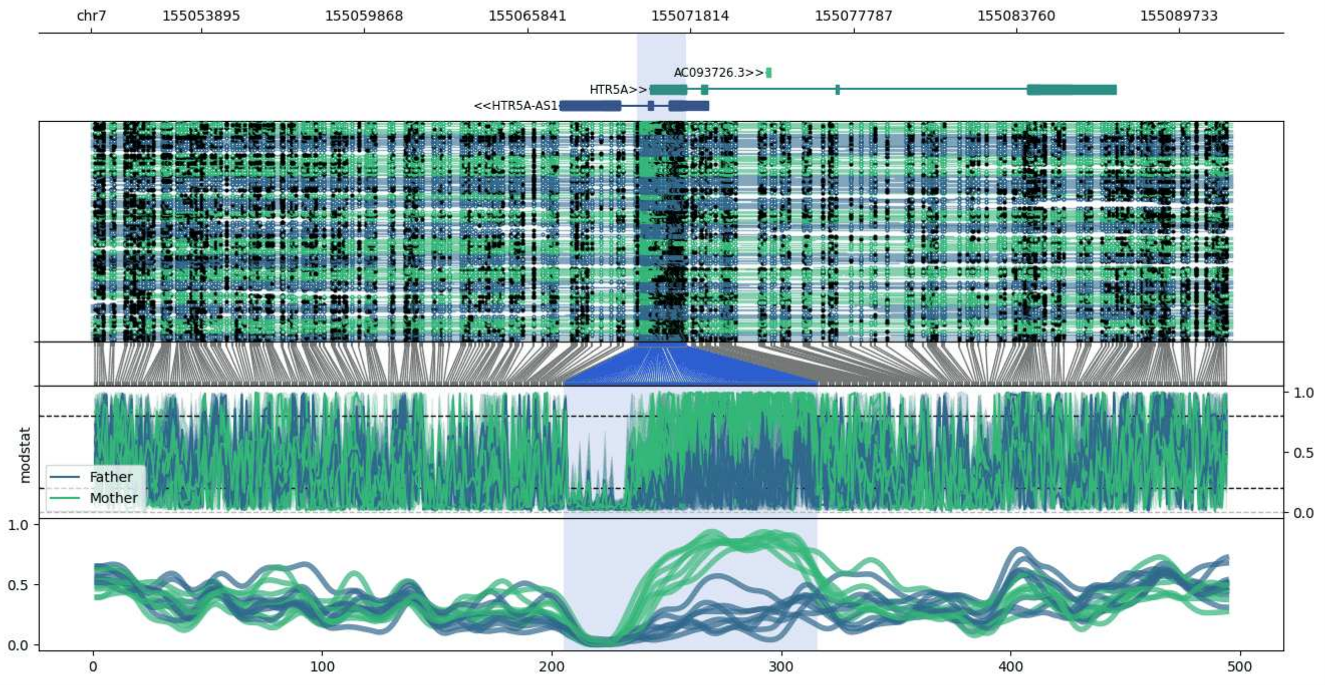


**chr7:155049859-155091672:** Region with 1 DMR containing genes: AC093726.3, HTR5A, HTR5A-AS1.

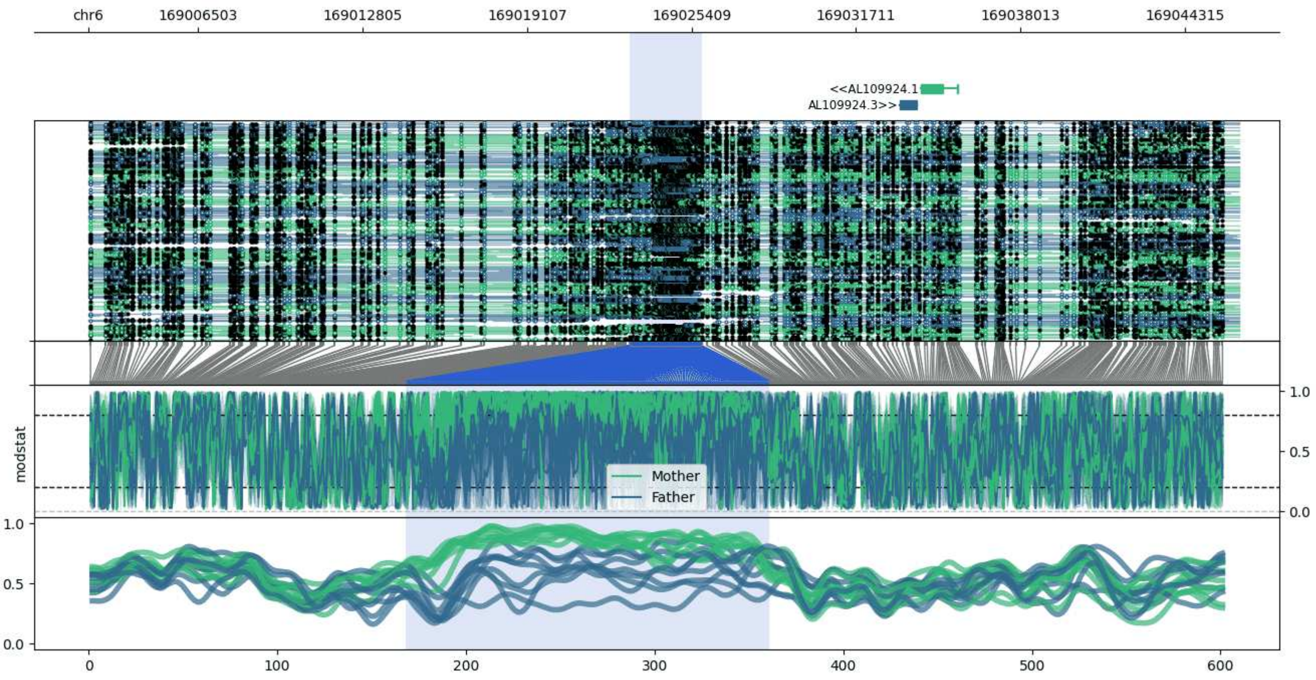


**chr6:169002272-169046391:** Region with 1 DMR containing genes: AL109924.1, AL109924.3, AL109924.4.

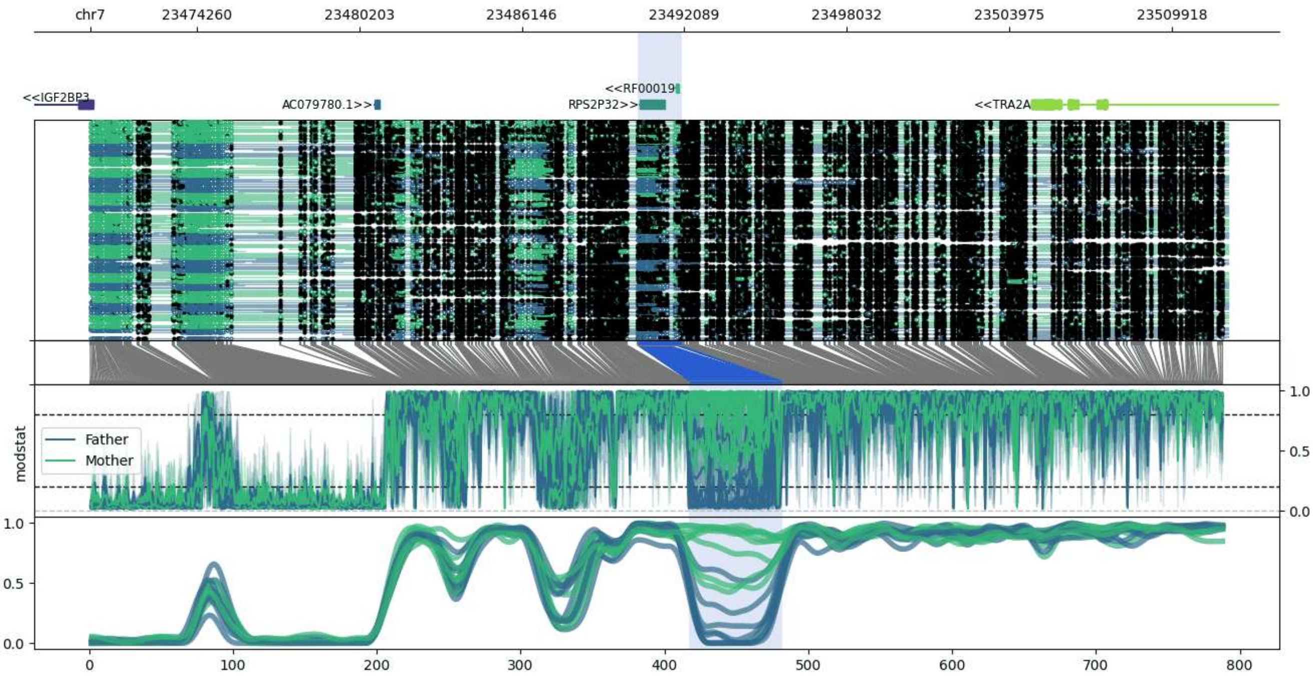


**chr7:23470379-23511983:** Region with 1 DMR containing genes: AC079780.1, IGF2BP3, RF00019, RPS2P32, TRA2A.

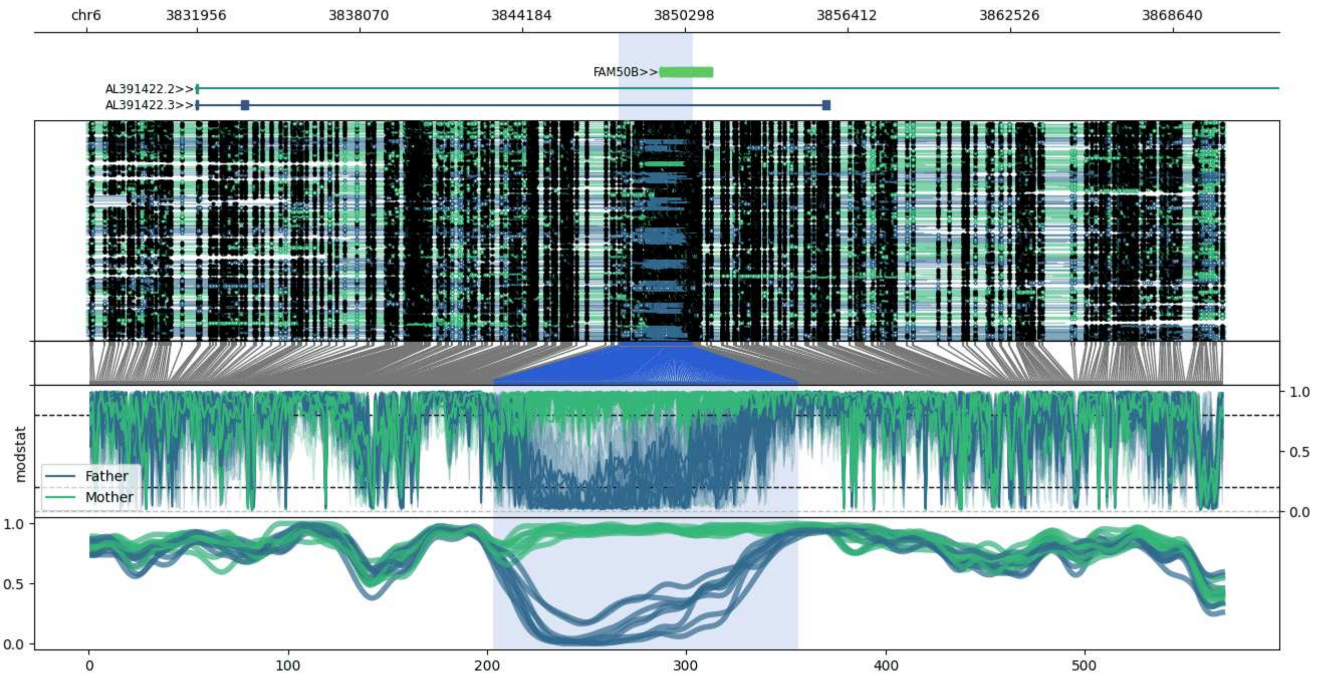


**chr6:3827802-3870601:** Region with 1 DMR containing genes: AL391422.2, AL391422.3, FAM50B.

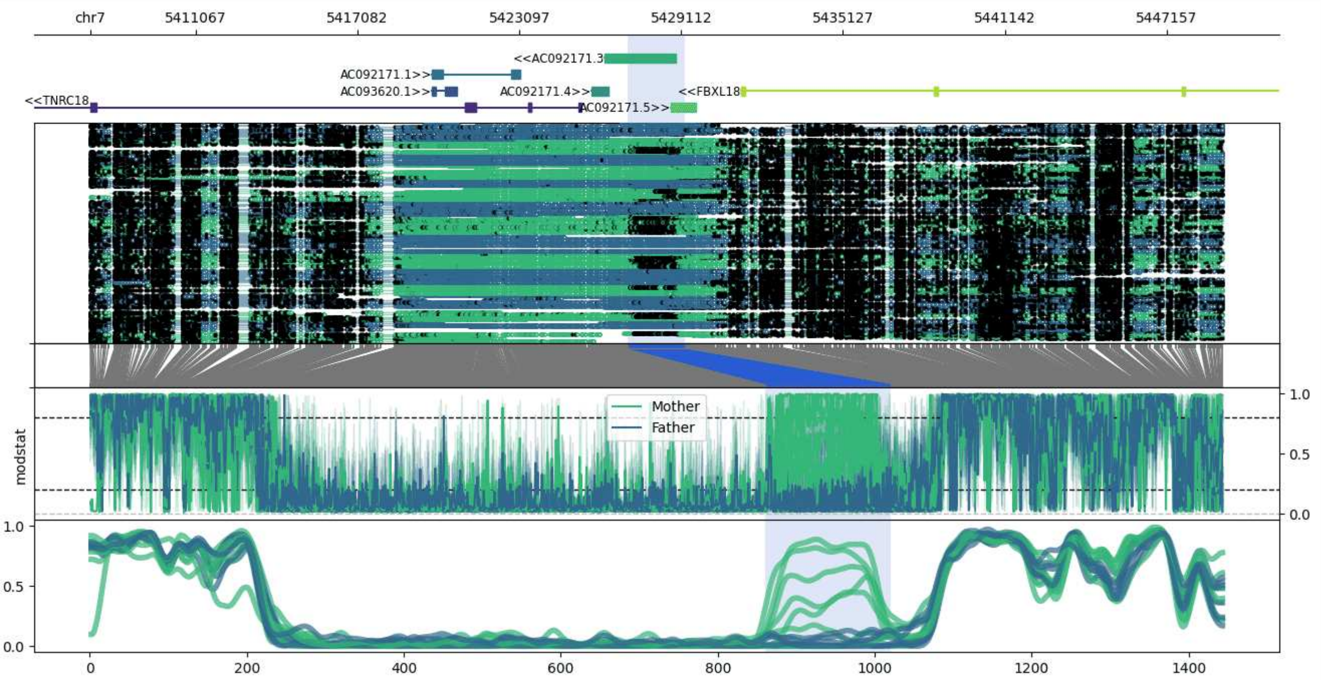


**chr7:5407134-5449242:** Region with 1 DMR containing genes: AC092171.1, AC092171.3, AC092171.4, AC092171.5, AC093620.1, FBXL18, TNRC18.

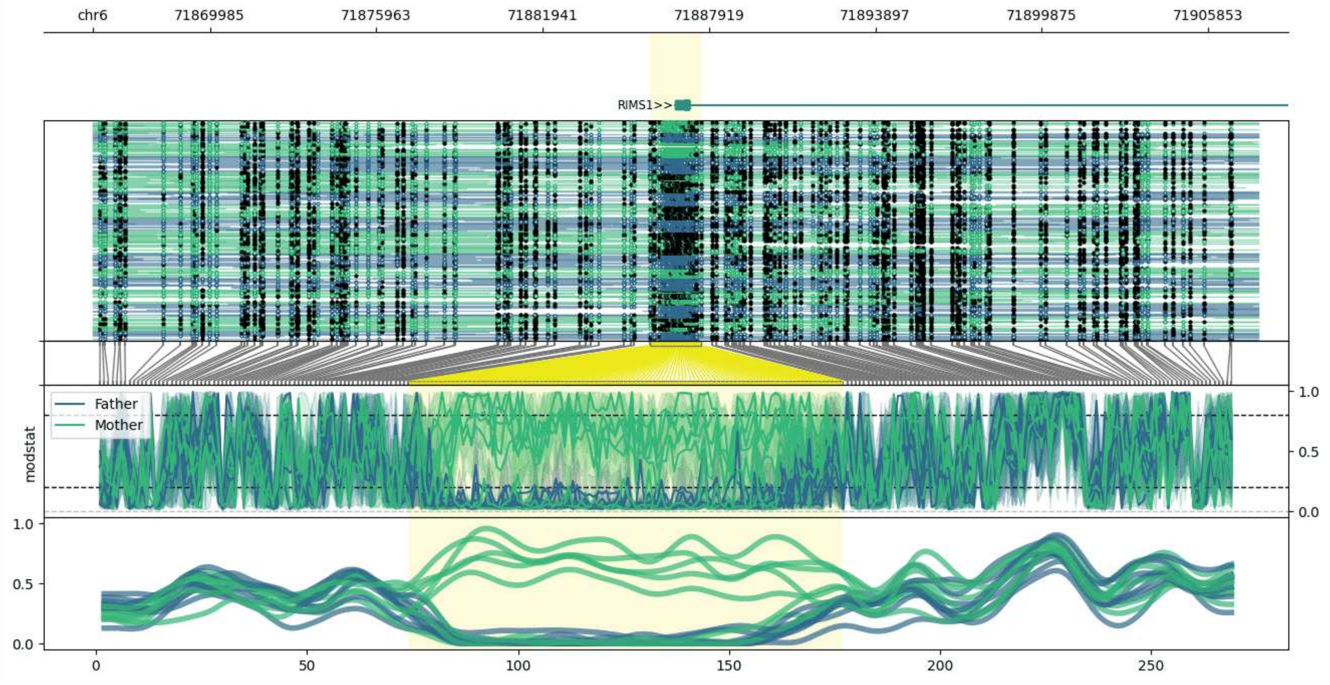


**chr6:71865782-71907634:** Region with 1 DMR containing genes: RIMS1.

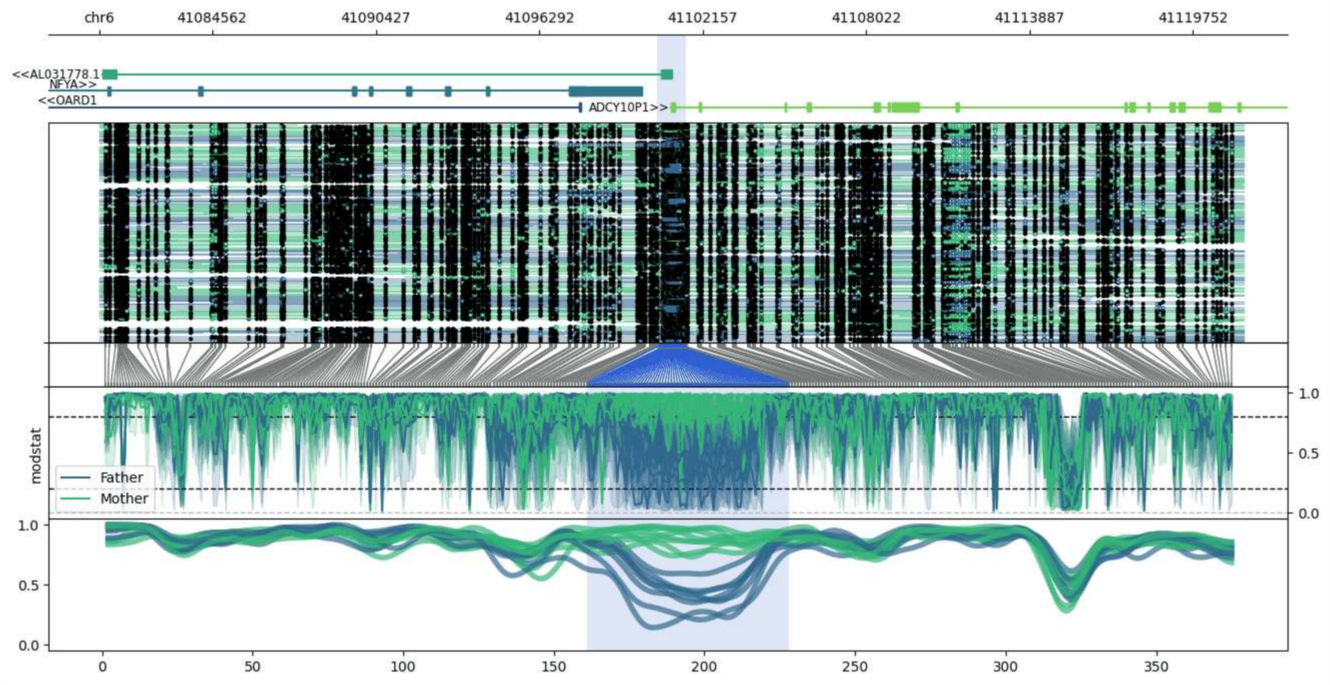


**chr6:41080505-41121563:** Region with 1 DMR containing genes: ADCY10P1, AL031778.1, NFYA, OARD1.

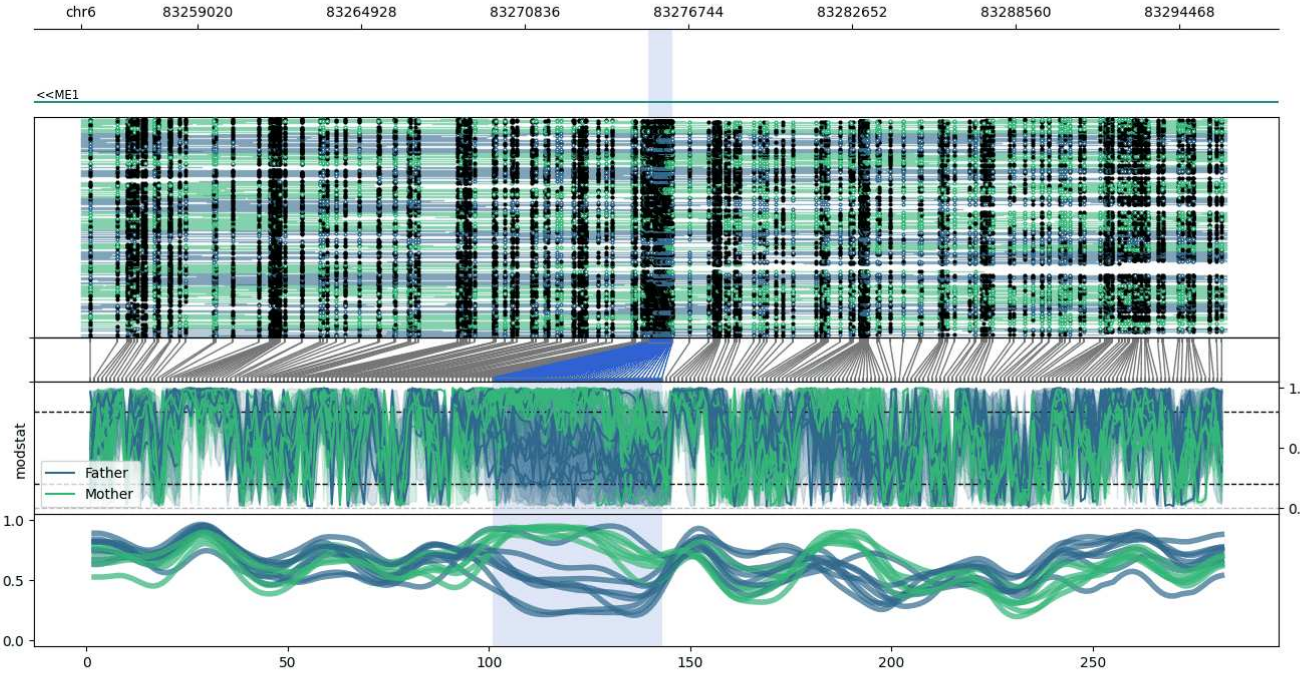


**chr6:83254817-83296173:** Region with 1 DMR containing genes: ME1.

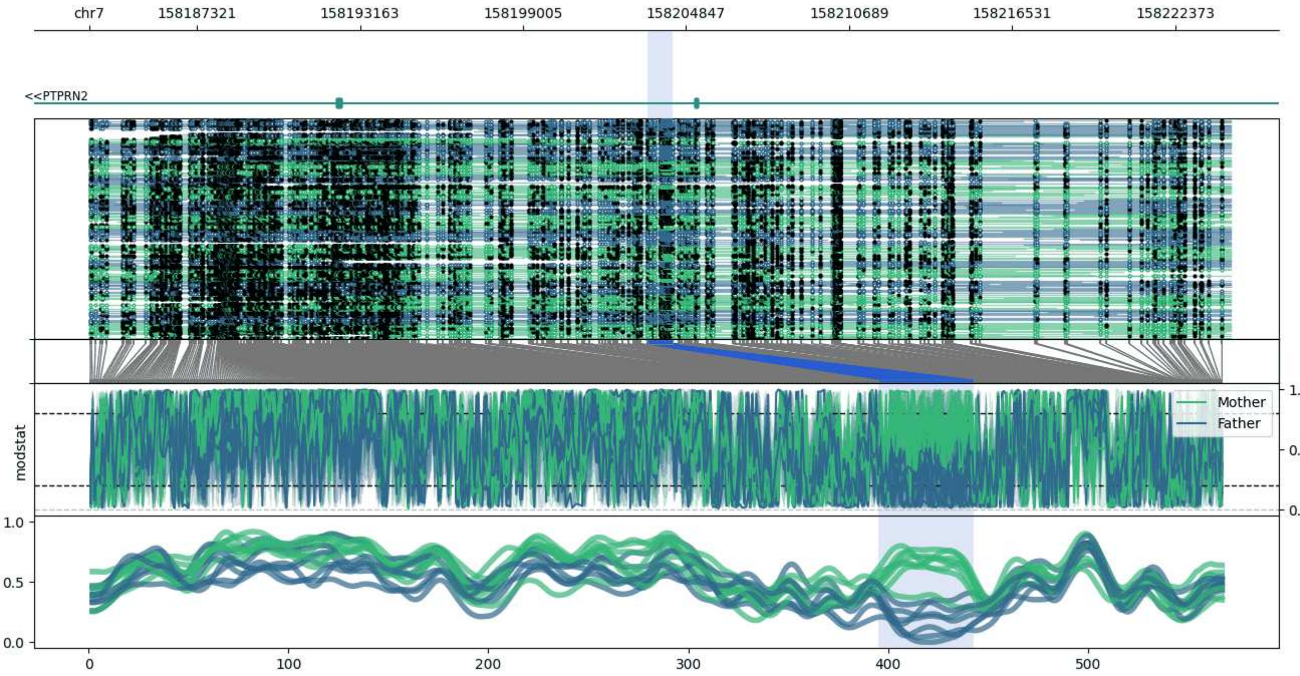


**chr7:158183453-158224352:** Region with 1 DMR containing genes: PTPRN2.

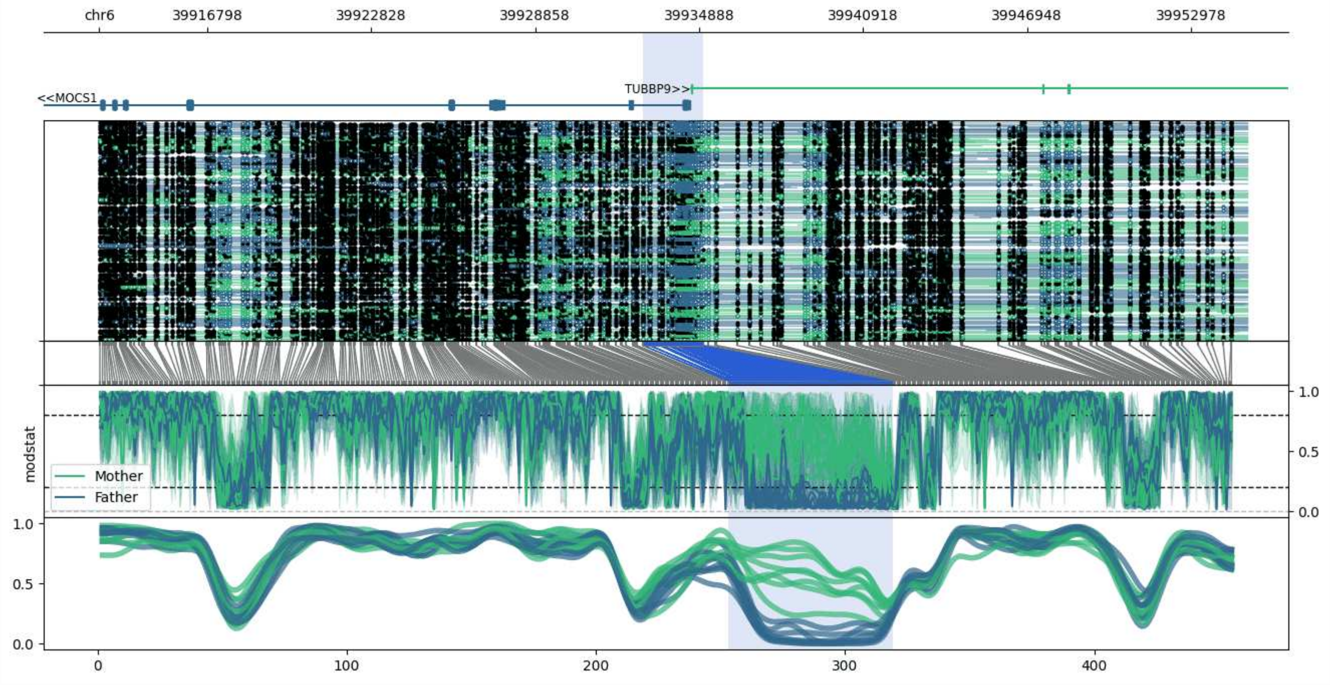


**chr6:39912819-39955035:** Region with 1 DMR containing genes: DAAM2, FO393411.1, MOCS1, TUBBP9.

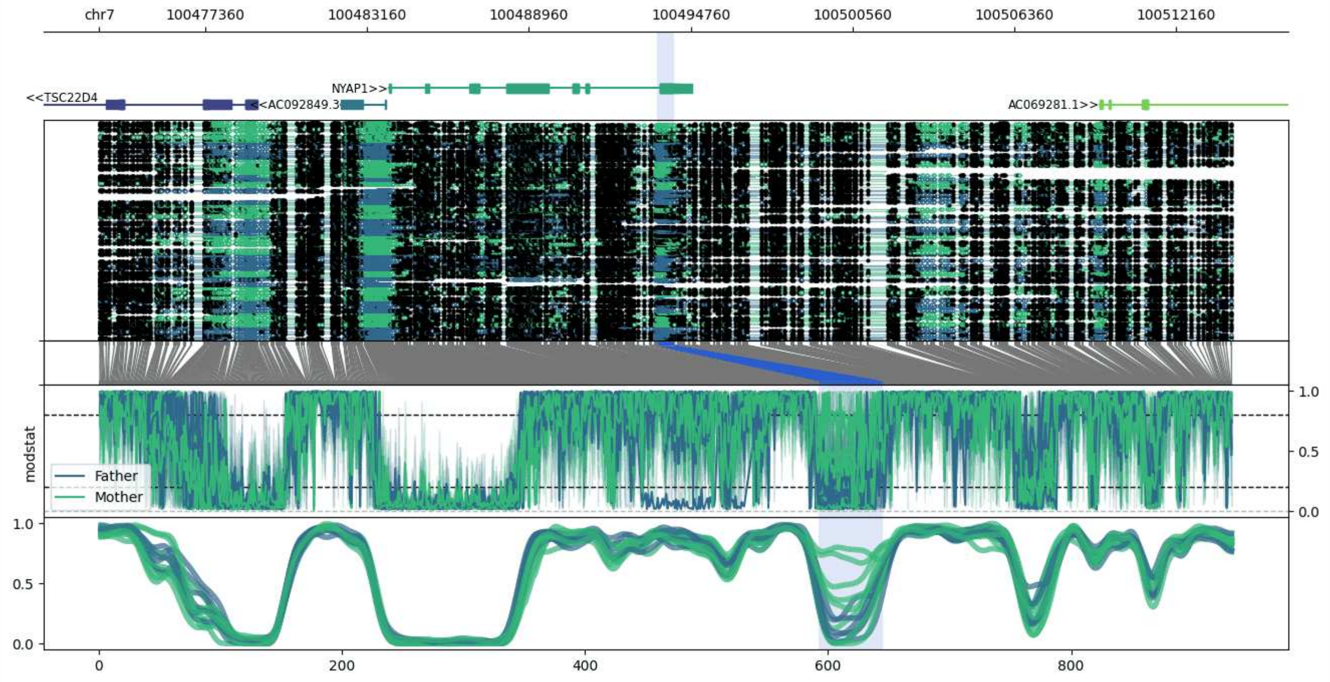


**chr7:100473558-100514163:** Region with 1 DMR containing genes: AC069281.1, AC092849.3, C7orf61, NYAP1, TSC22D4.

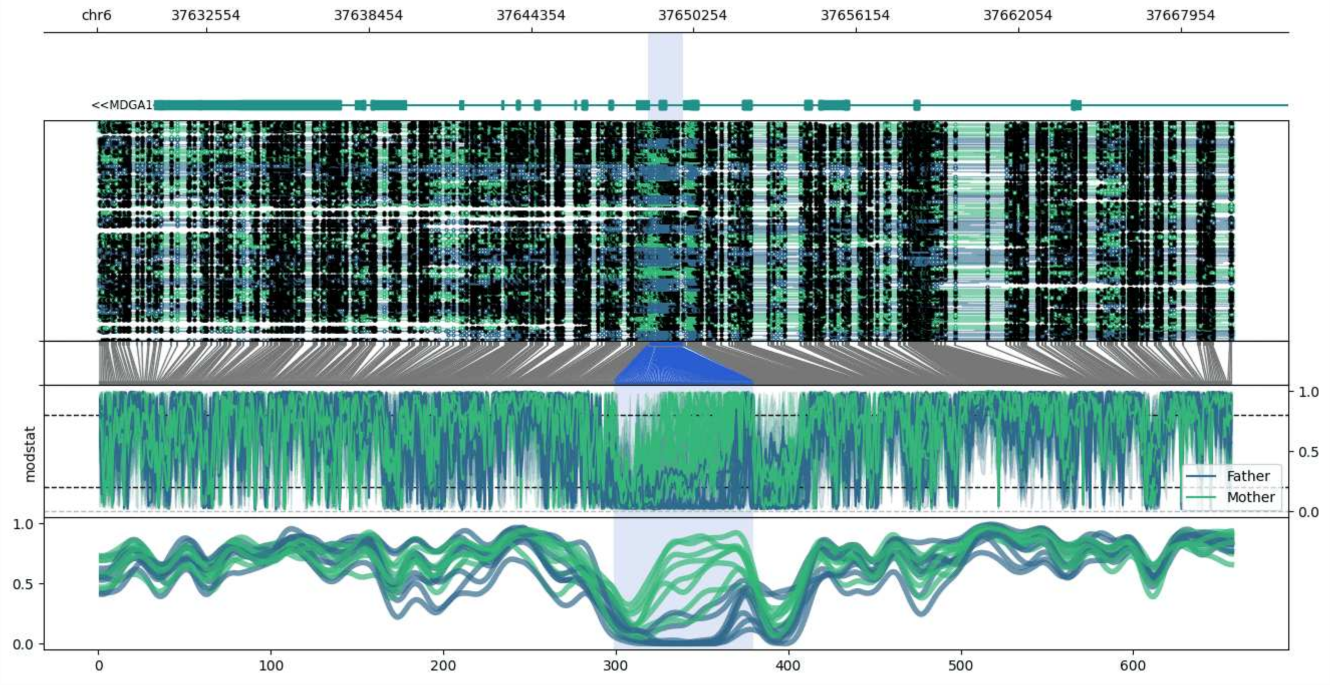


**chr6:37628581-37669886:** Region with 1 DMR containing genes: MDGA1.

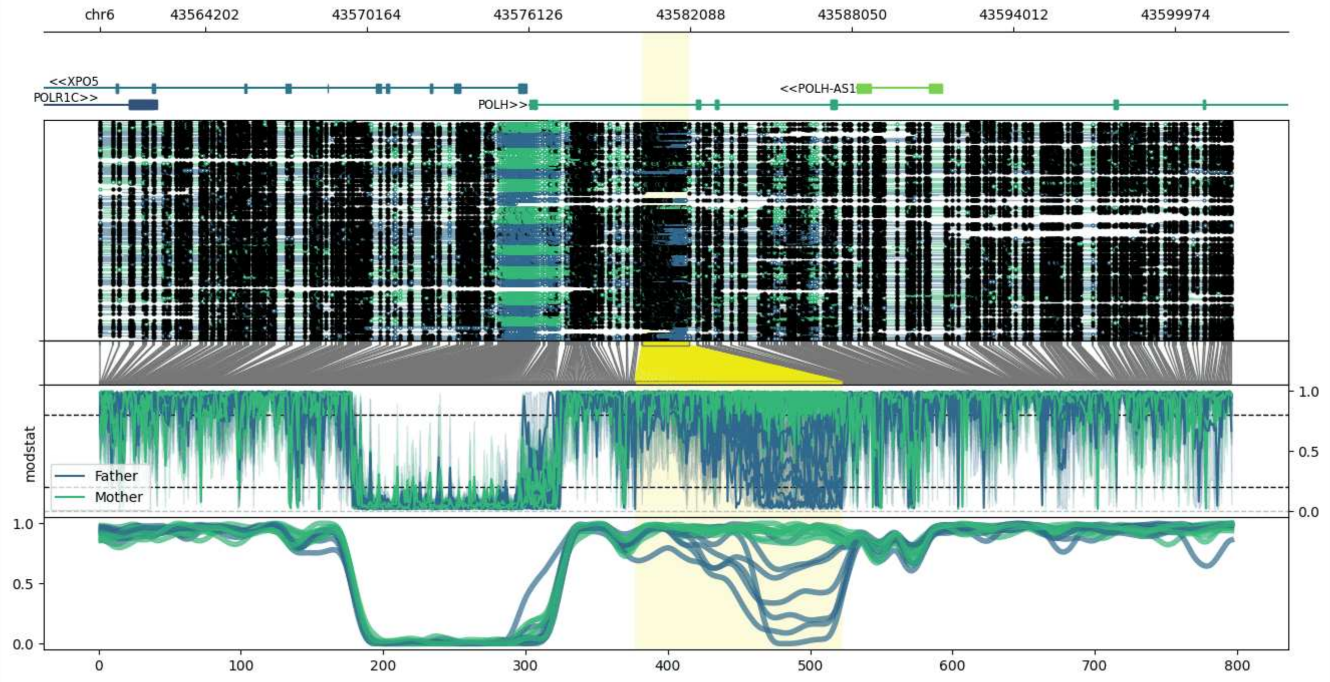


**chr6:43560323-43602063:** Region with 1 DMR containing genes: GTPBP2, POLH, POLH-AS1, POLR1C, XPO5.

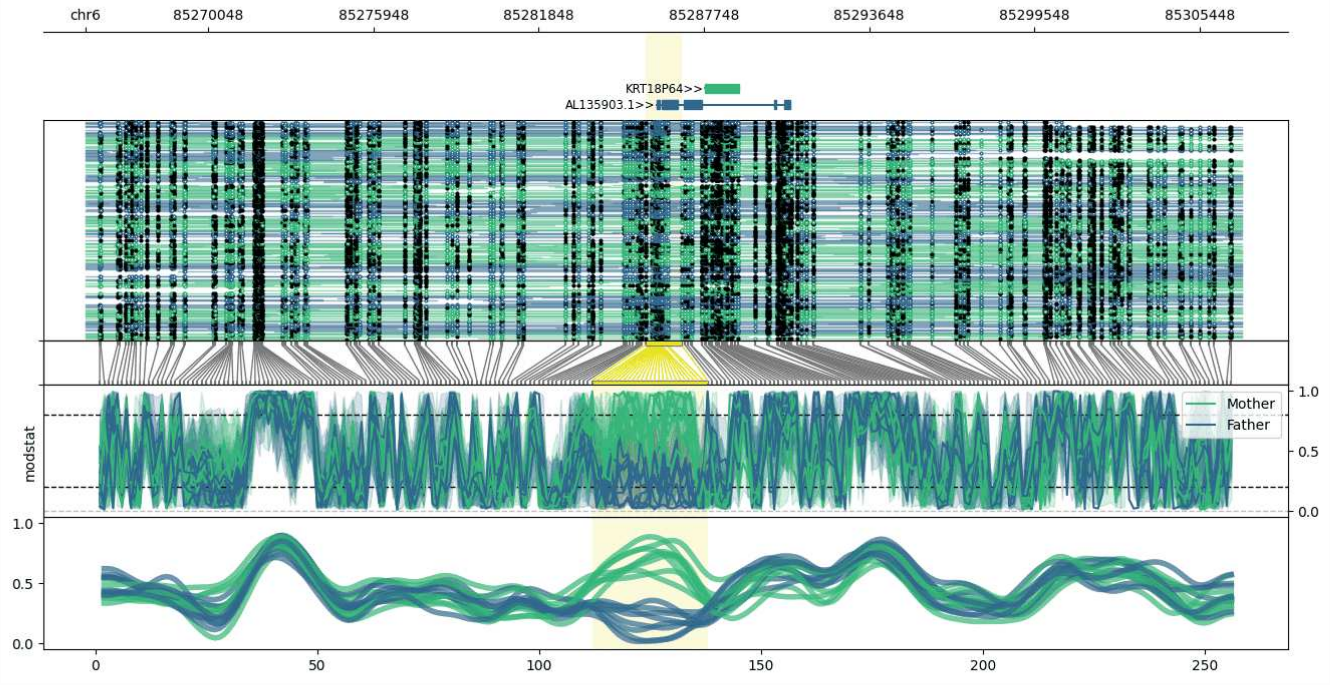


**chr6:85265655-85306957:** Region with 1 DMR containing genes: AL122016.1, AL135903.1, KRT18P64.

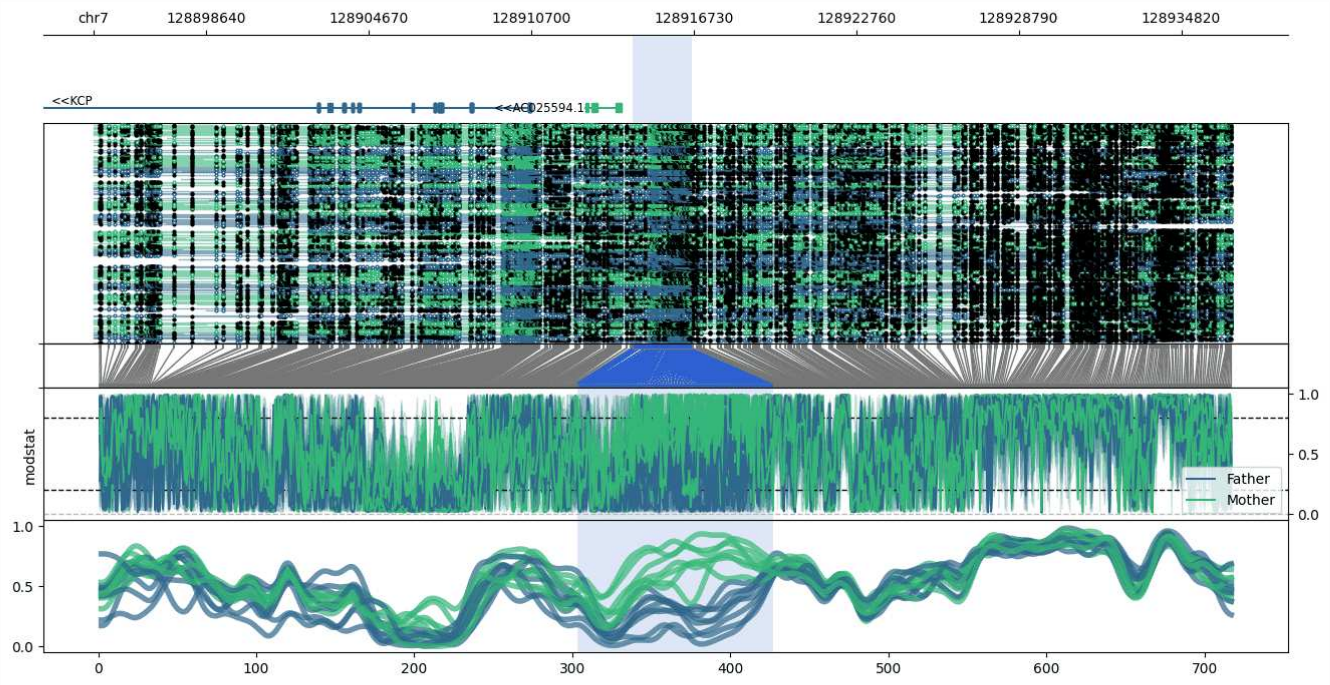


**chr7:128894468-128936679:** Region with 1 DMR containing genes: AC025594.1, IRF5, KCP.

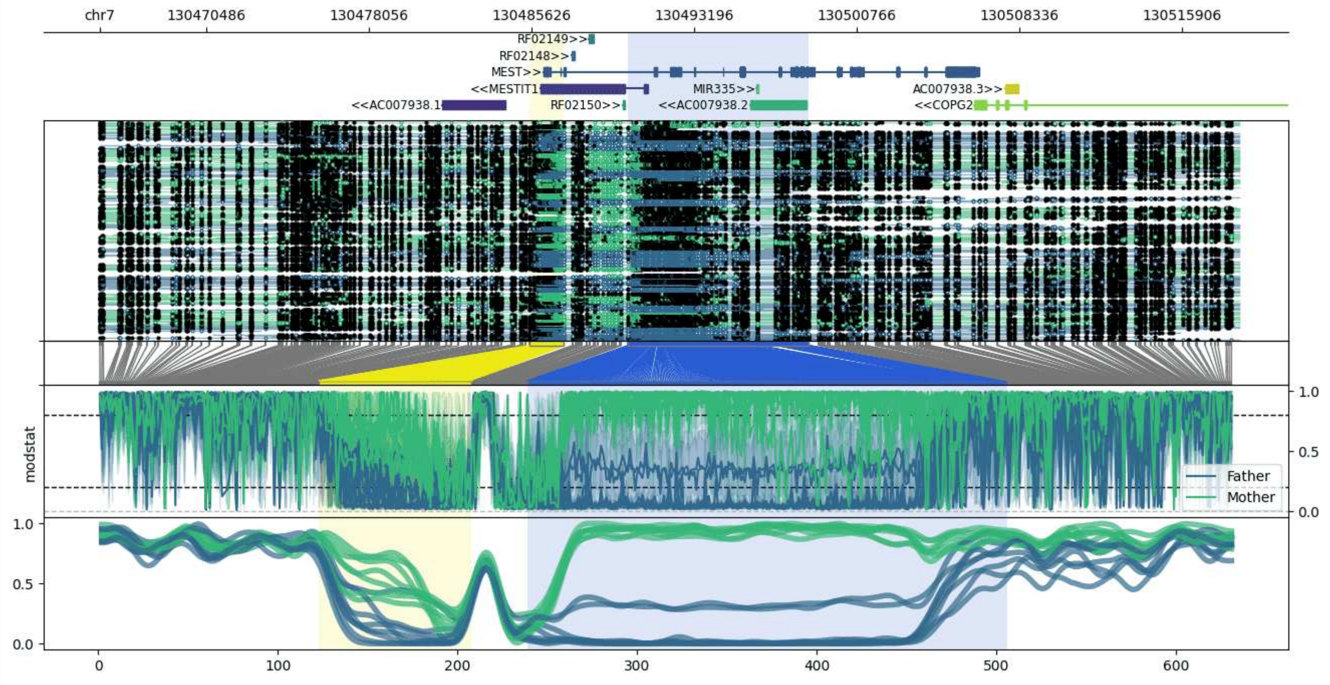


**chr7:130465548-130518539:** Region with 2 DMRs containing genes: AC007938.1, AC007938.2, AC007938.3, COPG2, MEST, MESTIT1, MIR335, RF02148, RF02149, RF02150.

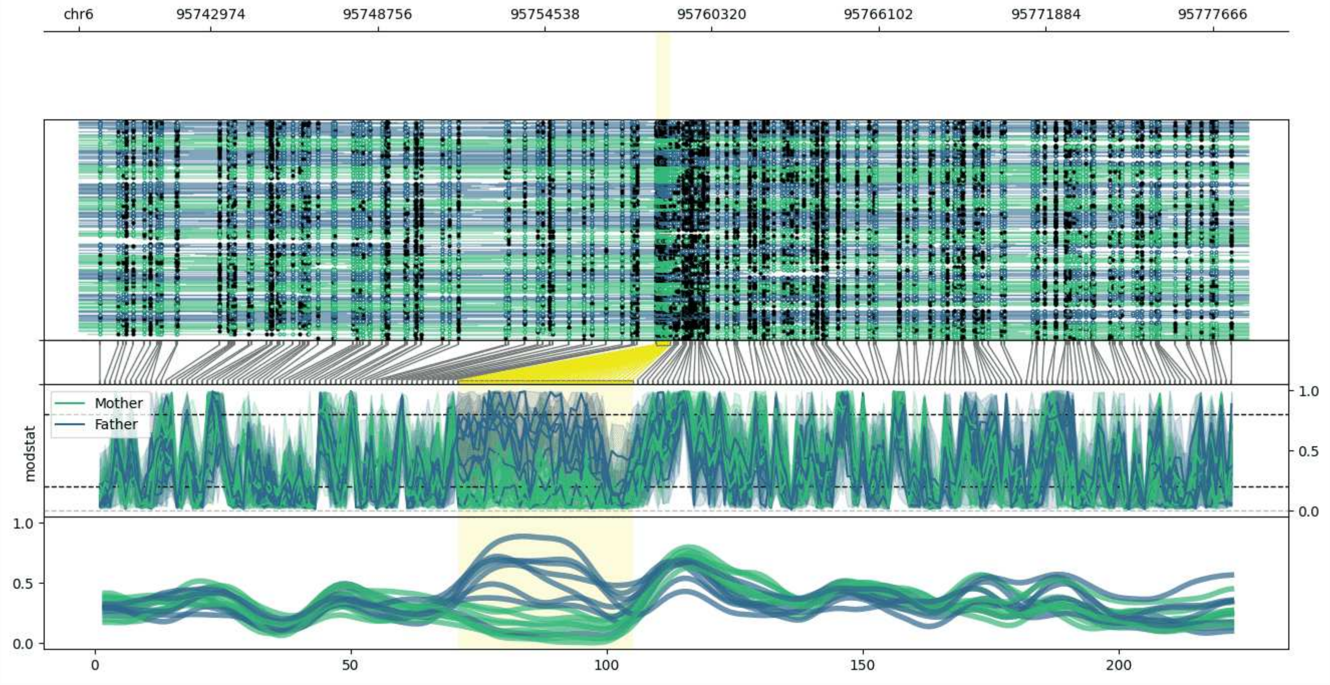


**chr6:95738405-95778885:** Region with 1 DMR containing no genes.

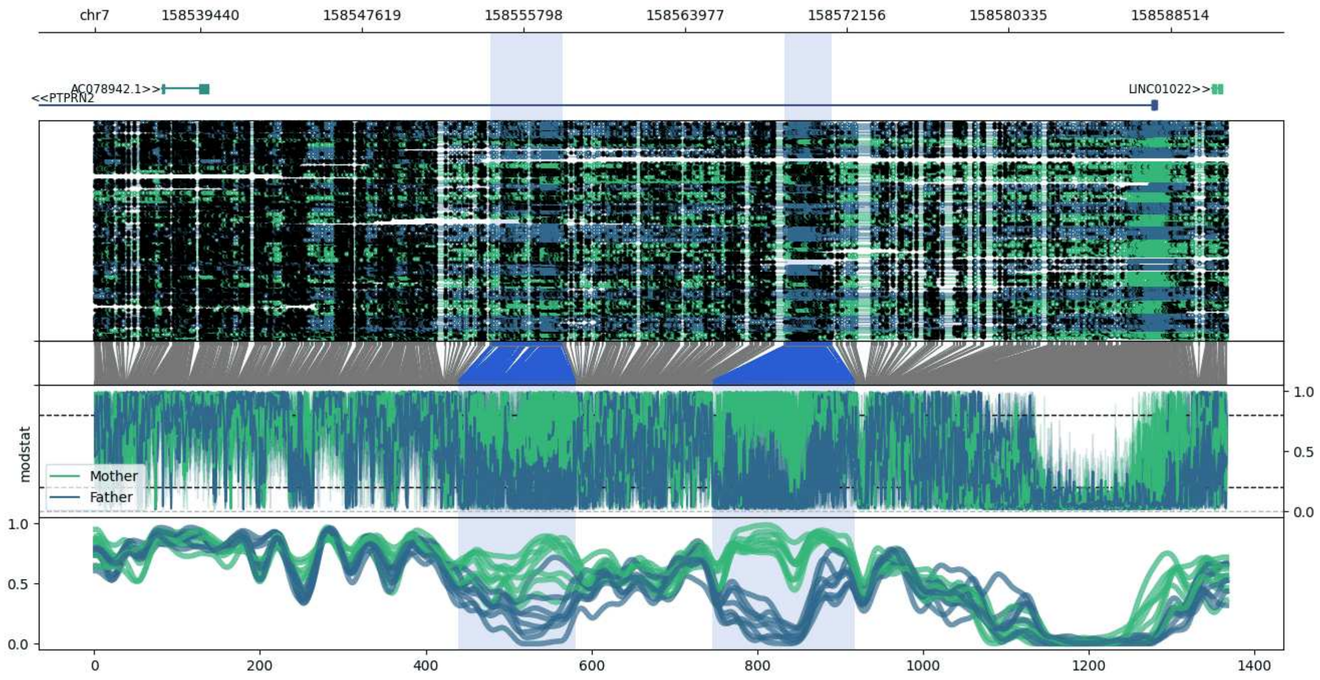


**chr7:158534121-158591374:** Region with 2 DMRs containing genes: AC078942.1, LINC01022, MIR5707, MIR595, PTPRN2.

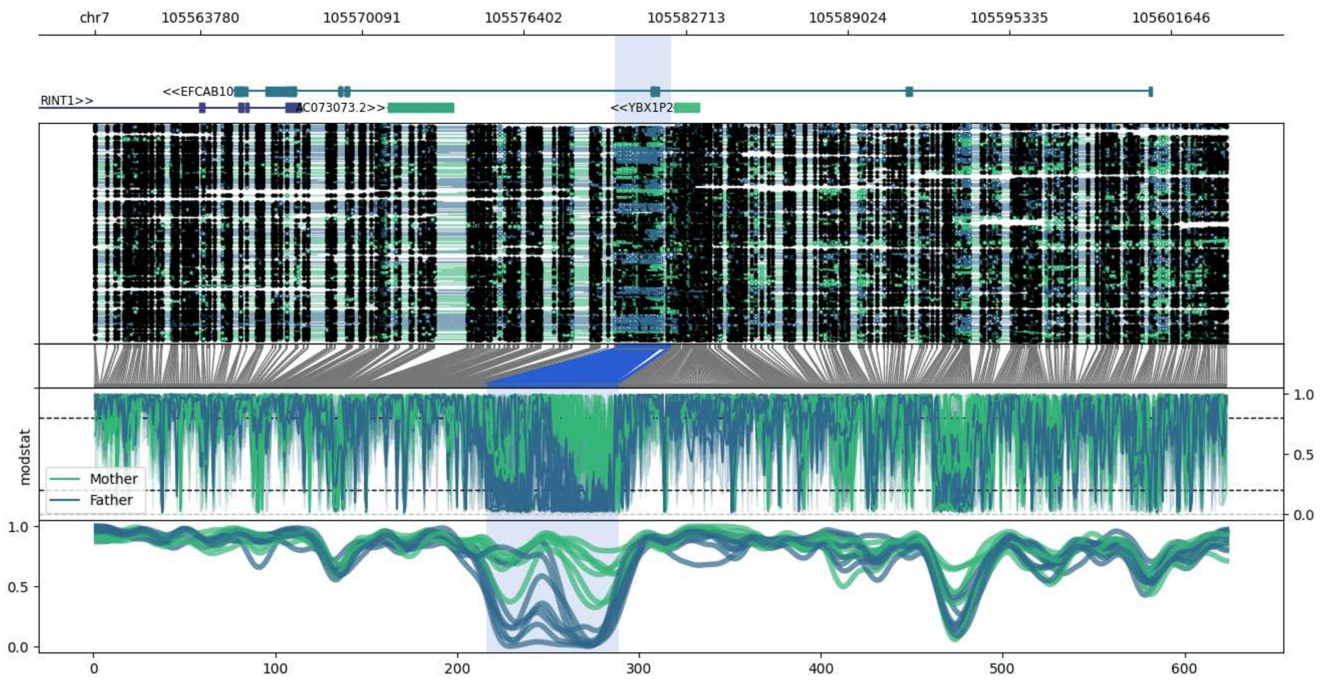


**chr7:105559668-105603850:** Region with 1 DMR containing genes: AC073073.2, ATXN7L1, EFCAB10, RINT1, YBX1P2.

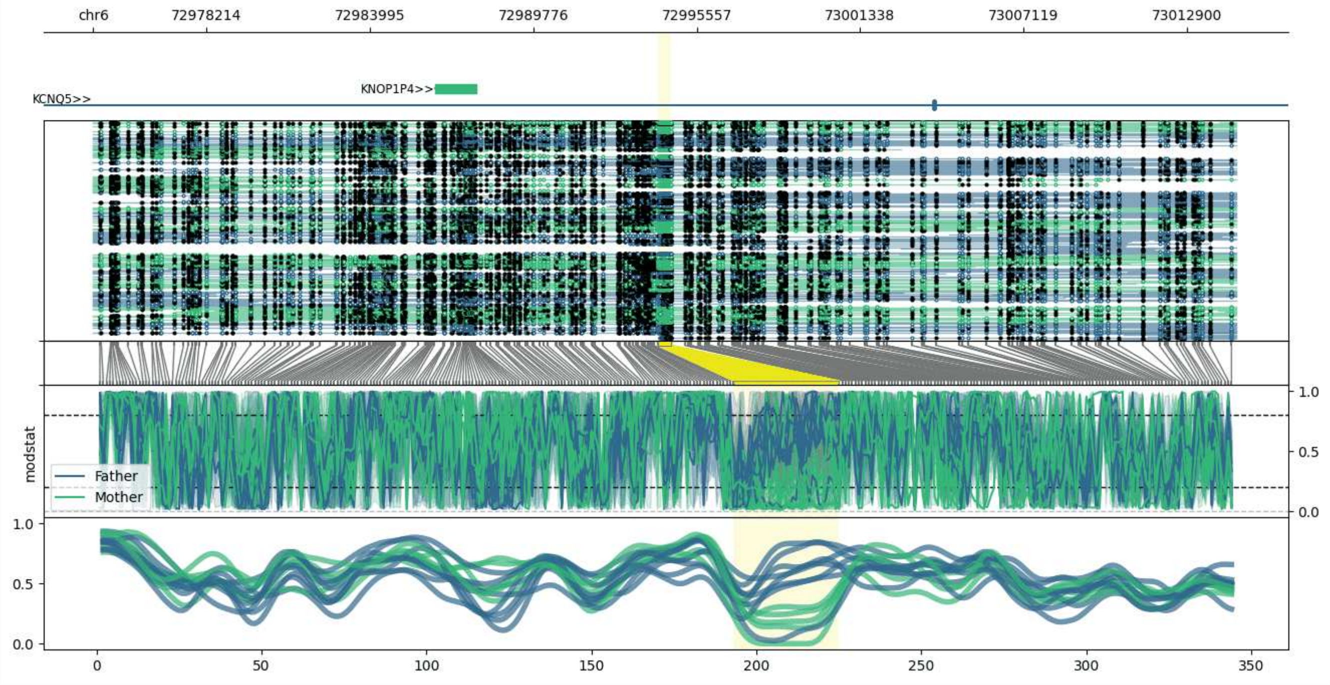


**chr6:72974193-73014664:** Region with 1 DMR containing genes: KCNQ5, KNOP1P4, MIR4282.

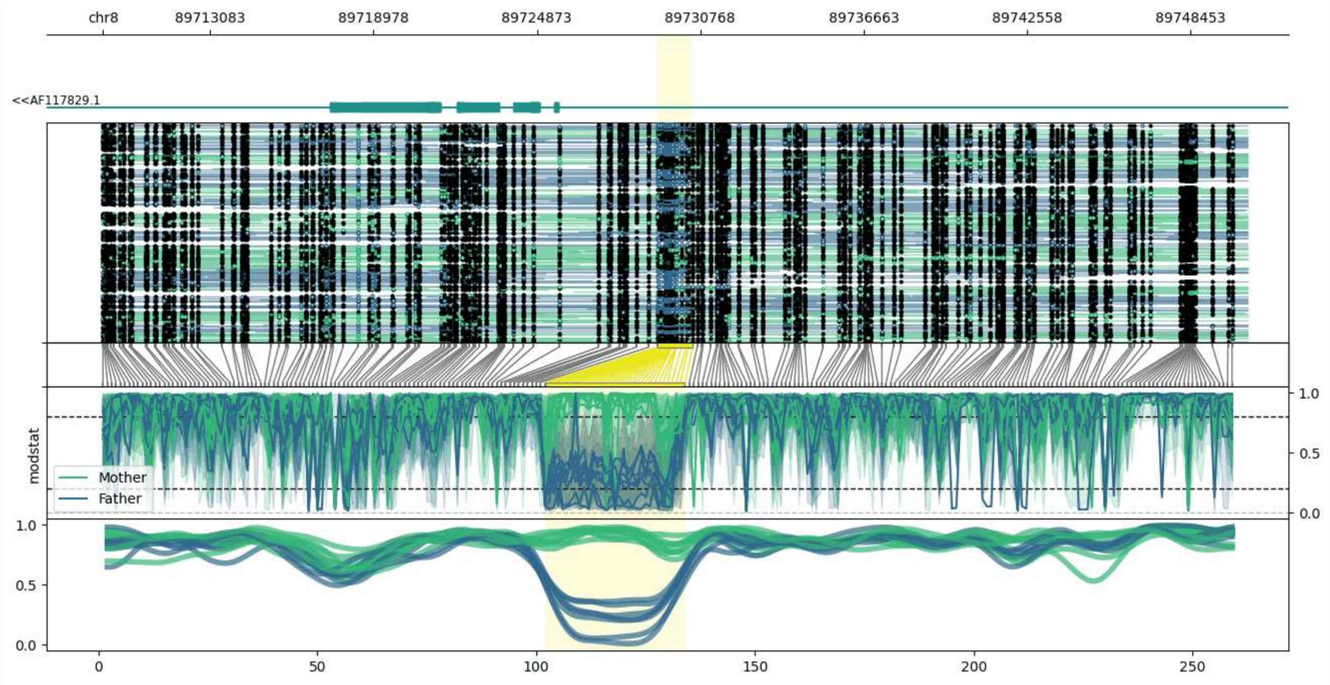


**chr8:89709220-89750491:** Region with 1 DMR containing genes: AF117829.1, RIPK2.

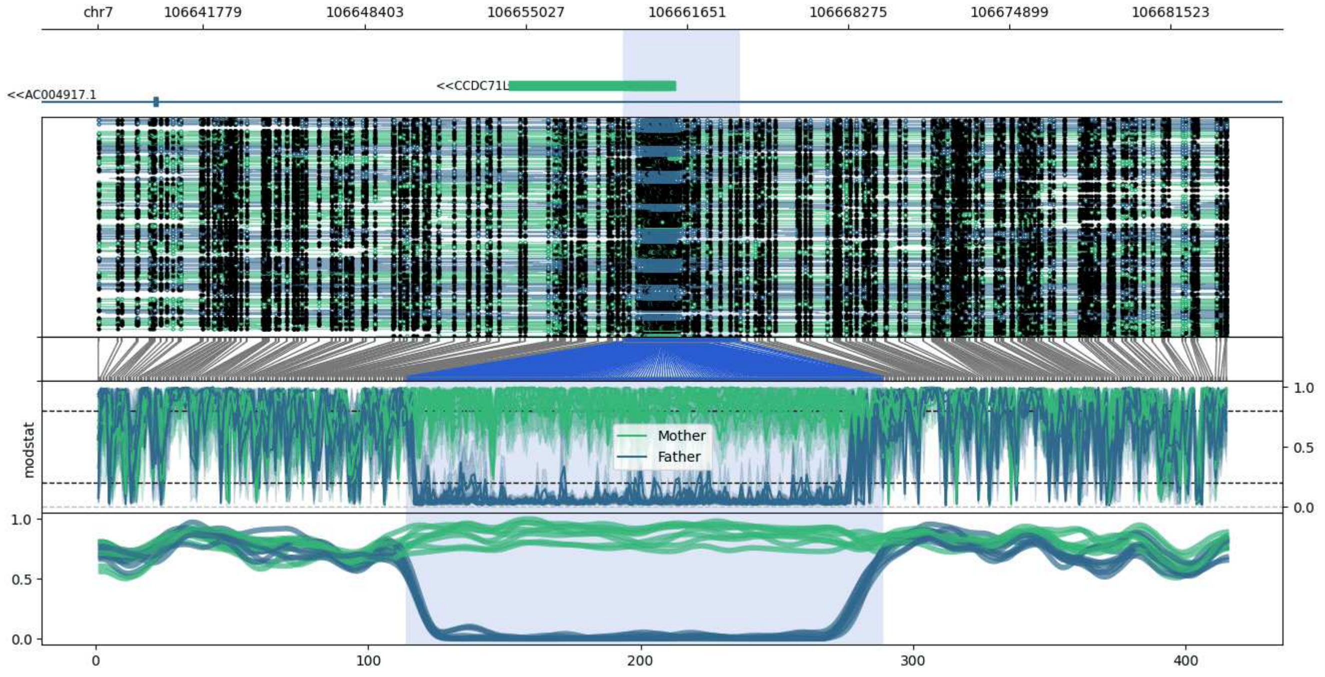


**chr7:106637459-106683830:** Region with 1 DMR containing genes: AC004917.1, AC005050.2, CCDC71L.

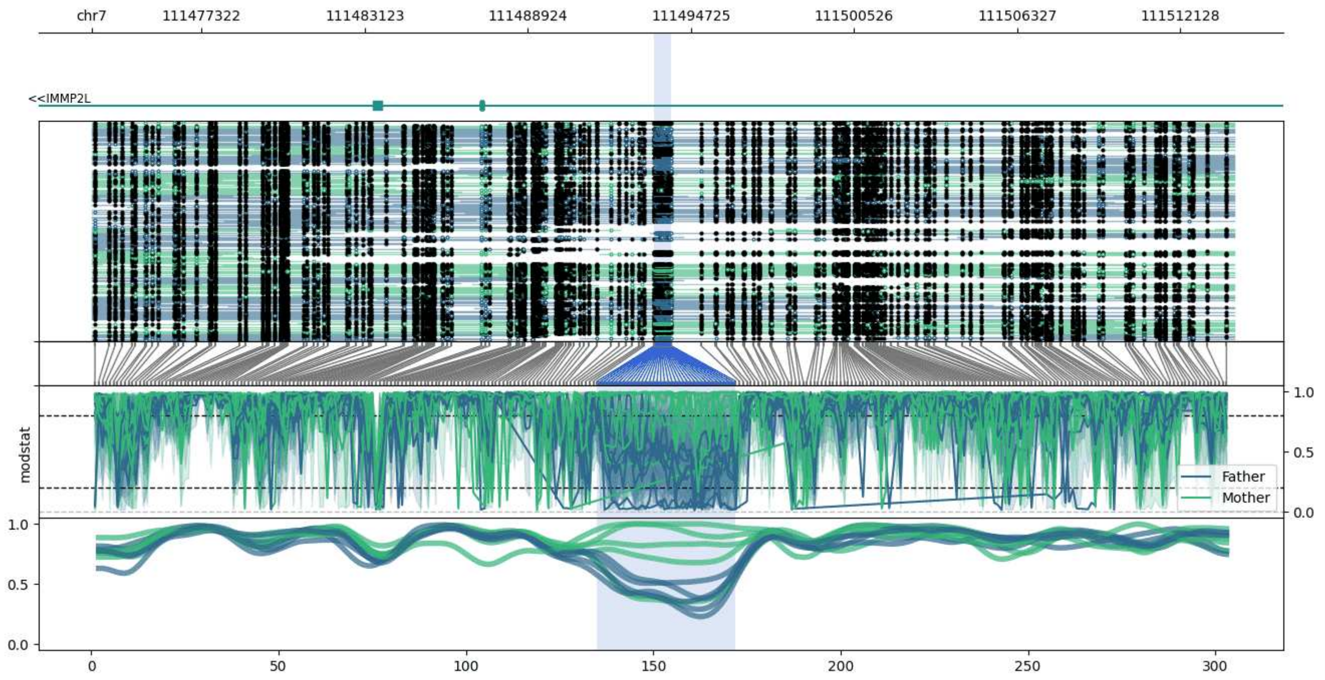


**chr7:111473423-111514035:** Region with 1 DMR containing genes: IMMP2L.

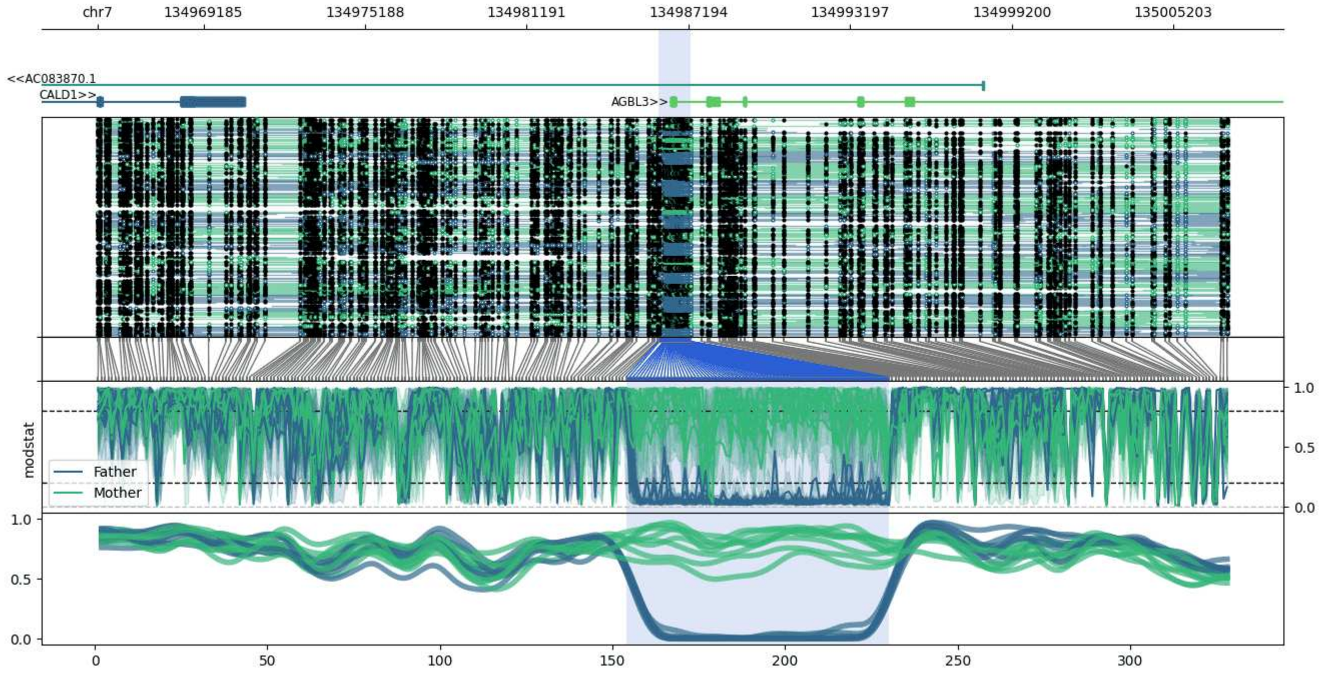


**chr7:134965261-135007282:** Region with 1 DMR containing genes: AC083870.1, AGBL3, CALD1.

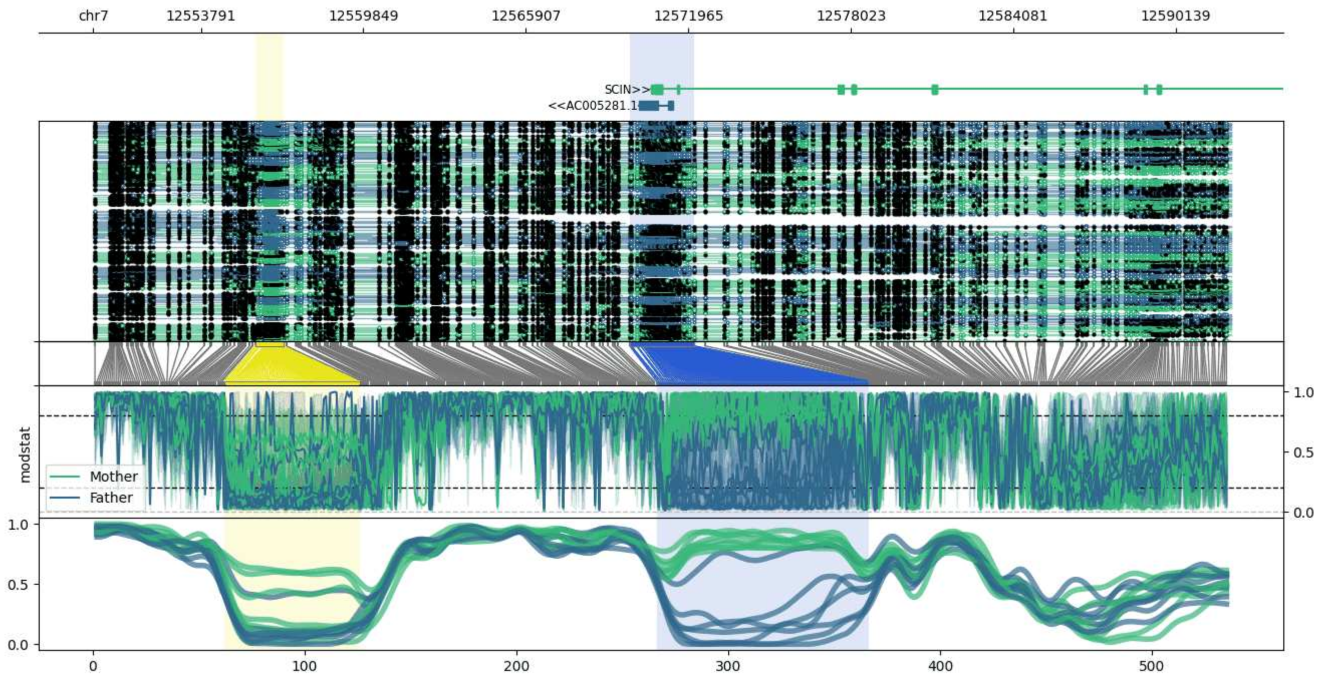


**chr7:12549780-12592190:** Region with 2 DMRs containing genes: AC005281.1, AC013470.2, SCIN.

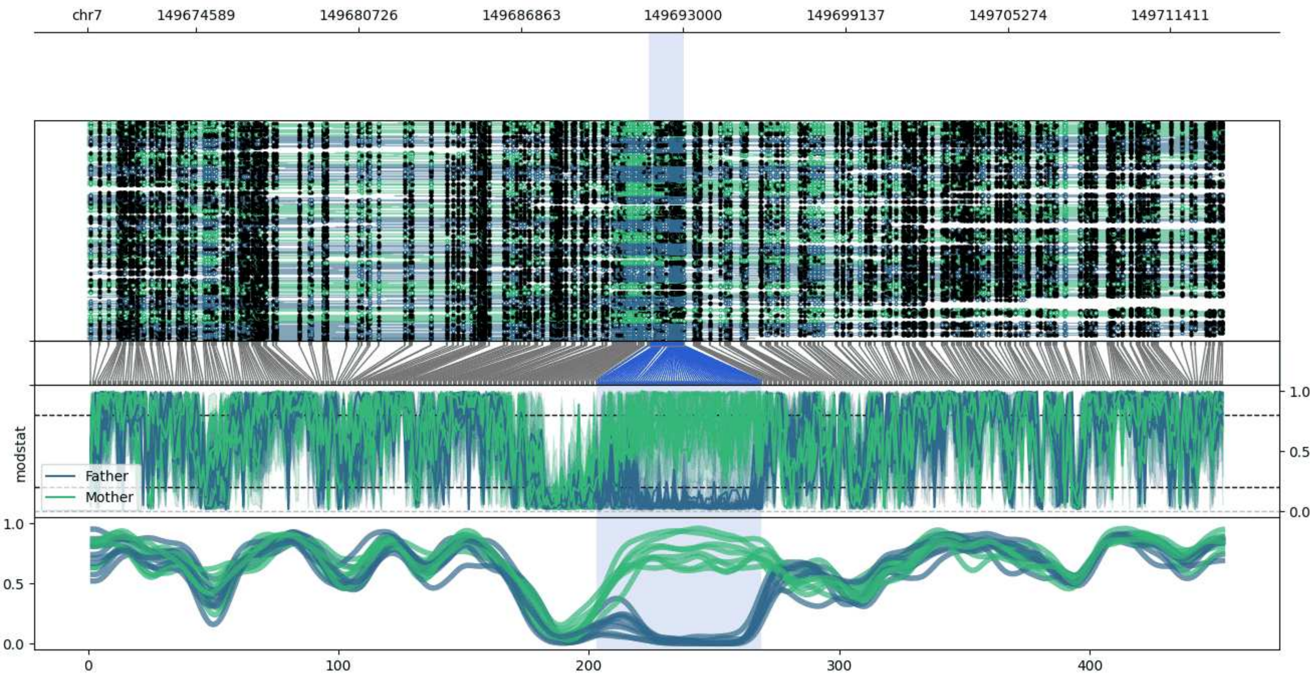


**chr7:149670462-149713422:** Region with 1 DMR containing genes: KRBA1.

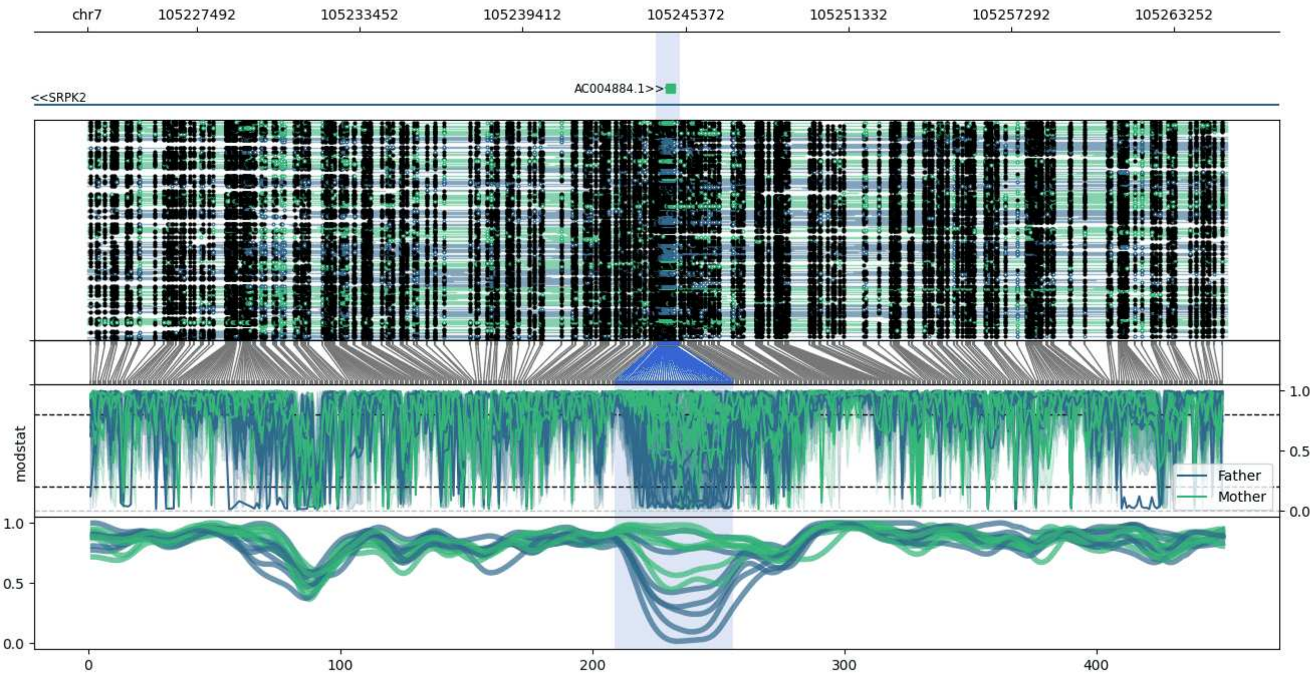


**chr7:105223484-105265206:** Region with 1 DMR containing genes: AC004884.1, SRPK2.

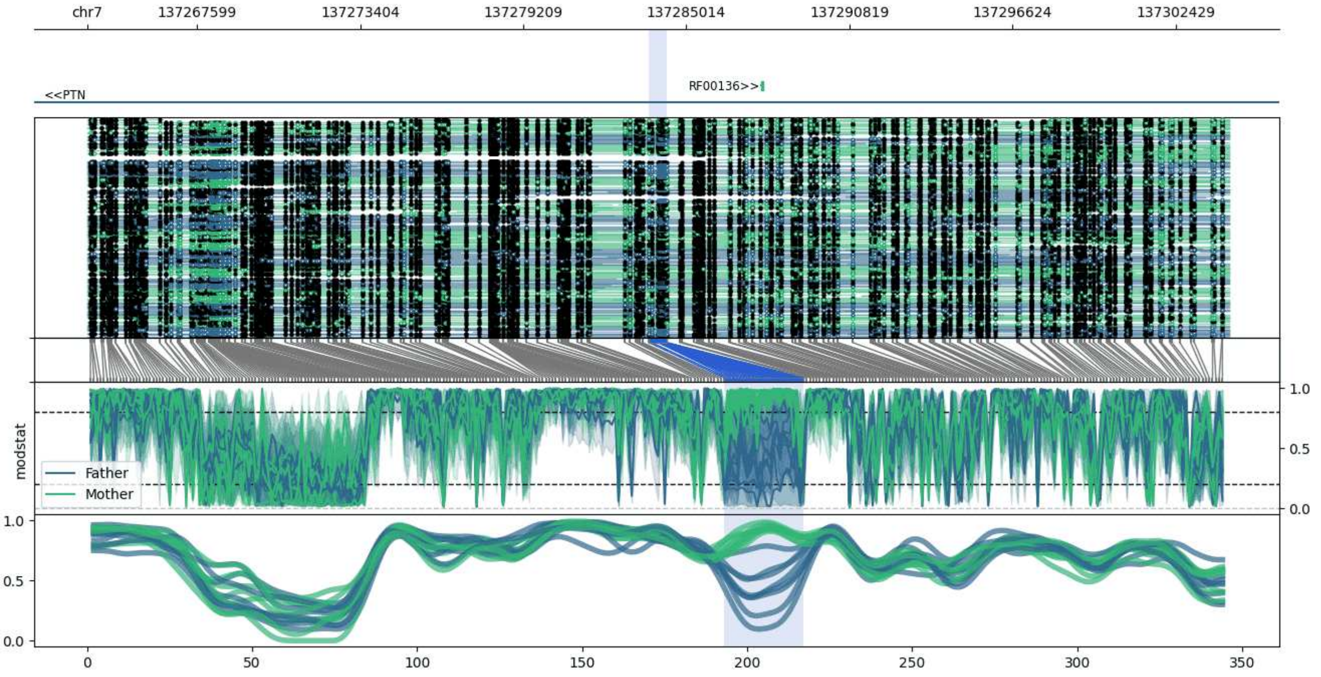


**chr7:137263680-137304321:** Region with 1 DMR containing genes: PTN, RF00136.

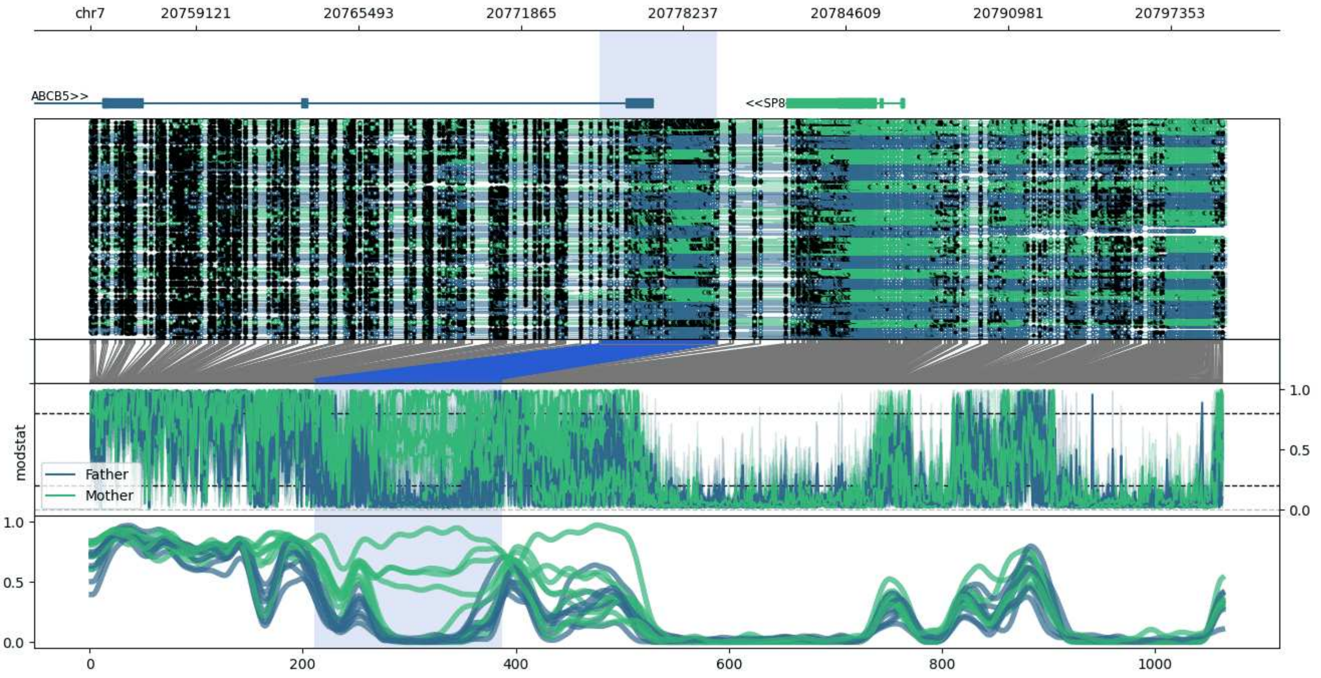


**chr7:20754948-20799554:** Region with 1 DMR containing genes: ABCB5, SP8.

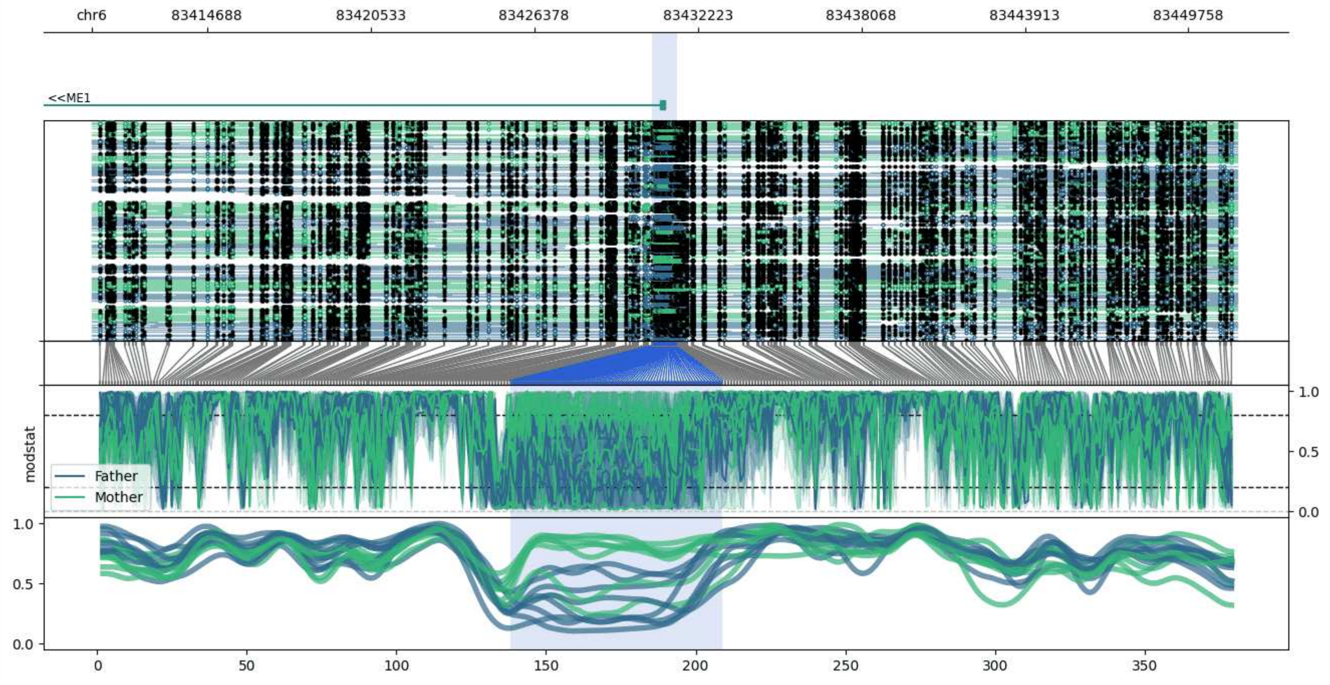


**chr6:83410580-83451499:** Region with 1 DMR containing genes: ME1.

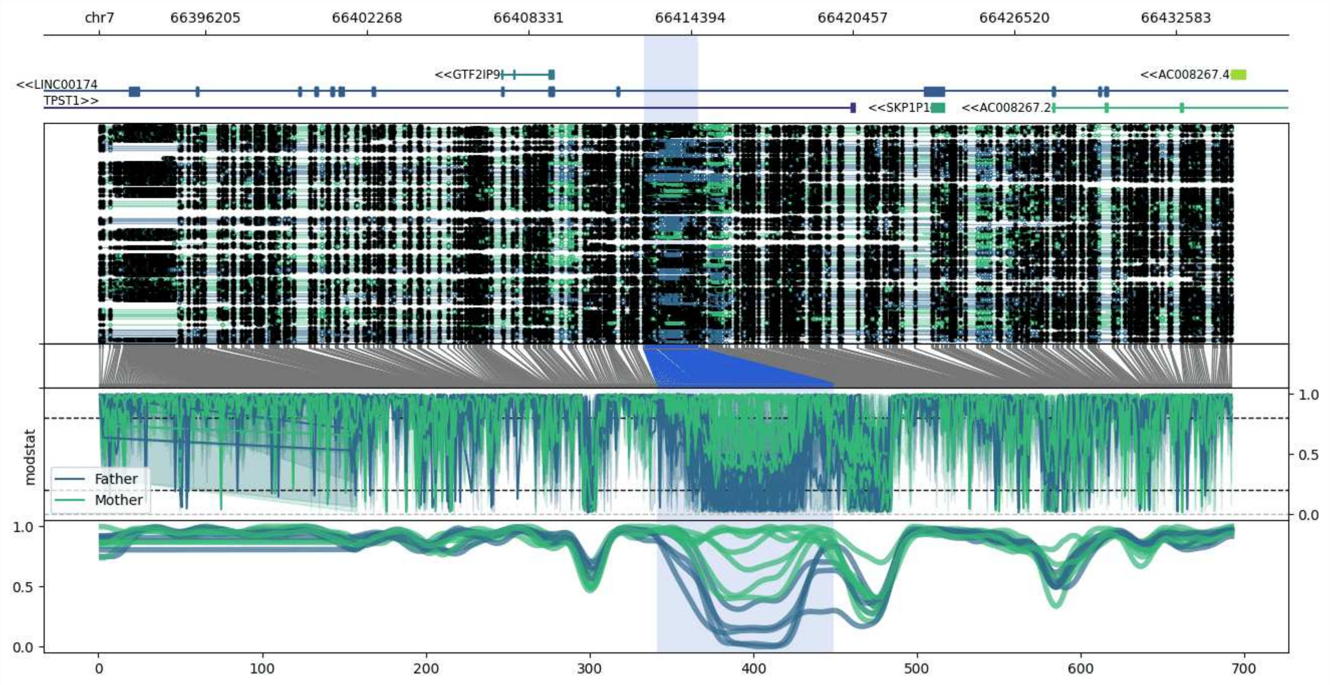


**chr7:66392235-66434682:** Region with 1 DMR containing genes: AC008267.2, AC008267.4, GTF2IP9, LINC00174, SKP1P1, TPST1.

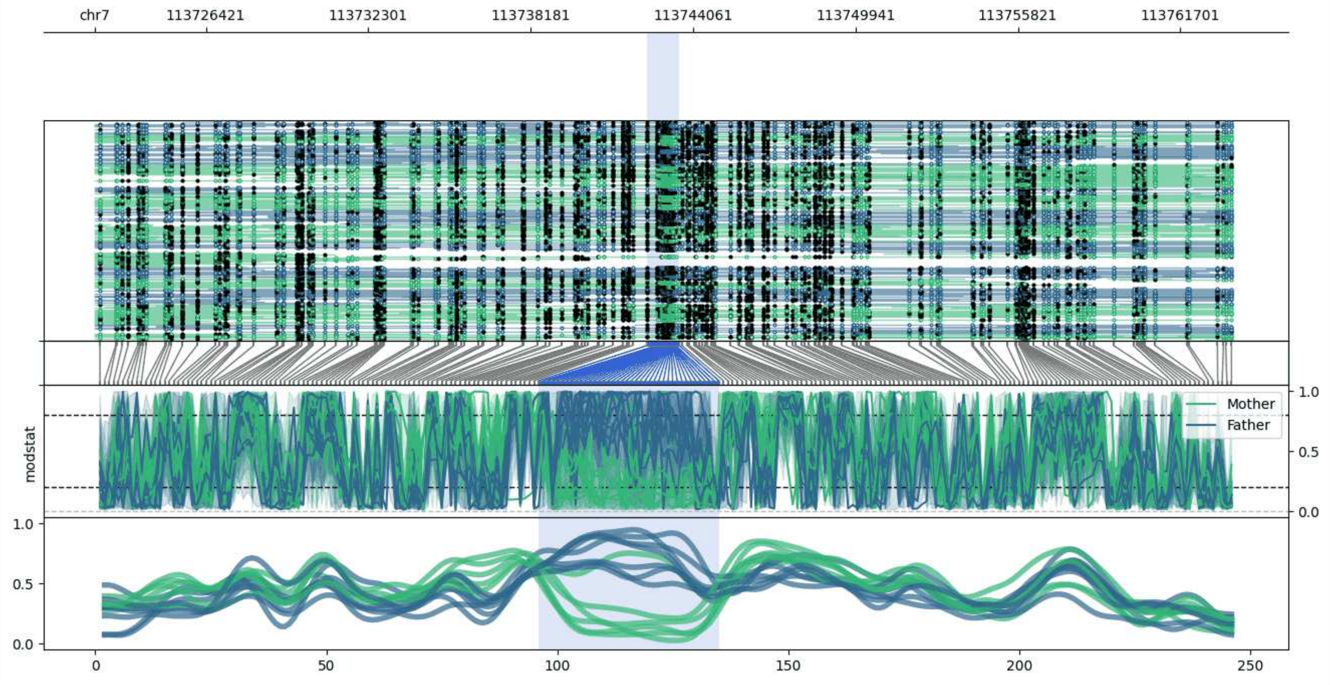


**chr7:113722398-113763559:** Region with 1 DMR containing no genes.

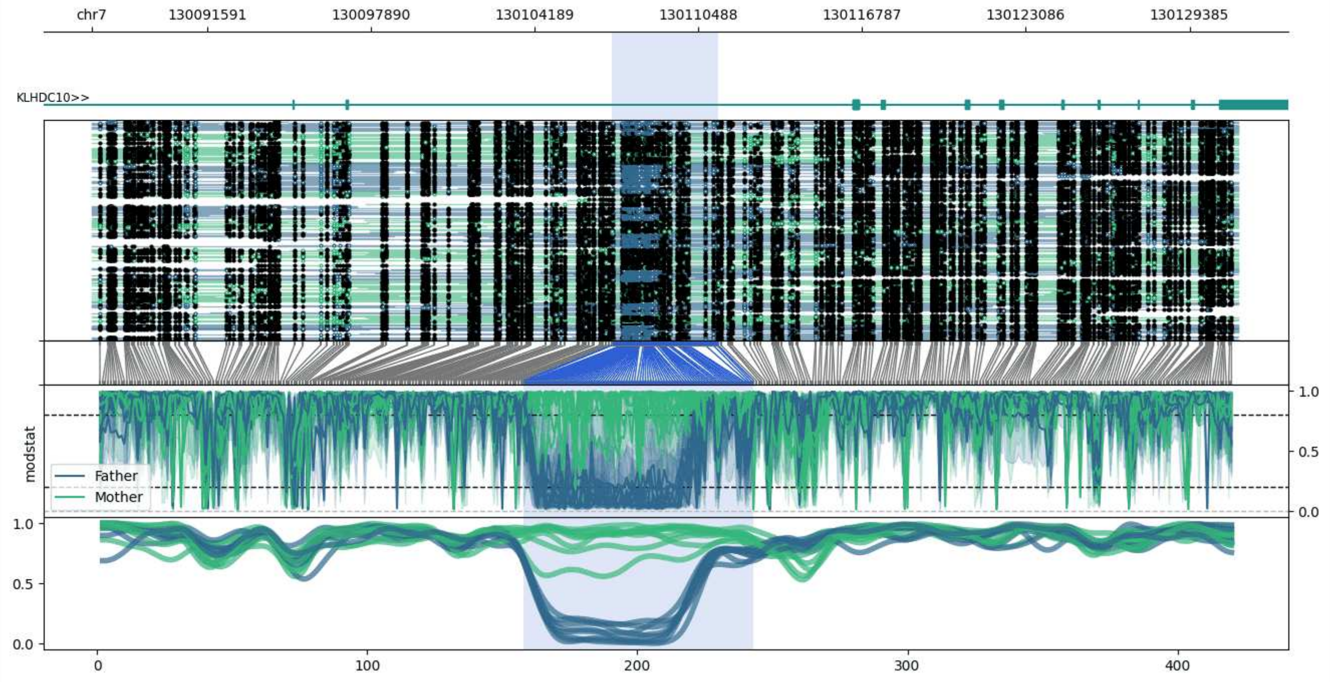


**chr7:130087146-130131244:** Region with 1 DMR containing genes: KLHDC10.

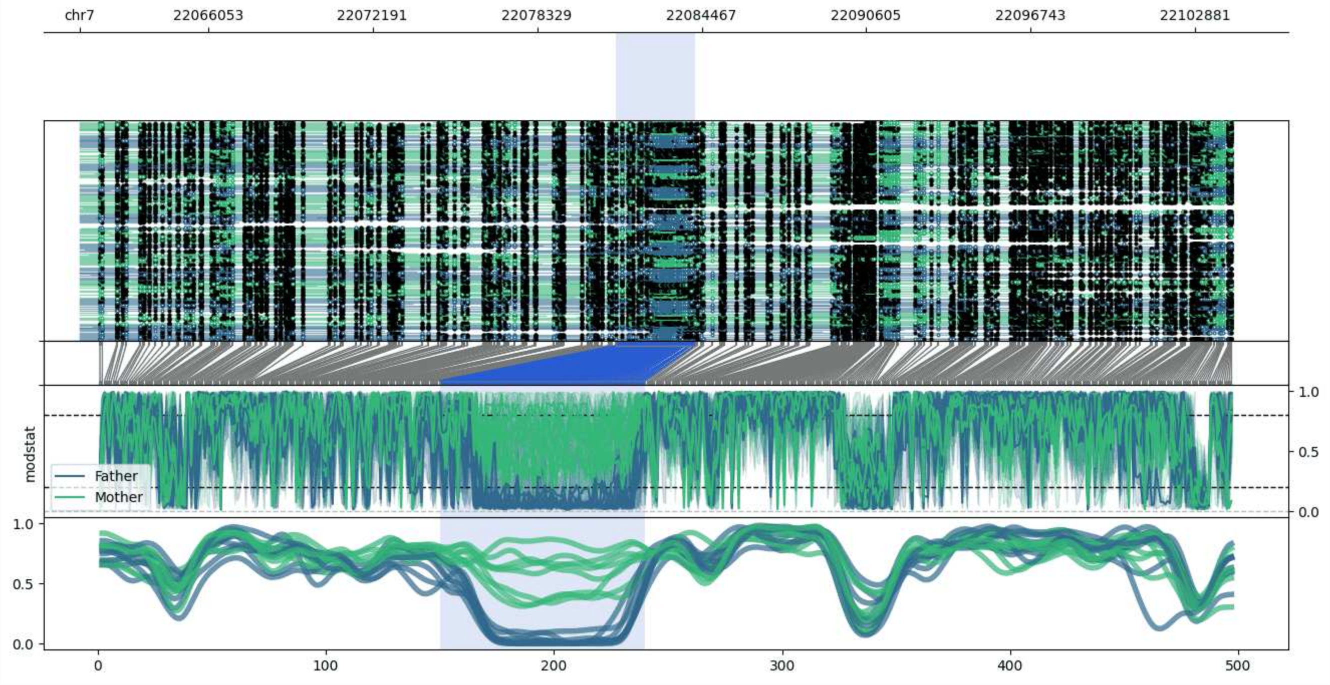


**chr7:22061258-22104230:** Region with 1 DMR containing no genes.

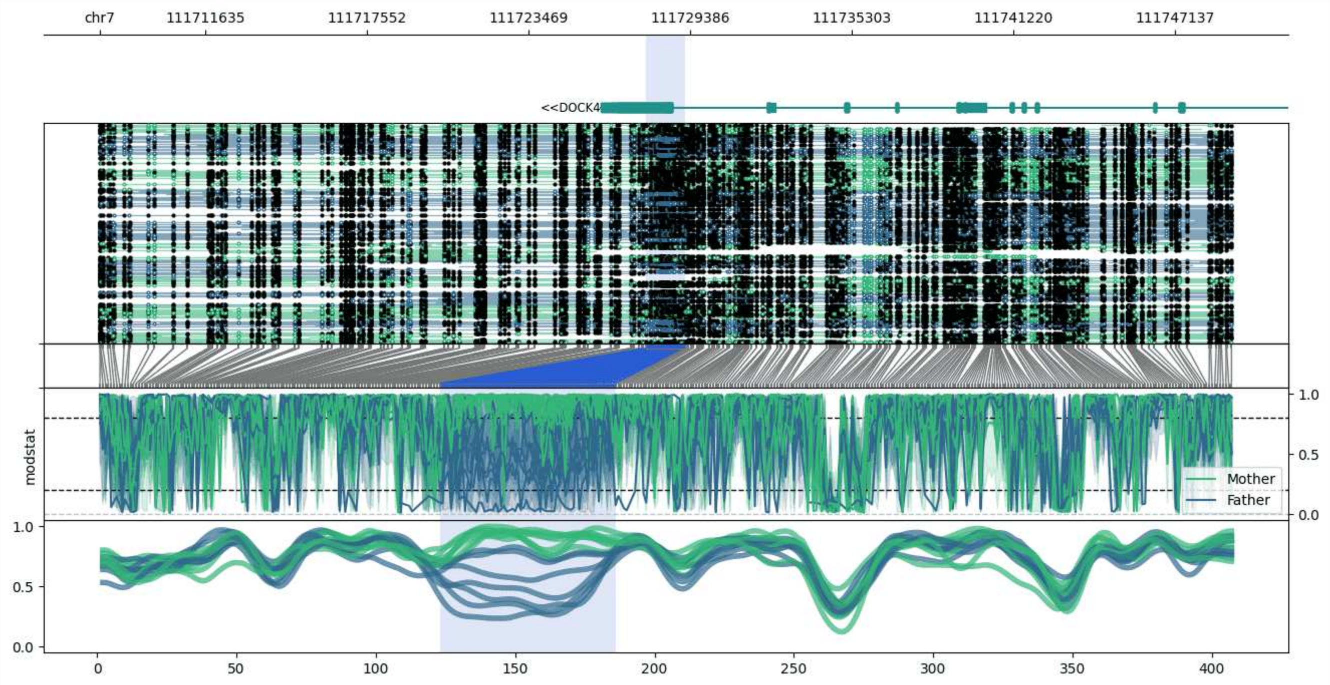


**chr7:111707781-111749204:** Region with 1 DMR containing genes: AC003077.1, DOCK4.

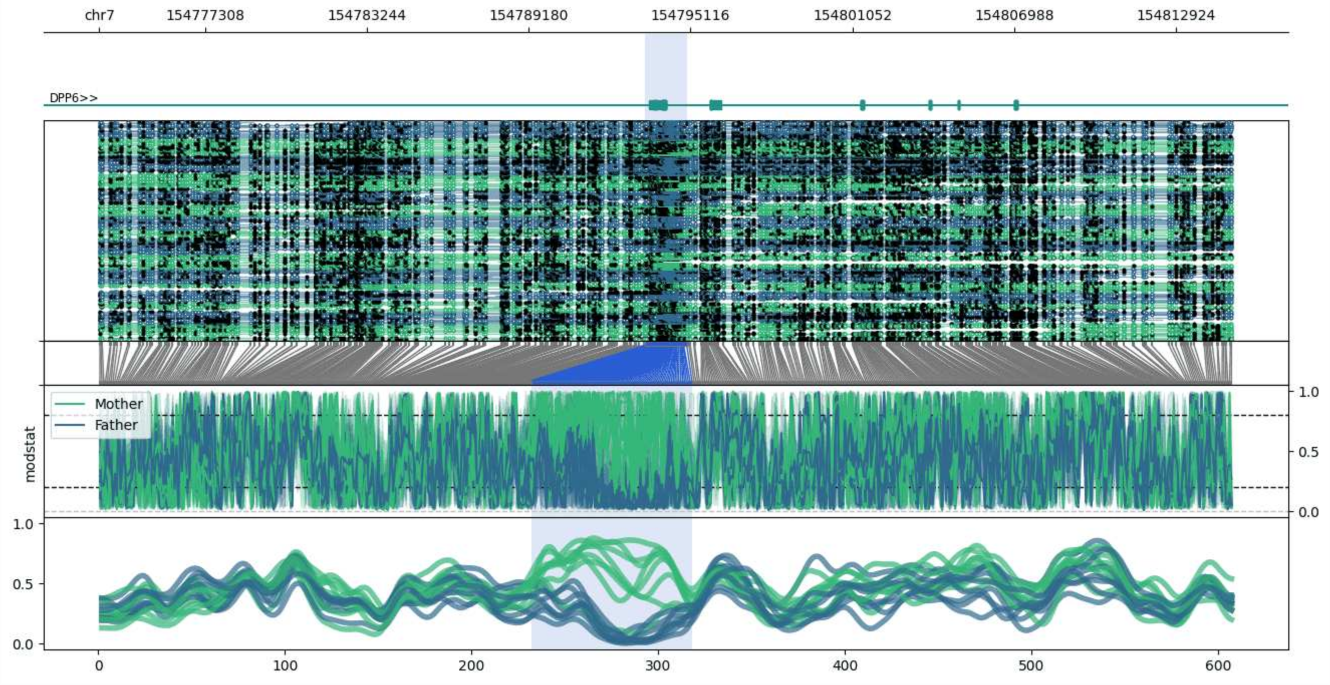


**chr7:154773425-154814979:** Region with 1 DMR containing genes: DPP6.

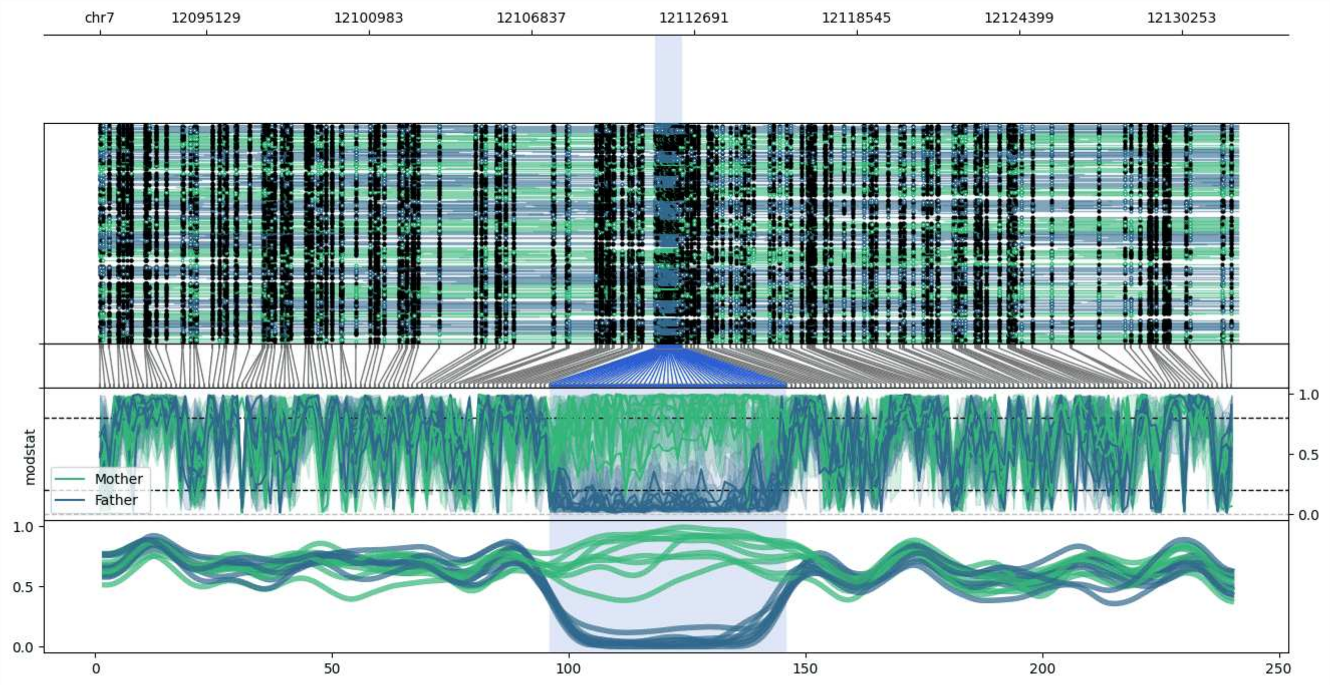


**chr7:12091300-12132280:** Region with 1 DMR containing no genes.

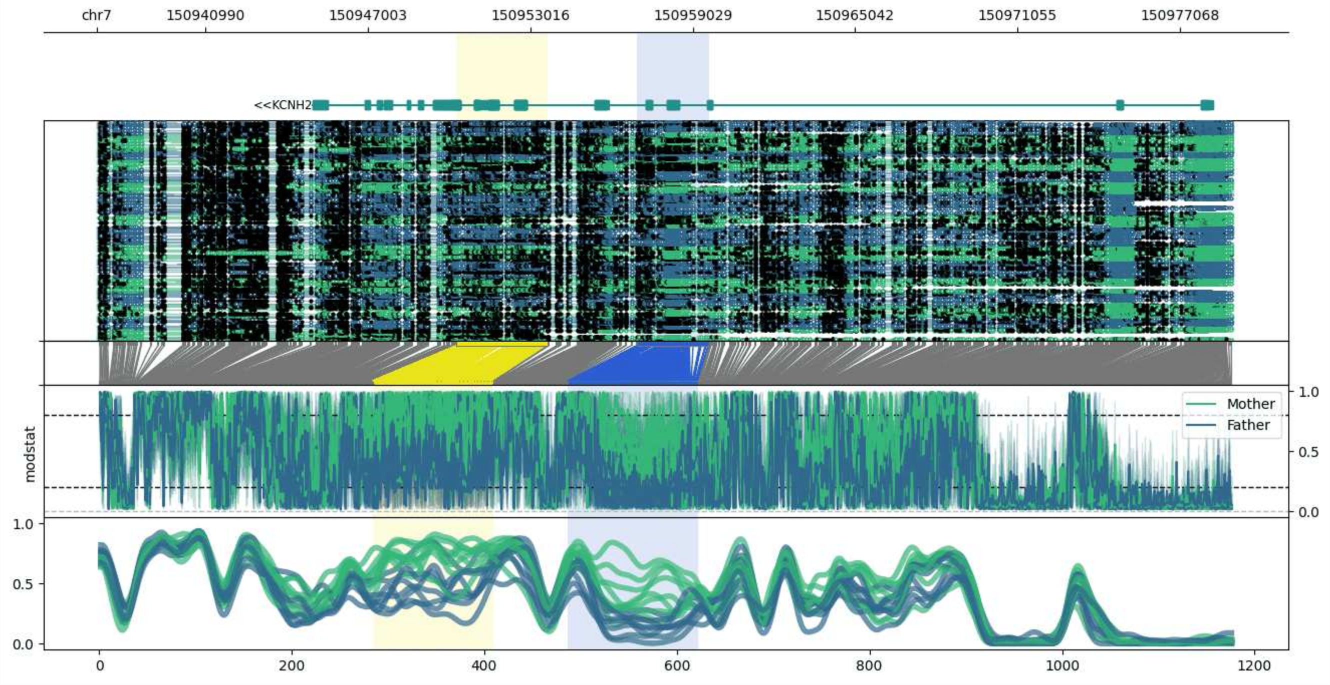


**chr7:150936948-150979041:** Region with 2 DMRs containing genes: KCNH2.

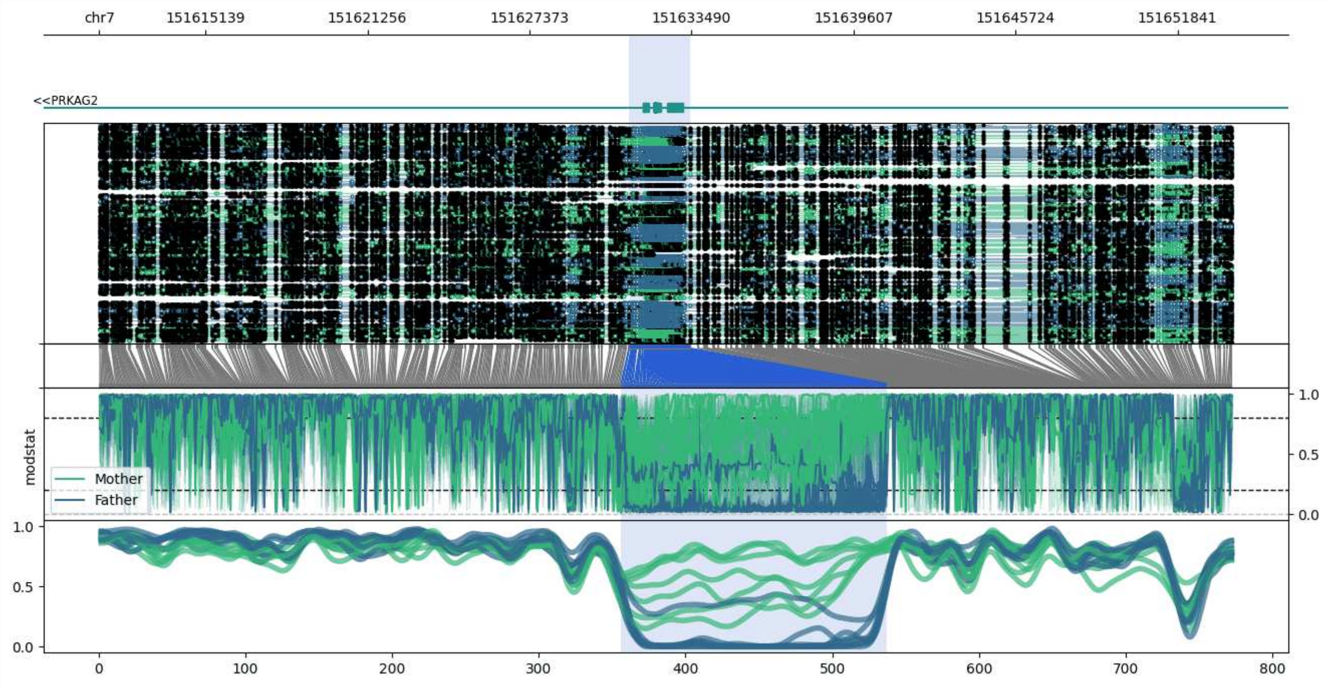


**chr7:151611130-151653952:** Region with 1 DMR containing genes: PRKAG2.

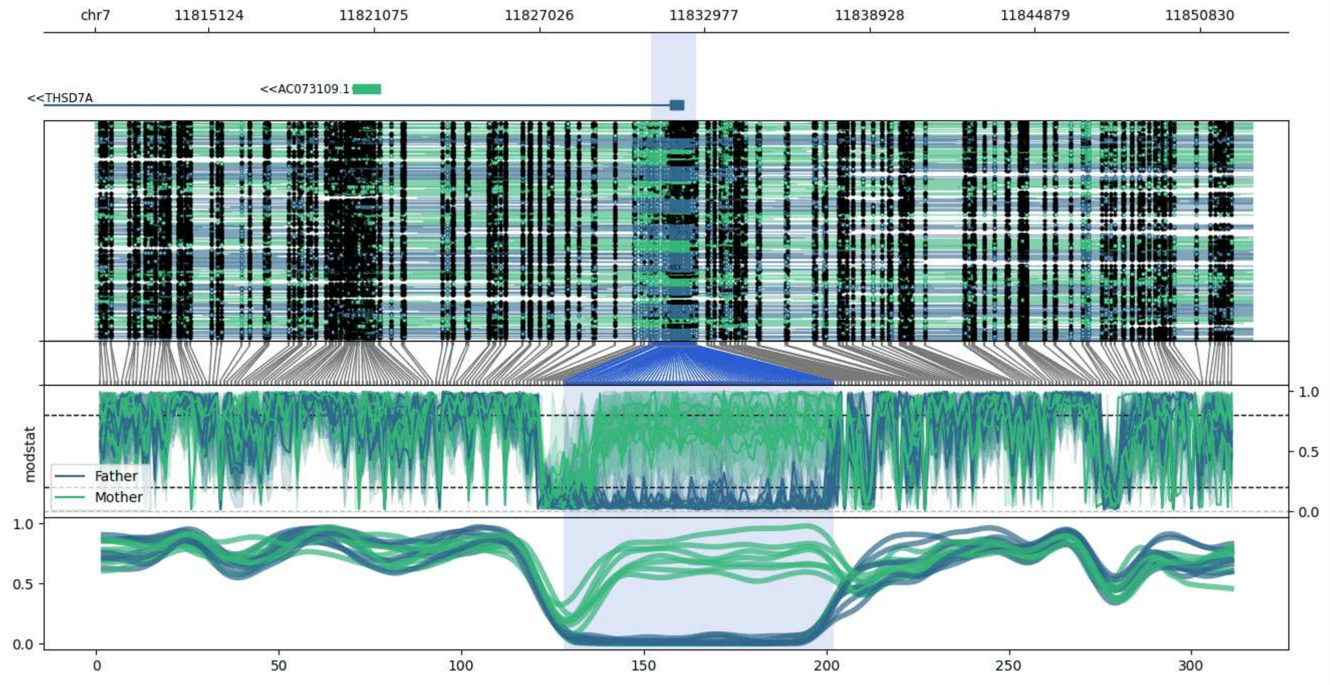


**chr7:11811037-11852699:** Region with 1 DMR containing genes: AC073109.1, THSD7A.

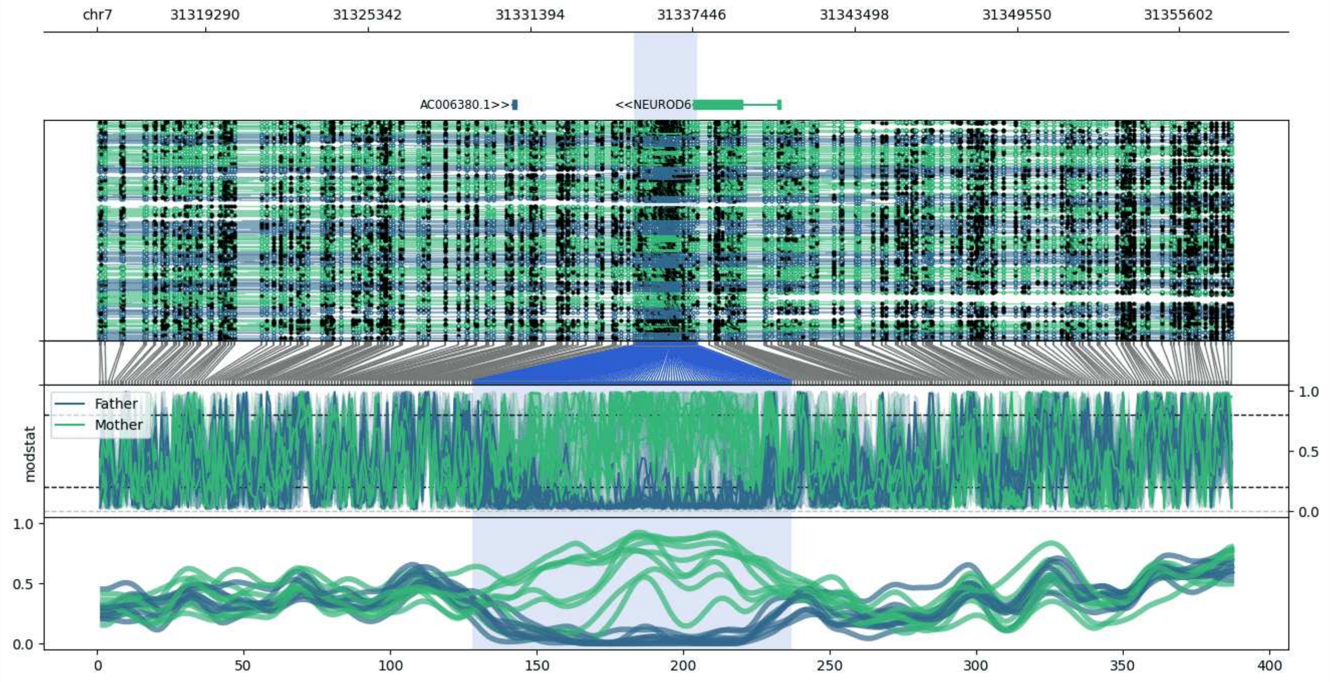


**chr7:31315247-31357617:** Region with 1 DMR containing genes: AC006380.1, NEUROD6.

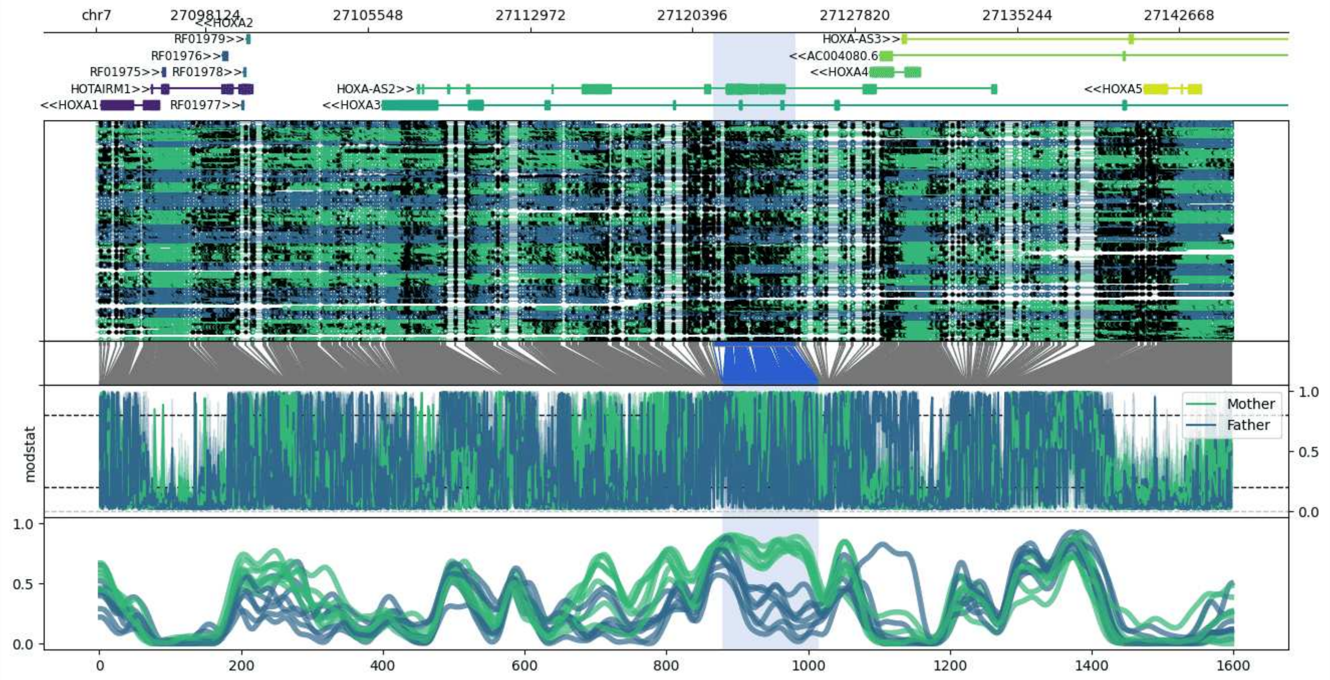


**chr7:27093118-27145089:** Region with 1 DMR containing genes: AC004080.5, AC004080.6, HOTAIRM1, HOXA-AS2, HOXA-AS3, HOXA1, HOXA2, HOXA3, HOXA4, HOXA5, HOXA6, HOXA7, RF01975, RF01976, RF01977, RF01978, RF01979.

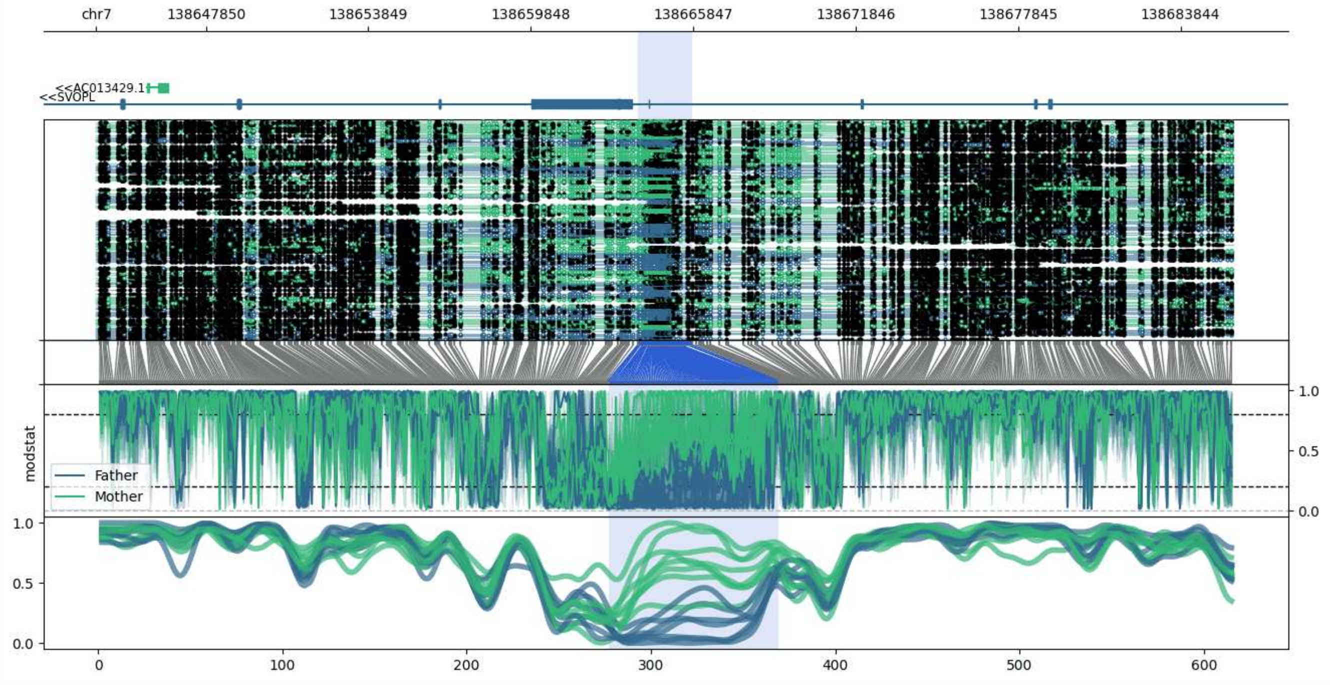


**chr7:138643806-138685803:** Region with 1 DMR containing genes: AC013429.1, AC020983.1, SVOPL.

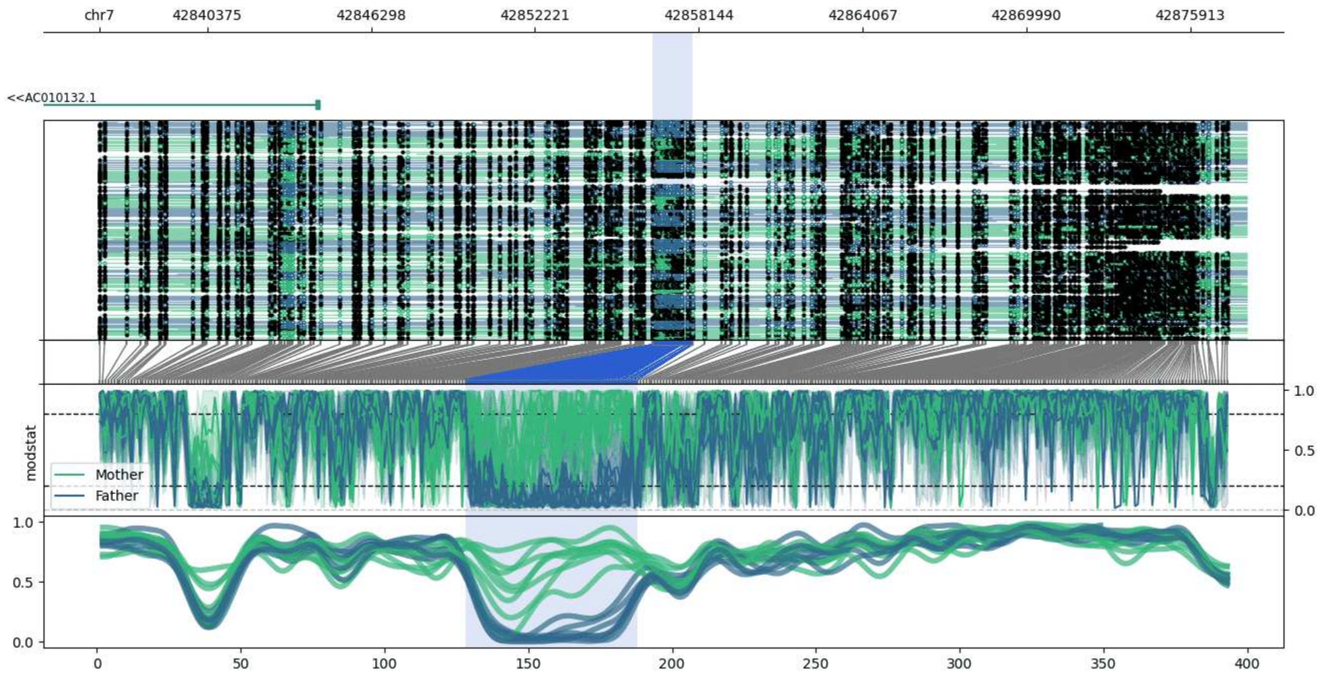


**chr7:42836469-42877930:** Region with 1 DMR containing genes: AC010132.1.

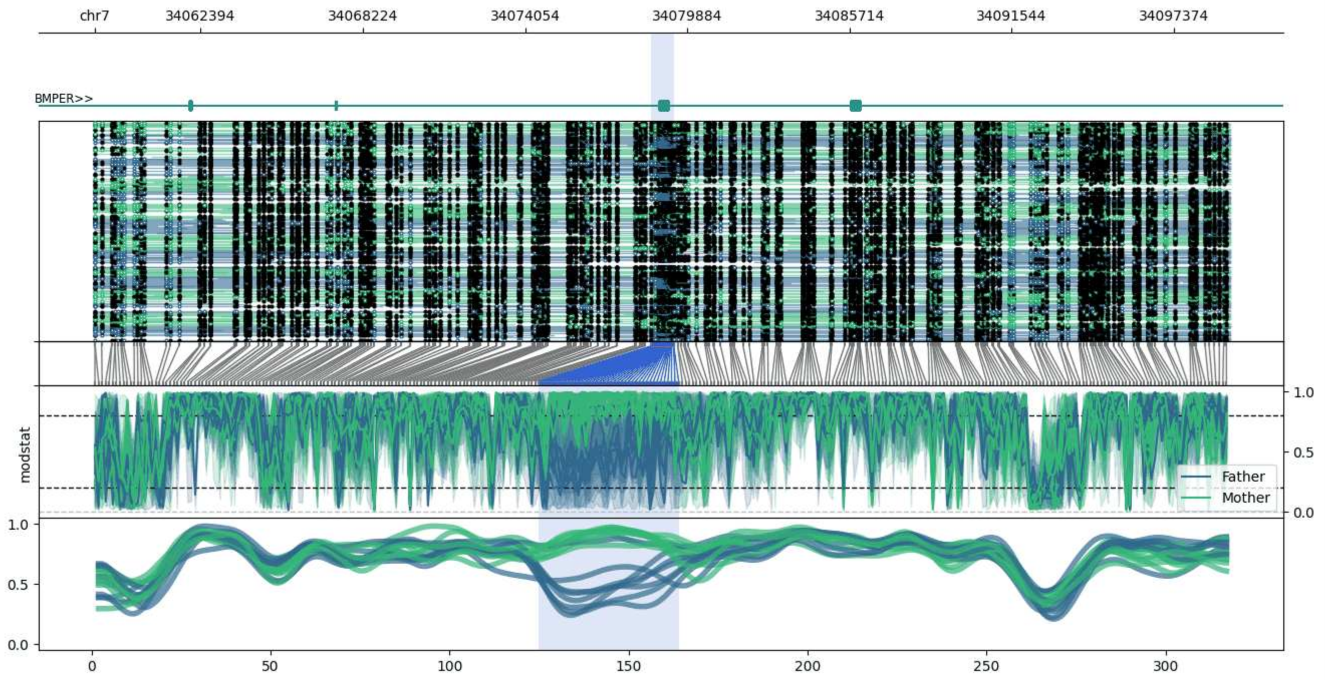


**chr7:34058580-34099396:** Region with 1 DMR containing genes: BMPER.

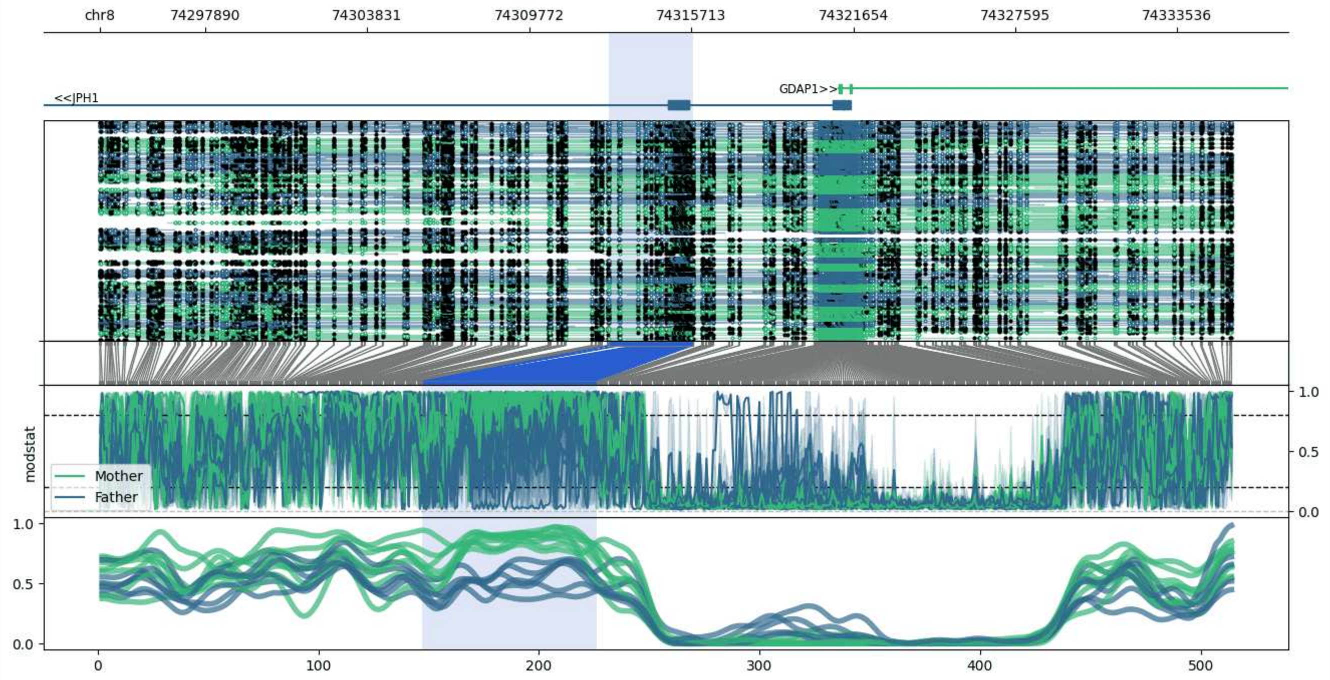


**chr8:74294006-74335597:** Region with 1 DMR containing genes: GDAP1, JPH1.

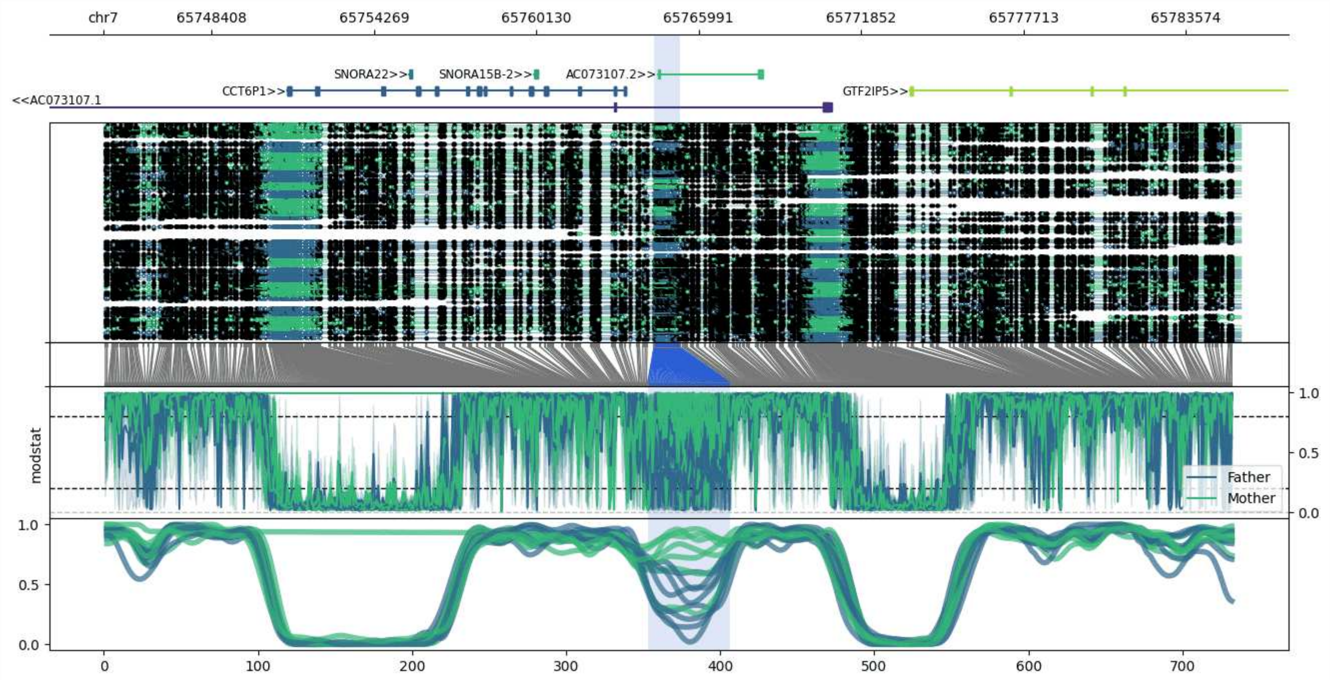


**chr7:65744512-65785539:** Region with 1 DMR containing genes: AC073107.1, AC073107.2, CCT6P1, GTF2IP5, SNORA15B-2, SNORA22.

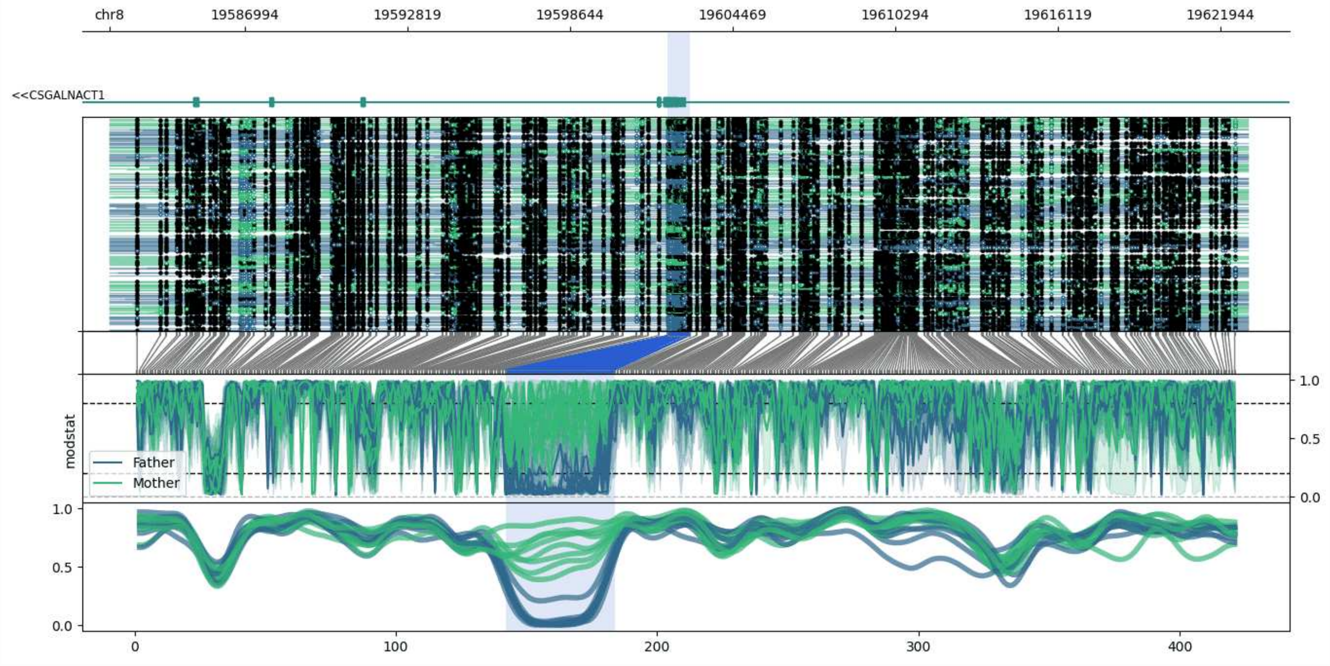


**chr8:19582154-19622931:** Region with 1 DMR containing genes: CSGALNACT1.

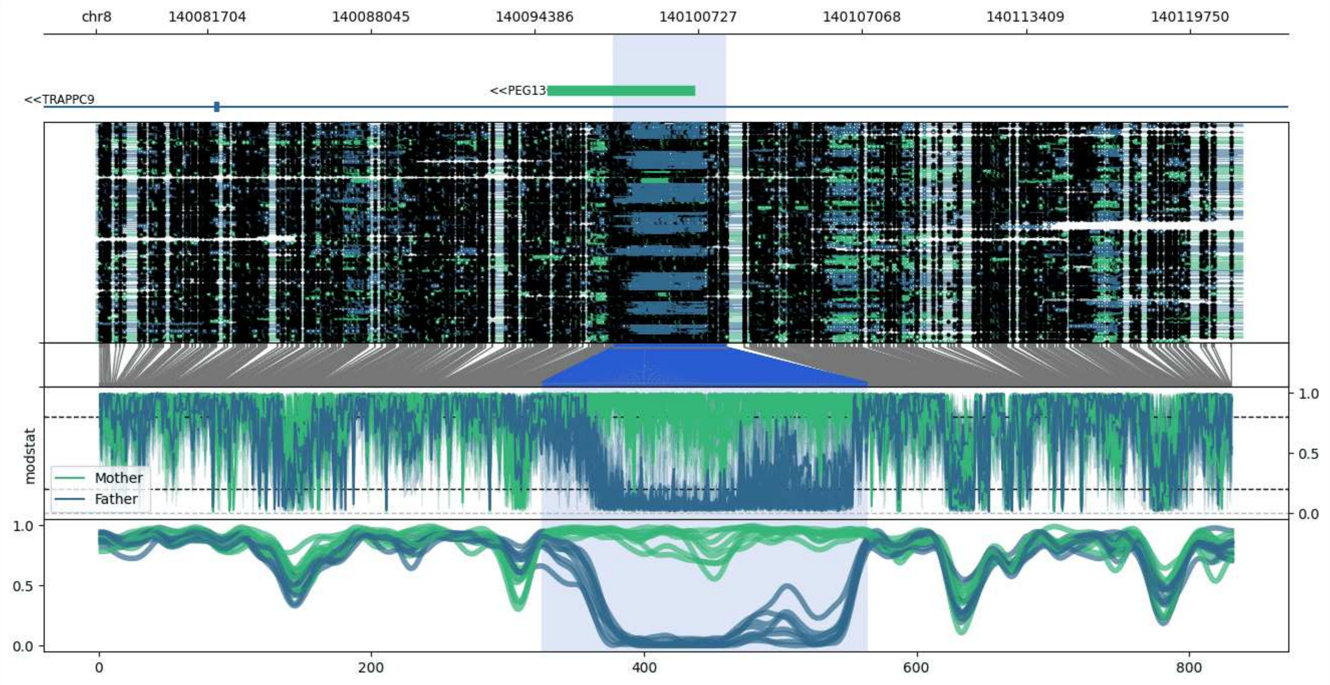


**chr8:140077405-140121793:** Region with 1 DMR containing genes: PEG13, TRAPPC9.

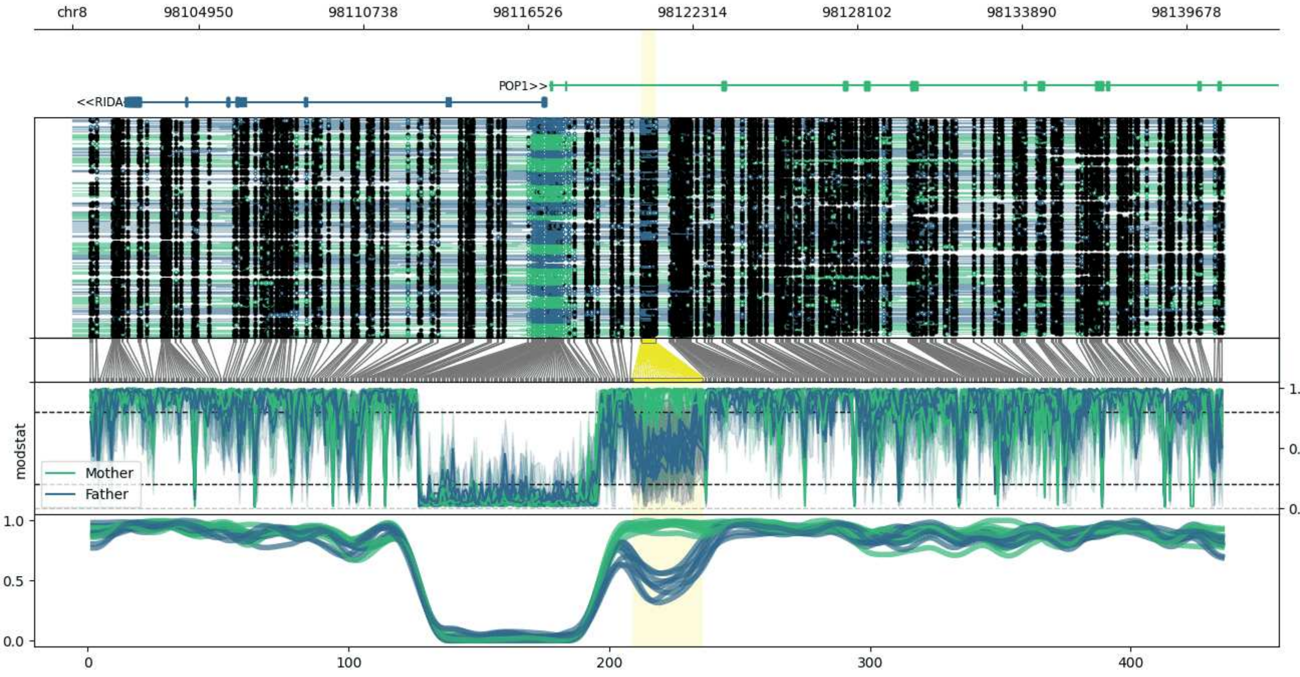


**chr8:98100509-98141028:** Region with 1 DMR containing genes: ERICH5, POP1, RIDA.

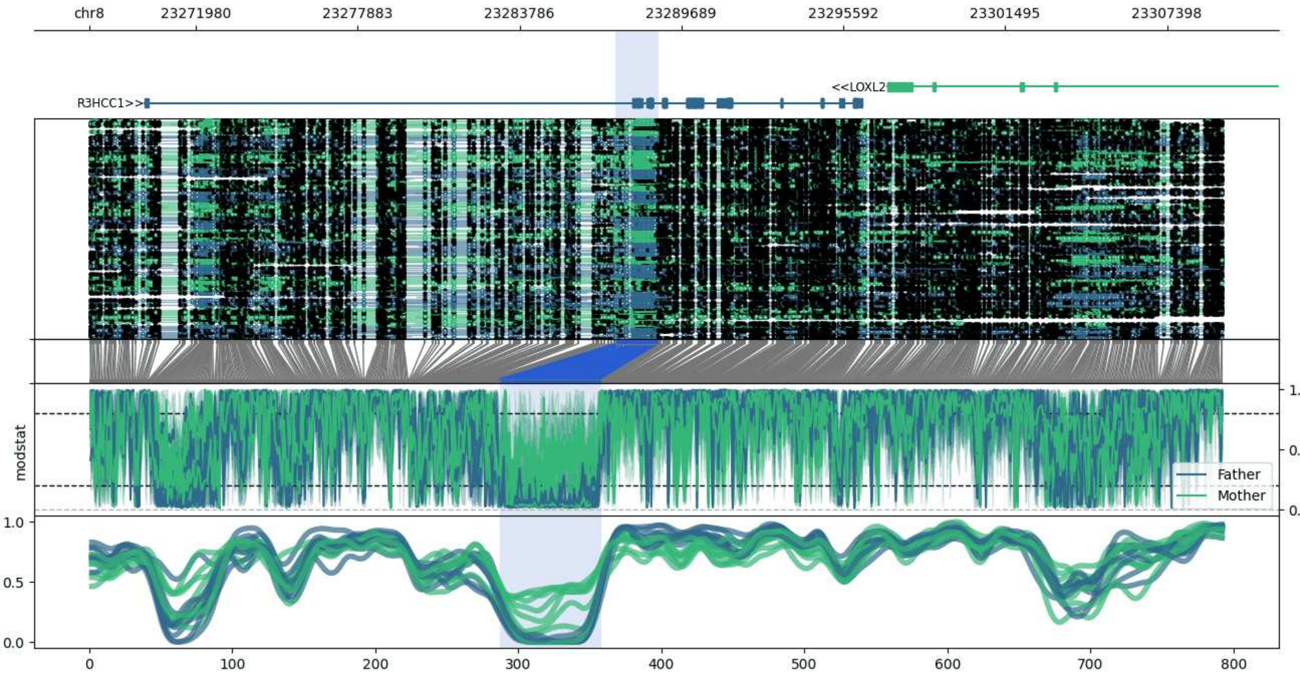


**chr8:23268097-23309418:** Region with 1 DMR containing genes: CHMP7, LOXL2, R3HCC1.

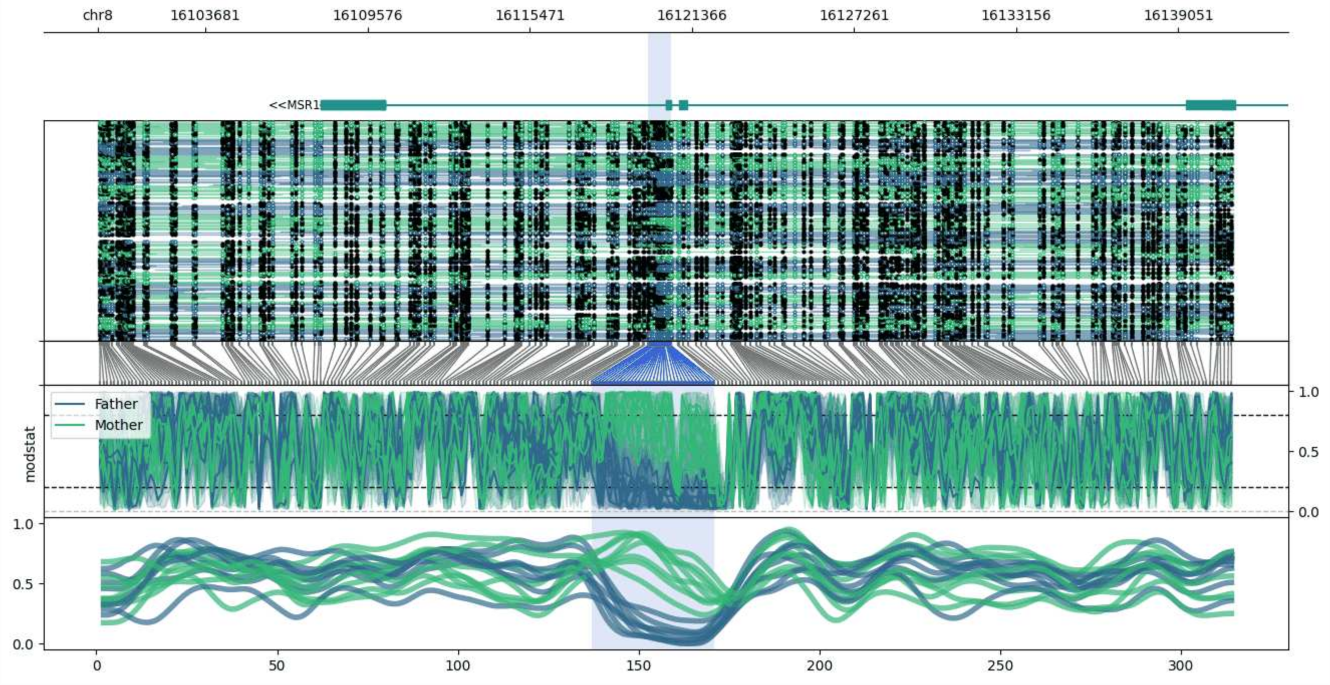


**chr8:16099768-16141037:** Region with 1 DMR containing genes: MSR1.

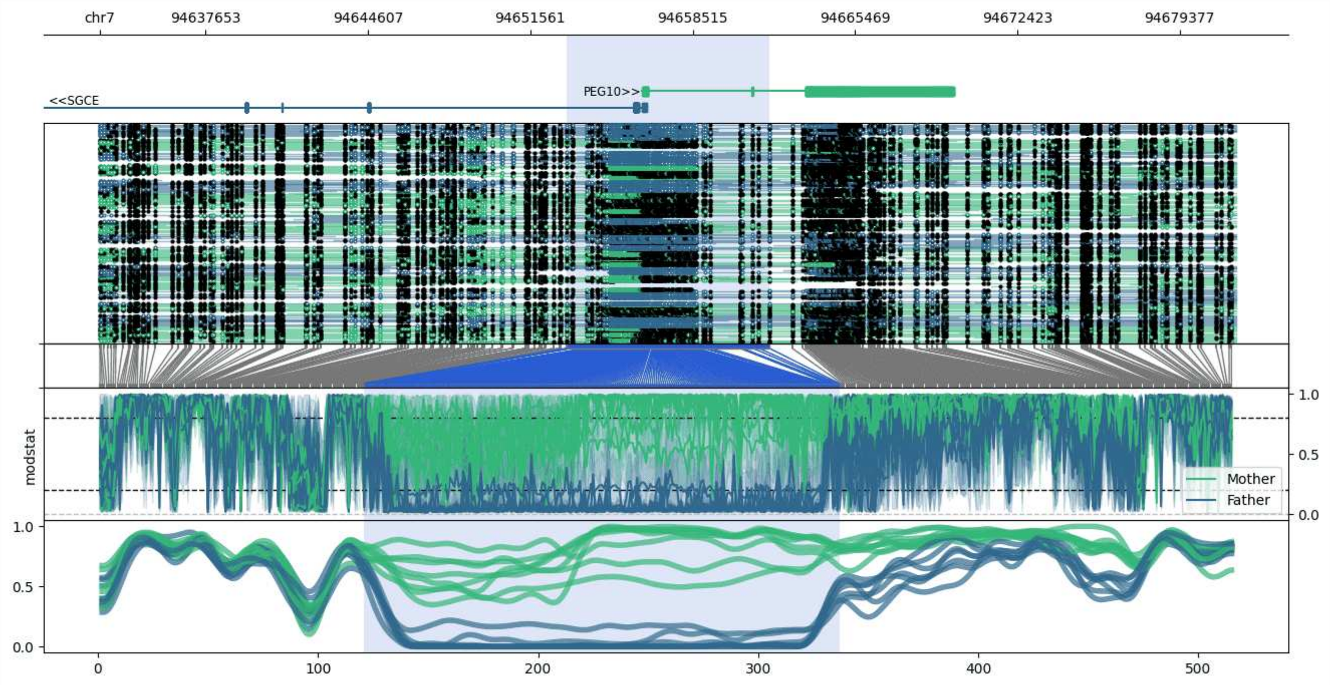


**chr7:94633112-94681791:** Region with 1 DMR containing genes: PEG10, SGCE.

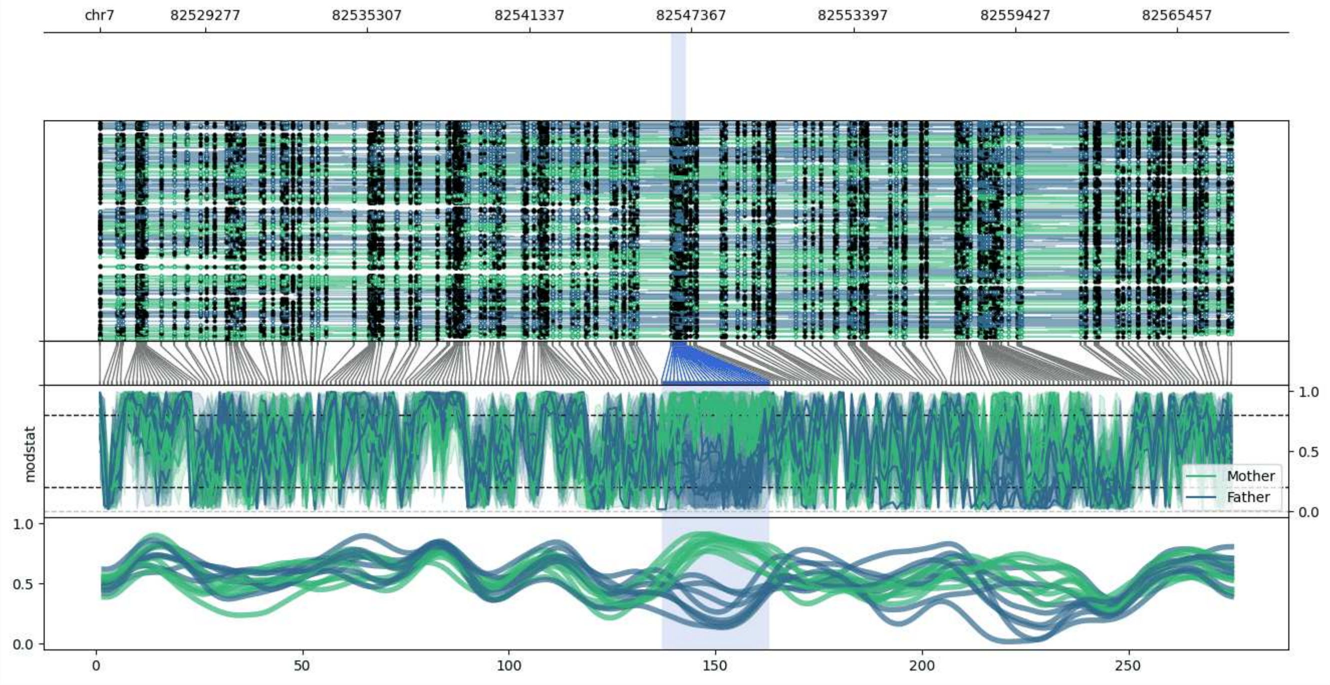


**chr7:82525340-82567552:** Region with 1 DMR containing no genes.

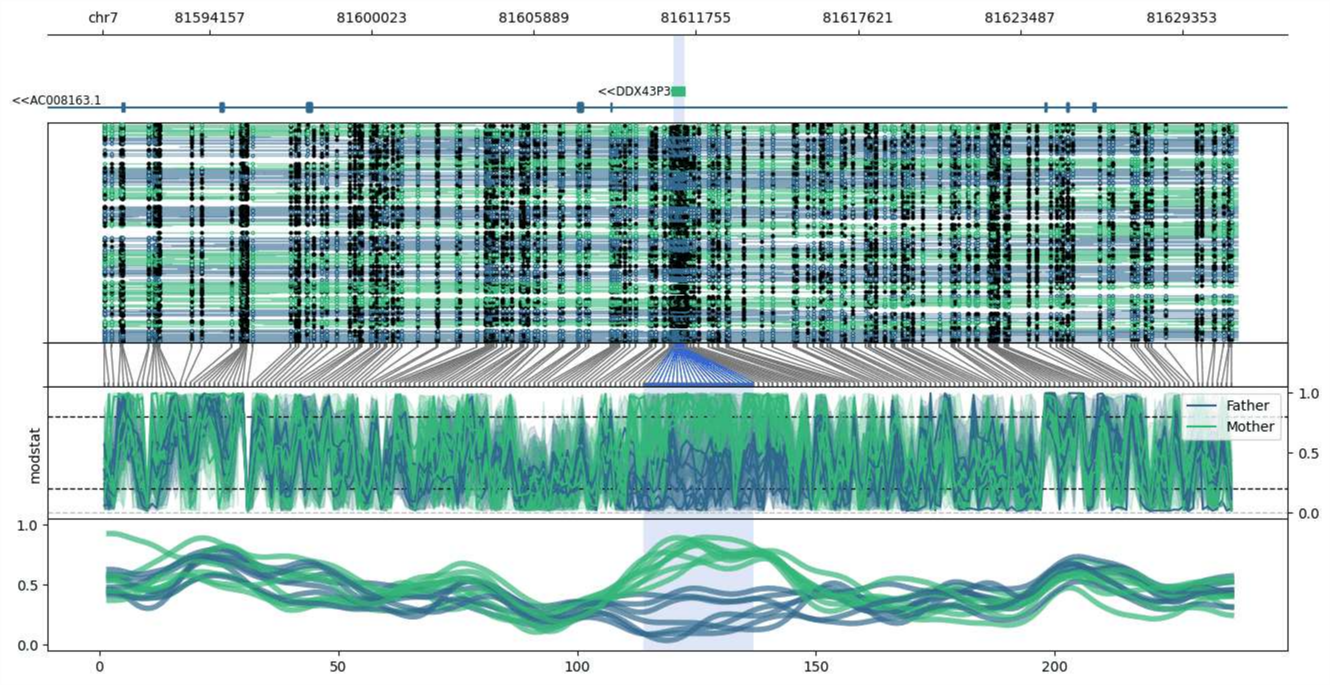


**chr7:81590287-81631351:** Region with 1 DMR containing genes: AC008163.1, DDX43P3.

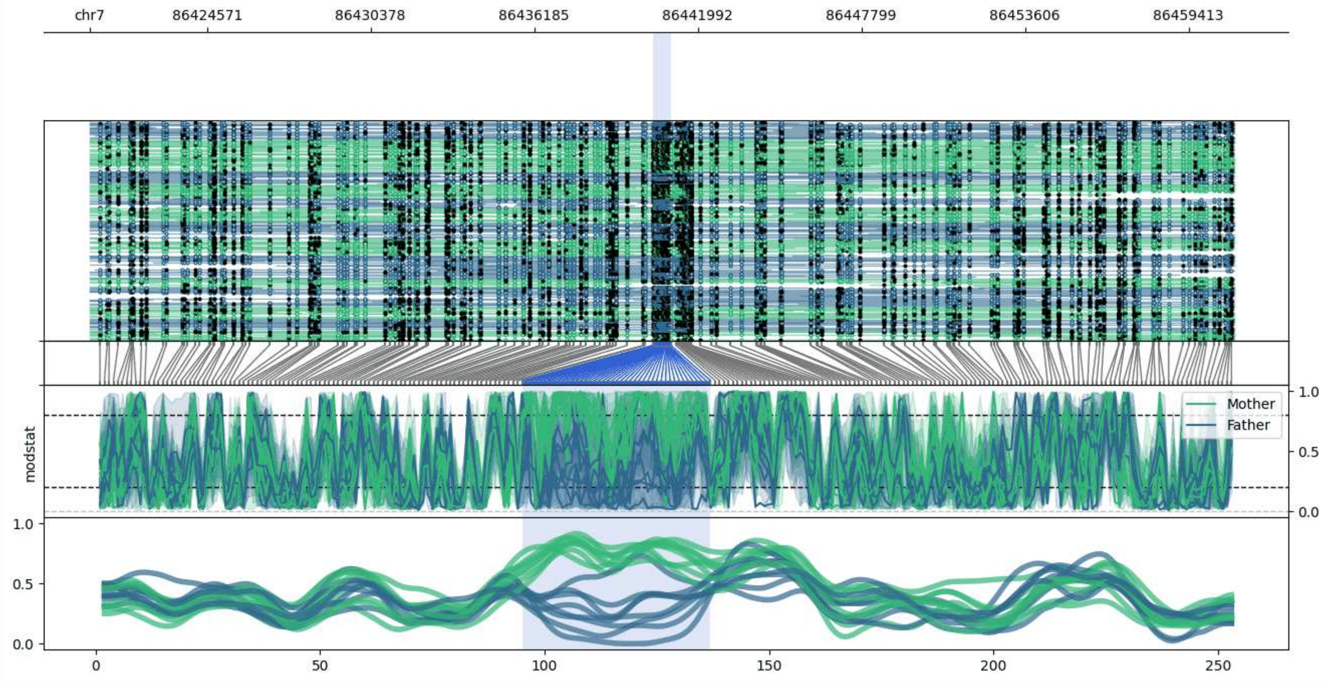


**chr7:86420398-86461047:** Region with 1 DMR containing no genes.

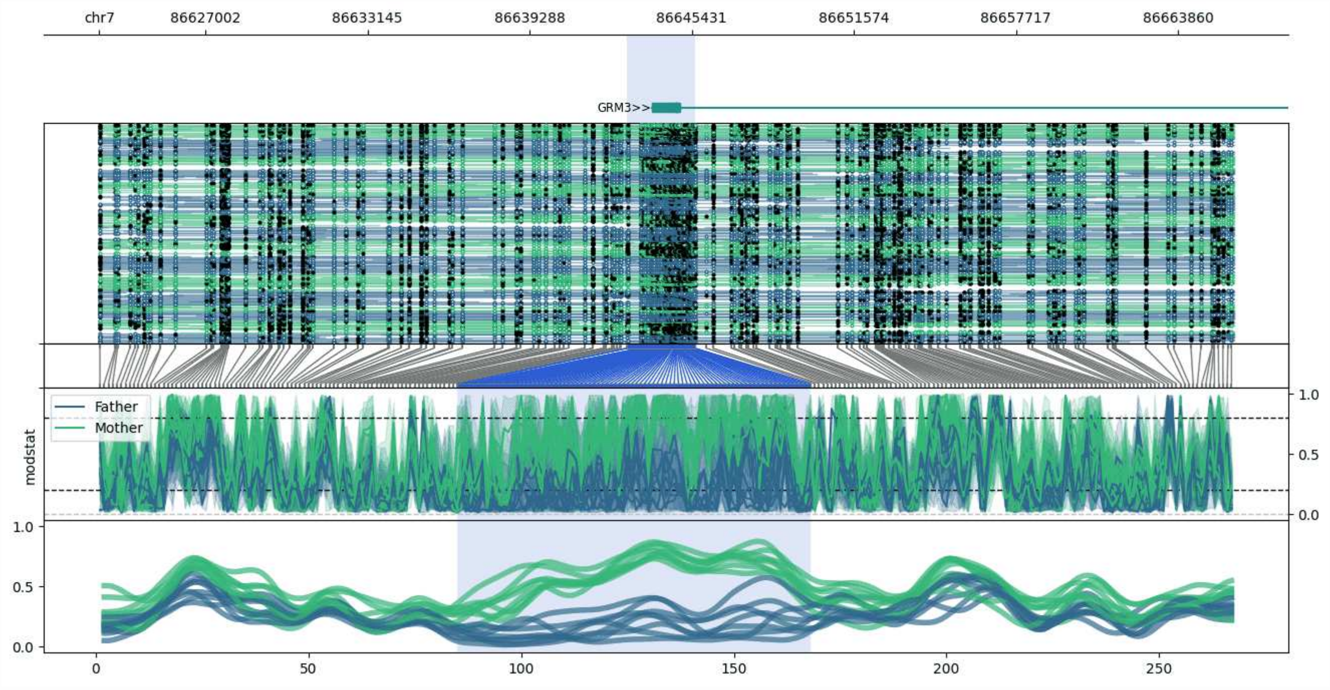


**chr7:86622977-86665984:** Region with 1 DMR containing genes: GRM3.

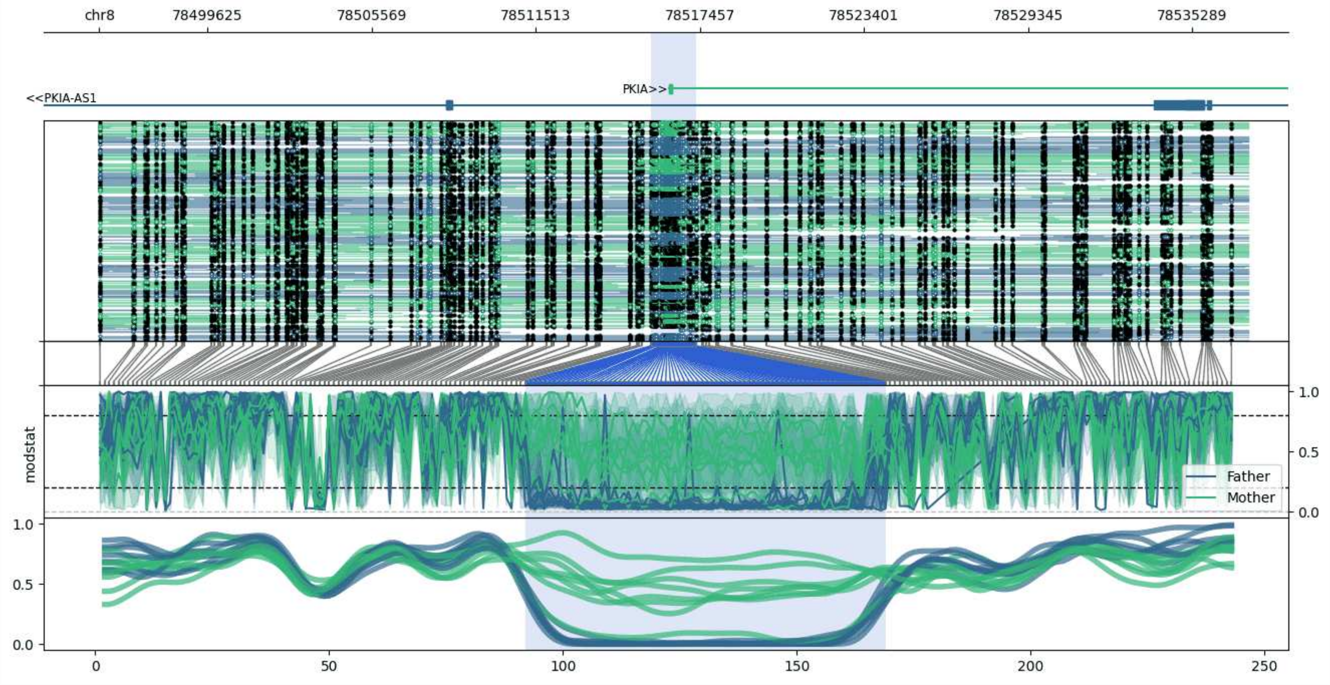


**chr8:78495701-78537314:** Region with 1 DMR containing genes: PKIA, PKIA-AS1.

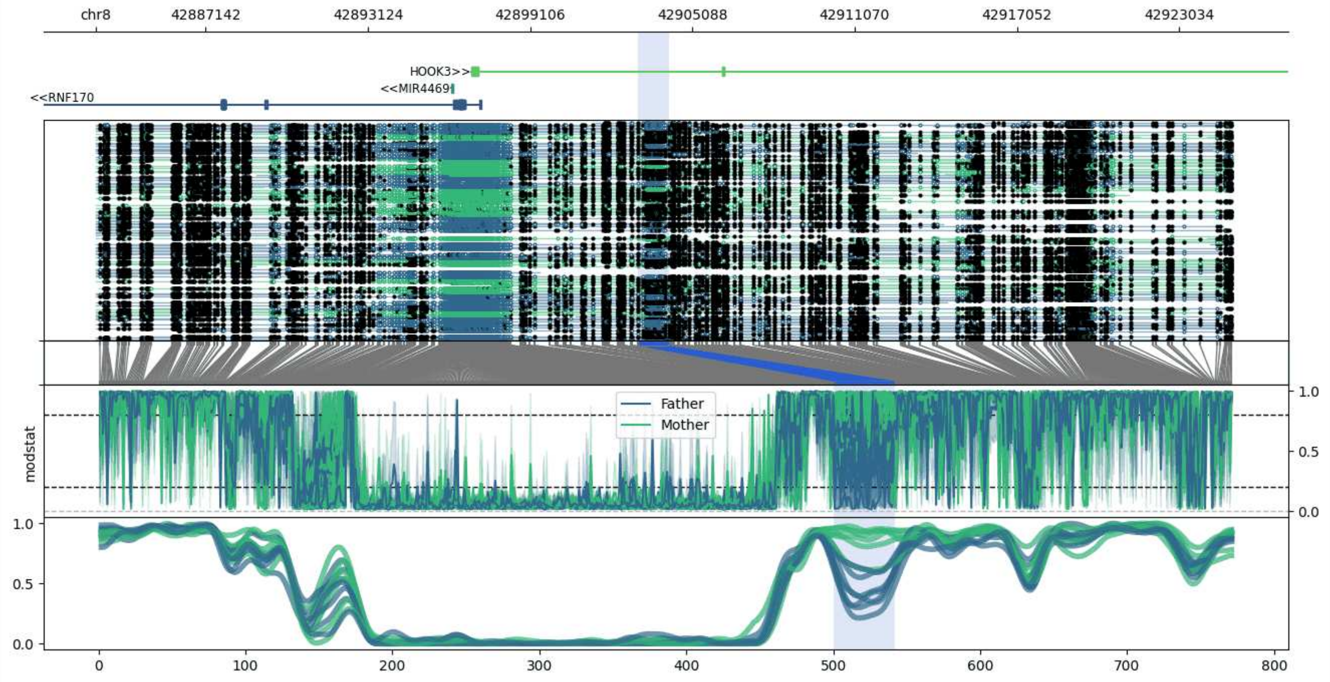


**chr8:42883084-42924964:** Region with 1 DMR containing genes: HOOK3, MIR4469, RN7SL806P, RNF170.

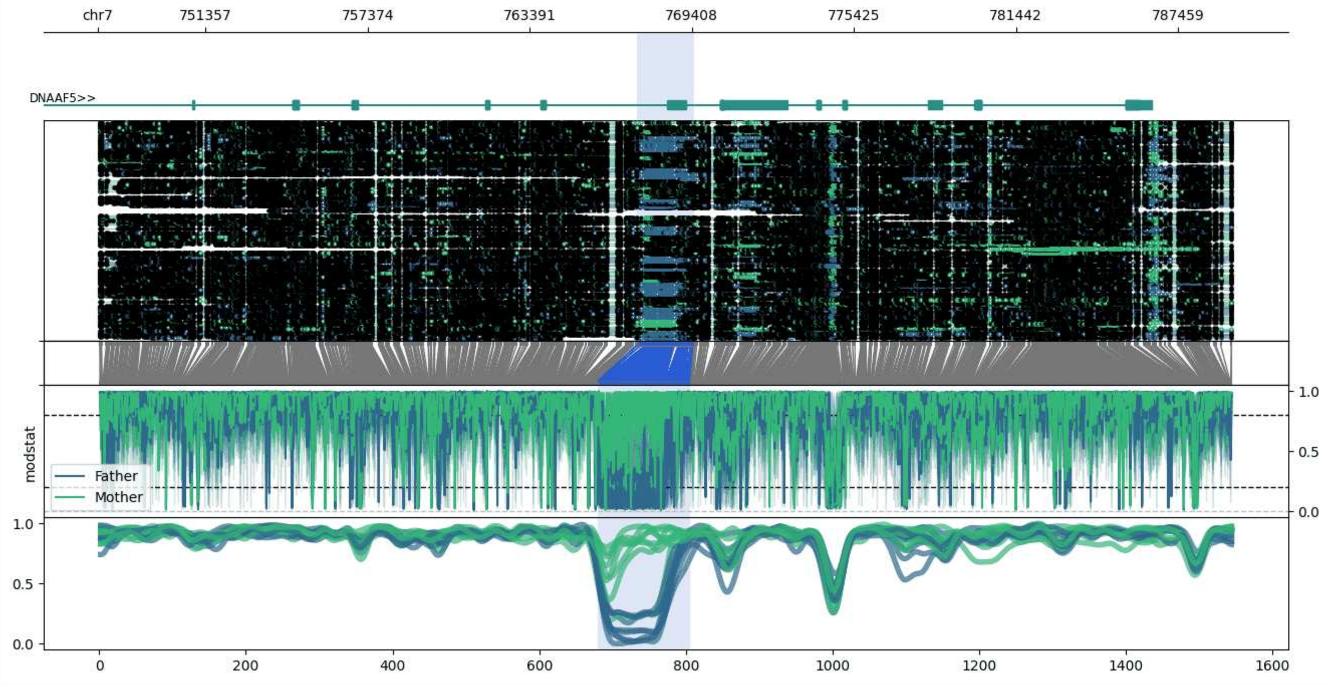


**chr7:747358-789480:** Region with 1 DMR containing genes: DNAAF5.

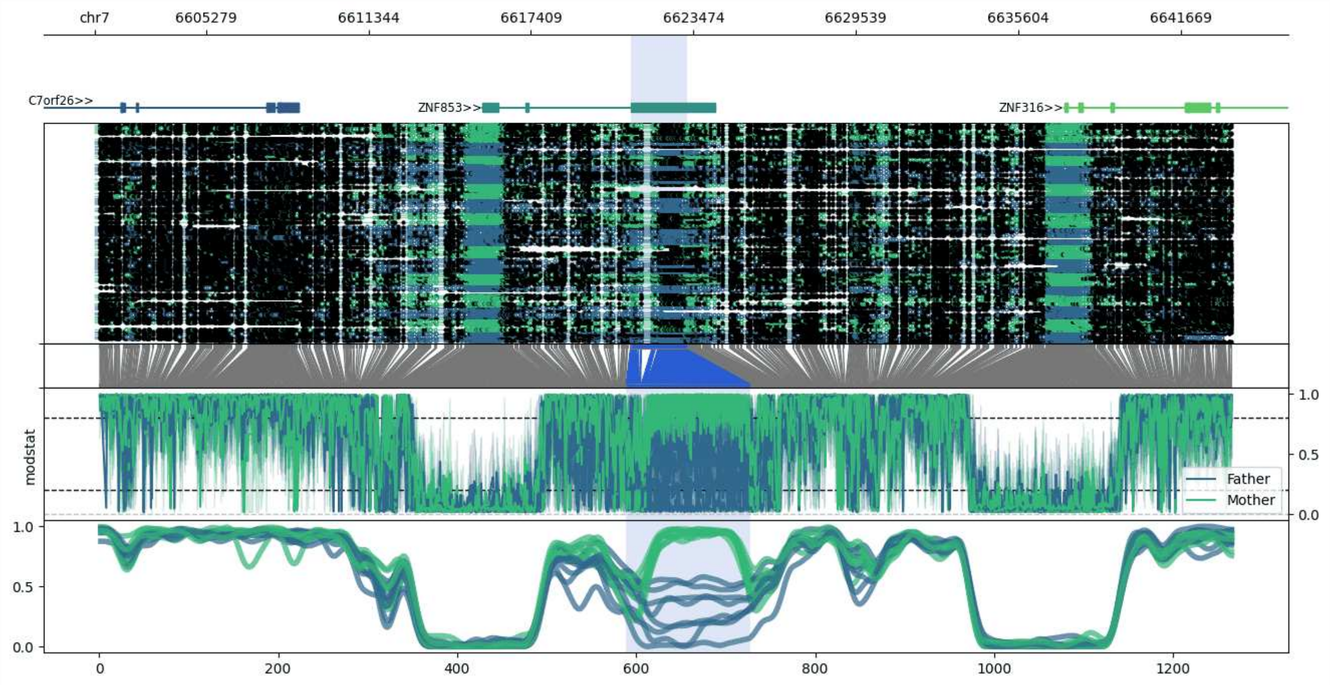


**chr7:6601128-6643584:** Region with 1 DMR containing genes: C7orf26, ZNF316, ZNF853.

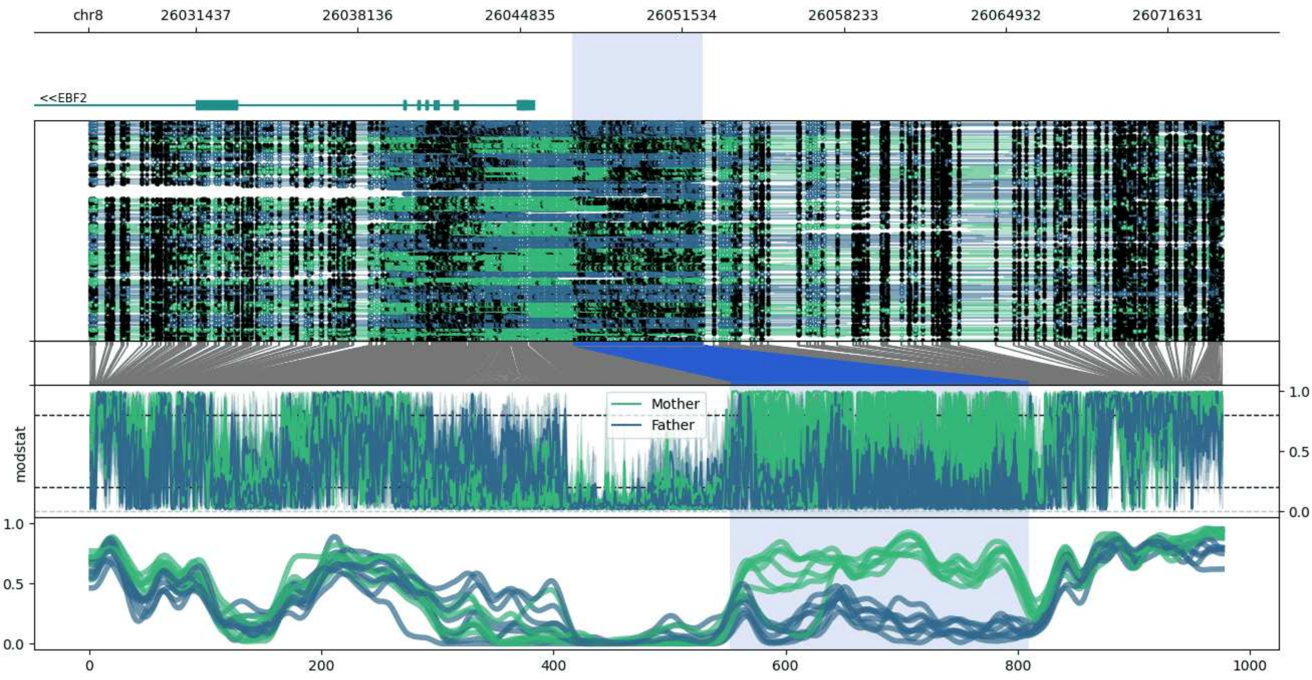


**chr8:26026994-26073891:** Region with 1 DMR containing genes: EBF2.

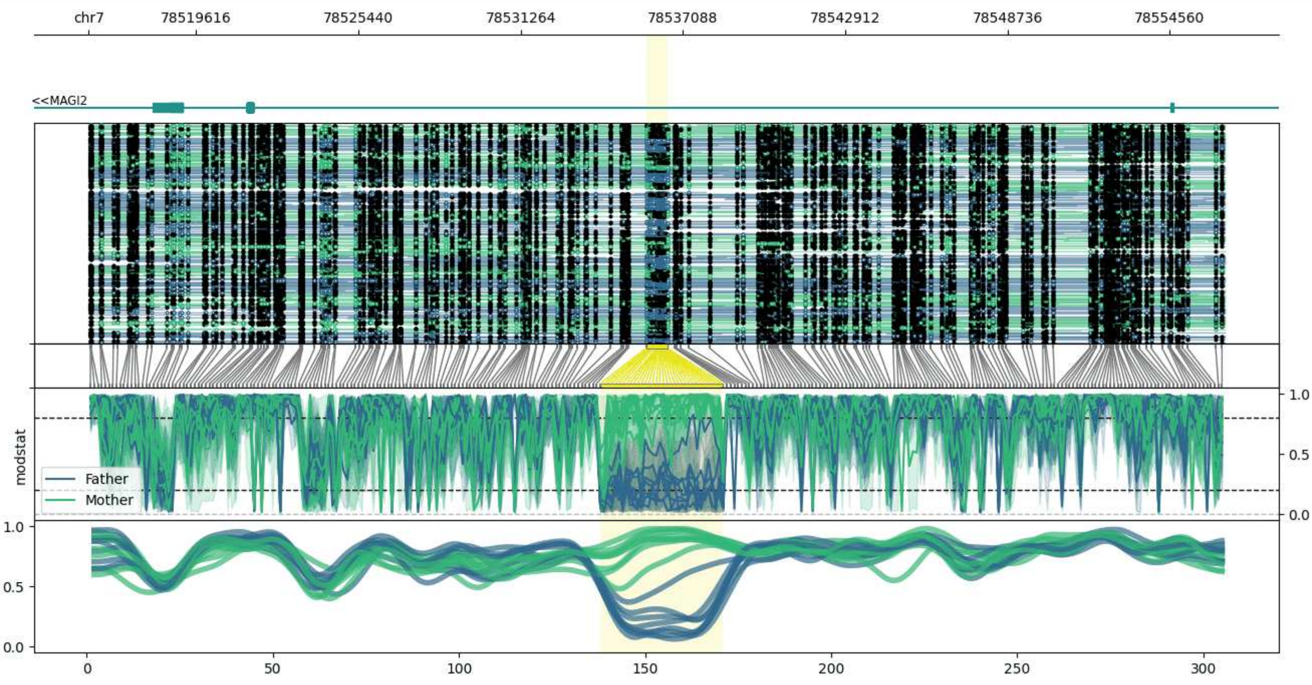


**chr7:78515763-78556533:** Region with 1 DMR containing genes: MAGI2.

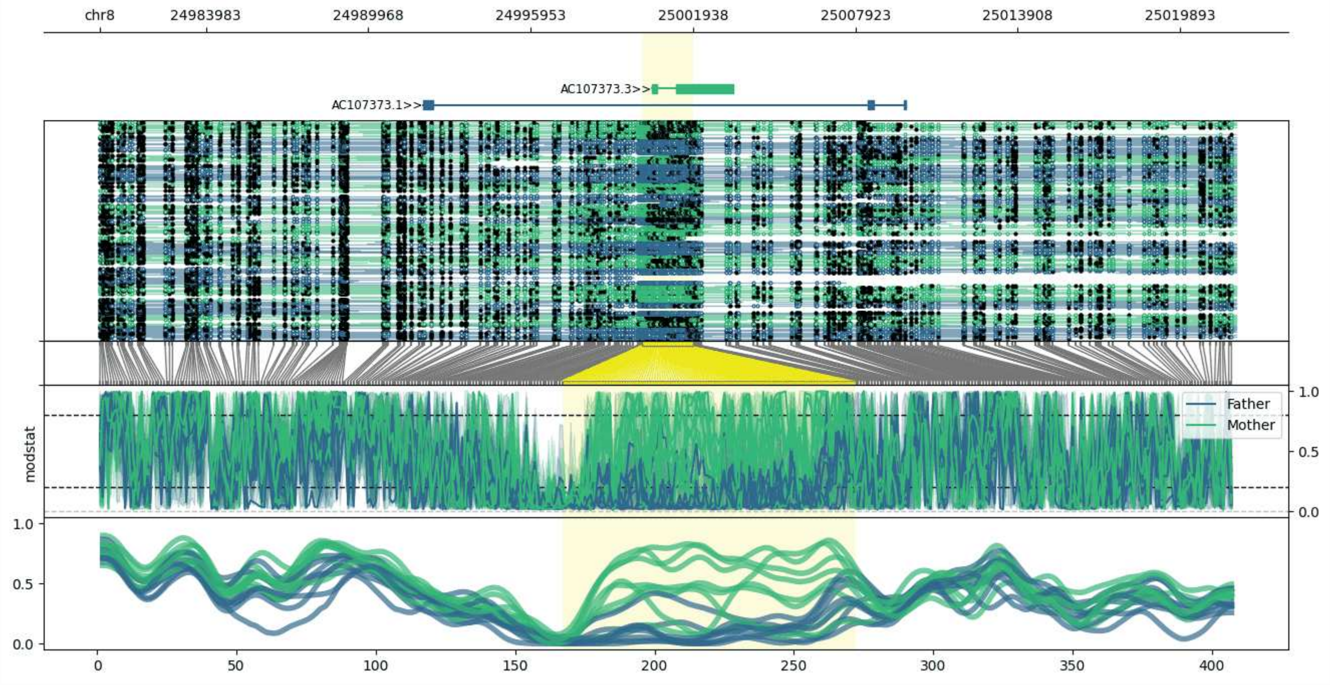


**chr8:24980063-25021963:** Region with 1 DMR containing genes: AC107373.1, AC107373.3.

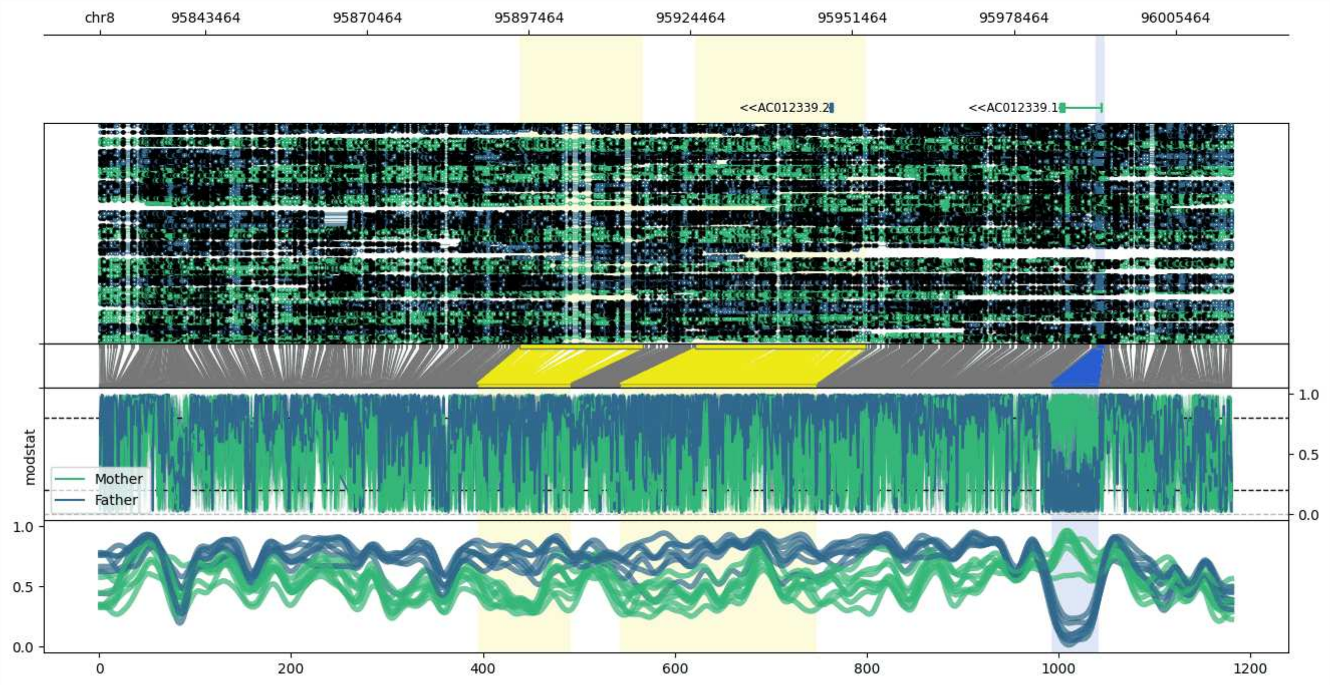


**chr8:95825862-96014865:** Region with 3 DMRs containing genes: AC012339.1, AC012339.2.

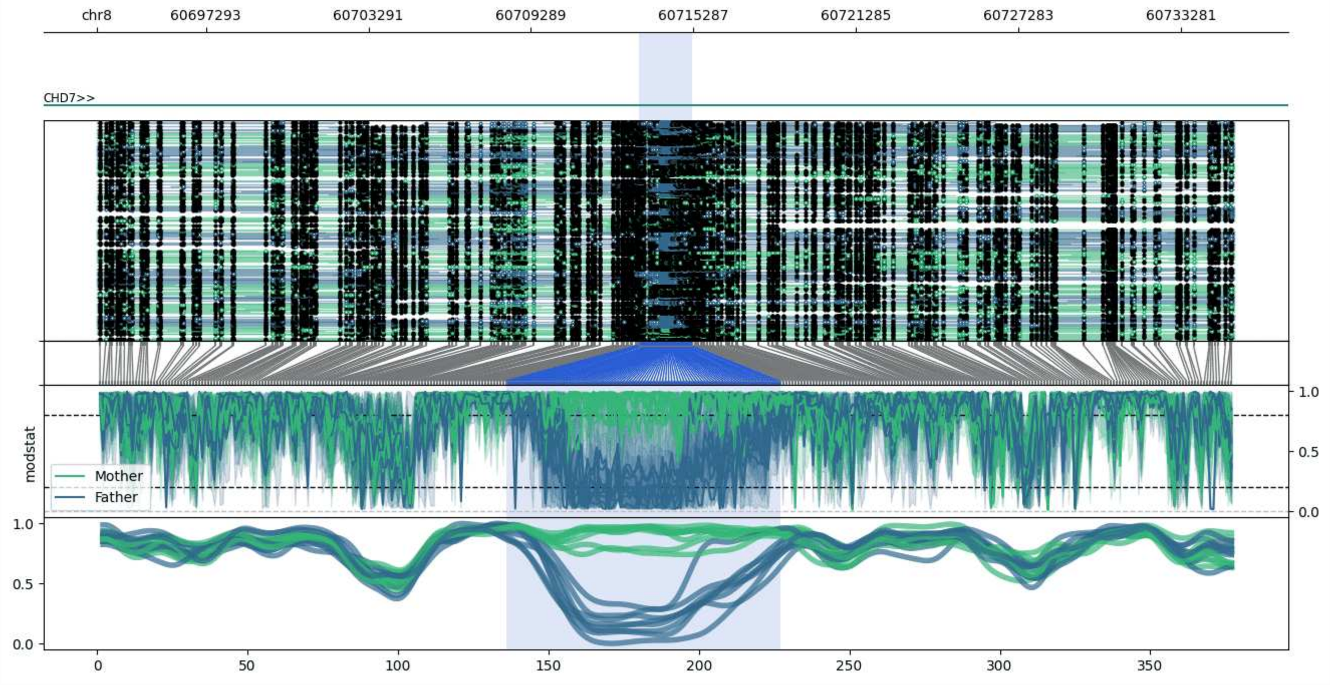


**chr8:60693255-60735241:** Region with 1 DMR containing genes: CHD7.

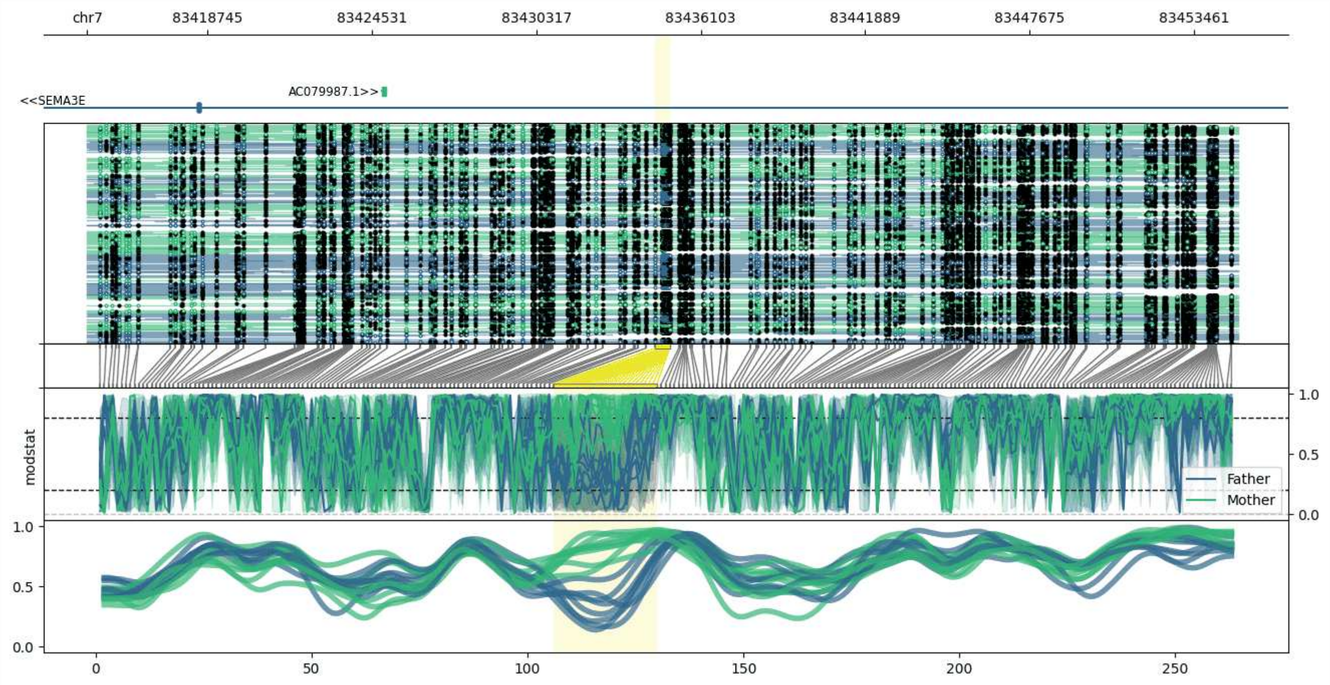


**chr7:83414505-83455007:** Region with 1 DMR containing genes: AC079987.1, SEMA3E.

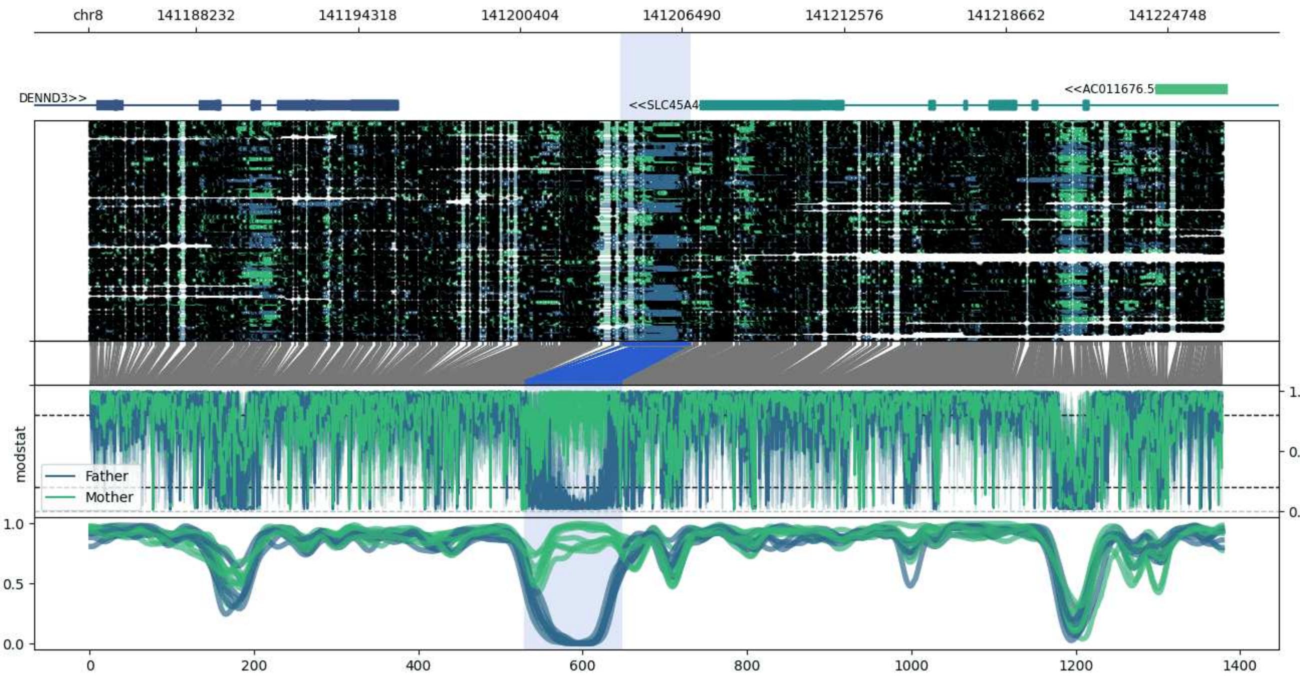


**chr8:141184196-141226799:** Region with 1 DMR containing genes: AC011676.5, DENND3, SLC45A4.

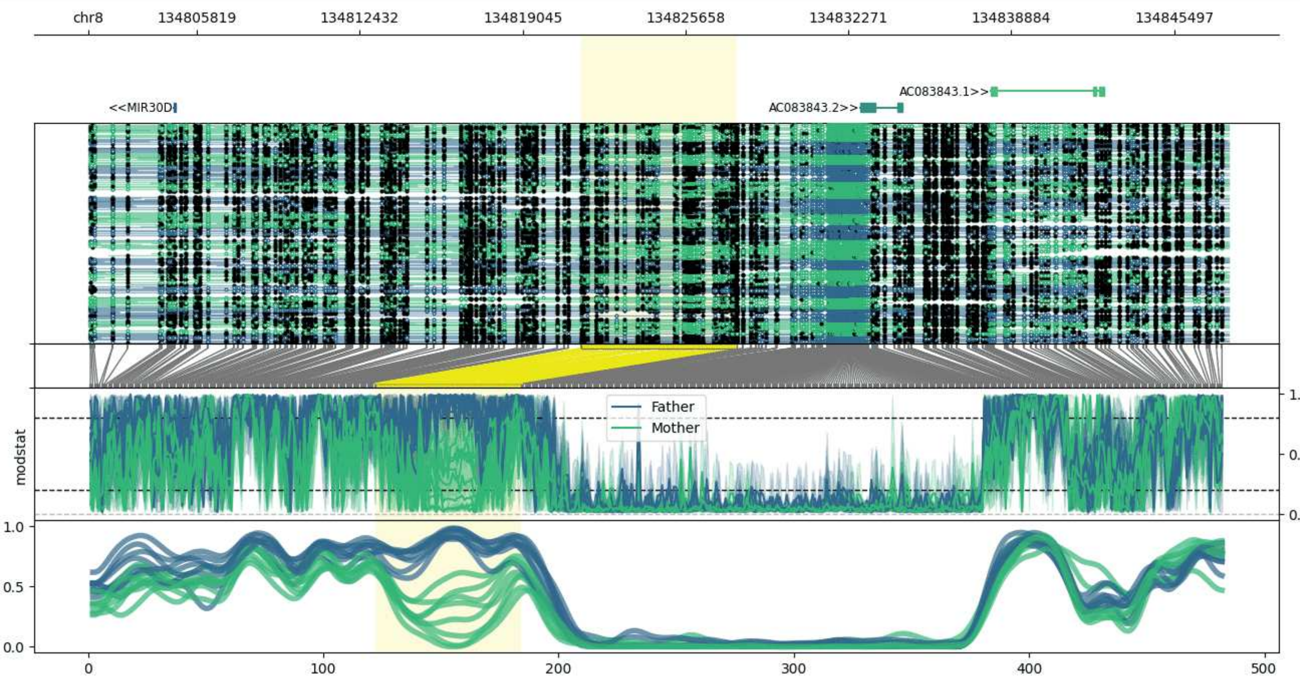


**chr8:134801427-134847723:** Region with 1 DMR containing genes: AC083843.1, AC083843.2, AC083843.3, AC103764.1, MIR30B, MIR30D.

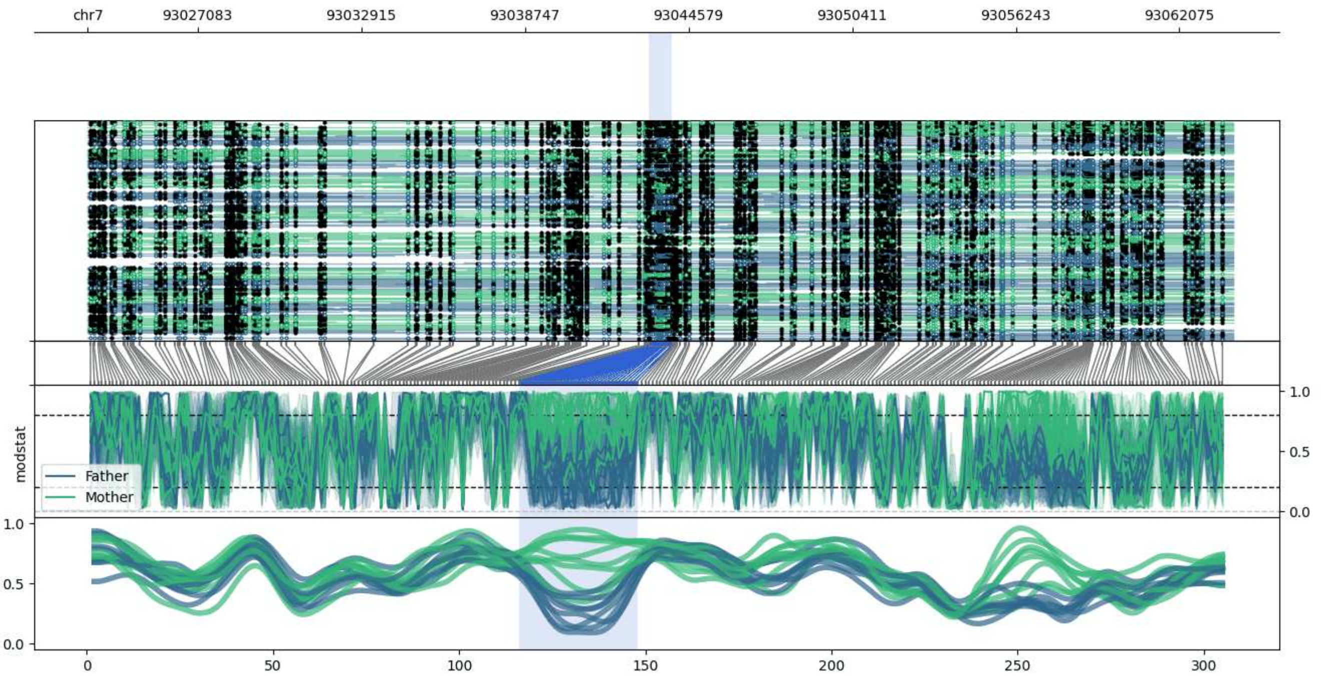


**chr7:93023169-93063999:** Region with 1 DMR containing no genes.

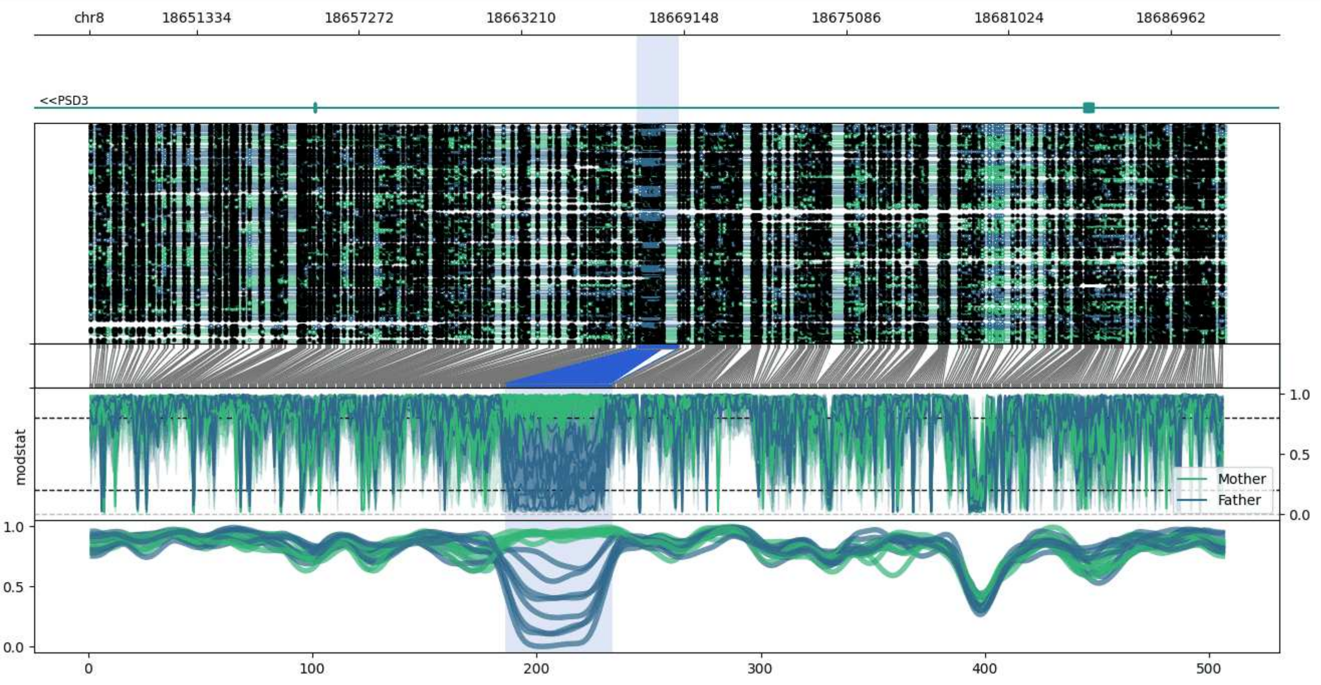


**chr8:18647417-18688987:** Region with 1 DMR containing genes: PSD3.

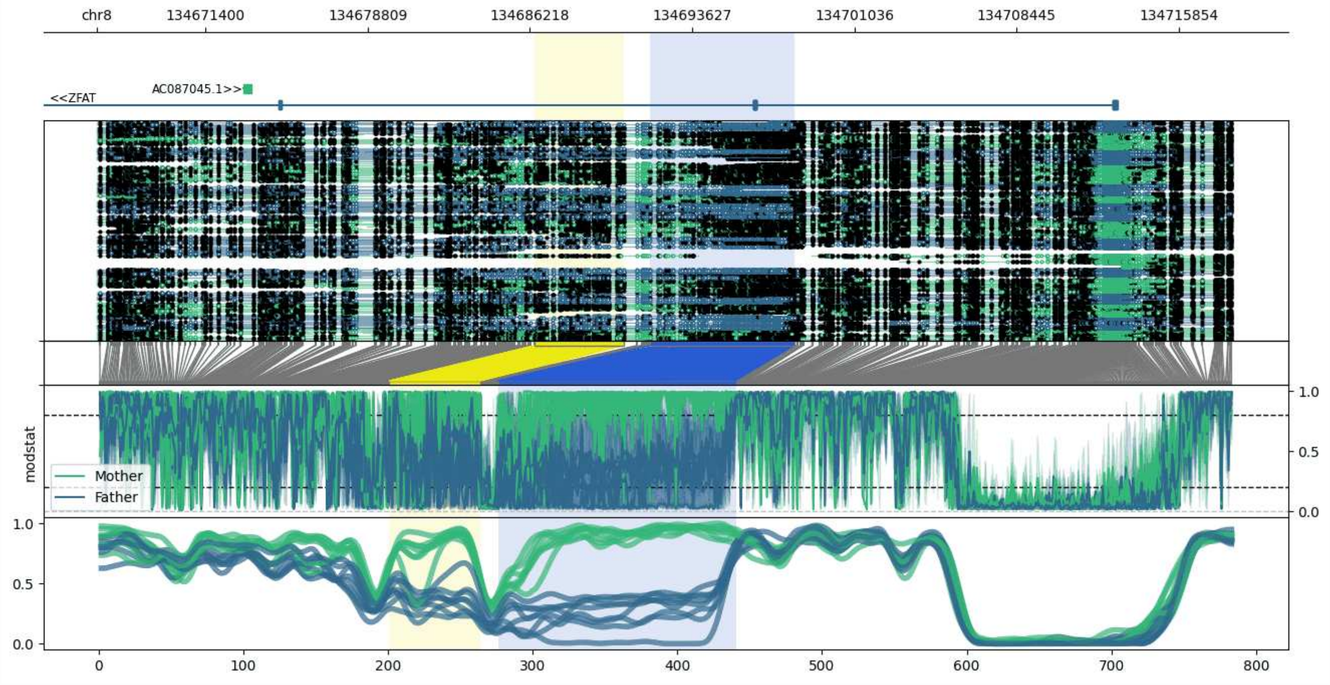


**chr8:134666444-134718309:** Region with 2 DMRs containing genes: AC087045.1, AC087045.2, ZFAT.

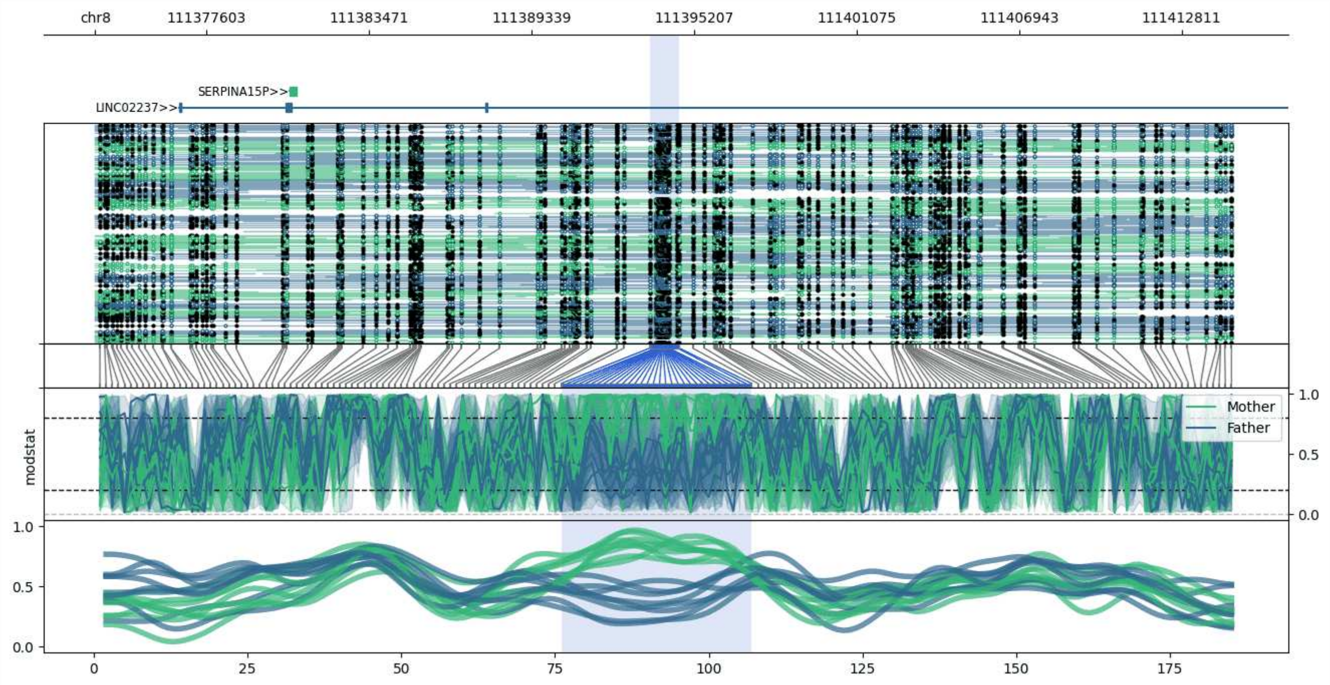


**chr8:111373609-111414686:** Region with 1 DMR containing genes: LINC02237, SERPINA15P.

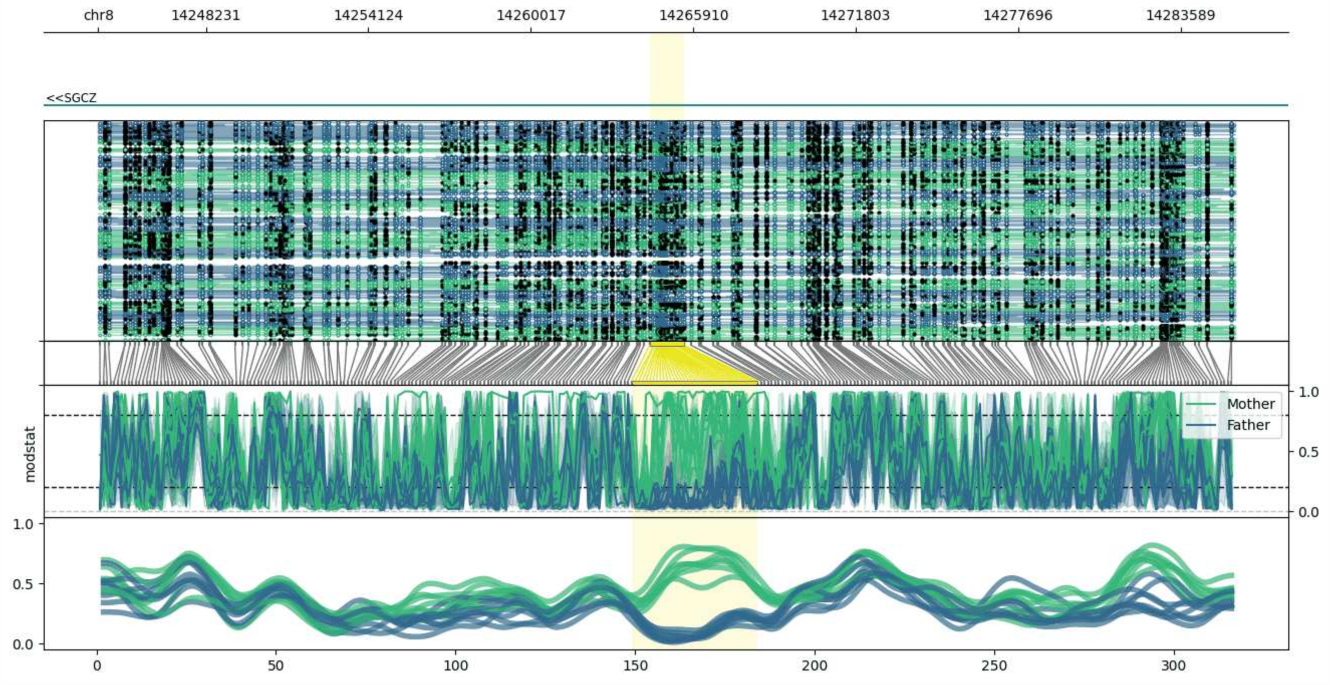


**chr8:14244324-14285578:** Region with 1 DMR containing genes: SGCZ.

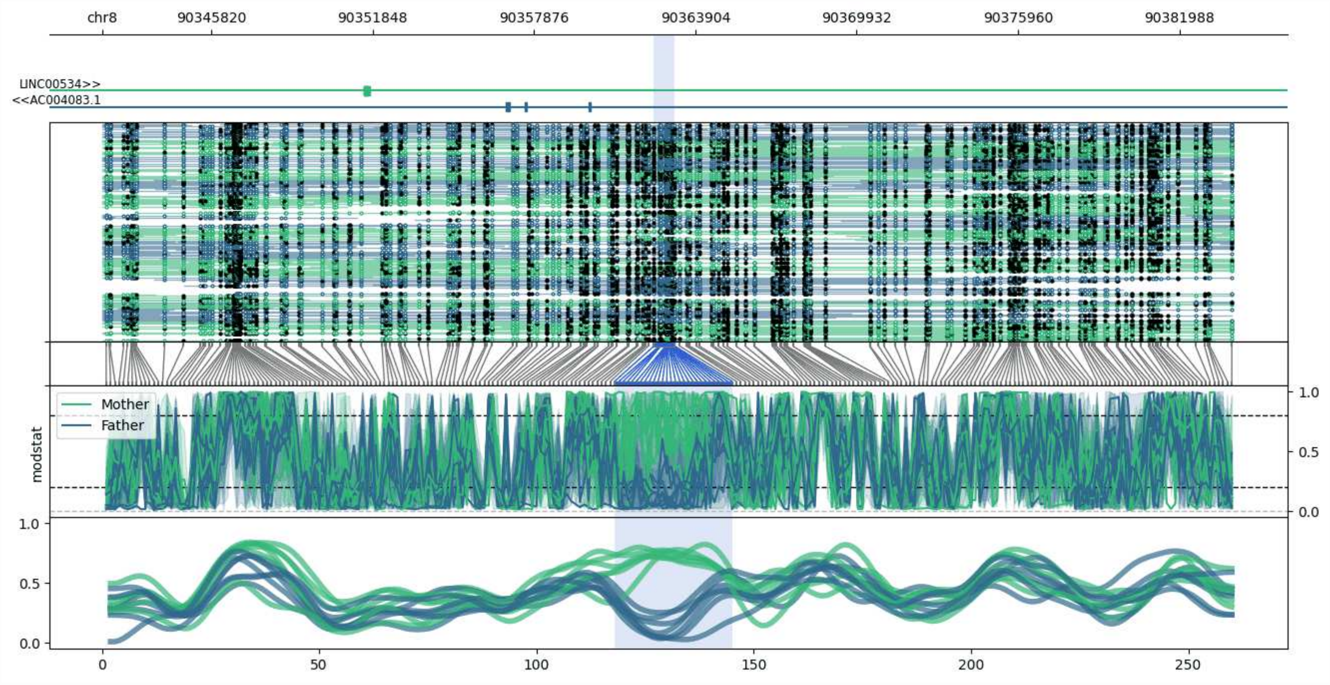


**chr8:90341754-90383951:** Region with 1 DMR containing genes: AC004083.1, LINC00534.

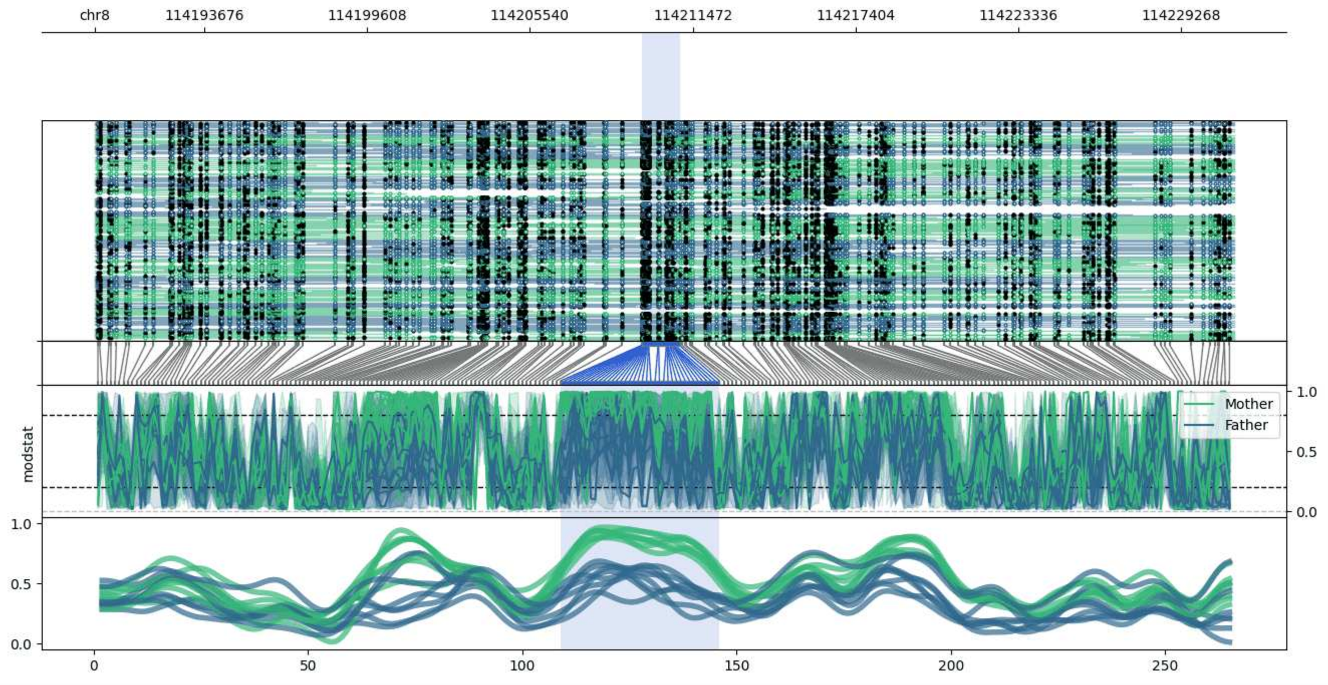


**chr8:114189674-114231204:** Region with 1 DMR containing no genes.

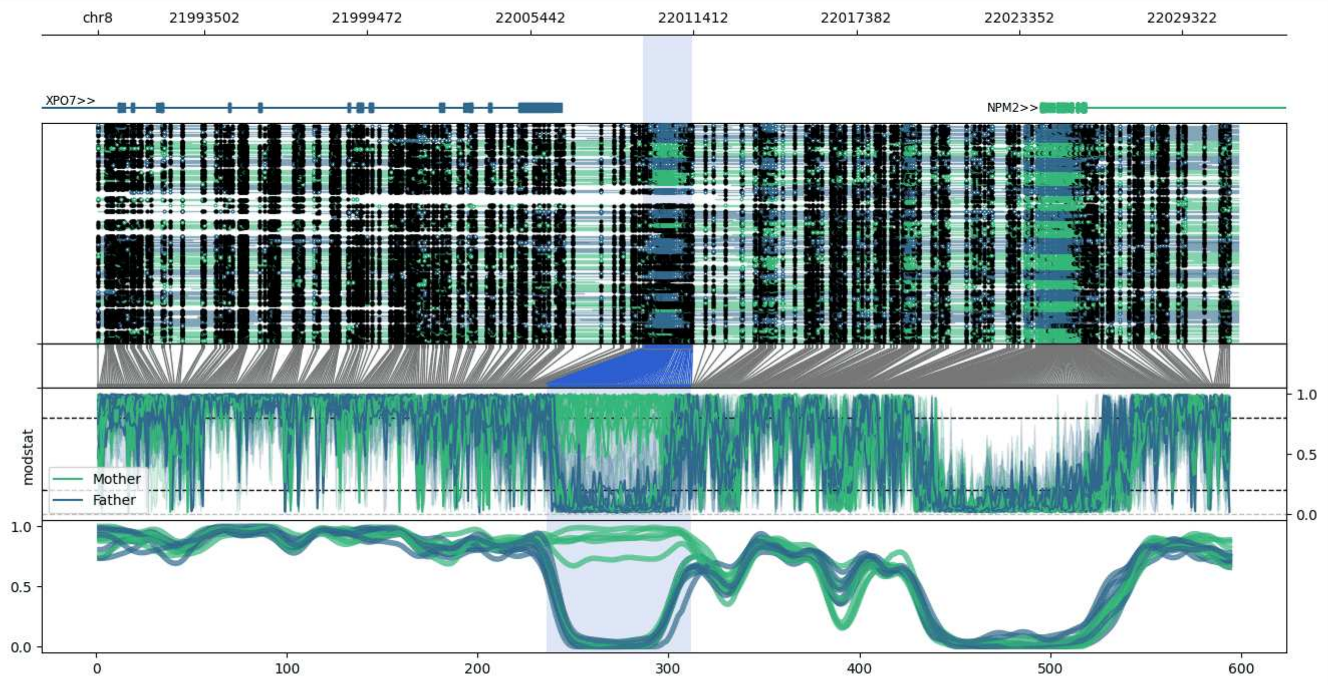


**chr8:21989583-22031378:** Region with 1 DMR containing genes: NPM2, XPO7.

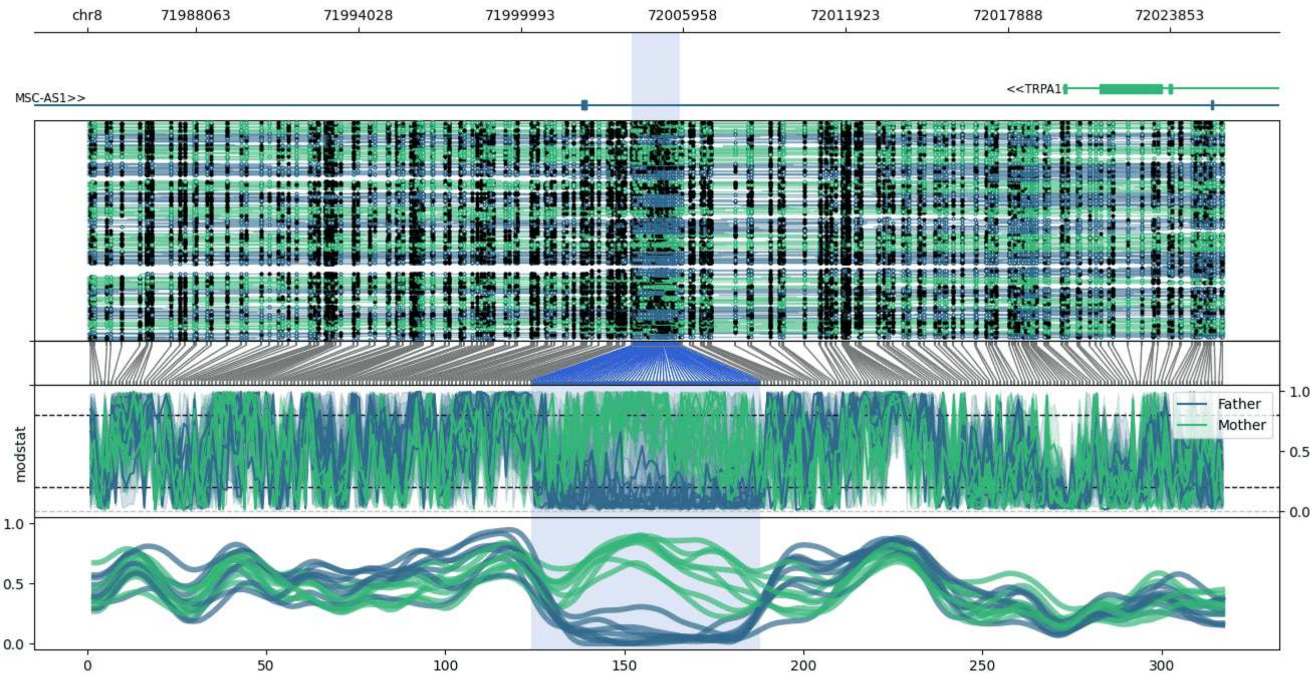


**chr8:71984066-72025823:** Region with 1 DMR containing genes: MSC-AS1, TRPA1.

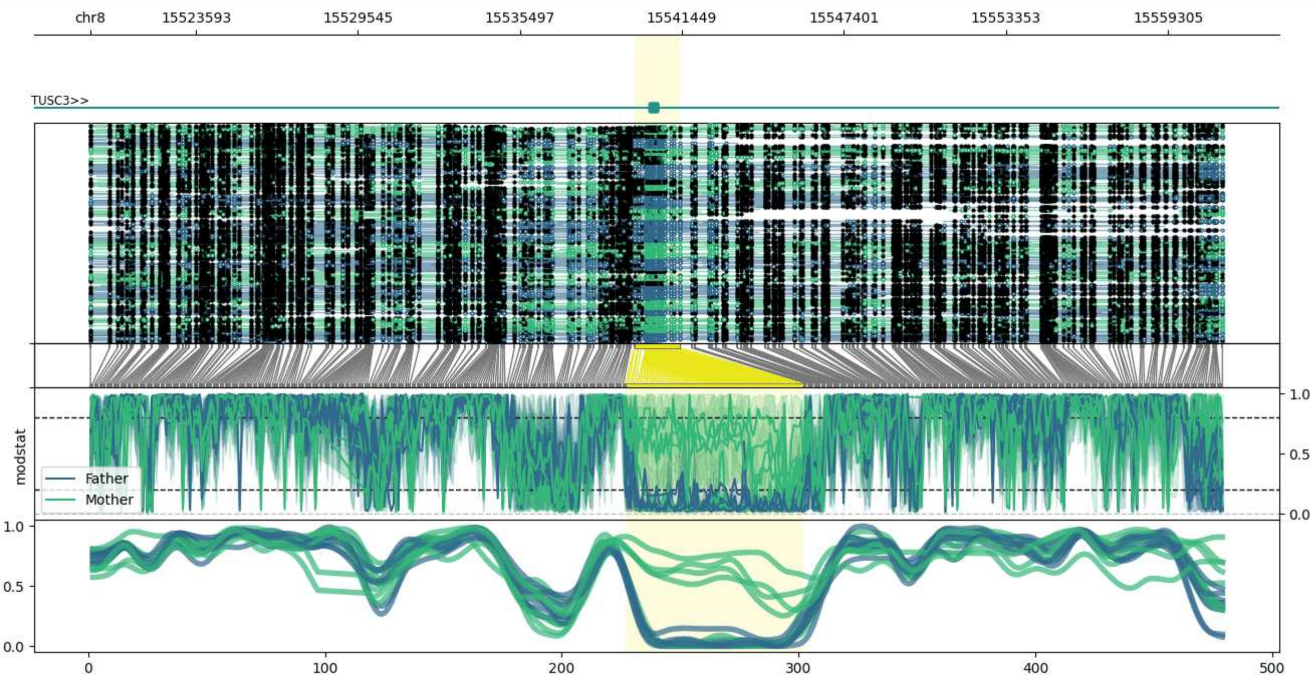


**chr8:15519701-15561367:** Region with 1 DMR containing genes: TUSC3.

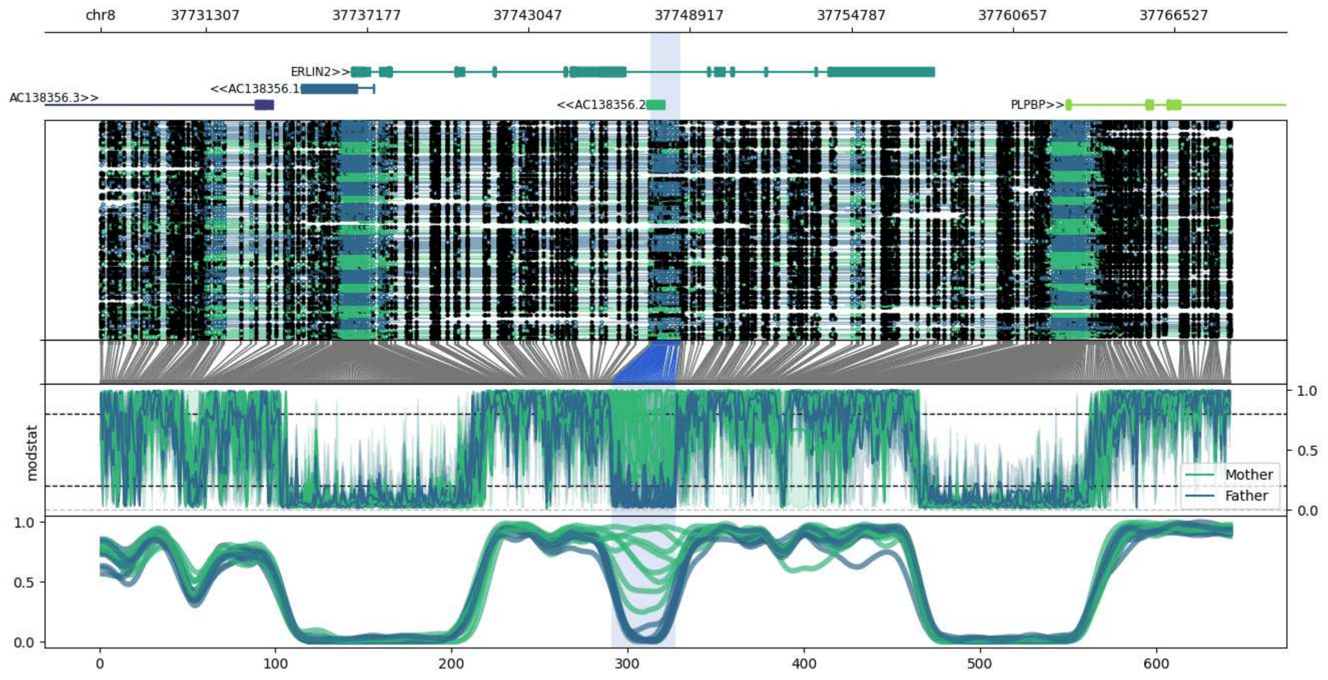


**chr8:37727474-37768570:** Region with 1 DMR containing genes: AC138356.1, AC138356.2, AC138356.3, ERLIN2, PLPBP.

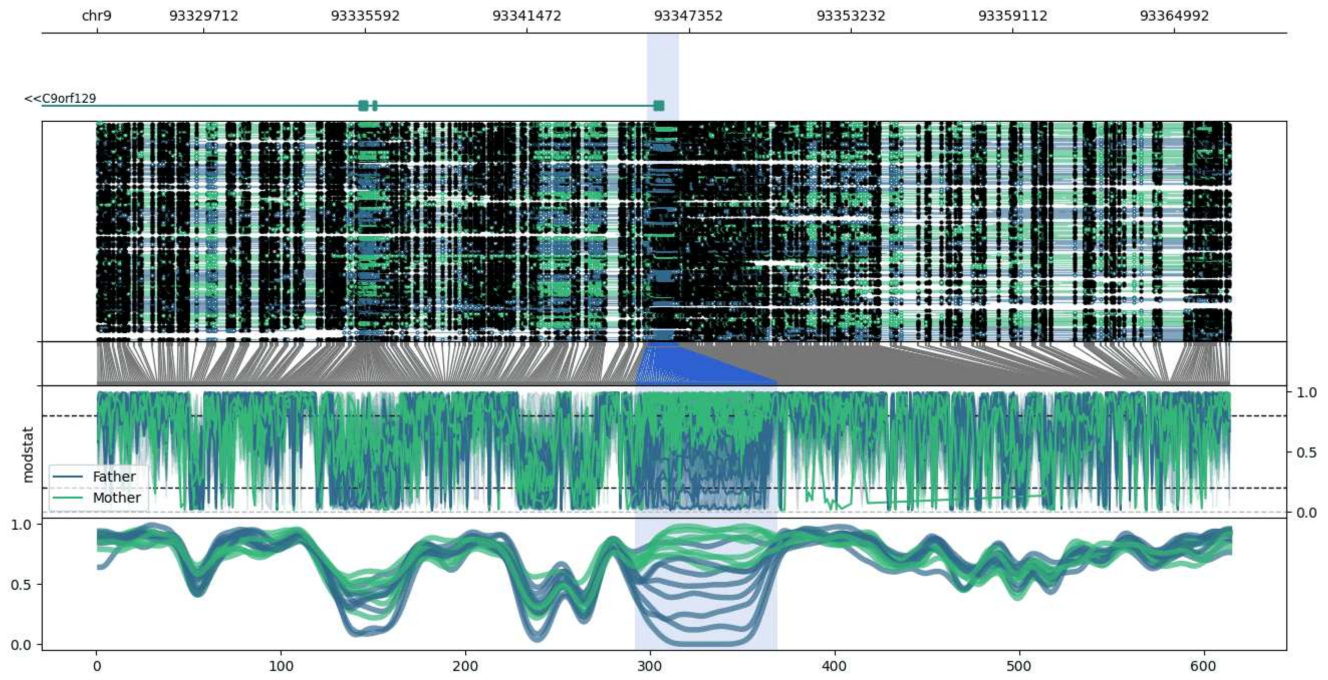


**chr9:93325846-93367006:** Region with 1 DMR containing genes: C9orf129, RNU6-829P, WNK2.

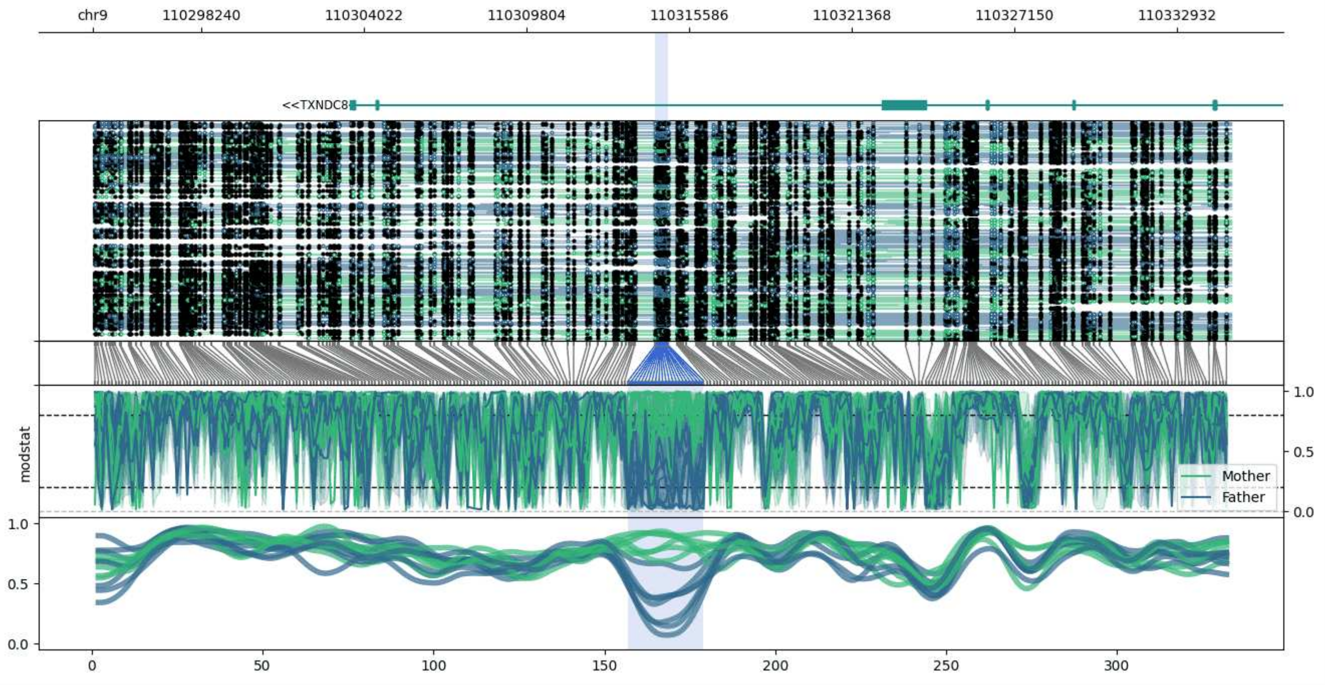


**chr9:110294385-110334860:** Region with 1 DMR containing genes: TXNDC8.

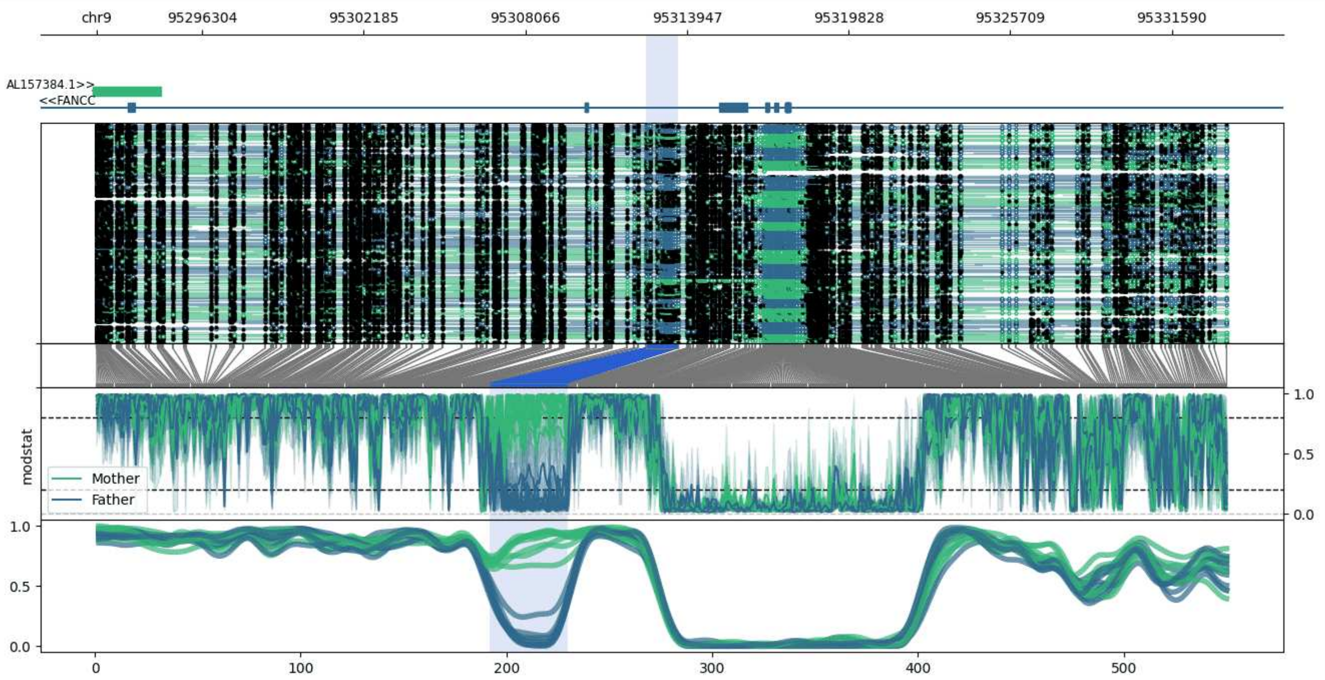


**chr9:95292464-95333632:** Region with 1 DMR containing genes: AL157384.1, FANCC.

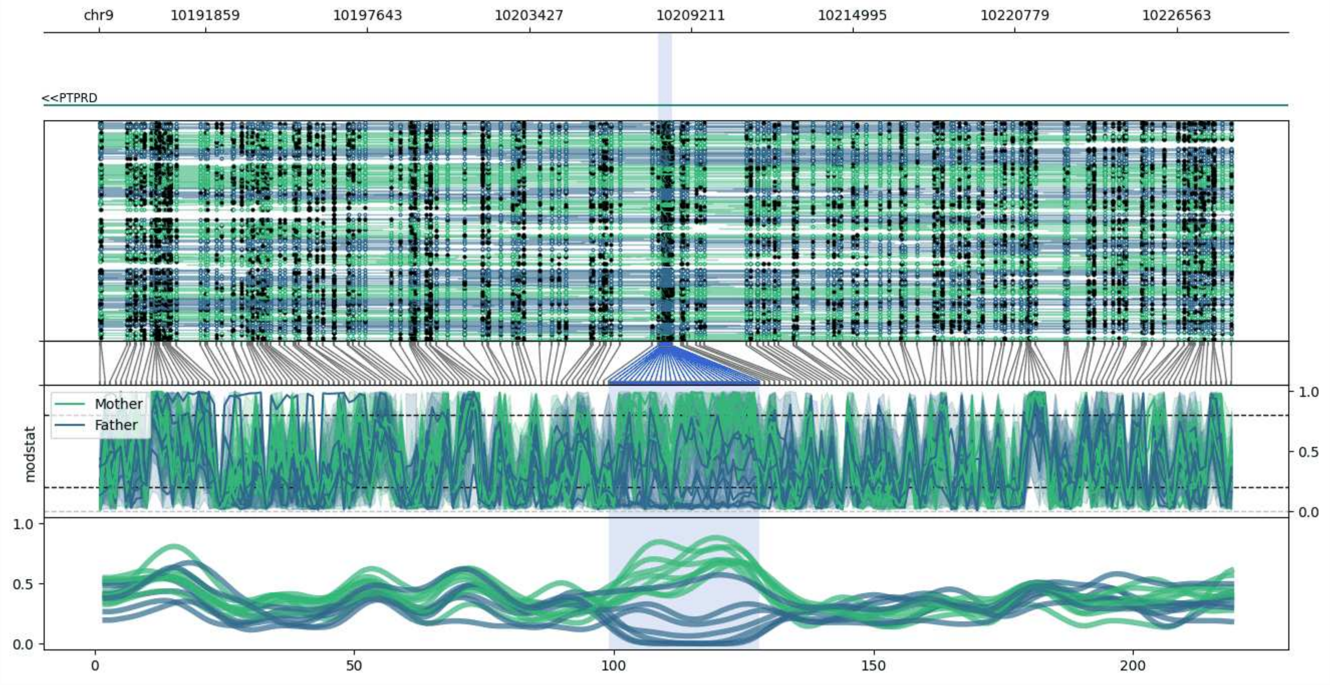


**chr9:10188043-10228534:** Region with 1 DMR containing genes: PTPRD.

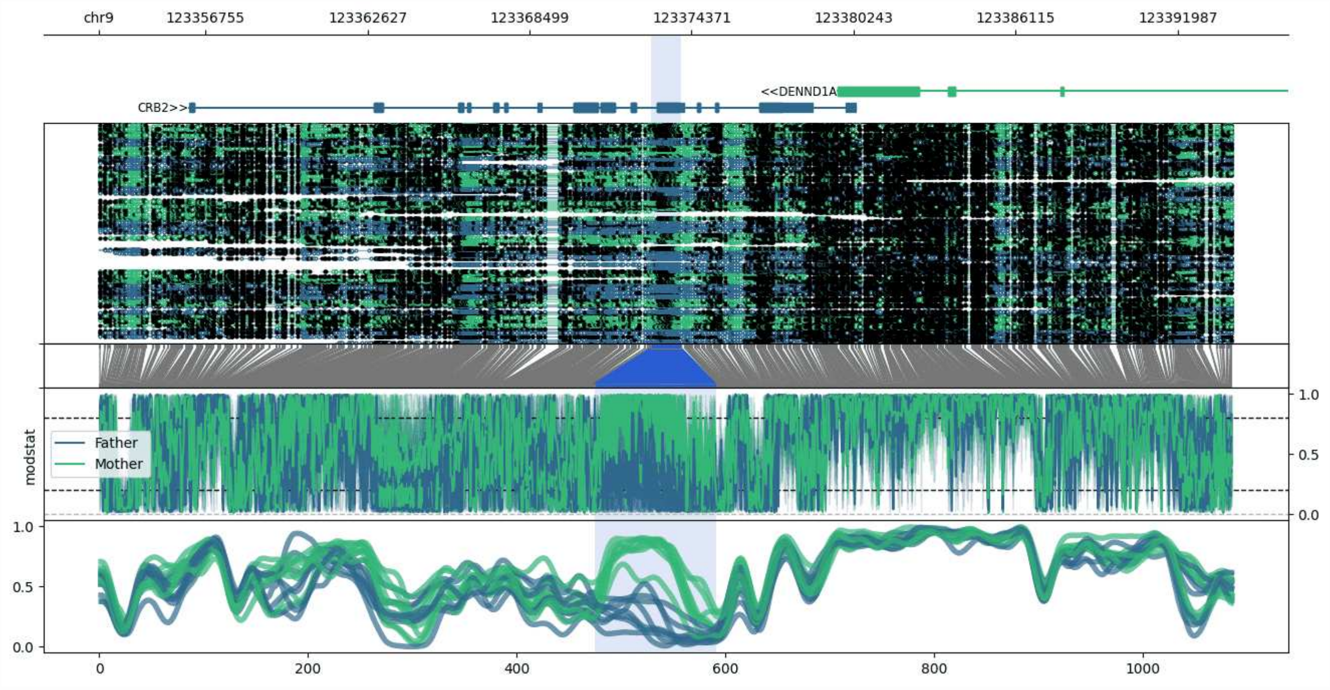


**chr9:123352902-123394006:** Region with 1 DMR containing genes: CRB2, DENND1A, MIR601.

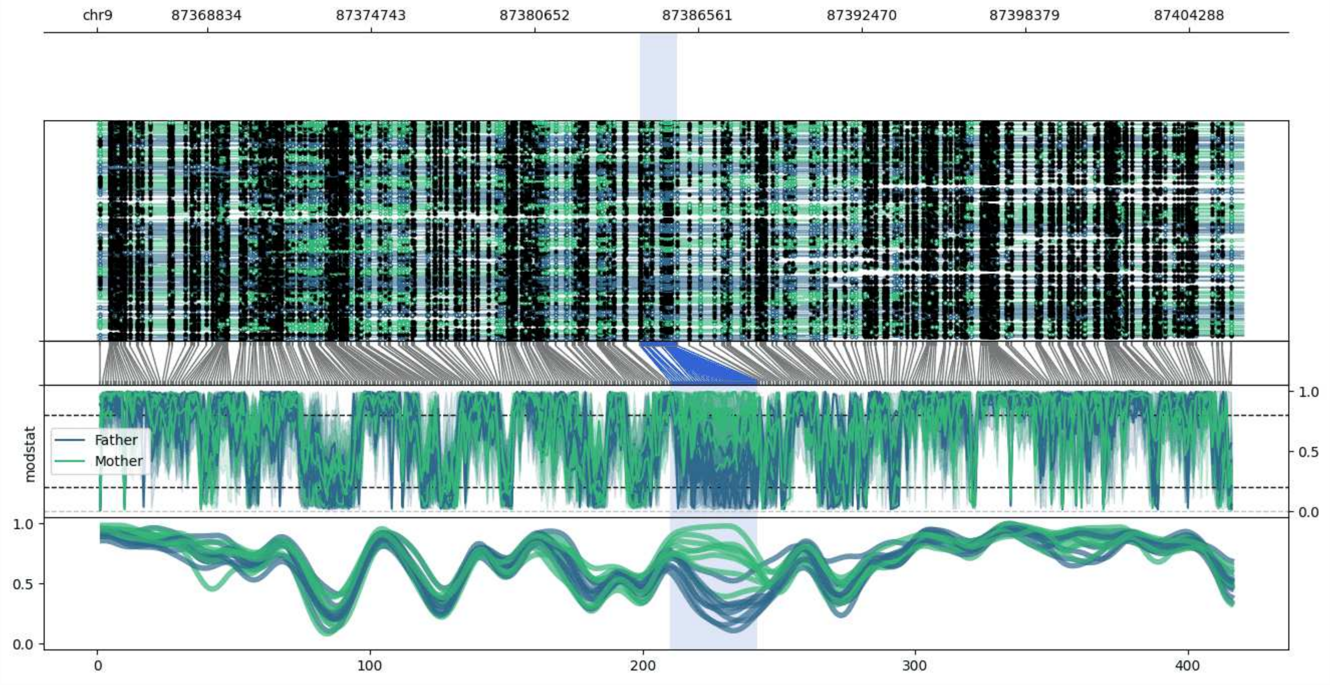


**chr9:87364872-87406238:** Region with 1 DMR containing no genes.

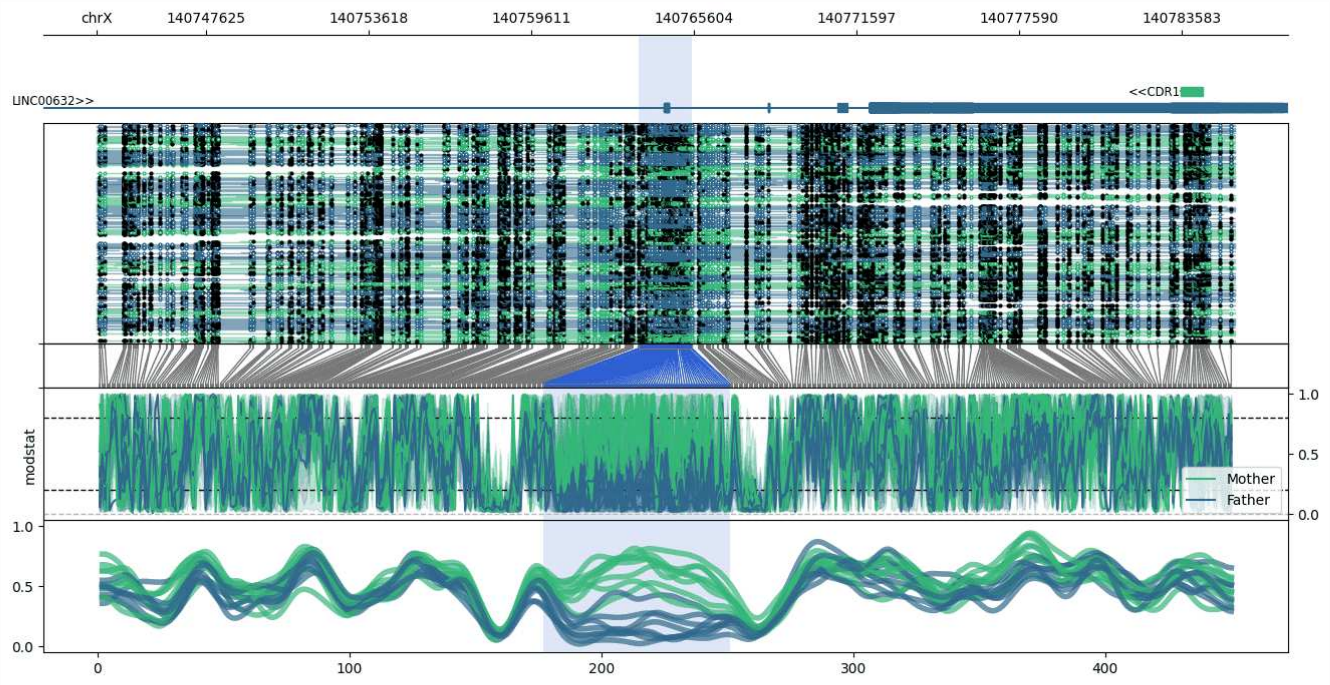


**chrX:140743590-140785543:** Region with 1 DMR containing genes: CDR1, LINC00632.

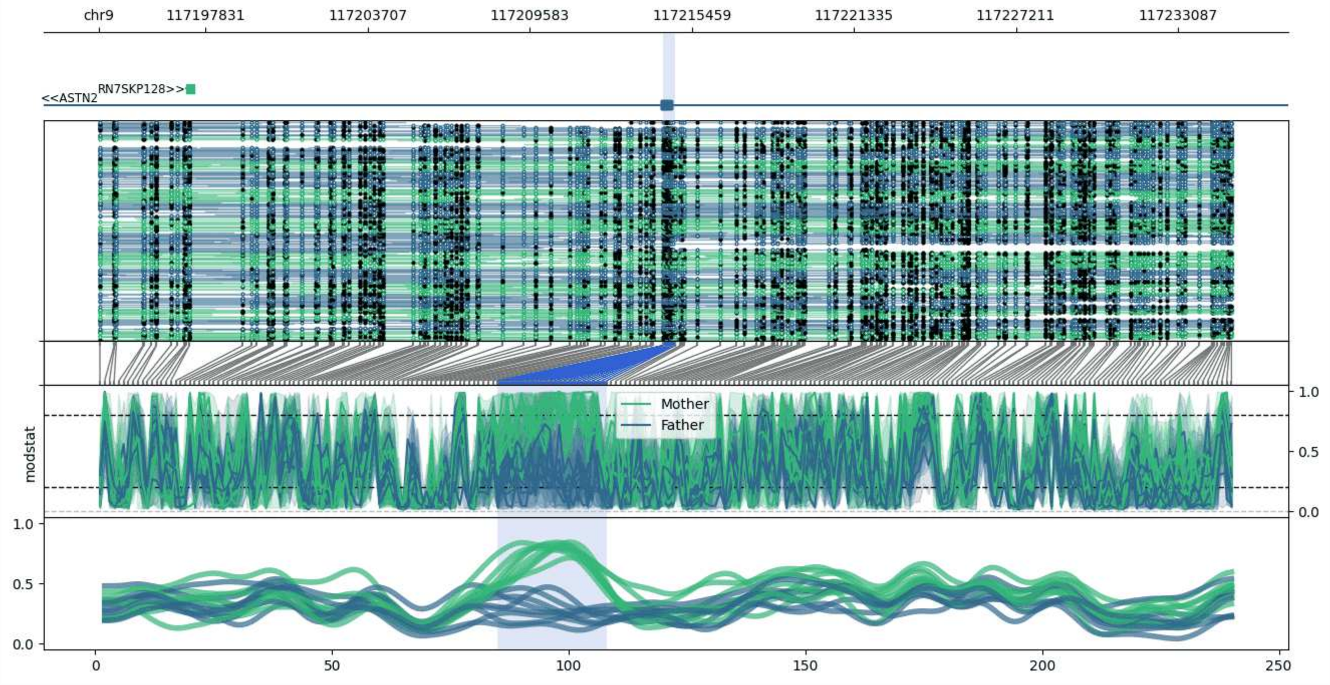


**chr9:117193975-117235107:** Region with 1 DMR containing genes: ASTN2, RN7SKP128.

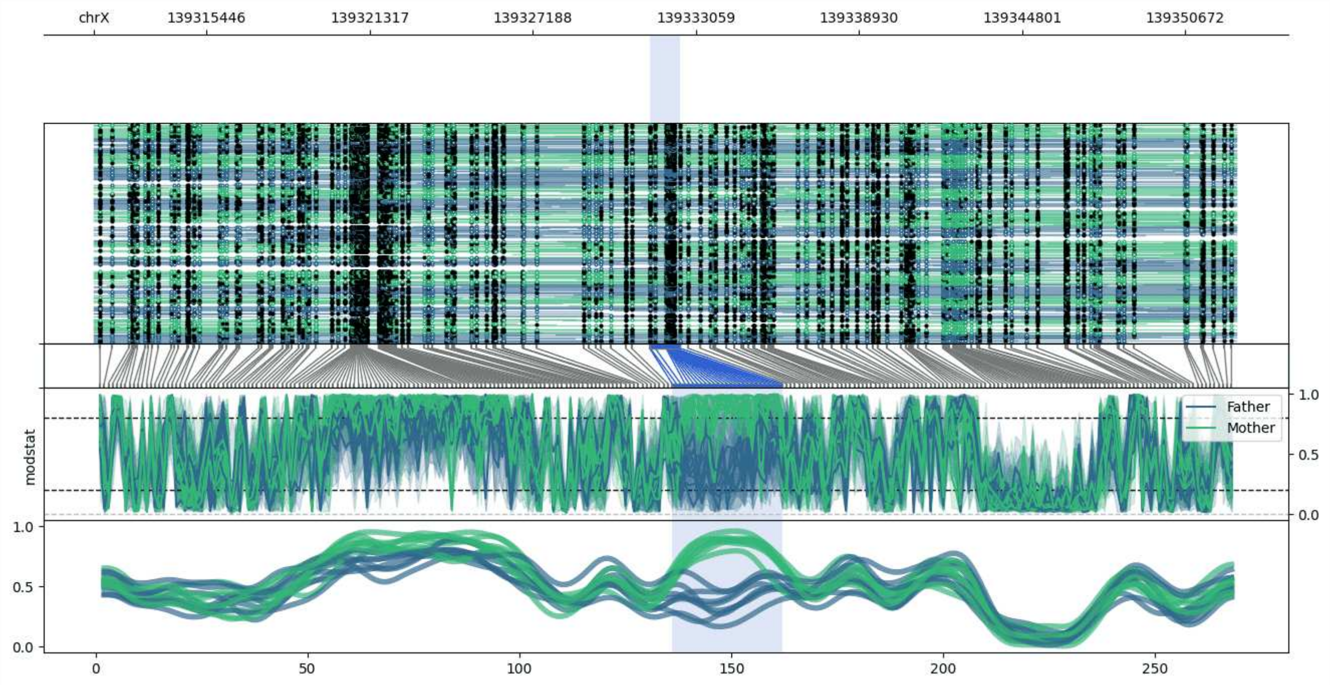


**chrX:139311393-139352491:** Region with 1 DMR containing genes: SNURFL.

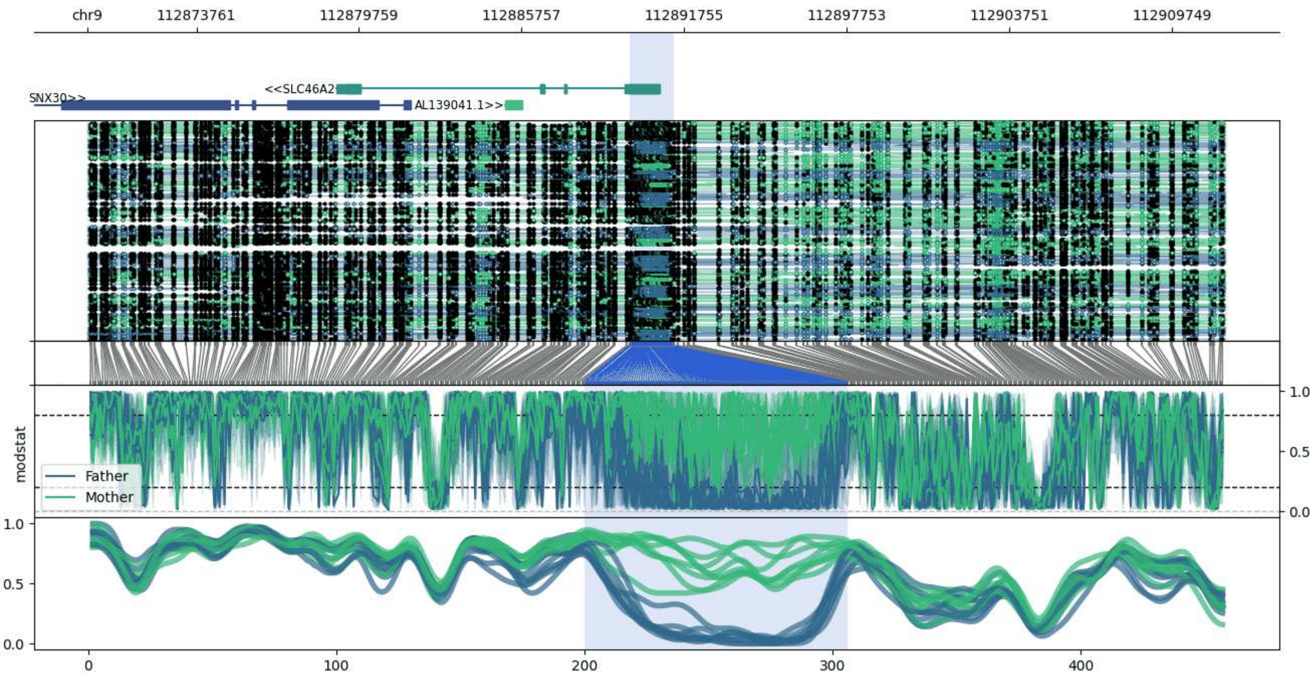


**chr9:112869741-112911733:** Region with 1 DMR containing genes: AL139041.1, SLC46A2, SNX30.

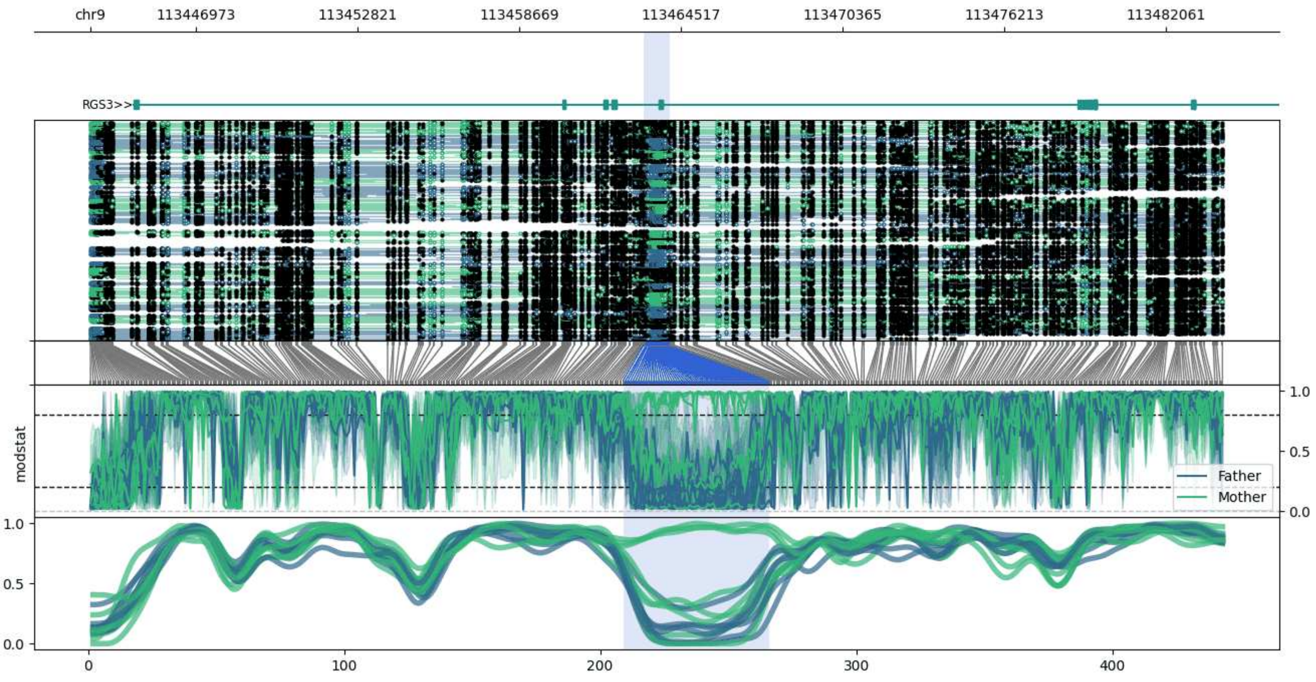


**chr9:113443171-113484109:** Region with 1 DMR containing genes: RGS3.

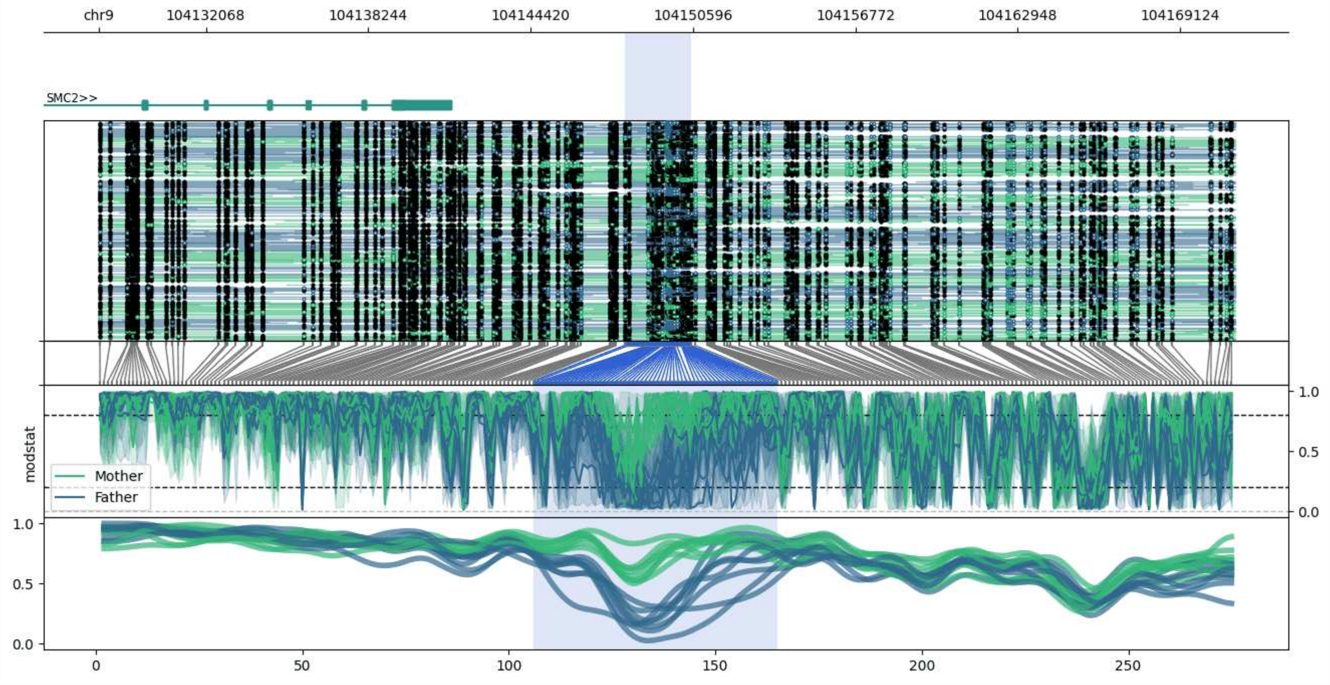


**chr9:104127995-104171227:** Region with 1 DMR containing genes: SMC2.

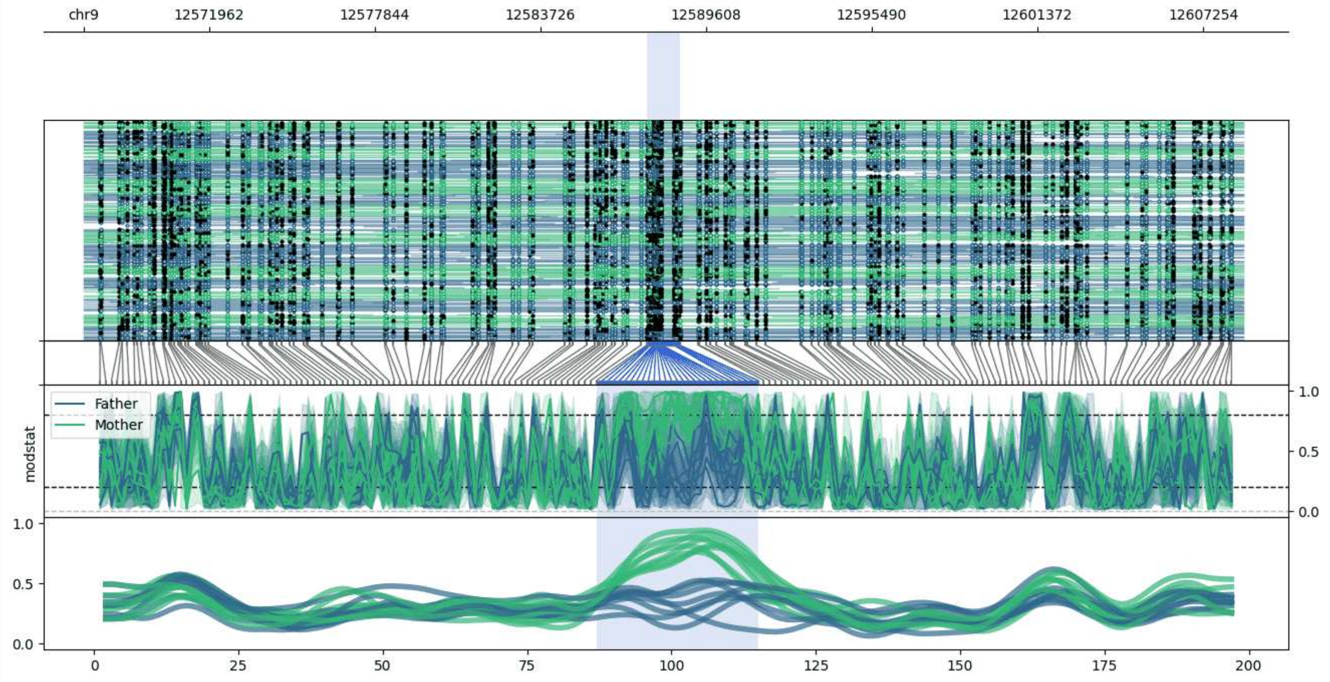


**chr9:12567500-12608674:** Region with 1 DMR containing no genes.

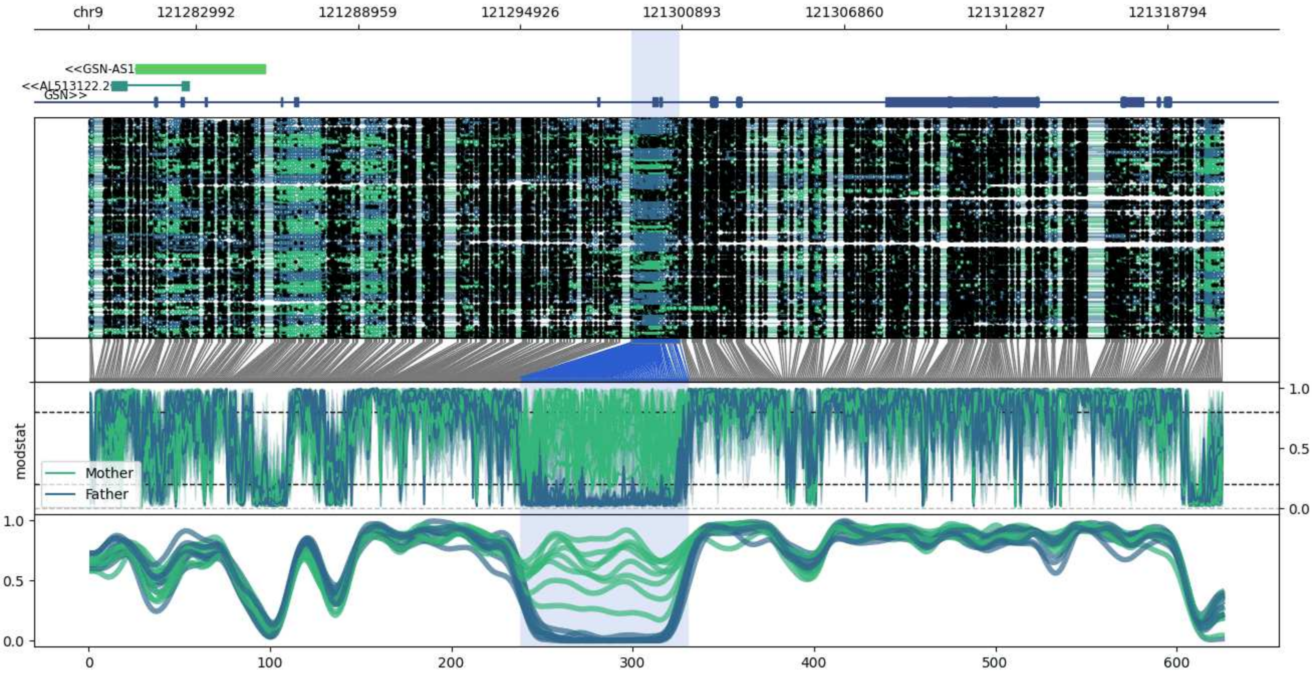


**chr9:121279031-121320804:** Region with 1 DMR containing genes: AL513122.2, GSN, GSN-AS1.

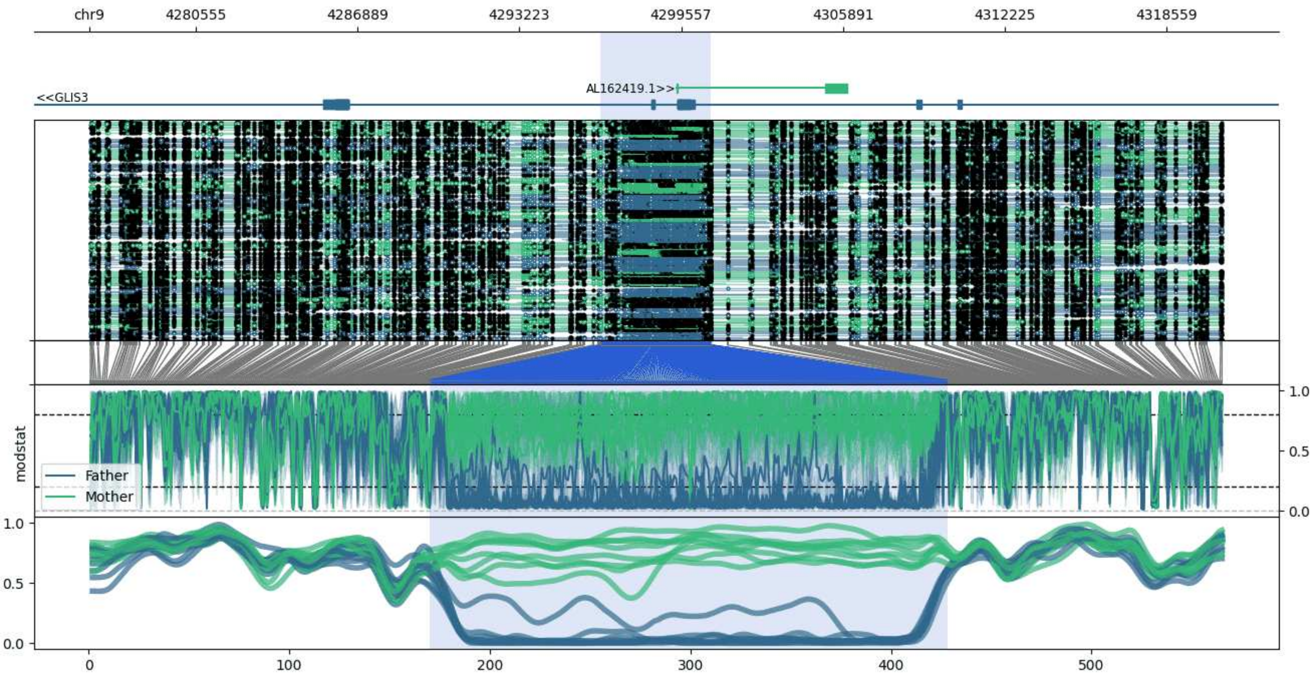


**chr9:4276391-4320730:** Region with 1 DMR containing genes: AL162419.1, GLIS3.

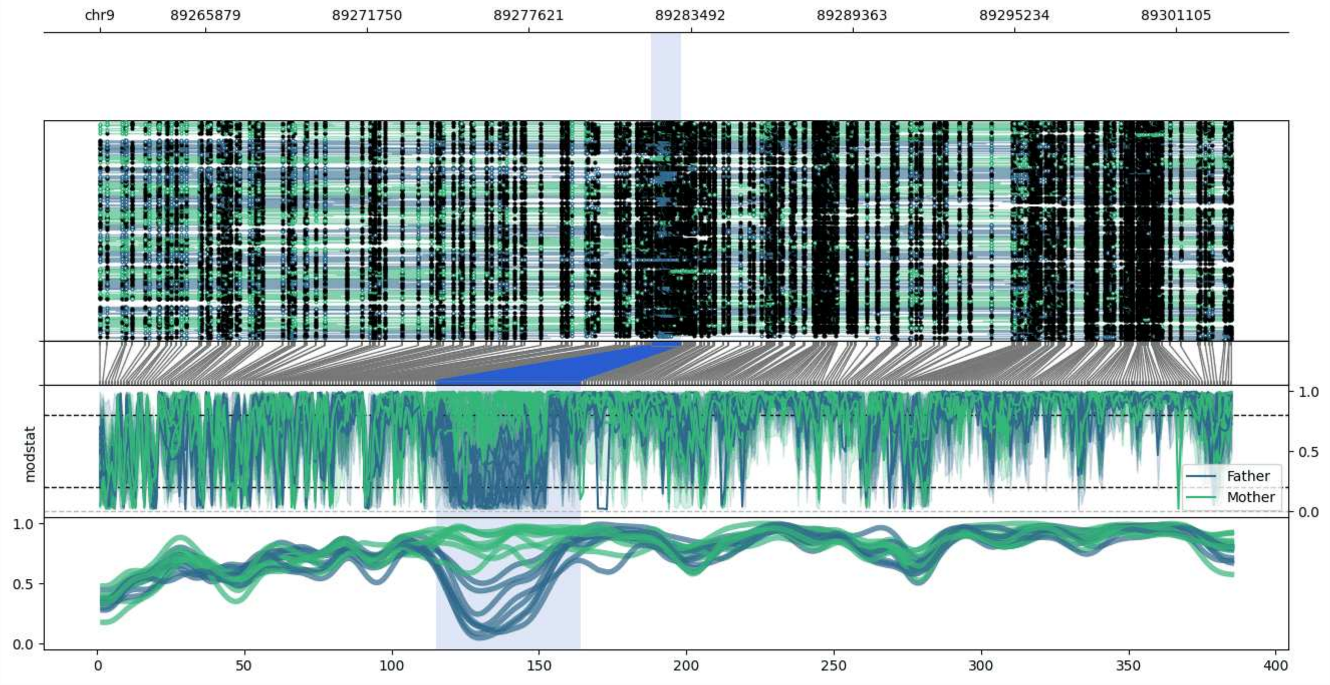


**chr9:89262048-89303146:** Region with 1 DMR containing genes: CKS2, MIR3153.

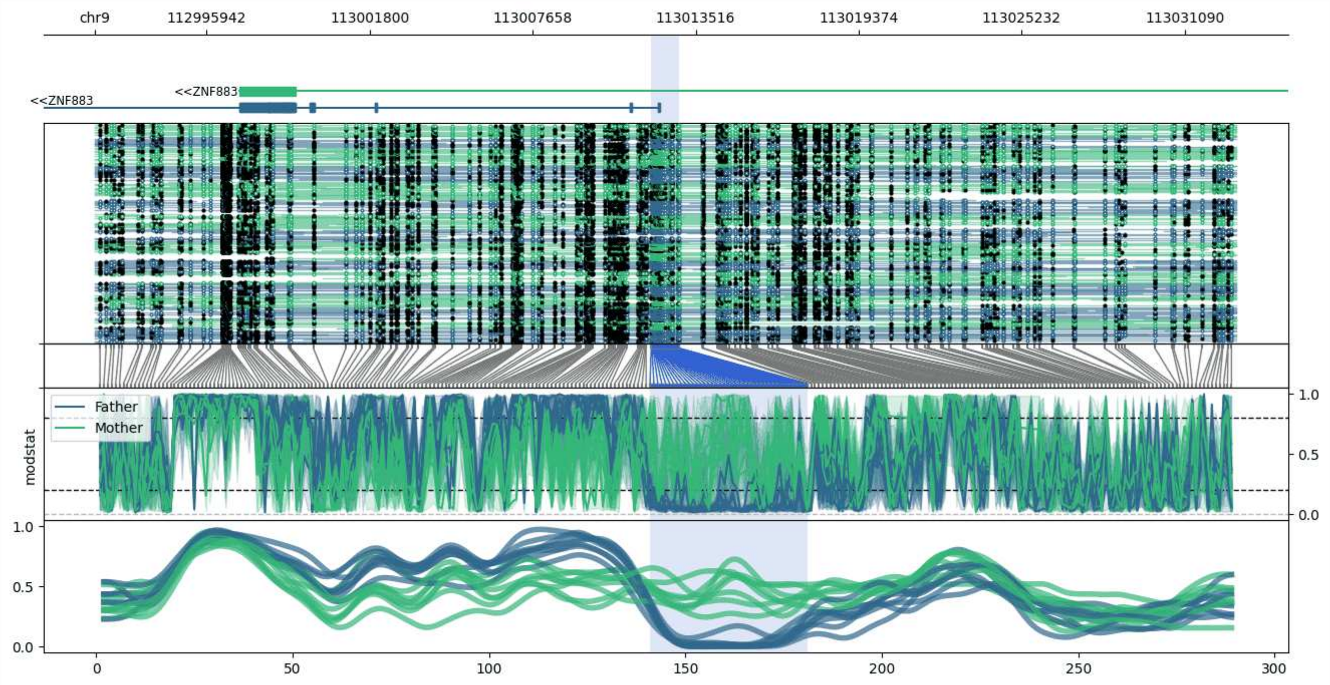


**chr9:112991919-113032931:** Region with 1 DMR containing genes: ZFP37, ZNF883.

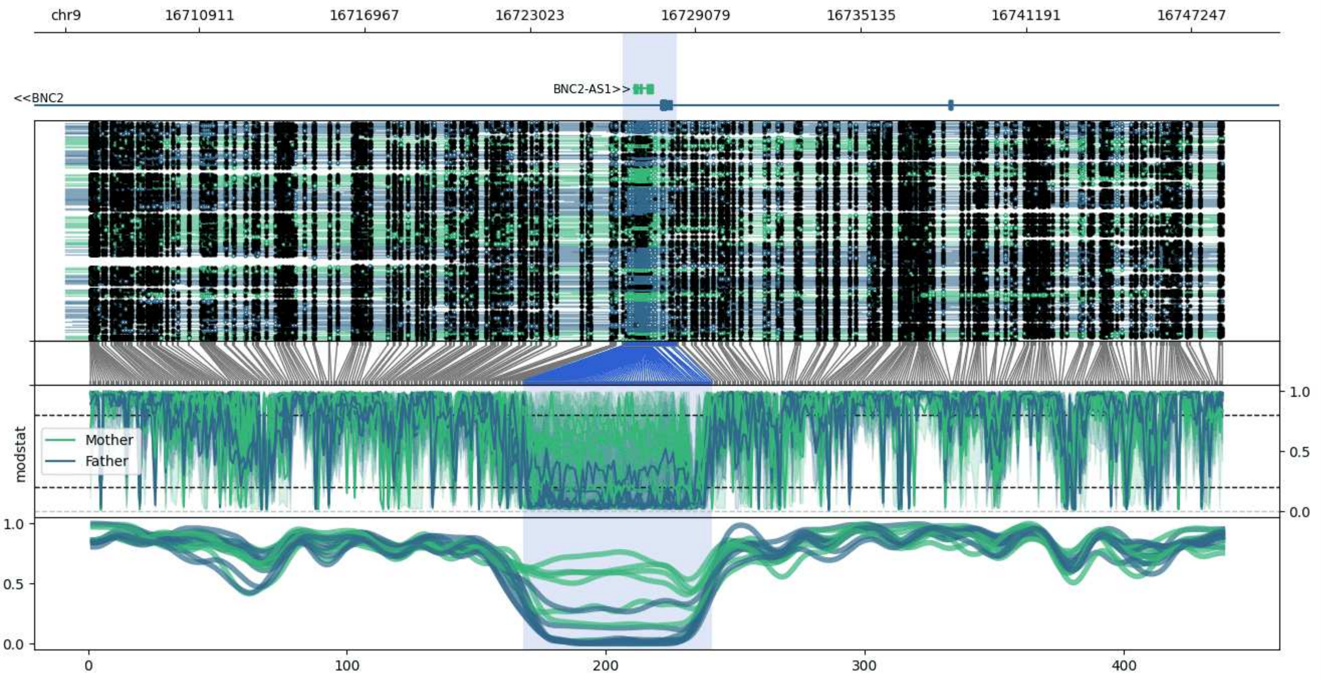


**chr9:16706017-16748412:** Region with 1 DMR containing genes: BNC2, BNC2-AS1.

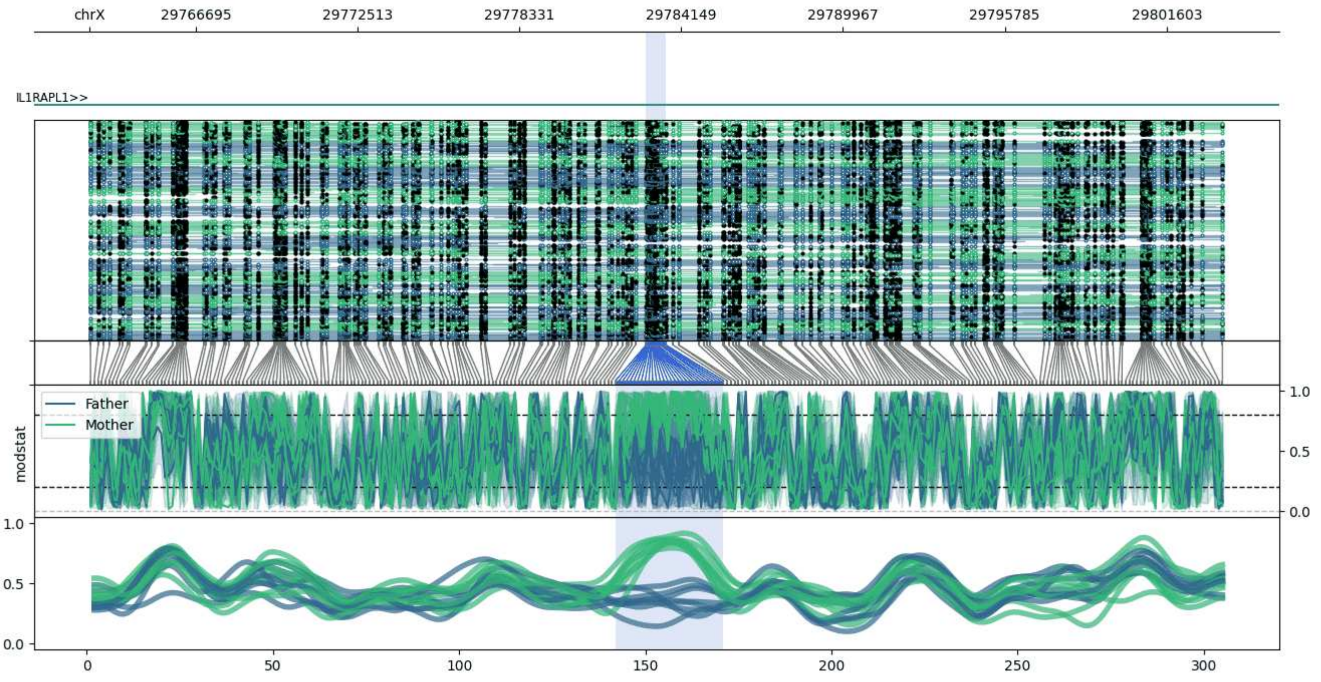


**chrX:29762881-29803612:** Region with 1 DMR containing genes: IL1RAPL1.

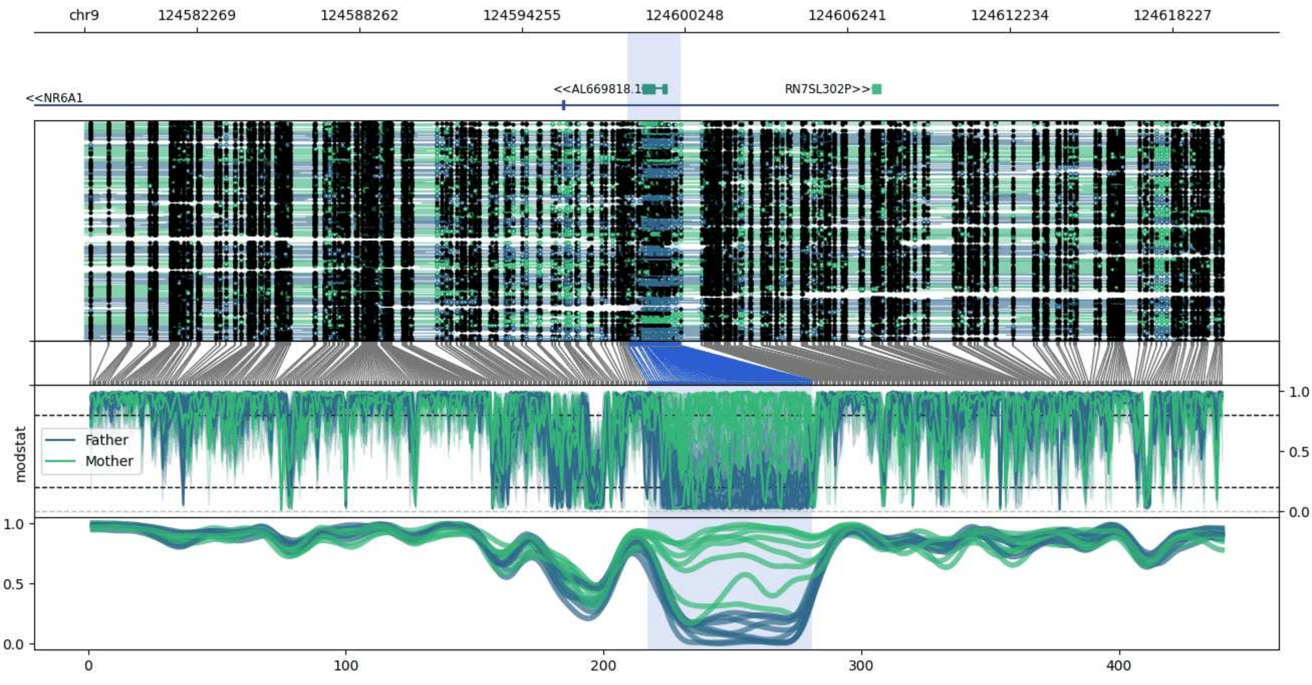


**chr9:124578143-124620095:** Region with 1 DMR containing genes: AL669818.1, NR6A1, RN7SL302P.

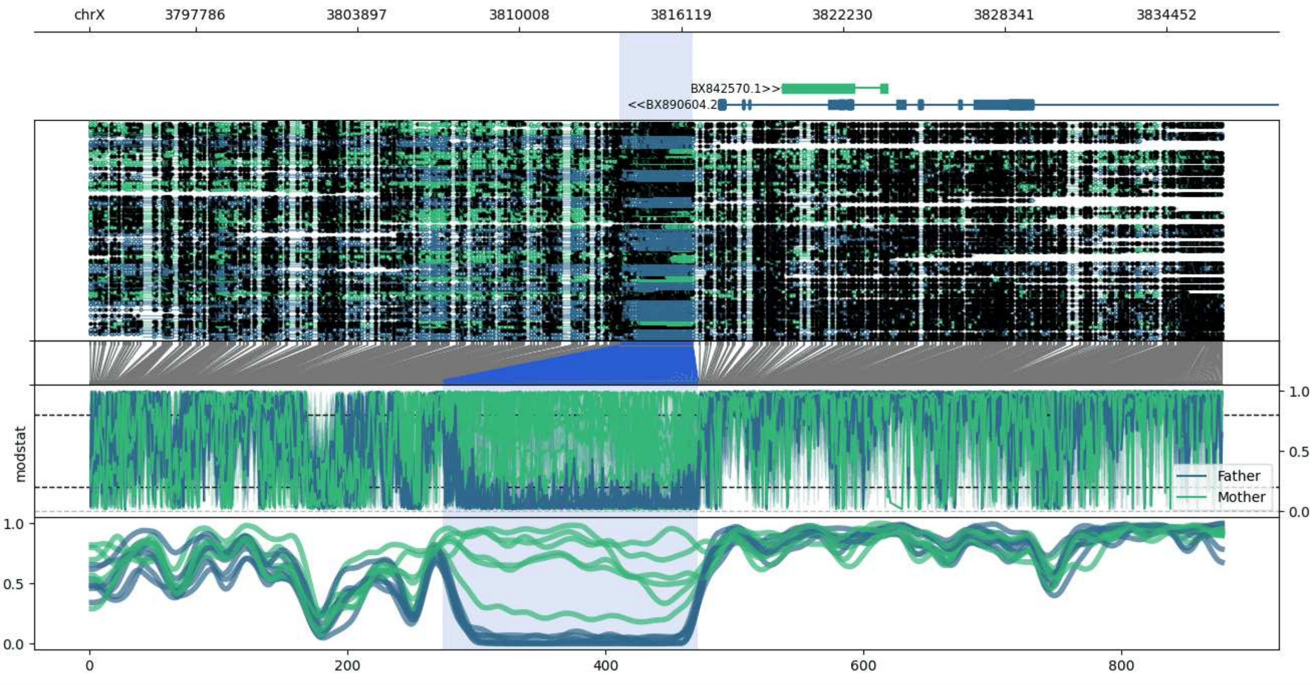


**chrX:3793778-3836559:** Region with 1 DMR containing genes: BX842570.1, BX890604.2.

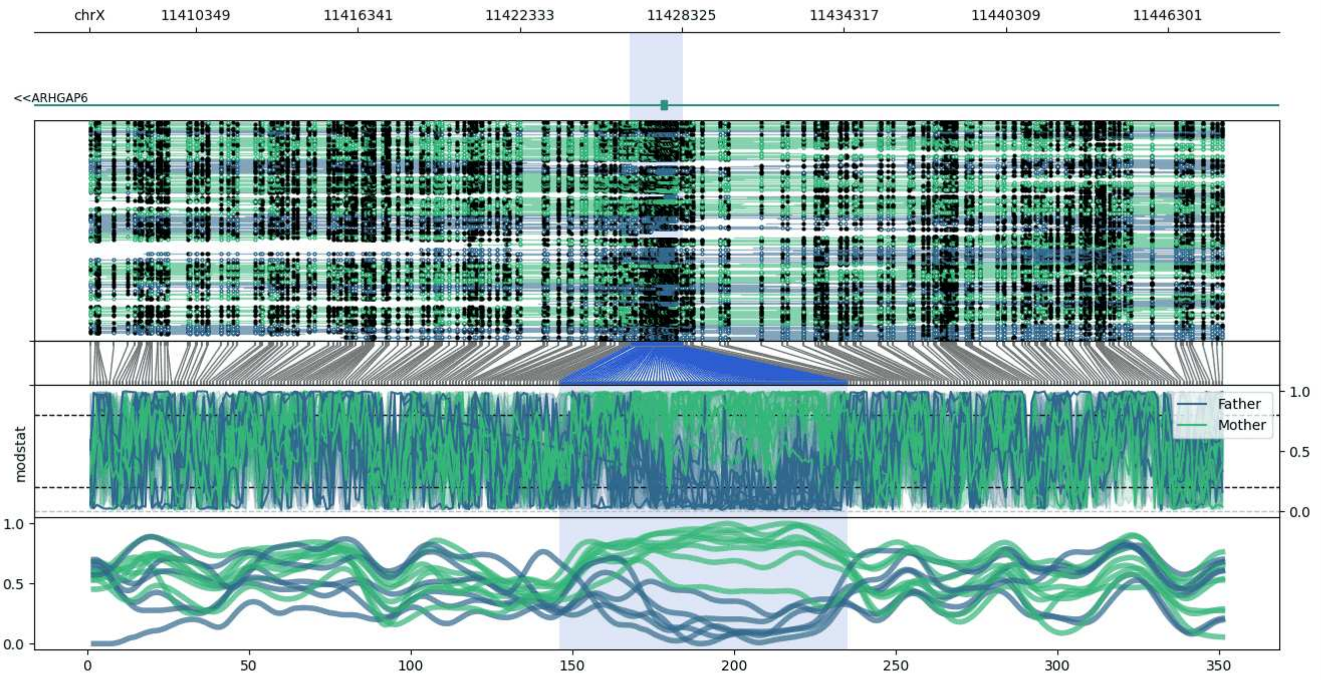


**chrX:11406400-11448345:** Region with 1 DMR containing genes: ARHGAP6.

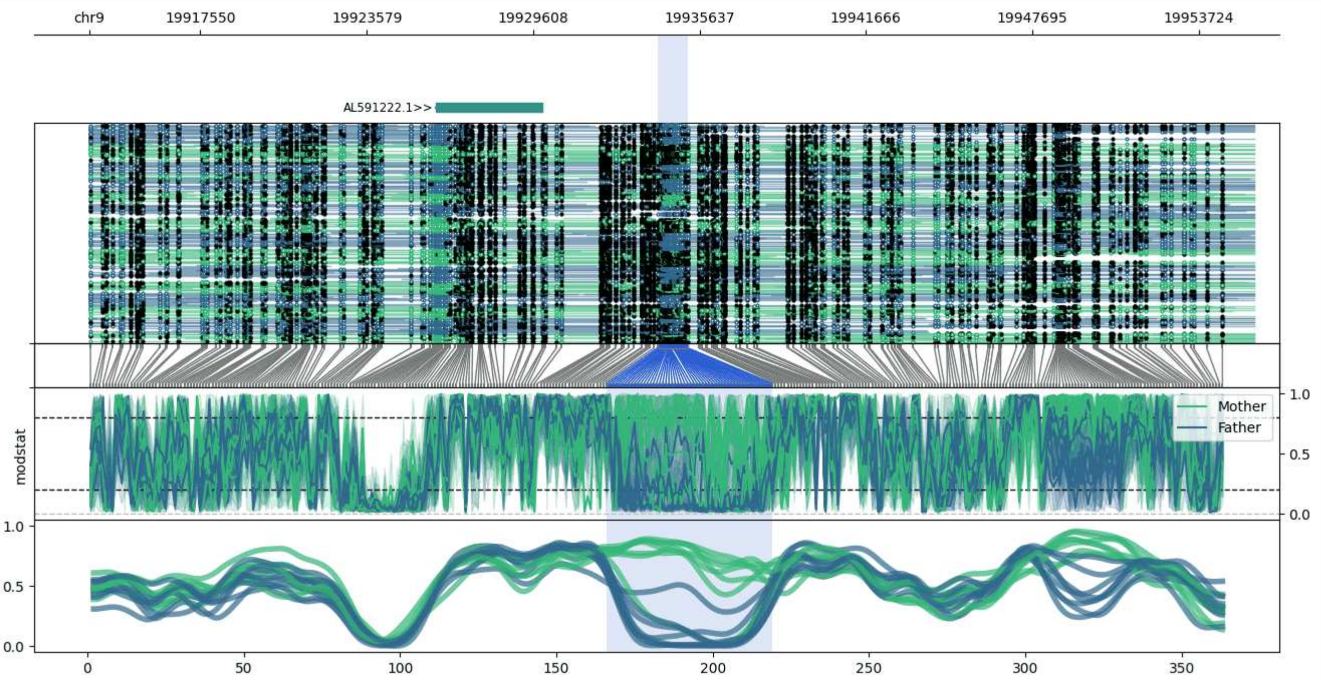


**chr9:19913526-19955732:** Region with 1 DMR containing genes: AL591222.1, AL591222.2.

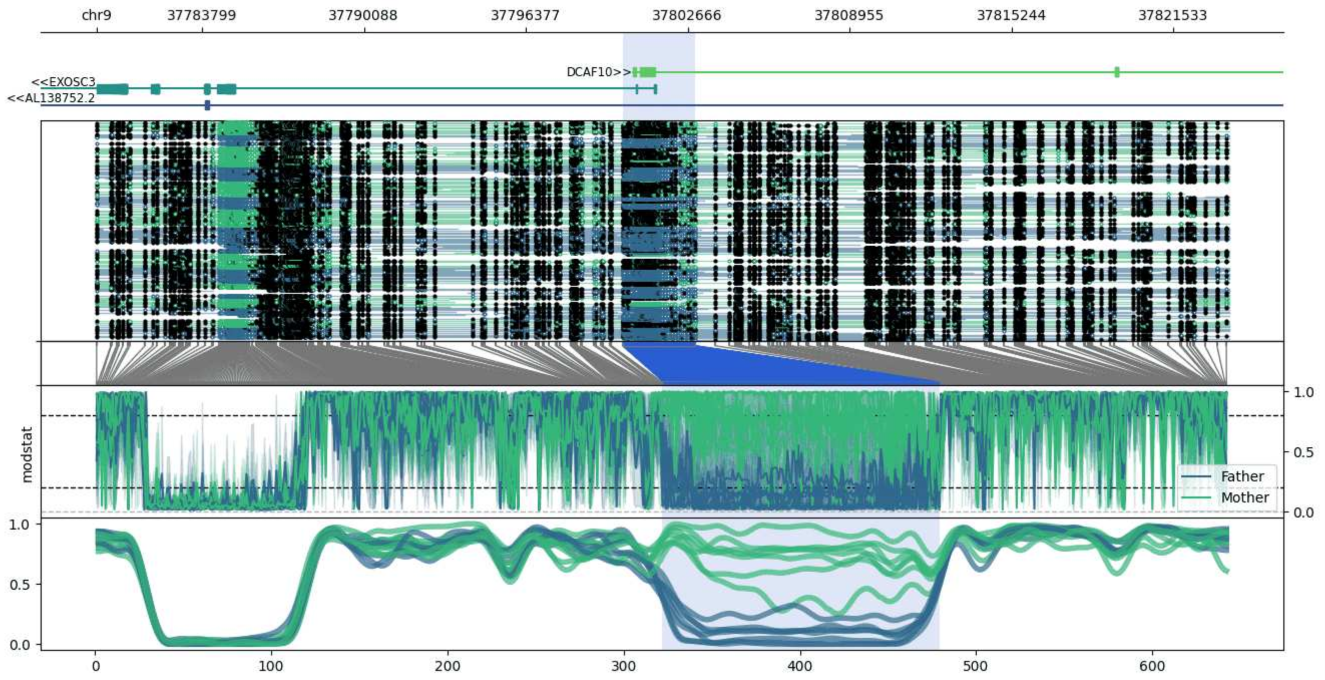


**chr9:37779696-37823722:** Region with 1 DMR containing genes: AL138752.2, DCAF10, EXOSC3, TRMT10B.

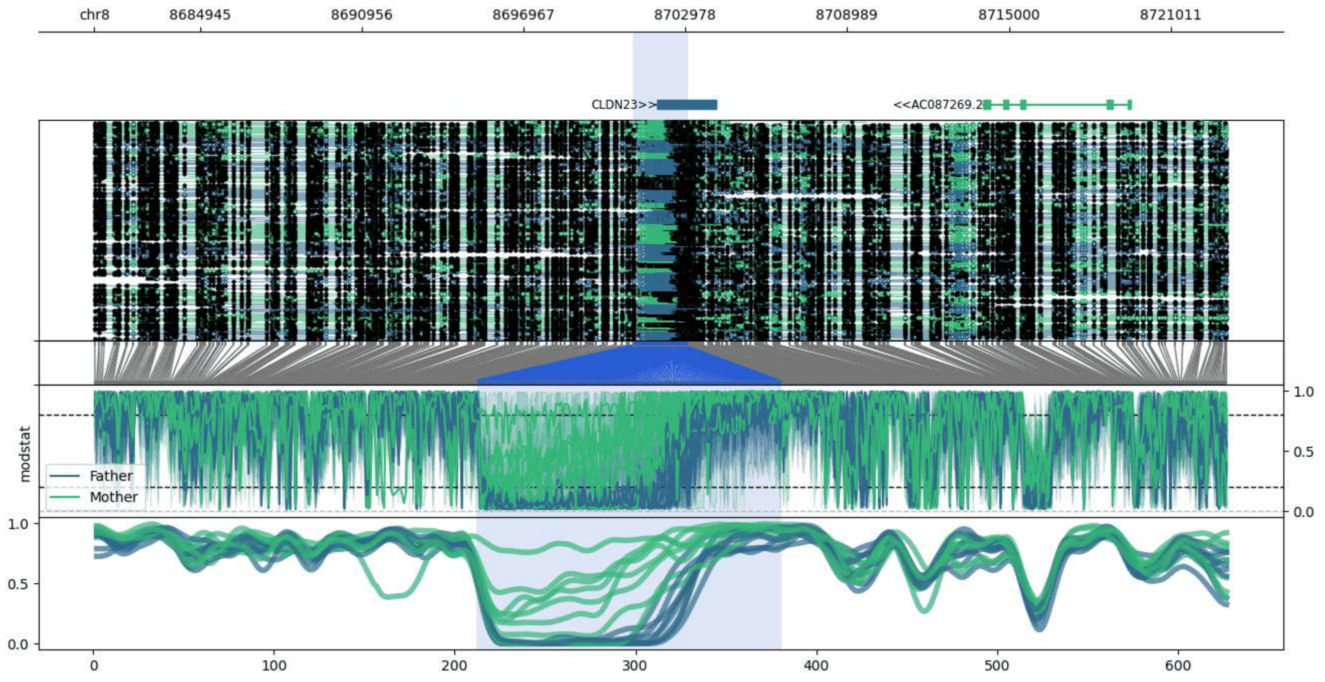


**chr8:8681011-8723089:** Region with 1 DMR containing genes: AC087269.1, AC087269.2, CLDN23.

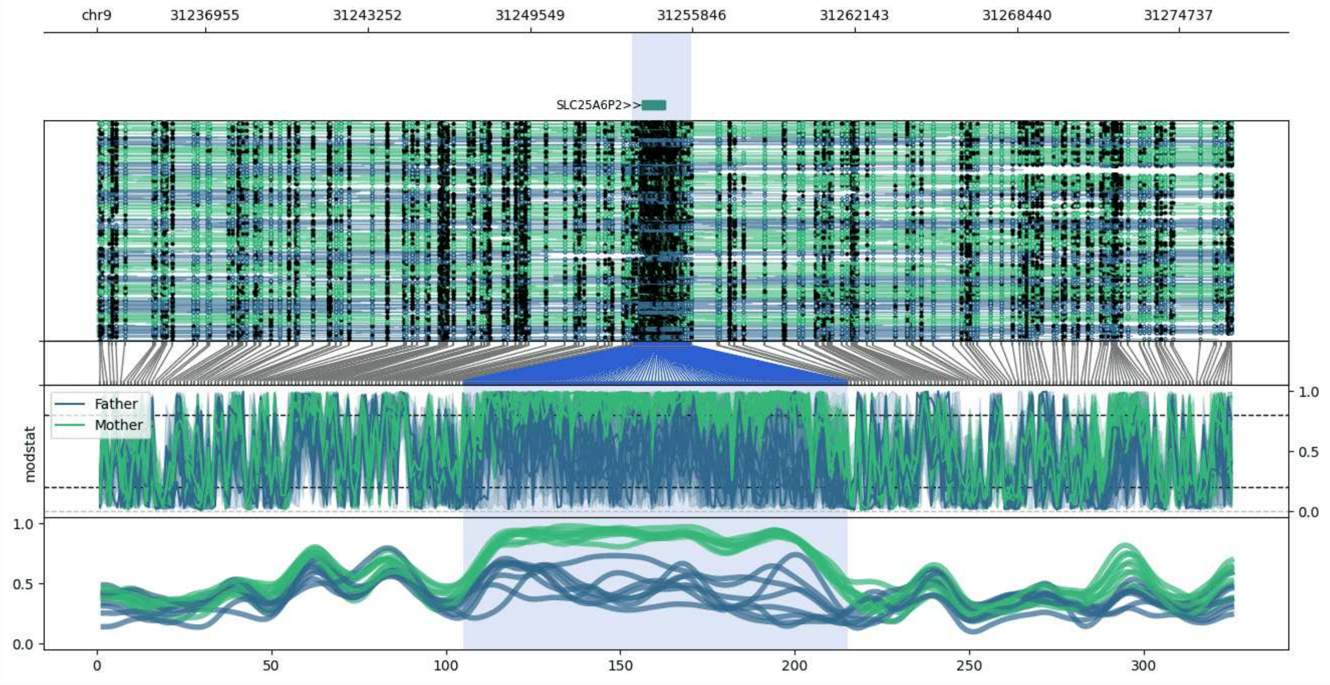


**chr9:31232732-31276817:** Region with 1 DMR containing genes: SLC25A6P2.

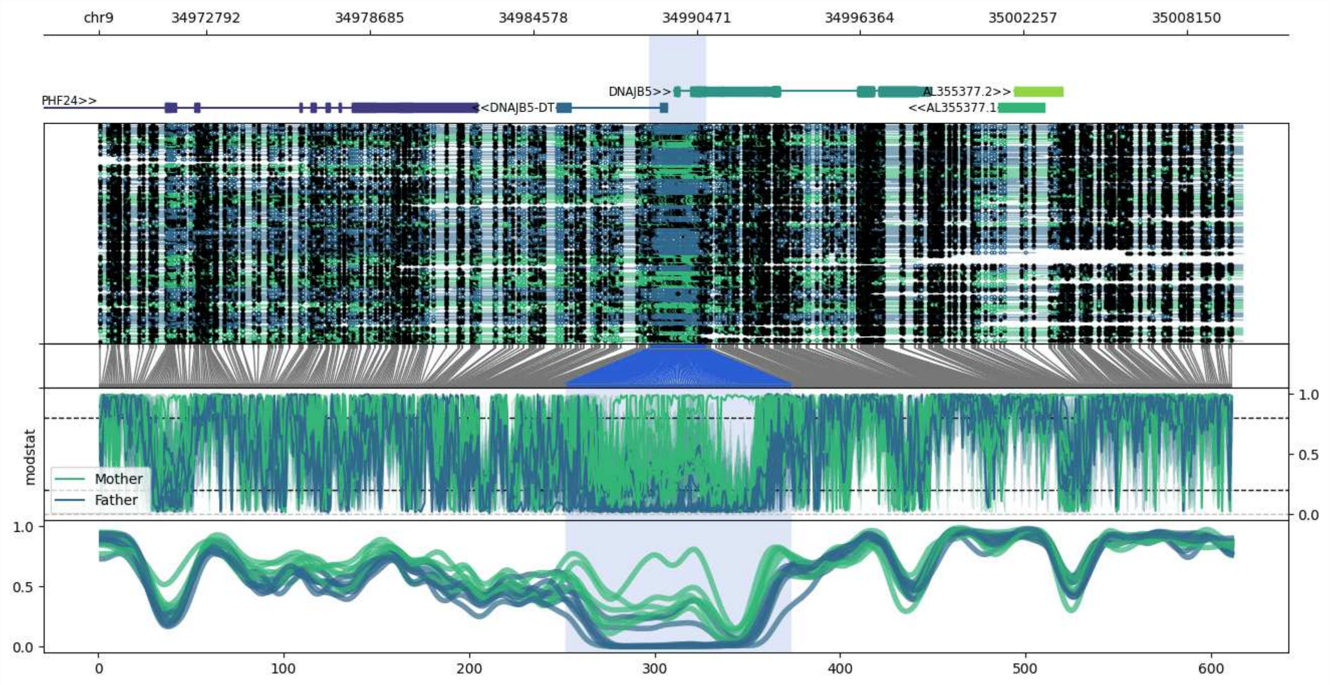


**chr9:34968895-35010147:** Region with 1 DMR containing genes: AL355377.1, AL355377.2, AL355377.3, DNAJB5, DNAJB5-DT, PHF24, SYF2P2.

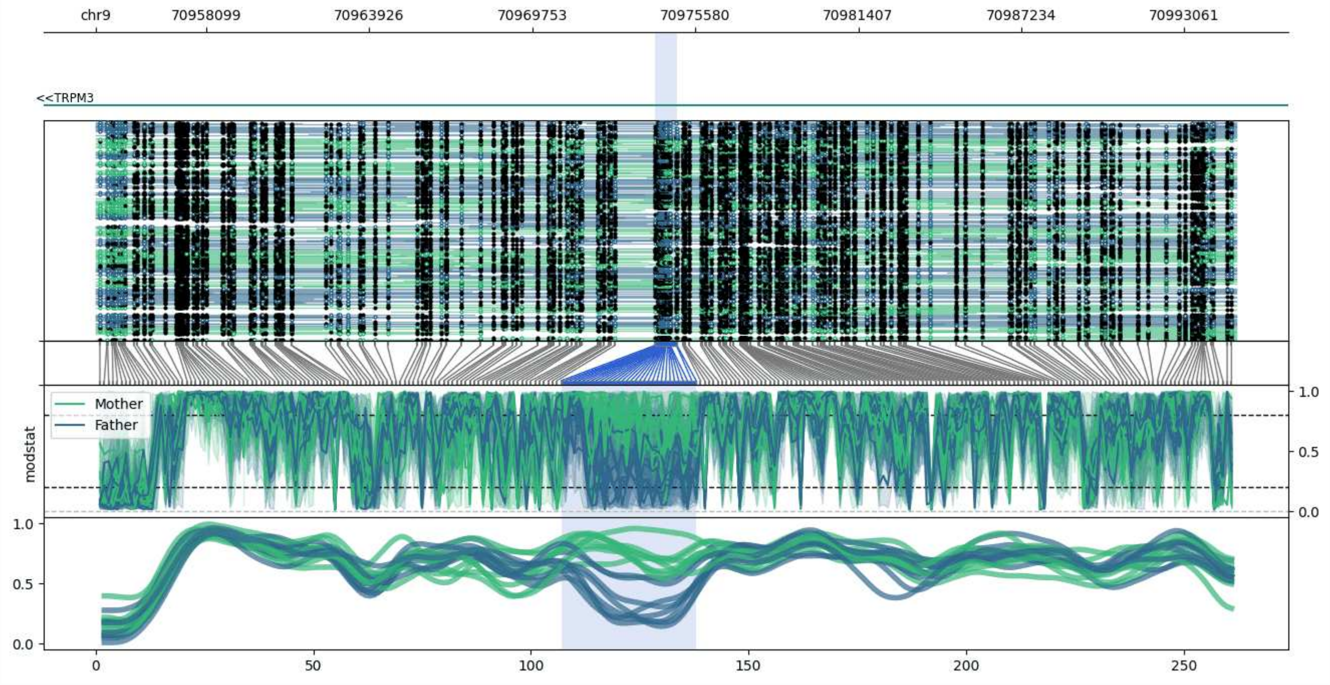


**chr9:70954134-70994927:** Region with 1 DMR containing genes: TRPM3.

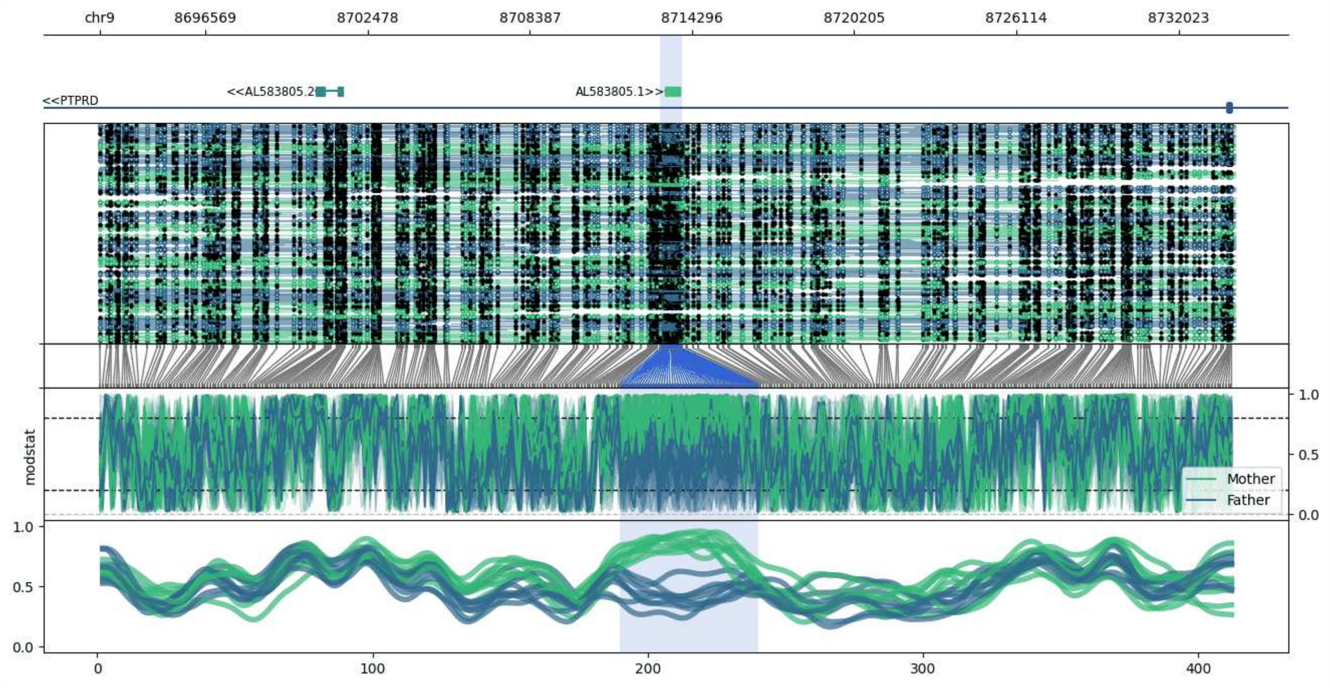


**chr9:8692707-8734076:** Region with 1 DMR containing genes: AL583805.1, AL583805.2, PTPRD.

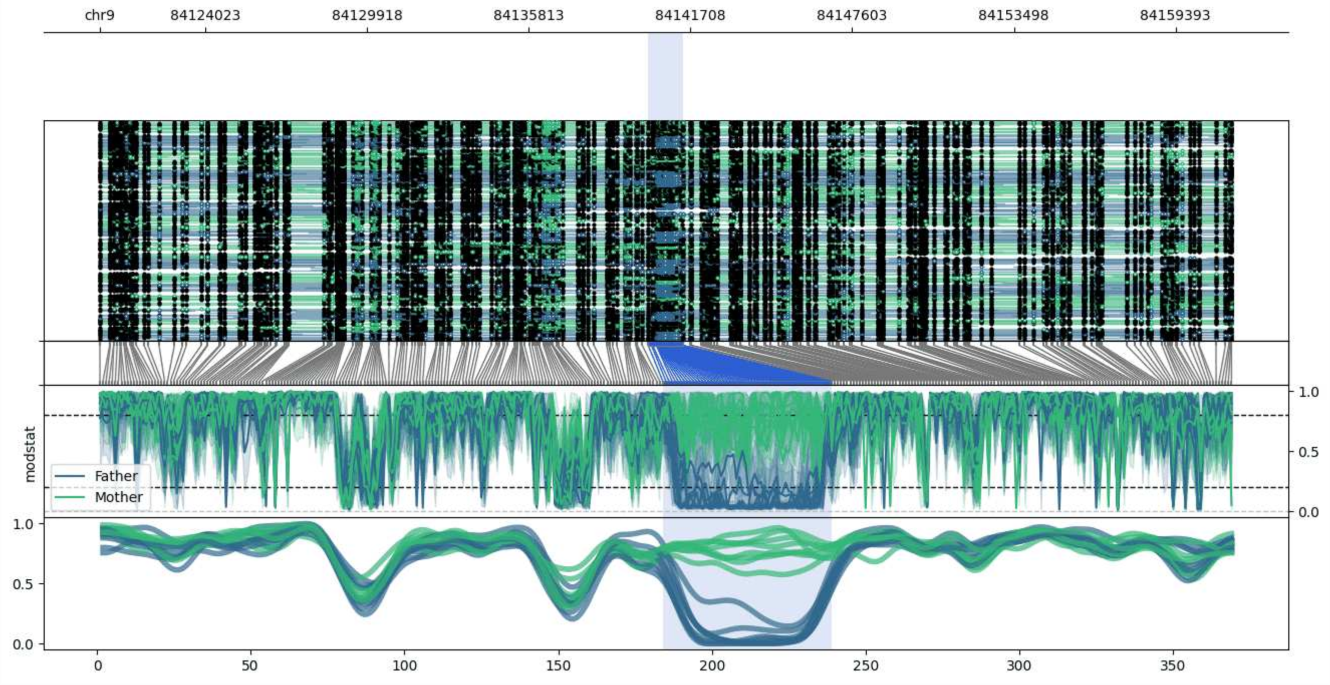


**chr9:84120170-84161441:** Region with 1 DMR containing no genes.

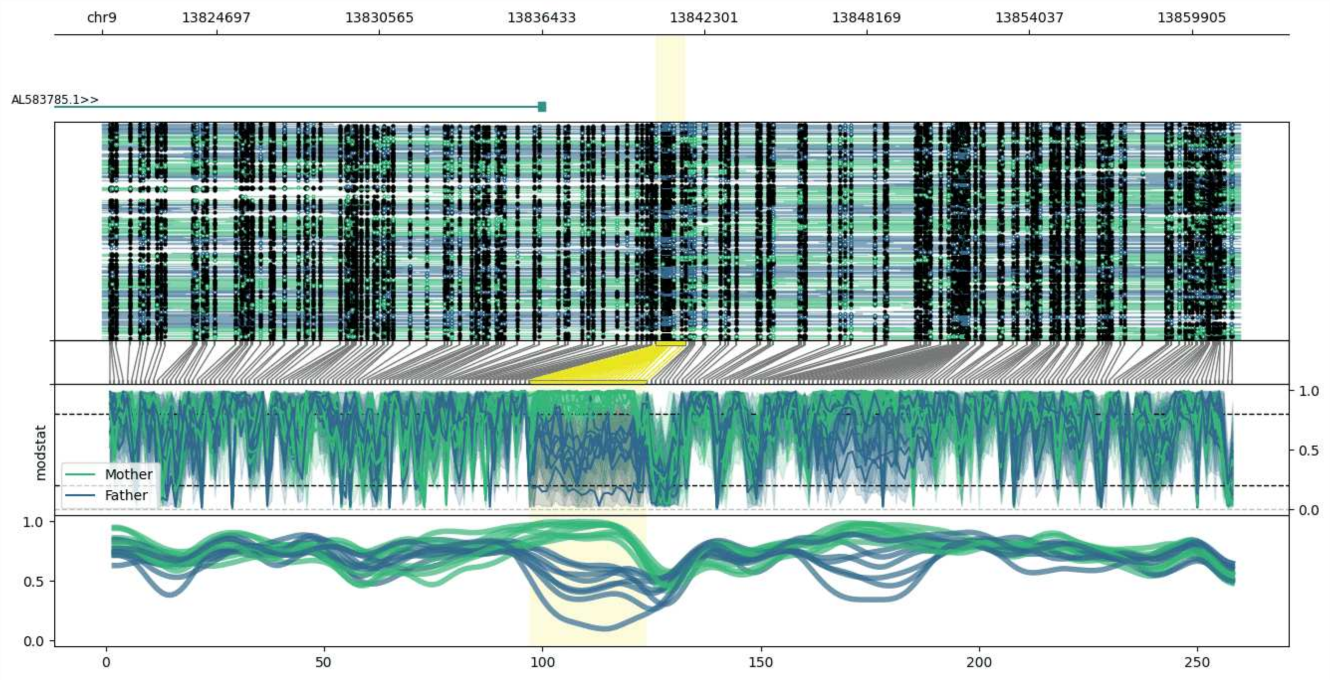


**chr9:13820551-13861628:** Region with 1 DMR containing genes: AL583785.1.

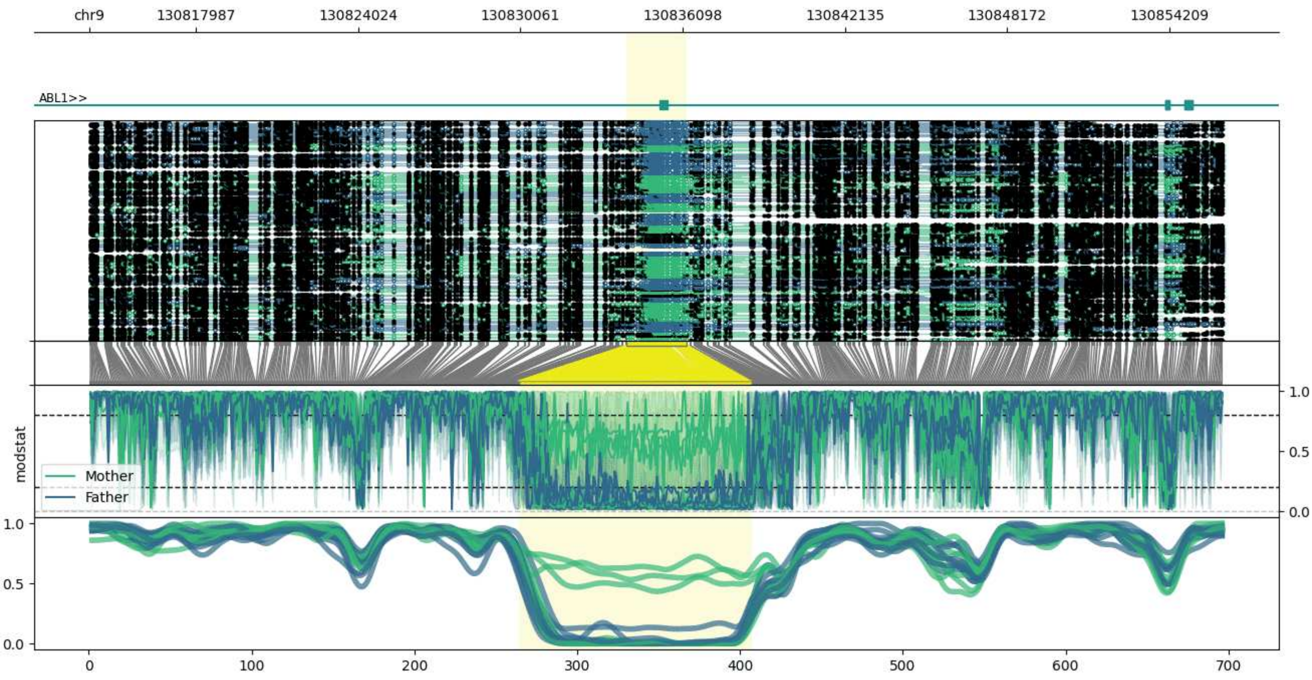


**chr9:130814009-130856268:** Region with 1 DMR containing genes: ABL1.

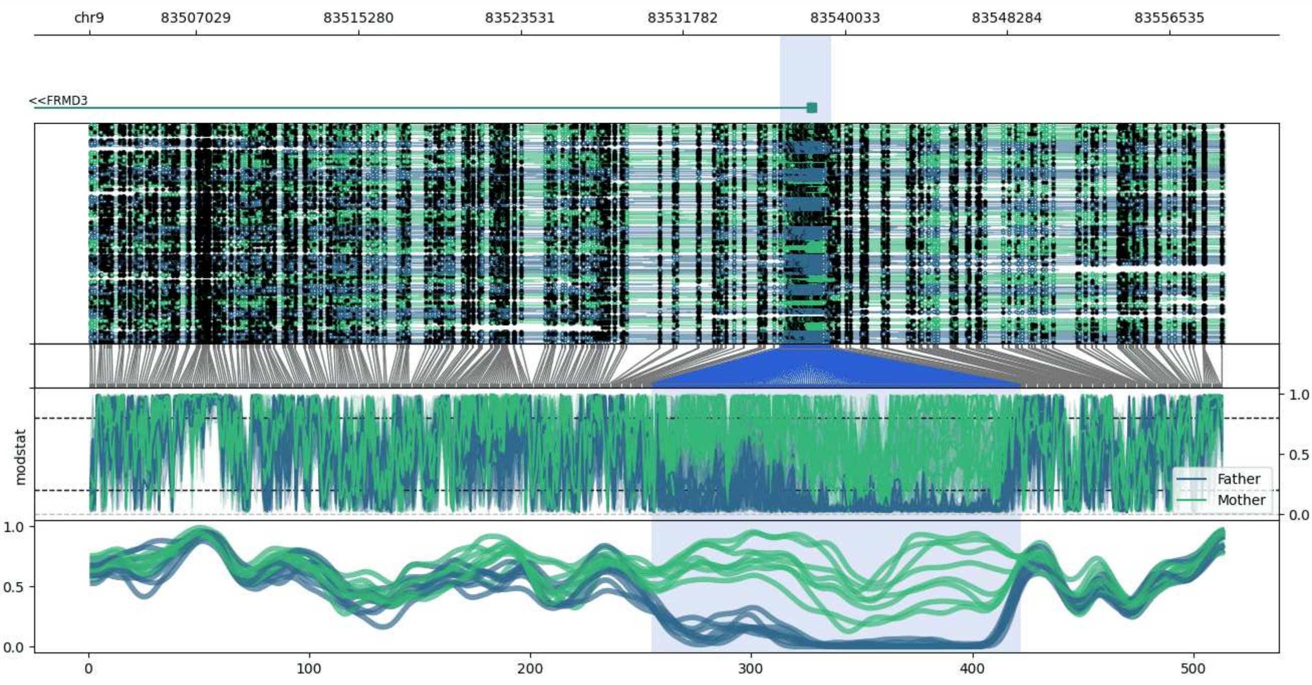


**chr9:83501586-83559345:** Region with 1 DMR containing genes: FRMD3.

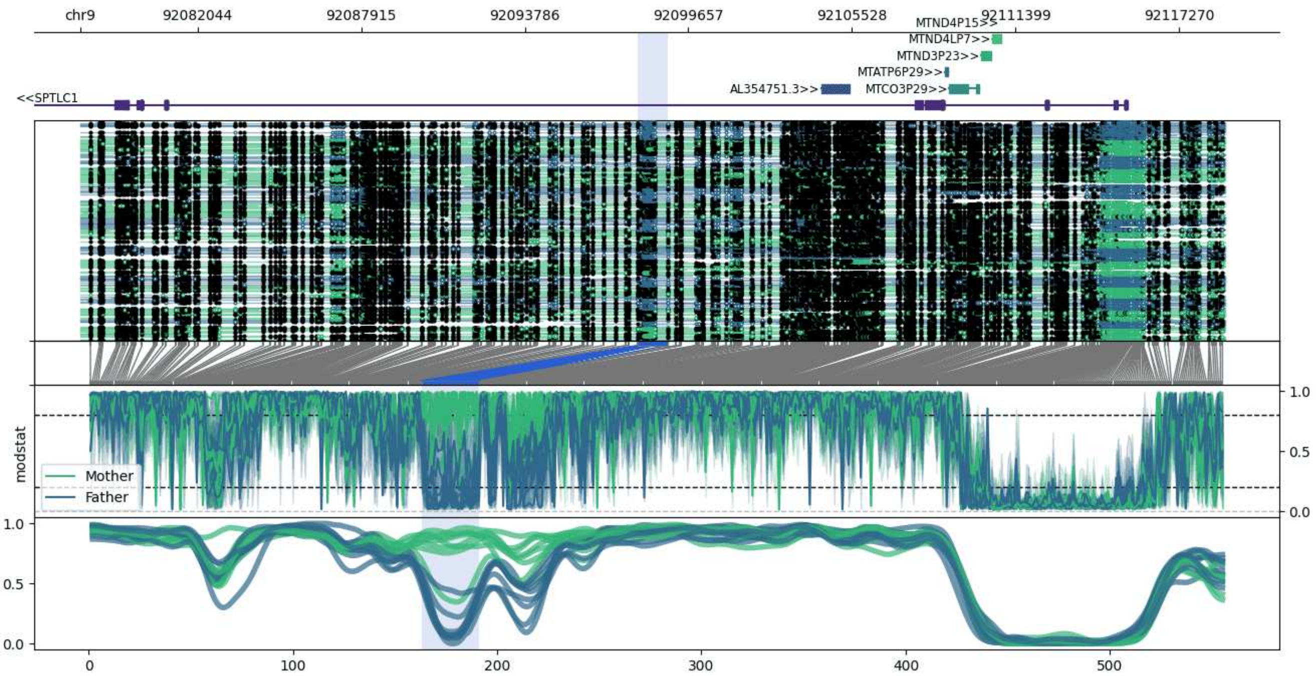


**chr9:92077826-92118923:** Region with 1 DMR containing genes: AL354751.3, MTATP6P29, MTCO3P29, MTND3P23, MTND4LP7, MTND4P15, SPTLC1.

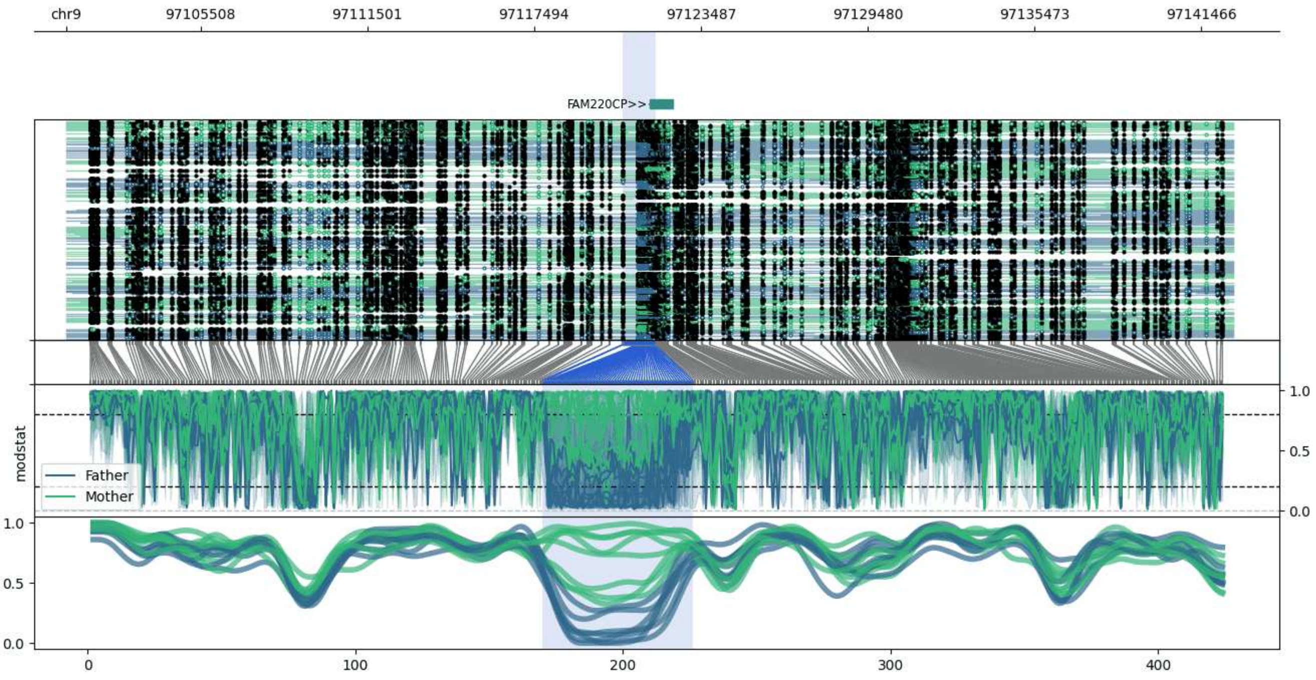


**chr9:97100676-97142627:** Region with 1 DMR containing genes: AL445670.1, FAM220CP, VN1R51P.
